# Multiple variants of the type VII secretion system in Gram-positive bacteria

**DOI:** 10.1101/2024.01.30.577966

**Authors:** Stephen R. Garrett, Andrew B. Higginson, Tracy Palmer

## Abstract

Type VII secretion systems (T7SS) are found in bacteria across the Bacillota and Actinomycetota phyla and have been well described in *Staphylococcus aureus*, *Bacillus subtilis* and pathogenic mycobacteria. The T7SS from Actinomycetota and Bacillota share two common components, a membrane-bound EccC/EssC ATPase and EsxA, a small helical hairpin protein of the WXG100 family. However, they also have additional phylum-specific components, and as a result they are termed the T7SSa (Actinomycetota) and T7SSb (Bacillota), respectively. Here we identify additional organisations of the T7SS across these two phyla and describe eight additional T7SS subtypes which we have named T7SSc – T7SSj. T7SSd is found exclusively in Actinomycetota including the *Olselnella* and *Bifodobacterium* genus, whereas the other seven are found only in Bacillota. All of the novel subtypes contain the canonical ATPase (TsxC) and the WXG100-family protein (TsxA). Most of them also contain a small ubiquitin-related protein, TsxB, related to the T7SSb EsaB/YukB component. Protein kinases, phosphatases and forkhead associated (FHA) proteins are often encoded in the novel T7SS gene clusters. Candidate substrates of these novel T7SS subtypes include LXG-domain and RHS proteins. Predicted substrates are frequently encoded alongside genes for additional small WXG100-related proteins that we speculate serve as co-secretion partners. Collectively our findings reveal unexpected diversity in the T7SS in Gram-positive bacteria.

## Introduction

Protein secretion systems are ubiquitous in prokaryotes. In Gram-negative bacteria there are at least ten distinct secretion systems (Filloux, 2022). Some of these, for example the type III secretion system, mediate translocation of substrate proteins directly across the cell envelope in a single step. Others are two-step pathways where the substrate is first exported to the periplasm by the general secretory (Sec) or twin arginine (Tat) transporters prior to passage across the outer membrane. Gram-positive bacteria generally have simpler cell envelope organisations and therefore lack the specialised systems found in Gram-negative bacteria (Filloux, 2022).

In 2003, a novel protein secretion system was described in pathogenic mycobacteria and was termed the type VII secretion system (T7SS) (Hsu *et al*., 2003, Pym *et al*., 2003, Stanley *et al*., 2003). The T7SS localises to the cytoplasmic (inner) membrane of mycobacteria and operates in parallel to Sec and Tat to mediate transport across this bilayer (Beckham *et al*., 2021, Bunduc *et al*., 2021). Some of the components of the T7SS were also shown to be present in many Gram-positive Bacillota including *Staphylococcus aureus* and *Bacillus subtilis* (Pallen, 2002). The T7SS is best characterised from *Mycobacterium tuberculosis*, where is it found in five distinct copies, termed ESX-1 – ESX-5. All five ESX systems comprise the membrane-bound components EccB, EccC, EccD and the mycosin protease MycP (Fig.1a). A fifth membrane protein, EccE, is also found in all ESX systems except for ESX-4, where it is largely absent (Bunduc *et al*., 2020, Lagune *et al*., 2021).

High resolution structures of purified ESX-5 have been reported (Beckham *et al*., 2021, Bunduc *et al*., 2021). The complex has a hexameric arrangement and a mass in excess of 2.3 MDa. Six copies of EccC are located at the centre of the machinery, forming the secretion pore. EccC is a AAA+ ATPase related to the DNA translocase FtsK and has two transmembrane domains (TMD) at its N-terminus followed by four nucleotide binding domains (Famelis *et al*., 2019). EccC forms interactions with the EccB and EccD subunits, while EccE located at the periphery of the complex (Beckham *et al*., 2017, Famelis *et al*., 2019, Poweleit *et al*., 2019, Beckham *et al*., 2021, Bunduc *et al*., 2021). MycP, which is loosely associated with the machinery, forms a trimeric cap at the periplasmic side, and may proteolytically process some substrates as they are secreted (Ohol *et al*., 2010, Bunduc *et al*., 2021) (Fig 1a). A second, cytoplasmic AAA+ ATPase, EccA, of unknown function is encoded alongside all ESX systems with the exception of ESX-4 (Gao *et al*., 2004, Converse & Cox, 2005, Crosskey *et al*., 2020).

**Figure 1.**
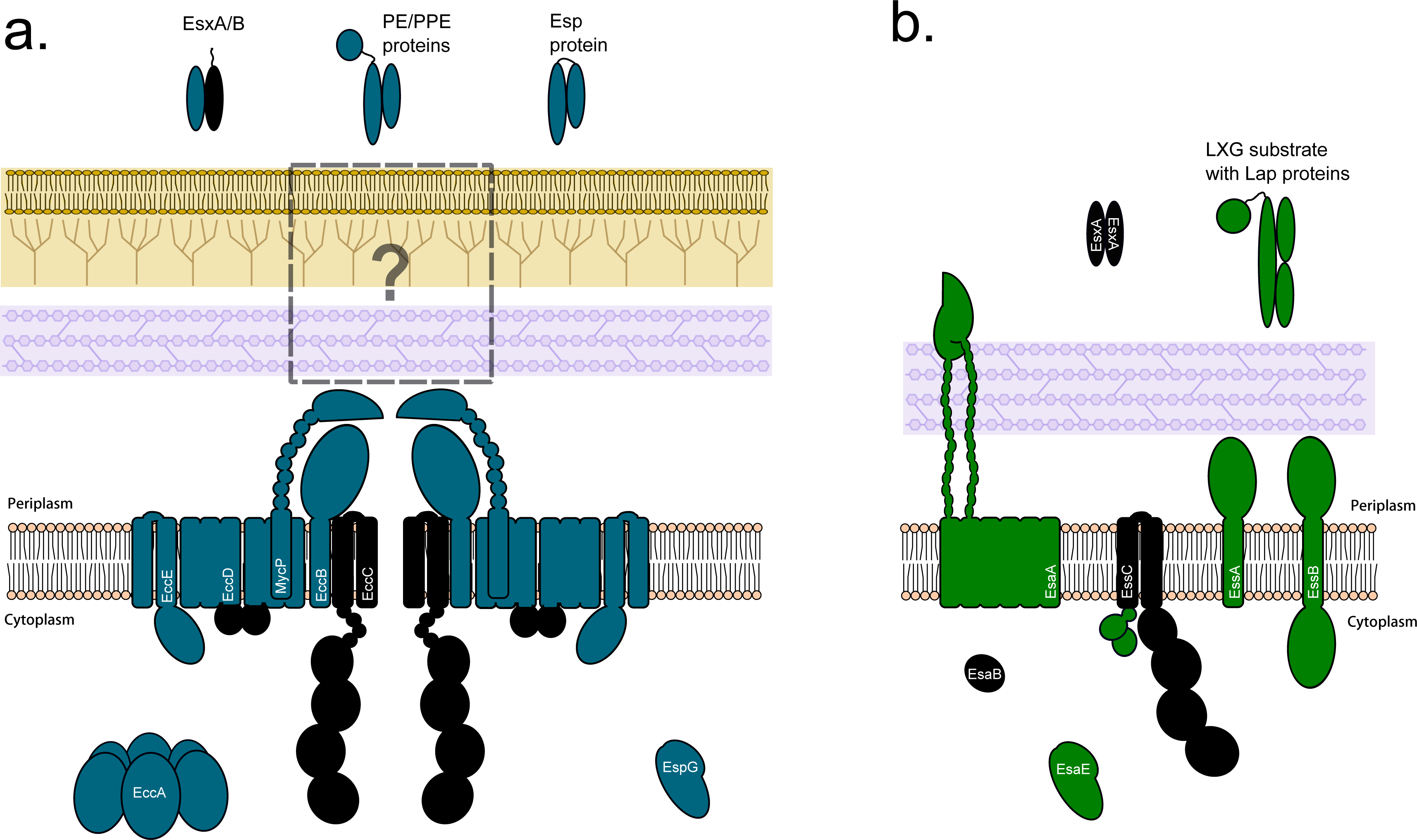
Schematic representation of (*left*) the T7SSa and (*right*) T7SSb systems. Core components that are common to the two systems are shaded in black. Note that EsaE is only found in some T7SSb systems.

The T7SS of Bacillota such as *S. aureus* is distantly related to mycobacterial ESX, with the AAA+ ATPase being the only common membrane component between the two systems (Pallen, 2002). This has led to the two systems being termed T7SSa (Actinomycetota) and T7SSb (Bacillota), respectively (Abdallah *et al*., 2007). In the T7SSb the ATPase is named EssC, and although it shares similar architecture to EccC, it is larger due to the presence of two forkhead associated (FHA) domains at the N-terminus (Tanaka *et al*., 2007, Zoltner *et al*., 2016). Three other membrane proteins, EssA, EssB and EsaA, dissimilar to ESX components in sequence and structure, are essential components of the *S. aureus* and *B. subtilis* T7SS (Fig. 1b) (Burts *et al*., 2005, Baptista *et al*., 2013, Huppert *et al*., 2014, Kneuper *et al*., 2014). EsaA has an extended extracellular domain that spans the cell wall raising the possibility that the Bacillota T7SS may form a conduit for release of substrates at the cell surface (Sao-Jose *et al*., 2006, Klein *et al*., 2021). A small cytoplasmic protein with a ubiquitin fold, EsaB (YukB in *Bacillus subtilis*), is a further essential component of the T7SSb (van den Ent & Lowe, 2005, Kneuper *et al*., 2014, Casabona *et al*., 2017). An EsaB-like domain is also found at the C-terminus of the T7SSa component EccD (Famelis *et al*., 2019, Poweleit *et al*., 2019), suggesting that this domain is a common feature of T7SS.

A further commonality between the T7SSa and T7SSb is in the requirement for proteins of the WXG100 family for function (Burts *et al*., 2005, Rosenberg *et al*., 2015). These are small helical hairpins of approximately 100 amino acids that are secreted as folded dimers by the T7SS (Renshaw *et al*., 2005, Sundaramoorthy *et al*., 2008, Sysoeva *et al*., 2014). A conserved Trp-Xaa-Gly motif is found at the hairpin hinge, giving the family its name (Pallen, 2002). In the T7SSb systems a single WXG100 protein, EsxA, is co-encoded with the secretion machinery and is essential for its activity (Huppert *et al*., 2014, Kneuper *et al*., 2014). In the T7SSa systems WXG100 proteins usually occur in pairs, which heterodimerise (Renshaw *et al*., 2002). Some WXG100 proteins carry a C-terminal ‘signal sequence’ that binds in a pocket on the ATPase domains of EccC/EssC, controlling ATPase activity and promoting interaction between protomers (Champion *et al*., 2006, Rosenberg *et al*., 2015, Mietrach *et al*., 2020).

Other substrates of the T7SSa are the PE-PPE protein families which are rich is proline and glutamate residues and are encoded by highly expanded gene clusters in pathogenic mycobacteria (Cole *et al*., 1998, Abdallah *et al*., 2006). These are also heterodimers of a PE and a PPE protein, which interact through their helical hairpin N-termini to form a four-helix bundle (Strong *et al*., 2006). A C-terminal signal sequence is present on the PE partner (Daleke *et al*., 2012). PE-PPE complexes bind dedicated EspG chaperones which both stabilise the complex by shielding a hydrophobic patch at the tip of the helical bundle and play a role in targeting the substrates to the cognate ESX system for secretion (Ekiert & Cox, 2014, Korotkova *et al*., 2014, Phan *et al*., 2017). Esp proteins are the third substrate family of the T7SSa. EspB adopts an N-terminal four helix bundle fold that carries both a WXG motif and a signal sequence and is secreted without an apparent binding partner (Solomonson *et al*., 2015). EspK was recently identified as a chaperone for EspB and binds to the tip of the EspB helical bundle, in an analogous fashion to EspG (Gijsbers *et al*., 2023).

PE-PPE and Esp proteins are not substrates for the T7SSb. Instead, this system secretes proteins of the LXG family (Whitney *et al*., 2017, Yang *et al*., 2023). LXG proteins are related to WXG100 proteins but are longer, with a helical LXG domain encompassing the first approximately 190 amino acids (Yang *et al*., 2023, Klein *et al*., 2024). LXG domains alone are not competent for secretion, and interact with small helical partner proteins, termed LXG- associated α-helical proteins (Laps) that are essential for export (Klein *et al*., 2022). Some Laps are proteins of the WXG100 family, whereas others share the same fold and similar size but lack the WXG motif (Klein *et al*., 2022, Yang *et al*., 2023). Structural analysis of LXG-Lap complexes reveal that they share striking similarity with the PE-PPE and Esp complexes of the T7SSa (Klein *et al*., 2024). Some LXG proteins, for example *S. aureus* EsaD, also require a chaperone, EsaE for export (Cao *et al*., 2016). Despite sharing very low sequence identity with EspG, EsaE is predicted to share the same EspG fold, and is also implicated in targeting of EsaD to the T7SSb (Cao *et al*., 2016, Yang *et al*., 2023). Very recently a second class of T7SSb substrate was identified. TslA is an antibacterial lipase toxin secreted by the *S. aureus* T7SS. TslA has a very unusual reverse arrangement of domains, with the helical LXG-like domain present at the C-terminus rather than the N-terminus. Two small Lap proteins interact with this C-terminal domain to generate a composite targeting sequence (Garrett *et al*., 2023). It is not yet known whether the T7SSa also secretes proteins with reverse domain organisation.

During our analysis of the T7SSb, we noted that the Bacillota bacterium *Bacillus anthracis*, which has a functional T7SS, lacked genes for the EssA, EssB and EsaA components. Instead, a different suite of conserved genes flanks *essC*. Despite the lack of these core components, the *B. anthracis* T7SS was still competent for the secretion of a WXG100 protein, EsxB (Garufi *et al*., 2008). This led us to investigate the diversity of EssC homologues and their encoding gene clusters among Gram-positive bacteria. From this we identify a further eight phylogenetically distinct EssC-like ATPases, each of which is encoded alongside unique sets of genes that are likely to code for further secretion system components. We propose that these represent T7SSc – T7SSj.

## Methods

To identify homologues of EssC across a diverse range of bacteria, an EssC homologue from *Brevibacillus brevis* was used to perform a BLASTp search against the RefSeq database (Altschul *et al*., 1990). An accession list generated from this BLASTp output was used to run FlaGs2 v1.1.2 (Saha *et al*., 2021). Further genetic neighbourhood analysis was performed following a similar pipeline, but with accession lists submitted to webFlaGs (Saha *et al*., 2021). Genetic neighbourhoods were displayed using Clinker (Gilchrist and Chooi, 2021).

Protein alignments were performed using MUSCLE v3.8.1551 (Edgar, 2004), and visualised with boxshade (https://github.com/mdbaron42/pyBoxshade). For the analysis of EssC diversity across the novel T7SS subtypes, a representative sample of EssC homologues were taken from the FlaGs2 output (Supplementary Data 1). These were aligned using MUSCLE, as described, and MEGA X was used to build a maximum likelihood tree, using the JTT matrix and 100 bootstraps (Kumar *et al*., 2018).

For the identification of domains and further analysis of proteins predicted to form the core components of each novel T7SS, amino acid sequences were submitted to a number of prediction software. For domain and function prediction based on sequence homology, proteins were submitted to BLASTp (Altschul *et al*., 1990) and the InterPro server (Jones *et al*., 2014, Paysan-Lafosse *et al*., 2023). A predicted structure of each component was also obtained from AlphaFold2 version 1.5.5 (Mirdita *et al*., 2022), which was then submitted to FoldSeek (van Kempen *et al*., 2023) for domain and functional predictions based on structural similarity. To identify predicted signal peptides, sequences were submitted to SignalP 5.0 (Almagro Armenteros *et al*., 2019) and to identify predicted transmembrane helices, sequences were submitted to DeepTMHMM (Hallgren *et al*., 2022). A summary of the output from these analyses can be found in Supplementary Data 2, and the AlphaFold2 models for each of the predicted components is included in Supplementary Data 3. Structural model alignments were performed on ChimeraX v1.4 (Pettersen *et al*., 2021) using the matchmaker tool.

For the identification of *esaB* and *esxA* homologues in T7SSd clusters, nucleotide profile Hidden Markov Models for these genes were generated using homologues from each of the other T7SS described and aligned using MAFFT v7.489 (Katoh *et al*., 2002). A custom bash script, (adapted from that used in (Garrett *et al*., 2022)) was used to extract copies of both *esaB* and *esxA* from T7SSd clusters, comprising 10,000 nucleotides upstream and downstream of *tsdC*, with an E-value cutoff of 0.05. Script available on Github (https://github.com/stephen-r-garrett/T7SS_variants).

## Results

### The Identification of novel T7SS arrangements

When we aligned EssC from *B. anthracis* with EssC sequences from the well-characterised *S. aureus* and *B. subtilis* T7SSb systems, we noted that while they shared clear sequence homology, the *B. anthracis* protein unexpectedly lacked FHA domains at the N-terminus (Fig S1). Further analysis revealed that EssC proteins from some other Bacillota members, for example *Brevibacillus brevis*, had high similarity to that of *B. anthracis* EssC and also lacked FHA sequences. Using the *B. brevis* EssC sequence we performed an extensive BLAST search against the RefSeq database to identify diverse EssC variants, which were then used to perform gene neighbourhood analysis (Saha *et al*., 2021). This generated a phylogenetic tree for all EssC accessions, along with the genetic neighbourhood of each *essC* based on the tree order (Supplementary Data 1). These data were used together to identify novel arrangements of the T7SS, allowing us to identify at least eight genetically distinct organisations of the T7SS.

To gain a clearer understanding of the genetic diversity of these novel T7SSs, a maximum likelihood phylogenetic tree was constructed using a subset of the EssC accessions from the genetic neighbourhood analysis in Supplementary Data 1, alongside T7SSa EccC and T7SSb EssC sequences (Fig 2). Again, a similar clustering of EssC homologues was observed as with the previous analysis, with eight groups of EssCs distinct from each other and from the homologues in the T7SSa and T7SSb systems. From this and the analysis below we propose the presence of an additional eight T7SS subtypes present in Gram-positive bacteria which we have named T7SSc – T7SSj. An alignment of the EccC/EssC homologues from T7SSc to T7SSj alongside EccC (T7SSa) and EssC (T7SSb) is shown in Fig S2, and all of the accessions we identified for EssC homologues from these systems are included in Supplementary Data 4. Typical genetic arrangements for each new system are shown in Fig. 3.

**Figure 2.**
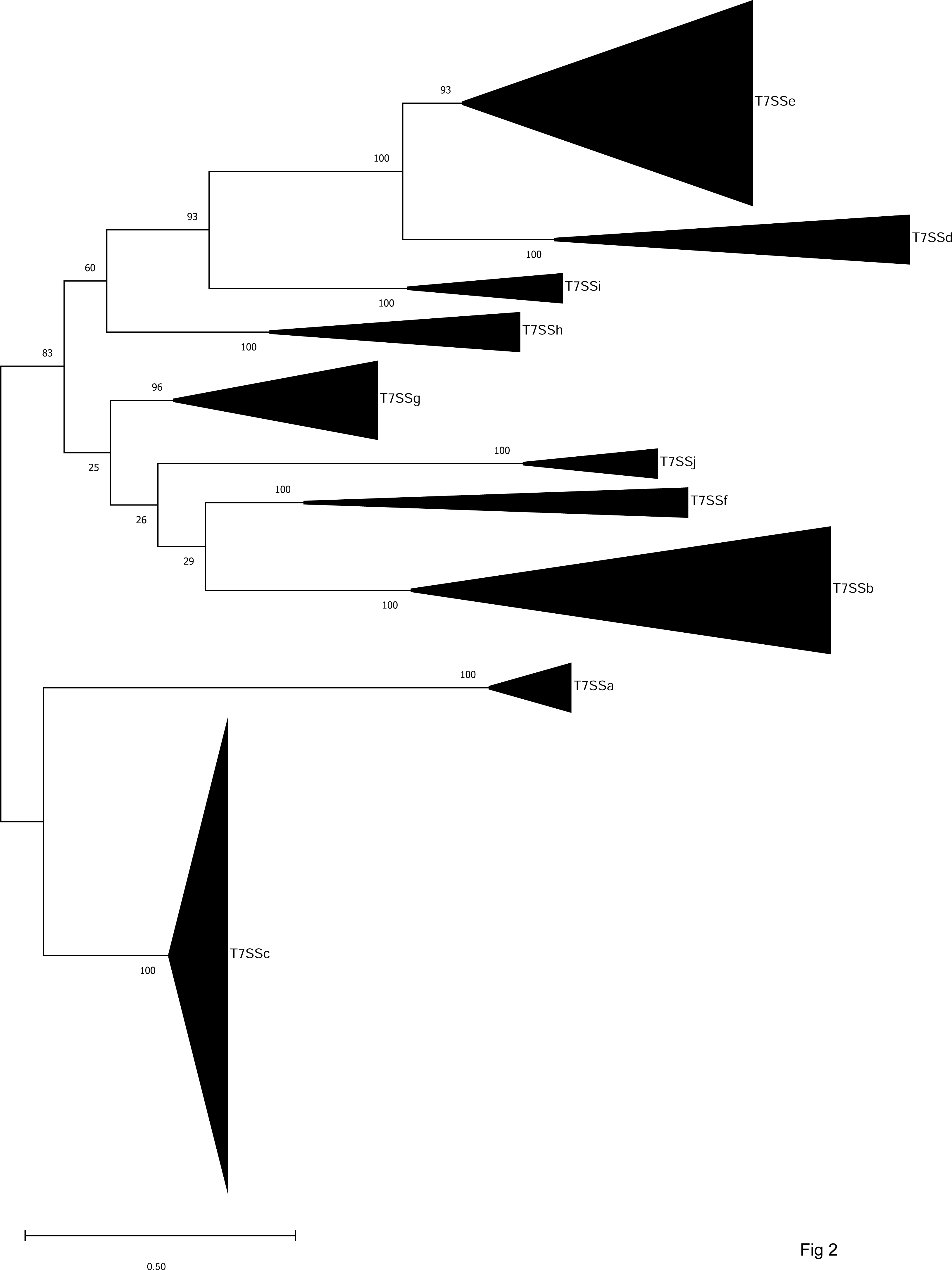
Maximum likelihood tree from an amino acid alignment of a representative sample of EssC/EccC homologues from each T7SS identified in Supplementary Data 1. The nodes are labelled with bootstrap values, based on 100 iterations. The scale bar depicts evolutionary distance by the number of amino acid substitutions per site.

**Figure 3.**
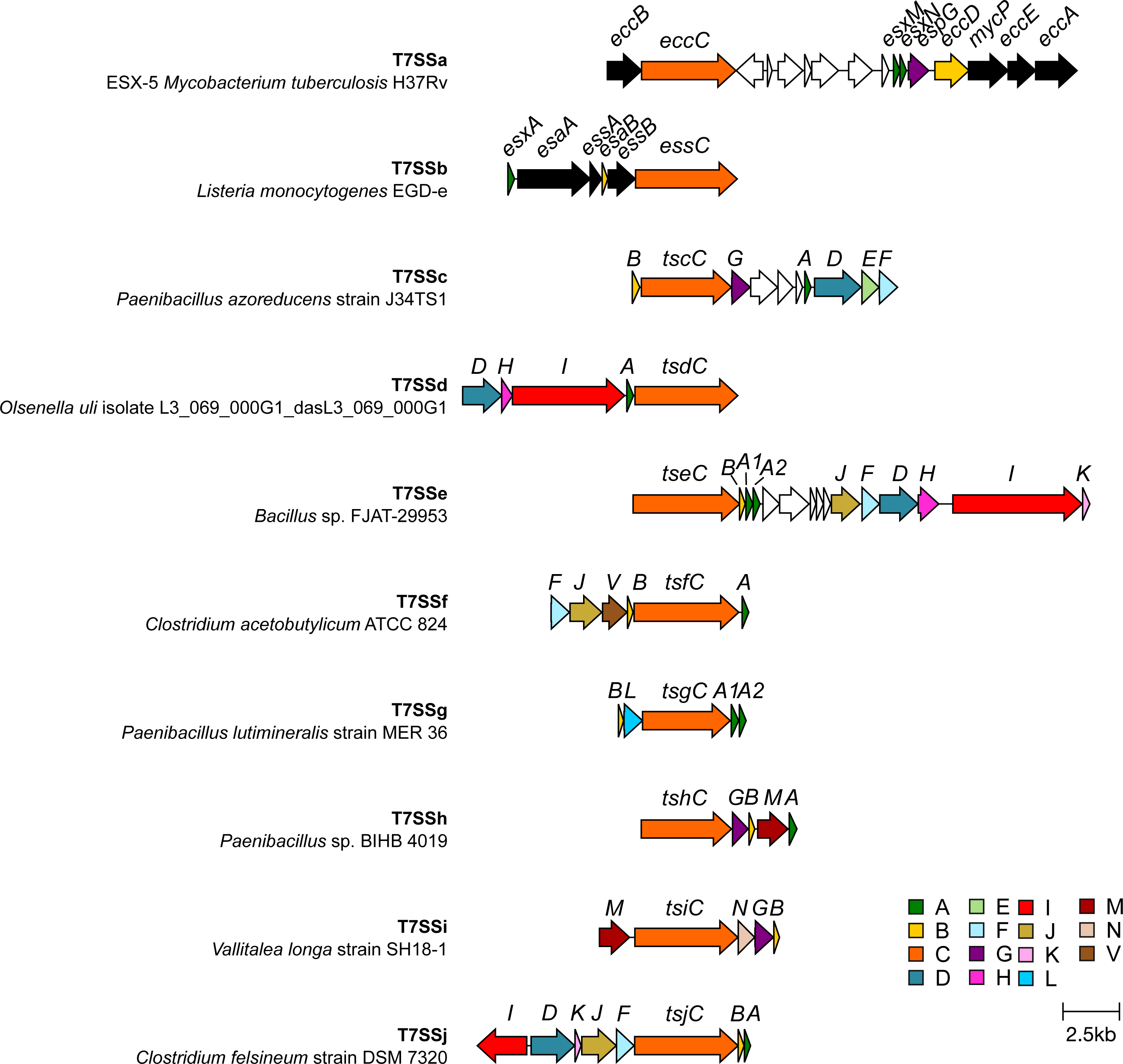
Genetic arrangements of the T7SS from the indicated organisms. Genes coding for components that are related between the different systems are shaded similarly. To simplify the nomenclature of the T7SSc – T7SSj systems we propose to name the components as Ts(x), where ‘x’ refers to the letter associated with that system, so for example Tsc components are found in the T7SSc. The TsxA, TsxB and TsxC components are (almost) universally conserved and are homologues of EsxA, EsaB and EccC/EssC, respectively. No TsxA-encoding gene is shown in the T7SSi gene cluster as we were unable to identify a likely candidate (see text).

### T7SSc

The T7SSc appears to be widespread in many Gram-positive bacteria, including *Paenibacillus, B. anthracis* and *B. brevis*, and after T7SSa and T7SSb had the most representative sequences in the Refseq database. The T7SSc EssC falls into two diverse clades, however the flanking genes, which encode putative additional T7SSc components, are conserved (Supplementary Data 1). While the EssC sequences from the T7SSc are homologous to EssC from the T7SSb, phylogenetically they cluster more closely with the EccC proteins from the T7SSa (Fig. 2). As mentioned above, T7SSc EssCs lacks N-terminal FHA domains which are also absent from the T7SSa EccC but present on the T7SSb EssC (Fig S1, Fig. S2). We propose to name EssC/EccC components as Ts(x)C, where x refers to the specific T7SS subtype, i.e. the EccC component would be TsaC, EssC from the T7SSb would be TsbC, etc (Table 1).

**Table 1.**
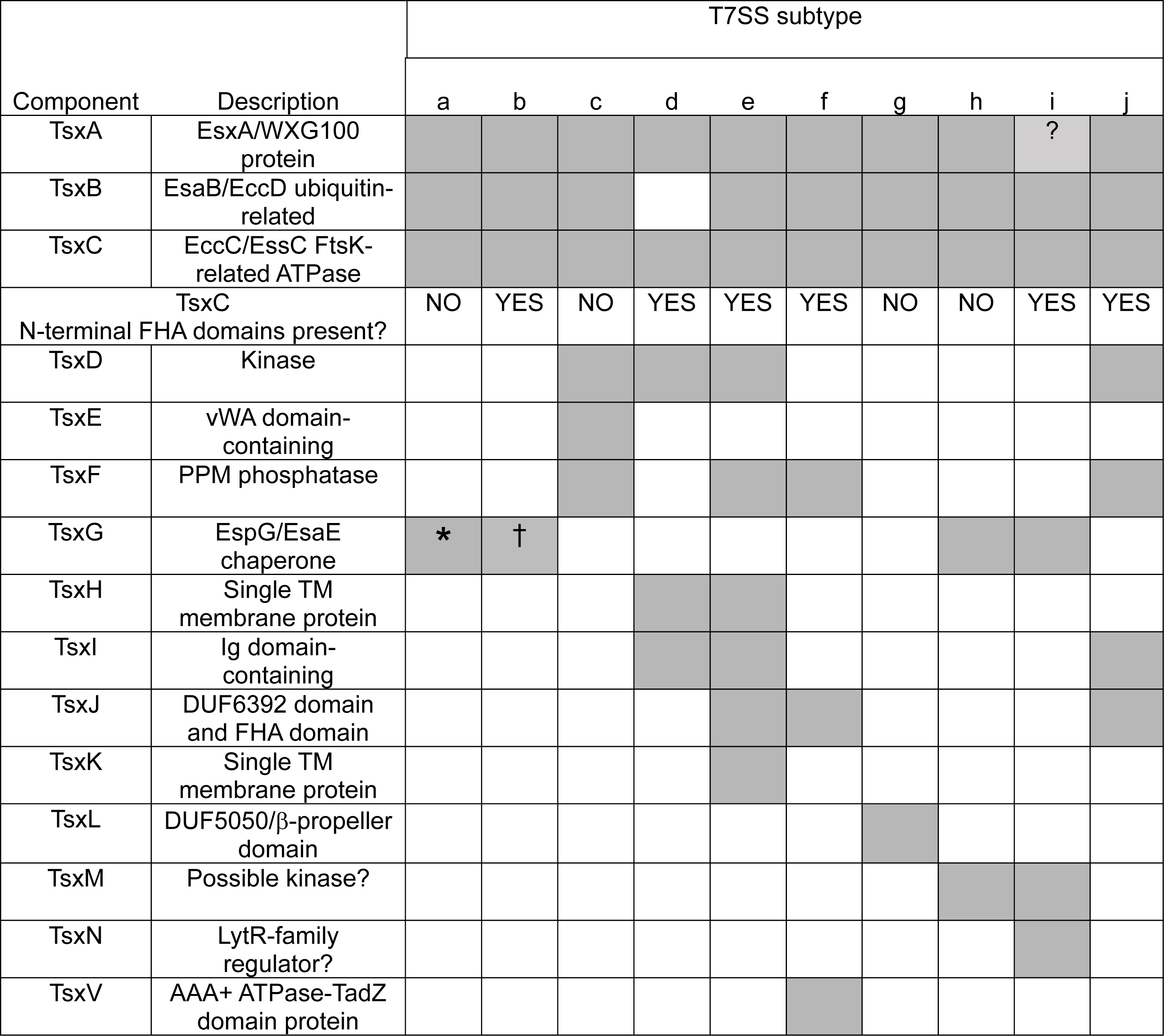
Summary of the components identified in the T7SSc – T7SSj systems and their distribution. * EspG (TsaG) is found in most T7SSa systems with the exception of ESX4. indicated subtype but not all of them. † The EsaE (TsbG) chaperone is found only in some T7SSb systems.indicates that no clear TsxA-encoding gene could be identified among the T7SSi gene clusters.

Based on genetic neighbourhood conservation, we predict that there are seven core components associated with the T7SSc (Fig. 3, Fig 4a, Table 1). To determine the putative domain composition and subcellular arrangement of each of these components, results from BLASTp, SignalP-5.0, DeepTMHMM and InterPro domain searches were collated (Supplementary Data 2). In addition, AlphaFold2 was used to obtain structural predictions (Supplementary Data 3) and these structural models were submitted to FoldSeek to obtain further predictions about possible domains present in these proteins.

**Figure 4.**
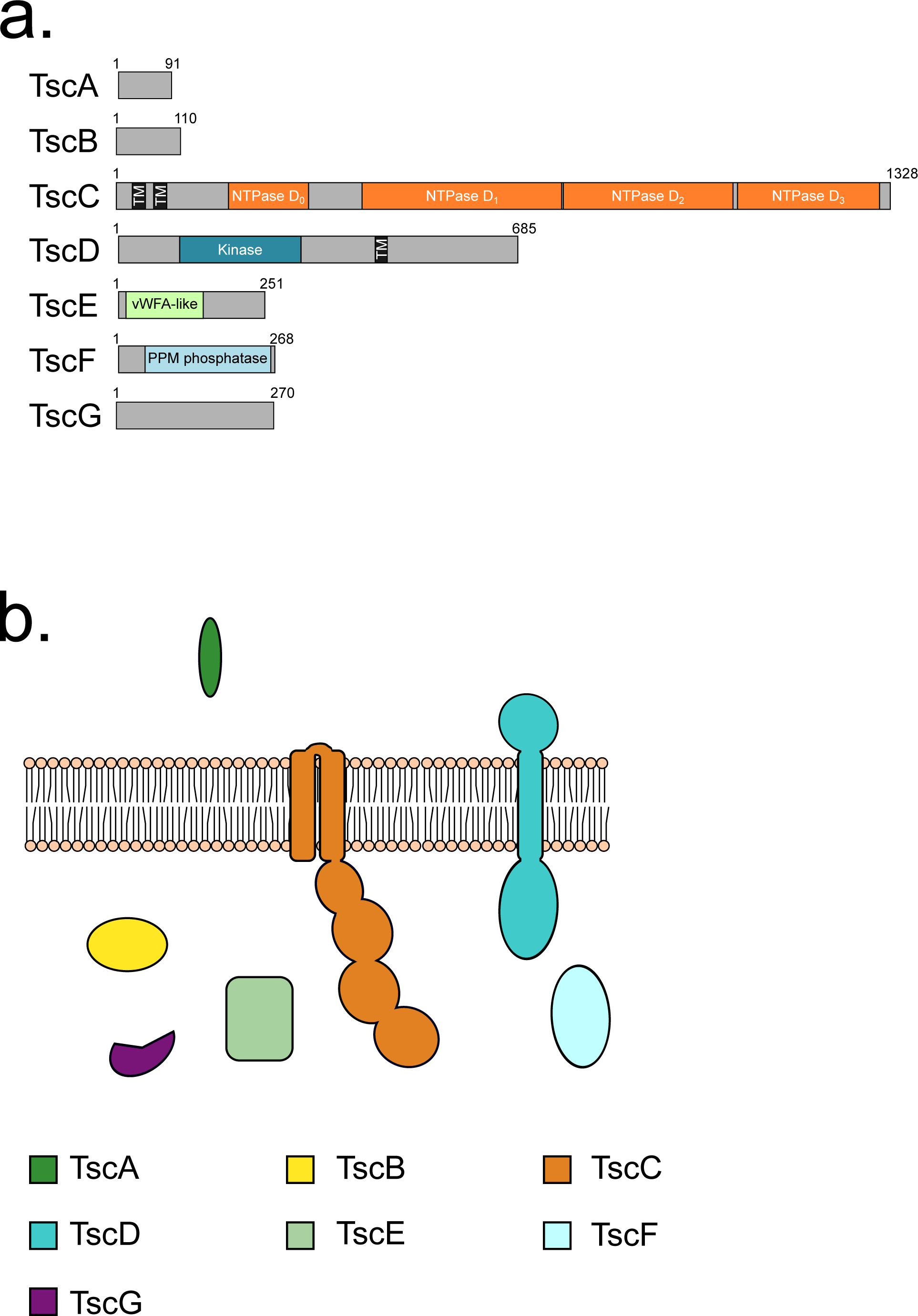
a. Predicted domain arrangement and b. predicted subcellular location of components of the T7SSc.

The T7SSc gene clusters each have a single gene encoding a WXG100 protein homologous to EsxA, which we have termed TscA. This is more akin to the T7SSb system which encodes a single WXG100 protein, EsxA, in the T7SSb gene cluster, whereas the T7SSa systems have a pair of non-identical proteins. The T7SSc gene cluster also encodes a small protein, TscB, related to EsaB/YukD, a ubiquitin-like protein found as a discrete protein in the T7SSb, and as a fused domain in the T7SSa (Fig.4a,b, Supplementary Data 2). Of the other likely T7SSc components, TscG has no recognisable domain predictions from either BLASTp or InterPro domain searches, however, FoldSeek searches using the AlphaFold2 structural prediction suggest it shares homology to the chaperone protein, EspG, from the T7SSa (Supplementary Data 2, Table 1).

There are three further conserved components encoded in T7SSc gene clusters which are not found in either the T7SSa or T7SSb. TscE is predicted to contain a von Willebrand factor type A (vWA) domain at its N-terminus and is likely localised to the cytoplasm due to the absence of any detectable signal peptide or transmembrane helices (Fig.4a,b, Supplementary Data 2). vWA domains often bind metal ions and frequently mediate protein-protein interactions (Whittaker & Hynes, 2002). The other two components, TscD and TscF carry a putative serine/threonine kinase domain and a PPM-phosphatase domain, respectively. While TscF is predicted to be localised to the cytoplasm, TscD is predicted to have at least one transmembrane helix, with the N-terminal kinase domain cytoplasmic (Fig.4a,b; Supplementary Data 2). TscD shares similar topology to the T7SSb component EssB, which also has a cytoplasmic pseudokinase domain (Zoltner *et al*., 2013, Tassinari *et al*., 2022). However, TscD shares no apparent sequence identity with EssB, and unlike EssB the kinase active site is conserved. Analysis of the genome sequences of bacteria that encode the T7SSc failed to identify any further homologues of T7SSa or T7SSb components and we propose that the T7SSc comprises the seven core components TscA – TscG (Table 1).

### T7SSd and T7SSe

All of the other EssC homologues we identified from our analysis clustered more closely with TsbC/EssC of the T7SSb system than with TsaC/EccC from the T7SSa (Fig 2). Of these, the two most distinct from the T7SSb, are those which we have assigned as TsdC and TseC from the T7SSd and T7SSe, respectively. The T7SSd and T7SSe systems are closely related to one another, but the T7SSd is found only in the Actinomycetota genus and the T7SSe only in Bacillota. Both TsdC and TseC have N-terminal FHA domains that are found on EssC/TsbC, but they share little sequence homology with the EssC FHA domains (Fig S2). FHA domains often bind phosphothreonines, linking protein phosphorylation with formation of protein complexes (Weiling *et al*., 2013). However, in the T7SSb, the FHA domains of EssC lack the conserved binding site for phosphothreonine, and form interactions with the cytoplasmic pseudokinase domain of EssB that is presumably not regulated by protein phosphorylation (Bobrovskyy *et al*., 2022, Tassinari *et al*., 2022). Interestingly, the phosphothreonine binding consensus sequence in the FHA domains of TsdC and TseC appears to be well conserved (Fig S2, Fig S3), raising the possibility that the assembly and/or activity of these systems is controlled by threonine phosphorylation.

The T7SSd is predicted to be composed of only five core components, as represented by the gene cluster from *Olsenella uli*: TsdA, TsdC, TsdD, TsdH and TsdI (Fig 3, Fig 5, Table 1). TsdA is a WXG100 protein with homology to other WXG100 family members (Fig.5a, Supplementary Data 2). Similar to the T7SSc, a predicted serine/threonine kinase, TsdD, is also found associated with the T7SSd, and in this system could potentially link phosphorylation events with TsdC-FHA domain interactions. The AlphaFold structural models of TscD and TsdD align closely at their N-terminal kinase domains (Fig 5b), however the C-terminal regions of these two proteins, while both predicted to be extensively helical, are quite diverse. This may potentially reflect differences in the organisation of the cell envelopes of these bacteria which is where the C-terminal regions are predicted to reside. The T7SSd has two further probable components that have not been described in any of the T7SSa-c systems. The first of these is TsdH, composed of a single transmembrane helix and a short, likely unstructured cytoplasmic region (Fig.5a,c, Supplementary Data 2). The second of these novel components is TsdI. TsdI is a large protein (1654 amino acids for *Olsenella uli* TsdI) that has a N-terminal Sec signal peptide and a C-terminal transmembrane helix (Fig.5a,c, Supplementary Data 2).

**Figure 5.**
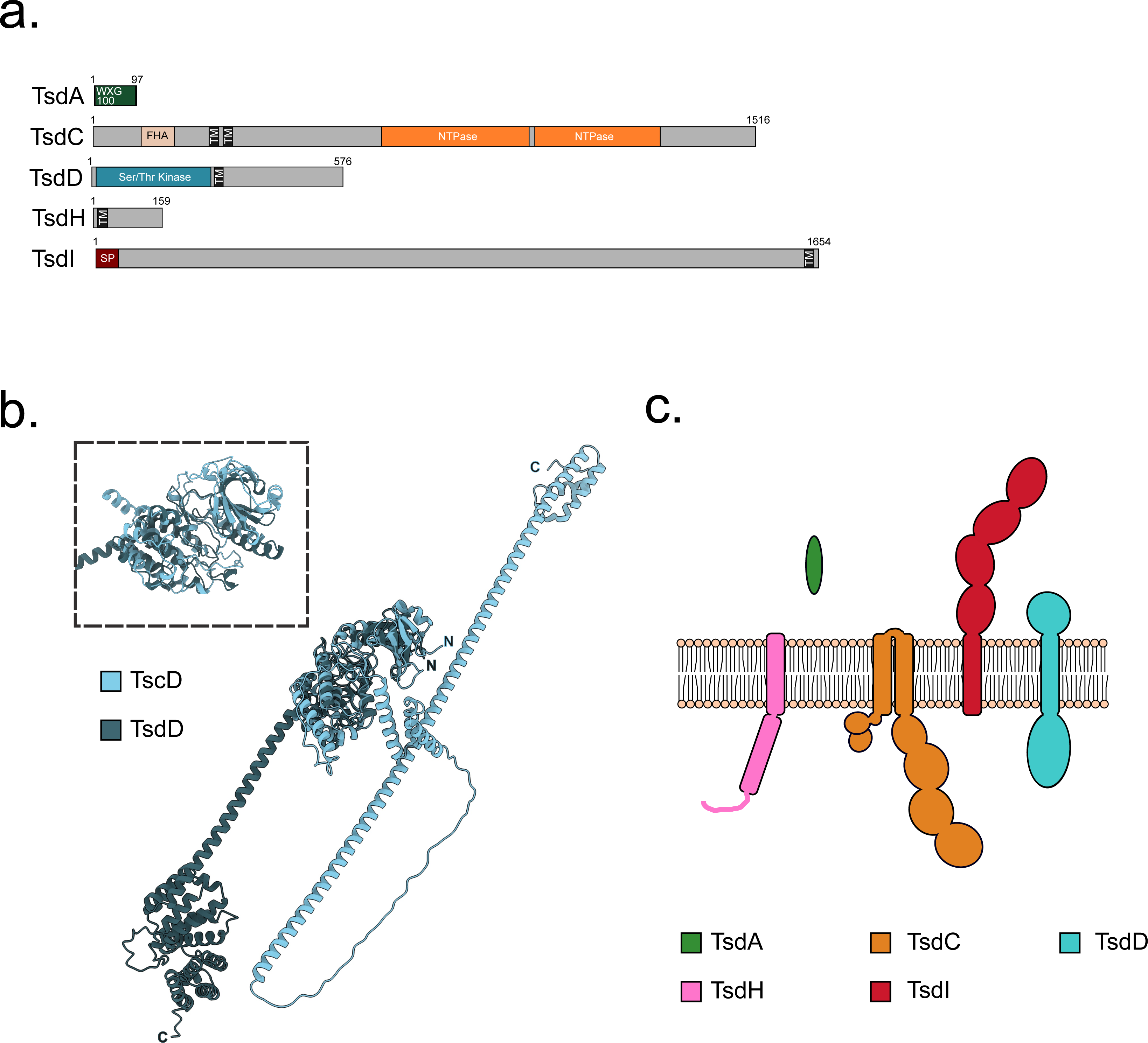
a. Predicted domain arrangement and c. predicted subcellular location of components of the T7SSd. b. Overlay of the structural models for the kinase domains of TscD and TseD. For the structural model, RMSD for the kinase domain between 61 pruned atom pairs is 1.034 angstroms; across all 514 pairs is 97.233.

Whilst not predicted by InterPro domain search, BLASTp and FoldSeek (using the AlphaFold2 model) identify several Ig-like domains along the length of TsdI (Fig 5c, Supplementary Data 2, Supplementary Data 3). We predict that TsdI may span the cell envelope akin to EsaA in the T7SSb system (Klein *et al*., 2021), potentially forming a conduit for substrates and/or playing a role in target recognition.

Surprisingly, no ubiquitin-like protein is found to be associated with the T7SSd, despite a homologue being found associated with all other T7SS (Fig 3 – annotated as gene ‘B’). To confirm whether the open reading frame (which would likely be very short) was present but not annotated, an HMM search was conducted using a custom script, against every known T7SSd locus identified in our analysis. However, no ubiquitin-like homologues were encoded at any T7SSd locus (Supplementary Data 1), and we were unable to identify a homologue encoded elsewhere in the genome by BLAST search.

All of the components we identified as being present in T7SSd are also found in T7SSe. However, we consistently identified additional core components in T7SSe which are not found in the T7SSd (Fig 3, Supplementary Data 1). Moreover, since the T7SSd is exclusively found in Actinomycetota, whereas the T7SSe is found in Bacillota we have assigned these as separate subsystems. In addition to the T7SSd-related components TseA, TseC, TseD, TseH and TseI, the T7SSe systems always encode an additional WXG100 protein (TseA2) as well as the ubiquitin-like protein TseB. TseF, a homologue of the TscF PPM phosphatase found in the T7SSc is also found in the T7SSe (Fig. 6a,b; Supplementary Data 2). Two further components, TseJ and TseK, are also encoded at all T7SSe loci. TseJ carries a DUF6382 domain at the N-terminus and an FHA domain at its C-terminus (Fig S4) and is predicted to be a globular protein localised to the cytoplasm. TseK is predicted to have a single transmembrane helix, with a short unstructured stretch of amino acids facing the cytoplasm (Fig. 6b, Supplementary Data 2).

**Figure 6.**
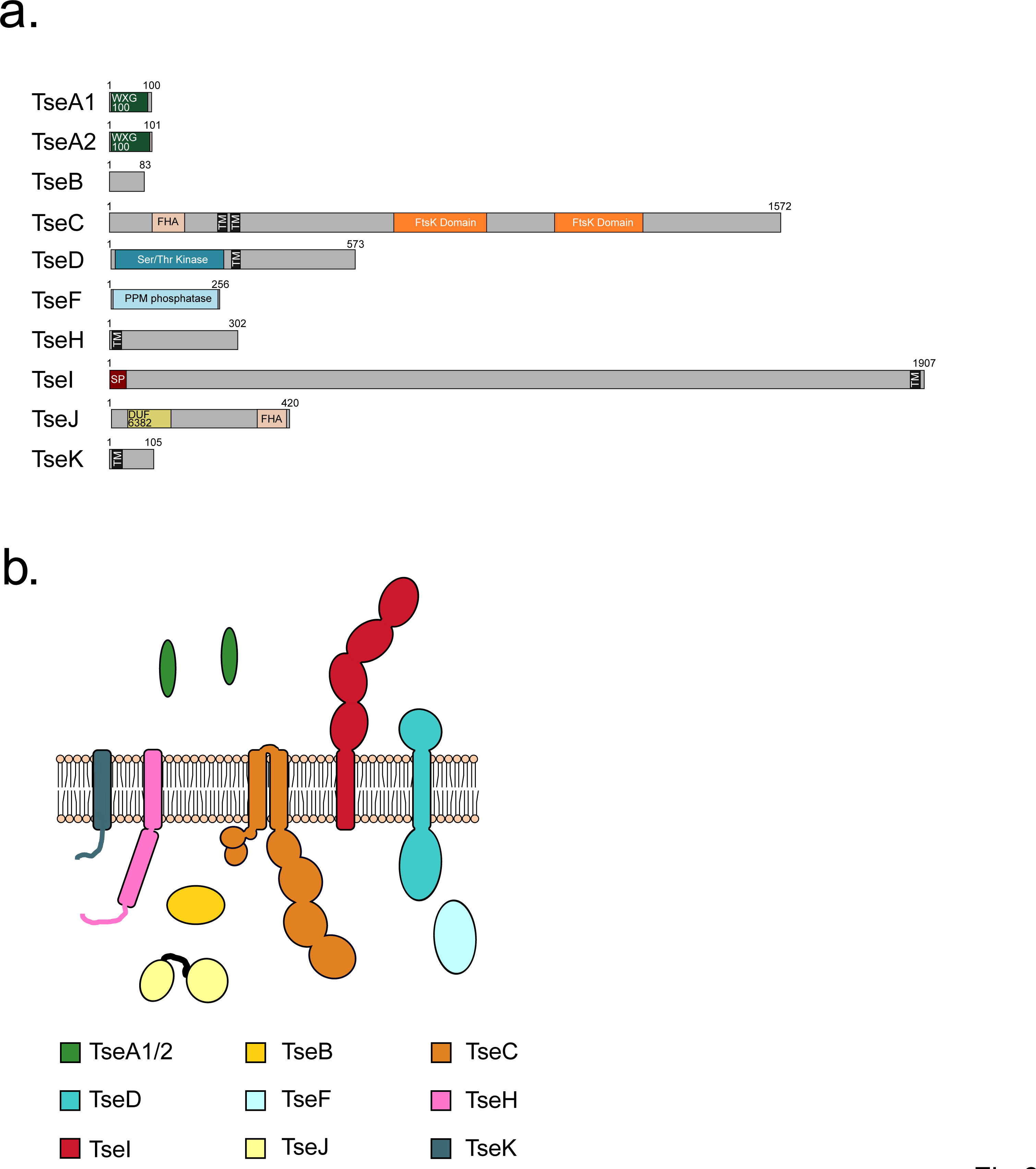
a. Predicted domain arrangement and b. predicted subcellular location of components of the T7SSe.

Five further novel T7SS clusters are found in a narrow species range.

The three novel T7SS organisations described above are present across a range of bacterial genera, allowing us to be relatively confident about the likely components based on clustering of genes. However, in addition to these, we have identified a further five novel T7SS genetic arrangements (T7SSf – T7SSj). Each of these five are found within a single or limited range of bacterial genera, and as a result, we are less confident about concluding the likely core components of each of these.

### T7SSf

The T7SSf is found in several species of *Clostridia*. T7SSf comprises the same three core components found in most T7SS described thus far: TsfA, a WXG100 protein; TsfB, a YukB- like ubiquitin homologue; and TsfC the FtsK-related membrane bound ATPase with N-terminal FHA domains (Fig. 7a, Supplementary Data 2). The phosphothreonine binding motifs in the TsfC FHA domain are only partially conserved (Fig S2). It also contains TsfF, a predicted PPM phosphatase found in the T7SSc and T7SSe systems, and a TsfJ component. The other T7SS systems that contain a PPM phosphatase domain also have a membrane-bound kinase. However, we could find no kinase genetically associated with the T7SSf system. TsfJ is similar to TseJ, harbouring an N-terminal DUF6382 domain and a C-terminal FHA domain (Fig S4), but unlike TseJ is also predicted to have two transmembrane helices, suggesting it is anchored in the membrane with both domains localised to the cytoplasmic side (Fig. 7a,b, Supplementary Data 2). The final component we identified is TsfV, a putative AAA+ ATPase protein. The EccA component found in many T7SSa systems is also a cytoplasmic AAA+ ATPase, however, it shares no detectable homology to TsfV. Unlike EccA, TsfV also has an N-terminal TadZ-like ARD domain (Fig. 7a; Supplementary Data 1). TadZ is an essential component of the tight-adherence (Tad) pilus machinery and has been implicated in its polar localisation (Tomich *et al*., 2007, Xu *et al*., 2012). As the *tsfV* gene is flanked on both sides by T7SS-associated genes we suggest that it is likely to encode a component of the T7SSf. It should be noted while other Tad components are not encoded at the T7SSf cluster, they are encoded at genetic loci of both the T7SSg and T7SSh.

**Figure 7.**
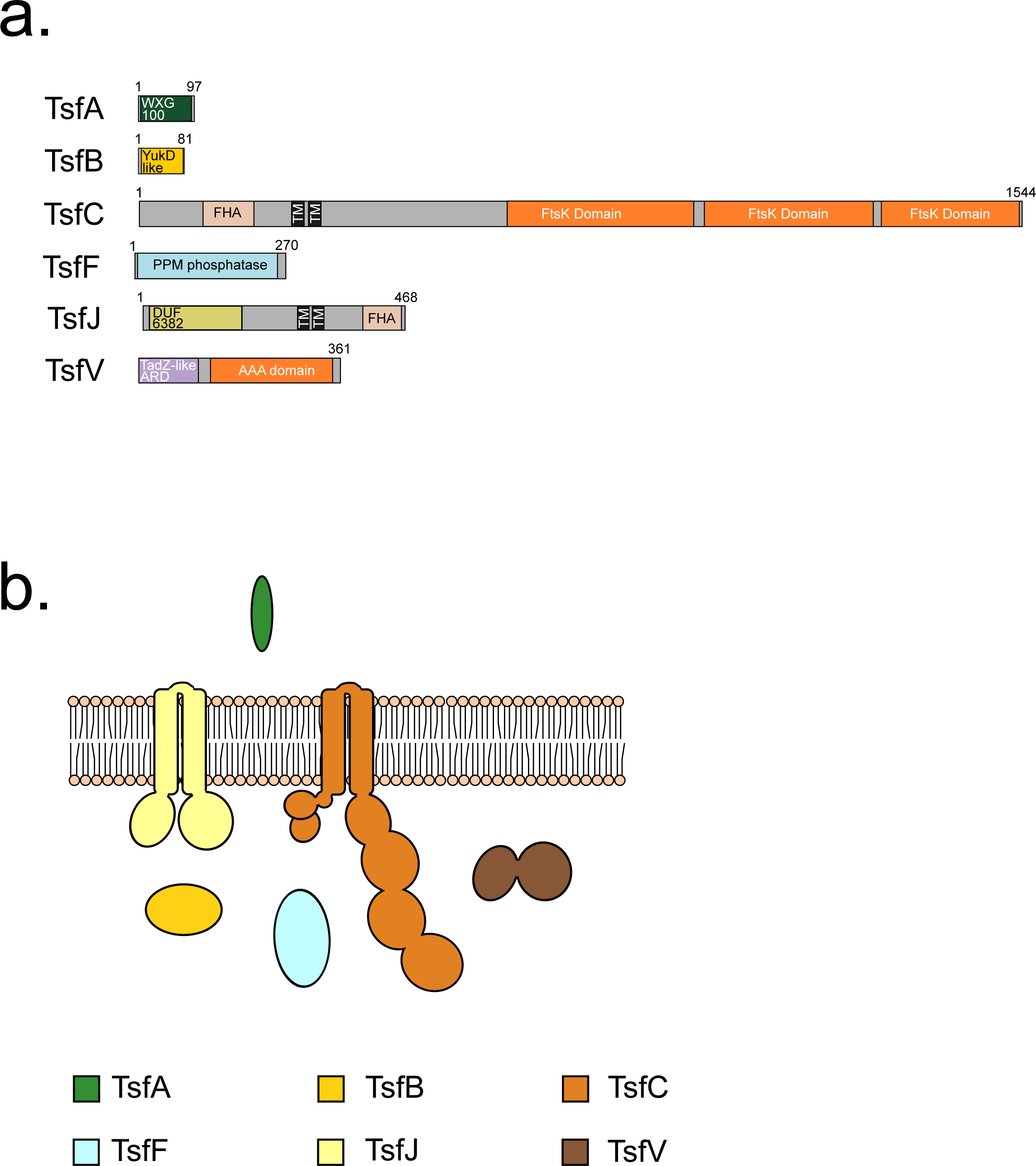
a. Predicted domain arrangement and b. predicted subcellular location of components of the T7SSf.

### T7SSg

The T7SSg system is found in strains of *Acetivibrio* and *Paenibacillus*. The system appears relatively simple - the putative core components of T7SSg are limited to two WXG100 proteins, TsgA1 and TsgA2, the ubiquitin-like TsgB, the membrane bound ATPase, TsgC, and TsgL (Fig 3, Fig. 8a, Supplementary Data 2). TsgC is most phylogenetically related to the EssC homologues from the T7SSj, T7SSf and T7SSb systems, but unlike all of these it lacks N-terminal FHA domains (Fig.2, Fig S2). TsgL is a DUF5050 domain-containing beta propeller protein (Fig. 8a, Supplementary Data 3). A full set of genes coding for a Tad pilus are located immediately adjacent to the T7SSg locus in all organisms (Supplementary Data 1). It is not clear whether these systems are related (for example by sharing a common mechanism of regulation) or whether their co-location is coincidental due to the limited genera in which this system is found.

**Figure 8.**
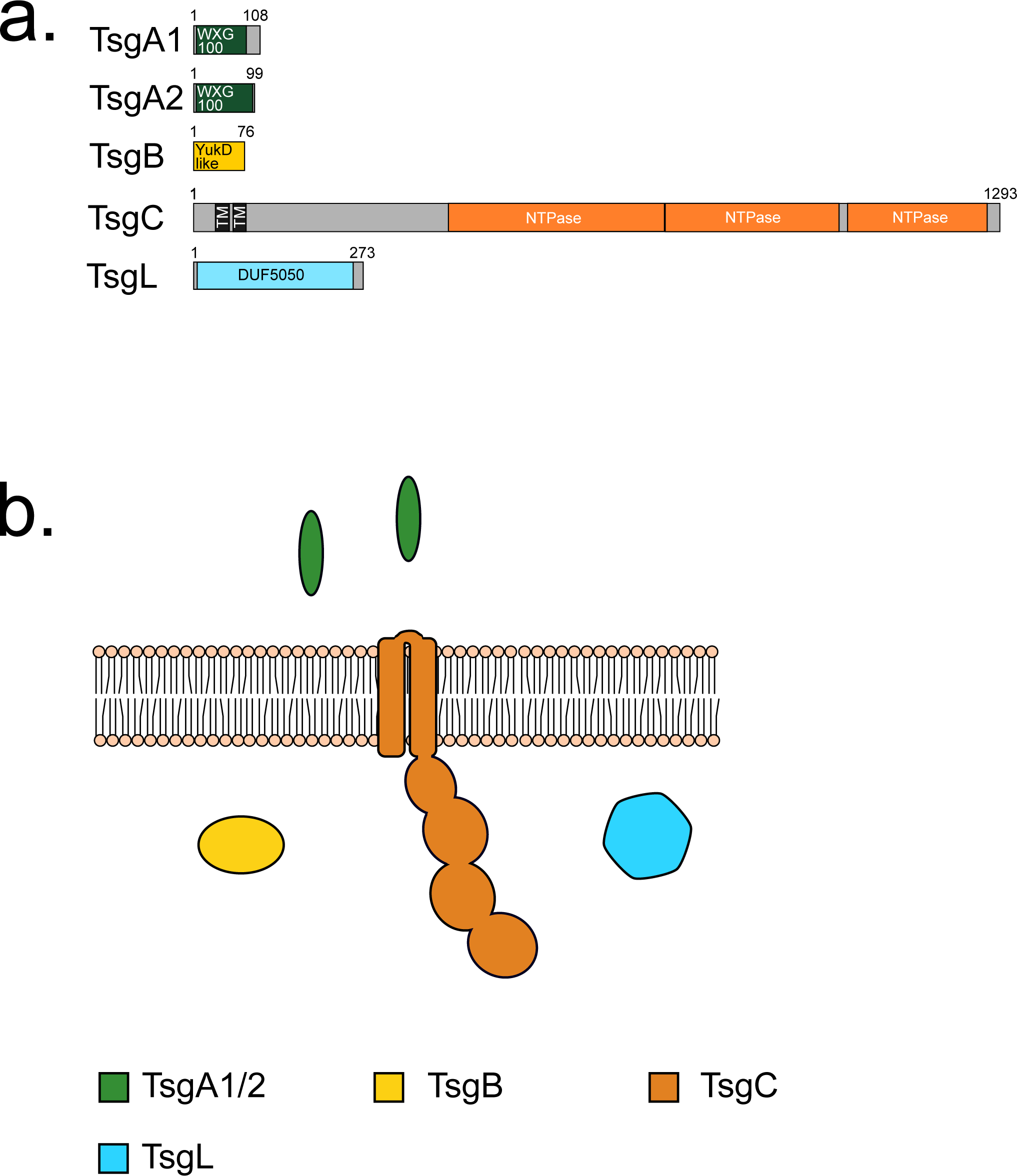
a. Predicted domain arrangement and b. predicted subcellular location of components of the T7SSg.

### T7SSh and T7SSi

T7SSh, which is also encoded next to Tad pilus genes, is comprised of five core components, TshA, TshB, TshC, TshG and TshM (Fig. 9a,b, Supplementary Data 2). TshA is a WXG100 protein, TshB a ubiquitin-related protein and TshC the membrane-bound ATPase that lacks N-terminal FHA domains. TshG, is homologous to the EspG chaperone from the T7SSa system. TshM is not confidently predicted but does share some potential homology with serine/threonine kinases. However, there is no sequence similarity with TscD, TsdD or TseD (not shown). Moreover, TshM is predicted to have a single transmembrane helix at its C-terminus and lacks an extracellular domain found in other kinase components, such as TscD and TsdD (Fig. 9b; Supplementary Data 2).

**Figure 9.**
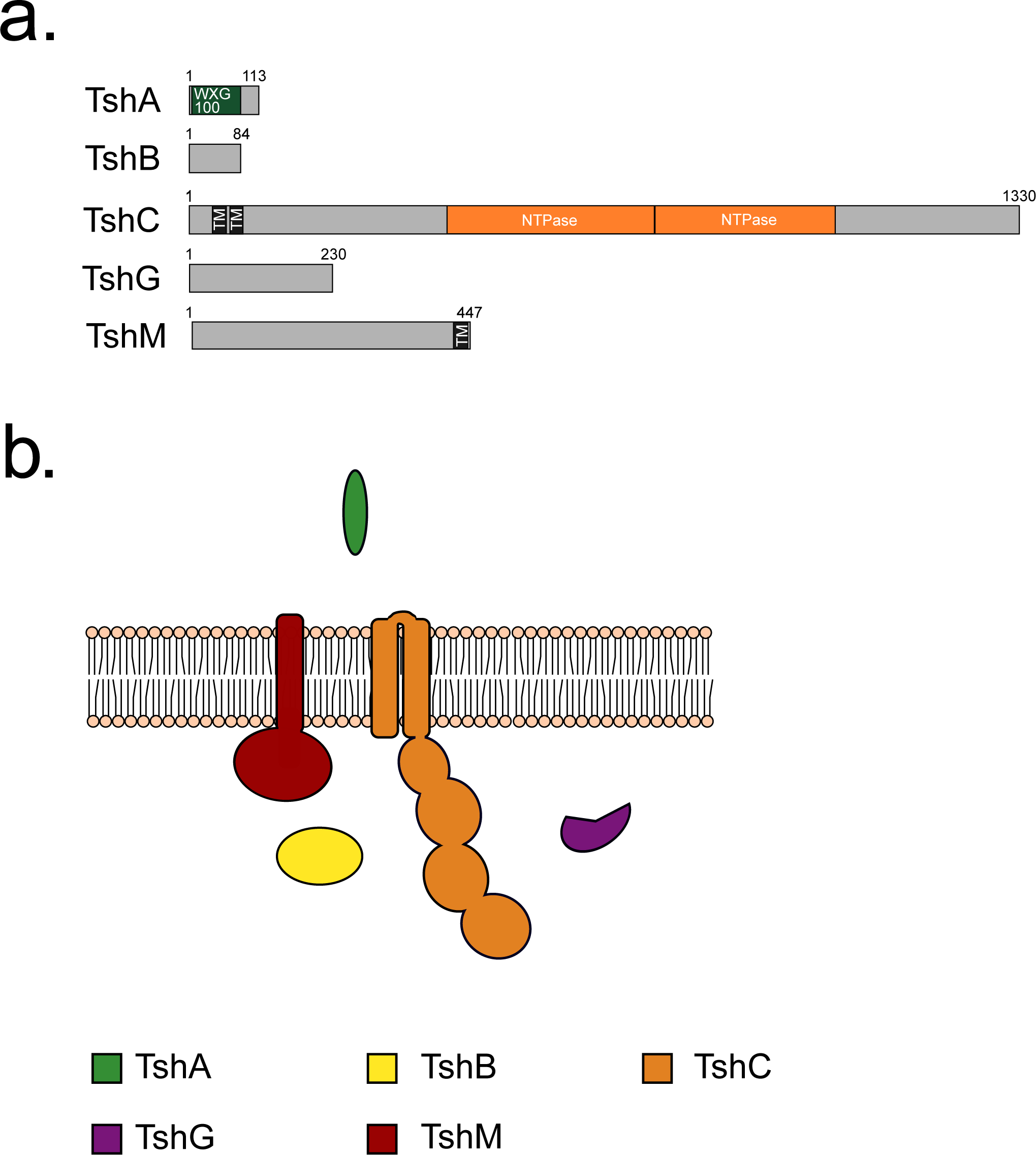
a. Predicted domain arrangement and b. predicted subcellular location of components of the T7SSh.

T7SSi is, to date, only found in *Vallitalea* species. While superficially it appears similar to T7SSh, sharing almost all of the same components, phylogenetically, the core ATPase, TsiC is distinct from that of the T7SSh (Supplementary Data 1). Moreover, TsiC has N-terminal FHA domains, unlike TshC, although they show poor conservation of the phosphothreonine recognition motifs (Fig 2, Fig S2). WXG100 proteins are encoded at the T7SSi locus, but we were not able to pinpoint which, if any, were likely to encode TsxA. This is because in the other T7SS genetic clusters *tsxA* is found in a defined position/order relative to the other core genes, whereas in the T7SSi clusters the WXG100 protein-encoding genes were more variably located, often clustering beside candidate substrate genes (Fig. 3, Fig 10, Supplementary Data 1). TsiN is unique to the T7SSi and is predicted to be a LytR-like transcriptional regulator that may potentially play a role in the regulation of T7SSi genes at the transcriptional level (Fig. 10a,b, Supplementary Data 2).

**Figure 10.**
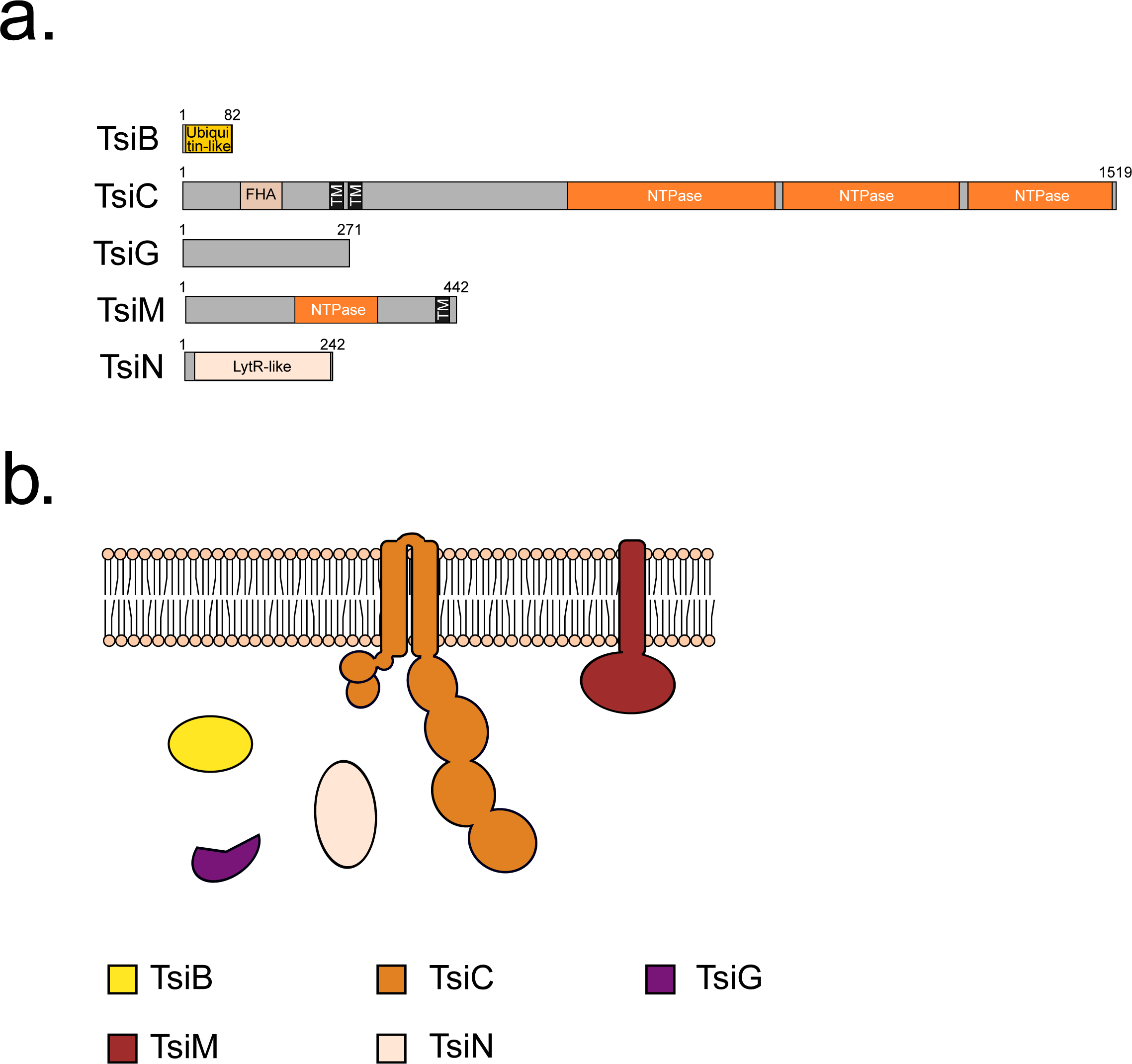
a. Predicted domain arrangement and b. predicted subcellular location of components of the T7SSi.

### T7SSj

The T7SSj was identified in a small subset of *Clostridium* species, and appears to represent a system similar to that of the T7SSd and T7SSe, sharing the homologous components TsjA, TsjB, TsjC, TsjD and TsjI (Fig. 3; 11a,b, Supplementary Data 2). However, the core component, TsjC is phylogenetically distinct from that of either TsdC or TseC, clustering more closely with the homologous component from T7SSf and T7SSb (Fig 2, Supplementary Data 1), and we have therefore opted to classify T7SSj as a unique system. TsjC has N-terminal FHA domains containing the conserved phosphothreonine recognition sequence. The C-terminal domain of TsjD is also distinct from the C-terminal domains of either TsdD or TseD and is predicted to contain a TPR domain (Fig. 11a, Supplementary Data 2). Much like the T7SSe, the T7SSj also contains the TsjF, TsjJ and TsjK components, however, it lacks a homologue of TseH, which is found in both the T7SSd and T7SSe.

**Figure 11.**
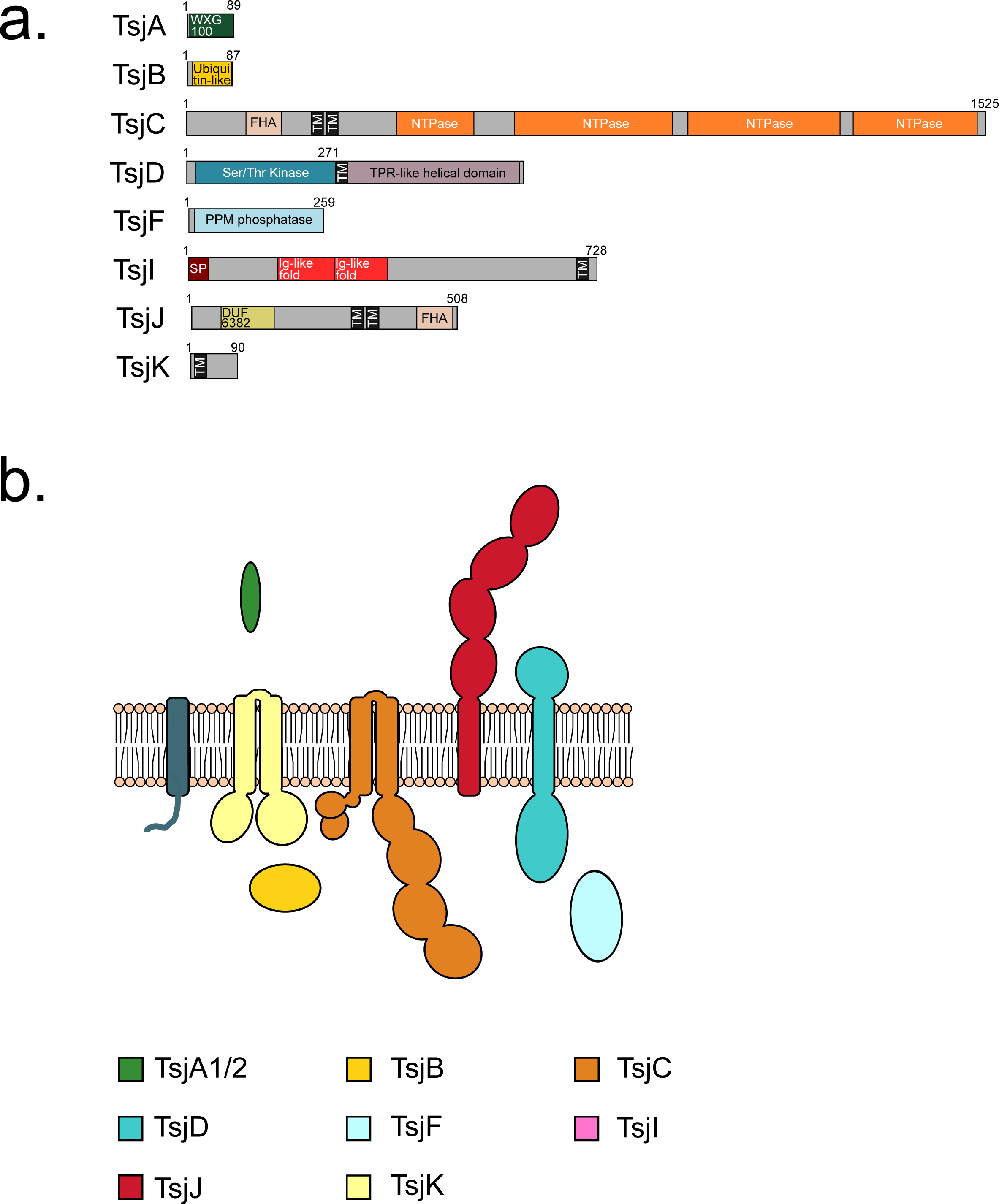
a. Predicted domain arrangement and b. predicted subcellular location of components of the T7SSj.

Candidate T7SS substrate proteins can be identified for all new T7SS variants.

Substrates of the T7SSa and T7SSb systems are often encoded adjacent to the T7SS locus in bacterial genomes (e.g. (Damen *et al*., 2020, Bowran & Palmer, 2021)). We therefore analysed the genetic loci of example strains encoding each of the novel T7SSs to determine whether we could identify candidate substrates. As shown in Fig. 12, likely substrate-encoding genes were present at the T7SS loci of some strains, as well as genes for candidate immunity proteins. For the T7SSc, genes for a predicted HNH endonuclease toxin (WP_080489494.1) and immunity protein are found at the T7SSc locus of *B. anthracis* strain MCCC1A02161, and genes for an Rhs protein also with a potential C-terminal nuclease domain (WP_024632681.1) alongside a candidate immunity protein are found at the same locus in *Paenibacillus* sp. MAEPY1. An Rhs-domain protein with an unknown C-terminal toxin domain and a predicted cytoplasmic immunity protein are also encoded next to the T7SSd in strain *Olsenella uli* DSM 7084. It should be noted that Rhs toxins have been genetically linked with the T7SSb (Bowran *et al*., 2023) and these observations suggest that they are more generally associated with type VII secretion systems. A gene for a smaller toxin like protein, with an N-terminal LXG-like domain is found at the T7SSd locus in *Olsenella* sp. HMSC062G07.

**Figure 12.**
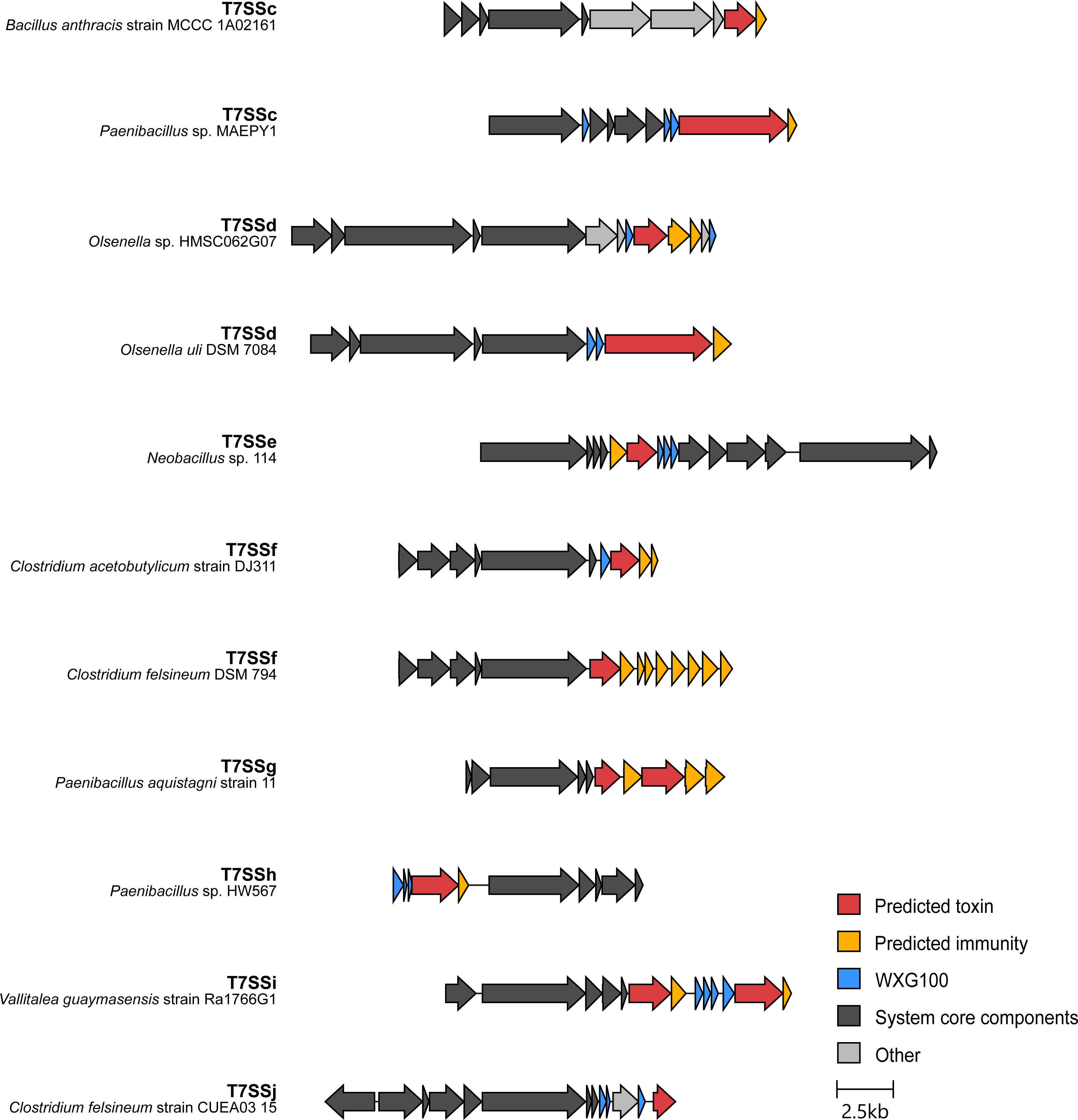
Strains that encode the T7SSc – T7SSj systems also variably encode candidate toxin substrates, immunity proteins and Laps at their T7SS loci. Clinker output showing system-representative loci with genes annotated: core components (dark grey), predicted T7SS-secreted substrate (red), predicted immunity protein (orange), WXG100 family protein/Lap (blue) and unknown (light grey). The loci are centred on the *tsxC* gene of each system. Two arrangements are shown for each of systems c, d and f, to highlight the variation seen is substrate clusters.

Recently it has been reported that the T7SSb can secrete a lipase that has a C-terminal helical targeting domain (Garrett *et al*., 2023). We observed similar ‘reverse’ lipase toxins encoded at the T7SSe gene cluster of *Neobacillus* sp. 114 (WP_284036497.1) and the T7SSg cluster of *Paenibacillus aquistagni* strain 11 (both WP_139829221.1 and WP_085499087.1), suggesting that the capacity to recognise C-terminal targeting domains is also likely to be a general feature of the T7SS. In the two examples of the T7SSf gene clusters shown in Fig 12, multiple candidate immunity genes are present downstream of the predicted substrates (WP_077893465.1 in strain *Clostridium felsineum* DSM 794 and WP_010963371.1 in *Clostridium acetobutylicum* strain DJ311), which has been classically observed for other polymorphic antibacterial toxins of the T7SSb (e.g. (Klein *et al*., 2018, Garrett *et al*., 2022)).

An NHN nuclease toxin (WP_019912159.1) alongside a SMI1/KNR4 family immunity protein is encoded at the T7SSh gene cluster from *Paenibacillus* sp. HW567 and two likely toxins, WP_212693237.1 of unknown function and WP_244971330.1, with a predicted C-terminal CdiA-related endonuclease domain are encoded at the T7SSi locus of *Vallitalea guaymasensis* strain Ra1766G1. A candidate immunity protein is encoded adjacent to each predicted toxin. Finally, an LXG domain containing protein, WP_257675215.1, is encoded at the T7SSj locus of *Clostridium felsineum* strain CUEA03 15.

For almost all of the candidate substrate protein genes identified at the novel T7SS loci we also observed that they were found next to one or two genes encoding small WXG100-related proteins. These were generally distinct from the TsxA/EsxA core component and likely represent Laps that interact with the helical targeting domains of their specific substrate partners as seen for the antibacterial T7SSb toxins (Klein *et al*., 2022, Garrett *et al*., 2023, Yang *et al*., 2023, Klein *et al*., 2024).

## Discussion

Here we describe eight novel arrangements of the T7SS in Gram-positive bacteria, based on phylogenetic and gene neighbourhood analysis, which we have named T7SSc – T7SSj. While the majority of these novel systems are encoded by members of the Bacillota phylum, the T7SSd, like the T7SSa, is found in Actinomycetota. Each of the T7SSc – T7SSj systems contains a homologue of the membrane-bound FtsK-related ATPase EccC/EssC, each with four predicted C-terminal NTPase domains. EssB from the T7SSb differs from EccC in the T7SSa by the presence of FHA domains at its N-terminus. Of the novel systems, the ATPases from the T7SSc, T7SSg and T7SSh also lack FHA domains whereas they are present on the ATPases from other five systems. However, there is no phylogenetic clustering of ATPases based on the presence or absence of FHA domains, and furthermore, the FHA domain sequences have very low sequence conservation between the different systems, probably because they mediate interactions with distinct protein partners.

Alongside the ATPase component, two small globular proteins are also (almost) universally conserved throughout the T7 systems. The first of these is EsxA/TsxA, a helical hairpin protein of the WXG100 family. EsxA is secreted by the T7SS, either as a homodimer by the T7SSb or as a heterodimer with a homologous partner protein, EsxB, in the T7SSa. EsxA secretion is essential for the secretion of other T7SS substrates (Fortune *et al*., 2005, Kneuper *et al*., 2014). However, the function/s of EsxA is unclear and it is unknown whether its activity is required both in the cytoplasm to support secretion, and extracellularly after secretion. A C-terminal sequence is present on the EsxA homodimer and the EsxAB heterodimer that interacts with EssC/EccC ATPase domains, regulating their conformation and activity (Champion *et al*., 2006, Rosenberg *et al*., 2015, Mietrach *et al*., 2020). This indicates a critical role inside the cell during the secretion process, but there is also evidence that at least some EsxA proteins have effector functions following secretion, through formation of pores in target membranes (Spencer *et al*., 2021, Tak *et al*., 2021). Of the eight novel systems identified, most resembled the T7SSb by encoding a single *esxA* homologue at the T7SS gene cluster. The exceptions to this are T7SSe and T7SSg where a non-identical pair of EsxA homologues are found, akin to EsxA and EsxB from the T7SSa system. Surprisingly, no clear *esxA* gene could be identified in the T7SS clusters of T7SSi-encoding strains. Given the essentiality of EsxA proteins it would be unexpected for the T7SSi system to function without one or a pair of core WXG100 proteins. It should be noted, however, that there are very few examples of T7SSi- enocding strains present in the Refseq database and availability of further genome sequences in future may allow us to be more confident in the identification of TsxA.

The second small conserved protein is EsaB/TsxB. The structure of the EsaB homologue from *B. subtilis*, YukB, has been solved and it adopts the same fold as ubiquitin but it lacks the C-terminal Gly-Gly motif that is essential for conjugation of ubiquitin to target lysines (van den Ent & Lowe, 2005). A ubiquitin-related protein, in each case lacking the C-terminal Gly-Gly motif, was identified for all of the novel systems, except the T7SSd. EsaB is essential for the function of the T7SSb, but its precise role is unclear (Casabona *et al*., 2017). Unexpectedly, cryoEM analysis of a protomer of the T7SSa ESX-3 complex revealed the presence of a cytoplasmic domain with the same fold as EsaB at the C-terminus of the polytopic EccD protein (Famelis *et al*., 2019, Poweleit *et al*., 2019). In the assembled ESX-5 system the EccD domain interacts with the first nucleotide binding domain of EccC (Beckham *et al*., 2021), and it is therefore likely that the globular EsaB/TsxB proteins similarly interact with their cognate ATPases.

Across the eight novel systems, predicted kinases and/or phosphatases were often found, and three of the systems (T7SSe, T7SSf and T7SSj) also had a separate FHA-domain containing component. In these latter three systems the phosphothreonine recognition motif on the TsxC ATPase is also well conserved. This raises the possibility that some of the T7SS subtypes may be regulated by phosphorylation/dephosphorylation. Assembly and activity of the Gram-negative type VI secretion system (T6SS) is also regulated by threonine phosphorylation. In the *Agrobacterium tumefaciens* T6SS, phosphorylation of one of the T6SS membrane proteins by a membrane-bound kinase promotes its interaction with an FHA domain protein and activates the secretion system (Lin et al., 2014). Interestingly, kinase-dependent regulation of the T7SSb has also been reported in *Enterococcus faecalis* (Chatterjee et al., 2020; 2021). Membrane damage mediated by incoming phage attack is detected by IreK, a membrane-bound serine–threonine kinase involved in cell envelope homeostasis, resulting in transcriptional activation of the T7SS locus (Kristich et al., 2007; Chatterjee et al., 2021).

EspG chaperones have been well characterised in the T7SSa system, and a related chaperone, EsaE, is found in some (but not all) T7SSb systems. A protein with predicted similarity to EspG was also encoded at the T7SSh and T7SSi loci. Each of the novel systems was also associated with substrate protein families related to those from the T7SSb, including LXG domain and RHS proteins, and lipases with reverse domain arrangement. Most substrates we identified were encoded at loci with genes for small Lap-related partner proteins, pointing to a common mechanism for substrate secretion.

The T7SSg and T7SSh clusters are always encoded adjacent to genes for Tad pilus components. Tad pili are part of the type 4 pili superfamily and have roles in cell adherence, biofilm formation and contact dependent bacterial killing (Kachlany *et al*., 2001; Seef *et al*., 2021). It is not clear whether they are linked with these T7 systems, for example through a common form of regulation or through shared biological functions, or whether their genetic co-location is coincidental. This underlines one of the shortfalls of genetic neighbourhood analysis when there are relatively few genome sequences available within a genus, and it remains possible that there are further T7 components for some of these systems that we have yet to identify. Nonetheless we anticipate that the findings we report here will underpin further investigation into the diversity of type VII secretion.

## Acknowledgements

We thank Kieran Bowran for his help with generating the boxshade figures and Dr Kate Beckham for critical reading of the manuscript. This work was supported by Wellcome through Investigator Awards 110183/A/15/Z and 224151/Z/21/Z TP. SG and AH are funded by the Newcastle-Liverpool-Durham BBSRC DTP2 Training Grant, project reference number BB/M011186/1.

## Notes

### Competing Interest Statement

The authors have declared no competing interest.

