## Supplemental Figures for "Multiple variants of the type VII secretion system in Gram-positive bacteria"

```
S. aureus       MHKLIIKYNKQLKMLNLRD---GKTYTISEDERADITLKSL---GEVIHLEQNNQG----   50
B. subtilis     MSLLWVFYQNNVQKLNLSNLPSSHPVTIGPDVKDSVTISTIPFNSGVISLKRKESGGQYE   60
B. anthracis    ------------------------------------------------------------   0

S. aureus       TWQANHT----SINKVLVRKGDLDDITLQLYTEADYASFAYPSIQDTMTIGPNAYD-DMV   105
B. subtilis     VFLGNDCLGTIETDISFTLQTDQQDIRLILTGSEPEKSVYFTGNRDEIVCSSEKTNADIY   120
B. anthracis    ------------------------------------------------------------   0

S. aureus       IQSLMNAIIIKDFQSIQESQYVRI-------VHDKNTDVYINYELQEQLTNKAYI--GDH   156
B. subtilis     LNP-------QDFAFAEQSTFSLLRAGGSWSVRPESGTIFLN---GEKINANTPLKPGDE   170
B. anthracis    ------------------------------------------------------------   0

S. aureus       IYVEGIWLEVQADGLNVLSQNTVASSLIRLTQEMPHAQADDYNTYHRSPRIIHREPTDDI   216
B. subtilis     IFWNFTQMRVTEQDLLEIVHYAQFETALTETVKPSTEMQKKYPQYRRTPRMVYDLPDDRV   230
B. anthracis    --------------------------------------MQKSPYFQRSPRVKVDIPTGDV   22

S. aureus       KIERPPQPIQKNNTVIWRSIIPPLVMIALTVVIFL----VRPIGIYILMMIGMSTVTI--   270
B. subtilis     SFSFPSQESDQTNRGLWLVILPPLVMLIVMGVVAI----IQPRGIFILVSLAMFMMTL--   284
B. anthracis    IIHDPPNIPEEPKFSIETMLLPAMMTVLTLVLYFVMMKFMKMNSSYMPLMMVSSIPMLGS   82

S. aureus       -VFGITTYFSEKKKYNKDVEKREKDYKAYLDNKSKEINKAIKAQRFSLNYHYPTVAEIKD   329
B. subtilis     -ITSTVQYFRDKNQRKKREEKRERVYKLYLDNKRKELQALAEKQKQVLEFHFPSFEQMKY   343
B. anthracis    YVITTMGHFRKKKEHRQMVEQLQTRYLEQIQKHRVELDTLKVEQAKYLITQNPSPLKSVQ   141

S. aureus       IVETKAPRIYEKTSHHHDFLHYKLGIANVEKSFKLDYQEEEFNQRR-DELFDDAKELYEF   388
B. subtilis     LTSEISDRIWEKSLESKDYLQLRLGTGTVPSSYEINMSGGDLANRDIDDLMEKSQHMQRV   403
B. anthracis    RIENRESNLWERTPESPDFLDIRIGTG--ERPFLVELKVPEQKGYEENPLVTEAQNVKRD   200

S. aureus       YTDVEQAPLINDLNHGP-IAYIGARHLILEELEKMLIQLSTFHSYHDLEFLFVTREDEVE   447
B. subtilis     YKDIRNAPVTVDLAEGP-MGLVGKSQIVKNEIHQLIGQLSFFNSYHDLRFVFIFHEEEYK   462
B. anthracis    FNTIPNGHISISLKKNDVIGVVGNKEDRLNFIRIVTTQIMTHHAPNEVKIAAFYHEKEKK   260

S. aureus       TLKWARWLPHMTLRGQNIRGFVYNQRTRDQILTSIYSMIKERIQAVRERSRSNEQIIFTP   507
B. subtilis     DWEWMKWLPQFQMPHIYAKGFIYNEQTRDQLLSSLYELIRER-----DLEDDKEKLQFKP   517
B. anthracis    QWDWMRWLPHVWDEQRSMRFLSENQQDAQKLAEVLFTPLNMR-RIYNSSAQADAKVPLIP   319

S. aureus       QLVFVITDMSLIIDHVILEYVNQDLSEYGISLIFVEDVIESLPEHVDTIIDIKSRTEGEL   567
B. subtilis     HFVFVITNQQLISEHVILEYLEGQHEHLGISTIVAAETKESLSENITTLVRYINEHEGDI   577
B. anthracis    MYVFFLSAREFLEDDPLTPMLLREGESVGASTFIFAEQKERLPMECDLVISLNGEN-GEL   378

S. aureus       I------TKEKELVQLKFTPENIDNVDKEYIARRLANLIHVEHLKNAIPDSITFLEMYNV   621
B. subtilis     L------IQKKKAVRIPFRLDHHQREDNERFSRTLRTLNHQVGITNSIPETVSFLELFHA   631
B. anthracis    VETFSSSAENSGTTRASFKVDRLSFERCELGARSIAPIRMKSSTAANIPKVLTFLDLFQA   438

S. aureus       KEVDQLDVVNRWRQNETYKTMAVPLGVRGKDDILSLNLHEKA----HGPHGLVAGTTGSG   677
B. subtilis     KEVKEIGIQQRWLTSESSKSLSVPIGYKGKDDIVYLNLHEKA----HGPHGLLAGTTGSG   687
B. anthracis    KNMEELQVLERWKENRYPTSLPVPIGVREGSKPVFLNIHDKIEKKGHGPHGLMAGTTGSG   498

S. aureus       KSEIIQSYILSLAINFHPHEVAFLLIDYKGGGMANLFKDLVHLVGTITNLDGDE--AMRA   735
B. subtilis     KSEFLQTYILSLAVHFHPHEAAFLLIDYKGGGMAQPFRNIPHLLGTITNIEGSKNFSMRA   747
B. anthracis    KSEVIQSIIAALAATYHPHEMAFMLIDYKGGGMSNTFAGLPHIIASITNLE-DPNLIERA   557

S. aureus       LTSIKAELRKRQRLFGEH-DVNHINQYHKLFKEGIATEPMPHLFIISDEFAELKSEQPDF   794
B. subtilis     LASIKSELKKRQRLFDQY-QVNHINDYTKLYKQGKAEVAMPHLFLISDEFAELKSEEPDF   806
B. anthracis    RISLKAELERRQKLFIQAGNVQHLDEYYE--TSWREKEPLPHLFIVIDEFAQMKKEQPEF   615

S. aureus       MKELVSTARIGRSLGIHLILATQKPSGVVDDQIWSNSKFKLALKVQDRQDSNEILKTPDA   854
B. subtilis     IRELVSAARIGRSLGVHLILATQKPGGIIDDQIWSNSRFKVALKVQDATDSKEILKNSDA   866
B. anthracis    MDELISVAAIGRTLGVHLLLATQKPSGVVNDKIWSNSRFRICLRVQDDADSREMLKIPDA   675

S. aureus       ADITLPGRAYLQVGNNEIYELFQSAWSGATYDIEGDKLEVEDKTIYMINDYGQLQAINKD   914
B. subtilis     ANITVTGRGYLQVGNNEVYELFQSAWSGAPYL---EEVYGTEDEIAIVTDTGLIPLSEVD   923
B. anthracis    SKINVPGRGYLQVGSNEVLELFQSAWSGAPYN-------PDEEKVLDIVDFTEVKLSGER   728
```

```
S. aureus     LSGLE-DEETKENQTELEAVIDHIESITTRLEIEEVVKRPWLPPLPENVYQEDLVETDFRK  973
B. subtilis   TE----DNAKKDVQTEIEAVVDEIERIQDEMGIEKLPSPWLPPLAERIPRTLF-------  972
B. anthracis  IKVKKRPKPMTNSPKQLQAFIQYVQSVSEKENIKALPGPWLDPLPEKLLLKEFYAMENWT  788

S. aureus     L--WSDDAKEVELTLGLKDVPEEQYQGPMVLQLKKAGHIALIGSPGYGRTTFLHNIIFDV  1031
B. subtilis   ---PSNEKDHFHFA--YVDEPDLQRQAPIAYKMMEDGNIGIFGSSGYGKSIAAATFLMSF  1027
B. anthracis  IADWNKPKEYLQVTVGLIDDVANQAQYPLKLDLQE-GHLNIYGMPGTGKTTMLQTIIMSL  847

S. aureus     ARHHRPDQAHMYLFDFGTNGLMPVTDIPHVADYFTVDQEDKIAKAIRIFNDEIDRRKKIL  1091
B. subtilis   ADVYTPEELHVYIFDFGNGTLLPLAKLPHTADYFLMDQSRKIEKFMIRIKEEIDRRKRLF  1087
B. anthracis  AVSHTPEEVNFYVIDFGR-MFLDFRDLPHIGGIIQEGENEKMKRLFGFLKQEITNRKESF  906

S. aureus     SQYRVTSISEYRKLTGETIPHVFILIDNFDAVKDSPFQEVFENMMIK-MTREGLALDMQV  1150
B. subtilis   REKEISHIKMYNALSEEELPFIFITIDNFDIVKDEMHE--LESEFVQ-LSRDGQSLGIYF  1144
B. anthracis  SNIGAKSFSMYNRMVEKKVPAIVVMVDGYMRFKNEFEK---ENEVLELLLRESSTYGVYF  963

S. aureus     TLTASRANAMKTPMYINMKTRIAMFLYDKSEVSNVVGQQKFAVKDVV-GRALLSSDDNVS  1209
B. subtilis   MLTATRVNAVRQSLLNNLKTKIVHYLMDQSEGYSIYGRPKFNLEPIP-GRVIIQKEELYF  1203
B. anthracis  YFSLNQTIDMFDRVRNNIPMALTFELQDGTEYHNLVGRPKFPLIEVPVGRGLMKGQPPEL  1023

S. aureus     FHIGQPFKHDETKSYNDQINDEVSAMT-EFYKGETPNDIPMMPDEIKYEDYRESLNLPDI  1268
B. subtilis   AQMFLPVDADDDIGMFNELKSDVQKLQGRFASMEQPAPIPMLPESLSTRELSIRFKLERK  1263
B. anthracis  FQAALPFIGESELEYSQQLKTIIQKMNGEW-NGEKAKTIPMVPKEIFVEHMVGKLETAQI  1082

S. aureus     VANGALPIGLDYEGVTLQKIKLTE--PAMISSENPREIAHIAEIMMKEID----------  1316
B. subtilis   PL--SVPIGLHEETVSPVYFDLGKHKHCLILGQTQRGKTNVLKVMLEHLI-DDETEMIGL  1320
B. anthracis  SA------GIETEDIRLQSFSLDEMSHIFIGGRIEGGKTSLLQTLLFTTTYQYSPEKVEL  1136

S. aureus     --ILNEKYAICIADSSGEFKAYRHQVANFAEERE------DIKAIHQLMI-EDLKQRE-M  1366
B. subtilis   FDSIDRGLSHYAKESDVSYLETKEDIEQWIETAE------DIFKTREAMYVEAVRQGDAQ  1374
B. anthracis  Y-LVDLGERPTGILALGDLPHVKKKVTDAIQLKEMLDELLELINNREAVM-PTLDPNVTV  1194

S. aureus     DGPFEKDSLYI--INDFKTFIDCTYIPEDDVKKLITKGPELGLNILFVGIHKELID-AYD  1423
B. subtilis   NLRFSQVVLMIDGITRFQQTIDTRI--QDRLANFMKSYAHLGFSFIPGGNHSEFSK-GYD  1431
B. anthracis  EFPYKRIMIAIDDIDQMLTVLSTDYEAKNKMELIVQNCKNKGVHFMTAATTSSLNSYSHE  1254

S. aureus     KQIDVARKMINQFSIGIRISDQQFF---KFRFIQREPVIKENEAYMVANQ------AYQK  1474
B. subtilis   SLTTEMKQIRHAILL-MKKSEQNVI---PLPYQRQEPEIQPGFGYVVENG------KEQK  1481
B. anthracis  KWFAEIRKRSIGYLLGTTQNNDVYFFNMKLPHTEMDQELLSGDGYCIRRKPIKIKCAYTP  1314

S. aureus     IRWFK----------------                                         1479
B. subtilis   VQIPLCSAERESAR-------                                         1495
B. anthracis  LHLLRTMIDKISVKWTELKVK                                         1335
```

Figure S1. Sequence alignment of the EssC sequences from *B. subtilis* and *S. aureus* with the protein annotated as EssC from *B. anthracis*. The region boxed in green is the FHA domains, in purple is the two transmembrane domains and in yellow the four NTPase domains.

```
EccC/1-1391 ------------------------------------------------------------
EssC/1-1479 -------MHKLIIKYNKQLKMLNLRDGKT--------------YTISEDERADITLKSLG
TscC/1-1328 ------------------------------------------------------------
TsdC/1-1563 ---MSVDAYTVTMIDAQRLENFALSQ-ES-----LGVSHLAFSESGPGYPCADVHIDGEH
TseC/1-1572 --------MNVTLINKERIHSISLPEKVR--------GQYWIYDSNGKSSRKLIGIEGVN
TsfC/1-1544 MEEKRNYNNVLILYNSDFYKEIDLDDYEKGTVVVGNSANCDIKLKLETEELVYIYFNKIN
TsgC/1-1293 ------------------------------------------------------------
TshC/1-1330 ------------------------------------------------------------
TsiC/1-1522 -----MKEMILYVYYNDTVKEYILSNESS---------------ICNIELSSLLPVVIVN
TsjC/1-1525 ------MNMKLLVYSDKKLYEVKLGNSKT-AVTIGNRKRFKHYIETEEKNRIDMSIKPIK

EccC/1-1391 ------------------------------------------------------------
EssC/1-1479 EVIHLEQNNQGTWQANHTSINKVL--------------------VRKGDLDDITLQLYTE
TscC/1-1328 ------------------------------------------------------------
TsdC/1-1563 MRLCAGCGSRLFDESGIIDSYDMHGVSEGHVLTLRTESTHSSGQSGNPGVASLLFIRPST
TseC/1-1572 GEWILKSNKDVKVLDNTGKPIRNTVL---------QPLSIYNLGLSTQDQKTIIFTEPTT
TsfC/1-1544 TSWQIKDGENAYCIVNDIKTPRKILVNG-------DQITIKNVKDKEELFKINFFIDFIV
TsgC/1-1293 ------------------------------------------------------------
TshC/1-1330 ------------------------------------------------------------
TsiC/1-1522 GEWFLAPTEDMAVSKKQINVGDKI-----------------SVSITDTDEKATIYVDRET
TsjC/1-1525 DIWVLKSKYYIGAEGQEVDIKLEHGV---------------VLQFGLNGKSSIMAIAFSE

EccC/1-1391 ------------------------------------------------------------
EssC/1-1479 ADYASFAYPSIQDT--MTIGPNAYDDMVIQSLMNAIIIKDFQSIQESQYVRIVHDKNTDV
TscC/1-1328 ------------------------------------------------------------
TsdC/1-1563 AGAREFRKIAFSSDADMTIGRDGSSALVYSLPFVSEHHAVLSYRDAE-FSIVDNGSVNGT
TseC/1-1572 DDRQIFTKYLVDRDIEITIGRTEQNDIVLANRFVSASHAKLSFHSGQ-WRIEDQNSSNGT
TsfC/1-1544 EKENYDKVISLKDVTYLSIGRDEDNNISIRDDLIDRKHCEIKCDSNNKFYVTDLKSKYGV
TsgC/1-1293 ------------------------------------------------------------
TshC/1-1330 ------------------------------------------------------------
TsiC/1-1522 NIELTYKKYSIEGEKQILIGNSEKNTIVYNGFDMKPNHAVISIENGR--ATISTKGNKCL
TsjC/1-1525 DKLLSKEKLMLKEKVKFTIGRGKFNDIVFNDIKVSEKHAEIFEDDGK-YVLVDLNSTNKT
                                 GR                SXXH                NG
EccC/1-1391 ------------------------------------------------------------
EssC/1-1479 YINYELQEQLT-NKAYIGDHIYVEGIWLEVQADGLNVLSQNTVA-------------SSL
TscC/1-1328 ------------------------------------------------------------
TsdC/1-1563 FINGVRLTSGRPVHLNAGDLVQIMDLIFTVGLRFISLNNPQGVRLAQSPAWHMVSHESIL
TseC/1-1572 FVDGERVKS---KALKIGDTIYIMGYKMIIGSYFVAFNNPDGTLTIKTHALKPFIPQTV-
TsfC/1-1544 YIDGKKVEDK--VYLKDNNFITICGHKILFMDNSLALSNLNENIVINGA-----------
TsgC/1-1293 ------------------------------------------------------------
TshC/1-1330 ------------------------------------------------------------
TsiC/1-1522 CVNGKMTES---TVLSYGDFIHIMGLKMFYLGKSLAINDSIHLGQCNLSPYTPIKPRE--
TsjC/1-1525 YLNGEMITR---AILARGDEINICGYIIQFNDESIVVQGALKN-----------------

EccC/1-1391 ------------MKRGFARPTPEKPPVIKPENIVLSTPLSIPPPEGKPWWLIVVGVVVVG
EssC/1-1479 IRLTQEMPHAQADDYNTYHRSPRIIHREPTDDIKIERPPQPIQKNNTVIWRSIIPPLVM-
TscC/1-1328 -------------MNVLYQRSPRIKPTLKEDSIEIPKPQNEPSKPSFSLISILLPAVMTI
TsdC/1-1563 AASPPAAQADDVEEAKPFYPAPRLMHTVHVKAFHVDEPPQAMKREEQPALMQLGPSFLMG
TseC/1-1572 -ETPDEEEEYEVPEVDYFYRSPRFKRDVEKAVIKIDSPPPNAIGEEMPMMLVIGPSMTMG
TsfC/1-1544 -TVYNKDRFDKYMSYPDFYRSPRFLVELPRDEIEIDAPPDKPSKVSIQYLLSVIPIVGT-
TsgC/1-1293 ----------MINEKYEFSRQPRFLPEMPRGELEIPNPPSAYDKPEIAWFALLAPPAVM-
TshC/1-1330 -------MDIKSASDTQIQRTPRIMPYYPDGELVLQDPARPLGKPTFSWLTILVPPLAM-
TsiC/1-1522 -DIKGKKKKKTKKEDLYFQRSPRIIKKLEEGMMDIDSPPSPPEQKEQPLILTIGPALTMG
TsjC/1-1525 -TKQKAGQEKVVTEYPFVQRAPRIYPEVKREDLKIENPPTLPNKPSIYGLMTVLPSLIF-

EccC/1-1391 LLGGMVAMVFASGSHVFGGIGSIFPLFMMVGIM---MMMFRGMGGGQQQMSRPKLDAMRA
EssC/1-1479 --IALTVVI----FLVRPIGIYILMMIGMSTV----TIVFGITTYFSEKKKYNKDVEKRE
TscC/1-1328 FSIGFYIYMNL--TGKTGNSSYMMFQMVSVMMMLT-SYTIPFFVYLGNKKKYKEQLKQRV
TsdC/1-1563 IGSVFMAISAVSNLSRGGNVLSAAPSLAMAVAMVGGTLIWPMISKRYTKKVEETKERRRQ
TseC/1-1572 MASMATGLFAVNHAMSTGDISKAIPSIVMSCSMLLGTVLWPILTKKFDMRRRRKKEKIRQ
TsfC/1-1544 --VVMMSLMS---SSMGSNKTYMIYSVGMVSL----SALVSVITFVTEFVSSKKYKKKRN
TsgC/1-1293 --LVISMLL-----AMTSNSRYMLLSISMTIM----TLIVSLTGAASQIKKYKKKKKERE
TshC/1-1330 --LVVAIFM----STLTKSHIMLISTIGMTTV----TVVVSILNYRSTVKKHEEQTKKQE
TsiC/1-1522 MAMMVSMMFTV--YSSRNNPFMVIPGIAMTGSMLLGAVLWPILSRRHKKKSKRKEEKKRI
TsjC/1-1525 --FVVTIIT----SMKSNNGSSTIIFAAGTGA----SALVYGGSYIGQKIVYNKKVKKRN
```

```
EccC/1-1391 Q-FMLMLDMLRETAQESADSMDANYRWFHPAPNTLAAAVG--SPRMWERKPDGKDLNFGV
EssC/1-1479 KDYKAYLDNKSKEINKAIKAQRFSLNYHYPTVAEIKDIVETKAPRIYEKTSHHHD--FLH
TscC/1-1328 RLYHEELDKYREQLIAGHKEQVDVLFEVHGDPDVCFQVVKNRSSVLWERSAHDDD--FLH
TsdC/1-1563 DKYTDYLNSITAKLLEERDRQSSILNENRPPLQNVLQRAYDLSPHLMDRNATQAD--FLQ
TseC/1-1572 EKYKDYLDRIAVVFNEEAEKQEEILRENHVPISDCINRIENVQRNLWERGPGQND--FLK
TsfC/1-1544 KFYRGYIAEKEKEITDNIELQKKNLKIMNPEINECFDFVREFDRRLWEKKPSHPD--FLN
TsgC/1-1293 KKYLQFIADSRSELQLAREQQIKAMNEMNPDPNACLQRIQELDNRLWERTPSSPD--FLS
TshC/1-1330 KKYLDYLFNIRYELLEASNQQREALRVISPSIDECISIVKQRQKQLWERTAIETD--FLS
TsiC/1-1522 KRYRKYMQNEYSRLEEKIAYNKKVLLGTYPEPTVLISRALNRDRRLWERMPSNKD--FLD
TsjC/1-1525 SKYTIYTNEIEKKLLKAEKNLREVLFDENPDLSEAIEIVKNLNMKLWSRSYGDED--FLI

EccC/1-1391 VRVGVGMTRPEVTWGEPQNMPTDIELEPVTGKALQEFGRYQSVVYNLPKMVSLLVEPWYA
EssC/1-1479 YKLGIANVEKSFKLDYQEE-EFNQRRDELFDDAKE-LYEFYTDVEQAPLINDL-NHGPIA
TscC/1-1328 TRIGTGNLPFYIEVKAPR--PEGYVKDSLIEAAQE-LAGEFQTVADAPVALPLFQAKVIG
TsdC/1-1563 LRVGMGEQRVEAQVQAPQR-RFSMENDALLEQAEA-LADKPTMMQ-APVAVDLAGPGTLG
TseC/1-1572 LRVGTGSGLLAADLTYSEK-KFSIDDDNLQGELYT-LVESPKVLKNIPITLSLYENHISG
TsfC/1-1544 VMLGHGDTSISFSVKIPDN-KNKMEIDDLERLAPE-LKKKYQKVEDVPLSLNLLKSSPIG
TsgC/1-1293 LRLGVGSVPAALRINYTKQ-AIIMETDPLLMEPQR-LALEFDRVQGVPVSVDLIASEICG
TshC/1-1330 VRMGNGMQPLALNPTYTSN-TKAVDEDNPLEKIASTICEEMAFVQDVPVQIPLQSISTLG
TsiC/1-1522 IRLGTGTVPSMINIQKPKD-RFTMRDDNLMGELKD-IGNNFDGVSGMPIAFSMMGNNMLG
TsjC/1-1525 LRVGLGNEEFPLKI-IKEK-RNFFDEDTLLDRLDK-ITEKYNIVDNVPICIPIKSNGITA

EccC/1-1391 LVGEREQVLGLMRAIICQLAFSHGPDHVQMIVVSS--DLDQWDWVKWLPH-FGDSRRHDA
EssC/1-1479 YIGARHLILEELEKMLIQLSTFHSYHDLEFLFVTREDEVETLKWARWLPHMTLRGQNIRG
TscC/1-1328 IVGDKEAVMNSLRVMIAQLAVRHSPDEVKLAAFYEEKDAEDWDWMRWLPHTWDEHRSQRY
TsdC/1-1563 VVGARPVVWDFVRGLLAQTCSFYGYNEVKIVAIVDPADQAQWAFLKDVPHARSDDGKMRY
TseC/1-1572 VIGTRKQTIEFAKGLIFQLSALYSYDEVKMIFIYDQDEEAQFSFAKWLPHVWSNDNKFRF
TsfC/1-1544 VVGKKQHTYKFISNLMVKLTAGHFYEDVKIAIIVSNEDKESLSWVRWVPHIWSRKKEIRF
TsgC/1-1293 IAGEEDKTFDMVTQMLLQLITHHGYDDVRVVILASEEGLEKWDWLKHLPHLWSEGYGIRF
TshC/1-1330 LFGKRAEVSEFLNALLVHLTTHHGFDDVKIVGLFDQAEMEMWGWTKWLPHTWDKDREIRF
TsiC/1-1522 MIGNRKTTIDTAMASIVHISALHSYDEVKIVCVYNAEESHKWEWVKQLPHVWAPEKALRF
TsjC/1-1525 VIGNRDKVVETVKNMLLQISINHTPDEVKLVVISDEKENKHWDWAKWLPHVWDDDKNMRF

EccC/1-1391 AGNARMVYTSVREFAAEQAELFAGR-GSFTPRHASSSAQTPTPHTVIIAD-----VDDPQ
EssC/1-1479 FVYNQ---RTRDQILTSIYSMIKERIQAVRERSRSNEQIIFTPQLVFVITDMSLIIDHVI
TscC/1-1328 MSDRR---SSAHQLADELLQRLSRR-KTARHE-QRKKTVEL-PVQVVLLSSGQLIEEEPL
TsdC/1-1563 LASDQ---AELLGLDSMLSKALAER-DGVRGD-AGKACI---PFYVVVCANRELAAASSS
TseC/1-1572 VATND---NEVKEVSAYIEKEIEIR-ANTNES--EMEDVK--PYYIVFALSKELAFKAEM
TsfC/1-1544 IGSDK---ESSHNVLNYLYDVARER-SDDNEN-SHEPKVNL-PHFIVFIADNKLTENEPI
TsgC/1-1293 LLCGK---AIAHAVFGELNTVFKER-----EM-KKLGGIPL-PHYVFIIEESSLLEEEPI
TshC/1-1330 LASNE---YEASILADVLLAEMKRR-DEDKPS-YGTKTIKL-PHYVFLMLAPTLWEDSDL
TsiC/1-1522 VASSR---DEVRDVFLYLKEVLSDR-DETKDNGYDKQVMHL-PHFLVYIADPELVEDELV
TsjC/1-1525 MAKNK---EQAHKLLSNLYDVLSER-KNKLEDKNNYDSYRFSPHFVFVLASRELIQNEAI

EccC/1-1391 WEYVISAEGVDGVTFFDLTGSSMWTDIPERKLQFDKTGVIEALPRDR-------------
EssC/1-1479 LEYVNQDLSEYGISLI------FVEDVIE-SLPEHVDTIIDIKSRTE-------------
TscC/1-1328 LPLLLEDAAEIDACTI------ILSERKE-TLPMQCHLIVDVGKE-K-------------
TsdC/1-1563 VSKLGNLEENKGFCVL------FLGDDLR-DLPRECSQVIELAPPDSSQAAKADYALGTR
TseC/1-1572 LKQVYAKKENLHISIV------TFYDELK-NLPKECSMVVELEEN-S-------------
TsfC/1-1544 MPYLENRN-NVNMTAV------FVSESIE-MLPKECTDVIEVSAGYN-------------
TsgC/1-1293 NKYLYNKSASLGISSV------YIARNQA-YLPMNCKQVILV----Q-------------
TshC/1-1330 MKYIVSNNDSLGITSI------FISERIDVALPLNSQVIIEVKNG-Q-------------
TsiC/1-1522 MKYLTNPNKQLGVSTI------FIYDKLN-QLPKECKAFIQCEET-E-------------
TsjC/1-1525 LKYLLNVDADMEISTI------FLSDKLG-NLPRNCNNIVQLNEG-E-------------

EccC/1-1391 ----DTWMVIDDK-----AWFFALTDQVSIAEAEEFAQKLAQWRLAEAYEEIGQRVAHIG
EssC/1-1479 ----GELITKEKELV---QLKFTPEN-IDNVDKEYIARRLANL-IHVEHLKNAIP----D
TscC/1-1328 ----GHYSLKREDGTFE-QMSFVPDN-LGLDKVDALSRYMAPIRLKRSLAS-DIP----E
TsdC/1-1563 GVLAGPSRMFDRSDVGGSERIFQPDILLSAAGIERFVLALAHARLATGRSEGAMP----E
TseC/1-1572 ----GKLFDK-NDITGK-TTSFTPDIYLNGADPYQLSVKLANVPLDTLANSFNLP----Q
TsfC/1-1544 ----GSSTETINRAH---FMQFAFDD-VNLRECYEYSKRMAPINVKSSFSESSLK----S
TsgC/1-1293 ----GKTGELADRETGE-KTTFNPDT-INLQSLSEAVRKMAPLRIKNSGGNFTLP----T
TshC/1-1330 ----GMTRKDQNNCMGQATYHFLSDS-VSCKRAEHFSRMLAPLTIKETKNNSFIP----P
TsiC/1-1522 ----SSLYNRDKPETG--LIKFNNDS-VIGKDLDSYSRALAGIKIKEIASSASLT----S
TsjC/1-1525 ----GIIYNIQNSSE---KSYVTPDK-MDNKMLMEYSRTMAPLRLEKDSYSNKLP----R
```

```
EccC/1-1391 ARDILSYYGIDDPGNIDFDSLWASRTDTMGRSRLRAPFGNRSDNGELLFLDMKSL-DEGG
EssC/1-1479 SITFLEMYNVKEVDQLDVVNRW---RQNETYKTMAVPLGVRG-KDDILSLNLHEK----A
TscC/1-1328 VLTLFEMLNIKTAADLDVERRW---RNNRYPESLPVPFGVRA-GGKNVNLNIHDKIERQG
TsdC/1-1563 SLGFMEMFQKGNVLQLNIADRW---KEHDSSRSLAAQVGIGA-KGEPVSLDLHEE----A
TseC/1-1572 MFTFLELYGVGKVEHLNALTRW---KDNDPTKSLEAPVGVDT-LGELFKLDLHEK----F
TsfC/1-1544 LITLFDLYNVKRTQDFNVLSNW---GKNKVYETMEVPIGVKA-GDEIEYLNLHEK----Y
TsgC/1-1293 SLTLLDKLQAKKTGEIDLLTRW---NMNKPYQGMSVPIGAKA-GGTLFHLDMHET----G
TshC/1-1330 MVTFLDSFNVKLVEELNITNRW---GENQANRTLAVPLGQAS-AGKTLLFDMHEK----N
TsiC/1-1522 MLTFLEMYKIGRIENLSIKSRW---KNNLSYRTLEAPLGIKA-GGTQFLLNIHEK----Y
TsjC/1-1525 SISLFEELDITNVEEIDFNSIW---SQAEAHKTLSVPVGIRE-NGEKFYLDLHQK----Y

EccC/1-1391 DGPHGVMSGTTGSGKSTLVRTVIESLMLSHPPEELQFVLADLKGGSAVKPFAG----VPH
EssC/1-1479 HGPHGLVAGTTGSGKSEIIQSYILSLAINFHPHEVAFLLIDYKGGGMANLFKD----LVH
TscC/1-1328 HGPHGLIAGTTGSGKSEVIQSIVASLAAEFHPHDLAFMLIDYKGGGMSNTFVH----LPH
TsdC/1-1563 HGPHGLIAGTTGSGKSEFIITWVLSMALNYSPDEVAFVLIDYKGGGLAGAFDNARLRLPH
TseC/1-1572 HGPHGLVAGMTGSGKSEFIISYILSLAVNYHPHEVAFILIDYKGGGMAKSFEK----LPH
TsfC/1-1544 HGPHGLVAGTTGSGKSEILQTYIISLAINYHPYDVALIIIDYKGGGMANLFKN----LPH
TsgC/1-1293 HGPHGLVAGTTGSGKSELLQSIIISLAINYHPHDVVFVLIDYKGGGMADVFQG----MPH
TshC/1-1330 YGPHGLVAGTTGSGKSELLQSLLLALAVNYHPHEIAFVLIDYKGGGMANAFAG----LPH
TsiC/1-1522 HGPHGLIAGMTGSGKSEFIQSYILSMAVNYHPHDVAFILIDYKGGGMANCFIG----LPH
TsjC/1-1525 HGPHGLVAGTTGAGKSELLETVIAALSFSYSPEYVNFLLIDYKGGSMANVFKN----LPH

EccC/1-1391 VSRIITDLEEDQALMERFLDALWGEIARRKAICDSA----GVDDAK--EYNSVRARMR--
EssC/1-1479 LVGTITNLDGDE--AMRALTSIKAELRKRQRLFGEH----DVNHIN--QYH----KLF--
TscC/1-1328 VVATITNLSSSL--MERAKVSLKAELVRRQKILNDAG---NLQHID--EYY----KLL--
TsdC/1-1563 LAGTITNLDGAA--VARSMVSIKSELKRRQALFNKACEATGEATMDIGKYI----SYF--
TseC/1-1572 TAGIITNLDGAA--IKRSLVSIESELKRRQAIFADASKLVGESNIDIYKYQ----KLY--
TsfC/1-1544 LVGTITNLDGNQ--INRSLVSIKSELKRRQRIFAKC----NVNHID--AYI----KLY--
TsgC/1-1293 LVGTITNLGGNQ--TTRALLSIKSELMRRQRLFSEF----GVNNID--KYQ----KLYYS
TshC/1-1330 LVGTITNLGGNQ--INRALASIKSELLRRQRLFGEA----GVTSID--DYI----VLY--
TsiC/1-1522 ITGTITNLGGNQ--IRRSLVSLQSELKRRQRVFAEH----GVNHID--KYQ----QLY--
TsjC/1-1525 AVGTVTNLEGNG--SKRAIIAIDSEIKRREKILTDN----TYSNID--EYQ----KNY--

EccC/1-1391 -ARGQDMAPLPMLVVVIDEFYEWFRIMPTAVDVLDSIGRQGRAYWIHLMMASQTIESRAE
EssC/1-1479 -KEGIATEPMPHLFIISDEFAELKSEQPDFMKELVSTARIGRSLGIHLILATQKPSGVVD
TscC/1-1328 RREGG--QPLPHLVIIIDEFAQLKKDQPEFMDELISIAAIGRTLGVHLILATQKPAGVVD
TsdC/1-1563 -RQGVLSEPCPHLFVVADEFAELKQQEPTFLDELVSAARIGRSLGVHLVLATQKPTGVVS
TseC/1-1572 -REGVVREPLQHLFIISDEFAELKTQQPEFMTQLVSAARIGRSLGVHLILATQKPSGVVD
TsfC/1-1544 -KEKKVTEPIPHLIIIADEFAELKSDQPEFMAELVSTARIGRSLGVHLILATQKPAGVVD
TsgC/1-1293 RNAGGHMPAIPHLIMIADEFAELKQDQPDFMKELVSAARVGRSLGVHLILATQKPAGVVD
TshC/1-1330 -REHKVQLPLPHLIIVVDEFAELKSDQPEFMKELVSAARVGRSLGIHLILATQKPSGVVD
TsiC/1-1522 -KEGKADCPLPHLVIISDEFAELKAGQPEFMQELVSAARIGRSLGVHLILATQKPSGVVD
TsjC/1-1525 -KFGKHKMSLPHLIIIVDEFAQLKKNDPDFISQLVNVAVVGRSLGVHLILATQKPSGIVD

EccC/1-1391 -KLMENMGYRLVLKARTAGAAQAA-GVPNAVNLPAQAGLGYFRKSLEDIIR-FQAEFLWR
EssC/1-1479 DQIWSNSKFKLALKVQDRQDSNEILKTPDAADITLP-GRAYLQVGNNEIYELFQSAWSGA
TscC/1-1328 DKIWSNSRFRICLRVQDEGDSRDMLKIPNAAWINKP-GRGYFQVGSDELFEEVQFAWSGA
TsdC/1-1563 DQIASNARFRVCLKVADAADSREMIRRTDAAALTRP-GEFYLLVGYDELFTGGQAAYAGG
TseC/1-1572 DQIWSNSKFRVSLKVQERADSMDMLKRPDAAELTDT-GRFYLQVGYNELFEMGQSAWAGA
TsfC/1-1544 NQIWSNSKFKLCLKVQDAEDSKEVLKSSLAADIVEP-GRAYFQVGNNEIFELFQSAWSGA
TsgC/1-1293 DQIWSNSKFKICLKVQDERDSRDVIKRPDAAMIKEP-GRAFIQVGNDEIFELFQSTYSGA
TshC/1-1330 DQIWSNSRFKLCLKVQTPSDSQEMLKRPDAAEIKEK-GRGYLQVGNNEVFTLFQSSWSGA
TsiC/1-1522 DQIWSNTRFRICLKVLDKSDSNEMLKRPEAANIKEP-GRCYVQVGNNEIFELLQSGWSGG
TsjC/1-1525 PQIETNTNLKICLRVQDNEDSRTVIGKPDASSIANP-GRAYIKVGNDLVYELIQTAFAGE

EccC/1-1391 DYFQPGVSIDGEEAPALV--HSIDYIRPQLFTNSFTPLEVSVGGPDIEPVVAQPNGEVLE
EssC/1-1479 TYDIEGDKLEVE--DKTI--YMIND-----------------------------YGQLQA
TscC/1-1328 PYVEQTDS------PGRIVPTEIAL-----------------------------NGKREN
TsdC/1-1563 TYTERDVYEPKR--DVSV--ELVGL-----------------------------DGGAVS
TseC/1-1572 PYYPSDKVVVEK--DTSV--VVIDR-----------------------------NGRPIK
TsfC/1-1544 KKYDEDDVNKN---EVEI--FNVGI-----------------------------DGTRSL
TsgC/1-1293 DYDPDGELQKSENKTKRI--YEVSL-----------------------------NGRIEQ
TshC/1-1330 PYNNTDSEEEV---AIKA--NSLSI-----------------------------NGERKR
TsiC/1-1522 SYIPTDKIENDE--DNQI--SLIDG-----------------------------CGRAIQ
TsjC/1-1525 KYRKTAAKK-----DKKI--LMVDT-----------------------------DGERFD
```

```
EccC/1-1391 SDDIEGGEDEDEEGVRTPKVGTVIIDQLRKIKF-----EPYRI-WQPPLTQPVAIDDLVN
EssC/1-1479 INKD-LSGLEDEETKENQTELEAVIDHIESITTRLEI-EEVKRPWLPPLPENVYQEDLVE
TscC/1-1328 LLTT-SMSAANLQMEQPPKQLQVFIDKLAEAAKEEGI-PRLQGPWLPPLPDRLELEVLP-
TsdC/1-1563 RLRP-PSAGRSGGIS----ELNAVLEQVCEVARAVN--KSARPLWLEPLPERVLLDDLRQ
TseC/1-1572 QAKIDKKKGLFTNPKK---QLDVITDYLSNIAAEENI--KIRPLWLEPIPALILLADIRK
TsfC/1-1544 LYST-KDKEKDKKRIT---QLEAAIEEIDKVCSENHI-ERLDGPWLPPLEDEIYLDDLLK
TsgC/1-1293 IYPRYEEKIVKHELPS---QLGATVEHIVRTAERSNI-EPLKGPWLPPLPELVYIDGILE
TshC/1-1330 LLPK-VSNHNESKRLN---QLESVVKYINRVSEEQNIREAFQL-WLPPLPELLSLQELLV
TsiC/1-1522 TVSC-KPKPVKSSTT----QLTAIVDYISDLAKDEGI-NPLKL-WLDPLEEEVFIQDIED
TsjC/1-1525 IVER-EKEGHSEDVT----QLSKLIDSI-KVYCDLNMNYERKIPWLPPLKSHIYIDKLS-

EccC/1-1391 RFL----GRPWHKEYGS---------------------------------ACNLVFPIGI
EssC/1-1479 TDF----RKLWSDDAKE------------------------------------VELTLGL
TscC/1-1328 -------GIHIHPAAEE------------------------------------LEGLVGI
TsdC/1-1563 KFD-------FRADHAG------------------------------------LTAVVGE
TseC/1-1572 KYN--ADRGTWLSSENRPDVFQEAVSEIAAAAETVQEDVLKSFTTASRRASLKLNPVIGE
TsfC/1-1544 DSKTGFNGKEWIEDREW------------------------------------ICPTIGM
TsgC/1-1293 NKQ---ELGHWAKQENL------------------------------------LKVPAGI
TshC/1-1330 NEE-GWNGQQWTEPGDY------------------------------------LVAPVGL
TsiC/1-1522 RKN-GWNGDDWQEVDHW------------------------------------LCPTIGL
TsjC/1-1525 ----------YIPQDK-------------------------------------FKARIGV

EccC/1-1391 IDRPYKHDQPPWTVDTSGPGANVLILGAGGSGKTTALQTLICSAALTHTPQQVQFYCLAY
EssC/1-1479 KDVPEEQYQGPMVLQLKKAG-HIALIGSPGYGRTTFLHNIIFDVARHHRPDQAHMYLFDF
TscC/1-1328 IDDLPNQSQRPLHIPLQQ-G-HWAVYGMPGLGKTTFVQTLLMSLALRYPPQAWYGYIVDM
TsdC/1-1563 LDDPEHQRQDVFSVDVAEAG-NVALYGTPTSGVESLAMSMVCSLMAEYTPDELNVYVWDM
TseC/1-1572 YDDPVHQQQCLLRLPLTDEG-NTIIYGAAGTGKTTFLNTMVYSLIHEHTPEEVNLYLLDL
TsfC/1-1544 LDDPERQDQRPLKIDLGELG-HLLLVGAPGYGKTTFLQTLMISLMLNYTPEEVNMYILDF
TsgC/1-1293 MDDPRSQSQDLLEIDFANEG-NLFVYGASGTGKTEFLKTLCMSMAHHYSAEDVNIYIMDF
TshC/1-1330 IDNPSQQAQYQLKIDFGRDG-HLLIYGAPSSGKTTMLKTILMSLALKYNPDYVHFYILDF
TsiC/1-1522 IDDPANQNQFPLKLDIGNMG-HVLLVGAPGTGKTTFLQTMIYSLVTSYTPELVNLYLLDF
TsjC/1-1525 YDDPYHQSQEIFEIDIQKEN-HIALYGMAGTGKTTFLQTFILSLASKNSPRDINFYIADC

EccC/1-1391 SSTALTTVSRIPHVGEVAGPTDPYGVRRTVAELLALVRERKRSFLECGIASMEMFRRRKF
EssC/1-1479 GTNGLMPVTDIPHVADYFTVDQEDKIAKAIRIFNDEIDRRKKILSQYRVTSISEYRKLT-
TscC/1-1328 G-RMMRDFAGLPHIGGVMMAEEDDRIKRLFRFLMKTVSVRKEMFAETGVKTAAAYRRSQ-
TsdC/1-1563 GAGSLSALSKAPHVGGVVLVEDSERVDNLFLLLEQEIAHRRRLFSRSG-GSYESYNGLP-
TseC/1-1572 ASETLRAFSKAPHVGDVILSYESEKVSNLFKMLQGEVEKRKKLLADYG-GDHRSYVEAT-
TsfC/1-1544 GARTLKMYEKSAYVGGVVTSDDEEKLMNLIKYLHKEIDRRKKIFSDNGVGSLKAYREVG-
TsgC/1-1293 GGNSLRIFEKLPHVGGVMTIEEKTKIDQFIMFLFREIEERKVLFEQSGSNGFNAYRHS--
TshC/1-1330 GTRTLGVFNDIPHLGDVIYPEDEQKVDKLLQLLLTELEDRKRKFSKLGISNLVAYRSYT-
TsiC/1-1522 GGRTMGYYKYLPHTGGVIFSEESDKLDKLFKMLKKELDNRKKKFSEYGVGTLKAYMEAT-
TsjC/1-1525 DKGTLNMFKSLVHTGEVVLSDDTDKVKKLLKFIMKEIDKRKNALTSMRAISVNDYNYKT-

EccC/1-1391 GGEAGPVPDDGFGDVYLVIDNYRALAE--ENEVLIEQVNVIINQGPSFGVHVVVTADRES
EssC/1-1479 ---GETIP-----HVFILIDNFDAVKDSPFQEVFENMMIKMTREGLALDMQVTLTASRAN
TscC/1-1328 ---AGPLP-----EMLIVIDGYLNFRN--SYPDENEMLEFILREGGNLGITFVITSNRIS
TsdC/1-1563 AEDRDPVP-----RIVVVLADIAAFNE--GYEKYVDRLNTLARDAPRYGIHLIITASLWS
TseC/1-1572 ---GQKVP-----AIVVAIHNFSAFTE--IYDEKEEAVSYLSREGTKYGIYFVLTALGTG
TsfC/1-1544 ---NTLIP-----QIVIILDNYSALIE--FYQDLEDELIFLSREGGTLGISLVVTAGNYT
TsgC/1-1293 ---GRKLT-----SIVLMIDNYFALSE--SYEDLDAHMMLLAREGTKYGIYLVATASNAT
TshC/1-1330 ---KEEIP-----SIVVALDNYTGFAE--SYVEQVLELGKLVREGGNYGIYFVFTGNAIN
TsiC/1-1522 ---NAKIP-----AMIVILDNYEAFYE--LYSDYEGILTTFSREGGNYGIYMVITASNAN
TsjC/1-1525 ---GKVLP-----QIIFIIDDIVTLLT--VNDDFKEDIVKIVREGGALGVHMVYTANSSN

EccC/1-1391 ELRPPVRSGFGSRIELRLAAVEDAKLVRSRFAKDVPVKP-GRGMVAVNYVRLDSDPQAGL
EssC/1-1479 AMKTPMYINMKTRIAMFLYDKSEVSNVVGQQKFAVKDVV-GRALLSSD-DNVSFH-----
TscC/1-1328 DIFEKVRSNIASAVTFELADPSDYYYAVGRPARNPGSMPPGRGLVKGNVPPLEFQ-----
TsdC/1-1563 QVNMRLRASMGRTLVTSFSNNDDYNSVLNGMHGVTPPKGFLAGVFQNGKGVTVYQ-----
TseC/1-1572 AVRFRLLQNFKQLITLQLNDEADYATIVGKTDGLFPSKFKGRGLIKRD-AIYEFQ-----
TsfC/1-1544 SVRYKVTANFKLSIALTCVDKGEYSNVTGRV-RMEPENKKGRALINVD-NLYEFQ-----
TsgC/1-1293 FVRYKFSVNFKLAVTFQMTDKGDYAGVVGRTEGLEPMKVPGRGLVRAK-PPLEFQ-----
TshC/1-1330 TFPYRIAQNIKQSIVFQMVDSSDYSSIVGRTEGLEPAKVVGRGLVKDQ-MPLEFQ-----
TsiC/1-1522 SVKFKITQSIKLMVALQLNDKYDYVSIVGQTGGLEPEQVKGRGLFRIE-TPLEFQ-----
TsjC/1-1525 SVHMKIKENVAFNIAYNLNDTSEYREIFGRNNGIVPDKISGRGVLRNV-NLLEFQ-----
```

```
EccC/1-1391 HTLVARPALGSTPDNVFECDSVVAAVSRLTSA----QAPPVRRLPARFGVEQVRELASRD
EssC/1-1479 ---IGQPFKHDETKS--YNDQINDEVSAMTEFYKGETPNDIPMMPDEIKYEDYRESLNL-
TscC/1-1328 ---AALPAAGNDEAE--RSLALRSLIQGITVRYPDVHAPLIKKLPERIMLPELMPHAESY
TsdC/1-1563 ---GASLC--ADPSG--EVQAVRDFCGSLAAGWGGTCAKQIPVLPRHVVPSMFNTGEGYD
TseC/1-1572 ---VAQI---TAEHV--PYPSIQAACQKLQEAWKGTAAKRVPILPEQVDHEFLAEYAGPK
TsfC/1-1544 ---TALPVRGENEGA--RAANVKELIEELNSAWKGKHAKPIPIVPDILSLEEAVEELEGN
TsgC/1-1293 ---ASLP---EYEGL--TCEQLLDRFAGLETF----QAKPIPIMPSSIDIGQINKG----
TshC/1-1330 ---GATPVEGESDDQ--IAEGIRVLSGQMKDKWKKKQVKTIAIIPDELSIKEFIENVKSH
TsiC/1-1522 ---TALACKSVNEAE--RVQKLRDLFMKMDESWIGNRAKPIPFVPEKLTIDHLLEHEDSK
TsjC/1-1525 ---TALPVEAE-EFN--WSNEVSDKISSINYSYPNIKVKGIPVLTEVLPLYDFIHGNEFL

EccC/1-1391 TR---QGVG-AGGIAWAISELDLAPVYLNFAENSHLMVTGRRECGRTTTL-ATIMSEIGR
EssC/1-1479 -----PDIVANGALPIGLDYEGVTLQKIKLTEPAMISSENPREIAHIAEIMMKEIDILNE
TscC/1-1328 RPE--RGLNAAFKVPVGLFTDDLEPFELDLREGPHFIVASPMEGGKTSFL-LTWMLSLAY
TsdC/1-1563 ----------ARSIPVGYAKEGAFPVEFDGSRNAMIVSSDDGDAL------YGWLEGIRE
TseC/1-1572 G---------SLTIPIGVEKNSLNVHYYPFGKSYLNMILSAGTEH------LSFVHDLSE
TsfC/1-1544 -----EELE-KYSYTIGIDYTEIEYVYGNLIENPLITVVGKSQTGKSNTI-QSIAHTLAV
TsgC/1-1293 ----------AGKLAIGLANNDLQPVYVDLYTTPIIMVAGEAMSGKSTLL-RSWINVLDE
TshC/1-1330 AGG--QGEQ-KYAFPVGLDWDSTLPFAMDVSSTNNLLLSYAPSVNAELLF-ASLLHSMSA
TsiC/1-1522 -----EFID-NNLLPIGYDTDEAEIVAVDLTGLSSYLILGYEMTGKTNLI-KSLMNTIVD
TsjC/1-1525 DYGLEEEFVYNSRIPVGVNIDEMESVFANFLSIDNMLISGESGCGKTNFL-LSFIMTIAE

EccC/1-1391 LYAPGASSAPPPAPGRPSAQVWLVDPRR---QLLTALGSDYVE--RFAYNLDGVVAMMGE
EssC/1-1479 KY-----------------AICIADSS----GEFKA-YRHQVA--NFAEEREDIKAIHQL
TscC/1-1328 HY------------SPDHLHMYTIDSRYGSRGLAELRELPHVQ--GSASKEEELALLIGR
TsdC/1-1563 VL------------LVQKRNYVVLDAT----GIFSGVEDSRVL--G---QNDEIAEWIEG
TseC/1-1572 FM-----------AQQAGVDVTLIDAEQ---SFIQKNNAGF----QYYSTVKEFEELTGD
TsfC/1-1544 NS------------GVNESKIYLIDSENY--GLLKIKDSNLIK--AYGYKTDESRNIMLE
TsgC/1-1293 VQ------------------VYALDSNGM--GLFEIMSLPYVT--DLAEAEESFIDEMKE
TshC/1-1330 FL------------DSERVKINVFDSS----TVLKSSGNINLEHVHYMRTEGEFSLLTHS
TsiC/1-1522 KL---------------DWKVFVVEKGK---NLQKASSRYQVD--GYINTTEDFDSFIGG
TsjC/1-1525 TK------------NRENNKIYIIDSPER--NMIATSKLNCID--GYLENAEKVADSLAF

EccC/1-1391 LAAALAGR-----------EPPPGLSAEELLSRSWWSGPEIFLIVDDIQQLPPGFDSP--
EssC/1-1479 MIEDLKQR---------EMDGPFE--------------KDSLYIINDFKTFIDCTYIP--
TscC/1-1328 VYDQVQQRGSGEGN------------------------PAILLAIDDADILAKQLNDFM-
TsdC/1-1563 M---LSGRL-----------------------------HADVIVIPSVVQLMVGLTGSV-
TseC/1-1572 LFDLVVYRNNSFKE---ALEEGRE-------VEHF---KQKVIIINSVVALKNALSQLG-
TsfC/1-1544 IKNEIELRKEALNNA--RMTSTGVFNEKEF-LDNM---PLIALFIDDLNDFMLQFSSDVE
TsgC/1-1293 LLDERRSQ---------MAECRRGGGDIKQLMRSW---QQIVFVFDKLSEFTDGDNYAL-
TshC/1-1330 VIEQLQIRKNDSREYVANMDDPDQFNESDYILSRY---PLFIICLPDIKASIERMDSDS-
TsiC/1-1522 LVEEMKLRFKDLKQF--REEDEQVVKEHEY-MKKY---QRIVVLVDDYNEFFEMISDDS-
TsjC/1-1525 IKQEIGERKSILRNL--RMEYGRDANKKI--ISDK---GNIFLIIDVIENFKIRFSDEI-

EccC/1-1391 -LHKAVPFVNRAADV-GLHVIVTRTFGGWSSAGSDPML---RALHQANAPLLVMDADPDE
EssC/1-1479 -EDDVKKLITKGPEL-GLNILFVGIHKELIDAYDKQIDVARKMINQFSIGIRISDQQFFK
TscC/1-1328 IKEQLGTIVRLGRDR-NVHVLLSGVPAD-FPTFGVDWF---NDVKACQSGFLFGTLDPND
TsdC/1-1563 -SDGFKAFLESERYK-GAAVMILASESWRLRGVFDSWF---KAVTANPNGLWIGDGFTSE
TseC/1-1572 -TEKLGLVLEKGDAKYNVTIIFAEQSKNVSSVTFDKWF---KQHIHSGNGIWIGSGITEQ
TsfC/1-1544 IMQIFEGIVQKDKNL-KVALIVAGTEEN-INNYAYSYDFV-KELKSLNYGFFFDN--INN
TsgC/1-1293 -KELLERIVKQERGM-KVAVIAADA----ISDFNGNWDSLGKAIKEEQTGILLGS--IKE
TshC/1-1330 -LQHMERIARFGGGL-GVHLLIGGLAEEIDQLQLVTSLV--PLLAEGESKMLVGGKPFEH
TsiC/1-1522 -SRLVENIVRSGMGL-EVCFVFTADTDTLSPHIGSPLY---NSVFKGMNGILLGGKMDSQ
TsjC/1-1525 -AQELEWIIANCKST-GIHVIVA---DN-MTLFTRAWDGMEKAIKSWETGIIFST--SVD

EccC/1-1391 -GFIRGKMKG------------GPLPRGRGLLMAEDTGVFVQVAATEVRR----------
EssC/1-1479 FRFIQRE---------------PVIKENEAYMVANQAYQKIRWFK---------------
TscC/1-1328 LSFF----RIPFSESGNASGGLKLLPPGQGYYI-KRKFTRVK-AAMPFDAEWSGNRWVTE
TsdC/1-1563 TVFPHTNVAEPFK---------RPMGPHDGVMHGTGRNTLMRLIEPDGKQEQGENGGKGR
TseC/1-1572 FQLKPAKTTSEMR---------EEMPADFAYSLQNGKAVKVKLLNSQKENDDDDE-----
TsfC/1-1544 SRFY--EVNVPYSYKE------KKFENGDGYFIDRGEF---KMIKIPFFK----------
TsgC/1-1293 QGLF--SARLPYGMTE------KEMDNYDGYFIHKSKFVGMRCATIPANDATQSLVLTN-
TshC/1-1330 TTLIGDSYKGDFQELN------RKMQEGEARFVIGTEIKRLKMPTVL-------------
TsiC/1-1522 NIF---DVQMDYKQRS------VALNPGFGYLINRSDYITIKTPQL--------------
TsjC/1-1525 SVFS--SVNIGFENSK------RILGLGEGFLIVNRKAVPVK-LPAPFEGKVKFMNYIDK
```

```
EccC/1-1391 ---------
EssC/1-1479 ---------
TscC/1-1328 IRDRWHVVV
TsdC/1-1563 SVKS-----
TseC/1-1572 ---------
TsfC/1-1544 ---------
TsgC/1-1293 ---------
TshC/1-1330 ---------
TsiC/1-1522 ---------
TsjC/1-1525 LNEKLN---
```

Figure S2. Sequence alignment of the ten representative TsxC sequences, one from each of T7SSa – T7SSj. The region boxed in green is the FHA domains, in purple is the two transmembrane domains and in yellow the four NTPase domains. The consensus sequence for phosphothreonine recognition is given in red under the FHA domain region. The sequences used to generate this figure are WP_003408799.1 (EccC), WP_000549278.1 (EssC), WP_212980188.1 (TscC), WP_052118160.1 (TsdC), WP_213117477.1 (TseC), WP_010963368.1 (TsfC), WP_251417664.1 (TsgC), WP_172455381.1 (TshC), WP_212693241.1 (TsiC) and WP_257675217.1 (TsjC).

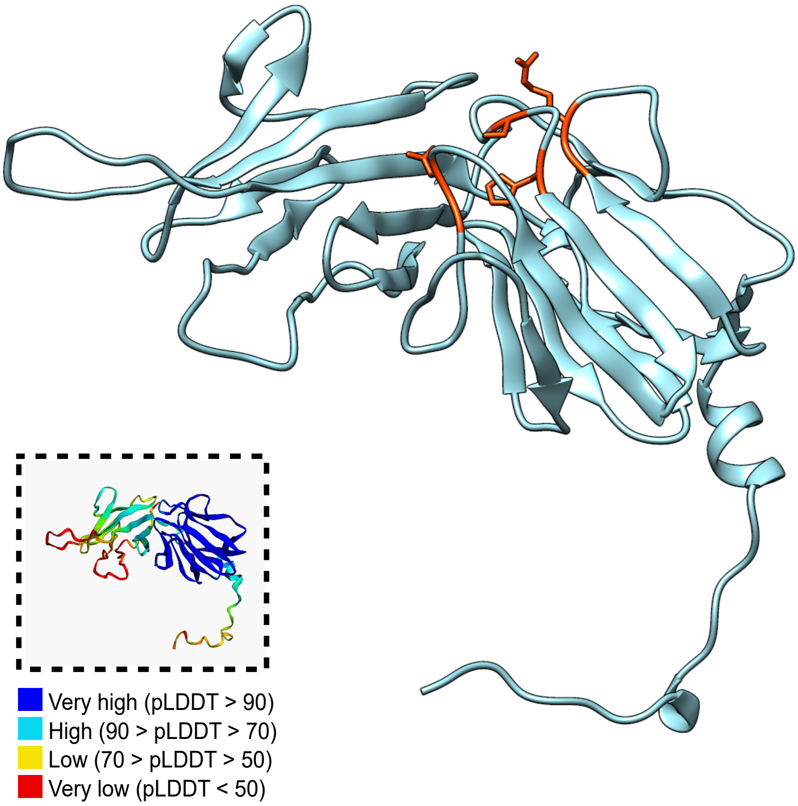


Figure S3. AlphaFold2 model of residues 1-250 of TsdC (WP_052118160.1), showing predicted FHA domains (cyan). Highlighted in red are conserved phosphopeptide-binding residues of the second FHA domain. Model coloured using UCSF Chimera. Inset: model coloured by pLDDT score (per-residue model confidence).

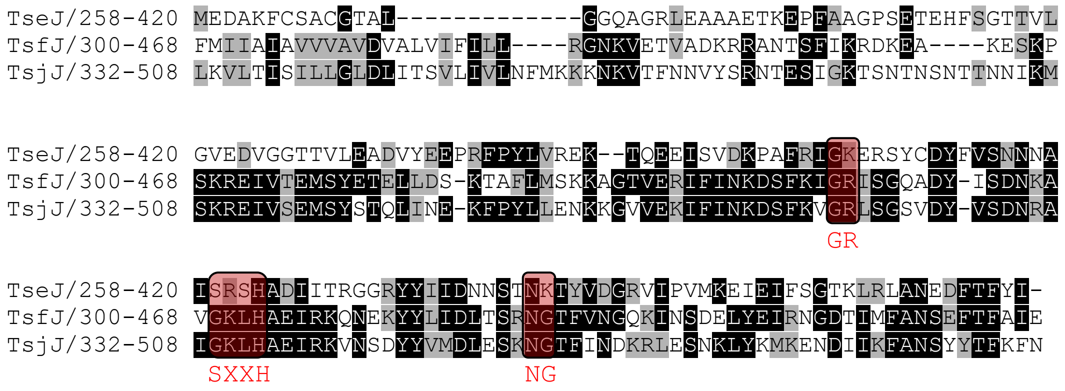


Figure S4. Amino acid alignment of predicted FHA domains of TsxJ components TseJ (WP_213117486.1), TsfJ (WP_010963365.1) and TsjJ (WP_257675219.1). Phosphopeptide-binding motifs are highlighted in red, with the phosphothreonine binding consensus sequence shown underneath.
