## Supplementary figures and images for "Multiple variants of the type VII secretion system in Gram-positive bacteria"

### Supplemental Data 3

**Supplementary Data 3**

AlphaFold2 models for each of the predicted T7SS components

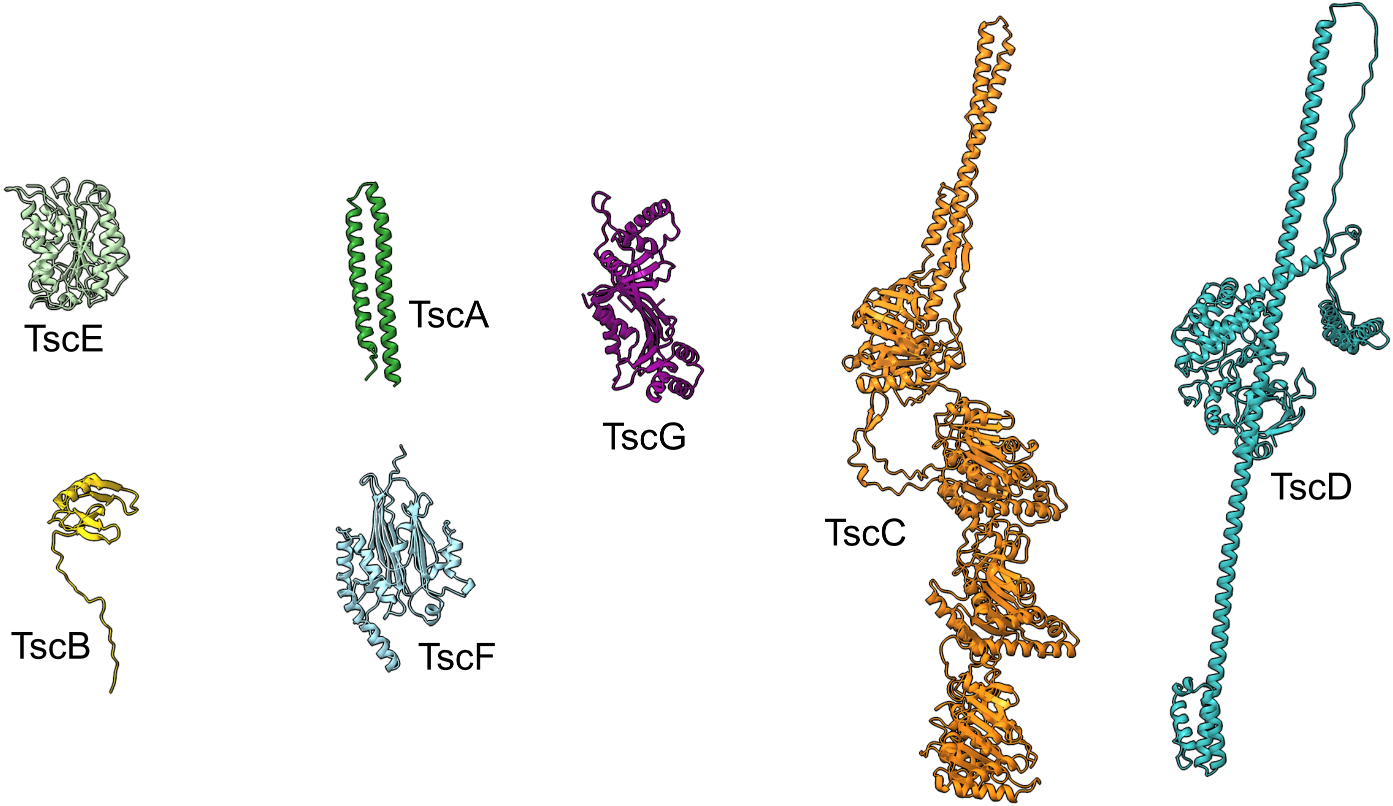

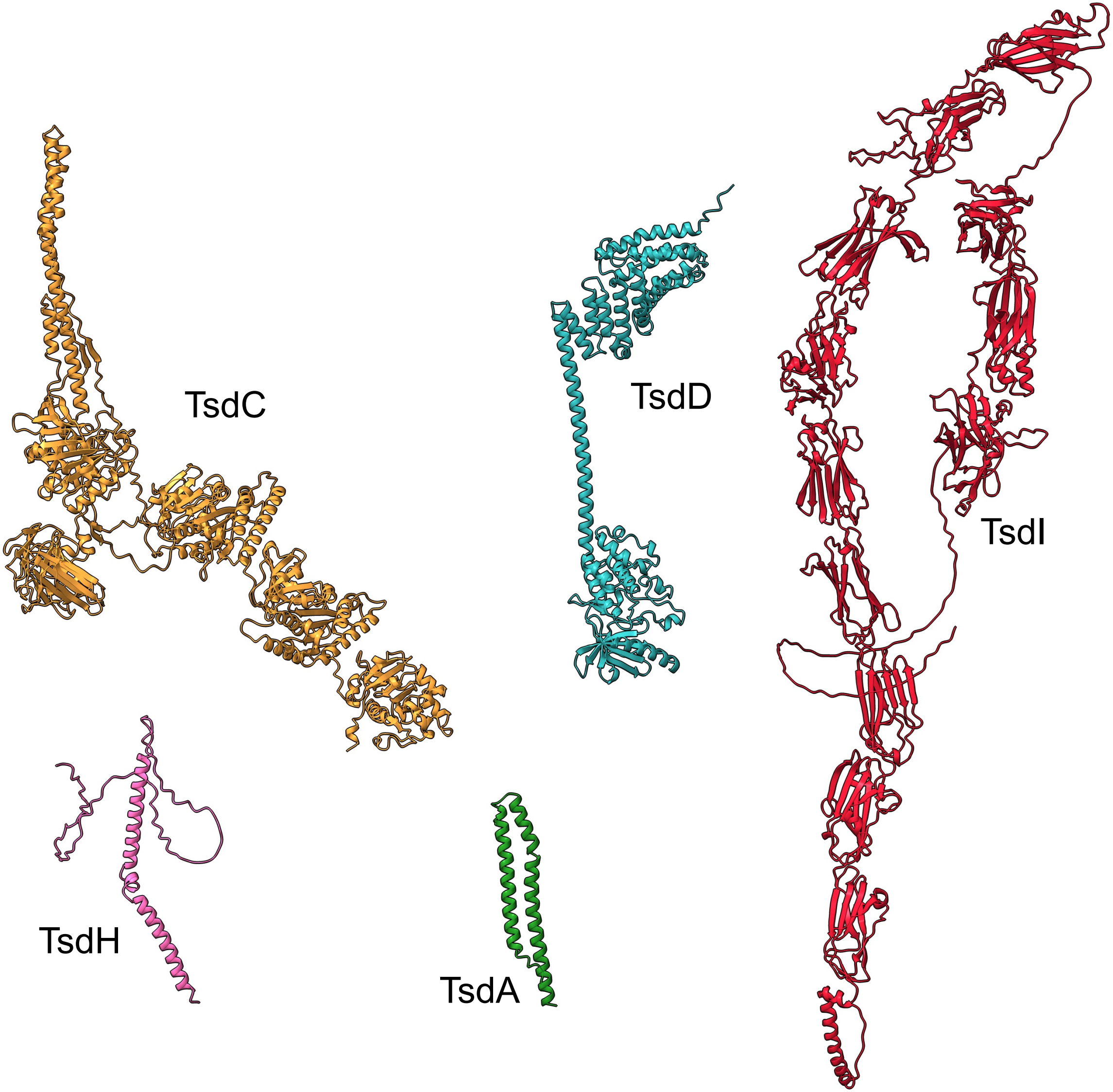

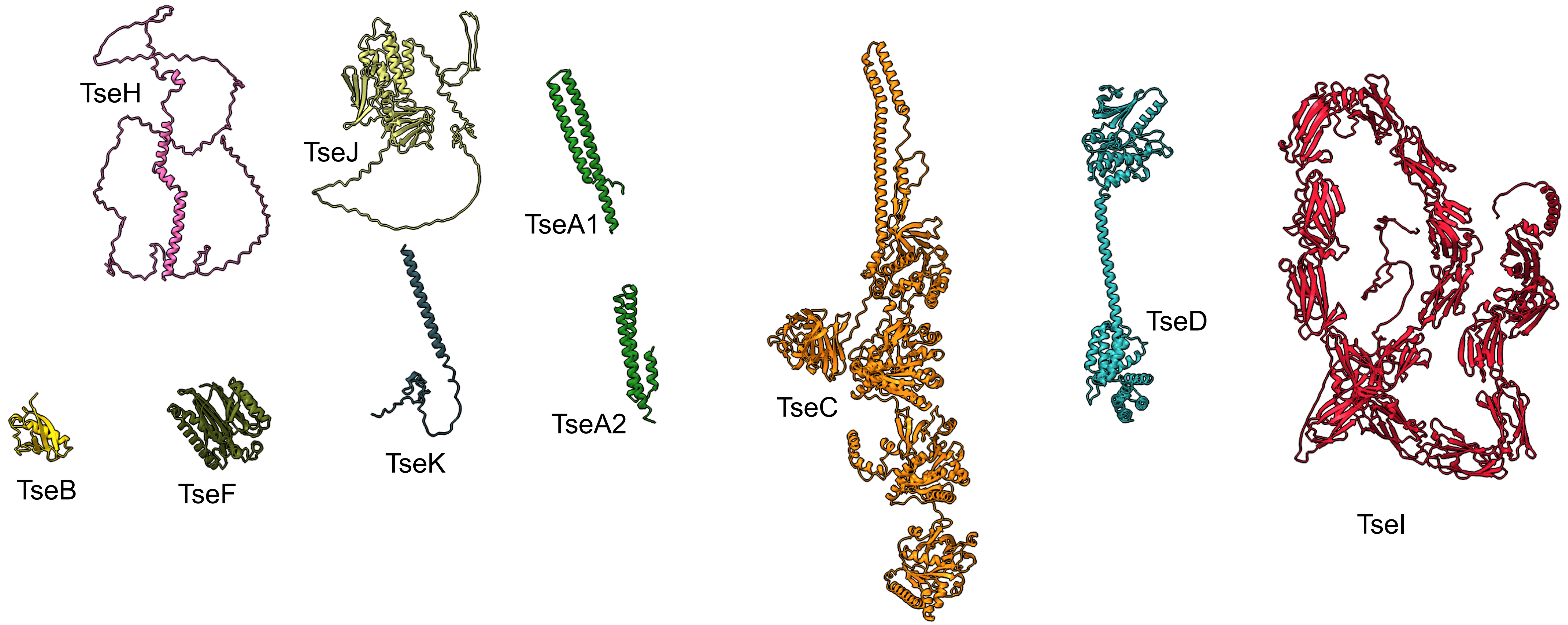

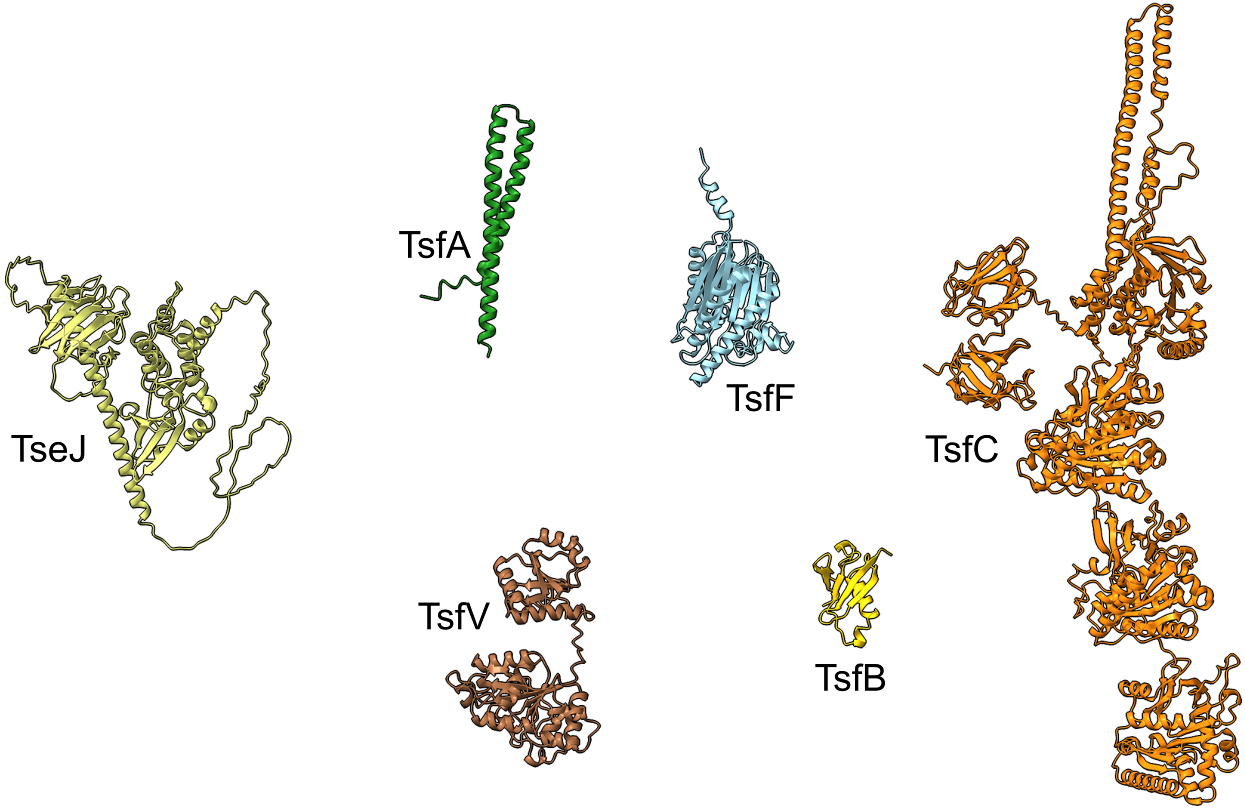

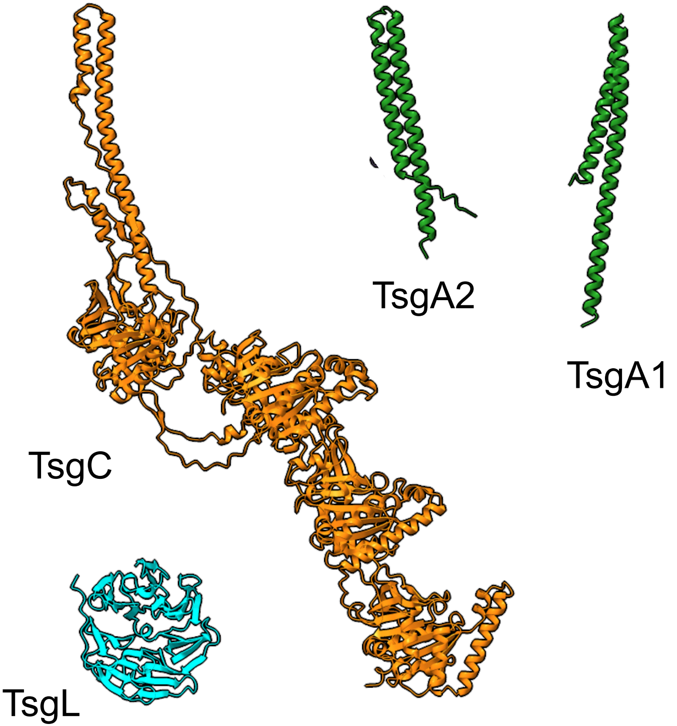

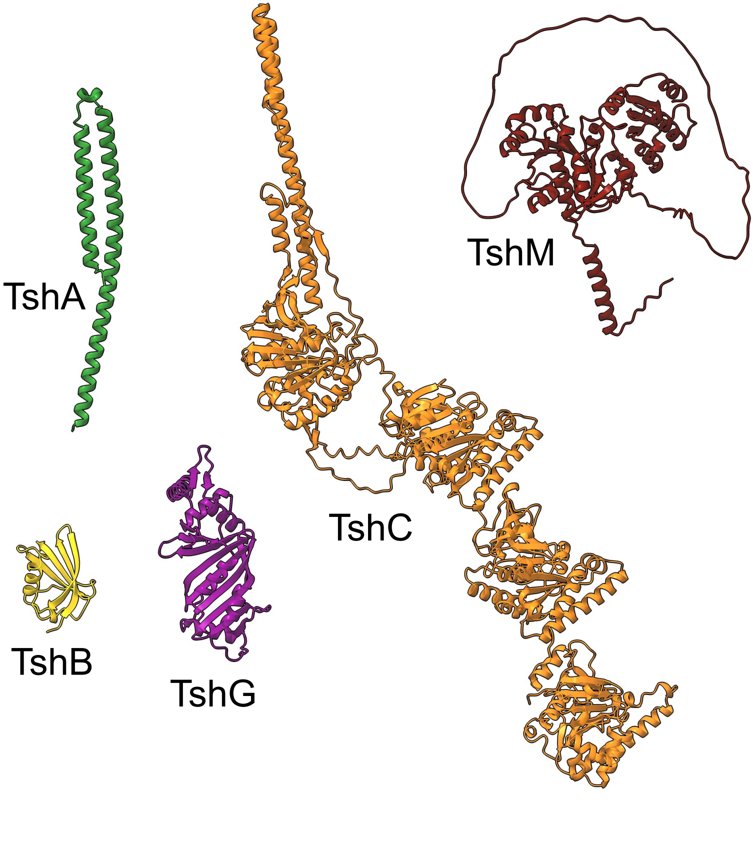

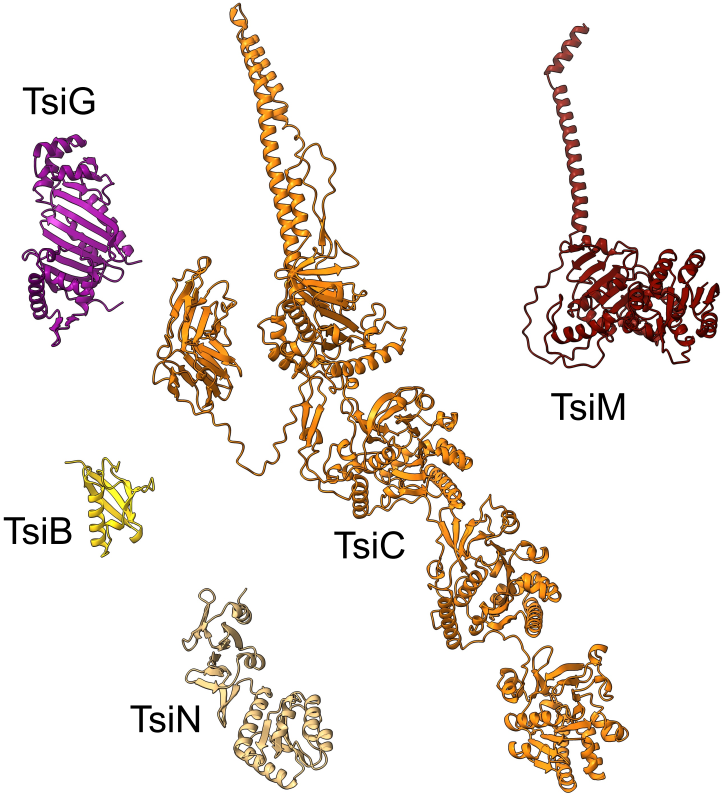

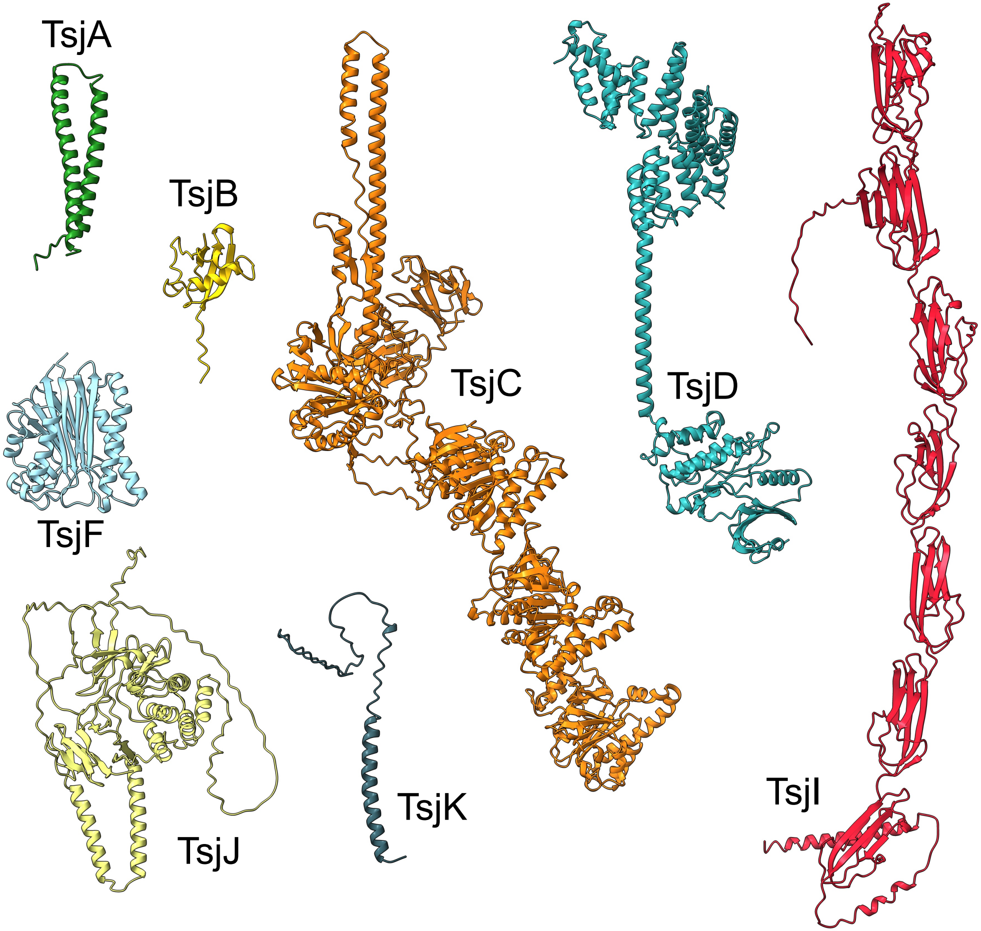
