## Supplemental Data 1 for "Multiple variants of the type VII secretion system in Gram-positive bacteria"

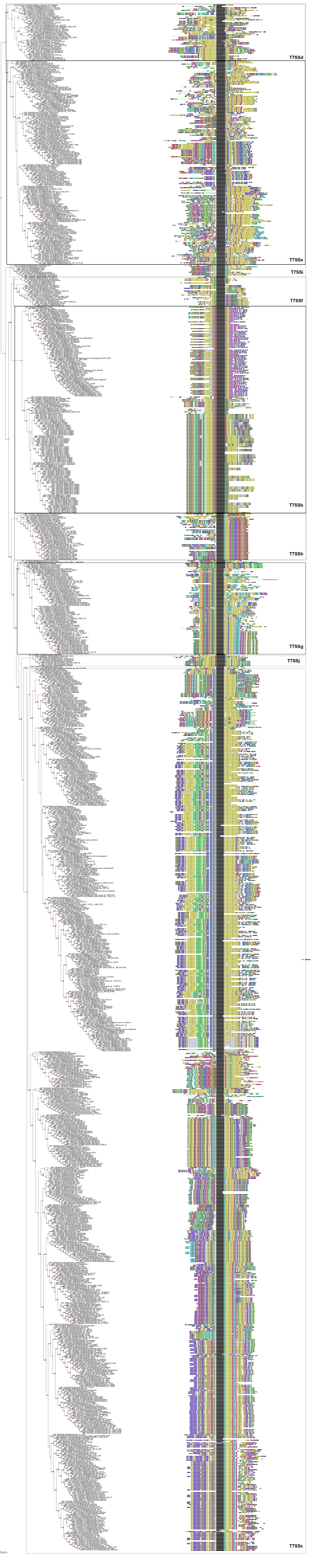
T7SSd
T7SSe
T7SSi
T7SSf
T7SSb
T7SSh
T7SSg
T7SSj
T7SSc

| | | |
|---|---|---|
| 1(1) | WP_199695248.1 | WP_199695248.1 bacteriocin-processing peptidase family protein |
| 1(1) | WP_283157901.1 | WP_283157901.1 DUF6382 domain-containing protein |
| 1(1) | WP_098255404.1 | WP_098255404.1 SseB family protein |
| 1(1) | WP_069193872.1 | WP_069193872.1 WXG100 family type VII secretion target |
| 1(1) | WP_251638762.1 | WP_251638762.1 FHA domain-containing protein |
| 1(1) | WP_010676857.1 | WP_010676857.1 serine/threonine-protein kinase |
| 1(1) | WP_077893585.1 | WP_077893585.1 Ig-like domain-containing protein |
| 1(1) | WP_307394626.1 | WP_307394626.1 RHS repeat-associated core domain-containing protein |
| 1(4) | WP_099354911.1 | WP_099354911.1 WXG100 family type VII secretion target |
| 1(1) | WP_027088432.1 | WP_027088432.1 hypothetical protein |
| 1(1) | WP_026478110.1 | WP_026478110.1 WXG100 family type VII secretion target |
| 1(2) | WP_212805121.1 | WP_212805121.1 hypothetical protein |
| 1(1) | WP_108465350.1 | WP_108465350.1 serine/threonine-protein kinase |
| 1(1) | WP_073008239.1 | WP_073008239.1 EndoU domain-containing protein |
| 1(2) | WP_036626770.1 | WP_036626770.1 WXG100 family type VII secretion target |
| 1(1) | WP_053492740.1 | WP_053492740.1 hypothetical protein |
| 1(1) | WP_283952947.1 | WP_283952947.1 DUF2974 domain-containing protein |
| 1(1) | WP_226679107.1 | WP_226679107.1 LysM peptidoglycan-binding domain-containing protein |
| 1(1) | WP_168976285.1 | WP_168976285.1 hypothetical protein |
| 1(1) | WP_098248510.1 | WP_098248510.1 hypothetical protein |
| 1(1) | WP_143316571.1 | WP_143316571.1 FHA domain-containing protein |
| 1(2) | WP_281813096.1 | WP_281813096.1 WXG100 family type VII secretion target |
| 1(1) | WP_081631380.1 | WP_081631380.1 type VII secretion protein EsaA |
| 1(1) | WP_101300059.1 | WP_101300059.1 copper amine oxidase N-terminal domain-containing protein |
| 1(1) | WP_048826559.1 | WP_048826559.1 WXG100 family type VII secretion target |
| 1(1) | WP_262748114.1 | WP_262748114.1 bacteriocin-processing peptidase family protein |
| 1(2) | WP_251426369.1 | WP_251426369.1 WXG100 family type VII secretion target |
| 1(1) | WP_133332986.1 | WP_133332986.1 Ig-like domain-containing protein |
| 1(1) | WP_211745828.1 | WP_211745828.1 hypothetical protein |
| 1(1) | WP_094593877.1 | WP_094593877.1 hypothetical protein |
| 1(2) | WP_061829413.1 | WP_061829413.1 S-layer homology domain-containing protein |
| 1(1) | WP_249410237.1 | WP_249410237.1 hypothetical protein |
| 1(1) | WP_040360538.1 | WP_040360538.1 WXG100 family type VII secretion target |
| 1(1) | WP_034675383.1 | WP_034675383.1 Ig-like domain-containing protein |
| 1(1) | WP_290403365.1 | WP_290403365.1 S-layer homology domain-containing protein |
| 1(1) | WP_144479047.1 | WP_144479047.1 DUF6531 domain-containing protein, partial |
| 1(1) | WP_090735413.1 | WP_090735413.1 copper amine oxidase N-terminal domain-containing protein |
| 1(1) | WP_254759304.1 | WP_254759304.1 serine/threonine-protein kinase |
| 1(1) | WP_281103269.1 | WP_281103269.1 tetratricopeptide repeat protein |
| 1(1) | WP_307192892.1 | WP_307192892.1 protein kinase |
| 1(2) | WP_195431284.1 | WP_195431284.1 WXG100 family type VII secretion target |
| 1(1) | WP_228463449.1 | WP_228463449.1 hypothetical protein |
| 1(1) | WP_308110115.1 | WP_308110115.1 DUF6382 domain-containing protein |
| 1(1) | WP_258296278.1 | WP_258296278.1 hypothetical protein |
| 1(1) | WP_088550044.1 | WP_088550044.1 EsaB/YukD family protein |
| 1(1) | WP_064639012.1 | WP_064639012.1 hypothetical protein |
| 1(1) | WP_050752347.1 | WP_050752347.1 Ig-like domain-containing protein |
| 1(2) | WP_194339780.1 | WP_194339780.1 type VII secretion protein EsaA |
| 1(1) | WP_098638818.1 | WP_098638818.1 bacteriocin-processing peptidase family protein |
| 1(1) | WP_167434148.1 | WP_167434148.1 protein kinase family protein |
| 1(1) | WP_253299366.1 | WP_253299366.1 hypothetical protein |
| 1(1) | WP_158701618.1 | WP_158701618.1 T7SS effector LXG polymorphic toxin |
| 1(2) | WP_213117479.1 | WP_213117479.1 WXG100 family type VII secretion target |
| 1(1) | WP_254759300.1 | WP_254759300.1 WXG100 family type VII secretion target |
| 1(2) | WP_271006643.1 | WP_271006643.1 glycohydrolase toxin TNT-related protein |
| 1(1) | WP_169509267.1 | WP_169509267.1 hypothetical protein |
| 1(1) | WP_013658714.1 | WP_013658714.1 EsaB/YukD family protein |
| 1(1) | WP_242251783.1 | WP_242251783.1 WXG100 family type VII secretion target |
| 1(1) | WP_098319879.1 | WP_098319879.1 bacteriocin-processing peptidase family protein |
| 1(1) | WP_064019316.1 | WP_064019316.1 type VII secretion target |
| 1(26) | WP_013370783.1 | WP_013370783.1 hypothetical protein |
| 1(2) | WP_139895506.1 | WP_139895506.1 hypothetical protein |
| 1(1) | WP_121610592.1 | WP_121610592.1 hypothetical protein |
| 1(1) | WP_064019322.1 | WP_064019322.1 WXG100 family type VII secretion target |
| 1(1) | WP_179034777.1 | WP_179034777.1 AAA family ATPase |
| 1(1) | WP_208926616.1 | WP_208926616.1 WXG100 family type VII secretion target |
| 1(1) | WP_190945006.1 | WP_190945006.1 WXG100 family type VII secretion target |
| 1(1) | WP_098836666.1 | WP_098836666.1 T7SS effector LXG polymorphic toxin |
| 1(2) | WP_100537977.1 | WP_100537977.1 hypothetical protein |
| 1(1) | WP_068619107.1 | WP_068619107.1 hypothetical protein |
| 1(1) | WP_150457766.1 | WP_150457766.1 DUF5050 domain-containing protein |
| 1(1) | WP_098288684.1 | WP_098288684.1 tetratricopeptide repeat protein |
| 1(1) | WP_087194489.1 | WP_087194489.1 WXG100 family type VII secretion target |
| 1(1) | WP_267414311.1 | WP_267414311.1 bacteriocin-processing peptidase family protein |
| 1(1) | WP_136604817.1 | WP_136604817.1 hypothetical protein |
| 1(2) | WP_185469290.1 | WP_185469290.1 RHS repeat protein |
| 1(3) | WP_076322205.1 | WP_076322205.1 hypothetical protein |
| 1(1) | WP_039273538.1 | WP_039273538.1 membrane protein |
| 1(1) | WP_276917313.1 | WP_276917313.1 hypothetical protein |
| 1(1) | WP_251638758.1 | WP_251638758.1 serine/threonine-protein kinase |
| 1(3) | WP_006674865.1 | WP_006674865.1 hypothetical protein |
| 1(2) | WP_110899364.1 | WP_110899364.1 WXG100 family type VII secretion target |
| 1(1) | WP_233530992.1 | WP_233530992.1 AAA family ATPase |
| 1(1) | WP_110934065.1 | WP_110934065.1 DUF2974 domain-containing protein |
| 1(1) | WP_307738727.1 | WP_307738727.1 hypothetical protein |
| 1(1) | WP_216791212.1 | WP_216791212.1 DUF2974 domain-containing protein |
| 1(1) | WP_212805123.1 | WP_212805123.1 tetratricopeptide repeat protein |
| 1(1) | WP_306007804.1 | WP_306007804.1 hypothetical protein |
| 1(1) | WP_216830395.1 | WP_216830395.1 hypothetical protein |
| 1(1) | WP_078419976.1 | WP_078419976.1 bacteriocin-processing peptidase family protein |
| 1(1) | WP_215038278.1 | WP_215038278.1 WXG100 family type VII secretion target |
| 1(1) | WP_304216874.1 | WP_304216874.1 DUF6382 domain-containing protein |
| 1(1) | WP_013914205.1 | WP_013914205.1 S-layer homology domain-containing protein |
| 1(1) | WP_007147976.1 | WP_007147976.1 YhgE/Pip domain-containing protein |
| 1(1) | WP_242264958.1 | WP_242264958.1 hypothetical protein |
| 1(7) | WP_025719059.1 | WP_025719059.1 WXG100 family type VII secretion target |
| 1(2) | WP_228469892.1 | WP_228469892.1 glycohydrolase toxin TNT-related protein |
| 1(1) | WP_181603070.1 | WP_181603070.1 WXG100 family type VII secretion target |
| 1(1) | WP_151208728.1 | WP_151208728.1 hypothetical protein |
| 1(2) | WP_271002888.1 | WP_271002888.1 type VII secretion protein EsaA |
| 1(1) | WP_210087125.1 | WP_210087125.1 hypothetical protein |
| 1(1) | WP_000506917.1 | WP_000506917.1 tetratricopeptide repeat protein |

| | | |
|---|---|---|
| 1(1) | WP_215598790.1 | WP_215598790.1 tetratricopeptide repeat protein |
| 1(1) | WP_251596670.1 | WP_251596670.1 WXG100 family type VII secretion target |
| 1(1) | WP_305022921.1 | WP_305022921.1 hypothetical protein |
| 1(1) | WP_195343923.1 | WP_195343923.1 Ig-like domain-containing protein |
| 1(1) | WP_080449897.1 | WP_080449897.1 WXG100 family type VII secretion target |
| 1(6) | WP_047909612.1 | WP_047909612.1 WXG100 family type VII secretion target |
| 1(1) | WP_024027632.1 | WP_024027632.1 WXG100 family type VII secretion target |
| 1(1) | WP_137052036.1 | WP_137052036.1 tetratricopeptide repeat protein |
| 1(23) | WP_023988263.1 | WP_023988263.1 hypothetical protein |
| 1(1) | WP_258875221.1 | WP_258875221.1 S-layer homology domain-containing protein |
| 1(2) | WP_023988253.1 | WP_023988253.1 S-layer homology domain-containing protein |
| 1(1) | WP_098284015.1 | WP_098284015.1 type VII secretion protein EsaA |
| 1(1) | WP_270699380.1 | WP_270699380.1 tetratricopeptide repeat protein |
| 1(1) | WP_141540984.1 | WP_141540984.1 WXG100 family type VII secretion target |
| 1(1) | WP_205001944.1 | WP_205001944.1 hypothetical protein |
| 1(1) | WP_251557374.1 | WP_251557374.1 serine/threonine-protein kinase |
| 1(2) | WP_045243840.1 | WP_045243840.1 S-layer homology domain-containing protein |
| 1(1) | WP_221875377.1 | WP_221875377.1 DUF2974 domain-containing protein |
| 1(1) | WP_283952940.1 | WP_283952940.1 Ig-like domain repeat protein |
| 1(1) | WP_247902246.1 | WP_247902246.1 hypothetical protein |
| 1(2) | WP_204412719.1 | WP_204412719.1 pore-forming ESAT-6 family protein |
| 1(1) | WP_223592686.1 | WP_223592686.1 WXG100 family type VII secretion target |
| 1(1) | WP_307199936.1 | WP_307199936.1 WXG100 family type VII secretion target |
| 1(1) | WP_028401642.1 | WP_028401642.1 hypothetical protein |
| 1(2) | WP_040418520.1 | WP_040418520.1 WXG100 family type VII secretion target |
| 1(2) | WP_185516169.1 | WP_185516169.1 pre-toxin TG domain-containing protein |
| 1(1) | WP_242805836.1 | WP_242805836.1 hypothetical protein |
| 1(1) | WP_073540573.1 | WP_073540573.1 tetratricopeptide repeat protein |
| 1(1) | WP_064473945.1 | WP_064473945.1 bacteriocin-processing peptidase family protein |
| 1(1) | WP_307399115.1 | WP_307399115.1 glucosaminidase domain-containing protein |
| 1(1) | WP_283927237.1 | WP_283927237.1 copper amine oxidase N-terminal domain-containing protein |
| 1(1) | WP_270065058.1 | WP_270065058.1 DUF2974 domain-containing protein |
| 1(1) | WP_013658705.1 | WP_013658705.1 copper amine oxidase N-terminal domain-containing protein |
| 1(1) | WP_283904343.1 | WP_283904343.1 FHA domain-containing protein |
| 1(2) | WP_072732442.1 | WP_072732442.1 hypothetical protein |
| 1(1) | WP_139829221.1 | WP_139829221.1 hypothetical protein |
| 1(1) | WP_026478108.1 | WP_026478108.1 EsaB/YukD family protein |
| 1(1) | WP_289344197.1 | WP_289344197.1 serine/threonine-protein kinase |
| 1(1) | WP_243864605.1 | WP_243864605.1 DUF2974 domain-containing protein |
| 1(1) | WP_126996285.1 | WP_126996285.1 peptidoglycan-binding protein |
| 1(2) | WP_143804315.1 | WP_143804315.1 hypothetical protein |
| 1(1) | WP_277142088.1 | WP_277142088.1 hypothetical protein |
| 1(1) | WP_116353257.1 | WP_116353257.1 hypothetical protein |
| 1(2) | WP_149051225.1 | WP_149051225.1 DUF6531 domain-containing protein |
| 1(1) | WP_181747869.1 | WP_181747869.1 hypothetical protein |
| 1(1) | WP_212693242.1 | WP_212693242.1 hypothetical protein |
| 1(1) | WP_121474478.1 | WP_121474478.1 hypothetical protein |
| 1(1) | WP_273383882.1 | WP_273383882.1 FHA domain-containing protein |
| 1(1) | WP_234414303.1 | WP_234414303.1 WXG100 family type VII secretion target |
| 1(1) | WP_113058719.1 | WP_113058719.1 hypothetical protein |
| 1(2) | WP_195343926.1 | WP_195343926.1 T7SS effector LXG polymorphic toxin |
| 1(5) | WP_071220203.1 | WP_071220203.1 hypothetical protein |
| 1(2) | WP_010963374.1 | WP_010963374.1 tetratricopeptide repeat protein |
| 1(1) | WP_013490123.1 | WP_013490123.1 FHA domain-containing protein |
| 1(1) | WP_289613139.1 | WP_289613139.1 T7SS effector LXG polymorphic toxin |
| 1(1) | WP_127592426.1 | WP_127592426.1 hypothetical protein |
| 1(1) | WP_137698106.1 | WP_137698106.1 copper amine oxidase N-terminal domain-containing protein |
| 1(2) | WP_194230401.1 | WP_194230401.1 WXG100 family type VII secretion target |
| 1(3) | WP_113755281.1 | WP_113755281.1 WXG100 family type VII secretion target |
| 1(1) | WP_289398701.1 | WP_289398701.1 DNA/RNA non-specific endonuclease |
| 1(1) | WP_048896847.1 | WP_048896847.1 tetratricopeptide repeat protein |
| 1(1) | WP_003516556.1 | WP_003516556.1 hypothetical protein |
| 1(1) | WP_159146505.1 | WP_159146505.1 tetratricopeptide repeat protein |
| 1(1) | WP_308082244.1 | WP_308082244.1 WXG100 family type VII secretion target |
| 1(1) | WP_229137196.1 | WP_229137196.1 WXG100 family type VII secretion target |
| 1(1) | WP_249897328.1 | WP_249897328.1 DUF5107 domain-containing protein |
| 1(1) | WP_000506909.1 | WP_000506909.1 tetratricopeptide repeat protein |
| 1(4) | WP_077833109.1 | WP_077833109.1 WXG100 family type VII secretion target |
| 1(1) | WP_285748179.1 | WP_285748179.1 hypothetical protein |
| 1(3) | WP_078181466.1 | WP_078181466.1 hypothetical protein |
| 1(1) | WP_242634582.1 | WP_242634582.1 Ig-like domain-containing protein |
| 1(2) | WP_018883271.1 | WP_018883271.1 WXG100 family type VII secretion target |
| 1(1) | WP_256872561.1 | WP_256872561.1 RHS repeat-associated core domain-containing protein |
| 1(1) | WP_150364198.1 | WP_150364198.1 RHS repeat-associated core domain-containing protein |
| 1(1) | WP_188535082.1 | WP_188535082.1 C39 family peptidase |
| 1(1) | WP_000391142.1 | WP_000391142.1 hypothetical protein |
| 1(1) | WP_058486874.1 | WP_058486874.1 stalk domain-containing protein |
| 1(1) | WP_026882391.1 | WP_026882391.1 WXG100 family type VII secretion target |
| 1(1) | WP_257675918.1 | WP_257675918.1 hypothetical protein |
| 1(1) | WP_019424149.1 | WP_019424149.1 hypothetical protein |
| 1(1) | WP_226086172.1 | WP_226086172.1 FHA domain-containing protein |
| 1(1) | WP_053361022.1 | WP_053361022.1 DUF2974 domain-containing protein |
| 1(1) | WP_014281180.1 | WP_014281180.1 hypothetical protein |
| 1(3) | WP_053403781.1 | WP_053403781.1 WXG100 family type VII secretion target |
| 1(1) | WP_277548444.1 | WP_277548444.1 WG repeat-containing protein |
| 1(1) | WP_307537578.1 | WP_307537578.1 hypothetical protein |
| 1(5) | WP_095287660.1 | WP_095287660.1 hypothetical protein |
| 1(1) | WP_086424321.1 | WP_086424321.1 T7SS effector LXG polymorphic toxin |
| 1(1) | WP_212330900.1 | WP_212330900.1 hypothetical protein |
| 1(1) | WP_196427068.1 | WP_196427068.1 hypothetical protein |
| 1(1) | WP_101195714.1 | WP_101195714.1 SseB family protein |
| 1(1) | WP_098679905.1 | WP_098679905.1 tetratricopeptide repeat protein |
| 1(1) | WP_119601486.1 | WP_119601486.1 WXG100 family type VII secretion target |
| 1(1) | WP_139796092.1 | WP_139796092.1 hypothetical protein |
| 1(1) | WP_140157198.1 | WP_140157198.1 WXG100 family type VII secretion target |
| 1(1) | WP_098961909.1 | WP_098961909.1 tetratricopeptide repeat protein |
| 1(2) | WP_103626233.1 | WP_103626233.1 WXG100 family type VII secretion target |
| 1(2) | WP_270993639.1 | WP_270993639.1 DUF6531 domain-containing protein |
| 1(1) | WP_120462952.1 | WP_120462952.1 EsaB/YukD family protein |
| 1(1) | WP_216531933.1 | WP_216531933.1 serine/threonine-protein kinase |
| 1(1) | WP_181754692.1 | WP_181754692.1 EsaB/YukD family protein |

1(1) WP_083599919.1 WP_083599919.1 pore-forming ESAT-6 family protein
1(1) WP_029517069.1 WP_029517069.1 stalk domain-containing protein
1(1) WP_021291836.1 WP_021291836.1 hypothetical protein
1(1) WP_230585140.1 WP_230585140.1 WXG100 family type VII secretion target
1(1) WP_063223277.1 WP_063223277.1 hypothetical protein
1(1) WP_207994160.1 WP_207994160.1 tetratricopeptide repeat protein
1(1) WP_148125759.1 WP_148125759.1 hypothetical protein, partial
1(1) WP_068782405.1 WP_068782405.1 pre-toxin TG domain-containing protein
1(1) WP_107734212.1 WP_107734212.1 S-layer homology domain-containing protein
1(1) WP_256227717.1 WP_256227717.1 hypothetical protein
1(1) WP_262787111.1 WP_262787111.1 hypothetical protein
1(2) WP_014368242.1 WP_014368242.1 EsaB/YukD family protein
1(1) WP_262725724.1 WP_262725724.1 WXG100 family type VII secretion target
1(1) WP_308110113.1 WP_308110113.1 protein kinase
1(2) WP_042134268.1 WP_042134268.1 hypothetical protein
1(1) WP_251638766.1 WP_251638766.1 hypothetical protein
1(1) WP_051569333.1 WP_051569333.1 hypothetical protein
1(1) WP_024632682.1 WP_024632682.1 WXG100 family type VII secretion target
1(2) WP_018670525.1 WP_018670525.1 WG repeat-containing protein
1(1) WP_071724729.1 WP_071724729.1 hypothetical protein
1(1) WP_179034789.1 WP_179034789.1 WXG100 family type VII secretion target
1(1) WP_307284672.1 WP_307284672.1 cadherin domain-containing protein
1(1) WP_098167594.1 WP_098167594.1 protein kinase
1(3) WP_044584389.1 WP_044584389.1 T7SS effector LXG polymorphic toxin
1(1) WP_155703924.1 WP_155703924.1 hypothetical protein
1(1) WP_110934060.1 WP_110934060.1 VWA domain-containing protein
1(1) WP_307284690.1 WP_307284690.1 FHA domain-containing protein
1(1) WP_251417667.1 WP_251417667.1 WXG100 family type VII secretion target
1(1) WP_270716432.1 WP_270716432.1 SseB family protein
1(1) WP_282742030.1 WP_282742030.1 serine/threonine-protein kinase
1(1) WP_110934071.1 WP_110934071.1 WXG100 family type VII secretion target
1(1) WP_141552654.1 WP_141552654.1 WXG100 family type VII secretion target, partial
1(1) WP_077320207.1 WP_077320207.1 T7SS effector LXG polymorphic toxin
1(3) WP_002022919.1 WP_002022919.1 hypothetical protein
1(1) WP_211023023.1 WP_211023023.1 WXG100 family type VII secretion target
1(1) WP_136372958.1 WP_136372958.1 hypothetical protein
1(1) WP_251638782.1 WP_251638782.1 WXG100 family type VII secretion target
1(1) WP_073008248.1 WP_073008248.1 TNT domain-containing protein
1(1) WP_223617608.1 WP_223617608.1 WXG100 family type VII secretion target
1(1) WP_026572744.1 WP_026572744.1 protein kinase
1(1) WP_169744750.1 WP_169744750.1 EsaB/YukD family protein
1(2) WP_160807257.1 WP_160807257.1 HNH endonuclease
1(1) WP_208926590.1 WP_208926590.1 serine/threonine-protein kinase
1(1) WP_215572755.1 WP_215572755.1 HNH endonuclease
1(1) WP_277166577.1 WP_277166577.1 TNT domain-containing protein
1(1) WP_098250527.1 WP_098250527.1 SseB family protein
1(1) WP_306077397.1 WP_306077397.1 hypothetical protein
1(1) WP_195325250.1 WP_195325250.1 tetratricopeptide repeat protein
1(1) WP_145156806.1 WP_145156806.1 WXG100 family type VII secretion target
1(1) WP_090761829.1 WP_090761829.1 FHA domain-containing protein
1(1) WP_098235623.1 WP_098235623.1 WXG100 family type VII secretion target
1(1) WP_066020842.1 WP_066020842.1 WXG100 family type VII secretion target
1(1) WP_141559744.1 WP_141559744.1 deaminase domain-containing protein
1(1) WP_307284697.1 WP_307284697.1 Ig-like domain-containing protein
1(1) WP_263324555.1 WP_263324555.1 serine/threonine-protein kinase
1(1) WP_293202260.1 WP_293202260.1 hypothetical protein
1(1) WP_125084780.1 WP_125084780.1 WXG100 family type VII secretion target
1(1) WP_270472030.1 WP_270472030.1 hypothetical protein
1(1) WP_214706011.1 WP_214706011.1 Ig-like domain-containing protein
1(1) WP_034526607.1 WP_034526607.1 hypothetical protein
1(2) WP_268599729.1 WP_268599729.1 WXG100 family type VII secretion target
1(1) WP_042125634.1 WP_042125634.1 DUF2974 domain-containing protein
1(1) WP_247902243.1 WP_247902243.1 hypothetical protein
1(1) WP_077852416.1 WP_077852416.1 WXG100 family type VII secretion target
1(1) WP_223592682.1 WP_223592682.1 Mbeg1-like protein
1(2) WP_185639019.1 WP_185639019.1 type VII secretion protein EsaA
1(1) WP_269667704.1 WP_269667704.1 HNH/ENDO VII family nuclease
1(1) WP_209872745.1 WP_209872745.1 hypothetical protein
1(1) WP_095287659.1 WP_095287659.1 hypothetical protein
1(1) WP_069355512.1 WP_069355512.1 SseB family protein
1(1) WP_116334077.1 WP_116334077.1 WG repeat-containing protein
1(1) WP_149190447.1 WP_149190447.1 bacteriocin-processing peptidase family protein
1(1) WP_106765958.1 WP_106765958.1 hypothetical protein
1(1) WP_013658710.1 WP_013658710.1 hypothetical protein
1(1) WP_210364455.1 WP_210364455.1 WXG100 family type VII secretion target
1(1) WP_199925555.1 WP_199925555.1 hypothetical protein
1(1) WP_289342267.1 WP_289342267.1 FHA domain-containing protein
1(1) WP_270065054.1 WP_270065054.1 WXG100 family type VII secretion target
1(2) WP_010963370.1 WP_010963370.1 WXG100 family type VII secretion target
1(1) WP_002141597.1 WP_002141597.1 SseB family protein
1(1) WP_246021131.1 WP_246021131.1 serine/threonine-protein kinase
1(1) WP_013370769.1 WP_013370769.1 S-layer homology domain-containing protein
1(1) WP_065228771.1 WP_065228771.1 SseB family protein
1(1) WP_223068047.1 WP_223068047.1 WXG100 family type VII secretion target
1(1) WP_295628337.1 WP_295628337.1 protein kinase
1(1) WP_149885341.1 WP_149885341.1 hypothetical protein
1(1) WP_306007835.1 WP_306007835.1 CdiA family toxin C-terminal domain-containing protein
1(1) WP_270784519.1 WP_270784519.1 WXG100 family type VII secretion target
1(1) WP_283910929.1 WP_283910929.1 DUF2974 domain-containing protein
1(4) WP_185474158.1 WP_185474158.1 type VII secretion protein EsaA
1(1) WP_287600008.1 WP_287600008.1 tetratricopeptide repeat protein
1(1) WP_268058020.1 WP_268058020.1 serine/threonine-protein kinase
1(1) WP_063223279.1 WP_063223279.1 tetratricopeptide repeat protein
1(1) WP_088004909.1 WP_088004909.1 hypothetical protein
1(1) WP_166244909.1 WP_166244909.1 hypothetical protein
1(1) WP_235715983.1 WP_235715983.1 phage tail tip lysozyme
1(1) WP_307388970.1 WP_307388970.1 DUF2974 domain-containing protein
1(3) WP_098266648.1 WP_098266648.1 EsaB/YukD family protein
1(1) WP_053783888.1 WP_053783888.1 hypothetical protein
1(1) WP_042462805.1 WP_042462805.1 DUF2974 domain-containing protein
1(1) WP_085499081.1 WP_085499081.1 copper amine oxidase N-terminal domain-containing protein
1(1) WP_196048853.1 WP_196048853.1 hypothetical protein
1(1) WP_115359660.1 WP_115359660.1 EsaB/YukD family protein

| | | |
|---|---|---|
| 1(1) | WP_289475118.1 | WP_289475118.1 hypothetical protein |
| 1(1) | WP_213506866.1 | WP_213506866.1 hypothetical protein |
| 1(1) | WP_024632686.1 | WP_024632686.1 hypothetical protein |
| 1(1) | WP_251426357.1 | WP_251426357.1 Ig-like domain-containing protein |
| 1(1) | WP_305166272.1 | WP_305166272.1 S-layer homology domain-containing protein |
| 1(2) | WP_068619109.1 | WP_068619109.1 WXG100 family type VII secretion target |
| 1(1) | WP_024027630.1 | WP_024027630.1 DUF2974 domain-containing protein |
| 1(1) | WP_069011424.1 | WP_069011424.1 copper amine oxidase N-terminal domain-containing protein |
| 1(1) | WP_098148061.1 | WP_098148061.1 SseB family protein |
| 1(1) | WP_087095005.1 | WP_087095005.1 hypothetical protein |
| 1(1) | WP_098659490.1 | WP_098659490.1 type VII secretion protein EsaA |
| 1(1) | WP_094870532.1 | WP_094870532.1 DUF2974 domain-containing protein |
| 1(1) | WP_012260837.1 | WP_012260837.1 hypothetical protein |
| 1(1) | WP_073008271.1 | WP_073008271.1 Ig-like domain-containing protein |
| 1(1) | WP_258425844.1 | WP_258425844.1 ESX-1 secretion-associated protein |
| 1(1) | WP_052352969.1 | WP_052352969.1 Ig-like domain-containing protein |
| 1(1) | WP_262725725.1 | WP_262725725.1 SseB family protein |
| 1(1) | WP_227774476.1 | WP_227774476.1 copper amine oxidase N-terminal domain-containing protein |
| 1(1) | WP_005546512.1 | WP_005546512.1 WXG100 family type VII secretion target |
| 1(2) | WP_003516579.1 | WP_003516579.1 WXG100 family type VII secretion target |
| 1(1) | WP_098252275.1 | WP_098252275.1 T7SS effector LXG polymorphic toxin |
| 1(1) | WP_115359657.1 | WP_115359657.1 WXG100 family type VII secretion target |
| 1(3) | WP_097999319.1 | WP_097999319.1 T7SS effector LXG polymorphic toxin |
| 1(1) | WP_207391261.1 | WP_207391261.1 HNH endonuclease |
| 1(45) | WP_013309990.1 | WP_013309990.1 WXG100 family type VII secretion target |
| 1(1) | WP_019392118.1 | WP_019392118.1 hypothetical protein |
| 1(1) | WP_281477416.1 | WP_281477416.1 WXG100 family type VII secretion target |
| 1(1) | WP_084117732.1 | WP_084117732.1 WXG100 family type VII secretion target |
| 1(2) | WP_185357254.1 | WP_185357254.1 T7SS effector LXG polymorphic toxin |
| 1(1) | WP_147786234.1 | WP_147786234.1 hypothetical protein |
| 1(1) | WP_077619645.1 | WP_077619645.1 WXG100 family type VII secretion target |
| 1(1) | WP_111268938.1 | WP_111268938.1 hypothetical protein |
| 1(2) | WP_001161863.1 | WP_001161863.1 DNA/RNA non-specific endonuclease |
| 1(1) | WP_277157944.1 | WP_277157944.1 hypothetical protein |
| 1(1) | WP_297183853.1 | WP_297183853.1 serine/threonine-protein kinase |
| 1(1) | WP_248063448.1 | WP_248063448.1 hypothetical protein |
| 1(1) | WP_119601488.1 | WP_119601488.1 WXG100 family type VII secretion target |
| 1(1) | WP_283927239.1 | WP_283927239.1 serine/threonine-protein kinase |
| 1(1) | WP_243182807.1 | WP_243182807.1 WXG100 family type VII secretion target |
| 1(1) | WP_122971537.1 | WP_122971537.1 DUF2974 domain-containing protein |
| 1(1) | WP_170880288.1 | WP_170880288.1 WXG100 family type VII secretion target |
| 1(1) | WP_232274504.1 | WP_232274504.1 LysM domain-containing protein |
| 1(1) | WP_163977058.1 | WP_163977058.1 hypothetical protein |
| 1(1) | WP_016100418.1 | WP_016100418.1 protein kinase |
| 1(1) | WP_155703944.1 | WP_155703944.1 S8 family serine peptidase |
| 1(1) | WP_251245807.1 | WP_251245807.1 DUF2974 domain-containing protein |
| 1(1) | WP_101195713.1 | WP_101195713.1 tetratricopeptide repeat protein |
| 1(5) | WP_002031623.1 | WP_002031623.1 hypothetical protein |
| 1(2) | WP_244879218.1 | WP_244879218.1 S-layer homology domain-containing protein |
| 1(1) | WP_149238904.1 | WP_149238904.1 hypothetical protein |
| 1(2) | WP_076281073.1 | WP_076281073.1 DUF2974 domain-containing protein |
| 1(1) | WP_247083047.1 | WP_247083047.1 hypothetical protein |
| 1(3) | WP_097952794.1 | WP_097952794.1 bacteriocin-processing peptidase family protein |
| 1(4) | WP_001030889.1 | WP_001030889.1 SMI1/KNR4 family protein |
| 1(1) | WP_301167498.1 | WP_301167498.1 hypothetical protein |
| 1(1) | WP_002031665.1 | WP_002031665.1 WXG100 family type VII secretion target |
| 1(1) | WP_195974874.1 | WP_195974874.1 WXG100 family type VII secretion target |
| 1(1) | WP_090805474.1 | WP_090805474.1 hypothetical protein |
| 1(1) | WP_194230400.1 | WP_194230400.1 cell envelope integrity protein TolA |
| 1(1) | WP_224839863.1 | WP_224839863.1 Mbeg1-like protein |
| 1(2) | WP_006289749.1 | WP_006289749.1 hypothetical protein |
| 1(1) | WP_223617611.1 | WP_223617611.1 tetratricopeptide repeat protein |
| 1(1) | WP_021291853.1 | WP_021291853.1 type VII secretion protein EsaA |
| 1(1) | WP_085499122.1 | WP_085499122.1 AAA family ATPase |
| 1(1) | WP_076119215.1 | WP_076119215.1 DUF2974 domain-containing protein |
| 1(1) | WP_199419768.1 | WP_199419768.1 DUF2974 domain-containing protein |
| 1(1) | WP_211022799.1 | WP_211022799.1 DUF87 domain-containing protein |
| 1(1) | WP_308083996.1 | WP_308083996.1 protein kinase |
| 1(1) | WP_101195709.1 | WP_101195709.1 bacteriocin-processing peptidase family protein |
| 1(1) | WP_062239852.1 | WP_062239852.1 hypothetical protein |
| 1(1) | WP_243255986.1 | WP_243255986.1 TNT domain-containing protein |
| 1(1) | WP_098148062.1 | WP_098148062.1 WXG100 family type VII secretion target |
| 1(1) | WP_197241536.1 | WP_197241536.1 bacteriocin-processing peptidase family protein |
| 1(3) | WP_068940679.1 | WP_068940679.1 hypothetical protein |
| 1(1) | WP_058271109.1 | WP_058271109.1 FHA domain-containing protein |
| 1(1) | WP_145021744.1 | WP_145021744.1 hypothetical protein |
| 1(1) | WP_210087114.1 | WP_210087114.1 WXG100 family type VII secretion target |
| 1(1) | WP_097821803.1 | WP_097821803.1 bacteriocin-processing peptidase family protein |
| 1(1) | WP_305022923.1 | WP_305022923.1 WXG100 family type VII secretion target |
| 1(3) | WP_103626231.1 | WP_103626231.1 tetratricopeptide repeat protein |
| 1(4) | WP_185462582.1 | WP_185462582.1 type VII secretion protein EsaA |
| 1(1) | WP_215185472.1 | WP_215185472.1 EsaB/YukD family protein |
| 1(1) | WP_084156396.1 | WP_084156396.1 tetratricopeptide repeat protein |
| 1(1) | WP_090735424.1 | WP_090735424.1 hypothetical protein |
| 1(1) | WP_262725727.1 | WP_262725727.1 tetratricopeptide repeat protein |
| 1(1) | WP_199419770.1 | WP_199419770.1 WXG100 family type VII secretion target |
| 1(1) | WP_064019323.1 | WP_064019323.1 WG repeat-containing protein |
| 1(2) | WP_056700790.1 | WP_056700790.1 WXG100 family type VII secretion target |
| 1(1) | WP_251217311.1 | WP_251217311.1 type VII secretion protein EsaA |
| 1(1) | WP_247083042.1 | WP_247083042.1 hypothetical protein |
| 1(1) | WP_021291862.1 | WP_021291862.1 polysaccharide deacetylase family protein |
| 1(1) | WP_076103993.1 | WP_076103993.1 DUF2974 domain-containing protein |
| 1(1) | WP_221875375.1 | WP_221875375.1 WXG100 family type VII secretion target |
| 1(1) | WP_094876486.1 | WP_094876486.1 WXG100 family type VII secretion target |
| 1(1) | WP_142287547.1 | WP_142287547.1 deaminase domain-containing protein |
| 1(1) | WP_162848533.1 | WP_162848533.1 DUF5050 domain-containing protein |
| 1(1) | WP_216830407.1 | WP_216830407.1 S8 family serine peptidase |
| 1(1) | WP_291647970.1 | WP_291647970.1 DUF6382 domain-containing protein |
| 1(2) | WP_071640567.1 | WP_071640567.1 S-layer homology domain-containing protein |
| 1(1) | WP_231094846.1 | WP_231094846.1 WXG100 family type VII secretion target |
| 1(1) | WP_054402905.1 | WP_054402905.1 hypothetical protein |
| 1(1) | WP_144524122.1 | WP_144524122.1 hypothetical protein |

| | | |
|---|---|---|
| 1(50) | WP_012984597.1 | WP_012984597.1 WXG100 family type VII secretion target |
| 1(1) | WP_223068048.1 | WP_223068048.1 hypothetical protein |
| 1(1) | WP_137062801.1 | WP_137062801.1 hypothetical protein |
| 1(1) | WP_189018624.1 | WP_189018624.1 J domain-containing protein |
| 1(1) | WP_307394649.1 | WP_307394649.1 hypothetical protein |
| 1(2) | WP_195431328.1 | WP_195431328.1 hypothetical protein |
| 1(3) | WP_077852413.1 | WP_077852413.1 WXG100 family type VII secretion target |
| 1(2) | WP_228478842.1 | WP_228478842.1 hypothetical protein |
| 1(1) | WP_272541591.1 | WP_272541591.1 hypothetical protein |
| 1(1) | WP_225999409.1 | WP_225999409.1 WXG100 family type VII secretion target |
| 1(1) | WP_102710722.1 | WP_102710722.1 WXG100 family type VII secretion target |
| 1(1) | WP_195571409.1 | WP_195571409.1 AAA family ATPase |
| 1(1) | WP_189031030.1 | WP_189031030.1 WXG100 family type VII secretion target |
| 1(1) | WP_166245129.1 | WP_166245129.1 DUF6382 domain-containing protein |
| 1(1) | WP_000392226.1 | WP_000392226.1 hypothetical protein |
| 1(1) | WP_291682713.1 | WP_291682713.1 DUF6382 domain-containing protein |
| 1(1) | WP_088298122.1 | WP_088298122.1 tetratricopeptide repeat protein |
| 1(1) | WP_274485148.1 | WP_274485148.1 hypothetical protein |
| 1(1) | WP_013309986.1 | WP_013309986.1 hypothetical protein |
| 1(1) | WP_133243505.1 | WP_133243505.1 hypothetical protein |
| 1(1) | WP_042462799.1 | WP_042462799.1 pore-forming ESAT-6 family protein |
| 1(1) | WP_000713827.1 | WP_000713827.1 type VII secretion protein EsaA |
| 1(1) | WP_188535097.1 | WP_188535097.1 DUF2974 domain-containing protein |
| 1(13) | WP_076167864.1 | WP_076167864.1 WXG100 family type VII secretion target |
| 1(1) | WP_133378771.1 | WP_133378771.1 WXG100 family type VII secretion target |
| 1(1) | WP_223592660.1 | WP_223592660.1 hypothetical protein |
| 1(1) | WP_077893468.1 | WP_077893468.1 DUF6382 domain-containing protein |
| 1(1) | WP_098379498.1 | WP_098379498.1 type VII secretion protein EsaA |
| 1(1) | WP_071724730.1 | WP_071724730.1 WXG100 family type VII secretion target |
| 1(1) | WP_061805364.1 | WP_061805364.1 hypothetical protein |
| 1(2) | WP_003759563.1 | WP_003759563.1 EsaB/YukD family protein |
| 1(1) | WP_307589690.1 | WP_307589690.1 glycosyltransferase |
| 1(1) | WP_179160612.1 | WP_179160612.1 Ig-like domain repeat protein |
| 1(1) | WP_209365671.1 | WP_209365671.1 phosphoserine aminotransferase |
| 1(1) | WP_264459877.1 | WP_264459877.1 HNH/ENDO VII family nuclease |
| 1(1) | WP_087982601.1 | WP_087982601.1 bacteriocin-processing peptidase family protein |
| 1(1) | WP_264929205.1 | WP_264929205.1 hypothetical protein |
| 1(1) | WP_066419991.1 | WP_066419991.1 DUF6382 domain-containing protein |
| 1(1) | WP_291647972.1 | WP_291647972.1 hypothetical protein, partial |
| 1(1) | WP_096886736.1 | WP_096886736.1 hypothetical protein |
| 1(1) | WP_142318161.1 | WP_142318161.1 WXG100 family type VII secretion target |
| 1(2) | WP_141439358.1 | WP_141439358.1 hypothetical protein |
| 1(1) | WP_101219118.1 | WP_101219118.1 bacteriocin-processing peptidase family protein |
| 1(1) | WP_155703934.1 | WP_155703934.1 hypothetical protein |
| 1(1) | WP_262748109.1 | WP_262748109.1 hypothetical protein |
| 1(1) | WP_240522514.1 | WP_240522514.1 CdiA family toxin C-terminal domain-containing protein |
| 1(1) | WP_243495554.1 | WP_243495554.1 zinc-ribbon domain-containing protein |
| 1(1) | WP_128631288.1 | WP_128631288.1 WXG100 family type VII secretion target |
| 1(2) | WP_003268548.1 | WP_003268548.1 hypothetical protein |
| 1(1) | WP_306049011.1 | WP_306049011.1 hypothetical protein |
| 1(1) | WP_291647966.1 | WP_291647966.1 EsaB/YukD family protein |
| 1(1) | WP_007430134.1 | WP_007430134.1 hypothetical protein |
| 1(1) | WP_013309972.1 | WP_013309972.1 S-layer homology domain-containing protein |
| 1(1) | WP_098556212.1 | WP_098556212.1 hypothetical protein |
| 1(1) | WP_147786233.1 | WP_147786233.1 SseB family protein |
| 1(1) | WP_307589596.1 | WP_307589596.1 WXG100 family type VII secretion target |
| 1(1) | WP_000506916.1 | WP_000506916.1 tetratricopeptide repeat protein |
| 1(1) | WP_079694646.1 | WP_079694646.1 hypothetical protein |
| 1(1) | WP_153978068.1 | WP_153978068.1 hypothetical protein |
| 1(1) | WP_193653019.1 | WP_193653019.1 SseB family protein |
| 1(1) | WP_273383866.1 | WP_273383866.1 serine/threonine-protein kinase |
| 1(1) | WP_215591202.1 | WP_215591202.1 SseB family protein |
| 1(1) | WP_145318692.1 | WP_145318692.1 hypothetical protein |
| 1(1) | WP_307394660.1 | WP_307394660.1 pre-toxin TG domain-containing protein |
| 1(2) | WP_000392229.1 | WP_000392229.1 hypothetical protein |
| 1(1) | WP_006292340.1 | WP_006292340.1 hypothetical protein |
| 1(1) | WP_007430132.1 | WP_007430132.1 WXG100 family type VII secretion target |
| 1(1) | WP_014281183.1 | WP_014281183.1 WXG100 family type VII secretion target |
| 1(2) | WP_166245127.1 | WP_166245127.1 pore-forming ESAT-6 family protein |
| 1(3) | WP_013914194.1 | WP_013914194.1 WXG100 family type VII secretion target |
| 1(1) | WP_213649971.1 | WP_213649971.1 WXG100 family type VII secretion target |
| 1(1) | WP_087952382.1 | WP_087952382.1 DNA/RNA non-specific endonuclease |
| 1(1) | WP_166244097.1 | WP_166244097.1 copper amine oxidase N-terminal domain-containing protein |
| 1(1) | WP_245850665.1 | WP_245850665.1 DNA/RNA non-specific endonuclease |
| 1(1) | WP_248251187.1 | WP_248251187.1 hypothetical protein |
| 1(1) | WP_257126953.1 | WP_257126953.1 S-layer homology domain-containing protein |
| 1(1) | WP_163245179.1 | WP_163245179.1 hypothetical protein |
| 1(1) | WP_215601060.1 | WP_215601060.1 tetratricopeptide repeat protein |
| 1(1) | WP_002036277.1 | WP_002036277.1 hypothetical protein |
| 1(1) | WP_235699830.1 | WP_235699830.1 serine/threonine-protein kinase |
| 1(3) | WP_098266650.1 | WP_098266650.1 WXG100 family type VII secretion target |
| 1(1) | WP_053403779.1 | WP_053403779.1 hypothetical protein |
| 1(1) | WP_073008257.1 | WP_073008257.1 WXG100 family type VII secretion target |
| 1(1) | WP_144479049.1 | WP_144479049.1 Ig-like domain-containing protein |
| 1(1) | WP_154118030.1 | WP_154118030.1 hypothetical protein |
| 1(1) | WP_069355511.1 | WP_069355511.1 hypothetical protein |
| 1(1) | WP_263324546.1 | WP_263324546.1 WXG100 family type VII secretion target |
| 1(1) | WP_157418138.1 | WP_157418138.1 tetratricopeptide repeat protein |
| 1(1) | WP_028401639.1 | WP_028401639.1 hypothetical protein |
| 1(1) | WP_098603904.1 | WP_098603904.1 WXG100 family type VII secretion target |
| 1(1) | WP_047842538.1 | WP_047842538.1 membrane protein |
| 1(2) | WP_200372542.1 | WP_200372542.1 hypothetical protein |
| 1(1) | WP_307214308.1 | WP_307214308.1 hypothetical protein |
| 1(2) | WP_240992978.1 | WP_240992978.1 WXG100 family type VII secretion target |
| 1(1) | WP_211022785.1 | WP_211022785.1 copper amine oxidase N-terminal domain-containing protein |
| 1(1) | WP_013490124.1 | WP_013490124.1 pore-forming ESAT-6 family protein |
| 1(1) | WP_277504675.1 | WP_277504675.1 hypothetical protein |
| 1(1) | WP_229137197.1 | WP_229137197.1 hypothetical protein |
| 1(1) | WP_183600889.1 | WP_183600889.1 WXG100 family type VII secretion target |
| 1(2) | WP_191798668.1 | WP_191798668.1 hypothetical protein |
| 1(1) | WP_005604633.1 | WP_005604633.1 pore-forming ESAT-6 family protein |
| 1(1) | WP_289475141.1 | WP_289475141.1 proline-rich domain-containing protein |

1(1) WP_139796091.1 WP_139796091.1 hypothetical protein, partial
1(1) WP_294953147.1 WP_294953147.1 hypothetical protein
1(1) WP_138322084.1 WP_138322084.1 hypothetical protein
1(1) WP_189031034.1 WP_189031034.1 DUF5050 domain-containing protein
1(1) WP_066020843.1 WP_066020843.1 WXG100 family type VII secretion target
1(1) WP_089522637.1 WP_089522637.1 hypothetical protein
1(1) WP_010963738.1 WP_010963738.1 tetratricopeptide repeat protein
1(1) WP_069193883.1 WP_069193883.1 hypothetical protein
1(1) WP_270696284.1 WP_270696284.1 hypothetical protein
1(1) WP_239617475.1 WP_239617475.1 hypothetical protein
1(1) WP_291672208.1 WP_291672208.1 FHA domain-containing protein
1(1) WP_055105802.1 WP_055105802.1 copper amine oxidase N-terminal domain-containing protein
1(1) WP_199795348.1 WP_199795348.1 bacteriocin-processing peptidase family protein
1(1) WP_257443662.1 WP_257443662.1 copper amine oxidase N-terminal domain-containing protein
1(3) WP_195431287.1 WP_195431287.1 WXG100 family type VII secretion target
1(1) WP_307349975.1 WP_307349975.1 hypothetical protein
1(1) WP_212695786.1 WP_212695786.1 glycohydrolase toxin TNT-related protein
1(4) WP_143801063.1 WP_143801063.1 hypothetical protein
1(1) WP_090887130.1 WP_090887130.1 hypothetical protein
1(2) WP_098360834.1 WP_098360834.1 tetratricopeptide repeat protein
1(1) WP_101195715.1 WP_101195715.1 WXG100 family type VII secretion target
1(1) WP_149871678.1 WP_149871678.1 FHA domain-containing protein
1(1) WP_238187885.1 WP_238187885.1 hypothetical protein
1(1) WP_136604813.1 WP_136604813.1 WXG100 family type VII secretion target
1(1) WP_102371672.1 WP_102371672.1 FHA domain-containing protein
1(1) WP_204517516.1 WP_204517516.1 hypothetical protein
1(1) WP_140969552.1 WP_140969552.1 tetratricopeptide repeat protein
1(1) WP_268058022.1 WP_268058022.1 DUF6382 domain-containing protein
1(1) WP_291647942.1 WP_291647942.1 WXG100 family type VII secretion target
1(1) WP_252188914.1 WP_252188914.1 hypothetical protein
1(1) WP_179603479.1 WP_179603479.1 DUF2974 domain-containing protein
1(1) WP_253411078.1 WP_253411078.1 WXG100 family type VII secretion target
1(1) WP_257675237.1 WP_257675237.1 serine/threonine-protein kinase
1(1) WP_274485136.1 WP_274485136.1 S-layer homology domain-containing protein
1(2) WP_003728976.1 WP_003728976.1 WXG100 family type VII secretion target
1(1) WP_035286447.1 WP_035286447.1 hypothetical protein
1(1) WP_308083989.1 WP_308083989.1 DUF2974 domain-containing protein
1(1) WP_268619937.1 WP_268619937.1 hypothetical protein
1(1) WP_054972091.1 WP_054972091.1 DUF2974 domain-containing protein
1(1) WP_307312561.1 WP_307312561.1 FHA domain-containing protein
1(1) WP_277142092.1 WP_277142092.1 WXG100 family type VII secretion target
1(1) WP_149096535.1 WP_149096535.1 WXG100 family type VII secretion target
1(6) WP_185340294.1 WP_185340294.1 type VII secretion protein EsaA
1(1) WP_090239232.1 WP_090239232.1 hypothetical protein
1(1) WP_276867629.1 WP_276867629.1 T7SS effector LXG polymorphic toxin
1(2) WP_281813092.1 WP_281813092.1 hypothetical protein
1(7) WP_061975586.1 WP_061975586.1 WXG100 family type VII secretion target
1(1) WP_149190450.1 WP_149190450.1 tetratricopeptide repeat protein
1(1) WP_110931547.1 WP_110931547.1 WXG100 family type VII secretion target
1(1) WP_033521276.1 WP_033521276.1 serine/threonine-protein kinase
1(1) WP_071719991.1 WP_071719991.1 bacteriocin-processing peptidase family protein
1(80) WP_000807831.1 WP_000807831.1 WXG100 family type VII secretion target
1(1) WP_231417866.1 WP_231417866.1 pore-forming ESAT-6 family protein
1(1) WP_217035857.1 WP_217035857.1 DUF2974 domain-containing protein
1(1) WP_290407038.1 WP_290407038.1 C39 family peptidase
1(1) WP_066419989.1 WP_066419989.1 WXG100 family type VII secretion target
1(1) WP_307399103.1 WP_307399103.1 tetratricopeptide repeat protein
1(1) WP_145406468.1 WP_145406468.1 ParB/Srx family N-terminal domain-containing protein
1(2) WP_057897627.1 WP_057897627.1 WXG100 family type VII secretion target
1(1) WP_127570429.1 WP_127570429.1 hypothetical protein
1(1) WP_209861643.1 WP_209861643.1 WXG100 family type VII secretion target
1(1) WP_055109371.1 WP_055109371.1 hypothetical protein
1(1) WP_121610596.1 WP_121610596.1 DUF2974 domain-containing protein
1(1) WP_301879145.1 WP_301879145.1 hypothetical protein
1(1) WP_151630603.1 WP_151630603.1 hypothetical protein
1(1) WP_208752247.1 WP_208752247.1 bacteriocin-processing peptidase family protein, partial
1(1) WP_237475773.1 WP_237475773.1 DNA/RNA non-specific endonuclease
1(3) WP_051590913.1 WP_051590913.1 pore-forming ESAT-6 family protein
1(1) WP_021261880.1 WP_021261880.1 hypothetical protein
1(1) WP_306077382.1 WP_306077382.1 Ig-like domain-containing protein
1(3) WP_048552313.1 WP_048552313.1 SseB family protein
1(1) WP_291288259.1 WP_291288259.1 serine/threonine-protein kinase
1(1) WP_000444462.1 WP_000444462.1 hypothetical protein
1(1) WP_066091981.1 WP_066091981.1 Ig-like domain-containing protein
1(1) WP_248565448.1 WP_248565448.1 hypothetical protein
1(1) WP_240461204.1 WP_240461204.1 type VII secretion protein EsaA
1(1) WP_211745830.1 WP_211745830.1 WXG100 family type VII secretion target
1(1) WP_263880216.1 WP_263880216.1 hypothetical protein
1(2) WP_219872455.1 WP_219872455.1 hypothetical protein
1(31) WP_000807833.1 WP_000807833.1 WXG100 family type VII secretion target
1(1) WP_212330882.1 WP_212330882.1 hypothetical protein
1(1) WP_163872763.1 WP_163872763.1 hypothetical protein
1(1) WP_215032112.1 WP_215032112.1 serine/threonine-protein kinase
1(1) WP_265408842.1 WP_265408842.1 WXG100 family type VII secretion target
1(1) WP_245864026.1 WP_245864026.1 Ig-like domain-containing protein
1(1) WP_265408846.1 WP_265408846.1 hypothetical protein
1(2) WP_194333107.1 WP_194333107.1 HNH endonuclease
1(1) WP_204412734.1 WP_204412734.1 beta-N-acetylhexosaminidase
1(1) WP_010250266.1 WP_010250266.1 hypothetical protein
1(1) WP_195571386.1 WP_195571386.1 WXG100 family type VII secretion target
1(2) WP_026593077.1 WP_026593077.1 tetratricopeptide repeat protein
1(1) WP_257808050.1 WP_257808050.1 WXG100 family type VII secretion target
1(1) WP_198053925.1 WP_198053925.1 EsaB/YukD family protein
1(1) WP_055105803.1 WP_055105803.1 copper amine oxidase N-terminal domain-containing protein
1(1) WP_137698099.1 WP_137698099.1 copper amine oxidase N-terminal domain-containing protein
1(1) WP_249897315.1 WP_249897315.1 DUF5050 domain-containing protein
1(1) WP_021261890.1 WP_021261890.1 YHYH domain-containing protein
1(3) WP_123067115.1 WP_123067115.1 WXG100 family type VII secretion target
1(1) WP_184183005.1 WP_184183005.1 EsaB/YukD family protein
1(1) WP_088032790.1 WP_088032790.1 WXG100 family type VII secretion target

1(1) WP_099361341.1 WP_099361341.1 serine/threonine-protein kinase
1(1) WP_213506888.1 WP_213506888.1 hypothetical protein
1(1) WP_013658707.1 WP_013658707.1 copper amine oxidase N-terminal domain-containing protein
1(1) WP_087442409.1 WP_087442409.1 hypothetical protein
1(1) WP_017149770.1 WP_017149770.1 WXG100 family type VII secretion target
1(1) WP_000883809.1 WP_000883809.1 bacteriocin-processing peptidase family protein
1(1) WP_251542625.1 WP_251542625.1 Ig-like domain-containing protein
1(1) WP_213618815.1 WP_213618815.1 WXG100 family type VII secretion target
1(1) WP_228467811.1 WP_228467811.1 DNA/RNA non-specific endonuclease
1(1) WP_215598789.1 WP_215598789.1 hypothetical protein
1(1) WP_114298768.1 WP_114298768.1 S-layer homology domain-containing protein
1(1) WP_101668485.1 WP_101668485.1 WG repeat-containing protein
1(3) WP_077832294.1 WP_077832294.1 WXG100 family type VII secretion target
1(1) WP_244723766.1 WP_244723766.1 copper amine oxidase N-terminal domain-containing protein
1(1) WP_229139922.1 WP_229139922.1 WXG100 family type VII secretion target
1(1) WP_236417209.1 WP_236417209.1 hypothetical protein
1(1) WP_307738712.1 WP_307738712.1 Stk1 family PASTA domain-containing Ser/Thr kinase
1(1) WP_169505041.1 WP_169505041.1 S-layer homology domain-containing protein
1(1) WP_098248516.1 WP_098248516.1 zinc-ribbon domain-containing protein
1(1) WP_003336002.1 WP_003336002.1 hypothetical protein
1(1) WP_126665666.1 WP_126665666.1 hypothetical protein
1(1) WP_137014714.1 WP_137014714.1 SseB family protein
1(1) WP_142306815.1 WP_142306815.1 WXG100 family type VII secretion target, partial
1(1) WP_290407041.1 WP_290407041.1 copper amine oxidase N-terminal domain-containing protein
1(1) WP_212981557.1 WP_212981557.1 WXG100 family type VII secretion target
1(1) WP_215598794.1 WP_215598794.1 C39 family peptidase
1(1) WP_077569025.1 WP_077569025.1 hypothetical protein
1(1) WP_165175592.1 WP_165175592.1 S-layer homology domain-containing protein
1(1) WP_213117486.1 WP_213117486.1 FHA domain-containing protein
1(1) WP_088078460.1 WP_088078460.1 tetratricopeptide repeat protein
1(1) WP_027088426.1 WP_027088426.1 pore-forming ESAT-6 family protein
1(1) WP_234121610.1 WP_234121610.1 FHA domain-containing protein
1(1) WP_307878873.1 WP_307878873.1 tetratricopeptide repeat protein
1(2) WP_076249774.1 WP_076249774.1 hypothetical protein
1(1) WP_136479366.1 WP_136479366.1 ParA family protein
1(1) WP_308082243.1 WP_308082243.1 WXG100 family type VII secretion target
1(1) WP_111147803.1 WP_111147803.1 WXG100 family type VII secretion target
1(1) WP_068619093.1 WP_068619093.1 glycosyl hydrolase family 18 protein
1(1) WP_068496253.1 WP_068496253.1 hypothetical protein
1(4) WP_103626226.1 WP_103626226.1 bacteriocin-processing peptidase family protein
1(1) WP_139356010.1 WP_139356010.1 glycoside hydrolase family 3 C-terminal domain-containing protein
1(1) WP_141558533.1 WP_141558533.1 hypothetical protein
1(1) WP_077619656.1 WP_077619656.1 hypothetical protein
1(1) WP_127199780.1 WP_127199780.1 YhgE/Pip domain-containing protein
1(1) WP_268619934.1 WP_268619934.1 hypothetical protein
1(3) WP_077833102.1 WP_077833102.1 tetratricopeptide repeat protein
1(1) WP_189031051.1 WP_189031051.1 serine hydrolase
1(1) WP_024630327.1 WP_024630327.1 hypothetical protein
1(2) WP_071734040.1 WP_071734040.1 CdiA family toxin C-terminal domain-containing protein
1(1) WP_039306372.1 WP_039306372.1 protein kinase
1(1) WP_098673108.1 WP_098673108.1 bacteriocin-processing peptidase family protein
1(1) WP_166704994.1 WP_166704994.1 tetratricopeptide repeat protein
1(1) WP_055109376.1 WP_055109376.1 WXG100 family type VII secretion target
1(1) WP_257443641.1 WP_257443641.1 hypothetical protein
1(1) WP_264387894.1 WP_264387894.1 hypothetical protein
1(2) WP_066419981.1 WP_066419981.1 WXG100 family type VII secretion target
1(2) WP_097898676.1 WP_097898676.1 hypothetical protein
1(1) WP_251481001.1 WP_251481001.1 DUF6382 domain-containing protein
1(1) WP_054958406.1 WP_054958406.1 hypothetical protein
1(1) WP_089965946.1 WP_089965946.1 DUF2974 domain-containing protein
1(1) WP_074533067.1 WP_074533067.1 tetratricopeptide repeat protein
1(1) WP_211082925.1 WP_211082925.1 protein kinase family protein
1(1) WP_088298128.1 WP_088298128.1 bacteriocin-processing peptidase family protein
1(1) WP_293755531.1 WP_293755531.1 hypothetical protein, partial
1(1) WP_116556146.1 WP_116556146.1 WXG100 family type VII secretion target
1(1) WP_277156192.1 WP_277156192.1 leucine-rich repeat domain-containing protein
1(1) WP_038245452.1 WP_038245452.1 type VII secretion protein EsaA
1(2) WP_194336456.1 WP_194336456.1 pre-toxin TG domain-containing protein
1(1) WP_113672701.1 WP_113672701.1 Fic family protein
1(6) WP_228467084.1 WP_228467084.1 hypothetical protein
1(1) WP_042462817.1 WP_042462817.1 serine/threonine protein kinase
1(1) WP_151630604.1 WP_151630604.1 tetratricopeptide repeat protein
1(1) WP_205635662.1 WP_205635662.1 hypothetical protein
1(1) WP_251375549.1 WP_251375549.1 WXG100 family type VII secretion target
1(1) WP_144611189.1 WP_144611189.1 tetratricopeptide repeat protein
1(1) WP_294386448.1 WP_294386448.1 pore-forming ESAT-6 family protein
1(1) WP_212981555.1 WP_212981555.1 pre-toxin TG domain-containing protein
1(1) WP_185436426.1 WP_185436426.1 type VII secretion protein EsaA
1(1) WP_307388956.1 WP_307388956.1 DUF2974 domain-containing protein
1(4) WP_000506930.1 WP_000506930.1 tetratricopeptide repeat protein
1(1) WP_042462796.1 WP_042462796.1 FHA domain-containing protein
1(1) WP_022586772.1 WP_022586772.1 WG repeat-containing protein
1(1) WP_097889498.1 WP_097889498.1 hypothetical protein
1(8) WP_185463135.1 WP_185463135.1 glycohydrolase toxin TNT-related protein
1(2) WP_044355259.1 WP_044355259.1 hypothetical protein
1(1) WP_072336492.1 WP_072336492.1 WXG100 family type VII secretion target
1(1) WP_010250272.1 WP_010250272.1 hypothetical protein
1(2) WP_010963737.1 WP_010963737.1 tetratricopeptide repeat protein
1(1) WP_307312572.1 WP_307312572.1 Ig-like domain-containing protein
1(1) WP_110892968.1 WP_110892968.1 hypothetical protein
1(2) WP_270995163.1 WP_270995163.1 hypothetical protein
1(3) WP_097820481.1 WP_097820481.1 T7SS effector LXG polymorphic toxin
1(1) WP_126996317.1 WP_126996317.1 copper amine oxidase N-terminal domain-containing protein
1(1) WP_049107938.1 WP_049107938.1 T7SS effector LXG polymorphic toxin
1(1) WP_019424144.1 WP_019424144.1 WXG100 family type VII secretion target
1(1) WP_208019352.1 WP_208019352.1 hypothetical protein
1(1) WP_234414304.1 WP_234414304.1 EndoU domain-containing protein
1(2) WP_003516573.1 WP_003516573.1 DUF5050 domain-containing protein
1(1) WP_235286203.1 WP_235286203.1 DUF2974 domain-containing protein

| | | |
|---|---|---|
| 1(1) | WP_055109378.1 | WP_055109378.1 WXG100 family type VII secretion target |
| 1(1) | WP_283892187.1 | WP_283892187.1 hypothetical protein |
| 1(1) | WP_179603476.1 | WP_179603476.1 FHA domain-containing protein |
| 1(2) | WP_208019349.1 | WP_208019349.1 EsaB/YukD family protein |
| 1(1) | WP_023957723.1 | WP_023957723.1 hypothetical protein |
| 1(1) | WP_207706234.1 | WP_207706234.1 hypothetical protein |
| 1(12) | WP_074094613.1 | WP_074094613.1 hypothetical protein |
| 1(1) | WP_307828226.1 | WP_307828226.1 AHH domain-containing protein |
| 1(1) | WP_216390032.1 | WP_216390032.1 EsaB/YukD family protein |
| 1(2) | WP_216518275.1 | WP_216518275.1 hypothetical protein |
| 1(1) | WP_108464091.1 | WP_108464091.1 copper amine oxidase N-terminal domain-containing protein |
| 1(1) | WP_006720321.1 | WP_006720321.1 FHA domain-containing protein |
| 1(1) | WP_149871673.1 | WP_149871673.1 VWA domain-containing protein |
| 1(1) | WP_122971535.1 | WP_122971535.1 DUF2974 domain-containing protein |
| 1(1) | WP_102372749.1 | WP_102372749.1 hypothetical protein |
| 1(1) | WP_189340002.1 | WP_189340002.1 hypothetical protein |
| 1(2) | WP_240463837.1 | WP_240463837.1 WXG100 family type VII secretion target |
| 1(1) | WP_180235365.1 | WP_180235365.1 hypothetical protein |
| 1(1) | WP_034675378.1 | WP_034675378.1 protein kinase |
| 1(1) | WP_143315254.1 | WP_143315254.1 DUF6382 domain-containing protein |
| 1(1) | WP_302579553.1 | WP_302579553.1 hypothetical protein |
| 1(6) | WP_036684488.1 | WP_036684488.1 DUF5050 domain-containing protein |
| 1(1) | WP_058829516.1 | WP_058829516.1 copper amine oxidase N-terminal domain-containing protein |
| 1(1) | WP_257443628.1 | WP_257443628.1 DUF2974 domain-containing protein |
| 1(1) | WP_159081751.1 | WP_159081751.1 copper amine oxidase N-terminal domain-containing protein |
| 1(1) | WP_270699381.1 | WP_270699381.1 hypothetical protein |
| 1(1) | WP_110899381.1 | WP_110899381.1 hypothetical protein |
| 1(1) | WP_183600890.1 | WP_183600890.1 glycine zipper domain-containing protein |
| 1(1) | WP_098948842.1 | WP_098948842.1 hypothetical protein |
| 1(1) | WP_179034786.1 | WP_179034786.1 EsaB/YukD family protein |
| 1(1) | WP_127199771.1 | WP_127199771.1 WXG100 family type VII secretion target |
| 1(1) | WP_200483410.1 | WP_200483410.1 hypothetical protein |
| 1(1) | WP_290407039.1 | WP_290407039.1 copper amine oxidase N-terminal domain-containing protein |
| 1(1) | WP_150364197.1 | WP_150364197.1 WXG100 family type VII secretion target |
| 1(1) | WP_062326149.1 | WP_062326149.1 hypothetical protein |
| 1(1) | WP_204445234.1 | WP_204445234.1 chitobiase/beta-hexosaminidase C-terminal domain-containing protein |
| 1(1) | WP_082187568.1 | WP_082187568.1 hypothetical protein |
| 1(6) | WP_090976429.1 | WP_090976429.1 tetratricopeptide repeat protein |
| 1(1) | WP_090761825.1 | WP_090761825.1 WXG100 family type VII secretion target |
| 1(1) | WP_000444463.1 | WP_000444463.1 DNA/RNA non-specific endonuclease |
| 1(3) | WP_098139259.1 | WP_098139259.1 T7SS effector LXG polymorphic toxin |
| 1(1) | WP_165867473.1 | WP_165867473.1 WXG100 family type VII secretion target |
| 1(2) | WP_129478243.1 | WP_129478243.1 hypothetical protein |
| 1(1) | WP_229136361.1 | WP_229136361.1 hypothetical protein |
| 1(1) | WP_104148793.1 | WP_104148793.1 hypothetical protein |
| 1(2) | WP_095398791.1 | WP_095398791.1 hypothetical protein |
| 1(3) | WP_098361432.1 | WP_098361432.1 SseB family protein |
| 1(1) | WP_058271105.1 | WP_058271105.1 hypothetical protein |
| 1(3) | WP_098266651.1 | WP_098266651.1 tetratricopeptide repeat protein |
| 1(1) | WP_251596694.1 | WP_251596694.1 hypothetical protein |
| 1(1) | WP_077832295.1 | WP_077832295.1 hypothetical protein |
| 1(1) | WP_103655004.1 | WP_103655004.1 bacteriocin-processing peptidase family protein |
| 1(1) | WP_308082254.1 | WP_308082254.1 Ig-like domain-containing protein |
| 1(1) | WP_206098980.1 | WP_206098980.1 copper amine oxidase N-terminal domain-containing protein |
| 1(1) | WP_251610752.1 | WP_251610752.1 hypothetical protein |
| 1(1) | WP_184183059.1 | WP_184183059.1 hypothetical protein |
| 1(1) | WP_139316657.1 | WP_139316657.1 hypothetical protein |
| 1(1) | WP_236804414.1 | WP_236804414.1 bacteriocin-processing peptidase family protein |
| 1(4) | WP_103626232.1 | WP_103626232.1 SseB family protein |
| 1(1) | WP_255443974.1 | WP_255443974.1 WXG100 family type VII secretion target |
| 1(1) | WP_165820923.1 | WP_165820923.1 hypothetical protein |
| 1(1) | WP_220775544.1 | WP_220775544.1 tetratricopeptide repeat protein |
| 1(1) | WP_251598145.1 | WP_251598145.1 DUF2974 domain-containing protein |
| 1(1) | WP_273321550.1 | WP_273321550.1 hypothetical protein |
| 1(1) | WP_079506388.1 | WP_079506388.1 hypothetical protein |
| 1(2) | WP_271839910.1 | WP_271839910.1 EsaB/YukD family protein |
| 1(1) | WP_268745614.1 | WP_268745614.1 EsaB/YukD family protein |
| 1(1) | WP_289355904.1 | WP_289355904.1 hypothetical protein |
| 1(2) | WP_213592863.1 | WP_213592863.1 WXG100 family type VII secretion target |
| 1(2) | WP_098204202.1 | WP_098204202.1 tetratricopeptide repeat protein |
| 1(1) | WP_225639697.1 | WP_225639697.1 AHH domain-containing protein |
| 1(1) | WP_079506404.1 | WP_079506404.1 FHA domain-containing protein |
| 1(1) | WP_263560235.1 | WP_263560235.1 hypothetical protein |
| 1(2) | WP_214169379.1 | WP_214169379.1 hypothetical protein |
| 1(1) | WP_088550049.1 | WP_088550049.1 AAA family ATPase |
| 1(1) | WP_197286653.1 | WP_197286653.1 WXG100 family type VII secretion target |
| 1(1) | WP_066022455.1 | WP_066022455.1 WXG100 family type VII secretion target |
| 1(1) | WP_301290465.1 | WP_301290465.1 RHS repeat-associated core domain-containing protein |
| 1(1) | WP_000713834.1 | WP_000713834.1 type VII secretion protein EsaA, partial |
| 1(1) | WP_015736241.1 | WP_015736241.1 serine/threonine protein kinase |
| 1(1) | WP_251480999.1 | WP_251480999.1 WXG100 family type VII secretion target |
| 1(1) | WP_113753093.1 | WP_113753093.1 hypothetical protein |
| 1(1) | WP_098961907.1 | WP_098961907.1 bacteriocin-processing peptidase family protein |
| 1(1) | WP_256680794.1 | WP_256680794.1 hypothetical protein |
| 1(1) | WP_211022779.1 | WP_211022779.1 DL-endopeptidase inhibitor IseA family protein |
| 1(1) | WP_143315240.1 | WP_143315240.1 hypothetical protein |
| 1(1) | WP_170880286.1 | WP_170880286.1 AAA family ATPase |
| 1(1) | WP_140168583.1 | WP_140168583.1 WXG100 family type VII secretion target, partial |
| 1(1) | WP_214328827.1 | WP_214328827.1 bacteriocin-processing peptidase family protein |
| 1(1) | WP_090688955.1 | WP_090688955.1 EsaB/YukD family protein |
| 1(1) | WP_076271932.1 | WP_076271932.1 AAA family ATPase |
| 1(1) | WP_283910923.1 | WP_283910923.1 DUF5050 domain-containing protein |
| 1(1) | WP_110931542.1 | WP_110931542.1 WXG100 family type VII secretion target |
| 1(1) | WP_112331579.1 | WP_112331579.1 WXG100 family type VII secretion target |
| 1(1) | WP_126996307.1 | WP_126996307.1 EsaB/YukD family protein |
| 1(1) | WP_255260041.1 | WP_255260041.1 pre-toxin TG domain-containing protein |
| 1(1) | WP_122971541.1 | WP_122971541.1 FHA domain-containing protein |
| 1(1) | WP_275175594.1 | WP_275175594.1 pre-toxin TG domain-containing protein |

| | | |
|---|---|---|
| 1(1) | WP_291289939.1 | WP_291289939.1 hypothetical protein |
| 1(1) | WP_134910020.1 | WP_134910020.1 S-layer homology domain-containing protein |
| 1(1) | WP_318245835.1 | WP_318245835.1 S8 family serine peptidase |
| 1(1) | WP_199613896.1 | WP_199613896.1 copper amine oxidase N-terminal domain-containing protein |
| 1(2) | WP_043067145.1 | WP_043067145.1 hypothetical protein |
| 1(1) | WP_262729431.1 | WP_262729431.1 tetratricopeptide repeat protein |
| 1(1) | WP_277142096.1 | WP_277142096.1 hypothetical protein |
| 1(1) | WP_076156851.1 | WP_076156851.1 hypothetical protein |
| 1(2) | WP_158326750.1 | WP_158326750.1 hypothetical protein |
| 1(1) | WP_301167506.1 | WP_301167506.1 hypothetical protein |
| 1(1) | WP_244294691.1 | WP_244294691.1 DNA/RNA non-specific endonuclease |
| 1(1) | WP_013658713.1 | WP_013658713.1 DUF5050 domain-containing protein |
| 1(1) | WP_088550041.1 | WP_088550041.1 WXG100 family type VII secretion target |
| 1(1) | WP_216830389.1 | WP_216830389.1 pore-forming ESAT-6 family protein |
| 1(1) | WP_098945330.1 | WP_098945330.1 tetratricopeptide repeat protein |
| 1(1) | WP_100910320.1 | WP_100910320.1 hypothetical protein |
| 1(1) | WP_271839944.1 | WP_271839944.1 WXG100 family type VII secretion target |
| 1(1) | WP_242303038.1 | WP_242303038.1 hypothetical protein |
| 1(2) | WP_071724723.1 | WP_071724723.1 hypothetical protein |
| 1(1) | WP_163255259.1 | WP_163255259.1 WXG100 family type VII secretion target |
| 1(1) | WP_010250256.1 | WP_010250256.1 WXG100 family type VII secretion target |
| 1(1) | WP_268599728.1 | WP_268599728.1 hypothetical protein |
| 1(1) | WP_283927240.1 | WP_283927240.1 VWA domain-containing protein |
| 1(1) | WP_217035862.1 | WP_217035862.1 WXG100 family type VII secretion target |
| 1(1) | WP_244723791.1 | WP_244723791.1 DUF5050 domain-containing protein |
| 1(3) | WP_000034622.1 | WP_000034622.1 T7SS effector LXG polymorphic toxin |
| 1(1) | WP_242264959.1 | WP_242264959.1 tetratricopeptide repeat protein |
| 1(1) | WP_268613678.1 | WP_268613678.1 hypothetical protein |
| 1(1) | WP_192266456.1 | WP_192266456.1 WXG100 family type VII secretion target |
| 1(25) | WP_001050360.1 | WP_001050360.1 SseB family protein |
| 1(1) | WP_013309999.1 | WP_013309999.1 copper amine oxidase N-terminal domain-containing protein |
| 1(1) | WP_206094036.1 | WP_206094036.1 hypothetical protein |
| 1(1) | WP_229139915.1 | WP_229139915.1 hypothetical protein |
| 1(1) | WP_002148297.1 | WP_002148297.1 polymorphic toxin type 15 domain-containing protein |
| 1(1) | WP_063188066.1 | WP_063188066.1 DUF2974 domain-containing protein |
| 1(1) | WP_164965496.1 | WP_164965496.1 WXG100 family type VII secretion target |
| 1(1) | WP_256556970.1 | WP_256556970.1 FHA domain-containing protein |
| 1(2) | WP_068782406.1 | WP_068782406.1 WXG100 family type VII secretion target |
| 1(1) | WP_305166260.1 | WP_305166260.1 hypothetical protein |
| 1(1) | WP_223067438.1 | WP_223067438.1 hypothetical protein |
| 1(1) | WP_017149769.1 | WP_017149769.1 tetratricopeptide repeat protein |
| 1(3) | WP_155609753.1 | WP_155609753.1 WXG100 family type VII secretion target |
| 1(1) | WP_111147815.1 | WP_111147815.1 glycoside hydrolase family 3 N-terminal domain-containing protein |
| 1(1) | WP_098776428.1 | WP_098776428.1 protein kinase |
| 1(1) | WP_150457759.1 | WP_150457759.1 copper amine oxidase N-terminal domain-containing protein |
| 1(1) | WP_216830386.1 | WP_216830386.1 serine/threonine-protein kinase |
| 1(2) | WP_025716876.1 | WP_025716876.1 WXG100 family type VII secretion target |
| 1(1) | WP_013246325.1 | WP_013246325.1 hypothetical protein |
| 1(1) | WP_106765957.1 | WP_106765957.1 WXG100 family type VII secretion target |
| 1(1) | WP_068496265.1 | WP_068496265.1 S-layer homology domain-containing protein |
| 1(1) | WP_212969035.1 | WP_212969035.1 hypothetical protein |
| 1(1) | WP_098226390.1 | WP_098226390.1 T7SS effector LXG polymorphic toxin |
| 1(1) | WP_098248514.1 | WP_098248514.1 zinc-ribbon domain-containing protein |
| 1(1) | WP_005604618.1 | WP_005604618.1 serine/threonine-protein kinase |
| 1(1) | WP_000506910.1 | WP_000506910.1 tetratricopeptide repeat protein |
| 1(1) | WP_251583292.1 | WP_251583292.1 hypothetical protein |
| 1(1) | WP_306077384.1 | WP_306077384.1 protein kinase |
| 1(1) | WP_120112256.1 | WP_120112256.1 pore-forming ESAT-6 family protein |
| 1(1) | WP_302878971.1 | WP_302878971.1 cadherin-like beta sandwich domain-containing protein |
| 1(1) | WP_223282660.1 | WP_223282660.1 WXG100 family type VII secretion target |
| 1(1) | WP_179283641.1 | WP_179283641.1 AAA family ATPase |
| 1(1) | WP_022586770.1 | WP_022586770.1 hypothetical protein |
| 1(1) | WP_179016427.1 | WP_179016427.1 RHS repeat-associated core domain-containing protein |
| 1(1) | WP_256992243.1 | WP_256992243.1 S-layer homology domain-containing protein |
| 1(1) | WP_066020835.1 | WP_066020835.1 tetratricopeptide repeat protein |
| 1(1) | WP_291672210.1 | WP_291672210.1 Ig-like domain-containing protein |
| 1(1) | WP_283910930.1 | WP_283910930.1 copper amine oxidase N-terminal domain-containing protein |
| 1(1) | WP_170839747.1 | WP_170839747.1 EsaB/YukD family protein |
| 1(1) | WP_319005631.1 | WP_319005631.1 protein kinase |
| 1(1) | WP_270364634.1 | WP_270364634.1 tetratricopeptide repeat protein |
| 1(1) | WP_076297591.1 | WP_076297591.1 DUF2974 domain-containing protein |
| 1(1) | WP_295628342.1 | WP_295628342.1 tetratricopeptide repeat protein |
| 1(1) | WP_156114750.1 | WP_156114750.1 hypothetical protein |
| 1(1) | WP_079506414.1 | WP_079506414.1 RHS repeat-associated core domain-containing protein |
| 1(2) | WP_107734208.1 | WP_107734208.1 hypothetical protein |
| 1(4) | WP_063219638.1 | WP_063219638.1 hypothetical protein |
| 1(1) | WP_019912152.1 | WP_019912152.1 hypothetical protein |
| 1(1) | WP_277175594.1 | WP_277175594.1 serine/threonine-protein kinase |
| 1(2) | WP_145329907.1 | WP_145329907.1 WXG100 family type VII secretion target |
| 1(1) | WP_094155406.1 | WP_094155406.1 hypothetical protein |
| 1(2) | WP_185469293.1 | WP_185469293.1 pre-toxin TG domain-containing protein |
| 1(1) | WP_142324454.1 | WP_142324454.1 hypothetical protein |
| 1(1) | WP_235801547.1 | WP_235801547.1 type VII secretion protein EsaA |
| 1(3) | WP_097961544.1 | WP_097961544.1 SseB family protein |
| 1(1) | WP_163853476.1 | WP_163853476.1 DUF2974 domain-containing protein |
| 1(1) | WP_160496930.1 | WP_160496930.1 WXG100 family type VII secretion target |
| 1(1) | WP_238655351.1 | WP_238655351.1 peptidoglycan-binding protein |
| 1(2) | WP_074085358.1 | WP_074085358.1 hypothetical protein |
| 1(1) | WP_204445204.1 | WP_204445204.1 pore-forming ESAT-6 family protein |
| 1(1) | WP_165097594.1 | WP_165097594.1 hypothetical protein |
| 1(1) | WP_237391783.1 | WP_237391783.1 hypothetical protein |
| 1(1) | WP_010250254.1 | WP_010250254.1 WXG100 family type VII secretion target |
| 1(1) | WP_272035323.1 | WP_272035323.1 hypothetical protein |
| 1(1) | WP_155703946.1 | WP_155703946.1 S-layer homology domain-containing protein |
| 1(1) | WP_270699383.1 | WP_270699383.1 WXG100 family type VII secretion target |
| 1(1) | WP_097911840.1 | WP_097911840.1 tetratricopeptide repeat protein |
| 1(2) | WP_010963728.1 | WP_010963728.1 FHA domain-containing protein |
| 1(1) | WP_242172371.1 | WP_242172371.1 tetratricopeptide repeat protein |
| 1(1) | WP_099223957.1 | WP_099223957.1 WXG100 family type VII secretion target |

1(1) WP_229522855.1 WP_229522855.1 hypothetical protein
1(1) WP_055109369.1 WP_055109369.1 copper amine oxidase N-terminal domain-containing protein
1(1) WP_142613000.1 WP_142613000.1 hypothetical protein
1(1) WP_244723793.1 WP_244723793.1 EsaB/YukD family protein
1(1) WP_163977054.1 WP_163977054.1 hypothetical protein
1(1) WP_213143947.1 WP_213143947.1 DUF2974 domain-containing protein
1(2) WP_197247507.1 WP_197247507.1 WG repeat-containing protein
1(1) WP_017689070.1 WP_017689070.1 hypothetical protein
1(1) WP_053783887.1 WP_053783887.1 hypothetical protein
1(1) WP_273383869.1 WP_273383869.1 hypothetical protein
1(1) WP_036671544.1 WP_036671544.1 membrane protein
1(1) WP_199925766.1 WP_199925766.1 copper amine oxidase N-terminal domain-containing protein
1(1) WP_307192890.1 WP_307192890.1 Ig-like domain-containing protein
1(1) WP_072336498.1 WP_072336498.1 hypothetical protein
1(1) WP_191272848.1 WP_191272848.1 Ig-like domain-containing protein
1(1) WP_304216869.1 WP_304216869.1 DUF2974 domain-containing protein
1(1) WP_142951721.1 WP_142951721.1 hypothetical protein
1(2) WP_085499094.1 WP_085499094.1 WXG100 family type VII secretion target
1(1) WP_273191248.1 WP_273191248.1 hypothetical protein
1(2) WP_112331583.1 WP_112331583.1 VWA domain-containing protein
1(1) WP_098250517.1 WP_098250517.1 bacteriocin-processing peptidase family protein
1(2) WP_185336682.1 WP_185336682.1 type VII secretion protein EsaA
1(1) WP_197937381.1 WP_197937381.1 hypothetical protein
1(1) WP_258113057.1 WP_258113057.1 type VII secretion protein EsaA
1(1) WP_125084778.1 WP_125084778.1 hypothetical protein
1(1) WP_066419987.1 WP_066419987.1 DUF2974 domain-containing protein
1(1) WP_102371675.1 WP_102371675.1 FHA domain-containing protein
1(1) WP_110931548.1 WP_110931548.1 WXG100 family type VII secretion target
1(1) WP_013309989.1 WP_013309989.1 WXG100 family type VII secretion target
1(1) WP_166244906.1 WP_166244906.1 WXG100 family type VII secretion target
1(1) WP_045518277.1 WP_045518277.1 protein kinase
1(1) WP_216390034.1 WP_216390034.1 DUF6382 domain-containing protein
1(1) WP_232284466.1 WP_232284466.1 Ig-like domain-containing protein
1(3) WP_033100017.1 WP_033100017.1 WXG100 family type VII secretion target
1(1) WP_256227720.1 WP_256227720.1 hypothetical protein
1(1) WP_165147497.1 WP_165147497.1 S-layer homology domain-containing protein
1(1) WP_006923820.1 WP_006923820.1 T7SS effector LXG polymorphic toxin
1(1) WP_226679102.1 WP_226679102.1 Mbeg1-like protein
1(4) WP_038406450.1 WP_038406450.1 GW domain-containing glycosaminoglycan-binding protein
1(1) WP_204412723.1 WP_204412723.1 DUF2974 domain-containing protein
1(1) WP_025685020.1 WP_025685020.1 S-layer homology domain-containing protein
1(2) WP_216518260.1 WP_216518260.1 WXG100 family type VII secretion target
1(1) WP_234813927.1 WP_234813927.1 hypothetical protein, partial
1(1) WP_176212751.1 WP_176212751.1 WXG100 family type VII secretion target
1(1) WP_106496016.1 WP_106496016.1 WXG100 family type VII secretion target
1(1) WP_242400481.1 WP_242400481.1 GH-E family nuclease
1(1) WP_305755037.1 WP_305755037.1 hypothetical protein
1(1) WP_036702401.1 WP_036702401.1 hypothetical protein, partial
1(1) WP_068782417.1 WP_068782417.1 S-layer homology domain-containing protein
1(1) WP_195542475.1 WP_195542475.1 Ig-like domain-containing protein
1(2) WP_066419982.1 WP_066419982.1 WXG100 family type VII secretion target
1(1) WP_012260836.1 WP_012260836.1 protein kinase
1(1) WP_065291246.1 WP_065291246.1 hypothetical protein
1(1) WP_244723780.1 WP_244723780.1 WXG100 family type VII secretion target
1(1) WP_214706007.1 WP_214706007.1 FHA domain-containing protein
1(1) WP_225999406.1 WP_225999406.1 hypothetical protein
1(1) WP_019912161.1 WP_019912161.1 hypothetical protein
1(1) WP_195974875.1 WP_195974875.1 chitobiase/beta-hexosaminidase C-terminal domain-containing protein
1(1) WP_276867643.1 WP_276867643.1 T7SS effector LXG polymorphic toxin
1(1) WP_213618812.1 WP_213618812.1 hypothetical protein
1(4) WP_064473932.1 WP_064473932.1 SseB family protein
1(1) WP_085499103.1 WP_085499103.1 EsaB/YukD family protein
1(1) WP_179160618.1 WP_179160618.1 WXG100 family type VII secretion target
1(1) WP_197225371.1 WP_197225371.1 tetratricopeptide repeat protein
1(2) WP_097952774.1 WP_097952774.1 SseB family protein
1(1) WP_098148051.1 WP_098148051.1 bacteriocin-processing peptidase family protein
1(1) WP_013658703.1 WP_013658703.1 WXG100 family type VII secretion target
1(1) WP_076219074.1 WP_076219074.1 hypothetical protein
1(1) WP_245693743.1 WP_245693743.1 DNA/RNA non-specific endonuclease
1(1) WP_044644803.1 WP_044644803.1 S-layer homology domain-containing protein
1(1) WP_307738733.1 WP_307738733.1 FHA domain-containing protein
1(1) WP_216531932.1 WP_216531932.1 hypothetical protein
1(1) WP_289475138.1 WP_289475138.1 tetratricopeptide repeat protein
1(1) WP_098256513.1 WP_098256513.1 bacteriocin-processing peptidase family protein
1(1) WP_227917891.1 WP_227917891.1 Ig-like domain-containing protein
1(1) WP_240379735.1 WP_240379735.1 SseB family protein
1(2) WP_010963733.1 WP_010963733.1 WXG100 family type VII secretion target
1(1) WP_257808055.1 WP_257808055.1 tetratricopeptide repeat protein
1(1) WP_243518759.1 WP_243518759.1 tetratricopeptide repeat protein
1(1) WP_045243838.1 WP_045243838.1 membrane protein
1(1) WP_098603959.1 WP_098603959.1 tetratricopeptide repeat protein
1(1) WP_270728497.1 WP_270728497.1 tetratricopeptide repeat protein
1(1) WP_282742026.1 WP_282742026.1 hypothetical protein
1(1) WP_193653020.1 WP_193653020.1 hypothetical protein
1(1) WP_144479466.1 WP_144479466.1 WXG100 family type VII secretion target
1(2) WP_053512321.1 WP_053512321.1 T7SS effector LXG polymorphic toxin
1(2) WP_068782401.1 WP_068782401.1 hypothetical protein
1(1) WP_076224118.1 WP_076224118.1 DUF2974 domain-containing protein
1(1) WP_249113668.1 WP_249113668.1 hypothetical protein
1(2) WP_063566321.1 WP_063566321.1 hypothetical protein
1(4) WP_064473935.1 WP_064473935.1 WXG100 family type VII secretion target
1(1) WP_179603474.1 WP_179603474.1 serine/threonine-protein kinase
1(1) WP_244161474.1 WP_244161474.1 type VII secretion protein EsaA
1(1) WP_244971330.1 WP_244971330.1 CdiA family toxin C-terminal domain-containing protein
1(1) WP_234121604.1 WP_234121604.1 pore-forming ESAT-6 family protein
1(1) WP_297183826.1 WP_297183826.1 DUF2974 domain-containing protein
1(1) WP_000506928.1 WP_000506928.1 tetratricopeptide repeat protein
1(1) WP_166245145.1 WP_166245145.1 vWA domain-containing protein
1(1) WP_252200283.1 WP_252200283.1 type VII secretion protein EsaA
1(1) WP_053361021.1 WP_053361021.1 WXG100 family type VII secretion target
1(1) WP_307589604.1 WP_307589604.1 hypothetical protein

| | | |
|---|---|---|
| 1(1) | WP_185165649.1 | WP_185165649.1 VWA domain-containing protein |
| 1(2) | WP_021878936.1 | WP_021878936.1 WXG100 family type VII secretion target |
| 1(2) | WP_076286997.1 | WP_076286997.1 hypothetical protein |
| 1(1) | WP_289342265.1 | WP_289342265.1 hypothetical protein |
| 1(1) | WP_272045577.1 | WP_272045577.1 hypothetical protein |
| 1(1) | WP_238655334.1 | WP_238655334.1 stalk domain-containing protein |
| 1(1) | WP_286458022.1 | WP_286458022.1 WXG100 family type VII secretion target |
| 1(1) | WP_274485154.1 | WP_274485154.1 hypothetical protein |
| 1(1) | WP_231094844.1 | WP_231094844.1 DUF6382 domain-containing protein |
| 1(1) | WP_280787396.1 | WP_280787396.1 DUF5050 domain-containing protein |
| 1(1) | WP_014368236.1 | WP_014368236.1 S-layer homology domain-containing protein |
| 1(1) | WP_002148317.1 | WP_002148317.1 hypothetical protein, partial |
| 1(2) | WP_048749324.1 | WP_048749324.1 EsaB/YukD family protein |
| 1(1) | WP_068782403.1 | WP_068782403.1 hypothetical protein |
| 1(1) | WP_098148059.1 | WP_098148059.1 hypothetical protein |
| 1(1) | WP_224839852.1 | WP_224839852.1 Ig-like domain-containing protein |
| 1(1) | WP_042125629.1 | WP_042125629.1 DUF2974 domain-containing protein |
| 1(3) | WP_000671527.1 | WP_000671527.1 polymorphic toxin type 15 domain-containing protein |
| 1(1) | WP_169505055.1 | WP_169505055.1 serine/threonine protein kinase |
| 1(1) | WP_100910324.1 | WP_100910324.1 bacteriocin-processing peptidase family protein |
| 1(1) | WP_243500461.1 | WP_243500461.1 hypothetical protein |
| 1(1) | WP_215572752.1 | WP_215572752.1 EsaB/YukD family protein |
| 1(1) | WP_098303175.1 | WP_098303175.1 WXG100 family type VII secretion target |
| 1(1) | WP_127570431.1 | WP_127570431.1 hypothetical protein |
| 1(1) | WP_257443624.1 | WP_257443624.1 copper amine oxidase N-terminal domain-containing protein |
| 1(1) | WP_145318696.1 | WP_145318696.1 WXG100 family type VII secretion target |
| 1(1) | WP_195571391.1 | WP_195571391.1 DUF5050 domain-containing protein |
| 1(1) | WP_289342263.1 | WP_289342263.1 DUF2974 domain-containing protein |
| 1(1) | WP_000506922.1 | WP_000506922.1 tetratricopeptide repeat protein |
| 1(4) | WP_080334524.1 | WP_080334524.1 WXG100 family type VII secretion target |
| 1(3) | WP_145406490.1 | WP_145406490.1 WXG100 family type VII secretion target |
| 1(1) | WP_113058716.1 | WP_113058716.1 hypothetical protein |
| 1(1) | WP_137698100.1 | WP_137698100.1 hypothetical protein |
| 1(1) | WP_145949000.1 | WP_145949000.1 WXG100 family type VII secretion target |
| 1(3) | WP_141499179.1 | WP_141499179.1 deaminase domain-containing protein |
| 1(1) | WP_197225372.1 | WP_197225372.1 hypothetical protein |
| 1(1) | WP_273061474.1 | WP_273061474.1 hypothetical protein |
| 1(1) | WP_212693233.1 | WP_212693233.1 WXG100 family type VII secretion target |
| 1(1) | WP_149871679.1 | WP_149871679.1 pore-forming ESAT-6 family protein |
| 1(1) | WP_279630442.1 | WP_279630442.1 RHS repeat-associated core domain-containing protein |
| 1(1) | WP_066020848.1 | WP_066020848.1 Ig-like domain-containing protein |
| 1(1) | WP_133332982.1 | WP_133332982.1 DUF6382 domain-containing protein |
| 1(1) | WP_227219209.1 | WP_227219209.1 hypothetical protein |
| 1(1) | WP_251596698.1 | WP_251596698.1 copper amine oxidase N-terminal domain-containing protein |
| 1(1) | WP_307738723.1 | WP_307738723.1 WXG100 family type VII secretion target |
| 1(1) | WP_306049030.1 | WP_306049030.1 DUF6382 domain-containing protein |
| 1(1) | WP_290407036.1 | WP_290407036.1 DL-endopeptidase inhibitor IseA family protein |
| 1(1) | WP_162616102.1 | WP_162616102.1 WXG100 family type VII secretion target |
| 1(1) | WP_090921052.1 | WP_090921052.1 hypothetical protein |
| 1(1) | WP_127570425.1 | WP_127570425.1 hypothetical protein |
| 1(1) | WP_283904345.1 | WP_283904345.1 hypothetical protein |
| 1(1) | WP_089965950.1 | WP_089965950.1 WXG100 family type VII secretion target |
| 1(2) | WP_185639020.1 | WP_185639020.1 WXG100 family type VII secretion target |
| 1(1) | WP_240379742.1 | WP_240379742.1 hypothetical protein |
| 1(1) | WP_279830800.1 | WP_279830800.1 hypothetical protein |
| 1(1) | WP_112331587.1 | WP_112331587.1 FHA domain-containing protein |
| 1(1) | WP_007083290.1 | WP_007083290.1 DUF2974 domain-containing protein |
| 1(1) | WP_068673700.1 | WP_068673700.1 hypothetical protein |
| 1(1) | WP_062239861.1 | WP_062239861.1 tetratricopeptide repeat protein |
| 1(1) | WP_268058014.1 | WP_268058014.1 hypothetical protein |
| 1(1) | WP_073008265.1 | WP_073008265.1 FHA domain-containing protein |
| 1(1) | WP_000404145.1 | WP_000404145.1 WXG100 family type VII secretion target |
| 1(1) | WP_241407463.1 | WP_241407463.1 hypothetical protein |
| 1(1) | WP_255263010.1 | WP_255263010.1 toxin |
| 1(2) | WP_037993169.1 | WP_037993169.1 WXG100 family type VII secretion target |
| 1(1) | WP_104495323.1 | WP_104495323.1 S-layer homology domain-containing protein |
| 1(1) | WP_307394621.1 | WP_307394621.1 hypothetical protein |
| 1(1) | WP_212693235.1 | WP_212693235.1 hypothetical protein |
| 1(1) | WP_213117482.1 | WP_213117482.1 DUF2974 domain-containing protein |
| 1(1) | WP_183600896.1 | WP_183600896.1 EsaB/YukD family protein |
| 1(1) | WP_209365673.1 | WP_209365673.1 hypothetical protein |
| 1(5) | WP_042125627.1 | WP_042125627.1 WXG100 family type VII secretion target |
| 1(1) | WP_157643666.1 | WP_157643666.1 hypothetical protein |
| 1(1) | WP_142543127.1 | WP_142543127.1 WXG100 family type VII secretion target |
| 1(1) | WP_100539445.1 | WP_100539445.1 hypothetical protein |
| 1(1) | WP_007147985.1 | WP_007147985.1 WXG100 family type VII secretion target |
| 1(1) | WP_282446361.1 | WP_282446361.1 RHS repeat-associated core domain-containing protein |
| 1(1) | WP_172460051.1 | WP_172460051.1 tetratricopeptide repeat protein |
| 1(1) | WP_138186691.1 | WP_138186691.1 hypothetical protein |
| 1(1) | WP_184183011.1 | WP_184183011.1 WXG100 family type VII secretion target |
| 1(1) | WP_258296408.1 | WP_258296408.1 copper amine oxidase N-terminal domain-containing protein |
| 1(1) | WP_211269593.1 | WP_211269593.1 lipopolysaccharide kinase InaA family protein |
| 1(1) | WP_242251758.1 | WP_242251758.1 bacteriocin-processing peptidase family protein |
| 1(1) | WP_179265138.1 | WP_179265138.1 hypothetical protein |
| 1(1) | WP_077833106.1 | WP_077833106.1 tetratricopeptide repeat protein |
| 1(1) | WP_095205826.1 | WP_095205826.1 DNA/RNA non-specific endonuclease |
| 1(1) | WP_216518254.1 | WP_216518254.1 hypothetical protein |
| 1(1) | WP_133059623.1 | WP_133059623.1 WXG100 family type VII secretion target |
| 1(1) | WP_087442414.1 | WP_087442414.1 WXG100 family type VII secretion target |
| 1(1) | WP_036944553.1 | WP_036944553.1 hypothetical protein |
| 1(1) | WP_098851260.1 | WP_098851260.1 hypothetical protein |
| 1(8) | WP_061402737.1 | WP_061402737.1 SseB family protein |
| 1(1) | WP_179160616.1 | WP_179160616.1 FHA domain-containing protein |
| 1(1) | WP_003287962.1 | WP_003287962.1 hypothetical protein |
| 1(1) | WP_158651616.1 | WP_158651616.1 DUF2974 domain-containing protein |
| 1(1) | WP_217035867.1 | WP_217035867.1 serine/threonine-protein kinase |
| 1(1) | WP_021725681.1 | WP_021725681.1 FHA domain-containing protein |
| 1(1) | WP_255264702.1 | WP_255264702.1 hypothetical protein |
| 1(1) | WP_098562453.1 | WP_098562453.1 SseB family protein |
| 1(1) | WP_170287043.1 | WP_170287043.1 hypothetical protein |
| 1(1) | WP_254759295.1 | WP_254759295.1 WXG100 family type VII secretion target |

| | | |
|---|---|---|
| 1(1) | WP_094936487.1 | WP_094936487.1 hypothetical protein |
| 1(1) | WP_224839858.1 | WP_224839858.1 hypothetical protein |
| 1(1) | WP_066022451.1 | WP_066022451.1 DUF2974 domain-containing protein |
| 1(1) | WP_129478239.1 | WP_129478239.1 tetratricopeptide repeat protein |
| 1(1) | WP_077832290.1 | WP_077832290.1 tetratricopeptide repeat protein |
| 1(1) | WP_051491340.1 | WP_051491340.1 copper amine oxidase N-terminal domain-containing protein |
| 1(1) | WP_063209142.1 | WP_063209142.1 WXG100 family type VII secretion target |
| 1(3) | WP_143801060.1 | WP_143801060.1 WXG100 family type VII secretion target |
| 1(1) | WP_066415462.1 | WP_066415462.1 Ig-like domain-containing protein |
| 1(1) | WP_255264701.1 | WP_255264701.1 hypothetical protein |
| 1(1) | WP_269055424.1 | WP_269055424.1 WXG100 family type VII secretion target |
| 1(1) | WP_226642868.1 | WP_226642868.1 LysM peptidoglycan-binding domain-containing protein |
| 1(1) | WP_069080931.1 | WP_069080931.1 hypothetical protein |
| 1(5) | WP_000506923.1 | WP_000506923.1 tetratricopeptide repeat protein |
| 1(1) | WP_256227719.1 | WP_256227719.1 tetratricopeptide repeat protein |
| 1(1) | WP_050753440.1 | WP_050753440.1 AAA family ATPase |
| 1(4) | WP_063479993.1 | WP_063479993.1 hypothetical protein |
| 1(1) | WP_120462944.1 | WP_120462944.1 hypothetical protein |
| 1(1) | WP_000360014.1 | WP_000360014.1 T7SS effector LXG polymorphic toxin |
| 1(1) | WP_053326559.1 | WP_053326559.1 WXG100 family type VII secretion target |
| 1(1) | WP_094878480.1 | WP_094878480.1 hypothetical protein |
| 1(1) | WP_258425839.1 | WP_258425839.1 WG repeat-containing protein |
| 1(1) | WP_063675262.1 | WP_063675262.1 EsaB/YukD family protein |
| 1(1) | WP_142613003.1 | WP_142613003.1 hypothetical protein |
| 1(1) | WP_240379729.1 | WP_240379729.1 bacteriocin-processing peptidase family protein |
| 1(1) | WP_132283080.1 | WP_132283080.1 AAA family ATPase |
| 1(1) | WP_186580070.1 | WP_186580070.1 hypothetical protein |
| 1(1) | WP_132008172.1 | WP_132008172.1 protein kinase |
| 1(1) | WP_024027642.1 | WP_024027642.1 VWA domain-containing protein |
| 1(1) | WP_044307210.1 | WP_044307210.1 T7SS effector LXG polymorphic toxin |
| 1(1) | WP_195515237.1 | WP_195515237.1 DUF2974 domain-containing protein |
| 1(1) | WP_165097628.1 | WP_165097628.1 DUF2974 domain-containing protein |
| 1(1) | WP_142306904.1 | WP_142306904.1 hypothetical protein |
| 1(1) | WP_152381395.1 | WP_152381395.1 hypothetical protein |
| 1(1) | WP_198310835.1 | WP_198310835.1 hypothetical protein |
| 1(1) | WP_285748172.1 | WP_285748172.1 hypothetical protein |
| 1(1) | WP_033521266.1 | WP_033521266.1 hypothetical protein |
| 1(1) | WP_036642163.1 | WP_036642163.1 serine/threonine protein kinase |
| 1(1) | WP_088032794.1 | WP_088032794.1 HNH/ENDO VII family nuclease |
| 1(1) | WP_285860874.1 | WP_285860874.1 DL-endopeptidase inhibitor IseA family protein |
| 1(7) | WP_162164278.1 | WP_162164278.1 EsaB/YukD family protein |
| 1(1) | WP_286017554.1 | WP_286017554.1 hypothetical protein |
| 1(1) | WP_251542616.1 | WP_251542616.1 FHA domain-containing protein |
| 1(1) | WP_077853215.1 | WP_077853215.1 DUF6382 domain-containing protein |
| 1(1) | WP_236417215.1 | WP_236417215.1 WXG100 family type VII secretion target |
| 1(1) | WP_159081753.1 | WP_159081753.1 peptidoglycan-binding protein |
| 1(1) | WP_141529617.1 | WP_141529617.1 CdiA family toxin C-terminal domain-containing protein |
| 1(1) | WP_243255989.1 | WP_243255989.1 hypothetical protein |
| 1(1) | WP_286124784.1 | WP_286124784.1 tetratricopeptide repeat protein |
| 1(35) | WP_000391141.1 | WP_000391141.1 hypothetical protein |
| 1(2) | WP_088063232.1 | WP_088063232.1 tetratricopeptide repeat protein |
| 1(1) | WP_042125605.1 | WP_042125605.1 AAA family ATPase |
| 1(1) | WP_307199944.1 | WP_307199944.1 WXG100 family type VII secretion target |
| 1(1) | WP_247902245.1 | WP_247902245.1 WXG100 family type VII secretion target |
| 1(1) | WP_141531119.1 | WP_141531119.1 hypothetical protein |
| 1(1) | WP_000392216.1 | WP_000392216.1 hypothetical protein |
| 1(2) | WP_185474155.1 | WP_185474155.1 pre-toxin TG domain-containing protein |
| 1(1) | WP_151255773.1 | WP_151255773.1 S-layer homology domain-containing protein |
| 1(2) | WP_061667206.1 | WP_061667206.1 T7SS effector LXG polymorphic toxin |
| 1(1) | WP_265444533.1 | WP_265444533.1 DUF5050 domain-containing protein |
| 1(4) | WP_007128628.1 | WP_007128628.1 hypothetical protein |
| 1(1) | WP_055109374.1 | WP_055109374.1 hypothetical protein |
| 1(1) | WP_024027636.1 | WP_024027636.1 serine/threonine-protein kinase |
| 1(1) | WP_155609757.1 | WP_155609757.1 hypothetical protein |
| 1(1) | WP_006720315.1 | WP_006720315.1 WXG100 family type VII secretion target |
| 1(1) | WP_249897305.1 | WP_249897305.1 hypothetical protein |
| 1(1) | WP_162848531.1 | WP_162848531.1 pre-toxin TG domain-containing protein |
| 1(1) | WP_286124785.1 | WP_286124785.1 hypothetical protein |
| 1(3) | WP_095266198.1 | WP_095266198.1 WXG100 family type VII secretion target |
| 1(1) | WP_291630880.1 | WP_291630880.1 EsaB/YukD family protein |
| 1(1) | WP_251480997.1 | WP_251480997.1 DUF2974 domain-containing protein |
| 1(1) | WP_211022782.1 | WP_211022782.1 NlpC/P60 family protein |
| 1(1) | WP_304216904.1 | WP_304216904.1 Ig-like domain-containing protein |
| 1(1) | WP_283157902.1 | WP_283157902.1 hypothetical protein |
| 1(1) | WP_077833115.1 | WP_077833115.1 Ig-like domain-containing protein |
| 1(1) | WP_010346282.1 | WP_010346282.1 WXG100 family type VII secretion target |
| 1(1) | WP_279830781.1 | WP_279830781.1 copper amine oxidase N-terminal domain-containing protein |
| 1(1) | WP_075395695.1 | WP_075395695.1 WXG100 family type VII secretion target |
| 1(1) | WP_073008251.1 | WP_073008251.1 hypothetical protein |
| 1(1) | WP_213509969.1 | WP_213509969.1 EsaB/YukD family protein |
| 1(1) | WP_268623242.1 | WP_268623242.1 hypothetical protein |
| 1(1) | WP_044037256.1 | WP_044037256.1 hypothetical protein |
| 1(1) | WP_058486868.1 | WP_058486868.1 WXG100 family type VII secretion target |
| 1(1) | WP_108464090.1 | WP_108464090.1 WXG100 family type VII secretion target |
| 1(1) | WP_270801949.1 | WP_270801949.1 hypothetical protein |
| 1(1) | WP_278740197.1 | WP_278740197.1 WXG100 family type VII secretion target |
| 1(1) | WP_076219073.1 | WP_076219073.1 hypothetical protein |
| 1(1) | WP_113052564.1 | WP_113052564.1 hypothetical protein |
| 1(1) | WP_068696887.1 | WP_068696887.1 WXG100 family type VII secretion target |
| 1(1) | WP_257675220.1 | WP_257675220.1 Ig-like domain-containing protein |
| 1(1) | WP_017149764.1 | WP_017149764.1 zincin-like metallopeptidase toxin domain-containing protein |
| 1(1) | WP_110931532.1 | WP_110931532.1 copper amine oxidase N-terminal domain-containing protein |
| 1(2) | WP_098248519.1 | WP_098248519.1 WXG100 family type VII secretion target |
| 1(1) | WP_291630876.1 | WP_291630876.1 FHA domain-containing protein |
| 1(1) | WP_220031161.1 | WP_220031161.1 hypothetical protein |
| 1(1) | WP_273061518.1 | WP_273061518.1 FHA domain-containing protein |
| 1(1) | WP_066022456.1 | WP_066022456.1 pore-forming ESAT-6 family protein |
| 1(1) | WP_041926414.1 | WP_041926414.1 HNH endonuclease |
| 1(1) | WP_283157898.1 | WP_283157898.1 tetratricopeptide repeat protein |
| 1(1) | WP_068694717.1 | WP_068694717.1 hypothetical protein |

| | | |
|---|---|---|
| 1(1) | WP_274485152.1 | WP_274485152.1 WXG100 family type VII secretion target |
| 1(1) | WP_291672215.1 | WP_291672215.1 T7SS effector LXG polymorphic toxin |
| 1(1) | WP_253440780.1 | WP_253440780.1 hypothetical protein |
| 1(1) | WP_036702442.1 | WP_036702442.1 stalk domain-containing protein |
| 1(1) | WP_277166583.1 | WP_277166583.1 Ig-like domain repeat protein |
| 1(1) | WP_214706004.1 | WP_214706004.1 DUF2974 domain-containing protein |
| 1(1) | WP_289342248.1 | WP_289342248.1 glycosyltransferase |
| 1(1) | WP_251548765.1 | WP_251548765.1 Ig-like domain-containing protein |
| 1(1) | WP_098670173.1 | WP_098670173.1 hypothetical protein |
| 1(1) | WP_234414302.1 | WP_234414302.1 hypothetical protein |
| 1(1) | WP_169238724.1 | WP_169238724.1 WXG100 family type VII secretion target |
| 1(1) | WP_307738715.1 | WP_307738715.1 DUF3662 domain-containing protein |
| 1(4) | WP_094699652.1 | WP_094699652.1 type VII secretion target |
| 1(2) | WP_018670522.1 | WP_018670522.1 hypothetical protein |
| 1(1) | WP_021878934.1 | WP_021878934.1 DUF5050 domain-containing protein |
| 1(1) | WP_132008178.1 | WP_132008178.1 Ig-like domain-containing protein |
| 1(1) | WP_024027633.1 | WP_024027633.1 pore-forming ESAT-6 family protein |
| 1(1) | WP_110934069.1 | WP_110934069.1 hypothetical protein |
| 1(1) | WP_192571318.1 | WP_192571318.1 hypothetical protein |
| 1(1) | WP_211023026.1 | WP_211023026.1 protein kinase family protein |
| 1(1) | WP_116231174.1 | WP_116231174.1 hypothetical protein |
| 1(1) | WP_240379734.1 | WP_240379734.1 hypothetical protein |
| 1(1) | WP_294386426.1 | WP_294386426.1 serine/threonine-protein kinase |
| 1(1) | WP_283910927.1 | WP_283910927.1 DUF2974 domain-containing protein |
| 1(3) | WP_110899367.1 | WP_110899367.1 hypothetical protein |
| 1(2) | WP_185349994.1 | WP_185349994.1 type VII secretion protein EsaA |
| 1(1) | WP_213532658.1 | WP_213532658.1 hypothetical protein |
| 1(1) | WP_160037485.1 | WP_160037485.1 WXG100 family type VII secretion target |
| 1(1) | WP_077852411.1 | WP_077852411.1 tetratricopeptide repeat protein |
| 1(1) | WP_090674366.1 | WP_090674366.1 WXG100 family type VII secretion target |
| 1(20) | WP_000391139.1 | WP_000391139.1 hypothetical protein |
| 1(1) | WP_195571393.1 | WP_195571393.1 EsaB/YukD family protein |
| 1(1) | WP_142325888.1 | WP_142325888.1 hypothetical protein |
| 1(1) | WP_048826600.1 | WP_048826600.1 Ig-like domain-containing protein |
| 1(1) | WP_246073058.1 | WP_246073058.1 hypothetical protein |
| 1(1) | WP_178420415.1 | WP_178420415.1 WXG100 family type VII secretion target |
| 1(2) | WP_281813075.1 | WP_281813075.1 S-layer homology domain-containing protein |
| 1(1) | WP_216830388.1 | WP_216830388.1 DUF6382 domain-containing protein |
| 1(1) | WP_028589293.1 | WP_028589293.1 membrane protein |
| 1(1) | WP_078181470.1 | WP_078181470.1 WXG100 family type VII secretion target |
| 1(1) | WP_127199772.1 | WP_127199772.1 WXG100 family type VII secretion target |
| 1(1) | WP_005210442.1 | WP_005210442.1 hypothetical protein |
| 1(1) | WP_244525363.1 | WP_244525363.1 hypothetical protein |
| 1(2) | WP_000054425.1 | WP_000054425.1 SseB family protein |
| 1(1) | WP_097956891.1 | WP_097956891.1 EsaB/YukD family protein |
| 1(1) | WP_206156779.1 | WP_206156779.1 WXG100 family type VII secretion target |
| 1(4) | WP_010496610.1 | WP_010496610.1 pore-forming ESAT-6 family protein |
| 1(2) | WP_185516173.1 | WP_185516173.1 type VII secretion protein EsaA |
| 1(1) | WP_283952944.1 | WP_283952944.1 FHA domain-containing protein |
| 1(1) | WP_257443646.1 | WP_257443646.1 WXG100 family type VII secretion target |
| 1(2) | WP_010963734.1 | WP_010963734.1 tetratricopeptide repeat protein |
| 1(1) | WP_303942546.1 | WP_303942546.1 FHA domain-containing protein |
| 1(1) | WP_159081750.1 | WP_159081750.1 WXG100 family type VII secretion target |
| 1(2) | WP_200372569.1 | WP_200372569.1 type VII secretion protein EsaA |
| 1(1) | WP_208019351.1 | WP_208019351.1 tetratricopeptide repeat protein |
| 1(1) | WP_163258051.1 | WP_163258051.1 WXG100 family type VII secretion target, partial |
| 1(16) | WP_017689072.1 | WP_017689072.1 WXG100 family type VII secretion target |
| 1(1) | WP_028405549.1 | WP_028405549.1 serine/threonine protein kinase |
| 1(1) | WP_251375541.1 | WP_251375541.1 YhgE/Pip domain-containing protein |
| 1(1) | WP_284566226.1 | WP_284566226.1 hypothetical protein |
| 1(1) | WP_216830390.1 | WP_216830390.1 WXG100 family type VII secretion target |
| 1(1) | WP_189018610.1 | WP_189018610.1 hypothetical protein |
| 1(1) | WP_046522786.1 | WP_046522786.1 DUF2974 domain-containing protein |
| 1(1) | WP_068696878.1 | WP_068696878.1 hypothetical protein |
| 1(1) | WP_039273523.1 | WP_039273523.1 stalk domain-containing protein |
| 1(1) | WP_283157896.1 | WP_283157896.1 hypothetical protein |
| 1(2) | WP_228465486.1 | WP_228465486.1 hypothetical protein |
| 1(2) | WP_094870545.1 | WP_094870545.1 AAA family ATPase |
| 1(2) | WP_065228764.1 | WP_065228764.1 bacteriocin-processing peptidase family protein |
| 1(1) | WP_090761837.1 | WP_090761837.1 Ig-like domain repeat protein |
| 1(1) | WP_155703927.1 | WP_155703927.1 hypothetical protein |
| 1(1) | WP_055109372.1 | WP_055109372.1 Mbeg1-like protein |
| 1(1) | WP_141522756.1 | WP_141522756.1 deaminase domain-containing protein |
| 1(1) | WP_242874737.1 | WP_242874737.1 serine/threonine-protein kinase |
| 1(1) | WP_210364459.1 | WP_210364459.1 serine/threonine-protein kinase |
| 1(1) | WP_139867376.1 | WP_139867376.1 WXG100 family type VII secretion target |
| 1(1) | WP_167383761.1 | WP_167383761.1 WXG100 family type VII secretion target |
| 1(2) | WP_185516167.1 | WP_185516167.1 glycohydrolase toxin TNT-related protein |
| 1(1) | WP_213618822.1 | WP_213618822.1 YhgE/Pip domain-containing protein |
| 1(1) | WP_137698110.1 | WP_137698110.1 WXG100 family type VII secretion target |
| 1(1) | WP_034675352.1 | WP_034675352.1 hypothetical protein |
| 1(1) | WP_144476563.1 | WP_144476563.1 bacteriocin-processing peptidase family protein |
| 1(1) | WP_149885345.1 | WP_149885345.1 tetratricopeptide repeat protein, partial |
| 1(1) | WP_025364674.1 | WP_025364674.1 S-layer homology domain-containing protein |
| 1(1) | WP_098558848.1 | WP_098558848.1 hypothetical protein |
| 1(1) | WP_188535088.1 | WP_188535088.1 hypothetical protein |
| 1(1) | WP_216791218.1 | WP_216791218.1 pore-forming ESAT-6 family protein |
| 1(1) | WP_270364633.1 | WP_270364633.1 hypothetical protein |
| 1(1) | WP_273061468.1 | WP_273061468.1 serine/threonine-protein kinase |
| 1(1) | WP_034773553.1 | WP_034773553.1 DUF2974 domain-containing protein |
| 1(1) | WP_242400031.1 | WP_242400031.1 hypothetical protein |
| 1(1) | WP_077619641.1 | WP_077619641.1 serine/threonine-protein kinase |
| 1(1) | WP_130810261.1 | WP_130810261.1 hypothetical protein |
| 1(1) | WP_165615085.1 | WP_165615085.1 WXG100 family type VII secretion target |
| 1(1) | WP_209861651.1 | WP_209861651.1 hypothetical protein |
| 1(1) | WP_215601068.1 | WP_215601068.1 DNA/RNA non-specific endonuclease |
| 1(1) | WP_051417620.1 | WP_051417620.1 DUF2974 domain-containing protein |
| 1(1) | WP_098291292.1 | WP_098291292.1 T7SS effector LXG polymorphic toxin |
| 1(1) | WP_055109384.1 | WP_055109384.1 DUF5050 domain-containing protein |
| 1(1) | WP_264456860.1 | WP_264456860.1 WXG100 family type VII secretion target |
| 1(6) | WP_003718141.1 | WP_003718141.1 EsaB/YukD family protein |
| 1(3) | WP_228463507.1 | WP_228463507.1 hypothetical protein |
| 1(3) | WP_171414519.1 | WP_171414519.1 hypothetical protein |
| 1(1) | WP_239617477.1 | WP_239617477.1 WXG100 family type VII secretion target |

1(2) WP_228464400.1 WP_228464400.1 pre-toxin TG domain-containing protein
1(1) WP_284566230.1 WP_284566230.1 HNH endonuclease
1(4) WP_013370787.1 WP_013370787.1 membrane protein
1(1) WP_308110124.1 WP_308110124.1 beta-N-acetylhexosaminidase
1(1) WP_216830400.1 WP_216830400.1 hypothetical protein
1(1) WP_196240856.1 WP_196240856.1 WXG100 family type VII secretion target
1(1) WP_254865042.1 WP_254865042.1 type VII secretion protein EsaA, partial
1(1) WP_211022783.1 WP_211022783.1 copper amine oxidase N-terminal domain-containing protein
1(1) WP_251466803.1 WP_251466803.1 pore-forming ESAT-6 family protein
1(1) WP_142307170.1 WP_142307170.1 WXG100 family type VII secretion target, partial
1(1) WP_041964569.1 WP_041964569.1 serine/threonine-protein kinase
1(1) WP_052106377.1 WP_052106377.1 FHA domain-containing protein
1(1) WP_307349941.1 WP_307349941.1 S-layer homology domain-containing protein
1(2) WP_000789101.1 WP_000789101.1 hypothetical protein
1(1) WP_305022922.1 WP_305022922.1 WXG100 family type VII secretion target
1(5) WP_025848611.1 WP_025848611.1 pore-forming ESAT-6 family protein
1(1) WP_213117488.1 WP_213117488.1 serine/threonine-protein kinase
1(1) WP_251596652.1 WP_251596652.1 serine hydrolase domain-containing protein
1(1) WP_212981562.1 WP_212981562.1 WXG100 family type VII secretion target
1(1) WP_254349520.1 WP_254349520.1 S-layer homology domain-containing protein
1(2) WP_142307417.1 WP_142307417.1 hypothetical protein
1(1) WP_309098784.1 WP_309098784.1 Ig-like domain-containing protein
1(1) WP_294386432.1 WP_294386432.1 FHA domain-containing protein
1(1) WP_123797110.1 WP_123797110.1 S-layer homology domain-containing protein
1(1) WP_174809198.1 WP_174809198.1 WXG100 family type VII secretion target
1(1) WP_098266652.1 WP_098266652.1 hypothetical protein
1(3) WP_097920541.1 WP_097920541.1 bacteriocin-processing peptidase family protein
1(1) WP_291289935.1 WP_291289935.1 FHA domain-containing protein
1(1) WP_086385917.1 WP_086385917.1 type VII secretion protein EsaA
1(1) WP_307399107.1 WP_307399107.1 serine/threonine-protein kinase
1(2) WP_227557631.1 WP_227557631.1 hypothetical protein
1(1) WP_270784521.1 WP_270784521.1 hypothetical protein
1(1) WP_127199773.1 WP_127199773.1 hypothetical protein
1(1) WP_077567017.1 WP_077567017.1 serine/threonine protein kinase
1(1) WP_052118161.1 WP_052118161.1 hypothetical protein
1(1) WP_076322113.1 WP_076322113.1 hypothetical protein
1(1) WP_251557376.1 WP_251557376.1 FHA domain-containing protein
1(1) WP_088119572.1 WP_088119572.1 WXG100 family type VII secretion target
1(1) WP_215601067.1 WP_215601067.1 WXG100 family type VII secretion target
1(2) WP_076135807.1 WP_076135807.1 hypothetical protein
1(1) WP_258425841.1 WP_258425841.1 hypothetical protein
1(1) WP_308723533.1 WP_308723533.1 hypothetical protein
1(1) WP_271839938.1 WP_271839938.1 WXG100 family type VII secretion target
1(1) WP_238187883.1 WP_238187883.1 pore-forming ESAT-6 family protein
1(2) WP_016512880.1 WP_016512880.1 WXG100 family type VII secretion target
1(1) WP_098360831.1 WP_098360831.1 bacteriocin-processing peptidase family protein
1(1) WP_205001928.1 WP_205001928.1 hypothetical protein
1(1) WP_213644229.1 WP_213644229.1 serine/threonine-protein kinase
1(1) WP_213506890.1 WP_213506890.1 NlpC/P60 family protein
1(1) WP_235286200.1 WP_235286200.1 WXG100 family type VII secretion target
1(1) WP_295628345.1 WP_295628345.1 tetratricopeptide repeat protein
1(1) WP_034675367.1 WP_034675367.1 WXG100 family type VII secretion target
1(1) WP_031463341.1 WP_031463341.1 S-layer homology domain-containing protein
1(1) WP_270574770.1 WP_270574770.1 FHA domain-containing protein
1(1) WP_244719645.1 WP_244719645.1 hypothetical protein
1(1) WP_270784520.1 WP_270784520.1 SseB family protein
1(1) WP_308083982.1 WP_308083982.1 glycoside hydrolase family 18 protein
1(1) WP_159081748.1 WP_159081748.1 EsaB/YukD family protein
1(1) WP_255295343.1 WP_255295343.1 deaminase domain-containing protein
1(1) WP_098976050.1 WP_098976050.1 SseB family protein
1(1) WP_013658711.1 WP_013658711.1 WXG100 family type VII secretion target
1(1) WP_098670174.1 WP_098670174.1 tetratricopeptide repeat protein
1(1) WP_071736273.1 WP_071736273.1 tetratricopeptide repeat protein
1(1) WP_183600893.1 WP_183600893.1 WXG100 family type VII secretion target
1(2) WP_057897628.1 WP_057897628.1 hypothetical protein
1(1) WP_097920537.1 WP_097920537.1 WXG100 family type VII secretion target
1(1) WP_251417659.1 WP_251417659.1 EsaB/YukD family protein
1(1) WP_069011418.1 WP_069011418.1 S-layer homology domain-containing protein
1(3) WP_018883275.1 WP_018883275.1 hypothetical protein
1(1) WP_142337621.1 WP_142337621.1 hypothetical protein
1(2) WP_087433609.1 WP_087433609.1 serine/threonine protein kinase
1(1) WP_306807292.1 WP_306807292.1 protein kinase
1(8) WP_001050354.1 WP_001050354.1 SseB family protein
1(1) WP_015595749.1 WP_015595749.1 YtxH domain-containing protein
1(2) WP_213592869.1 WP_213592869.1 hypothetical protein
1(1) WP_140168584.1 WP_140168584.1 hypothetical protein
1(1) WP_090722900.1 WP_090722900.1 hypothetical protein
1(2) WP_005546505.1 WP_005546505.1 hypothetical protein
1(1) WP_034773536.1 WP_034773536.1 DUF2974 domain-containing protein
1(1) WP_168976288.1 WP_168976288.1 hypothetical protein
1(1) WP_098768166.1 WP_098768166.1 hypothetical protein
1(1) WP_164515193.1 WP_164515193.1 DUF5050 domain-containing protein
1(1) WP_000243841.1 WP_000243841.1 hypothetical protein
1(1) WP_257149141.1 WP_257149141.1 hypothetical protein, partial
1(1) WP_213579161.1 WP_213579161.1 serine/threonine-protein kinase
1(1) WP_210364461.1 WP_210364461.1 Ig-like domain-containing protein
1(1) WP_216830399.1 WP_216830399.1 hypothetical protein
1(1) WP_126996309.1 WP_126996309.1 DUF5050 domain-containing protein
1(1) WP_277175592.1 WP_277175592.1 hypothetical protein
1(1) WP_207745874.1 WP_207745874.1 hypothetical protein
1(1) WP_251466802.1 WP_251466802.1 WXG100 family type VII secretion target
1(1) WP_077852741.1 WP_077852741.1 tetratricopeptide repeat protein
1(1) WP_098943210.1 WP_098943210.1 tetratricopeptide repeat protein
1(1) WP_283904338.1 WP_283904338.1 RHS repeat-associated core domain-containing protein
1(1) WP_066419975.1 WP_066419975.1 hypothetical protein
1(1) WP_065291254.1 WP_065291254.1 copper amine oxidase N-terminal domain-containing protein
1(1) WP_051624142.1 WP_051624142.1 Ig-like domain-containing protein
1(1) WP_180229742.1 WP_180229742.1 type VII secretion protein EsaA
1(2) WP_075702373.1 WP_075702373.1 type VII secretion protein EsaA
1(4) WP_102372753.1 WP_102372753.1 WXG100 family type VII secretion target
1(1) WP_286124792.1 WP_286124792.1 hypothetical protein
1(1) WP_021291841.1 WP_021291841.1 DNA/RNA non-specific endonuclease

| | | |
|---|---|---|
| 1(7) | WP_061805366.1 | WP_061805366.1 WXG100 family type VII secretion target |
| 1(1) | WP_000391138.1 | WP_000391138.1 EsaB/YukD family protein |
| 1(1) | WP_066419990.1 | WP_066419990.1 WXG100 family type VII secretion target |
| 1(1) | WP_088063234.1 | WP_088063234.1 WXG100 family type VII secretion target |
| 1(1) | WP_145046457.1 | WP_145046457.1 hypothetical protein |
| 1(1) | WP_271005907.1 | WP_271005907.1 hypothetical protein |
| 1(1) | WP_079506396.1 | WP_079506396.1 DUF2974 domain-containing protein |
| 1(1) | WP_075395492.1 | WP_075395492.1 bacteriocin-processing peptidase family protein |
| 1(1) | WP_034526602.1 | WP_034526602.1 serine/threonine-protein kinase |
| 1(1) | WP_047153271.1 | WP_047153271.1 WXG100 family type VII secretion target |
| 1(1) | WP_211023022.1 | WP_211023022.1 RHS repeat-associated core domain-containing protein |
| 1(1) | WP_108465345.1 | WP_108465345.1 hypothetical protein |
| 1(1) | WP_184182997.1 | WP_184182997.1 WXG100 family type VII secretion target |
| 1(1) | WP_087986804.1 | WP_087986804.1 tetratricopeptide repeat protein |
| 1(1) | WP_277175589.1 | WP_277175589.1 Ig-like domain repeat protein |
| 1(2) | WP_098087524.1 | WP_098087524.1 WXG100 family type VII secretion target, partial |
| 1(1) | WP_195515240.1 | WP_195515240.1 WXG100 family type VII secretion target |
| 1(1) | WP_102264548.1 | WP_102264548.1 WXG100 family type VII secretion target |
| 1(1) | WP_199925769.1 | WP_199925769.1 DUF5050 domain-containing protein |
| 1(1) | WP_026593080.1 | WP_026593080.1 EsaB/YukD family protein |
| 1(1) | WP_184183002.1 | WP_184183002.1 protein kinase family protein |
| 1(1) | WP_170839746.1 | WP_170839746.1 glycosyltransferase |
| 1(1) | WP_301167500.1 | WP_301167500.1 hypothetical protein |
| 1(9) | WP_036684485.1 | WP_036684485.1 WXG100 family type VII secretion target |
| 1(1) | WP_189031049.1 | WP_189031049.1 AAA family ATPase |
| 1(1) | WP_068694715.1 | WP_068694715.1 hypothetical protein |
| 1(1) | WP_180358490.1 | WP_180358490.1 WXG100 family type VII secretion target |
| 1(1) | WP_033100018.1 | WP_033100018.1 DUF2974 domain-containing protein |
| 1(2) | WP_145153643.1 | WP_145153643.1 WXG100 family type VII secretion target |
| 1(4) | WP_062833881.1 | WP_062833881.1 WXG100 family type VII secretion target |
| 1(1) | WP_291672200.1 | WP_291672200.1 DUF6382 domain-containing protein |
| 1(1) | WP_157441030.1 | WP_157441030.1 WXG100 family type VII secretion target |
| 1(1) | WP_307312566.1 | WP_307312566.1 protein kinase |
| 1(1) | WP_257142774.1 | WP_257142774.1 HNH endonuclease |
| 1(2) | WP_051614154.1 | WP_051614154.1 copper amine oxidase N-terminal domain-containing protein |
| 1(1) | WP_291288261.1 | WP_291288261.1 hypothetical protein |
| 1(1) | WP_150367640.1 | WP_150367640.1 hypothetical protein |
| 1(1) | WP_169028893.1 | WP_169028893.1 DUF5050 domain-containing protein |
| 1(1) | WP_158329533.1 | WP_158329533.1 WG repeat-containing protein |
| 1(2) | WP_098255403.1 | WP_098255403.1 WXG100 family type VII secretion target |
| 1(7) | WP_048556841.1 | WP_048556841.1 tetratricopeptide repeat protein |
| 1(1) | WP_066092000.1 | WP_066092000.1 DUF2974 domain-containing protein |
| 1(1) | WP_123067114.1 | WP_123067114.1 WXG100 family type VII secretion target |
| 1(1) | WP_215153839.1 | WP_215153839.1 hypothetical protein |
| 1(1) | WP_076322204.1 | WP_076322204.1 hypothetical protein |
| 1(1) | WP_039871650.1 | WP_039871650.1 hypothetical protein |
| 1(2) | WP_000883802.1 | WP_000883802.1 bacteriocin-processing peptidase family protein |
| 1(1) | WP_249897319.1 | WP_249897319.1 DUF2974 domain-containing protein |
| 1(1) | WP_116344882.1 | WP_116344882.1 type VII secretion protein EsaA, partial |
| 1(1) | WP_106382855.1 | WP_106382855.1 hypothetical protein |
| 1(1) | WP_150158012.1 | WP_150158012.1 type VII secretion protein EsaA |

| | | |
|---|---|---|
| 1(1) | WP_027088425.1 | WP_027088425.1 WXG100 family type VII secretion target |
| 1(1) | WP_251596676.1 | WP_251596676.1 EsaB/YukD family protein |
| 1(1) | WP_297183848.1 | WP_297183848.1 hypothetical protein |
| 1(1) | WP_238187882.1 | WP_238187882.1 WXG100 family type VII secretion target |
| 1(1) | WP_223868744.1 | WP_223868744.1 hypothetical protein |
| 1(1) | WP_165147476.1 | WP_165147476.1 hypothetical protein |
| 1(1) | WP_182301834.1 | WP_182301834.1 peptidoglycan-binding protein |
| 1(1) | WP_074565303.1 | WP_074565303.1 SseB family protein |
| 1(1) | WP_136379690.1 | WP_136379690.1 hypothetical protein |
| 1(4) | WP_090921054.1 | WP_090921054.1 WXG100 family type VII secretion target |
| 1(1) | WP_307394639.1 | WP_307394639.1 WXG100 family type VII secretion target |
| 1(1) | WP_227495882.1 | WP_227495882.1 HNH endonuclease |
| 1(1) | WP_289355900.1 | WP_289355900.1 hypothetical protein |
| 1(2) | WP_169850944.1 | WP_169850944.1 hypothetical protein |
| 1(1) | WP_213616453.1 | WP_213616453.1 serine/threonine-protein kinase |
| 1(1) | WP_257675924.1 | WP_257675924.1 tetratricopeptide repeat protein |
| 1(1) | WP_228463906.1 | WP_228463906.1 DUF6531 domain-containing protein |
| 1(2) | WP_213117485.1 | WP_213117485.1 WXG100 family type VII secretion target |
| 1(1) | WP_014281184.1 | WP_014281184.1 membrane protein |
| 1(1) | WP_120112230.1 | WP_120112230.1 pre-toxin TG domain-containing protein |
| 1(1) | WP_044307647.1 | WP_044307647.1 type VII secretion protein EsaA, partial |
| 1(1) | WP_256424317.1 | WP_256424317.1 hypothetical protein |
| 1(1) | WP_286124786.1 | WP_286124786.1 SseB family protein |
| 1(1) | WP_076210988.1 | WP_076210988.1 hypothetical protein |
| 1(1) | WP_208752242.1 | WP_208752242.1 tetratricopeptide repeat protein |
| 1(2) | WP_003516577.1 | WP_003516577.1 EsaB/YukD family protein |
| 1(1) | WP_301804672.1 | WP_301804672.1 type VII secretion target |
| 1(1) | WP_286017538.1 | WP_286017538.1 hypothetical protein |
| 1(1) | WP_295628334.1 | WP_295628334.1 tetratricopeptide repeat protein |
| 1(1) | WP_133332984.1 | WP_133332984.1 protein kinase |
| 1(2) | WP_048552310.1 | WP_048552310.1 tetratricopeptide repeat protein |
| 1(1) | WP_098264288.1 | WP_098264288.1 SseB family protein |
| 1(1) | WP_078205524.1 | WP_078205524.1 protein kinase |
| 1(1) | WP_094699656.1 | WP_094699656.1 hypothetical protein |
| 1(1) | WP_307349970.1 | WP_307349970.1 WXG100 family type VII secretion target |
| 1(1) | WP_251245804.1 | WP_251245804.1 DUF2974 domain-containing protein |
| 1(1) | WP_251466809.1 | WP_251466809.1 protein kinase |
| 1(1) | WP_137052045.1 | WP_137052045.1 bacteriocin-processing peptidase family protein, partial |
| 1(1) | WP_281667417.1 | WP_281667417.1 hypothetical protein |
| 1(1) | WP_243256001.1 | WP_243256001.1 FHA domain-containing protein |
| 1(1) | WP_270996003.1 | WP_270996003.1 hypothetical protein, partial |
| 1(1) | WP_306049034.1 | WP_306049034.1 protein kinase |
| 1(1) | WP_120462943.1 | WP_120462943.1 hypothetical protein |
| 1(1) | WP_295632050.1 | WP_295632050.1 FHA domain-containing protein |
| 1(1) | WP_017149766.1 | WP_017149766.1 WXG100 family type VII secretion target |
| 1(1) | WP_076210984.1 | WP_076210984.1 RHS repeat-associated core domain-containing protein |
| 1(1) | WP_106765961.1 | WP_106765961.1 hypothetical protein |
| 1(1) | WP_286124783.1 | WP_286124783.1 WXG100 family type VII secretion target |
| 1(1) | WP_152392575.1 | WP_152392575.1 hypothetical protein |
| 1(1) | WP_305163861.1 | WP_305163861.1 tetratricopeptide repeat protein |
| 1(1) | WP_124049171.1 | WP_124049171.1 hypothetical protein |

| | | |
|---|---|---|
| 1(1) | WP_137078369.1 | WP_137078369.1 copper amine oxidase N-terminal domain-containing protein |
| 1(1) | WP_197256860.1 | WP_197256860.1 hypothetical protein |
| 1(1) | WP_119601494.1 | WP_119601494.1 EsaB/YukD family protein |
| 1(1) | WP_229137982.1 | WP_229137982.1 bacteriocin-processing peptidase family protein |
| 1(1) | WP_258958953.1 | WP_258958953.1 pore-forming ESAT-6 family protein |
| 1(2) | WP_047909615.1 | WP_047909615.1 hypothetical protein |
| 1(1) | WP_255443973.1 | WP_255443973.1 CdiA family toxin C-terminal domain-containing protein |
| 1(1) | WP_111925422.1 | WP_111925422.1 hypothetical protein |
| 1(1) | WP_257535705.1 | WP_257535705.1 hypothetical protein |
| 1(1) | WP_211082927.1 | WP_211082927.1 WXG100 family type VII secretion target |
| 1(1) | WP_044792569.1 | WP_044792569.1 bacteriocin-processing peptidase family protein |
| 1(1) | WP_106382856.1 | WP_106382856.1 WXG100 family type VII secretion target |
| 1(1) | WP_255256110.1 | WP_255256110.1 GH-E family nuclease |
| 1(1) | WP_098013430.1 | WP_098013430.1 type VII secretion protein EsaA |
| 1(1) | WP_019912153.1 | WP_019912153.1 hypothetical protein |
| 1(1) | WP_268238743.1 | WP_268238743.1 WXG100 family type VII secretion target |
| 1(1) | WP_307349964.1 | WP_307349964.1 hypothetical protein |
| 1(5) | WP_102264547.1 | WP_102264547.1 pore-forming ESAT-6 family protein |
| 1(1) | WP_308082252.1 | WP_308082252.1 serine/threonine-protein kinase |
| 1(1) | WP_183578437.1 | WP_183578437.1 hypothetical protein |
| 1(1) | WP_169028901.1 | WP_169028901.1 AAA family ATPase |
| 1(1) | WP_270472026.1 | WP_270472026.1 T7SS effector LXG polymorphic toxin |
| 1(2) | WP_168178887.1 | WP_168178887.1 WXG100 family type VII secretion target |
| 1(1) | WP_017692335.1 | WP_017692335.1 hypothetical protein |
| 1(1) | WP_143881074.1 | WP_143881074.1 hypothetical protein |
| 1(1) | WP_010250282.1 | WP_010250282.1 hypothetical protein |
| 1(1) | WP_134750626.1 | WP_134750626.1 WXG100 family type VII secretion target |
| 1(1) | WP_070638653.1 | WP_070638653.1 FHA domain-containing protein |
| 1(1) | WP_208926587.1 | WP_208926587.1 tetratricopeptide repeat protein |
| 1(1) | WP_029390078.1 | WP_029390078.1 WXG100 family type VII secretion target |
| 1(1) | WP_143695844.1 | WP_143695844.1 hypothetical protein |
| 1(1) | WP_141533090.1 | WP_141533090.1 deaminase domain-containing protein |
| 1(1) | WP_207994161.1 | WP_207994161.1 WXG100 family type VII secretion target |
| 1(1) | WP_232274494.1 | WP_232274494.1 EsaB/YukD family protein |
| 1(1) | WP_102372744.1 | WP_102372744.1 FHA domain-containing protein |
| 1(1) | WP_066415457.1 | WP_066415457.1 serine/threonine-protein kinase |
| 1(2) | WP_000506915.1 | WP_000506915.1 tetratricopeptide repeat protein |
| 1(1) | WP_211023040.1 | WP_211023040.1 copper amine oxidase N-terminal domain-containing protein |
| 1(1) | WP_304142077.1 | WP_304142077.1 hypothetical protein |
| 1(1) | WP_102372754.1 | WP_102372754.1 hypothetical protein |
| 1(1) | WP_316324166.1 | WP_316324166.1 GBS Bsp-like repeat-containing protein |
| 1(1) | WP_281937375.1 | WP_281937375.1 DUF2974 domain-containing protein |
| 1(1) | WP_255260044.1 | WP_255260044.1 hypothetical protein |
| 1(1) | WP_016362249.1 | WP_016362249.1 hypothetical protein |
| 1(2) | WP_111268942.1 | WP_111268942.1 hypothetical protein |
| 1(2) | WP_109625242.1 | WP_109625242.1 WXG100 family type VII secretion target |
| 1(1) | WP_078185401.1 | WP_078185401.1 HNH/ENDO VII family nuclease |
| 1(1) | WP_260446636.1 | WP_260446636.1 hypothetical protein |
| 1(1) | WP_036684483.1 | WP_036684483.1 DUF2974 domain-containing protein |
| 1(1) | WP_289483787.1 | WP_289483787.1 hypothetical protein |
| 1(1) | WP_289613150.1 | WP_289613150.1 protein kinase |
| 1(1) | WP_142330389.1 | WP_142330389.1 WXG100 family type VII secretion target |
| 1(2) | WP_133332975.1 | WP_133332975.1 hypothetical protein |
| 1(1) | WP_197234720.1 | WP_197234720.1 tetratricopeptide repeat protein |
| 1(1) | WP_191272862.1 | WP_191272862.1 DUF2974 domain-containing protein |
| 1(1) | WP_098535853.1 | WP_098535853.1 T7SS effector LXG polymorphic toxin |
| 1(1) | WP_102710720.1 | WP_102710720.1 DUF5050 domain-containing protein |
| 1(1) | WP_113765179.1 | WP_113765179.1 hypothetical protein |
| 1(1) | WP_103141595.1 | WP_103141595.1 WXG100 family type VII secretion target |
| 1(1) | WP_148450945.1 | WP_148450945.1 WXG100 family type VII secretion target |
| 1(1) | WP_208203482.1 | WP_208203482.1 hypothetical protein |
| 1(1) | WP_074598759.1 | WP_074598759.1 EsaB/YukD family protein |
| 1(2) | WP_028542689.1 | WP_028542689.1 WXG100 family type VII secretion target |
| 1(1) | WP_006292355.1 | WP_006292355.1 hypothetical protein |
| 1(1) | WP_069355510.1 | WP_069355510.1 tetratricopeptide repeat protein |
| 1(1) | WP_027088427.1 | WP_027088427.1 hypothetical protein |
| 1(1) | WP_147306277.1 | WP_147306277.1 WXG100 family type VII secretion target |
| 1(1) | WP_179034794.1 | WP_179034794.1 copper amine oxidase N-terminal domain-containing protein |
| 1(1) | WP_006292336.1 | WP_006292336.1 serine/threonine-protein kinase |
| 1(1) | WP_199419771.1 | WP_199419771.1 WXG100 family type VII secretion target |
| 1(2) | WP_270993637.1 | WP_270993637.1 type VII secretion protein EsaA |
| 1(1) | WP_106822168.1 | WP_106822168.1 bacteriocin-processing peptidase family protein |
| 1(1) | WP_265408841.1 | WP_265408841.1 hypothetical protein |
| 1(1) | WP_116231179.1 | WP_116231179.1 hypothetical protein |
| 1(1) | WP_137698113.1 | WP_137698113.1 EsaB/YukD family protein |
| 1(1) | WP_308110111.1 | WP_308110111.1 Ig-like domain repeat protein |
| 1(1) | WP_039871651.1 | WP_039871651.1 hypothetical protein |
| 1(1) | WP_285748178.1 | WP_285748178.1 hypothetical protein |
| 1(3) | WP_012260834.1 | WP_012260834.1 hypothetical protein |
| 1(1) | WP_286017549.1 | WP_286017549.1 tetratricopeptide repeat protein |
| 1(1) | WP_286124794.1 | WP_286124794.1 hypothetical protein |
| 1(1) | WP_158318213.1 | WP_158318213.1 WXG100 family type VII secretion target, partial |
| 1(1) | WP_269055423.1 | WP_269055423.1 hypothetical protein |
| 1(1) | WP_119601484.1 | WP_119601484.1 hypothetical protein |
| 1(1) | WP_176551424.1 | WP_176551424.1 deaminase domain-containing protein |
| 1(1) | WP_132283088.1 | WP_132283088.1 hypothetical protein |
| 1(1) | WP_134268513.1 | WP_134268513.1 hypothetical protein |
| 1(3) | WP_175623547.1 | WP_175623547.1 AAA family ATPase |
| 1(1) | WP_155609754.1 | WP_155609754.1 WXG100 family type VII secretion target |
| 1(1) | WP_094593871.1 | WP_094593871.1 hypothetical protein |
| 1(1) | WP_068536379.1 | WP_068536379.1 WXG100 family type VII secretion target |
| 1(1) | WP_080489494.1 | WP_080489494.1 WXG100 family type VII secretion target |
| 1(1) | WP_025685034.1 | WP_025685034.1 membrane protein |
| 1(1) | WP_270065063.1 | WP_270065063.1 hypothetical protein |
| 1(2) | WP_063547142.1 | WP_063547142.1 bacteriocin-processing peptidase family protein |
| 1(1) | WP_145146964.1 | WP_145146964.1 hypothetical protein |
| 1(1) | WP_308912807.1 | WP_308912807.1 DUF6382 domain-containing protein |
| 1(1) | WP_307839770.1 | WP_307839770.1 type VII secretion protein EsaA |
| 1(1) | WP_215032119.1 | WP_215032119.1 WXG100 family type VII secretion target |
| 1(1) | WP_197937401.1 | WP_197937401.1 DUF2974 domain-containing protein |
| 1(2) | WP_068782414.1 | WP_068782414.1 hypothetical protein |

1(1) WP_068782426.1 WP_068782426.1 hypothetical protein
1(1) WP_066020840.1 WP_066020840.1 tetratricopeptide repeat protein
1(1) WP_270728529.1 WP_270728529.1 WXG100 family type VII secretion target
1(1) WP_000054424.1 WP_000054424.1 SseB family protein
1(1) WP_072180188.1 WP_072180188.1 bacteriocin-processing peptidase family protein
1(1) WP_307312532.1 WP_307312532.1 hypothetical protein
1(1) WP_111273166.1 WP_111273166.1 RHS repeat-associated core domain-containing protein, partial
1(1) WP_088550043.1 WP_088550043.1 DUF5050 domain-containing protein
1(1) WP_141538300.1 WP_141538300.1 hypothetical protein
1(1) WP_293755517.1 WP_293755517.1 hypothetical protein
1(1) WP_277156188.1 WP_277156188.1 Ig-like domain repeat protein
1(1) WP_185451817.1 WP_185451817.1 type VII secretion protein EsaA
1(1) WP_029438083.1 WP_029438083.1 bacteriocin-processing peptidase family protein
1(1) WP_095358546.1 WP_095358546.1 hypothetical protein
1(1) WP_183600897.1 WP_183600897.1 hypothetical protein
1(1) WP_213117490.1 WP_213117490.1 Ig-like domain repeat protein
1(7) WP_010496611.1 WP_010496611.1 WXG100 family type VII secretion target
1(1) WP_188535069.1 WP_188535069.1 DUF5050 domain-containing protein
1(1) WP_102710727.1 WP_102710727.1 copper amine oxidase N-terminal domain-containing protein
1(3) WP_168178787.1 WP_168178787.1 hypothetical protein
1(1) WP_290407043.1 WP_290407043.1 hypothetical protein
1(1) WP_170138181.1 WP_170138181.1 hypothetical protein
1(1) WP_052106376.1 WP_052106376.1 hypothetical protein
1(1) WP_077893465.1 WP_077893465.1 hypothetical protein
1(2) WP_095691010.1 WP_095691010.1 tetratricopeptide repeat protein
1(1) WP_141548206.1 WP_141548206.1 hypothetical protein
1(1) WP_060630648.1 WP_060630648.1 tetratricopeptide repeat protein
1(1) WP_212980186.1 WP_212980186.1 WXG100 family type VII secretion target
1(2) WP_197267873.1 WP_197267873.1 hypothetical protein
1(1) WP_308723525.1 WP_308723525.1 RHS repeat-associated core domain-containing protein
1(1) WP_304942558.1 WP_304942558.1 hypothetical protein
1(1) WP_127194957.1 WP_127194957.1 hypothetical protein
1(2) WP_010963369.1 WP_010963369.1 WXG100 family type VII secretion target
1(1) WP_270065057.1 WP_270065057.1 hypothetical protein
1(1) WP_076157820.1 WP_076157820.1 hypothetical protein
1(1) WP_243500440.1 WP_243500440.1 tetratricopeptide repeat protein
1(1) WP_042210995.1 WP_042210995.1 hypothetical protein
1(1) WP_208906814.1 WP_208906814.1 EsaB/YukD family protein
1(2) WP_212969038.1 WP_212969038.1 hypothetical protein
1(1) WP_076104000.1 WP_076104000.1 AAA family ATPase
1(1) WP_095398793.1 WP_095398793.1 serine/threonine protein kinase
1(6) WP_003718138.1 WP_003718138.1 WXG100 family type VII secretion target
1(1) WP_036642160.1 WP_036642160.1 hypothetical protein
1(1) WP_145023131.1 WP_145023131.1 S-layer homology domain-containing protein
1(1) WP_277156183.1 WP_277156183.1 hypothetical protein
1(1) WP_007083298.1 WP_007083298.1 Ig-like domain repeat protein
1(1) WP_019424143.1 WP_019424143.1 serine/threonine-protein kinase
1(1) WP_007083302.1 WP_007083302.1 vWA domain-containing protein
1(2) WP_251426363.1 WP_251426363.1 WXG100 family type VII secretion target
1(1) WP_057897626.1 WP_057897626.1 hypothetical protein
1(1) WP_119732861.1 WP_119732861.1 hypothetical protein
1(1) WP_040101532.1 WP_040101532.1 S-layer homology domain-containing protein
1(1) WP_076210948.1 WP_076210948.1 hypothetical protein
1(1) WP_145153686.1 WP_145153686.1 hypothetical protein
1(1) WP_211093659.1 WP_211093659.1 hypothetical protein
1(1) WP_123797114.1 WP_123797114.1 hypothetical protein
1(1) WP_058271103.1 WP_058271103.1 serine/threonine-protein kinase
1(1) WP_308083986.1 WP_308083986.1 pore-forming ESAT-6 family protein
1(1) WP_208919028.1 WP_208919028.1 hypothetical protein
1(1) WP_176542837.1 WP_176542837.1 CdiA family toxin C-terminal domain-containing protein
1(2) WP_087956294.1 WP_087956294.1 tetratricopeptide repeat protein
1(1) WP_055109380.1 WP_055109380.1 WXG100 family type VII secretion target
1(9) WP_000404139.1 WP_000404139.1 WXG100 family type VII secretion target
1(1) WP_070638649.1 WP_070638649.1 hypothetical protein
1(2) WP_098255411.1 WP_098255411.1 bacteriocin-processing peptidase family protein
1(1) WP_305022926.1 WP_305022926.1 hypothetical protein
1(17) WP_000054422.1 WP_000054422.1 SseB family protein
1(2) WP_066419974.1 WP_066419974.1 hypothetical protein
1(1) WP_139328520.1 WP_139328520.1 EsaB/YukD family protein
1(2) WP_204412728.1 WP_204412728.1 hypothetical protein
1(1) WP_210364457.1 WP_210364457.1 FHA domain-containing protein
1(1) WP_152597393.1 WP_152597393.1 serine protease
1(1) WP_306007803.1 WP_306007803.1 tetratricopeptide repeat protein
1(1) WP_277150355.1 WP_277150355.1 T7SS effector LXG polymorphic toxin
1(1) WP_096886734.1 WP_096886734.1 WG repeat-containing protein
1(3) WP_076279382.1 WP_076279382.1 hypothetical protein
1(1) WP_305755048.1 WP_305755048.1 protease complex subunit PrcB family protein
1(1) WP_307284689.1 WP_307284689.1 WXG100 family type VII secretion target
1(1) WP_019120055.1 WP_019120055.1 WXG100 family type VII secretion target
1(1) WP_244525357.1 WP_244525357.1 EndoU domain-containing protein
1(2) WP_194335279.1 WP_194335279.1 type VII secretion protein EsaA
1(17) WP_000391133.1 WP_000391133.1 hypothetical protein
1(1) WP_120112254.1 WP_120112254.1 hypothetical protein
1(1) WP_013913514.1 WP_013913514.1 tetratricopeptide repeat protein
1(1) WP_243518761.1 WP_243518761.1 hypothetical protein
1(1) WP_231417867.1 WP_231417867.1 hypothetical protein
1(1) WP_208752241.1 WP_208752241.1 hypothetical protein
1(1) WP_307192894.1 WP_307192894.1 FHA domain-containing protein
1(1) WP_077893579.1 WP_077893579.1 tetratricopeptide repeat protein
1(1) WP_217035864.1 WP_217035864.1 FHA domain-containing protein
1(2) WP_113672695.1 WP_113672695.1 WXG100 family type VII secretion target
1(1) WP_251426361.1 WP_251426361.1 FHA domain-containing protein
1(2) WP_271003052.1 WP_271003052.1 type VII secretion protein EsaA
1(1) WP_216830384.1 WP_216830384.1 Ig-like domain-containing protein
1(1) WP_192800073.1 WP_192800073.1 tetratricopeptide repeat protein
1(2) WP_076281074.1 WP_076281074.1 WXG100 family type VII secretion target
1(1) WP_111619121.1 WP_111619121.1 hypothetical protein
1(1) WP_242251935.1 WP_242251935.1 SseB family protein
1(1) WP_186812447.1 WP_186812447.1 WXG100 family type VII secretion target
1(1) WP_204517529.1 WP_204517529.1 DL-endopeptidase inhibitor IseA family protein
1(12) WP_003745040.1 WP_003745040.1 WXG100 family type VII secretion target
1(1) WP_195244535.1 WP_195244535.1 hypothetical protein

| | | |
|---|---|---|
| 1(1) | WP_208919034.1 | WP_208919034.1 hypothetical protein |
| 1(1) | WP_110899362.1 | WP_110899362.1 RHS repeat-associated core domain-containing protein |
| 1(2) | WP_175502002.1 | WP_175502002.1 WXG100 family type VII secretion target |
| 1(1) | WP_213143950.1 | WP_213143950.1 WXG100 family type VII secretion target |
| 1(1) | WP_070638631.1 | WP_070638631.1 glycosyltransferase family 2 protein |
| 1(1) | WP_090908280.1 | WP_090908280.1 serine/threonine protein kinase |
| 1(1) | WP_158329535.1 | WP_158329535.1 hypothetical protein |
| 1(1) | WP_258433017.1 | WP_258433017.1 hypothetical protein |
| 1(2) | WP_088023251.1 | WP_088023251.1 SseB family protein |
| 1(1) | WP_036684480.1 | WP_036684480.1 DUF2974 domain-containing protein |
| 1(1) | WP_226086176.1 | WP_226086176.1 hypothetical protein |
| 1(1) | WP_251426359.1 | WP_251426359.1 serine/threonine-protein kinase |
| 1(1) | WP_130810259.1 | WP_130810259.1 WXG100 family type VII secretion target |
| 1(1) | WP_212981448.1 | WP_212981448.1 copper amine oxidase N-terminal domain-containing protein |
| 1(1) | WP_048349286.1 | WP_048349286.1 YhgE/Pip domain-containing protein |
| 1(1) | WP_186327136.1 | WP_186327136.1 hypothetical protein |
| 1(1) | WP_251548768.1 | WP_251548768.1 serine/threonine-protein kinase |
| 1(1) | WP_068696867.1 | WP_068696867.1 WXG100 family type VII secretion target |
| 1(1) | WP_212695784.1 | WP_212695784.1 hypothetical protein |
| 1(1) | WP_098250526.1 | WP_098250526.1 tetratricopeptide repeat protein |
| 1(2) | WP_010963365.1 | WP_010963365.1 DUF6382 domain-containing protein |
| 1(1) | WP_063208776.1 | WP_063208776.1 hypothetical protein |
| 1(1) | WP_265153295.1 | WP_265153295.1 tetratricopeptide repeat protein |
| 1(1) | WP_048826567.1 | WP_048826567.1 WXG100 family type VII secretion target |
| 1(1) | WP_212981446.1 | WP_212981446.1 hypothetical protein |
| 1(1) | WP_132283092.1 | WP_132283092.1 hypothetical protein |
| 1(1) | WP_179950260.1 | WP_179950260.1 hypothetical protein |
| 1(1) | WP_145406444.1 | WP_145406444.1 S-layer homology domain-containing protein |
| 1(1) | WP_122971532.1 | WP_122971532.1 WXG100 family type VII secretion target |
| 1(1) | WP_098874963.1 | WP_098874963.1 type VII secretion protein EsaA |
| 1(1) | WP_016821032.1 | WP_016821032.1 membrane protein |
| 1(1) | WP_098337103.1 | WP_098337103.1 SseB family protein |
| 1(1) | WP_000883853.1 | WP_000883853.1 bacteriocin-processing peptidase family protein |
| 1(1) | WP_143316560.1 | WP_143316560.1 hypothetical protein |
| 1(2) | WP_108534959.1 | WP_108534959.1 WXG100 family type VII secretion target |
| 1(1) | WP_190284712.1 | WP_190284712.1 Ig-like domain repeat protein |
| 1(1) | WP_026478100.1 | WP_026478100.1 AAA family ATPase |
| 1(2) | WP_295628331.1 | WP_295628331.1 tetratricopeptide repeat protein |
| 1(1) | WP_128631286.1 | WP_128631286.1 DUF5050 domain-containing protein |
| 1(1) | WP_294953163.1 | WP_294953163.1 hypothetical protein |
| 1(1) | WP_213592856.1 | WP_213592856.1 DUF2974 domain-containing protein |
| 1(1) | WP_098519059.1 | WP_098519059.1 bacteriocin-processing peptidase family protein |
| 1(1) | WP_088830537.1 | WP_088830537.1 DUF2974 domain-containing protein |
| 1(1) | WP_110931551.1 | WP_110931551.1 S-layer homology domain-containing protein |
| 1(1) | WP_144564270.1 | WP_144564270.1 tetratricopeptide repeat protein |
| 1(2) | WP_166245125.1 | WP_166245125.1 WXG100 family type VII secretion target |
| 1(1) | WP_111273172.1 | WP_111273172.1 WXG100 family type VII secretion target |
| 1(1) | WP_189031036.1 | WP_189031036.1 EsaB/YukD family protein |
| 1(1) | WP_002002664.1 | WP_002002664.1 hypothetical protein |
| 1(1) | WP_058710505.1 | WP_058710505.1 S-layer homology domain-containing protein |
| 1(1) | WP_053403783.1 | WP_053403783.1 hypothetical protein |

| | | |
|---|---|---|
| 1(1) | WP_137013547.1 | WP_137013547.1 tetratricopeptide repeat protein |
| 1(4) | WP_071679492.1 | WP_071679492.1 WXG100 family type VII secretion target |
| 1(1) | WP_251542610.1 | WP_251542610.1 DUF2974 domain-containing protein |
| 1(1) | WP_028407771.1 | WP_028407771.1 hypothetical protein |
| 1(1) | WP_242950820.1 | WP_242950820.1 serine/threonine-protein kinase |
| 1(1) | WP_251638771.1 | WP_251638771.1 DUF2974 domain-containing protein |
| 1(1) | WP_052442079.1 | WP_052442079.1 LysM peptidoglycan-binding domain-containing protein |
| 1(1) | WP_077852419.1 | WP_077852419.1 FHA domain-containing protein |
| 1(1) | WP_158330731.1 | WP_158330731.1 WXG100 family type VII secretion target |
| 1(1) | WP_055105801.1 | WP_055105801.1 copper amine oxidase N-terminal domain-containing protein |
| 1(1) | WP_128631298.1 | WP_128631298.1 serine hydrolase domain-containing protein |
| 1(3) | WP_042512832.1 | WP_042512832.1 hypothetical protein |
| 1(1) | WP_192266449.1 | WP_192266449.1 hypothetical protein |
| 1(1) | WP_039306357.1 | WP_039306357.1 hypothetical protein |
| 1(1) | WP_262719888.1 | WP_262719888.1 tetratricopeptide repeat protein |
| 1(1) | WP_208926603.1 | WP_208926603.1 WXG100 family type VII secretion target |
| 1(1) | WP_272045573.1 | WP_272045573.1 WXG100 family type VII secretion target |
| 1(1) | WP_318999813.1 | WP_318999813.1 protein kinase |
| 1(1) | WP_144500999.1 | WP_144500999.1 hypothetical protein |
| 1(1) | WP_102371674.1 | WP_102371674.1 hypothetical protein |
| 1(1) | WP_307738735.1 | WP_307738735.1 FHA domain-containing protein |
| 1(1) | WP_189031026.1 | WP_189031026.1 copper amine oxidase N-terminal domain-containing protein |
| 1(3) | WP_063223282.1 | WP_063223282.1 SseB family protein |
| 1(3) | WP_256713669.1 | WP_256713669.1 hypothetical protein |
| 1(1) | WP_154118031.1 | WP_154118031.1 hypothetical protein |
| 1(1) | WP_227558286.1 | WP_227558286.1 tetratricopeptide repeat protein |
| 1(1) | WP_220027679.1 | WP_220027679.1 hypothetical protein |
| 1(1) | WP_251417675.1 | WP_251417675.1 NlpC/P60 family protein |
| 1(1) | WP_289398700.1 | WP_289398700.1 WXG100 family type VII secretion target |
| 1(2) | WP_001050356.1 | WP_001050356.1 SseB family protein |
| 1(1) | WP_231094845.1 | WP_231094845.1 pore-forming ESAT-6 family protein |
| 1(1) | WP_195571385.1 | WP_195571385.1 hypothetical protein |
| 1(3) | WP_168178784.1 | WP_168178784.1 hypothetical protein |
| 1(2) | WP_047910191.1 | WP_047910191.1 WXG100 family type VII secretion target |
| 1(1) | WP_108464087.1 | WP_108464087.1 DUF5050 domain-containing protein |
| 1(1) | WP_251596666.1 | WP_251596666.1 YhgE/Pip domain-containing protein |
| 1(1) | WP_250244819.1 | WP_250244819.1 WXG100 family type VII secretion target |
| 1(1) | WP_263703626.1 | WP_263703626.1 hypothetical protein |
| 1(1) | WP_223067436.1 | WP_223067436.1 hypothetical protein |
| 1(1) | WP_208019354.1 | WP_208019354.1 WXG100 family type VII secretion target, partial |
| 1(1) | WP_207743085.1 | WP_207743085.1 hypothetical protein |
| 1(1) | WP_166245137.1 | WP_166245137.1 Ig-like domain repeat protein |
| 1(1) | WP_242400026.1 | WP_242400026.1 hypothetical protein |
| 1(1) | WP_000883803.1 | WP_000883803.1 bacteriocin-processing peptidase family protein |
| 1(1) | WP_106822154.1 | WP_106822154.1 tetratricopeptide repeat protein |
| 1(1) | WP_262725726.1 | WP_262725726.1 hypothetical protein |
| 1(1) | WP_213509968.1 | WP_213509968.1 DUF5050 domain-containing protein |
| 1(1) | WP_095266199.1 | WP_095266199.1 serine/threonine protein kinase |
| 1(1) | WP_291647944.1 | WP_291647944.1 tetratricopeptide repeat protein |
| 1(1) | WP_098054113.1 | WP_098054113.1 T7SS effector LXG polymorphic toxin |

| | | |
|---|---|---|
| 1(1) | WP_215185479.1 | WP_215185479.1 WXG100 family type VII secretion target |
| 1(1) | WP_255807569.1 | WP_255807569.1 copper amine oxidase N-terminal domain-containing protein |
| 1(1) | WP_182301835.1 | WP_182301835.1 hypothetical protein |
| 1(1) | WP_187386577.1 | WP_187386577.1 copper amine oxidase N-terminal domain-containing protein |
| 1(1) | WP_179034790.1 | WP_179034790.1 WXG100 family type VII secretion target |
| 1(1) | WP_160037489.1 | WP_160037489.1 hypothetical protein |
| 1(1) | WP_112331585.1 | WP_112331585.1 serine/threonine-protein kinase |
| 1(1) | WP_143760315.1 | WP_143760315.1 S-layer homology domain-containing protein |
| 1(1) | WP_120462948.1 | WP_120462948.1 hypothetical protein |
| 1(1) | WP_283892185.1 | WP_283892185.1 tetratricopeptide repeat protein |
| 1(1) | WP_277166582.1 | WP_277166582.1 WXG100 family type VII secretion target |
| 1(1) | WP_078185404.1 | WP_078185404.1 protein kinase |
| 1(1) | WP_294386451.1 | WP_294386451.1 WXG100 family type VII secretion target |
| 1(1) | WP_098327467.1 | WP_098327467.1 type VII secretion protein EsaA |
| 1(1) | WP_003299545.1 | WP_003299545.1 hypothetical protein |
| 1(1) | WP_113753087.1 | WP_113753087.1 hypothetical protein |
| 1(1) | WP_212981565.1 | WP_212981565.1 hypothetical protein |
| 1(2) | WP_101300024.1 | WP_101300024.1 hypothetical protein |
| 1(1) | WP_068673699.1 | WP_068673699.1 hypothetical protein |
| 1(1) | WP_233697176.1 | WP_233697176.1 WXG100 family type VII secretion target |
| 1(1) | WP_098447527.1 | WP_098447527.1 bacteriocin-processing peptidase family protein |
| 1(1) | WP_243495557.1 | WP_243495557.1 beta-N-acetylhexosaminidase |
| 1(5) | WP_036609203.1 | WP_036609203.1 hypothetical protein |
| 1(1) | WP_049869167.1 | WP_049869167.1 DUF5050 domain-containing protein |
| 1(1) | WP_251417629.1 | WP_251417629.1 glycohydrolase toxin TNT-related protein |
| 1(1) | WP_098404091.1 | WP_098404091.1 hypothetical protein |
| 1(2) | WP_215570374.1 | WP_215570374.1 WXG100 family type VII secretion target |
| 1(1) | WP_138226022.1 | WP_138226022.1 hypothetical protein |
| 1(1) | WP_231094856.1 | WP_231094856.1 hypothetical protein |
| 1(1) | WP_079506402.1 | WP_079506402.1 pore-forming ESAT-6 family protein |
| 1(2) | WP_039273562.1 | WP_039273562.1 S-layer homology domain-containing protein |
| 1(1) | WP_247681891.1 | WP_247681891.1 RHS repeat-associated core domain-containing protein |
| 1(1) | WP_069011422.1 | WP_069011422.1 hypothetical protein |
| 1(1) | WP_270696896.1 | WP_270696896.1 tetratricopeptide repeat protein |
| 1(1) | WP_110822559.1 | WP_110822559.1 hypothetical protein |
| 1(1) | WP_289342261.1 | WP_289342261.1 WXG100 family type VII secretion target |
| 1(1) | WP_138322083.1 | WP_138322083.1 WXG100 family type VII secretion target |
| 1(1) | WP_150457762.1 | WP_150457762.1 hypothetical protein |
| 1(1) | WP_289483784.1 | WP_289483784.1 hypothetical protein |
| 1(1) | WP_099517104.1 | WP_099517104.1 hypothetical protein |
| 1(2) | WP_010963367.1 | WP_010963367.1 EsaB/YukD family protein |
| 1(1) | WP_245208023.1 | WP_245208023.1 DNA/RNA non-specific endonuclease |
| 1(2) | WP_010963725.1 | WP_010963725.1 Ig-like domain-containing protein |
| 1(1) | WP_307388963.1 | WP_307388963.1 hypothetical protein |
| 1(1) | WP_199925768.1 | WP_199925768.1 WXG100 family type VII secretion target |
| 1(1) | WP_108465349.1 | WP_108465349.1 WXG100 family type VII secretion target |
| 1(1) | WP_204445227.1 | WP_204445227.1 FHA domain-containing protein |
| 1(3) | WP_000883793.1 | WP_000883793.1 bacteriocin-processing peptidase family protein |
| 1(1) | WP_017692359.1 | WP_017692359.1 S-layer homology domain-containing protein |
| 1(1) | WP_186333848.1 | WP_186333848.1 pre-toxin TG domain-containing protein, partial |
| 1(1) | WP_242951947.1 | WP_242951947.1 serine/threonine-protein kinase |
| 1(1) | WP_255260043.1 | WP_255260043.1 hypothetical protein |
| 1(1) | WP_098991962.1 | WP_098991962.1 WXG100 family type VII secretion target |
| 1(1) | WP_072336489.1 | WP_072336489.1 glycohydrolase toxin TNT-related protein |
| 1(1) | WP_242261449.1 | WP_242261449.1 bacteriocin-processing peptidase family protein |
| 1(4) | WP_021291854.1 | WP_021291854.1 WXG100 family type VII secretion target |
| 1(1) | WP_079506384.1 | WP_079506384.1 LysM peptidoglycan-binding domain-containing protein |
| 1(1) | WP_016117669.1 | WP_016117669.1 T7SS effector LXG polymorphic toxin |
| 1(1) | WP_224839867.1 | WP_224839867.1 serine/threonine-protein kinase |
| 1(1) | WP_258427461.1 | WP_258427461.1 WG repeat-containing protein |
| 1(1) | WP_164515189.1 | WP_164515189.1 copper amine oxidase N-terminal domain-containing protein |
| 1(1) | WP_243500455.1 | WP_243500455.1 hypothetical protein |
| 1(1) | WP_098830139.1 | WP_098830139.1 WXG100 family type VII secretion target |
| 1(1) | WP_139236949.1 | WP_139236949.1 WXG100 family type VII secretion target |
| 1(1) | WP_076330723.1 | WP_076330723.1 hypothetical protein |
| 1(1) | WP_155609748.1 | WP_155609748.1 copper amine oxidase N-terminal domain-containing protein |
| 1(2) | WP_303660167.1 | WP_303660167.1 hypothetical protein |
| 1(1) | WP_077320202.1 | WP_077320202.1 EsaB/YukD family protein |
| 1(1) | WP_036626790.1 | WP_036626790.1 hypothetical protein |
| 1(1) | WP_064019320.1 | WP_064019320.1 hypothetical protein |
| 1(1) | WP_236417217.1 | WP_236417217.1 WXG100 family type VII secretion target |
| 1(2) | WP_112331580.1 | WP_112331580.1 pore-forming ESAT-6 family protein |
| 1(1) | WP_232274486.1 | WP_232274486.1 DUF6382 domain-containing protein |
| 1(1) | WP_066020853.1 | WP_066020853.1 glycosyltransferase |
| 1(1) | WP_076157833.1 | WP_076157833.1 WXG100 family type VII secretion target |
| 1(1) | WP_043067148.1 | WP_043067148.1 hypothetical protein |
| 1(3) | WP_000456052.1 | WP_000456052.1 CdiA family toxin C-terminal domain-containing protein |
| 1(1) | WP_094155417.1 | WP_094155417.1 S-layer homology domain-containing protein |
| 1(4) | WP_194333108.1 | WP_194333108.1 DNA/RNA non-specific endonuclease |
| 1(1) | WP_277713352.1 | WP_277713352.1 hypothetical protein |
| 1(1) | WP_068536386.1 | WP_068536386.1 hypothetical protein |
| 1(1) | WP_270574769.1 | WP_270574769.1 hypothetical protein |
| 1(1) | WP_137062797.1 | WP_137062797.1 hypothetical protein |
| 1(1) | WP_269058929.1 | WP_269058929.1 VWA domain-containing protein |
| 1(1) | WP_168419972.1 | WP_168419972.1 hypothetical protein |
| 1(1) | WP_159146499.1 | WP_159146499.1 C39 family peptidase |
| 1(1) | WP_036944542.1 | WP_036944542.1 EsaB/YukD family protein |
| 1(1) | WP_097846497.1 | WP_097846497.1 bacteriocin-processing peptidase family protein |
| 1(1) | WP_215185481.1 | WP_215185481.1 copper amine oxidase N-terminal domain-containing protein |
| 1(1) | WP_019912154.1 | WP_019912154.1 hypothetical protein |
| 1(1) | WP_148409437.1 | WP_148409437.1 DUF5050 domain-containing protein |
| 1(1) | WP_215559059.1 | WP_215559059.1 hypothetical protein |
| 1(1) | WP_274529062.1 | WP_274529062.1 S-layer homology domain-containing protein |
| 1(1) | WP_276867637.1 | WP_276867637.1 hypothetical protein |
| 1(1) | WP_066091992.1 | WP_066091992.1 DUF6382 domain-containing protein |
| 1(1) | WP_220559124.1 | WP_220559124.1 tetratricopeptide repeat protein |
| 1(1) | WP_136789994.1 | WP_136789994.1 hypothetical protein |
| 1(1) | WP_098961910.1 | WP_098961910.1 hypothetical protein |
| 1(1) | WP_136372957.1 | WP_136372957.1 pore-forming ESAT-6 family protein |

| | | |
|---|---|---|
| 1(1) | WP_102710719.1 | WP_102710719.1 EsaB/YukD family protein |
| 1(1) | WP_213143944.1 | WP_213143944.1 WXG100 family type VII secretion target |
| 1(2) | WP_251426362.1 | WP_251426362.1 pore-forming ESAT-6 family protein |
| 1(1) | WP_142327594.1 | WP_142327594.1 hypothetical protein |
| 1(2) | WP_038569548.1 | WP_038569548.1 DUF2974 domain-containing protein |
| 1(1) | WP_139359403.1 | WP_139359403.1 WXG100 family type VII secretion target, partial |
| 1(1) | WP_295628339.1 | WP_295628339.1 tetratricopeptide repeat protein |
| 1(1) | WP_270728520.1 | WP_270728520.1 tetratricopeptide repeat protein |
| 1(1) | WP_241929680.1 | WP_241929680.1 hypothetical protein |
| 1(1) | WP_184182977.1 | WP_184182977.1 hypothetical protein |
| 1(1) | WP_277150363.1 | WP_277150363.1 FHA domain-containing protein |
| 1(1) | WP_066091986.1 | WP_066091986.1 serine/threonine-protein kinase |
| 1(1) | WP_281961237.1 | WP_281961237.1 type VII secretion protein EsaA |
| 1(1) | WP_163768033.1 | WP_163768033.1 DUF2974 domain-containing protein |
| 1(1) | WP_198310830.1 | WP_198310830.1 pore-forming ESAT-6 family protein |
| 1(1) | WP_258126012.1 | WP_258126012.1 hypothetical protein |
| 1(1) | WP_098256950.1 | WP_098256950.1 WXG100 family type VII secretion target |
| 1(1) | WP_099361344.1 | WP_099361344.1 hypothetical protein |
| 1(1) | WP_063223268.1 | WP_063223268.1 bacteriocin-processing peptidase family protein |
| 1(1) | WP_036702406.1 | WP_036702406.1 hypothetical protein |
| 1(1) | WP_141097597.1 | WP_141097597.1 hypothetical protein |
| 1(1) | WP_308721987.1 | WP_308721987.1 hypothetical protein |
| 1(1) | WP_098952628.1 | WP_098952628.1 WXG100 family type VII secretion target |
| 1(1) | WP_078205519.1 | WP_078205519.1 bacteriocin-processing peptidase family protein |
| 1(2) | WP_119732858.1 | WP_119732858.1 type VII secretion target |
| 1(1) | WP_204412732.1 | WP_204412732.1 hypothetical protein |
| 1(2) | WP_243518755.1 | WP_243518755.1 hypothetical protein |
| 1(1) | WP_106496017.1 | WP_106496017.1 type VII secretion protein EsaA |
| 1(1) | WP_212981556.1 | WP_212981556.1 WXG100 family type VII secretion target |
| 1(1) | WP_159081747.1 | WP_159081747.1 hypothetical protein |
| 1(1) | WP_019392117.1 | WP_019392117.1 tetratricopeptide repeat protein |
| 1(9) | WP_063566325.1 | WP_063566325.1 hypothetical protein |
| 1(1) | WP_088032788.1 | WP_088032788.1 hypothetical protein |
| 1(1) | WP_099223954.1 | WP_099223954.1 EsaB/YukD family protein |
| 1(1) | WP_077832281.1 | WP_077832281.1 DUF6382 domain-containing protein |
| 1(1) | WP_099517118.1 | WP_099517118.1 LysM peptidoglycan-binding domain-containing protein |
| 1(1) | WP_138186695.1 | WP_138186695.1 WXG100 family type VII secretion target |
| 1(1) | WP_110934067.1 | WP_110934067.1 DUF2974 domain-containing protein |
| 1(1) | WP_068694716.1 | WP_068694716.1 WXG100 family type VII secretion target |
| 1(11) | WP_003335997.1 | WP_003335997.1 type VII secretion target |
| 1(1) | WP_190299131.1 | WP_190299131.1 S-layer homology domain-containing protein |
| 1(1) | WP_137698111.1 | WP_137698111.1 WXG100 family type VII secretion target |
| 1(1) | WP_270065055.1 | WP_270065055.1 WXG100 family type VII secretion target |
| 1(1) | WP_307601009.1 | WP_307601009.1 hypothetical protein |
| 1(1) | WP_098588023.1 | WP_098588023.1 bacteriocin-processing peptidase family protein |
| 1(2) | WP_271002891.1 | WP_271002891.1 HNH endonuclease |
| 1(1) | WP_036702415.1 | WP_036702415.1 DUF5050 domain-containing protein |
| 1(36) | WP_007430133.1 | WP_007430133.1 WXG100 family type VII secretion target |
| 1(2) | WP_010963735.1 | WP_010963735.1 WXG100 family type VII secretion target |
| 1(1) | WP_002148316.1 | WP_002148316.1 SseB family protein |
| 1(1) | WP_076271935.1 | WP_076271935.1 DUF5050 domain-containing protein |
| 1(1) | WP_229139926.1 | WP_229139926.1 serine/threonine-protein kinase |
| 1(1) | WP_076297592.1 | WP_076297592.1 DUF2974 domain-containing protein |
| 1(1) | WP_212693226.1 | WP_212693226.1 hypothetical protein |
| 1(2) | WP_058271111.1 | WP_058271111.1 WXG100 family type VII secretion target |
| 1(4) | WP_179160617.1 | WP_179160617.1 pore-forming ESAT-6 family protein |
| 1(1) | WP_223282659.1 | WP_223282659.1 DUF6382 domain-containing protein |
| 1(1) | WP_270728523.1 | WP_270728523.1 hypothetical protein |
| 1(1) | WP_307350420.1 | WP_307350420.1 stalk domain-containing protein |
| 1(1) | WP_155703933.1 | WP_155703933.1 WXG100 family type VII secretion target |
| 1(1) | WP_287600032.1 | WP_287600032.1 DUF6382 domain-containing protein |
| 1(1) | WP_113672694.1 | WP_113672694.1 WXG100 family type VII secretion target |
| 1(2) | WP_270994743.1 | WP_270994743.1 type VII secretion protein EsaA |
| 1(1) | WP_055105790.1 | WP_055105790.1 hypothetical protein |
| 1(1) | WP_155703945.1 | WP_155703945.1 S8 family serine peptidase |
| 1(1) | WP_186580083.1 | WP_186580083.1 WXG100 family type VII secretion target |
| 1(1) | WP_255443972.1 | WP_255443972.1 deaminase domain-containing protein |
| 1(1) | WP_227601482.1 | WP_227601482.1 hypothetical protein |
| 1(1) | WP_144479057.1 | WP_144479057.1 FHA domain-containing protein |
| 1(1) | WP_017689075.1 | WP_017689075.1 hypothetical protein |
| 1(1) | WP_227917893.1 | WP_227917893.1 hypothetical protein |
| 1(1) | WP_010496607.1 | WP_010496607.1 DUF2974 domain-containing protein |
| 1(1) | WP_185165632.1 | WP_185165632.1 WXG100 family type VII secretion target |
| 1(1) | WP_094876488.1 | WP_094876488.1 DUF5050 domain-containing protein |
| 1(1) | WP_307284678.1 | WP_307284678.1 WXG100 family type VII secretion target |
| 1(1) | WP_136372960.1 | WP_136372960.1 hypothetical protein |
| 1(1) | WP_286124787.1 | WP_286124787.1 WXG100 family type VII secretion target |
| 1(2) | WP_043924650.1 | WP_043924650.1 hypothetical protein |
| 1(1) | WP_290403367.1 | WP_290403367.1 hypothetical protein |
| 1(1) | WP_002148350.1 | WP_002148350.1 bacteriocin-processing peptidase family protein |
| 1(1) | WP_186580085.1 | WP_186580085.1 pore-forming ESAT-6 family protein |
| 1(1) | WP_145021732.1 | WP_145021732.1 hypothetical protein |
| 1(1) | WP_083180282.1 | WP_083180282.1 RHS repeat-associated core domain-containing protein |
| 1(1) | WP_047153273.1 | WP_047153273.1 hypothetical protein |
| 1(1) | WP_212930973.1 | WP_212930973.1 WG repeat-containing protein |
| 1(1) | WP_263703627.1 | WP_263703627.1 SseB family protein |
| 1(1) | WP_007430117.1 | WP_007430117.1 S-layer homology domain-containing protein |
| 1(4) | WP_228464160.1 | WP_228464160.1 pre-toxin TG domain-containing protein |
| 1(2) | WP_061678622.1 | WP_061678622.1 T7SS effector LXG polymorphic toxin |
| 1(1) | WP_097899492.1 | WP_097899492.1 hypothetical protein |
| 1(1) | WP_046522788.1 | WP_046522788.1 pore-forming ESAT-6 family protein |
| 1(1) | WP_109625239.1 | WP_109625239.1 DUF5050 domain-containing protein |
| 1(1) | WP_137698103.1 | WP_137698103.1 WXG100 family type VII secretion target |
| 1(2) | WP_076104575.1 | WP_076104575.1 hypothetical protein |
| 1(1) | WP_270691244.1 | WP_270691244.1 hypothetical protein |
| 1(1) | WP_297035102.1 | WP_297035102.1 hypothetical protein |
| 1(1) | WP_066020837.1 | WP_066020837.1 WXG100 family type VII secretion target |
| 1(1) | WP_029438096.1 | WP_029438096.1 hypothetical protein |
| 1(1) | WP_041964571.1 | WP_041964571.1 DUF6382 domain-containing protein |
| 1(1) | WP_007147994.1 | WP_007147994.1 hypothetical protein |
| 1(1) | WP_126665671.1 | WP_126665671.1 S-layer homology domain-containing protein |
| 1(1) | WP_035190312.1 | WP_035190312.1 WXG100 family type VII secretion target |
| 1(1) | WP_239617469.1 | WP_239617469.1 S-layer homology domain-containing protein |
| 1(1) | WP_195431285.1 | WP_195431285.1 Ig-like domain repeat protein, partial |

1(1) WP_226602910.1 WP_226602910.1 WXG100 family type VII secretion target
1(2) WP_133161642.1 WP_133161642.1 EsaB/YukD family protein
1(1) WP_064798160.1 WP_064798160.1 WXG100 family type VII secretion target
1(1) WP_168976286.1 WP_168976286.1 tetratricopeptide repeat protein
1(1) WP_199613903.1 WP_199613903.1 DUF5050 domain-containing protein
1(1) WP_236334334.1 WP_236334334.1 DUF5050 domain-containing protein
1(1) WP_291647938.1 WP_291647938.1 GBS Bsp-like repeat-containing protein
1(1) WP_199613901.1 WP_199613901.1 WXG100 family type VII secretion target
1(1) WP_270065051.1 WP_270065051.1 serine/threonine-protein kinase
1(3) WP_088609110.1 WP_088609110.1 bacteriocin-processing peptidase family protein
1(1) WP_223067437.1 WP_223067437.1 WXG100 family type VII secretion target
1(1) WP_009590694.1 WP_009590694.1 hypothetical protein
1(1) WP_137078366.1 WP_137078366.1 hypothetical protein
1(1) WP_213143945.1 WP_213143945.1 WXG100 family type VII secretion target
1(1) WP_000391134.1 WP_000391134.1 hypothetical protein
1(2) WP_010963376.1 WP_010963376.1 WXG100 family type VII secretion target
1(6) WP_291288262.1 WP_291288262.1 WXG100 family type VII secretion target
1(1) WP_283910925.1 WP_283910925.1 WXG100 family type VII secretion target
1(1) WP_044037280.1 WP_044037280.1 WXG100 family type VII secretion target
1(1) WP_251417670.1 WP_251417670.1 copper amine oxidase N-terminal domain-containing protein
1(2) WP_101300050.1 WP_101300050.1 WXG100 family type VII secretion target
1(1) WP_273321541.1 WP_273321541.1 hypothetical protein
1(1) WP_199613904.1 WP_199613904.1 EsaB/YukD family protein
1(1) WP_272801671.1 WP_272801671.1 hypothetical protein
1(1) WP_251456524.1 WP_251456524.1 hypothetical protein
1(1) WP_061184804.1 WP_061184804.1 bacteriocin-processing peptidase family protein
1(1) WP_215572746.1 WP_215572746.1 tetratricopeptide repeat protein
1(1) WP_087442412.1 WP_087442412.1 WXG100 family type VII secretion target
1(1) WP_169238723.1 WP_169238723.1 protein kinase
1(2) WP_200372547.1 WP_200372547.1 pre-toxin TG domain-containing protein
1(1) WP_197937380.1 WP_197937380.1 hypothetical protein
1(1) WP_098856762.1 WP_098856762.1 bacteriocin-processing peptidase family protein
1(1) WP_240379736.1 WP_240379736.1 WXG100 family type VII secretion target
1(1) WP_073540571.1 WP_073540571.1 bacteriocin-processing peptidase family protein
1(1) WP_304216882.1 WP_304216882.1 hypothetical protein
1(1) WP_073008269.1 WP_073008269.1 serine/threonine-protein kinase
1(1) WP_076101571.1 WP_076101571.1 DUF2974 domain-containing protein
1(1) WP_211745838.1 WP_211745838.1 serine hydrolase domain-containing protein
1(1) WP_251480992.1 WP_251480992.1 LysM peptidoglycan-binding domain-containing protein
1(1) WP_244719541.1 WP_244719541.1 protease complex subunit PrcB family protein
1(1) WP_209861624.1 WP_209861624.1 glycosyl hydrolase family 18 protein
1(2) WP_127812903.1 WP_127812903.1 tetratricopeptide repeat protein
1(2) WP_234288979.1 WP_234288979.1 pre-toxin TG domain-containing protein
1(2) WP_098678280.1 WP_098678280.1 bacteriocin-processing peptidase family protein
1(2) WP_145406488.1 WP_145406488.1 WXG100 family type VII secretion target
1(3) WP_094870399.1 WP_094870399.1 hypothetical protein
1(3) WP_094870529.1 WP_094870529.1 copper amine oxidase N-terminal domain-containing protein
1(2) WP_185463563.1 WP_185463563.1 T7SS effector LXG polymorphic toxin
1(2) WP_058271107.1 WP_058271107.1 WXG100 family type VII secretion target
1(1) WP_213117484.1 WP_213117484.1 WXG100 family type VII secretion target
1(1) WP_213426991.1 WP_213426991.1 WXG100 family type VII secretion target
1(4) WP_026572740.1 WP_026572740.1 WXG100 family type VII secretion target
1(1) WP_145156803.1 WP_145156803.1 S-layer homology domain-containing protein
1(1) WP_254366009.1 WP_254366009.1 pore-forming ESAT-6 family protein
1(1) WP_010250285.1 WP_010250285.1 copper amine oxidase N-terminal domain-containing protein
1(1) WP_289607495.1 WP_289607495.1 pore-forming ESAT-6 family protein
1(1) WP_144564265.1 WP_144564265.1 bacteriocin-processing peptidase family protein
1(1) WP_227558283.1 WP_227558283.1 tetratricopeptide repeat protein
1(1) WP_160037482.1 WP_160037482.1 hypothetical protein
1(1) WP_251426365.1 WP_251426365.1 DUF2974 domain-containing protein
1(5) WP_056700792.1 WP_056700792.1 WXG100 family type VII secretion target
1(1) WP_243500449.1 WP_243500449.1 EsaB/YukD family protein
1(1) WP_251378709.1 WP_251378709.1 S-layer homology domain-containing protein
1(1) WP_283892178.1 WP_283892178.1 hypothetical protein
1(1) WP_077833105.1 WP_077833105.1 WXG100 family type VII secretion target
1(1) WP_154985826.1 WP_154985826.1 hypothetical protein
1(1) WP_210364453.1 WP_210364453.1 DUF2974 domain-containing protein
1(1) WP_095266194.1 WP_095266194.1 hypothetical protein
1(1) WP_155703925.1 WP_155703925.1 hypothetical protein
1(1) WP_082790192.1 WP_082790192.1 DUF1308 domain-containing protein
1(1) WP_094870530.1 WP_094870530.1 DUF2974 domain-containing protein
1(1) WP_277157952.1 WP_277157952.1 serine/threonine-protein kinase
1(1) WP_000392230.1 WP_000392230.1 hypothetical protein
1(1) WP_295632013.1 WP_295632013.1 flavodoxin
1(1) WP_099354900.1 WP_099354900.1 hypothetical protein
1(1) WP_133378770.1 WP_133378770.1 hypothetical protein
1(5) WP_019392114.1 WP_019392114.1 hypothetical protein
1(2) WP_216518256.1 WP_216518256.1 hypothetical protein
1(1) WP_181603074.1 WP_181603074.1 hypothetical protein
1(1) WP_036609196.1 WP_036609196.1 membrane protein
1(1) WP_126996319.1 WP_126996319.1 hypothetical protein
1(1) WP_025716877.1 WP_025716877.1 membrane protein
1(1) WP_309421460.1 WP_309421460.1 type VII secretion protein EsaA, partial
1(1) WP_293755528.1 WP_293755528.1 hypothetical protein
1(1) WP_119794365.1 WP_119794365.1 hypothetical protein
1(1) WP_232339739.1 WP_232339739.1 WXG100 family type VII secretion target
1(1) WP_260983630.1 WP_260983630.1 WXG100 family type VII secretion target
1(1) WP_241993932.1 WP_241993932.1 pore-forming ESAT-6 family protein
1(1) WP_200633297.1 WP_200633297.1 WXG100 family type VII secretion target
1(1) WP_236804412.1 WP_236804412.1 serine hydrolase domain-containing protein
1(1) WP_051317150.1 WP_051317150.1 tetratricopeptide repeat protein
1(1) WP_301167505.1 WP_301167505.1 tetratricopeptide repeat protein
1(1) WP_076157838.1 WP_076157838.1 DUF5050 domain-containing protein
1(1) WP_181603072.1 WP_181603072.1 tetratricopeptide repeat protein
1(1) WP_215559057.1 WP_215559057.1 bacteriocin-processing peptidase family protein
1(1) WP_013490136.1 WP_013490136.1 hypothetical protein
1(1) WP_102997918.1 WP_102997918.1 hypothetical protein
1(1) WP_307389000.1 WP_307389000.1 T7SS effector LXG polymorphic toxin
1(1) WP_103047826.1 WP_103047826.1 hypothetical protein
1(1) WP_255294005.1 WP_255294005.1 deaminase domain-containing protein
1(1) WP_110931544.1 WP_110931544.1 hypothetical protein

| | | |
|---|---|---|
| 1(1) | WP_077832289.1 | WP_077832289.1 WXG100 family type VII secretion target |
| 1(1) | WP_143316573.1 | WP_143316573.1 serine/threonine-protein kinase |
| 1(1) | WP_189018628.1 | WP_189018628.1 leucine-rich repeat domain-containing protein |
| 1(1) | WP_273321560.1 | WP_273321560.1 hypothetical protein |
| 1(1) | WP_291289941.1 | WP_291289941.1 serine/threonine-protein kinase |
| 1(3) | WP_021253118.1 | WP_021253118.1 WXG100 family type VII secretion target |
| 1(1) | WP_215591203.1 | WP_215591203.1 WXG100 family type VII secretion target |
| 1(1) | WP_247681890.1 | WP_247681890.1 RHS repeat-associated core domain-containing protein |
| 1(1) | WP_000404138.1 | WP_000404138.1 WXG100 family type VII secretion target |
| 1(1) | WP_207738532.1 | WP_207738532.1 AHH domain-containing protein, partial |
| 1(1) | WP_308912806.1 | WP_308912806.1 protein kinase |
| 1(1) | WP_244719670.1 | WP_244719670.1 stalk domain-containing protein |
| 1(1) | WP_099853197.1 | WP_099853197.1 hypothetical protein |
| 1(1) | WP_211022786.1 | WP_211022786.1 WXG100 family type VII secretion target |
| 1(1) | WP_079506420.1 | WP_079506420.1 RHS repeat-associated core domain-containing protein |
| 1(1) | WP_010346281.1 | WP_010346281.1 membrane protein |
| 1(1) | WP_052520275.1 | WP_052520275.1 tetratricopeptide repeat protein |
| 1(1) | WP_215570372.1 | WP_215570372.1 SseB family protein |
| 1(1) | WP_112331584.1 | WP_112331584.1 chitobiase/beta-hexosaminidase C-terminal domain-containing protein |
| 1(1) | WP_254349527.1 | WP_254349527.1 hypothetical protein |
| 1(1) | WP_209365679.1 | WP_209365679.1 EsaB/YukD family protein |
| 1(1) | WP_034526646.1 | WP_034526646.1 FHA domain-containing protein |
| 1(1) | WP_018883270.1 | WP_018883270.1 hypothetical protein |
| 1(1) | WP_076157831.1 | WP_076157831.1 peptidoglycan-binding protein |
| 1(1) | WP_155703929.1 | WP_155703929.1 hypothetical protein |
| 1(1) | WP_068536390.1 | WP_068536390.1 hypothetical protein |
| 1(1) | WP_251674651.1 | WP_251674651.1 WXG100 family type VII secretion target |
| 1(1) | WP_000360011.1 | WP_000360011.1 T7SS effector LXG polymorphic toxin |
| 1(1) | WP_291289931.1 | WP_291289931.1 DUF2974 domain-containing protein |
| 1(1) | WP_236334318.1 | WP_236334318.1 copper amine oxidase N-terminal domain-containing protein |
| 1(1) | WP_251481003.1 | WP_251481003.1 serine/threonine-protein kinase |
| 1(1) | WP_211022781.1 | WP_211022781.1 hypothetical protein |
| 1(1) | WP_109999891.1 | WP_109999891.1 hypothetical protein |
| 1(1) | WP_098607334.1 | WP_098607334.1 EsaB/YukD family protein |
| 1(1) | WP_046522792.1 | WP_046522792.1 Ig-like domain-containing protein |
| 1(1) | WP_226086177.1 | WP_226086177.1 LysM peptidoglycan-binding domain-containing protein |
| 1(1) | WP_260443238.1 | WP_260443238.1 hypothetical protein |
| 1(1) | WP_139265505.1 | WP_139265505.1 hypothetical protein |
| 1(1) | WP_242264956.1 | WP_242264956.1 hypothetical protein |
| 1(1) | WP_181603073.1 | WP_181603073.1 WXG100 family type VII secretion target |
| 1(1) | WP_242475736.1 | WP_242475736.1 deaminase domain-containing protein |
| 1(1) | WP_289607502.1 | WP_289607502.1 carboxypeptidase-like regulatory domain-containing protein |
| 1(1) | WP_210095106.1 | WP_210095106.1 serine/threonine-protein kinase |
| 1(1) | WP_143315258.1 | WP_143315258.1 hypothetical protein |
| 1(1) | WP_042203646.1 | WP_042203646.1 hypothetical protein |
| 1(1) | WP_208203478.1 | WP_208203478.1 S-layer homology domain-containing protein |
| 1(1) | WP_090993623.1 | WP_090993623.1 hypothetical protein |
| 1(1) | WP_223067441.1 | WP_223067441.1 hypothetical protein |
| 1(2) | WP_219872457.1 | WP_219872457.1 WXG100 family type VII secretion target |
| 1(1) | WP_068496247.1 | WP_068496247.1 hypothetical protein |
| 1(1) | WP_275216786.1 | WP_275216786.1 type VII secretion protein EsaA |
| 1(1) | WP_280788631.1 | WP_280788631.1 hypothetical protein |
| 1(1) | WP_137698107.1 | WP_137698107.1 DUF2974 domain-containing protein |
| 1(1) | WP_197256864.1 | WP_197256864.1 WXG100 family type VII secretion target |
| 1(1) | WP_044037264.1 | WP_044037264.1 WXG100 family type VII secretion target |
| 1(1) | WP_249897314.1 | WP_249897314.1 EsaB/YukD family protein |
| 1(1) | WP_290407042.1 | WP_290407042.1 WXG100 family type VII secretion target |
| 1(1) | WP_090761815.1 | WP_090761815.1 WXG100 family type VII secretion target |
| 1(1) | WP_284036495.1 | WP_284036495.1 FHA domain-containing protein |
| 1(2) | WP_110899369.1 | WP_110899369.1 WXG100 family type VII secretion target |
| 1(1) | WP_140160092.1 | WP_140160092.1 hypothetical protein |
| 1(1) | WP_125084784.1 | WP_125084784.1 EsaB/YukD family protein |
| 1(2) | WP_130777023.1 | WP_130777023.1 WXG100 family type VII secretion target |
| 1(2) | WP_143801059.1 | WP_143801059.1 hypothetical protein |
| 1(1) | WP_215032123.1 | WP_215032123.1 DUF2974 domain-containing protein |
| 1(1) | WP_144061518.1 | WP_144061518.1 hypothetical protein |
| 1(1) | WP_068793928.1 | WP_068793928.1 WG repeat-containing protein |
| 1(1) | WP_223592674.1 | WP_223592674.1 Ig-like domain-containing protein |
| 1(1) | WP_048826573.1 | WP_048826573.1 WXG100 family type VII secretion target |
| 1(1) | WP_048749332.1 | WP_048749332.1 hypothetical protein |
| 1(1) | WP_128705703.1 | WP_128705703.1 DUF5050 domain-containing protein |
| 1(1) | WP_237100357.1 | WP_237100357.1 copper amine oxidase N-terminal domain-containing protein |
| 1(4) | WP_038406451.1 | WP_038406451.1 Ig-like domain-containing protein |
| 1(1) | WP_220031158.1 | WP_220031158.1 S-layer homology domain-containing protein |
| 1(1) | WP_199613900.1 | WP_199613900.1 WXG100 family type VII secretion target |
| 1(1) | WP_210087112.1 | WP_210087112.1 WXG100 family type VII secretion target |
| 1(1) | WP_207706235.1 | WP_207706235.1 hypothetical protein, partial |
| 1(1) | WP_045518282.1 | WP_045518282.1 DUF2974 domain-containing protein |
| 1(1) | WP_303942528.1 | WP_303942528.1 hypothetical protein |
| 1(1) | WP_002148319.1 | WP_002148319.1 tetratricopeptide repeat protein |
| 1(1) | WP_150158011.1 | WP_150158011.1 T7SS effector LXG polymorphic toxin |
| 1(2) | WP_149051230.1 | WP_149051230.1 LXG domain-containing protein |
| 1(1) | WP_195325252.1 | WP_195325252.1 tetratricopeptide repeat protein |
| 1(1) | WP_152392586.1 | WP_152392586.1 hypothetical protein |
| 1(1) | WP_226679096.1 | WP_226679096.1 Ig-like domain-containing protein |
| 1(1) | WP_293654397.1 | WP_293654397.1 WXG100 family type VII secretion target |
| 1(1) | WP_215032106.1 | WP_215032106.1 Ig-like domain-containing protein |
| 1(1) | WP_288161213.1 | WP_288161213.1 pore-forming ESAT-6 family protein |
| 1(1) | WP_247681893.1 | WP_247681893.1 hypothetical protein |
| 1(1) | WP_010346299.1 | WP_010346299.1 S-layer homology domain-containing protein |
| 1(1) | WP_270472004.1 | WP_270472004.1 WXG100 family type VII secretion target |
| 1(1) | WP_277473579.1 | WP_277473579.1 type VII secretion protein EsaA |
| 1(1) | WP_199677614.1 | WP_199677614.1 hypothetical protein |
| 1(1) | WP_078399858.1 | WP_078399858.1 type VII secretion protein EsaA |
| 1(1) | WP_283904335.1 | WP_283904335.1 hypothetical protein |
| 1(2) | WP_076311051.1 | WP_076311051.1 hypothetical protein |
| 1(1) | WP_191272856.1 | WP_191272856.1 pore-forming ESAT-6 family protein |
| 1(1) | WP_303942534.1 | WP_303942534.1 hypothetical protein |
| 1(1) | WP_144611190.1 | WP_144611190.1 SMI1/KNR4 family protein |
| 1(1) | WP_026882399.1 | WP_026882399.1 DUF6382 domain-containing protein |
| 1(6) | WP_116334079.1 | WP_116334079.1 hypothetical protein |

| | | |
|---|---|---|
| 1(1) | WP_270065053.1 | WP_270065053.1 DUF6382 domain-containing protein |
| 1(1) | WP_078181468.1 | WP_078181468.1 tetratricopeptide repeat protein |
| 1(1) | WP_211023038.1 | WP_211023038.1 pre-toxin TG domain-containing protein |
| 1(1) | WP_144479068.1 | WP_144479068.1 hypothetical protein |
| 1(3) | WP_104030439.1 | WP_104030439.1 type VII secretion target |
| 1(1) | WP_307284684.1 | WP_307284684.1 hypothetical protein |
| 1(1) | WP_215591197.1 | WP_215591197.1 bacteriocin-processing peptidase family protein |
| 1(1) | WP_169238722.1 | WP_169238722.1 FHA domain-containing protein |
| 1(1) | WP_049869171.1 | WP_049869171.1 copper amine oxidase N-terminal domain-containing protein |
| 1(5) | WP_021253121.1 | WP_021253121.1 hypothetical protein |
| 1(1) | WP_098592732.1 | WP_098592732.1 type VII secretion protein EsaA |
| 1(1) | WP_216518248.1 | WP_216518248.1 S-layer homology domain-containing protein |
| 1(1) | WP_040360539.1 | WP_040360539.1 hypothetical protein |
| 1(1) | WP_169505050.1 | WP_169505050.1 hypothetical protein |
| 1(1) | WP_101356552.1 | WP_101356552.1 EsaB/YukD family protein |
| 1(1) | WP_282103169.1 | WP_282103169.1 EsaB/YukD family protein |
| 1(3) | WP_076104574.1 | WP_076104574.1 hypothetical protein |
| 1(1) | WP_141436469.1 | WP_141436469.1 hypothetical protein |
| 1(1) | WP_068536377.1 | WP_068536377.1 WXG100 family type VII secretion target |
| 1(1) | WP_181737025.1 | WP_181737025.1 WXG100 family type VII secretion target |
| 1(1) | WP_229140039.1 | WP_229140039.1 Ig-like domain-containing protein |
| 1(1) | WP_319005030.1 | WP_319005030.1 protein kinase |
| 1(2) | WP_200152289.1 | WP_200152289.1 bacteriocin-processing peptidase family protein |
| 1(2) | WP_053325280.1 | WP_053325280.1 hypothetical protein |
| 1(1) | WP_134909999.1 | WP_134909999.1 WXG100 family type VII secretion target |
| 1(1) | WP_212981568.1 | WP_212981568.1 hypothetical protein |
| 1(3) | WP_010346287.1 | WP_010346287.1 hypothetical protein |
| 1(1) | WP_076286998.1 | WP_076286998.1 hypothetical protein |
| 1(1) | WP_198310827.1 | WP_198310827.1 protein kinase |
| 1(1) | WP_238333009.1 | WP_238333009.1 WG repeat-containing protein |
| 1(1) | WP_262800524.1 | WP_262800524.1 tetratricopeptide repeat protein |
| 1(1) | WP_249897318.1 | WP_249897318.1 WXG100 family type VII secretion target |
| 1(1) | WP_051569331.1 | WP_051569331.1 DUF5050 domain-containing protein |
| 1(1) | WP_116353251.1 | WP_116353251.1 Mbeg1-like protein |
| 1(2) | WP_007430129.1 | WP_007430129.1 hypothetical protein |
| 1(1) | WP_307589594.1 | WP_307589594.1 pore-forming ESAT-6 family protein |
| 1(1) | WP_098150443.1 | WP_098150443.1 hypothetical protein |
| 1(1) | WP_149871680.1 | WP_149871680.1 WXG100 family type VII secretion target |
| 1(1) | WP_148466640.1 | WP_148466640.1 hypothetical protein |
| 1(1) | WP_127570433.1 | WP_127570433.1 hypothetical protein |
| 1(1) | WP_187374157.1 | WP_187374157.1 serine/threonine-protein kinase |
| 1(1) | WP_058271112.1 | WP_058271112.1 hypothetical protein |
| 1(1) | WP_211022787.1 | WP_211022787.1 hypothetical protein |
| 1(5) | WP_001142463.1 | WP_001142463.1 hypothetical protein |
| 1(1) | WP_024027634.1 | WP_024027634.1 DUF6382 domain-containing protein |
| 1(1) | WP_125084775.1 | WP_125084775.1 DUF2974 domain-containing protein |
| 1(1) | WP_106496019.1 | WP_106496019.1 hypothetical protein |
| 1(2) | WP_098538859.1 | WP_098538859.1 tetratricopeptide repeat protein |
| 1(1) | WP_126996323.1 | WP_126996323.1 DL-endopeptidase inhibitor IseA family protein |
| 1(1) | WP_098945325.1 | WP_098945325.1 bacteriocin-processing peptidase family protein |
| 1(1) | WP_104065528.1 | WP_104065528.1 WG repeat-containing protein |
| 1(1) | WP_238655347.1 | WP_238655347.1 EsaB/YukD family protein |
| 1(1) | WP_085499084.1 | WP_085499084.1 copper amine oxidase N-terminal domain-containing protein |
| 1(1) | WP_246860932.1 | WP_246860932.1 LysM peptidoglycan-binding domain-containing protein |
| 1(1) | WP_241840573.1 | WP_241840573.1 hypothetical protein |
| 1(1) | WP_238655350.1 | WP_238655350.1 WXG100 family type VII secretion target |
| 1(1) | WP_058829513.1 | WP_058829513.1 hypothetical protein |
| 1(1) | WP_297183834.1 | WP_297183834.1 FHA domain-containing protein |
| 1(1) | WP_148450938.1 | WP_148450938.1 hypothetical protein |
| 1(1) | WP_015595760.1 | WP_015595760.1 hypothetical protein |
| 1(1) | WP_128631285.1 | WP_128631285.1 EsaB/YukD family protein |
| 1(1) | WP_279830837.1 | WP_279830837.1 hypothetical protein |
| 1(1) | WP_018664837.1 | WP_018664837.1 WXG100 family type VII secretion target |
| 1(1) | WP_099361337.1 | WP_099361337.1 DUF2974 domain-containing protein |
| 1(1) | WP_262725734.1 | WP_262725734.1 bacteriocin-processing peptidase family protein |
| 1(1) | WP_210364464.1 | WP_210364464.1 DUF2974 domain-containing protein |
| 1(2) | WP_044792559.1 | WP_044792559.1 tetratricopeptide repeat protein |
| 1(1) | WP_277156191.1 | WP_277156191.1 hypothetical protein |
| 1(1) | WP_019120052.1 | WP_019120052.1 WXG100 family type VII secretion target |
| 1(1) | WP_290401999.1 | WP_290401999.1 hypothetical protein |
| 1(1) | WP_043326247.1 | WP_043326247.1 WXG100 family type VII secretion target |
| 1(1) | WP_290409172.1 | WP_290409172.1 hypothetical protein |
| 1(1) | WP_098747409.1 | WP_098747409.1 hypothetical protein |
| 1(1) | WP_182301827.1 | WP_182301827.1 hypothetical protein |
| 1(1) | WP_098230734.1 | WP_098230734.1 bacteriocin-processing peptidase family protein |
| 1(4) | WP_077832293.1 | WP_077832293.1 WXG100 family type VII secretion target |
| 1(1) | WP_023521815.1 | WP_023521815.1 WXG100 family type VII secretion target |
| 1(1) | WP_251481005.1 | WP_251481005.1 Ig-like domain-containing protein |
| 1(1) | WP_103655289.1 | WP_103655289.1 WXG100 family type VII secretion target |
| 1(1) | WP_256872560.1 | WP_256872560.1 hypothetical protein |
| 1(2) | WP_143881073.1 | WP_143881073.1 tetratricopeptide repeat protein |
| 1(1) | WP_246120603.1 | WP_246120603.1 hypothetical protein |
| 1(1) | WP_033693878.1 | WP_033693878.1 type VII secretion protein EsaA, partial |
| 1(1) | WP_053361019.1 | WP_053361019.1 FHA domain-containing protein |
| 1(1) | WP_073008298.1 | WP_073008298.1 phosphodiester glycosidase family protein |
| 1(2) | WP_212956326.1 | WP_212956326.1 hypothetical protein |
| 1(1) | WP_283910926.1 | WP_283910926.1 WXG100 family type VII secretion target |
| 1(1) | WP_161109321.1 | WP_161109321.1 WXG100 family type VII secretion target, partial |
| 1(1) | WP_116353247.1 | WP_116353247.1 DUF6382 domain-containing protein |
| 1(1) | WP_128631274.1 | WP_128631274.1 copper amine oxidase N-terminal domain-containing protein |
| 1(2) | WP_025679790.1 | WP_025679790.1 S-layer homology domain-containing protein |
| 1(1) | WP_281068145.1 | WP_281068145.1 S-layer homology domain-containing protein |
| 1(1) | WP_212981560.1 | WP_212981560.1 EsaB/YukD family protein |
| 1(3) | WP_034773559.1 | WP_034773559.1 WXG100 family type VII secretion target |
| 1(1) | WP_242216126.1 | WP_242216126.1 hypothetical protein |
| 1(1) | WP_041184359.1 | WP_041184359.1 hypothetical protein |
| 1(2) | WP_144469660.1 | WP_144469660.1 hypothetical protein |
| 1(2) | WP_047909611.1 | WP_047909611.1 membrane protein |
| 1(1) | WP_308912804.1 | WP_308912804.1 Ig-like domain-containing protein |
| 1(1) | WP_002158055.1 | WP_002158055.1 bacteriocin-processing peptidase family protein |
| 1(1) | WP_221697948.1 | WP_221697948.1 hypothetical protein |

| | | |
|---|---|---|
| 1(1) | WP_142330022.1 | WP_142330022.1 WXG100 family type VII secretion target, partial |
| 1(1) | WP_155618100.1 | WP_155618100.1 serine/threonine protein kinase |
| 1(1) | WP_223068034.1 | WP_223068034.1 hypothetical protein |
| 1(1) | WP_211023024.1 | WP_211023024.1 WXG100 family type VII secretion target |
| 1(1) | WP_144660361.1 | WP_144660361.1 tetratricopeptide repeat protein |
| 1(1) | WP_207737247.1 | WP_207737247.1 hypothetical protein |
| 1(1) | WP_237690859.1 | WP_237690859.1 EsaB/YukD family protein |
| 1(1) | WP_211082917.1 | WP_211082917.1 hypothetical protein |
| 1(1) | WP_036575540.1 | WP_036575540.1 hypothetical protein |
| 1(1) | WP_007083296.1 | WP_007083296.1 protein kinase |
| 1(1) | WP_102997911.1 | WP_102997911.1 S-layer homology domain-containing protein |
| 1(6) | WP_185340297.1 | WP_185340297.1 hypothetical protein |
| 1(1) | WP_249897322.1 | WP_249897322.1 copper amine oxidase N-terminal domain-containing protein |
| 1(2) | WP_038569544.1 | WP_038569544.1 WXG100 family type VII secretion target |
| 1(2) | WP_273191254.1 | WP_273191254.1 hypothetical protein |
| 1(2) | WP_088609116.1 | WP_088609116.1 WXG100 family type VII secretion target |
| 1(1) | WP_251456526.1 | WP_251456526.1 tetratricopeptide repeat protein |
| 1(1) | WP_258417130.1 | WP_258417130.1 ESX-1 secretion-associated protein |
| 1(1) | WP_091017193.1 | WP_091017193.1 hypothetical protein |
| 1(1) | WP_125084773.1 | WP_125084773.1 DUF2974 domain-containing protein |
| 1(2) | WP_228464393.1 | WP_228464393.1 hypothetical protein |
| 1(1) | WP_262719887.1 | WP_262719887.1 hypothetical protein |
| 1(1) | WP_307192895.1 | WP_307192895.1 pore-forming ESAT-6 family protein |
| 1(1) | WP_079694647.1 | WP_079694647.1 hypothetical protein |
| 1(1) | WP_098365936.1 | WP_098365936.1 bacteriocin-processing peptidase family protein |
| 1(1) | WP_000804098.1 | WP_000804098.1 DNA/RNA non-specific endonuclease |
| 1(1) | WP_134750628.1 | WP_134750628.1 hypothetical protein |
| 1(1) | WP_235676370.1 | WP_235676370.1 hypothetical protein |
| 1(1) | WP_291289932.1 | WP_291289932.1 DUF2974 domain-containing protein |
| 1(1) | WP_278431123.1 | WP_278431123.1 WG repeat-containing protein |
| 1(2) | WP_133332976.1 | WP_133332976.1 hypothetical protein |
| 1(1) | WP_018883272.1 | WP_018883272.1 WXG100 family type VII secretion target |
| 1(1) | WP_188535067.1 | WP_188535067.1 EsaB/YukD family protein |
| 1(1) | WP_102710726.1 | WP_102710726.1 DUF2974 domain-containing protein |
| 1(1) | WP_144025354.1 | WP_144025354.1 WXG100 family type VII secretion target |
| 1(1) | WP_215601066.1 | WP_215601066.1 hypothetical protein |
| 1(13) | WP_000506911.1 | WP_000506911.1 tetratricopeptide repeat protein |
| 1(1) | WP_263324552.1 | WP_263324552.1 WXG100 family type VII secretion target |
| 1(1) | WP_257808053.1 | WP_257808053.1 hypothetical protein |
| 1(1) | WP_291630884.1 | WP_291630884.1 WXG100 family type VII secretion target |
| 1(2) | WP_002158039.1 | WP_002158039.1 SseB family protein |
| 1(1) | WP_145329908.1 | WP_145329908.1 hypothetical protein |
| 1(1) | WP_210087107.1 | WP_210087107.1 S-layer homology domain-containing protein |
| 1(1) | WP_235286206.1 | WP_235286206.1 hypothetical protein |
| 1(1) | WP_052016758.1 | WP_052016758.1 protein kinase family protein |
| 1(1) | WP_138755920.1 | WP_138755920.1 WXG100 family type VII secretion target |
| 1(2) | WP_024632683.1 | WP_024632683.1 WXG100 family type VII secretion target |
| 1(2) | WP_011700921.1 | WP_011700921.1 WXG100 family type VII secretion target |
| 1(1) | WP_094699658.1 | WP_094699658.1 WG repeat-containing protein |
| 1(1) | WP_169028890.1 | WP_169028890.1 hypothetical protein |
| 1(1) | WP_139186929.1 | WP_139186929.1 hypothetical protein |
| 1(1) | WP_062239868.1 | WP_062239868.1 EsaB/YukD family protein |
| 1(1) | WP_223592688.1 | WP_223592688.1 pore-forming ESAT-6 family protein |
| 1(1) | WP_141529615.1 | WP_141529615.1 hypothetical protein |
| 1(1) | WP_209872747.1 | WP_209872747.1 hypothetical protein |
| 1(1) | WP_094092622.1 | WP_094092622.1 WXG100 family type VII secretion target |
| 1(1) | WP_077833112.1 | WP_077833112.1 FHA domain-containing protein |
| 1(3) | WP_018670518.1 | WP_018670518.1 type VII secretion target |
| 1(1) | WP_165097616.1 | WP_165097616.1 DUF5050 domain-containing protein |
| 1(1) | WP_166244074.1 | WP_166244074.1 EsaB/YukD family protein |
| 1(1) | WP_120462947.1 | WP_120462947.1 copper amine oxidase N-terminal domain-containing protein |
| 1(1) | WP_273191252.1 | WP_273191252.1 hypothetical protein |
| 1(1) | WP_206094034.1 | WP_206094034.1 serine/threonine-protein kinase |
| 1(35) | WP_000136048.1 | WP_000136048.1 WXG100 family type VII secretion target |
| 1(1) | WP_251375548.1 | WP_251375548.1 serine/threonine-protein kinase |
| 1(1) | WP_069193873.1 | WP_069193873.1 EsaB/YukD family protein |
| 1(1) | WP_141669097.1 | WP_141669097.1 WXG100 family type VII secretion target, partial |
| 1(1) | WP_169744749.1 | WP_169744749.1 AAA family ATPase |
| 1(2) | WP_069080933.1 | WP_069080933.1 S-layer homology domain-containing protein |
| 1(1) | WP_243500457.1 | WP_243500457.1 WXG100 family type VII secretion target |
| 1(1) | WP_270728515.1 | WP_270728515.1 hypothetical protein |
| 1(1) | WP_282742032.1 | WP_282742032.1 DUF2974 domain-containing protein |
| 1(1) | WP_127484347.1 | WP_127484347.1 pore-forming ESAT-6 family protein |
| 1(1) | WP_090239241.1 | WP_090239241.1 tetratricopeptide repeat protein |
| 1(1) | WP_150364184.1 | WP_150364184.1 hypothetical protein |
| 1(3) | WP_094870401.1 | WP_094870401.1 hypothetical protein |
| 1(1) | WP_307589608.1 | WP_307589608.1 hypothetical protein |
| 1(20) | WP_013370786.1 | WP_013370786.1 WXG100 family type VII secretion target |
| 1(2) | WP_005210482.1 | WP_005210482.1 glycosyltransferase |
| 1(1) | WP_138755919.1 | WP_138755919.1 pore-forming ESAT-6 family protein |
| 1(2) | WP_097898675.1 | WP_097898675.1 tetratricopeptide repeat protein |
| 1(1) | WP_150457767.1 | WP_150457767.1 EsaB/YukD family protein |
| 1(1) | WP_127812908.1 | WP_127812908.1 bacteriocin-processing peptidase family protein |
| 1(1) | WP_243495549.1 | WP_243495549.1 zinc-ribbon domain-containing protein |
| 1(1) | WP_193726775.1 | WP_193726775.1 WXG100 family type VII secretion target |
| 1(1) | WP_144492100.1 | WP_144492100.1 hypothetical protein |
| 1(1) | WP_026882395.1 | WP_026882395.1 WXG100 family type VII secretion target |
| 1(1) | WP_213506886.1 | WP_213506886.1 copper amine oxidase N-terminal domain-containing protein |
| 1(1) | WP_226642878.1 | WP_226642878.1 Mbeg1-like protein |
| 1(1) | WP_160790577.1 | WP_160790577.1 S-layer homology domain-containing protein |
| 1(1) | WP_185165647.1 | WP_185165647.1 DUF6382 domain-containing protein |
| 1(1) | WP_000883810.1 | WP_000883810.1 bacteriocin-processing peptidase family protein |
| 1(1) | WP_277142100.1 | WP_277142100.1 serine/threonine-protein kinase |
| 1(1) | WP_307399106.1 | WP_307399106.1 tetratricopeptide repeat protein |
| 1(1) | WP_136789991.1 | WP_136789991.1 zinc-ribbon domain-containing protein |
| 1(1) | WP_076167862.1 | WP_076167862.1 hypothetical protein |
| 1(1) | WP_036707099.1 | WP_036707099.1 hypothetical protein |
| 1(1) | WP_096774454.1 | WP_096774454.1 hypothetical protein |
| 1(1) | WP_162944259.1 | WP_162944259.1 CHAP domain-containing protein |
| 1(1) | WP_155609749.1 | WP_155609749.1 DUF2974 domain-containing protein |
| 1(1) | WP_229137198.1 | WP_229137198.1 bacteriocin-processing peptidase family protein |

| | | |
|---|---|---|
| 1(1) | WP_307589612.1 | WP_307589612.1 S-layer homology domain-containing protein |
| 1(1) | WP_197248741.1 | WP_197248741.1 hypothetical protein |
| 1(1) | WP_309098616.1 | WP_309098616.1 protein kinase |
| 1(1) | WP_125084781.1 | WP_125084781.1 WXG100 family type VII secretion target |
| 1(1) | WP_044792554.1 | WP_044792554.1 HNH endonuclease |
| 1(4) | WP_099354912.1 | WP_099354912.1 pore-forming ESAT-6 family protein |
| 1(1) | WP_188593162.1 | WP_188593162.1 hypothetical protein |
| 1(6) | WP_024630324.1 | WP_024630324.1 hypothetical protein |
| 1(2) | WP_185475843.1 | WP_185475843.1 type VII secretion protein EsaA |
| 1(1) | WP_102371682.1 | WP_102371682.1 WXG100 family type VII secretion target |
| 1(1) | WP_049869158.1 | WP_049869158.1 hypothetical protein |
| 1(1) | WP_163977055.1 | WP_163977055.1 WXG100 family type VII secretion target |
| 1(1) | WP_151630613.1 | WP_151630613.1 bacteriocin-processing peptidase family protein |
| 1(1) | WP_253440814.1 | WP_253440814.1 hypothetical protein |
| 1(1) | WP_226602915.1 | WP_226602915.1 DUF2974 domain-containing protein |
| 1(1) | WP_074085356.1 | WP_074085356.1 hypothetical protein |
| 1(2) | WP_038584723.1 | WP_038584723.1 hypothetical protein |
| 1(1) | WP_139895513.1 | WP_139895513.1 bacteriocin-processing peptidase family protein |
| 1(2) | WP_097841432.1 | WP_097841432.1 hypothetical protein |
| 1(1) | WP_293202264.1 | WP_293202264.1 hypothetical protein |
| 1(1) | WP_251596674.1 | WP_251596674.1 DUF5050 domain-containing protein |
| 1(1) | WP_277480694.1 | WP_277480694.1 hypothetical protein |
| 1(2) | WP_158318106.1 | WP_158318106.1 bacteriocin-processing peptidase family protein |
| 1(1) | WP_263324553.1 | WP_263324553.1 FHA domain-containing protein |
| 1(1) | WP_076208840.1 | WP_076208840.1 hypothetical protein |
| 1(1) | WP_268821823.1 | WP_268821823.1 S-layer homology domain-containing protein |
| 1(1) | WP_104030443.1 | WP_104030443.1 hypothetical protein |
| 1(1) | WP_223868743.1 | WP_223868743.1 RHS repeat-associated core domain-containing protein |
| 1(1) | WP_188535086.1 | WP_188535086.1 hypothetical protein |
| 1(1) | WP_148450948.1 | WP_148450948.1 EsaB/YukD family protein |
| 1(1) | WP_234121602.1 | WP_234121602.1 hypothetical protein |
| 1(1) | WP_212330903.1 | WP_212330903.1 hypothetical protein |
| 1(1) | WP_260867010.1 | WP_260867010.1 S-layer homology domain-containing protein |
| 1(1) | WP_302153081.1 | WP_302153081.1 hypothetical protein |
| 1(2) | WP_095358548.1 | WP_095358548.1 hypothetical protein |
| 1(1) | WP_229137985.1 | WP_229137985.1 hypothetical protein |
| 1(1) | WP_121474476.1 | WP_121474476.1 WG repeat-containing protein |
| 1(1) | WP_160807261.1 | WP_160807261.1 type VII secretion protein EsaA |
| 1(1) | WP_007147977.1 | WP_007147977.1 YhgE/Pip domain-containing protein |
| 1(5) | WP_062326145.1 | WP_062326145.1 hypothetical protein |
| 1(1) | WP_270716427.1 | WP_270716427.1 tetratricopeptide repeat protein |
| 1(1) | WP_084117734.1 | WP_084117734.1 EsaB/YukD family protein |
| 1(1) | WP_241929676.1 | WP_241929676.1 hypothetical protein |
| 1(1) | WP_268613669.1 | WP_268613669.1 hypothetical protein |
| 1(1) | WP_128101489.1 | WP_128101489.1 hypothetical protein |
| 1(1) | WP_174551273.1 | WP_174551273.1 S-layer homology domain-containing protein, partial |
| 1(1) | WP_088550038.1 | WP_088550038.1 DUF2974 domain-containing protein |
| 1(4) | WP_007128627.1 | WP_007128627.1 WXG100 family type VII secretion target |
| 1(1) | WP_212956547.1 | WP_212956547.1 bacteriocin-processing peptidase family protein |
| 1(1) | WP_240968292.1 | WP_240968292.1 CdiA family toxin C-terminal domain-containing protein |
| 1(1) | WP_098264279.1 | WP_098264279.1 bacteriocin-processing peptidase family protein |
| 1(1) | WP_183578443.1 | WP_183578443.1 hypothetical protein |

| | | |
|---|---|---|
| 1(1) | WP_007148000.1 | WP_007148000.1 FHA domain-containing protein |
| 1(2) | WP_090921061.1 | WP_090921061.1 hypothetical protein |
| 1(1) | WP_306049006.1 | WP_306049006.1 LysM peptidoglycan-binding domain-containing protein |
| 1(1) | WP_053564630.1 | WP_053564630.1 tetratricopeptide repeat protein |
| 1(2) | WP_255289855.1 | WP_255289855.1 deaminase domain-containing protein |
| 1(1) | WP_015595771.1 | WP_015595771.1 glycosyltransferase |
| 1(1) | WP_018664840.1 | WP_018664840.1 EsaB/YukD family protein |
| 1(1) | WP_213592867.1 | WP_213592867.1 DUF5050 domain-containing protein |
| 1(1) | WP_308110105.1 | WP_308110105.1 VWA domain-containing protein |
| 1(1) | WP_208926618.1 | WP_208926618.1 WXG100 family type VII secretion target |
| 1(1) | WP_268613673.1 | WP_268613673.1 DUF2974 domain-containing protein |
| 1(1) | WP_283904339.1 | WP_283904339.1 Ig-like domain-containing protein |
| 1(1) | WP_141559431.1 | WP_141559431.1 hypothetical protein |
| 1(1) | WP_053564626.1 | WP_053564626.1 hypothetical protein |
| 1(1) | WP_199795353.1 | WP_199795353.1 hypothetical protein |
| 1(1) | WP_144476578.1 | WP_144476578.1 EsaB/YukD family protein |
| 1(1) | WP_097905189.1 | WP_097905189.1 bacteriocin-processing peptidase family protein |
| 1(1) | WP_148450935.1 | WP_148450935.1 DL-endopeptidase inhibitor IseA family protein |
| 1(1) | WP_145156808.1 | WP_145156808.1 hypothetical protein |
| 1(1) | WP_123067116.1 | WP_123067116.1 hypothetical protein |
| 1(1) | WP_290407045.1 | WP_290407045.1 DUF5050 domain-containing protein |
| 1(1) | WP_068619117.1 | WP_068619117.1 hypothetical protein |
| 1(1) | WP_077619644.1 | WP_077619644.1 pore-forming ESAT-6 family protein |
| 1(1) | WP_227219194.1 | WP_227219194.1 FHA domain-containing protein |
| 1(2) | WP_254366011.1 | WP_254366011.1 WXG100 family type VII secretion target |
| 1(1) | WP_211022789.1 | WP_211022789.1 DUF5050 domain-containing protein |
| 1(1) | WP_127570437.1 | WP_127570437.1 WXG100 family type VII secretion target |
| 1(1) | WP_224465615.1 | WP_224465615.1 tetratricopeptide repeat protein |
| 1(1) | WP_243495561.1 | WP_243495561.1 WXG100 family type VII secretion target |
| 1(2) | WP_098054117.1 | WP_098054117.1 type VII secretion protein EsaA |
| 1(1) | WP_163255257.1 | WP_163255257.1 hypothetical protein |
| 1(2) | WP_052096890.1 | WP_052096890.1 copper amine oxidase N-terminal domain-containing protein |
| 1(1) | WP_076150487.1 | WP_076150487.1 serine/threonine protein kinase |
| 1(1) | WP_121610601.1 | WP_121610601.1 serine/threonine-protein kinase |
| 1(1) | WP_231094842.1 | WP_231094842.1 serine/threonine-protein kinase |
| 1(1) | WP_272035310.1 | WP_272035310.1 WXG100 family type VII secretion target |
| 1(1) | WP_223068046.1 | WP_223068046.1 WXG100 family type VII secretion target |
| 1(1) | WP_251245814.1 | WP_251245814.1 WXG100 family type VII secretion target |
| 1(1) | WP_212695782.1 | WP_212695782.1 WXG100 family type VII secretion target |
| 1(1) | WP_229139923.1 | WP_229139923.1 pore-forming ESAT-6 family protein |
| 1(1) | WP_054958411.1 | WP_054958411.1 hypothetical protein |
| 1(1) | WP_235286201.1 | WP_235286201.1 pore-forming ESAT-6 family protein |
| 1(1) | WP_071719984.1 | WP_071719984.1 hypothetical protein |
| 1(1) | WP_143316563.1 | WP_143316563.1 WXG100 family type VII secretion target |
| 1(1) | WP_054958410.1 | WP_054958410.1 WXG100 family type VII secretion target |
| 1(1) | WP_137062800.1 | WP_137062800.1 WXG100 family type VII secretion target |
| 1(1) | WP_110899363.1 | WP_110899363.1 WXG100 family type VII secretion target |
| 1(1) | WP_234121603.1 | WP_234121603.1 hypothetical protein |
| 1(1) | WP_038584720.1 | WP_038584720.1 hypothetical protein |
| 1(1) | WP_134750594.1 | WP_134750594.1 hypothetical protein |
| 1(2) | WP_295628322.1 | WP_295628322.1 hypothetical protein |

| | | |
|---|---|---|
| 1(1) | WP_289607500.1 | WP_289607500.1 serine/threonine-protein kinase |
| 1(1) | WP_077567018.1 | WP_077567018.1 WXG100 family type VII secretion target |
| 1(1) | WP_197937399.1 | WP_197937399.1 DUF2974 domain-containing protein |
| 1(1) | WP_099223956.1 | WP_099223956.1 type VII secretion protein EsaA |
| 1(2) | WP_200457883.1 | WP_200457883.1 bacteriocin-processing peptidase family protein |
| 1(1) | WP_164515192.1 | WP_164515192.1 WXG100 family type VII secretion target |
| 1(1) | WP_101300063.1 | WP_101300063.1 copper amine oxidase N-terminal domain-containing protein |
| 1(1) | WP_239617472.1 | WP_239617472.1 DUF2974 domain-containing protein |
| 1(1) | WP_152402382.1 | WP_152402382.1 hypothetical protein |
| 1(1) | WP_068536366.1 | WP_068536366.1 WXG100 family type VII secretion target |
| 1(1) | WP_159439963.1 | WP_159439963.1 Ig-like domain-containing protein |
| 1(1) | WP_099517105.1 | WP_099517105.1 hypothetical protein |
| 1(1) | WP_158651617.1 | WP_158651617.1 LysM peptidoglycan-binding domain-containing protein |
| 1(5) | WP_000054421.1 | WP_000054421.1 SseB family protein |
| 1(1) | WP_080468522.1 | WP_080468522.1 WXG100 family type VII secretion target |
| 1(4) | WP_026593078.1 | WP_026593078.1 WXG100 family type VII secretion target |
| 1(1) | WP_212330897.1 | WP_212330897.1 WXG100 family type VII secretion target |
| 1(1) | WP_078181469.1 | WP_078181469.1 hypothetical protein |
| 1(1) | WP_304216878.1 | WP_304216878.1 protein kinase |
| 1(1) | WP_077320214.1 | WP_077320214.1 hypothetical protein |
| 1(1) | WP_263703625.1 | WP_263703625.1 tetratricopeptide repeat protein |
| 1(1) | WP_134909997.1 | WP_134909997.1 hypothetical protein |
| 1(1) | WP_244719642.1 | WP_244719642.1 WXG100 family type VII secretion target |
| 1(1) | WP_213506896.1 | WP_213506896.1 IseA DL-endopeptidase inhibitor family protein |
| 1(1) | WP_227011714.1 | WP_227011714.1 hypothetical protein |
| 1(2) | WP_094876485.1 | WP_094876485.1 DUF2974 domain-containing protein |
| 1(1) | WP_098836668.1 | WP_098836668.1 WXG100 family type VII secretion target, partial |
| 1(1) | WP_128631289.1 | WP_128631289.1 WXG100 family type VII secretion target |
| 1(1) | WP_076292682.1 | WP_076292682.1 hypothetical protein |
| 1(1) | WP_070638645.1 | WP_070638645.1 serine/threonine-protein kinase |
| 1(1) | WP_042210996.1 | WP_042210996.1 hypothetical protein |
| 1(1) | WP_136379692.1 | WP_136379692.1 hypothetical protein |
| 1(1) | WP_305163858.1 | WP_305163858.1 EsaB/YukD family protein |
| 1(1) | WP_289373793.1 | WP_289373793.1 hypothetical protein |
| 1(1) | WP_094092621.1 | WP_094092621.1 hypothetical protein |
| 1(1) | WP_099517092.1 | WP_099517092.1 WXG100 family type VII secretion target |
| 1(2) | WP_144025951.1 | WP_144025951.1 hypothetical protein |
| 1(1) | WP_061975588.1 | WP_061975588.1 tetratricopeptide repeat protein |
| 1(1) | WP_021261885.1 | WP_021261885.1 hypothetical protein |
| 1(1) | WP_188944152.1 | WP_188944152.1 type VII secretion protein EsaA |
| 1(1) | WP_142318160.1 | WP_142318160.1 hypothetical protein |
| 1(1) | WP_284036497.1 | WP_284036497.1 hypothetical protein |
| 1(1) | WP_136372966.1 | WP_136372966.1 hypothetical protein |
| 1(2) | WP_097898674.1 | WP_097898674.1 WXG100 family type VII secretion target |
| 1(2) | WP_038584744.1 | WP_038584744.1 pore-forming ESAT-6 family protein |
| 1(1) | WP_051590912.1 | WP_051590912.1 hypothetical protein |
| 1(1) | WP_279323593.1 | WP_279323593.1 tetratricopeptide repeat protein |
| 1(1) | WP_242400027.1 | WP_242400027.1 SseB family protein |
| 1(1) | WP_238655352.1 | WP_238655352.1 hypothetical protein |
| 1(1) | WP_146003684.1 | WP_146003684.1 serine/threonine-protein kinase |
| 1(2) | WP_240913010.1 | WP_240913010.1 S-layer homology domain-containing protein |
| 1(4) | WP_185337279.1 | WP_185337279.1 hypothetical protein |
| 1(2) | WP_087982586.1 | WP_087982586.1 tetratricopeptide repeat protein |
| 1(12) | WP_003336004.1 | WP_003336004.1 WXG100 family type VII secretion target |
| 1(1) | WP_202283184.1 | WP_202283184.1 hypothetical protein |
| 1(1) | WP_192266458.1 | WP_192266458.1 hypothetical protein |
| 1(1) | WP_077852421.1 | WP_077852421.1 Ig-like domain-containing protein |
| 1(1) | WP_213426989.1 | WP_213426989.1 WXG100 family type VII secretion target |
| 1(1) | WP_241929670.1 | WP_241929670.1 serine hydrolase domain-containing protein |
| 1(1) | WP_007083292.1 | WP_007083292.1 WXG100 family type VII secretion target |
| 1(3) | WP_036702412.1 | WP_036702412.1 WXG100 family type VII secretion target |
| 1(1) | WP_026882400.1 | WP_026882400.1 DUF6382 domain-containing protein |
| 1(1) | WP_212330888.1 | WP_212330888.1 FHA domain-containing protein |
| 1(1) | WP_128631291.1 | WP_128631291.1 YhgE/Pip domain-containing protein |
| 1(1) | WP_221791096.1 | WP_221791096.1 WXG100 family type VII secretion target |
| 1(1) | WP_038584767.1 | WP_038584767.1 glycoside hydrolase family 3 C-terminal domain-containing protein |
| 1(1) | WP_290401997.1 | WP_290401997.1 hypothetical protein |
| 1(4) | WP_141549175.1 | WP_141549175.1 WXG100 family type VII secretion target |
| 1(1) | WP_180233476.1 | WP_180233476.1 WXG100 family type VII secretion target |
| 1(1) | WP_098723009.1 | WP_098723009.1 type VII secretion protein EsaA |
| 1(1) | WP_120462941.1 | WP_120462941.1 hypothetical protein |
| 1(1) | WP_024632688.1 | WP_024632688.1 WXG100 family type VII secretion target |
| 1(1) | WP_139355768.1 | WP_139355768.1 glycoside hydrolase family 3 C-terminal domain-containing protein |
| 1(1) | WP_197937393.1 | WP_197937393.1 DUF5050 domain-containing protein |
| 1(1) | WP_159146507.1 | WP_159146507.1 WXG100 family type VII secretion target |
| 1(1) | WP_213506881.1 | WP_213506881.1 copper amine oxidase N-terminal domain-containing protein |
| 1(1) | WP_147807653.1 | WP_147807653.1 SseB family protein |
| 1(1) | WP_157942994.1 | WP_157942994.1 copper amine oxidase N-terminal domain-containing protein |
| 1(1) | WP_046522787.1 | WP_046522787.1 WXG100 family type VII secretion target |
| 1(1) | WP_173601442.1 | WP_173601442.1 bacteriocin-processing peptidase family protein |
| 1(1) | WP_116556145.1 | WP_116556145.1 WXG100 family type VII secretion target |
| 1(1) | WP_055105793.1 | WP_055105793.1 WXG100 family type VII secretion target |
| 1(1) | WP_254349548.1 | WP_254349548.1 copper amine oxidase N-terminal domain-containing protein |
| 1(1) | WP_048826569.1 | WP_048826569.1 WXG100 family type VII secretion target |
| 1(1) | WP_090896192.1 | WP_090896192.1 hypothetical protein |
| 1(1) | WP_268613677.1 | WP_268613677.1 hypothetical protein |
| 1(1) | WP_017692356.1 | WP_017692356.1 RHS repeat-associated core domain-containing protein |
| 1(1) | WP_288218586.1 | WP_288218586.1 hypothetical protein |
| 1(2) | WP_295628312.1 | WP_295628312.1 WXG100 family type VII secretion target |
| 1(1) | WP_220784812.1 | WP_220784812.1 hypothetical protein |
| 1(2) | WP_227601478.1 | WP_227601478.1 DNA/RNA non-specific endonuclease |
| 1(1) | WP_090761817.1 | WP_090761817.1 WXG100 family type VII secretion target |
| 1(2) | WP_185413347.1 | WP_185413347.1 type VII secretion protein EsaA |
| 1(1) | WP_270784523.1 | WP_270784523.1 hypothetical protein |
| 1(2) | WP_242951811.1 | WP_242951811.1 WXG100 family type VII secretion target |
| 1(1) | WP_204445202.1 | WP_204445202.1 WXG100 family type VII secretion target |
| 1(2) | WP_010963371.1 | WP_010963371.1 hypothetical protein |
| 1(1) | WP_302153083.1 | WP_302153083.1 hypothetical protein, partial |

| | | |
|---|---|---|
| 1(2) | WP_295628315.1 | WP_295628315.1 pore-forming ESAT-6 family protein |
| 1(1) | WP_271839906.1 | WP_271839906.1 DUF6382 domain-containing protein |
| 1(1) | WP_262787110.1 | WP_262787110.1 bacteriocin-processing peptidase family protein |
| 1(2) | WP_212805124.1 | WP_212805124.1 WXG100 family type VII secretion target |
| 1(1) | WP_277398343.1 | WP_277398343.1 serine/threonine-protein kinase |
| 1(2) | WP_149051220.1 | WP_149051220.1 type VII secretion protein EsaA |
| 1(1) | WP_167332615.1 | WP_167332615.1 hypothetical protein |
| 1(2) | WP_270995170.1 | WP_270995170.1 type VII secretion protein EsaA |
| 1(1) | WP_238187892.1 | WP_238187892.1 hypothetical protein |
| 1(1) | WP_142325887.1 | WP_142325887.1 WXG100 family type VII secretion target, partial |
| 1(1) | WP_289342266.1 | WP_289342266.1 WXG100 family type VII secretion target |
| 1(1) | WP_134750624.1 | WP_134750624.1 pore-forming ESAT-6 family protein |
| 1(1) | WP_071719979.1 | WP_071719979.1 SseB family protein |
| 1(1) | WP_270716430.1 | WP_270716430.1 hypothetical protein |
| 1(1) | WP_012260843.1 | WP_012260843.1 hypothetical protein |
| 1(1) | WP_290401995.1 | WP_290401995.1 WXG100 family type VII secretion target |
| 1(2) | WP_185469296.1 | WP_185469296.1 type VII secretion protein EsaA |
| 1(1) | WP_286124781.1 | WP_286124781.1 EsaB/YukD family protein |
| 1(1) | WP_068696869.1 | WP_068696869.1 WXG100 family type VII secretion target |
| 1(1) | WP_258208936.1 | WP_258208936.1 InlB B-repeat-containing protein |
| 1(1) | WP_144564314.1 | WP_144564314.1 hypothetical protein, partial |
| 1(3) | WP_240522517.1 | WP_240522517.1 HNH endonuclease |
| 1(26) | WP_000391135.1 | WP_000391135.1 hypothetical protein |
| 1(1) | WP_041964567.1 | WP_041964567.1 Ig-like domain-containing protein |
| 1(1) | WP_144488221.1 | WP_144488221.1 hypothetical protein |
| 1(3) | WP_071708777.1 | WP_071708777.1 SseB family protein |
| 1(1) | WP_088032763.1 | WP_088032763.1 bacteriocin-processing peptidase family protein |
| 1(6) | WP_291288265.1 | WP_291288265.1 hypothetical protein |
| 1(1) | WP_250244815.1 | WP_250244815.1 hypothetical protein |
| 1(1) | WP_270716533.1 | WP_270716533.1 WXG100 family type VII secretion target |
| 1(1) | WP_138755917.1 | WP_138755917.1 DUF456 domain-containing protein |
| 1(8) | WP_185364862.1 | WP_185364862.1 type VII secretion protein EsaA |
| 1(1) | WP_192510491.1 | WP_192510491.1 hypothetical protein |
| 1(1) | WP_125084793.1 | WP_125084793.1 hypothetical protein |
| 1(1) | WP_113755279.1 | WP_113755279.1 hypothetical protein |
| 1(1) | WP_013252380.1 | WP_013252380.1 DUF6531 domain-containing protein |
| 1(2) | WP_143801061.1 | WP_143801061.1 WXG100 family type VII secretion target |
| 1(1) | WP_188535076.1 | WP_188535076.1 WXG100 family type VII secretion target |
| 1(1) | WP_280925130.1 | WP_280925130.1 FHA domain-containing protein |
| 1(2) | WP_057897622.1 | WP_057897622.1 hypothetical protein |
| 1(1) | WP_155703921.1 | WP_155703921.1 stalk domain-containing protein |
| 1(1) | WP_025848605.1 | WP_025848605.1 DUF2974 domain-containing protein |
| 1(2) | WP_214169378.1 | WP_214169378.1 T7SS effector LXG polymorphic toxin |
| 1(1) | WP_211745833.1 | WP_211745833.1 hypothetical protein |
| 1(1) | WP_231417868.1 | WP_231417868.1 hypothetical protein |
| 1(1) | WP_225999410.1 | WP_225999410.1 hypothetical protein |
| 1(1) | WP_036944545.1 | WP_036944545.1 copper amine oxidase N-terminal domain-containing protein |
| 1(1) | WP_000392221.1 | WP_000392221.1 hypothetical protein |
| 1(4) | WP_003728973.1 | WP_003728973.1 EsaB/YukD family protein |
| 1(1) | WP_076297594.1 | WP_076297594.1 AAA family ATPase |
| 1(1) | WP_251466812.1 | WP_251466812.1 Ig-like domain-containing protein |
| 1(1) | WP_036944556.1 | WP_036944556.1 WXG100 family type VII secretion target |
| 1(1) | WP_258382319.1 | WP_258382319.1 copper amine oxidase N-terminal domain-containing protein |
| 1(2) | WP_013252387.1 | WP_013252387.1 serine/threonine-protein kinase |
| 1(1) | WP_257443652.1 | WP_257443652.1 EsaB/YukD family protein |
| 1(1) | WP_090510247.1 | WP_090510247.1 hypothetical protein |
| 1(1) | WP_111273182.1 | WP_111273182.1 hypothetical protein |
| 1(1) | WP_068694725.1 | WP_068694725.1 YhgE/Pip domain-containing protein |
| 1(1) | WP_039306353.1 | WP_039306353.1 hypothetical protein |
| 1(1) | WP_231417860.1 | WP_231417860.1 FHA domain-containing protein |
| 1(1) | WP_138755908.1 | WP_138755908.1 hypothetical protein |
| 1(1) | WP_208926594.1 | WP_208926594.1 hypothetical protein |
| 1(1) | WP_281103271.1 | WP_281103271.1 tetratricopeptide repeat protein |
| 1(1) | WP_215572754.1 | WP_215572754.1 hypothetical protein |
| 1(1) | WP_214706000.1 | WP_214706000.1 hypothetical protein |
| 1(1) | WP_111153300.1 | WP_111153300.1 hypothetical protein |
| 1(1) | WP_251417637.1 | WP_251417637.1 hypothetical protein |
| 1(1) | WP_085499075.1 | WP_085499075.1 hypothetical protein |
| 1(1) | WP_144655919.1 | WP_144655919.1 bacteriocin-processing peptidase family protein |
| 1(1) | WP_068619111.1 | WP_068619111.1 hypothetical protein |
| 1(1) | WP_279323735.1 | WP_279323735.1 tetratricopeptide repeat protein |
| 1(1) | WP_062239855.1 | WP_062239855.1 WXG100 family type VII secretion target |
| 1(1) | WP_102956433.1 | WP_102956433.1 T7SS effector LXG polymorphic toxin |
| 1(2) | WP_174820806.1 | WP_174820806.1 S-layer homology domain-containing protein |
| 1(1) | WP_025118713.1 | WP_025118713.1 WXG100 family type VII secretion target |
| 1(1) | WP_088313235.1 | WP_088313235.1 tetratricopeptide repeat protein |
| 1(1) | WP_273191245.1 | WP_273191245.1 serine/threonine-protein kinase |
| 1(1) | WP_263324549.1 | WP_263324549.1 DUF2974 domain-containing protein |
| 1(1) | WP_111153301.1 | WP_111153301.1 WXG100 family type VII secretion target |
| 1(2) | WP_076210969.1 | WP_076210969.1 WXG100 family type VII secretion target |
| 1(1) | WP_036575533.1 | WP_036575533.1 serine/threonine-protein kinase |
| 1(9) | WP_001007574.1 | WP_001007574.1 EsaB/YukD family protein |
| 1(2) | WP_023521814.1 | WP_023521814.1 SseB family protein |
| 1(1) | WP_231094840.1 | WP_231094840.1 Ig-like domain-containing protein |
| 1(1) | WP_098907646.1 | WP_098907646.1 tetratricopeptide repeat protein |
| 1(1) | WP_098508769.1 | WP_098508769.1 bacteriocin-processing peptidase family protein |
| 1(3) | WP_098248508.1 | WP_098248508.1 WXG100 family type VII secretion target |
| 1(1) | WP_021253125.1 | WP_021253125.1 copper amine oxidase N-terminal domain-containing protein |
| 1(1) | WP_089965942.1 | WP_089965942.1 hypothetical protein |
| 1(2) | WP_219872456.1 | WP_219872456.1 hypothetical protein |
| 1(1) | WP_088063227.1 | WP_088063227.1 bacteriocin-processing peptidase family protein |
| 1(1) | WP_003336005.1 | WP_003336005.1 WG repeat-containing protein |
| 1(1) | WP_212330879.1 | WP_212330879.1 hypothetical protein |
| 1(1) | WP_030025437.1 | WP_030025437.1 T7SS effector LXG polymorphic toxin |
| 1(2) | WP_239660300.1 | WP_239660300.1 DNA/RNA non-specific endonuclease |
| 1(1) | WP_094593876.1 | WP_094593876.1 WXG100 family type VII secretion target |
| 1(1) | WP_110931531.1 | WP_110931531.1 copper amine oxidase N-terminal domain-containing protein |
| 1(1) | WP_308082246.1 | WP_308082246.1 hypothetical protein |
| 1(1) | WP_019912159.1 | WP_019912159.1 HNH endonuclease |
| 1(1) | WP_127570436.1 | WP_127570436.1 pore-forming ESAT-6 family protein |

| | | |
|---|---|---|
| 1(1) | WP_029141777.1 | WP_029141777.1 EsaB/YukD family protein |
| 1(1) | WP_229696415.1 | WP_229696415.1 hypothetical protein |
| 1(1) | WP_291672203.1 | WP_291672203.1 protein kinase |
| 1(1) | WP_246565442.1 | WP_246565442.1 hypothetical protein |
| 1(1) | WP_136790007.1 | WP_136790007.1 glycosyltransferase family 2 protein |
| 1(1) | WP_134750588.1 | WP_134750588.1 glycosyltransferase |
| 1(1) | WP_257497519.1 | WP_257497519.1 serine/threonine-protein kinase |
| 1(1) | WP_171414513.1 | WP_171414513.1 hypothetical protein |
| 1(1) | WP_186325901.1 | WP_186325901.1 WXG100 family type VII secretion target |
| 1(1) | WP_307192896.1 | WP_307192896.1 WXG100 family type VII secretion target |
| 1(1) | WP_138186702.1 | WP_138186702.1 YHYH domain-containing protein |
| 1(1) | WP_002148315.1 | WP_002148315.1 WXG100 family type VII secretion target |
| 1(1) | WP_029438098.1 | WP_029438098.1 CdiA family toxin C-terminal domain-containing protein |
| 1(1) | WP_232274495.1 | WP_232274495.1 hypothetical protein |
| 1(4) | WP_023521807.1 | WP_023521807.1 bacteriocin-processing peptidase family protein |
| 1(1) | WP_090239229.1 | WP_090239229.1 hypothetical protein |
| 1(1) | WP_308082249.1 | WP_308082249.1 WXG100 family type VII secretion target |
| 1(1) | WP_143316565.1 | WP_143316565.1 hypothetical protein |
| 1(2) | WP_076281070.1 | WP_076281070.1 copper amine oxidase N-terminal domain-containing protein |
| 1(1) | WP_213426988.1 | WP_213426988.1 hypothetical protein |
| 1(2) | WP_195431322.1 | WP_195431322.1 EsaB/YukD family protein |
| 1(1) | WP_122971539.1 | WP_122971539.1 WXG100 family type VII secretion target |
| 1(1) | WP_233901879.1 | WP_233901879.1 copper amine oxidase N-terminal domain-containing protein |
| 1(1) | WP_166244896.1 | WP_166244896.1 hypothetical protein |
| 1(1) | WP_121610598.1 | WP_121610598.1 pore-forming ESAT-6 family protein |
| 1(4) | WP_185350818.1 | WP_185350818.1 hypothetical protein |
| 1(1) | WP_010250268.1 | WP_010250268.1 WXG100 family type VII secretion target |
| 1(1) | WP_257149140.1 | WP_257149140.1 HNH endonuclease |
| 1(1) | WP_098776438.1 | WP_098776438.1 bacteriocin-processing peptidase family protein |
| 1(1) | WP_163872769.1 | WP_163872769.1 hypothetical protein |
| 1(1) | WP_148500005.1 | WP_148500005.1 hypothetical protein |
| 1(1) | WP_071719981.1 | WP_071719981.1 tetratricopeptide repeat protein |
| 1(1) | WP_241929677.1 | WP_241929677.1 hypothetical protein |
| 1(1) | WP_148409439.1 | WP_148409439.1 DUF5050 domain-containing protein |
| 1(1) | WP_076311048.1 | WP_076311048.1 hypothetical protein |
| 1(1) | WP_029438093.1 | WP_029438093.1 WXG100 family type VII secretion target |
| 1(1) | WP_024632679.1 | WP_024632679.1 copper amine oxidase N-terminal domain-containing protein |
| 1(1) | WP_061113196.1 | WP_061113196.1 T7SS effector LXG polymorphic toxin |
| 1(1) | WP_080545873.1 | WP_080545873.1 type VII secretion protein EsaA, partial |
| 1(1) | WP_286017551.1 | WP_286017551.1 WXG100 family type VII secretion target |
| 1(1) | WP_197937402.1 | WP_197937402.1 stalk domain-containing protein |
| 1(1) | WP_098148057.1 | WP_098148057.1 EsaB/YukD family protein |
| 1(5) | WP_061829458.1 | WP_061829458.1 WXG100 family type VII secretion target |
| 1(1) | WP_307284680.1 | WP_307284680.1 WXG100 family type VII secretion target |
| 1(1) | WP_097841427.1 | WP_097841427.1 bacteriocin-processing peptidase family protein |
| 1(1) | WP_270364632.1 | WP_270364632.1 SseB family protein |
| 1(1) | WP_165443460.1 | WP_165443460.1 serine/threonine-protein kinase |
| 1(1) | WP_291672191.1 | WP_291672191.1 hypothetical protein, partial |
| 1(1) | WP_088058537.1 | WP_088058537.1 tetratricopeptide repeat protein |

| | | |
|---|---|---|
| 1(1) | WP_025991672.1 | WP_025991672.1 DNA/RNA non-specific endonuclease |
| 1(1) | WP_070638655.1 | WP_070638655.1 hypothetical protein |
| 1(1) | WP_099361339.1 | WP_099361339.1 DUF6382 domain-containing protein |
| 1(2) | WP_061667209.1 | WP_061667209.1 EsaB/YukD family protein |
| 1(1) | WP_251466805.1 | WP_251466805.1 DUF6382 domain-containing protein |
| 1(3) | WP_175623546.1 | WP_175623546.1 EsaB/YukD family protein |
| 1(1) | WP_044037257.1 | WP_044037257.1 WXG100 family type VII secretion target |
| 1(1) | WP_197937396.1 | WP_197937396.1 WXG100 family type VII secretion target |
| 1(1) | WP_071220488.1 | WP_071220488.1 hypothetical protein |
| 1(1) | WP_010676868.1 | WP_010676868.1 hypothetical protein |
| 1(1) | WP_056700783.1 | WP_056700783.1 hypothetical protein |
| 1(1) | WP_098284017.1 | WP_098284017.1 T7SS effector LXG polymorphic toxin |
| 1(3) | WP_071220206.1 | WP_071220206.1 hypothetical protein |
| 1(1) | WP_071558842.1 | WP_071558842.1 hypothetical protein |
| 1(1) | WP_229137986.1 | WP_229137986.1 WXG100 family type VII secretion target |
| 1(1) | WP_148362973.1 | WP_148362973.1 hypothetical protein |
| 1(1) | WP_220027682.1 | WP_220027682.1 WXG100 family type VII secretion target |
| 1(1) | WP_260410443.1 | WP_260410443.1 RHS repeat-associated core domain-containing protein |
| 1(1) | WP_270682701.1 | WP_270682701.1 tetratricopeptide repeat protein |
| 1(1) | WP_102264546.1 | WP_102264546.1 FHA domain-containing protein |
| 1(1) | WP_036702411.1 | WP_036702411.1 peptidoglycan-binding protein |
| 1(1) | WP_268871029.1 | WP_268871029.1 LysM domain-containing protein |
| 1(1) | WP_223617613.1 | WP_223617613.1 bacteriocin-processing peptidase family protein |
| 1(1) | WP_268613686.1 | WP_268613686.1 hypothetical protein |
| 1(1) | WP_274485610.1 | WP_274485610.1 copper amine oxidase N-terminal domain-containing protein |
| 1(1) | WP_036575566.1 | WP_036575566.1 hypothetical protein |
| 1(1) | WP_035318347.1 | WP_035318347.1 hypothetical protein |
| 1(1) | WP_152381385.1 | WP_152381385.1 S-layer homology domain-containing protein |
| 1(2) | WP_148450944.1 | WP_148450944.1 WXG100 family type VII secretion target |
| 1(1) | WP_223068050.1 | WP_223068050.1 DUF2974 domain-containing protein |
| 1(1) | WP_258207829.1 | WP_258207829.1 WXG100 family type VII secretion target |
| 1(1) | WP_113052568.1 | WP_113052568.1 hypothetical protein |
| 1(1) | WP_182301846.1 | WP_182301846.1 S-layer homology domain-containing protein |
| 1(1) | WP_236334353.1 | WP_236334353.1 hypothetical protein |
| 1(2) | WP_112331590.1 | WP_112331590.1 hypothetical protein |
| 1(1) | WP_054958409.1 | WP_054958409.1 hypothetical protein |
| 1(1) | WP_023957725.1 | WP_023957725.1 WXG100 family type VII secretion target |
| 1(1) | WP_229106295.1 | WP_229106295.1 DUF2974 domain-containing protein |
| 1(1) | WP_248251188.1 | WP_248251188.1 hypothetical protein |
| 1(1) | WP_053492737.1 | WP_053492737.1 hypothetical protein |
| 1(1) | WP_061402733.1 | WP_061402733.1 hypothetical protein |
| 1(1) | WP_208752240.1 | WP_208752240.1 SseB family protein |
| 1(1) | WP_257675216.1 | WP_257675216.1 tetratricopeptide repeat protein |
| 1(1) | WP_151630601.1 | WP_151630601.1 SMI1/KNR4 family protein |
| 1(1) | WP_215185474.1 | WP_215185474.1 DUF5050 domain-containing protein |
| 1(1) | WP_280787394.1 | WP_280787394.1 WXG100 family type VII secretion target |
| 1(1) | WP_233697177.1 | WP_233697177.1 serine/threonine-protein kinase |
| 1(1) | WP_098204197.1 | WP_098204197.1 bacteriocin-processing peptidase family protein |
| 1(1) | WP_098538860.1 | WP_098538860.1 DNA/RNA non-specific endonuclease |
| 1(1) | WP_257142775.1 | WP_257142775.1 CdiA family toxin C-terminal domain-containing protein |
| 1(1) | WP_291288271.1 | WP_291288271.1 FHA domain-containing protein |

| | | |
|---|---|---|
| 1(2) | WP_277746359.1 | WP_277746359.1 hypothetical protein |
| 1(1) | WP_251610749.1 | WP_251610749.1 tetratricopeptide repeat protein |
| 1(1) | WP_262748106.1 | WP_262748106.1 WXG100 family type VII secretion target |
| 1(1) | WP_204412717.1 | WP_204412717.1 FHA domain-containing protein |
| 1(1) | WP_076301096.1 | WP_076301096.1 hypothetical protein |
| 1(1) | WP_145412704.1 | WP_145412704.1 hypothetical protein |
| 1(2) | WP_230387341.1 | WP_230387341.1 CdiA family toxin C-terminal domain-containing protein |
| 1(1) | WP_242230446.1 | WP_242230446.1 WXG100 family type VII secretion target |
| 1(1) | WP_078993369.1 | WP_078993369.1 bacteriocin-processing peptidase family protein |
| 1(1) | WP_208926610.1 | WP_208926610.1 hypothetical protein |
| 1(1) | WP_010250291.1 | WP_010250291.1 hypothetical protein |
| 1(1) | WP_179034787.1 | WP_179034787.1 DUF5050 domain-containing protein |
| 1(1) | WP_254759296.1 | WP_254759296.1 WXG100 family type VII secretion target |
| 1(2) | WP_098148058.1 | WP_098148058.1 tetratricopeptide repeat protein |
| 1(1) | WP_028589294.1 | WP_028589294.1 WXG100 family type VII secretion target |
| 1(1) | WP_144596436.1 | WP_144596436.1 type VII secretion protein EsaA |
| 1(1) | WP_213650776.1 | WP_213650776.1 hypothetical protein |
| 1(1) | WP_097810735.1 | WP_097810735.1 hypothetical protein |
| 1(1) | WP_207302589.1 | WP_207302589.1 FHA domain-containing protein |
| 1(1) | WP_066022446.1 | WP_066022446.1 FHA domain-containing protein |
| 1(1) | WP_048826560.1 | WP_048826560.1 hypothetical protein |
| 1(1) | WP_248063417.1 | WP_248063417.1 hypothetical protein |
| 1(1) | WP_258126010.1 | WP_258126010.1 tetratricopeptide repeat protein |
| 1(1) | WP_089522635.1 | WP_089522635.1 hypothetical protein |
| 1(1) | WP_215601065.1 | WP_215601065.1 tetratricopeptide repeat protein |
| 1(1) | WP_224465614.1 | WP_224465614.1 hypothetical protein |
| 1(1) | WP_257443661.1 | WP_257443661.1 AAA family ATPase |
| 1(1) | WP_255289738.1 | WP_255289738.1 toxin |
| 1(1) | WP_044037275.1 | WP_044037275.1 EsaB/YukD family protein |
| 1(1) | WP_142286518.1 | WP_142286518.1 hypothetical protein |
| 1(1) | WP_297035105.1 | WP_297035105.1 serine/threonine-protein kinase |
| 1(3) | WP_006289744.1 | WP_006289744.1 WXG100 family type VII secretion target |
| 1(1) | WP_014281169.1 | WP_014281169.1 S-layer homology domain-containing protein |
| 1(2) | WP_098746865.1 | WP_098746865.1 hypothetical protein |
| 1(1) | WP_088032801.1 | WP_088032801.1 AHH domain-containing protein |
| 1(1) | WP_209941815.1 | WP_209941815.1 hypothetical protein |
| 1(1) | WP_116556140.1 | WP_116556140.1 tetratricopeptide repeat protein |
| 1(1) | WP_276917312.1 | WP_276917312.1 tetratricopeptide repeat protein |
| 1(1) | WP_273191261.1 | WP_273191261.1 FHA domain-containing protein |
| 1(1) | WP_147786242.1 | WP_147786242.1 bacteriocin-processing peptidase family protein |
| 1(1) | WP_255289857.1 | WP_255289857.1 WXG100 family type VII secretion target |
| 1(1) | WP_294386435.1 | WP_294386435.1 hypothetical protein |
| 1(1) | WP_045518279.1 | WP_045518279.1 FHA domain-containing protein |
| 1(1) | WP_140421210.1 | WP_140421210.1 WXG100 family type VII secretion target |
| 1(1) | WP_076113453.1 | WP_076113453.1 hypothetical protein |
| 1(1) | WP_307394628.1 | WP_307394628.1 WXG100 family type VII secretion target |
| 1(1) | WP_307601046.1 | WP_307601046.1 hypothetical protein |
| 1(1) | WP_013914195.1 | WP_013914195.1 hypothetical protein |
| 1(3) | WP_025685030.1 | WP_025685030.1 hypothetical protein |
| 1(1) | WP_115359658.1 | WP_115359658.1 type VII secretion protein EsaA |
| 1(1) | WP_290401996.1 | WP_290401996.1 tetratricopeptide repeat protein |
| 1(1) | WP_280753902.1 | WP_280753902.1 hypothetical protein |
| 1(1) | WP_000883808.1 | WP_000883808.1 bacteriocin-processing peptidase family protein |
| 1(1) | WP_062239858.1 | WP_062239858.1 hypothetical protein |
| 1(1) | WP_264456857.1 | WP_264456857.1 hypothetical protein |
| 1(3) | WP_110892970.1 | WP_110892970.1 WXG100 family type VII secretion target |
| 1(1) | WP_244525358.1 | WP_244525358.1 hypothetical protein |
| 1(1) | WP_251548755.1 | WP_251548755.1 WXG100 family type VII secretion target |
| 1(1) | WP_145134815.1 | WP_145134815.1 hypothetical protein |
| 1(1) | WP_097984055.1 | WP_097984055.1 hypothetical protein |
| 1(1) | WP_084156402.1 | WP_084156402.1 serine/threonine-protein kinase |
| 1(1) | WP_307738722.1 | WP_307738722.1 WXG100 family type VII secretion target |
| 1(1) | WP_076103995.1 | WP_076103995.1 DUF2974 domain-containing protein |
| 1(1) | WP_256227721.1 | WP_256227721.1 hypothetical protein |
| 1(1) | WP_026572742.1 | WP_026572742.1 DUF6382 domain-containing protein |
| 1(2) | WP_001007572.1 | WP_001007572.1 EsaB/YukD family protein |
| 1(1) | WP_283904341.1 | WP_283904341.1 protein kinase |
| 1(1) | WP_263880220.1 | WP_263880220.1 S-layer homology domain-containing protein |
| 1(1) | WP_021261882.1 | WP_021261882.1 WXG100 family type VII secretion target |
| 1(1) | WP_149190451.1 | WP_149190451.1 hypothetical protein |
| 1(1) | WP_307399111.1 | WP_307399111.1 WXG100 family type VII secretion target |
| 1(1) | WP_211082928.1 | WP_211082928.1 WXG100 family type VII secretion target |
| 1(1) | WP_060630649.1 | WP_060630649.1 hypothetical protein |
| 1(1) | WP_145412707.1 | WP_145412707.1 hypothetical protein |
| 1(1) | WP_023521817.1 | WP_023521817.1 CdiA family toxin C-terminal domain-containing protein |
| 1(1) | WP_031463347.1 | WP_031463347.1 membrane protein |
| 1(1) | WP_056700794.1 | WP_056700794.1 hypothetical protein |
| 1(1) | WP_071640580.1 | WP_071640580.1 hypothetical protein |
| 1(1) | WP_042203644.1 | WP_042203644.1 hypothetical protein |
| 1(1) | WP_251610747.1 | WP_251610747.1 hypothetical protein |
| 1(4) | WP_098266653.1 | WP_098266653.1 SseB family protein |
| 1(1) | WP_017149768.1 | WP_017149768.1 hypothetical protein |
| 1(1) | WP_138226037.1 | WP_138226037.1 S-layer homology domain-containing protein |
| 1(2) | WP_281813089.1 | WP_281813089.1 TNT domain-containing protein |
| 1(1) | WP_076119219.1 | WP_076119219.1 DUF2974 domain-containing protein |
| 1(1) | WP_289607494.1 | WP_289607494.1 WXG100 family type VII secretion target |
| 1(1) | WP_136379703.1 | WP_136379703.1 DUF6382 domain-containing protein |
| 1(1) | WP_235715985.1 | WP_235715985.1 LysM domain-containing protein |
| 1(1) | WP_090921056.1 | WP_090921056.1 WXG100 family type VII secretion target |
| 1(1) | WP_076297563.1 | WP_076297563.1 hypothetical protein |
| 1(1) | WP_012260839.1 | WP_012260839.1 WXG100 family type VII secretion target |
| 1(6) | WP_185463134.1 | WP_185463134.1 HNH endonuclease |
| 1(1) | WP_240522515.1 | WP_240522515.1 WXG100 family type VII secretion target |
| 1(1) | WP_270993641.1 | WP_270993641.1 hypothetical protein |
| 1(1) | WP_070805904.1 | WP_070805904.1 tetratricopeptide repeat protein |
| 1(1) | WP_272801673.1 | WP_272801673.1 zinc-ribbon domain-containing protein |
| 1(1) | WP_236533773.1 | WP_236533773.1 WXG100 family type VII secretion target |
| 1(1) | WP_258207830.1 | WP_258207830.1 pore-forming ESAT-6 family protein |
| 1(1) | WP_036702402.1 | WP_036702402.1 hypothetical protein |
| 1(1) | WP_251557372.1 | WP_251557372.1 Ig-like domain-containing protein |
| 1(1) | WP_114298762.1 | WP_114298762.1 hypothetical protein |
| 1(1) | WP_244719630.1 | WP_244719630.1 WXG100 family type VII secretion target |
| 1(1) | WP_098400683.1 | WP_098400683.1 toxin |
| 1(3) | WP_000506924.1 | WP_000506924.1 tetratricopeptide repeat protein |

1(1) WP_251674649.1 WP_251674649.1 WXG100 family type VII secretion target
1(2) WP_283655695.1 WP_283655695.1 hypothetical protein
1(1) WP_144469659.1 WP_144469659.1 WXG100 family type VII secretion target, partial
1(1) WP_142612994.1 WP_142612994.1 S-layer homology domain-containing protein
1(1) WP_242261454.1 WP_242261454.1 hypothetical protein
1(1) WP_185165643.1 WP_185165643.1 WXG100 family type VII secretion target
1(1) WP_195325259.1 WP_195325259.1 WXG100 family type VII secretion target
1(1) WP_022128272.1 WP_022128272.1 WXG100 family type VII secretion target
1(1) WP_272801676.1 WP_272801676.1 zinc-ribbon domain-containing protein
1(1) WP_254348217.1 WP_254348217.1 copper amine oxidase N-terminal domain-containing protein
1(1) WP_121610603.1 WP_121610603.1 Ig-like domain-containing protein
1(1) WP_268613680.1 WP_268613680.1 WXG100 family type VII secretion target
1(1) WP_114298764.1 WP_114298764.1 WXG100 family type VII secretion target
1(2) WP_194230407.1 WP_194230407.1 hypothetical protein
1(1) WP_089522648.1 WP_089522648.1 S-layer homology domain-containing protein
1(1) WP_084380659.1 WP_084380659.1 S8 family peptidase
1(1) WP_152826366.1 WP_152826366.1 serine/threonine protein kinase
1(1) WP_051922776.1 WP_051922776.1 hypothetical protein
1(2) WP_127484351.1 WP_127484351.1 DUF2974 domain-containing protein
1(1) WP_223592690.1 WP_223592690.1 DUF6382 domain-containing protein
1(1) WP_289613144.1 WP_289613144.1 FHA domain-containing protein
1(1) WP_249897321.1 WP_249897321.1 DUF2974 domain-containing protein
1(2) WP_022348880.1 WP_022348880.1 WXG100 family type VII secretion target
1(1) WP_159063398.1 WP_159063398.1 copper amine oxidase N-terminal domain-containing protein
1(1) WP_006292363.1 WP_006292363.1 YhgE/Pip domain-containing protein
1(1) WP_098645176.1 WP_098645176.1 T7SS effector LXG polymorphic toxin
1(1) WP_242172369.1 WP_242172369.1 hypothetical protein
1(1) WP_288161211.1 WP_288161211.1 hypothetical protein
1(1) WP_127199768.1 WP_127199768.1 hypothetical protein
1(2) WP_097952959.1 WP_097952959.1 DNA/RNA non-specific endonuclease
1(1) WP_144611180.1 WP_144611180.1 bacteriocin-processing peptidase family protein
1(1) WP_006292337.1 WP_006292337.1 FHA domain-containing protein
1(2) WP_160807262.1 WP_160807262.1 polysaccharide deacetylase family protein
1(1) WP_192510492.1 WP_192510492.1 WXG100 family type VII secretion target
1(2) WP_063188060.1 WP_063188060.1 WXG100 family type VII secretion target
1(1) WP_231417859.1 WP_231417859.1 protein kinase
1(1) WP_093291060.1 WP_093291060.1 hypothetical protein
1(1) WP_110934063.1 WP_110934063.1 copper amine oxidase N-terminal domain-containing protein
1(1) WP_204445206.1 WP_204445206.1 WXG100 family type VII secretion target
1(1) WP_167433840.1 WP_167433840.1 protein kinase family protein
1(1) WP_002118909.1 WP_002118909.1 WXG100 family type VII secretion target
1(1) WP_098319063.1 WP_098319063.1 bacteriocin-processing peptidase family protein
1(1) WP_134750602.1 WP_134750602.1 DUF2974 domain-containing protein
1(1) WP_068673705.1 WP_068673705.1 DUF2974 domain-containing protein
1(1) WP_006674862.1 WP_006674862.1 WXG100 family type VII secretion target
1(1) WP_044037270.1 WP_044037270.1 DUF6382 domain-containing protein
1(1) WP_164965495.1 WP_164965495.1 hypothetical protein
1(1) WP_137078365.1 WP_137078365.1 WXG100 family type VII secretion target
1(1) WP_065223574.1 WP_065223574.1 type VII secretion protein EsaA
1(1) WP_099517103.1 WP_099517103.1 WXG100 family type VII secretion target
1(1) WP_199677618.1 WP_199677618.1 bacteriocin-processing peptidase family protein
1(1) WP_077567019.1 WP_077567019.1 hypothetical protein
1(1) WP_215185459.1 WP_215185459.1 AAA family ATPase
1(1) WP_268058023.1 WP_268058023.1 hypothetical protein
1(3) WP_235349312.1 WP_235349312.1 DNA/RNA non-specific endonuclease
1(1) WP_066148103.1 WP_066148103.1 pore-forming ESAT-6 family protein
1(1) WP_307199964.1 WP_307199964.1 RHS repeat-associated core domain-containing protein
1(1) WP_169028888.1 WP_169028888.1 DUF2974 domain-containing protein
1(1) WP_065291248.1 WP_065291248.1 WXG100 family type VII secretion target
1(1) WP_307388976.1 WP_307388976.1 RHS repeat-associated core domain-containing protein
1(1) WP_097905194.1 WP_097905194.1 HNH endonuclease
1(1) WP_195431329.1 WP_195431329.1 hypothetical protein
1(1) WP_179160620.1 WP_179160620.1 DUF2974 domain-containing protein
1(2) WP_276565660.1 WP_276565660.1 hypothetical protein
1(1) WP_198065919.1 WP_198065919.1 WXG100 family type VII secretion target, partial
1(1) WP_227219192.1 WP_227219192.1 hypothetical protein
1(1) WP_242172374.1 WP_242172374.1 hypothetical protein
1(1) WP_076242357.1 WP_076242357.1 hypothetical protein
1(1) WP_277480696.1 WP_277480696.1 SseB family protein
1(1) WP_091260863.1 WP_091260863.1 hypothetical protein
1(1) WP_253411077.1 WP_253411077.1 pore-forming ESAT-6 family protein
1(1) WP_068782563.1 WP_068782563.1 WXG100 family type VII secretion target
1(1) WP_270721357.1 WP_270721357.1 hypothetical protein
1(1) WP_144655911.1 WP_144655911.1 hypothetical protein
1(1) WP_132283089.1 WP_132283089.1 DUF5050 domain-containing protein
1(1) WP_074094616.1 WP_074094616.1 hypothetical protein
1(1) WP_145408778.1 WP_145408778.1 hypothetical protein
1(1) WP_010250252.1 WP_010250252.1 polymorphic toxin type 44 domain-containing protein
1(1) WP_243500453.1 WP_243500453.1 tetratricopeptide repeat protein
1(1) WP_000788962.1 WP_000788962.1 hypothetical protein
1(3) WP_038978130.1 WP_038978130.1 membrane protein
1(1) WP_221875380.1 WP_221875380.1 WXG100 family type VII secretion target
1(1) WP_213618824.1 WP_213618824.1 glycosyl hydrolase family 18 protein
1(2) WP_281813078.1 WP_281813078.1 hypothetical protein
1(1) WP_235286205.1 WP_235286205.1 WXG100 family type VII secretion target
1(1) WP_145046455.1 WP_145046455.1 WXG100 family type VII secretion target
1(1) WP_017149758.1 WP_017149758.1 hypothetical protein
1(1) WP_084135370.1 WP_084135370.1 DUF2974 domain-containing protein
1(1) WP_141491316.1 WP_141491316.1 hypothetical protein, partial
1(1) WP_197069760.1 WP_197069760.1 WXG100 family type VII secretion target
1(1) WP_228596159.1 WP_228596159.1 EsaB/YukD family protein
1(1) WP_306077399.1 WP_306077399.1 glycosyltransferase
1(1) WP_141550442.1 WP_141550442.1 WXG100 family type VII secretion target, partial
1(1) WP_072727825.1 WP_072727825.1 hypothetical protein
1(1) WP_308615620.1 WP_308615620.1 hypothetical protein
1(11) WP_000788961.1 WP_000788961.1 hypothetical protein
1(10) WP_076104573.1 WP_076104573.1 hypothetical protein
1(1) WP_268058025.1 WP_268058025.1 WXG100 family type VII secretion target
1(1) WP_136789989.1 WP_136789989.1 zinc-ribbon domain-containing protein
1(1) WP_223592659.1 WP_223592659.1 WXG100 family type VII secretion target
1(1) WP_017692354.1 WP_017692354.1 RHS repeat-associated core domain-containing protein

| | | |
|---|---|---|
| 1(1) | WP_243495562.1 | WP_243495562.1 hypothetical protein |
| 1(1) | WP_306049032.1 | WP_306049032.1 FHA domain-containing protein |
| 1(1) | WP_224839869.1 | WP_224839869.1 Ig-like domain-containing protein |
| 1(37) | WP_001007571.1 | WP_001007571.1 EsaB/YukD family protein |
| 1(3) | WP_113713145.1 | WP_113713145.1 hypothetical protein |
| 1(1) | WP_094876484.1 | WP_094876484.1 DUF2974 domain-containing protein |
| 1(1) | WP_263324551.1 | WP_263324551.1 hypothetical protein |
| 1(2) | WP_204412721.1 | WP_204412721.1 WXG100 family type VII secretion target |
| 1(1) | WP_227219199.1 | WP_227219199.1 serine/threonine-protein kinase |
| 1(1) | WP_216830394.1 | WP_216830394.1 hypothetical protein |
| 1(1) | WP_088609115.1 | WP_088609115.1 tetratricopeptide repeat protein |
| 1(1) | WP_229139924.1 | WP_229139924.1 DUF6382 domain-containing protein |
| 1(1) | WP_013658723.1 | WP_013658723.1 hypothetical protein |
| 1(1) | WP_294953152.1 | WP_294953152.1 protein kinase |
| 1(1) | WP_226642887.1 | WP_226642887.1 Ig-like domain-containing protein |
| 1(1) | WP_243835626.1 | WP_243835626.1 HNH endonuclease |
| 1(1) | WP_063208810.1 | WP_063208810.1 S-layer homology domain-containing protein |
| 1(1) | WP_222861016.1 | WP_222861016.1 pre-toxin TG domain-containing protein |
| 1(1) | WP_270723907.1 | WP_270723907.1 tetratricopeptide repeat protein |
| 1(1) | WP_209861671.1 | WP_209861671.1 protease complex subunit PrcB family protein |
| 1(1) | WP_202283180.1 | WP_202283180.1 bacteriocin-processing peptidase family protein |
| 1(1) | WP_170287044.1 | WP_170287044.1 DUF5050 domain-containing protein |
| 1(1) | WP_024027638.1 | WP_024027638.1 Ig-like domain-containing protein |
| 1(2) | WP_017149772.1 | WP_017149772.1 EsaB/YukD family protein |
| 1(1) | WP_233697172.1 | WP_233697172.1 hypothetical protein |
| 1(1) | WP_015595778.1 | WP_015595778.1 glycosyltransferase |
| 1(1) | WP_076318551.1 | WP_076318551.1 hypothetical protein |
| 1(1) | WP_251548753.1 | WP_251548753.1 pore-forming ESAT-6 family protein |
| 1(1) | WP_149096534.1 | WP_149096534.1 hypothetical protein |
| 1(1) | WP_006720303.1 | WP_006720303.1 hypothetical protein |
| 1(1) | WP_272792286.1 | WP_272792286.1 WXG100 family type VII secretion target, partial |
| 1(4) | WP_049720635.1 | WP_049720635.1 WXG100 family type VII secretion target |
| 1(1) | WP_283952949.1 | WP_283952949.1 WXG100 family type VII secretion target |
| 1(1) | WP_002031616.1 | WP_002031616.1 tetratricopeptide repeat protein |
| 1(1) | WP_088078455.1 | WP_088078455.1 tetratricopeptide repeat protein |
| 1(1) | WP_247083045.1 | WP_247083045.1 WXG100 family type VII secretion target |
| 1(1) | WP_289342270.1 | WP_289342270.1 hypothetical protein |
| 1(1) | WP_013914183.1 | WP_013914183.1 S-layer homology domain-containing protein |
| 1(1) | WP_257675921.1 | WP_257675921.1 tetratricopeptide repeat protein |
| 1(9) | WP_001990254.1 | WP_001990254.1 T7SS effector LXG polymorphic toxin |
| 1(2) | WP_025719081.1 | WP_025719081.1 S-layer homology domain-containing protein |
| 1(1) | WP_006211203.1 | WP_006211203.1 hypothetical protein |
| 1(3) | WP_000506921.1 | WP_000506921.1 tetratricopeptide repeat protein |
| 1(1) | WP_119794362.1 | WP_119794362.1 WXG100 family type VII secretion target |
| 1(1) | WP_074598761.1 | WP_074598761.1 type VII secretion protein EsaA |
| 1(1) | WP_113755282.1 | WP_113755282.1 WG repeat-containing protein |
| 1(1) | WP_290409169.1 | WP_290409169.1 S-layer homology domain-containing protein |
| 1(5) | WP_023988266.1 | WP_023988266.1 WXG100 family type VII secretion target |
| 1(1) | WP_278773463.1 | WP_278773463.1 hypothetical protein |
| 1(1) | WP_212330909.1 | WP_212330909.1 hypothetical protein |
| 1(2) | WP_186313735.1 | WP_186313735.1 WXG100 family type VII secretion target |

| | | |
|---|---|---|
| 1(1) | WP_188535078.1 | WP_188535078.1 copper amine oxidase N-terminal domain-containing protein |
| 1(1) | WP_143315252.1 | WP_143315252.1 EsaB/YukD family protein |
| 1(1) | WP_024630328.1 | WP_024630328.1 membrane protein |
| 1(1) | WP_074085357.1 | WP_074085357.1 hypothetical protein |
| 1(2) | WP_005210448.1 | WP_005210448.1 pore-forming ESAT-6 family protein |
| 1(1) | WP_097987487.1 | WP_097987487.1 hypothetical protein |
| 1(1) | WP_049869169.1 | WP_049869169.1 hypothetical protein |
| 1(1) | WP_100537974.1 | WP_100537974.1 hypothetical protein |
| 1(1) | WP_139867392.1 | WP_139867392.1 bacteriocin-processing peptidase family protein |
| 1(1) | WP_005604621.1 | WP_005604621.1 FHA domain-containing protein |
| 1(1) | WP_213649969.1 | WP_213649969.1 hypothetical protein |
| 1(1) | WP_022128273.1 | WP_022128273.1 pore-forming ESAT-6 family protein |
| 1(1) | WP_127570435.1 | WP_127570435.1 hypothetical protein |
| 1(1) | WP_253440787.1 | WP_253440787.1 WXG100 family type VII secretion target |
| 1(4) | WP_098254023.1 | WP_098254023.1 SseB family protein |
| 1(2) | WP_185357238.1 | WP_185357238.1 type VII secretion protein EsaA |
| 1(2) | WP_186326816.1 | WP_186326816.1 DUF2974 domain-containing protein |
| 1(1) | WP_256872562.1 | WP_256872562.1 WXG100 family type VII secretion target |
| 1(1) | WP_294953141.1 | WP_294953141.1 pore-forming ESAT-6 family protein |
| 1(1) | WP_034675371.1 | WP_034675371.1 pore-forming ESAT-6 family protein |
| 1(1) | WP_195431324.1 | WP_195431324.1 DUF6382 domain-containing protein |
| 1(1) | WP_127459432.1 | WP_127459432.1 pore-forming ESAT-6 family protein |
| 1(1) | WP_212693237.1 | WP_212693237.1 hypothetical protein |
| 1(1) | WP_215591201.1 | WP_215591201.1 hypothetical protein |
| 1(1) | WP_048826563.1 | WP_048826563.1 DUF2974 domain-containing protein |
| 1(1) | WP_212693227.1 | WP_212693227.1 peptidoglycan-binding protein |
| 1(1) | WP_251417679.1 | WP_251417679.1 hypothetical protein |
| 1(1) | WP_179160614.1 | WP_179160614.1 protein kinase |
| 1(1) | WP_078419970.1 | WP_078419970.1 tetratricopeptide repeat protein |
| 1(3) | WP_076856200.1 | WP_076856200.1 bacteriocin-processing peptidase family protein |
| 1(1) | WP_305755032.1 | WP_305755032.1 hypothetical protein |
| 1(1) | WP_013658696.1 | WP_013658696.1 WXG100 family type VII secretion target |
| 1(2) | WP_041964581.1 | WP_041964581.1 hypothetical protein |
| 1(1) | WP_212930978.1 | WP_212930978.1 type VII secretion target |
| 1(2) | WP_213117476.1 | WP_213117476.1 serine hydrolase |
| 1(1) | WP_308082250.1 | WP_308082250.1 FHA domain-containing protein |
| 1(3) | WP_113755276.1 | WP_113755276.1 type VII secretion target |
| 1(1) | WP_110892978.1 | WP_110892978.1 hypothetical protein |
| 1(2) | WP_179603487.1 | WP_179603487.1 hypothetical protein |
| 1(1) | WP_076079061.1 | WP_076079061.1 hypothetical protein |
| 1(1) | WP_058829623.1 | WP_058829623.1 WXG100 family type VII secretion target |
| 1(1) | WP_090993621.1 | WP_090993621.1 hypothetical protein |
| 1(1) | WP_098991954.1 | WP_098991954.1 deaminase domain-containing protein |
| 1(1) | WP_309098618.1 | WP_309098618.1 hypothetical protein |
| 1(1) | WP_249897317.1 | WP_249897317.1 WXG100 family type VII secretion target |
| 1(1) | WP_007083294.1 | WP_007083294.1 DUF6382 domain-containing protein |
| 1(1) | WP_209861645.1 | WP_209861645.1 hypothetical protein |
| 1(2) | WP_075718895.1 | WP_075718895.1 tetratricopeptide repeat protein |
| 1(1) | WP_087952376.1 | WP_087952376.1 tetratricopeptide repeat protein |
| 1(2) | WP_258126014.1 | WP_258126014.1 WXG100 family type VII secretion target |
| 1(1) | WP_195325251.1 | WP_195325251.1 tetratricopeptide repeat protein |

1(1) WP_211082924.1 WP_211082924.1 WXG100 family type VII secretion target
1(3) WP_040305095.1 WP_040305095.1 WXG100 family type VII secretion target
1(1) WP_312853825.1 WP_312853825.1 EsaB/YukD family protein
1(2) WP_195951611.1 WP_195951611.1 hypothetical protein
1(1) WP_301167501.1 WP_301167501.1 hypothetical protein
1(1) WP_053403780.1 WP_053403780.1 tetratricopeptide repeat protein
1(1) WP_124049168.1 WP_124049168.1 hypothetical protein
1(1) WP_208752239.1 WP_208752239.1 WXG100 family type VII secretion target
1(8) WP_009590697.1 WP_009590697.1 hypothetical protein
1(1) WP_109625238.1 WP_109625238.1 DUF5050 domain-containing protein
1(1) WP_010250269.1 WP_010250269.1 WXG100 family type VII secretion target
1(1) WP_104495352.1 WP_104495352.1 hypothetical protein
1(1) WP_239617476.1 WP_239617476.1 pore-forming ESAT-6 family protein
1(1) WP_111147802.1 WP_111147802.1 pore-forming ESAT-6 family protein
1(1) WP_106822152.1 WP_106822152.1 WXG100 family type VII secretion target
1(2) WP_271839914.1 WP_271839914.1 WXG100 family type VII secretion target
1(1) WP_111273168.1 WP_111273168.1 WXG100 family type VII secretion target
1(1) WP_185463409.1 WP_185463409.1 hypothetical protein
1(1) WP_076279383.1 WP_076279383.1 hypothetical protein
1(1) WP_220775546.1 WP_220775546.1 SseB family protein
1(3) WP_002093976.1 WP_002093976.1 hypothetical protein
1(9) WP_042400994.1 WP_042400994.1 pore-forming ESAT-6 family protein
1(1) WP_110931549.1 WP_110931549.1 pre-toxin TG domain-containing protein
1(1) WP_205001936.1 WP_205001936.1 tetratricopeptide repeat protein
1(1) WP_158326752.1 WP_158326752.1 WG repeat-containing protein
1(1) WP_268623566.1 WP_268623566.1 copper amine oxidase N-terminal domain-containing
protein
1(1) WP_001050358.1 WP_001050358.1 SseB family protein
1(1) WP_010250288.1 WP_010250288.1 stalk domain-containing protein
1(1) WP_098167593.1 WP_098167593.1 WXG100 family type VII secretion target
1(1) WP_210364456.1 WP_210364456.1 pore-forming ESAT-6 family protein
1(1) WP_052091740.1 WP_052091740.1 hypothetical protein
1(1) WP_097905176.1 WP_097905176.1 SseB family protein
1(1) WP_182301831.1 WP_182301831.1 DUF2974 domain-containing protein
1(1) WP_281937370.1 WP_281937370.1 pore-forming ESAT-6 family protein
1(1) WP_098288678.1 WP_098288678.1 bacteriocin-processing peptidase family protein
1(1) WP_015595757.1 WP_015595757.1 WXG100 family type VII secretion target
1(1) WP_307601050.1 WP_307601050.1 glycohydrolase toxin TNT-related protein
1(1) WP_017560551.1 WP_017560551.1 DNA/RNA non-specific endonuclease
1(1) WP_283904346.1 WP_283904346.1 LysM peptidoglycan-binding domain-containing protein
1(1) WP_098148066.1 WP_098148066.1 polymorphic toxin type 15 domain-containing protein
1(1) WP_257215860.1 WP_257215860.1 Ig-like domain-containing protein
1(1) WP_000034619.1 WP_000034619.1 T7SS effector LXG polymorphic toxin
1(1) WP_277713351.1 WP_277713351.1 hypothetical protein
1(1) WP_126996321.1 WP_126996321.1 pesticin C-terminus-like muramidase
1(1) WP_042400992.1 WP_042400992.1 WXG100 family type VII secretion target
1(2) WP_271000081.1 WP_271000081.1 pre-toxin TG domain-containing protein
1(1) WP_129549124.1 WP_129549124.1 S-layer homology domain-containing protein
1(1) WP_046953664.1 WP_046953664.1 T7SS effector LXG polymorphic toxin
1(1) WP_170833626.1 WP_170833626.1 hypothetical protein
1(1) WP_220027683.1 WP_220027683.1 hypothetical protein
1(1) WP_257443651.1 WP_257443651.1 DUF5050 domain-containing protein
1(1) WP_213592854.1 WP_213592854.1 copper amine oxidase N-terminal domain-containing
protein
1(1) WP_111273167.1 WP_111273167.1 WXG100 family type VII secretion target
1(1) WP_088830548.1 WP_088830548.1 cadherin-like beta sandwich domain-containing protein
1(1) WP_076113457.1 WP_076113457.1 hypothetical protein
1(21) WP_002141592.1 WP_002141592.1 WXG100 family type VII secretion target
1(2) WP_185469291.1 WP_185469291.1 glycohydrolase toxin TNT-related protein
1(1) WP_242686561.1 WP_242686561.1 type VII secretion protein EsaA
1(1) WP_263880217.1 WP_263880217.1 WXG100 family type VII secretion target
1(1) WP_109999892.1 WP_109999892.1 hypothetical protein
1(1) WP_028542696.1 WP_028542696.1 S-layer homology domain-containing protein
1(1) WP_234489640.1 WP_234489640.1 S-layer homology domain-containing protein
1(1) WP_141435886.1 WP_141435886.1 hypothetical protein
1(1) WP_277480672.1 WP_277480672.1 tetratricopeptide repeat protein
1(1) WP_148246950.1 WP_148246950.1 serine/threonine protein kinase
1(1) WP_315968501.1 WP_315968501.1 toxin
1(1) WP_047153270.1 WP_047153270.1 hypothetical protein
1(1) WP_095691005.1 WP_095691005.1 polymorphic toxin type 15 domain-containing protein
1(2) WP_228469894.1 WP_228469894.1 glycohydrolase toxin TNT-related protein
1(1) WP_191272871.1 WP_191272871.1 hypothetical protein
1(1) WP_119601480.1 WP_119601480.1 DUF2974 domain-containing protein
1(1) WP_307388984.1 WP_307388984.1 DUF6531 domain-containing protein
1(1) WP_282321819.1 WP_282321819.1 hypothetical protein
1(4) WP_203573357.1 WP_203573357.1 type VII secretion protein EsaA
1(1) WP_243186738.1 WP_243186738.1 DNA/RNA non-specific endonuclease
1(1) WP_000392228.1 WP_000392228.1 hypothetical protein
1(1) WP_036702410.1 WP_036702410.1 hypothetical protein
1(1) WP_078993362.1 WP_078993362.1 toxin
1(1) WP_144655909.1 WP_144655909.1 HNH endonuclease
1(1) WP_054972106.1 WP_054972106.1 cadherin-like beta sandwich domain-containing protein
1(1) WP_077320213.1 WP_077320213.1 hypothetical protein
1(2) WP_069080956.1 WP_069080956.1 WXG100 family type VII secretion target
1(1) WP_068619096.1 WP_068619096.1 YhgE/Pip domain-containing protein
1(1) WP_036642155.1 WP_036642155.1 hypothetical protein
1(1) WP_270723902.1 WP_270723902.1 hypothetical protein
1(3) WP_098361431.1 WP_098361431.1 WXG100 family type VII secretion target
1(1) WP_232243181.1 WP_232243181.1 hypothetical protein
1(1) WP_142543125.1 WP_142543125.1 hypothetical protein
1(1) WP_166245133.1 WP_166245133.1 serine/threonine-protein kinase
1(1) WP_251583291.1 WP_251583291.1 WXG100 family type VII secretion target
1(1) WP_258208935.1 WP_258208935.1 hypothetical protein, partial
1(1) WP_128631276.1 WP_128631276.1 hypothetical protein
1(1) WP_258418573.1 WP_258418573.1 hypothetical protein
1(1) WP_089522636.1 WP_089522636.1 hypothetical protein
1(1) WP_229717041.1 WP_229717041.1 DL-endopeptidase inhibitor IseA family protein
1(1) WP_307192902.1 WP_307192902.1 hypothetical protein
1(1) WP_258417126.1 WP_258417126.1 WG repeat-containing protein
1(1) WP_148450939.1 WP_148450939.1 NlpC/P60 family protein
1(1) WP_124114119.1 WP_124114119.1 hypothetical protein
1(1) WP_278740194.1 WP_278740194.1 hypothetical protein
1(1) WP_127592427.1 WP_127592427.1 hypothetical protein
1(1) WP_136379705.1 WP_136379705.1 serine/threonine-protein kinase

| | | | |
|---|---|---|---|
| 1(2) | WP_097898660.1 | WP_097898660.1 | serine/threonine-protein kinase |
| 1(1) | WP_307394630.1 | WP_307394630.1 | WXG100 family type VII secretion target |
| 1(1) | WP_272045570.1 | WP_272045570.1 | hypothetical protein |
| 1(1) | WP_136372956.1 | WP_136372956.1 | WXG100 family type VII secretion target |
| 1(1) | WP_089546601.1 | WP_089546601.1 | hypothetical protein |
| 1(1) | WP_307738721.1 | WP_307738721.1 | Ig-like domain-containing protein |
| 1(1) | WP_182301836.1 | WP_182301836.1 | pore-forming ESAT-6 family protein |
| 1(1) | WP_147807661.1 | WP_147807661.1 | bacteriocin-processing peptidase family protein |
| 1(1) | WP_301804652.1 | WP_301804652.1 | WG repeat-containing protein |
| 1(1) | WP_084117736.1 | WP_084117736.1 | DUF6382 domain-containing protein |
| 1(2) | WP_068940669.1 | WP_068940669.1 | S-layer homology domain-containing protein |
| 1(1) | WP_152402378.1 | WP_152402378.1 | WXG100 family type VII secretion target |
| 1(1) | WP_110892972.1 | WP_110892972.1 | WXG100 family type VII secretion target |
| 1(1) | WP_015595751.1 | WP_015595751.1 | Ig-like domain-containing protein |
| 1(1) | WP_076157828.1 | WP_076157828.1 | hypothetical protein |
| 1(1) | WP_076208260.1 | WP_076208260.1 | hypothetical protein |
| 1(1) | WP_036707105.1 | WP_036707105.1 | WXG100 family type VII secretion target |
| 1(2) | WP_113672696.1 | WP_113672696.1 | hypothetical protein |
| 1(1) | WP_002158045.1 | WP_002158045.1 | hypothetical protein |
| 1(4) | WP_037993172.1 | WP_037993172.1 | EsaB/YukD family protein |
| 1(1) | WP_010676863.1 | WP_010676863.1 | DUF2974 domain-containing protein |
| 1(1) | WP_222655406.1 | WP_222655406.1 | HNH endonuclease |
| 1(1) | WP_038286703.1 | WP_038286703.1 | hypothetical protein |
| 1(2) | WP_137013298.1 | WP_137013298.1 | hypothetical protein |
| 1(1) | WP_294953149.1 | WP_294953149.1 | DUF6382 domain-containing protein |
| 1(1) | WP_090688978.1 | WP_090688978.1 | hypothetical protein |
| 1(1) | WP_301249699.1 | WP_301249699.1 | hypothetical protein |
| 1(1) | WP_098943212.1 | WP_098943212.1 | hypothetical protein |
| 1(1) | WP_163258050.1 | WP_163258050.1 | bacteriocin-processing peptidase family protein |
| 1(2) | WP_101300044.1 | WP_101300044.1 | DUF5050 domain-containing protein |
| 1(1) | WP_098834581.1 | WP_098834581.1 | hypothetical protein |
| 1(1) | WP_307284693.1 | WP_307284693.1 | serine/threonine-protein kinase |
| 1(2) | WP_194346728.1 | WP_194346728.1 | type VII secretion protein EsaA |
| 1(1) | WP_017692362.1 | WP_017692362.1 | DUF2974 domain-containing protein |
| 1(1) | WP_307399121.1 | WP_307399121.1 | FHA domain-containing protein |
| 1(2) | WP_169028887.1 | WP_169028887.1 | copper amine oxidase N-terminal domain-containing protein |
| 1(1) | WP_071558848.1 | WP_071558848.1 | S-layer homology domain-containing protein |
| 1(1) | WP_212981559.1 | WP_212981559.1 | protein kinase family protein |
| 1(1) | WP_098167601.1 | WP_098167601.1 | bacteriocin-processing peptidase family protein |
| 1(1) | WP_053361015.1 | WP_053361015.1 | Ig-like domain-containing protein |
| 1(1) | WP_238655357.1 | WP_238655357.1 | hypothetical protein |
| 1(1) | WP_144479070.1 | WP_144479070.1 | LysM peptidoglycan-binding domain-containing protein |
| 1(1) | WP_000392222.1 | WP_000392222.1 | hypothetical protein |
| 1(1) | WP_163245177.1 | WP_163245177.1 | WG repeat-containing protein |
| 1(1) | WP_257443621.1 | WP_257443621.1 | pre-toxin TG domain-containing protein |
| 1(1) | WP_210087110.1 | WP_210087110.1 | pre-toxin TG domain-containing protein |
| 1(2) | WP_195431325.1 | WP_195431325.1 | hypothetical protein |
| 1(1) | WP_141527795.1 | WP_141527795.1 | hypothetical protein |
| 1(1) | WP_036689404.1 | WP_036689404.1 | DUF2974 domain-containing protein |
| 1(1) | WP_242264963.1 | WP_242264963.1 | bacteriocin-processing peptidase family protein |
| 1(1) | WP_186580079.1 | WP_186580079.1 | DUF2974 domain-containing protein |
| 1(1) | WP_075395696.1 | WP_075395696.1 | tetratricopeptide repeat protein |
| 1(1) | WP_241752504.1 | WP_241752504.1 | hypothetical protein |
| 1(1) | WP_000918598.1 | WP_000918598.1 | WXG100 family type VII secretion target |
| 1(1) | WP_241973190.1 | WP_241973190.1 | hypothetical protein, partial |
| 1(1) | WP_241661647.1 | WP_241661647.1 | bacteriocin-processing peptidase family protein, partial |
| 1(1) | WP_076271937.1 | WP_076271937.1 | DUF2974 domain-containing protein |
| 1(1) | WP_297035092.1 | WP_297035092.1 | hypothetical protein, partial |
| 1(1) | WP_053361020.1 | WP_053361020.1 | pore-forming ESAT-6 family protein |
| 1(1) | WP_251638784.1 | WP_251638784.1 | WXG100 family type VII secretion target |
| 1(1) | WP_006292347.1 | WP_006292347.1 | WXG100 family type VII secretion target |
| 1(1) | WP_260065262.1 | WP_260065262.1 | hypothetical protein |
| 1(1) | WP_270574773.1 | WP_270574773.1 | WXG100 family type VII secretion target |
| 1(1) | WP_094870400.1 | WP_094870400.1 | hypothetical protein |
| 1(1) | WP_204445223.1 | WP_204445223.1 | hypothetical protein |
| 1(1) | WP_149885340.1 | WP_149885340.1 | WXG100 family type VII secretion target, partial |
| 1(2) | WP_076281071.1 | WP_076281071.1 | DUF2974 domain-containing protein |
| 1(1) | WP_143316564.1 | WP_143316564.1 | pore-forming ESAT-6 family protein |
| 1(1) | WP_289613149.1 | WP_289613149.1 | DUF2974 domain-containing protein |
| 1(1) | WP_283157900.1 | WP_283157900.1 | hypothetical protein |
| 1(1) | WP_270472014.1 | WP_270472014.1 | serine/threonine-protein kinase |
| 1(1) | WP_253411072.1 | WP_253411072.1 | DUF2974 domain-containing protein |
| 1(2) | WP_002118898.1 | WP_002118898.1 | WXG100 family type VII secretion target |
| 1(1) | WP_149871696.1 | WP_149871696.1 | serine hydrolase domain-containing protein |
| 1(1) | WP_183578452.1 | WP_183578452.1 | WXG100 family type VII secretion target |
| 1(1) | WP_279664453.1 | WP_279664453.1 | WXG100 family type VII secretion target |
| 1(1) | WP_280925141.1 | WP_280925141.1 | serine/threonine-protein kinase |
| 1(3) | WP_231564555.1 | WP_231564555.1 | EsaB/YukD family protein |
| 1(1) | WP_289475125.1 | WP_289475125.1 | hypothetical protein |
| 1(1) | WP_142254720.1 | WP_142254720.1 | hypothetical protein |
| 1(3) | WP_079346819.1 | WP_079346819.1 | hypothetical protein |
| 1(1) | WP_103048248.1 | WP_103048248.1 | WXG100 family type VII secretion target |
| 1(1) | WP_307199938.1 | WP_307199938.1 | WXG100 family type VII secretion target |
| 1(1) | WP_277480692.1 | WP_277480692.1 | tetratricopeptide repeat protein |
| 1(1) | WP_270472016.1 | WP_270472016.1 | Ig-like domain-containing protein |
| 1(1) | WP_058486870.1 | WP_058486870.1 | EsaB/YukD family protein |
| 1(1) | WP_000357378.1 | WP_000357378.1 | DNA/RNA non-specific endonuclease |
| 1(1) | WP_090761821.1 | WP_090761821.1 | DUF2974 domain-containing protein |
| 1(1) | WP_243495551.1 | WP_243495551.1 | WXG100 family type VII secretion target |
| 1(1) | WP_169509269.1 | WP_169509269.1 | WXG100 family type VII secretion target, partial |
| 1(25) | WP_000391132.1 | WP_000391132.1 | hypothetical protein |
| 1(1) | WP_255260183.1 | WP_255260183.1 | WXG100 family type VII secretion target |
| 1(1) | WP_229156364.1 | WP_229156364.1 | hypothetical protein, partial |
| 1(1) | WP_000404137.1 | WP_000404137.1 | WXG100 family type VII secretion target |
| 1(1) | WP_236334336.1 | WP_236334336.1 | EsaB/YukD family protein |
| 1(1) | WP_098856766.1 | WP_098856766.1 | hypothetical protein |
| 1(1) | WP_215570364.1 | WP_215570364.1 | tetratricopeptide repeat protein |
| 1(1) | WP_205602701.1 | WP_205602701.1 | GH-E family nuclease |
| 1(2) | WP_090896181.1 | WP_090896181.1 | hypothetical protein |
| 1(1) | WP_263324547.1 | WP_263324547.1 | WXG100 family type VII secretion target |
| 1(1) | WP_283952945.1 | WP_283952945.1 | WXG100 family type VII secretion target |
| 1(1) | WP_150457763.1 | WP_150457763.1 | WXG100 family type VII secretion target |
| 1(1) | WP_307199942.1 | WP_307199942.1 | EsaB/YukD family protein |

| | | |
|---|---|---|
| 1(1) | WP_284036496.1 | WP_284036496.1 WXG100 family type VII secretion target |
| 1(1) | WP_094870533.1 | WP_094870533.1 WXG100 family type VII secretion target |
| 1(1) | WP_141544305.1 | WP_141544305.1 hypothetical protein |
| 1(1) | WP_098776427.1 | WP_098776427.1 SseB family protein |
| 1(1) | WP_204666341.1 | WP_204666341.1 T7SS effector LXG polymorphic toxin |
| 1(1) | WP_230493912.1 | WP_230493912.1 copper amine oxidase N-terminal domain-containing protein |
| 1(1) | WP_077832298.1 | WP_077832298.1 tetratricopeptide repeat protein |
| 1(1) | WP_238799331.1 | WP_238799331.1 copper amine oxidase N-terminal domain-containing protein |
| 1(1) | WP_094092619.1 | WP_094092619.1 hypothetical protein |
| 1(1) | WP_305166261.1 | WP_305166261.1 WXG100 family type VII secretion target |
| 1(1) | WP_197234718.1 | WP_197234718.1 hypothetical protein |
| 1(1) | WP_102264542.1 | WP_102264542.1 Ig-like domain-containing protein |
| 1(1) | WP_291647936.1 | WP_291647936.1 hypothetical protein, partial |
| 1(1) | WP_042134269.1 | WP_042134269.1 hypothetical protein |
| 1(2) | WP_116353249.1 | WP_116353249.1 WXG100 family type VII secretion target |
| 1(1) | WP_199795354.1 | WP_199795354.1 WXG100 family type VII secretion target |
| 1(1) | WP_176124740.1 | WP_176124740.1 hypothetical protein |
| 1(1) | WP_289613134.1 | WP_289613134.1 WXG100 family type VII secretion target |
| 1(1) | WP_104030445.1 | WP_104030445.1 WG repeat-containing protein |
| 1(2) | WP_283655694.1 | WP_283655694.1 WXG100 family type VII secretion target |
| 1(1) | WP_241559289.1 | WP_241559289.1 serine/threonine-protein kinase |
| 1(1) | WP_083180284.1 | WP_083180284.1 DNA/RNA non-specific endonuclease |
| 1(1) | WP_013252385.1 | WP_013252385.1 hypothetical protein |
| 1(3) | WP_046325930.1 | WP_046325930.1 DUF6531 domain-containing protein |
| 1(1) | WP_159146506.1 | WP_159146506.1 hypothetical protein |
| 1(1) | WP_098303176.1 | WP_098303176.1 SseB family protein |
| 1(1) | WP_079506412.1 | WP_079506412.1 Ig-like domain-containing protein |
| 1(4) | WP_156558885.1 | WP_156558885.1 EsaB/YukD family protein |
| 1(1) | WP_273322274.1 | WP_273322274.1 CdiA family toxin C-terminal domain-containing protein |
| 1(1) | WP_291288266.1 | WP_291288266.1 hypothetical protein |
| 1(1) | WP_055277386.1 | WP_055277386.1 hypothetical protein |
| 1(1) | WP_213592850.1 | WP_213592850.1 serine/threonine-protein kinase |
| 1(1) | WP_214328825.1 | WP_214328825.1 hypothetical protein |
| 1(1) | WP_236334315.1 | WP_236334315.1 copper amine oxidase N-terminal domain-containing protein |
| 1(1) | WP_244723776.1 | WP_244723776.1 hypothetical protein |
| 1(2) | WP_121610597.1 | WP_121610597.1 WXG100 family type VII secretion target |
| 1(1) | WP_254366006.1 | WP_254366006.1 DUF2974 domain-containing protein |
| 1(1) | WP_154118032.1 | WP_154118032.1 hypothetical protein |
| 1(1) | WP_232699383.1 | WP_232699383.1 hypothetical protein |
| 1(1) | WP_199419777.1 | WP_199419777.1 peptidoglycan endopeptidase |
| 1(1) | WP_240379733.1 | WP_240379733.1 tetratricopeptide repeat protein |
| 1(7) | WP_001007566.1 | WP_001007566.1 EsaB/YukD family protein |
| 1(1) | WP_244719603.1 | WP_244719603.1 hypothetical protein |
| 1(2) | WP_026571199.1 | WP_026571199.1 WXG100 family type VII secretion target |
| 1(3) | WP_087433608.1 | WP_087433608.1 WXG100 family type VII secretion target |
| 1(1) | WP_210166179.1 | WP_210166179.1 FHA domain-containing protein |
| 1(1) | WP_076210975.1 | WP_076210975.1 hypothetical protein |
| 1(1) | WP_305755034.1 | WP_305755034.1 WXG100 family type VII secretion target |
| 1(13) | WP_002158043.1 | WP_002158043.1 WXG100 family type VII secretion target |
| 1(9) | WP_015736242.1 | WP_015736242.1 hypothetical protein |
| 1(1) | WP_157418143.1 | WP_157418143.1 bacteriocin-processing peptidase family protein |
| 1(1) | WP_208919044.1 | WP_208919044.1 serine hydrolase domain-containing protein |
| 1(1) | WP_302579552.1 | WP_302579552.1 WXG100 family type VII secretion target |
| 1(1) | WP_208926609.1 | WP_208926609.1 hypothetical protein |
| 1(1) | WP_289613142.1 | WP_289613142.1 T7SS effector LXG polymorphic toxin |
| 1(1) | WP_231690738.1 | WP_231690738.1 hypothetical protein |
| 1(3) | WP_077833108.1 | WP_077833108.1 WXG100 family type VII secretion target |
| 1(1) | WP_270472024.1 | WP_270472024.1 hypothetical protein |
| 1(1) | WP_144611187.1 | WP_144611187.1 hypothetical protein |
| 1(3) | WP_038569545.1 | WP_038569545.1 DUF2974 domain-containing protein |
| 1(1) | WP_068673712.1 | WP_068673712.1 WXG100 family type VII secretion target |
| 1(1) | WP_229136363.1 | WP_229136363.1 bacteriocin-processing peptidase family protein |
| 1(1) | WP_247681892.1 | WP_247681892.1 hypothetical protein |
| 1(1) | WP_302153073.1 | WP_302153073.1 FHA domain-containing protein |
| 1(1) | WP_098400684.1 | WP_098400684.1 WXG100 family type VII secretion target |
| 1(1) | WP_051421037.1 | WP_051421037.1 WXG100 family type VII secretion target |
| 1(7) | WP_162164277.1 | WP_162164277.1 WXG100 family type VII secretion target |
| 1(1) | WP_307738716.1 | WP_307738716.1 protein kinase |
| 1(1) | WP_267255471.1 | WP_267255471.1 type VII secretion protein EsaA |
| 1(1) | WP_223254474.1 | WP_223254474.1 Ig-like domain-containing protein |
| 1(1) | WP_248063442.1 | WP_248063442.1 S-layer homology domain-containing protein |
| 1(2) | WP_048749331.1 | WP_048749331.1 WXG100 family type VII secretion target |
| 1(1) | WP_077893455.1 | WP_077893455.1 tetratricopeptide repeat protein |
| 1(1) | WP_062833882.1 | WP_062833882.1 hypothetical protein |
| 1(1) | WP_127194954.1 | WP_127194954.1 penicillin-insensitive murein endopeptidase |
| 1(1) | WP_212930974.1 | WP_212930974.1 WXG100 family type VII secretion target |
| 1(2) | WP_194336457.1 | WP_194336457.1 pre-toxin TG domain-containing protein |
| 1(1) | WP_168419970.1 | WP_168419970.1 WG repeat-containing protein |
| 1(1) | WP_230493913.1 | WP_230493913.1 hypothetical protein |
| 1(1) | WP_078185403.1 | WP_078185403.1 hypothetical protein |
| 1(1) | WP_198059701.1 | WP_198059701.1 hypothetical protein |
| 1(1) | WP_265408843.1 | WP_265408843.1 hypothetical protein |
| 1(2) | WP_185461323.1 | WP_185461323.1 type VII secretion protein EsaA |
| 1(4) | WP_061660129.1 | WP_061660129.1 SseB family protein |
| 1(1) | WP_053102087.1 | WP_053102087.1 hypothetical protein |
| 1(1) | WP_163853473.1 | WP_163853473.1 WXG100 family type VII secretion target |
| 1(1) | WP_254759298.1 | WP_254759298.1 DUF2974 domain-containing protein |
| 1(1) | WP_267414296.1 | WP_267414296.1 hypothetical protein |
| 1(1) | WP_270711669.1 | WP_270711669.1 tetratricopeptide repeat protein |
| 1(1) | WP_094936488.1 | WP_094936488.1 hypothetical protein |
| 1(11) | WP_063566322.1 | WP_063566322.1 hypothetical protein |
| 1(1) | WP_308083985.1 | WP_308083985.1 WXG100 family type VII secretion target |
| 1(1) | WP_179160627.1 | WP_179160627.1 hypothetical protein |
| 1(1) | WP_243270452.1 | WP_243270452.1 FHA domain-containing protein |
| 1(1) | WP_170138180.1 | WP_170138180.1 WXG100 family type VII secretion target |
| 1(1) | WP_089546602.1 | WP_089546602.1 hypothetical protein |
| 1(1) | WP_295628347.1 | WP_295628347.1 tetratricopeptide repeat protein |
| 1(2) | WP_088020919.1 | WP_088020919.1 tetratricopeptide repeat protein |
| 1(1) | WP_169028894.1 | WP_169028894.1 EsaB/YukD family protein |
| 1(1) | WP_052414524.1 | WP_052414524.1 copper amine oxidase N-terminal domain-containing protein |

| | | |
|---|---|---|
| 1(1) | WP_208926592.1 | WP_208926592.1 hypothetical protein |
| 1(2) | WP_076101574.1 | WP_076101574.1 DUF2974 domain-containing protein |
| 1(2) | WP_036684506.1 | WP_036684506.1 AAA family ATPase |
| 1(1) | WP_220027672.1 | WP_220027672.1 S-layer homology domain-containing protein |
| 1(3) | WP_000034623.1 | WP_000034623.1 T7SS effector LXG polymorphic toxin |
| 1(1) | WP_145134817.1 | WP_145134817.1 WXG100 family type VII secretion target |
| 1(1) | WP_268613671.1 | WP_268613671.1 WXG100 family type VII secretion target |
| 1(2) | WP_270995166.1 | WP_270995166.1 DUF6531 domain-containing protein |
| 1(2) | WP_087433603.1 | WP_087433603.1 hypothetical protein |
| 1(1) | WP_077320199.1 | WP_077320199.1 WXG100 family type VII secretion target |
| 1(1) | WP_101668487.1 | WP_101668487.1 hypothetical protein |
| 1(1) | WP_306049008.1 | WP_306049008.1 LysM peptidoglycan-binding domain-containing protein |
| 1(1) | WP_242251781.1 | WP_242251781.1 hypothetical protein |
| 1(1) | WP_257209345.1 | WP_257209345.1 type VII secretion protein EsaA, partial |
| 1(1) | WP_291682717.1 | WP_291682717.1 hypothetical protein, partial |
| 1(1) | WP_181603061.1 | WP_181603061.1 WXG100 family type VII secretion target |
| 1(1) | WP_284566221.1 | WP_284566221.1 tetratricopeptide repeat protein |
| 1(1) | WP_306077386.1 | WP_306077386.1 FHA domain-containing protein |
| 1(1) | WP_278773427.1 | WP_278773427.1 DUF2974 domain-containing protein |
| 1(1) | WP_271839935.1 | WP_271839935.1 tetratricopeptide repeat protein |
| 1(2) | WP_133370061.1 | WP_133370061.1 WXG100 family type VII secretion target |
| 1(1) | WP_145153649.1 | WP_145153649.1 hypothetical protein |
| 1(1) | WP_268058024.1 | WP_268058024.1 pore-forming ESAT-6 family protein |
| 1(1) | WP_087995244.1 | WP_087995244.1 tetratricopeptide repeat protein |
| 1(1) | WP_085499087.1 | WP_085499087.1 DUF2974 domain-containing protein |
| 1(1) | WP_143881076.1 | WP_143881076.1 DNA/RNA non-specific endonuclease |
| 1(1) | WP_213143949.1 | WP_213143949.1 WXG100 family type VII secretion target |
| 1(1) | WP_257155372.1 | WP_257155372.1 hypothetical protein |
| 1(1) | WP_294953158.1 | WP_294953158.1 carboxypeptidase-like regulatory domain-containing protein |
| 1(2) | WP_145046449.1 | WP_145046449.1 hypothetical protein |
| 1(1) | WP_195949153.1 | WP_195949153.1 hypothetical protein |
| 1(2) | WP_001030887.1 | WP_001030887.1 SMI1/KNR4 family protein |
| 1(1) | WP_072727824.1 | WP_072727824.1 WXG100 family type VII secretion target |
| 1(3) | WP_029517061.1 | WP_029517061.1 hypothetical protein |
| 1(1) | WP_204412713.1 | WP_204412713.1 serine/threonine-protein kinase |
| 1(1) | WP_045518275.1 | WP_045518275.1 Ig-like domain-containing protein |
| 1(1) | WP_038286719.1 | WP_038286719.1 WXG100 family type VII secretion target |
| 1(1) | WP_179016429.1 | WP_179016429.1 hypothetical protein |
| 1(1) | WP_169505054.1 | WP_169505054.1 WXG100 family type VII secretion target |
| 1(2) | WP_104065530.1 | WP_104065530.1 hypothetical protein |
| 1(1) | WP_205001924.1 | WP_205001924.1 WXG100 family type VII secretion target |
| 1(1) | WP_166244902.1 | WP_166244902.1 hypothetical protein |
| 1(1) | WP_251548759.1 | WP_251548759.1 DUF2974 domain-containing protein |
| 1(1) | WP_289376844.1 | WP_289376844.1 RHS repeat-associated core domain-containing protein |
| 1(1) | WP_247083037.1 | WP_247083037.1 hypothetical protein |
| 1(1) | WP_251481000.1 | WP_251481000.1 pore-forming ESAT-6 family protein |
| 1(1) | WP_131231105.1 | WP_131231105.1 SseB family protein |
| 1(1) | WP_158329536.1 | WP_158329536.1 type VII secretion target |
| 1(1) | WP_251417661.1 | WP_251417661.1 DUF5050 domain-containing protein |
| 1(1) | WP_215032132.1 | WP_215032132.1 LysM peptidoglycan-binding domain-containing protein |
| 1(1) | WP_110931545.1 | WP_110931545.1 protein kinase family protein |
| 1(1) | WP_066020846.1 | WP_066020846.1 FHA domain-containing protein |
| 1(1) | WP_094092630.1 | WP_094092630.1 YhgE/Pip domain-containing protein |
| 1(1) | WP_309444467.1 | WP_309444467.1 hypothetical protein, partial |
| 1(4) | WP_031463346.1 | WP_031463346.1 WXG100 family type VII secretion target |
| 1(1) | WP_141528001.1 | WP_141528001.1 hypothetical protein |
| 1(1) | WP_195951609.1 | WP_195951609.1 pore-forming ESAT-6 family protein |
| 1(1) | WP_098562457.1 | WP_098562457.1 bacteriocin-processing peptidase family protein |
| 1(1) | WP_113052565.1 | WP_113052565.1 WXG100 family type VII secretion target |
| 1(1) | WP_063219634.1 | WP_063219634.1 SseB family protein |
| 1(4) | WP_021291851.1 | WP_021291851.1 EsaB/YukD family protein |
| 1(1) | WP_098854975.1 | WP_098854975.1 tetratricopeptide repeat protein |
| 1(1) | WP_226679100.1 | WP_226679100.1 FHA domain-containing protein |
| 1(1) | WP_211023032.1 | WP_211023032.1 hypothetical protein |
| 1(1) | WP_166742827.1 | WP_166742827.1 AAA family ATPase |
| 1(1) | WP_223592670.1 | WP_223592670.1 serine/threonine-protein kinase |
| 1(1) | WP_066091995.1 | WP_066091995.1 WXG100 family type VII secretion target |
| 1(1) | WP_232274491.1 | WP_232274491.1 WXG100 family type VII secretion target |
| 1(1) | WP_088032796.1 | WP_088032796.1 SMI1/KNR4 family protein |
| 1(1) | WP_308083995.1 | WP_308083995.1 serine/threonine-protein kinase |
| 1(2) | WP_189031028.1 | WP_189031028.1 WXG100 family type VII secretion target |
| 1(4) | WP_104065534.1 | WP_104065534.1 type VII secretion target |
| 1(1) | WP_068696906.1 | WP_068696906.1 copper amine oxidase N-terminal domain-containing protein |
| 1(1) | WP_195431288.1 | WP_195431288.1 tetratricopeptide repeat protein |
| 1(10) | WP_055277390.1 | WP_055277390.1 WXG100 family type VII secretion target |
| 1(1) | WP_280788636.1 | WP_280788636.1 HNH endonuclease |
| 1(1) | WP_145949001.1 | WP_145949001.1 hypothetical protein |
| 1(1) | WP_251241781.1 | WP_251241781.1 DUF2974 domain-containing protein, partial |
| 1(1) | WP_189018612.1 | WP_189018612.1 YHYH domain-containing protein |
| 1(1) | WP_090761833.1 | WP_090761833.1 serine/threonine-protein kinase |
| 1(1) | WP_308721989.1 | WP_308721989.1 hypothetical protein |
| 1(1) | WP_263053825.1 | WP_263053825.1 DNA/RNA non-specific endonuclease |
| 1(1) | WP_187292119.1 | WP_187292119.1 DUF2974 domain-containing protein |
| 1(1) | WP_232699381.1 | WP_232699381.1 WXG100 family type VII secretion target |
| 1(1) | WP_039306350.1 | WP_039306350.1 hypothetical protein |
| 1(1) | WP_280752667.1 | WP_280752667.1 WXG100 family type VII secretion target |
| 1(1) | WP_077893582.1 | WP_077893582.1 FHA domain-containing protein |
| 1(4) | WP_013252384.1 | WP_013252384.1 WXG100 family type VII secretion target |
| 1(1) | WP_185165633.1 | WP_185165633.1 tetratricopeptide repeat protein |
| 1(1) | WP_224839865.1 | WP_224839865.1 FHA domain-containing protein |
| 1(1) | WP_251375553.1 | WP_251375553.1 hypothetical protein |
| 1(1) | WP_061975581.1 | WP_061975581.1 hypothetical protein |
| 1(2) | WP_010963732.1 | WP_010963732.1 WXG100 family type VII secretion target |
| 1(1) | WP_098235617.1 | WP_098235617.1 bacteriocin-processing peptidase family protein |
| 1(1) | WP_170287045.1 | WP_170287045.1 stalk domain-containing protein |
| 1(1) | WP_066020834.1 | WP_066020834.1 tetratricopeptide repeat protein |
| 1(1) | WP_221825192.1 | WP_221825192.1 hypothetical protein |
| 1(1) | WP_209861641.1 | WP_209861641.1 serine/threonine-protein kinase |
| 1(1) | WP_126996325.1 | WP_126996325.1 hypothetical protein |
| 1(2) | WP_066419977.1 | WP_066419977.1 WXG100 family type VII secretion target |
| 1(1) | WP_262729433.1 | WP_262729433.1 hypothetical protein |
| 1(1) | WP_118991809.1 | WP_118991809.1 hypothetical protein |
| 1(1) | WP_257675916.1 | WP_257675916.1 hypothetical protein |

| | | |
|---|---|---|
| 1(1) | WP_142950820.1 | WP_142950820.1 type VII secretion protein EsaA |
| 1(1) | WP_232299892.1 | WP_232299892.1 DNA/RNA non-specific endonuclease |
| 1(1) | WP_002036279.1 | WP_002036279.1 hypothetical protein |
| 1(1) | WP_209861628.1 | WP_209861628.1 YhgE/Pip domain-containing protein |
| 1(2) | WP_076286996.1 | WP_076286996.1 hypothetical protein |
| 1(1) | WP_217035859.1 | WP_217035859.1 WXG100 family type VII secretion target |
| 1(1) | WP_240515074.1 | WP_240515074.1 polymorphic toxin type 15 domain-containing protein |
| 1(1) | WP_120112235.1 | WP_120112235.1 hypothetical protein |
| 1(1) | WP_155703922.1 | WP_155703922.1 copper amine oxidase N-terminal domain-containing protein |
| 1(1) | WP_137078359.1 | WP_137078359.1 S-layer homology domain-containing protein |
| 1(1) | WP_212949032.1 | WP_212949032.1 hypothetical protein |
| 1(1) | WP_169238728.1 | WP_169238728.1 FHA domain-containing protein |
| 1(1) | WP_284036492.1 | WP_284036492.1 Ig-like domain repeat protein |
| 1(1) | WP_251375550.1 | WP_251375550.1 hypothetical protein |
| 1(1) | WP_133384358.1 | WP_133384358.1 hypothetical protein |
| 1(1) | WP_280787393.1 | WP_280787393.1 DUF2974 domain-containing protein |
| 1(1) | WP_257675922.1 | WP_257675922.1 hypothetical protein |
| 1(1) | WP_191089523.1 | WP_191089523.1 WXG100 family type VII secretion target |
| 1(4) | WP_053356030.1 | WP_053356030.1 WXG100 family type VII secretion target |
| 1(1) | WP_243270457.1 | WP_243270457.1 DUF6382 domain-containing protein |
| 1(1) | WP_002148295.1 | WP_002148295.1 hypothetical protein |
| 1(1) | WP_090688968.1 | WP_090688968.1 WXG100 family type VII secretion target |
| 1(1) | WP_278046125.1 | WP_278046125.1 hypothetical protein |
| 1(1) | WP_230873710.1 | WP_230873710.1 serine/threonine-protein kinase |
| 1(3) | WP_013914191.1 | WP_013914191.1 WXG100 family type VII secretion target |
| 1(1) | WP_058271098.1 | WP_058271098.1 glycosyltransferase family 2 protein |
| 1(1) | WP_185165630.1 | WP_185165630.1 hypothetical protein |
| 1(1) | WP_213618816.1 | WP_213618816.1 hypothetical protein |
| 1(1) | WP_119794359.1 | WP_119794359.1 hypothetical protein |
| 1(1) | WP_010250258.1 | WP_010250258.1 WXG100 family type VII secretion target |
| 1(1) | WP_303942536.1 | WP_303942536.1 hypothetical protein |
| 1(1) | WP_283910914.1 | WP_283910914.1 AAA family ATPase |
| 1(2) | WP_270995165.1 | WP_270995165.1 hypothetical protein |
| 1(1) | WP_024632681.1 | WP_024632681.1 pre-toxin TG domain-containing protein |
| 1(1) | WP_190299137.1 | WP_190299137.1 hypothetical protein |
| 1(1) | WP_098579097.1 | WP_098579097.1 zinc-ribbon domain-containing protein |
| 1(1) | WP_077852415.1 | WP_077852415.1 hypothetical protein |
| 1(4) | WP_097952782.1 | WP_097952782.1 hypothetical protein |
| 1(1) | WP_214328824.1 | WP_214328824.1 WXG100 family type VII secretion target, partial |
| 1(1) | WP_284566229.1 | WP_284566229.1 HNH endonuclease |
| 1(1) | WP_120462951.1 | WP_120462951.1 DUF5050 domain-containing protein |
| 1(1) | WP_099354908.1 | WP_099354908.1 DUF2974 domain-containing protein |
| 1(1) | WP_275175502.1 | WP_275175502.1 hypothetical protein |
| 1(1) | WP_091155372.1 | WP_091155372.1 hypothetical protein |
| 1(1) | WP_280752668.1 | WP_280752668.1 DUF2974 domain-containing protein |
| 1(1) | WP_102710723.1 | WP_102710723.1 WXG100 family type VII secretion target |
| 1(10) | WP_053564628.1 | WP_053564628.1 SseB family protein |
| 1(2) | WP_051739195.1 | WP_051739195.1 ADP-ribosyltransferase |
| 1(1) | WP_152547782.1 | WP_152547782.1 cadherin-like beta sandwich domain-containing protein |
| 1(1) | WP_233457717.1 | WP_233457717.1 pre-toxin TG domain-containing protein |
| 1(1) | WP_213506885.1 | WP_213506885.1 copper amine oxidase N-terminal domain-containing protein |
| 1(2) | WP_076249775.1 | WP_076249775.1 WXG100 family type VII secretion target |
| 1(1) | WP_272035303.1 | WP_272035303.1 hypothetical protein |
| 1(1) | WP_166704992.1 | WP_166704992.1 hypothetical protein |
| 1(1) | WP_017149767.1 | WP_017149767.1 SseB family protein |
| 1(1) | WP_044037261.1 | WP_044037261.1 hypothetical protein |
| 1(1) | WP_309098609.1 | WP_309098609.1 hypothetical protein |
| 1(1) | WP_226086173.1 | WP_226086173.1 Mbeg1-like protein |
| 1(1) | WP_169028891.1 | WP_169028891.1 WXG100 family type VII secretion target |
| 1(1) | WP_289475128.1 | WP_289475128.1 hypothetical protein |
| 1(1) | WP_013490140.1 | WP_013490140.1 serine/threonine-protein kinase |
| 1(1) | WP_221875374.1 | WP_221875374.1 WXG100 family type VII secretion target |
| 1(1) | WP_307284688.1 | WP_307284688.1 WXG100 family type VII secretion target |
| 1(1) | WP_068696864.1 | WP_068696864.1 pre-toxin TG domain-containing protein |
| 1(1) | WP_081390252.1 | WP_081390252.1 copper amine oxidase N-terminal domain-containing protein |
| 1(2) | WP_213592859.1 | WP_213592859.1 hypothetical protein |
| 1(1) | WP_127536195.1 | WP_127536195.1 hypothetical protein |
| 1(1) | WP_208926607.1 | WP_208926607.1 hypothetical protein |
| 1(2) | WP_097952778.1 | WP_097952778.1 tetratricopeptide repeat protein |
| 1(3) | WP_142391903.1 | WP_142391903.1 hypothetical protein |
| 1(2) | WP_185397377.1 | WP_185397377.1 pre-toxin TG domain-containing protein |
| 1(3) | WP_098447519.1 | WP_098447519.1 SseB family protein |
| 1(1) | WP_098856768.1 | WP_098856768.1 hypothetical protein |
| 1(1) | WP_307601034.1 | WP_307601034.1 hypothetical protein |
| 1(2) | WP_281813098.1 | WP_281813098.1 WXG100 family type VII secretion target |
| 1(1) | WP_125902864.1 | WP_125902864.1 WXG100 family type VII secretion target |
| 1(1) | WP_306049012.1 | WP_306049012.1 FHA domain-containing protein |
| 1(1) | WP_144473247.1 | WP_144473247.1 hypothetical protein |
| 1(1) | WP_052947759.1 | WP_052947759.1 tetratricopeptide repeat protein |
| 1(1) | WP_000506914.1 | WP_000506914.1 tetratricopeptide repeat protein |
| 1(1) | WP_055109366.1 | WP_055109366.1 copper amine oxidase N-terminal domain-containing protein |
| 1(2) | WP_000404144.1 | WP_000404144.1 WXG100 family type VII secretion target |
| 1(1) | WP_149885342.1 | WP_149885342.1 tetratricopeptide repeat protein |
| 1(1) | WP_257443644.1 | WP_257443644.1 WXG100 family type VII secretion target |
| 1(1) | WP_119732863.1 | WP_119732863.1 WG repeat-containing protein |
| 1(1) | WP_281103266.1 | WP_281103266.1 DNA/RNA non-specific endonuclease |
| 1(1) | WP_195959145.1 | WP_195959145.1 hypothetical protein |
| 1(1) | WP_079506400.1 | WP_079506400.1 WXG100 family type VII secretion target |
| 1(1) | WP_100529434.1 | WP_100529434.1 hypothetical protein |
| 1(1) | WP_122971540.1 | WP_122971540.1 hypothetical protein |
| 1(1) | WP_273061465.1 | WP_273061465.1 Ig-like domain-containing protein |
| 1(2) | WP_028533181.1 | WP_028533181.1 hypothetical protein |
| 1(1) | WP_110899722.1 | WP_110899722.1 RHS repeat-associated core domain-containing protein |
| 1(1) | WP_236417220.1 | WP_236417220.1 hypothetical protein |
| 1(6) | WP_097821799.1 | WP_097821799.1 tetratricopeptide repeat protein |
| 1(2) | WP_250210512.1 | WP_250210512.1 hypothetical protein |
| 1(1) | WP_231094855.1 | WP_231094855.1 hypothetical protein |
| 1(2) | WP_194334461.1 | WP_194334461.1 type VII secretion protein EsaA |
| 1(1) | WP_290407047.1 | WP_290407047.1 DUF87 domain-containing protein |

| | | |
|---|---|---|
| 1(1) | WP_098329135.1 | WP_098329135.1 type VII secretion protein EsaA |
| 1(1) | WP_216975167.1 | WP_216975167.1 S-layer homology domain-containing protein |
| 1(1) | WP_055109367.1 | WP_055109367.1 hypothetical protein |
| 1(1) | WP_268238744.1 | WP_268238744.1 WXG100 family type VII secretion target |
| 1(1) | WP_165097630.1 | WP_165097630.1 copper amine oxidase N-terminal domain-containing protein |
| 1(1) | WP_076119209.1 | WP_076119209.1 DUF5050 domain-containing protein |
| 1(1) | WP_198310836.1 | WP_198310836.1 hypothetical protein |
| 1(1) | WP_307878869.1 | WP_307878869.1 tetratricopeptide repeat protein |
| 1(1) | WP_284036501.1 | WP_284036501.1 serine hydrolase |
| 1(1) | WP_307601032.1 | WP_307601032.1 hypothetical protein |
| 1(1) | WP_215153841.1 | WP_215153841.1 hypothetical protein |
| 1(1) | WP_042462801.1 | WP_042462801.1 WXG100 family type VII secretion target |
| 1(1) | WP_238655338.1 | WP_238655338.1 hypothetical protein |
| 1(1) | WP_046522790.1 | WP_046522790.1 serine/threonine-protein kinase |
| 1(1) | WP_145948998.1 | WP_145948998.1 WXG100 family type VII secretion target |
| 1(1) | WP_055109386.1 | WP_055109386.1 EsaB/YukD family protein |
| 1(3) | WP_097984057.1 | WP_097984057.1 tetratricopeptide repeat protein |
| 1(1) | WP_144476581.1 | WP_144476581.1 hypothetical protein |
| 1(1) | WP_015595777.1 | WP_015595777.1 glycosyltransferase family 2 protein |
| 1(1) | WP_242950781.1 | WP_242950781.1 hypothetical protein |
| 1(1) | WP_212695781.1 | WP_212695781.1 hypothetical protein |
| 1(1) | WP_147786235.1 | WP_147786235.1 tetratricopeptide repeat protein |
| 1(1) | WP_257126957.1 | WP_257126957.1 hypothetical protein |
| 1(2) | WP_109625241.1 | WP_109625241.1 pore-forming ESAT-6 family protein |
| 1(1) | WP_213506880.1 | WP_213506880.1 WXG100 family type VII secretion target |
| 1(1) | WP_010676854.1 | WP_010676854.1 hypothetical protein |
| 1(1) | WP_195325248.1 | WP_195325248.1 tetratricopeptide repeat protein |
| 1(1) | WP_212980299.1 | WP_212980299.1 serine hydrolase |
| 1(1) | WP_076157853.1 | WP_076157853.1 stalk domain-containing protein |
| 1(1) | WP_308615623.1 | WP_308615623.1 hypothetical protein |
| 1(1) | WP_233457712.1 | WP_233457712.1 hypothetical protein |
| 1(1) | WP_283746589.1 | WP_283746589.1 hypothetical protein, partial |
| 1(1) | WP_081097585.1 | WP_081097585.1 type VII secretion protein EsaA |
| 1(2) | WP_088550036.1 | WP_088550036.1 copper amine oxidase N-terminal domain-containing protein |
| 1(1) | WP_068940946.1 | WP_068940946.1 WXG100 family type VII secretion target |
| 1(1) | WP_290403369.1 | WP_290403369.1 hypothetical protein |
| 1(1) | WP_280752665.1 | WP_280752665.1 DUF5050 domain-containing protein |
| 1(1) | WP_226642880.1 | WP_226642880.1 FHA domain-containing protein |
| 1(1) | WP_223617610.1 | WP_223617610.1 hypothetical protein |
| 1(1) | WP_283910922.1 | WP_283910922.1 EsaB/YukD family protein |
| 1(1) | WP_204666336.1 | WP_204666336.1 hypothetical protein |
| 1(1) | WP_041618149.1 | WP_041618149.1 S-layer homology domain-containing protein |
| 1(1) | WP_086419835.1 | WP_086419835.1 WXG100 family type VII secretion target |
| 1(1) | WP_181603057.1 | WP_181603057.1 DUF6382 domain-containing protein |
| 1(3) | WP_213117480.1 | WP_213117480.1 WXG100 family type VII secretion target |
| 1(1) | WP_289398696.1 | WP_289398696.1 tetratricopeptide repeat protein |
| 1(1) | WP_002148322.1 | WP_002148322.1 hypothetical protein |
| 1(1) | WP_270731487.1 | WP_270731487.1 tetratricopeptide repeat protein |
| 1(1) | WP_157418137.1 | WP_157418137.1 hypothetical protein |
| 1(1) | WP_158287110.1 | WP_158287110.1 DUF2974 domain-containing protein |
| 1(1) | WP_166244926.1 | WP_166244926.1 S-layer homology domain-containing protein |
| 1(1) | WP_277166585.1 | WP_277166585.1 hypothetical protein |
| 1(1) | WP_257126959.1 | WP_257126959.1 hypothetical protein |
| 1(1) | WP_278773435.1 | WP_278773435.1 hypothetical protein |
| 1(1) | WP_229522859.1 | WP_229522859.1 hypothetical protein |
| 1(1) | WP_258113056.1 | WP_258113056.1 LXG domain-containing protein |
| 1(1) | WP_250244798.1 | WP_250244798.1 S-layer homology domain-containing protein |
| 1(1) | WP_195515241.1 | WP_195515241.1 pore-forming ESAT-6 family protein |
| 1(1) | WP_258433015.1 | WP_258433015.1 WG repeat-containing protein |
| 1(1) | WP_076210982.1 | WP_076210982.1 WXG100 family type VII secretion target |
| 1(1) | WP_026572734.1 | WP_026572734.1 hypothetical protein |
| 1(1) | WP_286124789.1 | WP_286124789.1 WXG100 family type VII secretion target |
| 1(1) | WP_028533177.1 | WP_028533177.1 membrane protein |
| 1(1) | WP_200633298.1 | WP_200633298.1 hypothetical protein |
| 1(1) | WP_152392572.1 | WP_152392572.1 hypothetical protein |
| 1(1) | WP_212969036.1 | WP_212969036.1 WXG100 family type VII secretion target |
| 1(1) | WP_088058518.1 | WP_088058518.1 bacteriocin-processing peptidase family protein |
| 1(1) | WP_242951634.1 | WP_242951634.1 serine/threonine-protein kinase |
| 1(1) | WP_026882390.1 | WP_026882390.1 WXG100 family type VII secretion target |
| 1(1) | WP_098303172.1 | WP_098303172.1 hypothetical protein |
| 1(1) | WP_079346825.1 | WP_079346825.1 hypothetical protein |
| 1(1) | WP_077852414.1 | WP_077852414.1 tetratricopeptide repeat protein |
| 1(1) | WP_143812903.1 | WP_143812903.1 WXG100 family type VII secretion target |
| 1(1) | WP_289392075.1 | WP_289392075.1 hypothetical protein |
| 1(1) | WP_143881071.1 | WP_143881071.1 hypothetical protein |
| 1(1) | WP_031415571.1 | WP_031415571.1 WXG100 family type VII secretion target |
| 1(1) | WP_074108296.1 | WP_074108296.1 hypothetical protein |
| 1(1) | WP_258296279.1 | WP_258296279.1 WXG100 family type VII secretion target |
| 1(1) | WP_283157903.1 | WP_283157903.1 AHH domain-containing protein, partial |
| 1(2) | WP_036702417.1 | WP_036702417.1 EsaB/YukD family protein |
| 1(4) | WP_041964574.1 | WP_041964574.1 WXG100 family type VII secretion target |
| 1(1) | WP_098655546.1 | WP_098655546.1 type VII secretion protein EsaA |
| 1(1) | WP_098264286.1 | WP_098264286.1 tetratricopeptide repeat protein |
| 1(1) | WP_227219205.1 | WP_227219205.1 Ig-like domain-containing protein |
| 1(1) | WP_277480698.1 | WP_277480698.1 WXG100 family type VII secretion target |
| 1(1) | WP_077833107.1 | WP_077833107.1 hypothetical protein |
| 1(1) | WP_138186699.1 | WP_138186699.1 hypothetical protein |
| 1(1) | WP_000034621.1 | WP_000034621.1 T7SS effector LXG polymorphic toxin |
| 1(1) | WP_058486875.1 | WP_058486875.1 stalk domain-containing protein |
| 1(1) | WP_212983255.1 | WP_212983255.1 WXG100 family type VII secretion target |
| 1(47) | WP_000918601.1 | WP_000918601.1 WXG100 family type VII secretion target |
| 1(4) | WP_000391137.1 | WP_000391137.1 EsaB/YukD family protein |
| 1(1) | WP_098882512.1 | WP_098882512.1 type VII secretion protein EsaA |
| 1(1) | WP_306077396.1 | WP_306077396.1 hypothetical protein |
| 1(1) | WP_139867381.1 | WP_139867381.1 tetratricopeptide repeat protein |
| 1(1) | WP_137698112.1 | WP_137698112.1 DUF5050 domain-containing protein |
| 1(8) | WP_006674860.1 | WP_006674860.1 WXG100 family type VII secretion target |
| 1(1) | WP_169238709.1 | WP_169238709.1 glycosyltransferase family 2 protein |
| 1(2) | WP_116353248.1 | WP_116353248.1 pore-forming ESAT-6 family protein |
| 1(2) | WP_251674645.1 | WP_251674645.1 S-layer homology domain-containing protein |
| 1(1) | WP_258418576.1 | WP_258418576.1 WG repeat-containing protein |
| 1(1) | WP_231417865.1 | WP_231417865.1 WXG100 family type VII secretion target |

| | | |
|---|---|---|
| 1(1) | WP_256704224.1 | WP_256704224.1 lipase family protein |
| 1(1) | WP_204517519.1 | WP_204517519.1 WG repeat-containing protein |
| 1(1) | WP_248085945.1 | WP_248085945.1 hypothetical protein |
| 1(1) | WP_208919038.1 | WP_208919038.1 quorum-sensing peptide PapR |
| 1(1) | WP_255443977.1 | WP_255443977.1 hypothetical protein, partial |
| 1(4) | WP_001007570.1 | WP_001007570.1 EsaB/YukD family protein |
| 1(1) | WP_068673693.1 | WP_068673693.1 WXG100 family type VII secretion target |
| 1(1) | WP_290407033.1 | WP_290407033.1 S-layer homology domain-containing protein |
| 1(1) | WP_307399101.1 | WP_307399101.1 copper amine oxidase N-terminal domain-containing protein |
| 1(1) | WP_077320200.1 | WP_077320200.1 type VII secretion protein EsaA |
| 1(1) | WP_099354913.1 | WP_099354913.1 DUF6382 domain-containing protein |
| 1(1) | WP_048558206.1 | WP_048558206.1 T7SS effector LXG polymorphic toxin |
| 1(1) | WP_111147800.1 | WP_111147800.1 DUF2974 domain-containing protein |
| 1(2) | WP_102372750.1 | WP_102372750.1 hypothetical protein |
| 1(2) | WP_208919036.1 | WP_208919036.1 WXG100 family type VII secretion target |
| 1(1) | WP_229190087.1 | WP_229190087.1 hypothetical protein |
| 1(1) | WP_043069422.1 | WP_043069422.1 hypothetical protein |
| 1(1) | WP_277548439.1 | WP_277548439.1 hypothetical protein |
| 1(1) | WP_277150347.1 | WP_277150347.1 WXG100 family type VII secretion target |
| 1(1) | WP_148450940.1 | WP_148450940.1 hypothetical protein |
| 1(1) | WP_242805838.1 | WP_242805838.1 bacteriocin-processing peptidase family protein |
| 1(1) | WP_169028881.1 | WP_169028881.1 hypothetical protein, partial |
| 1(1) | WP_055105787.1 | WP_055105787.1 WXG100 family type VII secretion target |
| 1(1) | WP_184182988.1 | WP_184182988.1 hypothetical protein |
| 1(1) | WP_222599448.1 | WP_222599448.1 hypothetical protein |
| 1(2) | WP_051494129.1 | WP_051494129.1 copper amine oxidase N-terminal domain-containing protein |
| 1(1) | WP_150364195.1 | WP_150364195.1 protein kinase family protein |
| 1(1) | WP_285848031.1 | WP_285848031.1 WXG100 family type VII secretion target |
| 1(1) | WP_137698102.1 | WP_137698102.1 phage tail tip lysozyme |
| 1(1) | WP_121610599.1 | WP_121610599.1 DUF6382 domain-containing protein |
| 1(1) | WP_307399125.1 | WP_307399125.1 protein kinase |
| 1(1) | WP_086405042.1 | WP_086405042.1 WXG100 family type VII secretion target |
| 1(2) | WP_017692347.1 | WP_017692347.1 WXG100 family type VII secretion target |
| 1(1) | WP_216975174.1 | WP_216975174.1 hypothetical protein |
| 1(1) | WP_212695791.1 | WP_212695791.1 hypothetical protein |
| 1(1) | WP_013490127.1 | WP_013490127.1 DUF2974 domain-containing protein |
| 1(1) | WP_195325254.1 | WP_195325254.1 FHA domain-containing protein |
| 1(1) | WP_218061148.1 | WP_218061148.1 hypothetical protein |
| 1(1) | WP_066091974.1 | WP_066091974.1 VWA domain-containing protein |
| 1(1) | WP_251583288.1 | WP_251583288.1 hypothetical protein |
| 1(1) | WP_039273530.1 | WP_039273530.1 copper amine oxidase N-terminal domain-containing protein |
| 1(1) | WP_234121608.1 | WP_234121608.1 FHA domain-containing protein |
| 1(1) | WP_087433606.1 | WP_087433606.1 WXG100 family type VII secretion target |
| 1(1) | WP_010676859.1 | WP_010676859.1 DUF6382 domain-containing protein |
| 1(1) | WP_204445213.1 | WP_204445213.1 DUF2974 domain-containing protein |
| 1(1) | WP_071640596.1 | WP_071640596.1 copper amine oxidase N-terminal domain-containing protein |
| 1(1) | WP_179034793.1 | WP_179034793.1 DUF2974 domain-containing protein |
| 1(1) | WP_127194956.1 | WP_127194956.1 hypothetical protein |
| 1(1) | WP_162848534.1 | WP_162848534.1 AAA family ATPase |
| 1(1) | WP_270711672.1 | WP_270711672.1 hypothetical protein |
| 1(2) | WP_040305090.1 | WP_040305090.1 hypothetical protein |
| 1(1) | WP_275175498.1 | WP_275175498.1 hypothetical protein |
| 1(1) | WP_119794364.1 | WP_119794364.1 WXG100 family type VII secretion target |
| 1(1) | WP_277156194.1 | WP_277156194.1 serine/threonine-protein kinase |
| 1(2) | WP_281813081.1 | WP_281813081.1 hypothetical protein |
| 1(1) | WP_145318704.1 | WP_145318704.1 hypothetical protein |
| 1(1) | WP_002031637.1 | WP_002031637.1 C39 family peptidase |
| 1(1) | WP_277175585.1 | WP_277175585.1 hypothetical protein |
| 1(1) | WP_127484329.1 | WP_127484329.1 cadherin-like beta sandwich domain-containing protein |
| 1(1) | WP_094092623.1 | WP_094092623.1 hypothetical protein |
| 1(1) | WP_068536370.1 | WP_068536370.1 EsaB/YukD family protein |
| 1(1) | WP_235286196.1 | WP_235286196.1 hypothetical protein |
| 1(1) | WP_051983561.1 | WP_051983561.1 serine/threonine-protein kinase |
| 1(1) | WP_234121613.1 | WP_234121613.1 Ig-like domain-containing protein |
| 1(2) | WP_057897624.1 | WP_057897624.1 WXG100 family type VII secretion target |
| 1(1) | WP_209365681.1 | WP_209365681.1 type VII secretion protein EsaA |
| 1(2) | WP_219872468.1 | WP_219872468.1 WXG100 family type VII secretion target |
| 1(1) | WP_139356158.1 | WP_139356158.1 glycoside hydrolase family 3 C-terminal domain-containing protein |
| 1(1) | WP_090761827.1 | WP_090761827.1 WXG100 family type VII secretion target |
| 1(1) | WP_016362248.1 | WP_016362248.1 S-layer homology domain-containing protein |
| 1(2) | WP_038569542.1 | WP_038569542.1 AAA family ATPase |
| 1(1) | WP_251542605.1 | WP_251542605.1 hypothetical protein |
| 1(1) | WP_216513490.1 | WP_216513490.1 WXG100 family type VII secretion target |
| 1(1) | WP_160789864.1 | WP_160789864.1 hypothetical protein |
| 1(1) | WP_181731447.1 | WP_181731447.1 WXG100 family type VII secretion target |
| 1(1) | WP_061829404.1 | WP_061829404.1 hypothetical protein |
| 1(2) | WP_195431318.1 | WP_195431318.1 WXG100 family type VII secretion target |
| 1(1) | WP_068673695.1 | WP_068673695.1 hypothetical protein |
| 1(3) | WP_203573360.1 | WP_203573360.1 hypothetical protein |
| 1(1) | WP_248251186.1 | WP_248251186.1 tetratricopeptide repeat protein |
| 1(2) | WP_213592885.1 | WP_213592885.1 hypothetical protein |
| 1(2) | WP_126996315.1 | WP_126996315.1 hypothetical protein |
| 1(1) | WP_242400016.1 | WP_242400016.1 bacteriocin-processing peptidase family protein |
| 1(1) | WP_213143951.1 | WP_213143951.1 FHA domain-containing protein |
| 1(5) | WP_006211205.1 | WP_006211205.1 hypothetical protein |
| 1(1) | WP_025685033.1 | WP_025685033.1 WXG100 family type VII secretion target |
| 1(1) | WP_220684640.1 | WP_220684640.1 serine/threonine-protein kinase |
| 1(2) | WP_002171736.1 | WP_002171736.1 hypothetical protein |
| 1(1) | WP_240459622.1 | WP_240459622.1 hypothetical protein, partial |
| 1(1) | WP_144473246.1 | WP_144473246.1 tetratricopeptide repeat protein |
| 1(1) | WP_256424316.1 | WP_256424316.1 WXG100 family type VII secretion target |
| 1(1) | WP_263324557.1 | WP_263324557.1 hypothetical protein |
| 1(1) | WP_104148791.1 | WP_104148791.1 WG repeat-containing protein |
| 1(1) | WP_251497193.1 | WP_251497193.1 hypothetical protein |
| 1(1) | WP_308082231.1 | WP_308082231.1 endonuclease |
| 1(1) | WP_277157950.1 | WP_277157950.1 hypothetical protein |
| 1(1) | WP_034526609.1 | WP_034526609.1 WXG100 family type VII secretion target |
| 1(1) | WP_090908276.1 | WP_090908276.1 hypothetical protein |
| 1(1) | WP_268583855.1 | WP_268583855.1 hypothetical protein |

| | | | |
|---|---|---|---|
| 1(3) | WP_095266197.1 | WP_095266197.1 | WXG100 family type VII secretion target |
| 1(1) | WP_145156804.1 | WP_145156804.1 | hypothetical protein |
| 1(1) | WP_269778087.1 | WP_269778087.1 | DNA/RNA non-specific endonuclease |
| 1(1) | WP_212695783.1 | WP_212695783.1 | WXG100 family type VII secretion target |
| 1(1) | WP_142316415.1 | WP_142316415.1 | hypothetical protein |
| 1(3) | WP_179603477.1 | WP_179603477.1 | WXG100 family type VII secretion target |
| 1(1) | WP_149871695.1 | WP_149871695.1 | glycoside hydrolase family 3 protein |
| 1(1) | WP_098841814.1 | WP_098841814.1 | tetratricopeptide repeat protein |
| 1(1) | WP_307399109.1 | WP_307399109.1 | FHA domain-containing protein |
| 1(1) | WP_150457776.1 | WP_150457776.1 | AAA family ATPase |
| 1(2) | WP_194335282.1 | WP_194335282.1 | hypothetical protein |
| 1(1) | WP_142306903.1 | WP_142306903.1 | WXG100 family type VII secretion target |
| 1(1) | WP_090239244.1 | WP_090239244.1 | hypothetical protein |
| 1(1) | WP_116556144.1 | WP_116556144.1 | hypothetical protein |
| 1(1) | WP_026572746.1 | WP_026572746.1 | Ig-like domain repeat protein |
| 1(1) | WP_087995250.1 | WP_087995250.1 | bacteriocin-processing peptidase family protein |
| 1(1) | WP_244723818.1 | WP_244723818.1 | hypothetical protein |
| 1(1) | WP_251557377.1 | WP_251557377.1 | DUF2974 domain-containing protein |
| 1(1) | WP_213532657.1 | WP_213532657.1 | WXG100 family type VII secretion target |
| 1(1) | WP_234415830.1 | WP_234415830.1 | SseB family protein |
| 1(1) | WP_268583856.1 | WP_268583856.1 | tetratricopeptide repeat protein |
| 1(1) | WP_165097625.1 | WP_165097625.1 | hypothetical protein |
| 1(1) | WP_283927236.1 | WP_283927236.1 | DUF2974 domain-containing protein |
| 1(1) | WP_000446751.1 | WP_000446751.1 | DNA/RNA non-specific endonuclease |
| 1(1) | WP_198310825.1 | WP_198310825.1 | Ig-like domain-containing protein |
| 1(1) | WP_181747865.1 | WP_181747865.1 | WG repeat-containing protein |
| 1(1) | WP_222591880.1 | WP_222591880.1 | hypothetical protein |
| 1(1) | WP_055109370.1 | WP_055109370.1 | hypothetical protein |
| 1(6) | WP_041964573.1 | WP_041964573.1 | pore-forming ESAT-6 family protein |
| 1(1) | WP_242951966.1 | WP_242951966.1 | hypothetical protein |
| 1(1) | WP_169238719.1 | WP_169238719.1 | serine/threonine-protein kinase |
| 1(2) | WP_031415573.1 | WP_031415573.1 | hypothetical protein |
| 1(1) | WP_132283091.1 | WP_132283091.1 | WXG100 family type VII secretion target |
| 1(1) | WP_149238907.1 | WP_149238907.1 | WXG100 family type VII secretion target |
| 1(62) | WP_003745045.1 | WP_003745045.1 | EsaB/YukD family protein |
| 1(1) | WP_169238718.1 | WP_169238718.1 | serine/threonine-protein kinase |
| 1(1) | WP_179603472.1 | WP_179603472.1 | Ig-like domain-containing protein |
| 1(2) | WP_185469029.1 | WP_185469029.1 | DNA/RNA non-specific endonuclease |
| 1(1) | WP_090674368.1 | WP_090674368.1 | pore-forming ESAT-6 family protein |
| 1(1) | WP_098264345.1 | WP_098264345.1 | CdiA family toxin C-terminal domain-containing protein |
| 1(2) | WP_053783884.1 | WP_053783884.1 | hypothetical protein |
| 1(1) | WP_098516211.1 | WP_098516211.1 | T7SS effector LXG polymorphic toxin |
| 1(1) | WP_088119567.1 | WP_088119567.1 | bacteriocin-processing peptidase family protein |
| 1(1) | WP_138755918.1 | WP_138755918.1 | hypothetical protein |
| 1(2) | WP_215570368.1 | WP_215570368.1 | protein kinase |
| 1(2) | WP_076264229.1 | WP_076264229.1 | hypothetical protein |
| 1(2) | WP_017692349.1 | WP_017692349.1 | EsaB/YukD family protein |
| 1(1) | WP_289373801.1 | WP_289373801.1 | hypothetical protein |
| 1(1) | WP_169028882.1 | WP_169028882.1 | hypothetical protein |
| 1(1) | WP_181603055.1 | WP_181603055.1 | Ig-like domain repeat protein |
| 1(1) | WP_013490137.1 | WP_013490137.1 | hypothetical protein |
| 1(1) | WP_197937392.1 | WP_197937392.1 | EsaB/YukD family protein |
| 1(8) | WP_000136046.1 | WP_000136046.1 | WXG100 family type VII secretion target |
| 1(1) | WP_277175591.1 | WP_277175591.1 | hypothetical protein |
| 1(1) | WP_307394633.1 | WP_307394633.1 | hypothetical protein |
| 1(1) | WP_159146508.1 | WP_159146508.1 | hypothetical protein |
| 1(1) | WP_236804424.1 | WP_236804424.1 | alanine and proline-rich secreted protein Apa |
| 1(1) | WP_166704995.1 | WP_166704995.1 | hypothetical protein |
| 1(4) | WP_076210973.1 | WP_076210973.1 | EsaB/YukD family protein |
| 1(1) | WP_283892177.1 | WP_283892177.1 | hypothetical protein |
| 1(1) | WP_270472005.1 | WP_270472005.1 | hypothetical protein |
| 1(1) | WP_307199940.1 | WP_307199940.1 | hypothetical protein |
| 1(1) | WP_251638765.1 | WP_251638765.1 | WXG100 family type VII secretion target |
| 1(1) | WP_283952946.1 | WP_283952946.1 | WXG100 family type VII secretion target |
| 1(1) | WP_095691017.1 | WP_095691017.1 | bacteriocin-processing peptidase family protein |
| 1(21) | WP_053564631.1 | WP_053564631.1 | WXG100 family type VII secretion target |
| 1(1) | WP_307399123.1 | WP_307399123.1 | FHA domain-containing protein |
| 1(1) | WP_287600001.1 | WP_287600001.1 | GBS Bsp-like repeat-containing protein, partial |
| 1(1) | WP_287600034.1 | WP_287600034.1 | hypothetical protein, partial |
| 1(1) | WP_098355767.1 | WP_098355767.1 | bacteriocin-processing peptidase family protein |
| 1(1) | WP_306077388.1 | WP_306077388.1 | DUF2974 domain-containing protein |
| 1(1) | WP_113050537.1 | WP_113050537.1 | hypothetical protein |
| 1(2) | WP_283655683.1 | WP_283655683.1 | S-layer homology domain-containing protein |
| 1(1) | WP_239209289.1 | WP_239209289.1 | WG repeat-containing protein |
| 1(9) | WP_021253116.1 | WP_021253116.1 | WXG100 family type VII secretion target |
| 1(1) | WP_101356554.1 | WP_101356554.1 | type VII secretion protein EsaA |
| 1(1) | WP_153064091.1 | WP_153064091.1 | hypothetical protein |
| 1(1) | WP_158318212.1 | WP_158318212.1 | tetratricopeptide repeat protein |
| 1(1) | WP_211022790.1 | WP_211022790.1 | EsaB/YukD family protein |
| 1(1) | WP_186580073.1 | WP_186580073.1 | hypothetical protein |
| 1(2) | WP_064473934.1 | WP_064473934.1 | tetratricopeptide repeat protein |
| 1(1) | WP_197241529.1 | WP_197241529.1 | WXG100 family type VII secretion target, partial |
| 1(1) | WP_071719980.1 | WP_071719980.1 | hypothetical protein |
| 1(1) | WP_238333010.1 | WP_238333010.1 | hypothetical protein |
| 1(1) | WP_062326148.1 | WP_062326148.1 | hypothetical protein |
| 1(1) | WP_023988267.1 | WP_023988267.1 | membrane protein |
| 1(1) | WP_163255249.1 | WP_163255249.1 | bacteriocin-processing peptidase family protein |
| 1(1) | WP_091260865.1 | WP_091260865.1 | tetratricopeptide repeat protein |
| 1(1) | WP_098747412.1 | WP_098747412.1 | hypothetical protein |
| 1(1) | WP_111147797.1 | WP_111147797.1 | hypothetical protein |
| 1(2) | WP_138322087.1 | WP_138322087.1 | bacteriocin-processing peptidase family protein |
| 1(1) | WP_166704981.1 | WP_166704981.1 | bacteriocin-processing peptidase family protein |
| 1(1) | WP_211745829.1 | WP_211745829.1 | WXG100 family type VII secretion target |
| 1(1) | WP_217035853.1 | WP_217035853.1 | WXG100 family type VII secretion target |
| 1(1) | WP_122971545.1 | WP_122971545.1 | Ig-like domain repeat protein |
| 1(1) | WP_179034774.1 | WP_179034774.1 | HNH endonuclease |
| 1(1) | WP_251480993.1 | WP_251480993.1 | hypothetical protein |
| 1(1) | WP_254759306.1 | WP_254759306.1 | Ig-like domain-containing protein |
| 1(1) | WP_188944162.1 | WP_188944162.1 | TNT domain-containing protein |
| 1(1) | WP_088032786.1 | WP_088032786.1 | tetratricopeptide repeat protein |
| 1(1) | WP_148450959.1 | WP_148450959.1 | Ig-like domain-containing protein |
| 1(1) | WP_283952942.1 | WP_283952942.1 | serine/threonine-protein kinase |
| 1(1) | WP_108534956.1 | WP_108534956.1 | Mbeg1-like protein |
| 1(1) | WP_102371668.1 | WP_102371668.1 | hypothetical protein |

| | | |
|---|---|---|
| 1(1) | WP_144506830.1 | WP_144506830.1 tetratricopeptide repeat protein |
| 1(1) | WP_044649165.1 | WP_044649165.1 WXG100 family type VII secretion target |
| 1(1) | WP_098991578.1 | WP_098991578.1 bacteriocin-processing peptidase family protein |
| 1(1) | WP_211023027.1 | WP_211023027.1 EsaB/YukD family protein |
| 1(1) | WP_141536470.1 | WP_141536470.1 hypothetical protein |
| 1(2) | WP_043067147.1 | WP_043067147.1 WXG100 family type VII secretion target |
| 1(1) | WP_313959731.1 | WP_313959731.1 protein kinase |
| 1(1) | WP_244988296.1 | WP_244988296.1 RHS repeat-associated core domain-containing protein |
| 1(1) | WP_091155382.1 | WP_091155382.1 hypothetical protein |
| 1(1) | WP_251638755.1 | WP_251638755.1 hypothetical protein |
| 1(1) | WP_150457760.1 | WP_150457760.1 Mbeg1-like protein |
| 1(1) | WP_303942524.1 | WP_303942524.1 serine/threonine-protein kinase |
| 1(2) | WP_000506918.1 | WP_000506918.1 tetratricopeptide repeat protein |
| 1(1) | WP_017692353.1 | WP_017692353.1 WXG100 family type VII secretion target |
| 1(1) | WP_027088430.1 | WP_027088430.1 hypothetical protein |
| 1(2) | WP_185474091.1 | WP_185474091.1 hypothetical protein |
| 1(1) | WP_289607498.1 | WP_289607498.1 FHA domain-containing protein |
| 1(1) | WP_185183366.1 | WP_185183366.1 tetratricopeptide repeat protein |
| 1(2) | WP_071736268.1 | WP_071736268.1 bacteriocin-processing peptidase family protein |
| 1(8) | WP_005546515.1 | WP_005546515.1 WXG100 family type VII secretion target |
| 1(1) | WP_083784186.1 | WP_083784186.1 serine/threonine-protein kinase |
| 1(1) | WP_198310829.1 | WP_198310829.1 DUF6382 domain-containing protein |
| 1(1) | WP_102371666.1 | WP_102371666.1 FHA domain-containing protein |
| 1(2) | WP_241747708.1 | WP_241747708.1 hypothetical protein |
| 1(1) | WP_211022780.1 | WP_211022780.1 copper amine oxidase N-terminal domain-containing protein |
| 1(1) | WP_268613679.1 | WP_268613679.1 pore-forming ESAT-6 family protein |
| 1(30) | WP_006211202.1 | WP_006211202.1 WXG100 family type VII secretion target |
| 1(1) | WP_138755910.1 | WP_138755910.1 hypothetical protein |
| 1(1) | WP_289398691.1 | WP_289398691.1 tetratricopeptide repeat protein |
| 1(1) | WP_113058864.1 | WP_113058864.1 hypothetical protein |
| 1(1) | WP_204666345.1 | WP_204666345.1 FHA domain-containing protein |
| 1(1) | WP_122971533.1 | WP_122971533.1 WXG100 family type VII secretion target |
| 1(2) | WP_082061566.1 | WP_082061566.1 YHYH domain-containing protein |
| 1(1) | WP_149238900.1 | WP_149238900.1 bacteriocin-processing peptidase family protein |
| 1(1) | WP_271840004.1 | WP_271840004.1 DNA/RNA non-specific endonuclease |
| 1(1) | WP_097853506.1 | WP_097853506.1 bacteriocin-processing peptidase family protein |
| 1(1) | WP_095691007.1 | WP_095691007.1 WXG100 family type VII secretion target |
| 1(1) | WP_282446277.1 | WP_282446277.1 RHS repeat-associated core domain-containing protein |
| 1(1) | WP_290401993.1 | WP_290401993.1 hypothetical protein |
| 1(1) | WP_189018605.1 | WP_189018605.1 hypothetical protein |
| 1(1) | WP_106765956.1 | WP_106765956.1 serine/threonine protein kinase |
| 1(1) | WP_055105797.1 | WP_055105797.1 hypothetical protein |
| 1(1) | WP_098562452.1 | WP_098562452.1 WXG100 family type VII secretion target |
| 1(1) | WP_097899491.1 | WP_097899491.1 WXG100 family type VII secretion target |
| 1(1) | WP_098948844.1 | WP_098948844.1 bacteriocin-processing peptidase family protein |
| 1(1) | WP_195959147.1 | WP_195959147.1 pore-forming ESAT-6 family protein |
| 1(1) | WP_000034620.1 | WP_000034620.1 T7SS effector LXG polymorphic toxin |
| 1(1) | WP_098952625.1 | WP_098952625.1 SseB family protein |
| 1(4) | WP_053325275.1 | WP_053325275.1 S-layer homology domain-containing protein |
| 1(1) | WP_277157948.1 | WP_277157948.1 WXG100 family type VII secretion target |
| 1(1) | WP_307601030.1 | WP_307601030.1 pore-forming ESAT-6 family protein |

| | | |
|---|---|---|
| 1(1) | WP_304216861.1 | WP_304216861.1 hypothetical protein |
| 1(1) | WP_270721356.1 | WP_270721356.1 WXG100 family type VII secretion target |
| 1(1) | WP_239079768.1 | WP_239079768.1 hypothetical protein |
| 1(1) | WP_272801674.1 | WP_272801674.1 zinc-ribbon domain-containing protein |
| 1(1) | WP_229139920.1 | WP_229139920.1 DUF2974 domain-containing protein |
| 1(1) | WP_055105788.1 | WP_055105788.1 WXG100 family type VII secretion target |
| 1(1) | WP_101356555.1 | WP_101356555.1 WXG100 family type VII secretion target |
| 1(1) | WP_036707112.1 | WP_036707112.1 WXG100 family type VII secretion target |
| 1(1) | WP_083677952.1 | WP_083677952.1 hypothetical protein |
| 1(1) | WP_270731495.1 | WP_270731495.1 WXG100 family type VII secretion target |
| 1(2) | WP_017692352.1 | WP_017692352.1 WXG100 family type VII secretion target |
| 1(1) | WP_108464076.1 | WP_108464076.1 hypothetical protein |
| 1(1) | WP_165097623.1 | WP_165097623.1 WXG100 family type VII secretion target |
| 1(1) | WP_208926612.1 | WP_208926612.1 DUF2974 domain-containing protein |
| 1(1) | WP_251548751.1 | WP_251548751.1 FHA domain-containing protein |
| 1(1) | WP_071724708.1 | WP_071724708.1 bacteriocin-processing peptidase family protein |
| 1(1) | WP_155703932.1 | WP_155703932.1 hypothetical protein |
| 1(1) | WP_036702413.1 | WP_036702413.1 WXG100 family type VII secretion target |
| 1(5) | WP_223849888.1 | WP_223849888.1 CdiA family toxin C-terminal domain-containing protein |
| 1(1) | WP_051624141.1 | WP_051624141.1 tetratricopeptide repeat protein |
| 1(1) | WP_098841819.1 | WP_098841819.1 bacteriocin-processing peptidase family protein |
| 1(1) | WP_306007797.1 | WP_306007797.1 tetratricopeptide repeat protein |
| 1(1) | WP_000883854.1 | WP_000883854.1 bacteriocin-processing peptidase family protein |
| 1(1) | WP_234121605.1 | WP_234121605.1 WXG100 family type VII secretion target |
| 1(1) | WP_068793930.1 | WP_068793930.1 hypothetical protein |
| 1(1) | WP_204517510.1 | WP_204517510.1 type VII secretion target |
| 1(1) | WP_006674859.1 | WP_006674859.1 hypothetical protein |
| 1(3) | WP_071719982.1 | WP_071719982.1 WXG100 family type VII secretion target |
| 1(1) | WP_126996289.1 | WP_126996289.1 hypothetical protein |
| 1(1) | WP_024632676.1 | WP_024632676.1 lipase family protein |
| 1(1) | WP_289376839.1 | WP_289376839.1 RHS repeat-associated core domain-containing protein |
| 1(1) | WP_305022939.1 | WP_305022939.1 S-layer homology domain-containing protein |
| 1(1) | WP_171003601.1 | WP_171003601.1 hypothetical protein |
| 1(1) | WP_214705999.1 | WP_214705999.1 LysM peptidoglycan-binding domain-containing protein |
| 1(1) | WP_254349526.1 | WP_254349526.1 WXG100 family type VII secretion target |
| 1(1) | WP_236334324.1 | WP_236334324.1 DUF2974 domain-containing protein |
| 1(1) | WP_000378750.1 | WP_000378750.1 DNA/RNA non-specific endonuclease |
| 1(4) | WP_062833878.1 | WP_062833878.1 hypothetical protein |
| 1(1) | WP_036689412.1 | WP_036689412.1 WXG100 family type VII secretion target |
| 1(1) | WP_021725634.1 | WP_021725634.1 WXG100 family type VII secretion target |
| 1(1) | WP_260859262.1 | WP_260859262.1 type VII secretion protein EsaA, partial |
| 1(1) | WP_286017557.1 | WP_286017557.1 hypothetical protein |
| 1(1) | WP_301804665.1 | WP_301804665.1 hypothetical protein |
| 1(1) | WP_248139696.1 | WP_248139696.1 hypothetical protein |
| 1(1) | WP_277156187.1 | WP_277156187.1 WXG100 family type VII secretion target |
| 1(1) | WP_050423257.1 | WP_050423257.1 GH-E family nuclease |
| 1(1) | WP_005546516.1 | WP_005546516.1 hypothetical protein |
| 1(1) | WP_251542603.1 | WP_251542603.1 LysM peptidoglycan-binding domain-containing protein |
| 1(1) | WP_189018603.1 | WP_189018603.1 WXG100 family type VII secretion target |
| 1(1) | WP_226642870.1 | WP_226642870.1 hypothetical protein |
| 1(1) | WP_033521269.1 | WP_033521269.1 WXG100 family type VII secretion target |
| 1(1) | WP_098556208.1 | WP_098556208.1 tetratricopeptide repeat protein |

| | | |
|---|---|---|
| 1(1) | WP_060630640.1 | WP_060630640.1 bacteriocin-processing peptidase family protein |
| 1(1) | WP_068536338.1 | WP_068536338.1 copper amine oxidase N-terminal domain-containing protein |
| 1(2) | WP_281813094.1 | WP_281813094.1 WXG100 family type VII secretion target |
| 1(1) | WP_052812188.1 | WP_052812188.1 tetratricopeptide repeat protein |
| 1(1) | WP_251674653.1 | WP_251674653.1 hypothetical protein |
| 1(1) | WP_143316570.1 | WP_143316570.1 leucine-rich repeat domain-containing protein, partial |
| 1(2) | WP_046325924.1 | WP_046325924.1 type VII secretion protein EsaA |
| 1(1) | WP_242622143.1 | WP_242622143.1 T7SS effector LXG polymorphic toxin |
| 1(1) | WP_102371670.1 | WP_102371670.1 serine/threonine-protein kinase |
| 1(2) | WP_076281076.1 | WP_076281076.1 DUF5050 domain-containing protein |
| 1(1) | WP_076119221.1 | WP_076119221.1 copper amine oxidase N-terminal domain-containing protein |
| 1(1) | WP_280752669.1 | WP_280752669.1 DUF2974 domain-containing protein |
| 1(4) | WP_023521818.1 | WP_023521818.1 hypothetical protein |
| 1(1) | WP_316324167.1 | WP_316324167.1 hypothetical protein |
| 1(1) | WP_013914189.1 | WP_013914189.1 EsaB/YukD family protein |
| 1(1) | WP_157763986.1 | WP_157763986.1 hypothetical protein |
| 1(2) | WP_000391140.1 | WP_000391140.1 hypothetical protein |
| 1(1) | WP_290409175.1 | WP_290409175.1 hypothetical protein |
| 1(1) | WP_127194960.1 | WP_127194960.1 hypothetical protein |
| 1(1) | WP_077893456.1 | WP_077893456.1 WXG100 family type VII secretion target |
| 1(1) | WP_183578458.1 | WP_183578458.1 hypothetical protein |
| 1(1) | WP_108465334.1 | WP_108465334.1 S-layer homology domain-containing protein |
| 1(1) | WP_015595756.1 | WP_015595756.1 pore-forming ESAT-6 family protein |
| 1(1) | WP_137062787.1 | WP_137062787.1 S-layer homology domain-containing protein |
| 1(1) | WP_180358375.1 | WP_180358375.1 hypothetical protein |
| 1(1) | WP_098866667.1 | WP_098866667.1 hypothetical protein |
| 1(1) | WP_232699387.1 | WP_232699387.1 hypothetical protein |
| 1(1) | WP_213592848.1 | WP_213592848.1 VWA domain-containing protein |
| 1(1) | WP_150364200.1 | WP_150364200.1 hypothetical protein |
| 1(1) | WP_307537580.1 | WP_307537580.1 WXG100 family type VII secretion target |
| 1(2) | WP_240326484.1 | WP_240326484.1 hypothetical protein |
| 1(1) | WP_270574772.1 | WP_270574772.1 WXG100 family type VII secretion target |
| 1(1) | WP_144660363.1 | WP_144660363.1 hypothetical protein |
| 1(1) | WP_036716969.1 | WP_036716969.1 hypothetical protein |
| 1(2) | WP_179283640.1 | WP_179283640.1 EsaB/YukD family protein |
| 1(1) | WP_098230737.1 | WP_098230737.1 hypothetical protein |
| 1(1) | WP_090908279.1 | WP_090908279.1 hypothetical protein |
| 1(1) | WP_208203483.1 | WP_208203483.1 hypothetical protein |
| 1(1) | WP_253440789.1 | WP_253440789.1 hypothetical protein |
| 1(1) | WP_028407767.1 | WP_028407767.1 zinc-ribbon domain-containing protein |
| 1(1) | WP_270472017.1 | WP_270472017.1 WXG100 family type VII secretion target |
| 1(1) | WP_138226024.1 | WP_138226024.1 hypothetical protein |
| 1(1) | WP_216083867.1 | WP_216083867.1 WXG100 family type VII secretion target |
| 1(1) | WP_098248507.1 | WP_098248507.1 hypothetical protein |
| 1(1) | WP_148450957.1 | WP_148450957.1 hypothetical protein |
| 1(2) | WP_128101488.1 | WP_128101488.1 WXG100 family type VII secretion target |
| 1(4) | WP_077833103.1 | WP_077833103.1 tetratricopeptide repeat protein |
| 1(1) | WP_213506874.1 | WP_213506874.1 EsaB/YukD family protein |
| 1(1) | WP_244169193.1 | WP_244169193.1 S-layer homology domain-containing protein |
| 1(3) | WP_076135805.1 | WP_076135805.1 hypothetical protein |
| 1(1) | WP_010676860.1 | WP_010676860.1 pore-forming ESAT-6 family protein |
| 1(1) | WP_048826577.1 | WP_048826577.1 DUF2974 domain-containing protein |
| 1(1) | WP_051354801.1 | WP_051354801.1 tetratricopeptide repeat protein |
| 1(1) | WP_198056462.1 | WP_198056462.1 DUF2974 domain-containing protein |
| 1(2) | WP_138226027.1 | WP_138226027.1 hypothetical protein |
| 1(1) | WP_242400028.1 | WP_242400028.1 WXG100 family type VII secretion target |
| 1(1) | WP_212693234.1 | WP_212693234.1 WXG100 family type VII secretion target |
| 1(1) | WP_280752670.1 | WP_280752670.1 copper amine oxidase N-terminal domain-containing protein |
| 1(1) | WP_277544341.1 | WP_277544341.1 type VII secretion protein EsaA |
| 1(4) | WP_077832292.1 | WP_077832292.1 WXG100 family type VII secretion target |
| 1(1) | WP_162174145.1 | WP_162174145.1 WXG100 family type VII secretion target |
| 1(1) | WP_260065260.1 | WP_260065260.1 hypothetical protein |
| 1(1) | WP_290401998.1 | WP_290401998.1 WXG100 family type VII secretion target |
| 1(1) | WP_216390030.1 | WP_216390030.1 WXG100 family type VII secretion target |
| 1(1) | WP_213506867.1 | WP_213506867.1 hypothetical protein |
| 1(1) | WP_164965500.1 | WP_164965500.1 S-layer homology domain-containing protein |
| 1(1) | WP_155703928.1 | WP_155703928.1 cysteine peptidase family C39 domain-containing protein |
| 1(1) | WP_289355901.1 | WP_289355901.1 WXG100 family type VII secretion target |
| 1(1) | WP_184182994.1 | WP_184182994.1 RHS repeat-associated core domain-containing protein |
| 1(1) | WP_232620907.1 | WP_232620907.1 DNA/RNA non-specific endonuclease |
| 1(1) | WP_112331588.1 | WP_112331588.1 hypothetical protein |
| 1(2) | WP_063219636.1 | WP_063219636.1 protein kinase |
| 1(1) | WP_015595755.1 | WP_015595755.1 DUF6382 domain-containing protein |
| 1(1) | WP_226086168.1 | WP_226086168.1 Ig-like domain-containing protein |
| 1(1) | WP_133378773.1 | WP_133378773.1 hypothetical protein |
| 1(1) | WP_097905180.1 | WP_097905180.1 hypothetical protein |
| 1(2) | WP_195431327.1 | WP_195431327.1 hypothetical protein |
| 1(1) | WP_013490125.1 | WP_013490125.1 WXG100 family type VII secretion target |
| 1(1) | WP_144027204.1 | WP_144027204.1 hypothetical protein |
| 1(2) | WP_185468945.1 | WP_185468945.1 type VII secretion protein EsaA |
| 1(1) | WP_120462961.1 | WP_120462961.1 hypothetical protein |
| 1(1) | WP_064477393.1 | WP_064477393.1 HNH endonuclease |
| 1(1) | WP_307738719.1 | WP_307738719.1 protein kinase |
| 1(53) | WP_017689071.1 | WP_017689071.1 WXG100 family type VII secretion target |
| 1(2) | WP_068782411.1 | WP_068782411.1 WXG100 family type VII secretion target |
| 1(3) | WP_098266641.1 | WP_098266641.1 bacteriocin-processing peptidase family protein |
| 1(1) | WP_152966007.1 | WP_152966007.1 hypothetical protein |
| 1(1) | WP_210087111.1 | WP_210087111.1 WXG100 family type VII secretion target |
| 1(1) | WP_163247639.1 | WP_163247639.1 bacteriocin-processing peptidase family protein |
| 1(1) | WP_110899360.1 | WP_110899360.1 S-layer homology domain-containing protein |
| 1(1) | WP_071724727.1 | WP_071724727.1 tetratricopeptide repeat protein |
| 1(1) | WP_097821798.1 | WP_097821798.1 hypothetical protein |
| 1(1) | WP_090674372.1 | WP_090674372.1 DUF2974 domain-containing protein |
| 1(1) | WP_055105791.1 | WP_055105791.1 hypothetical protein |
| 1(1) | WP_125084770.1 | WP_125084770.1 VWA domain-containing protein |
| 1(2) | WP_111268939.1 | WP_111268939.1 WXG100 family type VII secretion target |
| 1(1) | WP_197937383.1 | WP_197937383.1 AAA family ATPase |
| 1(1) | WP_098088249.1 | WP_098088249.1 EsaB/YukD family protein |
| 1(1) | WP_235957152.1 | WP_235957152.1 EsaB/YukD family protein |
| 1(1) | WP_289483786.1 | WP_289483786.1 tetratricopeptide repeat protein |
| 1(1) | WP_132131924.1 | WP_132131924.1 T7SS effector LXG polymorphic toxin |

| | | |
|---|---|---|
| 1(1) | WP_106496024.1 | WP_106496024.1 T7SS effector LXG polymorphic toxin |
| 1(1) | WP_199613899.1 | WP_199613899.1 hypothetical protein |
| 1(1) | WP_236334326.1 | WP_236334326.1 WXG100 family type VII secretion target |
| 1(1) | WP_097889499.1 | WP_097889499.1 tetratricopeptide repeat protein |
| 1(1) | WP_162178970.1 | WP_162178970.1 FHA domain-containing protein |
| 1(2) | WP_136604816.1 | WP_136604816.1 WXG100 family type VII secretion target |
| 1(1) | WP_195542465.1 | WP_195542465.1 hypothetical protein |
| 1(1) | WP_229522856.1 | WP_229522856.1 WXG100 family type VII secretion target |
| 1(1) | WP_309098612.1 | WP_309098612.1 DUF2974 domain-containing protein |
| 1(1) | WP_154985829.1 | WP_154985829.1 WXG100 family type VII secretion target |
| 1(1) | WP_251466790.1 | WP_251466790.1 WXG100 family type VII secretion target |
| 1(1) | WP_307413599.1 | WP_307413599.1 hypothetical protein |
| 1(2) | WP_016091014.1 | WP_016091014.1 T7SS effector LXG polymorphic toxin |
| 1(1) | WP_090887144.1 | WP_090887144.1 S-layer homology domain-containing protein |
| 1(1) | WP_188535092.1 | WP_188535092.1 hypothetical protein |
| 1(1) | WP_257443640.1 | WP_257443640.1 zincin-like metallopeptidase toxin domain-containing protein |
| 1(1) | WP_262729420.1 | WP_262729420.1 bacteriocin-processing peptidase family protein |
| 1(1) | WP_195974877.1 | WP_195974877.1 serine/threonine-protein kinase |
| 1(1) | WP_021253115.1 | WP_021253115.1 hypothetical protein |
| 1(1) | WP_277746357.1 | WP_277746357.1 hypothetical protein |
| 1(1) | WP_167527257.1 | WP_167527257.1 S-layer homology domain-containing protein |
| 1(2) | WP_076312504.1 | WP_076312504.1 WXG100 family type VII secretion target |
| 1(1) | WP_170220305.1 | WP_170220305.1 WXG100 family type VII secretion target |
| 1(1) | WP_016821019.1 | WP_016821019.1 S-layer homology domain-containing protein |
| 1(1) | WP_097877417.1 | WP_097877417.1 hypothetical protein |
| 1(1) | WP_053361017.1 | WP_053361017.1 serine/threonine-protein kinase |
| 1(1) | WP_076303952.1 | WP_076303952.1 hypothetical protein |
| 1(1) | WP_036670436.1 | WP_036670436.1 membrane protein |
| 1(1) | WP_277142094.1 | WP_277142094.1 hypothetical protein |
| 1(1) | WP_179160626.1 | WP_179160626.1 hypothetical protein |
| 1(1) | WP_191798527.1 | WP_191798527.1 WXG100 family type VII secretion target |
| 1(1) | WP_302153068.1 | WP_302153068.1 DUF2974 domain-containing protein |
| 1(1) | WP_242400025.1 | WP_242400025.1 tetratricopeptide repeat protein |
| 1(1) | WP_291682711.1 | WP_291682711.1 tetratricopeptide repeat protein, partial |
| 1(2) | WP_068782409.1 | WP_068782409.1 EsaB/YukD family protein |
| 1(1) | WP_286017556.1 | WP_286017556.1 AHH domain-containing protein |
| 1(1) | WP_149871682.1 | WP_149871682.1 Mbeg1-like protein |
| 1(1) | WP_208019353.1 | WP_208019353.1 SseB family protein |
| 1(1) | WP_212980184.1 | WP_212980184.1 hypothetical protein |
| 1(1) | WP_213506876.1 | WP_213506876.1 DUF5050 domain-containing protein |
| 1(1) | WP_103655360.1 | WP_103655360.1 HNH endonuclease |
| 1(1) | WP_179603486.1 | WP_179603486.1 hypothetical protein |
| 1(1) | WP_052427421.1 | WP_052427421.1 pore-forming ESAT-6 family protein |
| 1(1) | WP_025679807.1 | WP_025679807.1 membrane protein |
| 1(3) | WP_071736274.1 | WP_071736274.1 hypothetical protein |
| 1(1) | WP_285079440.1 | WP_285079440.1 hypothetical protein |
| 1(1) | WP_132008166.1 | WP_132008166.1 DUF6382 domain-containing protein |
| 1(1) | WP_112331581.1 | WP_112331581.1 WXG100 family type VII secretion target |
| 1(1) | WP_120112251.1 | WP_120112251.1 hypothetical protein |
| 1(1) | WP_270784525.1 | WP_270784525.1 tetratricopeptide repeat protein |
| 1(1) | WP_277157949.1 | WP_277157949.1 Ig-like domain repeat protein |
| 1(1) | WP_268058017.1 | WP_268058017.1 immunoglobulin domain-containing protein |
| 1(1) | WP_042125624.1 | WP_042125624.1 DUF5050 domain-containing protein |
| 1(1) | WP_090239224.1 | WP_090239224.1 hypothetical protein |
| 1(2) | WP_128101485.1 | WP_128101485.1 hypothetical protein |
| 1(1) | WP_160037488.1 | WP_160037488.1 WXG100 family type VII secretion target |
| 1(1) | WP_149871676.1 | WP_149871676.1 serine/threonine-protein kinase |
| 1(1) | WP_025719058.1 | WP_025719058.1 membrane protein |
| 1(1) | WP_286124774.1 | WP_286124774.1 tetratricopeptide repeat protein |
| 1(2) | WP_138226023.1 | WP_138226023.1 WXG100 family type VII secretion target |
| 1(1) | WP_019912163.1 | WP_019912163.1 hypothetical protein |
| 1(1) | WP_021959910.1 | WP_021959910.1 hypothetical protein |
| 1(1) | WP_238655348.1 | WP_238655348.1 DUF5050 domain-containing protein |
| 1(2) | WP_098508789.1 | WP_098508789.1 EsaB/YukD family protein |
| 1(4) | WP_012260838.1 | WP_012260838.1 SseB family protein |
| 1(1) | WP_140421209.1 | WP_140421209.1 hypothetical protein |
| 1(2) | WP_141526281.1 | WP_141526281.1 deaminase domain-containing protein |
| 1(1) | WP_235801707.1 | WP_235801707.1 type VII secretion protein EsaA |
| 1(1) | WP_110934084.1 | WP_110934084.1 hypothetical protein |
| 1(1) | WP_263703628.1 | WP_263703628.1 WXG100 family type VII secretion target |
| 1(1) | WP_102264544.1 | WP_102264544.1 serine/threonine-protein kinase |
| 1(1) | WP_286017550.1 | WP_286017550.1 hypothetical protein |
| 1(8) | WP_076167874.1 | WP_076167874.1 hypothetical protein |
| 1(1) | WP_182301837.1 | WP_182301837.1 WXG100 family type VII secretion target |
| 1(2) | WP_029517064.1 | WP_029517064.1 WXG100 family type VII secretion target |
| 1(1) | WP_277504687.1 | WP_277504687.1 tetratricopeptide repeat protein |
| 1(1) | WP_197241530.1 | WP_197241530.1 hypothetical protein |
| 1(3) | WP_077832283.1 | WP_077832283.1 EsaB/YukD family protein |
| 1(1) | WP_143881066.1 | WP_143881066.1 bacteriocin-processing peptidase family protein |
| 1(1) | WP_304502594.1 | WP_304502594.1 WG repeat-containing protein |
| 1(1) | WP_005604627.1 | WP_005604627.1 hypothetical protein |
| 1(1) | WP_094870536.1 | WP_094870536.1 EsaB/YukD family protein |
| 1(1) | WP_234405012.1 | WP_234405012.1 serine/threonine-protein kinase |
| 1(4) | WP_075702369.1 | WP_075702369.1 DUF6531 domain-containing protein |
| 1(1) | WP_193774468.1 | WP_193774468.1 hypothetical protein |
| 1(1) | WP_087956293.1 | WP_087956293.1 EsaB/YukD family protein |
| 1(1) | WP_166244076.1 | WP_166244076.1 hypothetical protein |
| 1(1) | WP_063208781.1 | WP_063208781.1 hypothetical protein |
| 1(1) | WP_210364448.1 | WP_210364448.1 hypothetical protein |
| 1(1) | WP_108465348.1 | WP_108465348.1 hypothetical protein |
| 1(1) | WP_270696281.1 | WP_270696281.1 tetratricopeptide repeat protein |
| 1(1) | WP_136379700.1 | WP_136379700.1 DUF2974 domain-containing protein |
| 1(1) | WP_308615624.1 | WP_308615624.1 serine/threonine-protein kinase |
| 1(1) | WP_021624825.1 | WP_021624825.1 hypothetical protein |
| 1(1) | WP_234447084.1 | WP_234447084.1 DNA/RNA non-specific endonuclease |
| 1(7) | WP_019392116.1 | WP_019392116.1 WXG100 family type VII secretion target |
| 1(1) | WP_098395975.1 | WP_098395975.1 WXG100 family type VII secretion target |
| 1(1) | WP_068673714.1 | WP_068673714.1 WXG100 family type VII secretion target |
| 1(1) | WP_293202261.1 | WP_293202261.1 WXG100 family type VII secretion target |
| 1(1) | WP_114298765.1 | WP_114298765.1 AHH domain-containing protein |
| 1(1) | WP_254759302.1 | WP_254759302.1 FHA domain-containing protein |
| 1(1) | WP_262800522.1 | WP_262800522.1 hypothetical protein |
| 1(1) | WP_077619643.1 | WP_077619643.1 DUF6382 domain-containing protein |

| | | |
|---|---|---|
| 1(1) | WP_166244078.1 | WP_166244078.1 hypothetical protein |
| 1(1) | WP_270684464.1 | WP_270684464.1 hypothetical protein |
| 1(1) | WP_090735442.1 | WP_090735442.1 S-layer homology domain-containing protein |
| 1(1) | WP_272640516.1 | WP_272640516.1 hypothetical protein |
| 1(1) | WP_028542688.1 | WP_028542688.1 membrane protein |
| 1(1) | WP_254325792.1 | WP_254325792.1 hypothetical protein |
| 1(1) | WP_216513491.1 | WP_216513491.1 WXG100 family type VII secretion target |
| 1(1) | WP_127570430.1 | WP_127570430.1 hypothetical protein |
| 1(1) | WP_099354898.1 | WP_099354898.1 hypothetical protein |
| 1(2) | WP_194346730.1 | WP_194346730.1 DUF6531 domain-containing protein |
| 1(1) | WP_289129507.1 | WP_289129507.1 hypothetical protein |
| 1(1) | WP_094870535.1 | WP_094870535.1 DUF5050 domain-containing protein |
| 1(1) | WP_098400686.1 | WP_098400686.1 bacteriocin-processing peptidase family protein |
| 1(1) | WP_309098614.1 | WP_309098614.1 DUF6382 domain-containing protein |
| 1(1) | WP_284036494.1 | WP_284036494.1 serine/threonine-protein kinase |
| 1(1) | WP_038245460.1 | WP_038245460.1 polysaccharide deacetylase family protein |
| 1(1) | WP_007148001.1 | WP_007148001.1 serine/threonine-protein kinase |
| 1(1) | WP_098451709.1 | WP_098451709.1 bacteriocin-processing peptidase family protein |
| 1(3) | WP_042401386.1 | WP_042401386.1 glycosyltransferase family 2 protein |
| 1(1) | WP_182301829.1 | WP_182301829.1 serine/threonine-protein kinase |
| 1(1) | WP_199695253.1 | WP_199695253.1 hypothetical protein |
| 1(5) | WP_060630646.1 | WP_060630646.1 hypothetical protein |
| 1(1) | WP_221875379.1 | WP_221875379.1 hypothetical protein |
| 1(1) | WP_307601028.1 | WP_307601028.1 WXG100 family type VII secretion target |
| 1(2) | WP_098248513.1 | WP_098248513.1 glycoside hydrolase family 3 N-terminal domain-containing protein |
| 1(1) | WP_251417677.1 | WP_251417677.1 hypothetical protein |
| 1(1) | WP_232243180.1 | WP_232243180.1 hypothetical protein |
| 1(1) | WP_235308161.1 | WP_235308161.1 hypothetical protein |
| 1(1) | WP_191272852.1 | WP_191272852.1 protein kinase |
| 1(1) | WP_119847553.1 | WP_119847553.1 serine/threonine protein kinase |
| 1(1) | WP_141097595.1 | WP_141097595.1 hypothetical protein, partial |
| 1(1) | WP_003340541.1 | WP_003340541.1 WG repeat-containing protein |
| 1(1) | WP_033521260.1 | WP_033521260.1 hypothetical protein, partial |
| 1(1) | WP_242261455.1 | WP_242261455.1 WXG100 family type VII secretion target |
| 1(5) | WP_076101575.1 | WP_076101575.1 copper amine oxidase N-terminal domain-containing protein |
| 1(1) | WP_248063439.1 | WP_248063439.1 WXG100 family type VII secretion target |
| 1(1) | WP_111147807.1 | WP_111147807.1 hypothetical protein |
| 1(4) | WP_228464159.1 | WP_228464159.1 hypothetical protein |
| 1(1) | WP_064473933.1 | WP_064473933.1 hypothetical protein |
| 1(1) | WP_205001938.1 | WP_205001938.1 hypothetical protein |
| 1(1) | WP_143315250.1 | WP_143315250.1 WXG100 family type VII secretion target |
| 1(1) | WP_028401644.1 | WP_028401644.1 WXG100 family type VII secretion target |
| 1(1) | WP_098866666.1 | WP_098866666.1 bacteriocin-processing peptidase family protein |
| 1(1) | WP_238655353.1 | WP_238655353.1 copper amine oxidase N-terminal domain-containing protein |
| 1(1) | WP_195325258.1 | WP_195325258.1 pore-forming ESAT-6 family protein |
| 1(1) | WP_251241783.1 | WP_251241783.1 WXG100 family type VII secretion target |
| 1(1) | WP_205665441.1 | WP_205665441.1 beta-N-acetylhexosaminidase |
| 1(1) | WP_192800072.1 | WP_192800072.1 hypothetical protein |
| 1(1) | WP_251466792.1 | WP_251466792.1 hypothetical protein |
| 1(1) | WP_172461027.1 | WP_172461027.1 tetratricopeptide repeat protein |
| 1(1) | WP_042203642.1 | WP_042203642.1 hypothetical protein |
| 1(1) | WP_197262004.1 | WP_197262004.1 WG repeat-containing protein |
| 1(1) | WP_188535050.1 | WP_188535050.1 AAA family ATPase |
| 1(1) | WP_215032117.1 | WP_215032117.1 DUF6382 domain-containing protein |
| 1(1) | WP_185966791.1 | WP_185966791.1 leucine-rich repeat domain-containing protein |
| 1(3) | WP_051989214.1 | WP_051989214.1 WXG100 family type VII secretion target |
| 1(1) | WP_000340946.1 | WP_000340946.1 hypothetical protein |
| 1(1) | WP_141528994.1 | WP_141528994.1 DNA/RNA non-specific endonuclease |
| 1(1) | WP_098150447.1 | WP_098150447.1 bacteriocin-processing peptidase family protein |
| 1(1) | WP_000506926.1 | WP_000506926.1 tetratricopeptide repeat protein |
| 1(4) | WP_007128631.1 | WP_007128631.1 hypothetical protein |
| 1(1) | WP_277175588.1 | WP_277175588.1 WXG100 family type VII secretion target |
| 1(1) | WP_049869825.1 | WP_049869825.1 HNH endonuclease |
| 1(1) | WP_058486872.1 | WP_058486872.1 WXG100 family type VII secretion target |
| 1(1) | WP_256716440.1 | WP_256716440.1 HNH endonuclease |
| 1(2) | WP_213592861.1 | WP_213592861.1 WXG100 family type VII secretion target |
| 1(1) | WP_155609751.1 | WP_155609751.1 hypothetical protein |
| 1(1) | WP_145329904.1 | WP_145329904.1 hypothetical protein |
| 1(1) | WP_264453872.1 | WP_264453872.1 hypothetical protein |
| 1(1) | WP_103047834.1 | WP_103047834.1 S-layer homology domain-containing protein |
| 1(1) | WP_289398697.1 | WP_289398697.1 hypothetical protein |
| 1(1) | WP_236334320.1 | WP_236334320.1 DUF2974 domain-containing protein |
| 1(1) | WP_084411334.1 | WP_084411334.1 RHS repeat-associated core domain-containing protein |
| 1(1) | WP_258418575.1 | WP_258418575.1 WXG100 family type VII secretion target |
| 1(1) | WP_007083293.1 | WP_007083293.1 pore-forming ESAT-6 family protein |
| 1(1) | WP_190299136.1 | WP_190299136.1 WXG100 family type VII secretion target |
| 1(1) | WP_229753216.1 | WP_229753216.1 nucleotidyltransferase domain-containing protein |
| 1(1) | WP_070805897.1 | WP_070805897.1 bacteriocin-processing peptidase family protein |
| 1(1) | WP_204445230.1 | WP_204445230.1 serine/threonine-protein kinase |
| 1(1) | WP_279323592.1 | WP_279323592.1 DUF6382 domain-containing protein |
| 1(2) | WP_185327600.1 | WP_185327600.1 hypothetical protein |
| 1(2) | WP_076210980.1 | WP_076210980.1 WXG100 family type VII secretion target |
| 1(1) | WP_238655355.1 | WP_238655355.1 hypothetical protein |
| 1(1) | WP_287550708.1 | WP_287550708.1 hypothetical protein, partial |
| 1(1) | WP_158318217.1 | WP_158318217.1 SseB family protein, partial |
| 1(1) | WP_051483716.1 | WP_051483716.1 serine/threonine-protein kinase |
| 1(1) | WP_242251779.1 | WP_242251779.1 tetratricopeptide repeat protein |
| 1(1) | WP_307388959.1 | WP_307388959.1 serine/threonine-protein kinase |
| 1(1) | WP_132283093.1 | WP_132283093.1 hypothetical protein |
| 1(1) | WP_139867379.1 | WP_139867379.1 hypothetical protein |
| 1(1) | WP_222599447.1 | WP_222599447.1 hypothetical protein |
| 1(1) | WP_013370792.1 | WP_013370792.1 copper amine oxidase N-terminal domain-containing protein |
| 1(1) | WP_015595753.1 | WP_015595753.1 protein kinase |
| 1(1) | WP_097911852.1 | WP_097911852.1 bacteriocin-processing peptidase family protein |
| 1(1) | WP_204445237.1 | WP_204445237.1 VWA domain-containing protein |
| 1(1) | WP_120046145.1 | WP_120046145.1 hypothetical protein |
| 1(5) | WP_064473937.1 | WP_064473937.1 hypothetical protein |
| 1(1) | WP_111147798.1 | WP_111147798.1 WXG100 family type VII secretion target |
| 1(1) | WP_099361334.1 | WP_099361334.1 hypothetical protein |
| 1(2) | WP_236804428.1 | WP_236804428.1 CdiA family toxin C-terminal domain-containing protein |

| | | | |
|---|---|---|---|
| 1(1) | WP_064472912.1 | WP_064472912.1 | bacteriocin-processing peptidase family protein |
| 1(1) | WP_000506929.1 | WP_000506929.1 | tetratricopeptide repeat protein |
| 1(1) | WP_098976051.1 | WP_098976051.1 | tetratricopeptide repeat protein |
| 1(1) | WP_077852740.1 | WP_077852740.1 | WXG100 family type VII secretion target |
| 1(1) | WP_248251184.1 | WP_248251184.1 | EsaB/YukD family protein |
| 1(1) | WP_204412709.1 | WP_204412709.1 | Ig-like domain-containing protein |
| 1(1) | WP_305755033.1 | WP_305755033.1 | WXG100 family type VII secretion target |
| 1(1) | WP_242264957.1 | WP_242264957.1 | WXG100 family type VII secretion target |
| 1(1) | WP_307192901.1 | WP_307192901.1 | hypothetical protein |
| 1(1) | WP_149238906.1 | WP_149238906.1 | tetratricopeptide repeat protein |
| 1(1) | WP_214706009.1 | WP_214706009.1 | serine/threonine-protein kinase |
| 1(1) | WP_074565304.1 | WP_074565304.1 | tetratricopeptide repeat protein |
| 1(1) | WP_251038449.1 | WP_251038449.1 | hypothetical protein |
| 1(1) | WP_265408844.1 | WP_265408844.1 | tetratricopeptide repeat protein |
| 1(1) | WP_229136360.1 | WP_229136360.1 | WXG100 family type VII secretion target |
| 1(2) | WP_200458887.1 | WP_200458887.1 | WXG100 family type VII secretion target, partial |
| 1(1) | WP_306802042.1 | WP_306802042.1 | CdiA family toxin C-terminal domain-containing protein |
| 1(1) | WP_172463486.1 | WP_172463486.1 | tetratricopeptide repeat protein |
| 1(1) | WP_267414293.1 | WP_267414293.1 | WXG100 family type VII secretion target |
| 1(1) | WP_209941813.1 | WP_209941813.1 | WXG100 family type VII secretion target |
| 1(1) | WP_254366019.1 | WP_254366019.1 | S-layer homology domain-containing protein, partial |
| 1(1) | WP_136789996.1 | WP_136789996.1 | hypothetical protein |
| 1(1) | WP_182301845.1 | WP_182301845.1 | S-layer homology domain-containing protein |
| 1(1) | WP_250244820.1 | WP_250244820.1 | hypothetical protein |
| 1(1) | WP_229522869.1 | WP_229522869.1 | S-layer homology domain-containing protein |
| 1(1) | WP_273061485.1 | WP_273061485.1 | WXG100 family type VII secretion target |
| 1(1) | WP_005604632.1 | WP_005604632.1 | WXG100 family type VII secretion target |
| 1(3) | WP_191798528.1 | WP_191798528.1 | WXG100 family type VII secretion target |
| 1(1) | WP_143316575.1 | WP_143316575.1 | Ig-like domain-containing protein |
| 1(1) | WP_090688963.1 | WP_090688963.1 | SseB family protein |
| 1(4) | WP_229521099.1 | WP_229521099.1 | hypothetical protein |
| 1(1) | WP_098088246.1 | WP_098088246.1 | tetratricopeptide repeat protein |
| 1(1) | WP_215570370.1 | WP_215570370.1 | hypothetical protein |
| 1(2) | WP_038584741.1 | WP_038584741.1 | WXG100 family type VII secretion target |
| 1(1) | WP_148450943.1 | WP_148450943.1 | copper amine oxidase N-terminal domain-containing protein |
| 1(1) | WP_068536382.1 | WP_068536382.1 | RHS repeat-associated core domain-containing protein |
| 1(1) | WP_127543698.1 | WP_127543698.1 | hypothetical protein |
| 1(1) | WP_021291844.1 | WP_021291844.1 | HNH endonuclease |
| 1(1) | WP_253299365.1 | WP_253299365.1 | RHS repeat-associated core domain-containing protein |
| 1(1) | WP_019120053.1 | WP_019120053.1 | SseB family protein |
| 1(1) | WP_063479992.1 | WP_063479992.1 | serine/threonine protein kinase |
| 1(1) | WP_243495556.1 | WP_243495556.1 | zinc-ribbon domain-containing protein |
| 1(1) | WP_226679106.1 | WP_226679106.1 | hypothetical protein |
| 1(1) | WP_200510207.1 | WP_200510207.1 | hypothetical protein |
| 1(1) | WP_257675920.1 | WP_257675920.1 | WXG100 family type VII secretion target |
| 1(1) | WP_181603059.1 | WP_181603059.1 | EsaB/YukD family protein |
| 1(1) | WP_195431333.1 | WP_195431333.1 | hypothetical protein, partial |
| 1(1) | WP_193653018.1 | WP_193653018.1 | WXG100 family type VII secretion target |
| 1(1) | WP_215598788.1 | WP_215598788.1 | WXG100 family type VII secretion target |
| 1(1) | WP_068497049.1 | WP_068497049.1 | WXG100 family type VII secretion target |
| 1(1) | WP_308082248.1 | WP_308082248.1 | WXG100 family type VII secretion target |

| | | | |
|---|---|---|---|
| 1(1) | WP_189018601.1 | WP_189018601.1 | stalk domain-containing protein |
| 1(1) | WP_052948651.1 | WP_052948651.1 | FHA domain-containing protein |
| 1(1) | WP_148450941.1 | WP_148450941.1 | copper amine oxidase N-terminal domain-containing protein |
| 1(1) | WP_278278680.1 | WP_278278680.1 | copper amine oxidase N-terminal domain-containing protein |
| 1(1) | WP_165175575.1 | WP_165175575.1 | hypothetical protein |
| 1(1) | WP_098952611.1 | WP_098952611.1 | bacteriocin-processing peptidase family protein |
| 1(1) | WP_163255255.1 | WP_163255255.1 | tetratricopeptide repeat protein |
| 1(2) | WP_242663055.1 | WP_242663055.1 | serine/threonine-protein kinase |
| 1(2) | WP_185397392.1 | WP_185397392.1 | glycohydrolase toxin TNT-related protein |
| 1(1) | WP_098204524.1 | WP_098204524.1 | SseB family protein |
| 1(1) | WP_216830404.1 | WP_216830404.1 | Ig-like domain-containing protein |
| 1(1) | WP_141551181.1 | WP_141551181.1 | hypothetical protein |
| 1(1) | WP_176581008.1 | WP_176581008.1 | bacteriocin-processing peptidase family protein |
| 1(3) | WP_134910005.1 | WP_134910005.1 | hypothetical protein |
| 1(1) | WP_257215859.1 | WP_257215859.1 | serine/threonine-protein kinase |
| 1(1) | WP_212693232.1 | WP_212693232.1 | hypothetical protein |
| 1(1) | WP_238187890.1 | WP_238187890.1 | serine/threonine-protein kinase |
| 1(1) | WP_211023029.1 | WP_211023029.1 | WXG100 family type VII secretion target |
| 1(1) | WP_236334329.1 | WP_236334329.1 | WXG100 family type VII secretion target |
| 1(1) | WP_286182604.1 | WP_286182604.1 | LysM peptidoglycan-binding domain-containing protein |
| 1(16) | WP_018670524.1 | WP_018670524.1 | WXG100 family type VII secretion target |
| 1(1) | WP_307394635.1 | WP_307394635.1 | hypothetical protein |
| 1(1) | WP_013658695.1 | WP_013658695.1 | WXG100 family type VII secretion target |
| 1(1) | WP_078185402.1 | WP_078185402.1 | WXG100 family type VII secretion target |
| 1(1) | WP_034526625.1 | WP_034526625.1 | DUF2974 domain-containing protein |
| 1(1) | WP_087442415.1 | WP_087442415.1 | hypothetical protein |
| 1(1) | WP_295628326.1 | WP_295628326.1 | FHA domain-containing protein |
| 1(1) | WP_098952623.1 | WP_098952623.1 | tetratricopeptide repeat protein |
| 1(1) | WP_074108295.1 | WP_074108295.1 | hypothetical protein |
| 1(1) | WP_063223280.1 | WP_063223280.1 | hypothetical protein |
| 1(1) | WP_204517518.1 | WP_204517518.1 | WXG100 family type VII secretion target |
| 1(1) | WP_162944260.1 | WP_162944260.1 | WXG100 family type VII secretion target |
| 1(1) | WP_231094848.1 | WP_231094848.1 | DUF2974 domain-containing protein |
| 1(1) | WP_226887710.1 | WP_226887710.1 | copper amine oxidase N-terminal domain-containing protein |
| 1(6) | WP_029438092.1 | WP_029438092.1 | SseB family protein |
| 1(1) | WP_142293491.1 | WP_142293491.1 | hypothetical protein |
| 1(1) | WP_270065066.1 | WP_270065066.1 | S8 family peptidase |
| 1(1) | WP_275540581.1 | WP_275540581.1 | hypothetical protein |
| 1(1) | WP_283927234.1 | WP_283927234.1 | DUF5050 domain-containing protein |
| 1(1) | WP_176229017.1 | WP_176229017.1 | DUF5050 domain-containing protein |
| 1(1) | WP_097905178.1 | WP_097905178.1 | tetratricopeptide repeat protein |
| 1(2) | WP_014518838.1 | WP_014518838.1 | hypothetical protein |
| 1(2) | WP_078384707.1 | WP_078384707.1 | T7SS effector LXG polymorphic toxin |
| 1(1) | WP_005604634.1 | WP_005604634.1 | WXG100 family type VII secretion target |
| 1(1) | WP_086419829.1 | WP_086419829.1 | bacteriocin-processing peptidase family protein |
| 1(1) | WP_194321178.1 | WP_194321178.1 | pre-toxin TG domain-containing protein |
| 1(1) | WP_212983257.1 | WP_212983257.1 | hypothetical protein |
| 1(1) | WP_290425832.1 | WP_290425832.1 | type VII secretion protein EsaA, partial |
| 1(1) | WP_191798529.1 | WP_191798529.1 | hypothetical protein |

1(1) WP_013309991.1 WP_013309991.1 membrane protein
1(1) WP_155609752.1 WP_155609752.1 WXG100 family type VII secretion target
1(1) WP_142330021.1 WP_142330021.1 hypothetical protein
1(1) WP_068536344.1 WP_068536344.1 hypothetical protein
1(2) WP_200458888.1 WP_200458888.1 SseB family protein
1(1) WP_280138894.1 WP_280138894.1 hypothetical protein
1(1) WP_098319061.1 WP_098319061.1 tetratricopeptide repeat protein
1(1) WP_063219646.1 WP_063219646.1 bacteriocin-processing peptidase family protein
1(1) WP_217035871.1 WP_217035871.1 Ig-like domain repeat protein
1(1) WP_066020839.1 WP_066020839.1 tetratricopeptide repeat protein
1(1) WP_098601636.1 WP_098601636.1 type VII secretion protein EsaA
1(1) WP_045518280.1 WP_045518280.1 WXG100 family type VII secretion target
1(1) WP_102371677.1 WP_102371677.1 T7SS effector LXG polymorphic toxin
1(1) WP_237728629.1 WP_237728629.1 WXG100 family type VII secretion target
1(1) WP_165097641.1 WP_165097641.1 glycosyltransferase family 2 protein
1(1) WP_061688406.1 WP_061688406.1 SseB family protein
1(1) WP_142332781.1 WP_142332781.1 WXG100 family type VII secretion target
1(1) WP_193653016.1 WP_193653016.1 hypothetical protein
1(1) WP_077619648.1 WP_077619648.1 DUF2974 domain-containing protein
1(1) WP_076309113.1 WP_076309113.1 hypothetical protein
1(1) WP_000404141.1 WP_000404141.1 WXG100 family type VII secretion target
1(2) WP_194336458.1 WP_194336458.1 polymorphic toxin type 15 domain-containing protein
1(1) WP_013490138.1 WP_013490138.1 Ig-like domain-containing protein
1(1) WP_194543497.1 WP_194543497.1 serine/threonine protein kinase
1(1) WP_205001920.1 WP_205001920.1 hypothetical protein
1(1) WP_097954754.1 WP_097954754.1 T7SS effector LXG polymorphic toxin
1(1) WP_048826571.1 WP_048826571.1 WXG100 family type VII secretion target
1(1) WP_085499095.1 WP_085499095.1 WXG100 family type VII secretion target
1(1) WP_185165645.1 WP_185165645.1 EsaB/YukD family protein
1(1) WP_179034791.1 WP_179034791.1 hypothetical protein
1(1) WP_102264553.1 WP_102264553.1 WXG100 family type VII secretion target
1(1) WP_255256111.1 WP_255256111.1 hypothetical protein, partial
1(1) WP_251542620.1 WP_251542620.1 serine/threonine-protein kinase
1(1) WP_281103268.1 WP_281103268.1 hypothetical protein
1(1) WP_233697174.1 WP_233697174.1 WXG100 family type VII secretion target
1(1) WP_145037674.1 WP_145037674.1 hypothetical protein
1(1) WP_307312553.1 WP_307312553.1 DUF2974 domain-containing protein
1(1) WP_038286717.1 WP_038286717.1 EsaB/YukD family protein
1(1) WP_024632699.1 WP_024632699.1 hypothetical protein
1(1) WP_160496929.1 WP_160496929.1 hypothetical protein
1(1) WP_195325249.1 WP_195325249.1 serine/threonine-protein kinase
1(1) WP_244723783.1 WP_244723783.1 WXG100 family type VII secretion target
1(1) WP_012260826.1 WP_012260826.1 tetratricopeptide repeat protein
1(1) WP_010676861.1 WP_010676861.1 WXG100 family type VII secretion target
1(1) WP_113713148.1 WP_113713148.1 hypothetical protein
1(1) WP_151208749.1 WP_151208749.1 protein kinase family protein
1(3) WP_088063233.1 WP_088063233.1 SseB family protein
1(1) WP_152402374.1 WP_152402374.1 hypothetical protein
1(1) WP_002036276.1 WP_002036276.1 hypothetical protein
1(1) WP_294953138.1 WP_294953138.1 WXG100 family type VII secretion target
1(1) WP_169505053.1 WP_169505053.1 hypothetical protein
1(1) WP_098856767.1 WP_098856767.1 tetratricopeptide repeat protein
1(3) WP_167555100.1 WP_167555100.1 tetratricopeptide repeat protein
1(1) WP_116353245.1 WP_116353245.1 serine/threonine-protein kinase
1(1) WP_240459621.1 WP_240459621.1 DNA/RNA non-specific endonuclease
1(1) WP_172801681.1 WP_172801681.1 hypothetical protein
1(1) WP_087956285.1 WP_087956285.1 bacteriocin-processing peptidase family protein
1(1) WP_281477420.1 WP_281477420.1 hypothetical protein
1(1) WP_187128405.1 WP_187128405.1 hypothetical protein
1(2) WP_281813107.1 WP_281813107.1 hypothetical protein
1(1) WP_013252388.1 WP_013252388.1 DUF2974 domain-containing protein
1(3) WP_291289937.1 WP_291289937.1 hypothetical protein
1(1) WP_027088433.1 WP_027088433.1 pre-toxin TG domain-containing protein
1(1) WP_308083991.1 WP_308083991.1 trypsin-like peptidase domain-containing protein
1(1) WP_071679490.1 WP_071679490.1 hypothetical protein
1(1) WP_110934075.1 WP_110934075.1 EsaB/YukD family protein
1(1) WP_041548999.1 WP_041548999.1 hypothetical protein
1(1) WP_291647963.1 WP_291647963.1 WXG100 family type VII secretion target
1(1) WP_076103997.1 WP_076103997.1 DUF5050 domain-containing protein
1(1) WP_137698109.1 WP_137698109.1 hypothetical protein
1(1) WP_017692350.1 WP_017692350.1 protein kinase family protein
1(1) WP_142543133.1 WP_142543133.1 copper amine oxidase N-terminal domain-containing protein
1(1) WP_284566227.1 WP_284566227.1 WXG100 family type VII secretion target
1(3) WP_071679486.1 WP_071679486.1 bacteriocin-processing peptidase family protein
1(2) WP_185639012.1 WP_185639012.1 DUF6531 domain-containing protein
1(1) WP_088007981.1 WP_088007981.1 SseB family protein
1(1) WP_110934072.1 WP_110934072.1 WXG100 family type VII secretion target
1(1) WP_088058539.1 WP_088058539.1 hypothetical protein
1(1) WP_257675219.1 WP_257675219.1 FHA domain-containing protein

2(1) WP_307349977.1 WP_307349977.1 vWA domain-containing protein
2(2) WP_062326150.1 WP_062326150.1 VWA domain-containing protein
2(1) WP_100539444.1 WP_100539444.1 VWA domain-containing protein
2(4) WP_007430135.1 WP_007430135.1 VWA domain-containing protein
2(1) WP_047909610.1 WP_047909610.1 vWA domain-containing protein
2(1) WP_305022920.1 WP_305022920.1 vWA domain-containing protein
2(1) WP_071745230.1 WP_071745230.1 vWA domain-containing protein
2(1) WP_211745827.1 WP_211745827.1 VWA domain-containing protein
2(1) WP_243896000.1 WP_243896000.1 VWA domain-containing protein
2(2) WP_242172379.1 WP_242172379.1 VWA domain-containing protein
2(2) WP_127543696.1 WP_127543696.1 VWA domain-containing protein
2(1) WP_197234725.1 WP_197234725.1 vWA domain-containing protein
2(4) WP_098768169.1 WP_098768169.1 vWA domain-containing protein
2(1) WP_064019317.1 WP_064019317.1 vWA domain-containing protein
2(1) WP_301804671.1 WP_301804671.1 vWA domain-containing protein
2(1) WP_289483782.1 WP_289483782.1 VWA domain-containing protein
2(1) WP_268583861.1 WP_268583861.1 VWA domain-containing protein
2(1) WP_223067435.1 WP_223067435.1 VWA domain-containing protein
2(2) WP_065228768.1 WP_065228768.1 vWA domain-containing protein
2(1) WP_286124780.1 WP_286124780.1 vWA domain-containing protein
2(2) WP_061805362.1 WP_061805362.1 vWA domain-containing protein
2(1) WP_147807656.1 WP_147807656.1 vWA domain-containing protein

| | | |
|---|---|---|
| 2(1) | WP_236417222.1 | WP_236417222.1 VWA domain-containing protein |
| 2(1) | WP_014281185.1 | WP_014281185.1 VWA domain-containing protein |
| 2(1) | WP_094699653.1 | WP_094699653.1 vWA domain-containing protein |
| 2(1) | WP_000966841.1 | WP_000966841.1 vWA domain-containing protein |
| 2(1) | WP_263703622.1 | WP_263703622.1 vWA domain-containing protein, partial |
| 2(2) | WP_071640582.1 | WP_071640582.1 VWA domain-containing protein |
| 2(1) | WP_251537230.1 | WP_251537230.1 VWA domain-containing protein |
| 2(1) | WP_124049173.1 | WP_124049173.1 vWA domain-containing protein |
| 2(4) | WP_076330724.1 | WP_076330724.1 VWA domain-containing protein |
| 2(1) | WP_230986884.1 | WP_230986884.1 vWA domain-containing protein |
| 2(1) | WP_098670171.1 | WP_098670171.1 vWA domain-containing protein |
| 2(1) | WP_003340551.1 | WP_003340551.1 vWA domain-containing protein |
| 2(1) | WP_017149774.1 | WP_017149774.1 vWA domain-containing protein |
| 2(2) | WP_097898670.1 | WP_097898670.1 vWA domain-containing protein |
| 2(1) | WP_233697178.1 | WP_233697178.1 vWA domain-containing protein |
| 2(15) | WP_017689069.1 | WP_017689069.1 VWA domain-containing protein |
| 2(1) | WP_104030440.1 | WP_104030440.1 vWA domain-containing protein |
| 2(3) | WP_095266200.1 | WP_095266200.1 VWA domain-containing protein |
| 2(1) | WP_251375547.1 | WP_251375547.1 vWA domain-containing protein |
| 2(1) | WP_094593878.1 | WP_094593878.1 VWA domain-containing protein |
| 2(1) | WP_047153274.1 | WP_047153274.1 VWA domain-containing protein |
| 2(1) | WP_002158047.1 | WP_002158047.1 vWA domain-containing protein |
| 2(1) | WP_258425843.1 | WP_258425843.1 vWA domain-containing protein |
| 2(5) | WP_056700796.1 | WP_056700796.1 VWA domain-containing protein |
| 2(2) | WP_119732859.1 | WP_119732859.1 vWA domain-containing protein |
| 2(1) | WP_208019347.1 | WP_208019347.1 vWA domain-containing protein |
| 2(1) | WP_290401991.1 | WP_290401991.1 vWA domain-containing protein |
| 2(1) | WP_021253114.1 | WP_021253114.1 vWA domain-containing protein |
| 2(1) | WP_108465351.1 | WP_108465351.1 vWA domain-containing protein |
| 2(1) | WP_061535953.1 | WP_061535953.1 vWA domain-containing protein |
| 2(2) | WP_194230399.1 | WP_194230399.1 VWA domain-containing protein |
| 2(1) | WP_232274492.1 | WP_232274492.1 vWA domain-containing protein |
| 2(1) | WP_258418571.1 | WP_258418571.1 vWA domain-containing protein |
| 2(28) | WP_013370788.1 | WP_013370788.1 VWA domain-containing protein |
| 2(1) | WP_025719056.1 | WP_025719056.1 VWA domain-containing protein |
| 2(1) | WP_129478246.1 | WP_129478246.1 VWA domain-containing protein |
| 2(8) | WP_005546518.1 | WP_005546518.1 vWA domain-containing protein |
| 2(1) | WP_137052039.1 | WP_137052039.1 vWA domain-containing protein |
| 2(10) | WP_036661656.1 | WP_036661656.1 VWA domain-containing protein |
| 2(2) | WP_018670519.1 | WP_018670519.1 vWA domain-containing protein |
| 2(4) | WP_024630329.1 | WP_024630329.1 VWA domain-containing protein |
| 2(1) | WP_243518750.1 | WP_243518750.1 vWA domain-containing protein |
| 2(1) | WP_213618817.1 | WP_213618817.1 vWA domain-containing protein |
| 2(1) | WP_160496928.1 | WP_160496928.1 VWA domain-containing protein |
| 2(2) | WP_076311047.1 | WP_076311047.1 vWA domain-containing protein |
| 2(2) | WP_063566320.1 | WP_063566320.1 VWA domain-containing protein |
| 2(1) | WP_213427024.1 | WP_213427024.1 vWA domain-containing protein |
| 2(1) | WP_053783889.1 | WP_053783889.1 VWA domain-containing protein |
| 2(1) | WP_002171724.1 | WP_002171724.1 vWA domain-containing protein |
| 2(1) | WP_305755031.1 | WP_305755031.1 VWA domain-containing protein |
| 2(2) | WP_143801058.1 | WP_143801058.1 vWA domain-containing protein |
| 2(1) | WP_021624819.1 | WP_021624819.1 vWA domain-containing protein |

| | | |
|---|---|---|
| 2(1) | WP_258126007.1 | WP_258126007.1 vWA domain-containing protein |
| 2(1) | WP_013914187.1 | WP_013914187.1 vWA domain-containing protein |
| 2(2) | WP_006674858.1 | WP_006674858.1 vWA domain-containing protein |
| 2(1) | WP_031415577.1 | WP_031415577.1 vWA domain-containing protein |
| 2(2) | WP_145329909.1 | WP_145329909.1 VWA domain-containing protein |
| 2(1) | WP_090993625.1 | WP_090993625.1 VWA domain-containing protein |
| 2(1) | WP_061975577.1 | WP_061975577.1 VWA domain-containing protein |
| 2(1) | WP_248251182.1 | WP_248251182.1 vWA domain-containing protein |
| 2(1) | WP_053325281.1 | WP_053325281.1 VWA domain-containing protein |
| 2(1) | WP_053403785.1 | WP_053403785.1 vWA domain-containing protein |
| 2(1) | WP_193653023.1 | WP_193653023.1 vWA domain-containing protein |
| 2(1) | WP_256227715.1 | WP_256227715.1 vWA domain-containing protein |
| 2(2) | WP_076322112.1 | WP_076322112.1 VWA domain-containing protein |
| 2(1) | WP_138226021.1 | WP_138226021.1 VWA domain-containing protein |
| 2(1) | WP_028401641.1 | WP_028401641.1 vWA domain-containing protein |
| 2(5) | WP_002031626.1 | WP_002031626.1 vWA domain-containing protein |
| 2(1) | WP_307589688.1 | WP_307589688.1 vWA domain-containing protein |
| 2(1) | WP_068782410.1 | WP_068782410.1 vWA domain-containing protein |
| 2(1) | WP_036707113.1 | WP_036707113.1 vWA domain-containing protein |
| 2(1) | WP_097961542.1 | WP_097961542.1 vWA domain-containing protein |
| 2(1) | WP_251583294.1 | WP_251583294.1 vWA domain-containing protein |
| 2(1) | WP_239617669.1 | WP_239617669.1 VWA domain-containing protein |
| 2(1) | WP_245948227.1 | WP_245948227.1 VWA domain-containing protein |
| 2(5) | WP_071220489.1 | WP_071220489.1 VWA domain-containing protein |
| 2(1) | WP_301879144.1 | WP_301879144.1 VWA domain-containing protein |
| 2(5) | WP_026009532.1 | WP_026009532.1 vWA domain-containing protein |
| 2(1) | WP_098266646.1 | WP_098266646.1 vWA domain-containing protein |
| 2(1) | WP_113058865.1 | WP_113058865.1 VWA domain-containing protein |
| 2(3) | WP_000966843.1 | WP_000966843.1 vWA domain-containing protein |
| 2(4) | WP_168178886.1 | WP_168178886.1 vWA domain-containing protein |
| 2(1) | WP_104148795.1 | WP_104148795.1 vWA domain-containing protein |
| 2(1) | WP_024632687.1 | WP_024632687.1 vWA domain-containing protein |
| 2(1) | WP_091155385.1 | WP_091155385.1 vWA domain-containing protein |
| 2(1) | WP_062239875.1 | WP_062239875.1 vWA domain-containing protein |
| 2(2) | WP_124114120.1 | WP_124114120.1 VWA domain-containing protein |
| 2(2) | WP_028533176.1 | WP_028533176.1 vWA domain-containing protein |
| 2(1) | WP_017692348.1 | WP_017692348.1 vWA domain-containing protein |
| 2(1) | WP_229522854.1 | WP_229522854.1 VWA domain-containing protein |
| 2(1) | WP_183578463.1 | WP_183578463.1 VWA domain-containing protein |
| 2(2) | WP_197262007.1 | WP_197262007.1 vWA domain-containing protein |
| 2(1) | WP_068694718.1 | WP_068694718.1 vWA domain-containing protein |
| 2(1) | WP_113713143.1 | WP_113713143.1 vWA domain-containing protein |
| 2(3) | WP_002148332.1 | WP_002148332.1 VWA domain-containing protein |
| 2(2) | WP_104065533.1 | WP_104065533.1 vWA domain-containing protein |
| 2(1) | WP_213649968.1 | WP_213649968.1 VWA domain-containing protein |
| 2(2) | WP_025716878.1 | WP_025716878.1 VWA domain-containing protein |
| 2(1) | WP_077567016.1 | WP_077567016.1 VWA domain-containing protein |
| 2(27) | WP_013309992.1 | WP_013309992.1 VWA domain-containing protein |
| 2(6) | WP_012260832.1 | WP_012260832.1 VWA domain-containing protein |
| 2(1) | WP_301167496.1 | WP_301167496.1 VWA domain-containing protein |
| 2(1) | WP_256872565.1 | WP_256872565.1 vWA domain-containing protein |
| 2(2) | WP_144611185.1 | WP_144611185.1 vWA domain-containing protein |

| | | |
|---|---|---|
| 2(1) | WP_068696882.1 | WP_068696882.1 vWA domain-containing protein |
| 2(13) | WP_076167860.1 | WP_076167860.1 VWA domain-containing protein |
| 2(1) | WP_204517513.1 | WP_204517513.1 vWA domain-containing protein |
| 2(1) | WP_098508784.1 | WP_098508784.1 vWA domain-containing protein |
| 2(1) | WP_229263935.1 | WP_229263935.1 VWA domain-containing protein |
| 2(1) | WP_068619105.1 | WP_068619105.1 vWA domain-containing protein |
| 2(1) | WP_055105792.1 | WP_055105792.1 vWA domain-containing protein |
| 2(1) | WP_262748112.1 | WP_262748112.1 vWA domain-containing protein |
| 2(3) | WP_088020916.1 | WP_088020916.1 vWA domain-containing protein |
| 2(1) | WP_265408848.1 | WP_265408848.1 VWA domain-containing protein |
| 2(1) | WP_246096656.1 | WP_246096656.1 vWA domain-containing protein |
| 2(1) | WP_010346280.1 | WP_010346280.1 VWA domain-containing protein |
| 2(4) | WP_021261879.1 | WP_021261879.1 vWA domain-containing protein |
| 2(1) | WP_243644269.1 | WP_243644269.1 vWA domain-containing protein |
| 2(1) | WP_276917306.1 | WP_276917306.1 VWA domain-containing protein |
| 2(1) | WP_212981561.1 | WP_212981561.1 vWA domain-containing protein |
| 2(1) | WP_098303179.1 | WP_098303179.1 vWA domain-containing protein |
| 2(1) | WP_003335998.1 | WP_003335998.1 vWA domain-containing protein |
| 2(5) | WP_000966845.1 | WP_000966845.1 VWA domain-containing protein |
| 2(1) | WP_065291245.1 | WP_065291245.1 vWA domain-containing protein |
| 2(1) | WP_183600894.1 | WP_183600894.1 vWA domain-containing protein |
| 2(1) | WP_116334081.1 | WP_116334081.1 vWA domain-containing protein |
| 2(1) | WP_097905182.1 | WP_097905182.1 vWA domain-containing protein |
| 2(1) | WP_208919040.1 | WP_208919040.1 VWA domain-containing protein |
| 2(1) | WP_268623243.1 | WP_268623243.1 VWA domain-containing protein |
| 2(1) | WP_188593163.1 | WP_188593163.1 vWA domain-containing protein |
| 2(1) | WP_243495558.1 | WP_243495558.1 vWA domain-containing protein |
| 2(1) | WP_248063434.1 | WP_248063434.1 vWA domain-containing protein |
| 2(1) | WP_205001916.1 | WP_205001916.1 VWA domain-containing protein |
| 2(2) | WP_111925420.1 | WP_111925420.1 vWA domain-containing protein |
| 2(1) | WP_057897620.1 | WP_057897620.1 vWA domain-containing protein |
| 2(1) | WP_241929675.1 | WP_241929675.1 VWA domain-containing protein |
| 2(1) | WP_090805476.1 | WP_090805476.1 VWA domain-containing protein |
| 2(1) | WP_116556150.1 | WP_116556150.1 vWA domain-containing protein |
| 2(1) | WP_098746862.1 | WP_098746862.1 VWA domain-containing protein |
| 2(2) | WP_197247515.1 | WP_197247515.1 vWA domain-containing protein |
| 2(1) | WP_282103174.1 | WP_282103174.1 vWA domain-containing protein |
| 2(38) | WP_044792564.1 | WP_044792564.1 vWA domain-containing protein |
| 2(1) | WP_116231173.1 | WP_116231173.1 VWA domain-containing protein |
| 2(1) | WP_110899368.1 | WP_110899368.1 vWA domain-containing protein |
| 2(1) | WP_268613695.1 | WP_268613695.1 VWA domain-containing protein |
| 2(1) | WP_207994164.1 | WP_207994164.1 vWA domain-containing protein |
| 2(2) | WP_063223275.1 | WP_063223275.1 vWA domain-containing protein |
| 2(1) | WP_246246293.1 | WP_246246293.1 VWA domain-containing protein |
| 2(1) | WP_109999893.1 | WP_109999893.1 VWA domain-containing protein |
| 2(1) | WP_238188556.1 | WP_238188556.1 vWA domain-containing protein |
| 2(2) | WP_098248512.1 | WP_098248512.1 vWA domain-containing protein |
| 2(1) | WP_151630607.1 | WP_151630607.1 vWA domain-containing protein |
| 2(1) | WP_023957731.1 | WP_023957731.1 vWA domain-containing protein |
| 2(1) | WP_088078458.1 | WP_088078458.1 vWA domain-containing protein |
| 2(1) | WP_250244821.1 | WP_250244821.1 VWA domain-containing protein |
| 2(1) | WP_242400022.1 | WP_242400022.1 vWA domain-containing protein |
| 2(1) | WP_145318689.1 | WP_145318689.1 VWA domain-containing protein |
| 2(1) | WP_160037490.1 | WP_160037490.1 vWA domain-containing protein |
| 2(1) | WP_036642471.1 | WP_036642471.1 VWA domain-containing protein |
| 2(1) | WP_160789865.1 | WP_160789865.1 VWA domain-containing protein, partial |
| 2(3) | WP_064473939.1 | WP_064473939.1 vWA domain-containing protein |
| 2(37) | WP_000966846.1 | WP_000966846.1 vWA domain-containing protein |
| 2(1) | WP_136789992.1 | WP_136789992.1 vWA domain-containing protein |
| 2(1) | WP_213532659.1 | WP_213532659.1 VWA domain-containing protein |
| 2(1) | WP_213644230.1 | WP_213644230.1 VWA domain-containing protein |
| 2(1) | WP_061402731.1 | WP_061402731.1 vWA domain-containing protein |
| 2(1) | WP_022586767.1 | WP_022586767.1 vWA domain-containing protein |
| 2(1) | WP_220031162.1 | WP_220031162.1 VWA domain-containing protein |
| 2(1) | WP_068536368.1 | WP_068536368.1 VWA domain-containing protein |
| 2(1) | WP_242255494.1 | WP_242255494.1 vWA domain-containing protein |
| 2(1) | WP_184183008.1 | WP_184183008.1 vWA domain-containing protein |
| 2(1) | WP_054958412.1 | WP_054958412.1 VWA domain-containing protein |
| 2(2) | WP_212956328.1 | WP_212956328.1 vWA domain-containing protein |
| 2(1) | WP_136604818.1 | WP_136604818.1 VWA domain-containing protein |
| 2(1) | WP_244719648.1 | WP_244719648.1 VWA domain-containing protein |
| 2(11) | WP_064639011.1 | WP_064639011.1 VWA domain-containing protein |
| 2(1) | WP_102997919.1 | WP_102997919.1 VWA domain-containing protein |
| 2(6) | WP_096886738.1 | WP_096886738.1 vWA domain-containing protein |
| 2(1) | WP_098776431.1 | WP_098776431.1 vWA domain-containing protein |
| 2(1) | WP_166244966.1 | WP_166244966.1 vWA domain-containing protein |
| 2(3) | WP_110892966.1 | WP_110892966.1 VWA domain-containing protein |
| 2(2) | WP_283892173.1 | WP_283892173.1 VWA domain-containing protein |
| 2(1) | WP_258433019.1 | WP_258433019.1 vWA domain-containing protein |
| 2(1) | WP_086419833.1 | WP_086419833.1 vWA domain-containing protein |
| 2(1) | WP_068793932.1 | WP_068793932.1 vWA domain-containing protein |
| 2(1) | WP_111273171.1 | WP_111273171.1 vWA domain-containing protein |
| 2(2) | WP_052092410.1 | WP_052092410.1 vWA domain-containing protein |
| 2(1) | WP_090239219.1 | WP_090239219.1 VWA domain-containing protein |
| 2(1) | WP_127199774.1 | WP_127199774.1 vWA domain-containing protein |
| 2(5) | WP_006211200.1 | WP_006211200.1 VWA domain-containing protein |
| 2(2) | WP_026593082.1 | WP_026593082.1 vWA domain-containing protein |
| 2(1) | WP_127592425.1 | WP_127592425.1 VWA domain-containing protein |
| 2(2) | WP_014368240.1 | WP_014368240.1 vWA domain-containing protein |
| 2(4) | WP_145153652.1 | WP_145153652.1 vWA domain-containing protein |
| 2(1) | WP_215572751.1 | WP_215572751.1 VWA domain-containing protein |
| 2(38) | WP_000966842.1 | WP_000966842.1 vWA domain-containing protein |
| 2(1) | WP_307394637.1 | WP_307394637.1 vWA domain-containing protein |
| 2(1) | WP_191798530.1 | WP_191798530.1 vWA domain-containing protein |
| 2(6) | WP_095287658.1 | WP_095287658.1 VWA domain-containing protein |
| 2(2) | WP_087442416.1 | WP_087442416.1 vWA domain-containing protein |
| 2(1) | WP_220559125.1 | WP_220559125.1 vWA domain-containing protein |
| 2(1) | WP_106765955.1 | WP_106765955.1 VWA domain-containing protein |
| 2(3) | WP_087956291.1 | WP_087956291.1 vWA domain-containing protein |
| 2(3) | WP_087433610.1 | WP_087433610.1 VWA domain-containing protein |
| 2(1) | WP_163245185.1 | WP_163245185.1 vWA domain-containing protein |
| 2(1) | WP_090908775.1 | WP_090908775.1 VWA domain-containing protein |
| 2(3) | WP_007128625.1 | WP_007128625.1 VWA domain-containing protein |
| 2(45) | WP_000966848.1 | WP_000966848.1 vWA domain-containing protein |

| | | | |
|---|---|---|---|
| 2(7) | WP_025685035.1 | WP_025685035.1 | VWA domain-containing protein |
| 2(3) | WP_088058533.1 | WP_088058533.1 | vWA domain-containing protein |
| 2(4) | WP_113755277.1 | WP_113755277.1 | vWA domain-containing protein |
| 2(2) | WP_216518258.1 | WP_216518258.1 | vWA domain-containing protein |
| 2(1) | WP_076210971.1 | WP_076210971.1 | vWA domain-containing protein |
| 2(1) | WP_119794366.1 | WP_119794366.1 | vWA domain-containing protein |
| 2(2) | WP_272035328.1 | WP_272035328.1 | VWA domain-containing protein |
| 2(1) | WP_019424142.1 | WP_019424142.1 | vWA domain-containing protein |
| 2(1) | WP_285748181.1 | WP_285748181.1 | vWA domain-containing protein |
| 2(2) | WP_090921050.1 | WP_090921050.1 | VWA domain-containing protein |
| 2(1) | WP_242251774.1 | WP_242251774.1 | vWA domain-containing protein |
| 2(1) | WP_155703931.1 | WP_155703931.1 | vWA domain-containing protein |
| 2(1) | WP_137062802.1 | WP_137062802.1 | vWA domain-containing protein |
| 2(1) | WP_247083040.1 | WP_247083040.1 | vWA domain-containing protein |
| 2(1) | WP_230632628.1 | WP_230632628.1 | VWA domain-containing protein |
| 2(2) | WP_043067143.1 | WP_043067143.1 | VWA domain-containing protein |
| 2(1) | WP_232699386.1 | WP_232699386.1 | vWA domain-containing protein |
| 2(1) | WP_245590366.1 | WP_245590366.1 | VWA domain-containing protein |
| 2(1) | WP_123067117.1 | WP_123067117.1 | VWA domain-containing protein |
| 2(1) | WP_168976290.1 | WP_168976290.1 | VWA domain-containing protein |
| 2(1) | WP_094092624.1 | WP_094092624.1 | vWA domain-containing protein |
| 2(1) | WP_154985825.1 | WP_154985825.1 | vWA domain-containing protein |
| 2(2) | WP_018883269.1 | WP_018883269.1 | VWA domain-containing protein |
| 2(1) | WP_053492742.1 | WP_053492742.1 | VWA domain-containing protein |
| 2(1) | WP_110931543.1 | WP_110931543.1 | vWA domain-containing protein |
| 2(1) | WP_246027917.1 | WP_246027917.1 | VWA domain-containing protein |
| 2(2) | WP_133378769.1 | WP_133378769.1 | vWA domain-containing protein |
| 2(1) | WP_235286298.1 | WP_235286298.1 | VWA domain-containing protein |
| 2(1) | WP_278431127.1 | WP_278431127.1 | VWA domain-containing protein |
| 2(1) | WP_209861639.1 | WP_209861639.1 | vWA domain-containing protein |
| 2(2) | WP_029438089.1 | WP_029438089.1 | vWA domain-containing protein |

| | | | |
|---|---|---|---|
| 3(1) | WP_095206168.1 | WP_095206168.1 | hypothetical protein |
| 3(1) | WP_094155405.1 | WP_094155405.1 | hypothetical protein |
| 3(4) | WP_069355508.1 | WP_069355508.1 | hypothetical protein |
| 3(1) | WP_160038439.1 | WP_160038439.1 | hypothetical protein |
| 3(1) | WP_190299138.1 | WP_190299138.1 | hypothetical protein |
| 3(1) | WP_258875220.1 | WP_258875220.1 | hypothetical protein |
| 3(1) | WP_144024451.1 | WP_144024451.1 | hypothetical protein |
| 3(1) | WP_000241767.1 | WP_000241767.1 | hypothetical protein |
| 3(1) | WP_085117882.1 | WP_085117882.1 | hypothetical protein |
| 3(1) | WP_091017192.1 | WP_091017192.1 | hypothetical protein |
| 3(1) | WP_098303178.1 | WP_098303178.1 | hypothetical protein |
| 3(1) | WP_090921048.1 | WP_090921048.1 | hypothetical protein |
| 3(1) | WP_272045567.1 | WP_272045567.1 | hypothetical protein |
| 3(1) | WP_234032639.1 | WP_234032639.1 | hypothetical protein |
| 3(1) | WP_278431126.1 | WP_278431126.1 | hypothetical protein |
| 3(1) | WP_213426987.1 | WP_213426987.1 | hypothetical protein |
| 3(1) | WP_098150444.1 | WP_098150444.1 | hypothetical protein |
| 3(1) | WP_136789993.1 | WP_136789993.1 | PP2C family serine/threonine-protein phosphatase |
| 3(8) | WP_013309993.1 | WP_013309993.1 | hypothetical protein |
| 3(1) | WP_192510490.1 | WP_192510490.1 | hypothetical protein |
| 3(1) | WP_036670437.1 | WP_036670437.1 | hypothetical protein |
| 3(1) | WP_272035330.1 | WP_272035330.1 | hypothetical protein |
| 3(1) | WP_232699385.1 | WP_232699385.1 | PP2C family serine/threonine-protein phosphatase |
| 3(1) | WP_094699654.1 | WP_094699654.1 | hypothetical protein |
| 3(1) | WP_012260833.1 | WP_012260833.1 | hypothetical protein |
| 3(1) | WP_035119749.1 | WP_035119749.1 | protein phosphatase 2C domain-containing protein |
| 3(2) | WP_262866587.1 | WP_262866587.1 | hypothetical protein |
| 3(1) | WP_301249700.1 | WP_301249700.1 | hypothetical protein |
| 3(1) | WP_243518753.1 | WP_243518753.1 | hypothetical protein |
| 3(1) | WP_272640517.1 | WP_272640517.1 | hypothetical protein |
| 3(1) | WP_155703930.1 | WP_155703930.1 | hypothetical protein |
| 3(1) | WP_236417426.1 | WP_236417426.1 | hypothetical protein |
| 3(1) | WP_098670172.1 | WP_098670172.1 | hypothetical protein |
| 3(1) | WP_091260869.1 | WP_091260869.1 | hypothetical protein |
| 3(2) | WP_006211198.1 | WP_006211198.1 | protein phosphatase 2C domain-containing protein |
| 3(1) | WP_078993365.1 | WP_078993365.1 | hypothetical protein |
| 3(1) | WP_000878001.1 | WP_000878001.1 | hypothetical protein |
| 3(1) | WP_098562454.1 | WP_098562454.1 | hypothetical protein |
| 3(2) | WP_091031765.1 | WP_091031765.1 | hypothetical protein |
| 3(1) | WP_272801672.1 | WP_272801672.1 | PP2C family serine/threonine-protein phosphatase |
| 3(1) | WP_124049172.1 | WP_124049172.1 | hypothetical protein |
| 3(1) | WP_260985319.1 | WP_260985319.1 | hypothetical protein |
| 3(1) | WP_238333011.1 | WP_238333011.1 | hypothetical protein |
| 3(1) | WP_228100912.1 | WP_228100912.1 | hypothetical protein |
| 3(1) | WP_116334080.1 | WP_116334080.1 | hypothetical protein |
| 3(5) | WP_000241762.1 | WP_000241762.1 | hypothetical protein |
| 3(1) | WP_127592424.1 | WP_127592424.1 | hypothetical protein |
| 3(1) | WP_098250524.1 | WP_098250524.1 | hypothetical protein |
| 3(2) | WP_233280993.1 | WP_233280993.1 | PP2C family serine/threonine-protein phosphatase |
| 3(7) | WP_024927870.1 | WP_024927870.1 | hypothetical protein |
| 3(1) | WP_134750596.1 | WP_134750596.1 | hypothetical protein |
| 3(1) | WP_038584747.1 | WP_038584747.1 | hypothetical protein |
| 3(1) | WP_076322111.1 | WP_076322111.1 | hypothetical protein |
| 3(1) | WP_068619104.1 | WP_068619104.1 | hypothetical protein |
| 3(1) | WP_142613004.1 | WP_142613004.1 | hypothetical protein |
| 3(1) | WP_144611186.1 | WP_144611186.1 | hypothetical protein |
| 3(1) | WP_199795351.1 | WP_199795351.1 | protein phosphatase 2C domain-containing protein |
| 3(1) | WP_197234723.1 | WP_197234723.1 | hypothetical protein |
| 3(1) | WP_104148794.1 | WP_104148794.1 | hypothetical protein |
| 3(2) | WP_088327111.1 | WP_088327111.1 | hypothetical protein |
| 3(1) | WP_057897621.1 | WP_057897621.1 | hypothetical protein |
| 3(1) | WP_025719054.1 | WP_025719054.1 | hypothetical protein |
| 3(6) | WP_025118303.1 | WP_025118303.1 | hypothetical protein |
| 3(1) | WP_021624820.1 | WP_021624820.1 | hypothetical protein |
| 3(3) | WP_088063231.1 | WP_088063231.1 | hypothetical protein |
| 3(1) | WP_139867385.1 | WP_139867385.1 | hypothetical protein |
| 3(5) | WP_023988268.1 | WP_023988268.1 | hypothetical protein |
| 3(1) | WP_262729428.1 | WP_262729428.1 | hypothetical protein |
| 3(1) | WP_192266461.1 | WP_192266461.1 | hypothetical protein |
| 3(1) | WP_282321818.1 | WP_282321818.1 | hypothetical protein |
| 3(1) | WP_077567015.1 | WP_077567015.1 | hypothetical protein |
| 3(1) | WP_152381396.1 | WP_152381396.1 | hypothetical protein |

| | | |
|---|---|---|
| 3(2) | WP_097898671.1 | WP_097898671.1 hypothetical protein |
| 3(1) | WP_113052563.1 | WP_113052563.1 hypothetical protein |
| 3(12) | WP_000241759.1 | WP_000241759.1 hypothetical protein |
| 3(2) | WP_143801057.1 | WP_143801057.1 hypothetical protein |
| 3(1) | WP_206773213.1 | WP_206773213.1 hypothetical protein |
| 3(2) | WP_098768168.1 | WP_098768168.1 hypothetical protein |
| 3(1) | WP_274566441.1 | WP_274566441.1 hypothetical protein |
| 3(2) | WP_098746860.1 | WP_098746860.1 hypothetical protein |
| 3(1) | WP_090949168.1 | WP_090949168.1 hypothetical protein |
| 3(1) | WP_216513493.1 | WP_216513493.1 protein phosphatase 2C domain-containing protein |
| 3(1) | WP_088032776.1 | WP_088032776.1 hypothetical protein |
| 3(1) | WP_136372965.1 | WP_136372965.1 hypothetical protein |
| 3(1) | WP_064639010.1 | WP_064639010.1 hypothetical protein |
| 3(1) | WP_307214589.1 | WP_307214589.1 hypothetical protein |
| 3(1) | WP_251497190.1 | WP_251497190.1 hypothetical protein |
| 3(1) | WP_147786237.1 | WP_147786237.1 hypothetical protein |
| 3(1) | WP_308721986.1 | WP_308721986.1 hypothetical protein |
| 3(1) | WP_265408847.1 | WP_265408847.1 PP2C family serine/threonine-protein phosphatase |
| 3(1) | WP_098227655.1 | WP_098227655.1 hypothetical protein |
| 3(2) | WP_098925580.1 | WP_098925580.1 hypothetical protein |
| 3(1) | WP_111925421.1 | WP_111925421.1 hypothetical protein |
| 3(1) | WP_166244900.1 | WP_166244900.1 protein phosphatase 2C domain-containing protein |
| 3(2) | WP_014368241.1 | WP_014368241.1 hypothetical protein |
| 3(2) | WP_197267872.1 | WP_197267872.1 hypothetical protein |
| 3(1) | WP_098266647.1 | WP_098266647.1 protein phosphatase 2C domain-containing protein |
| 3(1) | WP_168976289.1 | WP_168976289.1 hypothetical protein |
| 3(1) | WP_243495559.1 | WP_243495559.1 protein phosphatase 2C domain-containing protein |
| 3(1) | WP_076318552.1 | WP_076318552.1 hypothetical protein |
| 3(1) | WP_149238903.1 | WP_149238903.1 protein phosphatase 2C domain-containing protein |
| 3(1) | WP_139070690.1 | WP_139070690.1 hypothetical protein |
| 3(1) | WP_023957733.1 | WP_023957733.1 hypothetical protein |
| 3(1) | WP_155987324.1 | WP_155987324.1 hypothetical protein |
| 3(1) | WP_096774453.1 | WP_096774453.1 hypothetical protein |
| 3(4) | WP_078181465.1 | WP_078181465.1 hypothetical protein |
| 3(1) | WP_241929674.1 | WP_241929674.1 hypothetical protein |
| 3(11) | WP_000241763.1 | WP_000241763.1 hypothetical protein |
| 3(1) | WP_051317149.1 | WP_051317149.1 protein phosphatase 2C domain-containing protein |
| 3(1) | WP_113713144.1 | WP_113713144.1 hypothetical protein |
| 3(1) | WP_144488219.1 | WP_144488219.1 hypothetical protein |
| 3(1) | WP_000241765.1 | WP_000241765.1 hypothetical protein |
| 3(2) | WP_242172377.1 | WP_242172377.1 hypothetical protein |
| 3(1) | WP_110892964.1 | WP_110892964.1 protein phosphatase 2C domain-containing protein |
| 3(1) | WP_208919042.1 | WP_208919042.1 hypothetical protein |
| 3(1) | WP_098254025.1 | WP_098254025.1 hypothetical protein |
| 3(1) | WP_270699378.1 | WP_270699378.1 hypothetical protein |
| 3(1) | WP_160496927.1 | WP_160496927.1 hypothetical protein |
| 3(1) | WP_054402904.1 | WP_054402904.1 hypothetical protein |
| 3(1) | WP_134902007.1 | WP_134902007.1 hypothetical protein, partial |
| 3(1) | WP_100537973.1 | WP_100537973.1 hypothetical protein |
| 3(1) | WP_116556149.1 | WP_116556149.1 hypothetical protein |
| 3(1) | WP_281477423.1 | WP_281477423.1 hypothetical protein |
| 3(1) | WP_152826368.1 | WP_152826368.1 hypothetical protein |
| 3(1) | WP_263703623.1 | WP_263703623.1 hypothetical protein |
| 3(1) | WP_284566363.1 | WP_284566363.1 hypothetical protein |
| 3(2) | WP_083584548.1 | WP_083584548.1 hypothetical protein |
| 3(1) | WP_248565449.1 | WP_248565449.1 protein phosphatase 2C domain-containing protein |
| 3(1) | WP_099853193.1 | WP_099853193.1 hypothetical protein |
| 3(1) | WP_213649967.1 | WP_213649967.1 hypothetical protein |
| 3(1) | WP_212805120.1 | WP_212805120.1 hypothetical protein |
| 3(1) | WP_188593164.1 | WP_188593164.1 hypothetical protein |
| 3(1) | WP_087995245.1 | WP_087995245.1 hypothetical protein |
| 3(4) | WP_000878002.1 | WP_000878002.1 hypothetical protein |
| 3(2) | WP_098082496.1 | WP_098082496.1 hypothetical protein |
| 3(1) | WP_313955164.1 | WP_313955164.1 hypothetical protein |
| 3(2) | WP_090896196.1 | WP_090896196.1 hypothetical protein |
| 3(1) | WP_007430136.1 | WP_007430136.1 hypothetical protein |
| 3(1) | WP_078419971.1 | WP_078419971.1 hypothetical protein |
| 3(2) | WP_268619938.1 | WP_268619938.1 hypothetical protein |
| 3(1) | WP_149096533.1 | WP_149096533.1 hypothetical protein |
| 3(2) | WP_003340548.1 | WP_003340548.1 hypothetical protein |
| 3(1) | WP_305166259.1 | WP_305166259.1 hypothetical protein |
| 3(1) | WP_129478244.1 | WP_129478244.1 hypothetical protein |
| 3(1) | WP_258433018.1 | WP_258433018.1 hypothetical protein |
| 3(1) | WP_242400023.1 | WP_242400023.1 hypothetical protein |
| 3(1) | WP_111153299.1 | WP_111153299.1 hypothetical protein |
| 3(1) | WP_286459867.1 | WP_286459867.1 protein phosphatase 2C domain-containing protein |
| 3(1) | WP_168178783.1 | WP_168178783.1 PP2C family serine/threonine-protein phosphatase |
| 3(2) | WP_071220207.1 | WP_071220207.1 hypothetical protein |
| 3(1) | WP_107734207.1 | WP_107734207.1 hypothetical protein |
| 3(1) | WP_090908281.1 | WP_090908281.1 hypothetical protein |
| 3(1) | WP_242251776.1 | WP_242251776.1 hypothetical protein |
| 3(2) | WP_098264284.1 | WP_098264284.1 hypothetical protein |
| 3(1) | WP_240379732.1 | WP_240379732.1 hypothetical protein |
| 3(2) | WP_098558849.1 | WP_098558849.1 hypothetical protein |
| 3(1) | WP_024630330.1 | WP_024630330.1 hypothetical protein |
| 3(1) | WP_256227716.1 | WP_256227716.1 PP2C family serine/threonine-protein phosphatase |
| 3(1) | WP_285852595.1 | WP_285852595.1 hypothetical protein |
| 3(1) | WP_036642165.1 | WP_036642165.1 protein phosphatase 2C domain-containing protein |
| 3(1) | WP_237100033.1 | WP_237100033.1 hypothetical protein |
| 3(1) | WP_254327966.1 | WP_254327966.1 hypothetical protein |
| 3(1) | WP_071724718.1 | WP_071724718.1 hypothetical protein |
| 3(1) | WP_148246951.1 | WP_148246951.1 hypothetical protein |
| 3(1) | WP_064473938.1 | WP_064473938.1 hypothetical protein |
| 3(1) | WP_019424141.1 | WP_019424141.1 hypothetical protein |
| 3(1) | WP_010346279.1 | WP_010346279.1 hypothetical protein |
| 3(1) | WP_098976052.1 | WP_098976052.1 hypothetical protein |
| 3(1) | WP_289475116.1 | WP_289475116.1 PP2C family serine/threonine-protein phosphatase |
| 3(1) | WP_301804669.1 | WP_301804669.1 hypothetical protein |
| 3(1) | WP_088046781.1 | WP_088046781.1 hypothetical protein |
| 3(1) | WP_283892174.1 | WP_283892174.1 hypothetical protein |
| 3(1) | WP_064019318.1 | WP_064019318.1 hypothetical protein |
| 3(2) | WP_044792562.1 | WP_044792562.1 hypothetical protein |
| 3(1) | WP_276917308.1 | WP_276917308.1 hypothetical protein |
| 3(1) | WP_307589592.1 | WP_307589592.1 hypothetical protein |

| | | |
|---|---|---|
| 3(1) | WP_098248511.1 | WP_098248511.1 PP2C family serine/threonine-protein phosphatase |
| 3(1) | WP_139295272.1 | WP_139295272.1 hypothetical protein |
| 3(2) | WP_145329910.1 | WP_145329910.1 hypothetical protein |
| 3(2) | WP_052511637.1 | WP_052511637.1 hypothetical protein |
| 3(1) | WP_250261570.1 | WP_250261570.1 hypothetical protein |
| 3(1) | WP_063208774.1 | WP_063208774.1 hypothetical protein |
| 3(1) | WP_090688950.1 | WP_090688950.1 hypothetical protein |
| 3(1) | WP_076150488.1 | WP_076150488.1 hypothetical protein |
| 3(1) | WP_043069424.1 | WP_043069424.1 hypothetical protein |
| 3(1) | WP_061402732.1 | WP_061402732.1 hypothetical protein |
| 3(1) | WP_151630606.1 | WP_151630606.1 hypothetical protein |
| 3(1) | WP_025679809.1 | WP_025679809.1 hypothetical protein |
| 3(1) | WP_136604819.1 | WP_136604819.1 hypothetical protein |
| 3(1) | WP_258398817.1 | WP_258398817.1 hypothetical protein |
| 3(10) | WP_016821033.1 | WP_016821033.1 hypothetical protein |
| 3(2) | WP_212956327.1 | WP_212956327.1 hypothetical protein |
| 3(1) | WP_091155387.1 | WP_091155387.1 hypothetical protein |
| 3(1) | WP_098451706.1 | WP_098451706.1 hypothetical protein |
| 3(1) | WP_054958413.1 | WP_054958413.1 hypothetical protein |
| 3(1) | WP_137052038.1 | WP_137052038.1 hypothetical protein |
| 3(1) | WP_127812905.1 | WP_127812905.1 hypothetical protein |
| 3(1) | WP_262800526.1 | WP_262800526.1 hypothetical protein |
| 3(1) | WP_262351734.1 | WP_262351734.1 hypothetical protein |
| 3(2) | WP_015736240.1 | WP_015736240.1 hypothetical protein |
| 3(1) | WP_123067118.1 | WP_123067118.1 hypothetical protein |
| 3(1) | WP_251456530.1 | WP_251456530.1 hypothetical protein |
| 3(1) | WP_250244822.1 | WP_250244822.1 hypothetical protein |
| 3(1) | WP_216531934.1 | WP_216531934.1 hypothetical protein |
| 3(1) | WP_062326151.1 | WP_062326151.1 hypothetical protein |
| 3(1) | WP_014281186.1 | WP_014281186.1 hypothetical protein |
| 3(1) | WP_248251183.1 | WP_248251183.1 PP2C family serine/threonine-protein phosphatase |
| 3(1) | WP_103655008.1 | WP_103655008.1 hypothetical protein |
| 3(1) | WP_304142076.1 | WP_304142076.1 hypothetical protein |
| 3(1) | WP_209861637.1 | WP_209861637.1 hypothetical protein |
| 3(1) | WP_060630645.1 | WP_060630645.1 hypothetical protein |
| 3(1) | WP_270784524.1 | WP_270784524.1 hypothetical protein |
| 3(1) | WP_206094033.1 | WP_206094033.1 hypothetical protein |
| 3(1) | WP_144506828.1 | WP_144506828.1 hypothetical protein |
| 3(2) | WP_003335999.1 | WP_003335999.1 hypothetical protein |
| 3(1) | WP_097810736.1 | WP_097810736.1 hypothetical protein |
| 3(1) | WP_263299718.1 | WP_263299718.1 hypothetical protein |
| 3(1) | WP_293654341.1 | WP_293654341.1 hypothetical protein |
| 3(8) | WP_000241761.1 | WP_000241761.1 hypothetical protein |
| 3(3) | WP_095266201.1 | WP_095266201.1 hypothetical protein |
| 3(1) | WP_048552305.1 | WP_048552305.1 hypothetical protein |
| 3(1) | WP_039273537.1 | WP_039273537.1 hypothetical protein |
| 3(1) | WP_305022919.1 | WP_305022919.1 hypothetical protein |
| 3(9) | WP_000241758.1 | WP_000241758.1 hypothetical protein |
| 3(1) | WP_017149773.1 | WP_017149773.1 hypothetical protein |
| 3(1) | WP_212930977.1 | WP_212930977.1 hypothetical protein |
| 3(1) | WP_165175574.1 | WP_165175574.1 hypothetical protein |
| 3(1) | WP_286125243.1 | WP_286125243.1 hypothetical protein |
| 3(1) | WP_306007801.1 | WP_306007801.1 hypothetical protein |
| 3(1) | WP_163977053.1 | WP_163977053.1 hypothetical protein |
| 3(2) | WP_022586768.1 | WP_022586768.1 hypothetical protein |
| 3(1) | WP_098841816.1 | WP_098841816.1 hypothetical protein |
| 3(1) | WP_270723904.1 | WP_270723904.1 hypothetical protein |
| 3(1) | WP_145146965.1 | WP_145146965.1 hypothetical protein |
| 3(1) | WP_000241766.1 | WP_000241766.1 hypothetical protein |
| 3(1) | WP_123797115.1 | WP_123797115.1 hypothetical protein |
| 3(1) | WP_036707117.1 | WP_036707117.1 hypothetical protein |
| 3(1) | WP_242255496.1 | WP_242255496.1 hypothetical protein |
| 3(1) | WP_163245183.1 | WP_163245183.1 hypothetical protein |
| 3(1) | WP_213579160.1 | WP_213579160.1 hypothetical protein |
| 3(1) | WP_275441508.1 | WP_275441508.1 hypothetical protein |
| 3(1) | WP_097920539.1 | WP_097920539.1 hypothetical protein |
| 3(1) | WP_235074862.1 | WP_235074862.1 hypothetical protein |
| 3(1) | WP_223067434.1 | WP_223067434.1 hypothetical protein |
| 3(1) | WP_017689068.1 | WP_017689068.1 hypothetical protein |
| 3(3) | WP_002031625.1 | WP_002031625.1 hypothetical protein |
| 3(2) | WP_033656038.1 | WP_033656038.1 hypothetical protein |
| 3(1) | WP_251583296.1 | WP_251583296.1 hypothetical protein |
| 3(1) | WP_120112237.1 | WP_120112237.1 hypothetical protein |
| 3(1) | WP_104495354.1 | WP_104495354.1 hypothetical protein |
| 3(1) | WP_257155373.1 | WP_257155373.1 hypothetical protein |
| 3(1) | WP_013914188.1 | WP_013914188.1 hypothetical protein |
| 3(1) | WP_062239871.1 | WP_062239871.1 PP2C family serine/threonine-protein phosphatase |
| 3(1) | WP_258394502.1 | WP_258394502.1 hypothetical protein |
| 3(1) | WP_111147792.1 | WP_111147792.1 hypothetical protein |
| 3(3) | WP_103626230.1 | WP_103626230.1 hypothetical protein |
| 3(4) | WP_098255407.1 | WP_098255407.1 hypothetical protein |
| 3(4) | WP_097907629.1 | WP_097907629.1 hypothetical protein |
| 3(1) | WP_247083041.1 | WP_247083041.1 hypothetical protein |
| 3(1) | WP_053061522.1 | WP_053061522.1 hypothetical protein |
| 3(1) | WP_205001918.1 | WP_205001918.1 hypothetical protein |
| 3(1) | WP_289483783.1 | WP_289483783.1 hypothetical protein |
| 3(2) | WP_083655681.1 | WP_083655681.1 hypothetical protein |
| 3(1) | WP_000241764.1 | WP_000241764.1 hypothetical protein |
| 3(1) | WP_183600895.1 | WP_183600895.1 protein phosphatase 2C domain-containing protein |
| 3(1) | WP_133378768.1 | WP_133378768.1 hypothetical protein |
| 3(1) | WP_137062803.1 | WP_137062803.1 hypothetical protein |
| 3(1) | WP_174820808.1 | WP_174820808.1 hypothetical protein |
| 3(1) | WP_090239222.1 | WP_090239222.1 hypothetical protein |
| 3(1) | WP_268583859.1 | WP_268583859.1 hypothetical protein |
| 3(1) | WP_068940684.1 | WP_068940684.1 hypothetical protein |
| 3(2) | WP_072180907.1 | WP_072180907.1 hypothetical protein |
| 3(1) | WP_227495883.1 | WP_227495883.1 hypothetical protein |
| 3(1) | WP_151585739.1 | WP_151585739.1 hypothetical protein |
| 3(2) | WP_029438090.1 | WP_029438090.1 hypothetical protein |
| 3(1) | WP_213468111.1 | WP_213468111.1 PP2C family serine/threonine-protein phosphatase |
| 3(1) | WP_058829514.1 | WP_058829514.1 hypothetical protein |
| 3(1) | WP_098835433.1 | WP_098835433.1 hypothetical protein |
| 3(1) | WP_243500447.1 | WP_243500447.1 hypothetical protein |
| 3(1) | WP_293202259.1 | WP_293202259.1 hypothetical protein |

| | | |
|---|---|---|
| 3(1) | WP_307878871.1 | WP_307878871.1 hypothetical protein |
| 3(1) | WP_127199775.1 | WP_127199775.1 hypothetical protein |
| 3(1) | WP_232274493.1 | WP_232274493.1 hypothetical protein |
| 3(1) | WP_000241757.1 | WP_000241757.1 hypothetical protein |
| 3(3) | WP_078185406.1 | WP_078185406.1 hypothetical protein |
| 3(2) | WP_098991573.1 | WP_098991573.1 hypothetical protein |
| 3(1) | WP_124114121.1 | WP_124114121.1 hypothetical protein |
| 3(1) | WP_268623244.1 | WP_268623244.1 hypothetical protein |
| 3(2) | WP_071679489.1 | WP_071679489.1 hypothetical protein |
| 3(5) | WP_053564633.1 | WP_053564633.1 hypothetical protein |
| 3(2) | WP_087433611.1 | WP_087433611.1 hypothetical protein |
| 3(1) | WP_094936489.1 | WP_094936489.1 hypothetical protein |
| 3(1) | WP_211745826.1 | WP_211745826.1 PP2C family serine/threonine-protein phosphatase |
| 3(1) | WP_036671541.1 | WP_036671541.1 hypothetical protein |
| 3(1) | WP_018883268.1 | WP_018883268.1 hypothetical protein |
| 3(1) | WP_264453871.1 | WP_264453871.1 hypothetical protein |
| 3(3) | WP_137078367.1 | WP_137078367.1 hypothetical protein |
| 3(1) | WP_256992148.1 | WP_256992148.1 hypothetical protein |
| 3(1) | WP_249281039.1 | WP_249281039.1 protein phosphatase 2C domain-containing protein |
| 3(1) | WP_062833883.1 | WP_062833883.1 hypothetical protein |
| 3(1) | WP_270682702.1 | WP_270682702.1 hypothetical protein |
| 3(1) | WP_260865830.1 | WP_260865830.1 hypothetical protein |
| 3(1) | WP_098205152.1 | WP_098205152.1 hypothetical protein |
| 3(1) | WP_106822158.1 | WP_106822158.1 hypothetical protein |
| 3(1) | WP_153596468.1 | WP_153596468.1 hypothetical protein |
| 3(1) | WP_104030441.1 | WP_104030441.1 hypothetical protein |
| 3(1) | WP_076330725.1 | WP_076330725.1 hypothetical protein |
| 3(1) | WP_096886737.1 | WP_096886737.1 hypothetical protein |
| 3(1) | WP_090993627.1 | WP_090993627.1 hypothetical protein |
| 3(1) | WP_290401992.1 | WP_290401992.1 PP2C family serine/threonine-protein phosphatase |
| 3(5) | WP_065228769.1 | WP_065228769.1 hypothetical protein |
| 3(1) | WP_244364319.1 | WP_244364319.1 hypothetical protein |
| 3(1) | WP_244719651.1 | WP_244719651.1 hypothetical protein |
| 3(1) | WP_166704991.1 | WP_166704991.1 hypothetical protein |
| 3(1) | WP_242264960.1 | WP_242264960.1 hypothetical protein |
| 3(1) | WP_204517514.1 | WP_204517514.1 hypothetical protein |
| 3(2) | WP_255247862.1 | WP_255247862.1 protein phosphatase 2C domain-containing protein |
| 3(1) | WP_171414511.1 | WP_171414511.1 hypothetical protein |
| 3(1) | WP_070805902.1 | WP_070805902.1 hypothetical protein |
| 3(1) | WP_222661111.1 | WP_222661111.1 hypothetical protein |
| 3(1) | WP_152523858.1 | WP_152523858.1 hypothetical protein |
| 3(1) | WP_090976424.1 | WP_090976424.1 hypothetical protein |
| 3(2) | WP_063223276.1 | WP_063223276.1 hypothetical protein |
| 3(1) | WP_134909995.1 | WP_134909995.1 hypothetical protein |
| 3(1) | WP_207994163.1 | WP_207994163.1 hypothetical protein |
| 3(1) | WP_208019348.1 | WP_208019348.1 hypothetical protein |
| 3(1) | WP_215598792.1 | WP_215598792.1 hypothetical protein |
| 3(1) | WP_121474480.1 | WP_121474480.1 hypothetical protein |
| 3(1) | WP_108465352.1 | WP_108465352.1 hypothetical protein |
| 3(3) | WP_000922386.1 | WP_000922386.1 hypothetical protein |
| 3(1) | WP_094092625.1 | WP_094092625.1 hypothetical protein |
| 3(2) | WP_283655696.1 | WP_283655696.1 hypothetical protein |
| 3(1) | WP_119794367.1 | WP_119794367.1 hypothetical protein |
| 3(2) | WP_137013296.1 | WP_137013296.1 hypothetical protein |
| 3(1) | WP_002148328.1 | WP_002148328.1 hypothetical protein |
| 3(1) | WP_235286210.1 | WP_235286210.1 hypothetical protein |
| 3(1) | WP_071719985.1 | WP_071719985.1 hypothetical protein |
| 3(2) | WP_206762969.1 | WP_206762969.1 hypothetical protein |
| 3(1) | WP_138755911.1 | WP_138755911.1 hypothetical protein |
| 3(1) | WP_247902247.1 | WP_247902247.1 hypothetical protein |
| 3(1) | WP_019392113.1 | WP_019392113.1 hypothetical protein |
| 3(1) | WP_305755030.1 | WP_305755030.1 protein phosphatase 2C domain-containing protein |
| 3(5) | WP_028542687.1 | WP_028542687.1 hypothetical protein |
| 3(1) | WP_028589292.1 | WP_028589292.1 hypothetical protein |
| 3(1) | WP_119847554.1 | WP_119847554.1 hypothetical protein |
| 3(1) | WP_079346814.1 | WP_079346814.1 hypothetical protein |
| 3(1) | WP_142543124.1 | WP_142543124.1 hypothetical protein |
| 3(2) | WP_104065532.1 | WP_104065532.1 hypothetical protein |
| 3(1) | WP_236871199.1 | WP_236871199.1 hypothetical protein |
| 3(1) | WP_111619120.1 | WP_111619120.1 hypothetical protein |
| 3(1) | WP_246246256.1 | WP_246246256.1 hypothetical protein |
| 3(1) | WP_025685036.1 | WP_025685036.1 hypothetical protein |
| 3(1) | WP_254349528.1 | WP_254349528.1 hypothetical protein |
| 3(1) | WP_087982588.1 | WP_087982588.1 hypothetical protein |
| 3(1) | WP_258425842.1 | WP_258425842.1 hypothetical protein |
| 3(1) | WP_061975579.1 | WP_061975579.1 hypothetical protein |
| 3(1) | WP_258418572.1 | WP_258418572.1 hypothetical protein |
| 3(1) | WP_113753085.1 | WP_113753085.1 hypothetical protein |
| 3(1) | WP_213618818.1 | WP_213618818.1 hypothetical protein |
| 3(1) | WP_215153843.1 | WP_215153843.1 hypothetical protein |
| 3(1) | WP_043067144.1 | WP_043067144.1 hypothetical protein |
| 3(1) | WP_269055422.1 | WP_269055422.1 hypothetical protein |
| 3(1) | WP_087442417.1 | WP_087442417.1 hypothetical protein |
| 3(4) | WP_036609194.1 | WP_036609194.1 hypothetical protein |
| 3(1) | WP_098943206.1 | WP_098943206.1 hypothetical protein |
| 3(1) | WP_002080126.1 | WP_002080126.1 hypothetical protein |
| 3(1) | WP_301167497.1 | WP_301167497.1 hypothetical protein |
| 3(2) | WP_234465759.1 | WP_234465759.1 hypothetical protein |
| 3(1) | WP_225999411.1 | WP_225999411.1 hypothetical protein |
| 3(4) | WP_013370789.1 | WP_013370789.1 hypothetical protein |
| 3(1) | WP_025716879.1 | WP_025716879.1 hypothetical protein |
| 3(1) | WP_002158046.1 | WP_002158046.1 hypothetical protein |
| 3(2) | WP_076249773.1 | WP_076249773.1 hypothetical protein |
| 3(1) | WP_277480687.1 | WP_277480687.1 hypothetical protein |
| 3(1) | WP_227779378.1 | WP_227779378.1 protein phosphatase 2C domain-containing protein |
| 3(2) | WP_234489641.1 | WP_234489641.1 hypothetical protein |
| 3(1) | WP_301879143.1 | WP_301879143.1 hypothetical protein |
| 3(2) | WP_244152662.1 | WP_244152662.1 hypothetical protein |
| 3(1) | WP_257126958.1 | WP_257126958.1 hypothetical protein |
| 3(2) | WP_087956292.1 | WP_087956292.1 hypothetical protein |
| 3(1) | WP_277746356.1 | WP_277746356.1 hypothetical protein |
| 3(1) | WP_251384336.1 | WP_251384336.1 hypothetical protein |
| 3(1) | WP_197256858.1 | WP_197256858.1 hypothetical protein |
| 3(1) | WP_068496245.1 | WP_068496245.1 hypothetical protein |

| | | |
|---|---|---|
| 3(1) | WP_139170640.1 | WP_139170640.1 hypothetical protein |
| 3(1) | WP_071640584.1 | WP_071640584.1 hypothetical protein |
| 3(1) | WP_101668488.1 | WP_101668488.1 hypothetical protein |
| 3(1) | WP_265105625.1 | WP_265105625.1 hypothetical protein |
| 3(1) | WP_053783890.1 | WP_053783890.1 hypothetical protein |
| 3(1) | WP_069011423.1 | WP_069011423.1 hypothetical protein |
| 3(9) | WP_240913011.1 | WP_240913011.1 hypothetical protein |
| 3(3) | WP_048896848.1 | WP_048896848.1 hypothetical protein |
| 3(2) | WP_098148056.1 | WP_098148056.1 hypothetical protein |
| 3(1) | WP_206704280.1 | WP_206704280.1 hypothetical protein |
| 3(1) | WP_138186689.1 | WP_138186689.1 hypothetical protein |
| 3(1) | WP_018670520.1 | WP_018670520.1 hypothetical protein |
| 3(1) | WP_206701993.1 | WP_206701993.1 hypothetical protein |
| 3(1) | WP_194543496.1 | WP_194543496.1 hypothetical protein |
| 3(1) | WP_220684641.1 | WP_220684641.1 hypothetical protein |
| 3(1) | WP_191798531.1 | WP_191798531.1 hypothetical protein |
| 3(1) | WP_079694645.1 | WP_079694645.1 hypothetical protein |
| 3(1) | WP_159146503.1 | WP_159146503.1 hypothetical protein |
| 3(1) | WP_268613670.1 | WP_268613670.1 protein phosphatase 2C domain-containing protein |
| 3(1) | WP_145318686.1 | WP_145318686.1 hypothetical protein |
| 3(1) | WP_312887140.1 | WP_312887140.1 hypothetical protein |
| 3(1) | WP_100910321.1 | WP_100910321.1 hypothetical protein |
| 3(1) | WP_286017547.1 | WP_286017547.1 hypothetical protein |
| 3(1) | WP_235074860.1 | WP_235074860.1 hypothetical protein |
| 3(2) | WP_102997920.1 | WP_102997920.1 hypothetical protein |
| 3(1) | WP_246058095.1 | WP_246058095.1 protein phosphatase 2C domain-containing protein |
| 3(1) | WP_251375546.1 | WP_251375546.1 hypothetical protein |
| 3(2) | WP_076208839.1 | WP_076208839.1 hypothetical protein |
| 3(1) | WP_307413597.1 | WP_307413597.1 hypothetical protein |
| 3(1) | WP_192571319.1 | WP_192571319.1 hypothetical protein |
| 3(1) | WP_143881070.1 | WP_143881070.1 hypothetical protein |
| 3(1) | WP_074094617.1 | WP_074094617.1 hypothetical protein |
| 3(1) | WP_025364682.1 | WP_025364682.1 hypothetical protein |
| 3(1) | WP_086419834.1 | WP_086419834.1 hypothetical protein |
| 3(1) | WP_056701167.1 | WP_056701167.1 hypothetical protein |
| 3(1) | WP_147807655.1 | WP_147807655.1 hypothetical protein |
| 3(1) | WP_260986749.1 | WP_260986749.1 hypothetical protein |
| 3(1) | WP_006674857.1 | WP_006674857.1 hypothetical protein |
| 3(2) | WP_240515072.1 | WP_240515072.1 hypothetical protein |
| 3(1) | WP_165147473.1 | WP_165147473.1 hypothetical protein |
| 3(1) | WP_270716422.1 | WP_270716422.1 hypothetical protein |
| 3(1) | WP_182301828.1 | WP_182301828.1 hypothetical protein |
| 3(3) | WP_097952784.1 | WP_097952784.1 hypothetical protein |
| 3(1) | WP_111268937.1 | WP_111268937.1 protein phosphatase 2C domain-containing protein |
| 3(1) | WP_258417129.1 | WP_258417129.1 hypothetical protein |
| 3(1) | WP_109999894.1 | WP_109999894.1 hypothetical protein |
| 3(1) | WP_044649163.1 | WP_044649163.1 hypothetical protein |
| 3(1) | WP_213532660.1 | WP_213532660.1 hypothetical protein |
| 3(1) | WP_274485155.1 | WP_274485155.1 hypothetical protein |
| 3(1) | WP_206767019.1 | WP_206767019.1 hypothetical protein |
| 3(1) | WP_285748182.1 | WP_285748182.1 hypothetical protein |
| 3(1) | WP_052947760.1 | WP_052947760.1 hypothetical protein |

| | | |
|---|---|---|
| 3(1) | WP_229522853.1 | WP_229522853.1 hypothetical protein |
| 3(1) | WP_233697179.1 | WP_233697179.1 hypothetical protein |
| 3(1) | WP_098508787.1 | WP_098508787.1 hypothetical protein |
| 3(2) | WP_205828385.1 | WP_205828385.1 hypothetical protein |
| 3(1) | WP_239617467.1 | WP_239617467.1 hypothetical protein |
| 3(1) | WP_053403784.1 | WP_053403784.1 hypothetical protein |
| 3(1) | WP_206765452.1 | WP_206765452.1 hypothetical protein |
| 3(1) | WP_068694719.1 | WP_068694719.1 hypothetical protein |
| 3(1) | WP_264929204.1 | WP_264929204.1 hypothetical protein |

| | | |
|---|---|---|
| 4(1) | WP_101300019.1 | WP_101300019.1 ABC transporter ATP-binding protein |
| 4(1) | WP_293202275.1 | WP_293202275.1 ABC transporter ATP-binding protein |
| 4(1) | WP_042134260.1 | WP_042134260.1 ABC transporter ATP-binding protein |
| 4(2) | WP_095358556.1 | WP_095358556.1 ABC transporter transmembrane domain-containing protein |
| 4(1) | WP_138226036.1 | WP_138226036.1 ABC transporter ATP-binding protein |
| 4(2) | WP_023988254.1 | WP_023988254.1 ABC transporter ATP-binding protein |
| 4(1) | WP_111153291.1 | WP_111153291.1 ATP-binding cassette domain-containing protein |
| 4(1) | WP_145408784.1 | WP_145408784.1 ABC transporter ATP-binding protein |
| 4(1) | WP_290407048.1 | WP_290407048.1 ABC transporter ATP-binding protein |
| 4(1) | WP_150367647.1 | WP_150367647.1 ABC transporter transmembrane domain-containing protein |
| 4(2) | WP_145329885.1 | WP_145329885.1 ABC transporter transmembrane domain-containing protein |
| 4(1) | WP_213618803.1 | WP_213618803.1 ABC transporter ATP-binding protein |
| 4(1) | WP_291289934.1 | WP_291289934.1 ATP-binding cassette domain-containing protein |
| 4(5) | WP_145134827.1 | WP_145134827.1 ABC transporter ATP-binding protein |
| 4(1) | WP_153978060.1 | WP_153978060.1 ABC transporter transmembrane domain-containing protein |
| 4(1) | WP_123067102.1 | WP_123067102.1 ABC transporter ATP-binding protein |
| 4(1) | WP_102997913.1 | WP_102997913.1 ABC transporter transmembrane domain-containing protein |
| 4(1) | WP_166244889.1 | WP_166244889.1 glycine betaine/L-proline ABC transporter ATP-binding protein ProV |
| 4(1) | WP_250244800.1 | WP_250244800.1 ABC transporter ATP-binding protein |
| 4(1) | WP_243895990.1 | WP_243895990.1 ABC transporter ATP-binding protein |
| 4(1) | WP_036944574.1 | WP_036944574.1 ABC transporter ATP-binding protein |
| 4(2) | WP_268599737.1 | WP_268599737.1 sugar ABC transporter ATP-binding protein |
| 4(2) | WP_195951624.1 | WP_195951624.1 ABC transporter ATP-binding protein |
| 4(1) | WP_076318544.1 | WP_076318544.1 ABC transporter ATP-binding protein |
| 4(1) | WP_090761807.1 | WP_090761807.1 ABC transporter ATP-binding protein |
| 4(1) | WP_272035280.1 | WP_272035280.1 ABC transporter transmembrane domain-containing protein |
| 4(1) | WP_245339175.1 | WP_245339175.1 ABC transporter ATP-binding protein |
| 4(1) | WP_194230542.1 | WP_194230542.1 ABC transporter ATP-binding protein |
| 4(2) | WP_069291749.1 | WP_069291749.1 ABC transporter transmembrane domain-containing protein |
| 4(1) | WP_195953745.1 | WP_195953745.1 ABC transporter ATP-binding protein |
| 4(1) | WP_056700757.1 | WP_056700757.1 ABC transporter ATP-binding protein |
| 4(1) | WP_197247518.1 | WP_197247518.1 ABC transporter ATP-binding protein |
| 4(1) | WP_110822568.1 | WP_110822568.1 ABC transporter ATP-binding protein |
| 4(1) | WP_274485138.1 | WP_274485138.1 ABC transporter transmembrane domain-containing protein |
| 4(1) | WP_216975168.1 | WP_216975168.1 ABC transporter ATP-binding protein |
| 4(3) | WP_291288256.1 | WP_291288256.1 ATP-binding cassette domain-containing protein |
| 4(1) | WP_039273557.1 | WP_039273557.1 ABC transporter transmembrane domain-containing protein |
| 4(1) | WP_116231184.1 | WP_116231184.1 ABC transporter ATP-binding protein |
| 4(1) | WP_076167891.1 | WP_076167891.1 ABC transporter ATP-binding protein |

| | | |
|---|---|---|
| 4(1) | WP_013309974.1 | WP_013309974.1 ABC transporter transmembrane domain-containing protein |
| 4(1) | WP_127592438.1 | WP_127592438.1 ABC transporter ATP-binding protein |
| 4(1) | WP_090510221.1 | WP_090510221.1 sugar ABC transporter ATP-binding protein |
| 4(1) | WP_071220190.1 | WP_071220190.1 ABC transporter ATP-binding protein |
| 4(1) | WP_099853208.1 | WP_099853208.1 ABC transporter ATP-binding protein |
| 4(1) | WP_053361029.1 | WP_053361029.1 ABC transporter ATP-binding protein |
| 4(1) | WP_113052575.1 | WP_113052575.1 ABC transporter ATP-binding protein |
| 4(1) | WP_098746889.1 | WP_098746889.1 ABC transporter ATP-binding protein |
| 4(1) | WP_195953769.1 | WP_195953769.1 ABC transporter ATP-binding protein |
| 4(1) | WP_240470183.1 | WP_240470183.1 ABC transporter ATP-binding protein |
| 4(1) | WP_192510497.1 | WP_192510497.1 ABC transporter transmembrane domain-containing protein |
| 4(1) | WP_210087108.1 | WP_210087108.1 hypothetical protein |
| 4(1) | WP_145408783.1 | WP_145408783.1 ABC transporter transmembrane domain-containing protein |
| 4(1) | WP_220027673.1 | WP_220027673.1 ABC transporter ATP-binding protein |
| 4(1) | WP_301804687.1 | WP_301804687.1 ABC transporter ATP-binding protein |
| 4(1) | WP_175415415.1 | WP_175415415.1 ABC transporter ATP-binding protein |
| 4(1) | WP_077893477.1 | WP_077893477.1 ABC transporter ATP-binding protein |
| 4(1) | WP_133384353.1 | WP_133384353.1 ABC transporter ATP-binding protein |
| 4(1) | WP_145412697.1 | WP_145412697.1 ABC transporter ATP-binding protein |
| 4(1) | WP_236417195.1 | WP_236417195.1 ABC transporter transmembrane domain-containing protein |
| 4(2) | WP_043067152.1 | WP_043067152.1 ABC transporter ATP-binding protein |
| 4(1) | WP_258417133.1 | WP_258417133.1 ABC transporter ATP-binding protein |
| 4(1) | WP_235308152.1 | WP_235308152.1 ABC transporter ATP-binding protein |
| 4(1) | WP_142543117.1 | WP_142543117.1 dipeptide ABC transporter ATP-binding protein |
| 4(1) | WP_305755047.1 | WP_305755047.1 ABC transporter ATP-binding protein |
| 4(1) | WP_076264234.1 | WP_076264234.1 ABC transporter ATP-binding protein |
| 4(1) | WP_196048858.1 | WP_196048858.1 ABC transporter ATP-binding protein |
| 4(1) | WP_017689088.1 | WP_017689088.1 ABC transporter ATP-binding protein |
| 4(1) | WP_221791106.1 | WP_221791106.1 ABC transporter transmembrane domain-containing protein |
| 4(1) | WP_128101473.1 | WP_128101473.1 ABC transporter ATP-binding protein |
| 4(1) | WP_307413616.1 | WP_307413616.1 ABC transporter ATP-binding protein |
| 4(1) | WP_308721993.1 | WP_308721993.1 ABC transporter ATP-binding protein |
| 4(1) | WP_235286192.1 | WP_235286192.1 phosphonate ABC transporter ATP-binding protein |
| 4(1) | WP_107734211.1 | WP_107734211.1 ABC transporter ATP-binding protein |
| 4(1) | WP_074094607.1 | WP_074094607.1 ABC transporter transmembrane domain-containing protein |
| 4(2) | WP_042134258.1 | WP_042134258.1 ABC transporter ATP-binding protein |
| 4(1) | WP_217642381.1 | WP_217642381.1 peptidase domain-containing ABC transporter |
| 4(1) | WP_160496949.1 | WP_160496949.1 ABC transporter ATP-binding protein |
| 4(1) | WP_262351735.1 | WP_262351735.1 ABC transporter ATP-binding protein |
| 4(2) | WP_181022611.1 | WP_181022611.1 ABC transporter ATP-binding protein |
| 4(2) | WP_058710503.1 | WP_058710503.1 ABC transporter transmembrane domain-containing protein |
| 4(1) | WP_077893476.1 | WP_077893476.1 ABC transporter ATP-binding protein |
| 4(1) | WP_111268953.1 | WP_111268953.1 ABC transporter transmembrane domain-containing protein |
| 4(1) | WP_145046432.1 | WP_145046432.1 ABC transporter ATP-binding protein |
| 4(1) | WP_003335992.1 | WP_003335992.1 ABC transporter ATP-binding protein |
| 4(1) | WP_110893000.1 | WP_110893000.1 ABC transporter ATP-binding protein |
| 4(1) | WP_128101474.1 | WP_128101474.1 ABC transporter transmembrane domain-containing protein |
| 4(1) | WP_137078360.1 | WP_137078360.1 ABC transporter ATP-binding protein |
| 4(1) | WP_169505095.1 | WP_169505095.1 ABC transporter ATP-binding protein |
| 4(1) | WP_179160635.1 | WP_179160635.1 ABC transporter ATP-binding protein |
| 4(1) | WP_211745839.1 | WP_211745839.1 ATP-binding cassette domain-containing protein |
| 4(1) | WP_278773479.1 | WP_278773479.1 ABC transporter ATP-binding protein |
| 4(1) | WP_213532647.1 | WP_213532647.1 ABC transporter ATP-binding protein |
| 4(1) | WP_102997912.1 | WP_102997912.1 ABC transporter ATP-binding protein |
| 4(1) | WP_254068636.1 | WP_254068636.1 ABC transporter ATP-binding protein |
| 4(1) | WP_302579550.1 | WP_302579550.1 ABC transporter ATP-binding protein |
| 4(1) | WP_212949020.1 | WP_212949020.1 ABC transporter ATP-binding protein |
| 4(1) | WP_095398788.1 | WP_095398788.1 ABC transporter ATP-binding protein |
| 4(1) | WP_025685021.1 | WP_025685021.1 ABC transporter ATP-binding protein |
| 4(1) | WP_076156862.1 | WP_076156862.1 ABC transporter ATP-binding protein |
| 4(1) | WP_094155416.1 | WP_094155416.1 ABC transporter ATP-binding protein |
| 4(1) | WP_091017210.1 | WP_091017210.1 ABC transporter ATP-binding protein |
| 4(1) | WP_285079448.1 | WP_285079448.1 ABC transporter transmembrane domain-containing protein |
| 4(1) | WP_045243839.1 | WP_045243839.1 ABC transporter ATP-binding protein |
| 4(1) | WP_090993599.1 | WP_090993599.1 ABC transporter transmembrane domain-containing protein |
| 4(1) | WP_137062789.1 | WP_137062789.1 ABC transporter transmembrane domain-containing protein |
| 4(1) | WP_269320832.1 | WP_269320832.1 ABC transporter transmembrane domain-containing protein, partial |
| 4(1) | WP_145046434.1 | WP_145046434.1 ABC transporter transmembrane domain-containing protein |
| 4(1) | WP_164965499.1 | WP_164965499.1 ABC transporter ATP-binding protein |
| 4(1) | WP_239586182.1 | WP_239586182.1 ABC transporter ATP-binding protein |
| 4(1) | WP_061829412.1 | WP_061829412.1 ABC transporter ATP-binding protein |
| 4(1) | WP_058829506.1 | WP_058829506.1 ABC transporter ATP-binding protein |
| 4(1) | WP_231929221.1 | WP_231929221.1 ABC transporter ATP-binding protein |
| 4(1) | WP_307214298.1 | WP_307214298.1 ABC transporter transmembrane domain-containing protein |
| 4(1) | WP_254325799.1 | WP_254325799.1 ABC transporter ATP-binding protein |
| 4(1) | WP_290407049.1 | WP_290407049.1 ABC transporter ATP-binding protein |
| 4(1) | WP_255253070.1 | WP_255253070.1 ABC transporter ATP-binding protein |
| 4(1) | WP_233697186.1 | WP_233697186.1 ABC transporter ATP-binding protein |
| 4(1) | WP_040101530.1 | WP_040101530.1 ABC transporter transmembrane domain-containing protein |
| 4(1) | WP_142612995.1 | WP_142612995.1 ABC transporter ATP-binding protein |
| 4(1) | WP_285228770.1 | WP_285228770.1 ABC transporter transmembrane domain-containing protein |
| 4(1) | WP_152381387.1 | WP_152381387.1 ABC transporter transmembrane domain-containing protein |
| 4(1) | WP_308615626.1 | WP_308615626.1 ATP-binding cassette domain-containing protein |
| 4(1) | WP_090887141.1 | WP_090887141.1 ABC transporter ATP-binding protein |
| 4(1) | WP_302153066.1 | WP_302153066.1 ABC transporter ATP-binding protein |
| 4(1) | WP_160037471.1 | WP_160037471.1 ABC transporter transmembrane domain-containing protein |
| 4(1) | WP_094870390.1 | WP_094870390.1 ABC transporter ATP-binding protein |
| 4(1) | WP_010346297.1 | WP_010346297.1 ABC transporter transmembrane domain-containing protein |
| 4(1) | WP_307349944.1 | WP_307349944.1 ABC transporter transmembrane domain-containing protein |
| 4(1) | WP_164965498.1 | WP_164965498.1 ABC transporter transmembrane domain-containing protein |
| 4(1) | WP_305166271.1 | WP_305166271.1 ABC transporter ATP-binding protein |
| 4(1) | WP_076289564.1 | WP_076289564.1 ABC transporter transmembrane domain-containing protein |
| 4(1) | WP_169505094.1 | WP_169505094.1 ABC transporter ATP-binding protein |
| 4(1) | WP_190299132.1 | WP_190299132.1 ABC transporter ATP-binding protein |
| 4(1) | WP_077832271.1 | WP_077832271.1 ABC transporter ATP-binding protein |
| 4(1) | WP_129549126.1 | WP_129549126.1 ABC transporter transmembrane domain-containing protein |
| 4(2) | WP_269468913.1 | WP_269468913.1 ABC transporter ATP-binding protein |
| 4(1) | WP_016821020.1 | WP_016821020.1 ABC transporter ATP-binding protein |
| 4(1) | WP_068496263.1 | WP_068496263.1 ABC transporter transmembrane domain-containing protein |

| | | |
|---|---|---|
| 4(1) | WP_165079988.1 | WP_165079988.1 ABC transporter transmembrane domain-containing protein |
| 4(1) | WP_297183831.1 | WP_297183831.1 ATP-binding cassette domain-containing protein |
| 4(2) | WP_241752093.1 | WP_241752093.1 sugar ABC transporter ATP-binding protein |
| 4(1) | WP_042210985.1 | WP_042210985.1 ABC transporter ATP-binding protein |
| 4(1) | WP_289392064.1 | WP_289392064.1 ABC transporter ATP-binding protein |
| 4(1) | WP_232699391.1 | WP_232699391.1 ABC transporter ATP-binding protein |
| 4(1) | WP_007430119.1 | WP_007430119.1 ABC transporter transmembrane domain-containing protein |
| 4(1) | WP_028533188.1 | WP_028533188.1 sugar ABC transporter ATP-binding protein |
| 4(1) | WP_062833870.1 | WP_062833870.1 ABC transporter ATP-binding protein |
| 4(2) | WP_076292676.1 | WP_076292676.1 ABC transporter ATP-binding protein |
| 4(1) | WP_036609222.1 | WP_036609222.1 ABC transporter ATP-binding protein |
| 4(1) | WP_196048902.1 | WP_196048902.1 ABC transporter ATP-binding protein |
| 4(1) | WP_308110123.1 | WP_308110123.1 ABC transporter ATP-binding protein |
| 4(1) | WP_212980230.1 | WP_212980230.1 ABC transporter transmembrane domain-containing protein |
| 4(1) | WP_047842530.1 | WP_047842530.1 ABC transporter ATP-binding protein |
| 4(1) | WP_145146959.1 | WP_145146959.1 ABC transporter ATP-binding protein |
| 4(1) | WP_206767020.1 | WP_206767020.1 ABC transporter ATP-binding protein |
| 4(1) | WP_160496948.1 | WP_160496948.1 ABC transporter transmembrane domain-containing protein |
| 4(1) | WP_013370770.1 | WP_013370770.1 ABC transporter ATP-binding protein |
| 4(1) | WP_302153072.1 | WP_302153072.1 hypothetical protein |
| 4(1) | WP_306077379.1 | WP_306077379.1 ABC transporter ATP-binding protein |
| 4(1) | WP_136604804.1 | WP_136604804.1 ABC transporter ATP-binding protein |
| 4(1) | WP_301249687.1 | WP_301249687.1 ABC transporter ATP-binding protein |
| 4(1) | WP_307537599.1 | WP_307537599.1 ABC transporter ATP-binding protein |
| 4(5) | WP_089546596.1 | WP_089546596.1 ABC transporter ATP-binding protein |
| 4(5) | WP_072732433.1 | WP_072732433.1 ABC transporter transmembrane domain-containing protein |
| 4(1) | WP_211022800.1 | WP_211022800.1 ABC transporter ATP-binding protein |
| 4(1) | WP_065291259.1 | WP_065291259.1 sugar ABC transporter ATP-binding protein |
| 4(1) | WP_056700759.1 | WP_056700759.1 ABC transporter transmembrane domain-containing protein |
| 4(1) | WP_305755046.1 | WP_305755046.1 ABC transporter transmembrane domain-containing protein |
| 4(1) | WP_229522867.1 | WP_229522867.1 ABC transporter transmembrane domain-containing protein |
| 4(1) | WP_152392566.1 | WP_152392566.1 ABC transporter transmembrane domain-containing protein |
| 4(1) | WP_244719565.1 | WP_244719565.1 ABC transporter ATP-binding protein |
| 4(2) | WP_145037676.1 | WP_145037676.1 ABC transporter ATP-binding protein |
| 4(1) | WP_244965663.1 | WP_244965663.1 sugar ABC transporter ATP-binding protein |
| 4(1) | WP_071640569.1 | WP_071640569.1 ABC transporter ATP-binding protein |
| 4(12) | WP_023988255.1 | WP_023988255.1 ABC transporter transmembrane domain-containing protein |
| 4(1) | WP_054958420.1 | WP_054958420.1 ABC transporter ATP-binding protein |
| 4(1) | WP_076264233.1 | WP_076264233.1 ABC transporter transmembrane domain-containing protein |
| 4(1) | WP_264929217.1 | WP_264929217.1 ABC transporter ATP-binding protein/permease |
| 4(1) | WP_053492725.1 | WP_053492725.1 ABC transporter ATP-binding protein |
| 4(1) | WP_197256842.1 | WP_197256842.1 ATP-binding cassette domain-containing protein |
| 4(1) | WP_242858509.1 | WP_242858509.1 ABC transporter ATP-binding protein |
| 4(1) | WP_076208847.1 | WP_076208847.1 ABC transporter transmembrane domain-containing protein |
| 4(1) | WP_258418569.1 | WP_258418569.1 ABC transporter ATP-binding protein |
| 4(1) | WP_308116289.1 | WP_308116289.1 ABC transporter ATP-binding protein |
| 4(1) | WP_232243183.1 | WP_232243183.1 ABC transporter ATP-binding protein |
| 4(1) | WP_017689087.1 | WP_017689087.1 ABC transporter transmembrane domain-containing protein |
| 4(1) | WP_069011419.1 | WP_069011419.1 ABC transporter ATP-binding protein |
| 4(1) | WP_241929672.1 | WP_241929672.1 ABC transporter ATP-binding protein |
| 4(1) | WP_095287673.1 | WP_095287673.1 ABC transporter transmembrane domain-containing protein |
| 4(1) | WP_123067103.1 | WP_123067103.1 ABC transporter transmembrane domain-containing protein |
| 4(1) | WP_018883290.1 | WP_018883290.1 ABC transporter ATP-binding protein |
| 4(1) | WP_268623236.1 | WP_268623236.1 sugar ABC transporter ATP-binding protein |
| 4(1) | WP_200633293.1 | WP_200633293.1 ABC transporter ATP-binding protein |
| 4(1) | WP_054972083.1 | WP_054972083.1 amino acid ABC transporter ATP-binding protein |
| 4(2) | WP_179603469.1 | WP_179603469.1 ABC transporter ATP-binding protein |
| 4(1) | WP_288218592.1 | WP_288218592.1 ABC transporter ATP-binding protein |
| 4(1) | WP_018883285.1 | WP_018883285.1 ABC transporter ATP-binding protein |
| 4(1) | WP_091017208.1 | WP_091017208.1 ABC transporter transmembrane domain-containing protein |
| 4(2) | WP_262866582.1 | WP_262866582.1 sugar ABC transporter ATP-binding protein |
| 4(2) | WP_109999882.1 | WP_109999882.1 ABC transporter transmembrane domain-containing protein |
| 4(1) | WP_109999881.1 | WP_109999881.1 ABC transporter ATP-binding protein |
| 4(1) | WP_288218589.1 | WP_288218589.1 ABC transporter ATP-binding protein |
| 4(1) | WP_290409170.1 | WP_290409170.1 ABC transporter ATP-binding protein |
| 4(1) | WP_221697951.1 | WP_221697951.1 ABC transporter ATP-binding protein |
| 4(1) | WP_251548746.1 | WP_251548746.1 ABC transporter ATP-binding protein |
| 4(1) | WP_231417848.1 | WP_231417848.1 dipeptide ABC transporter ATP-binding protein |
| 4(1) | WP_165175591.1 | WP_165175591.1 ABC transporter ATP-binding protein |
| 4(1) | WP_103047833.1 | WP_103047833.1 ABC transporter ATP-binding protein |
| 4(1) | WP_024630312.1 | WP_024630312.1 ABC transporter transmembrane domain-containing protein |
| 4(1) | WP_028589305.1 | WP_028589305.1 ABC transporter ATP-binding protein |
| 4(1) | WP_220027674.1 | WP_220027674.1 ABC transporter transmembrane domain-containing protein |
| 4(1) | WP_068496264.1 | WP_068496264.1 ABC transporter ATP-binding protein |
| 4(1) | WP_244719546.1 | WP_244719546.1 ABC transporter ATP-binding protein |
| 4(1) | WP_285079449.1 | WP_285079449.1 ABC transporter ATP-binding protein |
| 4(1) | WP_076208848.1 | WP_076208848.1 ABC transporter ATP-binding protein |
| 4(1) | WP_291288272.1 | WP_291288272.1 ATP-binding cassette domain-containing protein, partial |
| 4(1) | WP_307199950.1 | WP_307199950.1 ABC transporter ATP-binding protein |
| 4(1) | WP_213650777.1 | WP_213650777.1 ABC transporter transmembrane domain-containing protein |
| 4(1) | WP_134910017.1 | WP_134910017.1 ABC transporter ATP-binding protein |
| 4(1) | WP_237690857.1 | WP_237690857.1 ABC transporter ATP-binding protein |
| 4(1) | WP_149871693.1 | WP_149871693.1 ABC transporter ATP-binding protein |
| 4(1) | WP_071640571.1 | WP_071640571.1 ABC transporter transmembrane domain-containing protein |
| 4(1) | WP_090921082.1 | WP_090921082.1 ABC transporter transmembrane domain-containing protein |
| 4(2) | WP_017427962.1 | WP_017427962.1 ABC transporter transmembrane domain-containing protein |
| 4(1) | WP_136606868.1 | WP_136606868.1 ABC transporter transmembrane domain-containing protein |
| 4(1) | WP_221697954.1 | WP_221697954.1 ABC transporter ATP-binding protein |
| 4(1) | WP_024630311.1 | WP_024630311.1 ABC transporter ATP-binding protein |
| 4(1) | WP_096774452.1 | WP_096774452.1 ABC transporter ATP-binding protein |
| 4(1) | WP_127543710.1 | WP_127543710.1 ABC transporter ATP-binding protein |
| 4(1) | WP_066022464.1 | WP_066022464.1 ABC transporter ATP-binding protein |
| 4(1) | WP_142543137.1 | WP_142543137.1 sugar ABC transporter ATP-binding protein |
| 4(2) | WP_068940671.1 | WP_068940671.1 ABC transporter ATP-binding protein |
| 4(1) | WP_225999397.1 | WP_225999397.1 ABC transporter transmembrane domain-containing protein |

4(1) WP_165147491.1 WP_165147491.1 ABC transporter transmembrane domain-containing protein
4(1) WP_208918995.1 WP_208918995.1 ABC transporter ATP-binding protein
4(2) WP_090896164.1 WP_090896164.1 ABC transporter ATP-binding protein
4(1) WP_021261896.1 WP_021261896.1 sugar ABC transporter ATP-binding protein
4(1) WP_111147811.1 WP_111147811.1 phosphonate ABC transporter ATP-binding protein
4(1) WP_258433022.1 WP_258433022.1 ABC transporter ATP-binding protein
4(3) WP_176452351.1 WP_176452351.1 ABC transporter ATP-binding protein
4(1) WP_289373822.1 WP_289373822.1 ABC transporter ATP-binding protein
4(2) WP_044355254.1 WP_044355254.1 sugar ABC transporter ATP-binding protein
4(1) WP_152392567.1 WP_152392567.1 ABC transporter ATP-binding protein/permease
4(1) WP_034675338.1 WP_034675338.1 ABC transporter ATP-binding protein
4(1) WP_015736238.1 WP_015736238.1 ABC transporter ATP-binding protein
4(1) WP_070638623.1 WP_070638623.1 ABC transporter ATP-binding protein
4(1) WP_134910015.1 WP_134910015.1 ABC transporter transmembrane domain-containing protein
4(1) WP_225999396.1 WP_225999396.1 ABC transporter ATP-binding protein
4(1) WP_076322126.1 WP_076322126.1 ABC transporter ATP-binding protein
4(1) WP_165079989.1 WP_165079989.1 ABC transporter ATP-binding protein
4(1) WP_110892998.1 WP_110892998.1 ABC transporter transmembrane domain-containing protein
4(1) WP_120462964.1 WP_120462964.1 ATP-binding cassette domain-containing protein
4(1) WP_212980193.1 WP_212980193.1 ABC transporter ATP-binding protein
4(1) WP_039273560.1 WP_039273560.1 ABC transporter ATP-binding protein
4(1) WP_258296285.1 WP_258296285.1 ABC transporter transmembrane domain-containing protein
4(1) WP_036661601.1 WP_036661601.1 ABC transporter ATP-binding protein
4(1) WP_036707086.1 WP_036707086.1 ABC transporter ATP-binding protein
4(2) WP_070098862.1 WP_070098862.1 ABC transporter ATP-binding protein
4(2) WP_138226035.1 WP_138226035.1 ABC transporter ATP-binding protein
4(1) WP_301249688.1 WP_301249688.1 ABC transporter transmembrane domain-containing protein
4(1) WP_307537597.1 WP_307537597.1 ABC transporter transmembrane domain-containing protein
4(1) WP_013309973.1 WP_013309973.1 ABC transporter ATP-binding protein
4(1) WP_206763598.1 WP_206763598.1 ABC transporter ATP-binding protein
4(1) WP_040101531.1 WP_040101531.1 ABC transporter ATP-binding protein
4(1) WP_209861663.1 WP_209861663.1 ABC transporter ATP-binding protein
4(1) WP_244850161.1 WP_244850161.1 ABC transporter ATP-binding protein
4(1) WP_031463342.1 WP_031463342.1 ABC transporter ATP-binding protein
4(1) WP_210087128.1 WP_210087128.1 ABC transporter ATP-binding protein
4(1) WP_013370771.1 WP_013370771.1 ABC transporter transmembrane domain-containing protein
4(1) WP_022586762.1 WP_022586762.1 ABC transporter ATP-binding protein
4(1) WP_213649983.1 WP_213649983.1 ABC transporter ATP-binding protein
4(1) WP_254349521.1 WP_254349521.1 ABC transporter ATP-binding protein
4(1) WP_006290906.1 WP_006290906.1 ABC-F family ATP-binding cassette domain-containing protein
4(1) WP_183578418.1 WP_183578418.1 ABC transporter ATP-binding protein
4(1) WP_129549125.1 WP_129549125.1 ABC transporter ATP-binding protein
4(1) WP_101356563.1 WP_101356563.1 ABC-F family ATP-binding cassette domain-containing protein
4(1) WP_039871628.1 WP_039871628.1 ABC transporter ATP-binding protein
4(2) WP_090896166.1 WP_090896166.1 ABC transporter transmembrane domain-containing protein
4(1) WP_073008283.1 WP_073008283.1 ABC transporter ATP-binding protein
4(1) WP_035311915.1 WP_035311915.1 ABC transporter ATP-binding protein
4(1) WP_076330713.1 WP_076330713.1 ABC transporter ATP-binding protein
4(3) WP_168178775.1 WP_168178775.1 ATP-binding cassette domain-containing protein
4(1) WP_104495327.1 WP_104495327.1 ABC transporter transmembrane domain-containing protein
4(2) WP_021624833.1 WP_021624833.1 ABC transporter ATP-binding protein
4(1) WP_238333012.1 WP_238333012.1 ABC transporter ATP-binding protein
4(1) WP_113052574.1 WP_113052574.1 ABC transporter transmembrane domain-containing protein
4(1) WP_169505045.1 WP_169505045.1 ABC transporter ATP-binding protein
4(2) WP_283655685.1 WP_283655685.1 ABC transporter transmembrane domain-containing protein
4(1) WP_293202274.1 WP_293202274.1 ABC transporter transmembrane domain-containing protein
4(3) WP_025719077.1 WP_025719077.1 ABC transporter transmembrane domain-containing protein
4(1) WP_094593865.1 WP_094593865.1 ABC transporter ATP-binding protein
4(1) WP_028405563.1 WP_028405563.1 ABC transporter ATP-binding protein
4(1) WP_113058705.1 WP_113058705.1 ABC transporter ATP-binding protein
4(1) WP_190299133.1 WP_190299133.1 ABC transporter transmembrane domain-containing protein
4(1) WP_016821021.1 WP_016821021.1 ABC transporter transmembrane domain-containing protein
4(1) WP_195949157.1 WP_195949157.1 ABC transporter ATP-binding protein
4(1) WP_138976937.1 WP_138976937.1 ABC transporter ATP-binding protein
4(1) WP_066020851.1 WP_066020851.1 peptidase domain-containing ABC transporter
4(1) WP_236871198.1 WP_236871198.1 ABC transporter ATP-binding protein
4(1) WP_028589301.1 WP_028589301.1 ABC transporter ATP-binding protein
4(1) WP_120462937.1 WP_120462937.1 amino acid ABC transporter ATP-binding protein
4(1) WP_209941805.1 WP_209941805.1 ABC transporter ATP-binding protein
4(1) WP_195949159.1 WP_195949159.1 ABC transporter ATP-binding protein
4(1) WP_247902233.1 WP_247902233.1 ABC transporter ATP-binding protein
4(1) WP_124114112.1 WP_124114112.1 ABC transporter transmembrane domain-containing protein
4(1) WP_077567034.1 WP_077567034.1 ABC transporter ATP-binding protein
4(8) WP_036609220.1 WP_036609220.1 ABC transporter transmembrane domain-containing protein
4(1) WP_072732432.1 WP_072732432.1 ABC transporter ATP-binding protein
4(1) WP_213426980.1 WP_213426980.1 ATP-binding cassette domain-containing protein
4(1) WP_014281171.1 WP_014281171.1 ABC transporter transmembrane domain-containing protein
4(1) WP_137062788.1 WP_137062788.1 ABC transporter ATP-binding protein
4(1) WP_227793448.1 WP_227793448.1 ABC-F type ribosomal protection protein
4(1) WP_077832270.1 WP_077832270.1 ABC transporter ATP-binding protein
4(2) WP_063566335.1 WP_063566335.1 ABC transporter transmembrane domain-containing protein
4(1) WP_042210986.1 WP_042210986.1 ABC transporter ATP-binding protein
4(1) WP_058710504.1 WP_058710504.1 ABC transporter ATP-binding protein
4(1) WP_171414505.1 WP_171414505.1 dipeptide ABC transporter ATP-binding protein
4(1) WP_108465335.1 WP_108465335.1 ABC transporter ATP-binding protein
4(1) WP_196426423.1 WP_196426423.1 ABC transporter ATP-binding protein
4(6) WP_042400984.1 WP_042400984.1 ABC transporter ATP-binding protein
4(1) WP_160496944.1 WP_160496944.1 ABC transporter ATP-binding protein
4(1) WP_305022937.1 WP_305022937.1 ABC transporter transmembrane domain-containing protein
4(1) WP_094155415.1 WP_094155415.1 ABC transporter transmembrane domain-containing protein
4(1) WP_311200612.1 WP_311200612.1 ABC transporter ATP-binding protein
4(1) WP_025679793.1 WP_025679793.1 ABC transporter transmembrane domain-containing protein
4(1) WP_250251634.1 WP_250251634.1 ABC transporter ATP-binding protein
4(1) WP_076292677.1 WP_076292677.1 ABC transporter transmembrane domain-containing protein
4(1) WP_185165656.1 WP_185165656.1 ABC transporter ATP-binding protein
4(1) WP_025685022.1 WP_025685022.1 ABC transporter transmembrane domain-containing protein
4(1) WP_169505038.1 WP_169505038.1 ABC transporter ATP-binding protein

| | | |
|---|---|---|
| 4(1) | WP_063208806.1 | WP_063208806.1 ABC transporter transmembrane domain-containing protein |
| 4(1) | WP_062833871.1 | WP_062833871.1 ABC transporter transmembrane domain-containing protein |
| 4(1) | WP_064639019.1 | WP_064639019.1 ABC transporter ATP-binding protein |
| 4(1) | WP_165147494.1 | WP_165147494.1 ABC transporter ATP-binding protein |
| 4(1) | WP_169478428.1 | WP_169478428.1 ABC transporter transmembrane domain-containing protein |
| 4(1) | WP_025364675.1 | WP_025364675.1 ABC transporter ATP-binding protein |
| 4(1) | WP_005546532.1 | WP_005546532.1 dipeptide ABC transporter ATP-binding protein |
| 4(1) | WP_104065538.1 | WP_104065538.1 ABC transporter ATP-binding protein |
| 4(1) | WP_007430118.1 | WP_007430118.1 ABC transporter ATP-binding protein |
| 4(1) | WP_095265649.1 | WP_095265649.1 ABC transporter transmembrane domain-containing protein |
| 4(1) | WP_194230417.1 | WP_194230417.1 ABC transporter ATP-binding protein |
| 4(1) | WP_305022938.1 | WP_305022938.1 ABC transporter ATP-binding protein |
| 4(1) | WP_277883669.1 | WP_277883669.1 ABC transporter ATP-binding protein |
| 4(1) | WP_216791192.1 | WP_216791192.1 amino acid ABC transporter ATP-binding protein |
| 4(1) | WP_110822567.1 | WP_110822567.1 ABC transporter transmembrane domain-containing protein |
| 4(1) | WP_036670427.1 | WP_036670427.1 ABC transporter ATP-binding protein |
| 4(1) | WP_155703949.1 | WP_155703949.1 ABC transporter ATP-binding protein |
| 4(1) | WP_247681909.1 | WP_247681909.1 ATP-binding cassette domain-containing protein |
| 4(1) | WP_096774464.1 | WP_096774464.1 ABC transporter ATP-binding protein |
| 4(12) | WP_025364676.1 | WP_025364676.1 ABC transporter transmembrane domain-containing protein |
| 4(1) | WP_068619138.1 | WP_068619138.1 ABC transporter ATP-binding protein |
| 4(1) | WP_192266429.1 | WP_192266429.1 ABC transporter transmembrane domain-containing protein |
| 4(1) | WP_254660713.1 | WP_254660713.1 ABC transporter ATP-binding protein |
| 4(1) | WP_009590673.1 | WP_009590673.1 ABC transporter ATP-binding protein |
| 4(1) | WP_191798517.1 | WP_191798517.1 ABC transporter ATP-binding protein |
| 4(1) | WP_039871627.1 | WP_039871627.1 ABC transporter ATP-binding protein |
| 4(1) | WP_106765971.1 | WP_106765971.1 ABC transporter ATP-binding protein |
| 4(1) | WP_181603048.1 | WP_181603048.1 ABC transporter ATP-binding protein |
| 4(1) | WP_155703948.1 | WP_155703948.1 ABC transporter ATP-binding protein |
| 4(1) | WP_105368169.1 | WP_105368169.1 ABC transporter ATP-binding protein |
| 4(1) | WP_108465353.1 | WP_108465353.1 ABC transporter ATP-binding protein |
| 4(1) | WP_136604800.1 | WP_136604800.1 ABC transporter ATP-binding protein |
| 4(1) | WP_153978059.1 | WP_153978059.1 ABC transporter ATP-binding protein |
| 4(1) | WP_105368170.1 | WP_105368170.1 ABC transporter ATP-binding protein |
| 4(1) | WP_213426979.1 | WP_213426979.1 ABC transporter ATP-binding protein |
| 4(1) | WP_241734701.1 | WP_241734701.1 sugar ABC transporter ATP-binding protein |
| 4(1) | WP_064639018.1 | WP_064639018.1 ABC transporter transmembrane domain-containing protein |
| 4(2) | WP_076249785.1 | WP_076249785.1 ABC transporter ATP-binding protein |
| 4(1) | WP_181747923.1 | WP_181747923.1 ABC transporter ATP-binding protein |
| 4(1) | WP_233882484.1 | WP_233882484.1 ABC transporter ATP-binding protein |
| 4(1) | WP_062326134.1 | WP_062326134.1 ABC transporter ATP-binding protein |
| 4(1) | WP_280530303.1 | WP_280530303.1 ABC transporter ATP-binding protein |
| 4(1) | WP_094870389.1 | WP_094870389.1 ABC transporter ATP-binding protein |
| 4(1) | WP_160037470.1 | WP_160037470.1 ABC transporter ATP-binding protein |
| 4(1) | WP_247902234.1 | WP_247902234.1 ABC transporter transmembrane domain-containing protein |
| 4(1) | WP_074108306.1 | WP_074108306.1 ABC transporter ATP-binding protein |
| 4(1) | WP_042462832.1 | WP_042462832.1 ABC transporter ATP-binding protein |
| 4(1) | WP_074085371.1 | WP_074085371.1 ABC transporter ATP-binding protein |
| 4(1) | WP_076330714.1 | WP_076330714.1 ABC transporter transmembrane domain-containing protein |
| 4(1) | WP_213649979.1 | WP_213649979.1 ABC transporter ATP-binding protein |
| 4(1) | WP_007147972.1 | WP_007147972.1 ABC-F type ribosomal protection protein |
| 4(1) | WP_018883289.1 | WP_018883289.1 ABC transporter transmembrane domain-containing protein |
| 4(1) | WP_192510498.1 | WP_192510498.1 ABC transporter ATP-binding protein |
| 4(1) | WP_277548432.1 | WP_277548432.1 ABC transporter ATP-binding protein |
| 4(1) | WP_160037475.1 | WP_160037475.1 ABC transporter ATP-binding protein |
| 4(1) | WP_212949022.1 | WP_212949022.1 ABC transporter transmembrane domain-containing protein |
| 4(1) | WP_206762968.1 | WP_206762968.1 ABC transporter ATP-binding protein |
| 4(1) | WP_307589585.1 | WP_307589585.1 methionine ABC transporter ATP-binding protein |
| 4(1) | WP_258398816.1 | WP_258398816.1 ABC transporter ATP-binding protein |
| 4(1) | WP_069011420.1 | WP_069011420.1 ABC transporter transmembrane domain-containing protein |
| 4(1) | WP_179283615.1 | WP_179283615.1 ABC transporter ATP-binding protein |
| 4(1) | WP_229522868.1 | WP_229522868.1 ABC transporter ATP-binding protein |
| 4(1) | WP_199910013.1 | WP_199910013.1 ABC transporter ATP-binding protein |
| 4(1) | WP_034675396.1 | WP_034675396.1 amino acid ABC transporter ATP-binding protein |
| 4(1) | WP_150367648.1 | WP_150367648.1 ABC transporter ATP-binding protein |
| 4(1) | WP_104495325.1 | WP_104495325.1 ABC transporter ATP-binding protein |
| 4(1) | WP_055277406.1 | WP_055277406.1 ABC transporter ATP-binding protein |
| 4(1) | WP_280788639.1 | WP_280788639.1 ABC transporter ATP-binding protein |
| 4(1) | WP_042462872.1 | WP_042462872.1 ABC transporter ATP-binding protein |
| 4(1) | WP_275441509.1 | WP_275441509.1 ABC transporter ATP-binding protein |
| 4(1) | WP_290403366.1 | WP_290403366.1 ABC transporter ATP-binding protein |
| 4(1) | WP_263560233.1 | WP_263560233.1 ABC transporter ATP-binding protein |
| 4(3) | WP_025848587.1 | WP_025848587.1 amino acid ABC transporter ATP-binding protein |
| 4(1) | WP_279591710.1 | WP_279591710.1 ABC transporter ATP-binding protein |
| 4(1) | WP_244159321.1 | WP_244159321.1 ABC transporter ATP-binding protein |
| 4(1) | WP_127199757.1 | WP_127199757.1 ABC transporter ATP-binding protein |
| 4(1) | WP_062326135.1 | WP_062326135.1 ABC transporter transmembrane domain-containing protein |
| 4(1) | WP_025679791.1 | WP_025679791.1 ABC transporter ATP-binding protein |
| 4(1) | WP_028589304.1 | WP_028589304.1 ABC transporter transmembrane domain-containing protein |
| 4(1) | WP_216975169.1 | WP_216975169.1 ABC transporter transmembrane domain-containing protein |
| 4(1) | WP_108465336.1 | WP_108465336.1 ABC transporter transmembrane domain-containing protein |
| 4(1) | WP_211022801.1 | WP_211022801.1 ABC transporter ATP-binding protein |
| 4(1) | WP_221697953.1 | WP_221697953.1 ABC transporter ATP-binding protein |
| 4(1) | WP_133384354.1 | WP_133384354.1 ABC transporter transmembrane domain-containing protein |
| 4(3) | WP_165175590.1 | WP_165175590.1 ABC transporter transmembrane domain-containing protein |
| 4(2) | WP_053783876.1 | WP_053783876.1 ABC transporter transmembrane domain-containing protein |
| 4(1) | WP_136379711.1 | WP_136379711.1 cell division ATP-binding protein FtsE |
| 4(1) | WP_268619942.1 | WP_268619942.1 dipeptide ABC transporter ATP-binding protein |
| 4(1) | WP_124114111.1 | WP_124114111.1 ABC transporter ATP-binding protein |
| 4(1) | WP_212980192.1 | WP_212980192.1 ABC transporter ATP-binding protein |
| 4(1) | WP_074094606.1 | WP_074094606.1 ABC transporter ATP-binding protein |
| 4(1) | WP_229139931.1 | WP_229139931.1 cell division ATP-binding protein FtsE |
| 4(2) | WP_025719079.1 | WP_025719079.1 ABC transporter ATP-binding protein |
| 4(1) | WP_231094838.1 | WP_231094838.1 ABC transporter ATP-binding protein |
| 4(1) | WP_010496599.1 | WP_010496599.1 amino acid ABC transporter ATP-binding protein |
| 4(1) | WP_285748164.1 | WP_285748164.1 ABC transporter ATP-binding protein |
| 4(1) | WP_188593155.1 | WP_188593155.1 ABC transporter ATP-binding protein |
| 4(1) | WP_090908267.1 | WP_090908267.1 ABC transporter ATP-binding protein |

| | | |
|---|---|---|
| 4(1) | WP_276575510.1 | WP_276575510.1 ABC transporter ATP-binding protein |
| 4(1) | WP_006289735.1 | WP_006289735.1 ABC-F family ATP-binding cassette domain-containing protein |
| 4(1) | WP_305166270.1 | WP_305166270.1 ABC transporter transmembrane domain-containing protein |
| 4(1) | WP_019424133.1 | WP_019424133.1 ABC transporter ATP-binding protein |
| 4(1) | WP_119847541.1 | WP_119847541.1 ABC transporter ATP-binding protein |
| 4(1) | WP_195951786.1 | WP_195951786.1 ABC transporter ATP-binding protein |
| 4(1) | WP_179160629.1 | WP_179160629.1 amino acid ABC transporter ATP-binding protein |
| 4(1) | WP_133378761.1 | WP_133378761.1 ATP-binding cassette domain-containing protein |
| 4(1) | WP_100537987.1 | WP_100537987.1 ABC transporter ATP-binding protein |
| 4(1) | WP_058829507.1 | WP_058829507.1 ABC transporter transmembrane domain-containing protein |
| 4(1) | WP_145318725.1 | WP_145318725.1 ABC transporter transmembrane domain-containing protein |
| 4(1) | WP_010346298.1 | WP_010346298.1 ABC transporter ATP-binding protein |
| 4(1) | WP_152381386.1 | WP_152381386.1 ABC transporter ATP-binding protein |
| 4(1) | WP_073008280.1 | WP_073008280.1 ABC transporter ATP-binding protein |
| 4(1) | WP_145318728.1 | WP_145318728.1 ABC transporter ATP-binding protein |
| 4(3) | WP_028542695.1 | WP_028542695.1 ABC transporter ATP-binding protein |
| 4(1) | WP_186580051.1 | WP_186580051.1 ABC transporter ATP-binding protein |
| 4(3) | WP_163977046.1 | WP_163977046.1 dipeptide ABC transporter ATP-binding protein |
| 4(2) | WP_076249784.1 | WP_076249784.1 ABC transporter transmembrane domain-containing protein |
| 4(1) | WP_274485137.1 | WP_274485137.1 ABC transporter ATP-binding protein |
| 4(1) | WP_204517528.1 | WP_204517528.1 ABC transporter ATP-binding protein |
| 4(1) | WP_251375558.1 | WP_251375558.1 ABC transporter transmembrane domain-containing protein |
| 4(1) | WP_194230416.1 | WP_194230416.1 ABC transporter transmembrane domain-containing protein |
| 4(1) | WP_145329883.1 | WP_145329883.1 ABC transporter ATP-binding protein |
| 4(1) | WP_061829411.1 | WP_061829411.1 ABC transporter transmembrane domain-containing protein |
| 4(1) | WP_301879178.1 | WP_301879178.1 ABC transporter transmembrane domain-containing protein |
| 4(1) | WP_216513481.1 | WP_216513481.1 ABC transporter transmembrane domain-containing protein |
| 4(1) | WP_213579158.1 | WP_213579158.1 ABC transporter ATP-binding protein |
| 4(1) | WP_076242351.1 | WP_076242351.1 ABC transporter ATP-binding protein |
| 4(1) | WP_251375559.1 | WP_251375559.1 ABC transporter ATP-binding protein |
| 4(1) | WP_281667409.1 | WP_281667409.1 ABC-F family ATP-binding cassette domain-containing protein |
| 4(1) | WP_072727814.1 | WP_072727814.1 sugar ABC transporter ATP-binding protein |
| 4(1) | WP_094936480.1 | WP_094936480.1 ABC transporter ATP-binding protein |
| 4(2) | WP_069080932.1 | WP_069080932.1 ABC transporter ATP-binding protein |
| 4(1) | WP_192571321.1 | WP_192571321.1 ABC transporter ATP-binding protein |
| 4(1) | WP_302153070.1 | WP_302153070.1 ATP-binding cassette domain-containing protein |
| 4(1) | WP_234032638.1 | WP_234032638.1 ABC transporter ATP-binding protein |
| 4(1) | WP_277746355.1 | WP_277746355.1 ABC transporter ATP-binding protein |
| 4(1) | WP_258296286.1 | WP_258296286.1 ABC transporter ATP-binding protein |
| 4(1) | WP_304142102.1 | WP_304142102.1 ABC transporter ATP-binding protein |
| 4(1) | WP_145021756.1 | WP_145021756.1 ABC transporter ATP-binding protein |
| 4(1) | WP_272035278.1 | WP_272035278.1 ABC transporter ATP-binding protein |
| 4(1) | WP_138186706.1 | WP_138186706.1 sugar ABC transporter ATP-binding protein |
| 4(1) | WP_307192905.1 | WP_307192905.1 ABC transporter ATP-binding protein |
| 4(1) | WP_244719549.1 | WP_244719549.1 ABC transporter ATP-binding protein |
| 4(1) | WP_100529428.1 | WP_100529428.1 ABC transporter ATP-binding protein |
| 4(1) | WP_044037309.1 | WP_044037309.1 ABC transporter ATP-binding protein |
| 4(1) | WP_240753674.1 | WP_240753674.1 ABC transporter ATP-binding protein |
| 4(2) | WP_268599727.1 | WP_268599727.1 dipeptide ABC transporter ATP-binding protein |

| | | |
|---|---|---|
| 4(1) | WP_014281170.1 | WP_014281170.1 ABC transporter ATP-binding protein |
| 4(1) | WP_018670513.1 | WP_018670513.1 ABC transporter ATP-binding protein |
| 4(1) | WP_103047832.1 | WP_103047832.1 ABC transporter transmembrane domain-containing protein |
| 4(1) | WP_192266428.1 | WP_192266428.1 ABC transporter ATP-binding protein |
| 4(1) | WP_282321828.1 | WP_282321828.1 ABC transporter ATP-binding protein |
| 4(1) | WP_063208808.1 | WP_063208808.1 ABC transporter ATP-binding protein |
| 4(2) | WP_283655684.1 | WP_283655684.1 ABC transporter ATP-binding protein |
| 4(1) | WP_111619125.1 | WP_111619125.1 ABC transporter ATP-binding protein |
| 4(1) | WP_127536203.1 | WP_127536203.1 ABC transporter ATP-binding protein |
| 4(1) | WP_003340561.1 | WP_003340561.1 ABC transporter ATP-binding protein |
| 4(1) | WP_236417194.1 | WP_236417194.1 ABC transporter ATP-binding protein |
| 4(2) | WP_017427961.1 | WP_017427961.1 ABC transporter ATP-binding protein |
| 4(1) | WP_231094861.1 | WP_231094861.1 ABC transporter ATP-binding protein |
| 4(1) | WP_205828386.1 | WP_205828386.1 ABC transporter ATP-binding protein |
| 4(1) | WP_091031779.1 | WP_091031779.1 ABC transporter transmembrane domain-containing protein |
| 4(1) | WP_021253131.1 | WP_021253131.1 sugar ABC transporter ATP-binding protein |
| 4(1) | WP_126665670.1 | WP_126665670.1 ABC transporter ATP-binding protein |
| 4(1) | WP_174820807.1 | WP_174820807.1 ABC transporter ATP-binding protein |
| 4(1) | WP_188593156.1 | WP_188593156.1 ABC transporter transmembrane domain-containing protein |
| 4(1) | WP_142612996.1 | WP_142612996.1 ABC transporter transmembrane domain-containing protein |
| 4(1) | WP_209861669.1 | WP_209861669.1 ABC transporter transmembrane domain-containing protein |
| 4(1) | WP_211745840.1 | WP_211745840.1 ABC transporter ATP-binding protein |
| 4(1) | WP_145146960.1 | WP_145146960.1 ABC transporter transmembrane domain-containing protein |
| 5(1) | WP_099354901.1 | WP_099354901.1 hypothetical protein |
| 5(1) | WP_163095403.1 | WP_163095403.1 hypothetical protein |
| 5(2) | WP_270993643.1 | WP_270993643.1 hypothetical protein |
| 5(1) | WP_102956432.1 | WP_102956432.1 hypothetical protein |
| 5(1) | WP_086424318.1 | WP_086424318.1 hypothetical protein |
| 5(1) | WP_257141301.1 | WP_257141301.1 hypothetical protein |
| 5(1) | WP_098291288.1 | WP_098291288.1 hypothetical protein |
| 5(1) | WP_111147794.1 | WP_111147794.1 DUF5085 family protein |
| 5(4) | WP_000267114.1 | WP_000267114.1 hypothetical protein |
| 5(5) | WP_016109855.1 | WP_016109855.1 hypothetical protein |
| 5(1) | WP_000267119.1 | WP_000267119.1 hypothetical protein |
| 5(1) | WP_106496025.1 | WP_106496025.1 hypothetical protein |
| 5(4) | WP_033732157.1 | WP_033732157.1 hypothetical protein |
| 5(1) | WP_061529908.1 | WP_061529908.1 hypothetical protein |
| 5(3) | WP_000701787.1 | WP_000701787.1 DUF5085 family protein |
| 5(1) | WP_098142227.1 | WP_098142227.1 hypothetical protein |
| 5(1) | WP_016134261.1 | WP_016134261.1 DUF5085 family protein |
| 5(1) | WP_234121599.1 | WP_234121599.1 DUF5085 family protein |
| 5(1) | WP_059303903.1 | WP_059303903.1 hypothetical protein |
| 5(1) | WP_098327464.1 | WP_098327464.1 hypothetical protein |
| 5(1) | WP_229139916.1 | WP_229139916.1 hypothetical protein |
| 5(1) | WP_290425833.1 | WP_290425833.1 hypothetical protein |
| 5(5) | WP_000582719.1 | WP_000582719.1 DUF5085 family protein |
| 5(1) | WP_098284021.1 | WP_098284021.1 hypothetical protein |
| 5(2) | WP_000868165.1 | WP_000868165.1 DUF5085 family protein |
| 5(1) | WP_138755915.1 | WP_138755915.1 hypothetical protein |
| 5(15) | WP_000582716.1 | WP_000582716.1 DUF5085 family protein |

| | | |
|---|---|---|
| 5(1) | WP_053512317.1 | WP_053512317.1 hypothetical protein |
| 5(2) | WP_098139251.1 | WP_098139251.1 hypothetical protein |
| 5(1) | WP_185451813.1 | WP_185451813.1 hypothetical protein |
| 5(4) | WP_221633036.1 | WP_221633036.1 hypothetical protein |
| 5(1) | WP_046953660.1 | WP_046953660.1 hypothetical protein |
| 5(1) | WP_098128487.1 | WP_098128487.1 hypothetical protein |
| 5(3) | WP_065223579.1 | WP_065223579.1 hypothetical protein |
| 5(1) | WP_053512319.1 | WP_053512319.1 DUF5085 family protein |
| 5(1) | WP_132131928.1 | WP_132131928.1 DUF5085 family protein |
| 5(1) | WP_000267118.1 | WP_000267118.1 hypothetical protein |
| 5(1) | WP_235286207.1 | WP_235286207.1 hypothetical protein |
| 5(1) | WP_061113314.1 | WP_061113314.1 hypothetical protein |
| 5(1) | WP_044584392.1 | WP_044584392.1 hypothetical protein |
| 5(1) | WP_025921922.1 | WP_025921922.1 hypothetical protein |
| 5(1) | WP_000267163.1 | WP_000267163.1 hypothetical protein |
| 5(3) | WP_001990241.1 | WP_001990241.1 hypothetical protein |
| 5(1) | WP_033690143.1 | WP_033690143.1 hypothetical protein |
| 5(1) | WP_038584726.1 | WP_038584726.1 hypothetical protein |
| 5(3) | WP_097954750.1 | WP_097954750.1 hypothetical protein |
| 5(2) | WP_221635811.1 | WP_221635811.1 hypothetical protein |
| 5(1) | WP_277493890.1 | WP_277493890.1 hypothetical protein, partial |
| 5(3) | WP_086406039.1 | WP_086406039.1 DUF5085 family protein |
| 5(2) | WP_234288978.1 | WP_234288978.1 hypothetical protein |
| 5(1) | WP_251217308.1 | WP_251217308.1 hypothetical protein |
| 5(3) | WP_001990253.1 | WP_001990253.1 hypothetical protein |
| 5(1) | WP_061667201.1 | WP_061667201.1 hypothetical protein |
| 5(1) | WP_098226389.1 | WP_098226389.1 DUF5085 family protein |
| 5(1) | WP_000582722.1 | WP_000582722.1 DUF5085 family protein |
| 5(1) | WP_079519269.1 | WP_079519269.1 hypothetical protein |
| 5(1) | WP_143316568.1 | WP_143316568.1 DUF5085 family protein |
| 5(1) | WP_116353255.1 | WP_116353255.1 hypothetical protein |
| 5(1) | WP_307601043.1 | WP_307601043.1 hypothetical protein |
| 5(1) | WP_116344880.1 | WP_116344880.1 hypothetical protein |
| 5(2) | WP_046953663.1 | WP_046953663.1 hypothetical protein |
| 5(1) | WP_150158010.1 | WP_150158010.1 hypothetical protein |
| 5(1) | WP_000912763.1 | WP_000912763.1 hypothetical protein |
| 5(3) | WP_048567944.1 | WP_048567944.1 hypothetical protein |
| 5(8) | WP_086406040.1 | WP_086406040.1 DUF5085 family protein |
| 5(1) | WP_228463448.1 | WP_228463448.1 hypothetical protein |
| 5(2) | WP_097999317.1 | WP_097999317.1 hypothetical protein |
| 5(1) | WP_235286208.1 | WP_235286208.1 DUF5085 family protein |
| 5(8) | WP_061667205.1 | WP_061667205.1 hypothetical protein |
| 5(2) | WP_000263483.1 | WP_000263483.1 hypothetical protein |
| 5(1) | WP_163095409.1 | WP_163095409.1 hypothetical protein |
| 5(5) | WP_000701786.1 | WP_000701786.1 DUF5085 family protein |
| 5(1) | WP_221638284.1 | WP_221638284.1 hypothetical protein |
| 5(1) | WP_098882511.1 | WP_098882511.1 hypothetical protein |
| 5(1) | WP_102956430.1 | WP_102956430.1 hypothetical protein |
| 5(1) | WP_000267105.1 | WP_000267105.1 hypothetical protein |
| 5(1) | WP_231094852.1 | WP_231094852.1 DUF5085 family protein |
| 5(1) | WP_098612034.1 | WP_098612034.1 hypothetical protein |
| 5(1) | WP_245799805.1 | WP_245799805.1 hypothetical protein |
| 5(2) | WP_098139255.1 | WP_098139255.1 DUF5085 family protein |
| 5(1) | WP_076113456.1 | WP_076113456.1 hypothetical protein |
| 5(1) | WP_086412584.1 | WP_086412584.1 hypothetical protein |
| 5(2) | WP_097954751.1 | WP_097954751.1 hypothetical protein |
| 5(1) | WP_006923834.1 | WP_006923834.1 hypothetical protein |
| 5(2) | WP_062821209.1 | WP_062821209.1 hypothetical protein |
| 5(1) | WP_098874964.1 | WP_098874964.1 hypothetical protein |
| 5(1) | WP_098157699.1 | WP_098157699.1 hypothetical protein |
| 5(1) | WP_275216784.1 | WP_275216784.1 hypothetical protein |
| 5(1) | WP_078384705.1 | WP_078384705.1 hypothetical protein |
| 5(8) | WP_016109854.1 | WP_016109854.1 hypothetical protein |
| 5(1) | WP_038584729.1 | WP_038584729.1 DUF5085 family protein |
| 5(2) | WP_053512320.1 | WP_053512320.1 hypothetical protein |
| 5(1) | WP_000267133.1 | WP_000267133.1 hypothetical protein |
| 5(3) | WP_016108262.1 | WP_016108262.1 DUF5085 family protein |
| 5(4) | WP_000868164.1 | WP_000868164.1 DUF5085 family protein |
| 5(1) | WP_099354904.1 | WP_099354904.1 DUF5085 family protein |
| 5(2) | WP_062821207.1 | WP_062821207.1 hypothetical protein |
| 5(1) | WP_088230460.1 | WP_088230460.1 DUF5085 family protein |
| 5(1) | WP_195343925.1 | WP_195343925.1 DUF5085 family protein |
| 5(1) | WP_208906818.1 | WP_208906818.1 hypothetical protein |
| 5(1) | WP_270065064.1 | WP_270065064.1 DUF5085 family protein |
| 5(1) | WP_120112242.1 | WP_120112242.1 DUF5085 family protein |
| 5(2) | WP_066419970.1 | WP_066419970.1 DUF5085 family protein |
| 5(1) | WP_098882508.1 | WP_098882508.1 hypothetical protein |
| 5(1) | WP_076876365.1 | WP_076876365.1 hypothetical protein |
| 5(1) | WP_136379696.1 | WP_136379696.1 DUF5085 family protein |
| 5(1) | WP_141527192.1 | WP_141527192.1 hypothetical protein |
| 5(1) | WP_000267172.1 | WP_000267172.1 hypothetical protein |
| 5(3) | WP_098291291.1 | WP_098291291.1 hypothetical protein |
| 5(1) | WP_041580743.1 | WP_041580743.1 hypothetical protein |
| 5(1) | WP_234121597.1 | WP_234121597.1 hypothetical protein |
| 5(2) | WP_030025439.1 | WP_030025439.1 hypothetical protein |
| 5(1) | WP_102956429.1 | WP_102956429.1 hypothetical protein |
| 5(1) | WP_136372962.1 | WP_136372962.1 DUF5085 domain-containing protein |
| 5(2) | WP_194335284.1 | WP_194335284.1 hypothetical protein |
| 5(3) | WP_030025438.1 | WP_030025438.1 hypothetical protein |
| 5(4) | WP_000267169.1 | WP_000267169.1 hypothetical protein |
| 5(2) | WP_098529293.1 | WP_098529293.1 hypothetical protein |
| 5(1) | WP_228596157.1 | WP_228596157.1 hypothetical protein, partial |
| 5(11) | WP_000701784.1 | WP_000701784.1 DUF5085 family protein |
| 5(10) | WP_061667204.1 | WP_061667204.1 DUF5085 family protein |
| 5(1) | WP_111147795.1 | WP_111147795.1 DUF5085 family protein |
| 5(1) | WP_000267138.1 | WP_000267138.1 hypothetical protein |
| 5(1) | WP_098645177.1 | WP_098645177.1 hypothetical protein |
| 5(1) | WP_098819903.1 | WP_098819903.1 DUF5085 family protein |
| 5(3) | WP_000267193.1 | WP_000267193.1 hypothetical protein |
| 5(4) | WP_097999318.1 | WP_097999318.1 hypothetical protein |
| 5(1) | WP_048558208.1 | WP_048558208.1 DUF5085 family protein |
| 5(1) | WP_016117674.1 | WP_016117674.1 hypothetical protein |
| 5(2) | WP_000582721.1 | WP_000582721.1 DUF5085 family protein |
| 5(1) | WP_076113454.1 | WP_076113454.1 DUF5085 family protein |

| | | |
|---|---|---|
| 5(1) | WP_291672213.1 | WP_291672213.1 DUF5085 family protein |
| 5(7) | WP_000634597.1 | WP_000634597.1 DUF5085 family protein |
| 5(3) | WP_033665476.1 | WP_033665476.1 hypothetical protein |
| 5(1) | WP_000267149.1 | WP_000267149.1 hypothetical protein |
| 5(1) | WP_000267131.1 | WP_000267131.1 hypothetical protein |
| 5(1) | WP_006923827.1 | WP_006923827.1 hypothetical protein |
| 5(1) | WP_078384706.1 | WP_078384706.1 hypothetical protein |
| 5(1) | WP_098284018.1 | WP_098284018.1 hypothetical protein |
| 5(1) | WP_163095405.1 | WP_163095405.1 hypothetical protein |
| 5(2) | WP_185413337.1 | WP_185413337.1 hypothetical protein |
| 5(2) | WP_061678621.1 | WP_061678621.1 hypothetical protein |
| 5(2) | WP_030025443.1 | WP_030025443.1 hypothetical protein |
| 5(1) | WP_006923828.1 | WP_006923828.1 hypothetical protein |
| 5(1) | WP_204335718.1 | WP_204335718.1 hypothetical protein |
| 5(1) | WP_290102343.1 | WP_290102343.1 hypothetical protein |
| 5(1) | WP_234121601.1 | WP_234121601.1 hypothetical protein |
| 5(1) | WP_132131926.1 | WP_132131926.1 hypothetical protein |
| 5(1) | WP_016117681.1 | WP_016117681.1 hypothetical protein |
| 5(2) | WP_098646938.1 | WP_098646938.1 hypothetical protein |
| 5(1) | WP_136379693.1 | WP_136379693.1 hypothetical protein |
| 5(1) | WP_013490133.1 | WP_013490133.1 DUF5085 family protein |
| 5(2) | WP_046953662.1 | WP_046953662.1 DUF5085 family protein |
| 5(1) | WP_245799806.1 | WP_245799806.1 hypothetical protein |
| 5(1) | WP_000267121.1 | WP_000267121.1 hypothetical protein |
| 5(1) | WP_231094854.1 | WP_231094854.1 hypothetical protein |
| 5(2) | WP_221638507.1 | WP_221638507.1 hypothetical protein |
| 5(1) | WP_006923845.1 | WP_006923845.1 hypothetical protein |
| 5(1) | WP_000267194.1 | WP_000267194.1 hypothetical protein |
| 5(1) | WP_048538776.1 | WP_048538776.1 hypothetical protein |
| 5(1) | WP_106496026.1 | WP_106496026.1 DUF5085 family protein |
| 5(1) | WP_077320209.1 | WP_077320209.1 hypothetical protein |
| 5(1) | WP_143316567.1 | WP_143316567.1 DUF5085 family protein |
| 5(1) | WP_076113455.1 | WP_076113455.1 DUF5085 family protein |
| 5(1) | WP_163095407.1 | WP_163095407.1 hypothetical protein |
| 5(2) | WP_000267116.1 | WP_000267116.1 hypothetical protein |
| 5(2) | WP_098327465.1 | WP_098327465.1 hypothetical protein |
| 5(1) | WP_238187887.1 | WP_238187887.1 DUF5085 family protein |
| 5(1) | WP_234121596.1 | WP_234121596.1 hypothetical protein |
| 5(1) | WP_098522176.1 | WP_098522176.1 hypothetical protein |
| 5(1) | WP_243644245.1 | WP_243644245.1 DUF5085 family protein |
| 5(1) | WP_186580059.1 | WP_186580059.1 hypothetical protein |
| 5(1) | WP_153589098.1 | WP_153589098.1 hypothetical protein |
| 5(1) | WP_136372959.1 | WP_136372959.1 hypothetical protein |
| 5(1) | WP_136372961.1 | WP_136372961.1 hypothetical protein |
| 5(1) | WP_000267166.1 | WP_000267166.1 hypothetical protein |
| 5(1) | WP_000265485.1 | WP_000265485.1 hypothetical protein |
| 5(1) | WP_195542476.1 | WP_195542476.1 hypothetical protein |
| 5(1) | WP_098013427.1 | WP_098013427.1 hypothetical protein |
| 5(3) | WP_000267168.1 | WP_000267168.1 hypothetical protein |
| 5(1) | WP_000267113.1 | WP_000267113.1 hypothetical protein |
| 5(1) | WP_000267160.1 | WP_000267160.1 hypothetical protein |
| 5(2) | WP_251217307.1 | WP_251217307.1 hypothetical protein |
| 5(4) | WP_064774891.1 | WP_064774891.1 hypothetical protein, partial |
| 5(2) | WP_185413335.1 | WP_185413335.1 DUF5085 family protein |
| 5(1) | WP_098142226.1 | WP_098142226.1 hypothetical protein |
| 5(1) | WP_255289251.1 | WP_255289251.1 hypothetical protein |
| 5(1) | WP_235286209.1 | WP_235286209.1 DUF5085 family protein |
| 5(1) | WP_153589097.1 | WP_153589097.1 hypothetical protein |
| 5(2) | WP_066419972.1 | WP_066419972.1 hypothetical protein |
| 5(1) | WP_044584390.1 | WP_044584390.1 hypothetical protein |
| 5(2) | WP_016717319.1 | WP_016717319.1 DUF5085 family protein |
| 5(1) | WP_098435165.1 | WP_098435165.1 hypothetical protein |
| 5(1) | WP_153589099.1 | WP_153589099.1 DUF5085 family protein |
| 5(1) | WP_098413328.1 | WP_098413328.1 hypothetical protein |
| 5(1) | WP_098882510.1 | WP_098882510.1 DUF5085 family protein |
| 5(1) | WP_090761789.1 | WP_090761789.1 AraC family transcriptional regulator |
| 5(1) | WP_258113119.1 | WP_258113119.1 hypothetical protein |
| 5(1) | WP_061667203.1 | WP_061667203.1 DUF5085 family protein |
| 5(3) | WP_228469156.1 | WP_228469156.1 hypothetical protein |
| 5(1) | WP_103597527.1 | WP_103597527.1 hypothetical protein |
| 5(2) | WP_098481731.1 | WP_098481731.1 hypothetical protein |
| 5(4) | WP_000582718.1 | WP_000582718.1 DUF5085 family protein |
| 5(1) | WP_136372963.1 | WP_136372963.1 DUF5085 family protein |
| 5(1) | WP_231094853.1 | WP_231094853.1 DUF5085 family protein |
| 5(1) | WP_234121600.1 | WP_234121600.1 DUF5085 family protein |
| 5(2) | WP_012984616.1 | WP_012984616.1 hypothetical protein |
| 5(2) | WP_016091016.1 | WP_016091016.1 DUF5085 family protein |
| 5(1) | WP_016134260.1 | WP_016134260.1 DUF5085 family protein |
| 5(3) | WP_003745060.1 | WP_003745060.1 hypothetical protein |
| 5(1) | WP_098522177.1 | WP_098522177.1 hypothetical protein |
| 5(1) | WP_066419971.1 | WP_066419971.1 DUF5085 family protein |
| 5(1) | WP_000267135.1 | WP_000267135.1 hypothetical protein |
| 5(1) | WP_038584731.1 | WP_038584731.1 DUF5085 family protein |
| 5(2) | WP_251217305.1 | WP_251217305.1 hypothetical protein |
| 5(3) | WP_097837319.1 | WP_097837319.1 hypothetical protein |
| 5(1) | WP_086424320.1 | WP_086424320.1 DUF5085 family protein |
| 5(1) | WP_238187886.1 | WP_238187886.1 hypothetical protein |
| 5(1) | WP_136379695.1 | WP_136379695.1 DUF5085 family protein |
| 5(3) | WP_195343924.1 | WP_195343924.1 DUF5085 family protein |
| 5(2) | WP_066419973.1 | WP_066419973.1 hypothetical protein |
| 5(4) | WP_001990243.1 | WP_001990243.1 hypothetical protein |
| 5(1) | WP_053512318.1 | WP_053512318.1 hypothetical protein |
| 5(2) | WP_228465488.1 | WP_228465488.1 hypothetical protein |
| 5(1) | WP_046392644.1 | WP_046392644.1 hypothetical protein |
| 5(1) | WP_048558207.1 | WP_048558207.1 hypothetical protein |
| 5(1) | WP_000267391.1 | WP_000267391.1 hypothetical protein |
| 5(1) | WP_238187888.1 | WP_238187888.1 DUF5085 family protein |
| 5(1) | WP_142255385.1 | WP_142255385.1 hypothetical protein |
| 5(1) | WP_141557692.1 | WP_141557692.1 hypothetical protein |
| 5(1) | WP_097796440.1 | WP_097796440.1 hypothetical protein |
| 5(1) | WP_016117671.1 | WP_016117671.1 hypothetical protein |
| 5(3) | WP_065223576.1 | WP_065223576.1 hypothetical protein |
| 5(2) | WP_098139257.1 | WP_098139257.1 DUF5085 family protein |
| 5(2) | WP_016091015.1 | WP_016091015.1 hypothetical protein |

| | | |
|---|---|---|
| 5(1) | WP_000912767.1 | WP_000912767.1 hypothetical protein |
| 5(1) | WP_098210154.1 | WP_098210154.1 hypothetical protein |
| 5(1) | WP_307601045.1 | WP_307601045.1 DUF5085 family protein |
| 5(1) | WP_138755913.1 | WP_138755913.1 DUF5085 family protein |
| 5(1) | WP_076876363.1 | WP_076876363.1 hypothetical protein |
| 5(9) | WP_001990250.1 | WP_001990250.1 DUF5085 family protein |
| 5(1) | WP_049131598.1 | WP_049131598.1 hypothetical protein, partial |
| 5(5) | WP_030025441.1 | WP_030025441.1 hypothetical protein |
| 5(1) | WP_016117672.1 | WP_016117672.1 DUF5085 family protein |
| 5(1) | WP_078384704.1 | WP_078384704.1 hypothetical protein |
| 5(3) | WP_240514391.1 | WP_240514391.1 hypothetical protein |
| 5(2) | WP_228465487.1 | WP_228465487.1 hypothetical protein |
| 5(2) | WP_000267115.1 | WP_000267115.1 hypothetical protein |
| 5(1) | WP_111147796.1 | WP_111147796.1 hypothetical protein |
| 5(2) | WP_063264138.1 | WP_063264138.1 hypothetical protein |
| 5(1) | WP_120112248.1 | WP_120112248.1 hypothetical protein |
| 5(6) | WP_086406042.1 | WP_086406042.1 hypothetical protein |
| 5(1) | WP_138755914.1 | WP_138755914.1 DUF5085 family protein |
| 5(1) | WP_251217306.1 | WP_251217306.1 hypothetical protein |
| 5(1) | WP_099354902.1 | WP_099354902.1 hypothetical protein |
| 5(1) | WP_098882509.1 | WP_098882509.1 hypothetical protein |
| 5(1) | WP_013490135.1 | WP_013490135.1 hypothetical protein |
| 5(1) | WP_136379694.1 | WP_136379694.1 hypothetical protein |
| 5(2) | WP_000267093.1 | WP_000267093.1 hypothetical protein |
| 5(1) | WP_044584395.1 | WP_044584395.1 hypothetical protein |
| 5(1) | WP_200462052.1 | WP_200462052.1 hypothetical protein |
| 5(3) | WP_061113066.1 | WP_061113066.1 DUF5085 family protein |
| 5(1) | WP_098819900.1 | WP_098819900.1 DUF5085 family protein |
| 5(5) | WP_016109857.1 | WP_016109857.1 hypothetical protein |
| 5(1) | WP_000267161.1 | WP_000267161.1 hypothetical protein |
| 5(1) | WP_098284022.1 | WP_098284022.1 hypothetical protein |
| 5(1) | WP_099354903.1 | WP_099354903.1 DUF5085 family protein |
| 5(2) | WP_097999316.1 | WP_097999316.1 hypothetical protein |
| 5(1) | WP_098226388.1 | WP_098226388.1 DUF5085 family protein |
| 5(2) | WP_046959244.1 | WP_046959244.1 DUF5085 family protein |
| 5(1) | WP_138755916.1 | WP_138755916.1 hypothetical protein |
| 5(1) | WP_258113055.1 | WP_258113055.1 DUF5085 family protein |
| 5(1) | WP_106496027.1 | WP_106496027.1 DUF5085 family protein |
| 5(1) | WP_098327463.1 | WP_098327463.1 hypothetical protein |
| 5(1) | WP_234121595.1 | WP_234121595.1 hypothetical protein |
| 5(1) | WP_063261712.1 | WP_063261712.1 hypothetical protein |
| 5(1) | WP_077320210.1 | WP_077320210.1 DUF5085 family protein |
| 5(1) | WP_016117673.1 | WP_016117673.1 hypothetical protein |
| 5(3) | WP_000912760.1 | WP_000912760.1 hypothetical protein |
| 5(4) | WP_000868166.1 | WP_000868166.1 DUF5085 family protein |
| 5(1) | WP_013490134.1 | WP_013490134.1 DUF5085 family protein |
| 5(2) | WP_016117670.1 | WP_016117670.1 hypothetical protein |

| | | |
|---|---|---|
| 6(1) | WP_224839859.1 | WP_224839859.1 hypothetical protein |
| 6(2) | WP_079694648.1 | WP_079694648.1 hypothetical protein |
| 6(1) | WP_076322114.1 | WP_076322114.1 hypothetical protein |
| 6(1) | WP_019912155.1 | WP_019912155.1 hypothetical protein |
| 6(1) | WP_150364196.1 | WP_150364196.1 hypothetical protein |
| 6(1) | WP_076279384.1 | WP_076279384.1 hypothetical protein |
| 6(1) | WP_014281182.1 | WP_014281182.1 hypothetical protein |
| 6(1) | WP_168178888.1 | WP_168178888.1 hypothetical protein |
| 6(1) | WP_213618814.1 | WP_213618814.1 hypothetical protein |
| 6(1) | WP_111619122.1 | WP_111619122.1 hypothetical protein |
| 6(1) | WP_068782407.1 | WP_068782407.1 hypothetical protein |
| 6(1) | WP_111268940.1 | WP_111268940.1 hypothetical protein |
| 6(1) | WP_028401640.1 | WP_028401640.1 hypothetical protein |
| 6(1) | WP_209941811.1 | WP_209941811.1 hypothetical protein |
| 6(1) | WP_022586769.1 | WP_022586769.1 hypothetical protein |
| 6(1) | WP_251417627.1 | WP_251417627.1 hypothetical protein |
| 6(1) | WP_039306360.1 | WP_039306360.1 hypothetical protein |
| 6(1) | WP_051249878.1 | WP_051249878.1 hypothetical protein |
| 6(1) | WP_301804667.1 | WP_301804667.1 hypothetical protein |
| 6(1) | WP_306077392.1 | WP_306077392.1 hypothetical protein |
| 6(1) | WP_273321547.1 | WP_273321547.1 hypothetical protein |
| 6(1) | WP_280788632.1 | WP_280788632.1 hypothetical protein |
| 6(5) | WP_003340546.1 | WP_003340546.1 hypothetical protein |
| 6(1) | WP_274485151.1 | WP_274485151.1 hypothetical protein |
| 6(1) | WP_010250276.1 | WP_010250276.1 hypothetical protein |
| 6(2) | WP_025679804.1 | WP_025679804.1 hypothetical protein |
| 6(1) | WP_091260862.1 | WP_091260862.1 hypothetical protein |
| 6(1) | WP_212695788.1 | WP_212695788.1 hypothetical protein |
| 6(1) | WP_134902009.1 | WP_134902009.1 hypothetical protein |
| 6(1) | WP_053325279.1 | WP_053325279.1 hypothetical protein |
| 6(1) | WP_232699384.1 | WP_232699384.1 hypothetical protein |
| 6(1) | WP_110931546.1 | WP_110931546.1 hypothetical protein |
| 6(2) | WP_197262006.1 | WP_197262006.1 hypothetical protein |
| 6(1) | WP_272035308.1 | WP_272035308.1 hypothetical protein |
| 6(1) | WP_113050536.1 | WP_113050536.1 hypothetical protein |
| 6(1) | WP_286458023.1 | WP_286458023.1 hypothetical protein |
| 6(1) | WP_121474479.1 | WP_121474479.1 hypothetical protein |
| 6(1) | WP_094878479.1 | WP_094878479.1 hypothetical protein |
| 6(1) | WP_250268130.1 | WP_250268130.1 hypothetical protein |
| 6(1) | WP_028589295.1 | WP_028589295.1 hypothetical protein |
| 6(1) | WP_209872748.1 | WP_209872748.1 hypothetical protein |
| 6(1) | WP_250244818.1 | WP_250244818.1 hypothetical protein |
| 6(1) | WP_166244930.1 | WP_166244930.1 hypothetical protein |
| 6(1) | WP_126996283.1 | WP_126996283.1 hypothetical protein |
| 6(1) | WP_194230405.1 | WP_194230405.1 hypothetical protein |
| 6(1) | WP_003336001.1 | WP_003336001.1 hypothetical protein |
| 6(1) | WP_213532656.1 | WP_213532656.1 hypothetical protein |
| 6(2) | WP_236628610.1 | WP_236628610.1 hypothetical protein |
| 6(1) | WP_094155408.1 | WP_094155408.1 hypothetical protein |
| 6(1) | WP_160496933.1 | WP_160496933.1 hypothetical protein |
| 6(4) | WP_029517063.1 | WP_029517063.1 hypothetical protein |
| 6(1) | WP_301167502.1 | WP_301167502.1 hypothetical protein |
| 6(1) | WP_289475131.1 | WP_289475131.1 hypothetical protein |
| 6(1) | WP_028542690.1 | WP_028542690.1 hypothetical protein |
| 6(1) | WP_094593873.1 | WP_094593873.1 hypothetical protein |
| 6(1) | WP_192266453.1 | WP_192266453.1 hypothetical protein |

| | | |
|---|---|---|
| 6(1) | WP_114298761.1 | WP_114298761.1 hypothetical protein |
| 6(1) | WP_076219072.1 | WP_076219072.1 hypothetical protein |
| 6(2) | WP_019424147.1 | WP_019424147.1 hypothetical protein |
| 6(1) | WP_307312542.1 | WP_307312542.1 hypothetical protein |
| 6(1) | WP_204517512.1 | WP_204517512.1 hypothetical protein |
| 6(1) | WP_198310833.1 | WP_198310833.1 hypothetical protein |
| 6(1) | WP_163977056.1 | WP_163977056.1 hypothetical protein |
| 6(1) | WP_145318698.1 | WP_145318698.1 hypothetical protein |
| 6(1) | WP_278431125.1 | WP_278431125.1 hypothetical protein |
| 6(1) | WP_113672703.1 | WP_113672703.1 hypothetical protein |
| 6(1) | WP_110892974.1 | WP_110892974.1 hypothetical protein |
| 6(1) | WP_212930976.1 | WP_212930976.1 hypothetical protein |
| 6(1) | WP_091017195.1 | WP_091017195.1 hypothetical protein |
| 6(1) | WP_076289573.1 | WP_076289573.1 hypothetical protein |
| 6(1) | WP_028405550.1 | WP_028405550.1 hypothetical protein |
| 6(1) | WP_179016428.1 | WP_179016428.1 hypothetical protein |
| 6(1) | WP_307537582.1 | WP_307537582.1 hypothetical protein |
| 6(1) | WP_307214307.1 | WP_307214307.1 hypothetical protein |
| 6(1) | WP_208203481.1 | WP_208203481.1 hypothetical protein |
| 6(1) | WP_189018606.1 | WP_189018606.1 hypothetical protein |
| 6(1) | WP_145046453.1 | WP_145046453.1 hypothetical protein |
| 6(1) | WP_216531931.1 | WP_216531931.1 hypothetical protein |
| 6(1) | WP_055105789.1 | WP_055105789.1 hypothetical protein |
| 6(1) | WP_068496249.1 | WP_068496249.1 hypothetical protein |
| 6(2) | WP_109999890.1 | WP_109999890.1 hypothetical protein |
| 6(1) | WP_123067113.1 | WP_123067113.1 hypothetical protein |
| 6(1) | WP_251375551.1 | WP_251375551.1 hypothetical protein |
| 6(1) | WP_248063438.1 | WP_248063438.1 hypothetical protein |
| 6(1) | WP_204517515.1 | WP_204517515.1 hypothetical protein |
| 6(1) | WP_290402000.1 | WP_290402000.1 hypothetical protein |
| 6(1) | WP_025685032.1 | WP_025685032.1 hypothetical protein |
| 6(1) | WP_036944557.1 | WP_036944557.1 hypothetical protein |
| 6(1) | WP_212693239.1 | WP_212693239.1 hypothetical protein |
| 6(1) | WP_305022924.1 | WP_305022924.1 hypothetical protein |
| 6(1) | WP_153978067.1 | WP_153978067.1 hypothetical protein |
| 6(1) | WP_076210978.1 | WP_076210978.1 hypothetical protein |
| 6(1) | WP_025719062.1 | WP_025719062.1 hypothetical protein |
| 6(1) | WP_211082926.1 | WP_211082926.1 hypothetical protein |
| 6(1) | WP_184183000.1 | WP_184183000.1 hypothetical protein |
| 6(1) | WP_024632684.1 | WP_024632684.1 hypothetical protein |
| 6(2) | WP_024630326.1 | WP_024630326.1 hypothetical protein |
| 6(2) | WP_007128629.1 | WP_007128629.1 hypothetical protein |
| 6(1) | WP_014649151.1 | WP_014649151.1 hypothetical protein |
| 6(1) | WP_157763985.1 | WP_157763985.1 hypothetical protein |
| 6(1) | WP_190299135.1 | WP_190299135.1 hypothetical protein |
| 6(1) | WP_248251189.1 | WP_248251189.1 hypothetical protein |
| 6(1) | WP_305166262.1 | WP_305166262.1 hypothetical protein |
| 6(1) | WP_268619936.1 | WP_268619936.1 hypothetical protein |
| 6(2) | WP_216518253.1 | WP_216518253.1 hypothetical protein |
| 6(1) | WP_265408840.1 | WP_265408840.1 hypothetical protein |
| 6(5) | WP_068940682.1 | WP_068940682.1 hypothetical protein |
| 6(1) | WP_254327965.1 | WP_254327965.1 hypothetical protein |
| 6(1) | WP_023957721.1 | WP_023957721.1 hypothetical protein |
| 6(1) | WP_265408839.1 | WP_265408839.1 hypothetical protein |
| 6(3) | WP_158326749.1 | WP_158326749.1 hypothetical protein |
| 6(1) | WP_003193393.1 | WP_003193393.1 hypothetical protein |
| 6(1) | WP_258427458.1 | WP_258427458.1 hypothetical protein |
| 6(2) | WP_283655693.1 | WP_283655693.1 hypothetical protein |
| 6(1) | WP_137062799.1 | WP_137062799.1 hypothetical protein |
| 6(2) | WP_197247511.1 | WP_197247511.1 hypothetical protein |
| 6(1) | WP_213468414.1 | WP_213468414.1 hypothetical protein |
| 6(1) | WP_090239234.1 | WP_090239234.1 hypothetical protein |
| 6(1) | WP_077567020.1 | WP_077567020.1 hypothetical protein |
| 6(1) | WP_241929678.1 | WP_241929678.1 hypothetical protein |
| 6(1) | WP_220027681.1 | WP_220027681.1 hypothetical protein |
| 6(1) | WP_160037484.1 | WP_160037484.1 hypothetical protein |
| 6(1) | WP_209861647.1 | WP_209861647.1 hypothetical protein |
| 6(1) | WP_285748176.1 | WP_285748176.1 hypothetical protein |
| 6(1) | WP_010676852.1 | WP_010676852.1 hypothetical protein |
| 6(1) | WP_308912811.1 | WP_308912811.1 hypothetical protein |
| 6(1) | WP_258296280.1 | WP_258296280.1 hypothetical protein |
| 6(1) | WP_136604812.1 | WP_136604812.1 hypothetical protein |
| 6(1) | WP_247083038.1 | WP_247083038.1 hypothetical protein |
| 6(1) | WP_097899493.1 | WP_097899493.1 hypothetical protein |
| 6(1) | WP_152381394.1 | WP_152381394.1 hypothetical protein |
| 6(2) | WP_219872454.1 | WP_219872454.1 hypothetical protein |
| 6(1) | WP_076292681.1 | WP_076292681.1 hypothetical protein |
| 6(1) | WP_106765959.1 | WP_106765959.1 hypothetical protein |
| 6(1) | WP_116231177.1 | WP_116231177.1 hypothetical protein |
| 6(2) | WP_023988265.1 | WP_023988265.1 hypothetical protein |
| 6(1) | WP_091155377.1 | WP_091155377.1 hypothetical protein |
| 6(1) | WP_250261573.1 | WP_250261573.1 hypothetical protein |
| 6(1) | WP_116556143.1 | WP_116556143.1 hypothetical protein |
| 6(1) | WP_061829405.1 | WP_061829405.1 hypothetical protein |
| 6(1) | WP_307192899.1 | WP_307192899.1 hypothetical protein |
| 6(1) | WP_127194959.1 | WP_127194959.1 hypothetical protein |
| 6(1) | WP_244719627.1 | WP_244719627.1 hypothetical protein |
| 6(1) | WP_210087113.1 | WP_210087113.1 hypothetical protein |
| 6(2) | WP_076330722.1 | WP_076330722.1 hypothetical protein |
| 6(2) | WP_090896189.1 | WP_090896189.1 hypothetical protein |
| 6(1) | WP_152392574.1 | WP_152392574.1 hypothetical protein |
| 6(1) | WP_225999408.1 | WP_225999408.1 hypothetical protein |
| 6(1) | WP_258417128.1 | WP_258417128.1 hypothetical protein |
| 6(1) | WP_099361346.1 | WP_099361346.1 hypothetical protein |
| 6(1) | WP_277548438.1 | WP_277548438.1 hypothetical protein |
| 6(1) | WP_169505052.1 | WP_169505052.1 hypothetical protein |
| 6(1) | WP_017689073.1 | WP_017689073.1 hypothetical protein |
| 6(1) | WP_068793931.1 | WP_068793931.1 hypothetical protein |
| 6(1) | WP_212981558.1 | WP_212981558.1 hypothetical protein |
| 6(1) | WP_074108297.1 | WP_074108297.1 hypothetical protein |
| 6(1) | WP_062239849.1 | WP_062239849.1 hypothetical protein |
| 6(1) | WP_064019319.1 | WP_064019319.1 hypothetical protein |
| 6(2) | WP_128101487.1 | WP_128101487.1 hypothetical protein |
| 6(1) | WP_094096057.1 | WP_094096057.1 hypothetical protein |

| | | |
|---|---|---|
| 6(1) | WP_293654398.1 | WP_293654398.1 hypothetical protein |
| 6(1) | WP_102997917.1 | WP_102997917.1 hypothetical protein |
| 6(1) | WP_133332978.1 | WP_133332978.1 hypothetical protein |
| 6(1) | WP_264929206.1 | WP_264929206.1 hypothetical protein |
| 6(1) | WP_213649972.1 | WP_213649972.1 hypothetical protein |
| 6(1) | WP_183600891.1 | WP_183600891.1 hypothetical protein |
| 6(1) | WP_063208778.1 | WP_063208778.1 hypothetical protein |
| 6(1) | WP_304942560.1 | WP_304942560.1 hypothetical protein |
| 6(1) | WP_307399119.1 | WP_307399119.1 hypothetical protein |
| 6(2) | WP_087442411.1 | WP_087442411.1 hypothetical protein |
| 6(1) | WP_179603483.1 | WP_179603483.1 hypothetical protein |
| 6(1) | WP_304216885.1 | WP_304216885.1 hypothetical protein |
| 6(2) | WP_281813102.1 | WP_281813102.1 hypothetical protein |
| 6(2) | WP_094699655.1 | WP_094699655.1 hypothetical protein |
| 6(2) | WP_076303951.1 | WP_076303951.1 hypothetical protein |
| 6(1) | WP_074094615.1 | WP_074094615.1 hypothetical protein |
| 6(1) | WP_068536375.1 | WP_068536375.1 hypothetical protein |
| 6(1) | WP_283892179.1 | WP_283892179.1 hypothetical protein |
| 6(2) | WP_036670435.1 | WP_036670435.1 hypothetical protein |
| 6(1) | WP_256227722.1 | WP_256227722.1 hypothetical protein |
| 6(1) | WP_071558843.1 | WP_071558843.1 hypothetical protein |
| 6(1) | WP_127592428.1 | WP_127592428.1 hypothetical protein |
| 6(1) | WP_036609198.1 | WP_036609198.1 hypothetical protein |
| 6(1) | WP_145329906.1 | WP_145329906.1 hypothetical protein |
| 6(1) | WP_251466797.1 | WP_251466797.1 hypothetical protein |
| 6(1) | WP_293755524.1 | WP_293755524.1 hypothetical protein |
| 6(1) | WP_257126956.1 | WP_257126956.1 hypothetical protein |
| 6(5) | WP_063566323.1 | WP_063566323.1 hypothetical protein |
| 6(1) | WP_293202262.1 | WP_293202262.1 hypothetical protein |
| 6(1) | WP_104065531.1 | WP_104065531.1 hypothetical protein |
| 6(1) | WP_042203641.1 | WP_042203641.1 hypothetical protein |
| 6(1) | WP_307601037.1 | WP_307601037.1 hypothetical protein |
| 6(2) | WP_090921057.1 | WP_090921057.1 hypothetical protein |
| 6(1) | WP_039871649.1 | WP_039871649.1 hypothetical protein |
| 6(1) | WP_211745831.1 | WP_211745831.1 hypothetical protein |
| 6(1) | WP_026478109.1 | WP_026478109.1 hypothetical protein |
| 6(1) | WP_053783886.1 | WP_053783886.1 hypothetical protein |
| 6(1) | WP_104495349.1 | WP_104495349.1 hypothetical protein |
| 6(1) | WP_111273169.1 | WP_111273169.1 hypothetical protein |
| 6(1) | WP_145948997.1 | WP_145948997.1 hypothetical protein |
| 6(1) | WP_071640577.1 | WP_071640577.1 hypothetical protein |
| 6(1) | WP_213426992.1 | WP_213426992.1 hypothetical protein |
| 6(1) | WP_285079441.1 | WP_285079441.1 hypothetical protein |
| 6(1) | WP_100537975.1 | WP_100537975.1 hypothetical protein |
| 6(3) | WP_171414582.1 | WP_171414582.1 hypothetical protein |
| 6(2) | WP_119847552.1 | WP_119847552.1 hypothetical protein |
| 6(1) | WP_090993619.1 | WP_090993619.1 hypothetical protein |
| 6(3) | WP_005546509.1 | WP_005546509.1 hypothetical protein |
| 6(1) | WP_138186697.1 | WP_138186697.1 hypothetical protein |
| 6(1) | WP_026572736.1 | WP_026572736.1 hypothetical protein |
| 6(1) | WP_074085359.1 | WP_074085359.1 hypothetical protein |
| 6(1) | WP_229522857.1 | WP_229522857.1 hypothetical protein |
| 6(1) | WP_113755278.1 | WP_113755278.1 hypothetical protein |
| 6(1) | WP_051240451.1 | WP_051240451.1 hypothetical protein |
| 6(1) | WP_145412706.1 | WP_145412706.1 hypothetical protein |
| 6(3) | WP_076104572.1 | WP_076104572.1 hypothetical protein |
| 6(2) | WP_051614169.1 | WP_051614169.1 hypothetical protein |
| 6(4) | WP_095287661.1 | WP_095287661.1 hypothetical protein |
| 6(9) | WP_089546600.1 | WP_089546600.1 hypothetical protein |
| 6(1) | WP_289392074.1 | WP_289392074.1 hypothetical protein |
| 6(1) | WP_176524034.1 | WP_176524034.1 hypothetical protein |
| 6(1) | WP_119794361.1 | WP_119794361.1 hypothetical protein |
| 6(1) | WP_242210846.1 | WP_242210846.1 hypothetical protein |
| 6(1) | WP_160789863.1 | WP_160789863.1 hypothetical protein |
| 6(1) | WP_010346285.1 | WP_010346285.1 hypothetical protein |
| 6(3) | WP_095266196.1 | WP_095266196.1 hypothetical protein |
| 6(1) | WP_242210845.1 | WP_242210845.1 hypothetical protein |
| 6(1) | WP_269055425.1 | WP_269055425.1 hypothetical protein |
| 6(1) | WP_034675358.1 | WP_034675358.1 hypothetical protein |
| 6(1) | WP_057897629.1 | WP_057897629.1 hypothetical protein |
| 6(1) | WP_108465347.1 | WP_108465347.1 hypothetical protein |
| 6(2) | WP_137078364.1 | WP_137078364.1 hypothetical protein |
| 6(1) | WP_154985828.1 | WP_154985828.1 hypothetical protein |
| 6(1) | WP_142543128.1 | WP_142543128.1 hypothetical protein |
| 6(1) | WP_155703935.1 | WP_155703935.1 hypothetical protein |
| 6(1) | WP_007430131.1 | WP_007430131.1 hypothetical protein |
| 6(1) | WP_290409174.1 | WP_290409174.1 hypothetical protein |
| 6(1) | WP_208919032.1 | WP_208919032.1 hypothetical protein |
| 6(1) | WP_145402726.1 | WP_145402726.1 hypothetical protein |
| 6(1) | WP_216830393.1 | WP_216830393.1 hypothetical protein |
| 6(4) | WP_006674863.1 | WP_006674863.1 hypothetical protein |
| 6(1) | WP_251537225.1 | WP_251537225.1 hypothetical protein |
| 6(1) | WP_083545032.1 | WP_083545032.1 hypothetical protein |
| 6(1) | WP_058829512.1 | WP_058829512.1 hypothetical protein |
| 6(1) | WP_076167870.1 | WP_076167870.1 hypothetical protein |
| 6(4) | WP_018670521.1 | WP_018670521.1 hypothetical protein |
| 6(1) | WP_309098620.1 | WP_309098620.1 hypothetical protein |
| 6(1) | WP_076301097.1 | WP_076301097.1 hypothetical protein |
| 6(1) | WP_098248518.1 | WP_098248518.1 hypothetical protein |
| 6(1) | WP_018883273.1 | WP_018883273.1 hypothetical protein |
| 6(3) | WP_104030442.1 | WP_104030442.1 hypothetical protein |
| 6(1) | WP_053492739.1 | WP_053492739.1 hypothetical protein |
| 6(1) | WP_251583290.1 | WP_251583290.1 hypothetical protein |
| 6(1) | WP_094870398.1 | WP_094870398.1 hypothetical protein |
| 6(1) | WP_042134266.1 | WP_042134266.1 hypothetical protein |
| 6(1) | WP_229521539.1 | WP_229521539.1 hypothetical protein |
| 6(1) | WP_062326147.1 | WP_062326147.1 hypothetical protein |
| 6(1) | WP_211023025.1 | WP_211023025.1 hypothetical protein |
| 6(1) | WP_044649164.1 | WP_044649164.1 hypothetical protein |
| 6(1) | WP_056700788.1 | WP_056700788.1 hypothetical protein |
| 6(1) | WP_142613002.1 | WP_142613002.1 hypothetical protein |
| 6(1) | WP_212980187.1 | WP_212980187.1 hypothetical protein |
| 6(1) | WP_036707101.1 | WP_036707101.1 hypothetical protein |
| 6(3) | WP_076135808.1 | WP_076135808.1 hypothetical protein |

| | | |
|---|---|---|
| 6(3) | WP_134910001.1 | WP_134910001.1 hypothetical protein |
| 6(3) | WP_071220205.1 | WP_071220205.1 hypothetical protein |
| 6(1) | WP_076286995.1 | WP_076286995.1 hypothetical protein |
| 6(1) | WP_183578449.1 | WP_183578449.1 hypothetical protein |
| 6(1) | WP_260065259.1 | WP_260065259.1 hypothetical protein |
| 6(1) | WP_009590693.1 | WP_009590693.1 hypothetical protein |
| 6(1) | WP_237167830.1 | WP_237167830.1 hypothetical protein |
| 6(1) | WP_179160624.1 | WP_179160624.1 hypothetical protein |
| 6(2) | WP_087433605.1 | WP_087433605.1 hypothetical protein |
| 6(1) | WP_216513489.1 | WP_216513489.1 hypothetical protein |
| 6(1) | WP_145021738.1 | WP_145021738.1 hypothetical protein |
| 6(1) | WP_205001942.1 | WP_205001942.1 hypothetical protein |
| 6(1) | WP_223067439.1 | WP_223067439.1 hypothetical protein |
| 6(1) | WP_017692351.1 | WP_017692351.1 hypothetical protein |
| 6(2) | WP_098746867.1 | WP_098746867.1 hypothetical protein |
| 6(2) | WP_013309988.1 | WP_013309988.1 hypothetical protein |
| 6(1) | WP_165175576.1 | WP_165175576.1 hypothetical protein |
| 6(1) | WP_068619113.1 | WP_068619113.1 hypothetical protein |
| 6(2) | WP_163245181.1 | WP_163245181.1 hypothetical protein |
| 6(1) | WP_236417213.1 | WP_236417213.1 hypothetical protein |
| 6(1) | WP_254912183.1 | WP_254912183.1 hypothetical protein |
| 6(1) | WP_220031160.1 | WP_220031160.1 hypothetical protein |
| 6(1) | WP_301249698.1 | WP_301249698.1 hypothetical protein |
| 6(1) | WP_307394631.1 | WP_307394631.1 hypothetical protein |
| 6(1) | WP_253440788.1 | WP_253440788.1 hypothetical protein |
| 6(2) | WP_076311049.1 | WP_076311049.1 hypothetical protein |
| 6(5) | WP_006211204.1 | WP_006211204.1 hypothetical protein |
| 6(1) | WP_113052566.1 | WP_113052566.1 hypothetical protein |
| 6(1) | WP_263880218.1 | WP_263880218.1 hypothetical protein |
| 6(1) | WP_152826364.1 | WP_152826364.1 hypothetical protein |
| 6(12) | WP_016821030.1 | WP_016821030.1 hypothetical protein |
| 6(1) | WP_272640515.1 | WP_272640515.1 hypothetical protein |
| 6(2) | WP_047909613.1 | WP_047909613.1 hypothetical protein |
| 6(1) | WP_301879166.1 | WP_301879166.1 hypothetical protein |
| 6(1) | WP_141136632.1 | WP_141136632.1 hypothetical protein |
| 6(4) | WP_013370785.1 | WP_013370785.1 hypothetical protein |
| 6(1) | WP_268623241.1 | WP_268623241.1 hypothetical protein |
| 6(1) | WP_113058718.1 | WP_113058718.1 hypothetical protein |
| 6(2) | WP_143802750.1 | WP_143802750.1 hypothetical protein |
| 6(1) | WP_169823379.1 | WP_169823379.1 hypothetical protein |
| 6(1) | WP_213644228.1 | WP_213644228.1 hypothetical protein |
| 6(1) | WP_017427966.1 | WP_017427966.1 hypothetical protein |
| 6(1) | WP_145153646.1 | WP_145153646.1 hypothetical protein |
| 6(1) | WP_144025950.1 | WP_144025950.1 hypothetical protein |
| 6(2) | WP_124114118.1 | WP_124114118.1 hypothetical protein |
| 6(1) | WP_191798526.1 | WP_191798526.1 hypothetical protein |
| 6(1) | WP_025716875.1 | WP_025716875.1 hypothetical protein |
| 6(1) | WP_204412730.1 | WP_204412730.1 hypothetical protein |
| 6(1) | WP_054958408.1 | WP_054958408.1 hypothetical protein |
| 6(1) | WP_099517106.1 | WP_099517106.1 hypothetical protein |
| 6(1) | WP_110899365.1 | WP_110899365.1 hypothetical protein |
| 6(1) | WP_138226025.1 | WP_138226025.1 hypothetical protein |
| 6(1) | WP_090908278.1 | WP_090908278.1 hypothetical protein |
| 6(1) | WP_197256866.1 | WP_197256866.1 hypothetical protein |
| 6(1) | WP_279830833.1 | WP_279830833.1 hypothetical protein |
| 6(1) | WP_127199770.1 | WP_127199770.1 hypothetical protein |
| 6(1) | WP_083189152.1 | WP_083189152.1 hypothetical protein |
| 6(7) | WP_015736243.1 | WP_015736243.1 hypothetical protein |
| 6(9) | WP_047842537.1 | WP_047842537.1 hypothetical protein |
| 6(1) | WP_062833880.1 | WP_062833880.1 hypothetical protein |
| 6(1) | WP_305755035.1 | WP_305755035.1 hypothetical protein |
| 7(1) | WP_146259787.1 | WP_146259787.1 hypothetical protein |
| 7(1) | WP_272045599.1 | WP_272045599.1 sigma 54-interacting transcriptional regulator |
| 7(1) | WP_241929669.1 | WP_241929669.1 response regulator transcription factor |
| 7(1) | WP_235286184.1 | WP_235286184.1 iron-siderophore ABC transporter substrate-binding protein |
| 7(1) | WP_015595764.1 | WP_015595764.1 response regulator transcription factor |
| 7(1) | WP_212695789.1 | WP_212695789.1 LytTR family DNA-binding domain-containing protein |
| 7(1) | WP_090674361.1 | WP_090674361.1 methyl-accepting chemotaxis protein |
| 7(3) | WP_076135825.1 | WP_076135825.1 response regulator transcription factor |
| 7(1) | WP_039871633.1 | WP_039871633.1 HD-GYP domain-containing protein |
| 7(1) | WP_074085374.1 | WP_074085374.1 sensor histidine kinase |
| 7(1) | WP_166244877.1 | WP_166244877.1 ABC transporter substrate-binding protein |
| 7(1) | WP_040729701.1 | WP_040729701.1 sigma-54-dependent Fis family transcriptional regulator |
| 7(1) | WP_270993668.1 | WP_270993668.1 LytTR family DNA-binding domain-containing protein |
| 7(1) | WP_022128265.1 | WP_022128265.1 response regulator transcription factor |
| 7(1) | WP_099517114.1 | WP_099517114.1 histidine kinase |
| 7(1) | WP_010676879.1 | WP_010676879.1 methyl-accepting chemotaxis protein |
| 7(1) | WP_051775654.1 | WP_051775654.1 methyl-accepting chemotaxis protein |
| 7(1) | WP_160037468.1 | WP_160037468.1 AraC family transcriptional regulator |
| 7(1) | WP_127484343.1 | WP_127484343.1 methyl-accepting chemotaxis protein |
| 7(1) | WP_242069516.1 | WP_242069516.1 sigma 54-interacting transcriptional regulator |
| 7(1) | WP_042210983.1 | WP_042210983.1 helix-turn-helix domain-containing protein |
| 7(1) | WP_166244125.1 | WP_166244125.1 helix-turn-helix domain-containing protein |
| 7(1) | WP_283892158.1 | WP_283892158.1 response regulator |
| 7(2) | WP_076124801.1 | WP_076124801.1 HAMP domain-containing sensor histidine kinase |
| 7(1) | WP_046217702.1 | WP_046217702.1 sigma-54-dependent transcriptional regulator |
| 7(1) | WP_076187095.1 | WP_076187095.1 response regulator transcription factor |
| 7(1) | WP_139867369.1 | WP_139867369.1 response regulator transcription factor |
| 7(1) | WP_253440820.1 | WP_253440820.1 AraC family transcriptional regulator |
| 7(1) | WP_204412705.1 | WP_204412705.1 sensor histidine kinase |
| 7(1) | WP_207746128.1 | WP_207746128.1 EAL domain-containing protein |
| 7(1) | WP_258208940.1 | WP_258208940.1 sigma 54-interacting transcriptional regulator |
| 7(1) | WP_090722909.1 | WP_090722909.1 histidine kinase |
| 7(1) | WP_183578410.1 | WP_183578410.1 AraC family transcriptional regulator |
| 7(1) | WP_179093400.1 | WP_179093400.1 helix-turn-helix domain-containing protein |
| 7(1) | WP_223067429.1 | WP_223067429.1 response regulator transcription factor |
| 7(1) | WP_042210989.1 | WP_042210989.1 response regulator |
| 7(1) | WP_274359795.1 | WP_274359795.1 response regulator |
| 7(1) | WP_126996280.1 | WP_126996280.1 AraC family transcriptional regulator |
| 7(1) | WP_053361026.1 | WP_053361026.1 response regulator transcription factor |
| 7(1) | WP_111273185.1 | WP_111273185.1 AraC family transcriptional regulator |
| 7(1) | WP_217035845.1 | WP_217035845.1 PAS domain S-box protein |
| 7(1) | WP_026478097.1 | WP_026478097.1 response regulator transcription factor |

| | | |
|---|---|---|
| 7(1) | WP_094870394.1 | WP_094870394.1 AraC family transcriptional regulator |
| 7(1) | WP_212695780.1 | WP_212695780.1 HD-GYP domain-containing protein |
| 7(1) | WP_188535098.1 | WP_188535098.1 methyl-accepting chemotaxis protein |
| 7(1) | WP_268058032.1 | WP_268058032.1 EAL domain-containing protein |
| 7(1) | WP_042210990.1 | WP_042210990.1 histidine kinase |
| 7(1) | WP_304216847.1 | WP_304216847.1 methyl-accepting chemotaxis protein |
| 7(1) | WP_129478258.1 | WP_129478258.1 response regulator |
| 7(1) | WP_076187097.1 | WP_076187097.1 HAMP domain-containing sensor histidine kinase |
| 7(1) | WP_196048857.1 | WP_196048857.1 EAL domain-containing protein |
| 7(1) | WP_129478256.1 | WP_129478256.1 sensor histidine kinase |
| 7(2) | WP_295628304.1 | WP_295628304.1 LytTR family DNA-binding domain-containing protein |
| 7(17) | WP_003745030.1 | WP_003745030.1 LytTR family DNA-binding domain-containing protein |
| 7(1) | WP_099521441.1 | WP_099521441.1 helix-turn-helix domain-containing protein |
| 7(1) | WP_251548749.1 | WP_251548749.1 GGDEF domain-containing protein |
| 7(1) | WP_208919046.1 | WP_208919046.1 response regulator transcription factor |
| 7(1) | WP_052352967.1 | WP_052352967.1 GGDEF domain-containing protein |
| 7(1) | WP_076210942.1 | WP_076210942.1 AraC family transcriptional regulator |
| 7(1) | WP_308110119.1 | WP_308110119.1 response regulator transcription factor |
| 7(1) | WP_043067141.1 | WP_043067141.1 LytTR family DNA-binding domain-containing protein |
| 7(1) | WP_281937363.1 | WP_281937363.1 methyl-accepting chemotaxis protein |
| 7(1) | WP_197234712.1 | WP_197234712.1 sigma-54-dependent transcriptional regulator |
| 7(2) | WP_003728979.1 | WP_003728979.1 LytTR family DNA-binding domain-containing protein |
| 7(1) | WP_081943722.1 | WP_081943722.1 sigma 54-interacting transcriptional regulator |
| 7(1) | WP_253440816.1 | WP_253440816.1 hypothetical protein |
| 7(1) | WP_017692332.1 | WP_017692332.1 AraC family transcriptional regulator |
| 7(1) | WP_235714690.1 | WP_235714690.1 diguanylate cyclase |
| 7(1) | WP_251557386.1 | WP_251557386.1 ATP-binding protein |
| 7(1) | WP_227219176.1 | WP_227219176.1 HAMP domain-containing sensor histidine kinase |
| 7(1) | WP_251548771.1 | WP_251548771.1 diguanylate cyclase |
| 7(1) | WP_243495654.1 | WP_243495654.1 methyl-accepting chemotaxis protein |
| 7(1) | WP_103655293.1 | WP_103655293.1 HAMP domain-containing histidine kinase |
| 7(1) | WP_244194181.1 | WP_244194181.1 sigma 54-interacting transcriptional regulator |
| 7(1) | WP_024632700.1 | WP_024632700.1 hypothetical protein |
| 7(1) | WP_254366018.1 | WP_254366018.1 response regulator |
| 7(1) | WP_239458682.1 | WP_239458682.1 response regulator transcription factor |
| 7(1) | WP_212958792.1 | WP_212958792.1 ATP-binding protein |
| 7(1) | WP_076311055.1 | WP_076311055.1 histidine kinase |
| 7(1) | WP_244533154.1 | WP_244533154.1 response regulator |
| 7(1) | WP_242251798.1 | WP_242251798.1 response regulator transcription factor |
| 7(1) | WP_208926639.1 | WP_208926639.1 LytTR family DNA-binding domain-containing protein |
| 7(3) | WP_076100020.1 | WP_076100020.1 response regulator transcription factor |
| 7(1) | WP_174812168.1 | WP_174812168.1 hypothetical protein |
| 7(1) | WP_210087130.1 | WP_210087130.1 response regulator transcription factor |
| 7(1) | WP_068536342.1 | WP_068536342.1 diguanylate cyclase |
| 7(1) | WP_258208941.1 | WP_258208941.1 PucR family transcriptional regulator |
| 7(1) | WP_257535704.1 | WP_257535704.1 sensor histidine kinase |
| 7(1) | WP_143760312.1 | WP_143760312.1 hypothetical protein |
| 7(1) | WP_276917296.1 | WP_276917296.1 LytTR family DNA-binding domain-containing protein |
| 7(1) | WP_144611193.1 | WP_144611193.1 response regulator transcription factor |
| 7(1) | WP_026478098.1 | WP_026478098.1 HAMP domain-containing sensor histidine kinase |
| 7(3) | WP_076104568.1 | WP_076104568.1 HAMP domain-containing sensor histidine kinase |
| 7(1) | WP_205053319.1 | WP_205053319.1 histidine kinase |
| 7(1) | WP_251598142.1 | WP_251598142.1 methyl-accepting chemotaxis protein |
| 7(2) | WP_246873638.1 | WP_246873638.1 sigma 54-interacting transcriptional regulator |
| 7(1) | WP_017692334.1 | WP_017692334.1 hypothetical protein |
| 7(1) | WP_276867661.1 | WP_276867661.1 ATP-binding protein |
| 7(2) | WP_281813103.1 | WP_281813103.1 LytTR family DNA-binding domain-containing protein |
| 7(1) | WP_039871642.1 | WP_039871642.1 HD-GYP domain-containing protein |
| 7(1) | WP_149871690.1 | WP_149871690.1 response regulator transcription factor |
| 7(1) | WP_139867367.1 | WP_139867367.1 HAMP domain-containing histidine kinase |
| 7(1) | WP_121610581.1 | WP_121610581.1 response regulator |
| 7(1) | WP_249897327.1 | WP_249897327.1 AraC family transcriptional regulator |
| 7(1) | WP_301167488.1 | WP_301167488.1 histidine kinase |
| 7(44) | WP_185336684.1 | WP_185336684.1 LytTR family DNA-binding domain-containing protein |
| 7(1) | WP_224839857.1 | WP_224839857.1 PAS domain S-box protein |
| 7(1) | WP_106496009.1 | WP_106496009.1 response regulator |
| 7(1) | WP_290407050.1 | WP_290407050.1 helix-turn-helix domain-containing protein |
| 7(1) | WP_212981553.1 | WP_212981553.1 AraC family transcriptional regulator |
| 7(1) | WP_088830547.1 | WP_088830547.1 response regulator |
| 7(1) | WP_064477383.1 | WP_064477383.1 response regulator transcription factor |
| 7(1) | WP_089965926.1 | WP_089965926.1 sensor histidine kinase |
| 7(1) | WP_204666329.1 | WP_204666329.1 ATP-binding protein |
| 7(1) | WP_168976280.1 | WP_168976280.1 siderophore ABC transporter substrate-binding protein |
| 7(1) | WP_308082237.1 | WP_308082237.1 diguanylate cyclase |
| 7(1) | WP_198059698.1 | WP_198059698.1 sigma 54-interacting transcriptional regulator |
| 7(1) | WP_043069421.1 | WP_043069421.1 siderophore ABC transporter substrate-binding protein |
| 7(1) | WP_249724670.1 | WP_249724670.1 sigma 54-interacting transcriptional regulator |
| 7(1) | WP_199613915.1 | WP_199613915.1 AraC family transcriptional regulator |
| 7(1) | WP_244723724.1 | WP_244723724.1 helix-turn-helix domain-containing protein |
| 7(1) | WP_090761801.1 | WP_090761801.1 HAMP domain-containing sensor histidine kinase |
| 7(1) | WP_145153688.1 | WP_145153688.1 hypothetical protein |
| 7(1) | WP_099517113.1 | WP_099517113.1 response regulator |
| 7(1) | WP_210087129.1 | WP_210087129.1 HAMP domain-containing sensor histidine kinase |
| 7(1) | WP_290407030.1 | WP_290407030.1 HAMP domain-containing sensor histidine kinase |
| 7(1) | WP_247083065.1 | WP_247083065.1 sensor histidine kinase |
| 7(1) | WP_184183065.1 | WP_184183065.1 AraC family transcriptional regulator |
| 7(1) | WP_090239196.1 | WP_090239196.1 sensor histidine kinase |
| 7(1) | WP_235286187.1 | WP_235286187.1 AraC family transcriptional regulator |
| 7(1) | WP_248063415.1 | WP_248063415.1 hypothetical protein |
| 7(1) | WP_212693240.1 | WP_212693240.1 LytTR family DNA-binding domain-containing protein |
| 7(1) | WP_295628292.1 | WP_295628292.1 EAL domain-containing protein |
| 7(1) | WP_210087126.1 | WP_210087126.1 hypothetical protein |
| 7(1) | WP_181438750.1 | WP_181438750.1 iron-siderophore ABC transporter substrate-binding protein |
| 7(1) | WP_251596660.1 | WP_251596660.1 response regulator |
| 7(1) | WP_180271141.1 | WP_180271141.1 methyl-accepting chemotaxis protein |
| 7(4) | WP_003718132.1 | WP_003718132.1 LytTR family DNA-binding domain-containing protein |
| 7(1) | WP_053403842.1 | WP_053403842.1 helix-turn-helix domain-containing protein |
| 7(1) | WP_099361321.1 | WP_099361321.1 diguanylate cyclase |
| 7(1) | WP_241993929.1 | WP_241993929.1 sensor histidine kinase |
| 7(1) | WP_213117463.1 | WP_213117463.1 PAS domain S-box protein |
| 7(1) | WP_307284666.1 | WP_307284666.1 diguanylate cyclase |
| 7(1) | WP_029881472.1 | WP_029881472.1 AraC family transcriptional regulator |
| 7(1) | WP_045518290.1 | WP_045518290.1 diguanylate cyclase |
| 7(2) | WP_281813111.1 | WP_281813111.1 HD domain-containing phosphohydrolase |

| | | |
|---|---|---|
| 7(1) | WP_166704999.1 | WP_166704999.1 response regulator transcription factor |
| 7(1) | WP_211082916.1 | WP_211082916.1 hypothetical protein |
| 7(1) | WP_212958794.1 | WP_212958794.1 response regulator transcription factor |
| 7(1) | WP_231094837.1 | WP_231094837.1 EAL domain-containing protein |
| 7(1) | WP_251456518.1 | WP_251456518.1 sigma-54-dependent transcriptional regulator |
| 7(1) | WP_021624811.1 | WP_021624811.1 LytTR family DNA-binding domain-containing protein |
| 7(1) | WP_150364183.1 | WP_150364183.1 hypothetical protein |
| 7(1) | WP_290407031.1 | WP_290407031.1 response regulator transcription factor |
| 7(1) | WP_295632015.1 | WP_295632015.1 EAL domain-containing protein |
| 7(1) | WP_010496614.1 | WP_010496614.1 methyl-accepting chemotaxis protein |
| 7(1) | WP_183600881.1 | WP_183600881.1 AraC family transcriptional regulator |
| 7(1) | WP_234489639.1 | WP_234489639.1 helix-turn-helix domain-containing protein |
| 7(1) | WP_053564622.1 | WP_053564622.1 response regulator transcription factor |
| 7(1) | WP_087186832.1 | WP_087186832.1 response regulator transcription factor |
| 7(2) | WP_076191082.1 | WP_076191082.1 response regulator transcription factor |
| 7(1) | WP_071736288.1 | WP_071736288.1 response regulator transcription factor |
| 7(1) | WP_189031018.1 | WP_189031018.1 helix-turn-helix domain-containing protein |
| 7(1) | WP_240913180.1 | WP_240913180.1 helix-turn-helix domain-containing protein |
| 7(1) | WP_169238710.1 | WP_169238710.1 response regulator |
| 7(1) | WP_198056466.1 | WP_198056466.1 sigma 54-interacting transcriptional regulator |
| 7(1) | WP_281477391.1 | WP_281477391.1 sigma 54-interacting transcriptional regulator |
| 7(1) | WP_231417850.1 | WP_231417850.1 response regulator |
| 7(1) | WP_025848639.1 | WP_025848639.1 helix-turn-helix domain-containing protein |
| 7(1) | WP_255404589.1 | WP_255404589.1 sigma 54-interacting transcriptional regulator |
| 7(1) | WP_044037305.1 | WP_044037305.1 ABC transporter substrate-binding protein |
| 7(2) | WP_200372571.1 | WP_200372571.1 LytTR family DNA-binding domain-containing protein |
| 7(1) | WP_301292477.1 | WP_301292477.1 methyl-accepting chemotaxis protein |
| 7(1) | WP_213143932.1 | WP_213143932.1 PAS domain S-box protein |
| 7(1) | WP_207722184.1 | WP_207722184.1 response regulator transcription factor |
| 7(1) | WP_242172364.1 | WP_242172364.1 helix-turn-helix domain-containing protein |
| 7(1) | WP_090239193.1 | WP_090239193.1 response regulator |
| 7(1) | WP_088830541.1 | WP_088830541.1 methyl-accepting chemotaxis protein |
| 7(1) | WP_243895991.1 | WP_243895991.1 CHASE3 domain-containing protein |
| 7(1) | WP_240764707.1 | WP_240764707.1 sigma 54-interacting transcriptional regulator |
| 7(1) | WP_244867384.1 | WP_244867384.1 helix-turn-helix domain-containing protein |
| 7(1) | WP_000504574.1 | WP_000504574.1 response regulator transcription factor |
| 7(1) | WP_237096524.1 | WP_237096524.1 sigma 54-interacting transcriptional regulator |
| 7(1) | WP_150457777.1 | WP_150457777.1 HAMP domain-containing protein |
| 7(1) | WP_197074338.1 | WP_197074338.1 helix-turn-helix domain-containing protein |
| 7(1) | WP_199613893.1 | WP_199613893.1 sensor histidine kinase |
| 7(1) | WP_307192888.1 | WP_307192888.1 PucR family transcriptional regulator ligand-binding domain-containing protein |
| 7(1) | WP_077852410.1 | WP_077852410.1 AraC family transcriptional regulator |
| 7(3) | WP_076135823.1 | WP_076135823.1 HAMP domain-containing sensor histidine kinase |
| 7(1) | WP_042462791.1 | WP_042462791.1 GGDEF domain-containing protein |
| 7(1) | WP_244918732.1 | WP_244918732.1 sigma 54-interacting transcriptional regulator |
| 7(1) | WP_047909619.1 | WP_047909619.1 histidine kinase |
| 7(1) | WP_190945004.1 | WP_190945004.1 AraC family transcriptional regulator |
| 7(1) | WP_216791227.1 | WP_216791227.1 methyl-accepting chemotaxis protein |
| 7(1) | WP_211082915.1 | WP_211082915.1 AraC family transcriptional regulator |
| 7(1) | WP_260986795.1 | WP_260986795.1 helix-turn-helix domain-containing protein |
| 7(1) | WP_235988996.1 | WP_235988996.1 sensor histidine kinase |
| 7(1) | WP_074108307.1 | WP_074108307.1 sensor histidine kinase |
| 7(1) | WP_245859370.1 | WP_245859370.1 HAMP domain-containing sensor histidine kinase |
| 7(1) | WP_043067153.1 | WP_043067153.1 siderophore ABC transporter substrate-binding protein |
| 7(1) | WP_055277401.1 | WP_055277401.1 EAL domain-containing protein |
| 7(1) | WP_085117893.1 | WP_085117893.1 sigma-54-dependent transcriptional regulator |
| 7(1) | WP_245368174.1 | WP_245368174.1 HD-GYP domain-containing protein |
| 7(1) | WP_102371687.1 | WP_102371687.1 response regulator transcription factor |
| 7(1) | WP_280788637.1 | WP_280788637.1 AraC family transcriptional regulator |
| 7(1) | WP_251596662.1 | WP_251596662.1 sensor histidine kinase |
| 7(1) | WP_241929668.1 | WP_241929668.1 HAMP domain-containing sensor histidine kinase |
| 7(1) | WP_000694910.1 | WP_000694910.1 ATP-binding protein |
| 7(1) | WP_213649985.1 | WP_213649985.1 sensor histidine kinase |
| 7(1) | WP_259392003.1 | WP_259392003.1 sigma 54-interacting transcriptional regulator |
| 7(1) | WP_212949017.1 | WP_212949017.1 AraC family transcriptional regulator |
| 7(1) | WP_212695778.1 | WP_212695778.1 HD domain-containing phosphohydrolase |
| 7(1) | WP_221697952.1 | WP_221697952.1 EAL domain-containing protein |
| 7(1) | WP_251583286.1 | WP_251583286.1 histidine kinase |
| 7(1) | WP_199613892.1 | WP_199613892.1 response regulator |
| 7(1) | WP_214705996.1 | WP_214705996.1 HD domain-containing phosphohydrolase |
| 7(1) | WP_256209538.1 | WP_256209538.1 response regulator |
| 7(1) | WP_283892159.1 | WP_283892159.1 histidine kinase |
| 7(1) | WP_102371686.1 | WP_102371686.1 ATP-binding protein |
| 7(1) | WP_275540592.1 | WP_275540592.1 HD domain-containing phosphohydrolase |
| 7(1) | WP_111147816.1 | WP_111147816.1 helix-turn-helix domain-containing protein |
| 7(1) | WP_166705000.1 | WP_166705000.1 ATP-binding protein |
| 7(3) | WP_076187093.1 | WP_076187093.1 HD-GYP domain-containing protein |
| 7(1) | WP_256702678.1 | WP_256702678.1 response regulator |
| 7(1) | WP_061975557.1 | WP_061975557.1 response regulator |
| 7(1) | WP_127459438.1 | WP_127459438.1 methyl-accepting chemotaxis protein |
| 7(1) | WP_094870393.1 | WP_094870393.1 AraC family transcriptional regulator |
| 7(1) | WP_235286195.1 | WP_235286195.1 methyl-accepting chemotaxis protein |
| 7(1) | WP_249897325.1 | WP_249897325.1 helix-turn-helix domain-containing protein |
| 7(1) | WP_151630598.1 | WP_151630598.1 response regulator transcription factor |
| 7(1) | WP_251426380.1 | WP_251426380.1 ATP-binding protein |
| 7(1) | WP_021624834.1 | WP_021624834.1 siderophore ABC transporter substrate-binding protein |
| 7(1) | WP_307199916.1 | WP_307199916.1 GGDEF domain-containing protein |
| 7(1) | WP_035190288.1 | WP_035190288.1 PocR ligand-binding domain-containing protein |
| 7(1) | WP_216518276.1 | WP_216518276.1 hypothetical protein |
| 7(1) | WP_090674347.1 | WP_090674347.1 response regulator |
| 7(1) | WP_091155364.1 | WP_091155364.1 histidine kinase |
| 7(1) | WP_054972096.1 | WP_054972096.1 methyl-accepting chemotaxis protein |
| 7(1) | WP_074565298.1 | WP_074565298.1 response regulator transcription factor |
| 7(1) | WP_302058396.1 | WP_302058396.1 methyl-accepting chemotaxis protein |
| 7(1) | WP_213427003.1 | WP_213427003.1 AraC family transcriptional regulator |
| 7(1) | WP_239458685.1 | WP_239458685.1 HAMP domain-containing sensor histidine kinase |
| 7(3) | WP_273322271.1 | WP_273322271.1 HD domain-containing phosphohydrolase |
| 7(1) | WP_042462840.1 | WP_042462840.1 response regulator |
| 7(1) | WP_077833100.1 | WP_077833100.1 AraC family transcriptional regulator |
| 7(1) | WP_302878976.1 | WP_302878976.1 methyl-accepting chemotaxis protein |
| 7(1) | WP_151630597.1 | WP_151630597.1 ATP-binding protein |
| 7(1) | WP_128631290.1 | WP_128631290.1 methyl-accepting chemotaxis protein |
| 7(1) | WP_251610741.1 | WP_251610741.1 helix-turn-helix domain-containing protein |

| | | |
|---|---|---|
| 7(1) | WP_251548741.1 | WP_251548741.1 GGDEF domain-containing phosphodiesterase |
| 7(2) | WP_273321545.1 | WP_273321545.1 LytTR family DNA-binding domain-containing protein |
| 7(1) | WP_076113445.1 | WP_076113445.1 helix-turn-helix domain-containing protein |
| 7(1) | WP_178075824.1 | WP_178075824.1 sensor histidine kinase |
| 7(1) | WP_256992242.1 | WP_256992242.1 helix-turn-helix domain-containing protein |
| 7(1) | WP_208919048.1 | WP_208919048.1 HAMP domain-containing sensor histidine kinase |
| 7(1) | WP_061975559.1 | WP_061975559.1 sensor histidine kinase |
| 7(1) | WP_289376832.1 | WP_289376832.1 hypothetical protein |
| 7(1) | WP_289483772.1 | WP_289483772.1 response regulator |
| 7(1) | WP_108534962.1 | WP_108534962.1 methyl-accepting chemotaxis protein |
| 7(1) | WP_181603046.1 | WP_181603046.1 ATP-binding protein |
| 7(1) | WP_186333849.1 | WP_186333849.1 AraC family transcriptional regulator |
| 7(3) | WP_076100022.1 | WP_076100022.1 HD-GYP domain-containing protein |
| 7(1) | WP_178075825.1 | WP_178075825.1 response regulator |
| 7(1) | WP_053564621.1 | WP_053564621.1 ATP-binding protein |
| 7(1) | WP_169505066.1 | WP_169505066.1 response regulator |
| 7(1) | WP_087186830.1 | WP_087186830.1 response regulator transcription factor |
| 7(1) | WP_054972105.1 | WP_054972105.1 response regulator |
| 7(1) | WP_055277403.1 | WP_055277403.1 hypothetical protein |
| 7(1) | WP_231417849.1 | WP_231417849.1 histidine kinase |
| 7(1) | WP_180271577.1 | WP_180271577.1 methyl-accepting chemotaxis protein |
| 7(1) | WP_184183061.1 | WP_184183061.1 hypothetical protein |
| 7(1) | WP_076113460.1 | WP_076113460.1 AraC family transcriptional regulator |
| 7(1) | WP_290401982.1 | WP_290401982.1 hypothetical protein |
| 7(1) | WP_010250278.1 | WP_010250278.1 LytTR family DNA-binding domain-containing protein |
| 7(2) | WP_185639023.1 | WP_185639023.1 LytTR family DNA-binding domain-containing protein |
| 7(1) | WP_289483774.1 | WP_289483774.1 histidine kinase |
| 7(1) | WP_113672704.1 | WP_113672704.1 LytTR family DNA-binding domain-containing protein |
| 7(1) | WP_000504583.1 | WP_000504583.1 response regulator transcription factor |
| 7(1) | WP_276867711.1 | WP_276867711.1 ATP-binding protein |
| 7(1) | WP_132008136.1 | WP_132008136.1 response regulator |
| 7(1) | WP_251583287.1 | WP_251583287.1 response regulator |
| 7(1) | WP_138322080.1 | WP_138322080.1 response regulator transcription factor |
| 7(1) | WP_025848618.1 | WP_025848618.1 methyl-accepting chemotaxis protein |
| 7(1) | WP_111147808.1 | WP_111147808.1 methyl-accepting chemotaxis protein |
| 7(1) | WP_293755522.1 | WP_293755522.1 LytTR family DNA-binding domain-containing protein |
| 7(1) | WP_195383892.1 | WP_195383892.1 LytTR family DNA-binding domain-containing protein |
| 7(1) | WP_289475086.1 | WP_289475086.1 histidine kinase |
| 7(2) | WP_063188056.1 | WP_063188056.1 methyl-accepting chemotaxis protein |
| 7(1) | WP_284036506.1 | WP_284036506.1 PAS domain S-box protein |
| 7(1) | WP_168976293.1 | WP_168976293.1 LytTR family DNA-binding domain-containing protein |
| 7(1) | WP_301167487.1 | WP_301167487.1 response regulator |
| 7(1) | WP_288218591.1 | WP_288218591.1 EAL domain-containing protein |
| 7(2) | WP_000694900.1 | WP_000694900.1 ATP-binding protein |
| 7(1) | WP_308723534.1 | WP_308723534.1 hypothetical protein |
| 7(1) | WP_214705997.1 | WP_214705997.1 response regulator transcription factor |
| 7(1) | WP_069011428.1 | WP_069011428.1 helix-turn-helix domain-containing protein |
| 7(1) | WP_051624143.1 | WP_051624143.1 ATP-binding protein |
| 7(1) | WP_102371772.1 | WP_102371772.1 response regulator transcription factor |
| 7(1) | WP_233280991.1 | WP_233280991.1 CHASE3 domain-containing protein |
| 7(1) | WP_251596668.1 | WP_251596668.1 methyl-accepting chemotaxis protein |
| 7(1) | WP_211022803.1 | WP_211022803.1 helix-turn-helix domain-containing protein |

| | | |
|---|---|---|
| 7(2) | WP_077893578.1 | WP_077893578.1 AraC family transcriptional regulator |
| 7(1) | WP_212695776.1 | WP_212695776.1 hybrid sensor histidine kinase/response regulator |
| 7(2) | WP_057897618.1 | WP_057897618.1 HD-GYP domain-containing protein |
| 7(1) | WP_019127580.1 | WP_019127580.1 response regulator transcription factor |
| 7(1) | WP_181603051.1 | WP_181603051.1 HAMP domain-containing sensor histidine kinase |
| 7(1) | WP_144611194.1 | WP_144611194.1 ATP-binding protein |
| 7(1) | WP_308723536.1 | WP_308723536.1 AraC family transcriptional regulator |
| 7(1) | WP_274336972.1 | WP_274336972.1 response regulator |
| 7(2) | WP_240927214.1 | WP_240927214.1 sigma 54-interacting transcriptional regulator |
| 7(1) | WP_289475083.1 | WP_289475083.1 response regulator |
| 7(1) | WP_304216838.1 | WP_304216838.1 diguanylate cyclase |
| 7(2) | WP_011700917.1 | WP_011700917.1 LytTR family DNA-binding domain-containing protein |
| 8(1) | WP_216513494.1 | WP_216513494.1 SDR family NAD(P)-dependent oxidoreductase |
| 8(1) | WP_191798532.1 | WP_191798532.1 SDR family NAD(P)-dependent oxidoreductase |
| 8(1) | WP_016821034.1 | WP_016821034.1 SDR family NAD(P)-dependent oxidoreductase |
| 8(1) | WP_229522852.1 | WP_229522852.1 SDR family NAD(P)-dependent oxidoreductase |
| 8(2) | WP_076292683.1 | WP_076292683.1 SDR family NAD(P)-dependent oxidoreductase |
| 8(1) | WP_236417225.1 | WP_236417225.1 SDR family NAD(P)-dependent oxidoreductase |
| 8(1) | WP_090722897.1 | WP_090722897.1 SDR family NAD(P)-dependent oxidoreductase |
| 8(1) | WP_090993629.1 | WP_090993629.1 SDR family NAD(P)-dependent oxidoreductase |
| 8(1) | WP_251548739.1 | WP_251548739.1 SDR family NAD(P)-dependent oxidoreductase |
| 8(1) | WP_127199776.1 | WP_127199776.1 SDR family NAD(P)-dependent oxidoreductase |
| 8(1) | WP_054958414.1 | WP_054958414.1 SDR family NAD(P)-dependent oxidoreductase |
| 8(2) | WP_145329911.1 | WP_145329911.1 SDR family NAD(P)-dependent oxidoreductase |
| 8(2) | WP_033100005.1 | WP_033100005.1 SDR family oxidoreductase |
| 8(1) | WP_279830794.1 | WP_279830794.1 SDR family NAD(P)-dependent oxidoreductase |
| 8(2) | WP_063208772.1 | WP_063208772.1 SDR family NAD(P)-dependent oxidoreductase |
| 8(1) | WP_136604820.1 | WP_136604820.1 SDR family NAD(P)-dependent oxidoreductase |
| 8(2) | WP_090735419.1 | WP_090735419.1 SDR family NAD(P)-dependent oxidoreductase |
| 8(1) | WP_018664829.1 | WP_018664829.1 SDR family oxidoreductase |
| 8(2) | WP_124114122.1 | WP_124114122.1 SDR family NAD(P)-dependent oxidoreductase |
| 8(1) | WP_305022917.1 | WP_305022917.1 SDR family NAD(P)-dependent oxidoreductase |
| 8(1) | WP_234121617.1 | WP_234121617.1 SDR family NAD(P)-dependent oxidoreductase |
| 8(1) | WP_108465355.1 | WP_108465355.1 SDR family NAD(P)-dependent oxidoreductase |
| 8(1) | WP_076311046.1 | WP_076311046.1 SDR family NAD(P)-dependent oxidoreductase |
| 8(1) | WP_142613005.1 | WP_142613005.1 SDR family NAD(P)-dependent oxidoreductase |
| 8(2) | WP_283655697.1 | WP_283655697.1 SDR family NAD(P)-dependent oxidoreductase |
| 8(1) | WP_025716880.1 | WP_025716880.1 SDR family NAD(P)-dependent oxidoreductase |
| 8(2) | WP_056700798.1 | WP_056700798.1 SDR family NAD(P)-dependent oxidoreductase |
| 8(2) | WP_194230398.1 | WP_194230398.1 SDR family NAD(P)-dependent oxidoreductase |
| 8(1) | WP_192510489.1 | WP_192510489.1 SDR family NAD(P)-dependent oxidoreductase |
| 8(1) | WP_236334359.1 | WP_236334359.1 SDR family oxidoreductase |
| 8(1) | WP_036670438.1 | WP_036670438.1 SDR family NAD(P)-dependent oxidoreductase |
| 8(1) | WP_308721985.1 | WP_308721985.1 SDR family NAD(P)-dependent oxidoreductase |
| 8(2) | WP_090896199.1 | WP_090896199.1 SDR family NAD(P)-dependent oxidoreductase |
| 8(3) | WP_028542686.1 | WP_028542686.1 SDR family NAD(P)-dependent oxidoreductase |
| 8(2) | WP_138226020.1 | WP_138226020.1 SDR family NAD(P)-dependent oxidoreductase |
| 8(1) | WP_305755028.1 | WP_305755028.1 SDR family NAD(P)-dependent oxidoreductase |
| 8(3) | WP_210095105.1 | WP_210095105.1 SDR family NAD(P)-dependent oxidoreductase |
| 8(3) | WP_039273536.1 | WP_039273536.1 SDR family NAD(P)-dependent oxidoreductase |
| 8(1) | WP_241933975.1 | WP_241933975.1 SDR family NAD(P)-dependent oxidoreductase |

| | | |
|---|---|---|
| 8(3) | WP_113753073.1 | WP_113753073.1 SDR family oxidoreductase |
| 8(1) | WP_043067157.1 | WP_043067157.1 glucose 1-dehydrogenase |
| 8(1) | WP_183600900.1 | WP_183600900.1 SDR family oxidoreductase |
| 8(1) | WP_068496243.1 | WP_068496243.1 SDR family NAD(P)-dependent oxidoreductase |
| 8(1) | WP_091155389.1 | WP_091155389.1 SDR family NAD(P)-dependent oxidoreductase |
| 8(1) | WP_198059697.1 | WP_198059697.1 SDR family oxidoreductase |
| 8(1) | WP_102997921.1 | WP_102997921.1 SDR family NAD(P)-dependent oxidoreductase |
| 8(1) | WP_076289576.1 | WP_076289576.1 SDR family NAD(P)-dependent oxidoreductase |
| 8(1) | WP_010346278.1 | WP_010346278.1 SDR family NAD(P)-dependent oxidoreductase |
| 8(1) | WP_137062804.1 | WP_137062804.1 SDR family NAD(P)-dependent oxidoreductase |
| 8(1) | WP_109999895.1 | WP_109999895.1 SDR family NAD(P)-dependent oxidoreductase |
| 8(1) | WP_244719654.1 | WP_244719654.1 SDR family NAD(P)-dependent oxidoreductase |
| 8(1) | WP_095266202.1 | WP_095266202.1 SDR family NAD(P)-dependent oxidoreductase |
| 8(1) | WP_076113461.1 | WP_076113461.1 SDR family NAD(P)-dependent oxidoreductase |
| 8(1) | WP_307537575.1 | WP_307537575.1 SDR family NAD(P)-dependent oxidoreductase |
| 8(1) | WP_254759284.1 | WP_254759284.1 SDR family NAD(P)-dependent oxidoreductase |
| 8(4) | WP_113058720.1 | WP_113058720.1 SDR family NAD(P)-dependent oxidoreductase |
| 8(1) | WP_039273521.1 | WP_039273521.1 SDR family oxidoreductase |
| 8(1) | WP_025685037.1 | WP_025685037.1 SDR family NAD(P)-dependent oxidoreductase |
| 8(1) | WP_074094618.1 | WP_074094618.1 SDR family NAD(P)-dependent oxidoreductase |
| 8(1) | WP_144424633.1 | WP_144424633.1 SDR family NAD(P)-dependent oxidoreductase |
| 8(1) | WP_207383566.1 | WP_207383566.1 SDR family NAD(P)-dependent oxidoreductase |
| 8(1) | WP_110892962.1 | WP_110892962.1 SDR family NAD(P)-dependent oxidoreductase |
| 8(1) | WP_094593879.1 | WP_094593879.1 SDR family NAD(P)-dependent oxidoreductase |
| 8(1) | WP_044649162.1 | WP_044649162.1 SDR family NAD(P)-dependent oxidoreductase |
| 8(1) | WP_018664832.1 | WP_018664832.1 SDR family oxidoreductase |
| 8(1) | WP_285079439.1 | WP_285079439.1 SDR family NAD(P)-dependent oxidoreductase |
| 8(1) | WP_152381397.1 | WP_152381397.1 SDR family NAD(P)-dependent oxidoreductase |
| 8(1) | WP_149096532.1 | WP_149096532.1 SDR family NAD(P)-dependent oxidoreductase |
| 8(1) | WP_190299139.1 | WP_190299139.1 SDR family NAD(P)-dependent oxidoreductase |
| 8(1) | WP_204445243.1 | WP_204445243.1 SDR family oxidoreductase |
| 8(1) | WP_272640518.1 | WP_272640518.1 SDR family NAD(P)-dependent oxidoreductase |
| 8(4) | WP_019392102.1 | WP_019392102.1 SDR family oxidoreductase |
| 8(13) | WP_023988269.1 | WP_023988269.1 SDR family NAD(P)-dependent oxidoreductase |
| 8(1) | WP_169505058.1 | WP_169505058.1 SDR family NAD(P)-dependent oxidoreductase |
| 8(1) | WP_160037491.1 | WP_160037491.1 SDR family NAD(P)-dependent oxidoreductase |
| 8(3) | WP_031335648.1 | WP_031335648.1 SDR family oxidoreductase |
| 8(1) | WP_263324543.1 | WP_263324543.1 SDR family NAD(P)-dependent oxidoreductase |
| 8(1) | WP_091017190.1 | WP_091017190.1 SDR family NAD(P)-dependent oxidoreductase |
| 8(1) | WP_250244823.1 | WP_250244823.1 SDR family NAD(P)-dependent oxidoreductase |
| 8(1) | WP_264929203.1 | WP_264929203.1 SDR family oxidoreductase |
| 8(1) | WP_209861635.1 | WP_209861635.1 SDR family NAD(P)-dependent oxidoreductase |
| 8(1) | WP_200894944.1 | WP_200894944.1 SDR family NAD(P)-dependent oxidoreductase |
| 8(1) | WP_212949034.1 | WP_212949034.1 SDR family NAD(P)-dependent oxidoreductase |
| 8(1) | WP_090921046.1 | WP_090921046.1 SDR family NAD(P)-dependent oxidoreductase |
| 8(1) | WP_134909994.1 | WP_134909994.1 SDR family NAD(P)-dependent oxidoreductase |
| 8(1) | WP_268583871.1 | WP_268583871.1 SDR family oxidoreductase |
| 8(1) | WP_111619119.1 | WP_111619119.1 SDR family NAD(P)-dependent oxidoreductase |
| 8(1) | WP_243518737.1 | WP_243518737.1 SDR family oxidoreductase |
| 8(1) | WP_069011427.1 | WP_069011427.1 SDR family oxidoreductase |
| 8(11) | WP_017689067.1 | WP_017689067.1 SDR family NAD(P)-dependent oxidoreductase |
| 8(1) | WP_113713138.1 | WP_113713138.1 SDR family oxidoreductase |
| 8(4) | WP_163872772.1 | WP_163872772.1 SDR family NAD(P)-dependent oxidoreductase |
| 8(1) | WP_160496926.1 | WP_160496926.1 SDR family NAD(P)-dependent oxidoreductase |
| 8(4) | WP_006211197.1 | WP_006211197.1 SDR family NAD(P)-dependent oxidoreductase |
| 8(11) | WP_017427968.1 | WP_017427968.1 SDR family NAD(P)-dependent oxidoreductase |
| 8(1) | WP_090761791.1 | WP_090761791.1 SDR family NAD(P)-dependent oxidoreductase |
| 8(2) | WP_272035332.1 | WP_272035332.1 SDR family NAD(P)-dependent oxidoreductase |
| 8(5) | WP_095287656.1 | WP_095287656.1 SDR family NAD(P)-dependent oxidoreductase |
| 8(1) | WP_068619103.1 | WP_068619103.1 SDR family NAD(P)-dependent oxidoreductase |
| 8(1) | WP_051289608.1 | WP_051289608.1 SDR family NAD(P)-dependent oxidoreductase |
| 8(1) | WP_127536193.1 | WP_127536193.1 SDR family NAD(P)-dependent oxidoreductase |
| 8(1) | WP_046217710.1 | WP_046217710.1 SDR family oxidoreductase |
| 8(1) | WP_209872752.1 | WP_209872752.1 SDR family NAD(P)-dependent oxidoreductase |
| 8(1) | WP_285748184.1 | WP_285748184.1 SDR family NAD(P)-dependent oxidoreductase |
| 8(1) | WP_268752841.1 | WP_268752841.1 SDR family oxidoreductase |
| 8(1) | WP_123067119.1 | WP_123067119.1 SDR family NAD(P)-dependent oxidoreductase |
| 8(1) | WP_116231172.1 | WP_116231172.1 SDR family NAD(P)-dependent oxidoreductase |
| 8(5) | WP_013370790.1 | WP_013370790.1 SDR family NAD(P)-dependent oxidoreductase |
| 8(1) | WP_301249732.1 | WP_301249732.1 SDR family NAD(P)-dependent oxidoreductase |
| 8(1) | WP_251583298.1 | WP_251583298.1 SDR family NAD(P)-dependent oxidoreductase |
| 8(6) | WP_063566319.1 | WP_063566319.1 SDR family NAD(P)-dependent oxidoreductase |
| 8(2) | WP_213579157.1 | WP_213579157.1 SDR family NAD(P)-dependent oxidoreductase |
| 8(1) | WP_192266464.1 | WP_192266464.1 SDR family NAD(P)-dependent oxidoreductase |
| 8(2) | WP_119847555.1 | WP_119847555.1 SDR family NAD(P)-dependent oxidoreductase |
| 8(1) | WP_211745825.1 | WP_211745825.1 SDR family NAD(P)-dependent oxidoreductase |
| 8(1) | WP_079694644.1 | WP_079694644.1 SDR family NAD(P)-dependent oxidoreductase |
| 8(1) | WP_251375545.1 | WP_251375545.1 SDR family NAD(P)-dependent oxidoreductase |
| 8(1) | WP_250251681.1 | WP_250251681.1 SDR family NAD(P)-dependent oxidoreductase |
| 8(1) | WP_076322110.1 | WP_076322110.1 SDR family NAD(P)-dependent oxidoreductase |
| 8(1) | WP_007430137.1 | WP_007430137.1 SDR family NAD(P)-dependent oxidoreductase |
| 8(1) | WP_220684642.1 | WP_220684642.1 SDR family NAD(P)-dependent oxidoreductase |
| 8(1) | WP_187370474.1 | WP_187370474.1 SDR family oxidoreductase |
| 8(1) | WP_094092626.1 | WP_094092626.1 SDR family NAD(P)-dependent oxidoreductase |
| 8(1) | WP_113050538.1 | WP_113050538.1 SDR family NAD(P)-dependent oxidoreductase |
| 8(1) | WP_014281187.1 | WP_014281187.1 SDR family NAD(P)-dependent oxidoreductase |
| 8(1) | WP_071640586.1 | WP_071640586.1 SDR family NAD(P)-dependent oxidoreductase |
| 8(1) | WP_113052562.1 | WP_113052562.1 SDR family NAD(P)-dependent oxidoreductase |
| 8(1) | WP_220027684.1 | WP_220027684.1 SDR family NAD(P)-dependent oxidoreductase |
| 8(1) | WP_068694722.1 | WP_068694722.1 SDR family NAD(P)-dependent oxidoreductase |
| 8(1) | WP_225999412.1 | WP_225999412.1 SDR family NAD(P)-dependent oxidoreductase |
| 8(1) | WP_215153845.1 | WP_215153845.1 SDR family NAD(P)-dependent oxidoreductase |
| 8(1) | WP_104495356.1 | WP_104495356.1 SDR family NAD(P)-dependent oxidoreductase |
| 8(6) | WP_009590728.1 | WP_009590728.1 SDR family NAD(P)-dependent oxidoreductase |
| 8(1) | WP_111925413.1 | WP_111925413.1 SDR family oxidoreductase |
| 8(1) | WP_124049181.1 | WP_124049181.1 SDR family oxidoreductase |
| 8(1) | WP_013309994.1 | WP_013309994.1 SDR family NAD(P)-dependent oxidoreductase |
| 8(1) | WP_199794792.1 | WP_199794792.1 SDR family NAD(P)-dependent oxidoreductase |
| 8(1) | WP_076208838.1 | WP_076208838.1 SDR family NAD(P)-dependent oxidoreductase |
| 8(2) | WP_036642472.1 | WP_036642472.1 SDR family NAD(P)-dependent oxidoreductase |
| 8(1) | WP_058829515.1 | WP_058829515.1 SDR family NAD(P)-dependent oxidoreductase |
| 8(1) | WP_213649966.1 | WP_213649966.1 SDR family NAD(P)-dependent oxidoreductase |
| 8(2) | WP_095398794.1 | WP_095398794.1 SDR family NAD(P)-dependent oxidoreductase |
| 8(2) | WP_145046461.1 | WP_145046461.1 SDR family NAD(P)-dependent oxidoreductase |

| | | |
|---|---|---|
| 8(1) | WP_145318684.1 | WP_145318684.1 SDR family NAD(P)-dependent oxidoreductase |
| 8(1) | WP_047842539.1 | WP_047842539.1 SDR family NAD(P)-dependent oxidoreductase |
| 8(1) | WP_061805354.1 | WP_061805354.1 SDR family oxidoreductase |
| 8(1) | WP_223067433.1 | WP_223067433.1 SDR family NAD(P)-dependent oxidoreductase |
| 8(4) | WP_061829403.1 | WP_061829403.1 SDR family NAD(P)-dependent oxidoreductase |
| 8(1) | WP_269055434.1 | WP_269055434.1 SDR family NAD(P)-dependent oxidoreductase, partial |
| 8(3) | WP_062833884.1 | WP_062833884.1 SDR family NAD(P)-dependent oxidoreductase |
| 8(1) | WP_062326152.1 | WP_062326152.1 SDR family NAD(P)-dependent oxidoreductase |
| 8(1) | WP_213532661.1 | WP_213532661.1 SDR family NAD(P)-dependent oxidoreductase |
| 8(1) | WP_208920201.1 | WP_208920201.1 SDR family NAD(P)-dependent oxidoreductase |
| 8(1) | WP_307192885.1 | WP_307192885.1 SDR family oxidoreductase |
| 8(1) | WP_018883267.1 | WP_018883267.1 SDR family NAD(P)-dependent oxidoreductase |
| 8(1) | WP_071220208.1 | WP_071220208.1 SDR family NAD(P)-dependent oxidoreductase |
| 8(1) | WP_080869111.1 | WP_080869111.1 SDR family NAD(P)-dependent oxidoreductase |
| 8(1) | WP_098746857.1 | WP_098746857.1 SDR family NAD(P)-dependent oxidoreductase |
| 8(1) | WP_039306366.1 | WP_039306366.1 SDR family NAD(P)-dependent oxidoreductase |
| 8(1) | WP_165079983.1 | WP_165079983.1 SDR family NAD(P)-dependent oxidoreductase |
| 8(1) | WP_189636047.1 | WP_189636047.1 SDR family NAD(P)-dependent oxidoreductase |
| 8(1) | WP_301879550.1 | WP_301879550.1 SDR family NAD(P)-dependent oxidoreductase |
| 8(9) | WP_076167858.1 | WP_076167858.1 SDR family NAD(P)-dependent oxidoreductase |
| 8(1) | WP_213618819.1 | WP_213618819.1 SDR family NAD(P)-dependent oxidoreductase |
| 8(2) | WP_047909609.1 | WP_047909609.1 SDR family NAD(P)-dependent oxidoreductase |
| 8(1) | WP_188593165.1 | WP_188593165.1 SDR family NAD(P)-dependent oxidoreductase |
| 8(2) | WP_024630331.1 | WP_024630331.1 SDR family NAD(P)-dependent oxidoreductase |
| 8(1) | WP_221875364.1 | WP_221875364.1 oxidoreductase |
| 8(1) | WP_183578466.1 | WP_183578466.1 SDR family NAD(P)-dependent oxidoreductase |
| 8(1) | WP_023957735.1 | WP_023957735.1 SDR family NAD(P)-dependent oxidoreductase |
| 8(2) | WP_107734206.1 | WP_107734206.1 SDR family NAD(P)-dependent oxidoreductase |
| 8(1) | WP_263324532.1 | WP_263324532.1 2-dehydro-3-deoxy-D-gluconate 5-dehydrogenase KduD |
| 8(1) | WP_090908282.1 | WP_090908282.1 SDR family NAD(P)-dependent oxidoreductase |
| 8(1) | WP_307349980.1 | WP_307349980.1 3-oxoacyl-ACP reductase FabG |
| 8(1) | WP_111268936.1 | WP_111268936.1 SDR family NAD(P)-dependent oxidoreductase |
| 8(1) | WP_213644231.1 | WP_213644231.1 SDR family NAD(P)-dependent oxidoreductase |
| 8(2) | WP_100537972.1 | WP_100537972.1 SDR family NAD(P)-dependent oxidoreductase |
| 8(1) | WP_094155404.1 | WP_094155404.1 SDR family NAD(P)-dependent oxidoreductase |
| 8(1) | WP_247902248.1 | WP_247902248.1 SDR family NAD(P)-dependent oxidoreductase |
| 8(1) | WP_102710709.1 | WP_102710709.1 SDR family oxidoreductase |

| | | |
|---|---|---|
| 9(2) | WP_239209286.1 | WP_239209286.1 ABC transporter permease |
| 9(1) | WP_096886728.1 | WP_096886728.1 ABC transporter permease |
| 9(1) | WP_074598765.1 | WP_074598765.1 sugar ABC transporter permease |
| 9(2) | WP_242068642.1 | WP_242068642.1 ABC transporter permease |
| 9(1) | WP_090674355.1 | WP_090674355.1 carbohydrate ABC transporter permease |
| 9(1) | WP_262351733.1 | WP_262351733.1 ABC transporter permease |
| 9(2) | WP_238333008.1 | WP_238333008.1 ABC transporter substrate-binding protein |
| 9(1) | WP_077619663.1 | WP_077619663.1 carbohydrate ABC transporter permease |
| 9(1) | WP_166244891.1 | WP_166244891.1 proline/glycine betaine ABC transporter permease |
| 9(1) | WP_258417122.1 | WP_258417122.1 ABC transporter permease |
| 9(1) | WP_121474469.1 | WP_121474469.1 ABC transporter substrate-binding protein |
| 9(2) | WP_163245188.1 | WP_163245188.1 ABC transporter substrate-binding protein |
| 9(1) | WP_096886740.1 | WP_096886740.1 ABC transporter substrate-binding protein |
| 9(1) | WP_249724673.1 | WP_249724673.1 hypothetical protein |
| 9(1) | WP_258398819.1 | WP_258398819.1 ABC transporter permease |
| 9(1) | WP_258417131.1 | WP_258417131.1 ABC transporter substrate-binding protein |
| 9(1) | WP_089965930.1 | WP_089965930.1 carbohydrate ABC transporter permease |
| 9(1) | WP_083208863.1 | WP_083208863.1 ABC transporter substrate-binding protein |
| 9(1) | WP_014368233.1 | WP_014368233.1 carbohydrate ABC transporter permease |
| 9(1) | WP_277548458.1 | WP_277548458.1 ABC transporter permease |
| 9(1) | WP_076242354.1 | WP_076242354.1 ABC transporter permease subunit |
| 9(1) | WP_223068058.1 | WP_223068058.1 ABC transporter permease |
| 9(2) | WP_113755272.1 | WP_113755272.1 ABC transporter permease |
| 9(2) | WP_119732872.1 | WP_119732872.1 ABC transporter substrate-binding protein |
| 9(2) | WP_036689464.1 | WP_036689464.1 ABC transporter permease subunit |
| 9(1) | WP_204413409.1 | WP_204413409.1 sugar ABC transporter permease |
| 9(1) | WP_244723721.1 | WP_244723721.1 ABC transporter permease subunit |
| 9(1) | WP_014649144.1 | WP_014649144.1 sugar ABC transporter permease |
| 9(1) | WP_212930980.1 | WP_212930980.1 ABC transporter substrate-binding protein |
| 9(1) | WP_258073318.1 | WP_258073318.1 ABC transporter permease |
| 9(1) | WP_121474481.1 | WP_121474481.1 ABC transporter substrate-binding protein |
| 9(1) | WP_309098595.1 | WP_309098595.1 carbohydrate ABC transporter permease |
| 9(1) | WP_205053320.1 | WP_205053320.1 carbohydrate ABC transporter permease |
| 9(1) | WP_244723756.1 | WP_244723756.1 ABC transporter permease subunit |
| 9(1) | WP_014649145.1 | WP_014649145.1 carbohydrate ABC transporter permease |
| 9(1) | WP_251598137.1 | WP_251598137.1 ABC transporter permease subunit |
| 9(1) | WP_094092613.1 | WP_094092613.1 sugar ABC transporter permease |
| 9(1) | WP_251583285.1 | WP_251583285.1 carbohydrate ABC transporter permease |
| 9(1) | WP_181023552.1 | WP_181023552.1 ABC transporter permease |
| 9(1) | WP_231097267.1 | WP_231097267.1 ABC transporter permease |
| 9(1) | WP_209365685.1 | WP_209365685.1 sugar ABC transporter permease |
| 9(2) | WP_036689458.1 | WP_036689458.1 carbohydrate ABC transporter permease |
| 9(1) | WP_034675332.1 | WP_034675332.1 ABC transporter permease |
| 9(1) | WP_258425831.1 | WP_258425831.1 ABC transporter permease |
| 9(1) | WP_199794087.1 | WP_199794087.1 ABC transporter substrate-binding protein |
| 9(1) | WP_158330734.1 | WP_158330734.1 ABC transporter permease |
| 9(1) | WP_209365684.1 | WP_209365684.1 carbohydrate ABC transporter permease |
| 9(1) | WP_101668490.1 | WP_101668490.1 ABC transporter permease |
| 9(1) | WP_094092614.1 | WP_094092614.1 carbohydrate ABC transporter permease |
| 9(1) | WP_232274505.1 | WP_232274505.1 sulfate ABC transporter permease subunit CysT |
| 9(1) | WP_231094839.1 | WP_231094839.1 ABC transporter substrate-binding protein |
| 9(5) | WP_094699648.1 | WP_094699648.1 ABC transporter permease |
| 9(1) | WP_242072814.1 | WP_242072814.1 ABC transporter permease |
| 9(1) | WP_018670516.1 | WP_018670516.1 ABC transporter substrate-binding protein |
| 9(9) | WP_018670514.1 | WP_018670514.1 ABC transporter permease |
| 9(1) | WP_197262008.1 | WP_197262008.1 ABC transporter substrate-binding protein |
| 9(1) | WP_054972101.1 | WP_054972101.1 ABC transporter permease subunit |
| 9(1) | WP_104030453.1 | WP_104030453.1 ABC transporter permease |
| 9(1) | WP_168178800.1 | WP_168178800.1 DUF2877 domain-containing protein |
| 9(1) | WP_213468105.1 | WP_213468105.1 DUF2877 domain-containing protein |
| 9(1) | WP_223254471.1 | WP_223254471.1 carbohydrate ABC transporter permease |
| 9(1) | WP_110931556.1 | WP_110931556.1 amino acid ABC transporter permease |
| 9(1) | WP_042210992.1 | WP_042210992.1 sugar ABC transporter permease |
| 9(2) | WP_003340519.1 | WP_003340519.1 hypothetical protein |
| 9(1) | WP_091155358.1 | WP_091155358.1 ABC transporter permease subunit |

| | | |
|---|---|---|
| 9(2) | WP_236871201.1 | WP_236871201.1 ABC transporter permease |
| 9(1) | WP_211745841.1 | WP_211745841.1 ABC transporter permease |
| 9(1) | WP_003340559.1 | WP_003340559.1 ABC transporter permease |
| 9(1) | WP_076113446.1 | WP_076113446.1 carbohydrate ABC transporter permease |
| 9(1) | WP_104030437.1 | WP_104030437.1 ABC transporter substrate-binding protein |
| 9(1) | WP_236871202.1 | WP_236871202.1 ABC transporter substrate-binding protein |
| 9(1) | WP_132008130.1 | WP_132008130.1 carbohydrate ABC transporter permease |
| 9(3) | WP_010496619.1 | WP_010496619.1 carbohydrate ABC transporter permease |
| 9(3) | WP_025848629.1 | WP_025848629.1 ABC transporter permease subunit |
| 9(1) | WP_258425830.1 | WP_258425830.1 ABC transporter substrate-binding protein |
| 9(1) | WP_258433010.1 | WP_258433010.1 ABC transporter permease |
| 9(1) | WP_101668479.1 | WP_101668479.1 ABC transporter substrate-binding protein |
| 9(1) | WP_293654348.1 | WP_293654348.1 DUF2877 domain-containing protein |
| 9(1) | WP_090722913.1 | WP_090722913.1 ABC transporter permease subunit |
| 9(1) | WP_258425845.1 | WP_258425845.1 ABC transporter substrate-binding protein |
| 9(1) | WP_077619664.1 | WP_077619664.1 sugar ABC transporter permease |
| 9(1) | WP_099517112.1 | WP_099517112.1 ABC transporter permease subunit |
| 9(1) | WP_254660712.1 | WP_254660712.1 ABC transporter permease |
| 9(1) | WP_281937354.1 | WP_281937354.1 carbohydrate ABC transporter permease |
| 9(2) | WP_163245168.1 | WP_163245168.1 ABC transporter permease |
| 9(1) | WP_013914177.1 | WP_013914177.1 carbohydrate ABC transporter permease |
| 9(1) | WP_183600877.1 | WP_183600877.1 proline/glycine betaine ABC transporter permease |
| 9(1) | WP_281378770.1 | WP_281378770.1 carbohydrate ABC transporter permease |
| 9(2) | WP_010496620.1 | WP_010496620.1 ABC transporter permease subunit |
| 9(1) | WP_302878973.1 | WP_302878973.1 carbohydrate ABC transporter permease |
| 9(1) | WP_179034797.1 | WP_179034797.1 ABC transporter permease |
| 9(2) | WP_121613840.1 | WP_121613840.1 sugar ABC transporter permease |
| 9(1) | WP_179160630.1 | WP_179160630.1 amino acid ABC transporter permease |
| 9(1) | WP_197256889.1 | WP_197256889.1 DUF2877 domain-containing protein |
| 9(1) | WP_307192906.1 | WP_307192906.1 ABC transporter permease |
| 9(2) | WP_025848628.1 | WP_025848628.1 carbohydrate ABC transporter permease |
| 9(1) | WP_301804622.1 | WP_301804622.1 ABC transporter substrate-binding protein |
| 9(1) | WP_244723753.1 | WP_244723753.1 carbohydrate ABC transporter permease |
| 9(1) | WP_186580049.1 | WP_186580049.1 ABC transporter permease |
| 9(1) | WP_094699650.1 | WP_094699650.1 ABC transporter substrate-binding protein |
| 9(1) | WP_121474483.1 | WP_121474483.1 ABC transporter permease |
| 9(1) | WP_052421700.1 | WP_052421700.1 ABC transporter permease subunit |
| 9(4) | WP_096886742.1 | WP_096886742.1 ABC transporter permease |
| 9(1) | WP_240962055.1 | WP_240962055.1 ABC transporter permease |
| 9(1) | WP_128631296.1 | WP_128631296.1 sugar ABC transporter permease |
| 9(1) | WP_035318342.1 | WP_035318342.1 ABC transporter substrate-binding protein |
| 9(1) | WP_258418570.1 | WP_258418570.1 ABC transporter permease |
| 9(1) | WP_229696414.1 | WP_229696414.1 ABC transporter permease subunit |
| 9(1) | WP_116353264.1 | WP_116353264.1 sugar ABC transporter permease |
| 9(1) | WP_165097588.1 | WP_165097588.1 carbohydrate ABC transporter permease |
| 9(1) | WP_101668489.1 | WP_101668489.1 ABC transporter substrate-binding protein |
| 9(1) | WP_229140036.1 | WP_229140036.1 sugar ABC transporter permease |
| 9(1) | WP_042210993.1 | WP_042210993.1 carbohydrate ABC transporter permease |
| 9(1) | WP_253411094.1 | WP_253411094.1 carbohydrate ABC transporter permease |
| 9(1) | WP_307589583.1 | WP_307589583.1 methionine ABC transporter permease |
| 9(2) | WP_197247495.1 | WP_197247495.1 ABC transporter substrate-binding protein |
| 9(2) | WP_088830545.1 | WP_088830545.1 carbohydrate ABC transporter permease |
| 9(1) | WP_247083063.1 | WP_247083063.1 carbohydrate ABC transporter permease |
| 9(1) | WP_101668480.1 | WP_101668480.1 ABC transporter permease |
| 9(3) | WP_113755274.1 | WP_113755274.1 ABC transporter substrate-binding protein |
| 9(1) | WP_034675335.1 | WP_034675335.1 ABC transporter permease |
| 9(1) | WP_090722912.1 | WP_090722912.1 carbohydrate ABC transporter permease |
| 9(2) | WP_013914176.1 | WP_013914176.1 sugar ABC transporter permease |
| 9(3) | WP_197262001.1 | WP_197262001.1 ABC transporter substrate-binding protein |
| 9(1) | WP_251596656.1 | WP_251596656.1 sugar ABC transporter permease |
| 9(1) | WP_301804677.1 | WP_301804677.1 ABC transporter substrate-binding protein |
| 9(1) | WP_158330733.1 | WP_158330733.1 ABC transporter substrate-binding protein |
| 9(1) | WP_166244050.1 | WP_166244050.1 sugar ABC transporter permease |
| 9(3) | WP_127459445.1 | WP_127459445.1 ABC transporter permease subunit |
| 9(1) | WP_121610580.1 | WP_121610580.1 carbohydrate ABC transporter permease |
| 9(1) | WP_212930963.1 | WP_212930963.1 ABC transporter substrate-binding protein |
| 9(1) | WP_031309868.1 | WP_031309868.1 ABC transporter permease |
| 9(1) | WP_235286191.1 | WP_235286191.1 phosphonate ABC transporter, permease protein PhnE |
| 9(1) | WP_179603471.1 | WP_179603471.1 ABC transporter permease |
| 9(2) | WP_063188039.1 | WP_063188039.1 carbohydrate ABC transporter permease |
| 9(1) | WP_003335993.1 | WP_003335993.1 ABC transporter permease |
| 9(1) | WP_104065522.1 | WP_104065522.1 ABC transporter substrate-binding protein |
| 9(1) | WP_244527439.1 | WP_244527439.1 ABC transporter permease subunit |
| 9(1) | WP_181747871.1 | WP_181747871.1 ABC transporter substrate-binding protein |
| 9(1) | WP_199613890.1 | WP_199613890.1 sugar ABC transporter permease |
| 9(1) | WP_003335995.1 | WP_003335995.1 ABC transporter substrate-binding protein |
| 9(1) | WP_308615625.1 | WP_308615625.1 methionine ABC transporter permease |
| 9(1) | WP_239582650.1 | WP_239582650.1 carbohydrate ABC transporter permease |
| 9(1) | WP_158326747.1 | WP_158326747.1 ABC transporter permease |
| 9(1) | WP_140574909.1 | WP_140574909.1 ABC transporter substrate-binding protein |
| 9(2) | WP_119732857.1 | WP_119732857.1 ABC transporter substrate-binding protein |
| 9(1) | WP_099517111.1 | WP_099517111.1 carbohydrate ABC transporter permease |
| 9(1) | WP_076311057.1 | WP_076311057.1 ABC transporter permease subunit |
| 9(1) | WP_116334071.1 | WP_116334071.1 ABC transporter substrate-binding protein |
| 9(1) | WP_076113447.1 | WP_076113447.1 ABC transporter permease subunit |
| 9(1) | WP_181747850.1 | WP_181747850.1 ABC transporter substrate-binding protein |
| 9(1) | WP_090674354.1 | WP_090674354.1 ABC transporter permease subunit |
| 9(1) | WP_242066306.1 | WP_242066306.1 ABC transporter permease |
| 9(1) | WP_074598764.1 | WP_074598764.1 carbohydrate ABC transporter permease |
| 9(1) | WP_081621591.1 | WP_081621591.1 ABC transporter permease subunit |
| 9(1) | WP_205053321.1 | WP_205053321.1 ABC transporter permease subunit |
| 9(1) | WP_068793934.1 | WP_068793934.1 ABC transporter substrate-binding protein |
| 9(1) | WP_307192907.1 | WP_307192907.1 ABC transporter permease |
| 9(1) | WP_239586179.1 | WP_239586179.1 ABC transporter permease |
| 9(1) | WP_104148796.1 | WP_104148796.1 ABC transporter substrate-binding protein |
| 9(1) | WP_251596654.1 | WP_251596654.1 carbohydrate ABC transporter permease |
| 9(1) | WP_163245190.1 | WP_163245190.1 ABC transporter permease |
| 9(1) | WP_199613889.1 | WP_199613889.1 carbohydrate ABC transporter permease |
| 9(1) | WP_003340555.1 | WP_003340555.1 ABC transporter substrate-binding protein |
| 9(1) | WP_068694707.1 | WP_068694707.1 sugar ABC transporter permease |
| 9(2) | WP_119732856.1 | WP_119732856.1 ABC transporter permease |
| 9(1) | WP_179603470.1 | WP_179603470.1 ABC transporter permease |
| 9(1) | WP_258417132.1 | WP_258417132.1 ABC transporter permease |
| 9(2) | WP_204412708.1 | WP_204412708.1 sugar ABC transporter permease |

| | | |
|---|---|---|
| 9(1) | WP_089965963.1 | WP_089965963.1 sugar ABC transporter permease |
| 9(2) | WP_306077381.1 | WP_306077381.1 ABC transporter permease |
| 9(1) | WP_253411096.1 | WP_253411096.1 ABC transporter permease subunit |
| 9(1) | WP_186580048.1 | WP_186580048.1 aliphatic sulfonate ABC transporter substrate-binding protein |
| 9(1) | WP_068793935.1 | WP_068793935.1 ABC transporter permease |
| 9(1) | WP_128631297.1 | WP_128631297.1 carbohydrate ABC transporter permease |
| 9(1) | WP_022586765.1 | WP_022586765.1 ABC transporter substrate-binding protein |
| 9(1) | WP_258433020.1 | WP_258433020.1 ABC transporter substrate-binding protein |
| 9(1) | WP_189031017.1 | WP_189031017.1 carbohydrate ABC transporter permease |
| 9(1) | WP_239209292.1 | WP_239209292.1 ABC transporter substrate-binding protein |
| 9(1) | WP_034675393.1 | WP_034675393.1 amino acid ABC transporter permease |
| 9(1) | WP_223068057.1 | WP_223068057.1 ABC transporter substrate-binding protein |
| 9(1) | WP_258418580.1 | WP_258418580.1 ABC transporter substrate-binding protein |
| 9(1) | WP_091155361.1 | WP_091155361.1 carbohydrate ABC transporter permease |
| 9(1) | WP_163245166.1 | WP_163245166.1 ABC transporter substrate-binding protein |
| 9(1) | WP_277548436.1 | WP_277548436.1 ABC transporter substrate-binding protein |
| 9(1) | WP_111147812.1 | WP_111147812.1 phosphonate ABC transporter, permease protein PhnE |
| 9(1) | WP_007430146.1 | WP_007430146.1 sugar ABC transporter permease |
| 9(1) | WP_232699390.1 | WP_232699390.1 ABC transporter permease |
| 9(5) | WP_116334082.1 | WP_116334082.1 ABC transporter substrate-binding protein |
| 9(1) | WP_229139908.1 | WP_229139908.1 carbohydrate ABC transporter permease |
| 9(1) | WP_136379688.1 | WP_136379688.1 sugar ABC transporter permease |
| 9(1) | WP_212930981.1 | WP_212930981.1 ABC transporter permease |
| 9(2) | WP_018670537.1 | WP_018670537.1 hypothetical protein |
| 9(1) | WP_064019313.1 | WP_064019313.1 ABC transporter permease |
| 9(1) | WP_297183855.1 | WP_297183855.1 methionine ABC transporter permease |
| 9(2) | WP_108534966.1 | WP_108534966.1 ABC transporter permease subunit |
| 9(1) | WP_236645787.1 | WP_236645787.1 ABC transporter permease |
| 9(1) | WP_232699388.1 | WP_232699388.1 ABC transporter substrate-binding protein |
| 9(1) | WP_047909621.1 | WP_047909621.1 ABC transporter permease subunit |
| 9(1) | WP_301804685.1 | WP_301804685.1 ABC transporter permease |
| 9(1) | WP_068694708.1 | WP_068694708.1 carbohydrate ABC transporter permease |
| 9(1) | WP_249724675.1 | WP_249724675.1 sugar ABC transporter permease |
| 9(1) | WP_120462938.1 | WP_120462938.1 amino acid ABC transporter permease |
| 9(1) | WP_307312574.1 | WP_307312574.1 ABC transporter permease |
| 9(2) | WP_204412707.1 | WP_204412707.1 carbohydrate ABC transporter permease |
| 9(1) | WP_047909620.1 | WP_047909620.1 carbohydrate ABC transporter permease |
| 9(1) | WP_064019314.1 | WP_064019314.1 ABC transporter substrate-binding protein |
| 9(1) | WP_306077380.1 | WP_306077380.1 ABC transporter permease |
| 9(1) | WP_054972100.1 | WP_054972100.1 carbohydrate ABC transporter permease |
| 9(1) | WP_244850159.1 | WP_244850159.1 ABC transporter permease |
| 9(1) | WP_022586784.1 | WP_022586784.1 ABC transporter substrate-binding protein |
| 9(1) | WP_258417121.1 | WP_258417121.1 ABC transporter substrate-binding protein |
| 9(1) | WP_136379687.1 | WP_136379687.1 sugar ABC transporter permease |
| 9(2) | WP_291288257.1 | WP_291288257.1 methionine ABC transporter permease |
| 9(1) | WP_076311056.1 | WP_076311056.1 carbohydrate ABC transporter permease |
| 9(1) | WP_251583283.1 | WP_251583283.1 ABC transporter permease subunit |
| 9(1) | WP_176452352.1 | WP_176452352.1 ABC transporter permease |
| 9(1) | WP_104065536.1 | WP_104065536.1 ABC transporter substrate-binding protein |
| 9(1) | WP_242061309.1 | WP_242061309.1 ABC transporter permease subunit |
| 9(1) | WP_166244052.1 | WP_166244052.1 carbohydrate ABC transporter permease |
| 9(1) | WP_022586763.1 | WP_022586763.1 ABC transporter permease |
| 9(1) | WP_247083062.1 | WP_247083062.1 sugar ABC transporter permease |
| 9(1) | WP_301804867.1 | WP_301804867.1 ABC transporter permease |
| 10(1) | WP_220027677.1 | WP_220027677.1 YpdA family putative bacillithiol disulfide reductase |
| 10(1) | WP_233697169.1 | WP_233697169.1 CoA-disulfide reductase |
| 10(1) | WP_123067107.1 | WP_123067107.1 YpdA family putative bacillithiol disulfide reductase |
| 10(2) | WP_036609210.1 | WP_036609210.1 YpdA family putative bacillithiol disulfide reductase |
| 10(1) | WP_258296284.1 | WP_258296284.1 YpdA family putative bacillithiol disulfide reductase |
| 10(1) | WP_095287668.1 | WP_095287668.1 YpdA family putative bacillithiol disulfide reductase |
| 10(1) | WP_260511138.1 | WP_260511138.1 YpdA family putative bacillithiol disulfide reductase |
| 10(1) | WP_058829509.1 | WP_058829509.1 YpdA family putative bacillithiol disulfide reductase |
| 10(1) | WP_064639016.1 | WP_064639016.1 YpdA family putative bacillithiol disulfide reductase |
| 10(1) | WP_307413611.1 | WP_307413611.1 YpdA family putative bacillithiol disulfide reductase |
| 10(2) | WP_109999886.1 | WP_109999886.1 YpdA family putative bacillithiol disulfide reductase |
| 10(1) | WP_205053823.1 | WP_205053823.1 YpdA family putative bacillithiol disulfide reductase |
| 10(1) | WP_074094609.1 | WP_074094609.1 YpdA family putative bacillithiol disulfide reductase |
| 10(1) | WP_145412700.1 | WP_145412700.1 YpdA family putative bacillithiol disulfide reductase |
| 10(1) | WP_079346844.1 | WP_079346844.1 YpdA family putative bacillithiol disulfide reductase |
| 10(1) | WP_307349951.1 | WP_307349951.1 YpdA family putative bacillithiol disulfide reductase |
| 10(1) | WP_068619136.1 | WP_068619136.1 YpdA family putative bacillithiol disulfide reductase |
| 10(1) | WP_094155412.1 | WP_094155412.1 YpdA family putative bacillithiol disulfide reductase |
| 10(1) | WP_188593157.1 | WP_188593157.1 YpdA family putative bacillithiol disulfide reductase |
| 10(3) | WP_036642150.1 | WP_036642150.1 YpdA family putative bacillithiol disulfide reductase |
| 10(1) | WP_165079986.1 | WP_165079986.1 YpdA family putative bacillithiol disulfide reductase |
| 10(2) | WP_029517056.1 | WP_029517056.1 YpdA family putative bacillithiol disulfide reductase |
| 10(1) | WP_137062793.1 | WP_137062793.1 YpdA family putative bacillithiol disulfide reductase |
| 10(2) | WP_152402365.1 | WP_152402365.1 YpdA family putative bacillithiol disulfide reductase |
| 10(1) | WP_305755042.1 | WP_305755042.1 YpdA family putative bacillithiol disulfide reductase |
| 10(1) | WP_099853206.1 | WP_099853206.1 YpdA family putative bacillithiol disulfide reductase |
| 10(1) | WP_152381390.1 | WP_152381390.1 YpdA family putative bacillithiol disulfide reductase |
| 10(1) | WP_127536200.1 | WP_127536200.1 YpdA family putative bacillithiol disulfide reductase |
| 10(1) | WP_127592432.1 | WP_127592432.1 YpdA family putative bacillithiol disulfide reductase |
| 10(1) | WP_019424152.1 | WP_019424152.1 CoA-disulfide reductase |
| 10(2) | WP_076330716.1 | WP_076330716.1 YpdA family putative bacillithiol disulfide reductase |
| 10(1) | WP_014281175.1 | WP_014281175.1 YpdA family putative bacillithiol disulfide reductase |
| 10(2) | WP_071220197.1 | WP_071220197.1 YpdA family putative bacillithiol disulfide reductase |
| 10(1) | WP_160037477.1 | WP_160037477.1 YpdA family putative bacillithiol disulfide reductase |
| 10(1) | WP_090723798.1 | WP_090723798.1 YpdA family putative bacillithiol disulfide reductase |
| 10(1) | WP_134910011.1 | WP_134910011.1 YpdA family putative bacillithiol disulfide reductase |
| 10(1) | WP_076208845.1 | WP_076208845.1 YpdA family putative bacillithiol disulfide reductase |
| 10(1) | WP_063566332.1 | WP_063566332.1 YpdA family putative bacillithiol disulfide reductase |
| 10(1) | WP_223067451.1 | WP_223067451.1 YpdA family putative bacillithiol disulfide reductase |
| 10(1) | WP_254327963.1 | WP_254327963.1 YpdA family putative bacillithiol disulfide reductase |
| 10(1) | WP_024630317.1 | WP_024630317.1 YpdA family putative bacillithiol disulfide reductase |
| 10(1) | WP_301249692.1 | WP_301249692.1 YpdA family putative bacillithiol disulfide reductase |
| 10(1) | WP_145318716.1 | WP_145318716.1 YpdA family putative bacillithiol disulfide reductase |
| 10(1) | WP_244159320.1 | WP_244159320.1 YpdA family putative bacillithiol disulfide reductase |
| 10(1) | WP_063479996.1 | WP_063479996.1 YpdA family putative bacillithiol disulfide reductase |
| 10(2) | WP_025719070.1 | WP_025719070.1 YpdA family putative bacillithiol disulfide reductase |
| 10(1) | WP_113052572.1 | WP_113052572.1 YpdA family putative bacillithiol disulfide reductase |
| 10(1) | WP_079694651.1 | WP_079694651.1 YpdA family putative bacillithiol disulfide reductase |

| | | |
|---|---|---|
| 10(3) | WP_090805465.1 | WP_090805465.1 YpdA family putative bacillithiol disulfide reductase |
| 10(1) | WP_145329895.1 | WP_145329895.1 YpdA family putative bacillithiol disulfide reductase |
| 10(1) | WP_036707093.1 | WP_036707093.1 YpdA family putative bacillithiol disulfide reductase |
| 10(1) | WP_305022933.1 | WP_305022933.1 YpdA family putative bacillithiol disulfide reductase |
| 10(1) | WP_013309979.1 | WP_013309979.1 YpdA family putative bacillithiol disulfide reductase |
| 10(1) | WP_103047829.1 | WP_103047829.1 YpdA family putative bacillithiol disulfide reductase |
| 10(1) | WP_111268948.1 | WP_111268948.1 YpdA family putative bacillithiol disulfide reductase |
| 10(1) | WP_213618804.1 | WP_213618804.1 YpdA family putative bacillithiol disulfide reductase |
| 10(1) | WP_077567027.1 | WP_077567027.1 YpdA family putative bacillithiol disulfide reductase |
| 10(1) | WP_108465340.1 | WP_108465340.1 YpdA family putative bacillithiol disulfide reductase |
| 10(1) | WP_208919008.1 | WP_208919008.1 YpdA family putative bacillithiol disulfide reductase |
| 10(1) | WP_145021753.1 | WP_145021753.1 YpdA family putative bacillithiol disulfide reductase |
| 10(2) | WP_145134825.1 | WP_145134825.1 YpdA family putative bacillithiol disulfide reductase |
| 10(1) | WP_071640574.1 | WP_071640574.1 YpdA family putative bacillithiol disulfide reductase |
| 10(1) | WP_094593867.1 | WP_094593867.1 YpdA family putative bacillithiol disulfide reductase |
| 10(1) | WP_090949159.1 | WP_090949159.1 YpdA family putative bacillithiol disulfide reductase |
| 10(1) | WP_192571315.1 | WP_192571315.1 YpdA family putative bacillithiol disulfide reductase |
| 10(2) | WP_283655687.1 | WP_283655687.1 YpdA family putative bacillithiol disulfide reductase |
| 10(1) | WP_056700769.1 | WP_056700769.1 YpdA family putative bacillithiol disulfide reductase |
| 10(1) | WP_106765964.1 | WP_106765964.1 YpdA family putative bacillithiol disulfide reductase |
| 10(1) | WP_145156800.1 | WP_145156800.1 YpdA family putative bacillithiol disulfide reductase |
| 10(1) | WP_169505047.1 | WP_169505047.1 YpdA family putative bacillithiol disulfide reductase |
| 10(1) | WP_102710731.1 | WP_102710731.1 dihydrolipoyl dehydrogenase |
| 10(2) | WP_076292678.1 | WP_076292678.1 YpdA family putative bacillithiol disulfide reductase |
| 10(1) | WP_209861661.1 | WP_209861661.1 YpdA family putative bacillithiol disulfide reductase |
| 10(1) | WP_136604807.1 | WP_136604807.1 YpdA family putative bacillithiol disulfide reductase |
| 10(1) | WP_076264232.1 | WP_076264232.1 YpdA family putative bacillithiol disulfide reductase |
| 10(1) | WP_025685025.1 | WP_025685025.1 YpdA family putative bacillithiol disulfide reductase |
| 10(8) | WP_025364677.1 | WP_025364677.1 YpdA family putative bacillithiol disulfide reductase |
| 10(1) | WP_028405556.1 | WP_028405556.1 YpdA family putative bacillithiol disulfide reductase |
| 10(1) | WP_254349522.1 | WP_254349522.1 YpdA family putative bacillithiol disulfide reductase |
| 10(4) | WP_076249782.1 | WP_076249782.1 YpdA family putative bacillithiol disulfide reductase |
| 10(2) | WP_013370776.1 | WP_013370776.1 YpdA family putative bacillithiol disulfide reductase |
| 10(1) | WP_256019202.1 | WP_256019202.1 YpdA family putative bacillithiol disulfide reductase |
| 10(1) | WP_072732436.1 | WP_072732436.1 YpdA family putative bacillithiol disulfide reductase |
| 10(1) | WP_145948993.1 | WP_145948993.1 YpdA family putative bacillithiol disulfide reductase |
| 10(1) | WP_047910193.1 | WP_047910193.1 YpdA family putative bacillithiol disulfide reductase |
| 10(1) | WP_127199758.1 | WP_127199758.1 YpdA family putative bacillithiol disulfide reductase |
| 10(1) | WP_241929685.1 | WP_241929685.1 YpdA family putative bacillithiol disulfide reductase |
| 10(1) | WP_272035290.1 | WP_272035290.1 YpdA family putative bacillithiol disulfide reductase |
| 10(2) | WP_153978062.1 | WP_153978062.1 YpdA family putative bacillithiol disulfide reductase |
| 10(1) | WP_307537593.1 | WP_307537593.1 YpdA family putative bacillithiol disulfide reductase |
| 10(1) | WP_094092608.1 | WP_094092608.1 YpdA family putative bacillithiol disulfide reductase |
| 10(1) | WP_293202270.1 | WP_293202270.1 YpdA family putative bacillithiol disulfide reductase |
| 10(1) | WP_028589300.1 | WP_028589300.1 YpdA family putative bacillithiol disulfide reductase |
| 10(1) | WP_236417200.1 | WP_236417200.1 YpdA family putative bacillithiol disulfide reductase |
| 10(1) | WP_100537981.1 | WP_100537981.1 YpdA family putative bacillithiol disulfide reductase |
| 10(1) | WP_068694700.1 | WP_068694700.1 YpdA family putative bacillithiol disulfide reductase |
| 10(1) | WP_289373810.1 | WP_289373810.1 YpdA family putative bacillithiol disulfide reductase |
| 10(1) | WP_280753910.1 | WP_280753910.1 YpdA family putative bacillithiol disulfide reductase |
| 10(1) | WP_264929215.1 | WP_264929215.1 YpdA family putative bacillithiol disulfide reductase |
| 10(1) | WP_110822564.1 | WP_110822564.1 YpdA family putative bacillithiol disulfide reductase |

| | | |
|---|---|---|
| 10(2) | WP_063208789.1 | WP_063208789.1 YpdA family putative bacillithiol disulfide reductase |
| 10(1) | WP_133384356.1 | WP_133384356.1 YpdA family putative bacillithiol disulfide reductase |
| 10(1) | WP_113058711.1 | WP_113058711.1 YpdA family putative bacillithiol disulfide reductase |
| 10(1) | WP_244719571.1 | WP_244719571.1 YpdA family putative bacillithiol disulfide reductase |
| 10(1) | WP_191798513.1 | WP_191798513.1 YpdA family putative bacillithiol disulfide reductase |
| 10(1) | WP_104495334.1 | WP_104495334.1 YpdA family putative bacillithiol disulfide reductase |
| 10(1) | WP_145046441.1 | WP_145046441.1 YpdA family putative bacillithiol disulfide reductase |
| 10(3) | WP_087433600.1 | WP_087433600.1 CoA-disulfide reductase |
| 10(2) | WP_127543706.1 | WP_127543706.1 YpdA family putative bacillithiol disulfide reductase |
| 10(1) | WP_009590683.1 | WP_009590683.1 YpdA family putative bacillithiol disulfide reductase |
| 10(1) | WP_094936482.1 | WP_094936482.1 YpdA family putative bacillithiol disulfide reductase |
| 10(1) | WP_307214301.1 | WP_307214301.1 YpdA family putative bacillithiol disulfide reductase |
| 10(1) | WP_091159074.1 | WP_091159074.1 YpdA family putative bacillithiol disulfide reductase |
| 10(1) | WP_053325276.1 | WP_053325276.1 YpdA family putative bacillithiol disulfide reductase |
| 10(1) | WP_192510496.1 | WP_192510496.1 YpdA family putative bacillithiol disulfide reductase |
| 10(1) | WP_134902676.1 | WP_134902676.1 YpdA family putative bacillithiol disulfide reductase |
| 10(1) | WP_010346294.1 | WP_010346294.1 YpdA family putative bacillithiol disulfide reductase |
| 10(2) | WP_025679797.1 | WP_025679797.1 YpdA family putative bacillithiol disulfide reductase |
| 10(1) | WP_090993607.1 | WP_090993607.1 YpdA family putative bacillithiol disulfide reductase |
| 10(1) | WP_054402908.1 | WP_054402908.1 YpdA family putative bacillithiol disulfide reductase |
| 10(1) | WP_116231183.1 | WP_116231183.1 YpdA family putative bacillithiol disulfide reductase |
| 10(5) | WP_076167886.1 | WP_076167886.1 YpdA family putative bacillithiol disulfide reductase |
| 10(2) | WP_138226029.1 | WP_138226029.1 YpdA family putative bacillithiol disulfide reductase |
| 10(1) | WP_023957701.1 | WP_023957701.1 YpdA family putative bacillithiol disulfide reductase |
| 10(3) | WP_007128639.1 | WP_007128639.1 YpdA family putative bacillithiol disulfide reductase |
| 10(1) | WP_274485607.1 | WP_274485607.1 YpdA family putative bacillithiol disulfide reductase |
| 10(1) | WP_018883283.1 | WP_018883283.1 YpdA family putative bacillithiol disulfide reductase |
| 10(3) | WP_061829409.1 | WP_061829409.1 YpdA family putative bacillithiol disulfide reductase |
| 10(1) | WP_183578424.1 | WP_183578424.1 YpdA family putative bacillithiol disulfide reductase |
| 10(3) | WP_095266188.1 | WP_095266188.1 YpdA family putative bacillithiol disulfide reductase |
| 10(3) | WP_096774459.1 | WP_096774459.1 YpdA family putative bacillithiol disulfide reductase |
| 10(1) | WP_251375555.1 | WP_251375555.1 YpdA family putative bacillithiol disulfide reductase |
| 10(1) | WP_054958400.1 | WP_054958400.1 YpdA family putative bacillithiol disulfide reductase |
| 10(5) | WP_015736250.1 | WP_015736250.1 YpdA family putative bacillithiol disulfide reductase |
| 10(3) | WP_017689082.1 | WP_017689082.1 YpdA family putative bacillithiol disulfide reductase |
| 10(1) | WP_053783879.1 | WP_053783879.1 YpdA family putative bacillithiol disulfide reductase |
| 10(1) | WP_194543502.1 | WP_194543502.1 YpdA family putative bacillithiol disulfide reductase |
| 10(1) | WP_229522863.1 | WP_229522863.1 YpdA family putative bacillithiol disulfide reductase |
| 10(1) | WP_040101529.1 | WP_040101529.1 YpdA family putative bacillithiol disulfide reductase |
| 10(1) | WP_269744427.1 | WP_269744427.1 YpdA family putative bacillithiol disulfide reductase |
| 10(1) | WP_110892988.1 | WP_110892988.1 YpdA family putative bacillithiol disulfide reductase |
| 10(1) | WP_152826356.1 | WP_152826356.1 YpdA family putative bacillithiol disulfide reductase |
| 10(1) | WP_058710501.1 | WP_058710501.1 YpdA family putative bacillithiol disulfide reductase |
| 10(1) | WP_150367646.1 | WP_150367646.1 YpdA family putative bacillithiol disulfide reductase |
| 10(1) | WP_235308154.1 | WP_235308154.1 YpdA family putative bacillithiol disulfide reductase |
| 10(1) | WP_213649977.1 | WP_213649977.1 YpdA family putative bacillithiol disulfide reductase |
| 10(1) | WP_036670429.1 | WP_036670429.1 YpdA family putative bacillithiol disulfide reductase |
| 10(1) | WP_091017204.1 | WP_091017204.1 YpdA family putative bacillithiol disulfide reductase |
| 10(1) | WP_305166268.1 | WP_305166268.1 YpdA family putative bacillithiol disulfide reductase |
| 10(1) | WP_225999558.1 | WP_225999558.1 YpdA family putative bacillithiol disulfide reductase |
| 10(1) | WP_194230410.1 | WP_194230410.1 YpdA family putative bacillithiol disulfide reductase |
| 10(1) | WP_212980190.1 | WP_212980190.1 YpdA family putative bacillithiol disulfide reductase |

| | | |
|---|---|---|
| 10(1) | WP_213532643.1 | WP_213532643.1 YpdA family putative bacillithiol disulfide reductase |
| 10(1) | WP_126665668.1 | WP_126665668.1 YpdA family putative bacillithiol disulfide reductase |
| 10(1) | WP_062833874.1 | WP_062833874.1 YpdA family putative bacillithiol disulfide reductase |
| 10(1) | WP_095358553.1 | WP_095358553.1 YpdA family putative bacillithiol disulfide reductase |
| 10(1) | WP_215153829.1 | WP_215153829.1 YpdA family putative bacillithiol disulfide reductase |
| 10(1) | WP_129549127.1 | WP_129549127.1 YpdA family putative bacillithiol disulfide reductase |
| 10(1) | WP_285748156.1 | WP_285748156.1 YpdA family putative bacillithiol disulfide reductase |
| 10(1) | WP_090908273.1 | WP_090908273.1 YpdA family putative bacillithiol disulfide reductase |
| 10(2) | WP_090896170.1 | WP_090896170.1 YpdA family putative bacillithiol disulfide reductase |
| 10(1) | WP_098746879.1 | WP_098746879.1 YpdA family putative bacillithiol disulfide reductase |
| 10(1) | WP_076312506.1 | WP_076312506.1 YpdA family putative bacillithiol disulfide reductase |
| 10(1) | WP_216975170.1 | WP_216975170.1 YpdA family putative bacillithiol disulfide reductase |
| 10(1) | WP_220684636.1 | WP_220684636.1 YpdA family putative bacillithiol disulfide reductase |
| 10(6) | WP_016821023.1 | WP_016821023.1 YpdA family putative bacillithiol disulfide reductase |
| 10(1) | WP_247902237.1 | WP_247902237.1 YpdA family putative bacillithiol disulfide reductase |
| 10(1) | WP_160496940.1 | WP_160496940.1 YpdA family putative bacillithiol disulfide reductase |
| 10(10) | WP_023988257.1 | WP_023988257.1 YpdA family putative bacillithiol disulfide reductase |
| 10(1) | WP_257126954.1 | WP_257126954.1 YpdA family putative bacillithiol disulfide reductase |
| 10(1) | WP_076322119.1 | WP_076322119.1 YpdA family putative bacillithiol disulfide reductase |
| 10(1) | WP_285079446.1 | WP_285079446.1 YpdA family putative bacillithiol disulfide reductase |
| 10(1) | WP_062326140.1 | WP_062326140.1 YpdA family putative bacillithiol disulfide reductase |
| 10(1) | WP_194233599.1 | WP_194233599.1 NAD(P)/FAD-dependent oxidoreductase |
| 10(2) | WP_128101479.1 | WP_128101479.1 YpdA family putative bacillithiol disulfide reductase |
| 10(1) | WP_216531927.1 | WP_216531927.1 YpdA family putative bacillithiol disulfide reductase |
| 10(1) | WP_192266439.1 | WP_192266439.1 YpdA family putative bacillithiol disulfide reductase |
| 10(1) | WP_213644222.1 | WP_213644222.1 YpdA family putative bacillithiol disulfide reductase |
| 10(1) | WP_251583654.1 | WP_251583654.1 YpdA family putative bacillithiol disulfide reductase |
| 10(1) | WP_025716868.1 | WP_025716868.1 YpdA family putative bacillithiol disulfide reductase |
| 10(1) | WP_165147485.1 | WP_165147485.1 YpdA family putative bacillithiol disulfide reductase |
| 10(1) | WP_212969039.1 | WP_212969039.1 YpdA family putative bacillithiol disulfide reductase |

| | | |
|---|---|---|
| 11(1) | WP_301249690.1 | WP_301249690.1 DUF441 domain-containing protein |
| 11(1) | WP_289373815.1 | WP_289373815.1 DUF441 domain-containing protein |
| 11(4) | WP_128101478.1 | WP_128101478.1 DUF441 domain-containing protein |
| 11(1) | WP_068496261.1 | WP_068496261.1 DUF441 domain-containing protein |
| 11(1) | WP_058710502.1 | WP_058710502.1 DUF441 domain-containing protein |
| 11(6) | WP_006211214.1 | WP_006211214.1 DUF441 domain-containing protein |
| 11(1) | WP_285749224.1 | WP_285749224.1 DUF441 domain-containing protein |
| 11(1) | WP_269055428.1 | WP_269055428.1 DUF441 domain-containing protein |
| 11(1) | WP_072732435.1 | WP_072732435.1 DUF441 domain-containing protein |
| 11(2) | WP_138226030.1 | WP_138226030.1 DUF441 domain-containing protein |
| 11(12) | WP_024630316.1 | WP_024630316.1 DUF441 domain-containing protein |
| 11(1) | WP_095358692.1 | WP_095358692.1 DUF441 domain-containing protein |
| 11(1) | WP_076289566.1 | WP_076289566.1 DUF441 domain-containing protein |
| 11(1) | WP_036707089.1 | WP_036707089.1 DUF441 domain-containing protein |
| 11(1) | WP_145156799.1 | WP_145156799.1 DUF441 domain-containing protein |
| 11(1) | WP_090722920.1 | WP_090722920.1 DUF441 domain-containing protein |
| 11(1) | WP_108465339.1 | WP_108465339.1 DUF441 domain-containing protein |
| 11(1) | WP_137062792.1 | WP_137062792.1 DUF441 domain-containing protein |
| 11(1) | WP_106765965.1 | WP_106765965.1 DUF441 domain-containing protein |
| 11(1) | WP_169505046.1 | WP_169505046.1 DUF441 domain-containing protein |
| 11(1) | WP_160496941.1 | WP_160496941.1 DUF441 domain-containing protein |
| 11(2) | WP_025716867.1 | WP_025716867.1 DUF441 domain-containing protein |
| 11(1) | WP_213649978.1 | WP_213649978.1 DUF441 domain-containing protein |
| 11(1) | WP_077567028.1 | WP_077567028.1 DUF441 domain-containing protein |
| 11(2) | WP_017689083.1 | WP_017689083.1 DUF441 domain-containing protein |
| 11(1) | WP_025685024.1 | WP_025685024.1 DUF441 domain-containing protein |
| 11(1) | WP_109999885.1 | WP_109999885.1 DUF441 domain-containing protein |
| 11(1) | WP_305755043.1 | WP_305755043.1 DUF441 domain-containing protein |
| 11(1) | WP_076311061.1 | WP_076311061.1 DUF441 domain-containing protein |
| 11(1) | WP_127199866.1 | WP_127199866.1 DUF441 domain-containing protein |
| 11(1) | WP_054402909.1 | WP_054402909.1 DUF441 domain-containing protein |
| 11(1) | WP_090735435.1 | WP_090735435.1 DUF441 domain-containing protein |
| 11(1) | WP_244719568.1 | WP_244719568.1 DUF441 domain-containing protein |
| 11(1) | WP_047909625.1 | WP_047909625.1 DUF441 domain-containing protein |
| 11(6) | WP_029517055.1 | WP_029517055.1 DUF441 domain-containing protein |
| 11(1) | WP_192266437.1 | WP_192266437.1 DUF441 domain-containing protein |
| 11(1) | WP_023957699.1 | WP_023957699.1 DUF441 domain-containing protein |
| 11(1) | WP_223067452.1 | WP_223067452.1 DUF441 domain-containing protein |
| 11(4) | WP_095287669.1 | WP_095287669.1 DUF441 domain-containing protein |
| 11(1) | WP_194230411.1 | WP_194230411.1 DUF441 domain-containing protein |
| 11(3) | WP_071220196.1 | WP_071220196.1 DUF441 domain-containing protein |
| 11(1) | WP_251378707.1 | WP_251378707.1 DUF441 domain-containing protein |
| 11(1) | WP_160037476.1 | WP_160037476.1 DUF441 domain-containing protein |
| 11(1) | WP_152402363.1 | WP_152402363.1 DUF441 domain-containing protein |
| 11(2) | WP_076322120.1 | WP_076322120.1 DUF441 domain-containing protein |
| 11(1) | WP_236417199.1 | WP_236417199.1 DUF441 domain-containing protein |
| 11(1) | WP_058829508.1 | WP_058829508.1 DUF441 domain-containing protein |
| 11(1) | WP_307349949.1 | WP_307349949.1 DUF441 family protein |
| 11(1) | WP_136604806.1 | WP_136604806.1 DUF441 domain-containing protein |
| 11(1) | WP_142612998.1 | WP_142612998.1 DUF441 domain-containing protein |
| 11(2) | WP_090896169.1 | WP_090896169.1 DUF441 domain-containing protein |
| 11(1) | WP_245339174.1 | WP_245339174.1 DUF441 domain-containing protein |
| 11(2) | WP_095266187.1 | WP_095266187.1 DUF441 domain-containing protein |
| 11(1) | WP_090993605.1 | WP_090993605.1 DUF441 domain-containing protein |
| 11(1) | WP_056700767.1 | WP_056700767.1 DUF441 domain-containing protein |
| 11(1) | WP_145329893.1 | WP_145329893.1 DUF441 domain-containing protein |
| 11(4) | WP_090921075.1 | WP_090921075.1 DUF441 domain-containing protein |
| 11(1) | WP_094593866.1 | WP_094593866.1 DUF441 domain-containing protein |
| 11(1) | WP_254348136.1 | WP_254348136.1 DUF441 domain-containing protein |
| 11(4) | WP_134910013.1 | WP_134910013.1 DUF441 domain-containing protein |
| 11(1) | WP_110822712.1 | WP_110822712.1 DUF441 domain-containing protein |
| 11(2) | WP_068622365.1 | WP_068622365.1 DUF441 domain-containing protein |
| 11(1) | WP_212980191.1 | WP_212980191.1 DUF441 domain-containing protein |
| 11(1) | WP_165175580.1 | WP_165175580.1 DUF441 domain-containing protein |
| 11(1) | WP_209941807.1 | WP_209941807.1 DUF441 domain-containing protein |
| 11(23) | WP_013370775.1 | WP_013370775.1 DUF441 domain-containing protein |
| 11(1) | WP_225999400.1 | WP_225999400.1 DUF441 domain-containing protein |
| 11(1) | WP_247903124.1 | WP_247903124.1 DUF441 domain-containing protein |
| 11(1) | WP_028405557.1 | WP_028405557.1 DUF441 domain-containing protein |
| 11(1) | WP_205053326.1 | WP_205053326.1 DUF441 domain-containing protein |
| 11(4) | WP_062835986.1 | WP_062835986.1 DUF441 domain-containing protein |
| 11(2) | WP_018883284.1 | WP_018883284.1 DUF441 domain-containing protein |
| 11(4) | WP_009590654.1 | WP_009590654.1 DUF441 domain-containing protein |

| | | |
|---|---|---|
| 11(1) | WP_229522864.1 | WP_229522864.1 DUF441 domain-containing protein |
| 11(1) | WP_305022934.1 | WP_305022934.1 DUF441 domain-containing protein |
| 11(1) | WP_183578421.1 | WP_183578421.1 DUF441 domain-containing protein |
| 11(1) | WP_014281174.1 | WP_014281174.1 DUF441 domain-containing protein |
| 11(1) | WP_047842532.1 | WP_047842532.1 DUF441 domain-containing protein |
| 11(1) | WP_213644221.1 | WP_213644221.1 DUF441 domain-containing protein |
| 11(1) | WP_095398789.1 | WP_095398789.1 DUF441 domain-containing protein |
| 11(6) | WP_039273550.1 | WP_039273550.1 DUF441 domain-containing protein |
| 11(3) | WP_007430122.1 | WP_007430122.1 DUF441 domain-containing protein |
| 11(1) | WP_054958399.1 | WP_054958399.1 DUF441 domain-containing protein |
| 11(1) | WP_110892990.1 | WP_110892990.1 DUF441 domain-containing protein |
| 11(11) | WP_076167888.1 | WP_076167888.1 DUF441 domain-containing protein |
| 11(1) | WP_215153827.1 | WP_215153827.1 DUF441 domain-containing protein |
| 11(17) | WP_036609212.1 | WP_036609212.1 DUF441 domain-containing protein |
| 11(2) | WP_208919007.1 | WP_208919007.1 DUF441 domain-containing protein |
| 11(1) | WP_220684635.1 | WP_220684635.1 DUF441 domain-containing protein |
| 11(1) | WP_062326139.1 | WP_062326139.1 DUF441 domain-containing protein |
| 11(10) | WP_015736251.1 | WP_015736251.1 DUF441 domain-containing protein |
| 11(18) | WP_013309978.1 | WP_013309978.1 DUF441 domain-containing protein |
| 11(1) | WP_064639017.1 | WP_064639017.1 DUF441 domain-containing protein |
| 11(2) | WP_111268949.1 | WP_111268949.1 DUF441 domain-containing protein |
| 11(1) | WP_212969040.1 | WP_212969040.1 DUF441 domain-containing protein |
| 11(3) | WP_007128640.1 | WP_007128640.1 DUF441 domain-containing protein |
| 11(2) | WP_063208791.1 | WP_063208791.1 DUF441 domain-containing protein |
| 11(1) | WP_216531926.1 | WP_216531926.1 DUF441 domain-containing protein |
| 11(2) | WP_272035288.1 | WP_272035288.1 DUF441 domain-containing protein |
| 12(1) | WP_090921073.1 | WP_090921073.1 metalloregulator ArsR/SmtB family transcription factor |
| 12(1) | WP_054958401.1 | WP_054958401.1 metalloregulator ArsR/SmtB family transcription factor |
| 12(1) | WP_145408782.1 | WP_145408782.1 metalloregulator ArsR/SmtB family transcription factor |
| 12(2) | WP_071220198.1 | WP_071220198.1 DUF2087 domain-containing protein |
| 12(1) | WP_010346293.1 | WP_010346293.1 metalloregulator ArsR/SmtB family transcription factor |
| 12(1) | WP_260867008.1 | WP_260867008.1 DUF2087 domain-containing protein |
| 12(1) | WP_123067108.1 | WP_123067108.1 metalloregulator ArsR/SmtB family transcription factor |
| 12(1) | WP_293202293.1 | WP_293202293.1 metalloregulator ArsR/SmtB family transcription factor |
| 12(1) | WP_269055435.1 | WP_269055435.1 metalloregulator ArsR/SmtB family transcription factor |
| 12(1) | WP_260065252.1 | WP_260065252.1 DUF2087 domain-containing protein |
| 12(1) | WP_068496257.1 | WP_068496257.1 metalloregulator ArsR/SmtB family transcription factor |
| 12(1) | WP_225999401.1 | WP_225999401.1 metalloregulator ArsR/SmtB family transcription factor |
| 12(1) | WP_304142095.1 | WP_304142095.1 DUF2087 domain-containing protein |
| 12(1) | WP_119847547.1 | WP_119847547.1 DUF2087 domain-containing protein |
| 12(1) | WP_036609208.1 | WP_036609208.1 metalloregulator ArsR/SmtB family transcription factor |
| 12(1) | WP_013309980.1 | WP_013309980.1 metalloregulator ArsR/SmtB family transcription factor |
| 12(1) | WP_223067450.1 | WP_223067450.1 metalloregulator ArsR/SmtB family transcription factor |
| 12(1) | WP_236417201.1 | WP_236417201.1 metalloregulator ArsR/SmtB family transcription factor |
| 12(1) | WP_250244803.1 | WP_250244803.1 metalloregulator ArsR/SmtB family transcription factor |
| 12(1) | WP_244879216.1 | WP_244879216.1 DUF2087 domain-containing protein |
| 12(1) | WP_062326141.1 | WP_062326141.1 metalloregulator ArsR/SmtB family transcription factor |
| 12(3) | WP_015736249.1 | WP_015736249.1 DUF2087 domain-containing protein |
| 12(1) | WP_103047828.1 | WP_103047828.1 metalloregulator ArsR/SmtB family transcription factor |
| 12(2) | WP_071640575.1 | WP_071640575.1 metalloregulator ArsR/SmtB family transcription factor |
| 12(1) | WP_305022931.1 | WP_305022931.1 metalloregulator ArsR/SmtB family transcription factor |
| 12(1) | WP_152826358.1 | WP_152826358.1 DUF2087 domain-containing protein |
| 12(2) | WP_025719068.1 | WP_025719068.1 metalloregulator ArsR/SmtB family transcription factor |
| 12(1) | WP_152381391.1 | WP_152381391.1 metalloregulator ArsR/SmtB family transcription factor |
| 12(1) | WP_079346842.1 | WP_079346842.1 metalloregulator ArsR/SmtB family transcription factor |
| 12(1) | WP_272036452.1 | WP_272036452.1 metalloregulator ArsR/SmtB family transcription factor |
| 12(1) | WP_205053325.1 | WP_205053325.1 metalloregulator ArsR/SmtB family transcription factor |
| 12(1) | WP_029514391.1 | WP_029514391.1 metalloregulator ArsR/SmtB family transcription factor |
| 12(2) | WP_104495336.1 | WP_104495336.1 metalloregulator ArsR/SmtB family transcription factor |
| 12(2) | WP_091155345.1 | WP_091155345.1 metalloregulator ArsR/SmtB family transcription factor |
| 12(1) | WP_307416930.1 | WP_307416930.1 metalloregulator ArsR/SmtB family transcription factor |
| 12(1) | WP_215075559.1 | WP_215075559.1 metalloregulator ArsR/SmtB family transcription factor |
| 12(1) | WP_241929684.1 | WP_241929684.1 metalloregulator ArsR/SmtB family transcription factor |
| 12(1) | WP_025685026.1 | WP_025685026.1 metalloregulator ArsR/SmtB family transcription factor |
| 12(1) | WP_260869266.1 | WP_260869266.1 DUF2087 domain-containing protein |
| 12(1) | WP_094092609.1 | WP_094092609.1 metalloregulator ArsR/SmtB family transcription factor |
| 12(1) | WP_264930681.1 | WP_264930681.1 metalloregulator ArsR/SmtB family transcription factor |
| 12(2) | WP_244867301.1 | WP_244867301.1 DUF2087 domain-containing protein |
| 12(1) | WP_127199759.1 | WP_127199759.1 metalloregulator ArsR/SmtB family transcription factor |
| 12(1) | WP_209861659.1 | WP_209861659.1 metalloregulator ArsR/SmtB family transcription factor |
| 12(1) | WP_184180389.1 | WP_184180389.1 metalloregulator ArsR/SmtB family transcription factor |
| 12(1) | WP_231888192.1 | WP_231888192.1 DUF2087 domain-containing protein |
| 12(1) | WP_307214587.1 | WP_307214587.1 metalloregulator ArsR/SmtB family transcription factor |
| 12(3) | WP_076249781.1 | WP_076249781.1 metalloregulator ArsR/SmtB family transcription factor |
| 12(1) | WP_165079985.1 | WP_165079985.1 metalloregulator ArsR/SmtB family transcription factor |
| 12(1) | WP_072732437.1 | WP_072732437.1 metalloregulator ArsR/SmtB family transcription factor |
| 12(1) | WP_160496939.1 | WP_160496939.1 metalloregulator ArsR/SmtB family transcription factor |
| 12(1) | WP_251035345.1 | WP_251035345.1 DUF2087 domain-containing protein |
| 12(1) | WP_113052571.1 | WP_113052571.1 metalloregulator ArsR/SmtB family transcription factor |
| 12(1) | WP_076289568.1 | WP_076289568.1 metalloregulator ArsR/SmtB family transcription factor |
| 12(1) | WP_110892986.1 | WP_110892986.1 metalloregulator ArsR/SmtB family transcription factor |
| 12(1) | WP_307349953.1 | WP_307349953.1 metalloregulator ArsR/SmtB family transcription factor |
| 12(2) | WP_237566520.1 | WP_237566520.1 DUF2087 domain-containing protein |
| 12(1) | WP_145046542.1 | WP_145046542.1 metalloregulator ArsR/SmtB family transcription factor |
| 12(1) | WP_213649976.1 | WP_213649976.1 metalloregulator ArsR/SmtB family transcription factor |
| 12(1) | WP_094936483.1 | WP_094936483.1 metalloregulator ArsR/SmtB family transcription factor |
| 12(1) | WP_192510495.1 | WP_192510495.1 metalloregulator ArsR/SmtB family transcription factor |
| 12(1) | WP_213532644.1 | WP_213532644.1 metalloregulator ArsR/SmtB family transcription factor |
| 12(1) | WP_256720487.1 | WP_256720487.1 DUF2087 domain-containing protein |
| 12(1) | WP_305755041.1 | WP_305755041.1 metalloregulator ArsR/SmtB family transcription factor |
| 12(1) | WP_232239344.1 | WP_232239344.1 DUF2087 domain-containing protein |
| 12(5) | WP_095358552.1 | WP_095358552.1 metalloregulator ArsR/SmtB family transcription factor |
| 12(1) | WP_250272063.1 | WP_250272063.1 metalloregulator ArsR/SmtB family transcription factor |
| 12(1) | WP_009590655.1 | WP_009590655.1 DUF2087 domain-containing protein |
| 12(1) | WP_285079612.1 | WP_285079612.1 metalloregulator ArsR/SmtB family transcription factor |
| 12(1) | WP_102997915.1 | WP_102997915.1 metalloregulator ArsR/SmtB family transcription factor |
| 12(1) | WP_213618805.1 | WP_213618805.1 metalloregulator ArsR/SmtB family transcription factor |
| 12(1) | WP_053783880.1 | WP_053783880.1 metalloregulator ArsR/SmtB family transcription factor |
| 12(1) | WP_165147482.1 | WP_165147482.1 metalloregulator ArsR/SmtB family transcription factor |
| 12(2) | WP_076311060.1 | WP_076311060.1 metalloregulator ArsR/SmtB family transcription factor |
| 12(1) | WP_116231761.1 | WP_116231761.1 DUF2087 domain-containing protein |
| 12(1) | WP_285748158.1 | WP_285748158.1 metalloregulator ArsR/SmtB family transcription factor |
| 12(1) | WP_260986747.1 | WP_260986747.1 DUF2087 domain-containing protein |

| | | |
|---|---|---|
| 12(1) | WP_062835987.1 | WP_062835987.1 metalloregulator ArsR/SmtB family transcription factor |
| 12(1) | WP_127544002.1 | WP_127544002.1 metalloregulator ArsR/SmtB family transcription factor |
| 12(1) | WP_053325277.1 | WP_053325277.1 metalloregulator ArsR/SmtB family transcription factor |
| 12(2) | WP_090896171.1 | WP_090896171.1 metalloregulator ArsR/SmtB family transcription factor |
| 12(2) | WP_063208787.1 | WP_063208787.1 metalloregulator ArsR/SmtB family transcription factor |
| 12(1) | WP_220684637.1 | WP_220684637.1 DUF2087 domain-containing protein |
| 12(1) | WP_006290886.1 | WP_006290886.1 DUF5937 family protein |
| 12(1) | WP_025716869.1 | WP_025716869.1 metalloregulator ArsR/SmtB family transcription factor |
| 12(1) | WP_044644797.1 | WP_044644797.1 metalloregulator ArsR/SmtB family transcription factor |
| 12(5) | WP_068940675.1 | WP_068940675.1 metalloregulator ArsR/SmtB family transcription factor |
| 12(1) | WP_208919010.1 | WP_208919010.1 metalloregulator ArsR/SmtB family transcription factor |
| 12(3) | WP_095266272.1 | WP_095266272.1 metalloregulator ArsR/SmtB family transcription factor |
| 12(1) | WP_192266441.1 | WP_192266441.1 metalloregulator ArsR/SmtB family transcription factor |
| 12(1) | WP_206703918.1 | WP_206703918.1 metalloregulator ArsR/SmtB family transcription factor |
| 12(1) | WP_254325798.1 | WP_254325798.1 metalloregulator ArsR/SmtB family transcription factor |
| 12(1) | WP_145321353.1 | WP_145321353.1 metalloregulator ArsR/SmtB family transcription factor |
| 12(1) | WP_018883282.1 | WP_018883282.1 metalloregulator ArsR/SmtB family transcription factor |
| 12(1) | WP_094155411.1 | WP_094155411.1 metalloregulator ArsR/SmtB family transcription factor |
| 12(1) | WP_026081135.1 | WP_026081135.1 metalloregulator ArsR/SmtB family transcription factor |
| 12(1) | WP_039273548.1 | WP_039273548.1 metalloregulator ArsR/SmtB family transcription factor |
| 12(1) | WP_029517057.1 | WP_029517057.1 metalloregulator ArsR/SmtB family transcription factor |
| 12(1) | WP_106765963.1 | WP_106765963.1 metalloregulator ArsR/SmtB family transcription factor |
| 12(1) | WP_234489612.1 | WP_234489612.1 DUF2087 domain-containing protein |
| 12(1) | WP_256992146.1 | WP_256992146.1 DUF2087 domain-containing protein |
| 12(1) | WP_111268947.1 | WP_111268947.1 metalloregulator ArsR/SmtB family transcription factor |
| 12(19) | WP_013370777.1 | WP_013370777.1 metalloregulator ArsR/SmtB family transcription factor |
| 12(1) | WP_090908274.1 | WP_090908274.1 DUF2087 domain-containing protein |
| 12(1) | WP_145329897.1 | WP_145329897.1 metalloregulator ArsR/SmtB family transcription factor |
| 12(1) | WP_305166267.1 | WP_305166267.1 metalloregulator ArsR/SmtB family transcription factor |
| 12(1) | WP_244364315.1 | WP_244364315.1 DUF2087 domain-containing protein |
| 12(1) | WP_216531928.1 | WP_216531928.1 DUF2087 domain-containing protein |
| 12(5) | WP_061829408.1 | WP_061829408.1 metalloregulator ArsR/SmtB family transcription factor |
| 12(1) | WP_076331278.1 | WP_076331278.1 metalloregulator ArsR/SmtB family transcription factor |
| 12(1) | WP_154958921.1 | WP_154958921.1 metalloregulator ArsR/SmtB family transcription factor |
| 12(5) | WP_091034982.1 | WP_091034982.1 metalloregulator ArsR/SmtB family transcription factor |
| 12(1) | WP_282322192.1 | WP_282322192.1 metalloregulator ArsR/SmtB family transcription factor |
| 12(1) | WP_260845579.1 | WP_260845579.1 DUF2087 domain-containing protein |
| 12(3) | WP_025679798.1 | WP_025679798.1 metalloregulator ArsR/SmtB family transcription factor |
| 12(1) | WP_036707095.1 | WP_036707095.1 metalloregulator ArsR/SmtB family transcription factor |
| 12(3) | WP_014281176.1 | WP_014281176.1 metalloregulator ArsR/SmtB family transcription factor |
| 12(1) | WP_036671025.1 | WP_036671025.1 metalloregulator ArsR/SmtB family transcription factor |
| 12(4) | WP_235187430.1 | WP_235187430.1 DUF2087 domain-containing protein |
| 12(1) | WP_244719575.1 | WP_244719575.1 metalloregulator ArsR/SmtB family transcription factor |
| 12(1) | WP_127592543.1 | WP_127592543.1 DUF2087 domain-containing protein |
| 12(2) | WP_024630318.1 | WP_024630318.1 metalloregulator ArsR/SmtB family transcription factor |
| 12(1) | WP_074096853.1 | WP_074096853.1 metalloregulator ArsR/SmtB family transcription factor |
| 12(1) | WP_047909624.1 | WP_047909624.1 metalloregulator ArsR/SmtB family transcription factor |
| 12(1) | WP_113058712.1 | WP_113058712.1 DUF2087 domain-containing protein |
| 12(3) | WP_240913008.1 | WP_240913008.1 DUF2087 domain-containing protein |
| 12(1) | WP_160037478.1 | WP_160037478.1 metalloregulator ArsR/SmtB family transcription factor |
| 12(1) | WP_095287667.1 | WP_095287667.1 metalloregulator ArsR/SmtB family transcription factor |
| 12(1) | WP_128101480.1 | WP_128101480.1 metalloregulator ArsR/SmtB family transcription factor |

| | | |
|---|---|---|
| 12(3) | WP_007128638.1 | WP_007128638.1 DUF2087 domain-containing protein |
| 12(1) | WP_068694701.1 | WP_068694701.1 metalloregulator ArsR/SmtB family transcription factor |
| 12(1) | WP_108465341.1 | WP_108465341.1 metalloregulator ArsR/SmtB family transcription factor |
| 12(1) | WP_165175578.1 | WP_165175578.1 metalloregulator ArsR/SmtB family transcription factor |
| 12(1) | WP_244169194.1 | WP_244169194.1 DUF2087 domain-containing protein |
| 12(1) | WP_006289759.1 | WP_006289759.1 metalloregulator ArsR/SmtB family transcription factor |
| 12(1) | WP_047844253.1 | WP_047844253.1 metalloregulator ArsR/SmtB family transcription factor |
| 12(1) | WP_145412701.1 | WP_145412701.1 metalloregulator ArsR/SmtB family transcription factor |
| 12(1) | WP_188593158.1 | WP_188593158.1 metalloregulator ArsR/SmtB family transcription factor |
| 12(1) | WP_068619134.1 | WP_068619134.1 metalloregulator ArsR/SmtB family transcription factor |
| 12(1) | WP_056700772.1 | WP_056700772.1 metalloregulator ArsR/SmtB family transcription factor |
| 12(1) | WP_246426404.1 | WP_246426404.1 DUF2087 domain-containing protein |
| 12(1) | WP_076209482.1 | WP_076209482.1 metalloregulator ArsR/SmtB family transcription factor |
| 12(4) | WP_096774458.1 | WP_096774458.1 DUF2087 domain-containing protein |
| 12(1) | WP_247903125.1 | WP_247903125.1 metalloregulator ArsR/SmtB family transcription factor |
| 12(1) | WP_083669797.1 | WP_083669797.1 metalloregulator ArsR/SmtB family transcription factor |
| 12(1) | WP_077567026.1 | WP_077567026.1 DUF2087 domain-containing protein |
| 12(9) | WP_063566331.1 | WP_063566331.1 metalloregulator ArsR/SmtB family transcription factor |
| 12(1) | WP_137062794.1 | WP_137062794.1 metalloregulator ArsR/SmtB family transcription factor |
| 12(1) | WP_243390761.1 | WP_243390761.1 DUF2087 domain-containing protein |
| 12(1) | WP_028589299.1 | WP_028589299.1 metalloregulator ArsR/SmtB family transcription factor |
| 12(1) | WP_007430124.1 | WP_007430124.1 metalloregulator ArsR/SmtB family transcription factor |
| 12(1) | WP_100529429.1 | WP_100529429.1 metalloregulator ArsR/SmtB family transcription factor |
| 12(1) | WP_023957703.1 | WP_023957703.1 DUF2087 domain-containing protein |
| 12(7) | WP_023988258.1 | WP_023988258.1 metalloregulator ArsR/SmtB family transcription factor |
| 12(1) | WP_136604808.1 | WP_136604808.1 metalloregulator ArsR/SmtB family transcription factor |
| 12(1) | WP_090761803.1 | WP_090761803.1 metalloregulator ArsR/SmtB family transcription factor |

| | | |
|---|---|---|
| 13(2) | WP_138226028.1 | WP_138226028.1 hypothetical protein |
| 13(1) | WP_160037481.1 | WP_160037481.1 hypothetical protein |
| 13(2) | WP_136604810.1 | WP_136604810.1 hypothetical protein |
| 13(3) | WP_018883276.1 | WP_018883276.1 hypothetical protein |
| 13(1) | WP_010346289.1 | WP_010346289.1 hypothetical protein |
| 13(6) | WP_095287663.1 | WP_095287663.1 hypothetical protein |
| 13(1) | WP_104495345.1 | WP_104495345.1 hypothetical protein |
| 13(4) | WP_105443571.1 | WP_105443571.1 hypothetical protein |
| 13(4) | WP_025716873.1 | WP_025716873.1 hypothetical protein |
| 13(1) | WP_183578440.1 | WP_183578440.1 hypothetical protein |
| 13(2) | WP_091155370.1 | WP_091155370.1 hypothetical protein |
| 13(2) | WP_047909616.1 | WP_047909616.1 hypothetical protein |
| 13(3) | WP_095266193.1 | WP_095266193.1 hypothetical protein |
| 13(1) | WP_305755038.1 | WP_305755038.1 hypothetical protein |
| 13(7) | WP_081473730.1 | WP_081473730.1 hypothetical protein |
| 13(1) | WP_106765962.1 | WP_106765962.1 hypothetical protein |
| 13(10) | WP_024630323.1 | WP_024630323.1 hypothetical protein |
| 13(2) | WP_083837565.1 | WP_083837565.1 hypothetical protein |
| 13(2) | WP_128101484.1 | WP_128101484.1 hypothetical protein |
| 13(1) | WP_237690775.1 | WP_237690775.1 hypothetical protein |
| 13(1) | WP_307349961.1 | WP_307349961.1 hypothetical protein |
| 13(2) | WP_076311052.1 | WP_076311052.1 hypothetical protein |
| 13(2) | WP_068694704.1 | WP_068694704.1 hypothetical protein |
| 13(1) | WP_076322116.1 | WP_076322116.1 hypothetical protein |

| | | | |
|---|---|---|---|
| 13(5) | WP_007128632.1 | WP_007128632.1 | hypothetical protein |
| 13(1) | WP_160496936.1 | WP_160496936.1 | hypothetical protein |
| 13(4) | WP_090921063.1 | WP_090921063.1 | hypothetical protein |
| 13(1) | WP_023957715.1 | WP_023957715.1 | hypothetical protein |
| 13(1) | WP_152402372.1 | WP_152402372.1 | hypothetical protein |
| 13(53) | WP_013370782.1 | WP_013370782.1 | hypothetical protein |
| 13(1) | WP_150367642.1 | WP_150367642.1 | hypothetical protein |
| 13(10) | WP_076167876.1 | WP_076167876.1 | hypothetical protein |
| 13(1) | WP_054958405.1 | WP_054958405.1 | hypothetical protein |
| 13(2) | WP_068619119.1 | WP_068619119.1 | hypothetical protein |
| 13(7) | WP_110892980.1 | WP_110892980.1 | hypothetical protein |
| 13(1) | WP_090993615.1 | WP_090993615.1 | hypothetical protein |
| 13(1) | WP_169505049.1 | WP_169505049.1 | hypothetical protein |
| 13(24) | WP_006211206.1 | WP_006211206.1 | hypothetical protein |
| 13(2) | WP_194230408.1 | WP_194230408.1 | hypothetical protein |
| 13(2) | WP_208919026.1 | WP_208919026.1 | hypothetical protein |
| 13(1) | WP_212949028.1 | WP_212949028.1 | hypothetical protein |
| 13(1) | WP_211745834.1 | WP_211745834.1 | hypothetical protein |
| 13(1) | WP_244719600.1 | WP_244719600.1 | hypothetical protein |
| 13(13) | WP_063566326.1 | WP_063566326.1 | hypothetical protein |
| 13(1) | WP_236417207.1 | WP_236417207.1 | hypothetical protein |
| 13(3) | WP_191798524.1 | WP_191798524.1 | hypothetical protein |
| 13(2) | WP_127199767.1 | WP_127199767.1 | hypothetical protein |
| 13(1) | WP_305022927.1 | WP_305022927.1 | hypothetical protein |
| 13(4) | WP_076330720.1 | WP_076330720.1 | hypothetical protein |
| 13(3) | WP_082560509.1 | WP_082560509.1 | hypothetical protein |
| 13(1) | WP_053492736.1 | WP_053492736.1 | hypothetical protein |
| 13(3) | WP_098746871.1 | WP_098746871.1 | hypothetical protein |
| 13(1) | WP_108465344.1 | WP_108465344.1 | hypothetical protein |
| 13(1) | WP_229522860.1 | WP_229522860.1 | hypothetical protein |
| 13(19) | WP_074094612.1 | WP_074094612.1 | hypothetical protein |

| | | | |
|---|---|---|---|
| 14(1) | WP_088327109.1 | WP_088327109.1 | aspartate--tRNA(Asn) ligase |
| 14(3) | WP_138322085.1 | WP_138322085.1 | aspartate--tRNA(Asn) ligase |
| 14(1) | WP_158318210.1 | WP_158318210.1 | aspartate--tRNA(Asn) ligase |
| 14(2) | WP_064473940.1 | WP_064473940.1 | aspartate--tRNA(Asn) ligase |
| 14(1) | WP_242805837.1 | WP_242805837.1 | aspartate--tRNA(Asn) ligase |
| 14(1) | WP_098776432.1 | WP_098776432.1 | aspartate--tRNA(Asn) ligase |
| 14(1) | WP_098856765.1 | WP_098856765.1 | aspartate--tRNA(Asn) ligase |
| 14(1) | WP_097952786.1 | WP_097952786.1 | aspartate--tRNA(Asn) ligase |
| 14(1) | WP_143881069.1 | WP_143881069.1 | aspartate--tRNA(Asn) ligase |
| 14(1) | WP_098562455.1 | WP_098562455.1 | aspartate--tRNA(Asn) ligase |
| 14(1) | WP_000529810.1 | WP_000529810.1 | aspartate--tRNA(Asn) ligase |
| 14(1) | WP_097852391.1 | WP_097852391.1 | aspartate--tRNA(Asn) ligase |
| 14(1) | WP_098235620.1 | WP_098235620.1 | aspartate--tRNA(Asn) ligase |
| 14(3) | WP_088609114.1 | WP_088609114.1 | aspartate--tRNA(Asn) ligase |
| 14(1) | WP_147786238.1 | WP_147786238.1 | aspartate--tRNA(Asn) ligase |
| 14(2) | WP_098487123.1 | WP_098487123.1 | aspartate--tRNA(Asn) ligase |
| 14(1) | WP_098201421.1 | WP_098201421.1 | aspartate--tRNA(Asn) ligase |
| 14(1) | WP_000529809.1 | WP_000529809.1 | aspartate--tRNA(Asn) ligase |
| 14(1) | WP_098250523.1 | WP_098250523.1 | aspartate--tRNA(Asn) ligase |
| 14(1) | WP_242251772.1 | WP_242251772.1 | aspartate--tRNA(Asn) ligase |
| 14(3) | WP_000529808.1 | WP_000529808.1 | aspartate--tRNA(Asn) ligase |
| 14(1) | WP_088058531.1 | WP_088058531.1 | aspartate--tRNA(Asn) ligase |
| 14(2) | WP_212956329.1 | WP_212956329.1 | aspartate--tRNA(Asn) ligase |
| 14(1) | WP_131231109.1 | WP_131231109.1 | aspartate--tRNA(Asn) ligase |
| 14(1) | WP_101219120.1 | WP_101219120.1 | aspartate--tRNA(Asn) ligase |
| 14(1) | WP_098508781.1 | WP_098508781.1 | aspartate--tRNA(Asn) ligase |
| 14(3) | WP_000529803.1 | WP_000529803.1 | aspartate--tRNA(Asn) ligase |
| 14(1) | WP_078035540.1 | WP_078035540.1 | aspartate--tRNA(Asn) ligase |
| 14(1) | WP_098952619.1 | WP_098952619.1 | aspartate--tRNA(Asn) ligase |
| 14(1) | WP_098925582.1 | WP_098925582.1 | aspartate--tRNA(Asn) ligase |
| 14(1) | WP_017149775.1 | WP_017149775.1 | aspartate--tRNA(Asn) ligase |
| 14(4) | WP_000377348.1 | WP_000377348.1 | aspartate--tRNA(Asn) ligase |
| 14(1) | WP_071736272.1 | WP_071736272.1 | aspartate--tRNA(Asn) ligase |
| 14(1) | WP_098519065.1 | WP_098519065.1 | aspartate--tRNA(Asn) ligase |
| 14(1) | WP_002158048.1 | WP_002158048.1 | aspartate--tRNA(Asn) ligase |
| 14(2) | WP_098588024.1 | WP_098588024.1 | aspartate--tRNA(Asn) ligase |
| 14(1) | WP_042512833.1 | WP_042512833.1 | aspartate--tRNA(Asn) ligase |
| 14(1) | WP_286125242.1 | WP_286125242.1 | aspartate--tRNA(Asn) ligase |
| 14(1) | WP_270728511.1 | WP_270728511.1 | aspartate--tRNA(Asn) ligase |
| 14(1) | WP_098230736.1 | WP_098230736.1 | aspartate--tRNA(Asn) ligase |
| 14(2) | WP_048552301.1 | WP_048552301.1 | aspartate--tRNA(Asn) ligase |
| 14(1) | WP_098259060.1 | WP_098259060.1 | aspartate--tRNA(Asn) ligase |
| 14(1) | WP_088119571.1 | WP_088119571.1 | aspartate--tRNA(Asn) ligase |
| 14(1) | WP_097961541.1 | WP_097961541.1 | aspartate--tRNA(Asn) ligase |
| 14(1) | WP_166704990.1 | WP_166704990.1 | aspartate--tRNA(Asn) ligase |
| 14(2) | WP_088063230.1 | WP_088063230.1 | aspartate--tRNA(Asn) ligase |
| 14(1) | WP_270364635.1 | WP_270364635.1 | aspartate--tRNA(Asn) ligase |
| 14(1) | WP_207994165.1 | WP_207994165.1 | aspartate--tRNA(Asn) ligase |
| 14(1) | WP_087995246.1 | WP_087995246.1 | aspartate--tRNA(Asn) ligase |
| 14(2) | WP_097911843.1 | WP_097911843.1 | aspartate--tRNA(Asn) ligase |
| 14(1) | WP_157418140.1 | WP_157418140.1 | aspartate--tRNA(Asn) ligase |
| 14(1) | WP_098991574.1 | WP_098991574.1 | aspartate--tRNA(Asn) ligase |
| 14(1) | WP_098082495.1 | WP_098082495.1 | aspartate--tRNA(Asn) ligase |
| 14(1) | WP_305163859.1 | WP_305163859.1 | aspartate--tRNA(Asn) ligase |
| 14(1) | WP_144506827.1 | WP_144506827.1 | aspartate--tRNA(Asn) ligase |
| 14(8) | WP_000443908.1 | WP_000443908.1 | aspartate--tRNA(Asn) ligase |
| 14(2) | WP_087931729.1 | WP_087931729.1 | aspartate--tRNA(Asn) ligase |
| 14(1) | WP_199695251.1 | WP_199695251.1 | aspartate--tRNA(Asn) ligase |
| 14(1) | WP_262725729.1 | WP_262725729.1 | aspartate--tRNA(Asn) ligase |
| 14(1) | WP_098907647.1 | WP_098907647.1 | aspartate--tRNA(Asn) ligase |
| 14(1) | WP_242261453.1 | WP_242261453.1 | aspartate--tRNA(Asn) ligase |
| 14(1) | WP_308902527.1 | WP_308902527.1 | aspartate--tRNA(Asn) ligase |
| 14(1) | WP_098266645.1 | WP_098266645.1 | aspartate--tRNA(Asn) ligase |
| 14(1) | WP_002148335.1 | WP_002148335.1 | aspartate--tRNA(Asn) ligase |
| 14(4) | WP_000443907.1 | WP_000443907.1 | aspartate--tRNA(Asn) ligase |
| 14(1) | WP_193653024.1 | WP_193653024.1 | aspartate--tRNA(Asn) ligase |
| 14(1) | WP_023521812.1 | WP_023521812.1 | aspartate--tRNA(Asn) ligase |
| 14(10) | WP_000377339.1 | WP_000377339.1 | aspartate--tRNA(Asn) ligase |
| 14(1) | WP_071719986.1 | WP_071719986.1 | aspartate--tRNA(Asn) ligase |
| 14(2) | WP_103655007.1 | WP_103655007.1 | aspartate--tRNA(Asn) ligase |
| 14(1) | WP_000529811.1 | WP_000529811.1 | aspartate--tRNA(Asn) ligase |
| 14(4) | WP_103626229.1 | WP_103626229.1 | aspartate--tRNA(Asn) ligase |

| | | |
|---|---|---|
| 14(4) | WP_071679488.1 | WP_071679488.1 aspartate--tRNA(Asn) ligase |
| 14(1) | WP_097905183.1 | WP_097905183.1 aspartate--tRNA(Asn) ligase |
| 14(1) | WP_270699376.1 | WP_270699376.1 aspartate--tRNA(Asn) ligase |
| 14(1) | WP_098768170.1 | WP_098768170.1 aspartate--tRNA(Asn) ligase |
| 14(3) | WP_048540908.1 | WP_048540908.1 aspartate--tRNA(Asn) ligase |
| 14(1) | WP_063219640.1 | WP_063219640.1 aspartate--tRNA(Asn) ligase |
| 14(1) | WP_098288682.1 | WP_098288682.1 aspartate--tRNA(Asn) ligase |
| 14(1) | WP_078993366.1 | WP_078993366.1 aspartate--tRNA(Asn) ligase |
| 14(1) | WP_306007800.1 | WP_306007800.1 aspartate--tRNA(Asn) ligase |
| 14(1) | WP_227558285.1 | WP_227558285.1 aspartate--tRNA(Asn) ligase |
| 14(1) | WP_286017546.1 | WP_286017546.1 aspartate--tRNA(Asn) ligase |
| 14(1) | WP_106822160.1 | WP_106822160.1 aspartate--tRNA(Asn) ligase |
| 14(1) | WP_098148055.1 | WP_098148055.1 aspartate--tRNA(Asn) ligase |
| 14(1) | WP_071724716.1 | WP_071724716.1 aspartate--tRNA(Asn) ligase |
| 14(10) | WP_000529807.1 | WP_000529807.1 aspartate--tRNA(Asn) ligase |
| 14(1) | WP_144492098.1 | WP_144492098.1 aspartate--tRNA(Asn) ligase |
| 14(1) | WP_098435998.1 | WP_098435998.1 aspartate--tRNA(Asn) ligase |
| 14(1) | WP_000529813.1 | WP_000529813.1 aspartate--tRNA(Asn) ligase |
| 14(1) | WP_075666790.1 | WP_075666790.1 aspartate--tRNA(Asn) ligase |
| 14(1) | WP_012260831.1 | WP_012260831.1 aspartate--tRNA(Asn) ligase |
| 14(6) | WP_000443906.1 | WP_000443906.1 aspartate--tRNA(Asn) ligase |
| 14(1) | WP_000529812.1 | WP_000529812.1 aspartate--tRNA(Asn) ligase |
| 14(1) | WP_097821800.1 | WP_097821800.1 aspartate--tRNA(Asn) ligase |
| 14(3) | WP_017561046.1 | WP_017561046.1 aspartate--tRNA(Asn) ligase |
| 14(1) | WP_087982590.1 | WP_087982590.1 aspartate--tRNA(Asn) ligase |
| 14(1) | WP_144564268.1 | WP_144564268.1 aspartate--tRNA(Asn) ligase |
| 14(1) | WP_090688947.1 | WP_090688947.1 aspartate--tRNA(Asn) ligase |
| 14(1) | WP_033709023.1 | WP_033709023.1 aspartate--tRNA(Asn) ligase |
| 14(1) | WP_139867388.1 | WP_139867388.1 aspartate--tRNA(Asn) ligase |
| 14(1) | WP_000529815.1 | WP_000529815.1 aspartate--tRNA(Asn) ligase |
| 14(1) | WP_240379731.1 | WP_240379731.1 aspartate--tRNA(Asn) ligase |
| 14(1) | WP_078205523.1 | WP_078205523.1 aspartate--tRNA(Asn) ligase |
| 14(1) | WP_087956290.1 | WP_087956290.1 aspartate--tRNA(Asn) ligase |
| 14(1) | WP_143315261.1 | WP_143315261.1 aspartate--tRNA(Asn) ligase |
| 14(1) | WP_000443905.1 | WP_000443905.1 aspartate--tRNA(Asn) ligase |
| 14(1) | WP_029438088.1 | WP_029438088.1 aspartate--tRNA(Asn) ligase |
| 14(1) | WP_262729426.1 | WP_262729426.1 aspartate--tRNA(Asn) ligase |
| 14(2) | WP_098447522.1 | WP_098447522.1 aspartate--tRNA(Asn) ligase |
| 14(1) | WP_097846500.1 | WP_097846500.1 aspartate--tRNA(Asn) ligase |
| 14(3) | WP_200152287.1 | WP_200152287.1 aspartate--tRNA(Asn) ligase |
| 14(1) | WP_242264961.1 | WP_242264961.1 aspartate--tRNA(Asn) ligase |
| 14(1) | WP_144611184.1 | WP_144611184.1 aspartate--tRNA(Asn) ligase |
| 14(4) | WP_000377342.1 | WP_000377342.1 aspartate--tRNA(Asn) ligase |
| 14(1) | WP_098558850.1 | WP_098558850.1 aspartate--tRNA(Asn) ligase |
| 14(1) | WP_100910322.1 | WP_100910322.1 aspartate--tRNA(Asn) ligase |
| 14(2) | WP_078181899.1 | WP_078181899.1 aspartate--tRNA(Asn) ligase |
| 14(1) | WP_144488218.1 | WP_144488218.1 aspartate--tRNA(Asn) ligase |
| 14(1) | WP_215598793.1 | WP_215598793.1 aspartate--tRNA(Asn) ligase |
| 14(1) | WP_070805901.1 | WP_070805901.1 aspartate--tRNA(Asn) ligase |
| 14(1) | WP_098684013.1 | WP_098684013.1 aspartate--tRNA(Asn) ligase |
| 14(1) | WP_199677615.1 | WP_199677615.1 aspartate--tRNA(Asn) ligase |
| 14(1) | WP_277480685.1 | WP_277480685.1 aspartate--tRNA(Asn) ligase |
| 14(2) | WP_257808062.1 | WP_257808062.1 aspartate--tRNA(Asn) ligase |
| 14(1) | WP_098303180.1 | WP_098303180.1 aspartate--tRNA(Asn) ligase |
| 14(2) | WP_000377338.1 | WP_000377338.1 aspartate--tRNA(Asn) ligase |
| 14(2) | WP_000529805.1 | WP_000529805.1 aspartate--tRNA(Asn) ligase |
| 14(1) | WP_208019346.1 | WP_208019346.1 aspartate--tRNA(Asn) ligase |
| 14(1) | WP_147807657.1 | WP_147807657.1 aspartate--tRNA(Asn) ligase |
| 14(3) | WP_061688402.1 | WP_061688402.1 aspartate--tRNA(Asn) ligase |
| 14(1) | WP_078419972.1 | WP_078419972.1 aspartate--tRNA(Asn) ligase |
| 14(1) | WP_088313231.1 | WP_088313231.1 aspartate--tRNA(Asn) ligase |
| 14(1) | WP_289398694.1 | WP_289398694.1 aspartate--tRNA(Asn) ligase |
| 14(1) | WP_151630608.1 | WP_151630608.1 aspartate--tRNA(Asn) ligase |
| 14(1) | WP_063223274.1 | WP_063223274.1 aspartate--tRNA(Asn) ligase |
| 14(1) | WP_098167596.1 | WP_098167596.1 aspartate--tRNA(Asn) ligase |
| 14(1) | WP_098264283.1 | WP_098264283.1 aspartate--tRNA(Asn) ligase |
| 14(1) | WP_242400021.1 | WP_242400021.1 aspartate--tRNA(Asn) ligase |
| 14(3) | WP_060630644.1 | WP_060630644.1 aspartate--tRNA(Asn) ligase |
| 14(1) | WP_000443911.1 | WP_000443911.1 aspartate--tRNA(Asn) ligase |
| 14(3) | WP_053564634.1 | WP_053564634.1 aspartate--tRNA(Asn) ligase |
| 14(2) | WP_090976423.1 | WP_090976423.1 aspartate--tRNA(Asn) ligase |
| 14(1) | WP_275175508.1 | WP_275175508.1 aspartate--tRNA(Asn) ligase |
| 14(1) | WP_137052040.1 | WP_137052040.1 aspartate--tRNA(Asn) ligase |
| 14(1) | WP_098319062.1 | WP_098319062.1 aspartate--tRNA(Asn) ligase |
| 14(1) | WP_159146502.1 | WP_159146502.1 aspartate--tRNA(Asn) ligase |
| 14(1) | WP_097906565.1 | WP_097906565.1 aspartate--tRNA(Asn) ligase |
| 14(1) | WP_215591200.1 | WP_215591200.1 aspartate--tRNA(Asn) ligase |
| 14(3) | WP_065228767.1 | WP_065228767.1 aspartate--tRNA(Asn) ligase |
| 15(1) | WP_094155418.1 | WP_094155418.1 arginine--tRNA ligase |
| 15(2) | WP_270699370.1 | WP_270699370.1 arginine--tRNA ligase |
| 15(3) | WP_000384894.1 | WP_000384894.1 arginine--tRNA ligase |
| 15(1) | WP_098400687.1 | WP_098400687.1 arginine--tRNA ligase |
| 15(1) | WP_000385017.1 | WP_000385017.1 arginine--tRNA ligase |
| 15(1) | WP_215572744.1 | WP_215572744.1 arginine--tRNA ligase |
| 15(1) | WP_074570403.1 | WP_074570403.1 arginine--tRNA ligase |
| 15(1) | WP_098365935.1 | WP_098365935.1 arginine--tRNA ligase |
| 15(1) | WP_010346300.1 | WP_010346300.1 arginine--tRNA ligase |
| 15(1) | WP_103047835.1 | WP_103047835.1 arginine--tRNA ligase |
| 15(1) | WP_139867397.1 | WP_139867397.1 arginine--tRNA ligase |
| 15(1) | WP_098519057.1 | WP_098519057.1 arginine--tRNA ligase |
| 15(1) | WP_137078358.1 | WP_137078358.1 arginine--tRNA ligase |
| 15(1) | WP_262787109.1 | WP_262787109.1 arginine--tRNA ligase |
| 15(1) | WP_088021924.1 | WP_088021924.1 arginine--tRNA ligase |
| 15(1) | WP_139895514.1 | WP_139895514.1 arginine--tRNA ligase |
| 15(1) | WP_229137980.1 | WP_229137980.1 arginine--tRNA ligase |
| 15(1) | WP_144488215.1 | WP_144488215.1 arginine--tRNA ligase |
| 15(1) | WP_098296559.1 | WP_098296559.1 arginine--tRNA ligase |
| 15(3) | WP_097911856.1 | WP_097911856.1 arginine--tRNA ligase |
| 15(1) | WP_029438081.1 | WP_029438081.1 arginine--tRNA ligase |
| 15(1) | WP_286124771.1 | WP_286124771.1 arginine--tRNA ligase |
| 15(1) | WP_060630638.1 | WP_060630638.1 arginine--tRNA ligase |
| 15(1) | WP_212956549.1 | WP_212956549.1 arginine--tRNA ligase |
| 15(1) | WP_147807662.1 | WP_147807662.1 arginine--tRNA ligase |

| | | |
|---|---|---|
| 15(1) | WP_086419827.1 | WP_086419827.1 arginine--tRNA ligase |
| 15(1) | WP_071719994.1 | WP_071719994.1 arginine--tRNA ligase |
| 15(1) | WP_098204196.1 | WP_098204196.1 arginine--tRNA ligase |
| 15(1) | WP_289398689.1 | WP_289398689.1 arginine--tRNA ligase |
| 15(1) | WP_087956283.1 | WP_087956283.1 arginine--tRNA ligase |
| 15(1) | WP_262719891.1 | WP_262719891.1 arginine--tRNA ligase |
| 15(1) | WP_045243841.1 | WP_045243841.1 arginine--tRNA ligase |
| 15(1) | WP_070805895.1 | WP_070805895.1 arginine--tRNA ligase |
| 15(1) | WP_258126005.1 | WP_258126005.1 arginine--tRNA ligase |
| 15(1) | WP_166704978.1 | WP_166704978.1 arginine--tRNA ligase |
| 15(1) | WP_143881064.1 | WP_143881064.1 arginine--tRNA ligase |
| 15(1) | WP_242400013.1 | WP_242400013.1 arginine--tRNA ligase |
| 15(1) | WP_216975166.1 | WP_216975166.1 arginine--tRNA ligase |
| 15(1) | WP_068496267.1 | WP_068496267.1 arginine--tRNA ligase |
| 15(1) | WP_270728490.1 | WP_270728490.1 arginine--tRNA ligase |
| 15(1) | WP_199695246.1 | WP_199695246.1 arginine--tRNA ligase |
| 15(1) | WP_126665672.1 | WP_126665672.1 arginine--tRNA ligase |
| 15(2) | WP_087982605.1 | WP_087982605.1 arginine--tRNA ligase |
| 15(2) | WP_065228752.1 | WP_065228752.1 arginine--tRNA ligase |
| 15(1) | WP_098254032.1 | WP_098254032.1 arginine--tRNA ligase |
| 15(1) | WP_061673874.1 | WP_061673874.1 arginine--tRNA ligase |
| 15(3) | WP_098266637.1 | WP_098266637.1 arginine--tRNA ligase |
| 15(1) | WP_242251754.1 | WP_242251754.1 arginine--tRNA ligase |
| 15(2) | WP_098431646.1 | WP_098431646.1 arginine--tRNA ligase |
| 15(1) | WP_098508766.1 | WP_098508766.1 arginine--tRNA ligase |
| 15(1) | WP_242261447.1 | WP_242261447.1 arginine--tRNA ligase |
| 15(1) | WP_127812909.1 | WP_127812909.1 arginine--tRNA ligase |
| 15(1) | WP_002148359.1 | WP_002148359.1 arginine--tRNA ligase |
| 15(2) | WP_071679484.1 | WP_071679484.1 arginine--tRNA ligase |
| 15(1) | WP_212805117.1 | WP_212805117.1 arginine--tRNA ligase |
| 15(1) | WP_007430116.1 | WP_007430116.1 arginine--tRNA ligase |
| 15(1) | WP_215598795.1 | WP_215598795.1 arginine--tRNA ligase |
| 15(1) | WP_098288676.1 | WP_098288676.1 arginine--tRNA ligase |
| 15(1) | WP_229522870.1 | WP_229522870.1 arginine--tRNA ligase |
| 15(1) | WP_071717040.1 | WP_071717040.1 arginine--tRNA ligase |
| 15(1) | WP_002158057.1 | WP_002158057.1 arginine--tRNA ligase |
| 15(1) | WP_227558287.1 | WP_227558287.1 arginine--tRNA ligase |
| 15(2) | WP_098264277.1 | WP_098264277.1 arginine--tRNA ligase |
| 15(1) | WP_129549123.1 | WP_129549123.1 arginine--tRNA ligase |
| 15(1) | WP_208752248.1 | WP_208752248.1 arginine--tRNA ligase |
| 15(1) | WP_157418145.1 | WP_157418145.1 arginine--tRNA ligase |
| 15(1) | WP_012260823.1 | WP_012260823.1 arginine--tRNA ligase |
| 15(1) | WP_097952798.1 | WP_097952798.1 arginine--tRNA ligase |
| 15(1) | WP_000384895.1 | WP_000384895.1 arginine--tRNA ligase |
| 15(1) | WP_097844625.1 | WP_097844625.1 arginine--tRNA ligase |
| 15(1) | WP_088032759.1 | WP_088032759.1 arginine--tRNA ligase |
| 15(1) | WP_307349939.1 | WP_307349939.1 arginine--tRNA ligase |
| 15(1) | WP_151630615.1 | WP_151630615.1 arginine--tRNA ligase |
| 15(1) | WP_000384891.1 | WP_000384891.1 arginine--tRNA ligase |
| 15(6) | WP_000384853.1 | WP_000384853.1 arginine--tRNA ligase |
| 15(1) | WP_098230733.1 | WP_098230733.1 arginine--tRNA ligase |
| 15(1) | WP_250244796.1 | WP_250244796.1 arginine--tRNA ligase |
| 15(4) | WP_023521805.1 | WP_023521805.1 arginine--tRNA ligase |
| 15(1) | WP_025685019.1 | WP_025685019.1 arginine--tRNA ligase |
| 15(1) | WP_088609108.1 | WP_088609108.1 arginine--tRNA ligase |
| 15(1) | WP_088119565.1 | WP_088119565.1 arginine--tRNA ligase |
| 15(5) | WP_000384898.1 | WP_000384898.1 arginine--tRNA ligase |
| 15(5) | WP_000384855.1 | WP_000384855.1 arginine--tRNA ligase |
| 15(1) | WP_190299130.1 | WP_190299130.1 arginine--tRNA ligase |
| 15(2) | WP_103626224.1 | WP_103626224.1 arginine--tRNA ligase |
| 15(3) | WP_000384892.1 | WP_000384892.1 arginine--tRNA ligase |
| 15(1) | WP_277480666.1 | WP_277480666.1 arginine--tRNA ligase |
| 15(1) | WP_256424319.1 | WP_256424319.1 arginine--tRNA ligase |
| 15(1) | WP_223617615.1 | WP_223617615.1 arginine--tRNA ligase |
| 15(1) | WP_137076284.1 | WP_137076284.1 arginine--tRNA ligase |
| 15(5) | WP_097821804.1 | WP_097821804.1 arginine--tRNA ligase |
| 15(3) | WP_016512888.1 | WP_016512888.1 arginine--tRNA ligase |
| 15(5) | WP_000384925.1 | WP_000384925.1 arginine--tRNA ligase |
| 15(1) | WP_163255245.1 | WP_163255245.1 arginine--tRNA ligase |
| 15(1) | WP_215591193.1 | WP_215591193.1 arginine--tRNA ligase |
| 15(1) | WP_087995252.1 | WP_087995252.1 arginine--tRNA ligase |
| 15(1) | WP_074601327.1 | WP_074601327.1 arginine--tRNA ligase |
| 15(1) | WP_025679789.1 | WP_025679789.1 arginine--tRNA ligase |
| 15(1) | WP_098678279.1 | WP_098678279.1 arginine--tRNA ligase |
| 15(1) | WP_069011417.1 | WP_069011417.1 arginine--tRNA ligase |
| 15(1) | WP_000385012.1 | WP_000385012.1 arginine--tRNA ligase |
| 15(2) | WP_098077002.1 | WP_098077002.1 arginine--tRNA ligase |
| 15(1) | WP_064473947.1 | WP_064473947.1 arginine--tRNA ligase |
| 15(2) | WP_097920543.1 | WP_097920543.1 arginine--tRNA ligase |
| 15(2) | WP_073542132.1 | WP_073542132.1 arginine--tRNA ligase |
| 15(1) | WP_137062786.1 | WP_137062786.1 arginine--tRNA ligase |
| 15(1) | WP_088078454.1 | WP_088078454.1 arginine--tRNA ligase |
| 15(1) | WP_088298130.1 | WP_088298130.1 arginine--tRNA ligase |
| 15(3) | WP_063547141.1 | WP_063547141.1 arginine--tRNA ligase |
| 15(1) | WP_002141577.1 | WP_002141577.1 arginine--tRNA ligase |
| 15(1) | WP_002189450.1 | WP_002189450.1 arginine--tRNA ligase |
| 15(1) | WP_071724705.1 | WP_071724705.1 arginine--tRNA ligase |
| 15(1) | WP_042512834.1 | WP_042512834.1 arginine--tRNA ligase |
| 15(1) | WP_149238899.1 | WP_149238899.1 arginine--tRNA ligase |
| 15(1) | WP_144473243.1 | WP_144473243.1 arginine--tRNA ligase |
| 15(1) | WP_262725737.1 | WP_262725737.1 arginine--tRNA ligase |
| 15(5) | WP_053564640.1 | WP_053564640.1 arginine--tRNA ligase |
| 15(1) | WP_106822172.1 | WP_106822172.1 arginine--tRNA ligase |
| 15(1) | WP_078205516.1 | WP_078205516.1 arginine--tRNA ligase |
| 15(1) | WP_013309971.1 | WP_013309971.1 arginine--tRNA ligase |
| 15(2) | WP_017561869.1 | WP_017561869.1 arginine--tRNA ligase |
| 15(1) | WP_144611178.1 | WP_144611178.1 arginine--tRNA ligase |
| 15(1) | WP_207994173.1 | WP_207994173.1 arginine--tRNA ligase |
| 15(1) | WP_098776440.1 | WP_098776440.1 arginine--tRNA ligase |
| 15(3) | WP_088063225.1 | WP_088063225.1 arginine--tRNA ligase |
| 15(2) | WP_098148050.1 | WP_098148050.1 arginine--tRNA ligase |
| 15(1) | WP_098907648.1 | WP_098907648.1 arginine--tRNA ligase |
| 15(9) | WP_000385013.1 | WP_000385013.1 arginine--tRNA ligase |
| 15(1) | WP_063223266.1 | WP_063223266.1 arginine--tRNA ligase |

| | | | |
|---|---|---|---|
| 15(1) | WP_002031641.1 | WP_002031641.1 | arginine--tRNA ligase |
| 15(1) | WP_270716414.1 | WP_270716414.1 | arginine--tRNA ligase |
| 15(1) | WP_305755050.1 | WP_305755050.1 | arginine--tRNA ligase |
| 15(2) | WP_000384852.1 | WP_000384852.1 | arginine--tRNA ligase |
| 15(1) | WP_017149783.1 | WP_017149783.1 | arginine--tRNA ligase |
| 15(1) | WP_100910325.1 | WP_100910325.1 | arginine--tRNA ligase |
| 15(1) | WP_000385007.1 | WP_000385007.1 | arginine--tRNA ligase |
| 15(2) | WP_197241540.1 | WP_197241540.1 | arginine--tRNA ligase |
| 15(1) | WP_071736265.1 | WP_071736265.1 | arginine--tRNA ligase |
| 15(1) | WP_075395494.1 | WP_075395494.1 | arginine--tRNA ligase |
| 15(1) | WP_283655682.1 | WP_283655682.1 | arginine--tRNA ligase |
| 15(1) | WP_095691019.1 | WP_095691019.1 | arginine--tRNA ligase |
| 15(1) | WP_098360829.1 | WP_098360829.1 | arginine--tRNA ligase |
| 15(1) | WP_270723908.1 | WP_270723908.1 | arginine--tRNA ligase |
| 15(1) | WP_290403364.1 | WP_290403364.1 | arginine--tRNA ligase |
| 15(1) | WP_199677619.1 | WP_199677619.1 | arginine--tRNA ligase |
| 15(2) | WP_098133698.1 | WP_098133698.1 | arginine--tRNA ligase |
| 15(1) | WP_088058511.1 | WP_088058511.1 | arginine--tRNA ligase |
| 15(1) | WP_149190446.1 | WP_149190446.1 | arginine--tRNA ligase |
| 15(1) | WP_103655003.1 | WP_103655003.1 | arginine--tRNA ligase |
| 15(1) | WP_098088262.1 | WP_098088262.1 | arginine--tRNA ligase |
| 15(1) | WP_078419978.1 | WP_078419978.1 | arginine--tRNA ligase |
| 15(1) | WP_044792571.1 | WP_044792571.1 | arginine--tRNA ligase |
| 15(1) | WP_014281168.1 | WP_014281168.1 | arginine--tRNA ligase |
| 15(1) | WP_243500436.1 | WP_243500436.1 | arginine--tRNA ligase |
| 15(1) | WP_098377168.1 | WP_098377168.1 | arginine--tRNA ligase |
| 15(4) | WP_000384897.1 | WP_000384897.1 | arginine--tRNA ligase |
| 15(1) | WP_093291047.1 | WP_093291047.1 | arginine--tRNA ligase |
| 15(1) | WP_173601441.1 | WP_173601441.1 | arginine--tRNA ligase |
| 15(1) | WP_000385003.1 | WP_000385003.1 | arginine--tRNA ligase |
| 15(1) | WP_192510499.1 | WP_192510499.1 | arginine--tRNA ligase |
| 15(1) | WP_098991583.1 | WP_098991583.1 | arginine--tRNA ligase |

| | | | |
|---|---|---|---|
| 16(1) | WP_284566223.1 | WP_284566223.1 | M36 family metallopeptidase |
| 16(2) | WP_063223271.1 | WP_063223271.1 | M36 family metallopeptidase |
| 16(1) | WP_098856763.1 | WP_098856763.1 | M36 family metallopeptidase |
| 16(1) | WP_086419831.1 | WP_086419831.1 | M36 family metallopeptidase |
| 16(2) | WP_064473943.1 | WP_064473943.1 | M36 family metallopeptidase |
| 16(2) | WP_017561871.1 | WP_017561871.1 | M36 family metallopeptidase |
| 16(1) | WP_098239631.1 | WP_098239631.1 | M36 family metallopeptidase |
| 16(1) | WP_147786240.1 | WP_147786240.1 | M36 family metallopeptidase |
| 16(3) | WP_000482268.1 | WP_000482268.1 | M36 family metallopeptidase |
| 16(3) | WP_097952790.1 | WP_097952790.1 | M36 family metallopeptidase |
| 16(1) | WP_098319881.1 | WP_098319881.1 | M36 family metallopeptidase |
| 16(1) | WP_002031632.1 | WP_002031632.1 | M36 family metallopeptidase |
| 16(10) | WP_000482298.1 | WP_000482298.1 | M36 family metallopeptidase |
| 16(1) | WP_097889494.1 | WP_097889494.1 | M36 family metallopeptidase |
| 16(1) | WP_144506824.1 | WP_144506824.1 | M36 family metallopeptidase |
| 16(1) | WP_070805899.1 | WP_070805899.1 | M36 family metallopeptidase |
| 16(1) | WP_147807659.1 | WP_147807659.1 | M36 family metallopeptidase |
| 16(1) | WP_097925473.1 | WP_097925473.1 | M36 family metallopeptidase |
| 16(7) | WP_000482297.1 | WP_000482297.1 | M36 family metallopeptidase |
| 16(1) | WP_002148345.1 | WP_002148345.1 | M36 family metallopeptidase |
| 16(6) | WP_063219642.1 | WP_063219642.1 | M36 family metallopeptidase |
| 16(2) | WP_098204199.1 | WP_098204199.1 | M36 family metallopeptidase |
| 16(1) | WP_166704984.1 | WP_166704984.1 | M36 family metallopeptidase |
| 16(31) | WP_000482274.1 | WP_000482274.1 | M36 family metallopeptidase |
| 16(1) | WP_098840610.1 | WP_098840610.1 | M36 family metallopeptidase |
| 16(1) | WP_098264280.1 | WP_098264280.1 | M36 family metallopeptidase |
| 16(1) | WP_098638816.1 | WP_098638816.1 | M36 family metallopeptidase |
| 16(10) | WP_000482291.1 | WP_000482291.1 | M36 family metallopeptidase |
| 16(1) | WP_098199508.1 | WP_098199508.1 | M36 family metallopeptidase |
| 16(1) | WP_000932732.1 | WP_000932732.1 | M36 family metallopeptidase |
| 16(1) | WP_144473244.1 | WP_144473244.1 | M36 family metallopeptidase |
| 16(3) | WP_044792566.1 | WP_044792566.1 | M36 family metallopeptidase |
| 16(1) | WP_262719890.1 | WP_262719890.1 | M36 family metallopeptidase |
| 16(1) | WP_242261451.1 | WP_242261451.1 | M36 family metallopeptidase |
| 16(1) | WP_097987494.1 | WP_097987494.1 | M36 family metallopeptidase |
| 16(3) | WP_097961540.1 | WP_097961540.1 | M36 family metallopeptidase |
| 16(1) | WP_098670169.1 | WP_098670169.1 | M36 family metallopeptidase |
| 16(1) | WP_215572749.1 | WP_215572749.1 | M36 family metallopeptidase |
| 16(1) | WP_236804418.1 | WP_236804418.1 | M36 family metallopeptidase |
| 16(1) | WP_262725731.1 | WP_262725731.1 | M36 family metallopeptidase |
| 16(2) | WP_000482281.1 | WP_000482281.1 | M36 family metallopeptidase |
| 16(1) | WP_243500443.1 | WP_243500443.1 | M36 family metallopeptidase |
| 16(1) | WP_286017543.1 | WP_286017543.1 | M36 family metallopeptidase |
| 16(1) | WP_088078457.1 | WP_088078457.1 | M36 family metallopeptidase |
| 16(2) | WP_087956287.1 | WP_087956287.1 | M36 family metallopeptidase |
| 16(1) | WP_137035710.1 | WP_137035710.1 | M36 family metallopeptidase |
| 16(1) | WP_202283182.1 | WP_202283182.1 | M36 family metallopeptidase |
| 16(1) | WP_002141586.1 | WP_002141586.1 | M36 family metallopeptidase |
| 16(1) | WP_100910323.1 | WP_100910323.1 | M36 family metallopeptidase |
| 16(2) | WP_173601444.1 | WP_173601444.1 | M36 family metallopeptidase |
| 16(1) | WP_286124777.1 | WP_286124777.1 | M36 family metallopeptidase |
| 16(1) | WP_090688940.1 | WP_090688940.1 | M36 family metallopeptidase |
| 16(1) | WP_277480678.1 | WP_277480678.1 | M36 family metallopeptidase |
| 16(1) | WP_097905185.1 | WP_097905185.1 | M36 family metallopeptidase |
| 16(1) | WP_071719987.1 | WP_071719987.1 | M36 family metallopeptidase |
| 16(1) | WP_002158051.1 | WP_002158051.1 | M36 family metallopeptidase |
| 16(1) | WP_012260829.1 | WP_012260829.1 | M36 family metallopeptidase |
| 16(1) | WP_207994167.1 | WP_207994167.1 | M36 family metallopeptidase |
| 16(1) | WP_088058527.1 | WP_088058527.1 | M36 family metallopeptidase |
| 16(1) | WP_017149777.1 | WP_017149777.1 | M36 family metallopeptidase |
| 16(1) | WP_229136362.1 | WP_229136362.1 | M36 family metallopeptidase |
| 16(1) | WP_242251766.1 | WP_242251766.1 | M36 family metallopeptidase |
| 16(1) | WP_098288680.1 | WP_098288680.1 | M36 family metallopeptidase |
| 16(2) | WP_098296563.1 | WP_098296563.1 | M36 family metallopeptidase |
| 16(2) | WP_098111903.1 | WP_098111903.1 | M36 family metallopeptidase |
| 16(6) | WP_073542128.1 | WP_073542128.1 | M36 family metallopeptidase |
| 16(2) | WP_000482300.1 | WP_000482300.1 | M36 family metallopeptidase |
| 16(1) | WP_305163860.1 | WP_305163860.1 | M36 family metallopeptidase |
| 16(1) | WP_215601062.1 | WP_215601062.1 | M36 family metallopeptidase |
| 16(2) | WP_000482296.1 | WP_000482296.1 | M36 family metallopeptidase |
| 16(1) | WP_242255487.1 | WP_242255487.1 | M36 family metallopeptidase |

| | | |
|---|---|---|
| 16(14) | WP_000482266.1 | WP_000482266.1 M36 family metallopeptidase |
| 16(1) | WP_144611182.1 | WP_144611182.1 M36 family metallopeptidase |
| 16(1) | WP_060630642.1 | WP_060630642.1 M36 family metallopeptidase |
| 16(1) | WP_029438086.1 | WP_029438086.1 M36 family metallopeptidase |
| 16(1) | WP_208752245.1 | WP_208752245.1 M36 family metallopeptidase |
| 16(1) | WP_000482269.1 | WP_000482269.1 M36 family metallopeptidase |
| 16(1) | WP_097906562.1 | WP_097906562.1 M36 family metallopeptidase |
| 16(1) | WP_098150445.1 | WP_098150445.1 M36 family metallopeptidase |
| 16(1) | WP_078419974.1 | WP_078419974.1 M36 family metallopeptidase |
| 16(2) | WP_127812907.1 | WP_127812907.1 M36 family metallopeptidase |
| 16(1) | WP_242400019.1 | WP_242400019.1 M36 family metallopeptidase |
| 16(1) | WP_144655915.1 | WP_144655915.1 M36 family metallopeptidase |
| 16(1) | WP_193653025.1 | WP_193653025.1 M36 family metallopeptidase |
| 16(1) | WP_088032772.1 | WP_088032772.1 M36 family metallopeptidase |
| 16(1) | WP_212956543.1 | WP_212956543.1 M36 family metallopeptidase |
| 16(1) | WP_098519063.1 | WP_098519063.1 M36 family metallopeptidase |
| 16(4) | WP_000482282.1 | WP_000482282.1 M36 family metallopeptidase |
| 16(1) | WP_088119569.1 | WP_088119569.1 M36 family metallopeptidase |
| 16(1) | WP_000482289.1 | WP_000482289.1 M36 family metallopeptidase |
| 16(1) | WP_098776434.1 | WP_098776434.1 M36 family metallopeptidase |
| 16(9) | WP_046959291.1 | WP_046959291.1 M36 family metallopeptidase |
| 16(1) | WP_097846499.1 | WP_097846499.1 M36 family metallopeptidase |
| 16(3) | WP_098266643.1 | WP_098266643.1 M36 family metallopeptidase |
| 16(1) | WP_098250521.1 | WP_098250521.1 M36 family metallopeptidase |
| 16(3) | WP_098360833.1 | WP_098360833.1 M36 family metallopeptidase |
| 16(5) | WP_000482267.1 | WP_000482267.1 M36 family metallopeptidase |
| 16(1) | WP_137052043.1 | WP_137052043.1 M36 family metallopeptidase |
| 16(1) | WP_106822162.1 | WP_106822162.1 M36 family metallopeptidase |
| 16(1) | WP_131231112.1 | WP_131231112.1 M36 family metallopeptidase |
| 16(7) | WP_000482302.1 | WP_000482302.1 M36 family metallopeptidase |
| 16(1) | WP_098167597.1 | WP_098167597.1 M36 family metallopeptidase |
| 16(1) | WP_208019343.1 | WP_208019343.1 M36 family metallopeptidase |
| 16(1) | WP_151630611.1 | WP_151630611.1 M36 family metallopeptidase |
| 16(2) | WP_098991576.1 | WP_098991576.1 M36 family metallopeptidase |
| 17(1) | WP_216390040.1 | WP_216390040.1 leucine--tRNA ligase |
| 17(1) | WP_214328826.1 | WP_214328826.1 isoleucine--tRNA ligase |
| 17(1) | WP_002158054.1 | WP_002158054.1 isoleucine--tRNA ligase |
| 17(1) | WP_208019342.1 | WP_208019342.1 isoleucine--tRNA ligase |
| 17(2) | WP_078185407.1 | WP_078185407.1 isoleucine--tRNA ligase |
| 17(3) | WP_053564637.1 | WP_053564637.1 isoleucine--tRNA ligase |
| 17(1) | WP_098312840.1 | WP_098312840.1 isoleucine--tRNA ligase |
| 17(1) | WP_151630612.1 | WP_151630612.1 isoleucine--tRNA ligase |
| 17(1) | WP_075395491.1 | WP_075395491.1 isoleucine--tRNA ligase |
| 17(1) | WP_098952613.1 | WP_098952613.1 isoleucine--tRNA ligase |
| 17(1) | WP_098230735.1 | WP_098230735.1 isoleucine--tRNA ligase |
| 17(1) | WP_242400017.1 | WP_242400017.1 isoleucine--tRNA ligase |
| 17(1) | WP_078419975.1 | WP_078419975.1 isoleucine--tRNA ligase |
| 17(1) | WP_240379730.1 | WP_240379730.1 isoleucine--tRNA ligase |
| 17(1) | WP_098991577.1 | WP_098991577.1 isoleucine--tRNA ligase |
| 17(1) | WP_284566222.1 | WP_284566222.1 isoleucine--tRNA ligase |
| 17(2) | WP_000754906.1 | WP_000754906.1 isoleucine--tRNA ligase |
| 17(1) | WP_088032765.1 | WP_088032765.1 isoleucine--tRNA ligase |
| 17(1) | WP_000754910.1 | WP_000754910.1 isoleucine--tRNA ligase |
| 17(1) | WP_088078456.1 | WP_088078456.1 isoleucine--tRNA ligase |
| 17(2) | WP_071736269.1 | WP_071736269.1 isoleucine--tRNA ligase |
| 17(1) | WP_199695249.1 | WP_199695249.1 isoleucine--tRNA ligase |
| 17(1) | WP_102371692.1 | WP_102371692.1 methionine--tRNA ligase |
| 17(1) | WP_098235618.1 | WP_098235618.1 isoleucine--tRNA ligase |
| 17(2) | WP_098678281.1 | WP_098678281.1 isoleucine--tRNA ligase |
| 17(1) | WP_088119568.1 | WP_088119568.1 isoleucine--tRNA ligase |
| 17(1) | WP_071719990.1 | WP_071719990.1 isoleucine--tRNA ligase |
| 17(1) | WP_144488217.1 | WP_144488217.1 isoleucine--tRNA ligase |
| 17(1) | WP_215559058.1 | WP_215559058.1 isoleucine--tRNA ligase |
| 17(1) | WP_289398692.1 | WP_289398692.1 isoleucine--tRNA ligase |
| 17(1) | WP_106822166.1 | WP_106822166.1 isoleucine--tRNA ligase |
| 17(1) | WP_229137983.1 | WP_229137983.1 isoleucine--tRNA ligase |
| 17(1) | WP_144476566.1 | WP_144476566.1 isoleucine--tRNA ligase |
| 17(6) | WP_000754927.1 | WP_000754927.1 isoleucine--tRNA ligase |
| 17(1) | WP_199795349.1 | WP_199795349.1 isoleucine--tRNA ligase |
| 17(1) | WP_097905188.1 | WP_097905188.1 isoleucine--tRNA ligase |
| 17(1) | WP_262729421.1 | WP_262729421.1 isoleucine--tRNA ligase |
| 17(1) | WP_097987497.1 | WP_097987497.1 isoleucine--tRNA ligase |
| 17(1) | WP_166704982.1 | WP_166704982.1 isoleucine--tRNA ligase |
| 17(1) | WP_144506822.1 | WP_144506822.1 isoleucine--tRNA ligase |
| 17(1) | WP_088609111.1 | WP_088609111.1 isoleucine--tRNA ligase |
| 17(1) | WP_000754914.1 | WP_000754914.1 isoleucine--tRNA ligase |
| 17(1) | WP_098239633.1 | WP_098239633.1 isoleucine--tRNA ligase |
| 17(1) | WP_078993368.1 | WP_078993368.1 isoleucine--tRNA ligase |
| 17(4) | WP_065483396.1 | WP_065483396.1 isoleucine--tRNA ligase |
| 17(1) | WP_087956286.1 | WP_087956286.1 isoleucine--tRNA ligase |
| 17(2) | WP_163255251.1 | WP_163255251.1 isoleucine--tRNA ligase |
| 17(1) | WP_000754935.1 | WP_000754935.1 isoleucine--tRNA ligase |
| 17(1) | WP_071679487.1 | WP_071679487.1 isoleucine--tRNA ligase |
| 17(1) | WP_277480675.1 | WP_277480675.1 isoleucine--tRNA ligase |
| 17(2) | WP_098082494.1 | WP_098082494.1 isoleucine--tRNA ligase |
| 17(2) | WP_071714359.1 | WP_071714359.1 isoleucine--tRNA ligase |
| 17(1) | WP_017149778.1 | WP_017149778.1 isoleucine--tRNA ligase |
| 17(1) | WP_098519060.1 | WP_098519060.1 isoleucine--tRNA ligase |
| 17(2) | WP_097952792.1 | WP_097952792.1 isoleucine--tRNA ligase |
| 17(5) | WP_000754934.1 | WP_000754934.1 isoleucine--tRNA ligase |
| 17(1) | WP_000754917.1 | WP_000754917.1 isoleucine--tRNA ligase |
| 17(1) | WP_286124775.1 | WP_286124775.1 isoleucine--tRNA ligase |
| 17(1) | WP_103655005.1 | WP_103655005.1 isoleucine--tRNA ligase |
| 17(1) | WP_098854978.1 | WP_098854978.1 isoleucine--tRNA ligase |
| 17(1) | WP_215591198.1 | WP_215591198.1 isoleucine--tRNA ligase |
| 17(3) | WP_000754905.1 | WP_000754905.1 isoleucine--tRNA ligase |
| 17(3) | WP_000754919.1 | WP_000754919.1 isoleucine--tRNA ligase |
| 17(1) | WP_270691241.1 | WP_270691241.1 isoleucine--tRNA ligase |
| 17(1) | WP_078205520.1 | WP_078205520.1 isoleucine--tRNA ligase |
| 17(1) | WP_270728500.1 | WP_270728500.1 isoleucine--tRNA ligase |
| 17(2) | WP_098360832.1 | WP_098360832.1 isoleucine--tRNA ligase |
| 17(1) | WP_012260827.1 | WP_012260827.1 isoleucine--tRNA ligase |
| 17(1) | WP_215572747.1 | WP_215572747.1 isoleucine--tRNA ligase |

| | | |
|---|---|---|
| 17(1) | WP_098365937.1 | WP_098365937.1 isoleucine--tRNA ligase |
| 17(1) | WP_097911848.1 | WP_097911848.1 isoleucine--tRNA ligase |
| 17(2) | WP_098447526.1 | WP_098447526.1 isoleucine--tRNA ligase |
| 17(1) | WP_086419830.1 | WP_086419830.1 isoleucine--tRNA ligase |
| 17(1) | WP_137052044.1 | WP_137052044.1 isoleucine--tRNA ligase |
| 17(1) | WP_098211173.1 | WP_098211173.1 isoleucine--tRNA ligase |
| 17(1) | WP_060630641.1 | WP_060630641.1 isoleucine--tRNA ligase |
| 17(1) | WP_215570366.1 | WP_215570366.1 isoleucine--tRNA ligase |
| 17(1) | WP_163247638.1 | WP_163247638.1 isoleucine--tRNA ligase |
| 17(2) | WP_149885344.1 | WP_149885344.1 isoleucine--tRNA ligase |
| 17(3) | WP_000416527.1 | WP_000416527.1 isoleucine--tRNA ligase |
| 17(1) | WP_098250518.1 | WP_098250518.1 isoleucine--tRNA ligase |
| 17(1) | WP_144655917.1 | WP_144655917.1 isoleucine--tRNA ligase |
| 17(1) | WP_270716416.1 | WP_270716416.1 isoleucine--tRNA ligase |
| 17(1) | WP_070805898.1 | WP_070805898.1 isoleucine--tRNA ligase |
| 17(1) | WP_215601061.1 | WP_215601061.1 isoleucine--tRNA ligase |
| 17(1) | WP_137036031.1 | WP_137036031.1 isoleucine--tRNA ligase |
| 17(1) | WP_141506912.1 | WP_141506912.1 isoleucine--tRNA ligase |
| 17(1) | WP_286017539.1 | WP_286017539.1 isoleucine--tRNA ligase |
| 17(1) | WP_262800527.1 | WP_262800527.1 isoleucine--tRNA ligase |
| 17(1) | WP_173601443.1 | WP_173601443.1 isoleucine--tRNA ligase |
| 17(2) | WP_143881067.1 | WP_143881067.1 isoleucine--tRNA ligase |
| 17(2) | WP_023521808.1 | WP_023521808.1 isoleucine--tRNA ligase |
| 17(1) | WP_157418142.1 | WP_157418142.1 isoleucine--tRNA ligase |
| 17(1) | WP_098355766.1 | WP_098355766.1 isoleucine--tRNA ligase |
| 17(1) | WP_208752246.1 | WP_208752246.1 isoleucine--tRNA ligase |
| 17(1) | WP_098948843.1 | WP_098948843.1 isoleucine--tRNA ligase |
| 17(1) | WP_106382853.1 | WP_106382853.1 isoleucine--tRNA ligase |
| 17(1) | WP_064473944.1 | WP_064473944.1 isoleucine--tRNA ligase |
| 17(1) | WP_002031635.1 | WP_002031635.1 isoleucine--tRNA ligase |
| 17(1) | WP_262725733.1 | WP_262725733.1 isoleucine--tRNA ligase |
| 17(1) | WP_048552298.1 | WP_048552298.1 isoleucine--tRNA ligase |
| 17(1) | WP_098150446.1 | WP_098150446.1 isoleucine--tRNA ligase |
| 17(1) | WP_258126006.1 | WP_258126006.1 isoleucine--tRNA ligase |
| 17(1) | WP_144492097.1 | WP_144492097.1 isoleucine--tRNA ligase |
| 17(1) | WP_264456847.1 | WP_264456847.1 isoleucine--tRNA ligase |
| 17(1) | WP_207994168.1 | WP_207994168.1 isoleucine--tRNA ligase |
| 17(2) | WP_137013290.1 | WP_137013290.1 isoleucine--tRNA ligase |
| 17(2) | WP_098451708.1 | WP_098451708.1 isoleucine--tRNA ligase |
| 17(1) | WP_097846498.1 | WP_097846498.1 isoleucine--tRNA ligase |
| 17(2) | WP_200457882.1 | WP_200457882.1 isoleucine--tRNA ligase |
| 17(1) | WP_088058520.1 | WP_088058520.1 isoleucine--tRNA ligase |
| 17(1) | WP_212956546.1 | WP_212956546.1 isoleucine--tRNA ligase |
| 17(1) | WP_202283181.1 | WP_202283181.1 isoleucine--tRNA ligase |
| 17(1) | WP_270723906.1 | WP_270723906.1 isoleucine--tRNA ligase |
| 17(2) | WP_063223269.1 | WP_063223269.1 isoleucine--tRNA ligase |
| 17(1) | WP_267414310.1 | WP_267414310.1 isoleucine--tRNA ligase |
| 17(1) | WP_061658097.1 | WP_061658097.1 isoleucine--tRNA ligase |
| 17(1) | WP_306007798.1 | WP_306007798.1 isoleucine--tRNA ligase |
| 17(1) | WP_147786241.1 | WP_147786241.1 isoleucine--tRNA ligase |
| 17(6) | WP_000754931.1 | WP_000754931.1 isoleucine--tRNA ligase |
| 17(7) | WP_000754882.1 | WP_000754882.1 isoleucine--tRNA ligase |
| 17(2) | WP_065228765.1 | WP_065228765.1 isoleucine--tRNA ligase |
| 17(1) | WP_098288679.1 | WP_098288679.1 isoleucine--tRNA ligase |
| 17(1) | WP_242251760.1 | WP_242251760.1 isoleucine--tRNA ligase |
| 17(1) | WP_243500441.1 | WP_243500441.1 isoleucine--tRNA ligase |
| 17(2) | WP_204666325.1 | WP_204666325.1 methionine--tRNA ligase |
| 17(3) | WP_097984058.1 | WP_097984058.1 isoleucine--tRNA ligase |
| 17(1) | WP_000754907.1 | WP_000754907.1 isoleucine--tRNA ligase |
| 17(1) | WP_000754909.1 | WP_000754909.1 isoleucine--tRNA ligase |
| 17(1) | WP_087982599.1 | WP_087982599.1 isoleucine--tRNA ligase |
| 17(1) | WP_101195710.1 | WP_101195710.1 isoleucine--tRNA ligase |
| 17(1) | WP_098776437.1 | WP_098776437.1 isoleucine--tRNA ligase |
| 17(1) | WP_149190448.1 | WP_149190448.1 isoleucine--tRNA ligase |
| 17(2) | WP_098508771.1 | WP_098508771.1 isoleucine--tRNA ligase |
| 17(1) | WP_236804416.1 | WP_236804416.1 isoleucine--tRNA ligase |
| 17(2) | WP_098255410.1 | WP_098255410.1 isoleucine--tRNA ligase |
| 17(1) | WP_275175512.1 | WP_275175512.1 isoleucine--tRNA ligase |
| 17(1) | WP_073542129.1 | WP_073542129.1 isoleucine--tRNA ligase |
| 17(1) | WP_095691016.1 | WP_095691016.1 isoleucine--tRNA ligase |
| 17(1) | WP_044792568.1 | WP_044792568.1 isoleucine--tRNA ligase |
| 17(1) | WP_098558851.1 | WP_098558851.1 isoleucine--tRNA ligase |
| 17(1) | WP_159146500.1 | WP_159146500.1 isoleucine--tRNA ligase |
| 17(1) | WP_098296562.1 | WP_098296562.1 isoleucine--tRNA ligase |
| 17(1) | WP_000754912.1 | WP_000754912.1 isoleucine--tRNA ligase |
| 17(1) | WP_098204198.1 | WP_098204198.1 isoleucine--tRNA ligase |
| 17(1) | WP_088298126.1 | WP_088298126.1 isoleucine--tRNA ligase |
| 17(2) | WP_098266642.1 | WP_098266642.1 isoleucine--tRNA ligase |
| 17(1) | WP_002148348.1 | WP_002148348.1 isoleucine--tRNA ligase |
| 17(2) | WP_098148052.1 | WP_098148052.1 isoleucine--tRNA ligase |
| 17(1) | WP_147807660.1 | WP_147807660.1 isoleucine--tRNA ligase |
| 17(1) | WP_090688937.1 | WP_090688937.1 isoleucine--tRNA ligase |
| 17(1) | WP_144611181.1 | WP_144611181.1 isoleucine--tRNA ligase |
| 17(1) | WP_071724710.1 | WP_071724710.1 isoleucine--tRNA ligase |
| 17(1) | WP_074570406.1 | WP_074570406.1 isoleucine--tRNA ligase |
| 17(1) | WP_087995249.1 | WP_087995249.1 isoleucine--tRNA ligase |
| 17(1) | WP_227219180.1 | WP_227219180.1 methionine--tRNA ligase |
| 17(4) | WP_103626227.1 | WP_103626227.1 isoleucine--tRNA ligase |
| 17(1) | WP_000754915.1 | WP_000754915.1 isoleucine--tRNA ligase |
| 17(1) | WP_098638817.1 | WP_098638817.1 isoleucine--tRNA ligase |
| 17(1) | WP_139867390.1 | WP_139867390.1 isoleucine--tRNA ligase |
| 17(1) | WP_242261450.1 | WP_242261450.1 isoleucine--tRNA ligase |
| 17(1) | WP_098201422.1 | WP_098201422.1 isoleucine--tRNA ligase |
| 17(1) | WP_222661107.1 | WP_222661107.1 isoleucine--tRNA ligase |
| 17(3) | WP_000754884.1 | WP_000754884.1 isoleucine--tRNA ligase |
| 17(1) | WP_029438084.1 | WP_029438084.1 isoleucine--tRNA ligase |
| 17(1) | WP_307878870.1 | WP_307878870.1 isoleucine--tRNA ligase |

| | | |
|---|---|---|
| 18(5) | WP_087982595.1 | WP_087982595.1 hypothetical protein |
| 18(1) | WP_144655913.1 | WP_144655913.1 hypothetical protein |
| 18(1) | WP_236804420.1 | WP_236804420.1 hypothetical protein |
| 18(1) | WP_208019345.1 | WP_208019345.1 hypothetical protein |
| 18(1) | WP_106382854.1 | WP_106382854.1 hypothetical protein |
| 18(1) | WP_000709317.1 | WP_000709317.1 hypothetical protein |

| | | |
|---|---|---|
| 18(1) | WP_149238901.1 | WP_149238901.1 hypothetical protein |
| 18(2) | WP_000709319.1 | WP_000709319.1 hypothetical protein |
| 18(1) | WP_000751334.1 | WP_000751334.1 hypothetical protein |
| 18(1) | WP_002141587.1 | WP_002141587.1 hypothetical protein |
| 18(1) | WP_284566224.1 | WP_284566224.1 hypothetical protein |
| 18(1) | WP_098638815.1 | WP_098638815.1 hypothetical protein |
| 18(8) | WP_001167915.1 | WP_001167915.1 hypothetical protein |
| 18(1) | WP_242251768.1 | WP_242251768.1 hypothetical protein |
| 18(1) | WP_070805900.1 | WP_070805900.1 hypothetical protein |
| 18(1) | WP_097906563.1 | WP_097906563.1 hypothetical protein |
| 18(2) | WP_088609112.1 | WP_088609112.1 hypothetical protein |
| 18(1) | WP_078181900.1 | WP_078181900.1 hypothetical protein |
| 18(2) | WP_000709321.1 | WP_000709321.1 hypothetical protein |
| 18(1) | WP_215601063.1 | WP_215601063.1 hypothetical protein |
| 18(11) | WP_000709318.1 | WP_000709318.1 hypothetical protein |
| 18(1) | WP_215591199.1 | WP_215591199.1 hypothetical protein |
| 18(1) | WP_137052042.1 | WP_137052042.1 hypothetical protein |
| 18(1) | WP_277504684.1 | WP_277504684.1 hypothetical protein |
| 18(1) | WP_306007799.1 | WP_306007799.1 hypothetical protein |
| 18(1) | WP_061531032.1 | WP_061531032.1 hypothetical protein |
| 18(1) | WP_270723905.1 | WP_270723905.1 hypothetical protein |
| 18(4) | WP_074601317.1 | WP_074601317.1 hypothetical protein |
| 18(1) | WP_098519064.1 | WP_098519064.1 hypothetical protein |
| 18(1) | WP_098991575.1 | WP_098991575.1 hypothetical protein |
| 18(1) | WP_098776433.1 | WP_098776433.1 hypothetical protein |
| 18(1) | WP_243500445.1 | WP_243500445.1 hypothetical protein |
| 18(1) | WP_097810738.1 | WP_097810738.1 hypothetical protein |
| 18(1) | WP_017149776.1 | WP_017149776.1 hypothetical protein |
| 18(1) | WP_207994166.1 | WP_207994166.1 hypothetical protein |
| 18(1) | WP_098229136.1 | WP_098229136.1 hypothetical protein |
| 18(1) | WP_159146501.1 | WP_159146501.1 hypothetical protein |
| 18(1) | WP_144506825.1 | WP_144506825.1 hypothetical protein |
| 18(2) | WP_098204200.1 | WP_098204200.1 hypothetical protein |
| 18(2) | WP_103655006.1 | WP_103655006.1 hypothetical protein |
| 18(5) | WP_103626228.1 | WP_103626228.1 hypothetical protein |
| 18(1) | WP_048552299.1 | WP_048552299.1 hypothetical protein |
| 18(1) | WP_088032774.1 | WP_088032774.1 hypothetical protein |
| 18(1) | WP_215572750.1 | WP_215572750.1 hypothetical protein |
| 18(1) | WP_144532287.1 | WP_144532287.1 hypothetical protein |
| 18(8) | WP_001167903.1 | WP_001167903.1 hypothetical protein |
| 18(11) | WP_001167914.1 | WP_001167914.1 hypothetical protein |
| 18(1) | WP_088023232.1 | WP_088023232.1 hypothetical protein |
| 18(1) | WP_098303181.1 | WP_098303181.1 hypothetical protein |
| 18(1) | WP_090688944.1 | WP_090688944.1 hypothetical protein |
| 18(1) | WP_029438087.1 | WP_029438087.1 hypothetical protein |
| 18(2) | WP_064473942.1 | WP_064473942.1 hypothetical protein |
| 18(1) | WP_000751333.1 | WP_000751333.1 hypothetical protein |
| 18(3) | WP_086419832.1 | WP_086419832.1 hypothetical protein |
| 18(1) | WP_088058529.1 | WP_088058529.1 hypothetical protein |
| 18(2) | WP_200457880.1 | WP_200457880.1 hypothetical protein |
| 18(1) | WP_098254027.1 | WP_098254027.1 hypothetical protein |
| 18(1) | WP_002158050.1 | WP_002158050.1 hypothetical protein |
| 18(1) | WP_144611183.1 | WP_144611183.1 hypothetical protein |
| 18(1) | WP_166704986.1 | WP_166704986.1 hypothetical protein |
| 18(1) | WP_098830137.1 | WP_098830137.1 hypothetical protein |
| 18(1) | WP_097911845.1 | WP_097911845.1 hypothetical protein |
| 18(1) | WP_071736270.1 | WP_071736270.1 hypothetical protein |
| 18(1) | WP_061680953.1 | WP_061680953.1 hypothetical protein |
| 18(2) | WP_098447523.1 | WP_098447523.1 hypothetical protein |
| 18(6) | WP_053564635.1 | WP_053564635.1 hypothetical protein |
| 18(5) | WP_000709322.1 | WP_000709322.1 hypothetical protein |
| 18(1) | WP_002148342.1 | WP_002148342.1 hypothetical protein |
| 18(1) | WP_075666789.1 | WP_075666789.1 hypothetical protein |
| 18(1) | WP_063223272.1 | WP_063223272.1 hypothetical protein |
| 18(1) | WP_286017545.1 | WP_286017545.1 hypothetical protein |
| 18(1) | WP_289398693.1 | WP_289398693.1 hypothetical protein |
| 18(1) | WP_097821801.1 | WP_097821801.1 hypothetical protein |
| 18(2) | WP_098508776.1 | WP_098508776.1 hypothetical protein |
| 18(2) | WP_002031631.1 | WP_002031631.1 hypothetical protein |
| 18(1) | WP_097987493.1 | WP_097987493.1 hypothetical protein |
| 18(1) | WP_270728508.1 | WP_270728508.1 hypothetical protein |
| 18(3) | WP_078205522.1 | WP_078205522.1 hypothetical protein |
| 18(1) | WP_208752244.1 | WP_208752244.1 hypothetical protein |
| 18(2) | WP_098255408.1 | WP_098255408.1 hypothetical protein |
| 18(1) | WP_000709327.1 | WP_000709327.1 hypothetical protein |
| 18(2) | WP_113770806.1 | WP_113770806.1 hypothetical protein |
| 18(1) | WP_098952615.1 | WP_098952615.1 hypothetical protein |
| 18(4) | WP_098264281.1 | WP_098264281.1 hypothetical protein |
| 18(3) | WP_060630643.1 | WP_060630643.1 hypothetical protein |
| 18(7) | WP_000709320.1 | WP_000709320.1 hypothetical protein |
| 18(2) | WP_087956288.1 | WP_087956288.1 hypothetical protein |
| 18(1) | WP_097905184.1 | WP_097905184.1 hypothetical protein |
| 18(1) | WP_063219641.1 | WP_063219641.1 hypothetical protein |
| 18(1) | WP_098115927.1 | WP_098115927.1 hypothetical protein |
| 18(1) | WP_087995248.1 | WP_087995248.1 hypothetical protein |
| 18(5) | WP_001174424.1 | WP_001174424.1 hypothetical protein |
| 18(1) | WP_131231111.1 | WP_131231111.1 hypothetical protein |
| 18(1) | WP_088308504.1 | WP_088308504.1 hypothetical protein |
| 18(1) | WP_151630610.1 | WP_151630610.1 hypothetical protein |
| 18(2) | WP_098148053.1 | WP_098148053.1 hypothetical protein |
| 18(31) | WP_001167912.1 | WP_001167912.1 hypothetical protein |
| 18(1) | WP_098199509.1 | WP_098199509.1 hypothetical protein |
| 18(1) | WP_025149358.1 | WP_025149358.1 hypothetical protein |
| 18(3) | WP_137013292.1 | WP_137013292.1 hypothetical protein |
| 18(1) | WP_098556215.1 | WP_098556215.1 hypothetical protein |
| 18(1) | WP_242400020.1 | WP_242400020.1 hypothetical protein |
| 18(3) | WP_098266644.1 | WP_098266644.1 hypothetical protein |
| 18(1) | WP_000709316.1 | WP_000709316.1 hypothetical protein |
| 18(1) | WP_212956542.1 | WP_212956542.1 hypothetical protein |
| 19(1) | WP_088550039.1 | WP_088550039.1 hypothetical protein |
| 19(1) | WP_217035855.1 | WP_217035855.1 hypothetical protein |
| 19(1) | WP_046522785.1 | WP_046522785.1 hypothetical protein |
| 19(1) | WP_210364462.1 | WP_210364462.1 hypothetical protein |

| | | |
|---|---|---|
| 19(1) | WP_090674371.1 | WP_090674371.1 hypothetical protein |
| 19(1) | WP_068673707.1 | WP_068673707.1 hypothetical protein |
| 19(1) | WP_156103297.1 | WP_156103297.1 hypothetical protein |
| 19(1) | WP_156520718.1 | WP_156520718.1 hypothetical protein |
| 19(1) | WP_197937398.1 | WP_197937398.1 hypothetical protein |
| 19(1) | WP_208926614.1 | WP_208926614.1 hypothetical protein |
| 19(1) | WP_010496608.1 | WP_010496608.1 hypothetical protein |
| 19(1) | WP_306049021.1 | WP_306049021.1 hypothetical protein |
| 19(1) | WP_188535095.1 | WP_188535095.1 hypothetical protein |
| 19(1) | WP_309098611.1 | WP_309098611.1 hypothetical protein |
| 19(1) | WP_048826579.1 | WP_048826579.1 hypothetical protein |
| 19(1) | WP_308082245.1 | WP_308082245.1 hypothetical protein |
| 19(1) | WP_066022450.1 | WP_066022450.1 hypothetical protein |
| 19(1) | WP_226642876.1 | WP_226642876.1 hypothetical protein |
| 19(1) | WP_119601482.1 | WP_119601482.1 hypothetical protein |
| 19(1) | WP_251245805.1 | WP_251245805.1 hypothetical protein |
| 19(1) | WP_132008157.1 | WP_132008157.1 hypothetical protein |
| 19(2) | WP_213592857.1 | WP_213592857.1 hypothetical protein |
| 19(4) | WP_076103994.1 | WP_076103994.1 hypothetical protein |
| 19(2) | WP_156103296.1 | WP_156103296.1 hypothetical protein |
| 19(1) | WP_156520719.1 | WP_156520719.1 hypothetical protein |
| 19(1) | WP_308083988.1 | WP_308083988.1 hypothetical protein |
| 19(1) | WP_307589610.1 | WP_307589610.1 hypothetical protein |
| 19(1) | WP_251245813.1 | WP_251245813.1 hypothetical protein |
| 19(1) | WP_249897320.1 | WP_249897320.1 hypothetical protein |
| 19(1) | WP_157076702.1 | WP_157076702.1 hypothetical protein |
| 19(1) | WP_307312550.1 | WP_307312550.1 hypothetical protein |
| 19(1) | WP_268613674.1 | WP_268613674.1 hypothetical protein |
| 19(1) | WP_269920256.1 | WP_269920256.1 hypothetical protein, partial |
| 19(1) | WP_186580067.1 | WP_186580067.1 hypothetical protein |
| 19(1) | WP_066022459.1 | WP_066022459.1 hypothetical protein |
| 19(1) | WP_254366007.1 | WP_254366007.1 hypothetical protein |
| 19(1) | WP_216791214.1 | WP_216791214.1 hypothetical protein |
| 19(1) | WP_146022700.1 | WP_146022700.1 hypothetical protein |
| 19(1) | WP_258207831.1 | WP_258207831.1 hypothetical protein |
| 19(1) | WP_099361336.1 | WP_099361336.1 hypothetical protein |
| 19(1) | WP_089965940.1 | WP_089965940.1 hypothetical protein |
| 19(1) | WP_213143946.1 | WP_213143946.1 hypothetical protein |
| 19(1) | WP_251245809.1 | WP_251245809.1 hypothetical protein |
| 19(1) | WP_013490130.1 | WP_013490130.1 hypothetical protein |
| 19(1) | WP_221875371.1 | WP_221875371.1 hypothetical protein |
| 19(1) | WP_066022460.1 | WP_066022460.1 hypothetical protein |
| 19(1) | WP_226602917.1 | WP_226602917.1 hypothetical protein |
| 19(1) | WP_195571377.1 | WP_195571377.1 hypothetical protein |
| 19(1) | WP_150457757.1 | WP_150457757.1 hypothetical protein |
| 19(1) | WP_251542608.1 | WP_251542608.1 hypothetical protein |
| 19(1) | WP_302878978.1 | WP_302878978.1 hypothetical protein |
| 19(1) | WP_063188064.1 | WP_063188064.1 hypothetical protein |
| 19(1) | WP_169028889.1 | WP_169028889.1 hypothetical protein |
| 19(1) | WP_251245810.1 | WP_251245810.1 hypothetical protein |
| 19(1) | WP_066022458.1 | WP_066022458.1 hypothetical protein |
| 19(1) | WP_090761819.1 | WP_090761819.1 hypothetical protein |
| 19(1) | WP_102264550.1 | WP_102264550.1 hypothetical protein |
| 19(1) | WP_251245803.1 | WP_251245803.1 hypothetical protein |
| 19(1) | WP_186580072.1 | WP_186580072.1 hypothetical protein |
| 19(1) | WP_034773542.1 | WP_034773542.1 hypothetical protein |
| 19(1) | WP_182301832.1 | WP_182301832.1 hypothetical protein |
| 19(1) | WP_210364452.1 | WP_210364452.1 hypothetical protein |
| 19(1) | WP_179034792.1 | WP_179034792.1 hypothetical protein |
| 19(1) | WP_066022449.1 | WP_066022449.1 hypothetical protein |
| 19(1) | WP_088830538.1 | WP_088830538.1 hypothetical protein |
| 19(1) | WP_283910928.1 | WP_283910928.1 hypothetical protein |
| 19(1) | WP_036689406.1 | WP_036689406.1 hypothetical protein |
| 19(2) | WP_144479063.1 | WP_144479063.1 hypothetical protein |
| 19(1) | WP_033100020.1 | WP_033100020.1 hypothetical protein |
| 19(1) | WP_251548761.1 | WP_251548761.1 hypothetical protein |
| 19(1) | WP_053361023.1 | WP_053361023.1 hypothetical protein |
| 19(1) | WP_122971534.1 | WP_122971534.1 hypothetical protein |
| 19(1) | WP_085499089.1 | WP_085499089.1 hypothetical protein |
| 19(1) | WP_045518283.1 | WP_045518283.1 hypothetical protein |
| 19(1) | WP_077619647.1 | WP_077619647.1 hypothetical protein |
| 19(1) | WP_304216870.1 | WP_304216870.1 hypothetical protein |
| 19(1) | WP_186580075.1 | WP_186580075.1 hypothetical protein |
| 19(1) | WP_235286202.1 | WP_235286202.1 hypothetical protein |
| 19(1) | WP_269920255.1 | WP_269920255.1 hypothetical protein |
| 19(1) | WP_253411073.1 | WP_253411073.1 hypothetical protein |
| 19(1) | WP_152681108.1 | WP_152681108.1 hypothetical protein |
| 19(1) | WP_263324548.1 | WP_263324548.1 hypothetical protein |
| 19(1) | WP_239617473.1 | WP_239617473.1 hypothetical protein |
| 19(1) | WP_150457761.1 | WP_150457761.1 hypothetical protein |
| 19(1) | WP_198056460.1 | WP_198056460.1 hypothetical protein |
| 19(1) | WP_251638773.1 | WP_251638773.1 hypothetical protein |
| 19(1) | WP_127459429.1 | WP_127459429.1 hypothetical protein |
| 19(1) | WP_254759297.1 | WP_254759297.1 hypothetical protein |
| 19(1) | WP_278740190.1 | WP_278740190.1 hypothetical protein |
| 19(1) | WP_034773545.1 | WP_034773545.1 hypothetical protein |
| 19(1) | WP_251245806.1 | WP_251245806.1 hypothetical protein |
| 19(1) | WP_163853475.1 | WP_163853475.1 hypothetical protein |
| 19(1) | WP_121610595.1 | WP_121610595.1 hypothetical protein |
| 19(2) | WP_038569547.1 | WP_038569547.1 hypothetical protein |
| 19(3) | WP_143812142.1 | WP_143812142.1 hypothetical protein |
| 19(1) | WP_013490129.1 | WP_013490129.1 hypothetical protein |
| 19(1) | WP_236334322.1 | WP_236334322.1 hypothetical protein |
| 19(1) | WP_076119217.1 | WP_076119217.1 hypothetical protein |
| 19(1) | WP_148500004.1 | WP_148500004.1 hypothetical protein |
| 19(1) | WP_224839862.1 | WP_224839862.1 hypothetical protein |
| 19(1) | WP_188535090.1 | WP_188535090.1 hypothetical protein |
| 19(1) | WP_198135581.1 | WP_198135581.1 hypothetical protein |
| 19(1) | WP_048826581.1 | WP_048826581.1 hypothetical protein |
| 19(1) | WP_306049023.1 | WP_306049023.1 hypothetical protein |
| 19(1) | WP_112331577.1 | WP_112331577.1 hypothetical protein |
| 19(1) | WP_307284682.1 | WP_307284682.1 hypothetical protein |
| 19(1) | WP_215032125.1 | WP_215032125.1 hypothetical protein |
| 19(1) | WP_223592680.1 | WP_223592680.1 hypothetical protein |

| | | |
|---|---|---|
| 19(1) | WP_199419767.1 | WP_199419767.1 hypothetical protein |
| 19(1) | WP_226086174.1 | WP_226086174.1 hypothetical protein |
| 19(1) | WP_155609750.1 | WP_155609750.1 hypothetical protein |
| 19(1) | WP_036684482.1 | WP_036684482.1 hypothetical protein |
| 19(1) | WP_125084776.1 | WP_125084776.1 hypothetical protein |
| 19(1) | WP_270065060.1 | WP_270065060.1 hypothetical protein |
| 19(1) | WP_066092003.1 | WP_066092003.1 hypothetical protein |
| 19(1) | WP_283952948.1 | WP_283952948.1 hypothetical protein |
| 19(1) | WP_186580065.1 | WP_186580065.1 hypothetical protein |
| 19(1) | WP_179603480.1 | WP_179603480.1 hypothetical protein |
| 19(1) | WP_110934068.1 | WP_110934068.1 hypothetical protein |
| 19(1) | WP_251245811.1 | WP_251245811.1 hypothetical protein |
| 19(1) | WP_284036498.1 | WP_284036498.1 hypothetical protein |
| 19(1) | WP_199419769.1 | WP_199419769.1 hypothetical protein |
| 19(1) | WP_191272864.1 | WP_191272864.1 hypothetical protein |
| 19(2) | WP_034773555.1 | WP_034773555.1 hypothetical protein |
| 19(1) | WP_116353252.1 | WP_116353252.1 hypothetical protein |
| 19(1) | WP_148362806.1 | WP_148362806.1 hypothetical protein |
| 19(1) | WP_122971536.1 | WP_122971536.1 hypothetical protein |
| 19(3) | WP_076281072.1 | WP_076281072.1 hypothetical protein |
| 19(1) | WP_251557378.1 | WP_251557378.1 hypothetical protein |
| 19(1) | WP_186580069.1 | WP_186580069.1 hypothetical protein |
| 19(1) | WP_278327423.1 | WP_278327423.1 hypothetical protein |
| 19(1) | WP_199925767.1 | WP_199925767.1 hypothetical protein |
| 19(1) | WP_139179441.1 | WP_139179441.1 hypothetical protein |
| 19(1) | WP_137698108.1 | WP_137698108.1 hypothetical protein |
| 19(1) | WP_213117481.1 | WP_213117481.1 hypothetical protein |
| 19(1) | WP_304216867.1 | WP_304216867.1 hypothetical protein |
| 19(1) | WP_221875376.1 | WP_221875376.1 hypothetical protein |
| 19(1) | WP_223068049.1 | WP_223068049.1 hypothetical protein |
| 19(1) | WP_226679103.1 | WP_226679103.1 hypothetical protein |
| 19(1) | WP_025848608.1 | WP_025848608.1 hypothetical protein |
| 19(1) | WP_232243182.1 | WP_232243182.1 hypothetical protein |
| 19(1) | WP_251480996.1 | WP_251480996.1 hypothetical protein |
| 19(1) | WP_204412724.1 | WP_204412724.1 hypothetical protein |
| 19(1) | WP_289342262.1 | WP_289342262.1 hypothetical protein |
| 19(1) | WP_154663560.1 | WP_154663560.1 hypothetical protein |
| 19(1) | WP_034773537.1 | WP_034773537.1 hypothetical protein |
| 19(1) | WP_281937374.1 | WP_281937374.1 hypothetical protein |
| 19(1) | WP_054972092.1 | WP_054972092.1 hypothetical protein |
| 19(1) | WP_099354907.1 | WP_099354907.1 hypothetical protein |
| 19(1) | WP_010676864.1 | WP_010676864.1 hypothetical protein |
| 19(1) | WP_077619649.1 | WP_077619649.1 hypothetical protein |
| 19(1) | WP_034773547.1 | WP_034773547.1 hypothetical protein |
| 19(1) | WP_214706003.1 | WP_214706003.1 hypothetical protein |
| 19(1) | WP_251242448.1 | WP_251242448.1 hypothetical protein |
| 19(1) | WP_066022448.1 | WP_066022448.1 hypothetical protein |
| 19(1) | WP_111147801.1 | WP_111147801.1 hypothetical protein |
| 19(1) | WP_195571376.1 | WP_195571376.1 hypothetical protein |
| 19(1) | WP_079506394.1 | WP_079506394.1 hypothetical protein |
| 19(1) | WP_251426366.1 | WP_251426366.1 hypothetical protein |
| 19(1) | WP_306077389.1 | WP_306077389.1 hypothetical protein |

| | | |
|---|---|---|
| 19(1) | WP_191272865.1 | WP_191272865.1 hypothetical protein |
| 19(1) | WP_231094849.1 | WP_231094849.1 hypothetical protein |
| 19(1) | WP_099354909.1 | WP_099354909.1 hypothetical protein |
| 19(1) | WP_222599449.1 | WP_222599449.1 hypothetical protein |
| 19(1) | WP_156114752.1 | WP_156114752.1 hypothetical protein |
| 19(1) | WP_251245812.1 | WP_251245812.1 hypothetical protein |
| 19(1) | WP_270065059.1 | WP_270065059.1 hypothetical protein |
| 19(1) | WP_278740192.1 | WP_278740192.1 hypothetical protein |
| 19(1) | WP_270065061.1 | WP_270065061.1 hypothetical protein |
| 19(1) | WP_204445211.1 | WP_204445211.1 hypothetical protein |
| 19(1) | WP_149871683.1 | WP_149871683.1 hypothetical protein |
| 19(1) | WP_066022447.1 | WP_066022447.1 hypothetical protein |
| 19(1) | WP_108534957.1 | WP_108534957.1 hypothetical protein |
| 19(1) | WP_033100019.1 | WP_033100019.1 hypothetical protein |
| 19(1) | WP_195515236.1 | WP_195515236.1 hypothetical protein |
| 19(1) | WP_134750604.1 | WP_134750604.1 hypothetical protein |
| 19(1) | WP_066022457.1 | WP_066022457.1 hypothetical protein |
| 19(1) | WP_144061519.1 | WP_144061519.1 hypothetical protein |
| 19(1) | WP_164515190.1 | WP_164515190.1 hypothetical protein |
| 19(1) | WP_251598144.1 | WP_251598144.1 hypothetical protein |
| 19(1) | WP_195571383.1 | WP_195571383.1 hypothetical protein |
| 19(1) | WP_257443630.1 | WP_257443630.1 hypothetical protein |
| 19(1) | WP_034773539.1 | WP_034773539.1 hypothetical protein |
| 19(1) | WP_251245808.1 | WP_251245808.1 hypothetical protein |
| 19(1) | WP_007083289.1 | WP_007083289.1 hypothetical protein |
| 19(1) | WP_198056461.1 | WP_198056461.1 hypothetical protein |
| 19(1) | WP_179160621.1 | WP_179160621.1 hypothetical protein |
| 19(1) | WP_198059702.1 | WP_198059702.1 hypothetical protein |
| 19(1) | WP_306049025.1 | WP_306049025.1 hypothetical protein |
| 19(1) | WP_136379699.1 | WP_136379699.1 hypothetical protein |
| 19(1) | WP_156520717.1 | WP_156520717.1 hypothetical protein |
| 19(1) | WP_041964578.1 | WP_041964578.1 hypothetical protein |
| 19(1) | WP_066419983.1 | WP_066419983.1 hypothetical protein |
| 19(1) | WP_309098613.1 | WP_309098613.1 hypothetical protein |
| 19(1) | WP_229139919.1 | WP_229139919.1 hypothetical protein |
| 19(1) | WP_199419766.1 | WP_199419766.1 hypothetical protein |
| 19(1) | WP_076101573.1 | WP_076101573.1 hypothetical protein |
| 19(1) | WP_251241782.1 | WP_251241782.1 hypothetical protein |
| 19(1) | WP_041808389.1 | WP_041808389.1 hypothetical protein |
| 19(1) | WP_165097626.1 | WP_165097626.1 hypothetical protein |
| 19(1) | WP_156103295.1 | WP_156103295.1 hypothetical protein |
| 19(1) | WP_034773549.1 | WP_034773549.1 hypothetical protein |
| 19(1) | WP_127484349.1 | WP_127484349.1 hypothetical protein |
| 20(1) | WP_185897352.1 | WP_185897352.1 hypothetical protein |
| 20(2) | WP_167301694.1 | WP_167301694.1 hypothetical protein |
| 20(1) | WP_179087805.1 | WP_179087805.1 hypothetical protein |
| 20(1) | WP_181014173.1 | WP_181014173.1 hypothetical protein |
| 20(1) | WP_251583281.1 | WP_251583281.1 hypothetical protein |
| 20(1) | WP_293202265.1 | WP_293202265.1 hypothetical protein |
| 20(2) | WP_255223059.1 | WP_255223059.1 hypothetical protein |
| 20(1) | WP_167666085.1 | WP_167666085.1 hypothetical protein |

| | | |
|---|---|---|
| 20(1) | WP_251378792.1 | WP_251378792.1 hypothetical protein |
| 20(1) | WP_191798523.1 | WP_191798523.1 hypothetical protein |
| 20(2) | WP_167527256.1 | WP_167527256.1 hypothetical protein |
| 20(3) | WP_167350850.1 | WP_167350850.1 hypothetical protein |
| 20(1) | WP_307214305.1 | WP_307214305.1 hypothetical protein |
| 20(3) | WP_007128633.1 | WP_007128633.1 hypothetical protein |
| 20(2) | WP_177185675.1 | WP_177185675.1 hypothetical protein |
| 20(1) | WP_169823406.1 | WP_169823406.1 hypothetical protein |
| 20(1) | WP_173405222.1 | WP_173405222.1 hypothetical protein |
| 20(2) | WP_265589731.1 | WP_265589731.1 hypothetical protein |
| 20(1) | WP_166653172.1 | WP_166653172.1 hypothetical protein |
| 20(1) | WP_170948036.1 | WP_170948036.1 hypothetical protein |
| 20(1) | WP_199794793.1 | WP_199794793.1 hypothetical protein |
| 20(1) | WP_281281438.1 | WP_281281438.1 hypothetical protein |
| 20(1) | WP_167518739.1 | WP_167518739.1 hypothetical protein |
| 20(4) | WP_179132691.1 | WP_179132691.1 hypothetical protein |
| 20(1) | WP_174704634.1 | WP_174704634.1 hypothetical protein |
| 20(1) | WP_133384357.1 | WP_133384357.1 hypothetical protein |
| 20(3) | WP_167375738.1 | WP_167375738.1 hypothetical protein |
| 20(1) | WP_216513487.1 | WP_216513487.1 hypothetical protein |
| 20(1) | WP_197029854.1 | WP_197029854.1 hypothetical protein |
| 20(2) | WP_283655690.1 | WP_283655690.1 hypothetical protein |
| 20(1) | WP_015736245.1 | WP_015736245.1 hypothetical protein |
| 20(2) | WP_018883278.1 | WP_018883278.1 hypothetical protein |
| 20(45) | WP_023988262.1 | WP_023988262.1 hypothetical protein |
| 20(1) | WP_017689076.1 | WP_017689076.1 hypothetical protein |
| 20(16) | WP_016821028.1 | WP_016821028.1 hypothetical protein |
| 20(1) | WP_186327703.1 | WP_186327703.1 hypothetical protein |
| 20(1) | WP_165593396.1 | WP_165593396.1 hypothetical protein |
| 20(1) | WP_186333579.1 | WP_186333579.1 hypothetical protein |
| 20(1) | WP_213532654.1 | WP_213532654.1 hypothetical protein |
| 20(1) | WP_282321824.1 | WP_282321824.1 hypothetical protein |
| 20(2) | WP_187386141.1 | WP_187386141.1 hypothetical protein |
| 20(1) | WP_192266446.1 | WP_192266446.1 hypothetical protein |
| 20(1) | WP_170970831.1 | WP_170970831.1 hypothetical protein |
| 20(2) | WP_193745964.1 | WP_193745964.1 hypothetical protein |
| 20(1) | WP_272035301.1 | WP_272035301.1 hypothetical protein |
| 20(2) | WP_167433928.1 | WP_167433928.1 hypothetical protein |
| 20(1) | WP_167372839.1 | WP_167372839.1 hypothetical protein |
| 20(2) | WP_175415284.1 | WP_175415284.1 hypothetical protein |
| 20(14) | WP_164783094.1 | WP_164783094.1 hypothetical protein |
| 20(1) | WP_013370781.1 | WP_013370781.1 hypothetical protein |
| 20(1) | WP_169505048.1 | WP_169505048.1 hypothetical protein |
| 20(1) | WP_199909991.1 | WP_199909991.1 hypothetical protein |
| 20(1) | WP_281177661.1 | WP_281177661.1 hypothetical protein |
| 20(2) | WP_167434743.1 | WP_167434743.1 hypothetical protein |
| 20(1) | WP_162834829.1 | WP_162834829.1 hypothetical protein |
| 20(2) | WP_186380865.1 | WP_186380865.1 hypothetical protein |
| 20(1) | WP_225999405.1 | WP_225999405.1 hypothetical protein |
| 20(2) | WP_181429970.1 | WP_181429970.1 hypothetical protein |
| 20(12) | WP_163872756.1 | WP_163872756.1 hypothetical protein |
| 20(1) | WP_170862375.1 | WP_170862375.1 hypothetical protein |
| 20(1) | WP_247902242.1 | WP_247902242.1 hypothetical protein |
| 20(2) | WP_164848677.1 | WP_164848677.1 hypothetical protein |
| 20(9) | WP_170867771.1 | WP_170867771.1 hypothetical protein |
| 20(1) | WP_254325794.1 | WP_254325794.1 hypothetical protein |
| 20(1) | WP_211745835.1 | WP_211745835.1 hypothetical protein |
| 20(1) | WP_264929210.1 | WP_264929210.1 hypothetical protein |
| 20(1) | WP_285748170.1 | WP_285748170.1 hypothetical protein |
| 20(1) | WP_236417205.1 | WP_236417205.1 hypothetical protein |
| 20(2) | WP_187644933.1 | WP_187644933.1 hypothetical protein |
| 20(1) | WP_162833918.1 | WP_162833918.1 hypothetical protein |
| 20(2) | WP_162842451.1 | WP_162842451.1 hypothetical protein |
| 20(1) | WP_167433560.1 | WP_167433560.1 hypothetical protein |
| 20(4) | WP_260871389.1 | WP_260871389.1 hypothetical protein |
| 20(1) | WP_179280988.1 | WP_179280988.1 hypothetical protein |
| 20(2) | WP_177178615.1 | WP_177178615.1 hypothetical protein |
| 20(1) | WP_009590662.1 | WP_009590662.1 hypothetical protein |
| 20(1) | WP_164764155.1 | WP_164764155.1 hypothetical protein |

| | | |
|---|---|---|
| 21(1) | WP_206094028.1 | WP_206094028.1 GNAT family N-acetyltransferase |
| 21(1) | WP_083189145.1 | WP_083189145.1 GNAT family N-acetyltransferase |
| 21(5) | WP_000635650.1 | WP_000635650.1 GNAT family N-acetyltransferase |
| 21(5) | WP_000635649.1 | WP_000635649.1 GNAT family N-acetyltransferase |
| 21(1) | WP_301249695.1 | WP_301249695.1 GNAT family N-acetyltransferase |
| 21(1) | WP_268629203.1 | WP_268629203.1 GNAT family N-acetyltransferase |
| 21(1) | WP_000584237.1 | WP_000584237.1 GNAT family N-acetyltransferase |
| 21(1) | WP_307214303.1 | WP_307214303.1 GNAT family N-acetyltransferase |
| 21(1) | WP_145149075.1 | WP_145149075.1 GNAT family N-acetyltransferase |
| 21(1) | WP_177178616.1 | WP_177178616.1 GNAT family N-acetyltransferase |
| 21(26) | WP_000221048.1 | WP_000221048.1 GNAT family N-acetyltransferase |
| 21(1) | WP_308082239.1 | WP_308082239.1 GNAT family N-acetyltransferase |
| 21(2) | WP_145329902.1 | WP_145329902.1 GNAT family N-acetyltransferase |
| 21(1) | WP_076330718.1 | WP_076330718.1 GNAT family N-acetyltransferase |
| 21(1) | WP_111268944.1 | WP_111268944.1 GNAT family N-acetyltransferase |
| 21(1) | WP_097954757.1 | WP_097954757.1 GNAT family N-acetyltransferase |
| 21(2) | WP_087433615.1 | WP_087433615.1 GNAT family N-acetyltransferase |
| 21(1) | WP_240661945.1 | WP_240661945.1 GNAT family N-acetyltransferase |
| 21(2) | WP_005546528.1 | WP_005546528.1 GNAT family N-acetyltransferase |
| 21(1) | WP_021253108.1 | WP_021253108.1 GNAT family N-acetyltransferase |
| 21(1) | WP_074593264.1 | WP_074593264.1 GNAT family N-acetyltransferase |
| 21(1) | WP_000584239.1 | WP_000584239.1 GNAT family N-acetyltransferase |
| 21(1) | WP_062833876.1 | WP_062833876.1 GNAT family N-acetyltransferase |
| 21(1) | WP_098141777.1 | WP_098141777.1 GNAT family N-acetyltransferase |
| 21(1) | WP_223869144.1 | WP_223869144.1 GNAT family N-acetyltransferase |
| 21(1) | WP_174811616.1 | WP_174811616.1 GNAT family N-acetyltransferase |
| 21(1) | WP_098874962.1 | WP_098874962.1 GNAT family N-acetyltransferase |
| 21(1) | WP_090896174.1 | WP_090896174.1 GNAT family N-acetyltransferase |
| 21(1) | WP_244213647.1 | WP_244213647.1 GNAT family N-acetyltransferase |
| 21(1) | WP_289392068.1 | WP_289392068.1 GNAT family N-acetyltransferase |
| 21(1) | WP_061667212.1 | WP_061667212.1 GNAT family N-acetyltransferase |
| 21(1) | WP_293202267.1 | WP_293202267.1 GNAT family N-acetyltransferase |
| 21(1) | WP_090921067.1 | WP_090921067.1 GNAT family N-acetyltransferase |
| 21(1) | WP_308721990.1 | WP_308721990.1 GNAT family N-acetyltransferase |

| | | |
|---|---|---|
| 21(1) | WP_138186682.1 | WP_138186682.1 GNAT family N-acetyltransferase |
| 21(1) | WP_132131918.1 | WP_132131918.1 GNAT family N-acetyltransferase |
| 21(1) | WP_128101482.1 | WP_128101482.1 GNAT family N-acetyltransferase |
| 21(1) | WP_098139271.1 | WP_098139271.1 GNAT family N-acetyltransferase |
| 21(1) | WP_289355898.1 | WP_289355898.1 GNAT family N-acetyltransferase |
| 21(1) | WP_301879170.1 | WP_301879170.1 hypothetical protein |
| 21(1) | WP_216513485.1 | WP_216513485.1 GNAT family N-acetyltransferase |
| 21(1) | WP_179265141.1 | WP_179265141.1 GNAT family N-acetyltransferase |
| 21(1) | WP_254777283.1 | WP_254777283.1 hypothetical protein |
| 21(1) | WP_024630321.1 | WP_024630321.1 GNAT family N-acetyltransferase |
| 21(1) | WP_211745845.1 | WP_211745845.1 GNAT family N-acetyltransferase |
| 21(1) | WP_264929212.1 | WP_264929212.1 GNAT family N-acetyltransferase |
| 21(1) | WP_251583278.1 | WP_251583278.1 GNAT family N-acetyltransferase |
| 21(1) | WP_247902240.1 | WP_247902240.1 GNAT family N-acetyltransferase |
| 21(1) | WP_036609432.1 | WP_036609432.1 GNAT family N-acetyltransferase |
| 21(1) | WP_142543121.1 | WP_142543121.1 GNAT family N-acetyltransferase |
| 21(3) | WP_000222822.1 | WP_000222822.1 GNAT family N-acetyltransferase |
| 21(1) | WP_272035296.1 | WP_272035296.1 GNAT family N-acetyltransferase |
| 21(1) | WP_197267870.1 | WP_197267870.1 GNAT family N-acetyltransferase |
| 21(1) | WP_205053322.1 | WP_205053322.1 GNAT family N-acetyltransferase |
| 21(1) | WP_000635652.1 | WP_000635652.1 GNAT family N-acetyltransferase |
| 21(1) | WP_091017200.1 | WP_091017200.1 GNAT family N-acetyltransferase |
| 21(1) | WP_110822562.1 | WP_110822562.1 GNAT family N-acetyltransferase |
| 21(5) | WP_000584245.1 | WP_000584245.1 GNAT family N-acetyltransferase |
| 21(1) | WP_086412797.1 | WP_086412797.1 GNAT family N-acetyltransferase |
| 21(3) | WP_090949161.1 | WP_090949161.1 GNAT family N-acetyltransferase |
| 21(1) | WP_192266444.1 | WP_192266444.1 GNAT family N-acetyltransferase |
| 21(1) | WP_095287665.1 | WP_095287665.1 GNAT family N-acetyltransferase |
| 21(2) | WP_063566328.1 | WP_063566328.1 GNAT family N-acetyltransferase |
| 21(1) | WP_076318548.1 | WP_076318548.1 GNAT family N-acetyltransferase |
| 21(1) | WP_285748168.1 | WP_285748168.1 GNAT family N-acetyltransferase |
| 21(1) | WP_017689078.1 | WP_017689078.1 GNAT family N-acetyltransferase |
| 21(2) | WP_162009925.1 | WP_162009925.1 GNAT family N-acetyltransferase |
| 21(1) | WP_100529431.1 | WP_100529431.1 GNAT family N-acetyltransferase |
| 21(1) | WP_076311058.1 | WP_076311058.1 GNAT family N-acetyltransferase |
| 21(1) | WP_021261873.1 | WP_021261873.1 GNAT family N-acetyltransferase |
| 21(1) | WP_060852181.1 | WP_060852181.1 GNAT family N-acetyltransferase |
| 21(1) | WP_072732439.1 | WP_072732439.1 GNAT family N-acetyltransferase |
| 21(1) | WP_070757480.1 | WP_070757480.1 GNAT family N-acetyltransferase |
| 21(1) | WP_098226395.1 | WP_098226395.1 GNAT family N-acetyltransferase |
| 21(1) | WP_056700777.1 | WP_056700777.1 GNAT family N-acetyltransferase |
| 21(1) | WP_091155352.1 | WP_091155352.1 GNAT family N-acetyltransferase |
| 21(1) | WP_064639015.1 | WP_064639015.1 GNAT family N-acetyltransferase |
| 21(1) | WP_006923801.1 | WP_006923801.1 GNAT family N-acetyltransferase |
| 21(1) | WP_188593159.1 | WP_188593159.1 GNAT family N-acetyltransferase |
| 21(1) | WP_053783882.1 | WP_053783882.1 GNAT family N-acetyltransferase |
| 21(1) | WP_111268945.1 | WP_111268945.1 hypothetical protein |
| 21(1) | WP_098284014.1 | WP_098284014.1 GNAT family N-acetyltransferase |
| 21(1) | WP_145408781.1 | WP_145408781.1 GNAT family N-acetyltransferase |
| 21(1) | WP_028533174.1 | WP_028533174.1 GNAT family N-acetyltransferase |
| 21(1) | WP_072727831.1 | WP_072727831.1 GNAT family N-acetyltransferase |
| 21(1) | WP_047909622.1 | WP_047909622.1 GNAT family N-acetyltransferase |
| 21(1) | WP_301879169.1 | WP_301879169.1 GNAT family N-acetyltransferase |
| 21(1) | WP_086388651.1 | WP_086388651.1 GNAT family N-acetyltransferase |
| 21(1) | WP_113052570.1 | WP_113052570.1 GNAT family N-acetyltransferase |
| 21(1) | WP_268619940.1 | WP_268619940.1 GNAT family N-acetyltransferase |
| 21(1) | WP_098139269.1 | WP_098139269.1 GNAT family N-acetyltransferase |
| 21(1) | WP_091031773.1 | WP_091031773.1 GNAT family N-acetyltransferase |
| 21(1) | WP_109999887.1 | WP_109999887.1 GNAT family N-acetyltransferase |
| 21(1) | WP_086392197.1 | WP_086392197.1 GNAT family N-acetyltransferase |
| 21(3) | WP_000570294.1 | WP_000570294.1 GNAT family N-acetyltransferase |
| 21(1) | WP_016108250.1 | WP_016108250.1 GNAT family N-acetyltransferase |
| 21(1) | WP_251480986.1 | WP_251480986.1 GNAT family N-acetyltransferase |
| 21(1) | WP_191798522.1 | WP_191798522.1 GNAT family N-acetyltransferase |
| 21(2) | WP_036670432.1 | WP_036670432.1 GNAT family N-acetyltransferase |
| 21(2) | WP_070179256.1 | WP_070179256.1 GNAT family N-acetyltransferase |
| 21(1) | WP_163061525.1 | WP_163061525.1 GNAT family N-acetyltransferase |
| 21(1) | WP_094936484.1 | WP_094936484.1 GNAT family N-acetyltransferase |
| 21(1) | WP_213532653.1 | WP_213532653.1 GNAT family N-acetyltransferase |
| 21(1) | WP_236417204.1 | WP_236417204.1 GNAT family N-acetyltransferase |
| 21(1) | WP_098612025.1 | WP_098612025.1 GNAT family N-acetyltransferase |
| 21(1) | WP_000221053.1 | WP_000221053.1 GNAT family N-acetyltransferase |
| 21(1) | WP_076208843.1 | WP_076208843.1 GNAT family N-acetyltransferase |
| 21(1) | WP_127536198.1 | WP_127536198.1 GNAT family N-acetyltransferase |
| 21(1) | WP_123067110.1 | WP_123067110.1 GNAT family N-acetyltransferase |
| 21(2) | WP_098055917.1 | WP_098055917.1 GNAT family N-acetyltransferase |
| 21(1) | WP_000221049.1 | WP_000221049.1 GNAT family N-acetyltransferase |
| 21(1) | WP_046392636.1 | WP_046392636.1 GNAT family N-acetyltransferase |
| 21(1) | WP_307413606.1 | WP_307413606.1 GNAT family N-acetyltransferase |
| 21(1) | WP_127543704.1 | WP_127543704.1 GNAT family N-acetyltransferase |
| 21(2) | WP_163977050.1 | WP_163977050.1 GNAT family N-acetyltransferase |
| 21(1) | WP_111619124.1 | WP_111619124.1 GNAT family N-acetyltransferase |
| 21(1) | WP_090805467.1 | WP_090805467.1 GNAT family N-acetyltransferase |
| 21(1) | WP_095358550.1 | WP_095358550.1 GNAT family N-acetyltransferase |
| 21(1) | WP_145318710.1 | WP_145318710.1 GNAT family N-acetyltransferase |
| 21(1) | WP_285079444.1 | WP_285079444.1 GNAT family N-acetyltransferase |
| 21(4) | WP_000222834.1 | WP_000222834.1 GNAT family N-acetyltransferase |
| 21(1) | WP_074094610.1 | WP_074094610.1 GNAT family N-acetyltransferase |
| 21(1) | WP_186380864.1 | WP_186380864.1 GNAT family N-acetyltransferase |
| 21(2) | WP_076249778.1 | WP_076249778.1 GNAT family N-acetyltransferase |
| 21(8) | WP_000584232.1 | WP_000584232.1 GNAT family N-acetyltransferase |
| 21(1) | WP_302848144.1 | WP_302848144.1 GNAT family N-acetyltransferase |
| 21(1) | WP_307543214.1 | WP_307543214.1 GNAT family N-acetyltransferase |
| 21(1) | WP_000584240.1 | WP_000584240.1 GNAT family N-acetyltransferase |
| 21(1) | WP_180287002.1 | WP_180287002.1 GNAT family N-acetyltransferase |
| 21(1) | WP_000221050.1 | WP_000221050.1 GNAT family N-acetyltransferase |
| 21(1) | WP_062326143.1 | WP_062326143.1 GNAT family N-acetyltransferase |
| 21(1) | WP_000221051.1 | WP_000221051.1 GNAT family N-acetyltransferase |
| 21(1) | WP_047842534.1 | WP_047842534.1 GNAT family N-acetyltransferase |
| 21(1) | WP_061676405.1 | WP_061676405.1 GNAT family N-acetyltransferase |
| 21(1) | WP_090993611.1 | WP_090993611.1 GNAT family N-acetyltransferase |
| 22(18) | WP_023988256.1 | WP_023988256.1 hypothetical protein |
| 22(1) | WP_251375557.1 | WP_251375557.1 ADP-heptose synthase |

| | | |
|---|---|---|
| 22(1) | WP_160496945.1 | WP_160496945.1 ADP-heptose synthase |
| 22(1) | WP_213618802.1 | WP_213618802.1 ADP-heptose synthase |
| 22(1) | WP_152381389.1 | WP_152381389.1 ADP-heptose synthase |
| 22(3) | WP_107734210.1 | WP_107734210.1 ADP-heptose synthase |
| 22(1) | WP_124114114.1 | WP_124114114.1 ADP-heptose synthase |
| 22(1) | WP_017689084.1 | WP_017689084.1 hypothetical protein |
| 22(1) | WP_090993603.1 | WP_090993603.1 ADP-heptose synthase |
| 22(1) | WP_108465337.1 | WP_108465337.1 ADP-heptose synthase |
| 22(1) | WP_137062791.1 | WP_137062791.1 ADP-heptose synthase |
| 22(1) | WP_134902677.1 | WP_134902677.1 ADP-heptose synthase |
| 22(1) | WP_209861665.1 | WP_209861665.1 ADP-heptose synthase |
| 22(1) | WP_145318719.1 | WP_145318719.1 ADP-heptose synthase |
| 22(1) | WP_028542694.1 | WP_028542694.1 hypothetical protein |
| 22(1) | WP_229522865.1 | WP_229522865.1 ADP-heptose synthase |
| 22(19) | WP_013370774.1 | WP_013370774.1 hypothetical protein |
| 22(1) | WP_010346295.1 | WP_010346295.1 hypothetical protein |
| 22(1) | WP_212949024.1 | WP_212949024.1 ADP-heptose synthase |
| 22(1) | WP_250268125.1 | WP_250268125.1 ADP-heptose synthase |
| 22(3) | WP_076311062.1 | WP_076311062.1 ADP-heptose synthase |
| 22(1) | WP_208918993.1 | WP_208918993.1 ADP-heptose synthase |
| 22(1) | WP_068496262.1 | WP_068496262.1 ADP-heptose synthase |
| 22(2) | WP_090896168.1 | WP_090896168.1 ADP-heptose synthase |
| 22(1) | WP_079346846.1 | WP_079346846.1 ADP-heptose synthase |
| 22(3) | WP_025679796.1 | WP_025679796.1 hypothetical protein |
| 22(1) | WP_056700766.1 | WP_056700766.1 hypothetical protein |
| 22(1) | WP_272035286.1 | WP_272035286.1 ADP-heptose synthase |
| 22(1) | WP_018883287.1 | WP_018883287.1 hypothetical protein |
| 22(3) | WP_062833873.1 | WP_062833873.1 hypothetical protein |
| 22(3) | WP_063208793.1 | WP_063208793.1 hypothetical protein |
| 22(1) | WP_053783878.1 | WP_053783878.1 hypothetical protein |
| 22(2) | WP_145329891.1 | WP_145329891.1 ADP-heptose synthase |
| 22(1) | WP_169505044.1 | WP_169505044.1 ADP-heptose synthase |
| 22(1) | WP_127199756.1 | WP_127199756.1 ADP-heptose synthase |
| 22(1) | WP_014281173.1 | WP_014281173.1 hypothetical protein |
| 22(1) | WP_307214300.1 | WP_307214300.1 ADP-heptose synthase |
| 22(1) | WP_103047830.1 | WP_103047830.1 ADP-heptose synthase |
| 22(1) | WP_062326138.1 | WP_062326138.1 hypothetical protein |
| 22(2) | WP_095398787.1 | WP_095398787.1 ADP-heptose synthase |
| 22(1) | WP_285748154.1 | WP_285748154.1 ADP-heptose synthase |
| 22(1) | WP_110892992.1 | WP_110892992.1 ADP-heptose synthase |
| 22(1) | WP_165175582.1 | WP_165175582.1 ADP-heptose synthase |
| 22(6) | WP_063566333.1 | WP_063566333.1 hypothetical protein |
| 22(1) | WP_068622367.1 | WP_068622367.1 ADP-heptose synthase |
| 22(1) | WP_165147488.1 | WP_165147488.1 ADP-heptose synthase |
| 22(1) | WP_305022935.1 | WP_305022935.1 ADP-heptose synthase |
| 22(2) | WP_044644800.1 | WP_044644800.1 hypothetical protein |
| 22(2) | WP_220027676.1 | WP_220027676.1 ADP-heptose synthase |
| 22(1) | WP_145046439.1 | WP_145046439.1 ADP-heptose synthase |
| 22(1) | WP_106765972.1 | WP_106765972.1 ADP-heptose synthase |
| 22(1) | WP_213649980.1 | WP_213649980.1 ADP-heptose synthase |
| 22(1) | WP_025685023.1 | WP_025685023.1 hypothetical protein |
| 22(5) | WP_095287670.1 | WP_095287670.1 ADP-heptose synthase |
| 22(4) | WP_024630315.1 | WP_024630315.1 hypothetical protein |
| 22(2) | WP_194230414.1 | WP_194230414.1 ADP-heptose synthase |
| 22(1) | WP_023957691.1 | WP_023957691.1 hypothetical protein |
| 22(15) | WP_076167894.1 | WP_076167894.1 ADP-heptose synthase |
| 22(2) | WP_283655686.1 | WP_283655686.1 ADP-heptose synthase |
| 22(1) | WP_025716866.1 | WP_025716866.1 hypothetical protein |
| 22(2) | WP_128101477.1 | WP_128101477.1 ADP-heptose synthase |
| 22(3) | WP_071640572.1 | WP_071640572.1 ADP-heptose synthase |
| 22(1) | WP_123067106.1 | WP_123067106.1 ADP-heptose synthase |
| 22(1) | WP_192266435.1 | WP_192266435.1 ADP-heptose synthase |
| 22(3) | WP_039273552.1 | WP_039273552.1 hypothetical protein |
| 22(1) | WP_160037474.1 | WP_160037474.1 ADP-heptose synthase |
| 22(1) | WP_236417198.1 | WP_236417198.1 ADP-heptose synthase |
| 22(1) | WP_013309977.1 | WP_013309977.1 hypothetical protein |
| 22(2) | WP_007430121.1 | WP_007430121.1 hypothetical protein |
| 22(1) | WP_247902236.1 | WP_247902236.1 ADP-heptose synthase |
| 22(2) | WP_138226032.1 | WP_138226032.1 ADP-heptose synthase |
| 22(17) | WP_036609214.1 | WP_036609214.1 hypothetical protein |
| 22(1) | WP_244719560.1 | WP_244719560.1 ADP-heptose synthase |
| 22(1) | WP_109999884.1 | WP_109999884.1 ADP-heptose synthase |
| 22(1) | WP_047909626.1 | WP_047909626.1 hypothetical protein |
| 22(1) | WP_241929690.1 | WP_241929690.1 ADP-heptose synthase |
| 22(1) | WP_179265142.1 | WP_179265142.1 ADP-heptose synthase |
| 22(1) | WP_094593864.1 | WP_094593864.1 ADP-heptose synthase |
| 22(1) | WP_076330715.1 | WP_076330715.1 ADP-heptose synthase |
| 22(1) | WP_286458030.1 | WP_286458030.1 ADP-heptose synthase |
| 22(3) | WP_111268950.1 | WP_111268950.1 ADP-heptose synthase |
| 22(1) | WP_126665669.1 | WP_126665669.1 ADP-heptose synthase |
| 22(2) | WP_090921076.1 | WP_090921076.1 ADP-heptose synthase |
| 22(1) | WP_274485142.1 | WP_274485142.1 ADP-heptose synthase |
| 22(1) | WP_225999399.1 | WP_225999399.1 ADP-heptose synthase |
| 22(1) | WP_028589302.1 | WP_028589302.1 hypothetical protein |
| 22(4) | WP_072732434.1 | WP_072732434.1 ADP-heptose synthase |
| 22(1) | WP_305755044.1 | WP_305755044.1 ADP-heptose synthase |
| 22(1) | WP_293202271.1 | WP_293202271.1 ADP-heptose synthase |
| 22(1) | WP_136604803.1 | WP_136604803.1 ADP-heptose synthase |
| 23(1) | WP_097905191.1 | WP_097905191.1 class I SAM-dependent methyltransferase |
| 23(1) | WP_144488216.1 | WP_144488216.1 methyltransferase domain-containing protein |
| 23(1) | WP_000451293.1 | WP_000451293.1 class I SAM-dependent methyltransferase |
| 23(1) | WP_097987498.1 | WP_097987498.1 class I SAM-dependent methyltransferase |
| 23(1) | WP_139365511.1 | WP_139365511.1 class I SAM-dependent methyltransferase |
| 23(2) | WP_098266640.1 | WP_098266640.1 class I SAM-dependent methyltransferase |
| 23(1) | WP_176551423.1 | WP_176551423.1 class I SAM-dependent methyltransferase |
| 23(2) | WP_098288677.1 | WP_098288677.1 class I SAM-dependent methyltransferase |
| 23(8) | WP_053564639.1 | WP_053564639.1 class I SAM-dependent methyltransferase |
| 23(3) | WP_000451288.1 | WP_000451288.1 class I SAM-dependent methyltransferase |
| 23(1) | WP_242261448.1 | WP_242261448.1 class I SAM-dependent methyltransferase |
| 23(2) | WP_000451272.1 | WP_000451272.1 class I SAM-dependent methyltransferase |
| 23(2) | WP_197241537.1 | WP_197241537.1 class I SAM-dependent methyltransferase |
| 23(1) | WP_002148353.1 | WP_002148353.1 class I SAM-dependent methyltransferase |
| 23(1) | WP_088063226.1 | WP_088063226.1 class I SAM-dependent methyltransferase |

| | | |
|---|---|---|
| 23(1) | WP_086419828.1 | WP_086419828.1 class I SAM-dependent methyltransferase |
| 23(12) | WP_000451291.1 | WP_000451291.1 class I SAM-dependent methyltransferase |
| 23(1) | WP_098533160.1 | WP_098533160.1 class I SAM-dependent methyltransferase |
| 23(1) | WP_242400015.1 | WP_242400015.1 class I SAM-dependent methyltransferase |
| 23(1) | WP_212805119.1 | WP_212805119.1 class I SAM-dependent methyltransferase |
| 23(3) | WP_061660801.1 | WP_061660801.1 class I SAM-dependent methyltransferase |
| 23(1) | WP_144611179.1 | WP_144611179.1 class I SAM-dependent methyltransferase |
| 23(1) | WP_243500438.1 | WP_243500438.1 methyltransferase domain-containing protein |
| 23(1) | WP_262725735.1 | WP_262725735.1 class I SAM-dependent methyltransferase |
| 23(3) | WP_000451281.1 | WP_000451281.1 class I SAM-dependent methyltransferase |
| 23(1) | WP_012260825.1 | WP_012260825.1 class I SAM-dependent methyltransferase |
| 23(1) | WP_139895592.1 | WP_139895592.1 class I SAM-dependent methyltransferase |
| 23(1) | WP_098199505.1 | WP_098199505.1 methyltransferase domain-containing protein |
| 23(1) | WP_098638819.1 | WP_098638819.1 class I SAM-dependent methyltransferase |
| 23(1) | WP_199695247.1 | WP_199695247.1 class I SAM-dependent methyltransferase |
| 23(1) | WP_098256514.1 | WP_098256514.1 class I SAM-dependent methyltransferase |
| 23(2) | WP_000451275.1 | WP_000451275.1 class I SAM-dependent methyltransferase |
| 23(1) | WP_095691018.1 | WP_095691018.1 class I SAM-dependent methyltransferase |
| 23(1) | WP_098296560.1 | WP_098296560.1 class I SAM-dependent methyltransferase |
| 23(1) | WP_214328828.1 | WP_214328828.1 class I SAM-dependent methyltransferase |
| 23(1) | WP_212956548.1 | WP_212956548.1 class I SAM-dependent methyltransferase |
| 23(2) | WP_087995251.1 | WP_087995251.1 class I SAM-dependent methyltransferase |
| 23(4) | WP_023521806.1 | WP_023521806.1 class I SAM-dependent methyltransferase |
| 23(1) | WP_081257078.1 | WP_081257078.1 class I SAM-dependent methyltransferase |
| 23(1) | WP_270728493.1 | WP_270728493.1 class I SAM-dependent methyltransferase |
| 23(1) | WP_042874973.1 | WP_042874973.1 class I SAM-dependent methyltransferase |
| 23(1) | WP_000451267.1 | WP_000451267.1 class I SAM-dependent methyltransferase |
| 23(1) | WP_163255247.1 | WP_163255247.1 class I SAM-dependent methyltransferase |
| 23(1) | WP_202283179.1 | WP_202283179.1 class I SAM-dependent methyltransferase |
| 23(1) | WP_215601123.1 | WP_215601123.1 methyltransferase domain-containing protein |
| 23(1) | WP_098562458.1 | WP_098562458.1 class I SAM-dependent methyltransferase |
| 23(1) | WP_144655996.1 | WP_144655996.1 class I SAM-dependent methyltransferase |
| 23(2) | WP_098264278.1 | WP_098264278.1 class I SAM-dependent methyltransferase |
| 23(1) | WP_016512887.1 | WP_016512887.1 class I SAM-dependent methyltransferase |
| 23(2) | WP_090976414.1 | WP_090976414.1 class I SAM-dependent methyltransferase |
| 23(1) | WP_242251757.1 | WP_242251757.1 class I SAM-dependent methyltransferase |
| 23(1) | WP_207994170.1 | WP_207994170.1 methyltransferase domain-containing protein |
| 23(1) | WP_060630639.1 | WP_060630639.1 class I SAM-dependent methyltransferase |
| 23(2) | WP_098088261.1 | WP_098088261.1 methyltransferase domain-containing protein |
| 23(1) | WP_106822170.1 | WP_106822170.1 class I SAM-dependent methyltransferase |
| 23(1) | WP_215591195.1 | WP_215591195.1 class I SAM-dependent methyltransferase |
| 23(1) | WP_074565311.1 | WP_074565311.1 class I SAM-dependent methyltransferase |
| 23(1) | WP_075395493.1 | WP_075395493.1 class I SAM-dependent methyltransferase |
| 23(2) | WP_098360830.1 | WP_098360830.1 class I SAM-dependent methyltransferase |
| 23(4) | WP_033709779.1 | WP_033709779.1 class I SAM-dependent methyltransferase |
| 23(1) | WP_098519058.1 | WP_098519058.1 class I SAM-dependent methyltransferase |
| 23(1) | WP_088609109.1 | WP_088609109.1 class I SAM-dependent methyltransferase |
| 23(1) | WP_270696895.1 | WP_270696895.1 class I SAM-dependent methyltransferase |
| 23(1) | WP_070805896.1 | WP_070805896.1 class I SAM-dependent methyltransferase |
| 23(1) | WP_270711667.1 | WP_270711667.1 methyltransferase domain-containing protein |
| 23(1) | WP_017149780.1 | WP_017149780.1 methyltransferase domain-containing protein |
| 23(1) | WP_144564262.1 | WP_144564262.1 class I SAM-dependent methyltransferase |
| 23(1) | WP_144492096.1 | WP_144492096.1 class I SAM-dependent methyltransferase |
| 23(1) | WP_061535956.1 | WP_061535956.1 class I SAM-dependent methyltransferase |
| 23(1) | WP_143881065.1 | WP_143881065.1 class I SAM-dependent methyltransferase |
| 23(1) | WP_098451710.1 | WP_098451710.1 class I SAM-dependent methyltransferase |
| 23(2) | WP_098991580.1 | WP_098991580.1 class I SAM-dependent methyltransferase |
| 23(1) | WP_071719992.1 | WP_071719992.1 class I SAM-dependent methyltransferase |
| 23(1) | WP_098214777.1 | WP_098214777.1 class I SAM-dependent methyltransferase |
| 23(2) | WP_157827839.1 | WP_157827839.1 class I SAM-dependent methyltransferase |
| 23(3) | WP_097952796.1 | WP_097952796.1 class I SAM-dependent methyltransferase |
| 23(2) | WP_078205525.1 | WP_078205525.1 class I SAM-dependent methyltransferase |
| 23(1) | WP_044792570.1 | WP_044792570.1 class I SAM-dependent methyltransferase |
| 23(1) | WP_262748115.1 | WP_262748115.1 class I SAM-dependent methyltransferase |
| 23(1) | WP_192797662.1 | WP_192797662.1 class I SAM-dependent methyltransferase |
| 23(5) | WP_000451274.1 | WP_000451274.1 class I SAM-dependent methyltransferase |
| 23(1) | WP_270710641.1 | WP_270710641.1 class I SAM-dependent methyltransferase |
| 23(1) | WP_166704979.1 | WP_166704979.1 class I SAM-dependent methyltransferase |
| 23(1) | WP_208019339.1 | WP_208019339.1 class I SAM-dependent methyltransferase |
| 23(1) | WP_000451289.1 | WP_000451289.1 class I SAM-dependent methyltransferase |
| 23(1) | WP_000451280.1 | WP_000451280.1 class I SAM-dependent methyltransferase |
| 23(3) | WP_176546139.1 | WP_176546139.1 class I SAM-dependent methyltransferase |
| 23(4) | WP_087982603.1 | WP_087982603.1 class I SAM-dependent methyltransferase |
| 23(1) | WP_286017536.1 | WP_286017536.1 class I SAM-dependent methyltransferase |
| 23(1) | WP_242264964.1 | WP_242264964.1 class I SAM-dependent methyltransferase |
| 23(1) | WP_081252594.1 | WP_081252594.1 class I SAM-dependent methyltransferase |
| 23(1) | WP_139867394.1 | WP_139867394.1 class I SAM-dependent methyltransferase |
| 23(1) | WP_088032761.1 | WP_088032761.1 class I SAM-dependent methyltransferase |
| 23(1) | WP_001980122.1 | WP_001980122.1 class I SAM-dependent methyltransferase |
| 23(1) | WP_189339432.1 | WP_189339432.1 class I SAM-dependent methyltransferase |
| 23(1) | WP_140160091.1 | WP_140160091.1 class I SAM-dependent methyltransferase |
| 23(1) | WP_098377167.1 | WP_098377167.1 class I SAM-dependent methyltransferase |
| 23(2) | WP_071679485.1 | WP_071679485.1 class I SAM-dependent methyltransferase |
| 23(1) | WP_229137981.1 | WP_229137981.1 class I SAM-dependent methyltransferase |
| 23(1) | WP_215570360.1 | WP_215570360.1 methyltransferase domain-containing protein |
| 23(1) | WP_087956284.1 | WP_087956284.1 methyltransferase domain-containing protein |
| 23(1) | WP_098167604.1 | WP_098167604.1 class I SAM-dependent methyltransferase |
| 23(1) | WP_048552295.1 | WP_048552295.1 class I SAM-dependent methyltransferase |
| 23(3) | WP_000451279.1 | WP_000451279.1 class I SAM-dependent methyltransferase |
| 23(3) | WP_097920542.1 | WP_097920542.1 class I SAM-dependent methyltransferase |
| 23(1) | WP_157418144.1 | WP_157418144.1 class I SAM-dependent methyltransferase |
| 23(1) | WP_098834579.1 | WP_098834579.1 class I SAM-dependent methyltransferase |
| 23(1) | WP_000417079.1 | WP_000417079.1 class I SAM-dependent methyltransferase |
| 23(4) | WP_103626225.1 | WP_103626225.1 class I SAM-dependent methyltransferase |
| 23(3) | WP_000451285.1 | WP_000451285.1 class I SAM-dependent methyltransferase |
| 23(1) | WP_098841820.1 | WP_098841820.1 class I SAM-dependent methyltransferase |
| 23(1) | WP_000451270.1 | WP_000451270.1 class I SAM-dependent methyltransferase |
| 23(2) | WP_061184805.1 | WP_061184805.1 class I SAM-dependent methyltransferase |
| 23(1) | WP_284566220.1 | WP_284566220.1 class I SAM-dependent methyltransferase |
| 23(1) | WP_026593091.1 | WP_026593091.1 methyltransferase domain-containing protein |
| 23(12) | WP_011732838.1 | WP_011732838.1 class I SAM-dependent methyltransferase |
| 23(2) | WP_071724707.1 | WP_071724707.1 class I SAM-dependent methyltransferase |
| 23(1) | WP_029438082.1 | WP_029438082.1 class I SAM-dependent methyltransferase |

| | | |
|---|---|---|
| 24(1) | WP_195325260.1 | WP_195325260.1 methyltransferase |
| 24(1) | WP_294386454.1 | WP_294386454.1 methyltransferase |
| 24(2) | WP_034773561.1 | WP_034773561.1 hypothetical protein |
| 24(1) | WP_295628310.1 | WP_295628310.1 methyltransferase |
| 24(1) | WP_216830391.1 | WP_216830391.1 methyltransferase |
| 24(1) | WP_254759294.1 | WP_254759294.1 EsaB/YukD family protein |
| 24(1) | WP_026572739.1 | WP_026572739.1 methyltransferase |
| 24(1) | WP_079506392.1 | WP_079506392.1 methyltransferase |
| 24(1) | WP_214902935.1 | WP_214902935.1 methyltransferase |
| 24(1) | WP_195244533.1 | WP_195244533.1 methyltransferase |
| 24(1) | WP_179160622.1 | WP_179160622.1 methyltransferase |
| 24(1) | WP_238567604.1 | WP_238567604.1 hypothetical protein |
| 24(1) | WP_258207828.1 | WP_258207828.1 methyltransferase |
| 24(1) | WP_272537437.1 | WP_272537437.1 methyltransferase |
| 24(1) | WP_195515239.1 | WP_195515239.1 methyltransferase |
| 24(1) | WP_224839861.1 | WP_224839861.1 methyltransferase |
| 24(1) | WP_277166573.1 | WP_277166573.1 hypothetical protein |
| 24(1) | WP_138755921.1 | WP_138755921.1 methyltransferase |
| 24(4) | WP_291288270.1 | WP_291288270.1 hypothetical protein |
| 24(1) | WP_251426367.1 | WP_251426367.1 methyltransferase |
| 24(1) | WP_048826587.1 | WP_048826587.1 hypothetical protein |
| 24(3) | WP_102372745.1 | WP_102372745.1 hypothetical protein |
| 24(1) | WP_102264551.1 | WP_102264551.1 methyltransferase |
| 24(1) | WP_068673720.1 | WP_068673720.1 methyltransferase |
| 24(1) | WP_013490131.1 | WP_013490131.1 hypothetical protein |
| 24(1) | WP_005210439.1 | WP_005210439.1 hypothetical protein |
| 24(1) | WP_263324545.1 | WP_263324545.1 EsaB/YukD family protein |
| 24(1) | WP_045518284.1 | WP_045518284.1 hypothetical protein |
| 24(1) | WP_284036499.1 | WP_284036499.1 methyltransferase |
| 24(1) | WP_127570438.1 | WP_127570438.1 methyltransferase |
| 24(1) | WP_231094850.1 | WP_231094850.1 methyltransferase |
| 24(1) | WP_238187881.1 | WP_238187881.1 EsaB/YukD family protein |
| 24(1) | WP_270574774.1 | WP_270574774.1 hypothetical protein |
| 24(1) | WP_006290898.1 | WP_006290898.1 hypothetical protein |
| 24(1) | WP_231417864.1 | WP_231417864.1 methyltransferase |
| 24(1) | WP_307738731.1 | WP_307738731.1 hypothetical protein |
| 24(1) | WP_136379698.1 | WP_136379698.1 methyltransferase |
| 24(1) | WP_010676865.1 | WP_010676865.1 methyltransferase |
| 24(1) | WP_034675364.1 | WP_034675364.1 methyltransferase |
| 24(4) | WP_010496612.1 | WP_010496612.1 hypothetical protein |
| 24(1) | WP_227219186.1 | WP_227219186.1 EsaB/YukD family protein |
| 24(1) | WP_113672702.1 | WP_113672702.1 hypothetical protein |
| 24(1) | WP_073008259.1 | WP_073008259.1 methyltransferase |
| 24(1) | WP_254366012.1 | WP_254366012.1 EsaB/YukD family protein |
| 24(1) | WP_015595762.1 | WP_015595762.1 hypothetical protein |
| 24(1) | WP_268613681.1 | WP_268613681.1 methyltransferase |
| 24(1) | WP_277157934.1 | WP_277157934.1 hypothetical protein |
| 24(1) | WP_066022454.1 | WP_066022454.1 methyltransferase |
| 24(1) | WP_136372955.1 | WP_136372955.1 methyltransferase |
| 24(1) | WP_191272867.1 | WP_191272867.1 methyltransferase |
| 24(1) | WP_223592678.1 | WP_223592678.1 methyltransferase |
| 24(1) | WP_226602924.1 | WP_226602924.1 methyltransferase |
| 24(1) | WP_166245123.1 | WP_166245123.1 methyltransferase |
| 24(1) | WP_041964579.1 | WP_041964579.1 hypothetical protein |
| 24(3) | WP_144479064.1 | WP_144479064.1 methyltransferase |
| 24(1) | WP_163853472.1 | WP_163853472.1 methyltransferase |
| 24(1) | WP_077619652.1 | WP_077619652.1 DUF2604 domain-containing protein |
| 24(1) | WP_308084174.1 | WP_308084174.1 methyltransferase |
| 24(1) | WP_195953748.1 | WP_195953748.1 methyltransferase |
| 24(1) | WP_109625243.1 | WP_109625243.1 methyltransferase |
| 24(1) | WP_295632044.1 | WP_295632044.1 hypothetical protein |
| 24(1) | WP_215032127.1 | WP_215032127.1 methyltransferase |
| 24(1) | WP_213144477.1 | WP_213144477.1 methyltransferase |
| 24(1) | WP_213117478.1 | WP_213117478.1 methyltransferase |
| 24(1) | WP_303942544.1 | WP_303942544.1 hypothetical protein |
| 24(1) | WP_102371676.1 | WP_102371676.1 hypothetical protein |
| 24(1) | WP_199419772.1 | WP_199419772.1 methyltransferase |
| 24(1) | WP_007083288.1 | WP_007083288.1 hypothetical protein |
| 24(1) | WP_307589599.1 | WP_307589599.1 methyltransferase |
| 24(1) | WP_038584738.1 | WP_038584738.1 hypothetical protein |
| 24(1) | WP_212695787.1 | WP_212695787.1 hypothetical protein |
| 24(1) | WP_212330865.1 | WP_212330865.1 hypothetical protein |
| 24(1) | WP_108534960.1 | WP_108534960.1 methyltransferase |
| 24(1) | WP_204412725.1 | WP_204412725.1 methyltransferase |
| 24(1) | WP_235286199.1 | WP_235286199.1 EsaB/YukD family protein |
| 24(1) | WP_179603481.1 | WP_179603481.1 methyltransferase |
| 24(1) | WP_090674364.1 | WP_090674364.1 methyltransferase |
| 24(1) | WP_006720320.1 | WP_006720320.1 hypothetical protein |
| 24(1) | WP_036689415.1 | WP_036689415.1 hypothetical protein |
| 24(1) | WP_277156178.1 | WP_277156178.1 hypothetical protein |
| 24(1) | WP_054972094.1 | WP_054972094.1 hypothetical protein |
| 24(3) | WP_025848614.1 | WP_025848614.1 hypothetical protein |
| 24(2) | WP_112331591.1 | WP_112331591.1 methyltransferase |
| 24(1) | WP_283952950.1 | WP_283952950.1 methyltransferase |
| 24(1) | WP_120112262.1 | WP_120112262.1 methyltransferase |
| 24(1) | WP_277142076.1 | WP_277142076.1 hypothetical protein |
| 24(1) | WP_027088424.1 | WP_027088424.1 hypothetical protein |
| 24(1) | WP_239617478.1 | WP_239617478.1 methyltransferase |
| 24(1) | WP_148409443.1 | WP_148409443.1 methyltransferase |
| 24(1) | WP_204445221.1 | WP_204445221.1 methyltransferase |
| 24(1) | WP_288218587.1 | WP_288218587.1 methyltransferase |
| 24(1) | WP_293755526.1 | WP_293755526.1 hypothetical protein |
| 24(2) | WP_121610594.1 | WP_121610594.1 methyltransferase |
| 24(2) | WP_212693238.1 | WP_212693238.1 hypothetical protein |
| 24(1) | WP_006292344.1 | WP_006292344.1 hypothetical protein |
| 24(1) | WP_217035851.1 | WP_217035851.1 EsaB/YukD family protein |
| 24(1) | WP_046522784.1 | WP_046522784.1 hypothetical protein |
| 24(1) | WP_253411080.1 | WP_253411080.1 EsaB/YukD family protein |
| 24(1) | WP_208926620.1 | WP_208926620.1 methyltransferase |
| 24(1) | WP_221875373.1 | WP_221875373.1 methyltransferase |
| 24(1) | WP_127459434.1 | WP_127459434.1 methyltransferase |
| 24(1) | WP_070638663.1 | WP_070638663.1 hypothetical protein |
| 24(1) | WP_251638786.1 | WP_251638786.1 methyltransferase |
| 24(1) | WP_307312547.1 | WP_307312547.1 methyltransferase |

| | | |
|---|---|---|
| 24(1) | WP_024027628.1 | WP_024027628.1 hypothetical protein |
| 24(1) | WP_226642874.1 | WP_226642874.1 methyltransferase |
| 24(2) | WP_281813100.1 | WP_281813100.1 hypothetical protein |
| 24(1) | WP_130810267.1 | WP_130810267.1 hypothetical protein |
| 24(1) | WP_157076701.1 | WP_157076701.1 methyltransferase |
| 24(1) | WP_308082242.1 | WP_308082242.1 methyltransferase |
| 24(1) | WP_254782314.1 | WP_254782314.1 EsaB/YukD family protein |
| 24(1) | WP_204666344.1 | WP_204666344.1 hypothetical protein |
| 24(1) | WP_308110116.1 | WP_308110116.1 methyltransferase |
| 24(1) | WP_021725643.1 | WP_021725643.1 hypothetical protein |
| 24(1) | WP_214706002.1 | WP_214706002.1 hypothetical protein |
| 24(1) | WP_186580063.1 | WP_186580063.1 methyltransferase |
| 24(1) | WP_134750606.1 | WP_134750606.1 methyltransferase |
| 24(1) | WP_010250274.1 | WP_010250274.1 hypothetical protein |
| 24(1) | WP_167831135.1 | WP_167831135.1 methyltransferase |
| 24(1) | WP_268058026.1 | WP_268058026.1 methyltransferase |
| 24(1) | WP_182301838.1 | WP_182301838.1 methyltransferase |
| 24(2) | WP_133332981.1 | WP_133332981.1 methyltransferase |
| 24(1) | WP_005604635.1 | WP_005604635.1 hypothetical protein |
| 24(1) | WP_143316562.1 | WP_143316562.1 methyltransferase |
| 24(1) | WP_116353253.1 | WP_116353253.1 methyltransferase |
| 24(1) | WP_088830539.1 | WP_088830539.1 methyltransferase |
| 24(1) | WP_196048854.1 | WP_196048854.1 methyltransferase |
| 24(1) | WP_304216865.1 | WP_304216865.1 methyltransferase |
| 24(1) | WP_306077390.1 | WP_306077390.1 EsaB/YukD family protein |
| 24(1) | WP_229139918.1 | WP_229139918.1 methyltransferase |
| 24(1) | WP_216791220.1 | WP_216791220.1 methyltransferase |
| 24(2) | WP_006289746.1 | WP_006289746.1 hypothetical protein |
| 24(9) | WP_042400990.1 | WP_042400990.1 hypothetical protein |
| 24(2) | WP_277157940.1 | WP_277157940.1 hypothetical protein |
| 24(1) | WP_251466800.1 | WP_251466800.1 methyltransferase |
| 24(1) | WP_226679104.1 | WP_226679104.1 methyltransferase |
| 24(1) | WP_066419980.1 | WP_066419980.1 methyltransferase |
| 24(1) | WP_007147987.1 | WP_007147987.1 hypothetical protein |
| 24(1) | WP_251548763.1 | WP_251548763.1 methyltransferase |
| 24(1) | WP_111147804.1 | WP_111147804.1 methyltransferase |
| 24(1) | WP_076113450.1 | WP_076113450.1 methyltransferase |
| 24(3) | WP_099354906.1 | WP_099354906.1 methyltransferase |
| 24(1) | WP_251245815.1 | WP_251245815.1 methyltransferase |
| 24(1) | WP_289613143.1 | WP_289613143.1 hypothetical protein |
| 24(1) | WP_210364450.1 | WP_210364450.1 methyltransferase |
| 24(1) | WP_289342260.1 | WP_289342260.1 methyltransferase |
| 24(1) | WP_198310831.1 | WP_198310831.1 methyltransferase |
| 24(1) | WP_307601026.1 | WP_307601026.1 methyltransferase |
| 24(1) | WP_294953135.1 | WP_294953135.1 methyltransferase |
| 24(1) | WP_149871684.1 | WP_149871684.1 methyltransferase |
| 24(1) | WP_289607493.1 | WP_289607493.1 hypothetical protein |
| 24(1) | WP_277150361.1 | WP_277150361.1 hypothetical protein |
| 24(1) | WP_042462810.1 | WP_042462810.1 methyltransferase |
| 24(1) | WP_066092006.1 | WP_066092006.1 methyltransferase |
| 24(1) | WP_058271116.1 | WP_058271116.1 hypothetical protein |
| 24(1) | WP_234121606.1 | WP_234121606.1 methyltransferase |

| | | |
|---|---|---|
| 24(1) | WP_307192897.1 | WP_307192897.1 EsaB/YukD family protein |
| 24(1) | WP_122971531.1 | WP_122971531.1 EsaB/YukD family protein |
| 24(1) | WP_251480995.1 | WP_251480995.1 methyltransferase |
| 24(1) | WP_307284676.1 | WP_307284676.1 methyltransferase |
| 24(1) | WP_251557379.1 | WP_251557379.1 methyltransferase |
| 24(1) | WP_231687843.1 | WP_231687843.1 methyltransferase |
| 25(1) | WP_256704104.1 | WP_256704104.1 ribosome small subunit-dependent GTPase A |
| 25(1) | WP_231952578.1 | WP_231952578.1 ribosome small subunit-dependent GTPase A |
| 25(1) | WP_017689080.1 | WP_017689080.1 ribosome small subunit-dependent GTPase A |
| 25(2) | WP_023988259.1 | WP_023988259.1 ribosome small subunit-dependent GTPase A |
| 25(1) | WP_256975168.1 | WP_256975168.1 ribosome small subunit-dependent GTPase A |
| 25(1) | WP_228469271.1 | WP_228469271.1 ribosome small subunit-dependent GTPase A |
| 25(1) | WP_094155410.1 | WP_094155410.1 ribosome small subunit-dependent GTPase A |
| 25(2) | WP_013370778.1 | WP_013370778.1 ribosome small subunit-dependent GTPase A |
| 25(1) | WP_152381392.1 | WP_152381392.1 ribosome small subunit-dependent GTPase A |
| 25(1) | WP_017427963.1 | WP_017427963.1 ribosome small subunit-dependent GTPase A |
| 25(1) | WP_025716870.1 | WP_025716870.1 ribosome small subunit-dependent GTPase A |
| 25(1) | WP_031463343.1 | WP_031463343.1 ribosome small subunit-dependent GTPase A |
| 25(2) | WP_025679799.1 | WP_025679799.1 ribosome small subunit-dependent GTPase A |
| 25(1) | WP_013309982.1 | WP_013309982.1 ribosome small subunit-dependent GTPase A |
| 25(1) | WP_256221987.1 | WP_256221987.1 ribosome small subunit-dependent GTPase A |
| 25(1) | WP_221791103.1 | WP_221791103.1 ribosome small subunit-dependent GTPase A |
| 25(2) | WP_240913009.1 | WP_240913009.1 ribosome small subunit-dependent GTPase A |
| 25(1) | WP_076330717.1 | WP_076330717.1 ribosome small subunit-dependent GTPase A |
| 25(1) | WP_255247852.1 | WP_255247852.1 ribosome small subunit-dependent GTPase A |
| 25(1) | WP_254325797.1 | WP_254325797.1 ribosome small subunit-dependent GTPase A |
| 25(1) | WP_076156859.1 | WP_076156859.1 ribosome small subunit-dependent GTPase A |
| 25(2) | WP_228100913.1 | WP_228100913.1 ribosome small subunit-dependent GTPase A |
| 25(1) | WP_236146740.1 | WP_236146740.1 ribosome small subunit-dependent GTPase A |
| 25(1) | WP_090921071.1 | WP_090921071.1 ribosome small subunit-dependent GTPase A |
| 25(1) | WP_256702679.1 | WP_256702679.1 ribosome small subunit-dependent GTPase A |
| 25(1) | WP_244867300.1 | WP_244867300.1 ribosome small subunit-dependent GTPase A |
| 25(1) | WP_256992147.1 | WP_256992147.1 ribosome small subunit-dependent GTPase A |
| 25(1) | WP_244533153.1 | WP_244533153.1 ribosome small subunit-dependent GTPase A |
| 25(1) | WP_160790132.1 | WP_160790132.1 ribosome small subunit-dependent GTPase A |
| 25(1) | WP_246058077.1 | WP_246058077.1 ribosome small subunit-dependent GTPase A |
| 25(1) | WP_260870916.1 | WP_260870916.1 ribosome small subunit-dependent GTPase A |
| 25(1) | WP_240636540.1 | WP_240636540.1 ribosome small subunit-dependent GTPase A |
| 25(1) | WP_063208785.1 | WP_063208785.1 ribosome small subunit-dependent GTPase A |
| 25(2) | WP_076249780.1 | WP_076249780.1 ribosome small subunit-dependent GTPase A |
| 25(1) | WP_305166266.1 | WP_305166266.1 ribosome small subunit-dependent GTPase A |
| 25(1) | WP_134910008.1 | WP_134910008.1 ribosome small subunit-dependent GTPase A |
| 25(1) | WP_257219302.1 | WP_257219302.1 ribosome small subunit-dependent GTPase A |
| 25(2) | WP_113050532.1 | WP_113050532.1 ribosome small subunit-dependent GTPase A |
| 25(1) | WP_307349955.1 | WP_307349955.1 ribosome small subunit-dependent GTPase A |
| 25(1) | WP_235540753.1 | WP_235540753.1 ribosome small subunit-dependent GTPase A |
| 25(1) | WP_029517058.1 | WP_029517058.1 ribosome small subunit-dependent GTPase A |
| 25(1) | WP_293202269.1 | WP_293202269.1 ribosome small subunit-dependent GTPase A |
| 25(1) | WP_229729890.1 | WP_229729890.1 ribosome small subunit-dependent GTPase A |
| 25(1) | WP_258296283.1 | WP_258296283.1 ribosome small subunit-dependent GTPase A |
| 25(1) | WP_244898321.1 | WP_244898321.1 ribosome small subunit-dependent GTPase A |

| | | |
|---|---|---|
| 25(1) | WP_236703655.1 | WP_236703655.1 ribosome small subunit-dependent GTPase A |
| 25(1) | WP_305755040.1 | WP_305755040.1 ribosome small subunit-dependent GTPase A |
| 25(1) | WP_068496255.1 | WP_068496255.1 ribosome small subunit-dependent GTPase A |
| 25(1) | WP_258530583.1 | WP_258530583.1 ribosome small subunit-dependent GTPase A |
| 25(1) | WP_264929214.1 | WP_264929214.1 ribosome small subunit-dependent GTPase A |
| 25(1) | WP_240037246.1 | WP_240037246.1 ribosome small subunit-dependent GTPase A |
| 25(1) | WP_235200749.1 | WP_235200749.1 ribosome small subunit-dependent GTPase A |
| 25(1) | WP_301879172.1 | WP_301879172.1 ribosome small subunit-dependent GTPase A |
| 25(1) | WP_076264231.1 | WP_076264231.1 ribosome small subunit-dependent GTPase A |
| 25(2) | WP_244192856.1 | WP_244192856.1 ribosome small subunit-dependent GTPase A |
| 25(1) | WP_312887166.1 | WP_312887166.1 ribosome small subunit-dependent GTPase A |
| 25(1) | WP_251583277.1 | WP_251583277.1 ribosome small subunit-dependent GTPase A |
| 25(1) | WP_209941809.1 | WP_209941809.1 ribosome small subunit-dependent GTPase A |
| 25(1) | WP_237391670.1 | WP_237391670.1 ribosome small subunit-dependent GTPase A |
| 25(1) | WP_250268126.1 | WP_250268126.1 ribosome small subunit-dependent GTPase A |
| 25(2) | WP_254438536.1 | WP_254438536.1 ribosome small subunit-dependent GTPase A |
| 25(1) | WP_244552051.1 | WP_244552051.1 ribosome small subunit-dependent GTPase A |
| 25(1) | WP_256721270.1 | WP_256721270.1 ribosome small subunit-dependent GTPase A |
| 25(1) | WP_094936503.1 | WP_094936503.1 ribosome small subunit-dependent GTPase A |
| 25(1) | WP_014281177.1 | WP_014281177.1 ribosome small subunit-dependent GTPase A |
| 25(1) | WP_249717202.1 | WP_249717202.1 ribosome small subunit-dependent GTPase A |
| 25(1) | WP_102997916.1 | WP_102997916.1 ribosome small subunit-dependent GTPase A |
| 25(1) | WP_058710500.1 | WP_058710500.1 ribosome small subunit-dependent GTPase A |
| 25(2) | WP_283655688.1 | WP_283655688.1 ribosome small subunit-dependent GTPase A |
| 25(1) | WP_242210844.1 | WP_242210844.1 ribosome small subunit-dependent GTPase A |
| 25(1) | WP_127543999.1 | WP_127543999.1 ribosome small subunit-dependent GTPase A |
| 25(1) | WP_150367645.1 | WP_150367645.1 ribosome small subunit-dependent GTPase A |
| 25(1) | WP_024630319.1 | WP_024630319.1 ribosome small subunit-dependent GTPase A |
| 25(1) | WP_260869264.1 | WP_260869264.1 ribosome small subunit-dependent GTPase A |
| 25(1) | WP_285748160.1 | WP_285748160.1 ribosome small subunit-dependent GTPase A |
| 25(1) | WP_246072898.1 | WP_246072898.1 ribosome small subunit-dependent GTPase A |
| 25(1) | WP_058829510.1 | WP_058829510.1 ribosome small subunit-dependent GTPase A |
| 25(1) | WP_290409171.1 | WP_290409171.1 ribosome small subunit-dependent GTPase A |
| 25(1) | WP_110892984.1 | WP_110892984.1 ribosome small subunit-dependent GTPase A |
| 25(1) | WP_244719593.1 | WP_244719593.1 ribosome small subunit-dependent GTPase A |
| 25(1) | WP_301249693.1 | WP_301249693.1 ribosome small subunit-dependent GTPase A |
| 25(2) | WP_028542693.1 | WP_028542693.1 ribosome small subunit-dependent GTPase A |
| 25(1) | WP_249281029.1 | WP_249281029.1 ribosome small subunit-dependent GTPase A |
| 25(1) | WP_254349523.1 | WP_254349523.1 ribosome small subunit-dependent GTPase A |
| 25(1) | WP_010346291.1 | WP_010346291.1 ribosome small subunit-dependent GTPase A |
| 25(3) | WP_025364678.1 | WP_025364678.1 ribosome small subunit-dependent GTPase A |
| 25(1) | WP_289392067.1 | WP_289392067.1 ribosome small subunit-dependent GTPase A |
| 25(1) | WP_260985317.1 | WP_260985317.1 ribosome small subunit-dependent GTPase A |
| 25(1) | WP_051449744.1 | WP_051449744.1 ribosome small subunit-dependent GTPase A |
| 25(1) | WP_269055427.1 | WP_269055427.1 ribosome small subunit-dependent GTPase A |
| 25(1) | WP_272035292.1 | WP_272035292.1 ribosome small subunit-dependent GTPase A |
| 25(1) | WP_100529446.1 | WP_100529446.1 ribosome small subunit-dependent GTPase A |
| 25(1) | WP_068940677.1 | WP_068940677.1 ribosome small subunit-dependent GTPase A |
| 25(1) | WP_256336117.1 | WP_256336117.1 ribosome small subunit-dependent GTPase A |
| 25(1) | WP_236417202.1 | WP_236417202.1 ribosome small subunit-dependent GTPase A |
| 25(1) | WP_216513483.1 | WP_216513483.1 ribosome small subunit-dependent GTPase A |
| 25(1) | WP_208332486.1 | WP_208332486.1 ribosome small subunit-dependent GTPase A |
| 25(1) | WP_225999403.1 | WP_225999403.1 ribosome small subunit-dependent GTPase A |
| 25(1) | WP_145146961.1 | WP_145146961.1 ribosome small subunit-dependent GTPase A |
| 25(1) | WP_263880223.1 | WP_263880223.1 ribosome small subunit-dependent GTPase A |
| 25(1) | WP_090887138.1 | WP_090887138.1 ribosome small subunit-dependent GTPase A |
| 25(1) | WP_124115518.1 | WP_124115518.1 ribosome small subunit-dependent GTPase A |
| 25(10) | WP_025719067.1 | WP_025719067.1 ribosome small subunit-dependent GTPase A |
| 25(1) | WP_213649975.1 | WP_213649975.1 ribosome small subunit-dependent GTPase A |
| 25(1) | WP_104495343.1 | WP_104495343.1 ribosome small subunit-dependent GTPase A |
| 25(1) | WP_240767977.1 | WP_240767977.1 ribosome small subunit-dependent GTPase A |
| 25(1) | WP_018883280.1 | WP_018883280.1 ribosome small subunit-dependent GTPase A |
| 25(1) | WP_254790063.1 | WP_254790063.1 ribosome small subunit-dependent GTPase A |
| 25(1) | WP_307214302.1 | WP_307214302.1 ribosome small subunit-dependent GTPase A |
| 25(1) | WP_225229037.1 | WP_225229037.1 ribosome small subunit-dependent GTPase A |
| 25(1) | WP_238546254.1 | WP_238546254.1 ribosome small subunit-dependent GTPase A |
| 25(1) | WP_076167882.1 | WP_076167882.1 ribosome small subunit-dependent GTPase A |
| 25(2) | WP_256337825.1 | WP_256337825.1 ribosome small subunit-dependent GTPase A |
| 25(1) | WP_286458028.1 | WP_286458028.1 ribosome small subunit-dependent GTPase A |
| 25(1) | WP_076292679.1 | WP_076292679.1 ribosome small subunit-dependent GTPase A |
| 25(2) | WP_137078362.1 | WP_137078362.1 ribosome small subunit-dependent GTPase A |
| 25(1) | WP_286458029.1 | WP_286458029.1 hypothetical protein |
| 25(1) | WP_053061521.1 | WP_053061521.1 ribosome small subunit-dependent GTPase A |
| 25(1) | WP_145329899.1 | WP_145329899.1 ribosome small subunit-dependent GTPase A |
| 25(1) | WP_244364317.1 | WP_244364317.1 ribosome small subunit-dependent GTPase A |
| 25(1) | WP_285079610.1 | WP_285079610.1 ribosome small subunit-dependent GTPase A |
| 25(1) | WP_229522862.1 | WP_229522862.1 ribosome small subunit-dependent GTPase A |
| 25(1) | WP_257031449.1 | WP_257031449.1 ribosome small subunit-dependent GTPase A |
| 25(5) | WP_016821025.1 | WP_016821025.1 ribosome small subunit-dependent GTPase A |
| 25(1) | WP_039273545.1 | WP_039273545.1 ribosome small subunit-dependent GTPase A |
| 25(1) | WP_007430126.1 | WP_007430126.1 ribosome small subunit-dependent GTPase A |
| 25(1) | WP_153028179.1 | WP_153028179.1 ribosome small subunit-dependent GTPase A |
| 25(1) | WP_260867009.1 | WP_260867009.1 ribosome small subunit-dependent GTPase A |
| 25(1) | WP_216975171.1 | WP_216975171.1 ribosome small subunit-dependent GTPase A |
| 25(1) | WP_025685027.1 | WP_025685027.1 ribosome small subunit-dependent GTPase A |
| 25(2) | WP_240321499.1 | WP_240321499.1 ribosome small subunit-dependent GTPase A |
| 25(1) | WP_256209539.1 | WP_256209539.1 ribosome small subunit-dependent GTPase A |
| 25(1) | WP_258377837.1 | WP_258377837.1 ribosome small subunit-dependent GTPase A |
| 25(1) | WP_234489613.1 | WP_234489613.1 ribosome small subunit-dependent GTPase A |
| 25(1) | WP_192510494.1 | WP_192510494.1 ribosome small subunit-dependent GTPase A |
| 25(1) | WP_260986748.1 | WP_260986748.1 ribosome small subunit-dependent GTPase A |
| 25(1) | WP_307537590.1 | WP_307537590.1 ribosome small subunit-dependent GTPase A |
| 25(1) | WP_245683698.1 | WP_245683698.1 ribosome small subunit-dependent GTPase A |
| 25(1) | WP_244879217.1 | WP_244879217.1 ribosome small subunit-dependent GTPase A |
| 25(1) | WP_123797112.1 | WP_123797112.1 ribosome small subunit-dependent GTPase A |
| 25(1) | WP_250244809.1 | WP_250244809.1 ribosome small subunit-dependent GTPase A |
| 25(1) | WP_028589298.1 | WP_028589298.1 ribosome small subunit-dependent GTPase A |
| 25(1) | WP_160037479.1 | WP_160037479.1 ribosome small subunit-dependent GTPase A |
| 25(1) | WP_305022930.1 | WP_305022930.1 ribosome small subunit-dependent GTPase A |
| 25(1) | WP_247902238.1 | WP_247902238.1 ribosome small subunit-dependent GTPase A |
| 25(1) | WP_256992765.1 | WP_256992765.1 ribosome small subunit-dependent GTPase A |
| 25(1) | WP_258875222.1 | WP_258875222.1 ribosome small subunit-dependent GTPase A |
| 25(1) | WP_142612999.1 | WP_142612999.1 ribosome small subunit-dependent GTPase A |
| 25(1) | WP_145412702.1 | WP_145412702.1 ribosome small subunit-dependent GTPase A |

| | | |
|---|---|---|
| 25(1) | WP_260845578.1 | WP_260845578.1 ribosome small subunit-dependent GTPase A |
| 25(1) | WP_137062795.1 | WP_137062795.1 ribosome small subunit-dependent GTPase A |
| 25(2) | WP_061829407.1 | WP_061829407.1 ribosome small subunit-dependent GTPase A |
| 25(1) | WP_212949026.1 | WP_212949026.1 ribosome small subunit-dependent GTPase A |
| 25(1) | WP_274485145.1 | WP_274485145.1 ribosome small subunit-dependent GTPase A |
| 25(1) | WP_235193786.1 | WP_235193786.1 ribosome small subunit-dependent GTPase A |
| 25(1) | WP_274359794.1 | WP_274359794.1 ribosome small subunit-dependent GTPase A |
| 25(1) | WP_307413609.1 | WP_307413609.1 ribosome small subunit-dependent GTPase A |

| | | |
|---|---|---|
| 26(1) | WP_098150448.1 | WP_098150448.1 class I SAM-dependent methyltransferase |
| 26(1) | WP_088063224.1 | WP_088063224.1 class I SAM-dependent methyltransferase |
| 26(4) | WP_097952800.1 | WP_097952800.1 class I SAM-dependent methyltransferase |
| 26(1) | WP_098608145.1 | WP_098608145.1 class I SAM-dependent methyltransferase |
| 26(1) | WP_078419979.1 | WP_078419979.1 class I SAM-dependent methyltransferase |
| 26(2) | WP_065228751.1 | WP_065228751.1 class I SAM-dependent methyltransferase |
| 26(2) | WP_044792572.1 | WP_044792572.1 class I SAM-dependent methyltransferase |
| 26(1) | WP_000637447.1 | WP_000637447.1 class I SAM-dependent methyltransferase |
| 26(1) | WP_139888711.1 | WP_139888711.1 class I SAM-dependent methyltransferase |
| 26(1) | WP_088078453.1 | WP_088078453.1 class I SAM-dependent methyltransferase |
| 26(1) | WP_071736264.1 | WP_071736264.1 class I SAM-dependent methyltransferase |
| 26(1) | WP_074565314.1 | WP_074565314.1 class I SAM-dependent methyltransferase |
| 26(1) | WP_098856760.1 | WP_098856760.1 class I SAM-dependent methyltransferase |
| 26(6) | WP_000637461.1 | WP_000637461.1 class I SAM-dependent methyltransferase |
| 26(1) | WP_207994174.1 | WP_207994174.1 class I SAM-dependent methyltransferase |
| 26(1) | WP_098204195.1 | WP_098204195.1 class I SAM-dependent methyltransferase |
| 26(1) | WP_000637465.1 | WP_000637465.1 class I SAM-dependent methyltransferase |
| 26(1) | WP_098991584.1 | WP_098991584.1 class I SAM-dependent methyltransferase |
| 26(1) | WP_098377169.1 | WP_098377169.1 class I SAM-dependent methyltransferase |
| 26(1) | WP_071727414.1 | WP_071727414.1 class I SAM-dependent methyltransferase |
| 26(1) | WP_097821805.1 | WP_097821805.1 class I SAM-dependent methyltransferase |
| 26(1) | WP_098088263.1 | WP_098088263.1 class I SAM-dependent methyltransferase |
| 26(1) | WP_002158058.1 | WP_002158058.1 class I SAM-dependent methyltransferase |
| 26(1) | WP_088058509.1 | WP_088058509.1 class I SAM-dependent methyltransferase |
| 26(1) | WP_098077005.1 | WP_098077005.1 class I SAM-dependent methyltransferase |
| 26(1) | WP_000637469.1 | WP_000637469.1 class I SAM-dependent methyltransferase |
| 26(1) | WP_073540569.1 | WP_073540569.1 class I SAM-dependent methyltransferase |
| 26(5) | WP_000637435.1 | WP_000637435.1 class I SAM-dependent methyltransferase |
| 26(1) | WP_147786243.1 | WP_147786243.1 class I SAM-dependent methyltransferase |
| 26(1) | WP_270728486.1 | WP_270728486.1 class I SAM-dependent methyltransferase |
| 26(1) | WP_262787108.1 | WP_262787108.1 class I SAM-dependent methyltransferase |
| 26(3) | WP_262737301.1 | WP_262737301.1 class I SAM-dependent methyltransferase |
| 26(1) | WP_270691239.1 | WP_270691239.1 class I SAM-dependent methyltransferase |
| 26(3) | WP_098266636.1 | WP_098266636.1 class I SAM-dependent methyltransferase |
| 26(6) | WP_063547140.1 | WP_063547140.1 class I SAM-dependent methyltransferase |
| 26(11) | WP_000637460.1 | WP_000637460.1 class I SAM-dependent methyltransferase |
| 26(1) | WP_098288675.1 | WP_098288675.1 class I SAM-dependent methyltransferase |
| 26(2) | WP_087952365.1 | WP_087952365.1 class I SAM-dependent methyltransferase |
| 26(1) | WP_286124770.1 | WP_286124770.1 class I SAM-dependent methyltransferase |
| 26(8) | WP_000637441.1 | WP_000637441.1 class I SAM-dependent methyltransferase |
| 26(1) | WP_098776441.1 | WP_098776441.1 class I SAM-dependent methyltransferase |
| 26(1) | WP_097852781.1 | WP_097852781.1 class I SAM-dependent methyltransferase |
| 26(1) | WP_103655002.1 | WP_103655002.1 class I SAM-dependent methyltransferase |
| 26(2) | WP_098678278.1 | WP_098678278.1 class I SAM-dependent methyltransferase |
| 26(1) | WP_166704977.1 | WP_166704977.1 class I SAM-dependent methyltransferase |
| 26(2) | WP_098360828.1 | WP_098360828.1 class I SAM-dependent methyltransferase |
| 26(1) | WP_163255243.1 | WP_163255243.1 class I SAM-dependent methyltransferase |
| 26(1) | WP_000637475.1 | WP_000637475.1 class I SAM-dependent methyltransferase |
| 26(2) | WP_061184594.1 | WP_061184594.1 class I SAM-dependent methyltransferase |
| 26(1) | WP_098274926.1 | WP_098274926.1 class I SAM-dependent methyltransferase |
| 26(1) | WP_215601059.1 | WP_215601059.1 class I SAM-dependent methyltransferase |
| 26(2) | WP_087988147.1 | WP_087988147.1 class I SAM-dependent methyltransferase |
| 26(1) | WP_151630616.1 | WP_151630616.1 class I SAM-dependent methyltransferase |
| 26(5) | WP_087982606.1 | WP_087982606.1 class I SAM-dependent methyltransferase |
| 26(1) | WP_064473948.1 | WP_064473948.1 class I SAM-dependent methyltransferase |
| 26(10) | WP_000637445.1 | WP_000637445.1 class I SAM-dependent methyltransferase |
| 26(1) | WP_000637466.1 | WP_000637466.1 class I SAM-dependent methyltransferase |
| 26(1) | WP_113711527.1 | WP_113711527.1 class I SAM-dependent methyltransferase |
| 26(1) | WP_270716411.1 | WP_270716411.1 class I SAM-dependent methyltransferase |
| 26(1) | WP_147807663.1 | WP_147807663.1 class I SAM-dependent methyltransferase |
| 26(1) | WP_139867399.1 | WP_139867399.1 class I SAM-dependent methyltransferase |
| 26(1) | WP_098250516.1 | WP_098250516.1 class I SAM-dependent methyltransferase |
| 26(1) | WP_144473242.1 | WP_144473242.1 class I SAM-dependent methyltransferase |
| 26(2) | WP_002141576.1 | WP_002141576.1 class I SAM-dependent methyltransferase |
| 26(2) | WP_071679483.1 | WP_071679483.1 class I SAM-dependent methyltransferase |
| 26(1) | WP_242264965.1 | WP_242264965.1 class I SAM-dependent methyltransferase |
| 26(1) | WP_270364636.1 | WP_270364636.1 class I SAM-dependent methyltransferase |
| 26(1) | WP_106822174.1 | WP_106822174.1 class I SAM-dependent methyltransferase |
| 26(2) | WP_127812910.1 | WP_127812910.1 class I SAM-dependent methyltransferase |
| 26(1) | WP_098264276.1 | WP_098264276.1 class I SAM-dependent methyltransferase |
| 26(1) | WP_097844626.1 | WP_097844626.1 class I SAM-dependent methyltransferase |
| 26(1) | WP_241973189.1 | WP_241973189.1 class I SAM-dependent methyltransferase, partial |
| 26(10) | WP_053564641.1 | WP_053564641.1 class I SAM-dependent methyltransferase |
| 26(1) | WP_017149784.1 | WP_017149784.1 class I SAM-dependent methyltransferase |
| 26(1) | WP_098148049.1 | WP_098148049.1 class I SAM-dependent methyltransferase |
| 26(1) | WP_088609107.1 | WP_088609107.1 class I SAM-dependent methyltransferase |
| 26(1) | WP_074570402.1 | WP_074570402.1 class I SAM-dependent methyltransferase |
| 26(1) | WP_001980117.1 | WP_001980117.1 class I SAM-dependent methyltransferase |
| 26(1) | WP_215598796.1 | WP_215598796.1 class I SAM-dependent methyltransferase |
| 26(1) | WP_137014120.1 | WP_137014120.1 class I SAM-dependent methyltransferase |
| 26(1) | WP_098519056.1 | WP_098519056.1 class I SAM-dependent methyltransferase |
| 26(1) | WP_212956550.1 | WP_212956550.1 class I SAM-dependent methyltransferase |
| 26(1) | WP_088119564.1 | WP_088119564.1 class I SAM-dependent methyltransferase |
| 26(1) | WP_243500434.1 | WP_243500434.1 class I SAM-dependent methyltransferase |
| 26(3) | WP_098133697.1 | WP_098133697.1 class I SAM-dependent methyltransferase |
| 26(2) | WP_101195707.1 | WP_101195707.1 class I SAM-dependent methyltransferase |
| 26(4) | WP_000637444.1 | WP_000637444.1 class I SAM-dependent methyltransferase |
| 26(1) | WP_144611177.1 | WP_144611177.1 class I SAM-dependent methyltransferase |
| 26(1) | WP_143881063.1 | WP_143881063.1 class I SAM-dependent methyltransferase |
| 26(1) | WP_002031642.1 | WP_002031642.1 class I SAM-dependent methyltransferase |
| 26(1) | WP_289398688.1 | WP_289398688.1 class I SAM-dependent methyltransferase |
| 26(4) | WP_061680940.1 | WP_061680940.1 class I SAM-dependent methyltransferase |
| 26(1) | WP_098214779.1 | WP_098214779.1 class I SAM-dependent methyltransferase |
| 26(4) | WP_023521804.1 | WP_023521804.1 class I SAM-dependent methyltransferase |
| 26(1) | WP_063223265.1 | WP_063223265.1 class I SAM-dependent methyltransferase |

26(6) WP_000637436.1 WP_000637436.1 class I SAM-dependent methyltransferase
26(1) WP_060630637.1 WP_060630637.1 class I SAM-dependent methyltransferase

27(1) WP_307199948.1 WP_307199948.1 GntR family transcriptional regulator
27(1) WP_109625256.1 WP_109625256.1 aminotransferase class I/II-fold pyridoxal phosphate-dependent enzyme
27(1) WP_277150373.1 WP_277150373.1 LacI family DNA-binding transcriptional regulator
27(1) WP_307589590.1 WP_307589590.1 GntR family transcriptional regulator
27(1) WP_241929673.1 WP_241929673.1 GntR family transcriptional regulator
27(1) WP_010676869.1 WP_010676869.1 LacI family DNA-binding transcriptional regulator
27(1) WP_108467787.1 WP_108467787.1 GntR family transcriptional regulator
27(1) WP_165175573.1 WP_165175573.1 PLP-dependent aminotransferase family protein
27(1) WP_013309995.1 WP_013309995.1 PLP-dependent aminotransferase family protein
27(1) WP_229139906.1 WP_229139906.1 LacI family DNA-binding transcriptional regulator
27(2) WP_025848632.1 WP_025848632.1 FadR/GntR family transcriptional regulator
27(1) WP_021261897.1 WP_021261897.1 sugar ABC transporter substrate-binding protein
27(3) WP_036689469.1 WP_036689469.1 FadR/GntR family transcriptional regulator
27(1) WP_307312517.1 WP_307312517.1 LacI family DNA-binding transcriptional regulator
27(1) WP_054972104.1 WP_054972104.1 FadR/GntR family transcriptional regulator
27(2) WP_213579159.1 WP_213579159.1 GntR family transcriptional regulator
27(4) WP_163977066.1 WP_163977066.1 sugar ABC transporter substrate-binding protein
27(1) WP_077619666.1 WP_077619666.1 LacI family DNA-binding transcriptional regulator
27(1) WP_273191270.1 WP_273191270.1 GntR family transcriptional regulator
27(1) WP_000348039.1 WP_000348039.1 PLP-dependent aminotransferase family protein
27(1) WP_000348021.1 WP_000348021.1 PLP-dependent aminotransferase family protein
27(2) WP_219872452.1 WP_219872452.1 FadR/GntR family transcriptional regulator
27(1) WP_179603492.1 WP_179603492.1 LacI family DNA-binding transcriptional regulator
27(1) WP_094092612.1 WP_094092612.1 substrate-binding domain-containing protein
27(1) WP_017763059.1 WP_017763059.1 PLP-dependent aminotransferase family protein
27(1) WP_231094857.1 WP_231094857.1 aminotransferase
27(1) WP_163061527.1 WP_163061527.1 PLP-dependent aminotransferase family protein
27(1) WP_191798518.1 WP_191798518.1 GntR family transcriptional regulator
27(1) WP_216791245.1 WP_216791245.1 FadR/GntR family transcriptional regulator
27(1) WP_197937407.1 WP_197937407.1 PLP-dependent aminotransferase family protein
27(1) WP_000348042.1 WP_000348042.1 PLP-dependent aminotransferase family protein
27(1) WP_000348004.1 WP_000348004.1 PLP-dependent aminotransferase family protein
27(1) WP_039273533.1 WP_039273533.1 aminotransferase class I/II-fold pyridoxal phosphate-dependent enzyme
27(1) WP_273383895.1 WP_273383895.1 GntR family transcriptional regulator
27(1) WP_264453870.1 WP_264453870.1 aminotransferase class I/II-fold pyridoxal phosphate-dependent enzyme
27(1) WP_025685040.1 WP_025685040.1 aminotransferase class I/II-fold pyridoxal phosphate-dependent enzyme
27(1) WP_098284009.1 WP_098284009.1 PLP-dependent aminotransferase family protein
27(1) WP_039273535.1 WP_039273535.1 PLP-dependent aminotransferase family protein
27(1) WP_229139914.1 WP_229139914.1 LacI family DNA-binding transcriptional regulator
27(1) WP_307349984.1 WP_307349984.1 aminotransferase class I/II-fold pyridoxal phosphate-dependent enzyme
27(2) WP_219872446.1 WP_219872446.1 LacI family DNA-binding transcriptional regulator
27(1) WP_065223572.1 WP_065223572.1 PLP-dependent aminotransferase family protein
27(1) WP_274485157.1 WP_274485157.1 aminotransferase class I/II-fold pyridoxal phosphate-dependent enzyme
27(1) WP_254349530.1 WP_254349530.1 aminotransferase class I/II-fold pyridoxal phosphate-dependent enzyme
27(1) WP_098874958.1 WP_098874958.1 PLP-dependent aminotransferase family protein
27(1) WP_048535509.1 WP_048535509.1 PLP-dependent aminotransferase family protein
27(2) WP_254348142.1 WP_254348142.1 PLP-dependent aminotransferase family protein
27(2) WP_127459448.1 WP_127459448.1 FadR/GntR family transcriptional regulator
27(2) WP_006923789.1 WP_006923789.1 PLP-dependent aminotransferase family protein
27(1) WP_098013431.1 WP_098013431.1 PLP-dependent aminotransferase family protein
27(1) WP_305022912.1 WP_305022912.1 substrate-binding domain-containing protein
27(1) WP_071740424.1 WP_071740424.1 PLP-dependent aminotransferase family protein
27(1) WP_077619657.1 WP_077619657.1 LacI family DNA-binding transcriptional regulator
27(1) WP_285748166.1 WP_285748166.1 GntR family transcriptional regulator
27(1) WP_000348028.1 WP_000348028.1 PLP-dependent aminotransferase family protein
27(1) WP_053513139.1 WP_053513139.1 PLP-dependent aminotransferase family protein
27(2) WP_047386013.1 WP_047386013.1 PLP-dependent aminotransferase family protein
27(1) WP_070179254.1 WP_070179254.1 PLP-dependent aminotransferase family protein
27(1) WP_089965958.1 WP_089965958.1 LacI family DNA-binding transcriptional regulator
27(1) WP_053325284.1 WP_053325284.1 aminotransferase class I/II-fold pyridoxal phosphate-dependent enzyme
27(1) WP_106769789.1 WP_106769789.1 GntR family transcriptional regulator
27(1) WP_044037304.1 WP_044037304.1 threonine-phosphate decarboxylase CobD
27(1) WP_086410428.1 WP_086410428.1 PLP-dependent aminotransferase family protein
27(1) WP_001990271.1 WP_001990271.1 PLP-dependent aminotransferase family protein
27(1) WP_065291260.1 WP_065291260.1 sugar ABC transporter substrate-binding protein
27(1) WP_098287135.1 WP_098287135.1 PLP-dependent aminotransferase family protein
27(1) WP_254348145.1 WP_254348145.1 aminotransferase class I/II-fold pyridoxal phosphate-dependent enzyme
27(1) WP_290102339.1 WP_290102339.1 PLP-dependent aminotransferase family protein
27(1) WP_099354897.1 WP_099354897.1 LacI family DNA-binding transcriptional regulator
27(1) WP_134902879.1 WP_134902879.1 PLP-dependent aminotransferase family protein
27(1) WP_274485156.1 WP_274485156.1 PLP-dependent aminotransferase family protein
27(1) WP_136379686.1 WP_136379686.1 LacI family DNA-binding transcriptional regulator
27(1) WP_307349982.1 WP_307349982.1 aminotransferase class I/II-fold pyridoxal phosphate-dependent enzyme
27(1) WP_249897303.1 WP_249897303.1 GntR family transcriptional regulator
27(1) WP_098139275.1 WP_098139275.1 PLP-dependent aminotransferase family protein
27(1) WP_155703950.1 WP_155703950.1 GntR family transcriptional regulator
27(1) WP_192571320.1 WP_192571320.1 GntR family transcriptional regulator
27(1) WP_306077404.1 WP_306077404.1 LacI family DNA-binding transcriptional regulator
27(1) WP_010346277.1 WP_010346277.1 PLP-dependent aminotransferase family protein
27(1) WP_068940708.1 WP_068940708.1 aminotransferase class I/II-fold pyridoxal phosphate-dependent enzyme
27(1) WP_102371685.1 WP_102371685.1 aminotransferase
27(1) WP_290407034.1 WP_290407034.1 PLP-dependent aminotransferase family protein
27(1) WP_047956479.1 WP_047956479.1 PLP-dependent aminotransferase family protein
27(1) WP_077893587.1 WP_077893587.1 LacI family DNA-binding transcriptional regulator
27(1) WP_000348031.1 WP_000348031.1 PLP-dependent aminotransferase family protein
27(1) WP_138186707.1 WP_138186707.1 sugar ABC transporter substrate-binding protein
27(1) WP_281937348.1 WP_281937348.1 FadR/GntR family transcriptional regulator
27(1) WP_025685038.1 WP_025685038.1 PLP-dependent aminotransferase family protein
27(1) WP_253411100.1 WP_253411100.1 FadR/GntR family transcriptional regulator
27(1) WP_148450933.1 WP_148450933.1 PLP-dependent aminotransferase family protein

27(1) WP_103047823.1 WP_103047823.1 aminotransferase class I/II-fold pyridoxal phosphate-dependent enzyme
27(1) WP_304216859.1 WP_304216859.1 LacI family DNA-binding transcriptional regulator
27(1) WP_053325282.1 WP_053325282.1 PLP-dependent aminotransferase family protein
27(1) WP_090674349.1 WP_090674349.1 FadR/GntR family transcriptional regulator
27(1) WP_071558837.1 WP_071558837.1 PLP-dependent aminotransferase family protein
27(1) WP_127484331.1 WP_127484331.1 FadR/GntR family transcriptional regulator
27(1) WP_015736239.1 WP_015736239.1 GntR family transcriptional regulator
27(1) WP_148409458.1 WP_148409458.1 aminotransferase class I/II-fold pyridoxal phosphate-dependent enzyme
27(1) WP_250244825.1 WP_250244825.1 aminotransferase class I/II-fold pyridoxal phosphate-dependent enzyme
27(1) WP_153589153.1 WP_153589153.1 PLP-dependent aminotransferase family protein
27(1) WP_102710707.1 WP_102710707.1 LacI family DNA-binding transcriptional regulator
27(2) WP_086385914.1 WP_086385914.1 PLP-dependent aminotransferase family protein
27(1) WP_103047825.1 WP_103047825.1 PLP-dependent aminotransferase family protein
27(1) WP_213532648.1 WP_213532648.1 GntR family transcriptional regulator
27(1) WP_257675222.1 WP_257675222.1 LacI family DNA-binding transcriptional regulator
27(1) WP_189031062.1 WP_189031062.1 MalY/PatB family protein
27(1) WP_250244824.1 WP_250244824.1 PLP-dependent aminotransferase family protein
27(1) WP_143315256.1 WP_143315256.1 GntR family transcriptional regulator
27(1) WP_077833118.1 WP_077833118.1 LacI family DNA-binding transcriptional regulator
27(1) WP_263324540.1 WP_263324540.1 LacI family DNA-binding transcriptional regulator
27(1) WP_116353266.1 WP_116353266.1 LacI family DNA-binding transcriptional regulator
27(1) WP_098226399.1 WP_098226399.1 PLP-dependent aminotransferase family protein
27(2) WP_063188032.1 WP_063188032.1 FadR/GntR family transcriptional regulator
27(1) WP_220027688.1 WP_220027688.1 PLP-dependent aminotransferase family protein
27(1) WP_016512968.1 WP_016512968.1 PLP-dependent aminotransferase family protein
27(1) WP_099093068.1 WP_099093068.1 substrate-binding domain-containing protein
27(1) WP_099361333.1 WP_099361333.1 LacI family DNA-binding transcriptional regulator
27(2) WP_044355253.1 WP_044355253.1 sugar ABC transporter substrate-binding protein
27(4) WP_005546481.1 WP_005546481.1 sugar ABC transporter substrate-binding protein
27(1) WP_010346274.1 WP_010346274.1 aminotransferase class I/II-fold pyridoxal phosphate-dependent enzyme
27(1) WP_309098608.1 WP_309098608.1 LacI family DNA-binding transcriptional regulator
27(1) WP_136379685.1 WP_136379685.1 LacI family DNA-binding transcriptional regulator
27(1) WP_010496623.1 WP_010496623.1 FadR/GntR family transcriptional regulator
27(1) WP_134902880.1 WP_134902880.1 aminotransferase class I/II-fold pyridoxal phosphate-dependent enzyme
27(2) WP_186326458.1 WP_186326458.1 GntR family transcriptional regulator
27(1) WP_157411782.1 WP_157411782.1 LacI family DNA-binding transcriptional regulator
27(1) WP_268623235.1 WP_268623235.1 sugar ABC transporter substrate-binding protein
27(1) WP_254349529.1 WP_254349529.1 PLP-dependent aminotransferase family protein
27(1) WP_303942556.1 WP_303942556.1 GntR family transcriptional regulator
27(1) WP_223592697.1 WP_223592697.1 GntR family transcriptional regulator
27(1) WP_086405915.1 WP_086405915.1 PLP-dependent aminotransferase family protein
27(1) WP_098955456.1 WP_098955456.1 PLP-dependent aminotransferase family protein
27(1) WP_163095421.1 WP_163095421.1 PLP-dependent aminotransferase family protein
27(1) WP_254759285.1 WP_254759285.1 FadR/GntR family transcriptional regulator
27(1) WP_030025419.1 WP_030025419.1 PLP-dependent aminotransferase family protein
27(1) WP_098567746.1 WP_098567746.1 PLP-dependent aminotransferase family protein
27(1) WP_066020850.1 WP_066020850.1 LacI family DNA-binding transcriptional regulator
27(1) WP_069081266.1 WP_069081266.1 PLP-dependent aminotransferase family protein
27(1) WP_087967970.1 WP_087967970.1 PLP-dependent aminotransferase family protein
27(2) WP_000348037.1 WP_000348037.1 PLP-dependent aminotransferase family protein
27(1) WP_165175572.1 WP_165175572.1 aminotransferase class I/II-fold pyridoxal phosphate-dependent enzyme
27(1) WP_061667216.1 WP_061667216.1 PLP-dependent aminotransferase family protein
27(5) WP_021253132.1 WP_021253132.1 sugar ABC transporter substrate-binding protein
27(1) WP_077852424.1 WP_077852424.1 LacI family DNA-binding transcriptional regulator
27(3) WP_000348003.1 WP_000348003.1 PLP-dependent aminotransferase family protein
27(1) WP_069081268.1 WP_069081268.1 aminotransferase class I/II-fold pyridoxal phosphate-dependent enzyme
27(1) WP_010963715.1 WP_010963715.1 LacI family DNA-binding transcriptional regulator
27(2) WP_036671562.1 WP_036671562.1 GntR family transcriptional regulator
27(1) WP_116353258.1 WP_116353258.1 LacI family DNA-binding transcriptional regulator

28(1) WP_251674657.1 WP_251674657.1 ATP-dependent helicase HrpB
28(1) WP_169505059.1 WP_169505059.1 ATP-dependent helicase HrpB
28(1) WP_263880214.1 WP_263880214.1 ATP-dependent helicase HrpB
28(2) WP_076249772.1 WP_076249772.1 ATP-dependent helicase HrpB
28(1) WP_179265137.1 WP_179265137.1 ATP-dependent helicase HrpB
28(1) WP_018883264.1 WP_018883264.1 ATP-dependent helicase HrpB
28(1) WP_106765950.1 WP_106765950.1 ATP-dependent helicase HrpB
28(1) WP_305022916.1 WP_305022916.1 ATP-dependent helicase HrpB
28(1) WP_017689066.1 WP_017689066.1 ATP-dependent helicase HrpB
28(1) WP_264929202.1 WP_264929202.1 ATP-dependent helicase HrpB
28(1) WP_210095104.1 WP_210095104.1 ATP-dependent helicase HrpB
28(1) WP_215153846.1 WP_215153846.1 ATP-dependent helicase HrpB
28(1) WP_113058722.1 WP_113058722.1 ATP-dependent helicase HrpB
28(1) WP_076311045.1 WP_076311045.1 ATP-dependent helicase HrpB
28(1) WP_127199778.1 WP_127199778.1 ATP-dependent helicase HrpB
28(1) WP_036609192.1 WP_036609192.1 ATP-dependent helicase HrpB
28(1) WP_145318682.1 WP_145318682.1 ATP-dependent helicase HrpB
28(1) WP_090949170.1 WP_090949170.1 ATP-dependent helicase HrpB
28(1) WP_136604823.1 WP_136604823.1 ATP-dependent helicase HrpB
28(1) WP_213532662.1 WP_213532662.1 ATP-dependent helicase HrpB
28(1) WP_272035333.1 WP_272035333.1 ATP-dependent helicase HrpB
28(1) WP_062833885.1 WP_062833885.1 ATP-dependent helicase HrpB
28(1) WP_213616459.1 WP_213616459.1 ATP-dependent helicase HrpB
28(1) WP_058710498.1 WP_058710498.1 ATP-dependent helicase HrpB
28(1) WP_192266468.1 WP_192266468.1 ATP-dependent helicase HrpB
28(1) WP_077567011.1 WP_077567011.1 ATP-dependent helicase HrpB
28(1) WP_301879142.1 WP_301879142.1 ATP-dependent helicase HrpB
28(1) WP_209861633.1 WP_209861633.1 ATP-dependent helicase HrpB
28(1) WP_213579155.1 WP_213579155.1 ATP-dependent helicase HrpB
28(1) WP_054958416.1 WP_054958416.1 ATP-dependent helicase HrpB
28(2) WP_047909607.1 WP_047909607.1 ATP-dependent helicase HrpB
28(1) WP_076167852.1 WP_076167852.1 ATP-dependent helicase HrpB
28(1) WP_290409177.1 WP_290409177.1 ATP-dependent helicase HrpB
28(1) WP_280753898.1 WP_280753898.1 ATP-dependent helicase HrpB
28(1) WP_090908284.1 WP_090908284.1 ATP-dependent helicase HrpB
28(2) WP_145156810.1 WP_145156810.1 ATP-dependent helicase HrpB
28(1) WP_064639009.1 WP_064639009.1 ATP-dependent helicase HrpB

| | | |
|---|---|---|
| 28(1) | WP_307537573.1 | WP_307537573.1 ATP-dependent helicase HrpB |
| 28(1) | WP_290403372.1 | WP_290403372.1 ATP-dependent helicase HrpB |
| 28(1) | WP_247902249.1 | WP_247902249.1 ATP-dependent helicase HrpB |
| 28(1) | WP_212983259.1 | WP_212983259.1 ATP-dependent helicase HrpB |
| 28(1) | WP_062326153.1 | WP_062326153.1 ATP-dependent helicase HrpB |
| 28(1) | WP_142613010.1 | WP_142613010.1 ATP-dependent helicase HrpB |
| 28(1) | WP_160496923.1 | WP_160496923.1 ATP-dependent helicase HrpB |
| 28(1) | WP_090921044.1 | WP_090921044.1 ATP-dependent helicase HrpB |
| 28(1) | WP_074094619.1 | WP_074094619.1 ATP-dependent helicase HrpB |
| 28(1) | WP_119847557.1 | WP_119847557.1 ATP-dependent helicase HrpB |
| 28(1) | WP_100529435.1 | WP_100529435.1 ATP-dependent helicase HrpB |
| 28(1) | WP_251537233.1 | WP_251537233.1 ATP-dependent helicase HrpB |
| 28(1) | WP_096774449.1 | WP_096774449.1 ATP-dependent helicase HrpB |
| 28(1) | WP_016821040.1 | WP_016821040.1 ATP-dependent helicase HrpB |
| 28(1) | WP_095287655.1 | WP_095287655.1 ATP-dependent helicase HrpB |
| 28(1) | WP_289392076.1 | WP_289392076.1 ATP-dependent helicase HrpB |
| 28(1) | WP_191798533.1 | WP_191798533.1 ATP-dependent helicase HrpB |
| 28(1) | WP_133384359.1 | WP_133384359.1 ATP-dependent helicase HrpB |
| 28(1) | WP_123797119.1 | WP_123797119.1 ATP-dependent helicase HrpB |
| 28(1) | WP_063480516.1 | WP_063480516.1 ATP-dependent helicase HrpB |
| 28(1) | WP_236703656.1 | WP_236703656.1 ATP-dependent helicase HrpB, partial |
| 28(1) | WP_305755530.1 | WP_305755530.1 ATP-dependent helicase HrpB |
| 28(1) | WP_127592421.1 | WP_127592421.1 ATP-dependent helicase HrpB |
| 28(1) | WP_293202258.1 | WP_293202258.1 ATP-dependent helicase HrpB |
| 28(1) | WP_145046463.1 | WP_145046463.1 ATP-dependent helicase HrpB |
| 28(1) | WP_148246953.1 | WP_148246953.1 ATP-dependent helicase HrpB |
| 28(1) | WP_076322107.1 | WP_076322107.1 ATP-dependent helicase HrpB |
| 28(1) | WP_094092628.1 | WP_094092628.1 ATP-dependent helicase HrpB |
| 28(1) | WP_036670440.1 | WP_036670440.1 ATP-dependent helicase HrpB |
| 28(1) | WP_188593166.1 | WP_188593166.1 ATP-dependent helicase HrpB |
| 28(1) | WP_127543693.1 | WP_127543693.1 ATP-dependent helicase HrpB |
| 28(2) | WP_036642474.1 | WP_036642474.1 ATP-dependent helicase HrpB |
| 28(1) | WP_127536192.1 | WP_127536192.1 ATP-dependent helicase HrpB |
| 28(1) | WP_056700801.1 | WP_056700801.1 ATP-dependent helicase HrpB |
| 28(1) | WP_145329912.1 | WP_145329912.1 ATP-dependent helicase HrpB |
| 28(1) | WP_150367639.1 | WP_150367639.1 ATP-dependent helicase HrpB |
| 28(1) | WP_208919052.1 | WP_208919052.1 ATP-dependent helicase HrpB |
| 28(1) | WP_145949003.1 | WP_145949003.1 ATP-dependent helicase HrpB |
| 28(1) | WP_282321817.1 | WP_282321817.1 ATP-dependent helicase HrpB |
| 28(1) | WP_071220211.1 | WP_071220211.1 ATP-dependent helicase HrpB |
| 28(1) | WP_274566442.1 | WP_274566442.1 ATP-dependent helicase HrpB |
| 28(1) | WP_095266204.1 | WP_095266204.1 ATP-dependent helicase HrpB |
| 28(1) | WP_076330726.1 | WP_076330726.1 ATP-dependent helicase HrpB |
| 28(1) | WP_213649963.1 | WP_213649963.1 ATP-dependent helicase HrpB |
| 28(1) | WP_223067431.1 | WP_223067431.1 ATP-dependent helicase HrpB |
| 28(1) | WP_155618099.1 | WP_155618099.1 ATP-dependent helicase HrpB |
| 28(1) | WP_123067120.1 | WP_123067120.1 ATP-dependent helicase HrpB |
| 28(1) | WP_145408776.1 | WP_145408776.1 ATP-dependent helicase HrpB |
| 28(1) | WP_072732443.1 | WP_072732443.1 ATP-dependent helicase HrpB |
| 28(1) | WP_192571324.1 | WP_192571324.1 ATP-dependent helicase HrpB |
| 28(1) | WP_109999896.1 | WP_109999896.1 ATP-dependent helicase HrpB |
| 28(1) | WP_044785617.1 | WP_044785617.1 ATP-dependent helicase HrpB |

| | | |
|---|---|---|
| 28(1) | WP_194543493.1 | WP_194543493.1 ATP-dependent helicase HrpB |
| 28(1) | WP_024630332.1 | WP_024630332.1 ATP-dependent helicase HrpB |
| 28(1) | WP_009590720.1 | WP_009590720.1 ATP-dependent helicase HrpB |
| 28(1) | WP_260065266.1 | WP_260065266.1 ATP-dependent helicase HrpB |
| 28(1) | WP_090993631.1 | WP_090993631.1 ATP-dependent helicase HrpB |
| 28(1) | WP_241929666.1 | WP_241929666.1 ATP-dependent helicase HrpB |
| 28(1) | WP_076208837.1 | WP_076208837.1 ATP-dependent helicase HrpB |
| 28(1) | WP_285748187.1 | WP_285748187.1 ATP-dependent helicase HrpB |
| 28(2) | WP_107734203.1 | WP_107734203.1 ATP-dependent helicase HrpB |
| 28(1) | WP_213644234.1 | WP_213644234.1 ATP-dependent helicase HrpB |
| 28(1) | WP_194230538.1 | WP_194230538.1 ATP-dependent helicase HrpB |
| 28(1) | WP_216531937.1 | WP_216531937.1 ATP-dependent helicase HrpB |
| 28(1) | WP_098746849.1 | WP_098746849.1 ATP-dependent helicase HrpB |
| 28(1) | WP_244719664.1 | WP_244719664.1 ATP-dependent helicase HrpB |
| 28(1) | WP_054402901.1 | WP_054402901.1 ATP-dependent helicase HrpB |
| 28(1) | WP_234414397.1 | WP_234414397.1 ATP-dependent helicase HrpB |
| 28(1) | WP_045243834.1 | WP_045243834.1 ATP-dependent helicase HrpB |
| 28(1) | WP_213618820.1 | WP_213618820.1 ATP-dependent helicase HrpB |
| 28(1) | WP_094936490.1 | WP_094936490.1 ATP-dependent helicase HrpB |
| 28(1) | WP_160037494.1 | WP_160037494.1 ATP-dependent helicase HrpB |
| 28(1) | WP_251375543.1 | WP_251375543.1 ATP-dependent helicase HrpB |
| 28(1) | WP_152826370.1 | WP_152826370.1 ATP-dependent helicase HrpB |
| 28(1) | WP_152402386.1 | WP_152402386.1 ATP-dependent helicase HrpB |
| 28(1) | WP_090805480.1 | WP_090805480.1 ATP-dependent helicase HrpB |
| 28(1) | WP_028589288.1 | WP_028589288.1 ATP-dependent helicase HrpB |
| 28(1) | WP_145021726.1 | WP_145021726.1 ATP-dependent helicase HrpB |
| 28(1) | WP_040101524.1 | WP_040101524.1 ATP-dependent helicase HrpB |
| 28(1) | WP_251583301.1 | WP_251583301.1 ATP-dependent helicase HrpB |
| 28(1) | WP_307214309.1 | WP_307214309.1 ATP-dependent helicase HrpB |
| 28(1) | WP_269084577.1 | WP_269084577.1 hypothetical protein, partial |
| 28(1) | WP_153978070.1 | WP_153978070.1 ATP-dependent helicase HrpB |
| 28(1) | WP_212969033.1 | WP_212969033.1 ATP-dependent helicase HrpB |
| 28(1) | WP_145146966.1 | WP_145146966.1 ATP-dependent helicase HrpB |
| 28(1) | WP_110822558.1 | WP_110822558.1 ATP-dependent helicase HrpB |
| 28(1) | WP_100537969.1 | WP_100537969.1 ATP-dependent helicase HrpB |
| 28(1) | WP_301249702.1 | WP_301249702.1 ATP-dependent helicase HrpB |
| 28(1) | WP_110892960.1 | WP_110892960.1 ATP-dependent helicase HrpB |
| 28(1) | WP_090896202.1 | WP_090896202.1 ATP-dependent helicase HrpB |
| 28(1) | WP_236417427.1 | WP_236417427.1 ATP-dependent helicase HrpB |
| 28(1) | WP_183578473.1 | WP_183578473.1 ATP-dependent helicase HrpB |
| 28(1) | WP_211745823.1 | WP_211745823.1 ATP-dependent helicase HrpB |
| 28(1) | WP_068694724.1 | WP_068694724.1 ATP-dependent helicase HrpB |
| 28(1) | WP_138226019.1 | WP_138226019.1 ATP-dependent helicase HrpB |
| 28(1) | WP_263560237.1 | WP_263560237.1 ATP-dependent helicase HrpB |
| 28(1) | WP_212980183.1 | WP_212980183.1 ATP-dependent helicase HrpB |
| 28(1) | WP_124114123.1 | WP_124114123.1 ATP-dependent helicase HrpB |
| 28(1) | WP_091017188.1 | WP_091017188.1 ATP-dependent helicase HrpB |
| 28(1) | WP_145921941.1 | WP_145921941.1 helicase-related protein, partial |
| 28(1) | WP_145134811.1 | WP_145134811.1 ATP-dependent helicase HrpB |
| 28(1) | WP_285079437.1 | WP_285079437.1 ATP-dependent helicase HrpB |
| 28(1) | WP_307413595.1 | WP_307413595.1 ATP-dependent helicase HrpB |
| 28(1) | WP_015736235.1 | WP_015736235.1 ATP-dependent helicase HrpB |

28(1) WP_145037682.1 WP_145037682.1 ATP-dependent helicase HrpB
28(1) WP_090887116.1 WP_090887116.1 ATP-dependent helicase HrpB
28(1) WP_047842540.1 WP_047842540.1 ATP-dependent helicase HrpB
28(1) WP_277746353.1 WP_277746353.1 ATP-dependent helicase HrpB
28(1) WP_111619118.1 WP_111619118.1 ATP-dependent helicase HrpB
28(1) WP_251497181.1 WP_251497181.1 ATP-dependent helicase HrpB
28(1) WP_208203486.1 WP_208203486.1 ATP-dependent helicase HrpB
28(1) WP_094593881.1 WP_094593881.1 ATP-dependent helicase HrpB
28(1) WP_099853192.1 WP_099853192.1 ATP-dependent helicase HrpB
28(1) WP_289373788.1 WP_289373788.1 ATP-dependent helicase HrpB
28(1) WP_212949036.1 WP_212949036.1 ATP-dependent helicase HrpB
28(1) WP_095398796.1 WP_095398796.1 ATP-dependent helicase HrpB
28(1) WP_068619098.1 WP_068619098.1 ATP-dependent helicase HrpB
28(1) WP_091155394.1 WP_091155394.1 ATP-dependent helicase HrpB
28(1) WP_220684645.1 WP_220684645.1 ATP-dependent helicase HrpB
28(1) WP_090722896.1 WP_090722896.1 ATP-dependent helicase HrpB
28(1) WP_095358544.1 WP_095358544.1 ATP-dependent helicase HrpB
28(1) WP_076318553.1 WP_076318553.1 ATP-dependent helicase HrpB
28(1) WP_116231170.1 WP_116231170.1 ATP-dependent helicase HrpB
28(1) WP_079694643.1 WP_079694643.1 ATP-dependent helicase HrpB
28(1) WP_308721984.1 WP_308721984.1 ATP-dependent helicase HrpB
28(1) WP_113052561.1 WP_113052561.1 ATP-dependent helicase HrpB
28(1) WP_145412708.1 WP_145412708.1 ATP-dependent helicase HrpB
28(1) WP_089546604.1 WP_089546604.1 ATP-dependent helicase HrpB
28(1) WP_163872777.1 WP_163872777.1 ATP-dependent helicase HrpB

29(6) WP_017689086.1 WP_017689086.1 HesB/YadR/YfhF family protein
29(2) WP_152381388.1 WP_152381388.1 HesB/YadR/YfhF family protein
29(2) WP_095265648.1 WP_095265648.1 HesB/YadR/YfhF family protein
29(1) WP_192266431.1 WP_192266431.1 HesB/YadR/YfhF family protein
29(1) WP_064798065.1 WP_064798065.1 HesB/YadR/YfhF family protein
29(1) WP_123067104.1 WP_123067104.1 HesB/YadR/YfhF family protein
29(1) WP_154886324.1 WP_154886324.1 HesB/YadR/YfhF family protein
29(1) WP_095287672.1 WP_095287672.1 HesB/YadR/YfhF family protein
29(1) WP_137062790.1 WP_137062790.1 HesB/YadR/YfhF family protein
29(20) WP_016821022.1 WP_016821022.1 HesB/YadR/YfhF family protein
29(1) WP_110822566.1 WP_110822566.1 HesB/YadR/YfhF family protein
29(1) WP_025716865.1 WP_025716865.1 HesB/YadR/YfhF family protein
29(1) WP_250244801.1 WP_250244801.1 HesB/YadR/YfhF family protein
29(1) WP_160037472.1 WP_160037472.1 HesB/YadR/YfhF family protein
29(1) WP_062326136.1 WP_062326136.1 hypothetical protein
29(1) WP_236417196.1 WP_236417196.1 HesB/YadR/YfhF family protein
29(1) WP_194230543.1 WP_194230543.1 hypothetical protein
29(1) WP_269055429.1 WP_269055429.1 HesB/YadR/YfhF family protein
29(1) WP_010346296.1 WP_010346296.1 HesB/YadR/YfhF family protein
29(1) WP_036707077.1 WP_036707077.1 HesB/YadR/YfhF family protein
29(1) WP_209861667.1 WP_209861667.1 hypothetical protein
29(1) WP_108467785.1 WP_108467785.1 HesB/YadR/YfhF family protein
29(1) WP_293202273.1 WP_293202273.1 HesB/YadR/YfhF family protein
29(6) WP_014281172.1 WP_014281172.1 HesB/YadR/YfhF family protein
29(2) WP_184180381.1 WP_184180381.1 HesB/YadR/YfhF family protein
29(1) WP_250247160.1 WP_250247160.1 HesB/YadR/YfhF family protein
29(1) WP_160496947.1 WP_160496947.1 hypothetical protein
29(1) WP_028589303.1 WP_028589303.1 HesB/YadR/YfhF family protein
29(1) WP_251378708.1 WP_251378708.1 HesB/YadR/YfhF family protein
29(1) WP_145412698.1 WP_145412698.1 HesB/YadR/YfhF family protein
29(3) WP_095358555.1 WP_095358555.1 HesB/YadR/YfhF family protein
29(1) WP_165079987.1 WP_165079987.1 HesB/YadR/YfhF family protein
29(12) WP_013309976.1 WP_013309976.1 HesB/YadR/YfhF family protein
29(21) WP_024630313.1 WP_024630313.1 HesB/YadR/YfhF family protein
29(7) WP_068940673.1 WP_068940673.1 HesB/YadR/YfhF family protein
29(1) WP_305755045.1 WP_305755045.1 HesB/YadR/YfhF family protein
29(1) WP_023957687.1 WP_023957687.1 HesB/YadR/YfhF family protein
29(2) WP_138226034.1 WP_138226034.1 HesB/YadR/YfhF family protein
29(3) WP_013370773.1 WP_013370773.1 HesB/YadR/YfhF family protein
29(2) WP_007430120.1 WP_007430120.1 HesB/YadR/YfhF family protein
29(1) WP_104495332.1 WP_104495332.1 HesB/YadR/YfhF family protein
29(1) WP_018883288.1 WP_018883288.1 HesB/YadR/YfhF family protein
29(5) WP_061829410.1 WP_061829410.1 HesB/YadR/YfhF family protein
29(1) WP_220027675.1 WP_220027675.1 HesB/YadR/YfhF family protein
29(1) WP_127536202.1 WP_127536202.1 HesB/YadR/YfhF family protein
29(2) WP_145046436.1 WP_145046436.1 HesB/YadR/YfhF family protein
29(1) WP_110892996.1 WP_110892996.1 HesB/YadR/YfhF family protein
29(2) WP_025719075.1 WP_025719075.1 HesB/YadR/YfhF family protein
29(2) WP_128101475.1 WP_128101475.1 HesB/YadR/YfhF family protein
29(1) WP_212980229.1 WP_212980229.1 hypothetical protein
29(1) WP_090921080.1 WP_090921080.1 HesB/YadR/YfhF family protein
29(1) WP_094155413.1 WP_094155413.1 HesB/YadR/YfhF family protein
29(1) WP_111268952.1 WP_111268952.1 HesB/YadR/YfhF family protein
29(1) WP_169505042.1 WP_169505042.1 HesB/YadR/YfhF family protein
29(1) WP_145329887.1 WP_145329887.1 HesB/YadR/YfhF family protein
29(1) WP_274485141.1 WP_274485141.1 HesB/YadR/YfhF family protein
29(1) WP_305022936.1 WP_305022936.1 HesB/YadR/YfhF family protein
29(4) WP_039273555.1 WP_039273555.1 HesB/YadR/YfhF family protein
29(1) WP_244719554.1 WP_244719554.1 hypothetical protein
29(1) WP_272035282.1 WP_272035282.1 HesB/YadR/YfhF family protein
29(1) WP_307413614.1 WP_307413614.1 HesB/YadR/YfhF family protein
29(1) WP_136604801.1 WP_136604801.1 hypothetical protein
29(1) WP_137078361.1 WP_137078361.1 HesB/YadR/YfhF family protein
29(17) WP_036609218.1 WP_036609218.1 HesB/YadR/YfhF family protein
29(1) WP_056700762.1 WP_056700762.1 HesB/YadR/YfhF family protein
29(1) WP_229522866.1 WP_229522866.1 HesB/YadR/YfhF family protein
29(2) WP_063208796.1 WP_063208796.1 HesB/YadR/YfhF family protein
29(1) WP_213649982.1 WP_213649982.1 hypothetical protein

30(31) WP_000063712.1 WP_000063712.1 BrxA/BrxB family bacilliredoxin
30(2) WP_098360827.1 WP_098360827.1 BrxA/BrxB family bacilliredoxin
30(1) WP_087995253.1 WP_087995253.1 BrxA/BrxB family bacilliredoxin
30(1) WP_199695245.1 WP_199695245.1 BrxA/BrxB family bacilliredoxin
30(1) WP_098255412.1 WP_098255412.1 BrxA/BrxB family bacilliredoxin
30(45) WP_000063706.1 WP_000063706.1 BrxA/BrxB family bacilliredoxin
30(21) WP_000063705.1 WP_000063705.1 BrxA/BrxB family bacilliredoxin
30(4) WP_088609106.1 WP_088609106.1 BrxA/BrxB family bacilliredoxin

| | | |
|---|---|---|
| 30(1) | WP_088114875.1 | WP_088114875.1 BrxA/BrxB family bacilliredoxin |
| 30(42) | WP_000063713.1 | WP_000063713.1 BrxA/BrxB family bacilliredoxin |
| 30(2) | WP_017149785.1 | WP_017149785.1 BrxA/BrxB family bacilliredoxin |
| 30(2) | WP_098264275.1 | WP_098264275.1 BrxA/BrxB family bacilliredoxin |
| 30(4) | WP_023521803.1 | WP_023521803.1 BrxA/BrxB family bacilliredoxin |
| 30(1) | WP_000063710.1 | WP_000063710.1 BrxA/BrxB family bacilliredoxin |
| 30(6) | WP_000063703.1 | WP_000063703.1 BrxA/BrxB family bacilliredoxin |
| 30(1) | WP_243500432.1 | WP_243500432.1 BrxA/BrxB family bacilliredoxin |
| 30(1) | WP_262748116.1 | WP_262748116.1 BrxA/BrxB family bacilliredoxin |
| 31(7) | WP_019687026.1 | WP_019687026.1 L,D-transpeptidase |
| 31(4) | WP_062833875.1 | WP_062833875.1 L,D-transpeptidase |
| 31(1) | WP_254325796.1 | WP_254325796.1 L,D-transpeptidase |
| 31(1) | WP_091017202.1 | WP_091017202.1 L,D-transpeptidase |
| 31(1) | WP_062326142.1 | WP_062326142.1 L,D-transpeptidase |
| 31(1) | WP_250251635.1 | WP_250251635.1 L,D-transpeptidase |
| 31(1) | WP_145318713.1 | WP_145318713.1 L,D-transpeptidase |
| 31(1) | WP_100529430.1 | WP_100529430.1 L,D-transpeptidase |
| 31(1) | WP_053783881.1 | WP_053783881.1 L,D-transpeptidase |
| 31(1) | WP_025685028.1 | WP_025685028.1 L,D-transpeptidase |
| 31(1) | WP_264929213.1 | WP_264929213.1 L,D-transpeptidase |
| 31(1) | WP_220027678.1 | WP_220027678.1 L,D-transpeptidase |
| 31(1) | WP_191798514.1 | WP_191798514.1 L,D-transpeptidase |
| 31(1) | WP_145402730.1 | WP_145402730.1 L,D-transpeptidase |
| 31(2) | WP_029517059.1 | WP_029517059.1 L,D-transpeptidase |
| 31(5) | WP_076249779.1 | WP_076249779.1 L,D-transpeptidase |
| 31(1) | WP_305022929.1 | WP_305022929.1 L,D-transpeptidase |
| 31(9) | WP_013370779.1 | WP_013370779.1 L,D-transpeptidase |
| 31(1) | WP_127536199.1 | WP_127536199.1 L,D-transpeptidase |
| 31(1) | WP_213532645.1 | WP_213532645.1 L,D-transpeptidase |
| 31(1) | WP_250244811.1 | WP_250244811.1 L,D-transpeptidase |
| 31(1) | WP_272035294.1 | WP_272035294.1 L,D-transpeptidase |
| 31(1) | WP_090993609.1 | WP_090993609.1 L,D-transpeptidase |
| 31(2) | WP_063208783.1 | WP_063208783.1 L,D-transpeptidase |
| 31(1) | WP_216513484.1 | WP_216513484.1 L,D-transpeptidase |
| 31(2) | WP_091155350.1 | WP_091155350.1 L,D-transpeptidase |
| 31(1) | WP_285748161.1 | WP_285748161.1 L,D-transpeptidase |
| 31(1) | WP_090921069.1 | WP_090921069.1 L,D-transpeptidase |
| 31(1) | WP_137062796.1 | WP_137062796.1 L,D-transpeptidase |
| 31(1) | WP_110822563.1 | WP_110822563.1 L,D-transpeptidase |
| 31(2) | WP_013309983.1 | WP_013309983.1 L,D-transpeptidase |
| 31(1) | WP_111268946.1 | WP_111268946.1 L,D-transpeptidase |
| 31(1) | WP_305166265.1 | WP_305166265.1 L,D-transpeptidase |
| 31(1) | WP_229522861.1 | WP_229522861.1 L,D-transpeptidase |
| 31(1) | WP_036670430.1 | WP_036670430.1 L,D-transpeptidase |
| 31(1) | WP_247902239.1 | WP_247902239.1 L,D-transpeptidase |
| 31(3) | WP_025716871.1 | WP_025716871.1 L,D-transpeptidase |
| 31(1) | WP_076289569.1 | WP_076289569.1 L,D-transpeptidase |
| 31(1) | WP_076208844.1 | WP_076208844.1 L,D-transpeptidase |
| 31(3) | WP_056700775.1 | WP_056700775.1 L,D-transpeptidase |
| 31(1) | WP_036609207.1 | WP_036609207.1 L,D-transpeptidase |
| 31(1) | WP_039273543.1 | WP_039273543.1 L,D-transpeptidase |
| 31(8) | WP_063566329.1 | WP_063566329.1 L,D-transpeptidase |
| 31(2) | WP_128101481.1 | WP_128101481.1 L,D-transpeptidase |
| 31(1) | WP_236417203.1 | WP_236417203.1 L,D-transpeptidase |
| 31(2) | WP_145329901.1 | WP_145329901.1 L,D-transpeptidase |
| 31(1) | WP_285079445.1 | WP_285079445.1 L,D-transpeptidase |
| 31(1) | WP_119601461.1 | WP_119601461.1 L,D-transpeptidase |
| 31(2) | WP_017689079.1 | WP_017689079.1 L,D-transpeptidase |
| 31(1) | WP_110892982.1 | WP_110892982.1 L,D-transpeptidase |
| 31(2) | WP_076311059.1 | WP_076311059.1 L,D-transpeptidase |
| 31(1) | WP_047842533.1 | WP_047842533.1 L,D-transpeptidase |
| 31(1) | WP_301249694.1 | WP_301249694.1 L,D-transpeptidase |
| 31(1) | WP_018883279.1 | WP_018883279.1 L,D-transpeptidase |
| 31(1) | WP_095287666.1 | WP_095287666.1 L,D-transpeptidase |
| 31(1) | WP_293202268.1 | WP_293202268.1 L,D-transpeptidase |
| 31(2) | WP_044644795.1 | WP_044644795.1 L,D-transpeptidase |
| 31(1) | WP_123067109.1 | WP_123067109.1 L,D-transpeptidase |
| 31(1) | WP_216975172.1 | WP_216975172.1 L,D-transpeptidase |
| 31(1) | WP_076156856.1 | WP_076156856.1 L,D-transpeptidase |
| 31(8) | WP_016821026.1 | WP_016821026.1 L,D-transpeptidase |
| 31(1) | WP_235308157.1 | WP_235308157.1 L,D-transpeptidase |
| 31(1) | WP_305755039.1 | WP_305755039.1 L,D-transpeptidase |
| 31(1) | WP_047909623.1 | WP_047909623.1 L,D-transpeptidase |
| 31(1) | WP_308721991.1 | WP_308721991.1 L,D-transpeptidase |
| 31(5) | WP_076318547.1 | WP_076318547.1 L,D-transpeptidase |
| 31(1) | WP_014281178.1 | WP_014281178.1 L,D-transpeptidase |
| 31(1) | WP_010346290.1 | WP_010346290.1 L,D-transpeptidase |
| 31(1) | WP_095358551.1 | WP_095358551.1 L,D-transpeptidase |
| 31(2) | WP_025676976.1 | WP_025676976.1 L,D-transpeptidase |
| 31(24) | WP_023988260.1 | WP_023988260.1 L,D-transpeptidase |
| 31(1) | WP_279664547.1 | WP_279664547.1 L,D-transpeptidase family protein |
| 31(1) | WP_024630320.1 | WP_024630320.1 L,D-transpeptidase |
| 31(2) | WP_153978063.1 | WP_153978063.1 L,D-transpeptidase |
| 31(1) | WP_122971553.1 | WP_122971553.1 L,D-transpeptidase family protein |
| 31(8) | WP_079346840.1 | WP_079346840.1 L,D-transpeptidase |
| 31(3) | WP_007430127.1 | WP_007430127.1 L,D-transpeptidase |
| 31(1) | WP_205053323.1 | WP_205053323.1 L,D-transpeptidase |
| 31(2) | WP_134910006.1 | WP_134910006.1 L,D-transpeptidase |
| 32(1) | WP_083180134.1 | WP_083180134.1 general stress protein |
| 32(2) | WP_128101491.1 | WP_128101491.1 general stress protein |
| 32(1) | WP_211745822.1 | WP_211745822.1 general stress protein |
| 32(1) | WP_133384360.1 | WP_133384360.1 general stress protein |
| 32(1) | WP_145046465.1 | WP_145046465.1 general stress protein |
| 32(1) | WP_216531938.1 | WP_216531938.1 general stress protein |
| 32(1) | WP_108465359.1 | WP_108465359.1 general stress protein |
| 32(1) | WP_127592420.1 | WP_127592420.1 general stress protein |
| 32(1) | WP_009590725.1 | WP_009590725.1 general stress protein |
| 32(4) | WP_019687045.1 | WP_019687045.1 general stress protein |
| 32(4) | WP_096774448.1 | WP_096774448.1 general stress protein |
| 32(3) | WP_145329913.1 | WP_145329913.1 general stress protein |
| 32(1) | WP_239289421.1 | WP_239289421.1 general stress protein |
| 32(1) | WP_289373786.1 | WP_289373786.1 general stress protein |

| | | |
|---|---|---|
| 32(1) | WP_116231757.1 | WP_116231757.1 general stress protein |
| 32(1) | WP_148246954.1 | WP_148246954.1 general stress protein |
| 32(3) | WP_015736234.1 | WP_015736234.1 general stress protein |
| 32(1) | WP_220684646.1 | WP_220684646.1 general stress protein |
| 32(2) | WP_095358543.1 | WP_095358543.1 general stress protein |
| 32(1) | WP_183581380.1 | WP_183581380.1 general stress protein |
| 32(1) | WP_301879141.1 | WP_301879141.1 general stress protein |
| 32(1) | WP_090993633.1 | WP_090993633.1 general stress protein |
| 32(1) | WP_072732444.1 | WP_072732444.1 general stress protein |
| 32(1) | WP_079694642.1 | WP_079694642.1 general stress protein |
| 32(1) | WP_188593167.1 | WP_188593167.1 general stress protein |
| 32(1) | WP_023957743.1 | WP_023957743.1 general stress protein |
| 32(1) | WP_095287654.1 | WP_095287654.1 general stress protein |
| 32(1) | WP_056700804.1 | WP_056700804.1 general stress protein |
| 32(1) | WP_145318680.1 | WP_145318680.1 general stress protein |
| 32(1) | WP_028589287.1 | WP_028589287.1 general stress protein |
| 32(1) | WP_209861630.1 | WP_209861630.1 general stress protein |
| 32(1) | WP_106765949.1 | WP_106765949.1 general stress protein |
| 32(1) | WP_210095103.1 | WP_210095103.1 general stress protein |
| 32(1) | WP_123067121.1 | WP_123067121.1 general stress protein |
| 32(4) | WP_076169521.1 | WP_076169521.1 general stress protein |
| 32(1) | WP_213532663.1 | WP_213532663.1 general stress protein |
| 32(1) | WP_079346808.1 | WP_079346808.1 general stress protein |
| 32(2) | WP_145156811.1 | WP_145156811.1 general stress protein |
| 32(1) | WP_264929201.1 | WP_264929201.1 general stress protein |
| 32(1) | WP_054958417.1 | WP_054958417.1 general stress protein |
| 32(1) | WP_274566444.1 | WP_274566444.1 general stress protein |
| 32(1) | WP_064639008.1 | WP_064639008.1 general stress protein |
| 32(1) | WP_293202257.1 | WP_293202257.1 general stress protein |
| 32(1) | WP_077567010.1 | WP_077567010.1 general stress protein |
| 32(1) | WP_305022915.1 | WP_305022915.1 general stress protein |
| 32(1) | WP_285748188.1 | WP_285748188.1 general stress protein |
| 32(3) | WP_062833886.1 | WP_062833886.1 general stress protein |
| 32(1) | WP_053783892.1 | WP_053783892.1 general stress protein |
| 32(1) | WP_016821041.1 | WP_016821041.1 general stress protein |
| 32(1) | WP_094936491.1 | WP_094936491.1 general stress protein |
| 32(1) | WP_090921042.1 | WP_090921042.1 general stress protein |
| 32(1) | WP_063208742.1 | WP_063208742.1 general stress protein |
| 32(5) | WP_017428324.1 | WP_017428324.1 general stress protein |
| 32(1) | WP_127199779.1 | WP_127199779.1 general stress protein |
| 32(2) | WP_138226018.1 | WP_138226018.1 general stress protein |
| 32(1) | WP_091155396.1 | WP_091155396.1 general stress protein |
| 32(1) | WP_136606870.1 | WP_136606870.1 general stress protein |
| 32(1) | WP_215153847.1 | WP_215153847.1 general stress protein |
| 32(1) | WP_113058723.1 | WP_113058723.1 general stress protein |
| 32(1) | WP_094092629.1 | WP_094092629.1 general stress protein |
| 32(1) | WP_090949173.1 | WP_090949173.1 general stress protein |
| 32(1) | WP_169505060.1 | WP_169505060.1 general stress protein |
| 32(1) | WP_191798534.1 | WP_191798534.1 general stress protein |
| 32(2) | WP_083655680.1 | WP_083655680.1 general stress protein |
| 32(1) | WP_244719667.1 | WP_244719667.1 general stress protein |
| 32(2) | WP_047909606.1 | WP_047909606.1 general stress protein |

| | | |
|---|---|---|
| 32(1) | WP_145021723.1 | WP_145021723.1 general stress protein |
| 32(1) | WP_105444105.1 | WP_105444105.1 general stress protein |
| 32(1) | WP_071220212.1 | WP_071220212.1 general stress protein |
| 32(1) | WP_160496922.1 | WP_160496922.1 general stress protein |
| 32(1) | WP_251375542.1 | WP_251375542.1 general stress protein |
| 32(1) | WP_119847558.1 | WP_119847558.1 general stress protein |
| 32(1) | WP_068622361.1 | WP_068622361.1 general stress protein |
| 32(2) | WP_013370825.1 | WP_013370825.1 general stress protein |
| 32(1) | WP_100537968.1 | WP_100537968.1 general stress protein |
| 32(1) | WP_194543492.1 | WP_194543492.1 general stress protein |
| 32(1) | WP_307413593.1 | WP_307413593.1 general stress protein |
| 32(1) | WP_236417227.1 | WP_236417227.1 general stress protein |
| 32(1) | WP_251583303.1 | WP_251583303.1 general stress protein |
| 32(1) | WP_192266470.1 | WP_192266470.1 general stress protein |
| 32(1) | WP_024630333.1 | WP_024630333.1 general stress protein |
| 32(1) | WP_007128619.1 | WP_007128619.1 general stress protein |
| 32(1) | WP_251537234.1 | WP_251537234.1 general stress protein |
| 32(1) | WP_018883262.1 | WP_018883262.1 general stress protein |
| 32(1) | WP_285079436.1 | WP_285079436.1 general stress protein |
| 32(1) | WP_110892958.1 | WP_110892958.1 general stress protein |
| 32(16) | WP_017689064.1 | WP_017689064.1 general stress protein |
| 32(7) | WP_063563227.1 | WP_063563227.1 general stress protein |
| 32(2) | WP_098746846.1 | WP_098746846.1 general stress protein |
| 32(1) | WP_247902250.1 | WP_247902250.1 general stress protein |
| 32(1) | WP_305755026.1 | WP_305755026.1 general stress protein |
| 32(1) | WP_054402900.1 | WP_054402900.1 general stress protein |
| 32(1) | WP_213618821.1 | WP_213618821.1 general stress protein |
| 32(1) | WP_062326154.1 | WP_062326154.1 general stress protein |
| 32(1) | WP_090908285.1 | WP_090908285.1 general stress protein |
| 32(3) | WP_127536191.1 | WP_127536191.1 general stress protein |
| 32(4) | WP_145134809.1 | WP_145134809.1 general stress protein |
| 32(1) | WP_094593882.1 | WP_094593882.1 general stress protein |
| 32(1) | WP_272035334.1 | WP_272035334.1 general stress protein |
| 32(5) | WP_006211194.1 | WP_006211194.1 general stress protein |
| 32(1) | WP_213649962.1 | WP_213649962.1 general stress protein |
| 32(1) | WP_152403401.1 | WP_152403401.1 general stress protein |
| 32(1) | WP_076322106.1 | WP_076322106.1 general stress protein |

| | | |
|---|---|---|
| 33(19) | WP_000136611.1 | WP_000136611.1 type VII secretion protein EssB |
| 33(1) | WP_077320203.1 | WP_077320203.1 type VII secretion protein EssB |
| 33(14) | WP_185337267.1 | WP_185337267.1 type VII secretion protein EssB |
| 33(1) | WP_098179587.1 | WP_098179587.1 type VII secretion protein EssB |
| 33(1) | WP_074598758.1 | WP_074598758.1 type VII secretion protein EssB |
| 33(1) | WP_086424324.1 | WP_086424324.1 type VII secretion protein EssB |
| 33(2) | WP_185516171.1 | WP_185516171.1 type VII secretion protein EssB |
| 33(2) | WP_185563008.1 | WP_185563008.1 type VII secretion protein EssB |
| 33(2) | WP_046325926.1 | WP_046325926.1 type VII secretion protein EssB |
| 33(1) | WP_099223952.1 | WP_099223952.1 type VII secretion protein EssB |
| 33(1) | WP_063536166.1 | WP_063536166.1 type VII secretion protein EssB |
| 33(3) | WP_098281580.1 | WP_098281580.1 type VII secretion protein EssB |
| 33(1) | WP_065845444.1 | WP_065845444.1 type VII secretion protein EssB |
| 33(1) | WP_098908415.1 | WP_098908415.1 type VII secretion protein EssB |

| | | |
|---|---|---|
| 33(2) | WP_098139265.1 | WP_098139265.1 type VII secretion protein EssB |
| 33(2) | WP_012582211.1 | WP_012582211.1 type VII secretion protein EssB |
| 33(2) | WP_000136616.1 | WP_000136616.1 type VII secretion protein EssB |
| 33(1) | WP_102956436.1 | WP_102956436.1 type VII secretion protein EssB |
| 33(6) | WP_000136612.1 | WP_000136612.1 type VII secretion protein EssB |
| 33(2) | WP_270995169.1 | WP_270995169.1 type VII secretion protein EssB |
| 33(1) | WP_098291294.1 | WP_098291294.1 type VII secretion protein EssB |
| 33(1) | WP_144596435.1 | WP_144596435.1 type VII secretion protein EssB |
| 33(2) | WP_271002889.1 | WP_271002889.1 type VII secretion protein EssB |
| 33(1) | WP_021291850.1 | WP_021291850.1 type VII secretion protein EssB |
| 33(1) | WP_098252273.1 | WP_098252273.1 type VII secretion protein EssB |
| 33(1) | WP_098142231.1 | WP_098142231.1 type VII secretion protein EssB |
| 33(1) | WP_038245448.1 | WP_038245448.1 type VII secretion protein EssB |
| 33(1) | WP_098537276.1 | WP_098537276.1 type VII secretion protein EssB |
| 33(2) | WP_185639017.1 | WP_185639017.1 type VII secretion protein EssB |
| 33(2) | WP_160807260.1 | WP_160807260.1 type VII secretion protein EssB |
| 33(4) | WP_097954756.1 | WP_097954756.1 type VII secretion protein EssB |
| 33(4) | WP_185363042.1 | WP_185363042.1 type VII secretion protein EssB |
| 33(2) | WP_185384173.1 | WP_185384173.1 type VII secretion protein EssB |
| 33(1) | WP_115359661.1 | WP_115359661.1 type VII secretion protein EssB |
| 33(1) | WP_088066064.1 | WP_088066064.1 type VII secretion protein EssB |
| 33(1) | WP_106496020.1 | WP_106496020.1 type VII secretion protein EssB |
| 33(6) | WP_075702372.1 | WP_075702372.1 type VII secretion protein EssB |
| 33(1) | WP_163095417.1 | WP_163095417.1 type VII secretion protein EssB |
| 33(1) | WP_000136618.1 | WP_000136618.1 type VII secretion protein EssB |
| 33(2) | WP_194318784.1 | WP_194318784.1 type VII secretion protein EssB |
| 33(12) | WP_185413345.1 | WP_185413345.1 type VII secretion protein EssB |
| 33(1) | WP_061113197.1 | WP_061113197.1 type VII secretion protein EssB |
| 33(4) | WP_185349993.1 | WP_185349993.1 type VII secretion protein EssB |
| 33(2) | WP_000136617.1 | WP_000136617.1 type VII secretion protein EssB |
| 33(4) | WP_038408418.1 | WP_038408418.1 type VII secretion protein EssB |
| 33(2) | WP_270993638.1 | WP_270993638.1 type VII secretion protein EssB |
| 33(1) | WP_290102341.1 | WP_290102341.1 type VII secretion protein EssB |
| 33(2) | WP_139590297.1 | WP_139590297.1 type VII secretion protein EssB |
| 33(2) | WP_011700924.1 | WP_011700924.1 type VII secretion protein EssB |
| 33(1) | WP_097956890.1 | WP_097956890.1 type VII secretion protein EssB |
| 33(1) | WP_209365678.1 | WP_209365678.1 type VII secretion protein EssB |
| 33(1) | WP_000136620.1 | WP_000136620.1 type VII secretion protein EssB |
| 33(1) | WP_030025434.1 | WP_030025434.1 type VII secretion protein EssB |
| 33(5) | WP_001990259.1 | WP_001990259.1 type VII secretion protein EssB |
| 33(1) | WP_101356551.1 | WP_101356551.1 type VII secretion protein EssB |
| 33(1) | WP_046946382.1 | WP_046946382.1 type VII secretion protein EssB |
| 33(6) | WP_185336680.1 | WP_185336680.1 type VII secretion protein EssB |
| 33(1) | WP_000136609.1 | WP_000136609.1 type VII secretion protein EssB |
| 33(1) | WP_035190280.1 | WP_035190280.1 type VII secretion protein EssB |
| 33(1) | WP_070173069.1 | WP_070173069.1 type VII secretion protein EssB |
| 33(1) | WP_086385918.1 | WP_086385918.1 type VII secretion protein EssB |
| 33(1) | WP_155891585.1 | WP_155891585.1 type VII secretion protein EssB |
| 33(1) | WP_016122371.1 | WP_016122371.1 type VII secretion protein EssB |
| 34(1) | WP_098612029.1 | WP_098612029.1 type VII secretion protein EssA |
| 34(8) | WP_000595649.1 | WP_000595649.1 type VII secretion protein EssA |
| 34(1) | WP_077320201.1 | WP_077320201.1 type VII secretion protein EssA |
| 34(18) | WP_185336681.1 | WP_185336681.1 type VII secretion protein EssA |
| 34(1) | WP_061678626.1 | WP_061678626.1 type VII secretion protein EssA |
| 34(1) | WP_209365680.1 | WP_209365680.1 type VII secretion protein EssA |
| 34(2) | WP_185469295.1 | WP_185469295.1 type VII secretion protein EssA |
| 34(6) | WP_038406377.1 | WP_038406377.1 type VII secretion protein EssA |
| 34(2) | WP_194339778.1 | WP_194339778.1 type VII secretion protein EssA |
| 34(2) | WP_185516172.1 | WP_185516172.1 type VII secretion protein EssA |
| 34(1) | WP_081674787.1 | WP_081674787.1 type VII secretion protein EssA |
| 34(1) | WP_086389010.1 | WP_086389010.1 type VII secretion protein EssA |
| 34(2) | WP_011700923.1 | WP_011700923.1 type VII secretion protein EssA |
| 34(2) | WP_185489277.1 | WP_185489277.1 type VII secretion protein EssA |
| 34(5) | WP_000595653.1 | WP_000595653.1 type VII secretion protein EssA |
| 34(2) | WP_185337266.1 | WP_185337266.1 type VII secretion protein EssA |
| 34(1) | WP_088066063.1 | WP_088066063.1 type VII secretion protein EssA |
| 34(1) | WP_163061523.1 | WP_163061523.1 type VII secretion protein EssA |
| 34(2) | WP_000595658.1 | WP_000595658.1 type VII secretion protein EssA |
| 34(1) | WP_088230468.1 | WP_088230468.1 type VII secretion protein EssA |
| 34(1) | WP_074598760.1 | WP_074598760.1 type VII secretion protein EssA |
| 34(29) | WP_000595655.1 | WP_000595655.1 type VII secretion protein EssA |
| 34(8) | WP_000595661.1 | WP_000595661.1 type VII secretion protein EssA |
| 34(2) | WP_185639018.1 | WP_185639018.1 type VII secretion protein EssA |
| 34(1) | WP_098226393.1 | WP_098226393.1 type VII secretion protein EssA |
| 34(1) | WP_158701616.1 | WP_158701616.1 type VII secretion protein EssA |
| 34(10) | WP_003750475.1 | WP_003750475.1 type VII secretion protein EssA |
| 34(1) | WP_101356553.1 | WP_101356553.1 type VII secretion protein EssA |
| 34(1) | WP_018664839.1 | WP_018664839.1 type VII secretion protein EssA |
| 34(1) | WP_029141778.1 | WP_029141778.1 type VII secretion protein EssA |
| 34(1) | WP_098300887.1 | WP_098300887.1 type VII secretion protein EssA |
| 34(2) | WP_185563007.1 | WP_185563007.1 type VII secretion protein EssA |
| 34(2) | WP_046325925.1 | WP_046325925.1 type VII secretion protein EssA |
| 34(2) | WP_000595657.1 | WP_000595657.1 type VII secretion protein EssA |
| 34(14) | WP_185357241.1 | WP_185357241.1 type VII secretion protein EssA |
| 34(2) | WP_070214792.1 | WP_070214792.1 type VII secretion protein EssA |
| 34(4) | WP_194334462.1 | WP_194334462.1 type VII secretion protein EssA |
| 34(2) | WP_185461322.1 | WP_185461322.1 type VII secretion protein EssA |
| 34(1) | WP_240422345.1 | WP_240422345.1 type VII secretion protein EssA |
| 34(1) | WP_115359659.1 | WP_115359659.1 type VII secretion protein EssA |
| 34(2) | WP_033695489.1 | WP_033695489.1 type VII secretion protein EssA |
| 34(1) | WP_098819895.1 | WP_098819895.1 type VII secretion protein EssA |
| 34(3) | WP_081856149.1 | WP_081856149.1 type VII secretion protein EssA |
| 34(1) | WP_098291295.1 | WP_098291295.1 type VII secretion protein EssA |
| 35(2) | WP_081384060.1 | WP_081384060.1 ATPase, T2SS/T4P/T4SS family |
| 35(1) | WP_042211001.1 | WP_042211001.1 Flp pilus assembly complex ATPase component TadA |
| 35(1) | WP_145939626.1 | WP_145939626.1 ATPase, T2SS/T4P/T4SS family |
| 35(1) | WP_207952309.1 | WP_207952309.1 ATPase, T2SS/T4P/T4SS family |
| 35(1) | WP_021878928.1 | WP_021878928.1 ATPase, T2SS/T4P/T4SS family |
| 35(1) | WP_068782564.1 | WP_068782564.1 ATPase, T2SS/T4P/T4SS family |
| 35(2) | WP_215185465.1 | WP_215185465.1 ATPase, T2SS/T4P/T4SS family |
| 35(1) | WP_236334347.1 | WP_236334347.1 ATPase, T2SS/T4P/T4SS family |

| | | |
|---|---|---|
| 35(1) | WP_019912147.1 | WP_019912147.1 ATPase, T2SS/T4P/T4SS family |
| 35(1) | WP_113672713.1 | WP_113672713.1 ATPase, T2SS/T4P/T4SS family |
| 35(1) | WP_216518271.1 | WP_216518271.1 ATPase, T2SS/T4P/T4SS family |
| 35(1) | WP_074085352.1 | WP_074085352.1 ATPase, T2SS/T4P/T4SS family |
| 35(1) | WP_166244913.1 | WP_166244913.1 ATPase, T2SS/T4P/T4SS family |
| 35(1) | WP_055105970.1 | WP_055105970.1 Flp pilus assembly complex ATPase component TadA |
| 35(2) | WP_195431315.1 | WP_195431315.1 ATPase, T2SS/T4P/T4SS family |
| 35(1) | WP_099517099.1 | WP_099517099.1 ATPase, T2SS/T4P/T4SS family |
| 35(1) | WP_199613910.1 | WP_199613910.1 ATPase, T2SS/T4P/T4SS family |
| 35(1) | WP_254848774.1 | WP_254848774.1 ATPase, T2SS/T4P/T4SS family |
| 35(1) | WP_227793449.1 | WP_227793449.1 ATPase, T2SS/T4P/T4SS family |
| 35(1) | WP_251596688.1 | WP_251596688.1 ATPase, T2SS/T4P/T4SS family |
| 35(1) | WP_287600026.1 | WP_287600026.1 ATPase, T2SS/T4P/T4SS family |
| 35(1) | WP_289376835.1 | WP_289376835.1 ATPase, T2SS/T4P/T4SS family |
| 35(1) | WP_076242359.1 | WP_076242359.1 Flp pilus assembly complex ATPase component TadA |
| 35(1) | WP_013658720.1 | WP_013658720.1 ATPase, T2SS/T4P/T4SS family |
| 35(1) | WP_036626784.1 | WP_036626784.1 ATPase, T2SS/T4P/T4SS family |
| 35(1) | WP_094878482.1 | WP_094878482.1 ATPase, T2SS/T4P/T4SS family |
| 35(1) | WP_229521537.1 | WP_229521537.1 ATPase, T2SS/T4P/T4SS family |
| 35(2) | WP_003516561.1 | WP_003516561.1 ATPase, T2SS/T4P/T4SS family |
| 35(1) | WP_271839917.1 | WP_271839917.1 ATPase, T2SS/T4P/T4SS family |
| 35(1) | WP_108464080.1 | WP_108464080.1 ATPase, T2SS/T4P/T4SS family |
| 35(1) | WP_220388588.1 | WP_220388588.1 ATPase, T2SS/T4P/T4SS family |
| 35(2) | WP_081383100.1 | WP_081383100.1 ATPase, T2SS/T4P/T4SS family |
| 35(2) | WP_281813123.1 | WP_281813123.1 ATPase, T2SS/T4P/T4SS family |
| 35(1) | WP_283910917.1 | WP_283910917.1 ATPase, T2SS/T4P/T4SS family |
| 35(1) | WP_148450954.1 | WP_148450954.1 ATPase, T2SS/T4P/T4SS family |
| 35(1) | WP_143315249.1 | WP_143315249.1 ATPase, T2SS/T4P/T4SS family |
| 35(1) | WP_036944563.1 | WP_036944563.1 ATPase, T2SS/T4P/T4SS family |
| 35(1) | WP_293755501.1 | WP_293755501.1 ATPase, T2SS/T4P/T4SS family |
| 35(1) | WP_138755936.1 | WP_138755936.1 Flp pilus assembly complex ATPase component TadA |
| 35(1) | WP_052421247.1 | WP_052421247.1 Flp pilus assembly complex ATPase component TadA |
| 35(1) | WP_291647962.1 | WP_291647962.1 ATPase, T2SS/T4P/T4SS family |
| 35(1) | WP_240968294.1 | WP_240968294.1 ATPase, T2SS/T4P/T4SS family |
| 35(1) | WP_308723531.1 | WP_308723531.1 ATPase, T2SS/T4P/T4SS family |
| 35(1) | WP_027088409.1 | WP_027088409.1 hypothetical protein |
| 35(1) | WP_013914198.1 | WP_013914198.1 ATPase, T2SS/T4P/T4SS family |
| 35(1) | WP_188535056.1 | WP_188535056.1 ATPase, T2SS/T4P/T4SS family |
| 35(1) | WP_110899377.1 | WP_110899377.1 ATPase, T2SS/T4P/T4SS family |
| 35(1) | WP_081388403.1 | WP_081388403.1 ATPase, T2SS/T4P/T4SS family |
| 35(1) | WP_213506869.1 | WP_213506869.1 ATPase, T2SS/T4P/T4SS family |
| 35(1) | WP_185165642.1 | WP_185165642.1 ATPase, T2SS/T4P/T4SS family |
| 35(1) | WP_038286707.1 | WP_038286707.1 ATPase, T2SS/T4P/T4SS family |
| 35(1) | WP_076101564.1 | WP_076101564.1 ATPase, T2SS/T4P/T4SS family |
| 35(1) | WP_026478102.1 | WP_026478102.1 ATPase, T2SS/T4P/T4SS family |
| 35(1) | WP_189018593.1 | WP_189018593.1 ATP-binding cassette domain-containing protein |
| 35(1) | WP_127194965.1 | WP_127194965.1 ATPase, T2SS/T4P/T4SS family |
| 35(2) | WP_042125611.1 | WP_042125611.1 ATPase, T2SS/T4P/T4SS family |
| 35(1) | WP_232243201.1 | WP_232243201.1 ATPase, T2SS/T4P/T4SS family |
| 35(1) | WP_119601506.1 | WP_119601506.1 ATPase, T2SS/T4P/T4SS family |
| 35(1) | WP_111273178.1 | WP_111273178.1 ATPase, T2SS/T4P/T4SS family |
| 35(1) | WP_248063425.1 | WP_248063425.1 ATPase, T2SS/T4P/T4SS family |
| 35(1) | WP_229106380.1 | WP_229106380.1 ATPase, T2SS/T4P/T4SS family |
| 35(1) | WP_055109399.1 | WP_055109399.1 ATPase, T2SS/T4P/T4SS family |
| 35(2) | WP_017692339.1 | WP_017692339.1 ATPase, T2SS/T4P/T4SS family |
| 35(1) | WP_069193881.1 | WP_069193881.1 ATPase, T2SS/T4P/T4SS family |
| 35(1) | WP_132283083.1 | WP_132283083.1 ATPase, T2SS/T4P/T4SS family |
| 35(1) | WP_253440806.1 | WP_253440806.1 ATPase, T2SS/T4P/T4SS family |
| 35(3) | WP_094870542.1 | WP_094870542.1 ATPase, T2SS/T4P/T4SS family |
| 35(3) | WP_076281082.1 | WP_076281082.1 ATPase, T2SS/T4P/T4SS family |
| 35(1) | WP_242987673.1 | WP_242987673.1 ATPase, T2SS/T4P/T4SS family |
| 35(1) | WP_278278679.1 | WP_278278679.1 ATPase, T2SS/T4P/T4SS family |
| 35(1) | WP_089522641.1 | WP_089522641.1 ATPase, T2SS/T4P/T4SS family |
| 35(1) | WP_307199927.1 | WP_307199927.1 pilus assembly protein CpaF |
| 35(1) | WP_052380732.1 | WP_052380732.1 Flp pilus assembly complex ATPase component TadA |
| 35(1) | WP_239498768.1 | WP_239498768.1 ATPase, T2SS/T4P/T4SS family |
| 35(1) | WP_085499116.1 | WP_085499116.1 ATPase, T2SS/T4P/T4SS family |
| 35(2) | WP_246628092.1 | WP_246628092.1 ATPase, T2SS/T4P/T4SS family |
| 35(1) | WP_280788629.1 | WP_280788629.1 ATPase, T2SS/T4P/T4SS family |
| 35(1) | WP_094870404.1 | WP_094870404.1 ATPase, T2SS/T4P/T4SS family |
| 35(1) | WP_128631279.1 | WP_128631279.1 ATPase, T2SS/T4P/T4SS family |
| 35(1) | WP_150457773.1 | WP_150457773.1 ATPase, T2SS/T4P/T4SS family |
| 35(1) | WP_105368683.1 | WP_105368683.1 ATPase, T2SS/T4P/T4SS family |
| 35(1) | WP_212693247.1 | WP_212693247.1 ATPase, T2SS/T4P/T4SS family |
| 35(1) | WP_126996295.1 | WP_126996295.1 ATPase, T2SS/T4P/T4SS family |
| 35(1) | WP_120462958.1 | WP_120462958.1 ATPase, T2SS/T4P/T4SS family |
| 35(1) | WP_024632695.1 | WP_024632695.1 ATPase, T2SS/T4P/T4SS family |
| 35(1) | WP_165097602.1 | WP_165097602.1 ATPase, T2SS/T4P/T4SS family |
| 35(1) | WP_239617493.1 | WP_239617493.1 CpaF/VirB11 family protein |
| 35(1) | WP_120112297.1 | WP_120112297.1 Flp pilus assembly complex ATPase component TadA |
| 35(1) | WP_223068037.1 | WP_223068037.1 ATPase, T2SS/T4P/T4SS family |
| 35(1) | WP_102710713.1 | WP_102710713.1 ATPase, T2SS/T4P/T4SS family |
| 35(1) | WP_042134282.1 | WP_042134282.1 Flp pilus assembly complex ATPase component TadA |
| 35(1) | WP_273321531.1 | WP_273321531.1 ATPase, T2SS/T4P/T4SS family |
| 35(1) | WP_076210954.1 | WP_076210954.1 ATPase, T2SS/T4P/T4SS family |
| 35(1) | WP_181603062.1 | WP_181603062.1 ATPase, T2SS/T4P/T4SS family |
| 35(1) | WP_207651899.1 | WP_207651899.1 ATPase, T2SS/T4P/T4SS family |
| 35(1) | WP_081391808.1 | WP_081391808.1 ATPase, T2SS/T4P/T4SS family |
| 35(3) | WP_155609763.1 | WP_155609763.1 ATPase, T2SS/T4P/T4SS family |
| 35(1) | WP_238655341.1 | WP_238655341.1 ATPase, T2SS/T4P/T4SS family |
| 35(1) | WP_145153673.1 | WP_145153673.1 ATPase, T2SS/T4P/T4SS family |
| 35(1) | WP_212695797.1 | WP_212695797.1 ATPase, T2SS/T4P/T4SS family |
| 35(2) | WP_211022796.1 | WP_211022796.1 ATPase, T2SS/T4P/T4SS family |
| 35(1) | WP_110934081.1 | WP_110934081.1 ATPase, T2SS/T4P/T4SS family |
| 35(1) | WP_068696915.1 | WP_068696915.1 ATPase, T2SS/T4P/T4SS family |
| 35(1) | WP_137698119.1 | WP_137698119.1 ATPase, T2SS/T4P/T4SS family |
| 35(1) | WP_125084790.1 | WP_125084790.1 ATPase, T2SS/T4P/T4SS family |
| 35(1) | WP_210087121.1 | WP_210087121.1 ATPase, T2SS/T4P/T4SS family |
| 35(1) | WP_150364188.1 | WP_150364188.1 ATPase, T2SS/T4P/T4SS family |
| 35(1) | WP_039306340.1 | WP_039306340.1 Flp pilus assembly complex ATPase component TadA |
| 35(1) | WP_083606938.1 | WP_083606938.1 ATPase, T2SS/T4P/T4SS family |
| 35(1) | WP_127570454.1 | WP_127570454.1 hypothetical protein |
| 35(1) | WP_306812736.1 | WP_306812736.1 ATPase, T2SS/T4P/T4SS family |
| 35(1) | WP_280521249.1 | WP_280521249.1 ATPase, T2SS/T4P/T4SS family |

| | | |
|---|---|---|
| 35(1) | WP_081389321.1 | WP_081389321.1 ATPase, T2SS/T4P/T4SS family |
| 35(1) | WP_179034780.1 | WP_179034780.1 ATPase, T2SS/T4P/T4SS family |
| 35(1) | WP_249897308.1 | WP_249897308.1 ATPase, T2SS/T4P/T4SS family |
| 35(2) | WP_081389017.1 | WP_081389017.1 ATPase, T2SS/T4P/T4SS family |
| 35(1) | WP_081390655.1 | WP_081390655.1 ATPase, T2SS/T4P/T4SS family |
| 35(1) | WP_251417644.1 | WP_251417644.1 ATPase, T2SS/T4P/T4SS family |
| 35(2) | WP_014368246.1 | WP_014368246.1 ATPase, T2SS/T4P/T4SS family |
| 35(2) | WP_036702427.1 | WP_036702427.1 ATPase, T2SS/T4P/T4SS family |
| 35(2) | WP_212592205.1 | WP_212592205.1 ATPase, T2SS/T4P/T4SS family |
| 35(1) | WP_184183038.1 | WP_184183038.1 ATPase, T2SS/T4P/T4SS family |
| 35(1) | WP_209872733.1 | WP_209872733.1 ATPase, T2SS/T4P/T4SS family |
| 35(5) | WP_036684499.1 | WP_036684499.1 ATPase, T2SS/T4P/T4SS family |
| 36(1) | WP_277156179.1 | WP_277156179.1 DUF4176 domain-containing protein |
| 36(4) | WP_082425285.1 | WP_082425285.1 DUF4176 domain-containing protein |
| 36(1) | WP_022349189.1 | WP_022349189.1 DUF4176 domain-containing protein |
| 36(1) | WP_270472019.1 | WP_270472019.1 DUF4176 domain-containing protein |
| 36(1) | WP_302153078.1 | WP_302153078.1 DUF4176 domain-containing protein |
| 36(1) | WP_180965812.1 | WP_180965812.1 DUF4176 domain-containing protein |
| 36(1) | WP_307738730.1 | WP_307738730.1 DUF4176 domain-containing protein |
| 36(4) | WP_279325364.1 | WP_279325364.1 DUF4176 domain-containing protein |
| 36(2) | WP_180965896.1 | WP_180965896.1 DUF4176 domain-containing protein |
| 36(1) | WP_197256862.1 | WP_197256862.1 DUF4176 domain-containing protein |
| 36(2) | WP_005210446.1 | WP_005210446.1 DUF4176 domain-containing protein |
| 36(2) | WP_168163959.1 | WP_168163959.1 DUF4176 domain-containing protein |
| 36(1) | WP_065291247.1 | WP_065291247.1 DUF4176 domain-containing protein |
| 36(1) | WP_132008151.1 | WP_132008151.1 DUF4176 domain-containing protein |
| 36(1) | WP_307394652.1 | WP_307394652.1 DUF4176 domain-containing protein |
| 36(1) | WP_152402380.1 | WP_152402380.1 DUF4176 domain-containing protein |
| 36(1) | WP_291672196.1 | WP_291672196.1 DUF4176 domain-containing protein |
| 36(1) | WP_156558887.1 | WP_156558887.1 DUF4176 domain-containing protein |
| 36(1) | WP_223592662.1 | WP_223592662.1 DUF4176 domain-containing protein |
| 36(1) | WP_045518288.1 | WP_045518288.1 DUF4176 domain-containing protein |
| 36(1) | WP_303942533.1 | WP_303942533.1 DUF4176 domain-containing protein |
| 36(1) | WP_055277388.1 | WP_055277388.1 DUF4176 domain-containing protein |
| 36(2) | WP_002031668.1 | WP_002031668.1 DUF4176 domain-containing protein |
| 36(5) | WP_021253117.1 | WP_021253117.1 DUF4176 domain-containing protein |
| 36(1) | WP_205001930.1 | WP_205001930.1 DUF4176 domain-containing protein |
| 36(8) | WP_005546513.1 | WP_005546513.1 DUF4176 domain-containing protein |
| 36(1) | WP_270574767.1 | WP_270574767.1 DUF4176 domain-containing protein |
| 36(1) | WP_195343915.1 | WP_195343915.1 DUF4176 domain-containing protein |
| 36(3) | WP_087433607.1 | WP_087433607.1 DUF4176 domain-containing protein |
| 36(1) | WP_247083039.1 | WP_247083039.1 DUF4176 domain-containing protein |
| 36(2) | WP_087442413.1 | WP_087442413.1 DUF4176 domain-containing protein |
| 36(2) | WP_185413331.1 | WP_185413331.1 DUF4176 domain-containing protein |
| 36(4) | WP_006674861.1 | WP_006674861.1 DUF4176 domain-containing protein |
| 36(1) | WP_145156807.1 | WP_145156807.1 DUF4176 domain-containing protein |
| 36(1) | WP_277142078.1 | WP_277142078.1 DUF4176 domain-containing protein |
| 36(2) | WP_194230402.1 | WP_194230402.1 DUF4176 domain-containing protein |
| 36(1) | WP_244719638.1 | WP_244719638.1 DUF4176 domain-containing protein |
| 36(1) | WP_307601041.1 | WP_307601041.1 DUF4176 domain-containing protein |
| 36(1) | WP_277150351.1 | WP_277150351.1 DUF4176 domain-containing protein |
| 36(1) | WP_277157941.1 | WP_277157941.1 DUF4176 domain-containing protein |
| 36(2) | WP_102372751.1 | WP_102372751.1 DUF4176 domain-containing protein |
| 36(1) | WP_273383880.1 | WP_273383880.1 DUF4176 domain-containing protein |
| 36(1) | WP_183578455.1 | WP_183578455.1 DUF4176 domain-containing protein |
| 36(1) | WP_050754943.1 | WP_050754943.1 DUF4176 domain-containing protein |
| 36(2) | WP_082423918.1 | WP_082423918.1 DUF4176 domain-containing protein |
| 36(1) | WP_270472003.1 | WP_270472003.1 DUF4176 domain-containing protein |
| 36(1) | WP_019424145.1 | WP_019424145.1 DUF4176 domain-containing protein |
| 36(1) | WP_006720314.1 | WP_006720314.1 DUF4176 domain-containing protein |
| 36(1) | WP_231417870.1 | WP_231417870.1 DUF4176 domain-containing protein |
| 36(1) | WP_216830398.1 | WP_216830398.1 DUF4176 domain-containing protein |
| 36(1) | WP_073008254.1 | WP_073008254.1 DUF4176 domain-containing protein |
| 36(1) | WP_130810255.1 | WP_130810255.1 DUF4176 domain-containing protein |
| 36(4) | WP_076167866.1 | WP_076167866.1 DUF4176 domain-containing protein |
| 36(1) | WP_013658709.1 | WP_013658709.1 DUF4176 domain-containing protein |
| 36(1) | WP_160037487.1 | WP_160037487.1 DUF4176 domain-containing protein |
| 36(1) | WP_121610590.1 | WP_121610590.1 DUF4176 domain-containing protein |
| 36(1) | WP_145948999.1 | WP_145948999.1 DUF4176 domain-containing protein |
| 36(1) | WP_142543126.1 | WP_142543126.1 DUF4176 domain-containing protein |
| 36(1) | WP_273383873.1 | WP_273383873.1 DUF4176 domain-containing protein |
| 36(1) | WP_094593875.1 | WP_094593875.1 DUF4176 domain-containing protein |
| 36(1) | WP_294953166.1 | WP_294953166.1 DUF4176 domain-containing protein |
| 36(1) | WP_213426990.1 | WP_213426990.1 DUF4176 domain-containing protein |
| 36(1) | WP_212330868.1 | WP_212330868.1 DUF4176 domain-containing protein |
| 36(1) | WP_036944555.1 | WP_036944555.1 DUF4176 domain-containing protein |
| 36(1) | WP_277175581.1 | WP_277175581.1 DUF4176 domain-containing protein |
| 36(1) | WP_105443569.1 | WP_105443569.1 DUF4176 domain-containing protein |
| 36(4) | WP_291288269.1 | WP_291288269.1 DUF4176 domain-containing protein |
| 36(1) | WP_307738725.1 | WP_307738725.1 DUF4176 domain-containing protein |
| 36(1) | WP_212983256.1 | WP_212983256.1 DUF4176 domain-containing protein |
| 36(1) | WP_270472012.1 | WP_270472012.1 DUF4176 domain-containing protein |
| 36(1) | WP_212330885.1 | WP_212330885.1 DUF4176 domain-containing protein |
| 36(1) | WP_034275551.1 | WP_034275551.1 DUF4176 domain-containing protein |
| 36(2) | WP_136604815.1 | WP_136604815.1 DUF4176 domain-containing protein |
| 36(4) | WP_291288264.1 | WP_291288264.1 DUF4176 domain-containing protein |
| 36(1) | WP_106496032.1 | WP_106496032.1 DUF4176 domain-containing protein |
| 36(1) | WP_277166580.1 | WP_277166580.1 DUF4176 domain-containing protein |
| 36(1) | WP_036707110.1 | WP_036707110.1 DUF4176 domain-containing protein |
| 36(1) | WP_207302590.1 | WP_207302590.1 DUF4176 domain-containing protein |
| 36(1) | WP_277157935.1 | WP_277157935.1 DUF4176 domain-containing protein |
| 36(1) | WP_195542468.1 | WP_195542468.1 DUF4176 domain-containing protein |
| 36(3) | WP_168178785.1 | WP_168178785.1 DUF4176 domain-containing protein |
| 36(1) | WP_202945206.1 | WP_202945206.1 DUF4176 domain-containing protein |
| 36(1) | WP_260457842.1 | WP_260457842.1 DUF4176 domain-containing protein |
| 36(1) | WP_233697175.1 | WP_233697175.1 DUF4176 domain-containing protein |
| 36(1) | WP_289613136.1 | WP_289613136.1 DUF4176 domain-containing protein |
| 36(1) | WP_138186693.1 | WP_138186693.1 DUF4176 domain-containing protein |
| 36(1) | WP_058271115.1 | WP_058271115.1 DUF4176 domain-containing protein |
| 36(3) | WP_119794363.1 | WP_119794363.1 DUF4176 domain-containing protein |
| 36(1) | WP_308615621.1 | WP_308615621.1 DUF4176 domain-containing protein |
| 36(4) | WP_163872766.1 | WP_163872766.1 DUF4176 domain-containing protein |
| 36(1) | WP_070638661.1 | WP_070638661.1 DUF4176 domain-containing protein |

| | | |
|---|---|---|
| 36(1) | WP_303942542.1 | WP_303942542.1 DUF4176 domain-containing protein |
| 36(1) | WP_302153074.1 | WP_302153074.1 DUF4176 domain-containing protein |
| 36(1) | WP_268058016.1 | WP_268058016.1 DUF4176 domain-containing protein |
| 36(1) | WP_021261881.1 | WP_021261881.1 DUF4176 domain-containing protein |
| 36(1) | WP_116231175.1 | WP_116231175.1 DUF4176 domain-containing protein |
| 36(1) | WP_237391669.1 | WP_237391669.1 DUF4176 domain-containing protein |
| 37(1) | WP_094870406.1 | WP_094870406.1 hypothetical protein |
| 37(1) | WP_244964940.1 | WP_244964940.1 ABC transporter permease |
| 37(1) | WP_076274893.1 | WP_076274893.1 hypothetical protein |
| 37(1) | WP_211023043.1 | WP_211023043.1 ABC transporter permease |
| 37(1) | WP_166244088.1 | WP_166244088.1 hypothetical protein |
| 37(1) | WP_209872721.1 | WP_209872721.1 hypothetical protein |
| 37(1) | WP_114298756.1 | WP_114298756.1 hypothetical protein |
| 37(1) | WP_099517096.1 | WP_099517096.1 hypothetical protein |
| 37(1) | WP_089522644.1 | WP_089522644.1 hypothetical protein |
| 37(1) | WP_042203654.1 | WP_042203654.1 hypothetical protein |
| 37(1) | WP_042203657.1 | WP_042203657.1 hypothetical protein |
| 37(1) | WP_253440800.1 | WP_253440800.1 ABC transporter permease |
| 37(2) | WP_281813131.1 | WP_281813131.1 hypothetical protein |
| 37(1) | WP_039871660.1 | WP_039871660.1 hypothetical protein |
| 37(1) | WP_211082921.1 | WP_211082921.1 ABC transporter permease |
| 37(1) | WP_182301854.1 | WP_182301854.1 hypothetical protein |
| 37(1) | WP_211082920.1 | WP_211082920.1 ABC transporter permease |
| 37(1) | WP_019912143.1 | WP_019912143.1 hypothetical protein |
| 37(1) | WP_074108290.1 | WP_074108290.1 hypothetical protein |
| 37(1) | WP_076242360.1 | WP_076242360.1 hypothetical protein |
| 37(1) | WP_089522643.1 | WP_089522643.1 hypothetical protein |
| 37(1) | WP_127194968.1 | WP_127194968.1 hypothetical protein |
| 37(1) | WP_068696912.1 | WP_068696912.1 ABC transporter permease |
| 37(1) | WP_127570452.1 | WP_127570452.1 hypothetical protein |
| 37(1) | WP_152392582.1 | WP_152392582.1 hypothetical protein |
| 37(2) | WP_076104581.1 | WP_076104581.1 hypothetical protein |
| 37(2) | WP_219872464.1 | WP_219872464.1 hypothetical protein |
| 37(2) | WP_076287004.1 | WP_076287004.1 hypothetical protein |
| 37(1) | WP_076144384.1 | WP_076144384.1 hypothetical protein |
| 37(1) | WP_289376836.1 | WP_289376836.1 ABC transporter permease |
| 37(1) | WP_019912145.1 | WP_019912145.1 hypothetical protein |
| 37(1) | WP_039871661.1 | WP_039871661.1 hypothetical protein |
| 37(1) | WP_024632693.1 | WP_024632693.1 hypothetical protein |
| 37(1) | WP_076279377.1 | WP_076279377.1 hypothetical protein |
| 37(1) | WP_216518266.1 | WP_216518266.1 ABC transporter permease |
| 37(1) | WP_210087119.1 | WP_210087119.1 ABC transporter permease |
| 37(1) | WP_068782421.1 | WP_068782421.1 ABC transporter permease |
| 37(1) | WP_099517097.1 | WP_099517097.1 hypothetical protein |
| 37(3) | WP_076135794.1 | WP_076135794.1 hypothetical protein |
| 37(1) | WP_076303954.1 | WP_076303954.1 hypothetical protein |
| 37(3) | WP_076219081.1 | WP_076219081.1 hypothetical protein |
| 37(1) | WP_238187868.1 | WP_238187868.1 hypothetical protein |
| 37(1) | WP_238187867.1 | WP_238187867.1 hypothetical protein |
| 37(2) | WP_076287003.1 | WP_076287003.1 hypothetical protein |
| 37(2) | WP_042134289.1 | WP_042134289.1 hypothetical protein |
| 37(1) | WP_110931527.1 | WP_110931527.1 ABC transporter permease |
| 37(1) | WP_074108289.1 | WP_074108289.1 hypothetical protein |
| 37(1) | WP_145153664.1 | WP_145153664.1 ABC transporter permease |
| 37(1) | WP_074085350.1 | WP_074085350.1 hypothetical protein |
| 37(1) | WP_248063429.1 | WP_248063429.1 ABC transporter permease |
| 37(1) | WP_293755493.1 | WP_293755493.1 hypothetical protein |
| 37(1) | WP_068536357.1 | WP_068536357.1 ABC transporter permease |
| 37(1) | WP_111273176.1 | WP_111273176.1 ABC transporter permease |
| 37(1) | WP_216518268.1 | WP_216518268.1 ABC transporter permease |
| 37(2) | WP_113672716.1 | WP_113672716.1 hypothetical protein |
| 37(3) | WP_076104582.1 | WP_076104582.1 hypothetical protein |
| 37(1) | WP_110899375.1 | WP_110899375.1 ABC transporter permease |
| 37(2) | WP_076287002.1 | WP_076287002.1 hypothetical protein |
| 37(1) | WP_273321524.1 | WP_273321524.1 hypothetical protein |
| 37(1) | WP_042203655.1 | WP_042203655.1 hypothetical protein |
| 37(1) | WP_212695800.1 | WP_212695800.1 hypothetical protein |
| 37(1) | WP_076210959.1 | WP_076210959.1 ABC transporter permease |
| 37(1) | WP_076219080.1 | WP_076219080.1 hypothetical protein |
| 37(1) | WP_308723529.1 | WP_308723529.1 ABC transporter permease |
| 37(1) | WP_184183019.1 | WP_184183019.1 ABC transporter permease |
| 37(3) | WP_094870407.1 | WP_094870407.1 hypothetical protein |
| 37(3) | WP_094870408.1 | WP_094870408.1 hypothetical protein |
| 37(1) | WP_017692341.1 | WP_017692341.1 hypothetical protein |
| 37(1) | WP_042134286.1 | WP_042134286.1 hypothetical protein |
| 37(1) | WP_238187866.1 | WP_238187866.1 hypothetical protein |
| 37(1) | WP_042211003.1 | WP_042211003.1 hypothetical protein |
| 37(1) | WP_039306332.1 | WP_039306332.1 hypothetical protein |
| 37(2) | WP_076303955.1 | WP_076303955.1 hypothetical protein |
| 37(1) | WP_017692342.1 | WP_017692342.1 hypothetical protein |
| 37(1) | WP_039871663.1 | WP_039871663.1 hypothetical protein |
| 37(1) | WP_209872724.1 | WP_209872724.1 hypothetical protein |
| 37(1) | WP_166244089.1 | WP_166244089.1 hypothetical protein |
| 37(1) | WP_076208271.1 | WP_076208271.1 hypothetical protein |
| 37(2) | WP_219872463.1 | WP_219872463.1 hypothetical protein |
| 37(2) | WP_076104580.1 | WP_076104580.1 hypothetical protein |
| 37(1) | WP_307199924.1 | WP_307199924.1 ABC transporter permease |
| 37(1) | WP_150364190.1 | WP_150364190.1 ABC transporter permease |
| 37(1) | WP_154118022.1 | WP_154118022.1 hypothetical protein |
| 37(2) | WP_094878484.1 | WP_094878484.1 hypothetical protein |
| 37(1) | WP_138755934.1 | WP_138755934.1 hypothetical protein |
| 37(1) | WP_145153667.1 | WP_145153667.1 ABC transporter permease |
| 37(1) | WP_307600995.1 | WP_307600995.1 hypothetical protein |
| 37(1) | WP_308723528.1 | WP_308723528.1 ABC transporter permease |
| 37(1) | WP_076135795.1 | WP_076135795.1 hypothetical protein |
| 37(1) | WP_019912144.1 | WP_019912144.1 hypothetical protein |
| 37(1) | WP_154118024.1 | WP_154118024.1 hypothetical protein |
| 37(1) | WP_209872727.1 | WP_209872727.1 hypothetical protein |
| 37(1) | WP_055105807.1 | WP_055105807.1 hypothetical protein |
| 37(1) | WP_206098709.1 | WP_206098709.1 ABC transporter permease |
| 37(1) | WP_189018589.1 | WP_189018589.1 hypothetical protein |
| 37(1) | WP_042211004.1 | WP_042211004.1 hypothetical protein |
| 37(1) | WP_039306335.1 | WP_039306335.1 hypothetical protein |

37(1) WP_152392581.1 WP_152392581.1 hypothetical protein
37(2) WP_076219079.1 WP_076219079.1 hypothetical protein
37(1) WP_027088411.1 WP_027088411.1 hypothetical protein
37(1) WP_074085348.1 WP_074085348.1 hypothetical protein
37(1) WP_042211005.1 WP_042211005.1 hypothetical protein
37(1) WP_074108291.1 WP_074108291.1 hypothetical protein
37(1) WP_076309116.1 WP_076309116.1 hypothetical protein
37(1) WP_120112293.1 WP_120112293.1 hypothetical protein
37(1) WP_039306329.1 WP_039306329.1 hypothetical protein
37(1) WP_068536355.1 WP_068536355.1 ABC transporter permease
37(1) WP_074085349.1 WP_074085349.1 hypothetical protein
37(1) WP_154118023.1 WP_154118023.1 hypothetical protein
37(1) WP_239617491.1 WP_239617491.1 hypothetical protein
37(1) WP_238187865.1 WP_238187865.1 hypothetical protein
37(3) WP_076135792.1 WP_076135792.1 hypothetical protein
37(2) WP_042134287.1 WP_042134287.1 hypothetical protein
37(1) WP_127194967.1 WP_127194967.1 hypothetical protein
37(1) WP_189018587.1 WP_189018587.1 DUF1634 domain-containing protein
37(1) WP_253440802.1 WP_253440802.1 ABC transporter permease
37(1) WP_076279376.1 WP_076279376.1 hypothetical protein

38(1) WP_147786239.1 WP_147786239.1 UDP-N-acetylmuramoyl-tripeptide--D-alanyl-D-alanine ligase
38(1) WP_284566225.1 WP_284566225.1 UDP-N-acetylmuramoyl-tripeptide--D-alanyl-D-alanine ligase
38(1) WP_001049601.1 WP_001049601.1 UDP-N-acetylmuramoyl-tripeptide--D-alanyl-D-alanine ligase
38(2) WP_141439360.1 WP_141439360.1 UDP-N-acetylmuramoyl-tripeptide--D-alanyl-D-alanine ligase
38(4) WP_023521811.1 WP_023521811.1 UDP-N-acetylmuramoyl-tripeptide--D-alanyl-D-alanine ligase
38(2) WP_088298124.1 WP_088298124.1 UDP-N-acetylmuramoyl-tripeptide--D-alanyl-D-alanine ligase
38(2) WP_157827840.1 WP_157827840.1 UDP-N-acetylmuramoyl-tripeptide--D-alanyl-D-alanine ligase
38(1) WP_222655360.1 WP_222655360.1 UDP-N-acetylmuramoyl-tripeptide--D-alanyl-D-alanine ligase
38(1) WP_236804580.1 WP_236804580.1 UDP-N-acetylmuramoyl-tripeptide--D-alanyl-D-alanine ligase
38(4) WP_002170266.1 WP_002170266.1 UDP-N-acetylmuramoyl-tripeptide--D-alanyl-D-alanine ligase
38(1) WP_049770079.1 WP_049770079.1 UDP-N-acetylmuramoyl-tripeptide--D-alanyl-D-alanine ligase
38(1) WP_166704988.1 WP_166704988.1 UDP-N-acetylmuramoyl-tripeptide--D-alanyl-D-alanine ligase
38(2) WP_140155657.1 WP_140155657.1 UDP-N-acetylmuramoyl-tripeptide--D-alanyl-D-alanine ligase
38(4) WP_016512884.1 WP_016512884.1 UDP-N-acetylmuramoyl-tripeptide--D-alanyl-D-alanine ligase
38(2) WP_200152288.1 WP_200152288.1 UDP-N-acetylmuramoyl-tripeptide--D-alanyl-D-alanine ligase
38(1) WP_141541881.1 WP_141541881.1 UDP-N-acetylmuramoyl-tripeptide--D-alanyl-D-alanine ligase
38(1) WP_080451211.1 WP_080451211.1 UDP-N-acetylmuramoyl-tripeptide--D-alanyl-D-alanine ligase
38(1) WP_098235619.1 WP_098235619.1 UDP-N-acetylmuramoyl-tripeptide--D-alanyl-D-alanine ligase
38(3) WP_141542758.1 WP_141542758.1 UDP-N-acetylmuramoyl-tripeptide--D-alanyl-D-alanine ligase
38(1) WP_000615529.1 WP_000615529.1 UDP-N-acetylmuramoyl-tripeptide--D-alanyl-D-alanine ligase
38(1) WP_098841817.1 WP_098841817.1 UDP-N-acetylmuramoyl-tripeptide--D-alanyl-D-alanine ligase
38(1) WP_141559429.1 WP_141559429.1 UDP-N-acetylmuramoyl-tripeptide--D-alanyl-D-alanine ligase
38(1) WP_273061455.1 WP_273061455.1 UDP-N-acetylmuramoyl-tripeptide--D-alanyl-D-alanine ligase
38(2) WP_069355507.1 WP_069355507.1 UDP-N-acetylmuramoyl-tripeptide--D-alanyl-D-alanine ligase
38(1) WP_308902525.1 WP_308902525.1 UDP-N-acetylmuramoyl-tripeptide--D-alanyl-D-alanine ligase
38(1) WP_242261452.1 WP_242261452.1 UDP-N-acetylmuramoyl-tripeptide--D-alanyl-D-alanine ligase
38(1) WP_088119570.1 WP_088119570.1 UDP-N-acetylmuramoyl-tripeptide--D-alanyl-D-alanine ligase
38(1) WP_144476575.1 WP_144476575.1 UDP-N-acetylmuramoyl-tripeptide--D-alanyl-D-alanine ligase
38(1) WP_262748768.1 WP_262748768.1 UDP-N-acetylmuramoyl-tripeptide--D-alanyl-D-alanine ligase
38(1) WP_151630609.1 WP_151630609.1 UDP-N-acetylmuramoyl-tripeptide--D-alanyl-D-alanine ligase
38(3) WP_061660799.1 WP_061660799.1 UDP-N-acetylmuramoyl-tripeptide--D-alanyl-D-alanine ligase
38(1) WP_144506826.1 WP_144506826.1 UDP-N-acetylmuramoyl-tripeptide--D-alanyl-D-alanine ligase
38(1) WP_147807658.1 WP_147807658.1 UDP-N-acetylmuramoyl-tripeptide--D-alanyl-D-alanine ligase
38(1) WP_270716419.1 WP_270716419.1 UDP-N-acetylmuramoyl-tripeptide--D-alanyl-D-alanine ligase
38(1) WP_140350460.1 WP_140350460.1 UDP-N-acetylmuramoyl-tripeptide--D-alanyl-D-alanine ligase
38(1) WP_270691242.1 WP_270691242.1 UDP-N-acetylmuramoyl-tripeptide--D-alanyl-D-alanine ligase
38(1) WP_149238902.1 WP_149238902.1 UDP-N-acetylmuramoyl-tripeptide--D-alanyl-D-alanine ligase
38(1) WP_142327657.1 WP_142327657.1 UDP-N-acetylmuramoyl-tripeptide--D-alanyl-D-alanine ligase
38(2) WP_141540991.1 WP_141540991.1 UDP-N-acetylmuramoyl-tripeptide--D-alanyl-D-alanine ligase
38(1) WP_074565307.1 WP_074565307.1 UDP-N-acetylmuramoyl-tripeptide--D-alanyl-D-alanine ligase
38(1) WP_116777314.1 WP_116777314.1 UDP-N-acetylmuramoyl-tripeptide--D-alanyl-D-alanine ligase

| | | |
|---|---|---|
| 38(1) | WP_262729424.1 | WP_262729424.1 UDP-N-acetylmuramoyl-tripeptide--D-alanyl-D-alanine ligase |
| 38(2) | WP_141499183.1 | WP_141499183.1 UDP-N-acetylmuramoyl-tripeptide--D-alanyl-D-alanine ligase |
| 38(1) | WP_142310347.1 | WP_142310347.1 UDP-N-acetylmuramoyl-tripeptide--D-alanyl-D-alanine ligase |
| 38(1) | WP_139120446.1 | WP_139120446.1 UDP-N-acetylmuramoyl-tripeptide--D-alanyl-D-alanine ligase |
| 38(1) | WP_199677616.1 | WP_199677616.1 UDP-N-acetylmuramoyl-tripeptide--D-alanyl-D-alanine ligase |
| 38(1) | WP_141555541.1 | WP_141555541.1 UDP-N-acetylmuramoyl-tripeptide--D-alanyl-D-alanine ligase |
| 38(1) | WP_257808064.1 | WP_257808064.1 UDP-N-acetylmuramoyl-tripeptide--D-alanyl-D-alanine ligase |
| 38(1) | WP_040963159.1 | WP_040963159.1 UDP-N-acetylmuramoyl-tripeptide--D-alanyl-D-alanine ligase |
| 38(1) | WP_080444454.1 | WP_080444454.1 UDP-N-acetylmuramoyl-tripeptide--D-alanyl-D-alanine ligase |
| 38(2) | WP_002163677.1 | WP_002163677.1 UDP-N-acetylmuramoyl-tripeptide--D-alanyl-D-alanine ligase |
| 38(1) | WP_000615533.1 | WP_000615533.1 UDP-N-acetylmuramoyl-tripeptide--D-alanyl-D-alanine ligase |
| 38(1) | WP_142387344.1 | WP_142387344.1 UDP-N-acetylmuramoyl-tripeptide--D-alanyl-D-alanine ligase |
| 38(1) | WP_270728807.1 | WP_270728807.1 UDP-N-acetylmuramoyl-tripeptide--D-alanyl-D-alanine ligase |
| 38(1) | WP_083322703.1 | WP_083322703.1 UDP-N-acetylmuramoyl-tripeptide--D-alanyl-D-alanine ligase |
| 38(1) | WP_270699375.1 | WP_270699375.1 UDP-N-acetylmuramoyl-tripeptide--D-alanyl-D-alanine ligase |
| 38(1) | WP_139895512.1 | WP_139895512.1 UDP-N-acetylmuramoyl-tripeptide--D-alanyl-D-alanine ligase |
| 38(1) | WP_098250522.1 | WP_098250522.1 UDP-N-acetylmuramoyl-tripeptide--D-alanyl-D-alanine ligase |
| 38(2) | WP_002141588.1 | WP_002141588.1 UDP-N-acetylmuramoyl-tripeptide--D-alanyl-D-alanine ligase |
| 38(1) | WP_120449934.1 | WP_120449934.1 UDP-N-acetylmuramoyl-tripeptide--D-alanyl-D-alanine ligase |
| 38(1) | WP_098355764.1 | WP_098355764.1 UDP-N-acetylmuramoyl-tripeptide--D-alanyl-D-alanine ligase |
| 38(1) | WP_130983656.1 | WP_130983656.1 UDP-N-acetylmuramoyl-tripeptide--D-alanyl-D-alanine ligase |
| 38(3) | WP_142314201.1 | WP_142314201.1 UDP-N-acetylmuramoyl-tripeptide--D-alanyl-D-alanine ligase |
| 38(2) | WP_080776499.1 | WP_080776499.1 UDP-N-acetylmuramoyl-tripeptide--D-alanyl-D-alanine ligase |
| 38(1) | WP_063223273.1 | WP_063223273.1 UDP-N-acetylmuramoyl-tripeptide--D-alanyl-D-alanine ligase |
| 38(1) | WP_098629189.1 | WP_098629189.1 UDP-N-acetylmuramoyl-tripeptide--D-alanyl-D-alanine ligase |
| 38(2) | WP_131231110.1 | WP_131231110.1 UDP-N-acetylmuramoyl-tripeptide--D-alanyl-D-alanine ligase |
| 38(3) | WP_142391901.1 | WP_142391901.1 UDP-N-acetylmuramoyl-tripeptide--D-alanyl-D-alanine ligase |
| 38(1) | WP_262725730.1 | WP_262725730.1 UDP-N-acetylmuramoyl-tripeptide--D-alanyl-D-alanine ligase |
| 38(1) | WP_064473941.1 | WP_064473941.1 UDP-N-acetylmuramoyl-tripeptide--D-alanyl-D-alanine ligase |
| 38(5) | WP_065483398.1 | WP_065483398.1 UDP-N-acetylmuramoyl-tripeptide--D-alanyl-D-alanine ligase |
| 38(1) | WP_127812906.1 | WP_127812906.1 UDP-N-acetylmuramoyl-tripeptide--D-alanyl-D-alanine ligase |
| 38(1) | WP_002148339.1 | WP_002148339.1 UDP-N-acetylmuramoyl-tripeptide--D-alanyl-D-alanine ligase |
| 38(1) | WP_144473245.1 | WP_144473245.1 UDP-N-acetylmuramoyl-tripeptide--D-alanyl-D-alanine ligase |
| 38(1) | WP_002031629.1 | WP_002031629.1 UDP-N-acetylmuramoyl-tripeptide--D-alanyl-D-alanine ligase |
| 38(2) | WP_142328641.1 | WP_142328641.1 UDP-N-acetylmuramoyl-tripeptide--D-alanyl-D-alanine ligase |
| 38(1) | WP_142322063.1 | WP_142322063.1 UDP-N-acetylmuramoyl-tripeptide--D-alanyl-D-alanine ligase |
| 38(1) | WP_141559745.1 | WP_141559745.1 UDP-N-acetylmuramoyl-tripeptide--D-alanyl-D-alanine ligase |
| 38(1) | WP_277480682.1 | WP_277480682.1 UDP-N-acetylmuramoyl-tripeptide--D-alanyl-D-alanine ligase |
| 38(1) | WP_141519887.1 | WP_141519887.1 UDP-N-acetylmuramoyl-tripeptide--D-alanyl-D-alanine ligase |
| 38(1) | WP_143881068.1 | WP_143881068.1 UDP-N-acetylmuramoyl-tripeptide--D-alanyl-D-alanine ligase |
| 38(1) | WP_260869609.1 | WP_260869609.1 UDP-N-acetylmuramoyl-tripeptide--D-alanyl-D-alanine ligase |
| 38(1) | WP_098670170.1 | WP_098670170.1 UDP-N-acetylmuramoyl-tripeptide--D-alanyl-D-alanine ligase |
| 38(1) | WP_142305880.1 | WP_142305880.1 UDP-N-acetylmuramoyl-tripeptide--D-alanyl-D-alanine ligase |
| 38(1) | WP_137052041.1 | WP_137052041.1 UDP-N-acetylmuramoyl-tripeptide--D-alanyl-D-alanine ligase |
| 38(2) | WP_212956330.1 | WP_212956330.1 UDP-N-acetylmuramoyl-tripeptide--D-alanyl-D-alanine ligase |
| 38(2) | WP_141536472.1 | WP_141536472.1 UDP-N-acetylmuramoyl-tripeptide--D-alanyl-D-alanine ligase |
| 38(1) | WP_157418141.1 | WP_157418141.1 UDP-N-acetylmuramoyl-tripeptide--D-alanyl-D-alanine ligase |
| 38(1) | WP_066092021.1 | WP_066092021.1 Mur ligase family protein |
| 38(1) | WP_033710609.1 | WP_033710609.1 UDP-N-acetylmuramoyl-tripeptide--D-alanyl-D-alanine ligase |
| 38(3) | WP_003266747.1 | WP_003266747.1 UDP-N-acetylmuramoyl-tripeptide--D-alanyl-D-alanine ligase |
| 38(1) | WP_137013294.1 | WP_137013294.1 UDP-N-acetylmuramoyl-tripeptide--D-alanyl-D-alanine ligase |
| 38(1) | WP_073535976.1 | WP_073535976.1 UDP-N-acetylmuramoyl-tripeptide--D-alanyl-D-alanine ligase |

38(1) WP_306007834.1 WP_306007834.1 UDP-N-acetylmuramoyl-tripeptide--D-alanyl-D-alanine ligase
38(1) WP_087995247.1 WP_087995247.1 UDP-N-acetylmuramoyl-tripeptide--D-alanyl-D-alanine ligase
38(1) WP_222661109.1 WP_222661109.1 UDP-N-acetylmuramoyl-tripeptide--D-alanyl-D-alanine ligase
38(1) WP_141524400.1 WP_141524400.1 UDP-N-acetylmuramoyl-tripeptide--D-alanyl-D-alanine ligase
38(1) WP_199795350.1 WP_199795350.1 UDP-N-acetylmuramoyl-tripeptide--D-alanyl-D-alanine ligase
38(1) WP_139365524.1 WP_139365524.1 UDP-N-acetylmuramoyl-tripeptide--D-alanyl-D-alanine ligase
38(1) WP_242264962.1 WP_242264962.1 UDP-N-acetylmuramoyl-tripeptide--D-alanyl-D-alanine ligase
38(1) WP_242251770.1 WP_242251770.1 UDP-N-acetylmuramoyl-tripeptide--D-alanyl-D-alanine ligase
38(1) WP_199695250.1 WP_199695250.1 UDP-N-acetylmuramoyl-tripeptide--D-alanyl-D-alanine ligase
38(1) WP_087956289.1 WP_087956289.1 UDP-N-acetylmuramoyl-tripeptide--D-alanyl-D-alanine ligase

39(2) WP_090949178.1 WP_090949178.1 Gfo/Idh/MocA family oxidoreductase
39(1) WP_150367637.1 WP_150367637.1 Gfo/Idh/MocA family oxidoreductase
39(1) WP_199613916.1 WP_199613916.1 Gfo/Idh/MocA family oxidoreductase
39(1) WP_138226016.1 WP_138226016.1 Gfo/Idh/MocA family oxidoreductase
39(1) WP_145318667.1 WP_145318667.1 Gfo/Idh/MocA family oxidoreductase
39(1) WP_036609186.1 WP_036609186.1 Gfo/Idh/MocA family oxidoreductase
39(1) WP_188593170.1 WP_188593170.1 Gfo/Idh/MocA family oxidoreductase
39(1) WP_017692333.1 WP_017692333.1 Gfo/Idh/MocA family oxidoreductase
39(2) WP_076249771.1 WP_076249771.1 Gfo/Idh/MocA family oxidoreductase
39(1) WP_076322104.1 WP_076322104.1 Gfo/Idh/MocA family oxidoreductase
39(1) WP_213532667.1 WP_213532667.1 Gfo/Idh/MocA family oxidoreductase
39(1) WP_184183063.1 WP_184183063.1 Gfo/Idh/MocA family oxidoreductase
39(1) WP_046522771.1 WP_046522771.1 Gfo/Idh/MocA family oxidoreductase
39(1) WP_124114125.1 WP_124114125.1 Gfo/Idh/MocA family oxidoreductase
39(1) WP_062326157.1 WP_062326157.1 Gfo/Idh/MocA family oxidoreductase
39(1) WP_191798538.1 WP_191798538.1 Gfo/Idh/MocA family oxidoreductase
39(1) WP_053361037.1 WP_053361037.1 Gfo/Idh/MocA family oxidoreductase
39(1) WP_253300501.1 WP_253300501.1 Gfo/Idh/MocA family oxidoreductase
39(2) WP_098746841.1 WP_098746841.1 Gfo/Idh/MocA family oxidoreductase
39(1) WP_285748190.1 WP_285748190.1 Gfo/Idh/MocA family oxidoreductase
39(1) WP_285079434.1 WP_285079434.1 Gfo/Idh/MocA family oxidoreductase
39(1) WP_076318555.1 WP_076318555.1 Gfo/Idh/MocA family oxidoreductase
39(1) WP_099853186.1 WP_099853186.1 Gfo/Idh/MocA family oxidoreductase
39(1) WP_046522772.1 WP_046522772.1 Gfo/Idh/MocA family oxidoreductase
39(1) WP_077567006.1 WP_077567006.1 Gfo/Idh/MocA family oxidoreductase
39(1) WP_205053317.1 WP_205053317.1 Gfo/Idh/MocA family oxidoreductase
39(1) WP_251035375.1 WP_251035375.1 Gfo/Idh/MocA family oxidoreductase
39(1) WP_125084795.1 WP_125084795.1 Gfo/Idh/MocA family oxidoreductase
39(1) WP_054402898.1 WP_054402898.1 Gfo/Idh/MocA family oxidoreductase
39(1) WP_123067125.1 WP_123067125.1 Gfo/Idh/MocA family oxidoreductase
39(1) WP_282321814.1 WP_282321814.1 Gfo/Idh/MocA family oxidoreductase
39(1) WP_169505063.1 WP_169505063.1 Gfo/Idh/MocA family oxidoreductase
39(1) WP_127592418.1 WP_127592418.1 Gfo/Idh/MocA family oxidoreductase
39(1) WP_091155401.1 WP_091155401.1 Gfo/Idh/MocA family oxidoreductase
39(1) WP_024634133.1 WP_024634133.1 Gfo/Idh/MocA family oxidoreductase
39(1) WP_145046471.1 WP_145046471.1 Gfo/Idh/MocA family oxidoreductase
39(1) WP_111273184.1 WP_111273184.1 Gfo/Idh/MocA family oxidoreductase
39(2) WP_017689060.1 WP_017689060.1 Gfo/Idh/MocA family oxidoreductase
39(1) WP_145153690.1 WP_145153690.1 Gfo/Idh/MocA family oxidoreductase
39(1) WP_042210984.1 WP_042210984.1 Gfo/Idh/MocA family oxidoreductase
39(5) WP_006211191.1 WP_006211191.1 Gfo/Idh/MocA family oxidoreductase
39(2) WP_090896213.1 WP_090896213.1 Gfo/Idh/MocA family oxidoreductase
39(1) WP_127543689.1 WP_127543689.1 Gfo/Idh/MocA family oxidoreductase
39(1) WP_090993639.1 WP_090993639.1 Gfo/Idh/MocA family oxidoreductase
39(2) WP_013370829.1 WP_013370829.1 Gfo/Idh/MocA family oxidoreductase
39(2) WP_042134256.1 WP_042134256.1 Gfo/Idh/MocA family oxidoreductase
39(1) WP_071220215.1 WP_071220215.1 Gfo/Idh/MocA family oxidoreductase
39(1) WP_090908288.1 WP_090908288.1 Gfo/Idh/MocA family oxidoreductase
39(1) WP_018883259.1 WP_018883259.1 Gfo/Idh/MocA family oxidoreductase
39(1) WP_236417232.1 WP_236417232.1 Gfo/Idh/MocA family oxidoreductase
39(1) WP_293202292.1 WP_293202292.1 Gfo/Idh/MocA family oxidoreductase
39(1) WP_074085363.1 WP_074085363.1 Gfo/Idh/MocA family oxidoreductase
39(1) WP_007128616.1 WP_007128616.1 Gfo/Idh/MocA family oxidoreductase
39(1) WP_109999899.1 WP_109999899.1 Gfo/Idh/MocA family oxidoreductase
39(1) WP_099517116.1 WP_099517116.1 Gfo/Idh/MocA family oxidoreductase
39(3) WP_062833889.1 WP_062833889.1 Gfo/Idh/MocA family oxidoreductase
39(1) WP_076242349.1 WP_076242349.1 Gfo/Idh/MocA family oxidoreductase
39(3) WP_095266206.1 WP_095266206.1 Gfo/Idh/MocA family oxidoreductase
39(1) WP_247902253.1 WP_247902253.1 Gfo/Idh/MocA family oxidoreductase
39(1) WP_301879135.1 WP_301879135.1 Gfo/Idh/MocA family oxidoreductase
39(1) WP_211083020.1 WP_211083020.1 Gfo/Idh/MocA family oxidoreductase
39(1) WP_154886312.1 WP_154886312.1 Gfo/Idh/MocA family oxidoreductase
39(1) WP_047842542.1 WP_047842542.1 Gfo/Idh/MocA family oxidoreductase
39(1) WP_076210944.1 WP_076210944.1 Gfo/Idh/MocA family oxidoreductase
39(1) WP_090805484.1 WP_090805484.1 Gfo/Idh/MocA family oxidoreductase
39(1) WP_042134252.1 WP_042134252.1 Gfo/Idh/MocA family oxidoreductase
39(1) WP_095358540.1 WP_095358540.1 Gfo/Idh/MocA family oxidoreductase
39(1) WP_301249705.1 WP_301249705.1 Gfo/Idh/MocA family oxidoreductase
39(1) WP_024630337.1 WP_024630337.1 Gfo/Idh/MocA family oxidoreductase
39(1) WP_264929198.1 WP_264929198.1 Gfo/Idh/MocA family oxidoreductase
39(1) WP_090722894.1 WP_090722894.1 Gfo/Idh/MocA family oxidoreductase
39(1) WP_099517115.1 WP_099517115.1 Gfo/Idh/MocA family oxidoreductase
39(1) WP_152826374.1 WP_152826374.1 Gfo/Idh/MocA family oxidoreductase
39(1) WP_074108300.1 WP_074108300.1 Gfo/Idh/MocA family oxidoreductase
39(1) WP_251583655.1 WP_251583655.1 Gfo/Idh/MocA family oxidoreductase
39(1) WP_110822555.1 WP_110822555.1 Gfo/Idh/MocA family oxidoreductase
39(1) WP_110892952.1 WP_110892952.1 Gfo/Idh/MocA family oxidoreductase
39(1) WP_150364182.1 WP_150364182.1 Gfo/Idh/MocA family oxidoreductase
39(1) WP_076311043.1 WP_076311043.1 Gfo/Idh/MocA family oxidoreductase
39(1) WP_113058726.1 WP_113058726.1 Gfo/Idh/MocA family oxidoreductase
39(1) WP_053361038.1 WP_053361038.1 Gfo/Idh/MocA family oxidoreductase
39(1) WP_076208835.1 WP_076208835.1 Gfo/Idh/MocA family oxidoreductase
39(1) WP_090921034.1 WP_090921034.1 Gfo/Idh/MocA family oxidoreductase

| | | |
|---|---|---|
| 39(1) | WP_023957749.1 | WP_023957749.1 Gfo/Idh/MocA family oxidoreductase |
| 39(2) | WP_072732446.1 | WP_072732446.1 Gfo/Idh/MocA family oxidoreductase |
| 39(1) | WP_110934087.1 | WP_110934087.1 Gfo/Idh/MocA family oxidoreductase |
| 39(1) | WP_053783897.1 | WP_053783897.1 Gfo/Idh/MocA family oxidoreductase |
| 39(1) | WP_064639005.1 | WP_064639005.1 Gfo/Idh/MocA family oxidoreductase |
| 39(1) | WP_153978074.1 | WP_153978074.1 Gfo/Idh/MocA family oxidoreductase |
| 39(1) | WP_253440818.1 | WP_253440818.1 Gfo/Idh/MocA family oxidoreductase |
| 39(1) | WP_193721880.1 | WP_193721880.1 Gfo/Idh/MocA family oxidoreductase |
| 39(1) | WP_127536188.1 | WP_127536188.1 Gfo/Idh/MocA family oxidoreductase |
| 39(1) | WP_128101494.1 | WP_128101494.1 Gfo/Idh/MocA family oxidoreductase |
| 39(1) | WP_192266478.1 | WP_192266478.1 Gfo/Idh/MocA family oxidoreductase |
| 39(1) | WP_307537566.1 | WP_307537566.1 Gfo/Idh/MocA family oxidoreductase |
| 39(1) | WP_194543490.1 | WP_194543490.1 Gfo/Idh/MocA family oxidoreductase |
| 39(1) | WP_111619116.1 | WP_111619116.1 Gfo/Idh/MocA family oxidoreductase |
| 39(1) | WP_101313446.1 | WP_101313446.1 Gfo/Idh/MocA family oxidoreductase |
| 39(1) | WP_006720299.1 | WP_006720299.1 Gfo/Idh/MocA family oxidoreductase |
| 39(1) | WP_260065270.1 | WP_260065270.1 Gfo/Idh/MocA family oxidoreductase |
| 39(1) | WP_074094621.1 | WP_074094621.1 Gfo/Idh/MocA family oxidoreductase |
| 39(1) | WP_289373780.1 | WP_289373780.1 Gfo/Idh/MocA family oxidoreductase |
| 39(1) | WP_216531941.1 | WP_216531941.1 Gfo/Idh/MocA family oxidoreductase |
| 39(1) | WP_113052558.1 | WP_113052558.1 Gfo/Idh/MocA family oxidoreductase |
| 39(1) | WP_095287650.1 | WP_095287650.1 Gfo/Idh/MocA family oxidoreductase |
| 39(1) | WP_119847560.1 | WP_119847560.1 Gfo/Idh/MocA family oxidoreductase |
| 39(1) | WP_211745818.1 | WP_211745818.1 Gfo/Idh/MocA family oxidoreductase |
| 39(1) | WP_308723535.1 | WP_308723535.1 Gfo/Idh/MocA family oxidoreductase |
| 39(2) | WP_145329916.1 | WP_145329916.1 Gfo/Idh/MocA family oxidoreductase |
| 39(1) | WP_183600883.1 | WP_183600883.1 Gfo/Idh/MocA family oxidoreductase |
| 39(2) | WP_036670444.1 | WP_036670444.1 Gfo/Idh/MocA family oxidoreductase |
| 39(1) | WP_307214311.1 | WP_307214311.1 Gfo/Idh/MocA family oxidoreductase |
| 39(1) | WP_145408773.1 | WP_145408773.1 Gfo/Idh/MocA family oxidoreductase |
| 39(1) | WP_272035342.1 | WP_272035342.1 Gfo/Idh/MocA family oxidoreductase |
| 39(1) | WP_056700811.1 | WP_056700811.1 Gfo/Idh/MocA family oxidoreductase |
| 39(1) | WP_047909604.1 | WP_047909604.1 Gfo/Idh/MocA family oxidoreductase |
| 39(1) | WP_305755022.1 | WP_305755022.1 Gfo/Idh/MocA family oxidoreductase |
| 39(1) | WP_100537966.1 | WP_100537966.1 Gfo/Idh/MocA family oxidoreductase |
| 39(1) | WP_307413587.1 | WP_307413587.1 Gfo/Idh/MocA family oxidoreductase |
| 39(1) | WP_249897330.1 | WP_249897330.1 Gfo/Idh/MocA family oxidoreductase |
| 39(1) | WP_243896002.1 | WP_243896002.1 Gfo/Idh/MocA family oxidoreductase |
| 39(1) | WP_210095101.1 | WP_210095101.1 Gfo/Idh/MocA family oxidoreductase |

| | | |
|---|---|---|
| 40(1) | WP_145318670.1 | WP_145318670.1 helix-turn-helix transcriptional regulator |
| 40(1) | WP_236417231.1 | WP_236417231.1 helix-turn-helix transcriptional regulator |
| 40(13) | WP_063563231.1 | WP_063563231.1 helix-turn-helix transcriptional regulator |
| 40(1) | WP_208919058.1 | WP_208919058.1 helix-turn-helix transcriptional regulator |
| 40(3) | WP_024630336.1 | WP_024630336.1 helix-turn-helix transcriptional regulator |
| 40(2) | WP_053783896.1 | WP_053783896.1 helix-turn-helix transcriptional regulator |
| 40(3) | WP_095266205.1 | WP_095266205.1 helix-turn-helix transcriptional regulator |
| 40(1) | WP_307413589.1 | WP_307413589.1 helix-turn-helix transcriptional regulator |
| 40(1) | WP_009590710.1 | WP_009590710.1 helix-turn-helix transcriptional regulator |
| 40(1) | WP_301249704.1 | WP_301249704.1 helix-turn-helix transcriptional regulator |
| 40(1) | WP_184191565.1 | WP_184191565.1 helix-turn-helix transcriptional regulator |
| 40(1) | WP_272035340.1 | WP_272035340.1 helix-turn-helix transcriptional regulator |
| 40(1) | WP_077567007.1 | WP_077567007.1 helix-turn-helix transcriptional regulator |
| 40(1) | WP_090908287.1 | WP_090908287.1 helix-turn-helix transcriptional regulator |
| 40(1) | WP_305755023.1 | WP_305755023.1 helix-turn-helix transcriptional regulator |
| 40(1) | WP_205053318.1 | WP_205053318.1 helix-turn-helix transcriptional regulator |
| 40(1) | WP_169505062.1 | WP_169505062.1 helix-turn-helix transcriptional regulator |
| 40(1) | WP_017689061.1 | WP_017689061.1 helix-turn-helix transcriptional regulator |
| 40(2) | WP_018883260.1 | WP_018883260.1 helix-turn-helix transcriptional regulator |
| 40(1) | WP_023957747.1 | WP_023957747.1 helix-turn-helix transcriptional regulator |
| 40(20) | WP_036609188.1 | WP_036609188.1 helix-turn-helix transcriptional regulator |
| 40(2) | WP_128101493.1 | WP_128101493.1 helix-turn-helix transcriptional regulator |
| 40(2) | WP_007128617.1 | WP_007128617.1 helix-turn-helix transcriptional regulator |
| 40(1) | WP_053492748.1 | WP_053492748.1 helix-turn-helix transcriptional regulator |
| 40(2) | WP_110892954.1 | WP_110892954.1 helix-turn-helix transcriptional regulator |
| 40(1) | WP_191798537.1 | WP_191798537.1 helix-turn-helix transcriptional regulator |
| 40(2) | WP_076311044.1 | WP_076311044.1 helix-turn-helix transcriptional regulator |
| 40(3) | WP_056700809.1 | WP_056700809.1 helix-turn-helix transcriptional regulator |
| 40(14) | WP_019392100.1 | WP_019392100.1 helix-turn-helix transcriptional regulator |
| 40(2) | WP_013370828.1 | WP_013370828.1 helix-turn-helix transcriptional regulator |
| 40(6) | WP_095287651.1 | WP_095287651.1 helix-turn-helix transcriptional regulator |
| 40(1) | WP_113058725.1 | WP_113058725.1 helix-turn-helix transcriptional regulator |
| 40(1) | WP_211745819.1 | WP_211745819.1 helix-turn-helix transcriptional regulator |
| 40(5) | WP_090921036.1 | WP_090921036.1 helix-turn-helix transcriptional regulator |
| 40(1) | WP_301879137.1 | WP_301879137.1 helix-turn-helix transcriptional regulator |
| 40(1) | WP_192266476.1 | WP_192266476.1 helix-turn-helix transcriptional regulator |
| 40(1) | WP_047909605.1 | WP_047909605.1 helix-turn-helix transcriptional regulator |
| 40(1) | WP_223067428.1 | WP_223067428.1 helix-turn-helix transcriptional regulator |
| 40(2) | WP_091155398.1 | WP_091155398.1 helix-turn-helix transcriptional regulator |
| 40(1) | WP_123067124.1 | WP_123067124.1 helix-turn-helix transcriptional regulator |
| 40(7) | WP_013310011.1 | WP_013310011.1 helix-turn-helix transcriptional regulator |
| 40(1) | WP_216531940.1 | WP_216531940.1 helix-turn-helix transcriptional regulator |
| 40(6) | WP_006211192.1 | WP_006211192.1 helix-turn-helix transcriptional regulator |
| 40(2) | WP_138226017.1 | WP_138226017.1 helix-turn-helix transcriptional regulator |
| 40(10) | WP_071220214.1 | WP_071220214.1 helix-turn-helix transcriptional regulator |
| 40(1) | WP_106765947.1 | WP_106765947.1 helix-turn-helix transcriptional regulator |
| 40(2) | WP_213532666.1 | WP_213532666.1 helix-turn-helix transcriptional regulator |

| | | |
|---|---|---|
| 41(1) | WP_098266661.1 | WP_098266661.1 iron-containing alcohol dehydrogenase |
| 41(1) | WP_070805910.1 | WP_070805910.1 alcohol dehydrogenase EutG |
| 41(1) | WP_047842535.1 | WP_047842535.1 aldehyde dehydrogenase family protein |
| 41(1) | WP_289373804.1 | WP_289373804.1 aldehyde dehydrogenase family protein |
| 41(1) | WP_090805470.1 | WP_090805470.1 aldehyde dehydrogenase family protein |
| 41(1) | WP_074094611.1 | WP_074094611.1 aldehyde dehydrogenase family protein |
| 41(1) | WP_060633126.1 | WP_060633126.1 alcohol dehydrogenase EutG |
| 41(1) | WP_213427002.1 | WP_213427002.1 L-glutamate gamma-semialdehyde dehydrogenase |
| 41(1) | WP_111153312.1 | WP_111153312.1 L-glutamate gamma-semialdehyde dehydrogenase |
| 41(1) | WP_247902241.1 | WP_247902241.1 aldehyde dehydrogenase family protein |
| 41(1) | WP_091031771.1 | WP_091031771.1 aldehyde dehydrogenase family protein |
| 41(1) | WP_006674878.1 | WP_006674878.1 L-glutamate gamma-semialdehyde dehydrogenase |
| 41(1) | WP_274566438.1 | WP_274566438.1 aldehyde dehydrogenase family protein |
| 41(1) | WP_079346827.1 | WP_079346827.1 aldehyde dehydrogenase family protein |
| 41(1) | WP_076330719.1 | WP_076330719.1 aldehyde dehydrogenase family protein |
| 41(1) | WP_019424162.1 | WP_019424162.1 aldehyde dehydrogenase EutE |

| | | |
|---|---|---|
| 41(1) | WP_307537587.1 | WP_307537587.1 aldehyde dehydrogenase family protein |
| 41(1) | WP_145318707.1 | WP_145318707.1 aldehyde dehydrogenase family protein |
| 41(3) | WP_065228774.1 | WP_065228774.1 iron-containing alcohol dehydrogenase |
| 41(1) | WP_090896177.1 | WP_090896177.1 aldehyde dehydrogenase family protein |
| 41(1) | WP_036609205.1 | WP_036609205.1 aldehyde dehydrogenase family protein |
| 41(1) | WP_145329903.1 | WP_145329903.1 aldehyde dehydrogenase family protein |
| 41(1) | WP_272035299.1 | WP_272035299.1 aldehyde dehydrogenase family protein |
| 41(1) | WP_144660549.1 | WP_144660549.1 alcohol dehydrogenase EutG |
| 41(2) | WP_109999888.1 | WP_109999888.1 aldehyde dehydrogenase family protein |
| 41(1) | WP_144564319.1 | WP_144564319.1 alcohol dehydrogenase EutG |
| 41(1) | WP_127543702.1 | WP_127543702.1 aldehyde dehydrogenase family protein |
| 41(1) | WP_285079443.1 | WP_285079443.1 aldehyde dehydrogenase family protein |
| 41(1) | WP_076208842.1 | WP_076208842.1 aldehyde dehydrogenase family protein |
| 41(2) | WP_076249777.1 | WP_076249777.1 aldehyde dehydrogenase family protein |
| 41(1) | WP_079694650.1 | WP_079694650.1 aldehyde dehydrogenase family protein |
| 41(1) | WP_124114116.1 | WP_124114116.1 aldehyde dehydrogenase family protein |
| 41(1) | WP_091017198.1 | WP_091017198.1 aldehyde dehydrogenase family protein |
| 41(1) | WP_099853202.1 | WP_099853202.1 aldehyde dehydrogenase family protein |
| 41(1) | WP_110822561.1 | WP_110822561.1 aldehyde dehydrogenase family protein |
| 41(1) | WP_133378784.1 | WP_133378784.1 L-glutamate gamma-semialdehyde dehydrogenase |
| 41(5) | WP_048552329.1 | WP_048552329.1 alcohol dehydrogenase EutG |
| 41(1) | WP_264929211.1 | WP_264929211.1 aldehyde dehydrogenase family protein |
| 41(1) | WP_307589619.1 | WP_307589619.1 bifunctional acetaldehyde-CoA/alcohol dehydrogenase |
| 41(2) | WP_087956637.1 | WP_087956637.1 alcohol dehydrogenase EutG |
| 41(1) | WP_071724740.1 | WP_071724740.1 alcohol dehydrogenase EutG |
| 41(1) | WP_071719969.1 | WP_071719969.1 iron-containing alcohol dehydrogenase |
| 41(1) | WP_076318549.1 | WP_076318549.1 aldehyde dehydrogenase family protein |
| 41(1) | WP_149190452.1 | WP_149190452.1 iron-containing alcohol dehydrogenase |
| 41(1) | WP_072732440.1 | WP_072732440.1 aldehyde dehydrogenase family protein |
| 41(1) | WP_098666656.1 | WP_098666656.1 alcohol dehydrogenase EutG |
| 41(1) | WP_262725738.1 | WP_262725738.1 alcohol dehydrogenase EutG |
| 41(2) | WP_112331601.1 | WP_112331601.1 aldehyde dehydrogenase EutE |
| 41(1) | WP_094936485.1 | WP_094936485.1 aldehyde dehydrogenase family protein |
| 41(1) | WP_150367643.1 | WP_150367643.1 aldehyde dehydrogenase family protein |
| 41(2) | WP_087433591.1 | WP_087433591.1 aldehyde dehydrogenase EutE |
| 41(1) | WP_052154015.1 | WP_052154015.1 aldehyde dehydrogenase family protein |
| 41(2) | WP_001158502.1 | WP_001158502.1 alcohol dehydrogenase EutG |
| 41(1) | WP_024630322.1 | WP_024630322.1 aldehyde dehydrogenase family protein |
| 41(1) | WP_307192887.1 | WP_307192887.1 CoA-acylating methylmalonate-semialdehyde dehydrogenase |
| 41(1) | WP_289392070.1 | WP_289392070.1 aldehyde dehydrogenase family protein |
| 41(1) | WP_123067111.1 | WP_123067111.1 aldehyde dehydrogenase family protein |
| 41(1) | WP_145046447.1 | WP_145046447.1 aldehyde dehydrogenase family protein |
| 41(1) | WP_281477388.1 | WP_281477388.1 L-glutamate gamma-semialdehyde dehydrogenase |
| 41(1) | WP_272045603.1 | WP_272045603.1 L-glutamate gamma-semialdehyde dehydrogenase |
| 41(2) | WP_088313253.1 | WP_088313253.1 alcohol dehydrogenase EutG |
| 41(4) | WP_088063241.1 | WP_088063241.1 alcohol dehydrogenase EutG |
| 41(1) | WP_111268943.1 | WP_111268943.1 aldehyde dehydrogenase family protein |
| 41(1) | WP_119794349.1 | WP_119794349.1 L-glutamate gamma-semialdehyde dehydrogenase |
| 41(1) | WP_188593160.1 | WP_188593160.1 aldehyde dehydrogenase family protein |
| 41(1) | WP_145412703.1 | WP_145412703.1 aldehyde dehydrogenase family protein |
| 41(1) | WP_017689077.1 | WP_017689077.1 aldehyde dehydrogenase family protein |
| 41(2) | WP_001158667.1 | WP_001158667.1 iron-containing alcohol dehydrogenase |
| 41(1) | WP_100529432.1 | WP_100529432.1 aldehyde dehydrogenase family protein |
| 41(1) | WP_062833877.1 | WP_062833877.1 aldehyde dehydrogenase family protein |
| 41(1) | WP_149238909.1 | WP_149238909.1 alcohol dehydrogenase EutG |
| 41(3) | WP_168178798.1 | WP_168178798.1 L-glutamate gamma-semialdehyde dehydrogenase |
| 41(1) | WP_277481004.1 | WP_277481004.1 alcohol dehydrogenase EutG |
| 41(1) | WP_293202266.1 | WP_293202266.1 aldehyde dehydrogenase family protein |
| 41(1) | WP_179265140.1 | WP_179265140.1 aldehyde dehydrogenase family protein |
| 41(1) | WP_036670433.1 | WP_036670433.1 aldehyde dehydrogenase family protein |
| 41(1) | WP_063566327.1 | WP_063566327.1 aldehyde dehydrogenase family protein |
| 41(1) | WP_056700780.1 | WP_056700780.1 aldehyde dehydrogenase family protein |
| 41(1) | WP_206094042.1 | WP_206094042.1 aldehyde dehydrogenase EutE |
| 41(1) | WP_095358549.1 | WP_095358549.1 aldehyde dehydrogenase family protein |
| 41(1) | WP_286458026.1 | WP_286458026.1 aldehyde dehydrogenase family protein |
| 41(1) | WP_095287664.1 | WP_095287664.1 aldehyde dehydrogenase family protein |
| 41(1) | WP_307214304.1 | WP_307214304.1 aldehyde dehydrogenase family protein |
| 41(1) | WP_128101483.1 | WP_128101483.1 aldehyde dehydrogenase family protein |
| 41(1) | WP_145408780.1 | WP_145408780.1 aldehyde dehydrogenase family protein |
| 41(1) | WP_042874983.1 | WP_042874983.1 alcohol dehydrogenase EutG |
| 41(1) | WP_087442394.1 | WP_087442394.1 L-glutamate gamma-semialdehyde dehydrogenase |
| 41(1) | WP_233697162.1 | WP_233697162.1 aldehyde dehydrogenase EutE |
| 41(1) | WP_064639014.1 | WP_064639014.1 aldehyde dehydrogenase family protein |
| 41(1) | WP_282321825.1 | WP_282321825.1 aldehyde dehydrogenase family protein |
| 41(1) | WP_113052569.1 | WP_113052569.1 aldehyde dehydrogenase family protein |
| 41(1) | WP_153978065.1 | WP_153978065.1 aldehyde dehydrogenase family protein |
| 41(1) | WP_090921065.1 | WP_090921065.1 aldehyde dehydrogenase family protein |
| 41(1) | WP_053783883.1 | WP_053783883.1 aldehyde dehydrogenase family protein |
| 41(1) | WP_069355518.1 | WP_069355518.1 iron-containing alcohol dehydrogenase |
| 41(1) | WP_242252193.1 | WP_242252193.1 alcohol dehydrogenase EutG |
| 41(1) | WP_301249696.1 | WP_301249696.1 aldehyde dehydrogenase family protein |
| 41(1) | WP_088609123.1 | WP_088609123.1 alcohol dehydrogenase EutG |
| 41(2) | WP_090949163.1 | WP_090949163.1 aldehyde dehydrogenase family protein |
| 41(1) | WP_127536197.1 | WP_127536197.1 aldehyde dehydrogenase family protein |
| 41(1) | WP_197256887.1 | WP_197256887.1 L-glutamate gamma-semialdehyde dehydrogenase |
| 41(1) | WP_087982561.1 | WP_087982561.1 alcohol dehydrogenase EutG |
| 41(1) | WP_229137195.1 | WP_229137195.1 alcohol dehydrogenase EutG |
| 41(1) | WP_053564616.1 | WP_053564616.1 iron-containing alcohol dehydrogenase |
| 41(1) | WP_145146962.1 | WP_145146962.1 aldehyde dehydrogenase family protein |
| 41(1) | WP_307413604.1 | WP_307413604.1 aldehyde dehydrogenase family protein |
| 41(1) | WP_221791101.1 | WP_221791101.1 aldehyde dehydrogenase family protein |
| 41(1) | WP_001158665.1 | WP_001158665.1 iron-containing alcohol dehydrogenase |
| 42(1) | WP_071220213.1 | WP_071220213.1 acyltransferase |
| 42(1) | WP_127536189.1 | WP_127536189.1 acyltransferase |
| 42(1) | WP_058710497.1 | WP_058710497.1 acyltransferase |
| 42(1) | WP_047842541.1 | WP_047842541.1 acyltransferase |
| 42(1) | WP_094936492.1 | WP_094936492.1 acyltransferase |
| 42(1) | WP_247902252.1 | WP_247902252.1 acyltransferase |
| 42(1) | WP_263560238.1 | WP_263560238.1 acyltransferase |
| 42(1) | WP_224839850.1 | WP_224839850.1 acyltransferase family protein |
| 42(1) | WP_208919056.1 | WP_208919056.1 acyltransferase |
| 42(1) | WP_113058724.1 | WP_113058724.1 acyltransferase |

| | | |
|---|---|---|
| 42(1) | WP_009590678.1 | WP_009590678.1 acyltransferase |
| 42(1) | WP_169505061.1 | WP_169505061.1 acyltransferase |
| 42(1) | WP_180361715.1 | WP_180361715.1 acyltransferase |
| 42(1) | WP_192266474.1 | WP_192266474.1 acyltransferase |
| 42(1) | WP_289373784.1 | WP_289373784.1 acyltransferase |
| 42(1) | WP_286458006.1 | WP_286458006.1 acyltransferase |
| 42(1) | WP_280753897.1 | WP_280753897.1 acyltransferase |
| 42(1) | WP_279630379.1 | WP_279630379.1 acyltransferase |
| 42(1) | WP_090805482.1 | WP_090805482.1 acyltransferase |
| 42(1) | WP_213579154.1 | WP_213579154.1 acyltransferase |
| 42(1) | WP_145318673.1 | WP_145318673.1 acyltransferase |
| 42(1) | WP_293202255.1 | WP_293202255.1 acyltransferase |
| 42(1) | WP_076330728.1 | WP_076330728.1 acyltransferase |
| 42(1) | WP_213532664.1 | WP_213532664.1 acyltransferase |
| 42(1) | WP_251537235.1 | WP_251537235.1 acyltransferase |
| 42(1) | WP_145408774.1 | WP_145408774.1 acyltransferase |
| 42(1) | WP_190945695.1 | WP_190945695.1 acyltransferase |
| 42(1) | WP_165980145.1 | WP_165980145.1 acyltransferase |
| 42(1) | WP_192571325.1 | WP_192571325.1 acyltransferase |
| 42(1) | WP_040101523.1 | WP_040101523.1 acyltransferase |
| 42(1) | WP_220684647.1 | WP_220684647.1 acyltransferase |
| 42(1) | WP_119847559.1 | WP_119847559.1 acyltransferase |
| 42(1) | WP_307413591.1 | WP_307413591.1 acyltransferase |
| 42(1) | WP_152826372.1 | WP_152826372.1 acyltransferase |
| 42(1) | WP_100537967.1 | WP_100537967.1 acyltransferase |
| 42(1) | WP_226383528.1 | WP_226383528.1 acyltransferase |
| 42(1) | WP_090993637.1 | WP_090993637.1 acyltransferase |
| 42(1) | WP_301249703.1 | WP_301249703.1 acyltransferase |
| 42(1) | WP_090908286.1 | WP_090908286.1 acyltransferase |
| 42(1) | WP_044785618.1 | WP_044785618.1 acyltransferase |
| 42(1) | WP_072732445.1 | WP_072732445.1 acyltransferase |
| 42(1) | WP_077567008.1 | WP_077567008.1 acyltransferase |
| 42(1) | WP_164783092.1 | WP_164783092.1 acyltransferase |
| 42(1) | WP_099853189.1 | WP_099853189.1 acyltransferase |
| 42(1) | WP_191798535.1 | WP_191798535.1 acyltransferase |
| 42(1) | WP_167375737.1 | WP_167375737.1 acyltransferase |
| 42(1) | WP_107734202.1 | WP_107734202.1 acyltransferase |
| 42(1) | WP_285079435.1 | WP_285079435.1 acyltransferase |
| 42(1) | WP_213644235.1 | WP_213644235.1 acyltransferase |
| 42(1) | WP_307537568.1 | WP_307537568.1 acyltransferase |
| 42(1) | WP_307214310.1 | WP_307214310.1 acyltransferase |
| 42(1) | WP_074094620.1 | WP_074094620.1 acyltransferase |
| 42(1) | WP_110822556.1 | WP_110822556.1 acyltransferase |
| 42(1) | WP_283246410.1 | WP_283246410.1 acyltransferase |
| 42(1) | WP_290403373.1 | WP_290403373.1 acyltransferase |
| 42(1) | WP_113052559.1 | WP_113052559.1 acyltransferase |
| 42(1) | WP_127592419.1 | WP_127592419.1 acyltransferase |
| 42(1) | WP_005210466.1 | WP_005210466.1 acyltransferase |
| 42(1) | WP_213616461.1 | WP_213616461.1 acyltransferase |
| 42(1) | WP_210095102.1 | WP_210095102.1 acyltransferase |
| 42(1) | WP_145146968.1 | WP_145146968.1 acyltransferase |
| 42(1) | WP_017428325.1 | WP_017428325.1 acyltransferase |
| 42(1) | WP_128101492.1 | WP_128101492.1 acyltransferase |
| 42(1) | WP_054958418.1 | WP_054958418.1 acyltransferase |
| 42(1) | WP_231505357.1 | WP_231505357.1 acyltransferase |
| 42(1) | WP_095287652.1 | WP_095287652.1 acyltransferase |
| 42(1) | WP_036670442.1 | WP_036670442.1 acyltransferase |
| 42(1) | WP_186375402.1 | WP_186375402.1 acyltransferase |
| 42(1) | WP_016821042.1 | WP_016821042.1 acyltransferase |
| 42(1) | WP_106765948.1 | WP_106765948.1 acyltransferase |
| 42(1) | WP_076318554.1 | WP_076318554.1 acyltransferase |
| 42(1) | WP_186331576.1 | WP_186331576.1 acyltransferase |
| 42(1) | WP_017689062.1 | WP_017689062.1 acyltransferase |
| 42(1) | WP_264929200.1 | WP_264929200.1 acyltransferase |
| 42(1) | WP_153978073.1 | WP_153978073.1 acyltransferase |
| 42(1) | WP_251497178.1 | WP_251497178.1 acyltransferase |
| 42(1) | WP_109999898.1 | WP_109999898.1 acyltransferase |
| 42(1) | WP_179265136.1 | WP_179265136.1 acyltransferase |
| 42(1) | WP_062833888.1 | WP_062833888.1 acyltransferase |
| 42(1) | WP_260065268.1 | WP_260065268.1 acyltransferase |
| 42(1) | WP_111619117.1 | WP_111619117.1 acyltransferase |
| 42(1) | WP_305755025.1 | WP_305755025.1 acyltransferase |
| 42(1) | WP_054402899.1 | WP_054402899.1 acyltransferase |
| 42(1) | WP_188593169.1 | WP_188593169.1 acyltransferase |
| 42(1) | WP_090949176.1 | WP_090949176.1 acyltransferase |
| 42(1) | WP_301879139.1 | WP_301879139.1 acyltransferase |
| 42(1) | WP_179198554.1 | WP_179198554.1 acyltransferase |
| 42(1) | WP_076208836.1 | WP_076208836.1 acyltransferase |
| 42(1) | WP_090887110.1 | WP_090887110.1 acyltransferase |
| 42(2) | WP_036609190.1 | WP_036609190.1 acyltransferase |
| 42(1) | WP_194543491.1 | WP_194543491.1 acyltransferase |
| 42(1) | WP_216531939.1 | WP_216531939.1 acyltransferase |
| 42(1) | WP_145046469.1 | WP_145046469.1 acyltransferase |
| 42(1) | WP_289392079.1 | WP_289392079.1 acyltransferase |
| 42(1) | WP_062326156.1 | WP_062326156.1 acyltransferase |
| 42(1) | WP_056700808.1 | WP_056700808.1 acyltransferase |
| 42(1) | WP_096774447.1 | WP_096774447.1 acyltransferase |
| 42(1) | WP_308721983.1 | WP_308721983.1 acyltransferase |
| 42(1) | WP_282321815.1 | WP_282321815.1 acyltransferase |
| 42(1) | WP_215153850.1 | WP_215153850.1 acyltransferase |
| 42(1) | WP_053783894.1 | WP_053783894.1 acyltransferase |
| 42(1) | WP_274566446.1 | WP_274566446.1 acyltransferase |
| 42(1) | WP_023957745.1 | WP_023957745.1 acyltransferase |
| 42(1) | WP_095358541.1 | WP_095358541.1 acyltransferase |
| 42(2) | WP_036642170.1 | WP_036642170.1 acyltransferase |
| 42(1) | WP_161557066.1 | WP_161557066.1 acyltransferase |
| 42(1) | WP_024630335.1 | WP_024630335.1 acyltransferase |
| 42(1) | WP_290409178.1 | WP_290409178.1 acyltransferase |
| 42(1) | WP_272035338.1 | WP_272035338.1 acyltransferase |
| 42(1) | WP_025364690.1 | WP_025364690.1 acyltransferase |
| 42(1) | WP_290371463.1 | WP_290371463.1 acyltransferase |
| 42(1) | WP_090896208.1 | WP_090896208.1 acyltransferase |
| 42(1) | WP_076322105.1 | WP_076322105.1 acyltransferase |
| 42(1) | WP_079694641.1 | WP_079694641.1 acyltransferase |

42(2) WP_015736233.1 WP_015736233.1 acyltransferase
42(1) WP_064639007.1 WP_064639007.1 acyltransferase
42(2) WP_063479989.1 WP_063479989.1 acyltransferase
42(1) WP_277746352.1 WP_277746352.1 acyltransferase
42(1) WP_167483536.1 WP_167483536.1 acyltransferase
42(1) WP_098746843.1 WP_098746843.1 acyltransferase

43(1) WP_098856774.1 WP_098856774.1 sodium-dependent transporter
43(2) WP_071708776.1 WP_071708776.1 sodium-dependent transporter
43(4) WP_040056500.1 WP_040056500.1 sodium-dependent transporter
43(5) WP_000814557.1 WP_000814557.1 sodium-dependent transporter
43(2) WP_088609117.1 WP_088609117.1 sodium-dependent transporter
43(1) WP_151630594.1 WP_151630594.1 sodium-dependent transporter
43(2) WP_002136059.1 WP_002136059.1 sodium-dependent transporter
43(1) WP_098365942.1 WP_098365942.1 sodium-dependent transporter
43(2) WP_088020923.1 WP_088020923.1 sodium-dependent transporter
43(1) WP_097987485.1 WP_097987485.1 sodium-dependent transporter
43(1) WP_019392111.1 WP_019392111.1 sodium-dependent transporter
43(1) WP_000814515.1 WP_000814515.1 sodium-dependent transporter
43(1) WP_000814522.1 WP_000814522.1 sodium-dependent transporter
43(1) WP_166705003.1 WP_166705003.1 sodium-dependent transporter
43(1) WP_078181475.1 WP_078181475.1 sodium-dependent transporter
43(1) WP_088063237.1 WP_088063237.1 sodium-dependent transporter
43(2) WP_098319884.1 WP_098319884.1 sodium-dependent transporter
43(1) WP_000814518.1 WP_000814518.1 sodium-dependent transporter
43(6) WP_000814553.1 WP_000814553.1 sodium-dependent transporter
43(1) WP_098239613.1 WP_098239613.1 sodium-dependent transporter
43(1) WP_088078466.1 WP_088078466.1 sodium-dependent transporter
43(3) WP_097925469.1 WP_097925469.1 sodium-dependent transporter
43(1) WP_097905171.1 WP_097905171.1 sodium-dependent transporter
43(6) WP_023521821.1 WP_023521821.1 sodium-dependent transporter
43(1) WP_098088235.1 WP_098088235.1 sodium-dependent transporter
43(1) WP_270784518.1 WP_270784518.1 sodium-dependent transporter
43(1) WP_087952384.1 WP_087952384.1 sodium-dependent transporter
43(1) WP_088119573.1 WP_088119573.1 sodium-dependent transporter
43(1) WP_143881080.1 WP_143881080.1 sodium-dependent transporter
43(3) WP_061805361.1 WP_061805361.1 sodium-dependent transporter
43(1) WP_071719976.1 WP_071719976.1 sodium-dependent transporter
43(2) WP_098082498.1 WP_098082498.1 sodium-dependent transporter
43(2) WP_048552317.1 WP_048552317.1 sodium-dependent transporter
43(1) WP_243518749.1 WP_243518749.1 sodium-dependent transporter
43(1) WP_001980149.1 WP_001980149.1 sodium-dependent transporter
43(1) WP_000814554.1 WP_000814554.1 sodium-dependent transporter
43(1) WP_270711674.1 WP_270711674.1 sodium-dependent transporter
43(1) WP_220775548.1 WP_220775548.1 sodium-dependent transporter
43(4) WP_000814517.1 WP_000814517.1 sodium-dependent transporter
43(1) WP_098680262.1 WP_098680262.1 sodium-dependent transporter
43(1) WP_098148068.1 WP_098148068.1 sodium-dependent transporter
43(1) WP_268583862.1 WP_268583862.1 sodium-dependent transporter
43(3) WP_087982573.1 WP_087982573.1 sodium-dependent transporter
43(1) WP_072938291.1 WP_072938291.1 sodium-dependent transporter
43(1) WP_098250532.1 WP_098250532.1 sodium-dependent transporter
43(1) WP_071679493.1 WP_071679493.1 sodium-dependent transporter
43(2) WP_215570378.1 WP_215570378.1 sodium-dependent transporter
43(1) WP_270699385.1 WP_270699385.1 sodium-dependent transporter
43(1) WP_215601070.1 WP_215601070.1 sodium-dependent transporter
43(1) WP_262748105.1 WP_262748105.1 sodium-dependent transporter
43(2) WP_098976049.1 WP_098976049.1 sodium-dependent transporter
43(1) WP_262725723.1 WP_262725723.1 sodium-dependent transporter
43(2) WP_087956298.1 WP_087956298.1 sodium-dependent transporter
43(2) WP_098266655.1 WP_098266655.1 sodium-dependent transporter
43(1) WP_097810729.1 WP_097810729.1 sodium-dependent transporter
43(1) WP_070805905.1 WP_070805905.1 sodium-dependent transporter
43(1) WP_064477380.1 WP_064477380.1 sodium-dependent transporter
43(1) WP_262719884.1 WP_262719884.1 sodium-dependent transporter
43(2) WP_139895504.1 WP_139895504.1 sodium-dependent transporter
43(1) WP_071724734.1 WP_071724734.1 sodium-dependent transporter
43(1) WP_147807648.1 WP_147807648.1 sodium-dependent transporter
43(1) WP_242251954.1 WP_242251954.1 sodium-dependent transporter
43(1) WP_139867360.1 WP_139867360.1 sodium-dependent transporter
43(1) WP_069355514.1 WP_069355514.1 sodium-dependent transporter
43(1) WP_199695256.1 WP_199695256.1 sodium-dependent transporter
43(1) WP_144611197.1 WP_144611197.1 sodium-dependent transporter
43(1) WP_251456532.1 WP_251456532.1 sodium-dependent transporter
43(1) WP_060630652.1 WP_060630652.1 sodium-dependent transporter
43(2) WP_098264291.1 WP_098264291.1 sodium-dependent transporter
43(1) WP_088058543.1 WP_088058543.1 sodium-dependent transporter
43(1) WP_097852386.1 WP_097852386.1 sodium-dependent transporter
43(1) WP_106382857.1 WP_106382857.1 sodium-dependent transporter
43(2) WP_002031671.1 WP_002031671.1 sodium-dependent transporter
43(1) WP_197234727.1 WP_197234727.1 sodium-dependent transporter
43(1) WP_215598785.1 WP_215598785.1 sodium-dependent transporter
43(4) WP_103626236.1 WP_103626236.1 sodium-dependent transporter
43(1) WP_053564618.1 WP_053564618.1 sodium-dependent transporter
43(1) WP_063223288.1 WP_063223288.1 sodium-dependent transporter
43(1) WP_098776423.1 WP_098776423.1 sodium-dependent transporter
43(1) WP_124049174.1 WP_124049174.1 sodium-dependent transporter

44(1) WP_029438085.1 WP_029438085.1 TetR/AcrR family transcriptional regulator
44(1) WP_242255485.1 WP_242255485.1 TetR/AcrR family transcriptional regulator
44(1) WP_221875358.1 WP_221875358.1 TetR-like C-terminal domain-containing protein
44(1) WP_286125241.1 WP_286125241.1 TetR/AcrR family transcriptional regulator
44(1) WP_229521575.1 WP_229521575.1 TetR/AcrR family transcriptional regulator C-terminal domain-containing protein
44(1) WP_144744066.1 WP_144744066.1 TetR/AcrR family transcriptional regulator C-terminal domain-containing protein
44(2) WP_098167598.1 WP_098167598.1 TetR/AcrR family transcriptional regulator
44(1) WP_098776435.1 WP_098776435.1 TetR/AcrR family transcriptional regulator
44(1) WP_242251763.1 WP_242251763.1 TetR/AcrR family transcriptional regulator
44(7) WP_023521810.1 WP_023521810.1 TetR/AcrR family transcriptional regulator
44(2) WP_090976421.1 WP_090976421.1 TetR/AcrR family transcriptional regulator
44(1) WP_103047822.1 WP_103047822.1 TetR/AcrR family transcriptional regulator
44(1) WP_134902881.1 WP_134902881.1 TetR/AcrR family transcriptional regulator
44(1) WP_242400018.1 WP_242400018.1 TetR/AcrR family transcriptional regulator

44(1) WP_095691014.1 WP_095691014.1 TetR/AcrR family transcriptional regulator
44(1) WP_071727419.1 WP_071727419.1 TetR/AcrR family transcriptional regulator
44(1) WP_193653026.1 WP_193653026.1 TetR/AcrR family transcriptional regulator, partial
44(1) WP_215572748.1 WP_215572748.1 TetR/AcrR family transcriptional regulator
44(1) WP_036944572.1 WP_036944572.1 TetR/AcrR family transcriptional regulator
44(2) WP_002158052.1 WP_002158052.1 TetR/AcrR family transcriptional regulator
44(1) WP_098451707.1 WP_098451707.1 TetR/AcrR family transcriptional regulator
44(1) WP_232274503.1 WP_232274503.1 TetR/AcrR family transcriptional regulator
44(1) WP_190299140.1 WP_190299140.1 TetR/AcrR family transcriptional regulator
44(4) WP_000045218.1 WP_000045218.1 TetR/AcrR family transcriptional regulator
44(5) WP_000045217.1 WP_000045217.1 TetR/AcrR family transcriptional regulator
44(2) WP_098678282.1 WP_098678282.1 TetR/AcrR family transcriptional regulator
44(1) WP_000045225.1 WP_000045225.1 TetR/AcrR family transcriptional regulator
44(2) WP_273191240.1 WP_273191240.1 TetR/AcrR family transcriptional regulator C-terminal domain-containing protein
44(1) WP_078205521.1 WP_078205521.1 TetR/AcrR family transcriptional regulator
44(1) WP_088058525.1 WP_088058525.1 TetR/AcrR family transcriptional regulator
44(2) WP_010963361.1 WP_010963361.1 TetR/AcrR family transcriptional regulator
44(2) WP_098447524.1 WP_098447524.1 TetR/AcrR family transcriptional regulator
44(1) WP_144476572.1 WP_144476572.1 TetR/AcrR family transcriptional regulator
44(1) WP_098250520.1 WP_098250520.1 TetR/AcrR family transcriptional regulator
44(1) WP_088032770.1 WP_088032770.1 TetR/AcrR family transcriptional regulator
44(1) WP_016104718.1 WP_016104718.1 TetR/AcrR family transcriptional regulator
44(1) WP_212956544.1 WP_212956544.1 TetR/AcrR family transcriptional regulator
44(1) WP_097905186.1 WP_097905186.1 TetR/AcrR family transcriptional regulator
44(1) WP_165175571.1 WP_165175571.1 TetR/AcrR family transcriptional regulator
44(1) WP_307589627.1 WP_307589627.1 TetR/AcrR family transcriptional regulator
44(1) WP_098840609.1 WP_098840609.1 TetR/AcrR family transcriptional regulator
44(1) WP_144506823.1 WP_144506823.1 TetR/AcrR family transcriptional regulator
44(1) WP_063219643.1 WP_063219643.1 TetR/AcrR family transcriptional regulator
44(3) WP_097961539.1 WP_097961539.1 TetR/AcrR family transcriptional regulator
44(1) WP_066022441.1 WP_066022441.1 TetR/AcrR family transcriptional regulator
44(1) WP_154118039.1 WP_154118039.1 TetR/AcrR family transcriptional regulator
44(1) WP_149871688.1 WP_149871688.1 TetR/AcrR family transcriptional regulator
44(1) WP_286124778.1 WP_286124778.1 TetR/AcrR family transcriptional regulator
44(1) WP_244723823.1 WP_244723823.1 TetR/AcrR family transcriptional regulator
44(15) WP_016512885.1 WP_016512885.1 TetR/AcrR family transcriptional regulator
44(1) WP_199677617.1 WP_199677617.1 TetR/AcrR family transcriptional regulator
44(2) WP_071724714.1 WP_071724714.1 TetR/AcrR family transcriptional regulator
44(1) WP_043067156.1 WP_043067156.1 TetR/AcrR family transcriptional regulator
44(1) WP_019912172.1 WP_019912172.1 TetR/AcrR family transcriptional regulator
44(2) WP_251426379.1 WP_251426379.1 TetR/AcrR family transcriptional regulator
44(1) WP_102372760.1 WP_102372760.1 TetR/AcrR family transcriptional regulator
44(1) WP_071719988.1 WP_071719988.1 TetR/AcrR family transcriptional regulator
44(1) WP_248251176.1 WP_248251176.1 TetR/AcrR family transcriptional regulator
44(1) WP_063223270.1 WP_063223270.1 TetR/AcrR family transcriptional regulator
44(1) WP_098519062.1 WP_098519062.1 TetR/AcrR family transcriptional regulator
44(1) WP_234121614.1 WP_234121614.1 TetR/AcrR family transcriptional regulator
44(1) WP_192266427.1 WP_192266427.1 TetR/AcrR family transcriptional regulator
44(1) WP_286017542.1 WP_286017542.1 TetR/AcrR family transcriptional regulator
44(1) WP_255357620.1 WP_255357620.1 hypothetical protein
44(1) WP_097987495.1 WP_097987495.1 TetR/AcrR family transcriptional regulator
44(1) WP_098834580.1 WP_098834580.1 TetR/AcrR family transcriptional regulator
44(1) WP_236334300.1 WP_236334300.1 TetR/AcrR family transcriptional regulator
44(5) WP_002136053.1 WP_002136053.1 TetR/AcrR family transcriptional regulator
44(3) WP_000044993.1 WP_000044993.1 TetR/AcrR family transcriptional regulator
44(1) WP_247083029.1 WP_247083029.1 TetR/AcrR family transcriptional regulator
44(1) WP_012260828.1 WP_012260828.1 TetR/AcrR family transcriptional regulator

45(1) WP_110822554.1 WP_110822554.1 MFS transporter
45(1) WP_099517120.1 WP_099517120.1 MDR family MFS transporter
45(1) WP_045243836.1 WP_045243836.1 MFS transporter
45(1) WP_213618808.1 WP_213618808.1 MFS transporter
45(1) WP_247083030.1 WP_247083030.1 MFS transporter
45(1) WP_145046473.1 WP_145046473.1 MFS transporter
45(1) WP_110892950.1 WP_110892950.1 MFS transporter
45(1) WP_126665664.1 WP_126665664.1 MFS transporter
45(1) WP_164965494.1 WP_164965494.1 MFS transporter
45(1) WP_290403370.1 WP_290403370.1 MFS transporter
45(1) WP_216531942.1 WP_216531942.1 MFS transporter
45(2) WP_006211190.1 WP_006211190.1 MFS transporter
45(1) WP_188593171.1 WP_188593171.1 MFS transporter
45(1) WP_071220216.1 WP_071220216.1 MFS transporter
45(2) WP_111925415.1 WP_111925415.1 MFS transporter
45(1) WP_237167831.1 WP_237167831.1 MFS transporter
45(1) WP_026314618.1 WP_026314618.1 MFS transporter
45(2) WP_124049179.1 WP_124049179.1 MFS transporter
45(2) WP_207714829.1 WP_207714829.1 hypothetical protein
45(1) WP_155618134.1 WP_155618134.1 MFS transporter
45(1) WP_093291070.1 WP_093291070.1 MFS transporter
45(1) WP_068694703.1 WP_068694703.1 MFS transporter
45(1) WP_223067447.1 WP_223067447.1 MFS transporter
45(1) WP_077832272.1 WP_077832272.1 MFS transporter
45(3) WP_077833098.1 WP_077833098.1 MFS transporter
45(1) WP_119847561.1 WP_119847561.1 MFS transporter
45(1) WP_257675213.1 WP_257675213.1 MFS transporter
45(1) WP_280753896.1 WP_280753896.1 MFS transporter
45(1) WP_123797116.1 WP_123797116.1 MFS transporter
45(1) WP_251375554.1 WP_251375554.1 MFS transporter
45(1) WP_192266480.1 WP_192266480.1 MFS transporter
45(2) WP_036642172.1 WP_036642172.1 MFS transporter
45(1) WP_074598762.1 WP_074598762.1 MFS transporter
45(1) WP_286124772.1 WP_286124772.1 MFS transporter
45(1) WP_113753077.1 WP_113753077.1 MFS transporter
45(1) WP_308721982.1 WP_308721982.1 MFS transporter
45(1) WP_263560236.1 WP_263560236.1 MFS transporter
45(1) WP_209365682.1 WP_209365682.1 MFS transporter
45(1) WP_154118038.1 WP_154118038.1 MFS transporter
45(1) WP_289373779.1 WP_289373779.1 MFS transporter
45(1) WP_181737047.1 WP_181737047.1 MFS transporter
45(1) WP_236417239.1 WP_236417239.1 MFS transporter
45(1) WP_127536187.1 WP_127536187.1 MFS transporter
45(2) WP_207714781.1 WP_207714781.1 MFS transporter
45(1) WP_077853497.1 WP_077853497.1 MFS transporter

| | | |
|---|---|---|
| 45(1) | WP_254759286.1 | WP_254759286.1 MFS transporter |
| 45(1) | WP_164848678.1 | WP_164848678.1 MFS transporter |
| 45(1) | WP_243518741.1 | WP_243518741.1 MFS transporter |
| 45(1) | WP_213644236.1 | WP_213644236.1 MFS transporter |
| 45(1) | WP_194543489.1 | WP_194543489.1 MFS transporter |
| 45(1) | WP_028401636.1 | WP_028401636.1 MFS transporter |
| 45(1) | WP_095287649.1 | WP_095287649.1 MFS transporter |
| 45(1) | WP_077893475.1 | WP_077893475.1 MFS transporter |
| 45(1) | WP_289342245.1 | WP_289342245.1 multidrug effflux MFS transporter |
| 45(1) | WP_127592417.1 | WP_127592417.1 MFS transporter |
| 45(1) | WP_076322103.1 | WP_076322103.1 MFS transporter |
| 45(1) | WP_237741331.1 | WP_237741331.1 hypothetical protein |
| 45(2) | WP_219872451.1 | WP_219872451.1 MFS transporter |
| 45(1) | WP_063479988.1 | WP_063479988.1 MFS transporter |
| 45(1) | WP_108465343.1 | WP_108465343.1 MFS transporter |
| 45(2) | WP_090887124.1 | WP_090887124.1 MFS transporter |
| 45(1) | WP_113713140.1 | WP_113713140.1 MFS transporter |
| 45(1) | WP_307214312.1 | WP_307214312.1 MFS transporter |
| 45(1) | WP_268583868.1 | WP_268583868.1 MFS transporter |
| 45(1) | WP_094092611.1 | WP_094092611.1 MFS transporter |
| 45(1) | WP_128101495.1 | WP_128101495.1 MFS transporter |
| 45(1) | WP_272885663.1 | WP_272885663.1 DHA2 family efflux MFS transporter permease subunit |
| 45(1) | WP_198056469.1 | WP_198056469.1 MFS transporter |
| 45(1) | WP_272541593.1 | WP_272541593.1 MFS transporter |
| 45(1) | WP_293202254.1 | WP_293202254.1 MFS transporter |
| 45(1) | WP_208606793.1 | WP_208606793.1 MFS transporter |
| 45(1) | WP_215153852.1 | WP_215153852.1 MFS transporter |
| 45(1) | WP_205127958.1 | WP_205127958.1 MFS transporter |
| 45(1) | WP_113058727.1 | WP_113058727.1 MFS transporter |
| 45(1) | WP_056700813.1 | WP_056700813.1 MFS transporter |
| 45(1) | WP_007128615.1 | WP_007128615.1 MFS transporter |
| 45(1) | WP_072336472.1 | WP_072336472.1 MFS transporter |
| 45(1) | WP_260065272.1 | WP_260065272.1 MFS transporter |
| 45(1) | WP_095398797.1 | WP_095398797.1 MFS transporter |
| 45(1) | WP_068622363.1 | WP_068622363.1 MFS transporter |
| 45(1) | WP_236334302.1 | WP_236334302.1 MFS transporter |
| 45(1) | WP_220684648.1 | WP_220684648.1 MFS transporter |
| 45(1) | WP_272035344.1 | WP_272035344.1 MFS transporter |
| 45(2) | WP_186331577.1 | WP_186331577.1 MFS transporter |
| 45(2) | WP_033100007.1 | WP_033100007.1 MFS transporter |
| 45(1) | WP_251537238.1 | WP_251537238.1 MFS transporter |
| 45(1) | WP_095358539.1 | WP_095358539.1 MFS transporter |
| 45(1) | WP_308083971.1 | WP_308083971.1 MFS transporter |
| 45(1) | WP_016821036.1 | WP_016821036.1 MFS transporter |
| 45(1) | WP_258209122.1 | WP_258209122.1 MFS transporter |
| 45(1) | WP_100537965.1 | WP_100537965.1 MFS transporter |
| 45(1) | WP_145408772.1 | WP_145408772.1 MFS transporter |
| 45(1) | WP_127199763.1 | WP_127199763.1 MFS transporter |
| 45(1) | WP_247083034.1 | WP_247083034.1 MFS transporter |
| 45(1) | WP_077567005.1 | WP_077567005.1 MFS transporter |
| 45(1) | WP_244908217.1 | WP_244908217.1 MFS transporter |
| 45(1) | WP_061805356.1 | WP_061805356.1 MFS transporter |

| | | |
|---|---|---|
| 45(1) | WP_098746838.1 | WP_098746838.1 MFS transporter |
| 45(1) | WP_231638513.1 | WP_231638513.1 MFS transporter |
| 45(4) | WP_019392104.1 | WP_019392104.1 MFS transporter |
| 46(1) | WP_282742028.1 | WP_282742028.1 hypothetical protein |
| 46(1) | WP_214706010.1 | WP_214706010.1 hypothetical protein |
| 46(1) | WP_216830385.1 | WP_216830385.1 hypothetical protein |
| 46(1) | WP_251542622.1 | WP_251542622.1 hypothetical protein |
| 46(1) | WP_307388961.1 | WP_307388961.1 hypothetical protein |
| 46(1) | WP_251557373.1 | WP_251557373.1 hypothetical protein |
| 46(1) | WP_213143954.1 | WP_213143954.1 hypothetical protein |
| 46(1) | WP_251426358.1 | WP_251426358.1 hypothetical protein |
| 46(1) | WP_191272850.1 | WP_191272850.1 hypothetical protein |
| 46(1) | WP_077619640.1 | WP_077619640.1 hypothetical protein |
| 46(1) | WP_308912805.1 | WP_308912805.1 hypothetical protein |
| 46(1) | WP_179160613.1 | WP_179160613.1 hypothetical protein |
| 46(1) | WP_263324556.1 | WP_263324556.1 hypothetical protein |
| 46(1) | WP_015595752.1 | WP_015595752.1 hypothetical protein |
| 46(1) | WP_034675381.1 | WP_034675381.1 hypothetical protein |
| 46(1) | WP_278773432.1 | WP_278773432.1 hypothetical protein |
| 46(1) | WP_308082253.1 | WP_308082253.1 hypothetical protein |
| 46(1) | WP_204412711.1 | WP_204412711.1 hypothetical protein |
| 46(1) | WP_102264543.1 | WP_102264543.1 hypothetical protein |
| 46(1) | WP_090761835.1 | WP_090761835.1 hypothetical protein |
| 46(3) | WP_291289940.1 | WP_291289940.1 hypothetical protein |
| 46(1) | WP_070638647.1 | WP_070638647.1 hypothetical protein |
| 46(1) | WP_079506410.1 | WP_079506410.1 hypothetical protein |
| 46(1) | WP_277175593.1 | WP_277175593.1 hypothetical protein |
| 46(1) | WP_251548766.1 | WP_251548766.1 hypothetical protein |
| 46(1) | WP_273191246.1 | WP_273191246.1 hypothetical protein |
| 46(1) | WP_294386424.1 | WP_294386424.1 hypothetical protein, partial |
| 46(1) | WP_226086169.1 | WP_226086169.1 hypothetical protein |
| 46(1) | WP_058271104.1 | WP_058271104.1 hypothetical protein |
| 46(1) | WP_133332985.1 | WP_133332985.1 hypothetical protein |
| 46(1) | WP_303942526.1 | WP_303942526.1 hypothetical protein |
| 46(1) | WP_046522791.1 | WP_046522791.1 hypothetical protein |
| 46(1) | WP_144479051.1 | WP_144479051.1 hypothetical protein |
| 46(1) | WP_273383868.1 | WP_273383868.1 hypothetical protein |
| 46(1) | WP_013252386.1 | WP_013252386.1 hypothetical protein |
| 46(1) | WP_277166586.1 | WP_277166586.1 hypothetical protein |
| 46(1) | WP_024027637.1 | WP_024027637.1 hypothetical protein |
| 46(1) | WP_116353244.1 | WP_116353244.1 hypothetical protein |
| 46(1) | WP_007148002.1 | WP_007148002.1 hypothetical protein |
| 46(1) | WP_277142098.1 | WP_277142098.1 hypothetical protein |
| 46(1) | WP_136379706.1 | WP_136379706.1 hypothetical protein |
| 46(1) | WP_306077383.1 | WP_306077383.1 hypothetical protein |
| 46(1) | WP_304216880.1 | WP_304216880.1 hypothetical protein |
| 46(1) | WP_121610602.1 | WP_121610602.1 hypothetical protein |
| 46(1) | WP_307738720.1 | WP_307738720.1 hypothetical protein |
| 46(1) | WP_066091983.1 | WP_066091983.1 hypothetical protein |
| 46(1) | WP_198310826.1 | WP_198310826.1 hypothetical protein |
| 46(1) | WP_102372755.1 | WP_102372755.1 hypothetical protein |

| | | |
|---|---|---|
| 46(1) | WP_149871675.1 | WP_149871675.1 hypothetical protein |
| 46(1) | WP_291288260.1 | WP_291288260.1 hypothetical protein |
| 46(1) | WP_277157951.1 | WP_277157951.1 hypothetical protein |
| 46(1) | WP_273061466.1 | WP_273061466.1 hypothetical protein |
| 46(1) | WP_283904340.1 | WP_283904340.1 hypothetical protein |
| 46(1) | WP_268058019.1 | WP_268058019.1 hypothetical protein |
| 46(1) | WP_034526605.1 | WP_034526605.1 hypothetical protein |
| 46(1) | WP_210364460.1 | WP_210364460.1 hypothetical protein |
| 46(1) | WP_254759305.1 | WP_254759305.1 hypothetical protein |
| 46(1) | WP_036575536.1 | WP_036575536.1 hypothetical protein |
| 46(1) | WP_251481004.1 | WP_251481004.1 hypothetical protein |
| 46(1) | WP_143316574.1 | WP_143316574.1 hypothetical protein |
| 46(1) | WP_277156193.1 | WP_277156193.1 hypothetical protein |
| 46(1) | WP_166245135.1 | WP_166245135.1 hypothetical protein |
| 46(1) | WP_297183851.1 | WP_297183851.1 hypothetical protein |
| 46(1) | WP_309098617.1 | WP_309098617.1 hypothetical protein |
| 46(1) | WP_005604617.1 | WP_005604617.1 hypothetical protein |
| 46(1) | WP_215032109.1 | WP_215032109.1 hypothetical protein |
| 46(1) | WP_007083297.1 | WP_007083297.1 hypothetical protein |
| 46(1) | WP_308110112.1 | WP_308110112.1 hypothetical protein |
| 46(1) | WP_102371669.1 | WP_102371669.1 hypothetical protein |
| 46(1) | WP_006292335.1 | WP_006292335.1 hypothetical protein |
| 46(1) | WP_270472015.1 | WP_270472015.1 hypothetical protein |
| 46(1) | WP_226679097.1 | WP_226679097.1 hypothetical protein |
| 46(1) | WP_226602900.1 | WP_226602900.1 hypothetical protein |
| 46(1) | WP_099361342.1 | WP_099361342.1 hypothetical protein |
| 46(1) | WP_179603473.1 | WP_179603473.1 hypothetical protein |
| 46(1) | WP_251638757.1 | WP_251638757.1 hypothetical protein |
| 46(1) | WP_294953154.1 | WP_294953154.1 hypothetical protein |
| 46(1) | WP_130810263.1 | WP_130810263.1 hypothetical protein |
| 46(1) | WP_227219200.1 | WP_227219200.1 hypothetical protein |
| 46(1) | WP_053361016.1 | WP_053361016.1 hypothetical protein |
| 46(1) | WP_132008175.1 | WP_132008175.1 hypothetical protein |
| 46(1) | WP_041964568.1 | WP_041964568.1 hypothetical protein |
| 46(1) | WP_042462815.1 | WP_042462815.1 hypothetical protein |
| 46(1) | WP_231094841.1 | WP_231094841.1 hypothetical protein |
| 46(1) | WP_044037255.1 | WP_044037255.1 hypothetical protein |
| 46(1) | WP_217035869.1 | WP_217035869.1 hypothetical protein |
| 46(1) | WP_223592672.1 | WP_223592672.1 hypothetical protein |
| 46(1) | WP_224839868.1 | WP_224839868.1 hypothetical protein |
| 46(1) | WP_212330906.1 | WP_212330906.1 hypothetical protein |
| 46(1) | WP_307192891.1 | WP_307192891.1 hypothetical protein |
| 46(1) | WP_307312569.1 | WP_307312569.1 hypothetical protein |
| 46(1) | WP_026572745.1 | WP_026572745.1 hypothetical protein |
| 46(1) | WP_302153082.1 | WP_302153082.1 hypothetical protein |
| 46(1) | WP_226642884.1 | WP_226642884.1 hypothetical protein |
| 46(1) | WP_033521274.1 | WP_033521274.1 hypothetical protein |
| 46(1) | WP_229139927.1 | WP_229139927.1 hypothetical protein |
| 46(1) | WP_283952941.1 | WP_283952941.1 hypothetical protein |
| 46(1) | WP_289342269.1 | WP_289342269.1 hypothetical protein |
| 46(1) | WP_251466811.1 | WP_251466811.1 hypothetical protein |
| 46(1) | WP_010676856.1 | WP_010676856.1 hypothetical protein |
| 46(1) | WP_066415460.1 | WP_066415460.1 hypothetical protein |
| 46(1) | WP_284036493.1 | WP_284036493.1 hypothetical protein |
| 46(1) | WP_006720306.1 | WP_006720306.1 hypothetical protein |
| 46(1) | WP_099354916.1 | WP_099354916.1 hypothetical protein |
| 46(1) | WP_213117489.1 | WP_213117489.1 hypothetical protein |
| 46(1) | WP_270065050.1 | WP_270065050.1 hypothetical protein |
| 46(1) | WP_307284694.1 | WP_307284694.1 hypothetical protein |
| 46(1) | WP_122971544.1 | WP_122971544.1 hypothetical protein |
| 46(1) | WP_013490139.1 | WP_013490139.1 hypothetical protein |
| 46(1) | WP_289607501.1 | WP_289607501.1 hypothetical protein |
| 46(1) | WP_045518276.1 | WP_045518276.1 hypothetical protein |

| | | |
|---|---|---|
| 47(1) | WP_226602904.1 | WP_226602904.1 protein phosphatase 2C domain-containing protein |
| 47(1) | WP_227219197.1 | WP_227219197.1 PP2C family serine/threonine-protein phosphatase |
| 47(1) | WP_226642882.1 | WP_226642882.1 protein phosphatase 2C domain-containing protein |
| 47(1) | WP_077893581.1 | WP_077893581.1 serine/threonine-protein phosphatase |
| 47(1) | WP_254759303.1 | WP_254759303.1 protein phosphatase 2C domain-containing protein |
| 47(1) | WP_307399127.1 | WP_307399127.1 Stp1/IreP family PP2C-type Ser/Thr phosphatase |
| 47(1) | WP_283952943.1 | WP_283952943.1 protein phosphatase 2C domain-containing protein |
| 47(1) | WP_265444531.1 | WP_265444531.1 SpoIIE family protein phosphatase |
| 47(1) | WP_121610600.1 | WP_121610600.1 protein phosphatase 2C domain-containing protein |
| 47(1) | WP_308082251.1 | WP_308082251.1 protein phosphatase 2C domain-containing protein |
| 47(1) | WP_166245131.1 | WP_166245131.1 serine/threonine-protein phosphatase |
| 47(1) | WP_041964570.1 | WP_041964570.1 protein phosphatase 2C domain-containing protein |
| 47(1) | WP_077852418.1 | WP_077852418.1 serine/threonine-protein phosphatase |
| 47(1) | WP_304216876.1 | WP_304216876.1 protein phosphatase 2C domain-containing protein |
| 47(1) | WP_224839866.1 | WP_224839866.1 protein phosphatase 2C domain-containing protein |
| 47(1) | WP_046522789.1 | WP_046522789.1 protein phosphatase 2C domain-containing protein |
| 47(1) | WP_090761831.1 | WP_090761831.1 protein phosphatase 2C domain-containing protein |
| 47(1) | WP_102264545.1 | WP_102264545.1 protein phosphatase 2C domain-containing protein |
| 47(1) | WP_005604619.1 | WP_005604619.1 protein phosphatase 2C domain-containing protein |
| 47(1) | WP_069193874.1 | WP_069193874.1 SpoIIE family protein phosphatase |
| 47(1) | WP_195325253.1 | WP_195325253.1 protein phosphatase 2C domain-containing protein |
| 47(1) | WP_251557375.1 | WP_251557375.1 serine/threonine-protein phosphatase |
| 47(1) | WP_251638760.1 | WP_251638760.1 protein phosphatase 2C domain-containing protein |
| 47(1) | WP_307192893.1 | WP_307192893.1 protein phosphatase 2C domain-containing protein |
| 47(1) | WP_306077385.1 | WP_306077385.1 protein phosphatase 2C domain-containing protein |
| 47(1) | WP_270065052.1 | WP_270065052.1 protein phosphatase 2C domain-containing protein |
| 47(1) | WP_010676858.1 | WP_010676858.1 protein phosphatase 2C domain-containing protein |
| 47(1) | WP_024027635.1 | WP_024027635.1 serine/threonine-protein phosphatase |
| 47(1) | WP_231094843.1 | WP_231094843.1 serine/threonine-protein phosphatase |
| 47(1) | WP_044037267.1 | WP_044037267.1 protein phosphatase 2C domain-containing protein |
| 47(1) | WP_198310828.1 | WP_198310828.1 protein phosphatase 2C domain-containing protein |
| 47(1) | WP_026882398.1 | WP_026882398.1 serine/threonine-protein phosphatase |
| 47(1) | WP_144479055.1 | WP_144479055.1 protein phosphatase 2C domain-containing protein |
| 47(1) | WP_149871677.1 | WP_149871677.1 PP2C family serine/threonine-protein phosphatase |
| 47(1) | WP_045518278.1 | WP_045518278.1 serine/threonine-protein phosphatase |
| 47(1) | WP_099354914.1 | WP_099354914.1 protein phosphatase 2C domain-containing protein |
| 47(1) | WP_079506406.1 | WP_079506406.1 protein phosphatase 2C domain-containing protein |
| 47(1) | WP_294386428.1 | WP_294386428.1 protein phosphatase 2C domain-containing protein |
| 47(1) | WP_257675218.1 | WP_257675218.1 serine/threonine-protein phosphatase |
| 47(1) | WP_208926623.1 | WP_208926623.1 protein phosphatase 2C domain-containing protein |

| | | |
|---|---|---|
| 47(1) | WP_213117487.1 | WP_213117487.1 protein phosphatase 2C domain-containing protein |
| 47(1) | WP_226679099.1 | WP_226679099.1 protein phosphatase 2C domain-containing protein |
| 47(1) | WP_191272854.1 | WP_191272854.1 protein phosphatase 2C domain-containing protein |
| 47(1) | WP_195542473.1 | WP_195542473.1 serine/threonine-protein phosphatase |
| 47(1) | WP_007083295.1 | WP_007083295.1 serine/threonine-protein phosphatase |
| 47(1) | WP_251426360.1 | WP_251426360.1 serine/threonine-protein phosphatase |
| 47(1) | WP_204412715.1 | WP_204412715.1 protein phosphatase 2C domain-containing protein |
| 47(1) | WP_251542618.1 | WP_251542618.1 protein phosphatase 2C domain-containing protein |
| 47(2) | WP_133332983.1 | WP_133332983.1 protein phosphatase 2C domain-containing protein |
| 47(1) | WP_251548769.1 | WP_251548769.1 protein phosphatase 2C domain-containing protein |
| 47(3) | WP_077832280.1 | WP_077832280.1 serine/threonine-protein phosphatase |
| 47(1) | WP_179603475.1 | WP_179603475.1 protein phosphatase 2C domain-containing protein |
| 47(1) | WP_210364458.1 | WP_210364458.1 protein phosphatase 2C domain-containing protein |
| 47(1) | WP_234987242.1 | WP_234987242.1 protein phosphatase 2C domain-containing protein |
| 47(1) | WP_283904342.1 | WP_283904342.1 protein phosphatase 2C domain-containing protein |
| 47(1) | WP_015595754.1 | WP_015595754.1 serine/threonine-protein phosphatase |
| 47(1) | WP_307738714.1 | WP_307738714.1 Stp1/IreP family PP2C-type Ser/Thr phosphatase |
| 47(1) | WP_251466807.1 | WP_251466807.1 serine/threonine-protein phosphatase |
| 47(1) | WP_226086171.1 | WP_226086171.1 protein phosphatase 2C domain-containing protein |
| 47(2) | WP_010963729.1 | WP_010963729.1 serine/threonine-protein phosphatase |
| 47(1) | WP_066415453.1 | WP_066415453.1 protein phosphatase 2C domain-containing protein |
| 47(2) | WP_217035865.1 | WP_217035865.1 protein phosphatase 2C domain-containing protein |
| 47(1) | WP_289342268.1 | WP_289342268.1 protein phosphatase 2C domain-containing protein |
| 47(1) | WP_307312563.1 | WP_307312563.1 protein phosphatase 2C domain-containing protein |
| 47(1) | WP_180965811.1 | WP_180965811.1 protein phosphatase 2C domain-containing protein |
| 47(1) | WP_179160615.1 | WP_179160615.1 protein phosphatase 2C domain-containing protein |
| 47(1) | WP_294953215.1 | WP_294953215.1 serine/threonine-protein phosphatase |
| 47(2) | WP_195343921.1 | WP_195343921.1 serine/threonine-protein phosphatase |
| 47(1) | WP_307284691.1 | WP_307284691.1 protein phosphatase 2C domain-containing protein |
| 47(1) | WP_289607499.1 | WP_289607499.1 protein phosphatase 2C domain-containing protein |
| 47(1) | WP_122971542.1 | WP_122971542.1 protein phosphatase 2C domain-containing protein |
| 47(1) | WP_066020845.1 | WP_066020845.1 serine/threonine-protein phosphatase |
| 47(1) | WP_245546895.1 | WP_245546895.1 serine/threonine-protein phosphatase |
| 47(2) | WP_004463275.1 | WP_004463275.1 protein phosphatase 2C domain-containing protein |
| 47(2) | WP_010963364.1 | WP_010963364.1 serine/threonine-protein phosphatase |
| 47(1) | WP_223592668.1 | WP_223592668.1 protein phosphatase 2C domain-containing protein |
| 47(1) | WP_309098615.1 | WP_309098615.1 protein phosphatase 2C domain-containing protein |
| 47(1) | WP_073008263.1 | WP_073008263.1 serine/threonine-protein phosphatase |
| 47(2) | WP_295628328.1 | WP_295628328.1 protein phosphatase 2C domain-containing protein |
| 47(1) | WP_284526996.1 | WP_284526996.1 serine/threonine-protein phosphatase |
| 47(1) | WP_308110114.1 | WP_308110114.1 serine/threonine-protein phosphatase |
| 47(1) | WP_214706008.1 | WP_214706008.1 protein phosphatase 2C domain-containing protein |
| 47(1) | WP_026572743.1 | WP_026572743.1 protein phosphatase 2C domain-containing protein |
| 47(1) | WP_077833111.1 | WP_077833111.1 serine/threonine-protein phosphatase |
| 47(1) | WP_132008169.1 | WP_132008169.1 protein phosphatase 2C domain-containing protein |
| 47(1) | WP_234121609.1 | WP_234121609.1 serine/threonine-protein phosphatase |
| 47(1) | WP_308083992.1 | WP_308083992.1 protein phosphatase 2C domain-containing protein |
| 47(1) | WP_116353246.1 | WP_116353246.1 protein phosphatase 2C domain-containing protein |
| 47(1) | WP_099361340.1 | WP_099361340.1 serine/threonine-protein phosphatase |
| 47(1) | WP_268058021.1 | WP_268058021.1 protein phosphatase 2C domain-containing protein |
| 47(1) | WP_255464558.1 | WP_255464558.1 protein phosphatase 2C domain-containing protein |
| 47(1) | WP_034675375.1 | WP_034675375.1 protein phosphatase 2C domain-containing protein |
| 47(1) | WP_308083990.1 | WP_308083990.1 protein phosphatase 2C domain-containing protein |
| 47(1) | WP_143316572.1 | WP_143316572.1 protein phosphatase 2C domain-containing protein |
| 47(1) | WP_251481002.1 | WP_251481002.1 protein phosphatase 2C domain-containing protein |
| 47(1) | WP_136379704.1 | WP_136379704.1 protein phosphatase 2C domain-containing protein |
| 47(1) | WP_213143952.1 | WP_213143952.1 protein phosphatase 2C domain-containing protein |
| 47(1) | WP_215032115.1 | WP_215032115.1 protein phosphatase 2C domain-containing protein |
| 47(1) | WP_263324554.1 | WP_263324554.1 protein phosphatase 2C domain-containing protein |
| 47(1) | WP_042462820.1 | WP_042462820.1 protein phosphatase 2C domain-containing protein |
| 47(1) | WP_169238721.1 | WP_169238721.1 serine/threonine-protein phosphatase |
| 47(1) | WP_053361018.1 | WP_053361018.1 protein phosphatase 2C domain-containing protein |
| 47(1) | WP_066091989.1 | WP_066091989.1 protein phosphatase 2C domain-containing protein |

| | | |
|---|---|---|
| 48(1) | WP_098361429.1 | WP_098361429.1 alpha/beta hydrolase |
| 48(2) | WP_088020979.1 | WP_088020979.1 alpha/beta hydrolase |
| 48(1) | WP_084644562.1 | WP_084644562.1 alpha/beta hydrolase |
| 48(1) | WP_144488317.1 | WP_144488317.1 alpha/beta hydrolase |
| 48(1) | WP_197241461.1 | WP_197241461.1 alpha/beta hydrolase |
| 48(1) | WP_000584218.1 | WP_000584218.1 alpha/beta hydrolase |
| 48(1) | WP_144612051.1 | WP_144612051.1 alpha/beta hydrolase |
| 48(1) | WP_147807647.1 | WP_147807647.1 alpha/beta hydrolase |
| 48(1) | WP_306007807.1 | WP_306007807.1 alpha/beta hydrolase |
| 48(1) | WP_074620253.1 | WP_074620253.1 alpha/beta hydrolase |
| 48(2) | WP_098264292.1 | WP_098264292.1 alpha/beta hydrolase |
| 48(1) | WP_000774252.1 | WP_000774252.1 alpha/beta hydrolase |
| 48(1) | WP_097920535.1 | WP_097920535.1 alpha/beta hydrolase |
| 48(1) | WP_060630653.1 | WP_060630653.1 alpha/beta hydrolase |
| 48(1) | WP_098841813.1 | WP_098841813.1 alpha/beta hydrolase |
| 48(1) | WP_080776495.1 | WP_080776495.1 alpha/beta hydrolase |
| 48(2) | WP_098266656.1 | WP_098266656.1 alpha/beta hydrolase |
| 48(1) | WP_144564316.1 | WP_144564316.1 alpha/beta hydrolase |
| 48(1) | WP_088063238.1 | WP_088063238.1 alpha/beta fold hydrolase |
| 48(1) | WP_001284948.1 | WP_001284948.1 alpha/beta hydrolase |
| 48(1) | WP_215570622.1 | WP_215570622.1 alpha/beta hydrolase |
| 48(4) | WP_000584222.1 | WP_000584222.1 alpha/beta hydrolase |
| 48(1) | WP_098776422.1 | WP_098776422.1 alpha/beta hydrolase |
| 48(1) | WP_215598784.1 | WP_215598784.1 alpha/beta hydrolase |
| 48(1) | WP_151630593.1 | WP_151630593.1 alpha/beta fold hydrolase |
| 48(1) | WP_098255402.1 | WP_098255402.1 alpha/beta hydrolase |
| 48(1) | WP_149238908.1 | WP_149238908.1 alpha/beta hydrolase |
| 48(2) | WP_075719814.1 | WP_075719814.1 alpha/beta hydrolase |
| 48(1) | WP_097905170.1 | WP_097905170.1 alpha/beta hydrolase |
| 48(1) | WP_098856775.1 | WP_098856775.1 alpha/beta hydrolase |
| 48(1) | WP_270723901.1 | WP_270723901.1 alpha/beta hydrolase |
| 48(1) | WP_098365943.1 | WP_098365943.1 alpha/beta hydrolase |
| 48(1) | WP_199695257.1 | WP_199695257.1 alpha/beta hydrolase |
| 48(2) | WP_094194508.1 | WP_094194508.1 alpha/beta hydrolase |
| 48(1) | WP_277480703.1 | WP_277480703.1 alpha/beta hydrolase |
| 48(1) | WP_166705012.1 | WP_166705012.1 alpha/beta hydrolase |
| 48(4) | WP_103626237.1 | WP_103626237.1 alpha/beta hydrolase |
| 48(1) | WP_088119574.1 | WP_088119574.1 alpha/beta hydrolase |
| 48(1) | WP_078185399.1 | WP_078185399.1 alpha/beta hydrolase |
| 48(1) | WP_215591205.1 | WP_215591205.1 alpha/beta hydrolase |

| | | |
|---|---|---|
| 48(1) | WP_173601445.1 | WP_173601445.1 alpha/beta hydrolase |
| 48(1) | WP_145959662.1 | WP_145959662.1 alpha/beta hydrolase |
| 48(2) | WP_088609118.1 | WP_088609118.1 alpha/beta hydrolase |
| 48(1) | WP_080448755.1 | WP_080448755.1 alpha/beta hydrolase |
| 48(1) | WP_000584223.1 | WP_000584223.1 alpha/beta hydrolase |
| 48(1) | WP_262748103.1 | WP_262748103.1 alpha/beta hydrolase |
| 48(1) | WP_270728534.1 | WP_270728534.1 alpha/beta hydrolase |
| 48(1) | WP_002118924.1 | WP_002118924.1 alpha/beta hydrolase |
| 48(1) | WP_212957601.1 | WP_212957601.1 alpha/beta hydrolase |
| 48(1) | WP_087956299.1 | WP_087956299.1 alpha/beta hydrolase |
| 48(1) | WP_141558532.1 | WP_141558532.1 alpha/beta hydrolase |
| 48(1) | WP_270699387.1 | WP_270699387.1 alpha/beta hydrolase |
| 48(1) | WP_098943214.1 | WP_098943214.1 alpha/beta hydrolase |
| 48(1) | WP_143881764.1 | WP_143881764.1 alpha/beta hydrolase |
| 48(1) | WP_083322705.1 | WP_083322705.1 alpha/beta hydrolase |
| 48(1) | WP_098830141.1 | WP_098830141.1 alpha/beta hydrolase |
| 48(1) | WP_098254017.1 | WP_098254017.1 alpha/beta hydrolase |
| 48(5) | WP_087982571.1 | WP_087982571.1 alpha/beta fold hydrolase |
| 48(1) | WP_078181476.1 | WP_078181476.1 alpha/beta hydrolase |
| 48(1) | WP_071719975.1 | WP_071719975.1 alpha/beta hydrolase |
| 48(1) | WP_163247671.1 | WP_163247671.1 alpha/beta hydrolase |
| 48(1) | WP_082250274.1 | WP_082250274.1 alpha/beta hydrolase |
| 48(1) | WP_262725722.1 | WP_262725722.1 alpha/beta hydrolase |
| 48(1) | WP_000584225.1 | WP_000584225.1 alpha/beta hydrolase |
| 48(1) | WP_163255265.1 | WP_163255265.1 alpha/beta hydrolase |
| 48(1) | WP_072938294.1 | WP_072938294.1 alpha/beta hydrolase |
| 48(1) | WP_215572756.1 | WP_215572756.1 alpha/beta hydrolase |
| 48(1) | WP_097911834.1 | WP_097911834.1 alpha/beta hydrolase |
| 48(1) | WP_073540581.1 | WP_073540581.1 alpha/beta hydrolase |
| 48(2) | WP_139895591.1 | WP_139895591.1 alpha/beta hydrolase |
| 48(1) | WP_098150440.1 | WP_098150440.1 alpha/beta hydrolase |
| 48(2) | WP_048552319.1 | WP_048552319.1 alpha/beta hydrolase |
| 48(2) | WP_097925468.1 | WP_097925468.1 alpha/beta hydrolase |
| 48(6) | WP_000584227.1 | WP_000584227.1 alpha/beta hydrolase |
| 48(1) | WP_088078467.1 | WP_088078467.1 alpha/beta hydrolase |
| 48(1) | WP_098961911.1 | WP_098961911.1 alpha/beta hydrolase |
| 48(1) | WP_080468526.1 | WP_080468526.1 alpha/beta hydrolase |
| 48(1) | WP_098319058.1 | WP_098319058.1 alpha/beta hydrolase |
| 49(1) | WP_214706005.1 | WP_214706005.1 hypothetical protein |
| 49(1) | WP_024027631.1 | WP_024027631.1 hypothetical protein |
| 49(3) | WP_213117483.1 | WP_213117483.1 hypothetical protein |
| 49(1) | WP_308083987.1 | WP_308083987.1 hypothetical protein |
| 49(1) | WP_231094847.1 | WP_231094847.1 hypothetical protein |
| 49(1) | WP_221875378.1 | WP_221875378.1 hypothetical protein |
| 49(1) | WP_289342264.1 | WP_289342264.1 hypothetical protein |
| 49(1) | WP_307284686.1 | WP_307284686.1 hypothetical protein |
| 49(3) | WP_041964575.1 | WP_041964575.1 hypothetical protein |
| 49(2) | WP_251426364.1 | WP_251426364.1 hypothetical protein |
| 49(1) | WP_239617468.1 | WP_239617468.1 hypothetical protein |
| 49(1) | WP_099354910.1 | WP_099354910.1 hypothetical protein |
| 49(1) | WP_010676862.1 | WP_010676862.1 hypothetical protein |
| 49(1) | WP_066091998.1 | WP_066091998.1 hypothetical protein |
| 49(1) | WP_013490126.1 | WP_013490126.1 hypothetical protein |
| 49(1) | WP_179603478.1 | WP_179603478.1 hypothetical protein |
| 49(1) | WP_133370057.1 | WP_133370057.1 hypothetical protein |
| 49(1) | WP_223592684.1 | WP_223592684.1 hypothetical protein |
| 49(1) | WP_204445208.1 | WP_204445208.1 hypothetical protein |
| 49(1) | WP_251548757.1 | WP_251548757.1 hypothetical protein |
| 49(1) | WP_254759299.1 | WP_254759299.1 hypothetical protein |
| 49(1) | WP_068673702.1 | WP_068673702.1 hypothetical protein |
| 49(1) | WP_077619646.1 | WP_077619646.1 hypothetical protein |
| 49(1) | WP_210364454.1 | WP_210364454.1 hypothetical protein |
| 49(1) | WP_263324550.1 | WP_263324550.1 hypothetical protein |
| 49(1) | WP_122971538.1 | WP_122971538.1 hypothetical protein |
| 49(1) | WP_306077387.1 | WP_306077387.1 hypothetical protein |
| 49(1) | WP_254366008.1 | WP_254366008.1 hypothetical protein |
| 49(2) | WP_144479059.1 | WP_144479059.1 hypothetical protein |
| 49(1) | WP_307589606.1 | WP_307589606.1 hypothetical protein |
| 49(1) | WP_251638768.1 | WP_251638768.1 hypothetical protein |
| 49(1) | WP_182301830.1 | WP_182301830.1 hypothetical protein |
| 49(1) | WP_215032121.1 | WP_215032121.1 hypothetical protein |
| 49(2) | WP_204412722.1 | WP_204412722.1 hypothetical protein |
| 49(1) | WP_191272860.1 | WP_191272860.1 hypothetical protein |
| 49(3) | WP_025848609.1 | WP_025848609.1 hypothetical protein |
| 49(1) | WP_127459431.1 | WP_127459431.1 hypothetical protein |
| 49(1) | WP_108534958.1 | WP_108534958.1 hypothetical protein |
| 49(1) | WP_066419988.1 | WP_066419988.1 hypothetical protein |
| 49(1) | WP_241993931.1 | WP_241993931.1 hypothetical protein |
| 49(1) | WP_048826575.1 | WP_048826575.1 hypothetical protein |
| 49(1) | WP_179160619.1 | WP_179160619.1 hypothetical protein |
| 49(1) | WP_253411075.1 | WP_253411075.1 hypothetical protein |
| 49(1) | WP_208926605.1 | WP_208926605.1 hypothetical protein |
| 49(1) | WP_048826562.1 | WP_048826562.1 hypothetical protein |
| 49(1) | WP_157261479.1 | WP_157261479.1 hypothetical protein |
| 49(1) | WP_045518281.1 | WP_045518281.1 hypothetical protein |
| 49(1) | WP_306049027.1 | WP_306049027.1 hypothetical protein |
| 49(1) | WP_042462803.1 | WP_042462803.1 hypothetical protein |
| 49(1) | WP_195515238.1 | WP_195515238.1 hypothetical protein |
| 49(1) | WP_216791216.1 | WP_216791216.1 hypothetical protein |
| 49(1) | WP_116353250.1 | WP_116353250.1 hypothetical protein |
| 49(1) | WP_066022452.1 | WP_066022452.1 hypothetical protein |
| 49(1) | WP_251032277.1 | WP_251032277.1 hypothetical protein |
| 49(2) | WP_036689408.1 | WP_036689408.1 hypothetical protein |
| 49(1) | WP_015595758.1 | WP_015595758.1 DUF1732 domain-containing protein |
| 49(1) | WP_136379701.1 | WP_136379701.1 hypothetical protein |
| 49(1) | WP_134750600.1 | WP_134750600.1 hypothetical protein |
| 49(1) | WP_213143948.1 | WP_213143948.1 hypothetical protein |
| 49(2) | WP_163853474.1 | WP_163853474.1 hypothetical protein |
| 49(1) | WP_226679101.1 | WP_226679101.1 hypothetical protein |
| 49(1) | WP_090674369.1 | WP_090674369.1 hypothetical protein |
| 49(1) | WP_029191602.1 | WP_029191602.1 hypothetical protein |
| 49(1) | WP_167831138.1 | WP_167831138.1 hypothetical protein |
| 49(1) | WP_149871681.1 | WP_149871681.1 hypothetical protein |

| | | |
|---|---|---|
| 49(1) | WP_007083291.1 | WP_007083291.1 hypothetical protein |
| 49(1) | WP_090761823.1 | WP_090761823.1 hypothetical protein |
| 49(1) | WP_111147799.1 | WP_111147799.1 hypothetical protein |
| 49(1) | WP_270065056.1 | WP_270065056.1 hypothetical protein |
| 49(1) | WP_251542613.1 | WP_251542613.1 hypothetical protein |
| 49(1) | WP_307312556.1 | WP_307312556.1 hypothetical protein |
| 49(1) | WP_186580081.1 | WP_186580081.1 hypothetical protein |
| 49(1) | WP_231687842.1 | WP_231687842.1 hypothetical protein |
| 49(1) | WP_241481488.1 | WP_241481488.1 hypothetical protein |
| 49(1) | WP_281937372.1 | WP_281937372.1 hypothetical protein |
| 49(1) | WP_235286204.1 | WP_235286204.1 hypothetical protein |
| 49(3) | WP_099361338.1 | WP_099361338.1 hypothetical protein |
| 49(1) | WP_267497945.1 | WP_267497945.1 hypothetical protein |
| 49(1) | WP_079506398.1 | WP_079506398.1 hypothetical protein |
| 49(1) | WP_063188062.1 | WP_063188062.1 hypothetical protein |
| 49(1) | WP_308082247.1 | WP_308082247.1 hypothetical protein |
| 49(1) | WP_251480998.1 | WP_251480998.1 hypothetical protein |
| 49(2) | WP_146740512.1 | WP_146740512.1 hypothetical protein |

| | | |
|---|---|---|
| 50(1) | WP_270699388.1 | WP_270699388.1 nucleotidyltransferase family protein |
| 50(5) | WP_001025800.1 | WP_001025800.1 nucleotidyltransferase family protein |
| 50(1) | WP_098199515.1 | WP_098199515.1 nucleotidyltransferase family protein |
| 50(1) | WP_002012589.1 | WP_002012589.1 nucleotidyltransferase family protein |
| 50(1) | WP_098361428.1 | WP_098361428.1 nucleotidyltransferase family protein |
| 50(1) | WP_001022656.1 | WP_001022656.1 nucleotidyltransferase family protein |
| 50(2) | WP_069355515.1 | WP_069355515.1 nucleotidyltransferase family protein |
| 50(1) | WP_098266657.1 | WP_098266657.1 nucleotidyltransferase family protein |
| 50(1) | WP_088308515.1 | WP_088308515.1 nucleotidyltransferase family protein |
| 50(1) | WP_001025798.1 | WP_001025798.1 nucleotidyltransferase family protein |
| 50(2) | WP_065483400.1 | WP_065483400.1 nucleotidyltransferase family protein |
| 50(1) | WP_074565292.1 | WP_074565292.1 nucleotidyltransferase family protein |
| 50(3) | WP_060630654.1 | WP_060630654.1 nucleotidyltransferase family protein |
| 50(1) | WP_144611198.1 | WP_144611198.1 nucleotidyltransferase family protein |
| 50(4) | WP_103626238.1 | WP_103626238.1 nucleotidyltransferase family protein |
| 50(1) | WP_144564273.1 | WP_144564273.1 nucleotidyltransferase family protein |
| 50(1) | WP_097949254.1 | WP_097949254.1 nucleotidyltransferase family protein |
| 50(1) | WP_139867358.1 | WP_139867358.1 nucleotidyltransferase family protein |
| 50(8) | WP_000155289.1 | WP_000155289.1 nucleotidyltransferase family protein |
| 50(1) | WP_001025804.1 | WP_001025804.1 nucleotidyltransferase family protein |
| 50(1) | WP_002031673.1 | WP_002031673.1 nucleotidyltransferase family protein |
| 50(1) | WP_097987484.1 | WP_097987484.1 nucleotidyltransferase family protein |
| 50(1) | WP_262725721.1 | WP_262725721.1 nucleotidyltransferase family protein |
| 50(1) | WP_071719974.1 | WP_071719974.1 nucleotidyltransferase family protein |
| 50(1) | WP_106822146.1 | WP_106822146.1 nucleotidyltransferase family protein |
| 50(2) | WP_131231937.1 | WP_131231937.1 nucleotidyltransferase family protein |
| 50(7) | WP_001025803.1 | WP_001025803.1 nucleotidyltransferase family protein |
| 50(1) | WP_098400682.1 | WP_098400682.1 nucleotidyltransferase family protein |
| 50(1) | WP_097911832.1 | WP_097911832.1 nucleotidyltransferase family protein |
| 50(1) | WP_088609119.1 | WP_088609119.1 nucleotidyltransferase family protein |
| 50(1) | WP_098856776.1 | WP_098856776.1 nucleotidyltransferase family protein |
| 50(8) | WP_087982569.1 | WP_087982569.1 nucleotidyltransferase family protein |
| 50(1) | WP_166705004.1 | WP_166705004.1 nucleotidyltransferase family protein |

| | | |
|---|---|---|
| 50(1) | WP_088078468.1 | WP_088078468.1 nucleotidyltransferase family protein |
| 50(2) | WP_139895503.1 | WP_139895503.1 nucleotidyltransferase family protein |
| 50(1) | WP_000155290.1 | WP_000155290.1 nucleotidyltransferase family protein |
| 50(1) | WP_270728536.1 | WP_270728536.1 nucleotidyltransferase family protein |
| 50(1) | WP_088327136.1 | WP_088327136.1 nucleotidyltransferase family protein |
| 50(3) | WP_090976445.1 | WP_090976445.1 nucleotidyltransferase family protein |
| 50(1) | WP_098204518.1 | WP_098204518.1 nucleotidyltransferase family protein |
| 50(1) | WP_087980963.1 | WP_087980963.1 nucleotidyltransferase family protein |
| 50(1) | WP_097852385.1 | WP_097852385.1 nucleotidyltransferase family protein |
| 50(1) | WP_098680261.1 | WP_098680261.1 nucleotidyltransferase family protein |
| 50(2) | WP_087956300.1 | WP_087956300.1 nucleotidyltransferase family protein |
| 50(1) | WP_098365944.1 | WP_098365944.1 nucleotidyltransferase family protein |
| 50(1) | WP_088058545.1 | WP_088058545.1 nucleotidyltransferase family protein |
| 50(1) | WP_078181477.1 | WP_078181477.1 nucleotidyltransferase family protein |
| 50(1) | WP_061688410.1 | WP_061688410.1 nucleotidyltransferase family protein |
| 50(1) | WP_151630592.1 | WP_151630592.1 nucleotidyltransferase family protein |
| 50(2) | WP_097961547.1 | WP_097961547.1 nucleotidyltransferase family protein |
| 50(1) | WP_098088231.1 | WP_098088231.1 nucleotidyltransferase family protein |
| 50(3) | WP_048552321.1 | WP_048552321.1 nucleotidyltransferase family protein |
| 50(1) | WP_215598783.1 | WP_215598783.1 nucleotidyltransferase family protein |
| 50(1) | WP_070805906.1 | WP_070805906.1 nucleotidyltransferase family protein |
| 50(1) | WP_212957598.1 | WP_212957598.1 nucleotidyltransferase family protein |

| | | |
|---|---|---|
| 51(1) | WP_000288829.1 | WP_000288829.1 HAD family hydrolase |
| 51(1) | WP_215572757.1 | WP_215572757.1 HAD family hydrolase |
| 51(2) | WP_087956302.1 | WP_087956302.1 HAD family hydrolase |
| 51(1) | WP_242172387.1 | WP_242172387.1 beta-phosphoglucomutase |
| 51(1) | WP_088063240.1 | WP_088063240.1 HAD family hydrolase |
| 51(1) | WP_257675212.1 | WP_257675212.1 HAD family hydrolase |
| 51(1) | WP_098239605.1 | WP_098239605.1 HAD family hydrolase |
| 51(2) | WP_010963740.1 | WP_010963740.1 HAD family hydrolase |
| 51(1) | WP_147786232.1 | WP_147786232.1 HAD family hydrolase |
| 51(1) | WP_259392012.1 | WP_259392012.1 HAD family hydrolase |
| 51(1) | WP_197256844.1 | WP_197256844.1 HAD family hydrolase |
| 51(1) | WP_098841812.1 | WP_098841812.1 HAD family hydrolase |
| 51(1) | WP_097852383.1 | WP_097852383.1 HAD family hydrolase |
| 51(2) | WP_097961548.1 | WP_097961548.1 HAD family hydrolase |
| 51(1) | WP_223067444.1 | WP_223067444.1 HAD family hydrolase |
| 51(3) | WP_168178776.1 | WP_168178776.1 HAD family hydrolase |
| 51(1) | WP_087442423.1 | WP_087442423.1 HAD family hydrolase |
| 51(1) | WP_070805908.1 | WP_070805908.1 HAD family hydrolase |
| 51(1) | WP_013490143.1 | WP_013490143.1 pyrophosphatase PpaX |
| 51(1) | WP_098266659.1 | WP_098266659.1 HAD family hydrolase |
| 51(1) | WP_077852408.1 | WP_077852408.1 HAD family hydrolase |
| 51(1) | WP_262719883.1 | WP_262719883.1 HAD family hydrolase |
| 51(1) | WP_251548772.1 | WP_251548772.1 YjjG family noncanonical pyrimidine nucleotidase |
| 51(1) | WP_231094862.1 | WP_231094862.1 HAD hydrolase-like protein |
| 51(1) | WP_272045556.1 | WP_272045556.1 HAD family hydrolase |
| 51(1) | WP_077893576.1 | WP_077893576.1 HAD family hydrolase |
| 51(2) | WP_006674850.1 | WP_006674850.1 HAD family hydrolase |
| 51(2) | WP_048552325.1 | WP_048552325.1 HAD family hydrolase |
| 51(4) | WP_000278475.1 | WP_000278475.1 HAD family hydrolase |

| | | |
|---|---|---|
| 51(1) | WP_065228773.1 | WP_065228773.1 HAD family hydrolase |
| 51(1) | WP_098088229.1 | WP_098088229.1 HAD family hydrolase |
| 51(1) | WP_088058547.1 | WP_088058547.1 HAD family hydrolase |
| 51(1) | WP_053403790.1 | WP_053403790.1 beta-phosphoglucomutase |
| 51(1) | WP_071717054.1 | WP_071717054.1 HAD family hydrolase |
| 51(1) | WP_088609121.1 | WP_088609121.1 HAD family hydrolase |
| 51(1) | WP_165979807.1 | WP_165979807.1 HAD family hydrolase |
| 51(2) | WP_000288832.1 | WP_000288832.1 HAD family hydrolase |
| 51(1) | WP_098834783.1 | WP_098834783.1 HAD family hydrolase |
| 51(1) | WP_066020831.1 | WP_066020831.1 HAD family hydrolase |
| 51(1) | WP_270699389.1 | WP_270699389.1 HAD family hydrolase |
| 51(1) | WP_088667123.1 | WP_088667123.1 HAD family hydrolase |
| 51(3) | WP_002158031.1 | WP_002158031.1 HAD family hydrolase |
| 51(1) | WP_098264294.1 | WP_098264294.1 HAD family hydrolase |
| 51(4) | WP_087952388.1 | WP_087952388.1 HAD family hydrolase |
| 51(2) | WP_053564617.1 | WP_053564617.1 HAD family hydrolase |
| 51(1) | WP_108467786.1 | WP_108467786.1 HAD family hydrolase |
| 51(1) | WP_077833097.1 | WP_077833097.1 HAD family hydrolase |
| 51(2) | WP_000288833.1 | WP_000288833.1 HAD family hydrolase |
| 51(1) | WP_213426981.1 | WP_213426981.1 HAD family hydrolase |
| 51(1) | WP_061688411.1 | WP_061688411.1 HAD family hydrolase |
| 51(1) | WP_071719971.1 | WP_071719971.1 HAD family hydrolase |
| 51(5) | WP_069355516.1 | WP_069355516.1 HAD family hydrolase |
| 51(1) | WP_139895502.1 | WP_139895502.1 HAD family hydrolase |
| 51(1) | WP_251610753.1 | WP_251610753.1 beta-phosphoglucomutase |
| 51(1) | WP_098961912.1 | WP_098961912.1 HAD family hydrolase |
| 51(1) | WP_060630656.1 | WP_060630656.1 HAD family hydrolase |
| 51(1) | WP_098856777.1 | WP_098856777.1 HAD family hydrolase |
| 51(1) | WP_281477433.1 | WP_281477433.1 HAD family hydrolase |
| 51(1) | WP_270728540.1 | WP_270728540.1 HAD family hydrolase |
| 51(2) | WP_094870391.1 | WP_094870391.1 beta-phosphoglucomutase |
| 51(3) | WP_087982565.1 | WP_087982565.1 HAD family hydrolase |
| 51(1) | WP_012260851.1 | WP_012260851.1 HAD family hydrolase |
| 51(1) | WP_270711677.1 | WP_270711677.1 HAD family hydrolase |
| 51(2) | WP_143801052.1 | WP_143801052.1 HAD family hydrolase |
| 51(1) | WP_204445241.1 | WP_204445241.1 HAD family hydrolase |
| 51(1) | WP_042462835.1 | WP_042462835.1 YjjG family noncanonical pyrimidine nucleotidase |
| 51(1) | WP_237690764.1 | WP_237690764.1 HAD family phosphatase |

| | | |
|---|---|---|
| 52(1) | WP_199613911.1 | WP_199613911.1 hypothetical protein |
| 52(1) | WP_128631278.1 | WP_128631278.1 ATPase |
| 52(2) | WP_094870543.1 | WP_094870543.1 hypothetical protein |
| 52(1) | WP_136479368.1 | WP_136479368.1 ATPase |
| 52(1) | WP_102710712.1 | WP_102710712.1 ATPase |
| 52(2) | WP_156558884.1 | WP_156558884.1 AAA family ATPase |
| 52(1) | WP_036702429.1 | WP_036702429.1 hypothetical protein |
| 52(1) | WP_195343917.1 | WP_195343917.1 AAA family ATPase |
| 52(2) | WP_076271933.1 | WP_076271933.1 hypothetical protein |
| 52(1) | WP_271839908.1 | WP_271839908.1 AAA family ATPase |
| 52(1) | WP_077853216.1 | WP_077853216.1 AAA family ATPase |
| 52(2) | WP_101300029.1 | WP_101300029.1 ATPase |
| 52(1) | WP_077832282.1 | WP_077832282.1 AAA family ATPase |
| 52(1) | WP_197937385.1 | WP_197937385.1 hypothetical protein |
| 52(1) | WP_136790002.1 | WP_136790002.1 CpsD/CapB family tyrosine-protein kinase |
| 52(1) | WP_179034779.1 | WP_179034779.1 ATPase |
| 52(1) | WP_076157847.1 | WP_076157847.1 hypothetical protein |
| 52(1) | WP_143315253.1 | WP_143315253.1 AAA family ATPase |
| 52(2) | WP_195431323.1 | WP_195431323.1 AAA family ATPase |
| 52(1) | WP_238655340.1 | WP_238655340.1 hypothetical protein |
| 52(1) | WP_049869160.1 | WP_049869160.1 hypothetical protein |
| 52(1) | WP_150457774.1 | WP_150457774.1 ATPase |
| 52(1) | WP_291647968.1 | WP_291647968.1 AAA family ATPase |
| 52(1) | WP_084117735.1 | WP_084117735.1 AAA family ATPase |
| 52(2) | WP_076301354.1 | WP_076301354.1 hypothetical protein |
| 52(1) | WP_119601508.1 | WP_119601508.1 ATPase |
| 52(1) | WP_181603058.1 | WP_181603058.1 AAA family ATPase |
| 52(1) | WP_283910916.1 | WP_283910916.1 hypothetical protein |
| 52(1) | WP_189031047.1 | WP_189031047.1 hypothetical protein |
| 52(1) | WP_291682714.1 | WP_291682714.1 AAA family ATPase |
| 52(1) | WP_126996293.1 | WP_126996293.1 ATPase |
| 52(1) | WP_128705704.1 | WP_128705704.1 ATPase |
| 52(1) | WP_036626786.1 | WP_036626786.1 hypothetical protein |
| 52(6) | WP_036684503.1 | WP_036684503.1 hypothetical protein |
| 52(3) | WP_013914197.1 | WP_013914197.1 AAA family ATPase |
| 52(1) | WP_195571406.1 | WP_195571406.1 ATPase |
| 52(1) | WP_165097599.1 | WP_165097599.1 ATPase |
| 52(1) | WP_044037273.1 | WP_044037273.1 AAA family ATPase |
| 52(1) | WP_026478101.1 | WP_026478101.1 hypothetical protein |
| 52(1) | WP_169028899.1 | WP_169028899.1 ATPase |
| 52(2) | WP_213592881.1 | WP_213592881.1 hypothetical protein |
| 52(1) | WP_003516560.1 | WP_003516560.1 hypothetical protein |
| 52(1) | WP_243735260.1 | WP_243735260.1 AAA family ATPase |
| 52(1) | WP_013658721.1 | WP_013658721.1 hypothetical protein |
| 52(1) | WP_185165646.1 | WP_185165646.1 AAA family ATPase |
| 52(1) | WP_215185462.1 | WP_215185462.1 hypothetical protein |
| 52(1) | WP_244723806.1 | WP_244723806.1 hypothetical protein |
| 52(1) | WP_120462959.1 | WP_120462959.1 ATPase |
| 52(1) | WP_076119199.1 | WP_076119199.1 hypothetical protein |
| 52(1) | WP_249897307.1 | WP_249897307.1 hypothetical protein |
| 52(1) | WP_236334349.1 | WP_236334349.1 hypothetical protein |
| 52(2) | WP_148450955.1 | WP_148450955.1 CpsD/CapB family tyrosine-protein kinase |
| 52(1) | WP_077893467.1 | WP_077893467.1 AAA family ATPase |
| 52(1) | WP_155609764.1 | WP_155609764.1 ATPase |
| 52(1) | WP_055109401.1 | WP_055109401.1 hypothetical protein |
| 52(1) | WP_251417642.1 | WP_251417642.1 hypothetical protein |
| 52(1) | WP_287600030.1 | WP_287600030.1 AAA family ATPase |
| 52(2) | WP_010963366.1 | WP_010963366.1 AAA family ATPase |
| 52(1) | WP_125084791.1 | WP_125084791.1 ATPase |
| 52(1) | WP_188535054.1 | WP_188535054.1 ATPase |
| 52(1) | WP_108464079.1 | WP_108464079.1 ATPase |
| 52(1) | WP_036944562.1 | WP_036944562.1 ATPase |
| 52(1) | WP_216390033.1 | WP_216390033.1 AAA family ATPase |
| 52(1) | WP_223068036.1 | WP_223068036.1 hypothetical protein |

| | | |
|---|---|---|
| 52(1) | WP_280787398.1 | WP_280787398.1 hypothetical protein |
| 52(1) | WP_076281083.1 | WP_076281083.1 hypothetical protein |
| 52(1) | WP_251596690.1 | WP_251596690.1 hypothetical protein |
| 52(1) | WP_291630878.1 | WP_291630878.1 AAA family ATPase |
| 52(1) | WP_276020676.1 | WP_276020676.1 ATPase |
| 52(1) | WP_257443659.1 | WP_257443659.1 hypothetical protein |
| 52(2) | WP_085499118.1 | WP_085499118.1 hypothetical protein |
| 52(1) | WP_042125609.1 | WP_042125609.1 hypothetical protein |
| 52(2) | WP_211022797.1 | WP_211022797.1 hypothetical protein |
| 52(1) | WP_132283082.1 | WP_132283082.1 ATPase |
| 52(1) | WP_110934082.1 | WP_110934082.1 ATPase |

| | | |
|---|---|---|
| 53(1) | WP_290402003.1 | WP_290402003.1 DUF2085 domain-containing protein |
| 53(3) | WP_098266638.1 | WP_098266638.1 DUF2085 domain-containing protein |
| 53(44) | WP_000620803.1 | WP_000620803.1 DUF2085 domain-containing protein |
| 53(5) | WP_062821093.1 | WP_062821093.1 DUF2085 domain-containing protein |
| 53(1) | WP_289483790.1 | WP_289483790.1 DUF2085 domain-containing protein |
| 53(1) | WP_002148356.1 | WP_002148356.1 DUF2085 domain-containing protein |
| 53(1) | WP_088058513.1 | WP_088058513.1 DUF2085 domain-containing protein |
| 53(1) | WP_098319064.1 | WP_098319064.1 DUF2085 domain-containing protein |
| 53(1) | WP_163258049.1 | WP_163258049.1 DUF2085 domain-containing protein |
| 53(2) | WP_002141578.1 | WP_002141578.1 DUF2085 domain-containing protein |
| 53(1) | WP_242400014.1 | WP_242400014.1 DUF2085 domain-containing protein |
| 53(1) | WP_215591194.1 | WP_215591194.1 DUF2085 domain-containing protein |
| 53(1) | WP_242251755.1 | WP_242251755.1 DUF2085 domain-containing protein |
| 53(1) | WP_301167508.1 | WP_301167508.1 DUF2085 domain-containing protein |
| 53(1) | WP_087952367.1 | WP_087952367.1 DUF2085 domain-containing protein |
| 53(1) | WP_090239246.1 | WP_090239246.1 DUF2085 domain-containing protein |
| 53(1) | WP_062239843.1 | WP_062239843.1 DUF2085 domain-containing protein |
| 53(1) | WP_212805118.1 | WP_212805118.1 DUF2085 domain-containing protein |
| 53(1) | WP_232274497.1 | WP_232274497.1 DUF2085 domain-containing protein |
| 53(2) | WP_078181754.1 | WP_078181754.1 DUF2085 domain-containing protein |
| 53(1) | WP_289475149.1 | WP_289475149.1 DUF2085 domain-containing protein |
| 53(2) | WP_098991582.1 | WP_098991582.1 DUF2085 domain-containing protein |
| 53(1) | WP_071719993.1 | WP_071719993.1 DUF2085 domain-containing protein |
| 53(1) | WP_215572745.1 | WP_215572745.1 DUF2085 domain-containing protein |
| 53(1) | WP_074440643.1 | WP_074440643.1 DUF2085 domain-containing protein |
| 53(2) | WP_002127167.1 | WP_002127167.1 DUF2085 domain-containing protein |
| 53(1) | WP_248251192.1 | WP_248251192.1 DUF2085 domain-containing protein |
| 53(1) | WP_283892191.1 | WP_283892191.1 DUF2085 domain-containing protein |
| 53(1) | WP_265408836.1 | WP_265408836.1 DUF2085 domain-containing protein |
| 53(1) | WP_012260824.1 | WP_012260824.1 DUF2085 domain-containing protein |
| 53(1) | WP_033685962.1 | WP_033685962.1 DUF2085 domain-containing protein |
| 53(1) | WP_129478235.1 | WP_129478235.1 DUF2085 domain-containing protein |
| 53(1) | WP_000620801.1 | WP_000620801.1 DUF2085 domain-containing protein |
| 53(4) | WP_078205517.1 | WP_078205517.1 DUF2085 domain-containing protein |
| 53(1) | WP_256227724.1 | WP_256227724.1 DUF2085 domain-containing protein |

| | | |
|---|---|---|
| 54(1) | WP_160496946.1 | WP_160496946.1 NUDIX domain-containing protein |
| 54(1) | WP_244719557.1 | WP_244719557.1 NUDIX domain-containing protein |
| 54(1) | WP_307413613.1 | WP_307413613.1 NUDIX domain-containing protein |
| 54(1) | WP_145329889.1 | WP_145329889.1 NUDIX domain-containing protein |
| 54(1) | WP_023957689.1 | WP_023957689.1 NUDIX domain-containing protein |
| 54(1) | WP_062326137.1 | WP_062326137.1 NUDIX domain-containing protein |
| 54(1) | WP_307214299.1 | WP_307214299.1 NUDIX domain-containing protein |
| 54(1) | WP_056700764.1 | WP_056700764.1 NUDIX domain-containing protein |
| 54(1) | WP_229106294.1 | WP_229106294.1 NUDIX domain-containing protein |
| 54(1) | WP_247902235.1 | WP_247902235.1 NUDIX domain-containing protein |
| 54(1) | WP_074094608.1 | WP_074094608.1 NUDIX domain-containing protein |
| 54(1) | WP_293202272.1 | WP_293202272.1 NUDIX domain-containing protein |
| 54(1) | WP_095287671.1 | WP_095287671.1 NUDIX domain-containing protein |
| 54(1) | WP_095358554.1 | WP_095358554.1 NUDIX domain-containing protein |
| 54(1) | WP_138226033.1 | WP_138226033.1 NUDIX domain-containing protein |
| 54(1) | WP_210107808.1 | WP_210107808.1 NUDIX domain-containing protein |
| 54(1) | WP_090921078.1 | WP_090921078.1 NUDIX domain-containing protein |
| 54(1) | WP_110892994.1 | WP_110892994.1 NUDIX domain-containing protein |
| 54(1) | WP_091017206.1 | WP_091017206.1 NUDIX domain-containing protein |
| 54(1) | WP_076318545.1 | WP_076318545.1 NUDIX domain-containing protein |
| 54(3) | WP_062833872.1 | WP_062833872.1 NUDIX domain-containing protein |
| 54(1) | WP_127543708.1 | WP_127543708.1 NUDIX domain-containing protein |
| 54(1) | WP_301879175.1 | WP_301879175.1 NUDIX domain-containing protein |
| 54(2) | WP_076249783.1 | WP_076249783.1 NUDIX domain-containing protein |
| 54(2) | WP_167434744.1 | WP_167434744.1 NUDIX domain-containing protein |
| 54(1) | WP_017689085.1 | WP_017689085.1 NUDIX domain-containing protein |
| 54(1) | WP_186380863.1 | WP_186380863.1 NUDIX domain-containing protein |
| 54(1) | WP_110822565.1 | WP_110822565.1 NUDIX domain-containing protein |
| 54(1) | WP_234121620.1 | WP_234121620.1 NUDIX hydrolase |
| 54(1) | WP_145412699.1 | WP_145412699.1 NUDIX domain-containing protein |
| 54(1) | WP_106765973.1 | WP_106765973.1 NUDIX domain-containing protein |
| 54(1) | WP_282321827.1 | WP_282321827.1 NUDIX domain-containing protein |
| 54(8) | WP_063566334.1 | WP_063566334.1 NUDIX domain-containing protein |
| 54(1) | WP_076208846.1 | WP_076208846.1 NUDIX domain-containing protein |
| 54(1) | WP_047842531.1 | WP_047842531.1 NUDIX domain-containing protein |
| 54(1) | WP_208919054.1 | WP_208919054.1 NUDIX hydrolase |
| 54(1) | WP_264929216.1 | WP_264929216.1 NUDIX domain-containing protein |
| 54(1) | WP_153978061.1 | WP_153978061.1 NUDIX domain-containing protein |
| 54(1) | WP_236417197.1 | WP_236417197.1 NUDIX domain-containing protein |
| 54(1) | WP_136604802.1 | WP_136604802.1 NUDIX domain-containing protein |
| 54(2) | WP_095265647.1 | WP_095265647.1 NUDIX domain-containing protein |
| 54(1) | WP_169505043.1 | WP_169505043.1 NUDIX domain-containing protein |
| 54(1) | WP_286458031.1 | WP_286458031.1 NUDIX domain-containing protein |
| 54(3) | WP_036609216.1 | WP_036609216.1 NUDIX domain-containing protein |
| 54(2) | WP_090896167.1 | WP_090896167.1 NUDIX domain-containing protein |
| 54(1) | WP_133384355.1 | WP_133384355.1 NUDIX domain-containing protein |
| 54(1) | WP_076289565.1 | WP_076289565.1 NUDIX domain-containing protein |
| 54(1) | WP_145318722.1 | WP_145318722.1 NUDIX domain-containing protein |
| 54(1) | WP_208203479.1 | WP_208203479.1 NUDIX domain-containing protein |
| 54(1) | WP_308721992.1 | WP_308721992.1 NUDIX domain-containing protein |
| 54(2) | WP_164848676.1 | WP_164848676.1 NUDIX domain-containing protein |
| 54(2) | WP_145402733.1 | WP_145402733.1 NUDIX domain-containing protein |
| 54(4) | WP_024630314.1 | WP_024630314.1 NUDIX domain-containing protein |
| 54(1) | WP_036707081.1 | WP_036707081.1 NUDIX domain-containing protein |
| 54(1) | WP_000035005.1 | WP_000035005.1 NUDIX hydrolase |
| 54(2) | WP_124114113.1 | WP_124114113.1 NUDIX domain-containing protein |

| | | |
|---|---|---|
| 54(1) | WP_123067105.1 | WP_123067105.1 NUDIX domain-containing protein |
| 54(1) | WP_053783877.1 | WP_053783877.1 NUDIX domain-containing protein |
| 54(1) | WP_285748152.1 | WP_285748152.1 NUDIX domain-containing protein |
| 54(1) | WP_181586519.1 | WP_181586519.1 NUDIX domain-containing protein |
| 54(1) | WP_160037473.1 | WP_160037473.1 NUDIX domain-containing protein |
| 54(1) | WP_192266433.1 | WP_192266433.1 NUDIX domain-containing protein |
| 54(1) | WP_111268951.1 | WP_111268951.1 NUDIX domain-containing protein |
| 54(1) | WP_272035284.1 | WP_272035284.1 NUDIX domain-containing protein |
| 54(1) | WP_289392065.1 | WP_289392065.1 NUDIX domain-containing protein |
| 54(1) | WP_213649981.1 | WP_213649981.1 NUDIX domain-containing protein |
| 54(1) | WP_175480540.1 | WP_175480540.1 NUDIX domain-containing protein |
| 54(2) | WP_194230415.1 | WP_194230415.1 NUDIX domain-containing protein |
| 54(1) | WP_127536201.1 | WP_127536201.1 NUDIX domain-containing protein |

| | | |
|---|---|---|
| 55(1) | WP_090908776.1 | WP_090908776.1 tRNA glutamyl-Q(34) synthetase GluQRS |
| 55(1) | WP_123797118.1 | WP_123797118.1 tRNA glutamyl-Q(34) synthetase GluQRS |
| 55(2) | WP_145156809.1 | WP_145156809.1 tRNA glutamyl-Q(34) synthetase GluQRS |
| 55(1) | WP_098746852.1 | WP_098746852.1 tRNA glutamyl-Q(34) synthetase GluQRS |
| 55(1) | WP_145949002.1 | WP_145949002.1 tRNA glutamyl-Q(34) synthetase GluQRS |
| 55(1) | WP_095398846.1 | WP_095398846.1 tRNA glutamyl-Q(34) synthetase GluQRS |
| 55(1) | WP_163872774.1 | WP_163872774.1 tRNA glutamyl-Q(34) synthetase GluQRS |
| 55(1) | WP_230875697.1 | WP_230875697.1 tRNA glutamyl-Q(34) synthetase GluQRS |
| 55(1) | WP_068694723.1 | WP_068694723.1 tRNA glutamyl-Q(34) synthetase GluQRS |
| 55(1) | WP_015736236.1 | WP_015736236.1 tRNA glutamyl-Q(34) synthetase GluQRS |
| 55(1) | WP_244864498.1 | WP_244864498.1 tRNA glutamyl-Q(34) synthetase GluQRS |
| 55(1) | WP_108465357.1 | WP_108465357.1 tRNA glutamyl-Q(34) synthetase GluQRS |
| 55(1) | WP_089546603.1 | WP_089546603.1 tRNA glutamyl-Q(34) synthetase GluQRS |
| 55(1) | WP_128705698.1 | WP_128705698.1 glutamate--tRNA ligase |
| 55(1) | WP_290409176.1 | WP_290409176.1 tRNA glutamyl-Q(34) synthetase GluQRS |
| 55(1) | WP_246072899.1 | WP_246072899.1 tRNA glutamyl-Q(34) synthetase GluQRS |
| 55(1) | WP_263880215.1 | WP_263880215.1 tRNA glutamyl-Q(34) synthetase GluQRS |
| 55(1) | WP_244719660.1 | WP_244719660.1 tRNA glutamyl-Q(34) synthetase GluQRS |
| 55(2) | WP_148248753.1 | WP_148248753.1 tRNA glutamyl-Q(34) synthetase GluQRS |
| 55(1) | WP_213644233.1 | WP_213644233.1 tRNA glutamyl-Q(34) synthetase GluQRS |
| 55(1) | WP_113058721.1 | WP_113058721.1 tRNA glutamyl-Q(34) synthetase GluQRS |
| 55(1) | WP_076167855.1 | WP_076167855.1 tRNA glutamyl-Q(34) synthetase GluQRS |
| 55(1) | WP_080902257.1 | WP_080902257.1 tRNA glutamyl-Q(34) synthetase GluQRS |
| 55(1) | WP_043890952.1 | WP_043890952.1 tRNA glutamyl-Q(34) synthetase GluQRS |
| 55(1) | WP_183578470.1 | WP_183578470.1 tRNA glutamyl-Q(34) synthetase GluQRS |
| 55(1) | WP_251497184.1 | WP_251497184.1 tRNA glutamyl-Q(34) synthetase GluQRS |
| 55(2) | WP_036642168.1 | WP_036642168.1 tRNA glutamyl-Q(34) synthetase GluQRS |
| 55(1) | WP_210095134.1 | WP_210095134.1 tRNA glutamyl-Q(34) synthetase GluQRS |
| 55(1) | WP_280753899.1 | WP_280753899.1 tRNA glutamyl-Q(34) synthetase GluQRS |
| 55(1) | WP_246536780.1 | WP_246536780.1 tRNA glutamyl-Q(34) synthetase GluQRS |
| 55(1) | WP_212983258.1 | WP_212983258.1 tRNA glutamyl-Q(34) synthetase GluQRS |
| 55(1) | WP_076322108.1 | WP_076322108.1 tRNA glutamyl-Q(34) synthetase GluQRS |
| 55(1) | WP_100537970.1 | WP_100537970.1 tRNA glutamyl-Q(34) synthetase GluQRS |
| 55(1) | WP_090887118.1 | WP_090887118.1 tRNA glutamyl-Q(34) synthetase GluQRS |
| 55(1) | WP_277746354.1 | WP_277746354.1 tRNA glutamyl-Q(34) synthetase GluQRS |
| 55(1) | WP_028589290.1 | WP_028589290.1 tRNA glutamyl-Q(34) synthetase GluQRS |
| 55(1) | WP_077567012.1 | WP_077567012.1 tRNA glutamyl-Q(34) synthetase GluQRS |
| 55(1) | WP_116231171.1 | WP_116231171.1 tRNA glutamyl-Q(34) synthetase GluQRS |
| 55(1) | WP_142613009.1 | WP_142613009.1 tRNA glutamyl-Q(34) synthetase GluQRS |
| 55(1) | WP_212980227.1 | WP_212980227.1 tRNA glutamyl-Q(34) synthetase GluQRS |
| 55(1) | WP_213579156.1 | WP_213579156.1 tRNA glutamyl-Q(34) synthetase GluQRS |
| 55(1) | WP_194230539.1 | WP_194230539.1 tRNA glutamyl-Q(34) synthetase GluQRS |
| 55(1) | WP_305755027.1 | WP_305755027.1 tRNA glutamyl-Q(34) synthetase GluQRS |
| 55(1) | WP_160496924.1 | WP_160496924.1 tRNA glutamyl-Q(34) synthetase GluQRS |
| 55(1) | WP_080942654.1 | WP_080942654.1 tRNA glutamyl-Q(34) synthetase GluQRS |
| 55(1) | WP_063479990.1 | WP_063479990.1 tRNA glutamyl-Q(34) synthetase GluQRS |
| 55(1) | WP_192571323.1 | WP_192571323.1 tRNA glutamyl-Q(34) synthetase GluQRS |
| 55(1) | WP_009590630.1 | WP_009590630.1 tRNA glutamyl-Q(34) synthetase GluQRS |
| 55(1) | WP_220684644.1 | WP_220684644.1 tRNA glutamyl-Q(34) synthetase GluQRS |
| 55(2) | WP_040101525.1 | WP_040101525.1 tRNA glutamyl-Q(34) synthetase GluQRS |
| 55(1) | WP_215154572.1 | WP_215154572.1 tRNA glutamyl-Q(34) synthetase GluQRS |
| 55(1) | WP_094092627.1 | WP_094092627.1 tRNA glutamyl-Q(34) synthetase GluQRS |
| 55(1) | WP_213616457.1 | WP_213616457.1 tRNA glutamyl-Q(34) synthetase GluQRS |
| 55(1) | WP_003516596.1 | WP_003516596.1 glutamate--tRNA ligase |
| 55(1) | WP_212969034.1 | WP_212969034.1 tRNA glutamyl-Q(34) synthetase GluQRS |
| 55(1) | WP_279591712.1 | WP_279591712.1 tRNA glutamyl-Q(34) synthetase GluQRS |
| 55(1) | WP_018883266.1 | WP_018883266.1 tRNA glutamyl-Q(34) synthetase GluQRS |
| 55(1) | WP_290403371.1 | WP_290403371.1 tRNA glutamyl-Q(34) synthetase GluQRS |
| 55(1) | WP_249507421.1 | WP_249507421.1 tRNA glutamyl-Q(34) synthetase GluQRS |
| 55(1) | WP_096774450.1 | WP_096774450.1 tRNA glutamyl-Q(34) synthetase GluQRS |
| 55(1) | WP_251674655.1 | WP_251674655.1 tRNA glutamyl-Q(34) synthetase GluQRS |
| 55(1) | WP_223067432.1 | WP_223067432.1 tRNA glutamyl-Q(34) synthetase GluQRS |
| 55(2) | WP_107734204.1 | WP_107734204.1 tRNA glutamyl-Q(34) synthetase GluQRS |
| 55(1) | WP_023957737.1 | WP_023957737.1 tRNA glutamyl-Q(34) synthetase GluQRS |
| 55(1) | WP_194543494.1 | WP_194543494.1 tRNA glutamyl-Q(34) synthetase GluQRS |
| 55(1) | WP_094593880.1 | WP_094593880.1 tRNA glutamyl-Q(34) synthetase GluQRS |
| 55(1) | WP_251537232.1 | WP_251537232.1 tRNA glutamyl-Q(34) synthetase GluQRS |
| 55(1) | WP_145134813.1 | WP_145134813.1 tRNA glutamyl-Q(34) synthetase GluQRS |
| 55(1) | WP_145021729.1 | WP_145021729.1 tRNA glutamyl-Q(34) synthetase GluQRS |
| 55(1) | WP_305023327.1 | WP_305023327.1 tRNA glutamyl-Q(34) synthetase GluQRS |
| 55(1) | WP_071220210.1 | WP_071220210.1 tRNA glutamyl-Q(34) synthetase GluQRS |
| 55(1) | WP_216531936.1 | WP_216531936.1 tRNA glutamyl-Q(34) synthetase GluQRS |
| 55(1) | WP_080582305.1 | WP_080582305.1 tRNA glutamyl-Q(34) synthetase GluQRS |
| 55(1) | WP_127592422.1 | WP_127592422.1 tRNA glutamyl-Q(34) synthetase GluQRS |
| 55(1) | WP_152402384.1 | WP_152402384.1 tRNA glutamyl-Q(34) synthetase GluQRS |
| 55(1) | WP_233280992.1 | WP_233280992.1 tRNA glutamyl-Q(34) synthetase GluQRS |
| 55(1) | WP_095266273.1 | WP_095266273.1 tRNA glutamyl-Q(34) synthetase GluQRS |
| 55(1) | WP_260065265.1 | WP_260065265.1 tRNA glutamyl-Q(34) synthetase GluQRS |
| 55(2) | WP_119847556.1 | WP_119847556.1 tRNA glutamyl-Q(34) synthetase GluQRS |
| 55(1) | WP_145037680.1 | WP_145037680.1 tRNA glutamyl-Q(34) synthetase GluQRS |
| 55(1) | WP_265444516.1 | WP_265444516.1 glutamate--tRNA ligase |
| 55(1) | WP_236575325.1 | WP_236575325.1 tRNA glutamyl-Q(34) synthetase GluQRS |
| 55(1) | WP_054402902.1 | WP_054402902.1 tRNA glutamyl-Q(34) synthetase GluQRS |

| | | |
|---|---|---|
| 56(3) | WP_098088264.1 | WP_098088264.1 DUF393 domain-containing protein |
| 56(1) | WP_060630636.1 | WP_060630636.1 DCC1-like thiol-disulfide oxidoreductase family protein |
| 56(1) | WP_042874971.1 | WP_042874971.1 DCC1-like thiol-disulfide oxidoreductase family protein |
| 56(1) | WP_144488214.1 | WP_144488214.1 DCC1-like thiol-disulfide oxidoreductase family protein |
| 56(5) | WP_088063223.1 | WP_088063223.1 DUF393 domain-containing protein |
| 56(1) | WP_166704975.1 | WP_166704975.1 DUF393 domain-containing protein |

| | | | |
|---|---|---|---|
| 56(16) | WP_000634891.1 | WP_000634891.1 | DCC1-like thiol-disulfide oxidoreductase family protein |
| 56(1) | WP_087982608.1 | WP_087982608.1 | DUF393 domain-containing protein |
| 56(1) | WP_000634905.1 | WP_000634905.1 | DCC1-like thiol-disulfide oxidoreductase family protein |
| 56(1) | WP_000634871.1 | WP_000634871.1 | DCC1-like thiol-disulfide oxidoreductase family protein |
| 56(1) | WP_262748117.1 | WP_262748117.1 | DUF393 domain-containing protein |
| 56(1) | WP_207994175.1 | WP_207994175.1 | DUF393 domain-containing protein |
| 56(2) | WP_078419980.1 | WP_078419980.1 | DCC1-like thiol-disulfide oxidoreductase family protein |
| 56(1) | WP_000634865.1 | WP_000634865.1 | DCC1-like thiol-disulfide oxidoreductase family protein |
| 56(3) | WP_098296558.1 | WP_098296558.1 | DCC1-like thiol-disulfide oxidoreductase family protein |
| 56(1) | WP_098319877.1 | WP_098319877.1 | DCC1-like thiol-disulfide oxidoreductase family protein |
| 56(3) | WP_000634879.1 | WP_000634879.1 | DCC1-like thiol-disulfide oxidoreductase family protein |
| 56(2) | WP_017561866.1 | WP_017561866.1 | DCC1-like thiol-disulfide oxidoreductase family protein |
| 56(1) | WP_144564259.1 | WP_144564259.1 | DCC1-like thiol-disulfide oxidoreductase family protein |
| 56(1) | WP_097821806.1 | WP_097821806.1 | DUF393 domain-containing protein |
| 56(8) | WP_000634866.1 | WP_000634866.1 | DCC1-like thiol-disulfide oxidoreductase family protein |
| 56(1) | WP_098519055.1 | WP_098519055.1 | DCC1-like thiol-disulfide oxidoreductase family protein |
| 56(1) | WP_144611176.1 | WP_144611176.1 | DCC1-like thiol-disulfide oxidoreductase family protein |
| 56(3) | WP_000634875.1 | WP_000634875.1 | DCC1-like thiol-disulfide oxidoreductase family protein |
| 56(1) | WP_098952609.1 | WP_098952609.1 | DCC1-like thiol-disulfide oxidoreductase family protein |
| 56(4) | WP_000634876.1 | WP_000634876.1 | DCC1-like thiol-disulfide oxidoreductase family protein |
| 56(2) | WP_000634897.1 | WP_000634897.1 | DCC1-like thiol-disulfide oxidoreductase family protein |
| 56(1) | WP_098856759.1 | WP_098856759.1 | DUF393 domain-containing protein |
| 56(1) | WP_014893790.1 | WP_014893790.1 | DCC1-like thiol-disulfide oxidoreductase family protein |
| 56(1) | WP_262787107.1 | WP_262787107.1 | DUF393 domain-containing protein |
| 56(2) | WP_087952364.1 | WP_087952364.1 | DCC1-like thiol-disulfide oxidoreductase family protein |
| 56(2) | WP_098137211.1 | WP_098137211.1 | DUF393 domain-containing protein |
| 56(3) | WP_098264274.1 | WP_098264274.1 | DCC1-like thiol-disulfide oxidoreductase family protein |
| 56(1) | WP_199695244.1 | WP_199695244.1 | DUF393 domain-containing protein |
| 56(3) | WP_000634868.1 | WP_000634868.1 | DCC1-like thiol-disulfide oxidoreductase family protein |
| 56(1) | WP_103655001.1 | WP_103655001.1 | DCC1-like thiol-disulfide oxidoreductase family protein |
| 56(1) | WP_098198680.1 | WP_098198680.1 | DCC1-like thiol-disulfide oxidoreductase family protein |
| 56(1) | WP_061658093.1 | WP_061658093.1 | DCC1-like thiol-disulfide oxidoreductase family protein |
| 56(2) | WP_000634906.1 | WP_000634906.1 | DCC1-like thiol-disulfide oxidoreductase family protein |
| 56(1) | WP_097920544.1 | WP_097920544.1 | DCC1-like thiol-disulfide oxidoreductase family protein |
| 56(1) | WP_074570401.1 | WP_074570401.1 | DUF393 domain-containing protein |
| 57(3) | WP_098266639.1 | WP_098266639.1 | hypothetical protein |
| 57(2) | WP_063223267.1 | WP_063223267.1 | hypothetical protein |
| 57(3) | WP_078205518.1 | WP_078205518.1 | hypothetical protein |
| 57(2) | WP_000498527.1 | WP_000498527.1 | hypothetical protein |
| 57(3) | WP_033709025.1 | WP_033709025.1 | hypothetical protein |
| 57(3) | WP_080025011.1 | WP_080025011.1 | hypothetical protein |
| 57(2) | WP_016102108.1 | WP_016102108.1 | hypothetical protein |
| 57(1) | WP_248139694.1 | WP_248139694.1 | hypothetical protein |
| 57(2) | WP_002141579.1 | WP_002141579.1 | hypothetical protein |
| 57(2) | WP_098991581.1 | WP_098991581.1 | hypothetical protein |
| 57(1) | WP_264456843.1 | WP_264456843.1 | hypothetical protein |
| 57(2) | WP_002148354.1 | WP_002148354.1 | hypothetical protein |
| 57(2) | WP_000488505.1 | WP_000488505.1 | hypothetical protein |
| 57(5) | WP_062821094.1 | WP_062821094.1 | hypothetical protein |
| 57(1) | WP_242251756.1 | WP_242251756.1 | hypothetical protein |
| 57(7) | WP_000498607.1 | WP_000498607.1 | hypothetical protein |
| 57(14) | WP_000498526.1 | WP_000498526.1 | hypothetical protein |
| 57(1) | WP_098776439.1 | WP_098776439.1 | hypothetical protein |
| 57(1) | WP_002189452.1 | WP_002189452.1 | hypothetical protein |
| 57(18) | WP_000498599.1 | WP_000498599.1 | hypothetical protein |
| 57(1) | WP_000498600.1 | WP_000498600.1 | hypothetical protein |
| 57(1) | WP_088058516.1 | WP_088058516.1 | hypothetical protein |
| 57(1) | WP_002171713.1 | WP_002171713.1 | hypothetical protein |
| 57(2) | WP_000498596.1 | WP_000498596.1 | hypothetical protein |
| 57(1) | WP_087952368.1 | WP_087952368.1 | hypothetical protein |
| 57(1) | WP_277480669.1 | WP_277480669.1 | hypothetical protein |
| 57(1) | WP_270691240.1 | WP_270691240.1 | hypothetical protein |
| 58(14) | WP_000028911.1 | WP_000028911.1 | lipid II flippase Amj family protein |
| 58(1) | WP_097810727.1 | WP_097810727.1 | lipid II flippase Amj family protein |
| 58(1) | WP_000237160.1 | WP_000237160.1 | lipid II flippase Amj family protein |
| 58(1) | WP_270728538.1 | WP_270728538.1 | lipid II flippase Amj family protein |
| 58(1) | WP_070805907.1 | WP_070805907.1 | lipid II flippase Amj family protein |
| 58(8) | WP_087982567.1 | WP_087982567.1 | lipid II flippase Amj family protein |
| 58(1) | WP_026882386.1 | WP_026882386.1 | lipid II flippase Amj family protein |
| 58(4) | WP_048552323.1 | WP_048552323.1 | lipid II flippase Amj family protein |
| 58(1) | WP_106382858.1 | WP_106382858.1 | lipid II flippase Amj family protein |
| 58(2) | WP_071724735.1 | WP_071724735.1 | lipid II flippase Amj family protein |
| 58(1) | WP_061535951.1 | WP_061535951.1 | lipid II flippase Amj family protein |
| 58(2) | WP_098264293.1 | WP_098264293.1 | lipid II flippase Amj family protein |
| 58(1) | WP_215598782.1 | WP_215598782.1 | DUF2837 family protein |
| 58(2) | WP_000237162.1 | WP_000237162.1 | lipid II flippase Amj family protein |
| 58(1) | WP_087994850.1 | WP_087994850.1 | lipid II flippase Amj family protein |
| 58(2) | WP_139895590.1 | WP_139895590.1 | lipid II flippase Amj family protein |
| 58(1) | WP_144611199.1 | WP_144611199.1 | lipid II flippase Amj family protein |
| 58(1) | WP_098924913.1 | WP_098924913.1 | lipid II flippase Amj family protein |
| 58(4) | WP_087952387.1 | WP_087952387.1 | lipid II flippase Amj family protein |
| 58(1) | WP_060630655.1 | WP_060630655.1 | lipid II flippase Amj family protein |
| 58(1) | WP_000028921.1 | WP_000028921.1 | lipid II flippase Amj family protein |
| 58(1) | WP_071719973.1 | WP_071719973.1 | lipid II flippase Amj family protein |
| 58(1) | WP_097852384.1 | WP_097852384.1 | lipid II flippase Amj family protein |
| 58(1) | WP_197241463.1 | WP_197241463.1 | lipid II flippase Amj family protein |
| 58(1) | WP_071758132.1 | WP_071758132.1 | lipid II flippase Amj family protein |
| 58(1) | WP_071745224.1 | WP_071745224.1 | lipid II flippase Amj family protein |
| 58(4) | WP_002158032.1 | WP_002158032.1 | lipid II flippase Amj family protein |
| 58(1) | WP_098299073.1 | WP_098299073.1 | lipid II flippase Amj family protein |
| 58(3) | WP_072938300.1 | WP_072938300.1 | lipid II flippase Amj family protein |
| 58(1) | WP_098365945.1 | WP_098365945.1 | lipid II flippase Amj family protein |
| 58(1) | WP_002189473.1 | WP_002189473.1 | DUF2837 family protein |
| 58(1) | WP_098266658.1 | WP_098266658.1 | lipid II flippase Amj family protein |
| 58(1) | WP_223067443.1 | WP_223067443.1 | lipid II flippase Amj family protein |
| 58(2) | WP_087956301.1 | WP_087956301.1 | lipid II flippase Amj family protein |
| 58(2) | WP_088609120.1 | WP_088609120.1 | DUF2837 family protein |
| 58(1) | WP_098199516.1 | WP_098199516.1 | DUF2837 family protein |
| 58(1) | WP_151630591.1 | WP_151630591.1 | lipid II flippase Amj family protein |
| 58(7) | WP_000028910.1 | WP_000028910.1 | lipid II flippase Amj family protein |
| 58(1) | WP_002141610.1 | WP_002141610.1 | lipid II flippase Amj family protein |

| | | |
|---|---|---|
| 59(2) | WP_086406043.1 | WP_086406043.1 DUF3952 domain-containing protein |
| 59(2) | WP_174402543.1 | WP_174402543.1 DUF3952 domain-containing protein |
| 59(1) | WP_000768485.1 | WP_000768485.1 DUF3952 domain-containing protein |
| 59(1) | WP_098139249.1 | WP_098139249.1 DUF3952 domain-containing protein |
| 59(1) | WP_117616460.1 | WP_117616460.1 DUF3952 domain-containing protein |
| 59(1) | WP_116891186.1 | WP_116891186.1 hypothetical protein |
| 59(1) | WP_000628346.1 | WP_000628346.1 DUF3952 domain-containing protein |
| 59(1) | WP_000623464.1 | WP_000623464.1 DUF3952 domain-containing protein |
| 59(1) | WP_030025442.1 | WP_030025442.1 DUF3952 domain-containing protein |
| 59(1) | WP_006923850.1 | WP_006923850.1 DUF3952 domain-containing protein |
| 59(1) | WP_142326098.1 | WP_142326098.1 DUF3952 domain-containing protein |
| 59(1) | WP_306307679.1 | WP_306307679.1 hypothetical protein |
| 59(1) | WP_000044430.1 | WP_000044430.1 DUF3952 domain-containing protein |
| 59(4) | WP_001990247.1 | WP_001990247.1 DUF3952 domain-containing protein |
| 59(1) | WP_175879505.1 | WP_175879505.1 DUF3952 domain-containing protein |
| 59(1) | WP_098291287.1 | WP_098291287.1 DUF3952 domain-containing protein |
| 59(1) | WP_098529291.1 | WP_098529291.1 DUF3952 domain-containing protein |
| 59(3) | WP_000768487.1 | WP_000768487.1 DUF3952 domain-containing protein |
| 59(1) | WP_000628357.1 | WP_000628357.1 DUF3952 domain-containing protein |
| 59(3) | WP_044584393.1 | WP_044584393.1 DUF3952 domain-containing protein |
| 59(2) | WP_029442153.1 | WP_029442153.1 DUF3952 domain-containing protein |
| 59(1) | WP_063675265.1 | WP_063675265.1 DUF3952 domain-containing protein |
| 59(1) | WP_046946379.1 | WP_046946379.1 DUF3952 domain-containing protein |
| 59(1) | WP_000768465.1 | WP_000768465.1 DUF3952 domain-containing protein |
| 59(1) | WP_016117683.1 | WP_016117683.1 DUF3952 domain-containing protein |
| 59(1) | WP_097796438.1 | WP_097796438.1 DUF3952 domain-containing protein |
| 59(1) | WP_208906819.1 | WP_208906819.1 DUF3952 domain-containing protein |
| 59(1) | WP_142312406.1 | WP_142312406.1 DUF3952 domain-containing protein |
| 59(1) | WP_086412056.1 | WP_086412056.1 DUF3952 domain-containing protein |
| 59(1) | WP_141526900.1 | WP_141526900.1 DUF3952 domain-containing protein |
| 59(1) | WP_000044428.1 | WP_000044428.1 DUF3952 domain-containing protein |
| 59(1) | WP_087967959.1 | WP_087967959.1 DUF3952 domain-containing protein |
| 59(3) | WP_097954749.1 | WP_097954749.1 DUF3952 domain-containing protein |
| 59(1) | WP_074643593.1 | WP_074643593.1 DUF3952 domain-containing protein |
| 59(1) | WP_197323875.1 | WP_197323875.1 DUF3952 domain-containing protein |
| 59(2) | WP_119683232.1 | WP_119683232.1 DUF3952 domain-containing protein |
| 59(1) | WP_063675264.1 | WP_063675264.1 DUF3952 domain-containing protein |
| 59(4) | WP_030025444.1 | WP_030025444.1 DUF3952 domain-containing protein |
| 59(3) | WP_179204796.1 | WP_179204796.1 DUF3952 domain-containing protein |
| 59(1) | WP_153589096.1 | WP_153589096.1 DUF3952 domain-containing protein |
| 59(1) | WP_141557691.1 | WP_141557691.1 DUF3952 domain-containing protein |
| 59(1) | WP_033693851.1 | WP_033693851.1 DUF3952 domain-containing protein |
| 59(1) | WP_000768466.1 | WP_000768466.1 DUF3952 domain-containing protein |
| 59(2) | WP_098316708.1 | WP_098316708.1 DUF3952 domain-containing protein |
| 59(1) | WP_204319151.1 | WP_204319151.1 DUF3952 domain-containing protein |
| 59(3) | WP_000768482.1 | WP_000768482.1 DUF3952 domain-containing protein |
| 59(2) | WP_000628355.1 | WP_000628355.1 DUF3952 domain-containing protein |
| 59(1) | WP_098535854.1 | WP_098535854.1 DUF3952 domain-containing protein |
| 59(4) | WP_001990246.1 | WP_001990246.1 DUF3952 domain-containing protein |
| 59(1) | WP_016117677.1 | WP_016117677.1 DUF3952 domain-containing protein |
| 59(1) | WP_098819905.1 | WP_098819905.1 DUF3952 domain-containing protein |
| 59(1) | WP_098013428.1 | WP_098013428.1 DUF3952 domain-containing protein |
| 59(1) | WP_098516212.1 | WP_098516212.1 DUF3952 domain-containing protein |
| 59(2) | WP_061529909.1 | WP_061529909.1 DUF3952 domain-containing protein |
| 59(1) | WP_102956431.1 | WP_102956431.1 DUF3952 domain-containing protein |
| 60(1) | WP_016104717.1 | WP_016104717.1 heparan-alpha-glucosaminide N-acetyltransferase domain-containing protein |
| 60(1) | WP_242251762.1 | WP_242251762.1 heparan-alpha-glucosaminide N-acetyltransferase domain-containing protein |
| 60(2) | WP_141534299.1 | WP_141534299.1 heparan-alpha-glucosaminide N-acetyltransferase domain-containing protein |
| 60(1) | WP_142327656.1 | WP_142327656.1 heparan-alpha-glucosaminide N-acetyltransferase domain-containing protein |
| 60(2) | WP_141506911.1 | WP_141506911.1 heparan-alpha-glucosaminide N-acetyltransferase domain-containing protein |
| 60(1) | WP_141544659.1 | WP_141544659.1 heparan-alpha-glucosaminide N-acetyltransferase domain-containing protein |
| 60(1) | WP_087945119.1 | WP_087945119.1 heparan-alpha-glucosaminide N-acetyltransferase domain-containing protein |
| 60(3) | WP_002189456.1 | WP_002189456.1 heparan-alpha-glucosaminide N-acetyltransferase domain-containing protein |
| 60(1) | WP_145959664.1 | WP_145959664.1 heparan-alpha-glucosaminide N-acetyltransferase domain-containing protein |
| 60(5) | WP_141505484.1 | WP_141505484.1 heparan-alpha-glucosaminide N-acetyltransferase domain-containing protein |
| 60(1) | WP_141533136.1 | WP_141533136.1 heparan-alpha-glucosaminide N-acetyltransferase domain-containing protein |
| 60(2) | WP_131231113.1 | WP_131231113.1 heparan-alpha-glucosaminide N-acetyltransferase domain-containing protein |
| 60(1) | WP_002031634.1 | WP_002031634.1 heparan-alpha-glucosaminide N-acetyltransferase domain-containing protein |
| 60(1) | WP_080019206.1 | WP_080019206.1 DUF418 domain-containing protein |
| 60(1) | WP_002158053.1 | WP_002158053.1 heparan-alpha-glucosaminide N-acetyltransferase domain-containing protein |
| 60(4) | WP_053564636.1 | WP_053564636.1 heparan-alpha-glucosaminide N-acetyltransferase domain-containing protein |
| 60(1) | WP_002141585.1 | WP_002141585.1 heparan-alpha-glucosaminide N-acetyltransferase domain-containing protein |
| 60(1) | WP_141548208.1 | WP_141548208.1 heparan-alpha-glucosaminide N-acetyltransferase domain-containing protein |
| 60(2) | WP_197241533.1 | WP_197241533.1 heparan-alpha-glucosaminide N-acetyltransferase domain-containing protein |
| 60(2) | WP_063219644.1 | WP_063219644.1 heparan-alpha-glucosaminide N-acetyltransferase domain-containing protein |
| 60(1) | WP_142327337.1 | WP_142327337.1 heparan-alpha-glucosaminide N-acetyltransferase domain-containing protein |
| 60(1) | WP_001093110.1 | WP_001093110.1 heparan-alpha-glucosaminide N-acetyltransferase domain-containing protein |
| 60(1) | WP_071719989.1 | WP_071719989.1 heparan-alpha-glucosaminide N-acetyltransferase domain-containing protein |
| 60(1) | WP_098776436.1 | WP_098776436.1 heparan-alpha-glucosaminide N-acetyltransferase domain-containing protein |

60(1) WP_033658252.1 WP_033658252.1 heparan-alpha-glucosaminide N-acetyltransferase domain-containing protein
60(1) WP_140152374.1 WP_140152374.1 heparan-alpha-glucosaminide N-acetyltransferase domain-containing protein
60(1) WP_212956545.1 WP_212956545.1 heparan-alpha-glucosaminide N-acetyltransferase domain-containing protein
60(1) WP_286017541.1 WP_286017541.1 heparan-alpha-glucosaminide N-acetyltransferase domain-containing protein
60(1) WP_142306906.1 WP_142306906.1 heparan-alpha-glucosaminide N-acetyltransferase domain-containing protein
60(2) WP_142669931.1 WP_142669931.1 heparan-alpha-glucosaminide N-acetyltransferase domain-containing protein
60(2) WP_139213220.1 WP_139213220.1 heparan-alpha-glucosaminide N-acetyltransferase domain-containing protein
60(4) WP_001093112.1 WP_001093112.1 heparan-alpha-glucosaminide N-acetyltransferase domain-containing protein
60(1) WP_141436471.1 WP_141436471.1 heparan-alpha-glucosaminide N-acetyltransferase domain-containing protein
60(3) WP_001093111.1 WP_001093111.1 heparan-alpha-glucosaminide N-acetyltransferase domain-containing protein
60(1) WP_144476569.1 WP_144476569.1 heparan-alpha-glucosaminide N-acetyltransferase domain-containing protein
60(1) WP_270728503.1 WP_270728503.1 heparan-alpha-glucosaminide N-acetyltransferase domain-containing protein
60(1) WP_141531121.1 WP_141531121.1 heparan-alpha-glucosaminide N-acetyltransferase domain-containing protein, partial
60(1) WP_142306378.1 WP_142306378.1 heparan-alpha-glucosaminide N-acetyltransferase domain-containing protein
60(1) WP_142254722.1 WP_142254722.1 heparan-alpha-glucosaminide N-acetyltransferase domain-containing protein
60(1) WP_286124776.1 WP_286124776.1 heparan-alpha-glucosaminide N-acetyltransferase domain-containing protein
60(1) WP_142318158.1 WP_142318158.1 heparan-alpha-glucosaminide N-acetyltransferase domain-containing protein
60(1) WP_262725732.1 WP_262725732.1 DUF1624 domain-containing protein
60(1) WP_023521809.1 WP_023521809.1 heparan-alpha-glucosaminide N-acetyltransferase domain-containing protein
60(2) WP_071724712.1 WP_071724712.1 heparan-alpha-glucosaminide N-acetyltransferase domain-containing protein
60(3) WP_065228766.1 WP_065228766.1 heparan-alpha-glucosaminide N-acetyltransferase domain-containing protein
60(1) WP_142951152.1 WP_142951152.1 heparan-alpha-glucosaminide N-acetyltransferase domain-containing protein
60(1) WP_141552652.1 WP_141552652.1 heparan-alpha-glucosaminide N-acetyltransferase domain-containing protein
60(1) WP_088058523.1 WP_088058523.1 heparan-alpha-glucosaminide N-acetyltransferase domain-containing protein
60(2) WP_200457881.1 WP_200457881.1 heparan-alpha-glucosaminide N-acetyltransferase domain-containing protein
60(1) WP_141241078.1 WP_141241078.1 heparan-alpha-glucosaminide N-acetyltransferase domain-containing protein
60(4) WP_141538301.1 WP_141538301.1 heparan-alpha-glucosaminide N-acetyltransferase domain-containing protein

61(2) WP_194330191.1 WP_194330191.1 accessory gene regulator ArgB-like protein
61(58) WP_003750466.1 WP_003750466.1 accessory gene regulator ArgB-like protein
61(1) WP_021624813.1 WP_021624813.1 accessory gene regulator B family protein
61(2) WP_003728981.1 WP_003728981.1 accessory gene regulator ArgB-like protein
61(2) WP_011700914.1 WP_011700914.1 accessory gene regulator ArgB-like protein
61(1) WP_168976292.1 WP_168976292.1 accessory gene regulator B family protein
61(2) WP_185516175.1 WP_185516175.1 accessory gene regulator ArgB-like protein
61(2) WP_185639025.1 WP_185639025.1 accessory gene regulator ArgB-like protein
61(1) WP_276917299.1 WP_276917299.1 accessory gene regulator B family protein
61(4) WP_038406368.1 WP_038406368.1 accessory gene regulator ArgB-like protein
61(2) WP_200372573.1 WP_200372573.1 accessory gene regulator ArgB-like protein
61(1) WP_052812187.1 WP_052812187.1 accessory gene regulator B family protein

62(2) WP_071732575.1 WP_071732575.1 DUF5082 family protein
62(2) WP_006923816.1 WP_006923816.1 DUF5082 family protein
62(1) WP_106496022.1 WP_106496022.1 DUF5082 family protein
62(38) WP_000510565.1 WP_000510565.1 DUF5082 family protein
62(1) WP_088230464.1 WP_088230464.1 DUF5082 family protein
62(1) WP_289613140.1 WP_289613140.1 DUF5082 family protein
62(1) WP_053512323.1 WP_053512323.1 DUF5082 family protein
62(1) WP_077320205.1 WP_077320205.1 DUF5082 family protein
62(1) WP_000510564.1 WP_000510564.1 DUF5082 family protein
62(2) WP_016108254.1 WP_016108254.1 DUF5082 family protein
62(1) WP_277150359.1 WP_277150359.1 hypothetical protein
62(1) WP_270472020.1 WP_270472020.1 DUF5082 family protein
62(1) WP_046946380.1 WP_046946380.1 DUF5082 family protein
62(1) WP_130810253.1 WP_130810253.1 DUF5082 family protein
62(1) WP_098142229.1 WP_098142229.1 DUF5082 family protein
62(3) WP_098139261.1 WP_098139261.1 DUF5082 family protein
62(1) WP_102371679.1 WP_102371679.1 DUF5082 family protein
62(1) WP_098645175.1 WP_098645175.1 DUF5082 family protein
62(1) WP_277166574.1 WP_277166574.1 DUF5082 family protein
62(1) WP_277142058.1 WP_277142058.1 DUF5082 family protein
62(1) WP_102956434.1 WP_102956434.1 DUF5082 family protein
62(5) WP_097837316.1 WP_097837316.1 DUF5082 family protein
62(4) WP_001990256.1 WP_001990256.1 DUF5082 family protein
62(1) WP_163095413.1 WP_163095413.1 DUF5082 family protein
62(1) WP_289613137.1 WP_289613137.1 DUF5082 family protein
62(1) WP_086385920.1 WP_086385920.1 DUF5082 family protein
62(1) WP_098612031.1 WP_098612031.1 DUF5082 family protein
62(1) WP_061667207.1 WP_061667207.1 DUF5082 family protein
62(1) WP_022349190.1 WP_022349190.1 DUF5082 family protein
63(1) WP_001181853.1 WP_001181853.1 GNAT family protein
63(1) WP_022586776.1 WP_022586776.1 GNAT family protein
63(1) WP_098300892.1 WP_098300892.1 GNAT family protein
63(1) WP_087948421.1 WP_087948421.1 GNAT family protein
63(1) WP_232699373.1 WP_232699373.1 GNAT family protein

| | | |
|---|---|---|
| 63(1) | WP_064019327.1 | WP_064019327.1 GNAT family protein |
| 63(1) | WP_166244068.1 | WP_166244068.1 GNAT family protein |
| 63(1) | WP_301804641.1 | WP_301804641.1 GNAT family protein |
| 63(1) | WP_158329531.1 | WP_158329531.1 GNAT family protein |
| 63(2) | WP_113755285.1 | WP_113755285.1 GNAT family protein |
| 63(2) | WP_086405913.1 | WP_086405913.1 GNAT family protein |
| 63(1) | WP_272898223.1 | WP_272898223.1 hypothetical protein |
| 63(1) | WP_061676407.1 | WP_061676407.1 GNAT family protein |
| 63(3) | WP_097954760.1 | WP_097954760.1 GNAT family protein |
| 63(1) | WP_063536163.1 | WP_063536163.1 GNAT family protein |
| 63(1) | WP_001181860.1 | WP_001181860.1 GNAT family protein |
| 63(2) | WP_001089593.1 | WP_001089593.1 GNAT family protein |
| 63(8) | WP_001181857.1 | WP_001181857.1 GNAT family protein |
| 63(1) | WP_017763061.1 | WP_017763061.1 GNAT family protein |
| 63(2) | WP_001181781.1 | WP_001181781.1 GNAT family protein |
| 63(1) | WP_286119522.1 | WP_286119522.1 GNAT family protein |
| 63(1) | WP_030025427.1 | WP_030025427.1 GNAT family protein |
| 63(1) | WP_153589151.1 | WP_153589151.1 GNAT family protein |
| 63(1) | WP_158330729.1 | WP_158330729.1 GNAT family protein |
| 63(1) | WP_277548450.1 | WP_277548450.1 GNAT family protein |
| 63(1) | WP_001181852.1 | WP_001181852.1 GNAT family protein |
| 63(2) | WP_212930969.1 | WP_212930969.1 GNAT family protein |
| 63(1) | WP_084064600.1 | WP_084064600.1 GNAT family protein |
| 63(1) | WP_234121618.1 | WP_234121618.1 GNAT family N-acetyltransferase |
| 63(1) | WP_001181856.1 | WP_001181856.1 GNAT family protein |
| 63(1) | WP_239586178.1 | WP_239586178.1 GNAT family protein |
| 63(4) | WP_001181855.1 | WP_001181855.1 GNAT family protein |
| 63(1) | WP_097861167.1 | WP_097861167.1 GNAT family protein |
| 63(4) | WP_001181784.1 | WP_001181784.1 GNAT family protein |
| 63(1) | WP_081621590.1 | WP_081621590.1 GNAT family protein |
| 63(1) | WP_061667214.1 | WP_061667214.1 GNAT family protein |
| 63(3) | WP_001990268.1 | WP_001990268.1 GNAT family protein |
| 63(1) | WP_098607333.1 | WP_098607333.1 GNAT family protein |
| 63(2) | WP_242066307.1 | WP_242066307.1 hypothetical protein |
| 63(1) | WP_068793926.1 | WP_068793926.1 GNAT family protein |
| 63(2) | WP_242068644.1 | WP_242068644.1 GNAT family protein |
| 63(1) | WP_096886732.1 | WP_096886732.1 GNAT family protein |
| 63(1) | WP_016117663.1 | WP_016117663.1 GNAT family protein |
| 63(1) | WP_097998246.1 | WP_097998246.1 GNAT family protein |
| 63(1) | WP_150457778.1 | WP_150457778.1 GNAT family protein |
| 63(1) | WP_236645789.1 | WP_236645789.1 hypothetical protein |
| 63(7) | WP_003336010.1 | WP_003336010.1 GNAT family protein |
| 64(74) | WP_003721669.1 | WP_003721669.1 30S ribosomal protein S18 |
| 64(1) | WP_075876819.1 | WP_075876819.1 30S ribosomal protein S18 |
| 65(1) | WP_280753901.1 | WP_280753901.1 DUF1992 domain-containing protein |
| 65(1) | WP_091155391.1 | WP_091155391.1 DUF1992 domain-containing protein |
| 65(1) | WP_213644232.1 | WP_213644232.1 DUF1992 domain-containing protein |
| 65(1) | WP_029514392.1 | WP_029514392.1 DUF1992 domain-containing protein |
| 65(1) | WP_071220209.1 | WP_071220209.1 DUF1992 domain-containing protein |
| 65(1) | WP_100537971.1 | WP_100537971.1 DUF1992 domain-containing protein |
| 65(1) | WP_063479991.1 | WP_063479991.1 DUF1992 domain-containing protein |
| 65(1) | WP_260065264.1 | WP_260065264.1 DUF1992 domain-containing protein |
| 65(1) | WP_251375544.1 | WP_251375544.1 DUF1992 domain-containing protein |
| 65(1) | WP_127592423.1 | WP_127592423.1 DUF1992 domain-containing protein |
| 65(1) | WP_305023328.1 | WP_305023328.1 DUF1992 domain-containing protein |
| 65(1) | WP_208919050.1 | WP_208919050.1 DUF1992 domain-containing protein |
| 65(2) | WP_194230397.1 | WP_194230397.1 DUF1992 domain-containing protein |
| 65(1) | WP_076322109.1 | WP_076322109.1 DUF1992 domain-containing protein |
| 65(1) | WP_098746855.1 | WP_098746855.1 DUF1992 domain-containing protein |
| 65(2) | WP_040101526.1 | WP_040101526.1 DnaJ family domain-containing protein |
| 65(1) | WP_106765951.1 | WP_106765951.1 DUF1992 domain-containing protein |
| 65(1) | WP_007430140.1 | WP_007430140.1 DUF1992 domain-containing protein |
| 65(2) | WP_095266203.1 | WP_095266203.1 DUF1992 domain-containing protein |
| 65(1) | WP_054958415.1 | WP_054958415.1 DUF1992 domain-containing protein |
| 65(1) | WP_136604821.1 | WP_136604821.1 DUF1992 domain-containing protein |
| 65(1) | WP_194543495.1 | WP_194543495.1 DUF1992 domain-containing protein |
| 65(1) | WP_216531935.1 | WP_216531935.1 DUF1992 domain-containing protein |
| 65(1) | WP_244719657.1 | WP_244719657.1 DUF1992 domain-containing protein |
| 65(2) | WP_076312502.1 | WP_076312502.1 DUF1992 domain-containing protein |
| 65(1) | WP_251583300.1 | WP_251583300.1 DUF1992 domain-containing protein |
| 65(1) | WP_160037492.1 | WP_160037492.1 DUF1992 domain-containing protein |
| 65(2) | WP_007128622.1 | WP_007128622.1 DUF1992 domain-containing protein |
| 65(1) | WP_108465356.1 | WP_108465356.1 DUF1992 domain-containing protein |
| 65(1) | WP_251537231.1 | WP_251537231.1 DUF1992 domain-containing protein |
| 65(1) | WP_090908283.1 | WP_090908283.1 DUF1992 domain-containing protein |
| 65(1) | WP_054402903.1 | WP_054402903.1 DUF1992 domain-containing protein |
| 65(2) | WP_068619100.1 | WP_068619100.1 DnaJ family domain-containing protein |
| 65(1) | WP_241929667.1 | WP_241929667.1 DUF1992 domain-containing protein |
| 65(1) | WP_160496925.1 | WP_160496925.1 DUF1992 domain-containing protein |
| 65(1) | WP_123797117.1 | WP_123797117.1 DUF1992 domain-containing protein |
| 65(1) | WP_213649965.1 | WP_213649965.1 DUF1992 domain-containing protein |
| 65(2) | WP_148246952.1 | WP_148246952.1 DUF1992 domain-containing protein |
| 65(1) | WP_127199777.1 | WP_127199777.1 DUF1992 domain-containing protein |
| 65(2) | WP_036642167.1 | WP_036642167.1 DUF1992 domain-containing protein |
| 65(1) | WP_215154570.1 | WP_215154570.1 DUF1992 domain-containing protein |
| 65(1) | WP_220684643.1 | WP_220684643.1 DUF1992 domain-containing protein |
| 65(10) | WP_009590658.1 | WP_009590658.1 DUF1992 domain-containing protein |
| 65(1) | WP_142613008.1 | WP_142613008.1 DUF1992 domain-containing protein |
| 65(1) | WP_036707121.1 | WP_036707121.1 DUF1992 domain-containing protein |
| 65(1) | WP_077567013.1 | WP_077567013.1 DnaJ family domain-containing protein |
| 65(8) | WP_025364688.1 | WP_025364688.1 DUF1992 domain-containing protein |
| 65(1) | WP_285748186.1 | WP_285748186.1 DUF1992 domain-containing protein |
| 65(1) | WP_095398795.1 | WP_095398795.1 DUF1992 domain-containing protein |
| 65(2) | WP_047909608.1 | WP_047909608.1 DUF1992 domain-containing protein |
| 66(1) | WP_076311053.1 | WP_076311053.1 extracellular solute-binding protein |
| 66(1) | WP_090674357.1 | WP_090674357.1 extracellular solute-binding protein |
| 66(2) | WP_047909617.1 | WP_047909617.1 extracellular solute-binding protein |
| 66(1) | WP_251596658.1 | WP_251596658.1 ABC transporter substrate-binding protein |
| 66(1) | WP_179603468.1 | WP_179603468.1 ABC transporter substrate-binding protein |
| 66(1) | WP_281937356.1 | WP_281937356.1 extracellular solute-binding protein |
| 66(1) | WP_189031016.1 | WP_189031016.1 ABC transporter substrate-binding protein |

| | | |
|---|---|---|
| 66(1) | WP_034675342.1 | WP_034675342.1 ABC transporter substrate-binding protein |
| 66(1) | WP_206094038.1 | WP_206094038.1 extracellular solute-binding protein |
| 66(1) | WP_042210991.1 | WP_042210991.1 extracellular solute-binding protein |
| 66(2) | WP_219872448.1 | WP_219872448.1 extracellular solute-binding protein |
| 66(1) | WP_054972099.1 | WP_054972099.1 extracellular solute-binding protein |
| 66(1) | WP_036689455.1 | WP_036689455.1 extracellular solute-binding protein |
| 66(1) | WP_088830544.1 | WP_088830544.1 extracellular solute-binding protein |
| 66(1) | WP_077619665.1 | WP_077619665.1 extracellular solute-binding protein |
| 66(1) | WP_204412737.1 | WP_204412737.1 extracellular solute-binding protein |
| 66(1) | WP_241840574.1 | WP_241840574.1 ABC transporter substrate-binding protein |
| 66(1) | WP_251598138.1 | WP_251598138.1 extracellular solute-binding protein |
| 66(1) | WP_244723735.1 | WP_244723735.1 extracellular solute-binding protein |
| 66(1) | WP_247083064.1 | WP_247083064.1 ABC transporter substrate-binding protein |
| 66(2) | WP_013914178.1 | WP_013914178.1 ABC transporter substrate-binding protein |
| 66(1) | WP_307312578.1 | WP_307312578.1 ABC transporter substrate-binding protein |
| 66(1) | WP_019424153.1 | WP_019424153.1 hypothetical protein |
| 66(1) | WP_289483771.1 | WP_289483771.1 ABC transporter substrate-binding protein |
| 66(2) | WP_127484337.1 | WP_127484337.1 extracellular solute-binding protein |
| 66(1) | WP_010496618.1 | WP_010496618.1 extracellular solute-binding protein |
| 66(1) | WP_165097590.1 | WP_165097590.1 extracellular solute-binding protein |
| 66(1) | WP_090722904.1 | WP_090722904.1 extracellular solute-binding protein |
| 66(1) | WP_244520548.1 | WP_244520548.1 ABC transporter substrate-binding protein |
| 66(1) | WP_166244054.1 | WP_166244054.1 extracellular solute-binding protein |
| 66(1) | WP_249897326.1 | WP_249897326.1 ABC transporter substrate-binding protein |
| 66(1) | WP_128631295.1 | WP_128631295.1 ABC transporter substrate-binding protein |
| 66(1) | WP_007430145.1 | WP_007430145.1 sugar ABC transporter substrate-binding protein |
| 66(1) | WP_301167486.1 | WP_301167486.1 ABC transporter substrate-binding protein |
| 66(1) | WP_068694709.1 | WP_068694709.1 extracellular solute-binding protein |
| 66(1) | WP_306077378.1 | WP_306077378.1 ABC transporter substrate-binding protein |
| 66(1) | WP_307192904.1 | WP_307192904.1 ABC transporter substrate-binding protein |
| 66(1) | WP_289475080.1 | WP_289475080.1 ABC transporter substrate-binding protein |
| 66(1) | WP_309098596.1 | WP_309098596.1 ABC transporter substrate-binding protein |
| 66(1) | WP_229139907.1 | WP_229139907.1 sugar ABC transporter substrate-binding protein |
| 66(1) | WP_121610579.1 | WP_121610579.1 ABC transporter substrate-binding protein |
| 66(1) | WP_199613917.1 | WP_199613917.1 extracellular solute-binding protein |
| 66(1) | WP_245208024.1 | WP_245208024.1 ABC transporter substrate-binding protein |
| 66(1) | WP_179160634.1 | WP_179160634.1 ABC transporter substrate-binding protein |
| 66(1) | WP_025848626.1 | WP_025848626.1 extracellular solute-binding protein |
| 66(2) | WP_091155368.1 | WP_091155368.1 extracellular solute-binding protein |
| 66(1) | WP_089522624.1 | WP_089522624.1 extracellular solute-binding protein |
| 66(1) | WP_061975555.1 | WP_061975555.1 ABC transporter substrate-binding protein |
| 66(1) | WP_244527438.1 | WP_244527438.1 extracellular solute-binding protein |
| 66(1) | WP_253411092.1 | WP_253411092.1 extracellular solute-binding protein |
| 66(1) | WP_089965960.1 | WP_089965960.1 ABC transporter substrate-binding protein |
| 66(1) | WP_233697168.1 | WP_233697168.1 hypothetical protein |
| 66(2) | WP_087433599.1 | WP_087433599.1 extracellular solute-binding protein |
| 66(1) | WP_132008127.1 | WP_132008127.1 ABC transporter substrate-binding protein |
| 66(1) | WP_076113448.1 | WP_076113448.1 extracellular solute-binding protein |
| 66(1) | WP_216791238.1 | WP_216791238.1 extracellular solute-binding protein |
| 66(1) | WP_099517110.1 | WP_099517110.1 extracellular solute-binding protein |
| 66(1) | WP_014368234.1 | WP_014368234.1 ABC transporter substrate-binding protein |
| 66(1) | WP_116353265.1 | WP_116353265.1 sugar ABC transporter substrate-binding protein |

| | | |
|---|---|---|
| 66(1) | WP_094092615.1 | WP_094092615.1 extracellular solute-binding protein |
| 66(1) | WP_199613891.1 | WP_199613891.1 sugar ABC transporter substrate-binding protein |
| 66(2) | WP_204412706.1 | WP_204412706.1 ABC transporter substrate-binding protein |
| 66(1) | WP_127459444.1 | WP_127459444.1 extracellular solute-binding protein |
| 66(2) | WP_108534965.1 | WP_108534965.1 extracellular solute-binding protein |
| 66(1) | WP_136379730.1 | WP_136379730.1 extracellular solute-binding protein |
| 66(1) | WP_074598766.1 | WP_074598766.1 ABC transporter substrate-binding protein |
| 66(1) | WP_063188041.1 | WP_063188041.1 extracellular solute-binding protein |
| 67(43) | WP_012984596.1 | WP_012984596.1 50S ribosomal protein L9 |
| 67(6) | WP_038406372.1 | WP_038406372.1 50S ribosomal protein L9 |
| 67(2) | WP_185469297.1 | WP_185469297.1 50S ribosomal protein L9 |
| 67(16) | WP_185340293.1 | WP_185340293.1 50S ribosomal protein L9 |
| 67(2) | WP_003721676.1 | WP_003721676.1 50S ribosomal protein L9 |
| 67(2) | WP_011700919.1 | WP_011700919.1 50S ribosomal protein L9 |
| 67(1) | WP_185451818.1 | WP_185451818.1 50S ribosomal protein L9 |
| 67(2) | WP_185391721.1 | WP_185391721.1 50S ribosomal protein L9 |
| 68(62) | WP_003745028.1 | WP_003745028.1 cyclic lactone autoinducer peptide |
| 68(6) | WP_003718129.1 | WP_003718129.1 cyclic lactone autoinducer peptide |
| 68(2) | WP_185532432.1 | WP_185532432.1 cyclic lactone autoinducer peptide |
| 68(2) | WP_003721672.1 | WP_003721672.1 cyclic lactone autoinducer peptide |
| 68(2) | WP_011700915.1 | WP_011700915.1 cyclic lactone autoinducer peptide |
| 69(6) | WP_038408416.1 | WP_038408416.1 adenylosuccinate synthase |
| 69(2) | WP_185415312.1 | WP_185415312.1 adenylosuccinate synthase |
| 69(14) | WP_003745033.1 | WP_003745033.1 adenylosuccinate synthase |
| 69(2) | WP_003728977.1 | WP_003728977.1 adenylosuccinate synthase |
| 69(2) | WP_011700920.1 | WP_011700920.1 adenylosuccinate synthase |
| 69(2) | WP_185639021.1 | WP_185639021.1 adenylosuccinate synthase |
| 69(46) | WP_185336683.1 | WP_185336683.1 adenylosuccinate synthase |
| 70(2) | WP_185639026.1 | WP_185639026.1 PepSY domain-containing protein |
| 70(2) | WP_270994741.1 | WP_270994741.1 PepSY domain-containing protein |
| 70(60) | WP_003750463.1 | WP_003750463.1 PepSY domain-containing protein |
| 70(2) | WP_097350517.1 | WP_097350517.1 PepSY domain-containing protein |
| 70(2) | WP_003735137.1 | WP_003735137.1 PepSY domain-containing protein |
| 70(6) | WP_038406367.1 | WP_038406367.1 PepSY domain-containing protein |
| 71(1) | WP_185451819.1 | WP_185451819.1 sensor histidine kinase |
| 71(2) | WP_185363044.1 | WP_185363044.1 sensor histidine kinase |
| 71(15) | WP_003745029.1 | WP_003745029.1 sensor histidine kinase |
| 71(6) | WP_038406370.1 | WP_038406370.1 sensor histidine kinase |
| 71(38) | WP_185336685.1 | WP_185336685.1 sensor histidine kinase |
| 71(2) | WP_003728980.1 | WP_003728980.1 sensor histidine kinase |
| 71(2) | WP_097350518.1 | WP_097350518.1 sensor histidine kinase |
| 71(4) | WP_185474160.1 | WP_185474160.1 sensor histidine kinase |
| 71(2) | WP_270994742.1 | WP_270994742.1 sensor histidine kinase |
| 71(2) | WP_185639024.1 | WP_185639024.1 sensor histidine kinase |
| 72(24) | WP_012984595.1 | WP_012984595.1 cyclic-di-AMP phosphodiesterase PdeA |
| 72(2) | WP_185516174.1 | WP_185516174.1 cyclic-di-AMP phosphodiesterase PdeA |

| | | |
|---|---|---|
| 72(4) | WP_185474159.1 | WP_185474159.1 cyclic-di-AMP phosphodiesterase PdeA |
| 72(14) | WP_003750470.1 | WP_003750470.1 cyclic-di-AMP phosphodiesterase PdeA |
| 72(4) | WP_038406371.1 | WP_038406371.1 cyclic-di-AMP phosphodiesterase PdeA |
| 72(2) | WP_185639022.1 | WP_185639022.1 cyclic-di-AMP phosphodiesterase PdeA |
| 72(14) | WP_185337264.1 | WP_185337264.1 cyclic-di-AMP phosphodiesterase PdeA |
| 72(2) | WP_270995171.1 | WP_270995171.1 cyclic-di-AMP phosphodiesterase PdeA |
| 72(2) | WP_003721675.1 | WP_003721675.1 cyclic-di-AMP phosphodiesterase PdeA |
| 72(2) | WP_185430921.1 | WP_185430921.1 cyclic-di-AMP phosphodiesterase PdeA |
| 72(2) | WP_185460020.1 | WP_185460020.1 cyclic-di-AMP phosphodiesterase PdeA |
| 72(2) | WP_038408415.1 | WP_038408415.1 cyclic-di-AMP phosphodiesterase PdeA |

| | | |
|---|---|---|
| 73(7) | WP_000997785.1 | WP_000997785.1 immunity 22 family protein |
| 73(1) | WP_166705001.1 | WP_166705001.1 PepSY domain-containing protein |
| 73(1) | WP_286017555.1 | WP_286017555.1 immunity 22 family protein |
| 73(1) | WP_238655358.1 | WP_238655358.1 immunity 22 family protein |
| 73(1) | WP_053564620.1 | WP_053564620.1 PepSY domain-containing protein |
| 73(1) | WP_255293735.1 | WP_255293735.1 hypothetical protein |
| 73(1) | WP_000731148.1 | WP_000731148.1 PepSY domain-containing protein |
| 73(1) | WP_000786931.1 | WP_000786931.1 immunity 22 family protein |
| 73(4) | WP_025118714.1 | WP_025118714.1 immunity 22 family protein |
| 73(1) | WP_000731149.1 | WP_000731149.1 PepSY domain-containing protein |
| 73(1) | WP_053564625.1 | WP_053564625.1 immunity 22 family protein |
| 73(1) | WP_103141593.1 | WP_103141593.1 immunity 22 family protein |
| 73(1) | WP_163255261.1 | WP_163255261.1 immunity 22 family protein |
| 73(3) | WP_000414707.1 | WP_000414707.1 immunity 22 family protein |
| 73(1) | WP_119601474.1 | WP_119601474.1 immunity 22 family protein |
| 73(1) | WP_088046790.1 | WP_088046790.1 PepSY domain-containing protein |
| 73(1) | WP_012260847.1 | WP_012260847.1 immunity 22 family protein |
| 73(1) | WP_139867364.1 | WP_139867364.1 PepSY domain-containing protein |
| 73(1) | WP_242251792.1 | WP_242251792.1 immunity 22 family protein |
| 73(1) | WP_262719885.1 | WP_262719885.1 hypothetical protein |
| 73(1) | WP_088032797.1 | WP_088032797.1 immunity 22 family protein |
| 73(5) | WP_000414658.1 | WP_000414658.1 immunity 22 family protein |
| 73(2) | WP_185469292.1 | WP_185469292.1 immunity 22 family protein |
| 73(1) | WP_220775547.1 | WP_220775547.1 PepSY domain-containing protein |
| 73(1) | WP_098239615.1 | WP_098239615.1 hypothetical protein |
| 73(1) | WP_002170261.1 | WP_002170261.1 PepSY domain-containing protein |
| 73(2) | WP_185350822.1 | WP_185350822.1 immunity 22 family protein |
| 73(4) | WP_000997786.1 | WP_000997786.1 immunity 22 family protein |
| 73(1) | WP_215591204.1 | WP_215591204.1 immunity 22 family protein |
| 73(2) | WP_000786932.1 | WP_000786932.1 immunity 22 family protein |
| 73(1) | WP_291672220.1 | WP_291672220.1 immunity 22 family protein |
| 73(2) | WP_098148063.1 | WP_098148063.1 immunity 22 family protein |
| 73(3) | WP_255262854.1 | WP_255262854.1 hypothetical protein |
| 73(2) | WP_087956297.1 | WP_087956297.1 hypothetical protein |
| 73(1) | WP_151630596.1 | WP_151630596.1 PepSY domain-containing protein |
| 73(1) | WP_256933051.1 | WP_256933051.1 hypothetical protein |
| 73(5) | WP_000414798.1 | WP_000414798.1 immunity 22 family protein |
| 73(6) | WP_012094444.1 | WP_012094444.1 immunity 22 family protein |
| 73(1) | WP_029438094.1 | WP_029438094.1 immunity 22 family protein |
| 73(1) | WP_144611195.1 | WP_144611195.1 PepSY domain-containing protein |

| | | |
|---|---|---|
| 74(1) | WP_125084788.1 | WP_125084788.1 hypothetical protein |
| 74(1) | WP_013914200.1 | WP_013914200.1 hypothetical protein |
| 74(1) | WP_049869163.1 | WP_049869163.1 hypothetical protein |
| 74(1) | WP_155609761.1 | WP_155609761.1 hypothetical protein |
| 74(1) | WP_036626779.1 | WP_036626779.1 hypothetical protein |
| 74(1) | WP_223068039.1 | WP_223068039.1 hypothetical protein |
| 74(1) | WP_179034782.1 | WP_179034782.1 hypothetical protein |
| 74(1) | WP_088550048.1 | WP_088550048.1 hypothetical protein |
| 74(1) | WP_165097607.1 | WP_165097607.1 hypothetical protein |
| 74(1) | WP_218641693.1 | WP_218641693.1 hypothetical protein |
| 74(1) | WP_119601502.1 | WP_119601502.1 hypothetical protein |
| 74(1) | WP_026478104.1 | WP_026478104.1 hypothetical protein |
| 74(1) | WP_283910919.1 | WP_283910919.1 hypothetical protein |
| 74(1) | WP_038286710.1 | WP_038286710.1 hypothetical protein |
| 74(1) | WP_128631281.1 | WP_128631281.1 hypothetical protein |
| 74(2) | WP_014368248.1 | WP_014368248.1 hypothetical protein |
| 74(1) | WP_126996299.1 | WP_126996299.1 hypothetical protein |
| 74(1) | WP_236334343.1 | WP_236334343.1 hypothetical protein |
| 74(1) | WP_189031044.1 | WP_189031044.1 hypothetical protein |
| 74(1) | WP_251596684.1 | WP_251596684.1 hypothetical protein |
| 74(1) | WP_195571401.1 | WP_195571401.1 hypothetical protein |
| 74(1) | WP_166244916.1 | WP_166244916.1 hypothetical protein |
| 74(1) | WP_208414540.1 | WP_208414540.1 hypothetical protein |
| 74(1) | WP_283927232.1 | WP_283927232.1 hypothetical protein |
| 74(1) | WP_280787397.1 | WP_280787397.1 hypothetical protein |
| 74(5) | WP_036684496.1 | WP_036684496.1 hypothetical protein |
| 74(1) | WP_055109393.1 | WP_055109393.1 hypothetical protein |
| 74(2) | WP_211022794.1 | WP_211022794.1 hypothetical protein |
| 74(1) | WP_069193879.1 | WP_069193879.1 hypothetical protein |
| 74(3) | WP_076281080.1 | WP_076281080.1 hypothetical protein |
| 74(1) | WP_150457771.1 | WP_150457771.1 hypothetical protein |
| 74(1) | WP_108464082.1 | WP_108464082.1 hypothetical protein |
| 74(1) | WP_213592877.1 | WP_213592877.1 hypothetical protein |
| 74(2) | WP_042125615.1 | WP_042125615.1 hypothetical protein |
| 74(1) | WP_110934079.1 | WP_110934079.1 hypothetical protein |
| 74(1) | WP_076157844.1 | WP_076157844.1 hypothetical protein |
| 74(1) | WP_169028897.1 | WP_169028897.1 hypothetical protein |
| 74(1) | WP_238655343.1 | WP_238655343.1 hypothetical protein |
| 74(1) | WP_132283084.1 | WP_132283084.1 hypothetical protein |
| 74(1) | WP_215185467.1 | WP_215185467.1 hypothetical protein |
| 74(1) | WP_249897310.1 | WP_249897310.1 hypothetical protein |
| 74(1) | WP_148450952.1 | WP_148450952.1 hypothetical protein |
| 74(1) | WP_013658718.1 | WP_013658718.1 hypothetical protein |
| 74(2) | WP_101300033.1 | WP_101300033.1 hypothetical protein |
| 74(1) | WP_188535060.1 | WP_188535060.1 hypothetical protein |
| 74(1) | WP_036702424.1 | WP_036702424.1 hypothetical protein |
| 74(1) | WP_213506870.1 | WP_213506870.1 hypothetical protein |
| 74(1) | WP_257443657.1 | WP_257443657.1 hypothetical protein |
| 74(1) | WP_036944565.1 | WP_036944565.1 hypothetical protein |
| 74(1) | WP_251417649.1 | WP_251417649.1 hypothetical protein |
| 74(1) | WP_076146943.1 | WP_076146943.1 hypothetical protein |
| 74(1) | WP_199613908.1 | WP_199613908.1 hypothetical protein |

| | | |
|---|---|---|
| 74(1) | WP_136479369.1 | WP_136479369.1 hypothetical protein |
| 74(2) | WP_094870540.1 | WP_094870540.1 hypothetical protein |
| 74(1) | WP_197937388.1 | WP_197937388.1 hypothetical protein |
| 74(1) | WP_076271934.1 | WP_076271934.1 hypothetical protein |
| 74(1) | WP_003516566.1 | WP_003516566.1 hypothetical protein |
| 74(1) | WP_048749317.1 | WP_048749317.1 hypothetical protein, partial |
| 74(1) | WP_244723802.1 | WP_244723802.1 hypothetical protein |
| 74(1) | WP_137698117.1 | WP_137698117.1 hypothetical protein |
| 74(1) | WP_120462956.1 | WP_120462956.1 hypothetical protein |
| 74(1) | WP_085499112.1 | WP_085499112.1 hypothetical protein |

| | | |
|---|---|---|
| 75(1) | WP_273321516.1 | WP_273321516.1 hypothetical protein |
| 75(1) | WP_127570447.1 | WP_127570447.1 hypothetical protein |
| 75(2) | WP_074085346.1 | WP_074085346.1 hypothetical protein |
| 75(1) | WP_068782419.1 | WP_068782419.1 hypothetical protein |
| 75(1) | WP_042203659.1 | WP_042203659.1 hypothetical protein |
| 75(1) | WP_247681911.1 | WP_247681911.1 hypothetical protein |
| 75(1) | WP_301290466.1 | WP_301290466.1 hypothetical protein |
| 75(1) | WP_166244093.1 | WP_166244093.1 hypothetical protein |
| 75(1) | WP_039871667.1 | WP_039871667.1 hypothetical protein |
| 75(1) | WP_138755930.1 | WP_138755930.1 hypothetical protein |
| 75(1) | WP_019912140.1 | WP_019912140.1 hypothetical protein |
| 75(2) | WP_219872466.1 | WP_219872466.1 hypothetical protein |
| 75(2) | WP_076104584.1 | WP_076104584.1 hypothetical protein |
| 75(1) | WP_209872716.1 | WP_209872716.1 hypothetical protein |
| 75(2) | WP_076287006.1 | WP_076287006.1 hypothetical protein |
| 75(1) | WP_182301849.1 | WP_182301849.1 hypothetical protein |
| 75(1) | WP_152392584.1 | WP_152392584.1 hypothetical protein |
| 75(1) | WP_307199920.1 | WP_307199920.1 hypothetical protein |
| 75(1) | WP_244988295.1 | WP_244988295.1 hypothetical protein |
| 75(3) | WP_076135789.1 | WP_076135789.1 hypothetical protein |
| 75(1) | WP_017692344.1 | WP_017692344.1 hypothetical protein |
| 75(1) | WP_074048566.1 | WP_074048566.1 hypothetical protein |
| 75(1) | WP_134750632.1 | WP_134750632.1 hypothetical protein |
| 75(1) | WP_089522646.1 | WP_089522646.1 hypothetical protein |
| 75(1) | WP_212695804.1 | WP_212695804.1 hypothetical protein |
| 75(1) | WP_114298758.1 | WP_114298758.1 hypothetical protein |
| 75(1) | WP_068696908.1 | WP_068696908.1 hypothetical protein |
| 75(1) | WP_154118020.1 | WP_154118020.1 hypothetical protein |
| 75(1) | WP_076219083.1 | WP_076219083.1 hypothetical protein |
| 75(5) | WP_110899372.1 | WP_110899372.1 hypothetical protein |
| 75(1) | WP_307601005.1 | WP_307601005.1 hypothetical protein |
| 75(1) | WP_076210963.1 | WP_076210963.1 hypothetical protein |
| 75(1) | WP_076144386.1 | WP_076144386.1 hypothetical protein |
| 75(1) | WP_027088416.1 | WP_027088416.1 hypothetical protein |
| 75(1) | WP_150364192.1 | WP_150364192.1 hypothetical protein |
| 75(1) | WP_145153661.1 | WP_145153661.1 hypothetical protein |
| 75(1) | WP_024632691.1 | WP_024632691.1 hypothetical protein |
| 75(1) | WP_239617486.1 | WP_239617486.1 hypothetical protein |
| 75(3) | WP_094870410.1 | WP_094870410.1 hypothetical protein |
| 75(1) | WP_210087117.1 | WP_210087117.1 hypothetical protein |
| 75(1) | WP_039306323.1 | WP_039306323.1 hypothetical protein |
| 75(1) | WP_245700194.1 | WP_245700194.1 hypothetical protein |
| 75(1) | WP_042134293.1 | WP_042134293.1 hypothetical protein |
| 75(2) | WP_281813139.1 | WP_281813139.1 hypothetical protein |
| 75(1) | WP_293755484.1 | WP_293755484.1 hypothetical protein |
| 75(1) | WP_248063430.1 | WP_248063430.1 hypothetical protein |
| 75(1) | WP_099517094.1 | WP_099517094.1 hypothetical protein |
| 75(3) | WP_076279374.1 | WP_076279374.1 hypothetical protein |
| 75(1) | WP_110931529.1 | WP_110931529.1 hypothetical protein |
| 75(1) | WP_189018584.1 | WP_189018584.1 hypothetical protein |
| 75(1) | WP_212693252.1 | WP_212693252.1 hypothetical protein |
| 75(1) | WP_042211007.1 | WP_042211007.1 hypothetical protein |
| 75(1) | WP_238187872.1 | WP_238187872.1 hypothetical protein |
| 75(1) | WP_127194970.1 | WP_127194970.1 hypothetical protein |
| 75(1) | WP_120112284.1 | WP_120112284.1 hypothetical protein |
| 75(1) | WP_308723527.1 | WP_308723527.1 hypothetical protein |
| 75(1) | WP_076242362.1 | WP_076242362.1 hypothetical protein |
| 75(1) | WP_136372946.1 | WP_136372946.1 hypothetical protein |

| | | |
|---|---|---|
| 76(1) | WP_238655342.1 | WP_238655342.1 hypothetical protein |
| 76(1) | WP_101300031.1 | WP_101300031.1 hypothetical protein |
| 76(2) | WP_036702426.1 | WP_036702426.1 hypothetical protein |
| 76(3) | WP_076281081.1 | WP_076281081.1 hypothetical protein |
| 76(1) | WP_236334345.1 | WP_236334345.1 hypothetical protein |
| 76(1) | WP_125084789.1 | WP_125084789.1 hypothetical protein |
| 76(1) | WP_257443658.1 | WP_257443658.1 hypothetical protein |
| 76(1) | WP_102710714.1 | WP_102710714.1 hypothetical protein |
| 76(2) | WP_211022795.1 | WP_211022795.1 hypothetical protein |
| 76(1) | WP_166244914.1 | WP_166244914.1 hypothetical protein |
| 76(8) | WP_036684498.1 | WP_036684498.1 hypothetical protein |
| 76(2) | WP_189031046.1 | WP_189031046.1 hypothetical protein |
| 76(2) | WP_213592879.1 | WP_213592879.1 hypothetical protein |
| 76(1) | WP_165097605.1 | WP_165097605.1 hypothetical protein |
| 76(2) | WP_148450953.1 | WP_148450953.1 hypothetical protein |
| 76(1) | WP_036626782.1 | WP_036626782.1 hypothetical protein |
| 76(1) | WP_150457772.1 | WP_150457772.1 hypothetical protein |
| 76(2) | WP_042125613.1 | WP_042125613.1 hypothetical protein |
| 76(1) | WP_197937387.1 | WP_197937387.1 hypothetical protein |
| 76(1) | WP_110934080.1 | WP_110934080.1 hypothetical protein |
| 76(2) | WP_085499114.1 | WP_085499114.1 hypothetical protein |
| 76(1) | WP_126996297.1 | WP_126996297.1 hypothetical protein |
| 76(1) | WP_188535057.1 | WP_188535057.1 hypothetical protein |
| 76(1) | WP_179034781.1 | WP_179034781.1 hypothetical protein |
| 76(1) | WP_013914199.1 | WP_013914199.1 hypothetical protein |
| 76(1) | WP_026478103.1 | WP_026478103.1 hypothetical protein |
| 76(1) | WP_199613909.1 | WP_199613909.1 hypothetical protein |
| 76(1) | WP_013658719.1 | WP_013658719.1 hypothetical protein |
| 76(1) | WP_049869162.1 | WP_049869162.1 hypothetical protein |
| 76(1) | WP_038286709.1 | WP_038286709.1 hypothetical protein |
| 76(1) | WP_223068038.1 | WP_223068038.1 hypothetical protein |
| 76(1) | WP_234414301.1 | WP_234414301.1 hypothetical protein |
| 76(1) | WP_249897309.1 | WP_249897309.1 hypothetical protein |
| 76(1) | WP_069193880.1 | WP_069193880.1 hypothetical protein |

| | | |
|---|---|---|
| 76(1) | WP_242971696.1 | WP_242971696.1 hypothetical protein |
| 76(1) | WP_283910918.1 | WP_283910918.1 hypothetical protein |
| 76(1) | WP_119601504.1 | WP_119601504.1 hypothetical protein |
| 76(1) | WP_120462957.1 | WP_120462957.1 hypothetical protein |
| 76(1) | WP_244723804.1 | WP_244723804.1 hypothetical protein |
| 76(1) | WP_169028898.1 | WP_169028898.1 hypothetical protein |
| 76(2) | WP_003516564.1 | WP_003516564.1 hypothetical protein |
| 76(1) | WP_251417647.1 | WP_251417647.1 hypothetical protein |
| 76(2) | WP_094876490.1 | WP_094876490.1 hypothetical protein |
| 76(1) | WP_036944564.1 | WP_036944564.1 hypothetical protein |
| 76(1) | WP_195571402.1 | WP_195571402.1 hypothetical protein |
| 76(1) | WP_128631280.1 | WP_128631280.1 hypothetical protein |
| 76(1) | WP_243117052.1 | WP_243117052.1 hypothetical protein |
| 76(1) | WP_155609762.1 | WP_155609762.1 hypothetical protein |
| 76(1) | WP_094870541.1 | WP_094870541.1 hypothetical protein |
| 76(1) | WP_055109396.1 | WP_055109396.1 hypothetical protein |
| 76(2) | WP_014368247.1 | WP_014368247.1 hypothetical protein |
| 76(1) | WP_137698118.1 | WP_137698118.1 hypothetical protein |
| 76(1) | WP_251596686.1 | WP_251596686.1 hypothetical protein |
| 77(1) | WP_102710711.1 | WP_102710711.1 flagellar biosynthesis protein FlgA |
| 77(2) | WP_036702431.1 | WP_036702431.1 hypothetical protein |
| 77(1) | WP_244723809.1 | WP_244723809.1 hypothetical protein |
| 77(1) | WP_076281084.1 | WP_076281084.1 hypothetical protein |
| 77(1) | WP_155609765.1 | WP_155609765.1 flagellar biosynthesis protein FlgA |
| 77(1) | WP_094876491.1 | WP_094876491.1 hypothetical protein |
| 77(1) | WP_055109404.1 | WP_055109404.1 hypothetical protein |
| 77(1) | WP_251417640.1 | WP_251417640.1 hypothetical protein |
| 77(1) | WP_132283081.1 | WP_132283081.1 flagellar biosynthesis protein FlgA |
| 77(2) | WP_013914196.1 | WP_013914196.1 hypothetical protein |
| 77(2) | WP_094870544.1 | WP_094870544.1 hypothetical protein |
| 77(1) | WP_136479367.1 | WP_136479367.1 flagellar biosynthesis protein FlgA |
| 77(1) | WP_013658722.1 | WP_013658722.1 hypothetical protein |
| 77(2) | WP_085499120.1 | WP_085499120.1 hypothetical protein |
| 77(1) | WP_110934083.1 | WP_110934083.1 flagellar biosynthesis protein FlgA |
| 77(1) | WP_215185461.1 | WP_215185461.1 hypothetical protein |
| 77(1) | WP_213506868.1 | WP_213506868.1 hypothetical protein |
| 77(1) | WP_249897306.1 | WP_249897306.1 hypothetical protein |
| 77(1) | WP_125084792.1 | WP_125084792.1 flagellar biosynthesis protein FlgA |
| 77(1) | WP_148450956.1 | WP_148450956.1 flagellar biosynthesis protein FlgA |
| 77(1) | WP_257443660.1 | WP_257443660.1 hypothetical protein |
| 77(2) | WP_213592883.1 | WP_213592883.1 hypothetical protein |
| 77(7) | WP_036684504.1 | WP_036684504.1 hypothetical protein |
| 77(1) | WP_014368245.1 | WP_014368245.1 hypothetical protein |
| 77(1) | WP_108464078.1 | WP_108464078.1 flagellar biosynthesis protein FlgA |
| 77(1) | WP_021878926.1 | WP_021878926.1 hypothetical protein |
| 77(1) | WP_251596692.1 | WP_251596692.1 hypothetical protein |
| 77(1) | WP_238655339.1 | WP_238655339.1 hypothetical protein |
| 77(1) | WP_188535052.1 | WP_188535052.1 flagellar biosynthesis protein FlgA |
| 77(1) | WP_069196264.1 | WP_069196264.1 flagellar biosynthesis protein FlgA |
| 77(1) | WP_036944560.1 | WP_036944560.1 hypothetical protein |
| 77(1) | WP_128631277.1 | WP_128631277.1 flagellar biosynthesis protein FlgA |
| 77(1) | WP_195571407.1 | WP_195571407.1 flagellar biosynthesis protein FlgA |
| 77(1) | WP_165097596.1 | WP_165097596.1 flagellar biosynthesis protein FlgA |
| 77(1) | WP_150457775.1 | WP_150457775.1 flagellar biosynthesis protein FlgA |
| 77(1) | WP_036626788.1 | WP_036626788.1 hypothetical protein |
| 77(1) | WP_179034778.1 | WP_179034778.1 flagellar biosynthesis protein FlgA |
| 77(1) | WP_207952339.1 | WP_207952339.1 hypothetical protein |
| 77(1) | WP_119601510.1 | WP_119601510.1 flagellar biosynthesis protein FlgA |
| 77(2) | WP_211022798.1 | WP_211022798.1 hypothetical protein |
| 77(1) | WP_283910915.1 | WP_283910915.1 hypothetical protein |
| 77(1) | WP_169028900.1 | WP_169028900.1 flagellar biosynthesis protein FlgA |
| 77(1) | WP_197937384.1 | WP_197937384.1 hypothetical protein |
| 77(1) | WP_223068035.1 | WP_223068035.1 hypothetical protein |
| 77(1) | WP_189031048.1 | WP_189031048.1 flagellar biosynthesis protein FlgA |
| 77(1) | WP_038286704.1 | WP_038286704.1 hypothetical protein |
| 77(1) | WP_076103999.1 | WP_076103999.1 hypothetical protein |
| 77(2) | WP_076301355.1 | WP_076301355.1 hypothetical protein |
| 77(1) | WP_236334351.1 | WP_236334351.1 hypothetical protein |
| 77(1) | WP_051569330.1 | WP_051569330.1 hypothetical protein |
| 77(2) | WP_042125607.1 | WP_042125607.1 hypothetical protein |
| 77(1) | WP_003516558.1 | WP_003516558.1 hypothetical protein |
| 77(2) | WP_101300026.1 | WP_101300026.1 flagellar biosynthesis protein FlgA |
| 77(1) | WP_120462960.1 | WP_120462960.1 flagellar biosynthesis protein FlgA |
| 77(1) | WP_126996291.1 | WP_126996291.1 flagellar biosynthesis protein FlgA |
| 77(1) | WP_199613912.1 | WP_199613912.1 hypothetical protein |
| 78(1) | WP_279830843.1 | WP_279830843.1 flagellar motor protein MotB |
| 78(2) | WP_230877355.1 | WP_230877355.1 flagellar motor protein MotB |
| 78(2) | WP_238791006.1 | WP_238791006.1 flagellar motor protein MotB |
| 78(1) | WP_256715587.1 | WP_256715587.1 flagellar motor protein MotB |
| 78(1) | WP_025719065.1 | WP_025719065.1 hypothetical protein |
| 78(1) | WP_258957356.1 | WP_258957356.1 flagellar motor protein MotB |
| 78(1) | WP_254348137.1 | WP_254348137.1 flagellar motor protein MotB |
| 78(1) | WP_230585141.1 | WP_230585141.1 flagellar motor protein MotB |
| 78(1) | WP_058829622.1 | WP_058829622.1 hypothetical protein |
| 78(1) | WP_245233399.1 | WP_245233399.1 flagellar motor protein MotB |
| 78(1) | WP_258296282.1 | WP_258296282.1 flagellar motor protein MotB |
| 78(1) | WP_029517060.1 | WP_029517060.1 hypothetical protein |
| 78(1) | WP_250251636.1 | WP_250251636.1 flagellar motor protein MotB |
| 78(1) | WP_090887135.1 | WP_090887135.1 flagellar motor protein MotB |
| 78(1) | WP_013309984.1 | WP_013309984.1 hypothetical protein |
| 78(1) | WP_231887380.1 | WP_231887380.1 flagellar motor protein MotB |
| 78(1) | WP_245240471.1 | WP_245240471.1 flagellar motor protein MotB |
| 78(1) | WP_250244813.1 | WP_250244813.1 flagellar motor protein MotB |
| 78(1) | WP_244315815.1 | WP_244315815.1 flagellar motor protein MotB |
| 78(2) | WP_028542692.1 | WP_028542692.1 hypothetical protein |
| 78(1) | WP_240664150.1 | WP_240664150.1 flagellar motor protein MotB |
| 78(4) | WP_013370780.1 | WP_013370780.1 hypothetical protein |
| 78(1) | WP_223264158.1 | WP_223264158.1 flagellar motor protein MotB |
| 78(3) | WP_240753675.1 | WP_240753675.1 flagellar motor protein MotB |
| 78(1) | WP_252508782.1 | WP_252508782.1 flagellar motor protein MotB |
| 78(1) | WP_254349524.1 | WP_254349524.1 flagellar motor protein MotB |
| 78(2) | WP_023988261.1 | WP_023988261.1 hypothetical protein |

| | | |
|---|---|---|
| 78(1) | WP_225999404.1 | WP_225999404.1 flagellar motor protein MotB |
| 78(3) | WP_225164875.1 | WP_225164875.1 flagellar motor protein MotB |
| 78(1) | WP_237714611.1 | WP_237714611.1 flagellar motor protein MotB |
| 78(1) | WP_307349958.1 | WP_307349958.1 flagellar motor protein MotB |
| 78(1) | WP_226888622.1 | WP_226888622.1 flagellar motor protein MotB |
| 78(1) | WP_254325795.1 | WP_254325795.1 flagellar motor protein MotB |
| 78(1) | WP_238353837.1 | WP_238353837.1 flagellar motor protein MotB |
| 78(1) | WP_258383060.1 | WP_258383060.1 flagellar motor protein MotB |
| 78(1) | WP_231109417.1 | WP_231109417.1 flagellar motor protein MotB |
| 78(1) | WP_256703527.1 | WP_256703527.1 flagellar motor protein MotB |
| 78(1) | WP_258012393.1 | WP_258012393.1 flagellar motor protein MotB |
| 78(1) | WP_231503571.1 | WP_231503571.1 flagellar motor protein MotB |
| 78(1) | WP_305166264.1 | WP_305166264.1 flagellar motor protein MotB |
| 78(2) | WP_283655689.1 | WP_283655689.1 flagellar motor protein MotB |
| 78(1) | WP_231124575.1 | WP_231124575.1 flagellar motor protein MotB |
| 78(1) | WP_223822052.1 | WP_223822052.1 flagellar motor protein MotB |
| 78(3) | WP_230681838.1 | WP_230681838.1 flagellar motor protein MotB |
| 78(1) | WP_272640513.1 | WP_272640513.1 flagellar motor protein MotB |
| 78(1) | WP_274485146.1 | WP_274485146.1 flagellar motor protein MotB |
| 78(1) | WP_233901859.1 | WP_233901859.1 flagellar motor protein MotB |
| 78(1) | WP_025679801.1 | WP_025679801.1 hypothetical protein |
| 78(1) | WP_238807607.1 | WP_238807607.1 flagellar motor protein MotB |
| 78(1) | WP_254664445.1 | WP_254664445.1 flagellar motor protein MotB |
| 78(1) | WP_245248386.1 | WP_245248386.1 flagellar motor protein MotB |
| 78(2) | WP_256703236.1 | WP_256703236.1 flagellar motor protein MotB |
| 78(1) | WP_254327964.1 | WP_254327964.1 flagellar motor protein MotB |
| 78(1) | WP_209018285.1 | WP_209018285.1 flagellar motor protein MotB |
| 78(1) | WP_226887696.1 | WP_226887696.1 flagellar motor protein MotB |
| 78(1) | WP_244310717.1 | WP_244310717.1 flagellar motor protein MotB |
| 79(1) | WP_030025436.1 | WP_030025436.1 TIGR04197 family type VII secretion effector |
| 79(2) | WP_000503892.1 | WP_000503892.1 TIGR04197 family type VII secretion effector |
| 79(1) | WP_098226391.1 | WP_098226391.1 TIGR04197 family type VII secretion effector |
| 79(4) | WP_001990255.1 | WP_001990255.1 TIGR04197 family type VII secretion effector |
| 79(2) | WP_061113195.1 | WP_061113195.1 TIGR04197 family type VII secretion effector |
| 79(11) | WP_053512322.1 | WP_053512322.1 TIGR04197 family type VII secretion effector |
| 79(1) | WP_000503888.1 | WP_000503888.1 TIGR04197 family type VII secretion effector |
| 79(1) | WP_098287079.1 | WP_098287079.1 TIGR04197 family type VII secretion effector |
| 79(1) | WP_153589100.1 | WP_153589100.1 TIGR04197 family type VII secretion effector |
| 79(2) | WP_251217309.1 | WP_251217309.1 TIGR04197 family type VII secretion effector |
| 79(3) | WP_044307211.1 | WP_044307211.1 TIGR04197 family type VII secretion effector |
| 79(1) | WP_098592734.1 | WP_098592734.1 TIGR04197 family type VII secretion effector |
| 79(1) | WP_098157700.1 | WP_098157700.1 TIGR04197 family type VII secretion effector |
| 79(2) | WP_016117668.1 | WP_016117668.1 TIGR04197 family type VII secretion effector |
| 79(3) | WP_000503885.1 | WP_000503885.1 TIGR04197 family type VII secretion effector |
| 79(3) | WP_097837317.1 | WP_097837317.1 TIGR04197 family type VII secretion effector |
| 79(1) | WP_106496023.1 | WP_106496023.1 TIGR04197 family type VII secretion effector |
| 79(3) | WP_000503889.1 | WP_000503889.1 TIGR04197 family type VII secretion effector |
| 79(4) | WP_000503891.1 | WP_000503891.1 TIGR04197 family type VII secretion effector |
| 79(1) | WP_077320206.1 | WP_077320206.1 TIGR04197 family type VII secretion effector |
| 79(1) | WP_078384708.1 | WP_078384708.1 TIGR04197 family type VII secretion effector |
| 79(1) | WP_049107940.1 | WP_049107940.1 TIGR04197 family type VII secretion effector |

| | | |
|---|---|---|
| 79(13) | WP_000503886.1 | WP_000503886.1 TIGR04197 family type VII secretion effector |
| 79(4) | WP_000503887.1 | WP_000503887.1 TIGR04197 family type VII secretion effector |
| 79(1) | WP_163095411.1 | WP_163095411.1 TIGR04197 family type VII secretion effector |
| 79(1) | WP_006923818.1 | WP_006923818.1 TIGR04197 family type VII secretion effector |
| 80(1) | WP_026478105.1 | WP_026478105.1 hypothetical protein |
| 80(1) | WP_251596682.1 | WP_251596682.1 hypothetical protein |
| 80(1) | WP_055109392.1 | WP_055109392.1 hypothetical protein |
| 80(2) | WP_211022793.1 | WP_211022793.1 hypothetical protein |
| 80(1) | WP_199613907.1 | WP_199613907.1 hypothetical protein |
| 80(2) | WP_101300035.1 | WP_101300035.1 hypothetical protein |
| 80(1) | WP_110934078.1 | WP_110934078.1 hypothetical protein |
| 80(1) | WP_251417652.1 | WP_251417652.1 hypothetical protein |
| 80(1) | WP_213509995.1 | WP_213509995.1 hypothetical protein |
| 80(2) | WP_141097600.1 | WP_141097600.1 hypothetical protein |
| 80(1) | WP_132283085.1 | WP_132283085.1 hypothetical protein |
| 80(2) | WP_003516568.1 | WP_003516568.1 hypothetical protein |
| 80(1) | WP_249897311.1 | WP_249897311.1 hypothetical protein |
| 80(1) | WP_150457770.1 | WP_150457770.1 hypothetical protein |
| 80(1) | WP_069193878.1 | WP_069193878.1 hypothetical protein |
| 80(1) | WP_148450951.1 | WP_148450951.1 hypothetical protein |
| 80(1) | WP_257443656.1 | WP_257443656.1 hypothetical protein |
| 80(1) | WP_215185468.1 | WP_215185468.1 hypothetical protein |
| 80(1) | WP_137698116.1 | WP_137698116.1 hypothetical protein |
| 80(1) | WP_120462955.1 | WP_120462955.1 hypothetical protein |
| 80(1) | WP_076119201.1 | WP_076119201.1 hypothetical protein |
| 80(1) | WP_195571399.1 | WP_195571399.1 hypothetical protein |
| 80(1) | WP_236334341.1 | WP_236334341.1 hypothetical protein |
| 80(1) | WP_085499110.1 | WP_085499110.1 hypothetical protein |
| 80(1) | WP_238655344.1 | WP_238655344.1 hypothetical protein |
| 80(1) | WP_197937389.1 | WP_197937389.1 hypothetical protein |
| 80(1) | WP_036944567.1 | WP_036944567.1 hypothetical protein |
| 80(1) | WP_213506871.1 | WP_213506871.1 hypothetical protein |
| 80(1) | WP_108464083.1 | WP_108464083.1 hypothetical protein |
| 80(3) | WP_094870539.1 | WP_094870539.1 hypothetical protein |
| 80(4) | WP_076281079.1 | WP_076281079.1 hypothetical protein |
| 80(1) | WP_125084787.1 | WP_125084787.1 hypothetical protein |
| 80(2) | WP_213592875.1 | WP_213592875.1 hypothetical protein |
| 80(8) | WP_036684494.1 | WP_036684494.1 hypothetical protein |
| 80(1) | WP_189031042.1 | WP_189031042.1 hypothetical protein |
| 80(1) | WP_155609760.1 | WP_155609760.1 hypothetical protein |
| 80(1) | WP_179034783.1 | WP_179034783.1 hypothetical protein |
| 80(1) | WP_119601500.1 | WP_119601500.1 hypothetical protein |
| 80(1) | WP_126996301.1 | WP_126996301.1 hypothetical protein |
| 80(1) | WP_244723800.1 | WP_244723800.1 hypothetical protein |
| 80(1) | WP_042125617.1 | WP_042125617.1 hypothetical protein |
| 80(1) | WP_102710716.1 | WP_102710716.1 hypothetical protein |
| 80(1) | WP_188535062.1 | WP_188535062.1 hypothetical protein |
| 80(1) | WP_038286713.1 | WP_038286713.1 hypothetical protein |
| 80(1) | WP_223068040.1 | WP_223068040.1 hypothetical protein |
| 80(1) | WP_128631282.1 | WP_128631282.1 hypothetical protein |
| 80(1) | WP_165097608.1 | WP_165097608.1 hypothetical protein |

| | | |
|---|---|---|
| 80(2) | WP_048749319.1 | WP_048749319.1 hypothetical protein |
| 80(2) | WP_036702422.1 | WP_036702422.1 hypothetical protein |
| 80(1) | WP_013658717.1 | WP_013658717.1 hypothetical protein |
| 81(1) | WP_223068042.1 | WP_223068042.1 hypothetical protein |
| 81(1) | WP_213509970.1 | WP_213509970.1 hypothetical protein |
| 81(2) | WP_211022791.1 | WP_211022791.1 hypothetical protein |
| 81(1) | WP_026478107.1 | WP_026478107.1 hypothetical protein |
| 81(1) | WP_188535065.1 | WP_188535065.1 hypothetical protein |
| 81(2) | WP_048749321.1 | WP_048749321.1 hypothetical protein |
| 81(1) | WP_013658715.1 | WP_013658715.1 hypothetical protein |
| 81(1) | WP_195571395.1 | WP_195571395.1 hypothetical protein |
| 81(1) | WP_150457768.1 | WP_150457768.1 hypothetical protein |
| 81(2) | WP_076281078.1 | WP_076281078.1 hypothetical protein |
| 81(1) | WP_069193876.1 | WP_069193876.1 hypothetical protein |
| 81(1) | WP_055109388.1 | WP_055109388.1 hypothetical protein |
| 81(1) | WP_244723796.1 | WP_244723796.1 hypothetical protein |
| 81(1) | WP_156114749.1 | WP_156114749.1 hypothetical protein |
| 81(1) | WP_213592871.1 | WP_213592871.1 hypothetical protein |
| 81(1) | WP_132283087.1 | WP_132283087.1 hypothetical protein |
| 81(1) | WP_257443653.1 | WP_257443653.1 hypothetical protein |
| 81(1) | WP_236334338.1 | WP_236334338.1 hypothetical protein |
| 81(1) | WP_125084785.1 | WP_125084785.1 hypothetical protein |
| 81(1) | WP_238655346.1 | WP_238655346.1 hypothetical protein |
| 81(1) | WP_280752664.1 | WP_280752664.1 hypothetical protein |
| 81(1) | WP_251596678.1 | WP_251596678.1 hypothetical protein |
| 81(2) | WP_036702419.1 | WP_036702419.1 hypothetical protein |
| 81(1) | WP_139829223.1 | WP_139829223.1 hypothetical protein |
| 81(1) | WP_249897313.1 | WP_249897313.1 hypothetical protein |
| 81(1) | WP_102710718.1 | WP_102710718.1 hypothetical protein |
| 81(1) | WP_199613905.1 | WP_199613905.1 hypothetical protein |
| 81(2) | WP_003516571.1 | WP_003516571.1 hypothetical protein |
| 81(1) | WP_139328519.1 | WP_139328519.1 hypothetical protein |
| 81(1) | WP_197937391.1 | WP_197937391.1 hypothetical protein |
| 81(1) | WP_251417657.1 | WP_251417657.1 hypothetical protein |
| 81(8) | WP_036684491.1 | WP_036684491.1 hypothetical protein |
| 81(1) | WP_141097599.1 | WP_141097599.1 hypothetical protein |
| 81(1) | WP_169028895.1 | WP_169028895.1 hypothetical protein |
| 81(1) | WP_036944570.1 | WP_036944570.1 hypothetical protein |
| 81(1) | WP_189031038.1 | WP_189031038.1 hypothetical protein |
| 81(1) | WP_215185470.1 | WP_215185470.1 hypothetical protein |
| 81(1) | WP_165097612.1 | WP_165097612.1 hypothetical protein |
| 81(1) | WP_038286715.1 | WP_038286715.1 hypothetical protein |
| 81(3) | WP_094870537.1 | WP_094870537.1 hypothetical protein |
| 81(1) | WP_128631284.1 | WP_128631284.1 hypothetical protein |
| 81(1) | WP_213506873.1 | WP_213506873.1 hypothetical protein |
| 81(1) | WP_110934076.1 | WP_110934076.1 hypothetical protein |
| 81(1) | WP_283910921.1 | WP_283910921.1 hypothetical protein |
| 81(1) | WP_283927233.1 | WP_283927233.1 hypothetical protein |
| 81(1) | WP_179034785.1 | WP_179034785.1 hypothetical protein |
| 81(1) | WP_148450949.1 | WP_148450949.1 hypothetical protein |
| 81(1) | WP_108464085.1 | WP_108464085.1 hypothetical protein |
| 81(1) | WP_137698114.1 | WP_137698114.1 hypothetical protein |
| 81(1) | WP_120462953.1 | WP_120462953.1 hypothetical protein |
| 81(2) | WP_101300039.1 | WP_101300039.1 hypothetical protein |
| 81(1) | WP_126996305.1 | WP_126996305.1 hypothetical protein |
| 81(1) | WP_147381825.1 | WP_147381825.1 hypothetical protein |
| 81(1) | WP_196427067.1 | WP_196427067.1 hypothetical protein |
| 82(1) | WP_036944568.1 | WP_036944568.1 hypothetical protein |
| 82(1) | WP_055109390.1 | WP_055109390.1 hypothetical protein |
| 82(4) | WP_076103998.1 | WP_076103998.1 hypothetical protein |
| 82(1) | WP_076101566.1 | WP_076101566.1 hypothetical protein |
| 82(1) | WP_195571397.1 | WP_195571397.1 hypothetical protein |
| 82(1) | WP_126996303.1 | WP_126996303.1 hypothetical protein |
| 82(1) | WP_199613906.1 | WP_199613906.1 hypothetical protein |
| 82(1) | WP_102710717.1 | WP_102710717.1 hypothetical protein |
| 82(1) | WP_251596680.1 | WP_251596680.1 hypothetical protein |
| 82(1) | WP_211022792.1 | WP_211022792.1 hypothetical protein |
| 82(1) | WP_128631283.1 | WP_128631283.1 hypothetical protein |
| 82(2) | WP_003516570.1 | WP_003516570.1 hypothetical protein |
| 82(1) | WP_223068041.1 | WP_223068041.1 hypothetical protein |
| 82(1) | WP_213509971.1 | WP_213509971.1 hypothetical protein |
| 82(1) | WP_148450950.1 | WP_148450950.1 hypothetical protein |
| 82(1) | WP_169028896.1 | WP_169028896.1 hypothetical protein |
| 82(5) | WP_036684493.1 | WP_036684493.1 hypothetical protein |
| 82(1) | WP_188535064.1 | WP_188535064.1 hypothetical protein |
| 82(1) | WP_013658716.1 | WP_013658716.1 hypothetical protein |
| 82(1) | WP_283910920.1 | WP_283910920.1 hypothetical protein |
| 82(3) | WP_094870538.1 | WP_094870538.1 hypothetical protein |
| 82(1) | WP_238655345.1 | WP_238655345.1 hypothetical protein |
| 82(1) | WP_110934077.1 | WP_110934077.1 hypothetical protein |
| 82(1) | WP_280752663.1 | WP_280752663.1 hypothetical protein |
| 82(1) | WP_088550046.1 | WP_088550046.1 hypothetical protein |
| 82(1) | WP_108464084.1 | WP_108464084.1 hypothetical protein |
| 82(1) | WP_290407046.1 | WP_290407046.1 hypothetical protein |
| 82(1) | WP_036702420.1 | WP_036702420.1 hypothetical protein |
| 82(1) | WP_085499108.1 | WP_085499108.1 hypothetical protein |
| 82(1) | WP_165097610.1 | WP_165097610.1 hypothetical protein |
| 82(1) | WP_249897312.1 | WP_249897312.1 hypothetical protein |
| 82(1) | WP_076119203.1 | WP_076119203.1 hypothetical protein |
| 82(1) | WP_105368172.1 | WP_105368172.1 hypothetical protein |
| 82(1) | WP_179034784.1 | WP_179034784.1 hypothetical protein |
| 82(1) | WP_189031040.1 | WP_189031040.1 hypothetical protein |
| 82(1) | WP_213506872.1 | WP_213506872.1 hypothetical protein |
| 82(1) | WP_244723798.1 | WP_244723798.1 hypothetical protein |
| 82(1) | WP_069193877.1 | WP_069193877.1 hypothetical protein |
| 82(1) | WP_265444536.1 | WP_265444536.1 hypothetical protein |
| 82(1) | WP_120462954.1 | WP_120462954.1 hypothetical protein |
| 82(1) | WP_155609759.1 | WP_155609759.1 hypothetical protein |
| 82(1) | WP_132283086.1 | WP_132283086.1 hypothetical protein |
| 82(2) | WP_213592873.1 | WP_213592873.1 hypothetical protein |
| 82(1) | WP_076157841.1 | WP_076157841.1 hypothetical protein |
| 82(1) | WP_119601498.1 | WP_119601498.1 hypothetical protein |

| | | |
|---|---|---|
| 82(1) | WP_137698115.1 | WP_137698115.1 hypothetical protein |
| 82(1) | WP_026478106.1 | WP_026478106.1 hypothetical protein |
| 82(1) | WP_257443655.1 | WP_257443655.1 hypothetical protein |
| 82(1) | WP_048749320.1 | WP_048749320.1 hypothetical protein |
| 82(1) | WP_251417654.1 | WP_251417654.1 hypothetical protein |
| 82(1) | WP_150457769.1 | WP_150457769.1 hypothetical protein |
| 82(1) | WP_125084786.1 | WP_125084786.1 hypothetical protein |
| 82(1) | WP_049869165.1 | WP_049869165.1 hypothetical protein |
| 82(1) | WP_101300037.1 | WP_101300037.1 hypothetical protein |
| 82(1) | WP_215185469.1 | WP_215185469.1 hypothetical protein |
| 82(1) | WP_236334340.1 | WP_236334340.1 hypothetical protein |
| 82(1) | WP_042125619.1 | WP_042125619.1 hypothetical protein |
| 82(1) | WP_197937390.1 | WP_197937390.1 hypothetical protein |

| | | |
|---|---|---|
| 83(1) | WP_211082933.1 | WP_211082933.1 recombinase family protein |
| 83(1) | WP_098670175.1 | WP_098670175.1 recombinase family protein |
| 83(1) | WP_097987489.1 | WP_097987489.1 recombinase family protein |
| 83(1) | WP_000504229.1 | WP_000504229.1 recombinase family protein |
| 83(1) | WP_251241790.1 | WP_251241790.1 recombinase family protein |
| 83(1) | WP_000504227.1 | WP_000504227.1 recombinase family protein |
| 83(1) | WP_262787112.1 | WP_262787112.1 recombinase family protein |
| 83(1) | WP_139895508.1 | WP_139895508.1 recombinase family protein |
| 83(1) | WP_253440781.1 | WP_253440781.1 recombinase family protein |
| 83(1) | WP_046392896.1 | WP_046392896.1 recombinase family protein |
| 83(1) | WP_012260841.1 | WP_012260841.1 recombinase family protein |
| 83(2) | WP_061531027.1 | WP_061531027.1 recombinase family protein |
| 83(1) | WP_017560547.1 | WP_017560547.1 recombinase family protein |
| 83(2) | WP_000516940.1 | WP_000516940.1 recombinase family protein |
| 83(13) | WP_001151344.1 | WP_001151344.1 hypothetical protein |
| 83(1) | WP_098768165.1 | WP_098768165.1 recombinase family protein |
| 83(1) | WP_098103682.1 | WP_098103682.1 recombinase family protein |
| 83(1) | WP_075395693.1 | WP_075395693.1 recombinase family protein |
| 83(2) | WP_042512831.1 | WP_042512831.1 recombinase family protein |
| 83(1) | WP_063219633.1 | WP_063219633.1 recombinase family protein |
| 83(9) | WP_000391693.1 | WP_000391693.1 recombinase family protein |
| 83(1) | WP_000504230.1 | WP_000504230.1 recombinase family protein |
| 83(1) | WP_063223284.1 | WP_063223284.1 recombinase family protein |
| 83(2) | WP_087952381.1 | WP_087952381.1 recombinase family protein |
| 83(1) | WP_144492258.1 | WP_144492258.1 recombinase family protein |
| 83(2) | WP_029438095.1 | WP_029438095.1 recombinase family protein |
| 83(1) | WP_000412789.1 | WP_000412789.1 recombinase family protein |
| 83(1) | WP_260489572.1 | WP_260489572.1 recombinase family protein |
| 83(1) | WP_074598754.1 | WP_074598754.1 recombinase family protein |
| 83(1) | WP_242261457.1 | WP_242261457.1 recombinase family protein |
| 83(1) | WP_098250531.1 | WP_098250531.1 DNA recombinase |
| 83(1) | WP_078185400.1 | WP_078185400.1 recombinase family protein |
| 83(1) | WP_035795780.1 | WP_035795780.1 recombinase family protein, partial |
| 83(1) | WP_240379744.1 | WP_240379744.1 recombinase family protein |
| 83(1) | WP_086419724.1 | WP_086419724.1 recombinase family protein |
| 83(1) | WP_033717906.1 | WP_033717906.1 recombinase family protein |
| 83(1) | WP_179034776.1 | WP_179034776.1 recombinase family protein |
| 83(1) | WP_000412791.1 | WP_000412791.1 recombinase family protein |
| 83(1) | WP_098167592.1 | WP_098167592.1 recombinase family protein |
| 83(1) | WP_068696851.1 | WP_068696851.1 recombinase family protein |
| 83(1) | WP_000391696.1 | WP_000391696.1 recombinase family protein |
| 83(1) | WP_000504231.1 | WP_000504231.1 recombinase family protein |
| 83(1) | WP_088054824.1 | WP_088054824.1 recombinase family protein |

| | | |
|---|---|---|
| 84(2) | WP_017692345.1 | WP_017692345.1 hypothetical protein |
| 84(1) | WP_094878485.1 | WP_094878485.1 hypothetical protein |
| 84(1) | WP_074085345.1 | WP_074085345.1 hypothetical protein |
| 84(1) | WP_076242363.1 | WP_076242363.1 hypothetical protein |
| 84(1) | WP_166244095.1 | WP_166244095.1 hypothetical protein |
| 84(1) | WP_076144387.1 | WP_076144387.1 hypothetical protein |
| 84(1) | WP_138755929.1 | WP_138755929.1 hypothetical protein |
| 84(1) | WP_253440793.1 | WP_253440793.1 hypothetical protein |
| 84(1) | WP_076279373.1 | WP_076279373.1 hypothetical protein |
| 84(2) | WP_076104585.1 | WP_076104585.1 hypothetical protein |
| 84(1) | WP_019912138.1 | WP_019912138.1 hypothetical protein |
| 84(1) | WP_238187873.1 | WP_238187873.1 hypothetical protein |
| 84(1) | WP_114298759.1 | WP_114298759.1 hypothetical protein |
| 84(1) | WP_076309118.1 | WP_076309118.1 hypothetical protein |
| 84(1) | WP_068782418.1 | WP_068782418.1 F0F1-type ATP synthase |
| 84(1) | WP_076303957.1 | WP_076303957.1 hypothetical protein |
| 84(1) | WP_182301848.1 | WP_182301848.1 hypothetical protein |
| 84(1) | WP_134750630.1 | WP_134750630.1 hypothetical protein |
| 84(1) | WP_074108288.1 | WP_074108288.1 hypothetical protein |
| 84(1) | WP_209872713.1 | WP_209872713.1 hypothetical protein |
| 84(1) | WP_068536362.1 | WP_068536362.1 hypothetical protein |
| 84(1) | WP_127194971.1 | WP_127194971.1 hypothetical protein |
| 84(1) | WP_136372947.1 | WP_136372947.1 hypothetical protein |
| 84(1) | WP_145406509.1 | WP_145406509.1 hypothetical protein |
| 84(1) | WP_308723526.1 | WP_308723526.1 hypothetical protein |
| 84(2) | WP_145153658.1 | WP_145153658.1 hypothetical protein |
| 84(2) | WP_219872467.1 | WP_219872467.1 hypothetical protein |
| 84(1) | WP_193726777.1 | WP_193726777.1 hypothetical protein |
| 84(1) | WP_027088417.1 | WP_027088417.1 hypothetical protein |
| 84(1) | WP_089522647.1 | WP_089522647.1 hypothetical protein |
| 84(1) | WP_184183017.1 | WP_184183017.1 hypothetical protein |
| 84(1) | WP_111273174.1 | WP_111273174.1 hypothetical protein |
| 84(1) | WP_127570446.1 | WP_127570446.1 hypothetical protein |
| 84(1) | WP_189018582.1 | WP_189018582.1 hypothetical protein, partial |
| 84(1) | WP_110899371.1 | WP_110899371.1 hypothetical protein |
| 84(1) | WP_039306319.1 | WP_039306319.1 hypothetical protein |
| 84(1) | WP_076301095.1 | WP_076301095.1 hypothetical protein |
| 84(1) | WP_210087116.1 | WP_210087116.1 hypothetical protein |
| 84(1) | WP_293755482.1 | WP_293755482.1 hypothetical protein |
| 84(1) | WP_212695805.1 | WP_212695805.1 hypothetical protein |
| 84(1) | WP_150131506.1 | WP_150131506.1 hypothetical protein |
| 84(1) | WP_068696907.1 | WP_068696907.1 hypothetical protein |
| 84(1) | WP_239617485.1 | WP_239617485.1 hypothetical protein |
| 84(1) | WP_212981450.1 | WP_212981450.1 hypothetical protein |
| 84(1) | WP_120112282.1 | WP_120112282.1 hypothetical protein |
| 84(2) | WP_216518264.1 | WP_216518264.1 hypothetical protein |

| | | |
|---|---|---|
| 84(1) | WP_094870411.1 | WP_094870411.1 hypothetical protein |
| 84(1) | WP_076210965.1 | WP_076210965.1 hypothetical protein |
| 84(1) | WP_307199918.1 | WP_307199918.1 hypothetical protein |
| 84(1) | WP_055105804.1 | WP_055105804.1 hypothetical protein |
| 84(1) | WP_110931530.1 | WP_110931530.1 hypothetical protein |
| 84(1) | WP_156123798.1 | WP_156123798.1 hypothetical protein |
| 84(1) | WP_154118019.1 | WP_154118019.1 hypothetical protein |
| 84(1) | WP_076287007.1 | WP_076287007.1 hypothetical protein |
| 84(1) | WP_211023041.1 | WP_211023041.1 hypothetical protein |
| 84(1) | WP_076219084.1 | WP_076219084.1 hypothetical protein |
| 84(1) | WP_280788628.1 | WP_280788628.1 hypothetical protein |
| 84(1) | WP_024632690.1 | WP_024632690.1 hypothetical protein |
| 84(1) | WP_157763987.1 | WP_157763987.1 hypothetical protein |
| 84(1) | WP_307601007.1 | WP_307601007.1 hypothetical protein |
| 84(3) | WP_076135788.1 | WP_076135788.1 hypothetical protein |

| | | |
|---|---|---|
| 85(2) | WP_219872462.1 | WP_219872462.1 hypothetical protein |
| 85(1) | WP_024632694.1 | WP_024632694.1 hypothetical protein |
| 85(1) | WP_019912146.1 | WP_019912146.1 hypothetical protein |
| 85(1) | WP_184183022.1 | WP_184183022.1 hypothetical protein |
| 85(2) | WP_150364189.1 | WP_150364189.1 hypothetical protein |
| 85(1) | WP_042211002.1 | WP_042211002.1 hypothetical protein |
| 85(1) | WP_074108292.1 | WP_074108292.1 hypothetical protein |
| 85(1) | WP_209872730.1 | WP_209872730.1 hypothetical protein |
| 85(1) | WP_113672714.1 | WP_113672714.1 hypothetical protein |
| 85(1) | WP_068536353.1 | WP_068536353.1 hypothetical protein |
| 85(1) | WP_055105808.1 | WP_055105808.1 hypothetical protein |
| 85(1) | WP_076279378.1 | WP_076279378.1 hypothetical protein |
| 85(1) | WP_145153670.1 | WP_145153670.1 hypothetical protein |
| 85(1) | WP_189018591.1 | WP_189018591.1 hypothetical protein |
| 85(1) | WP_111273177.1 | WP_111273177.1 hypothetical protein |
| 85(1) | WP_308723530.1 | WP_308723530.1 hypothetical protein |
| 85(1) | WP_111503210.1 | WP_111503210.1 hypothetical protein |
| 85(1) | WP_127570453.1 | WP_127570453.1 hypothetical protein |
| 85(1) | WP_114298755.1 | WP_114298755.1 hypothetical protein |
| 85(1) | WP_089522642.1 | WP_089522642.1 hypothetical protein |
| 85(1) | WP_273321529.1 | WP_273321529.1 hypothetical protein |
| 85(1) | WP_166244086.1 | WP_166244086.1 hypothetical protein |
| 85(1) | WP_154118025.1 | WP_154118025.1 hypothetical protein |
| 85(1) | WP_253440804.1 | WP_253440804.1 hypothetical protein |
| 85(1) | WP_169733292.1 | WP_169733292.1 hypothetical protein |
| 85(2) | WP_076135797.1 | WP_076135797.1 hypothetical protein |
| 85(2) | WP_094878483.1 | WP_094878483.1 hypothetical protein |
| 85(1) | WP_239617492.1 | WP_239617492.1 hypothetical protein |
| 85(2) | WP_042134284.1 | WP_042134284.1 hypothetical protein |
| 85(2) | WP_216518270.1 | WP_216518270.1 hypothetical protein |
| 85(1) | WP_138755935.1 | WP_138755935.1 hypothetical protein |
| 85(1) | WP_076210957.1 | WP_076210957.1 hypothetical protein |
| 85(2) | WP_076104579.1 | WP_076104579.1 hypothetical protein |
| 85(1) | WP_210087120.1 | WP_210087120.1 hypothetical protein |
| 85(2) | WP_076219078.1 | WP_076219078.1 hypothetical protein |
| 85(1) | WP_248063427.1 | WP_248063427.1 hypothetical protein |
| 85(1) | WP_293755498.1 | WP_293755498.1 hypothetical protein |
| 85(1) | WP_110931526.1 | WP_110931526.1 hypothetical protein |
| 85(1) | WP_152392580.1 | WP_152392580.1 hypothetical protein |
| 85(1) | WP_127194966.1 | WP_127194966.1 hypothetical protein |
| 85(1) | WP_017692340.1 | WP_017692340.1 hypothetical protein |
| 85(1) | WP_039871658.1 | WP_039871658.1 hypothetical protein |
| 85(1) | WP_212693248.1 | WP_212693248.1 hypothetical protein |
| 85(1) | WP_110899376.1 | WP_110899376.1 hypothetical protein |
| 85(1) | WP_076208268.1 | WP_076208268.1 hypothetical protein |
| 85(1) | WP_068696914.1 | WP_068696914.1 hypothetical protein |
| 85(1) | WP_042203653.1 | WP_042203653.1 hypothetical protein |
| 85(1) | WP_094870405.1 | WP_094870405.1 hypothetical protein |
| 85(1) | WP_068782422.1 | WP_068782422.1 hypothetical protein |
| 85(1) | WP_212695798.1 | WP_212695798.1 hypothetical protein |
| 85(1) | WP_076309115.1 | WP_076309115.1 hypothetical protein |
| 85(1) | WP_039306337.1 | WP_039306337.1 hypothetical protein |
| 85(1) | WP_099517098.1 | WP_099517098.1 hypothetical protein |
| 85(2) | WP_281813126.1 | WP_281813126.1 hypothetical protein |
| 85(2) | WP_076287001.1 | WP_076287001.1 hypothetical protein |
| 85(1) | WP_074085351.1 | WP_074085351.1 hypothetical protein |
| 85(1) | WP_120112295.1 | WP_120112295.1 hypothetical protein |
| 85(1) | WP_307199925.1 | WP_307199925.1 hypothetical protein |

| | | |
|---|---|---|
| 86(1) | WP_199613895.1 | WP_199613895.1 hypothetical protein |
| 86(1) | WP_125084771.1 | WP_125084771.1 hypothetical protein |
| 86(1) | WP_094870528.1 | WP_094870528.1 hypothetical protein |
| 86(1) | WP_238655354.1 | WP_238655354.1 hypothetical protein |
| 86(1) | WP_052414525.1 | WP_052414525.1 hypothetical protein |
| 86(1) | WP_194434081.1 | WP_194434081.1 hypothetical protein |
| 86(1) | WP_249897323.1 | WP_249897323.1 hypothetical protein |
| 86(1) | WP_251417672.1 | WP_251417672.1 hypothetical protein |
| 86(1) | WP_076271938.1 | WP_076271938.1 hypothetical protein |
| 86(1) | WP_208414541.1 | WP_208414541.1 hypothetical protein |
| 86(1) | WP_137698096.1 | WP_137698096.1 hypothetical protein |
| 86(1) | WP_223068030.1 | WP_223068030.1 hypothetical protein |
| 86(1) | WP_280787392.1 | WP_280787392.1 hypothetical protein |
| 86(1) | WP_036944546.1 | WP_036944546.1 hypothetical protein |
| 86(1) | WP_283927238.1 | WP_283927238.1 hypothetical protein |
| 86(1) | WP_215185483.1 | WP_215185483.1 hypothetical protein |
| 86(1) | WP_169028886.1 | WP_169028886.1 hypothetical protein |
| 86(1) | WP_207654803.1 | WP_207654803.1 hypothetical protein |
| 86(1) | WP_148450942.1 | WP_148450942.1 hypothetical protein |
| 86(1) | WP_076103992.1 | WP_076103992.1 hypothetical protein |
| 86(1) | WP_076101577.1 | WP_076101577.1 hypothetical protein |
| 86(1) | WP_251596696.1 | WP_251596696.1 hypothetical protein |
| 86(1) | WP_157054019.1 | WP_157054019.1 hypothetical protein |
| 86(2) | WP_076281069.1 | WP_076281069.1 hypothetical protein |
| 86(1) | WP_076157826.1 | WP_076157826.1 hypothetical protein |
| 86(1) | WP_119601478.1 | WP_119601478.1 hypothetical protein |
| 86(1) | WP_105368175.1 | WP_105368175.1 hypothetical protein |
| 86(1) | WP_076297590.1 | WP_076297590.1 hypothetical protein |
| 86(2) | WP_101300055.1 | WP_101300055.1 hypothetical protein |

| | | |
|---|---|---|
| 86(1) | WP_211022784.1 | WP_211022784.1 hypothetical protein |
| 86(1) | WP_189031024.1 | WP_189031024.1 hypothetical protein |
| 86(1) | WP_290407040.1 | WP_290407040.1 hypothetical protein |
| 86(1) | WP_155609747.1 | WP_155609747.1 hypothetical protein |
| 86(1) | WP_257443623.1 | WP_257443623.1 hypothetical protein |
| 86(1) | WP_236334314.1 | WP_236334314.1 hypothetical protein |
| 86(1) | WP_159081752.1 | WP_159081752.1 hypothetical protein |
| 86(1) | WP_280752671.1 | WP_280752671.1 hypothetical protein |
| 86(1) | WP_232101760.1 | WP_232101760.1 hypothetical protein |
| 86(1) | WP_094876483.1 | WP_094876483.1 hypothetical protein |
| 86(1) | WP_165097633.1 | WP_165097633.1 hypothetical protein |
| 86(1) | WP_283910931.1 | WP_283910931.1 hypothetical protein |
| 86(1) | WP_120462946.1 | WP_120462946.1 hypothetical protein |
| 86(1) | WP_162616103.1 | WP_162616103.1 hypothetical protein |
| 86(1) | WP_076224116.1 | WP_076224116.1 hypothetical protein |
| 86(1) | WP_036944543.1 | WP_036944543.1 hypothetical protein |
| 86(1) | WP_195571379.1 | WP_195571379.1 hypothetical protein |
| 86(1) | WP_128631275.1 | WP_128631275.1 hypothetical protein |
| 86(1) | WP_137698097.1 | WP_137698097.1 hypothetical protein |
| 86(1) | WP_055109365.1 | WP_055109365.1 hypothetical protein |
| 86(1) | WP_036702408.1 | WP_036702408.1 hypothetical protein |
| 86(1) | WP_150457758.1 | WP_150457758.1 hypothetical protein |
| 86(1) | WP_052096894.1 | WP_052096894.1 hypothetical protein |
| 86(1) | WP_051491339.1 | WP_051491339.1 hypothetical protein |
| 86(1) | WP_141097596.1 | WP_141097596.1 hypothetical protein |
| 86(1) | WP_137698105.1 | WP_137698105.1 hypothetical protein |
| 86(1) | WP_076141380.1 | WP_076141380.1 hypothetical protein |
| 86(1) | WP_244723762.1 | WP_244723762.1 hypothetical protein |
| 86(1) | WP_085499078.1 | WP_085499078.1 hypothetical protein |
| 86(1) | WP_076119223.1 | WP_076119223.1 hypothetical protein |
| 86(1) | WP_013658704.1 | WP_013658704.1 hypothetical protein |
| 86(1) | WP_213506883.1 | WP_213506883.1 hypothetical protein |
| 86(1) | WP_213592852.1 | WP_213592852.1 hypothetical protein |
| 86(1) | WP_101300065.1 | WP_101300065.1 hypothetical protein |
| 86(1) | WP_110934062.1 | WP_110934062.1 hypothetical protein |
| 86(1) | WP_197937403.1 | WP_197937403.1 hypothetical protein |
| 87(1) | WP_158329528.1 | WP_158329528.1 thiamine phosphate synthase |
| 87(1) | WP_022586781.1 | WP_022586781.1 thiamine phosphate synthase |
| 87(1) | WP_239209287.1 | WP_239209287.1 thiamine phosphate synthase |
| 87(1) | WP_096886730.1 | WP_096886730.1 thiamine phosphate synthase |
| 87(1) | WP_301804627.1 | WP_301804627.1 thiamine phosphate synthase |
| 87(1) | WP_258425836.1 | WP_258425836.1 thiamine phosphate synthase |
| 87(1) | WP_003340532.1 | WP_003340532.1 thiamine phosphate synthase |
| 87(1) | WP_258418578.1 | WP_258418578.1 thiamine phosphate synthase |
| 87(2) | WP_121474471.1 | WP_121474471.1 thiamine phosphate synthase |
| 87(1) | WP_163245174.1 | WP_163245174.1 thiamine phosphate synthase |
| 87(1) | WP_277548453.1 | WP_277548453.1 thiamine phosphate synthase |
| 87(1) | WP_096886873.1 | WP_096886873.1 thiamine phosphate synthase |
| 87(1) | WP_258433012.1 | WP_258433012.1 thiamine phosphate synthase |
| 87(2) | WP_158326754.1 | WP_158326754.1 thiamine phosphate synthase |
| 87(1) | WP_232699369.1 | WP_232699369.1 thiamine phosphate synthase |
| 87(2) | WP_158326756.1 | WP_158326756.1 thiamine phosphate synthase |
| 87(1) | WP_181747857.1 | WP_181747857.1 thiamine phosphate synthase |
| 87(5) | WP_094699666.1 | WP_094699666.1 thiamine phosphate synthase |
| 87(1) | WP_283952957.1 | WP_283952957.1 thiamine phosphate synthase |
| 87(1) | WP_116334073.1 | WP_116334073.1 thiamine phosphate synthase |
| 87(1) | WP_068793958.1 | WP_068793958.1 thiamine phosphate synthase |
| 87(1) | WP_104030450.1 | WP_104030450.1 thiamine phosphate synthase |
| 87(1) | WP_003340524.1 | WP_003340524.1 thiamine phosphate synthase |
| 87(1) | WP_181747853.1 | WP_181747853.1 thiamine phosphate synthase |
| 87(1) | WP_113755289.1 | WP_113755289.1 thiamine phosphate synthase |
| 87(1) | WP_232699371.1 | WP_232699371.1 thiamine phosphate synthase |
| 87(1) | WP_188535044.1 | WP_188535044.1 thiamine phosphate synthase |
| 87(1) | WP_212930968.1 | WP_212930968.1 thiamine phosphate synthase |
| 87(1) | WP_158329530.1 | WP_158329530.1 thiamine phosphate synthase |
| 87(2) | WP_094699663.1 | WP_094699663.1 thiamine phosphate synthase |
| 87(1) | WP_104065524.1 | WP_104065524.1 thiamine phosphate synthase |
| 87(1) | WP_104148788.1 | WP_104148788.1 thiamine phosphate synthase |
| 87(1) | WP_068793923.1 | WP_068793923.1 thiamine phosphate synthase |
| 87(2) | WP_104148789.1 | WP_104148789.1 thiamine phosphate synthase |
| 87(2) | WP_119732870.1 | WP_119732870.1 thiamine phosphate synthase |
| 87(1) | WP_278431119.1 | WP_278431119.1 thiamine phosphate synthase |
| 87(1) | WP_101668484.1 | WP_101668484.1 thiamine phosphate synthase |
| 87(1) | WP_258417124.1 | WP_258417124.1 thiamine phosphate synthase |
| 87(1) | WP_258425833.1 | WP_258425833.1 thiamine phosphate synthase |
| 87(2) | WP_119732867.1 | WP_119732867.1 thiamine phosphate synthase |
| 87(2) | WP_018670533.1 | WP_018670533.1 thiamine phosphate synthase |
| 87(1) | WP_121474871.1 | WP_121474871.1 thiamine phosphate synthase |
| 87(1) | WP_064019329.1 | WP_064019329.1 thiamine phosphate synthase |
| 87(1) | WP_258417123.1 | WP_258417123.1 thiamine phosphate synthase |
| 87(1) | WP_212930965.1 | WP_212930965.1 thiamine phosphate synthase |
| 87(1) | WP_104030451.1 | WP_104030451.1 thiamine phosphate synthase |
| 87(1) | WP_199794086.1 | WP_199794086.1 thiamine phosphate synthase |
| 87(1) | WP_101668482.1 | WP_101668482.1 thiamine phosphate synthase |
| 87(1) | WP_113755287.1 | WP_113755287.1 thiamine phosphate synthase |
| 87(2) | WP_163245172.1 | WP_163245172.1 thiamine phosphate synthase |
| 87(1) | WP_301804636.1 | WP_301804636.1 thiamine phosphate synthase |
| 87(1) | WP_031309867.1 | WP_031309867.1 thiamine phosphate synthase |
| 87(2) | WP_197247497.1 | WP_197247497.1 thiamine phosphate synthase |
| 88(1) | WP_042134280.1 | WP_042134280.1 hypothetical protein |
| 88(1) | WP_248063423.1 | WP_248063423.1 hypothetical protein |
| 88(2) | WP_074085353.1 | WP_074085353.1 hypothetical protein |
| 88(1) | WP_042203651.1 | WP_042203651.1 hypothetical protein |
| 88(1) | WP_280788630.1 | WP_280788630.1 hypothetical protein |
| 88(1) | WP_068536350.1 | WP_068536350.1 hypothetical protein |
| 88(1) | WP_099517100.1 | WP_099517100.1 hypothetical protein |
| 88(1) | WP_212693246.1 | WP_212693246.1 hypothetical protein |
| 88(1) | WP_166244084.1 | WP_166244084.1 hypothetical protein |
| 88(3) | WP_076219077.1 | WP_076219077.1 hypothetical protein |
| 88(1) | WP_152679865.1 | WP_152679865.1 hypothetical protein |
| 88(2) | WP_076287000.1 | WP_076287000.1 hypothetical protein |
| 88(1) | WP_143767394.1 | WP_143767394.1 hypothetical protein |

88(3) WP_017692338.1 WP_017692338.1 hypothetical protein
88(1) WP_169823440.1 WP_169823440.1 hypothetical protein
88(1) WP_209872736.1 WP_209872736.1 hypothetical protein
88(1) WP_144025952.1 WP_144025952.1 hypothetical protein
88(2) WP_143766715.1 WP_143766715.1 hypothetical protein
88(2) WP_113672712.1 WP_113672712.1 hypothetical protein
88(1) WP_154985816.1 WP_154985816.1 hypothetical protein
88(2) WP_281813121.1 WP_281813121.1 hypothetical protein
88(1) WP_127194964.1 WP_127194964.1 hypothetical protein
88(1) WP_307199930.1 WP_307199930.1 hypothetical protein
88(1) WP_210087122.1 WP_210087122.1 AAA family ATPase
88(1) WP_156118549.1 WP_156118549.1 hypothetical protein
88(1) WP_120112299.1 WP_120112299.1 hypothetical protein
88(1) WP_189018595.1 WP_189018595.1 hypothetical protein
88(1) WP_145153676.1 WP_145153676.1 hypothetical protein
88(1) WP_110899378.1 WP_110899378.1 hypothetical protein
88(1) WP_152392579.1 WP_152392579.1 hypothetical protein
88(1) WP_289376834.1 WP_289376834.1 hypothetical protein
88(3) WP_076104578.1 WP_076104578.1 hypothetical protein
88(1) WP_150364187.1 WP_150364187.1 hypothetical protein
88(1) WP_089522640.1 WP_089522640.1 hypothetical protein
88(1) WP_076279379.1 WP_076279379.1 hypothetical protein
88(1) WP_239617494.1 WP_239617494.1 hypothetical protein
88(1) WP_138755937.1 WP_138755937.1 hypothetical protein
88(1) WP_229521100.1 WP_229521100.1 hypothetical protein
88(2) WP_184183041.1 WP_184183041.1 hypothetical protein
88(1) WP_293755503.1 WP_293755503.1 hypothetical protein
88(1) WP_229521538.1 WP_229521538.1 hypothetical protein
88(2) WP_094870403.1 WP_094870403.1 hypothetical protein
88(1) WP_114298754.1 WP_114298754.1 hypothetical protein
88(1) WP_027088408.1 WP_027088408.1 hypothetical protein
88(1) WP_111273179.1 WP_111273179.1 hypothetical protein
88(1) WP_024632696.1 WP_024632696.1 hypothetical protein
88(1) WP_167549523.1 WP_167549523.1 hypothetical protein
88(1) WP_039871657.1 WP_039871657.1 hypothetical protein
88(2) WP_219872460.1 WP_219872460.1 hypothetical protein
88(1) WP_253440808.1 WP_253440808.1 hypothetical protein
88(1) WP_304942554.1 WP_304942554.1 hypothetical protein
88(1) WP_212695796.1 WP_212695796.1 hypothetical protein
88(1) WP_019912148.1 WP_019912148.1 hypothetical protein

89(1) WP_123067122.1 WP_123067122.1 hypothetical protein
89(1) WP_257219296.1 WP_257219296.1 DUF202 domain-containing protein
89(1) WP_272035336.1 WP_272035336.1 DUF202 domain-containing protein
89(1) WP_090949174.1 WP_090949174.1 DUF202 domain-containing protein
89(1) WP_235431543.1 WP_235431543.1 DUF202 domain-containing protein
89(1) WP_282321816.1 WP_282321816.1 DUF202 domain-containing protein
89(1) WP_192266472.1 WP_192266472.1 DUF202 domain-containing protein
89(5) WP_063563228.1 WP_063563228.1 DUF202 domain-containing protein
89(2) WP_109999897.1 WP_109999897.1 DUF202 domain-containing protein
89(1) WP_254790064.1 WP_254790064.1 DUF202 domain-containing protein
89(1) WP_223869120.1 WP_223869120.1 DUF202 domain-containing protein
89(1) WP_307537570.1 WP_307537570.1 DUF202 domain-containing protein
89(1) WP_256704103.1 WP_256704103.1 DUF202 domain-containing protein
89(1) WP_256975167.1 WP_256975167.1 DUF202 domain-containing protein
89(1) WP_076330727.1 WP_076330727.1 DUF202 domain-containing protein
89(3) WP_029880936.1 WP_029880936.1 DUF202 domain-containing protein
89(1) WP_244898322.1 WP_244898322.1 DUF202 domain-containing protein
89(1) WP_110892957.1 WP_110892957.1 DUF202 domain-containing protein
89(1) WP_193721882.1 WP_193721882.1 DUF202 domain-containing protein
89(2) WP_145146967.1 WP_145146967.1 DUF202 domain-containing protein
89(2) WP_056700806.1 WP_056700806.1 DUF202 domain-containing protein
89(1) WP_235200748.1 WP_235200748.1 DUF202 domain-containing protein
89(1) WP_293202256.1 WP_293202256.1 DUF202 domain-containing protein
89(1) WP_289392078.1 WP_289392078.1 DUF202 domain-containing protein
89(1) WP_240661817.1 WP_240661817.1 DUF202 domain-containing protein
89(1) WP_181754682.1 WP_181754682.1 DUF202 domain-containing protein
89(1) WP_110822557.1 WP_110822557.1 DUF202 domain-containing protein
89(1) WP_062833887.1 WP_062833887.1 DUF202 domain-containing protein
89(1) WP_095287653.1 WP_095287653.1 DUF202 domain-containing protein
89(1) WP_188593168.1 WP_188593168.1 DUF202 domain-containing protein
89(1) WP_145046467.1 WP_145046467.1 DUF202 domain-containing protein
89(1) WP_090921040.1 WP_090921040.1 DUF202 domain-containing protein
89(1) WP_145329914.1 WP_145329914.1 DUF202 domain-containing protein
89(5) WP_017689063.1 WP_017689063.1 DUF202 domain-containing protein
89(1) WP_127536190.1 WP_127536190.1 DUF202 domain-containing protein
89(2) WP_145412709.1 WP_145412709.1 DUF202 domain-containing protein
89(1) WP_095358542.1 WP_095358542.1 DUF202 domain-containing protein
89(2) WP_293017544.1 WP_293017544.1 DUF202 domain-containing protein
89(1) WP_240767978.1 WP_240767978.1 DUF202 domain-containing protein
89(1) WP_113052560.1 WP_113052560.1 DUF202 domain-containing protein
89(1) WP_260865825.1 WP_260865825.1 hypothetical protein
89(1) WP_153978071.1 WP_153978071.1 hypothetical protein
89(2) WP_090896205.1 WP_090896205.1 DUF202 domain-containing protein
89(1) WP_145318677.1 WP_145318677.1 DUF202 domain-containing protein
89(1) WP_235193787.1 WP_235193787.1 DUF202 domain-containing protein
89(1) WP_243391412.1 WP_243391412.1 DUF202 domain-containing protein
89(1) WP_247902251.1 WP_247902251.1 DUF202 domain-containing protein
89(1) WP_062326155.1 WP_062326155.1 DUF202 domain-containing protein
89(1) WP_053783893.1 WP_053783893.1 DUF202 domain-containing protein

90(4) WP_003721677.1 WP_003721677.1 replicative DNA helicase
90(15) WP_221633040.1 WP_221633040.1 replicative DNA helicase
90(43) WP_003718135.1 WP_003718135.1 replicative DNA helicase
90(2) WP_228478846.1 WP_228478846.1 replicative DNA helicase

91(1) WP_033100011.1 WP_033100011.1 GntP family permease
91(5) WP_000424326.1 WP_000424326.1 GntP family permease
91(1) WP_088058550.1 WP_088058550.1 GntP family permease
91(1) WP_071724738.1 WP_071724738.1 GntP family permease
91(1) WP_262729438.1 WP_262729438.1 GntP family permease
91(1) WP_199695258.1 WP_199695258.1 GntP family permease
91(1) WP_061688412.1 WP_061688412.1 GntP family permease
91(1) WP_198056465.1 WP_198056465.1 GntP family permease

| | | |
|---|---|---|
| 91(1) | WP_270731498.1 | WP_270731498.1 GntP family permease |
| 91(1) | WP_098088228.1 | WP_098088228.1 GntP family permease |
| 91(1) | WP_098299072.1 | WP_098299072.1 GntP family permease |
| 91(1) | WP_071719970.1 | WP_071719970.1 GntP family permease |
| 91(2) | WP_048552327.1 | WP_048552327.1 GntP family permease |
| 91(1) | WP_097844610.1 | WP_097844610.1 GntP family permease |
| 91(1) | WP_087956303.1 | WP_087956303.1 GntP family permease |
| 91(2) | WP_070805909.1 | WP_070805909.1 GntP family permease |
| 91(1) | WP_147786231.1 | WP_147786231.1 GntP family permease |
| 91(4) | WP_000424331.1 | WP_000424331.1 GntP family permease |
| 91(1) | WP_000424304.1 | WP_000424304.1 GntP family permease |
| 91(1) | WP_060630657.1 | WP_060630657.1 GntP family permease |
| 91(1) | WP_198059699.1 | WP_198059699.1 GntP family permease |
| 91(4) | WP_088313251.1 | WP_088313251.1 GntP family permease |
| 91(1) | WP_144488222.1 | WP_144488222.1 GntP family permease |
| 91(1) | WP_098214715.1 | WP_098214715.1 GntP family permease |
| 91(6) | WP_087982563.1 | WP_087982563.1 GntP family permease |
| 91(1) | WP_061535950.1 | WP_061535950.1 GntP family permease |
| 91(3) | WP_000424306.1 | WP_000424306.1 GntP family permease |
| 91(1) | WP_098266660.1 | WP_098266660.1 GntP family permease |
| 91(1) | WP_263324542.1 | WP_263324542.1 GntP family permease |
| 91(3) | WP_097911830.1 | WP_097911830.1 GntP family permease |
| 91(3) | WP_069355517.1 | WP_069355517.1 GntP family permease |
| 91(1) | WP_002158030.1 | WP_002158030.1 GntP family permease |
| 91(1) | WP_098856778.1 | WP_098856778.1 GntP family permease |
| 91(1) | WP_088609122.1 | WP_088609122.1 GntP family permease |
| 91(3) | WP_078181478.1 | WP_078181478.1 GntP family permease |
| 91(1) | WP_061658108.1 | WP_061658108.1 GntP family permease |
| 92(1) | WP_210087124.1 | WP_210087124.1 SAF domain-containing protein |
| 92(1) | WP_212693245.1 | WP_212693245.1 SAF domain-containing protein |
| 92(1) | WP_209872742.1 | WP_209872742.1 SAF domain-containing protein |
| 92(1) | WP_293755508.1 | WP_293755508.1 SAF domain-containing protein |
| 92(1) | WP_289376833.1 | WP_289376833.1 SAF domain-containing protein |
| 92(6) | WP_076208263.1 | WP_076208263.1 SAF domain-containing protein |
| 92(1) | WP_110899380.1 | WP_110899380.1 SAF domain-containing protein |
| 92(1) | WP_150364185.1 | WP_150364185.1 SAF domain-containing protein |
| 92(1) | WP_076279381.1 | WP_076279381.1 SAF domain-containing protein |
| 92(1) | WP_273321534.1 | WP_273321534.1 SAF domain-containing protein |
| 92(1) | WP_076309114.1 | WP_076309114.1 SAF domain-containing protein |
| 92(1) | WP_042210998.1 | WP_042210998.1 SAF domain-containing protein |
| 92(1) | WP_304942555.1 | WP_304942555.1 SAF domain-containing protein |
| 92(1) | WP_076210950.1 | WP_076210950.1 SAF domain-containing protein |
| 92(1) | WP_039871655.1 | WP_039871655.1 SAF domain-containing protein |
| 92(1) | WP_145153681.1 | WP_145153681.1 SAF domain-containing protein |
| 92(1) | WP_068782425.1 | WP_068782425.1 SAF domain-containing protein |
| 92(1) | WP_114298752.1 | WP_114298752.1 hypothetical protein |
| 92(1) | WP_017692336.1 | WP_017692336.1 SAF domain-containing protein |
| 92(2) | WP_076135803.1 | WP_076135803.1 SAF domain-containing protein |
| 92(1) | WP_189018599.1 | WP_189018599.1 SAF domain-containing protein |
| 92(1) | WP_042203649.1 | WP_042203649.1 SAF domain-containing protein |
| 92(1) | WP_253440812.1 | WP_253440812.1 SAF domain-containing protein |
| 92(1) | WP_145406525.1 | WP_145406525.1 SAF domain-containing protein |
| 92(1) | WP_024632698.1 | WP_024632698.1 SAF domain-containing protein |
| 92(1) | WP_184183058.1 | WP_184183058.1 SAF domain-containing protein |
| 92(2) | WP_281813116.1 | WP_281813116.1 SAF domain-containing protein |
| 92(1) | WP_154118029.1 | WP_154118029.1 SAF domain-containing protein |
| 92(1) | WP_068536346.1 | WP_068536346.1 SAF domain-containing protein |
| 92(2) | WP_042134274.1 | WP_042134274.1 SAF domain-containing protein |
| 92(1) | WP_307199934.1 | WP_307199934.1 SAF domain-containing protein |
| 92(1) | WP_150131507.1 | WP_150131507.1 hypothetical protein |
| 92(1) | WP_113672710.1 | WP_113672710.1 SAF domain-containing protein |
| 92(1) | WP_167434149.1 | WP_167434149.1 SAF domain-containing protein |
| 92(1) | WP_127194962.1 | WP_127194962.1 hypothetical protein |
| 92(1) | WP_154985814.1 | WP_154985814.1 SAF domain-containing protein |
| 92(1) | WP_019912150.1 | WP_019912150.1 SAF domain-containing protein |
| 92(2) | WP_219872458.1 | WP_219872458.1 SAF domain-containing protein |
| 92(1) | WP_212695794.1 | WP_212695794.1 SAF domain-containing protein |
| 92(1) | WP_074085355.1 | WP_074085355.1 SAF domain-containing protein |
| 92(1) | WP_248063419.1 | WP_248063419.1 SAF domain-containing protein |
| 92(1) | WP_211082918.1 | WP_211082918.1 flagella basal body P-ring formation protein FlgA |
| 92(1) | WP_166244080.1 | WP_166244080.1 SAF domain-containing protein |
| 92(3) | WP_076104576.1 | WP_076104576.1 SAF domain-containing protein |
| 92(1) | WP_089522638.1 | WP_089522638.1 SAF domain-containing protein |
| 92(1) | WP_039306348.1 | WP_039306348.1 SAF domain-containing protein |
| 92(1) | WP_193726776.1 | WP_193726776.1 hypothetical protein |
| 92(1) | WP_074108294.1 | WP_074108294.1 SAF domain-containing protein |
| 92(2) | WP_076219075.1 | WP_076219075.1 SAF domain-containing protein |
| 93(1) | WP_076322118.1 | WP_076322118.1 FMN-dependent NADH-azoreductase |
| 93(1) | WP_200633294.1 | WP_200633294.1 FMN-dependent NADH-azoreductase |
| 93(2) | WP_071220199.1 | WP_071220199.1 FMN-dependent NADH-azoreductase |
| 93(1) | WP_145156801.1 | WP_145156801.1 FMN-dependent NADH-azoreductase |
| 93(1) | WP_183578430.1 | WP_183578430.1 FMN-dependent NADH-azoreductase |
| 93(1) | WP_096774457.1 | WP_096774457.1 FMN-dependent NADH-azoreductase |
| 93(1) | WP_163872753.1 | WP_163872753.1 FMN-dependent NADH-azoreductase |
| 93(1) | WP_280753908.1 | WP_280753908.1 FMN-dependent NADH-azoreductase |
| 93(1) | WP_077567025.1 | WP_077567025.1 FMN-dependent NADH-azoreductase |
| 93(1) | WP_098746877.1 | WP_098746877.1 FMN-dependent NADH-azoreductase |
| 93(3) | WP_006211211.1 | WP_006211211.1 FMN-dependent NADH-azoreductase |
| 93(1) | WP_028405555.1 | WP_028405555.1 FMN-dependent NADH-azoreductase |
| 93(3) | WP_194543501.1 | WP_194543501.1 FMN-dependent NADH-azoreductase |
| 93(1) | WP_015736248.1 | WP_015736248.1 FMN-dependent NADH-azoreductase |
| 93(1) | WP_263880222.1 | WP_263880222.1 FMN-dependent NADH-azoreductase |
| 93(1) | WP_145021750.1 | WP_145021750.1 FMN-dependent NADH-azoreductase |
| 93(1) | WP_053492733.1 | WP_053492733.1 FMN-dependent NADH-azoreductase |
| 93(1) | WP_042462794.1 | WP_042462794.1 FMN-dependent NADH-azoreductase |
| 93(2) | WP_210095108.1 | WP_210095108.1 FMN-dependent NADH-azoreductase |
| 93(1) | WP_063479995.1 | WP_063479995.1 FMN-dependent NADH-azoreductase |
| 93(1) | WP_023957705.1 | WP_023957705.1 FMN-dependent NADH-azoreductase |
| 93(1) | WP_127592431.1 | WP_127592431.1 FMN-dependent NADH-azoreductase |
| 93(1) | WP_007128637.1 | WP_007128637.1 FMN-dependent NADH-azoreductase |
| 93(2) | WP_119847548.1 | WP_119847548.1 FMN-dependent NADH-azoreductase |
| 93(1) | WP_094593868.1 | WP_094593868.1 FMN-dependent NADH-azoreductase |

| | | |
|---|---|---|
| 93(1) | WP_116231181.1 | WP_116231181.1 FMN-dependent NADH-azoreductase |
| 93(1) | WP_145948994.1 | WP_145948994.1 FMN-dependent NADH-azoreductase |
| 93(1) | WP_089546598.1 | WP_089546598.1 FMN-dependent NADH-azoreductase |
| 93(1) | WP_100537980.1 | WP_100537980.1 FMN-dependent NADH-azoreductase |
| 93(1) | WP_223067426.1 | WP_223067426.1 FMN-dependent NADH-azoreductase |
| 93(1) | WP_152826360.1 | WP_152826360.1 FMN-dependent NADH-azoreductase |
| 93(1) | WP_251548747.1 | WP_251548747.1 FMN-dependent NADH-azoreductase |
| 93(3) | WP_095266189.1 | WP_095266189.1 FMN-dependent NADH-azoreductase |
| 93(1) | WP_113058713.1 | WP_113058713.1 FMN-dependent NADH-azoreductase |
| 93(3) | WP_145134823.1 | WP_145134823.1 FMN-dependent NADH-azoreductase |
| 93(1) | WP_152402367.1 | WP_152402367.1 FMN-dependent NADH-azoreductase |
| 93(7) | WP_009590690.1 | WP_009590690.1 FMN-dependent NADH-azoreductase |
| 93(1) | WP_213644223.1 | WP_213644223.1 FMN-dependent NADH-azoreductase |
| 93(2) | WP_076167880.1 | WP_076167880.1 FMN-dependent NADH-azoreductase |
| 93(1) | WP_215153831.1 | WP_215153831.1 FMN-dependent NADH-azoreductase |

| | | |
|---|---|---|
| 94(2) | WP_270993640.1 | WP_270993640.1 hypothetical protein |
| 94(4) | WP_046325931.1 | WP_046325931.1 hypothetical protein |
| 94(2) | WP_149051226.1 | WP_149051226.1 hypothetical protein |
| 94(2) | WP_185639011.1 | WP_185639011.1 hypothetical protein |
| 94(3) | WP_194358207.1 | WP_194358207.1 hypothetical protein |
| 94(2) | WP_270995161.1 | WP_270995161.1 hypothetical protein |
| 94(4) | WP_228467086.1 | WP_228467086.1 hypothetical protein |
| 94(2) | WP_228463346.1 | WP_228463346.1 hypothetical protein |
| 94(3) | WP_228469160.1 | WP_228469160.1 hypothetical protein |
| 94(2) | WP_046325932.1 | WP_046325932.1 hypothetical protein |
| 94(3) | WP_194358203.1 | WP_194358203.1 hypothetical protein |
| 94(4) | WP_239079143.1 | WP_239079143.1 hypothetical protein |
| 94(4) | WP_185340299.1 | WP_185340299.1 hypothetical protein |
| 94(2) | WP_185474088.1 | WP_185474088.1 hypothetical protein |
| 94(1) | WP_185479300.1 | WP_185479300.1 hypothetical protein |
| 94(2) | WP_194334466.1 | WP_194334466.1 hypothetical protein |
| 94(2) | WP_270993642.1 | WP_270993642.1 hypothetical protein |
| 94(2) | WP_185365681.1 | WP_185365681.1 hypothetical protein |
| 94(2) | WP_194346731.1 | WP_194346731.1 hypothetical protein |
| 94(1) | WP_185436222.1 | WP_185436222.1 hypothetical protein |
| 94(2) | WP_270995164.1 | WP_270995164.1 hypothetical protein |
| 94(2) | WP_194335286.1 | WP_194335286.1 hypothetical protein |

| | | |
|---|---|---|
| 95(1) | WP_102372767.1 | WP_102372767.1 LysR family transcriptional regulator |
| 95(4) | WP_018670526.1 | WP_018670526.1 LysR family transcriptional regulator |
| 95(1) | WP_204445189.1 | WP_204445189.1 LysR family transcriptional regulator |
| 95(1) | WP_099354892.1 | WP_099354892.1 LysR family transcriptional regulator |
| 95(1) | WP_031415569.1 | WP_031415569.1 LysR family transcriptional regulator |
| 95(1) | WP_199794084.1 | WP_199794084.1 LysR family transcriptional regulator |
| 95(1) | WP_212930972.1 | WP_212930972.1 LysR family transcriptional regulator |
| 95(1) | WP_251638799.1 | WP_251638799.1 LysR family transcriptional regulator |
| 95(1) | WP_099361327.1 | WP_099361327.1 LysR family transcriptional regulator |
| 95(1) | WP_213656767.1 | WP_213656767.1 LysR family transcriptional regulator |
| 95(1) | WP_258398818.1 | WP_258398818.1 LysR family transcriptional regulator |
| 95(1) | WP_197247505.1 | WP_197247505.1 LysR family transcriptional regulator |
| 95(1) | WP_232699380.1 | WP_232699380.1 LysR family transcriptional regulator |
| 95(3) | WP_236871200.1 | WP_236871200.1 LysR family transcriptional regulator |
| 95(1) | WP_211022802.1 | WP_211022802.1 LysR family transcriptional regulator |
| 95(1) | WP_258425838.1 | WP_258425838.1 LysR family transcriptional regulator |
| 95(2) | WP_219872440.1 | WP_219872440.1 LysR family transcriptional regulator |
| 95(1) | WP_301804650.1 | WP_301804650.1 selenium metabolism-associated LysR family transcriptional regulator |
| 95(1) | WP_094699659.1 | WP_094699659.1 LysR family transcriptional regulator |
| 95(1) | WP_258427462.1 | WP_258427462.1 LysR family transcriptional regulator |
| 95(1) | WP_307192889.1 | WP_307192889.1 LysR family transcriptional regulator |
| 95(1) | WP_273383852.1 | WP_273383852.1 LysR family transcriptional regulator |
| 95(8) | WP_003336006.1 | WP_003336006.1 LysR family transcriptional regulator |
| 95(2) | WP_197262003.1 | WP_197262003.1 LysR family transcriptional regulator |
| 95(1) | WP_181747863.1 | WP_181747863.1 LysR family transcriptional regulator |
| 95(1) | WP_010676877.1 | WP_010676877.1 LysR family transcriptional regulator |
| 95(2) | WP_116334076.1 | WP_116334076.1 LysR family transcriptional regulator |
| 95(1) | WP_104065527.1 | WP_104065527.1 LysR family transcriptional regulator |
| 95(1) | WP_304216848.1 | WP_304216848.1 LysR family transcriptional regulator |
| 95(1) | WP_152392564.1 | WP_152392564.1 LysR family transcriptional regulator |
| 95(2) | WP_104030446.1 | WP_104030446.1 LysR family transcriptional regulator |
| 95(1) | WP_251638793.1 | WP_251638793.1 LysR family transcriptional regulator |
| 95(1) | WP_064019324.1 | WP_064019324.1 LysR family transcriptional regulator |
| 95(2) | WP_121474475.1 | WP_121474475.1 LysR family transcriptional regulator |
| 95(1) | WP_303942504.1 | WP_303942504.1 LysR family transcriptional regulator, partial |
| 95(1) | WP_258433014.1 | WP_258433014.1 LysR family transcriptional regulator |

| | | |
|---|---|---|
| 96(1) | WP_143881078.1 | WP_143881078.1 antitoxin YezG family protein |
| 96(1) | WP_215601069.1 | WP_215601069.1 DUF600 family protein |
| 96(1) | WP_119601468.1 | WP_119601468.1 antitoxin YezG family protein |
| 96(1) | WP_088078462.1 | WP_088078462.1 antitoxin YezG family protein |
| 96(1) | WP_181603056.1 | WP_181603056.1 antitoxin YezG family protein |
| 96(2) | WP_061663665.1 | WP_061663665.1 antitoxin YezG family protein |
| 96(2) | WP_000997352.1 | WP_000997352.1 DUF600 family protein |
| 96(1) | WP_000449604.1 | WP_000449604.1 DUF600 family protein |
| 96(5) | WP_000410096.1 | WP_000410096.1 antitoxin YezG family protein |
| 96(1) | WP_263703629.1 | WP_263703629.1 antitoxin YezG family protein, partial |
| 96(10) | WP_000997350.1 | WP_000997350.1 antitoxin YezG family protein |
| 96(1) | WP_272886474.1 | WP_272886474.1 antitoxin YezG family protein |
| 96(1) | WP_087952383.1 | WP_087952383.1 DUF600 family protein |
| 96(2) | WP_185350823.1 | WP_185350823.1 DUF600 family protein |
| 96(1) | WP_000543631.1 | WP_000543631.1 antitoxin YezG family protein |
| 96(4) | WP_000410097.1 | WP_000410097.1 antitoxin YezG family protein |
| 96(1) | WP_001091054.1 | WP_001091054.1 DUF600 family protein |
| 96(1) | WP_237522450.1 | WP_237522450.1 antitoxin YezG family protein |
| 96(1) | WP_271839900.1 | WP_271839900.1 DUF600 family protein |
| 96(1) | WP_242303039.1 | WP_242303039.1 DUF600 family protein |
| 96(10) | WP_001091052.1 | WP_001091052.1 DUF600 family protein |
| 96(1) | WP_098776426.1 | WP_098776426.1 antitoxin YezG family protein |
| 96(1) | WP_215598787.1 | WP_215598787.1 DUF600 family protein |
| 96(1) | WP_080494658.1 | WP_080494658.1 DUF600 family protein |

| | | |
|---|---|---|
| 97(2) | WP_214169375.1 | WP_214169375.1 DUF5081 family protein |

| | | |
|---|---|---|
| 97(4) | WP_185397375.1 | WP_185397375.1 DUF5081 family protein |
| 97(1) | WP_209365674.1 | WP_209365674.1 DUF5081 family protein |
| 97(1) | WP_099223944.1 | WP_099223944.1 DUF5081 family protein |
| 97(14) | WP_185336676.1 | WP_185336676.1 DUF5081 family protein |
| 97(1) | WP_021291846.1 | WP_021291846.1 DUF5081 family protein |
| 97(3) | WP_038245442.1 | WP_038245442.1 DUF5081 family protein |
| 97(1) | WP_115359664.1 | WP_115359664.1 DUF5081 family protein |
| 97(1) | WP_101356547.1 | WP_101356547.1 DUF5081 family protein |
| 97(1) | WP_216390029.1 | WP_216390029.1 DUF5081 family protein |
| 97(1) | WP_018664845.1 | WP_018664845.1 DUF5081 family protein |
| 97(1) | WP_035190270.1 | WP_035190270.1 DUF5081 family protein |
| 97(4) | WP_185375685.1 | WP_185375685.1 DUF5081 family protein |
| 97(2) | WP_234288981.1 | WP_234288981.1 DUF5081 family protein |
| 97(2) | WP_185461810.1 | WP_185461810.1 DUF5081 family protein |
| 97(1) | WP_074598756.1 | WP_074598756.1 DUF5081 family protein |
| 97(8) | WP_221633490.1 | WP_221633490.1 DUF5081 family protein |
| 97(2) | WP_185489274.1 | WP_185489274.1 DUF5081 family protein |
| 98(3) | WP_193373350.1 | WP_193373350.1 hypothetical protein |
| 98(1) | WP_253806479.1 | WP_253806479.1 hypothetical protein |
| 98(1) | WP_181127218.1 | WP_181127218.1 hypothetical protein |
| 98(5) | WP_231109418.1 | WP_231109418.1 hypothetical protein |
| 98(1) | WP_080582304.1 | WP_080582304.1 hypothetical protein |
| 98(2) | WP_225164874.1 | WP_225164874.1 hypothetical protein |
| 98(1) | WP_228552522.1 | WP_228552522.1 hypothetical protein |
| 98(1) | WP_244310716.1 | WP_244310716.1 hypothetical protein |
| 98(1) | WP_250247158.1 | WP_250247158.1 hypothetical protein |
| 98(1) | WP_274485140.1 | WP_274485140.1 hypothetical protein |
| 98(1) | WP_225999398.1 | WP_225999398.1 hypothetical protein |
| 98(8) | WP_231124967.1 | WP_231124967.1 hypothetical protein |
| 98(17) | WP_226887697.1 | WP_226887697.1 hypothetical protein |
| 98(1) | WP_258013832.1 | WP_258013832.1 hypothetical protein |
| 98(3) | WP_013370772.1 | WP_013370772.1 hypothetical protein |
| 98(1) | WP_231887381.1 | WP_231887381.1 hypothetical protein |
| 98(1) | WP_230877356.1 | WP_230877356.1 hypothetical protein |
| 98(1) | WP_305166269.1 | WP_305166269.1 hypothetical protein |
| 99(1) | WP_035190274.1 | WP_035190274.1 WXG100 family type VII secretion target |
| 99(22) | WP_185336677.1 | WP_185336677.1 WXG100 family type VII secretion target |
| 99(3) | WP_021291847.1 | WP_021291847.1 hypothetical protein |
| 99(1) | WP_099223946.1 | WP_099223946.1 WXG100 family type VII secretion target |
| 99(1) | WP_209365675.1 | WP_209365675.1 WXG100 family type VII secretion target |
| 99(1) | WP_026571204.1 | WP_026571204.1 WXG100 family type VII secretion target |
| 99(1) | WP_101356548.1 | WP_101356548.1 WXG100 family type VII secretion target |
| 99(1) | WP_018664844.1 | WP_018664844.1 WXG100 family type VII secretion target |
| 99(1) | WP_160807258.1 | WP_160807258.1 hypothetical protein |
| 99(1) | WP_115359663.1 | WP_115359663.1 WXG100 family type VII secretion target |
| 99(2) | WP_200372557.1 | WP_200372557.1 WXG100 family type VII secretion target |
| 99(14) | WP_185375683.1 | WP_185375683.1 WXG100 family type VII secretion target |
| 100(14) | WP_185336678.1 | WP_185336678.1 chorismate synthase |
| 100(2) | WP_200372559.1 | WP_200372559.1 chorismate synthase |
| 100(8) | WP_185349991.1 | WP_185349991.1 chorismate synthase |
| 100(1) | WP_037329432.1 | WP_037329432.1 hypothetical protein, partial |
| 100(12) | WP_185375681.1 | WP_185375681.1 chorismate synthase |
| 100(3) | WP_074436683.1 | WP_074436683.1 hypothetical protein |
| 100(1) | WP_101356549.1 | WP_101356549.1 chorismate synthase |
| 100(1) | WP_115359662.1 | WP_115359662.1 hypothetical protein |
| 100(1) | WP_099223948.1 | WP_099223948.1 chorismate synthase |
| 100(1) | WP_018664843.1 | WP_018664843.1 hypothetical protein |
| 100(1) | WP_035190276.1 | WP_035190276.1 hypothetical protein |
| 100(1) | WP_209365676.1 | WP_209365676.1 chorismate synthase |
| 100(2) | WP_194332265.1 | WP_194332265.1 chorismate synthase |
| 100(1) | WP_155974818.1 | WP_155974818.1 hypothetical protein |
| 101(2) | WP_194336873.1 | WP_194336873.1 hypothetical protein |
| 101(1) | WP_101356546.1 | WP_101356546.1 hypothetical protein |
| 101(22) | WP_185336675.1 | WP_185336675.1 hypothetical protein |
| 101(2) | WP_185489273.1 | WP_185489273.1 hypothetical protein |
| 101(1) | WP_035190267.1 | WP_035190267.1 hypothetical protein |
| 101(6) | WP_185375686.1 | WP_185375686.1 hypothetical protein |
| 101(4) | WP_021291845.1 | WP_021291845.1 hypothetical protein |
| 101(4) | WP_185397376.1 | WP_185397376.1 hypothetical protein |
| 101(2) | WP_038406383.1 | WP_038406383.1 hypothetical protein |
| 101(1) | WP_099223942.1 | WP_099223942.1 hypothetical protein |
| 101(1) | WP_115359665.1 | WP_115359665.1 hypothetical protein |
| 101(1) | WP_051382196.1 | WP_051382196.1 DUF5344 family protein |
| 101(1) | WP_083334785.1 | WP_083334785.1 DUF5344 family protein |
| 102(3) | WP_098296557.1 | WP_098296557.1 conserved virulence factor C family protein |
| 102(2) | WP_000692572.1 | WP_000692572.1 conserved virulence factor C family protein |
| 102(1) | WP_098920185.1 | WP_098920185.1 conserved virulence factor C family protein |
| 102(1) | WP_071679482.1 | WP_071679482.1 conserved virulence factor C family protein |
| 102(2) | WP_017561865.1 | WP_017561865.1 conserved virulence factor C family protein |
| 102(1) | WP_262719892.1 | WP_262719892.1 conserved virulence factor C family protein |
| 102(1) | WP_087959684.1 | WP_087959684.1 conserved virulence factor C family protein |
| 102(1) | WP_070805894.1 | WP_070805894.1 conserved virulence factor C family protein |
| 102(3) | WP_098264273.1 | WP_098264273.1 conserved virulence factor C family protein |
| 102(1) | WP_000692552.1 | WP_000692552.1 conserved virulence factor C family protein |
| 102(1) | WP_098230732.1 | WP_098230732.1 conserved virulence factor C family protein |
| 102(1) | WP_139867400.1 | WP_139867400.1 conserved virulence factor C family protein |
| 102(1) | WP_144564256.1 | WP_144564256.1 conserved virulence factor C family protein |
| 102(1) | WP_243500430.1 | WP_243500430.1 virulence factor |
| 102(6) | WP_087982610.1 | WP_087982610.1 conserved virulence factor C family protein |
| 102(1) | WP_087952363.1 | WP_087952363.1 conserved virulence factor C family protein |
| 102(1) | WP_144611175.1 | WP_144611175.1 conserved virulence factor C family protein |
| 102(1) | WP_060630635.1 | WP_060630635.1 conserved virulence factor C family protein |
| 102(1) | WP_098856758.1 | WP_098856758.1 conserved virulence factor C family protein |
| 102(1) | WP_078419981.1 | WP_078419981.1 conserved virulence factor C family protein |
| 102(1) | WP_173601440.1 | WP_173601440.1 conserved virulence factor C family protein |
| 102(1) | WP_097952804.1 | WP_097952804.1 conserved virulence factor C family protein |
| 102(1) | WP_098288674.1 | WP_098288674.1 conserved virulence factor C family protein |
| 102(3) | WP_064473949.1 | WP_064473949.1 conserved virulence factor C family protein |
| 102(3) | WP_098088265.1 | WP_098088265.1 conserved virulence factor C family protein |

| | | | |
|---|---|---|---|
| 102(1) | WP_000692555.1 | WP_000692555.1 | conserved virulence factor C family protein |
| 102(1) | WP_088312109.1 | WP_088312109.1 | conserved virulence factor C family protein |
| 102(1) | WP_144488213.1 | WP_144488213.1 | conserved virulence factor C family protein |
| 102(1) | WP_000692584.1 | WP_000692584.1 | conserved virulence factor C family protein |
| 102(1) | WP_151630617.1 | WP_151630617.1 | conserved virulence factor C family protein |
| 102(1) | WP_033692030.1 | WP_033692030.1 | conserved virulence factor C family protein |
| 102(2) | WP_000692577.1 | WP_000692577.1 | conserved virulence factor C family protein |
| 103(1) | WP_077836249.1 | WP_077836249.1 | IS110 family transposase |
| 103(1) | WP_253442817.1 | WP_253442817.1 | IS110 family transposase |
| 103(1) | WP_042512830.1 | WP_042512830.1 | IS110 family transposase |
| 103(1) | WP_139895507.1 | WP_139895507.1 | IS110 family transposase |
| 103(1) | WP_212330930.1 | WP_212330930.1 | transposase |
| 103(1) | WP_199677613.1 | WP_199677613.1 | IS110 family transposase |
| 103(1) | WP_000551906.1 | WP_000551906.1 | IS110 family transposase |
| 103(1) | WP_012260842.1 | WP_012260842.1 | IS110 family transposase |
| 103(1) | WP_215577251.1 | WP_215577251.1 | IS110 family transposase |
| 103(2) | WP_228469905.1 | WP_228469905.1 | transposase |
| 103(1) | WP_157418135.1 | WP_157418135.1 | IS110 family transposase |
| 103(1) | WP_242251790.1 | WP_242251790.1 | IS110 family transposase |
| 103(1) | WP_078993363.1 | WP_078993363.1 | IS110 family transposase |
| 103(2) | WP_200152285.1 | WP_200152285.1 | IS110 family transposase |
| 103(1) | WP_236804426.1 | WP_236804426.1 | IS110 family transposase |
| 103(1) | WP_212330927.1 | WP_212330927.1 | transposase |
| 103(1) | WP_068696849.1 | WP_068696849.1 | IS110 family transposase |
| 103(1) | WP_063219632.1 | WP_063219632.1 | IS110 family transposase |
| 103(1) | WP_098670176.1 | WP_098670176.1 | IS110 family transposase |
| 103(1) | WP_143881075.1 | WP_143881075.1 | IS110 family transposase |
| 103(1) | WP_095691006.1 | WP_095691006.1 | IS110 family transposase |
| 103(2) | WP_000551892.1 | WP_000551892.1 | IS110 family transposase |
| 103(1) | WP_098033305.1 | WP_098033305.1 | IS110 family transposase |
| 103(1) | WP_055105783.1 | WP_055105783.1 | IS110 family transposase |
| 103(1) | WP_144492101.1 | WP_144492101.1 | IS110 family transposase |
| 103(1) | WP_245859336.1 | WP_245859336.1 | IS110 family transposase |
| 103(1) | WP_074598753.1 | WP_074598753.1 | transposase |
| 103(1) | WP_000551891.1 | WP_000551891.1 | IS110 family transposase |
| 103(1) | WP_075395692.1 | WP_075395692.1 | IS110 family transposase |
| 103(1) | WP_197225373.1 | WP_197225373.1 | IS110 family transposase |
| 103(1) | WP_000546503.1 | WP_000546503.1 | IS110 family transposase |
| 103(1) | WP_063223285.1 | WP_063223285.1 | IS110 family transposase |
| 103(1) | WP_086419725.1 | WP_086419725.1 | IS110 family transposase |
| 103(1) | WP_097822701.1 | WP_097822701.1 | IS110 family transposase |
| 103(1) | WP_098768164.1 | WP_098768164.1 | IS110 family transposase |
| 103(1) | WP_211082934.1 | WP_211082934.1 | IS110 family transposase |
| 103(1) | WP_097987488.1 | WP_097987488.1 | IS110 family transposase |
| 103(2) | WP_228469893.1 | WP_228469893.1 | IS110 family transposase |
| 103(1) | WP_098148065.1 | WP_098148065.1 | IS110 family transposase |
| 103(1) | WP_237100167.1 | WP_237100167.1 | IS110 family transposase |
| 103(1) | WP_002148299.1 | WP_002148299.1 | IS110 family transposase |
| 103(1) | WP_000551901.1 | WP_000551901.1 | IS110 family transposase |
| 103(1) | WP_289398703.1 | WP_289398703.1 | IS110 family transposase |
| 103(1) | WP_228469272.1 | WP_228469272.1 | IS110 family transposase |

| | | | |
|---|---|---|---|
| 104(12) | WP_001087573.1 | WP_001087573.1 | hypothetical protein |
| 104(5) | WP_078181471.1 | WP_078181471.1 | hypothetical protein |
| 104(1) | WP_098204520.1 | WP_098204520.1 | hypothetical protein |
| 104(1) | WP_098167781.1 | WP_098167781.1 | hypothetical protein |
| 104(1) | WP_169028884.1 | WP_169028884.1 | ribonuclease toxin immunity protein CdiI |
| 104(7) | WP_001087574.1 | WP_001087574.1 | hypothetical protein |
| 104(1) | WP_253440785.1 | WP_253440785.1 | ribonuclease toxin immunity protein CdiI |
| 104(1) | WP_097846502.1 | WP_097846502.1 | hypothetical protein |
| 104(1) | WP_147807652.1 | WP_147807652.1 | hypothetical protein |
| 104(8) | WP_001087572.1 | WP_001087572.1 | hypothetical protein |
| 104(2) | WP_097952957.1 | WP_097952957.1 | hypothetical protein |
| 104(1) | WP_076157815.1 | WP_076157815.1 | ribonuclease toxin immunity protein CdiI |
| 104(1) | WP_098296565.1 | WP_098296565.1 | hypothetical protein |
| 104(1) | WP_306007806.1 | WP_306007806.1 | hypothetical protein |
| 104(2) | WP_061682591.1 | WP_061682591.1 | hypothetical protein |
| 104(2) | WP_097822667.1 | WP_097822667.1 | hypothetical protein |
| 105(3) | WP_001021920.1 | WP_001021920.1 | GNAT family N-acetyltransferase |
| 105(1) | WP_001101105.1 | WP_001101105.1 | GNAT family N-acetyltransferase |
| 105(3) | WP_001101115.1 | WP_001101115.1 | GNAT family N-acetyltransferase |
| 105(1) | WP_088066051.1 | WP_088066051.1 | GNAT family N-acetyltransferase |
| 105(3) | WP_001101116.1 | WP_001101116.1 | GNAT family N-acetyltransferase |
| 105(4) | WP_058840366.1 | WP_058840366.1 | GNAT family N-acetyltransferase |
| 105(3) | WP_001101111.1 | WP_001101111.1 | GNAT family N-acetyltransferase |
| 105(1) | WP_307284649.1 | WP_307284649.1 | GNAT family N-acetyltransferase |
| 105(5) | WP_001101112.1 | WP_001101112.1 | GNAT family N-acetyltransferase |
| 105(1) | WP_001101104.1 | WP_001101104.1 | GNAT family N-acetyltransferase |
| 105(2) | WP_001101109.1 | WP_001101109.1 | GNAT family N-acetyltransferase |
| 105(1) | WP_098874959.1 | WP_098874959.1 | GNAT family N-acetyltransferase |
| 105(1) | WP_086412795.1 | WP_086412795.1 | GNAT family N-acetyltransferase |
| 105(1) | WP_001101110.1 | WP_001101110.1 | GNAT family N-acetyltransferase |
| 105(3) | WP_001101103.1 | WP_001101103.1 | GNAT family N-acetyltransferase |
| 105(1) | WP_098284010.1 | WP_098284010.1 | GNAT family N-acetyltransferase |
| 105(1) | WP_308082227.1 | WP_308082227.1 | GNAT family N-acetyltransferase |
| 105(1) | WP_071721667.1 | WP_071721667.1 | GNAT family N-acetyltransferase |
| 105(1) | WP_001101108.1 | WP_001101108.1 | GNAT family N-acetyltransferase |
| 105(5) | WP_001101118.1 | WP_001101118.1 | GNAT family N-acetyltransferase |
| 105(1) | WP_001101124.1 | WP_001101124.1 | GNAT family N-acetyltransferase |
| 105(2) | WP_001101123.1 | WP_001101123.1 | GNAT family N-acetyltransferase |
| 105(2) | WP_086385915.1 | WP_086385915.1 | GNAT family N-acetyltransferase |
| 106(1) | WP_000270898.1 | WP_000270898.1 | 2'-5' RNA ligase family protein |
| 106(1) | WP_200462050.1 | WP_200462050.1 | 2'-5' RNA ligase family protein |
| 106(1) | WP_000270857.1 | WP_000270857.1 | 2'-5' RNA ligase family protein |
| 106(1) | WP_098141778.1 | WP_098141778.1 | 2'-5' RNA ligase family protein |
| 106(1) | WP_078384710.1 | WP_078384710.1 | 2'-5' RNA ligase family protein |
| 106(1) | WP_000270884.1 | WP_000270884.1 | 2'-5' RNA ligase family protein |
| 106(1) | WP_098139273.1 | WP_098139273.1 | 2'-5' RNA ligase family protein |
| 106(1) | WP_065211638.1 | WP_065211638.1 | 2'-5' RNA ligase family protein |
| 106(1) | WP_048655640.1 | WP_048655640.1 | 2'-5' RNA ligase family protein |
| 106(1) | WP_098955452.1 | WP_098955452.1 | 2'-5' RNA ligase family protein |

| | | |
|---|---|---|
| 106(1) | WP_025710185.1 | WP_025710185.1 2'-5' RNA ligase family protein |
| 106(6) | WP_006923797.1 | WP_006923797.1 2'-5' RNA ligase family protein |
| 106(1) | WP_065223573.1 | WP_065223573.1 2'-5' RNA ligase family protein |
| 106(1) | WP_016117664.1 | WP_016117664.1 2'-5' RNA ligase family protein |
| 106(1) | WP_061667213.1 | WP_061667213.1 2'-5' RNA ligase family protein |
| 106(3) | WP_000270885.1 | WP_000270885.1 2'-5' RNA ligase family protein |
| 106(1) | WP_098874961.1 | WP_098874961.1 2'-5' RNA ligase family protein |
| 106(2) | WP_000270888.1 | WP_000270888.1 2'-5' RNA ligase family protein |
| 106(2) | WP_000270887.1 | WP_000270887.1 2'-5' RNA ligase family protein |
| 106(1) | WP_098226397.1 | WP_098226397.1 2'-5' RNA ligase family protein |
| 106(3) | WP_097954759.1 | WP_097954759.1 2'-5' RNA ligase family protein |
| 106(1) | WP_088066053.1 | WP_088066053.1 2'-5' RNA ligase family protein |
| 106(1) | WP_000270890.1 | WP_000270890.1 2'-5' RNA ligase family protein |
| 106(3) | WP_000270892.1 | WP_000270892.1 2'-5' RNA ligase family protein |
| 106(1) | WP_208906813.1 | WP_208906813.1 2'-5' RNA ligase family protein |
| 106(3) | WP_001990267.1 | WP_001990267.1 2'-5' RNA ligase family protein |
| 106(1) | WP_098284012.1 | WP_098284012.1 2'-5' RNA ligase family protein |
| 106(1) | WP_132131914.1 | WP_132131914.1 2'-5' RNA ligase family protein |
| 106(3) | WP_086405912.1 | WP_086405912.1 2'-5' RNA ligase family protein |
| 107(3) | WP_213117467.1 | WP_213117467.1 PadR family transcriptional regulator |
| 107(3) | WP_100537978.1 | WP_100537978.1 PadR family transcriptional regulator |
| 107(3) | WP_007128634.1 | WP_007128634.1 PadR family transcriptional regulator |
| 107(1) | WP_054958404.1 | WP_054958404.1 PadR family transcriptional regulator |
| 107(1) | WP_023957713.1 | WP_023957713.1 PadR family transcriptional regulator |
| 107(1) | WP_028405552.1 | WP_028405552.1 PadR family transcriptional regulator |
| 107(1) | WP_077567022.1 | WP_077567022.1 PadR family transcriptional regulator |
| 107(8) | WP_009590638.1 | WP_009590638.1 PadR family transcriptional regulator |
| 107(1) | WP_053492735.1 | WP_053492735.1 PadR family transcriptional regulator |
| 107(8) | WP_071220202.1 | WP_071220202.1 PadR family transcriptional regulator |
| 107(1) | WP_241929681.1 | WP_241929681.1 PadR family transcriptional regulator |
| 107(6) | WP_006211208.1 | WP_006211208.1 PadR family transcriptional regulator |
| 107(1) | WP_208919024.1 | WP_208919024.1 PadR family transcriptional regulator |
| 107(3) | WP_095266192.1 | WP_095266192.1 PadR family transcriptional regulator |
| 107(2) | WP_119847550.1 | WP_119847550.1 PadR family transcriptional regulator |
| 107(1) | WP_213644226.1 | WP_213644226.1 PadR family transcriptional regulator |
| 108(1) | WP_256719594.1 | WP_256719594.1 DUF1700 domain-containing protein |
| 108(1) | WP_100537979.1 | WP_100537979.1 DUF1700 domain-containing protein |
| 108(1) | WP_256719595.1 | WP_256719595.1 DUF1700 domain-containing protein |
| 108(1) | WP_054958403.1 | WP_054958403.1 DUF1700 domain-containing protein |
| 108(1) | WP_145037670.1 | WP_145037670.1 DUF1700 domain-containing protein |
| 108(3) | WP_095266191.1 | WP_095266191.1 DUF1700 domain-containing protein |
| 108(1) | WP_054402907.1 | WP_054402907.1 DUF1700 domain-containing protein |
| 108(2) | WP_119847549.1 | WP_119847549.1 DUF1700 domain-containing protein |
| 108(1) | WP_023957709.1 | WP_023957709.1 DUF1700 domain-containing protein |
| 108(1) | WP_028405553.1 | WP_028405553.1 DUF1700 domain-containing protein |
| 108(3) | WP_096774456.1 | WP_096774456.1 DUF1700 domain-containing protein |
| 108(1) | WP_215153835.1 | WP_215153835.1 DUF1700 domain-containing protein |
| 108(1) | WP_113058715.1 | WP_113058715.1 DUF1700 domain-containing protein |
| 108(1) | WP_194543499.1 | WP_194543499.1 DUF1700 domain-containing protein |
| 108(1) | WP_213644225.1 | WP_213644225.1 DUF1700 domain-containing protein |
| 108(5) | WP_036642153.1 | WP_036642153.1 DUF1700 domain-containing protein |
| 108(1) | WP_127592430.1 | WP_127592430.1 DUF1700 domain-containing protein |
| 108(1) | WP_251497196.1 | WP_251497196.1 DUF1700 domain-containing protein |
| 108(1) | WP_090908275.1 | WP_090908275.1 DUF1700 domain-containing protein |
| 108(1) | WP_076322117.1 | WP_076322117.1 DUF1700 domain-containing protein |
| 108(1) | WP_009590652.1 | WP_009590652.1 DUF1700 domain-containing protein |
| 108(1) | WP_007128635.1 | WP_007128635.1 DUF1700 domain-containing protein |
| 108(1) | WP_098746874.1 | WP_098746874.1 DUF1700 domain-containing protein |
| 108(1) | WP_077567023.1 | WP_077567023.1 DUF1700 domain-containing protein |
| 108(1) | WP_210095107.1 | WP_210095107.1 DUF1700 domain-containing protein |
| 108(1) | WP_213616449.1 | WP_213616449.1 DUF1700 domain-containing protein |
| 108(1) | WP_208919014.1 | WP_208919014.1 DUF1700 domain-containing protein |
| 108(1) | WP_015736246.1 | WP_015736246.1 DUF1700 domain-containing protein |
| 108(1) | WP_213579163.1 | WP_213579163.1 DUF1700 domain-containing protein |
| 108(1) | WP_023957711.1 | WP_023957711.1 DUF1700 domain-containing protein |
| 108(1) | WP_241929682.1 | WP_241929682.1 DUF1700 domain-containing protein |
| 108(4) | WP_071220201.1 | WP_071220201.1 DUF1700 domain-containing protein |
| 109(2) | WP_199794078.1 | WP_199794078.1 AAA family ATPase |
| 109(1) | WP_101668494.1 | WP_101668494.1 AAA family ATPase |
| 109(1) | WP_278431133.1 | WP_278431133.1 AAA family ATPase |
| 109(1) | WP_277548425.1 | WP_277548425.1 AAA family ATPase |
| 109(1) | WP_115359672.1 | WP_115359672.1 ATP-dependent zinc metalloprotease FtsH |
| 109(1) | WP_239209295.1 | WP_239209295.1 AAA family ATPase |
| 109(1) | WP_168419975.1 | WP_168419975.1 AAA family ATPase |
| 109(1) | WP_212930983.1 | WP_212930983.1 AAA family ATPase |
| 109(1) | WP_258417134.1 | WP_258417134.1 AAA family ATPase |
| 109(1) | WP_197262011.1 | WP_197262011.1 AAA family ATPase |
| 109(1) | WP_096886746.1 | WP_096886746.1 AAA family ATPase |
| 109(1) | WP_003340568.1 | WP_003340568.1 AAA family ATPase |
| 109(1) | WP_158326745.1 | WP_158326745.1 AAA family ATPase |
| 109(1) | WP_140574906.1 | WP_140574906.1 AAA family ATPase |
| 109(1) | WP_158329540.1 | WP_158329540.1 AAA family ATPase |
| 109(1) | WP_258418567.1 | WP_258418567.1 AAA family ATPase |
| 109(1) | WP_232699396.1 | WP_232699396.1 AAA family ATPase |
| 109(1) | WP_003335987.1 | WP_003335987.1 AAA family ATPase |
| 109(1) | WP_158330736.1 | WP_158330736.1 AAA family ATPase |
| 109(1) | WP_163245196.1 | WP_163245196.1 AAA family ATPase |
| 109(1) | WP_134750586.1 | WP_134750586.1 AAA family ATPase |
| 109(1) | WP_116334086.1 | WP_116334086.1 AAA family ATPase |
| 109(1) | WP_113755268.1 | WP_113755268.1 AAA family ATPase |
| 109(1) | WP_121474484.1 | WP_121474484.1 AAA family ATPase |
| 109(1) | WP_018670509.1 | WP_018670509.1 AAA family ATPase |
| 109(1) | WP_064019309.1 | WP_064019309.1 AAA family ATPase |
| 109(1) | WP_181747879.1 | WP_181747879.1 AAA family ATPase |
| 109(1) | WP_094699644.1 | WP_094699644.1 AAA family ATPase |
| 109(2) | WP_104030432.1 | WP_104030432.1 AAA family ATPase |
| 109(1) | WP_197247526.1 | WP_197247526.1 AAA family ATPase |
| 109(1) | WP_104065541.1 | WP_104065541.1 AAA family ATPase |
| 109(3) | WP_213117474.1 | WP_213117474.1 AAA family ATPase |
| 109(1) | WP_068793939.1 | WP_068793939.1 AAA family ATPase |
| 109(1) | WP_213143940.1 | WP_213143940.1 AAA family ATPase |

109(1) WP_258425847.1 WP_258425847.1 AAA family ATPase
109(1) WP_249897304.1 WP_249897304.1 ATP-binding protein
109(1) WP_301804693.1 WP_301804693.1 AAA family ATPase
109(1) WP_119732852.1 WP_119732852.1 AAA family ATPase
109(1) WP_022586758.1 WP_022586758.1 AAA family ATPase
109(1) WP_258433026.1 WP_258433026.1 AAA family ATPase

110(3) WP_001990270.1 WP_001990270.1 metallophosphoesterase
110(1) WP_000721377.1 WP_000721377.1 metallophosphoesterase
110(1) WP_000411335.1 WP_000411335.1 metallophosphoesterase
110(1) WP_098226398.1 WP_098226398.1 metallophosphoesterase
110(1) WP_144596437.1 WP_144596437.1 metallophosphoesterase
110(1) WP_000721380.1 WP_000721380.1 metallophosphoesterase
110(1) WP_053513137.1 WP_053513137.1 metallophosphoesterase
110(1) WP_151525517.1 WP_151525517.1 metallophosphoesterase
110(1) WP_208906811.1 WP_208906811.1 metallophosphoesterase
110(1) WP_000721376.1 WP_000721376.1 metallophosphoesterase
110(1) WP_142325859.1 WP_142325859.1 metallophosphoesterase
110(1) WP_070179258.1 WP_070179258.1 metallophosphoesterase
110(2) WP_002105355.1 WP_002105355.1 metallophosphoesterase
110(1) WP_000721382.1 WP_000721382.1 metallophosphoesterase
110(1) WP_140335839.1 WP_140335839.1 metallophosphoesterase
110(1) WP_000721388.1 WP_000721388.1 metallophosphoesterase
110(2) WP_000721383.1 WP_000721383.1 metallophosphoesterase
110(1) WP_033703539.1 WP_033703539.1 metallophosphoesterase
110(1) WP_140337666.1 WP_140337666.1 metallophosphoesterase
110(1) WP_103597529.1 WP_103597529.1 metallophosphoesterase
110(2) WP_016122366.1 WP_016122366.1 metallophosphoesterase
110(1) WP_000721390.1 WP_000721390.1 metallophosphoesterase
110(1) WP_284996385.1 WP_284996385.1 metallophosphoesterase
110(1) WP_042972406.1 WP_042972406.1 metallophosphoesterase
110(1) WP_142327751.1 WP_142327751.1 metallophosphoesterase
110(1) WP_006923792.1 WP_006923792.1 metallophosphoesterase
110(1) WP_048535507.1 WP_048535507.1 metallophosphoesterase
110(1) WP_163095419.1 WP_163095419.1 metallophosphoesterase
110(2) WP_030025422.1 WP_030025422.1 metallophosphoesterase
110(1) WP_098157702.1 WP_098157702.1 metallophosphoesterase
110(1) WP_141538934.1 WP_141538934.1 metallophosphoesterase
110(5) WP_000721385.1 WP_000721385.1 metallophosphoesterase
110(1) WP_142329694.1 WP_142329694.1 metallophosphoesterase

111(1) WP_179090612.1 WP_179090612.1 DUF4097 family beta strand repeat-containing protein
111(1) WP_176502424.1 WP_176502424.1 DUF4097 family beta strand repeat-containing protein
111(1) WP_175495020.1 WP_175495020.1 DUF4097 family beta strand repeat-containing protein
111(1) WP_007128636.1 WP_007128636.1 DUF4097 family beta strand repeat-containing protein
111(1) WP_181469795.1 WP_181469795.1 DUF4097 family beta strand repeat-containing protein
111(1) WP_162487570.1 WP_162487570.1 DUF4097 family beta strand repeat-containing protein
111(1) WP_172963933.1 WP_172963933.1 DUF4097 family beta strand repeat-containing protein
111(1) WP_260065255.1 WP_260065255.1 DUF4097 domain-containing protein
111(1) WP_213616447.1 WP_213616447.1 DUF4097 family beta strand repeat-containing protein
111(1) WP_253300559.1 WP_253300559.1 DUF4097 family beta strand repeat-containing protein
111(2) WP_162939118.1 WP_162939118.1 DUF4097 family beta strand repeat-containing protein
111(1) WP_194543500.1 WP_194543500.1 DUF4097 family beta strand repeat-containing protein
111(1) WP_192571316.1 WP_192571316.1 DUF4097 family beta strand repeat-containing protein
111(1) WP_077567024.1 WP_077567024.1 DUF4097 family beta strand repeat-containing protein
111(1) WP_277746360.1 WP_277746360.1 DUF4097 family beta strand repeat-containing protein
111(1) WP_152826476.1 WP_152826476.1 DUF4097 family beta strand repeat-containing protein
111(1) WP_028405554.1 WP_028405554.1 DUF4097 family beta strand repeat-containing protein
111(1) WP_285857489.1 WP_285857489.1 DUF4097 family beta strand repeat-containing protein
111(1) WP_213644224.1 WP_213644224.1 DUF4097 family beta strand repeat-containing protein
111(1) WP_251035374.1 WP_251035374.1 DUF4097 family beta strand repeat-containing protein
111(1) WP_241929683.1 WP_241929683.1 DUF4097 family beta strand repeat-containing protein
111(1) WP_161952958.1 WP_161952958.1 DUF4097 family beta strand repeat-containing protein
111(1) WP_095398790.1 WP_095398790.1 DUF4097 family beta strand repeat-containing protein
111(1) WP_164817411.1 WP_164817411.1 DUF4097 family beta strand repeat-containing protein
111(1) WP_155618102.1 WP_155618102.1 DUF4097 family beta strand repeat-containing protein
111(1) WP_165789660.1 WP_165789660.1 DUF4097 family beta strand repeat-containing protein
111(1) WP_054958402.1 WP_054958402.1 DUF4097 family beta strand repeat-containing protein
111(1) WP_208919012.1 WP_208919012.1 DUF4097 family beta strand repeat-containing protein
111(4) WP_006211210.1 WP_006211210.1 DUF4097 family beta strand repeat-containing protein
111(2) WP_162164388.1 WP_162164388.1 DUF4097 family beta strand repeat-containing protein
111(1) WP_175320744.1 WP_175320744.1 DUF4097 family beta strand repeat-containing protein
111(1) WP_213579164.1 WP_213579164.1 DUF4097 family beta strand repeat-containing protein
111(1) WP_220684638.1 WP_220684638.1 DUF4097 family beta strand repeat-containing protein
111(1) WP_245256053.1 WP_245256053.1 DUF4097 family beta strand repeat-containing protein
111(1) WP_165593397.1 WP_165593397.1 DUF4097 family beta strand repeat-containing protein
111(1) WP_023957707.1 WP_023957707.1 DUF4097 family beta strand repeat-containing protein
111(1) WP_095266190.1 WP_095266190.1 DUF4097 family beta strand repeat-containing protein

112(1) WP_236417234.1 WP_236417234.1 ribonuclease domain-containing protein
112(1) WP_090993640.1 WP_090993640.1 ribonuclease domain-containing protein
112(1) WP_036670445.1 WP_036670445.1 ribonuclease
112(1) WP_024630338.1 WP_024630338.1 ribonuclease
112(1) WP_076208834.1 WP_076208834.1 ribonuclease domain-containing protein
112(3) WP_124114126.1 WP_124114126.1 ribonuclease domain-containing protein
112(2) WP_090896217.1 WP_090896217.1 ribonuclease domain-containing protein
112(1) WP_247902254.1 WP_247902254.1 ribonuclease domain-containing protein
112(1) WP_062326158.1 WP_062326158.1 ribonuclease domain-containing protein
112(1) WP_289392080.1 WP_289392080.1 ribonuclease domain-containing protein
112(3) WP_076318556.1 WP_076318556.1 ribonuclease domain-containing protein
112(1) WP_079694640.1 WP_079694640.1 ribonuclease domain-containing protein
112(1) WP_091017185.1 WP_091017185.1 ribonuclease domain-containing protein
112(1) WP_127543687.1 WP_127543687.1 ribonuclease domain-containing protein
112(3) WP_285079433.1 WP_285079433.1 ribonuclease domain-containing protein
112(1) WP_074094622.1 WP_074094622.1 ribonuclease domain-containing protein
112(5) WP_064639004.1 WP_064639004.1 ribonuclease domain-containing protein
112(1) WP_109999900.1 WP_109999900.1 ribonuclease domain-containing protein
112(1) WP_145318664.1 WP_145318664.1 ribonuclease domain-containing protein
112(1) WP_053783898.1 WP_053783898.1 ribonuclease
112(1) WP_047842543.1 WP_047842543.1 ribonuclease
112(1) WP_017689059.1 WP_017689059.1 ribonuclease domain-containing protein
112(2) WP_062833890.1 WP_062833890.1 ribonuclease domain-containing protein
112(1) WP_076330729.1 WP_076330729.1 ribonuclease domain-containing protein
112(3) WP_036609184.1 WP_036609184.1 ribonuclease domain-containing protein

| | | |
|---|---|---|
| 112(1) | WP_091031755.1 | WP_091031755.1 ribonuclease domain-containing protein |
| 112(1) | WP_264929197.1 | WP_264929197.1 ribonuclease |

| | | |
|---|---|---|
| 113(1) | WP_086412798.1 | WP_086412798.1 cupin |
| 113(2) | WP_000688080.1 | WP_000688080.1 hypothetical protein |
| 113(1) | WP_000688082.1 | WP_000688082.1 cupin |
| 113(3) | WP_000688078.1 | WP_000688078.1 hypothetical protein |
| 113(1) | WP_088066054.1 | WP_088066054.1 cupin |
| 113(1) | WP_098284013.1 | WP_098284013.1 cupin |
| 113(1) | WP_098287134.1 | WP_098287134.1 cupin |
| 113(1) | WP_000688064.1 | WP_000688064.1 cupin |
| 113(1) | WP_098955449.1 | WP_098955449.1 cupin |
| 113(1) | WP_098226396.1 | WP_098226396.1 cupin |
| 113(3) | WP_030025429.1 | WP_030025429.1 cupin |
| 113(1) | WP_098567748.1 | WP_098567748.1 cupin |
| 113(2) | WP_000688072.1 | WP_000688072.1 cupin |
| 113(1) | WP_046392635.1 | WP_046392635.1 cupin |
| 113(1) | WP_098381997.1 | WP_098381997.1 cupin |
| 113(5) | WP_000688076.1 | WP_000688076.1 cupin |
| 113(3) | WP_001990264.1 | WP_001990264.1 cupin |
| 113(6) | WP_000688074.1 | WP_000688074.1 cupin |
| 113(1) | WP_097954758.1 | WP_097954758.1 cupin |
| 113(1) | WP_098612026.1 | WP_098612026.1 cupin |
| 113(2) | WP_098055915.1 | WP_098055915.1 cupin |
| 113(1) | WP_000846433.1 | WP_000846433.1 hypothetical protein |

| | | |
|---|---|---|
| 114(3) | WP_001990269.1 | WP_001990269.1 DUF1648 domain-containing protein |
| 114(1) | WP_000797171.1 | WP_000797171.1 DUF1648 domain-containing protein |
| 114(1) | WP_098955454.1 | WP_098955454.1 DUF1648 domain-containing protein |
| 114(1) | WP_060852182.1 | WP_060852182.1 DUF1648 domain-containing protein |
| 114(1) | WP_098874960.1 | WP_098874960.1 DUF1648 domain-containing protein |
| 114(1) | WP_070179255.1 | WP_070179255.1 DUF1648 domain-containing protein |
| 114(2) | WP_001065171.1 | WP_001065171.1 DUF1648 domain-containing protein |
| 114(1) | WP_153589152.1 | WP_153589152.1 DUF1648 domain-containing protein |
| 114(2) | WP_000797172.1 | WP_000797172.1 DUF1648 domain-containing protein |
| 114(2) | WP_030025424.1 | WP_030025424.1 DUF1648 domain-containing protein |
| 114(2) | WP_086385916.1 | WP_086385916.1 DUF1648 domain-containing protein |
| 114(2) | WP_048535501.1 | WP_048535501.1 DUF1648 domain-containing protein |
| 114(1) | WP_001999660.1 | WP_001999660.1 DUF1648 domain-containing protein |
| 114(1) | WP_098284011.1 | WP_098284011.1 DUF1648 domain-containing protein |
| 114(1) | WP_098395626.1 | WP_098395626.1 DUF1648 domain-containing protein |
| 114(1) | WP_098592731.1 | WP_098592731.1 DUF1648 domain-containing protein |
| 114(1) | WP_017763060.1 | WP_017763060.1 DUF1648 domain-containing protein |
| 114(1) | WP_061667215.1 | WP_061667215.1 DUF1648 domain-containing protein |
| 114(2) | WP_003300956.1 | WP_003300956.1 DUF1648 domain-containing protein |
| 114(1) | WP_016117662.1 | WP_016117662.1 DUF1648 domain-containing protein |
| 114(3) | WP_006923794.1 | WP_006923794.1 DUF1648 domain-containing protein |
| 114(4) | WP_086405914.1 | WP_086405914.1 DUF1648 domain-containing protein |
| 114(1) | WP_086412796.1 | WP_086412796.1 DUF1648 domain-containing protein |
| 114(1) | WP_208906812.1 | WP_208906812.1 DUF1648 domain-containing protein |
| 114(2) | WP_053513135.1 | WP_053513135.1 DUF1648 domain-containing protein |

| | | |
|---|---|---|
| 115(1) | WP_015736253.1 | WP_015736253.1 HAD-IB family hydrolase |
| 115(4) | WP_006211216.1 | WP_006211216.1 HAD-IB family hydrolase |
| 115(1) | WP_251497202.1 | WP_251497202.1 HAD family hydrolase |
| 115(1) | WP_106765967.1 | WP_106765967.1 HAD family hydrolase |
| 115(1) | WP_028405559.1 | WP_028405559.1 HAD family hydrolase |
| 115(1) | WP_090908271.1 | WP_090908271.1 HAD family hydrolase |
| 115(1) | WP_152826351.1 | WP_152826351.1 HAD family hydrolase |
| 115(1) | WP_260065248.1 | WP_260065248.1 HAD family hydrolase |
| 115(1) | WP_071220194.1 | WP_071220194.1 HAD-IB family hydrolase |
| 115(1) | WP_215153824.1 | WP_215153824.1 HAD family hydrolase |
| 115(1) | WP_192571314.1 | WP_192571314.1 HAD family hydrolase |
| 115(1) | WP_194543503.1 | WP_194543503.1 HAD family hydrolase |
| 115(1) | WP_220684634.1 | WP_220684634.1 HAD family hydrolase |
| 115(2) | WP_096774460.1 | WP_096774460.1 HAD family hydrolase |
| 115(1) | WP_098746883.1 | WP_098746883.1 HAD family hydrolase |
| 115(1) | WP_077567030.1 | WP_077567030.1 HAD family hydrolase |
| 115(1) | WP_054958397.1 | WP_054958397.1 HAD family hydrolase |
| 115(1) | WP_241929686.1 | WP_241929686.1 HAD family hydrolase |
| 115(2) | WP_213616445.1 | WP_213616445.1 HAD family hydrolase |
| 115(1) | WP_113058709.1 | WP_113058709.1 HAD family hydrolase |
| 115(1) | WP_213644220.1 | WP_213644220.1 HAD family hydrolase |
| 115(1) | WP_009590646.1 | WP_009590646.1 HAD-IB family hydrolase |
| 115(2) | WP_119847545.1 | WP_119847545.1 HAD family hydrolase |
| 115(1) | WP_208919003.1 | WP_208919003.1 HAD-IB family hydrolase |
| 115(1) | WP_076322122.1 | WP_076322122.1 HAD-IB family hydrolase |
| 115(1) | WP_216531924.1 | WP_216531924.1 HAD family hydrolase |
| 115(1) | WP_100537983.1 | WP_100537983.1 HAD family hydrolase |
| 115(1) | WP_127592434.1 | WP_127592434.1 HAD family hydrolase |
| 115(1) | WP_277746362.1 | WP_277746362.1 HAD family hydrolase |
| 115(1) | WP_063479997.1 | WP_063479997.1 HAD-IB family hydrolase |
| 115(1) | WP_007128642.1 | WP_007128642.1 HAD family hydrolase |
| 115(1) | WP_054402911.1 | WP_054402911.1 HAD family hydrolase |
| 115(1) | WP_145037668.1 | WP_145037668.1 HAD family hydrolase |

| | | |
|---|---|---|
| 116(1) | WP_175320745.1 | WP_175320745.1 decaprenyl-phosphate phosphoribosyltransferase |
| 116(1) | WP_015736252.1 | WP_015736252.1 decaprenyl-phosphate phosphoribosyltransferase |
| 116(1) | WP_285857328.1 | WP_285857328.1 decaprenyl-phosphate phosphoribosyltransferase |
| 116(2) | WP_208919005.1 | WP_208919005.1 decaprenyl-phosphate phosphoribosyltransferase |
| 116(1) | WP_259445206.1 | WP_259445206.1 decaprenyl-phosphate phosphoribosyltransferase |
| 116(3) | WP_007128641.1 | WP_007128641.1 decaprenyl-phosphate phosphoribosyltransferase |
| 116(1) | WP_194544867.1 | WP_194544867.1 decaprenyl-phosphate phosphoribosyltransferase |
| 116(2) | WP_119847546.1 | WP_119847546.1 decaprenyl-phosphate phosphoribosyltransferase |
| 116(1) | WP_077567029.1 | WP_077567029.1 decaprenyl-phosphate phosphoribosyltransferase |
| 116(1) | WP_113058710.1 | WP_113058710.1 decaprenyl-phosphate phosphoribosyltransferase |
| 116(1) | WP_216531925.1 | WP_216531925.1 decaprenyl-phosphate phosphoribosyltransferase |
| 116(1) | WP_280530304.1 | WP_280530304.1 decaprenyl-phosphate phosphoribosyltransferase |
| 116(1) | WP_174819211.1 | WP_174819211.1 decaprenyl-phosphate phosphoribosyltransferase |
| 116(1) | WP_215153825.1 | WP_215153825.1 decaprenyl-phosphate phosphoribosyltransferase |
| 116(1) | WP_207797473.1 | WP_207797473.1 decaprenyl-phosphate phosphoribosyltransferase |
| 116(1) | WP_090908272.1 | WP_090908272.1 decaprenyl-phosphate phosphoribosyltransferase |
| 116(2) | WP_071220195.1 | WP_071220195.1 decaprenyl-phosphate phosphoribosyltransferase |

| | | |
|---|---|---|
| 116(1) | WP_249732810.1 | WP_249732810.1 decaprenyl-phosphate phosphoribosyltransferase |
| 116(1) | WP_245640235.1 | WP_245640235.1 decaprenyl-phosphate phosphoribosyltransferase |
| 116(1) | WP_034273406.1 | WP_034273406.1 decaprenyl-phosphate phosphoribosyltransferase |
| 116(1) | WP_152826354.1 | WP_152826354.1 decaprenyl-phosphate phosphoribosyltransferase |
| 116(1) | WP_054402910.1 | WP_054402910.1 decaprenyl-phosphate phosphoribosyltransferase |
| 116(5) | WP_036642148.1 | WP_036642148.1 decaprenyl-phosphate phosphoribosyltransferase |
| 116(1) | WP_098746881.1 | WP_098746881.1 decaprenyl-phosphate phosphoribosyltransferase |
| 116(1) | WP_199794898.1 | WP_199794898.1 decaprenyl-phosphate phosphoribosyltransferase |
| 116(3) | WP_176502425.1 | WP_176502425.1 decaprenyl-phosphate phosphoribosyltransferase |
| 116(1) | WP_076322121.1 | WP_076322121.1 decaprenyl-phosphate phosphoribosyltransferase |
| 116(1) | WP_277746361.1 | WP_277746361.1 decaprenyl-phosphate phosphoribosyltransferase |
| 117(5) | WP_006211217.1 | WP_006211217.1 DMT family transporter |
| 117(1) | WP_100537984.1 | WP_100537984.1 EamA family transporter |
| 117(1) | WP_260065247.1 | WP_260065247.1 EamA/RhaT family transporter |
| 117(1) | WP_106765968.1 | WP_106765968.1 EamA family transporter |
| 117(1) | WP_152826350.1 | WP_152826350.1 EamA family transporter |
| 117(1) | WP_119847544.1 | WP_119847544.1 EamA family transporter |
| 117(1) | WP_304142097.1 | WP_304142097.1 EamA family transporter |
| 117(2) | WP_076322123.1 | WP_076322123.1 EamA family transporter |
| 117(1) | WP_054958396.1 | WP_054958396.1 EamA family transporter |
| 117(1) | WP_077567031.1 | WP_077567031.1 EamA family transporter |
| 117(1) | WP_090908270.1 | WP_090908270.1 EamA family transporter |
| 117(2) | WP_009590672.1 | WP_009590672.1 EamA family transporter |
| 117(1) | WP_096774461.1 | WP_096774461.1 EamA family transporter |
| 117(1) | WP_241929687.1 | WP_241929687.1 EamA family transporter |
| 117(1) | WP_144866416.1 | WP_144866416.1 EamA family transporter |
| 117(1) | WP_054402912.1 | WP_054402912.1 EamA family transporter |
| 117(1) | WP_098746885.1 | WP_098746885.1 hypothetical protein |
| 117(1) | WP_213644219.1 | WP_213644219.1 EamA/RhaT family transporter |
| 117(2) | WP_213579165.1 | WP_213579165.1 EamA family transporter |
| 117(1) | WP_071220193.1 | WP_071220193.1 hypothetical protein |
| 117(2) | WP_213616444.1 | WP_213616444.1 EamA family transporter |
| 117(1) | WP_028405560.1 | WP_028405560.1 EamA family transporter |
| 117(1) | WP_208919001.1 | WP_208919001.1 EamA family transporter |
| 117(1) | WP_113058708.1 | WP_113058708.1 EamA family transporter |
| 117(3) | WP_145037666.1 | WP_145037666.1 EamA family transporter |
| 117(1) | WP_194543504.1 | WP_194543504.1 EamA/RhaT family transporter |
| 117(3) | WP_127592435.1 | WP_127592435.1 EamA family transporter |
| 118(1) | WP_090908269.1 | WP_090908269.1 EamA family transporter |
| 118(1) | WP_152826348.1 | WP_152826348.1 EamA family transporter |
| 118(5) | WP_006211218.1 | WP_006211218.1 EamA family transporter |
| 118(1) | WP_077567032.1 | WP_077567032.1 EamA family transporter |
| 118(1) | WP_192571313.1 | WP_192571313.1 EamA family transporter |
| 118(1) | WP_054402913.1 | WP_054402913.1 EamA family transporter |
| 118(2) | WP_194543505.1 | WP_194543505.1 EamA family transporter |
| 118(1) | WP_113058707.1 | WP_113058707.1 EamA family transporter |
| 118(1) | WP_106765969.1 | WP_106765969.1 EamA family transporter |
| 118(4) | WP_076322124.1 | WP_076322124.1 EamA family transporter |
| 118(1) | WP_015736254.1 | WP_015736254.1 EamA family transporter |
| 118(1) | WP_054958395.1 | WP_054958395.1 EamA family transporter |
| 118(3) | WP_096774462.1 | WP_096774462.1 EamA family transporter |
| 118(3) | WP_007128644.1 | WP_007128644.1 EamA family transporter |
| 118(1) | WP_216531923.1 | WP_216531923.1 EamA family transporter |
| 118(1) | WP_036661597.1 | WP_036661597.1 EamA family transporter |
| 118(1) | WP_127592436.1 | WP_127592436.1 EamA family transporter |
| 118(1) | WP_119847543.1 | WP_119847543.1 EamA family transporter |
| 118(1) | WP_100537985.1 | WP_100537985.1 EamA family transporter |
| 118(1) | WP_208918999.1 | WP_208918999.1 EamA family transporter |
| 118(1) | WP_304142099.1 | WP_304142099.1 EamA family transporter |
| 118(1) | WP_036671549.1 | WP_036671549.1 EamA family transporter |
| 118(2) | WP_071220192.1 | WP_071220192.1 EamA family transporter |
| 118(1) | WP_028405561.1 | WP_028405561.1 DMT family transporter |
| 118(1) | WP_241929688.1 | WP_241929688.1 EamA family transporter |
| 118(1) | WP_145037664.1 | WP_145037664.1 EamA family transporter |
| 119(1) | WP_220684633.1 | WP_220684633.1 DUF6080 domain-containing protein |
| 119(1) | WP_213579166.1 | WP_213579166.1 DUF6080 domain-containing protein |
| 119(1) | WP_096774463.1 | WP_096774463.1 DUF6080 domain-containing protein |
| 119(1) | WP_009590676.1 | WP_009590676.1 DUF6080 domain-containing protein |
| 119(1) | WP_028405562.1 | WP_028405562.1 DUF6080 domain-containing protein |
| 119(1) | WP_304142101.1 | WP_304142101.1 DUF6080 domain-containing protein |
| 119(1) | WP_054402914.1 | WP_054402914.1 DUF6080 domain-containing protein |
| 119(1) | WP_215153823.1 | WP_215153823.1 DUF6080 domain-containing protein |
| 119(5) | WP_006211219.1 | WP_006211219.1 DUF6080 domain-containing protein |
| 119(1) | WP_260065245.1 | WP_260065245.1 DUF6080 domain-containing protein |
| 119(1) | WP_015736255.1 | WP_015736255.1 DUF6080 domain-containing protein |
| 119(1) | WP_054958394.1 | WP_054958394.1 DUF6080 domain-containing protein |
| 119(1) | WP_194543506.1 | WP_194543506.1 DUF6080 domain-containing protein |
| 119(1) | WP_216531922.1 | WP_216531922.1 DUF6080 domain-containing protein |
| 119(1) | WP_076322125.1 | WP_076322125.1 DUF6080 domain-containing protein |
| 119(1) | WP_090908268.1 | WP_090908268.1 DUF6080 domain-containing protein |
| 119(1) | WP_119847542.1 | WP_119847542.1 DUF6080 domain-containing protein |
| 119(1) | WP_106765970.1 | WP_106765970.1 DUF6080 domain-containing protein |
| 119(1) | WP_098746887.1 | WP_098746887.1 DUF6080 domain-containing protein |
| 119(1) | WP_127592437.1 | WP_127592437.1 DUF6080 domain-containing protein |
| 119(1) | WP_251497205.1 | WP_251497205.1 DUF6080 domain-containing protein |
| 119(1) | WP_071220191.1 | WP_071220191.1 DUF6080 domain-containing protein |
| 119(1) | WP_145037662.1 | WP_145037662.1 DUF6080 domain-containing protein |
| 119(1) | WP_244563025.1 | WP_244563025.1 DUF6080 domain-containing protein |
| 119(1) | WP_077567033.1 | WP_077567033.1 DUF6080 domain-containing protein |
| 119(1) | WP_100537986.1 | WP_100537986.1 DUF6080 domain-containing protein |
| 119(1) | WP_213644218.1 | WP_213644218.1 DUF6080 domain-containing protein |
| 119(1) | WP_076150485.1 | WP_076150485.1 DUF6080 domain-containing protein |
| 119(3) | WP_210095109.1 | WP_210095109.1 DUF6080 domain-containing protein |
| 119(1) | WP_241929689.1 | WP_241929689.1 DUF6080 domain-containing protein |
| 119(1) | WP_277746363.1 | WP_277746363.1 DUF6080 domain-containing protein |
| 119(1) | WP_113058706.1 | WP_113058706.1 DUF6080 domain-containing protein |
| 119(1) | WP_192571312.1 | WP_192571312.1 DUF6080 domain-containing protein |
| 120(1) | WP_111273186.1 | WP_111273186.1 sigma factor |
| 120(1) | WP_293755514.1 | WP_293755514.1 sigma-70 family RNA polymerase sigma factor |
| 120(1) | WP_110899384.1 | WP_110899384.1 sigma-70 family RNA polymerase sigma factor |

120(1) WP_113672707.1 WP_113672707.1 sigma-70 family RNA polymerase sigma factor
120(1) WP_189018618.1 WP_189018618.1 RNA polymerase sigma factor
120(1) WP_109625257.1 WP_109625257.1 sigma-70 family RNA polymerase sigma factor
120(1) WP_191798520.1 WP_191798520.1 sigma-70 family RNA polymerase sigma factor
120(1) WP_108464075.1 WP_108464075.1 sigma-70 family RNA polymerase sigma factor
120(1) WP_307349988.1 WP_307349988.1 sigma factor
120(1) WP_075154086.1 WP_075154086.1 sigma-70 family RNA polymerase sigma factor
120(1) WP_150364179.1 WP_150364179.1 sigma-70 family RNA polymerase sigma factor
120(1) WP_274485160.1 WP_274485160.1 sigma-70 family RNA polymerase sigma factor
120(2) WP_281813109.1 WP_281813109.1 sigma-70 family RNA polymerase sigma factor
120(1) WP_076210938.1 WP_076210938.1 sigma-70 family RNA polymerase sigma factor
120(1) WP_211745846.1 WP_211745846.1 DUF4367 domain-containing protein
120(1) WP_245850666.1 WP_245850666.1 sigma-70 family RNA polymerase sigma factor
120(1) WP_185165658.1 WP_185165658.1 RNA polymerase sigma factor
120(1) WP_184183069.1 WP_184183069.1 sigma-70 family RNA polymerase sigma factor
120(1) WP_069011426.1 WP_069011426.1 hypothetical protein
120(1) WP_285749225.1 WP_285749225.1 DUF4367 domain-containing protein
120(1) WP_043069426.1 WP_043069426.1 RNA polymerase sigma-70 factor
120(1) WP_148409460.1 WP_148409460.1 sigma-70 family RNA polymerase sigma factor
120(1) WP_273321538.1 WP_273321538.1 sigma-70 family RNA polymerase sigma factor
120(2) WP_216518280.1 WP_216518280.1 sigma-70 family RNA polymerase sigma factor
120(1) WP_212695792.1 WP_212695792.1 sigma-70 family RNA polymerase sigma factor
120(1) WP_304942557.1 WP_304942557.1 sigma-70 family RNA polymerase sigma factor
120(1) WP_039273518.1 WP_039273518.1 sigma-70 family RNA polymerase sigma factor
120(1) WP_248063412.1 WP_248063412.1 sigma-70 family RNA polymerase sigma factor
120(1) WP_191798521.1 WP_191798521.1 DUF4367 domain-containing protein
120(1) WP_173405262.1 WP_173405262.1 sigma-70 family RNA polymerase sigma factor
120(3) WP_077832274.1 WP_077832274.1 sigma-70 family RNA polymerase sigma factor
120(1) WP_164965492.1 WP_164965492.1 sigma-70 family RNA polymerase sigma factor
120(1) WP_213532652.1 WP_213532652.1 DUF4367 domain-containing protein
120(1) WP_230585139.1 WP_230585139.1 hypothetical protein
120(1) WP_212693243.1 WP_212693243.1 sigma-70 family RNA polymerase sigma factor

121(7) WP_031309866.1 WP_031309866.1 ammonia-dependent NAD(+) synthetase
121(1) WP_304502593.1 WP_304502593.1 ammonia-dependent NAD(+) synthetase
121(1) WP_064019334.1 WP_064019334.1 ammonia-dependent NAD(+) synthetase
121(1) WP_000174889.1 WP_000174889.1 ammonia-dependent NAD(+) synthetase
121(2) WP_003340537.1 WP_003340537.1 ammonia-dependent NAD(+) synthetase
121(1) WP_199794085.1 WP_199794085.1 ammonia-dependent NAD(+) synthetase
121(5) WP_094699661.1 WP_094699661.1 ammonia-dependent NAD(+) synthetase
121(1) WP_003336008.1 WP_003336008.1 ammonia-dependent NAD(+) synthetase
121(2) WP_018670528.1 WP_018670528.1 ammonia-dependent NAD(+) synthetase
121(1) WP_158330199.1 WP_158330199.1 ammonia-dependent NAD(+) synthetase
121(2) WP_104030448.1 WP_104030448.1 ammonia-dependent NAD(+) synthetase
121(1) WP_119732865.1 WP_119732865.1 ammonia-dependent NAD(+) synthetase
121(1) WP_181747859.1 WP_181747859.1 ammonia-dependent NAD(+) synthetase
121(1) WP_158326784.1 WP_158326784.1 ammonia-dependent NAD(+) synthetase
121(2) WP_113755559.1 WP_113755559.1 ammonia-dependent NAD(+) synthetase
121(1) WP_068793959.1 WP_068793959.1 ammonia-dependent NAD(+) synthetase
121(1) WP_104065525.1 WP_104065525.1 ammonia-dependent NAD(+) synthetase
121(1) WP_301804646.1 WP_301804646.1 ammonia-dependent NAD(+) synthetase
121(2) WP_031415567.1 WP_031415567.1 ammonia-dependent NAD(+) synthetase
121(2) WP_104148790.1 WP_104148790.1 ammonia-dependent NAD(+) synthetase
121(2) WP_197247503.1 WP_197247503.1 ammonia-dependent NAD(+) synthetase

122(1) WP_099354919.1 WP_099354919.1 YitT family protein
122(1) WP_098226400.1 WP_098226400.1 YitT family protein
122(2) WP_086405916.1 WP_086405916.1 YitT family protein
122(3) WP_000831972.1 WP_000831972.1 YitT family protein
122(1) WP_136379709.1 WP_136379709.1 YitT family protein
122(1) WP_098284008.1 WP_098284008.1 YitT family protein
122(1) WP_229139928.1 WP_229139928.1 YitT family protein
122(1) WP_098157703.1 WP_098157703.1 YitT family protein
122(1) WP_098327468.1 WP_098327468.1 YitT family protein
122(1) WP_122971521.1 WP_122971521.1 YitT family protein
122(1) WP_098612021.1 WP_098612021.1 YitT family protein
122(8) WP_000755126.1 WP_000755126.1 YitT family protein
122(1) WP_098141780.1 WP_098141780.1 YitT family protein
122(1) WP_116353240.1 WP_116353240.1 YitT family protein
122(1) WP_000755127.1 WP_000755127.1 YitT family protein
122(1) WP_030025417.1 WP_030025417.1 YitT family protein
122(1) WP_061676411.1 WP_061676411.1 YitT family protein
122(1) WP_098535851.1 WP_098535851.1 YitT family protein
122(9) WP_000755125.1 WP_000755125.1 YitT family protein

123(4) WP_031415582.1 WP_031415582.1 MTH1187 family thiamine-binding protein
123(1) WP_232699389.1 WP_232699389.1 MTH1187 family thiamine-binding protein
123(4) WP_113755273.1 WP_113755273.1 MTH1187 family thiamine-binding protein
123(3) WP_022586764.1 WP_022586764.1 MTH1187 family thiamine-binding protein
123(1) WP_096886741.1 WP_096886741.1 MTH1187 family thiamine-binding protein
123(1) WP_258425846.1 WP_258425846.1 MTH1187 family thiamine-binding protein
123(2) WP_104030436.1 WP_104030436.1 MTH1187 family thiamine-binding protein
123(3) WP_094699649.1 WP_094699649.1 MTH1187 family thiamine-binding protein
123(1) WP_121474482.1 WP_121474482.1 MTH1187 family thiamine-binding protein
123(1) WP_181747873.1 WP_181747873.1 MTH1187 family thiamine-binding protein
123(1) WP_104065537.1 WP_104065537.1 MTH1187 family thiamine-binding protein
123(1) WP_258433021.1 WP_258433021.1 MTH1187 family thiamine-binding protein
123(4) WP_003340557.1 WP_003340557.1 MTH1187 family thiamine-binding protein
123(9) WP_018670515.1 WP_018670515.1 MTH1187 family thiamine-binding protein
123(1) WP_003335994.1 WP_003335994.1 MTH1187 family thiamine-binding protein

124(1) WP_049869174.1 WP_049869174.1 DUF2004 domain-containing protein
124(1) WP_270728532.1 WP_270728532.1 DUF2004 domain-containing protein
124(1) WP_063223283.1 WP_063223283.1 DUF2004 domain-containing protein
124(1) WP_258126016.1 WP_258126016.1 DUF2004 domain-containing protein
124(1) WP_144655907.1 WP_144655907.1 DUF2004 domain-containing protein
124(1) WP_071724732.1 WP_071724732.1 DUF2004 domain-containing protein
124(2) WP_002141600.1 WP_002141600.1 DUF2004 domain-containing protein
124(3) WP_071759431.1 WP_071759431.1 DUF2004 domain-containing protein
124(1) WP_000154329.1 WP_000154329.1 DUF2004 domain-containing protein
124(1) WP_103655290.1 WP_103655290.1 DUF2004 domain-containing protein
124(3) WP_021291843.1 WP_021291843.1 hypothetical protein
124(1) WP_000700048.1 WP_000700048.1 DUF2004 domain-containing protein
124(1) WP_193653015.1 WP_193653015.1 DUF2004 domain-containing protein

124(1) WP_044792549.1 WP_044792549.1 DUF2004 domain-containing protein
124(1) WP_025148811.1 WP_025148811.1 DUF2004 domain-containing protein
124(6) WP_048552315.1 WP_048552315.1 DUF2004 domain-containing protein
124(1) WP_076301100.1 WP_076301100.1 DUF2004 domain-containing protein
124(10) WP_185350825.1 WP_185350825.1 hypothetical protein

125(1) WP_098250529.1 WP_098250529.1 pentapeptide repeat-containing protein
125(1) WP_036702404.1 WP_036702404.1 pentapeptide repeat-containing protein
125(1) WP_282446270.1 WP_282446270.1 pentapeptide repeat-containing protein
125(1) WP_146236237.1 WP_146236237.1 pentapeptide repeat-containing protein
125(1) WP_184182980.1 WP_184182980.1 pentapeptide repeat-containing protein
125(2) WP_185340333.1 WP_185340333.1 pentapeptide repeat-containing protein
125(1) WP_098562448.1 WP_098562448.1 pentapeptide repeat-containing protein
125(2) WP_185475841.1 WP_185475841.1 pentapeptide repeat-containing protein
125(1) WP_081943723.1 WP_081943723.1 pentapeptide repeat-containing protein
125(4) WP_041176140.1 WP_041176140.1 pentapeptide repeat-containing protein
125(2) WP_270993669.1 WP_270993669.1 pentapeptide repeat-containing protein
125(2) WP_185639003.1 WP_185639003.1 pentapeptide repeat-containing protein
125(4) WP_203573364.1 WP_203573364.1 pentapeptide repeat-containing protein
125(1) WP_029438097.1 WP_029438097.1 pentapeptide repeat-containing protein
125(3) WP_046325947.1 WP_046325947.1 pentapeptide repeat-containing protein
125(4) WP_090976437.1 WP_090976437.1 pentapeptide repeat-containing protein
125(2) WP_149051229.1 WP_149051229.1 pentapeptide repeat-containing protein
125(1) WP_076157818.1 WP_076157818.1 pentapeptide repeat-containing protein
125(2) WP_185350592.1 WP_185350592.1 pentapeptide repeat-containing protein
125(1) WP_286119525.1 WP_286119525.1 pentapeptide repeat-containing protein

126(1) WP_121474474.1 WP_121474474.1 lipoate--protein ligase
126(1) WP_258418577.1 WP_258418577.1 lipoate--protein ligase
126(1) WP_181747861.1 WP_181747861.1 lipoate--protein ligase
126(3) WP_116334075.1 WP_116334075.1 lipoate--protein ligase
126(1) WP_096886733.1 WP_096886733.1 lipoate--protein ligase
126(4) WP_140574917.1 WP_140574917.1 lipoate--protein ligase
126(2) WP_158329532.1 WP_158329532.1 lipoate--protein ligase
126(2) WP_003340539.1 WP_003340539.1 lipoate--protein ligase
126(1) WP_197262002.1 WP_197262002.1 lipoate--protein ligase
126(2) WP_119732864.1 WP_119732864.1 lipoate--protein ligase
126(3) WP_104030447.1 WP_104030447.1 lipoate--protein ligase
126(2) WP_113755283.1 WP_113755283.1 lipoate--protein ligase
126(1) WP_301804648.1 WP_301804648.1 lipoate--protein ligase
126(2) WP_212930971.1 WP_212930971.1 lipoate--protein ligase
126(1) WP_232699379.1 WP_232699379.1 lipoate--protein ligase
126(1) WP_094699660.1 WP_094699660.1 lipoate--protein ligase
126(1) WP_277548448.1 WP_277548448.1 lipoate--protein ligase
126(1) WP_068793927.1 WP_068793927.1 lipoate--protein ligase
126(1) WP_018670527.1 WP_018670527.1 lipoate--protein ligase
126(1) WP_003336007.1 WP_003336007.1 lipoate--protein ligase
126(1) WP_163245176.1 WP_163245176.1 lipoate--protein ligase
126(1) WP_104065526.1 WP_104065526.1 lipoate--protein ligase
126(2) WP_022586773.1 WP_022586773.1 lipoate--protein ligase
126(1) WP_064019325.1 WP_064019325.1 lipoate--protein ligase

127(1) WP_163245192.1 WP_163245192.1 isoprenylcysteine carboxylmethyltransferase family protein
127(1) WP_158330735.1 WP_158330735.1 isoprenylcysteine carboxylmethyltransferase family protein
127(1) WP_003335991.1 WP_003335991.1 isoprenylcysteine carboxylmethyltransferase family protein
127(1) WP_258418568.1 WP_258418568.1 isoprenylcysteine carboxylmethyltransferase family protein
127(2) WP_113755271.1 WP_113755271.1 isoprenylcysteine carboxylmethyltransferase family protein
127(1) WP_104030435.1 WP_104030435.1 isoprenylcysteine carboxylmethyltransferase family protein
127(2) WP_068793936.1 WP_068793936.1 isoprenylcysteine carboxylmethyltransferase family protein
127(1) WP_104065539.1 WP_104065539.1 isoprenylcysteine carboxylmethyltransferase family protein
127(2) WP_018670512.1 WP_018670512.1 isoprenylcysteine carboxylmethyltransferase family protein
127(1) WP_239209293.1 WP_239209293.1 isoprenylcysteine carboxylmethyltransferase family protein
127(1) WP_197247520.1 WP_197247520.1 isoprenylcysteine carboxylmethyltransferase family protein
127(1) WP_031415586.1 WP_031415586.1 isoprenylcysteine carboxylmethyltransferase family protein
127(2) WP_119732855.1 WP_119732855.1 isoprenylcysteine carboxylmethyltransferase family protein
127(2) WP_094699647.1 WP_094699647.1 isoprenylcysteine carboxylmethyltransferase family protein
127(1) WP_022586761.1 WP_022586761.1 isoprenylcysteine carboxylmethyltransferase family protein
127(1) WP_232699393.1 WP_232699393.1 isoprenylcysteine carboxylmethyltransferase family protein
127(1) WP_199794080.1 WP_199794080.1 isoprenylcysteine carboxylmethyltransferase family protein
127(1) WP_301804690.1 WP_301804690.1 isoprenylcysteine carboxylmethyltransferase family protein
127(1) WP_116334083.1 WP_116334083.1 isoprenylcysteine carboxylmethyltransferase family protein
127(1) WP_104148797.1 WP_104148797.1 isoprenylcysteine carboxylmethyltransferase family protein
127(1) WP_212930982.1 WP_212930982.1 isoprenylcysteine carboxylmethyltransferase family protein
127(1) WP_064019312.1 WP_064019312.1 isoprenylcysteine carboxylmethyltransferase family protein
127(1) WP_158329538.1 WP_158329538.1 isoprenylcysteine carboxylmethyltransferase family protein
127(1) WP_101668491.1 WP_101668491.1 isoprenylcysteine carboxyl methyltransferase
127(2) WP_003340563.1 WP_003340563.1 isoprenylcysteine carboxylmethyltransferase family protein
127(1) WP_181747875.1 WP_181747875.1 isoprenylcysteine carboxylmethyltransferase family protein
127(1) WP_278431131.1 WP_278431131.1 isoprenylcysteine carboxylmethyltransferase family protein
127(1) WP_258427453.1 WP_258427453.1 isoprenylcysteine carboxylmethyltransferase family protein
127(1) WP_277548430.1 WP_277548430.1 isoprenylcysteine carboxylmethyltransferase family protein
127(1) WP_096886743.1 WP_096886743.1 isoprenylcysteine carboxylmethyltransferase family protein
127(1) WP_258433023.1 WP_258433023.1 isoprenylcysteine carboxylmethyltransferase family protein

128(1) WP_003340566.1 WP_003340566.1 branched-chain amino acid transport system II carrier protein
128(1) WP_278431132.1 WP_278431132.1 branched-chain amino acid transport system II carrier protein
128(2) WP_104030433.1 WP_104030433.1 branched-chain amino acid transport system II carrier protein
128(1) WP_022586759.1 WP_022586759.1 branched-chain amino acid transport system II carrier protein
128(1) WP_116334085.1 WP_116334085.1 branched-chain amino acid transport system II carrier protein
128(2) WP_158329539.1 WP_158329539.1 branched-chain amino acid transport system II carrier protein
128(2) WP_181747877.1 WP_181747877.1 branched-chain amino acid transport system II carrier protein
128(1) WP_197247524.1 WP_197247524.1 branched-chain amino acid transport system II carrier protein
128(1) WP_232699395.1 WP_232699395.1 branched-chain amino acid transport system II carrier protein
128(1) WP_094699645.1 WP_094699645.1 branched-chain amino acid transport system II carrier protein

128(2) WP_113755269.1 WP_113755269.1 branched-chain amino acid transport system II carrier protein
128(1) WP_258433025.1 WP_258433025.1 branched-chain amino acid transport system II carrier protein
128(4) WP_018670510.1 WP_018670510.1 branched-chain amino acid transport system II carrier protein
128(1) WP_068793938.1 WP_068793938.1 branched-chain amino acid transport system II carrier protein
128(2) WP_031415588.1 WP_031415588.1 branched-chain amino acid transport system II carrier protein
128(1) WP_101668493.1 WP_101668493.1 branched-chain amino acid transport system II carrier protein
128(1) WP_199794079.1 WP_199794079.1 branched-chain amino acid transport system II carrier protein
128(1) WP_064019310.1 WP_064019310.1 branched-chain amino acid transport system II carrier protein
128(1) WP_197262010.1 WP_197262010.1 branched-chain amino acid transport system II carrier protein
128(3) WP_096886745.1 WP_096886745.1 branched-chain amino acid transport system II carrier protein
128(1) WP_239209294.1 WP_239209294.1 branched-chain amino acid transport system II carrier protein
128(1) WP_003335988.1 WP_003335988.1 branched-chain amino acid transport system II carrier protein
128(2) WP_119732853.1 WP_119732853.1 branched-chain amino acid transport system II carrier protein
128(2) WP_163245194.1 WP_163245194.1 branched-chain amino acid transport system II carrier protein
128(1) WP_301804691.1 WP_301804691.1 branched-chain amino acid transport system II carrier protein

129(1) WP_277548428.1 WP_277548428.1 3-oxoacyl-
129(3) WP_197247522.1 WP_197247522.1 3-oxoacyl-
129(1) WP_104148798.1 WP_104148798.1 3-oxoacyl-
129(1) WP_116334084.1 WP_116334084.1 type III polyketide synthase
129(2) WP_212931076.1 WP_212931076.1 3-oxoacyl-
129(1) WP_119732854.1 WP_119732854.1 3-oxoacyl-
129(1) WP_104030434.1 WP_104030434.1 3-oxoacyl-
129(1) WP_101668492.1 WP_101668492.1 type III polyketide synthase
129(1) WP_232699394.1 WP_232699394.1 3-oxoacyl-
129(3) WP_094699646.1 WP_094699646.1 3-oxoacyl-
129(1) WP_104065540.1 WP_104065540.1 3-oxoacyl-
129(1) WP_181747925.1 WP_181747925.1 3-oxoacyl-
129(1) WP_064019311.1 WP_064019311.1 3-oxoacyl-
129(3) WP_096886744.1 WP_096886744.1 3-oxoacyl-
129(2) WP_018670511.1 WP_018670511.1 3-oxoacyl-
129(3) WP_158326746.1 WP_158326746.1 3-oxoacyl-
129(1) WP_258433024.1 WP_258433024.1 type III polyketide synthase
129(2) WP_003335989.1 WP_003335989.1 3-oxoacyl-
129(2) WP_022586760.1 WP_022586760.1 3-oxoacyl-
129(1) WP_035318350.1 WP_035318350.1 3-oxoacyl-
129(1) WP_113755270.1 WP_113755270.1 3-oxoacyl-
129(1) WP_278431166.1 WP_278431166.1 3-oxoacyl-
129(1) WP_068793937.1 WP_068793937.1 3-oxoacyl-
129(1) WP_301804869.1 WP_301804869.1 3-oxoacyl-
129(1) WP_168419974.1 WP_168419974.1 3-oxoacyl-

130(1) WP_034526657.1 WP_034526657.1 DeoR/GlpR family DNA-binding transcription regulator
130(1) WP_021291860.1 WP_021291860.1 DeoR family transcriptional regulator
130(1) WP_077320192.1 WP_077320192.1 DeoR family transcriptional regulator
130(2) WP_072727818.1 WP_072727818.1 DeoR/GlpR family DNA-binding transcription regulator
130(1) WP_065291255.1 WP_065291255.1 DeoR/GlpR family DNA-binding transcription regulator
130(1) WP_306465034.1 WP_306465034.1 DeoR family transcriptional regulator
130(1) WP_038245458.1 WP_038245458.1 DeoR family transcriptional regulator
130(1) WP_272541584.1 WP_272541584.1 DeoR/GlpR family DNA-binding transcription regulator
130(1) WP_268583851.1 WP_268583851.1 DeoR/GlpR family DNA-binding transcription regulator
130(3) WP_262866583.1 WP_262866583.1 DeoR/GlpR family DNA-binding transcription regulator
130(2) WP_231586854.1 WP_231586854.1 hypothetical protein
130(2) WP_019392121.1 WP_019392121.1 DeoR/GlpR family DNA-binding transcription regulator
130(1) WP_289355907.1 WP_289355907.1 DeoR/GlpR family DNA-binding transcription regulator
130(1) WP_286191345.1 WP_286191345.1 DeoR family transcriptional regulator
130(3) WP_021253127.1 WP_021253127.1 DeoR/GlpR family DNA-binding transcription regulator
130(1) WP_138186703.1 WP_138186703.1 DeoR/GlpR family DNA-binding transcription regulator
130(4) WP_163977063.1 WP_163977063.1 DeoR/GlpR family DNA-binding transcription regulator
130(1) WP_124049164.1 WP_124049164.1 DeoR/GlpR family DNA-binding transcription regulator
130(2) WP_231586855.1 WP_231586855.1 hypothetical protein
130(1) WP_243518767.1 WP_243518767.1 DeoR/GlpR family DNA-binding transcription regulator
130(1) WP_077833120.1 WP_077833120.1 DeoR/GlpR family DNA-binding transcription regulator
130(1) WP_090510235.1 WP_090510235.1 DeoR/GlpR family DNA-binding transcription regulator
130(2) WP_077893589.1 WP_077893589.1 DeoR/GlpR family DNA-binding transcription regulator
130(1) WP_021261892.1 WP_021261892.1 DeoR/GlpR family DNA-binding transcription regulator
130(1) WP_195953734.1 WP_195953734.1 DeoR/GlpR family DNA-binding transcription regulator

131(1) WP_206094032.1 WP_206094032.1 DUF402 domain-containing protein
131(4) WP_163977052.1 WP_163977052.1 DUF402 domain-containing protein
131(3) WP_168178782.1 WP_168178782.1 DUF402 domain-containing protein
131(1) WP_111153298.1 WP_111153298.1 DUF402 domain-containing protein
131(1) WP_281477424.1 WP_281477424.1 DUF402 domain-containing protein
131(1) WP_006674856.1 WP_006674856.1 DUF402 domain-containing protein
131(1) WP_233697180.1 WP_233697180.1 DUF402 domain-containing protein
131(2) WP_143801056.1 WP_143801056.1 DUF402 domain-containing protein
131(1) WP_138186687.1 WP_138186687.1 DUF402 domain-containing protein
131(1) WP_065291243.1 WP_065291243.1 DUF402 domain-containing protein
131(1) WP_272045566.1 WP_272045566.1 DUF402 domain-containing protein
131(2) WP_087433612.1 WP_087433612.1 DUF402 domain-containing protein
131(1) WP_244194182.1 WP_244194182.1 DUF402 domain-containing protein
131(1) WP_142543123.1 WP_142543123.1 DUF402 domain-containing protein
131(1) WP_019424140.1 WP_019424140.1 DUF402 domain-containing protein
131(4) WP_021253112.1 WP_021253112.1 DUF402 domain-containing protein
131(1) WP_268619939.1 WP_268619939.1 DUF402 domain-containing protein
131(1) WP_072727827.1 WP_072727827.1 DUF402 domain-containing protein
131(1) WP_021261877.1 WP_021261877.1 DUF402 domain-containing protein
131(3) WP_005546525.1 WP_005546525.1 DUF402 domain-containing protein
131(1) WP_213426986.1 WP_213426986.1 DUF402 domain-containing protein

131(1) WP_119794399.1 WP_119794399.1 DUF402 domain-containing protein
131(1) WP_133378767.1 WP_133378767.1 DUF402 domain-containing protein
131(1) WP_197256856.1 WP_197256856.1 DUF402 domain-containing protein

132(2) WP_019424157.1 WP_019424157.1 BMC domain-containing protein
132(1) WP_112332011.1 WP_112332011.1 BMC domain-containing protein
132(1) WP_019424160.1 WP_019424160.1 BMC domain-containing protein
132(1) WP_206094040.1 WP_206094040.1 BMC domain-containing protein
132(3) WP_087433597.1 WP_087433597.1 BMC domain-containing protein
132(2) WP_019424156.1 WP_019424156.1 BMC domain-containing protein
132(2) WP_112331599.1 WP_112331599.1 BMC domain-containing protein
132(1) WP_146740513.1 WP_146740513.1 BMC domain-containing protein
132(3) WP_087433595.1 WP_087433595.1 BMC domain-containing protein
132(1) WP_233697164.1 WP_233697164.1 BMC domain-containing protein
132(3) WP_087433593.1 WP_087433593.1 BMC domain-containing protein
132(1) WP_286206877.1 WP_286206877.1 BMC domain-containing protein
132(2) WP_112331594.1 WP_112331594.1 BMC domain-containing protein
132(1) WP_112331598.1 WP_112331598.1 BMC domain-containing protein
132(1) WP_233697166.1 WP_233697166.1 BMC domain-containing protein
132(2) WP_087433594.1 WP_087433594.1 BMC domain-containing protein
132(1) WP_019424158.1 WP_019424158.1 BMC domain-containing protein
132(1) WP_112332012.1 WP_112332012.1 BMC domain-containing protein
132(3) WP_087433596.1 WP_087433596.1 BMC domain-containing protein
132(1) WP_233697165.1 WP_233697165.1 BMC domain-containing protein
132(1) WP_019424159.1 WP_019424159.1 BMC domain-containing protein
132(2) WP_112332010.1 WP_112332010.1 BMC domain-containing protein

133(3) WP_113755560.1 WP_113755560.1 sulfur carrier protein ThiS
133(1) WP_277548455.1 WP_277548455.1 sulfur carrier protein ThiS
133(1) WP_068793925.1 WP_068793925.1 sulfur carrier protein ThiS
133(1) WP_301804633.1 WP_301804633.1 sulfur carrier protein ThiS
133(2) WP_242068643.1 WP_242068643.1 sulfur carrier protein ThiS
133(1) WP_242066321.1 WP_242066321.1 sulfur carrier protein ThiS
133(1) WP_258073319.1 WP_258073319.1 sulfur carrier protein ThiS
133(1) WP_064019330.1 WP_064019330.1 sulfur carrier protein ThiS, partial
133(3) WP_236645788.1 WP_236645788.1 sulfur carrier protein ThiS
133(1) WP_258425835.1 WP_258425835.1 sulfur carrier protein ThiS
133(7) WP_022586779.1 WP_022586779.1 sulfur carrier protein ThiS
133(1) WP_278431117.1 WP_278431117.1 sulfur carrier protein ThiS
133(1) WP_026315056.1 WP_026315056.1 sulfur carrier protein ThiS
133(1) WP_003340528.1 WP_003340528.1 sulfur carrier protein ThiS
133(2) WP_212930967.1 WP_212930967.1 sulfur carrier protein ThiS
133(1) WP_258074288.1 WP_258074288.1 sulfur carrier protein ThiS
133(2) WP_258072913.1 WP_258072913.1 sulfur carrier protein ThiS
133(1) WP_236847616.1 WP_236847616.1 sulfur carrier protein ThiS
133(1) WP_158330198.1 WP_158330198.1 sulfur carrier protein ThiS
133(1) WP_232699370.1 WP_232699370.1 sulfur carrier protein ThiS
133(2) WP_239586184.1 WP_239586184.1 sulfur carrier protein ThiS
133(1) WP_242061310.1 WP_242061310.1 sulfur carrier protein ThiS

134(9) WP_003336012.1 WP_003336012.1 hypothetical protein
134(5) WP_240962077.1 WP_240962077.1 hypothetical protein
134(2) WP_064019328.1 WP_064019328.1 hypothetical protein
134(2) WP_239586183.1 WP_239586183.1 hypothetical protein
134(1) WP_232699372.1 WP_232699372.1 hypothetical protein
134(2) WP_246329043.1 WP_246329043.1 hypothetical protein
134(1) WP_031415562.1 WP_031415562.1 hypothetical protein
134(4) WP_113755286.1 WP_113755286.1 hypothetical protein
134(1) WP_301804638.1 WP_301804638.1 hypothetical protein
134(9) WP_018670532.1 WP_018670532.1 hypothetical protein

135(1) WP_168419969.1 WP_168419969.1 UbiD family decarboxylase
135(1) WP_258433013.1 WP_258433013.1 UbiD family decarboxylase
135(1) WP_239209288.1 WP_239209288.1 UbiD family decarboxylase
135(1) WP_158326753.1 WP_158326753.1 UbiD family decarboxylase
135(1) WP_212930970.1 WP_212930970.1 UbiD family decarboxylase
135(1) WP_140574921.1 WP_140574921.1 UbiD family decarboxylase
135(1) WP_258425837.1 WP_258425837.1 UbiD family decarboxylase
135(1) WP_003340535.1 WP_003340535.1 UbiD family decarboxylase
135(2) WP_094699662.1 WP_094699662.1 UbiD family decarboxylase
135(1) WP_158330730.1 WP_158330730.1 UbiD family decarboxylase
135(6) WP_003336009.1 WP_003336009.1 UbiD family decarboxylase
135(2) WP_113755284.1 WP_113755284.1 UbiD family decarboxylase
135(3) WP_018670529.1 WP_018670529.1 UbiD family decarboxylase
135(2) WP_104030449.1 WP_104030449.1 UbiD family decarboxylase
135(1) WP_064019326.1 WP_064019326.1 UbiD family decarboxylase
135(2) WP_116334074.1 WP_116334074.1 UbiD family decarboxylase
135(2) WP_197247498.1 WP_197247498.1 UbiD family decarboxylase
135(1) WP_163245175.1 WP_163245175.1 UbiD family decarboxylase
135(1) WP_258417125.1 WP_258417125.1 UbiD family decarboxylase
135(1) WP_022586775.1 WP_022586775.1 UbiD family decarboxylase
135(1) WP_121474473.1 WP_121474473.1 UbiD family decarboxylase
135(1) WP_119732866.1 WP_119732866.1 UbiD family decarboxylase
135(1) WP_301804644.1 WP_301804644.1 UbiD family decarboxylase
135(1) WP_232699374.1 WP_232699374.1 UbiD family decarboxylase

136(1) WP_104065535.1 WP_104065535.1 Na+/H+ antiporter NhaC family protein
136(5) WP_094699651.1 WP_094699651.1 SLC13 family permease
136(1) WP_113755275.1 WP_113755275.1 Na+/H+ antiporter NhaC family protein
136(6) WP_096886739.1 WP_096886739.1 Na+/H+ antiporter NhaC family protein
136(1) WP_168419973.1 WP_168419973.1 Na+/H+ antiporter NhaC family protein
136(1) WP_068793933.1 WP_068793933.1 Na+/H+ antiporter NhaC family protein
136(2) WP_158329537.1 WP_158329537.1 Na+/H+ antiporter NhaC family protein
136(1) WP_022586766.1 WP_022586766.1 Na+/H+ antiporter NhaC family protein
136(1) WP_199794082.1 WP_199794082.1 Na+/H+ antiporter NhaC family protein
136(1) WP_003335996.1 WP_003335996.1 Na+/H+ antiporter NhaC family protein
136(2) WP_163245187.1 WP_163245187.1 Na+/H+ antiporter NhaC family protein
136(1) WP_197247516.1 WP_197247516.1 Na+/H+ antiporter NhaC family protein
136(1) WP_158326748.1 WP_158326748.1 Na+/H+ antiporter NhaC family protein
136(5) WP_104030438.1 WP_104030438.1 Na+/H+ antiporter NhaC family protein
136(1) WP_301804675.1 WP_301804675.1 Na+/H+ antiporter NhaC family protein
136(1) WP_018670517.1 WP_018670517.1 Na+/H+ antiporter NhaC family protein
136(1) WP_003340553.1 WP_003340553.1 Na+/H+ antiporter NhaC family protein

| | | |
|---|---|---|
| 136(1) | WP_258427457.1 | WP_258427457.1 sodium:proton antiporter |
| 136(2) | WP_212930979.1 | WP_212930979.1 Na+/H+ antiporter NhaC family protein |
| 136(1) | WP_064019315.1 | WP_064019315.1 Na+/H+ antiporter NhaC family protein |
| 137(1) | WP_000625109.1 | WP_000625109.1 DUF975 family protein |
| 137(1) | WP_097905177.1 | WP_097905177.1 DUF975 family protein |
| 137(6) | WP_097952776.1 | WP_097952776.1 DUF975 family protein |
| 137(1) | WP_000625107.1 | WP_000625107.1 DUF975 family protein |
| 137(1) | WP_087956295.1 | WP_087956295.1 DUF975 family protein |
| 137(1) | WP_098856769.1 | WP_098856769.1 DUF975 family protein |
| 137(1) | WP_088058541.1 | WP_088058541.1 DUF975 family protein |
| 137(1) | WP_060630650.1 | WP_060630650.1 DUF975 family protein |
| 137(1) | WP_306007805.1 | WP_306007805.1 DUF975 family protein |
| 137(1) | WP_242251782.1 | WP_242251782.1 DUF975 family protein |
| 137(1) | WP_087982582.1 | WP_087982582.1 DUF975 family protein |
| 137(2) | WP_098239620.1 | WP_098239620.1 DUF975 family protein |
| 137(1) | WP_097841433.1 | WP_097841433.1 DUF975 family protein |
| 137(1) | WP_289398699.1 | WP_289398699.1 DUF975 family protein |
| 137(1) | WP_262748107.1 | WP_262748107.1 DUF975 family protein |
| 137(1) | WP_166704996.1 | WP_166704996.1 DUF975 family protein |
| 137(1) | WP_147807654.1 | WP_147807654.1 DUF975 family protein |
| 137(2) | WP_063223281.1 | WP_063223281.1 DUF975 family protein |
| 137(1) | WP_151630602.1 | WP_151630602.1 DUF975 family protein |
| 137(1) | WP_071734019.1 | WP_071734019.1 DUF975 family protein |
| 137(1) | WP_098148060.1 | WP_098148060.1 DUF975 family protein |
| 137(4) | WP_097889501.1 | WP_097889501.1 DUF975 family protein |
| 137(1) | WP_098264287.1 | WP_098264287.1 DUF975 family protein |
| 137(1) | WP_002031664.1 | WP_002031664.1 DUF975 family protein |
| 137(1) | WP_098150442.1 | WP_098150442.1 DUF975 family protein |
| 137(1) | WP_212958766.1 | WP_212958766.1 DUF975 family protein |
| 138(1) | WP_000656075.1 | WP_000656075.1 Imm3 family immunity protein |
| 138(1) | WP_098088237.1 | WP_098088237.1 Imm3 family immunity protein |
| 138(1) | WP_097889504.1 | WP_097889504.1 Imm3 family immunity protein |
| 138(2) | WP_000390489.1 | WP_000390489.1 Imm3 family immunity protein |
| 138(1) | WP_147807650.1 | WP_147807650.1 Imm3 family immunity protein |
| 138(1) | WP_097822664.1 | WP_097822664.1 Imm3 family immunity protein |
| 138(1) | WP_098684015.1 | WP_098684015.1 Imm3 family immunity protein |
| 138(1) | WP_017561049.1 | WP_017561049.1 Imm3 family immunity protein |
| 138(12) | WP_000656077.1 | WP_000656077.1 Imm3 family immunity protein |
| 138(1) | WP_199695255.1 | WP_199695255.1 Imm3 family immunity protein |
| 138(1) | WP_098856773.1 | WP_098856773.1 Imm3 family immunity protein |
| 138(1) | WP_098347470.1 | WP_098347470.1 Imm3 family immunity protein |
| 138(3) | WP_098082497.1 | WP_098082497.1 Imm3 family immunity protein |
| 138(1) | WP_087982575.1 | WP_087982575.1 Imm3 family immunity protein |
| 138(5) | WP_000656074.1 | WP_000656074.1 Imm3 family immunity protein |
| 138(2) | WP_098264290.1 | WP_098264290.1 Imm3 family immunity protein |
| 139(1) | WP_076330730.1 | WP_076330730.1 barstar family protein |
| 139(1) | WP_285079431.1 | WP_285079431.1 barstar family protein |
| 139(1) | WP_076318557.1 | WP_076318557.1 barstar family protein |
| 139(1) | WP_236417237.1 | WP_236417237.1 barstar family protein |
| 139(2) | WP_024630339.1 | WP_024630339.1 barstar family protein |
| 139(1) | WP_101313447.1 | WP_101313447.1 barstar family protein |
| 139(1) | WP_036609182.1 | WP_036609182.1 barstar family protein |
| 139(1) | WP_091017183.1 | WP_091017183.1 barstar family protein |
| 139(1) | WP_247902255.1 | WP_247902255.1 barstar family protein |
| 139(1) | WP_133384362.1 | WP_133384362.1 barstar family protein |
| 139(1) | WP_264929196.1 | WP_264929196.1 barstar family protein |
| 139(1) | WP_109999901.1 | WP_109999901.1 barstar family protein |
| 139(1) | WP_017689058.1 | WP_017689058.1 barstar family protein |
| 139(1) | WP_124114127.1 | WP_124114127.1 barstar family protein |
| 139(1) | WP_105598244.1 | WP_105598244.1 barstar family protein |
| 139(1) | WP_113052556.1 | WP_113052556.1 barstar family protein |
| 139(1) | WP_076249770.1 | WP_076249770.1 barstar family protein |
| 139(2) | WP_076208833.1 | WP_076208833.1 barstar family protein |
| 139(1) | WP_150367636.1 | WP_150367636.1 barstar family protein |
| 139(2) | WP_090896220.1 | WP_090896220.1 barstar family protein |
| 139(1) | WP_062833891.1 | WP_062833891.1 barstar family protein |
| 139(2) | WP_127543685.1 | WP_127543685.1 barstar family protein |
| 139(1) | WP_145318661.1 | WP_145318661.1 barstar family protein |
| 139(1) | WP_111619115.1 | WP_111619115.1 barstar family protein |
| 139(1) | WP_079346794.1 | WP_079346794.1 barstar family protein |
| 139(1) | WP_036670446.1 | WP_036670446.1 barstar family protein |
| 139(1) | WP_099853184.1 | WP_099853184.1 barstar family protein |
| 139(1) | WP_074094623.1 | WP_074094623.1 barstar family protein |
| 139(1) | WP_062326159.1 | WP_062326159.1 barstar family protein |
| 139(1) | WP_064639003.1 | WP_064639003.1 barstar family protein |
| 139(1) | WP_145146969.1 | WP_145146969.1 barstar family protein |
| 140(1) | WP_181747855.1 | WP_181747855.1 thiazole synthase |
| 140(1) | WP_003340526.1 | WP_003340526.1 thiazole synthase |
| 140(1) | WP_022586780.1 | WP_022586780.1 thiazole synthase |
| 140(1) | WP_031415559.1 | WP_031415559.1 thiazole synthase |
| 140(1) | WP_113755288.1 | WP_113755288.1 thiazole synthase |
| 140(1) | WP_258425834.1 | WP_258425834.1 thiazole synthase |
| 140(1) | WP_094699665.1 | WP_094699665.1 thiazole synthase |
| 140(1) | WP_258418579.1 | WP_258418579.1 thiazole synthase |
| 140(1) | WP_101668483.1 | WP_101668483.1 thiazole synthase |
| 140(2) | WP_158326755.1 | WP_158326755.1 thiazole synthase |
| 140(2) | WP_119732869.1 | WP_119732869.1 thiazole synthase |
| 140(2) | WP_121474472.1 | WP_121474472.1 thiazole synthase |
| 140(1) | WP_232699441.1 | WP_232699441.1 thiazole synthase |
| 140(1) | WP_301804630.1 | WP_301804630.1 thiazole synthase |
| 140(11) | WP_018670535.1 | WP_018670535.1 thiazole synthase |
| 140(1) | WP_212930966.1 | WP_212930966.1 thiazole synthase |
| 140(1) | WP_163245173.1 | WP_163245173.1 thiazole synthase |
| 140(1) | WP_158329529.1 | WP_158329529.1 thiazole synthase |
| 140(1) | WP_068793924.1 | WP_068793924.1 thiazole synthase |
| 140(1) | WP_096886731.1 | WP_096886731.1 thiazole synthase |
| 140(2) | WP_197247496.1 | WP_197247496.1 thiazole synthase |
| 141(2) | WP_205828387.1 | WP_205828387.1 hypothetical protein |
| 141(4) | WP_085963725.1 | WP_085963725.1 hypothetical protein |

| | | |
|---|---|---|
| 141(1) | WP_220186166.1 | WP_220186166.1 hypothetical protein |
| 141(1) | WP_301804696.1 | WP_301804696.1 hypothetical protein |
| 141(1) | WP_308471519.1 | WP_308471519.1 hypothetical protein |
| 141(3) | WP_113755267.1 | WP_113755267.1 hypothetical protein |
| 141(1) | WP_308471724.1 | WP_308471724.1 hypothetical protein |
| 141(1) | WP_206767021.1 | WP_206767021.1 hypothetical protein |
| 141(3) | WP_018670508.1 | WP_018670508.1 hypothetical protein |
| 141(1) | WP_158329541.1 | WP_158329541.1 hypothetical protein |
| 141(2) | WP_064019307.1 | WP_064019307.1 hypothetical protein |
| 141(1) | WP_035311914.1 | WP_035311914.1 hypothetical protein |
| 141(2) | WP_026315054.1 | WP_026315054.1 hypothetical protein |
| 141(1) | WP_275441489.1 | WP_275441489.1 hypothetical protein |
| 141(1) | WP_255253071.1 | WP_255253071.1 hypothetical protein |
| 141(1) | WP_068793940.1 | WP_068793940.1 hypothetical protein |
| 141(4) | WP_041752217.1 | WP_041752217.1 hypothetical protein |
| 141(1) | WP_201747411.1 | WP_201747411.1 hypothetical protein |
| 141(1) | WP_276564936.1 | WP_276564936.1 hypothetical protein |
| 141(3) | WP_176452357.1 | WP_176452357.1 hypothetical protein |

| | | |
|---|---|---|
| 142(1) | WP_001030785.1 | WP_001030785.1 hypothetical protein |
| 142(1) | WP_002186029.1 | WP_002186029.1 hypothetical protein |
| 142(1) | WP_313894952.1 | WP_313894952.1 hypothetical protein |
| 142(7) | WP_000588631.1 | WP_000588631.1 hypothetical protein |
| 142(1) | WP_098226387.1 | WP_098226387.1 hypothetical protein |
| 142(1) | WP_033683858.1 | WP_033683858.1 hypothetical protein |
| 142(3) | WP_016108263.1 | WP_016108263.1 hypothetical protein |
| 142(1) | WP_017657489.1 | WP_017657489.1 hypothetical protein |
| 142(8) | WP_046392640.1 | WP_046392640.1 hypothetical protein |
| 142(2) | WP_097954752.1 | WP_097954752.1 hypothetical protein |
| 142(1) | WP_025921923.1 | WP_025921923.1 hypothetical protein |
| 142(1) | WP_001030784.1 | WP_001030784.1 hypothetical protein |
| 142(1) | WP_098284020.1 | WP_098284020.1 hypothetical protein |
| 142(1) | WP_016134262.1 | WP_016134262.1 hypothetical protein |
| 142(1) | WP_144596433.1 | WP_144596433.1 hypothetical protein |
| 142(1) | WP_277724854.1 | WP_277724854.1 hypothetical protein, partial |
| 142(2) | WP_064460228.1 | WP_064460228.1 hypothetical protein |
| 142(1) | WP_088230459.1 | WP_088230459.1 hypothetical protein |

| | | |
|---|---|---|
| 143(1) | WP_074108293.1 | WP_074108293.1 hypothetical protein |
| 143(1) | WP_099517101.1 | WP_099517101.1 hypothetical protein |
| 143(1) | WP_039871656.1 | WP_039871656.1 hypothetical protein |
| 143(1) | WP_042203650.1 | WP_042203650.1 hypothetical protein |
| 143(1) | WP_178381995.1 | WP_178381995.1 hypothetical protein |
| 143(1) | WP_019912149.1 | WP_019912149.1 hypothetical protein |
| 143(1) | WP_189018597.1 | WP_189018597.1 hypothetical protein |
| 143(2) | WP_219872459.1 | WP_219872459.1 hypothetical protein |
| 143(2) | WP_076286999.1 | WP_076286999.1 hypothetical protein |
| 143(1) | WP_209872739.1 | WP_209872739.1 hypothetical protein |
| 143(1) | WP_166244082.1 | WP_166244082.1 hypothetical protein |
| 143(2) | WP_094878481.1 | WP_094878481.1 hypothetical protein |
| 143(1) | WP_042210999.1 | WP_042210999.1 hypothetical protein |
| 143(2) | WP_042134276.1 | WP_042134276.1 hypothetical protein |
| 143(3) | WP_076135801.1 | WP_076135801.1 hypothetical protein |
| 143(3) | WP_076104577.1 | WP_076104577.1 hypothetical protein |
| 143(1) | WP_152392578.1 | WP_152392578.1 hypothetical protein |
| 143(1) | WP_076219076.1 | WP_076219076.1 hypothetical protein |
| 143(2) | WP_076303953.1 | WP_076303953.1 hypothetical protein |
| 143(1) | WP_094870402.1 | WP_094870402.1 hypothetical protein |
| 143(1) | WP_127194963.1 | WP_127194963.1 hypothetical protein |
| 143(1) | WP_039306346.1 | WP_039306346.1 hypothetical protein |
| 143(1) | WP_076279380.1 | WP_076279380.1 hypothetical protein |
| 143(1) | WP_089522639.1 | WP_089522639.1 hypothetical protein |
| 143(1) | WP_154118028.1 | WP_154118028.1 hypothetical protein |

| | | |
|---|---|---|
| 144(1) | WP_042203658.1 | WP_042203658.1 hypothetical protein |
| 144(1) | WP_076279375.1 | WP_076279375.1 hypothetical protein |
| 144(1) | WP_019912142.1 | WP_019912142.1 hypothetical protein |
| 144(2) | WP_076208273.1 | WP_076208273.1 hypothetical protein |
| 144(3) | WP_076104583.1 | WP_076104583.1 hypothetical protein |
| 144(1) | WP_076242361.1 | WP_076242361.1 hypothetical protein |
| 144(1) | WP_166244091.1 | WP_166244091.1 hypothetical protein |
| 144(3) | WP_143812135.1 | WP_143812135.1 hypothetical protein |
| 144(1) | WP_039871665.1 | WP_039871665.1 hypothetical protein |
| 144(1) | WP_089522645.1 | WP_089522645.1 hypothetical protein |
| 144(2) | WP_219872465.1 | WP_219872465.1 hypothetical protein |
| 144(1) | WP_042211006.1 | WP_042211006.1 hypothetical protein |
| 144(1) | WP_189018586.1 | WP_189018586.1 hypothetical protein |
| 144(1) | WP_209872719.1 | WP_209872719.1 hypothetical protein |
| 144(1) | WP_076303956.1 | WP_076303956.1 hypothetical protein |
| 144(1) | WP_154118021.1 | WP_154118021.1 hypothetical protein |
| 144(1) | WP_152392583.1 | WP_152392583.1 hypothetical protein |
| 144(1) | WP_099517095.1 | WP_099517095.1 hypothetical protein |
| 144(1) | WP_076135791.1 | WP_076135791.1 hypothetical protein |
| 144(1) | WP_042134291.1 | WP_042134291.1 hypothetical protein |
| 144(1) | WP_039306326.1 | WP_039306326.1 hypothetical protein |
| 144(1) | WP_076219082.1 | WP_076219082.1 hypothetical protein |
| 144(1) | WP_076309117.1 | WP_076309117.1 hypothetical protein |
| 144(2) | WP_074085347.1 | WP_074085347.1 hypothetical protein |
| 144(1) | WP_127194969.1 | WP_127194969.1 hypothetical protein |
| 144(2) | WP_076287005.1 | WP_076287005.1 hypothetical protein |

| | | |
|---|---|---|
| 145(1) | WP_014281191.1 | WP_014281191.1 CRISPR-associated helicase/endonuclease Cas3 |
| 145(1) | WP_025719049.1 | WP_025719049.1 CRISPR-associated helicase/endonuclease Cas3 |
| 145(1) | WP_220031163.1 | WP_220031163.1 CRISPR-associated helicase Cas3' |
| 145(2) | WP_063208768.1 | WP_063208768.1 CRISPR-associated helicase/endonuclease Cas3 |
| 145(1) | WP_025716881.1 | WP_025716881.1 CRISPR-associated helicase/endonuclease Cas3 |
| 145(1) | WP_061829401.1 | WP_061829401.1 CRISPR-associated helicase/endonuclease Cas3 |
| 145(1) | WP_225999413.1 | WP_225999413.1 CRISPR-associated helicase Cas3' |
| 145(2) | WP_017427970.1 | WP_017427970.1 CRISPR-associated helicase/endonuclease Cas3 |
| 145(1) | WP_165079981.1 | WP_165079981.1 CRISPR-associated helicase Cas3' |
| 145(1) | WP_071558840.1 | WP_071558840.1 CRISPR-associated helicase/endonuclease Cas3 |
| 145(1) | WP_028542685.1 | WP_028542685.1 CRISPR-associated helicase/endonuclease Cas3 |
| 145(1) | WP_192510486.1 | WP_192510486.1 CRISPR-associated helicase Cas3' |
| 145(1) | WP_104495358.1 | WP_104495358.1 CRISPR-associated helicase/endonuclease Cas3 |

| | | |
|---|---|---|
| 145(1) | WP_254327968.1 | WP_254327968.1 CRISPR-associated helicase Cas3' |
| 145(1) | WP_165147467.1 | WP_165147467.1 CRISPR-associated helicase Cas3' |
| 145(1) | WP_134909992.1 | WP_134909992.1 CRISPR-associated helicase Cas3' |
| 145(1) | WP_254325790.1 | WP_254325790.1 CRISPR-associated helicase Cas3' |
| 145(1) | WP_010346271.1 | WP_010346271.1 CRISPR-associated helicase/endonuclease Cas3 |
| 145(1) | WP_229522849.1 | WP_229522849.1 CRISPR-associated helicase Cas3' |
| 145(1) | WP_137062806.1 | WP_137062806.1 CRISPR-associated helicase Cas3' |
| 145(2) | WP_283655698.1 | WP_283655698.1 CRISPR-associated helicase Cas3' |
| 145(1) | WP_068940688.1 | WP_068940688.1 CRISPR-associated helicase/endonuclease Cas3 |
| 145(1) | WP_272640519.1 | WP_272640519.1 CRISPR-associated helicase Cas3' |
| 145(2) | WP_076264228.1 | WP_076264228.1 CRISPR-associated helicase/endonuclease Cas3 |
| 145(1) | WP_209941819.1 | WP_209941819.1 CRISPR-associated helicase Cas3' |
| 145(1) | WP_094155403.1 | WP_094155403.1 CRISPR-associated helicase/endonuclease Cas3 |
| 145(1) | WP_023988271.1 | WP_023988271.1 CRISPR-associated helicase/endonuclease Cas3 |
| 145(1) | WP_240753676.1 | WP_240753676.1 CRISPR-associated helicase Cas3' |
| 145(1) | WP_068496239.1 | WP_068496239.1 CRISPR-associated helicase/endonuclease Cas3 |
| 145(1) | WP_152381398.1 | WP_152381398.1 CRISPR-associated helicase Cas3' |

| | | |
|---|---|---|
| 146(1) | WP_094155402.1 | WP_094155402.1 type I-C CRISPR-associated protein Cas5c |
| 146(1) | WP_165147464.1 | WP_165147464.1 type I-C CRISPR-associated protein Cas5c |
| 146(1) | WP_229522848.1 | WP_229522848.1 type I-C CRISPR-associated protein Cas5c |
| 146(2) | WP_283655699.1 | WP_283655699.1 type I-C CRISPR-associated protein Cas5c |
| 146(1) | WP_014281192.1 | WP_014281192.1 type I-C CRISPR-associated protein Cas5c |
| 146(1) | WP_025719048.1 | WP_025719048.1 type I-C CRISPR-associated protein Cas5c |
| 146(1) | WP_225999414.1 | WP_225999414.1 type I-C CRISPR-associated protein Cas5c |
| 146(1) | WP_068940690.1 | WP_068940690.1 type I-C CRISPR-associated protein Cas5c |
| 146(1) | WP_152381399.1 | WP_152381399.1 type I-C CRISPR-associated protein Cas5c |
| 146(1) | WP_134909990.1 | WP_134909990.1 type I-C CRISPR-associated protein Cas5c |
| 146(3) | WP_017427971.1 | WP_017427971.1 type I-C CRISPR-associated protein Cas5c |
| 146(1) | WP_272640520.1 | WP_272640520.1 type I-C CRISPR-associated protein Cas5c |
| 146(2) | WP_063208766.1 | WP_063208766.1 type I-C CRISPR-associated protein Cas5c |
| 146(1) | WP_025716882.1 | WP_025716882.1 type I-C CRISPR-associated protein Cas5c |
| 146(1) | WP_149094180.1 | WP_149094180.1 type I-C CRISPR-associated protein Cas5c |
| 146(3) | WP_023988272.1 | WP_023988272.1 type I-C CRISPR-associated protein Cas5c |
| 146(1) | WP_165079980.1 | WP_165079980.1 type I-C CRISPR-associated protein Cas5c |
| 146(1) | WP_254327969.1 | WP_254327969.1 type I-C CRISPR-associated protein Cas5c |
| 146(1) | WP_137062807.1 | WP_137062807.1 type I-C CRISPR-associated protein Cas5c |
| 146(1) | WP_068496238.1 | WP_068496238.1 type I-C CRISPR-associated protein Cas5c |
| 146(1) | WP_010346270.1 | WP_010346270.1 type I-C CRISPR-associated protein Cas5c |
| 146(4) | WP_028542684.1 | WP_028542684.1 type I-C CRISPR-associated protein Cas5c |
| 146(1) | WP_076294183.1 | WP_076294183.1 type I-C CRISPR-associated protein Cas5c |
| 146(2) | WP_061829400.1 | WP_061829400.1 type I-C CRISPR-associated protein Cas5c |

| | | |
|---|---|---|
| 147(1) | WP_142543122.1 | WP_142543122.1 phosphotransferase |
| 147(1) | WP_111153304.1 | WP_111153304.1 phosphotransferase |
| 147(1) | WP_138186686.1 | WP_138186686.1 phosphotransferase |
| 147(2) | WP_021253111.1 | WP_021253111.1 phosphotransferase |
| 147(1) | WP_182914761.1 | WP_182914761.1 phosphotransferase |
| 147(1) | WP_065291242.1 | WP_065291242.1 phosphotransferase |
| 147(1) | WP_241734700.1 | WP_241734700.1 phosphotransferase |
| 147(1) | WP_021261876.1 | WP_021261876.1 phosphotransferase |
| 147(1) | WP_197256872.1 | WP_197256872.1 phosphotransferase |
| 147(1) | WP_272045580.1 | WP_272045580.1 phosphotransferase |
| 147(4) | WP_157266793.1 | WP_157266793.1 hypothetical protein |
| 147(1) | WP_195571411.1 | WP_195571411.1 phosphotransferase |
| 147(2) | WP_168178789.1 | WP_168178789.1 phosphotransferase |
| 147(1) | WP_174811342.1 | WP_174811342.1 phosphotransferase |
| 147(1) | WP_272827197.1 | WP_272827197.1 phosphotransferase |
| 147(1) | WP_169028902.1 | WP_169028902.1 phosphotransferase |
| 147(1) | WP_164776259.1 | WP_164776259.1 phosphotransferase |
| 147(1) | WP_293654343.1 | WP_293654343.1 phosphotransferase |
| 147(1) | WP_006674867.1 | WP_006674867.1 phosphotransferase |
| 147(1) | WP_244965662.1 | WP_244965662.1 phosphotransferase |
| 147(1) | WP_072727828.1 | WP_072727828.1 phosphotransferase |
| 147(1) | WP_088550050.1 | WP_088550050.1 phosphotransferase |
| 147(1) | WP_165643410.1 | WP_165643410.1 phosphotransferase |
| 147(1) | WP_281477409.1 | WP_281477409.1 phosphotransferase |
| 147(1) | WP_085499124.1 | WP_085499124.1 phosphotransferase |
| 147(2) | WP_197267871.1 | WP_197267871.1 phosphotransferase |
| 147(1) | WP_133378775.1 | WP_133378775.1 phosphotransferase |
| 147(1) | WP_081819937.1 | WP_081819937.1 phosphotransferase |

| | | |
|---|---|---|
| 148(1) | WP_281477401.1 | WP_281477401.1 alpha/beta fold hydrolase |
| 148(1) | WP_268623234.1 | WP_268623234.1 alpha/beta hydrolase |
| 148(1) | WP_119794355.1 | WP_119794355.1 alpha/beta fold hydrolase |
| 148(1) | WP_072727813.1 | WP_072727813.1 alpha/beta hydrolase |
| 148(1) | WP_142543138.1 | WP_142543138.1 alpha/beta hydrolase |
| 148(1) | WP_111153307.1 | WP_111153307.1 alpha/beta fold hydrolase |
| 148(1) | WP_138186708.1 | WP_138186708.1 alpha/beta hydrolase |
| 148(1) | WP_005546479.1 | WP_005546479.1 alpha/beta fold hydrolase |
| 148(1) | WP_163977067.1 | WP_163977067.1 alpha/beta hydrolase |
| 148(1) | WP_272045591.1 | WP_272045591.1 alpha/beta hydrolase |
| 148(1) | WP_087442401.1 | WP_087442401.1 alpha/beta hydrolase |
| 148(1) | WP_197256879.1 | WP_197256879.1 alpha/beta fold hydrolase |
| 148(2) | WP_143801070.1 | WP_143801070.1 alpha/beta hydrolase |
| 148(1) | WP_293654345.1 | WP_293654345.1 alpha/beta fold hydrolase |
| 148(2) | WP_028533189.1 | WP_028533189.1 alpha/beta hydrolase |
| 148(3) | WP_171414531.1 | WP_171414531.1 alpha/beta hydrolase |
| 148(1) | WP_204412742.1 | WP_204412742.1 alpha/beta fold hydrolase |
| 148(3) | WP_021261898.1 | WP_021261898.1 alpha/beta hydrolase |
| 148(1) | WP_213426998.1 | WP_213426998.1 alpha/beta fold hydrolase |
| 148(1) | WP_006674872.1 | WP_006674872.1 alpha/beta fold hydrolase |
| 148(1) | WP_021253133.1 | WP_021253133.1 alpha/beta hydrolase |
| 148(1) | WP_240764706.1 | WP_240764706.1 alpha/beta fold hydrolase |
| 148(3) | WP_268599738.1 | WP_268599738.1 alpha/beta fold hydrolase |
| 148(1) | WP_168178794.1 | WP_168178794.1 alpha/beta hydrolase |
| 148(1) | WP_065291261.1 | WP_065291261.1 alpha/beta hydrolase |
| 148(1) | WP_213468108.1 | WP_213468108.1 alpha/beta hydrolase |

| | | |
|---|---|---|
| 149(1) | WP_258425832.1 | WP_258425832.1 thiaminase II |
| 149(2) | WP_119732871.1 | WP_119732871.1 thiaminase II |
| 149(1) | WP_121474470.1 | WP_121474470.1 thiaminase II |
| 149(1) | WP_068793922.1 | WP_068793922.1 thiaminase II |

149(1) WP_104030452.1 WP_104030452.1 thiaminase II
149(2) WP_116334072.1 WP_116334072.1 thiaminase II
149(2) WP_094699667.1 WP_094699667.1 thiaminase II
149(1) WP_181747852.1 WP_181747852.1 thiaminase II
149(2) WP_163245170.1 WP_163245170.1 thiaminase II
149(1) WP_277548457.1 WP_277548457.1 thiaminase II
149(2) WP_113755290.1 WP_113755290.1 thiaminase II
149(2) WP_158326757.1 WP_158326757.1 thiaminase II
149(1) WP_101668481.1 WP_101668481.1 thiaminase II
149(1) WP_003340521.1 WP_003340521.1 thiaminase II
149(1) WP_096886729.1 WP_096886729.1 thiaminase II
149(1) WP_301804625.1 WP_301804625.1 thiaminase II
149(6) WP_026315068.1 WP_026315068.1 thiaminase II
149(2) WP_212930964.1 WP_212930964.1 thiaminase II
149(1) WP_104148787.1 WP_104148787.1 thiaminase II
149(1) WP_278431116.1 WP_278431116.1 thiaminase II
149(1) WP_022586782.1 WP_022586782.1 thiaminase II
149(1) WP_258433011.1 WP_258433011.1 thiaminase II

150(1) WP_094155401.1 WP_094155401.1 type I-C CRISPR-associated protein Cas8c/Csd1
150(1) WP_305166256.1 WP_305166256.1 type I-C CRISPR-associated protein Cas8c/Csd1
150(1) WP_137062808.1 WP_137062808.1 type I-C CRISPR-associated protein Cas8c/Csd1
150(1) WP_165147461.1 WP_165147461.1 type I-C CRISPR-associated protein Cas8c/Csd1
150(1) WP_272640521.1 WP_272640521.1 type I-C CRISPR-associated protein Cas8c/Csd1
150(3) WP_017427972.1 WP_017427972.1 type I-C CRISPR-associated protein Cas8c/Csd1
150(3) WP_076264227.1 WP_076264227.1 type I-C CRISPR-associated protein Cas8c/Csd1
150(1) WP_134909988.1 WP_134909988.1 type I-C CRISPR-associated protein Cas8c/Csd1
150(1) WP_023988273.1 WP_023988273.1 type I-C CRISPR-associated protein Cas8c/Csd1
150(1) WP_220031164.1 WP_220031164.1 type I-C CRISPR-associated protein Cas8c/Csd1
150(1) WP_209941821.1 WP_209941821.1 type I-C CRISPR-associated protein Cas8c/Csd1
150(1) WP_068496236.1 WP_068496236.1 type I-C CRISPR-associated protein Cas8c/Csd1
150(1) WP_014281193.1 WP_014281193.1 type I-C CRISPR-associated protein Cas8c/Csd1
150(1) WP_152381400.1 WP_152381400.1 type I-C CRISPR-associated protein Cas8c/Csd1
150(1) WP_254325789.1 WP_254325789.1 type I-C CRISPR-associated protein Cas8c/Csd1
150(1) WP_025719046.1 WP_025719046.1 type I-C CRISPR-associated protein Cas8c/Csd1
150(1) WP_028542683.1 WP_028542683.1 type I-C CRISPR-associated protein Cas8c/Csd1
150(1) WP_068940691.1 WP_068940691.1 type I-C CRISPR-associated protein Cas8c/Csd1
150(2) WP_283655700.1 WP_283655700.1 type I-C CRISPR-associated protein Cas8c/Csd1
150(1) WP_229522847.1 WP_229522847.1 type I-C CRISPR-associated protein Cas8c/Csd1
150(1) WP_025716883.1 WP_025716883.1 type I-C CRISPR-associated protein Cas8c/Csd1
150(1) WP_192510485.1 WP_192510485.1 type I-C CRISPR-associated protein Cas8c/Csd1
150(1) WP_254327970.1 WP_254327970.1 type I-C CRISPR-associated protein Cas8c/Csd1
150(1) WP_165079979.1 WP_165079979.1 type I-C CRISPR-associated protein Cas8c/Csd1
150(1) WP_225999415.1 WP_225999415.1 type I-C CRISPR-associated protein Cas8c/Csd1
150(1) WP_063208764.1 WP_063208764.1 type I-C CRISPR-associated protein Cas8c/Csd1
150(1) WP_061829399.1 WP_061829399.1 type I-C CRISPR-associated protein Cas8c/Csd1
150(1) WP_010346269.1 WP_010346269.1 type I-C CRISPR-associated protein Cas8c/Csd1
150(1) WP_076156848.1 WP_076156848.1 type I-C CRISPR-associated protein Cas8c/Csd1

151(2) WP_200372543.1 WP_200372543.1 DUF1672 family protein
151(2) WP_185365688.1 WP_185365688.1 DUF1672 family protein
151(2) WP_185413318.1 WP_185413318.1 DUF1672 family protein
151(2) WP_185474092.1 WP_185474092.1 DUF1672 family protein
151(2) WP_185639010.1 WP_185639010.1 DUF1672 family protein
151(2) WP_194335281.1 WP_194335281.1 DUF1672 family protein
151(2) WP_270995160.1 WP_270995160.1 hypothetical protein
151(6) WP_185340296.1 WP_185340296.1 DUF1672 family protein
151(4) WP_185337278.1 WP_185337278.1 DUF1672 family protein
151(3) WP_243285278.1 WP_243285278.1 hypothetical protein
151(4) WP_203573359.1 WP_203573359.1 DUF1672 family protein
151(2) WP_185639006.1 WP_185639006.1 DUF1672 family protein

152(1) WP_119794369.1 WP_119794369.1 DUF2164 domain-containing protein
152(1) WP_111153296.1 WP_111153296.1 DUF2164 domain-containing protein
152(7) WP_005546527.1 WP_005546527.1 DUF2164 domain-containing protein
152(1) WP_289355899.1 WP_289355899.1 DUF2164 domain-containing protein
152(1) WP_197256852.1 WP_197256852.1 DUF2164 domain-containing protein
152(5) WP_021253110.1 WP_021253110.1 DUF2164 domain-containing protein
152(1) WP_072727829.1 WP_072727829.1 DUF2164 domain-containing protein
152(1) WP_065291241.1 WP_065291241.1 DUF2164 domain-containing protein
152(3) WP_168178780.1 WP_168178780.1 DUF2164 domain-containing protein
152(2) WP_087433614.1 WP_087433614.1 DUF2164 domain-containing protein
152(3) WP_006674854.1 WP_006674854.1 DUF2164 domain-containing protein
152(2) WP_143801054.1 WP_143801054.1 DUF2164 domain-containing protein
152(1) WP_021261875.1 WP_021261875.1 DUF2164 domain-containing protein
152(1) WP_272045562.1 WP_272045562.1 DUF2164 domain-containing protein
152(1) WP_138186685.1 WP_138186685.1 DUF2164 domain-containing protein
152(1) WP_087442419.1 WP_087442419.1 DUF2164 domain-containing protein
152(1) WP_206094029.1 WP_206094029.1 DUF2164 domain-containing protein

153(1) WP_000657437.1 WP_000657437.1 DUF600 family protein
153(1) WP_089522626.1 WP_089522626.1 DUF600 domain-containing protein
153(3) WP_097952961.1 WP_097952961.1 DUF600 family protein
153(1) WP_289398702.1 WP_289398702.1 DUF600 family protein
153(1) WP_002148293.1 WP_002148293.1 DUF600 family protein
153(7) WP_088063235.1 WP_088063235.1 DUF600 family protein
153(1) WP_049112854.1 WP_049112854.1 DUF600 family protein
153(1) WP_086720022.1 WP_086720022.1 DUF600 family protein
153(4) WP_000657431.1 WP_000657431.1 DUF600 family protein
153(1) WP_083334784.1 WP_083334784.1 DUF600 domain-containing protein
153(1) WP_209365670.1 WP_209365670.1 DUF600 domain-containing protein
153(1) WP_086720020.1 WP_086720020.1 DUF600 family protein
153(1) WP_271839893.1 WP_271839893.1 hypothetical protein
153(1) WP_068696901.1 WP_068696901.1 DUF600 domain-containing protein
153(2) WP_021291840.1 WP_021291840.1 hypothetical protein
153(1) WP_202406552.1 WP_202406552.1 DUF600 domain-containing protein
153(3) WP_042512828.1 WP_042512828.1 DUF600 family protein
153(1) WP_000657433.1 WP_000657433.1 DUF600 family protein
153(1) WP_000657439.1 WP_000657439.1 DUF600 family protein

154(1) WP_305166255.1 WP_305166255.1 type I-C CRISPR-associated protein Cas7/Csd2
154(1) WP_165147458.1 WP_165147458.1 type I-C CRISPR-associated protein Cas7/Csd2
154(1) WP_137062809.1 WP_137062809.1 type I-C CRISPR-associated protein Cas7/Csd2
154(1) WP_025719044.1 WP_025719044.1 type I-C CRISPR-associated protein Cas7/Csd2

154(1) WP_068940693.1 WP_068940693.1 type I-C CRISPR-associated protein Cas7/Csd2
154(1) WP_192510484.1 WP_192510484.1 type I-C CRISPR-associated protein Cas7/Csd2
154(1) WP_068496234.1 WP_068496234.1 type I-C CRISPR-associated protein Cas7/Csd2
154(1) WP_063208759.1 WP_063208759.1 type I-C CRISPR-associated protein Cas7/Csd2
154(1) WP_229522846.1 WP_229522846.1 type I-C CRISPR-associated protein Cas7/Csd2
154(2) WP_025716884.1 WP_025716884.1 type I-C CRISPR-associated protein Cas7/Csd2
154(1) WP_242210847.1 WP_242210847.1 type I-C CRISPR-associated protein Cas7/Csd2
154(1) WP_134909986.1 WP_134909986.1 type I-C CRISPR-associated protein Cas7/Csd2
154(1) WP_014281194.1 WP_014281194.1 type I-C CRISPR-associated protein Cas7/Csd2
154(1) WP_076294181.1 WP_076294181.1 type I-C CRISPR-associated protein Cas7/Csd2
154(2) WP_076156845.1 WP_076156845.1 type I-C CRISPR-associated protein Cas7/Csd2
154(2) WP_017427973.1 WP_017427973.1 type I-C CRISPR-associated protein Cas7/Csd2
154(4) WP_028542682.1 WP_028542682.1 type I-C CRISPR-associated protein Cas7/Csd2
154(1) WP_094155400.1 WP_094155400.1 type I-C CRISPR-associated protein Cas7/Csd2
154(2) WP_023988274.1 WP_023988274.1 type I-C CRISPR-associated protein Cas7/Csd2
154(2) WP_061829398.1 WP_061829398.1 type I-C CRISPR-associated protein Cas7/Csd2
154(2) WP_283655701.1 WP_283655701.1 type I-C CRISPR-associated protein Cas7/Csd2
154(1) WP_272640522.1 WP_272640522.1 type I-C CRISPR-associated protein Cas7/Csd2
154(1) WP_225999416.1 WP_225999416.1 type I-C CRISPR-associated protein Cas7/Csd2
154(1) WP_165079978.1 WP_165079978.1 type I-C CRISPR-associated protein Cas7/Csd2

155(1) WP_302878975.1 WP_302878975.1 ribokinase
155(1) WP_088830542.1 WP_088830542.1 ribokinase
155(1) WP_066020849.1 WP_066020849.1 sugar kinase
155(1) WP_253411086.1 WP_253411086.1 ribokinase
155(1) WP_108534963.1 WP_108534963.1 ribokinase
155(1) WP_251598141.1 WP_251598141.1 ribokinase
155(1) WP_053403778.1 WP_053403778.1 pyridoxine/pyridoxal/pyridoxamine kinase
155(1) WP_254366017.1 WP_254366017.1 PfkB family carbohydrate kinase
155(1) WP_179603489.1 WP_179603489.1 carbohydrate kinase
155(1) WP_034675327.1 WP_034675327.1 carbohydrate kinase
155(1) WP_054972097.1 WP_054972097.1 ribokinase
155(1) WP_127459440.1 WP_127459440.1 ribokinase
155(1) WP_242172368.1 WP_242172368.1 pyridoxine/pyridoxal/pyridoxamine kinase
155(2) WP_010963717.1 WP_010963717.1 sugar kinase
155(1) WP_163853468.1 WP_163853468.1 ribokinase
155(1) WP_181737041.1 WP_181737041.1 sugar kinase
155(1) WP_063188053.1 WP_063188053.1 PfkB family carbohydrate kinase
155(1) WP_281937361.1 WP_281937361.1 ribokinase
155(1) WP_188535042.1 WP_188535042.1 bifunctional hydroxymethylpyrimidine kinase/phosphomethylpyrimidine kinase
155(1) WP_307312527.1 WP_307312527.1 carbohydrate kinase
155(1) WP_036689431.1 WP_036689431.1 ribokinase
155(1) WP_025848620.1 WP_025848620.1 ribokinase
155(1) WP_090674360.1 WP_090674360.1 ribokinase
155(1) WP_127484341.1 WP_127484341.1 ribokinase
155(1) WP_263324534.1 WP_263324534.1 sugar kinase
155(1) WP_204445200.1 WP_204445200.1 PfkB family carbohydrate kinase
155(1) WP_267904943.1 WP_267904943.1 PfkB family carbohydrate kinase
155(1) WP_306077401.1 WP_306077401.1 carbohydrate kinase
155(1) WP_010496615.1 WP_010496615.1 ribokinase
155(1) WP_063188047.1 WP_063188047.1 ribokinase
155(1) WP_216791230.1 WP_216791230.1 ribokinase
155(1) WP_251610746.1 WP_251610746.1 pyridoxine/pyridoxal/pyridoxamine kinase

156(2) WP_044355262.1 WP_044355262.1 HD domain-containing protein
156(1) WP_087442404.1 WP_087442404.1 HD domain-containing protein
156(2) WP_065291253.1 WP_065291253.1 HD domain-containing protein
156(3) WP_006674869.1 WP_006674869.1 HD domain-containing protein
156(3) WP_268599734.1 WP_268599734.1 HD domain-containing protein
156(3) WP_021253124.1 WP_021253124.1 HD domain-containing protein
156(1) WP_133378777.1 WP_133378777.1 HD domain-containing protein
156(2) WP_171414525.1 WP_171414525.1 HD domain-containing protein
156(2) WP_143801067.1 WP_143801067.1 HD domain-containing protein
156(2) WP_142543132.1 WP_142543132.1 HD domain-containing protein
156(1) WP_272045583.1 WP_272045583.1 HD domain-containing protein
156(3) WP_168178791.1 WP_168178791.1 HD domain-containing protein
156(1) WP_072727820.1 WP_072727820.1 HD domain-containing protein
156(1) WP_197256875.1 WP_197256875.1 HD domain-containing protein
156(1) WP_213426995.1 WP_213426995.1 HD domain-containing protein
156(1) WP_119794356.1 WP_119794356.1 HD domain-containing protein
156(1) WP_005546499.1 WP_005546499.1 HD domain-containing protein
156(2) WP_197267876.1 WP_197267876.1 HD domain-containing protein

157(1) WP_209941823.1 WP_209941823.1 CRISPR-associated protein Cas4
157(1) WP_305166253.1 WP_305166253.1 CRISPR-associated protein Cas4
157(1) WP_225999417.1 WP_225999417.1 CRISPR-associated protein Cas4
157(1) WP_068496232.1 WP_068496232.1 CRISPR-associated protein Cas4
157(3) WP_076264226.1 WP_076264226.1 CRISPR-associated protein Cas4
157(1) WP_202894728.1 WP_202894728.1 CRISPR-associated protein Cas4
157(1) WP_152381401.1 WP_152381401.1 CRISPR-associated protein Cas4
157(1) WP_063208757.1 WP_063208757.1 CRISPR-associated protein Cas4
157(2) WP_061829397.1 WP_061829397.1 CRISPR-associated protein Cas4
157(1) WP_272640523.1 WP_272640523.1 CRISPR-associated protein Cas4
157(1) WP_104495360.1 WP_104495360.1 CRISPR-associated protein Cas4
157(1) WP_014281195.1 WP_014281195.1 CRISPR-associated protein Cas4
157(1) WP_076294180.1 WP_076294180.1 CRISPR-associated protein Cas4
157(1) WP_025716885.1 WP_025716885.1 CRISPR-associated protein Cas4
157(1) WP_094155399.1 WP_094155399.1 CRISPR-associated protein Cas4
157(2) WP_023988275.1 WP_023988275.1 CRISPR-associated protein Cas4
157(1) WP_076156843.1 WP_076156843.1 CRISPR-associated protein Cas4
157(1) WP_068940694.1 WP_068940694.1 CRISPR-associated protein Cas4
157(1) WP_165079977.1 WP_165079977.1 CRISPR-associated protein Cas4
157(1) WP_134914326.1 WP_134914326.1 CRISPR-associated protein Cas4
157(2) WP_283655702.1 WP_283655702.1 CRISPR-associated protein Cas4
157(3) WP_017427974.1 WP_017427974.1 CRISPR-associated protein Cas4
157(1) WP_137062833.1 WP_137062833.1 CRISPR-associated protein Cas4
157(1) WP_025719043.1 WP_025719043.1 CRISPR-associated protein Cas4
157(1) WP_254327971.1 WP_254327971.1 CRISPR-associated protein Cas4

158(1) WP_047909603.1 WP_047909603.1 helix-turn-helix domain-containing protein
158(1) WP_213532668.1 WP_213532668.1 helix-turn-helix domain-containing protein
158(1) WP_110892948.1 WP_110892948.1 helix-turn-helix domain-containing protein
158(1) WP_307214313.1 WP_307214313.1 helix-turn-helix domain-containing protein

| | | |
|---|---|---|
| 158(1) | WP_145412711.1 | WP_145412711.1 helix-turn-helix domain-containing protein |
| 158(1) | WP_076311042.1 | WP_076311042.1 helix-turn-helix domain-containing protein |
| 158(1) | WP_230875701.1 | WP_230875701.1 helix-turn-helix domain-containing protein |
| 158(1) | WP_205053316.1 | WP_205053316.1 winged helix-turn-helix domain-containing protein |
| 158(1) | WP_251583312.1 | WP_251583312.1 helix-turn-helix domain-containing protein |
| 158(1) | WP_128101497.1 | WP_128101497.1 helix-turn-helix domain-containing protein |
| 158(1) | WP_091155403.1 | WP_091155403.1 helix-turn-helix domain-containing protein |
| 158(1) | WP_056700815.1 | WP_056700815.1 helix-turn-helix domain-containing protein |
| 158(1) | WP_308721981.1 | WP_308721981.1 helix-turn-helix domain-containing protein |
| 158(1) | WP_023957752.1 | WP_023957752.1 helix-turn-helix domain-containing protein |
| 158(1) | WP_145046475.1 | WP_145046475.1 helix-turn-helix domain-containing protein |
| 158(1) | WP_090722892.1 | WP_090722892.1 helix-turn-helix domain-containing protein |
| 158(2) | WP_095266274.1 | WP_095266274.1 helix-turn-helix domain-containing protein |
| 158(1) | WP_244159319.1 | WP_244159319.1 helix-turn-helix domain-containing protein |
| 158(1) | WP_272035346.1 | WP_272035346.1 helix-turn-helix domain-containing protein |
| 158(1) | WP_127536186.1 | WP_127536186.1 winged helix-turn-helix domain-containing protein |
| 158(1) | WP_145408771.1 | WP_145408771.1 helix-turn-helix domain-containing protein |
| 158(1) | WP_289373777.1 | WP_289373777.1 helix-turn-helix domain-containing protein |
| 158(1) | WP_095287648.1 | WP_095287648.1 helix-turn-helix domain-containing protein |
| 158(1) | WP_188593172.1 | WP_188593172.1 helix-turn-helix domain-containing protein |
| 158(1) | WP_191798539.1 | WP_191798539.1 helix-turn-helix domain-containing protein |
| 158(2) | WP_095358538.1 | WP_095358538.1 helix-turn-helix domain-containing protein |
| 158(1) | WP_145329920.1 | WP_145329920.1 helix-turn-helix domain-containing protein |
| 158(1) | WP_211745817.1 | WP_211745817.1 helix-turn-helix domain-containing protein |
| 158(1) | WP_285748191.1 | WP_285748191.1 helix-turn-helix domain-containing protein |
| 158(1) | WP_293202253.1 | WP_293202253.1 helix-turn-helix domain-containing protein |
| 159(2) | WP_283655703.1 | WP_283655703.1 type I-C CRISPR-associated endonuclease Cas1c |
| 159(1) | WP_238791009.1 | WP_238791009.1 CRISPR-associated endonuclease Cas1 |
| 159(1) | WP_068496229.1 | WP_068496229.1 type I-C CRISPR-associated endonuclease Cas1c |
| 159(2) | WP_068940696.1 | WP_068940696.1 type I-C CRISPR-associated endonuclease Cas1c |
| 159(1) | WP_061829396.1 | WP_061829396.1 type I-C CRISPR-associated endonuclease Cas1c |
| 159(1) | WP_209941825.1 | WP_209941825.1 type I-C CRISPR-associated endonuclease Cas1c |
| 159(3) | WP_025716886.1 | WP_025716886.1 type I-C CRISPR-associated endonuclease Cas1c |
| 159(1) | WP_305166252.1 | WP_305166252.1 type I-C CRISPR-associated endonuclease Cas1c |
| 159(1) | WP_272640524.1 | WP_272640524.1 type I-C CRISPR-associated endonuclease Cas1c |
| 159(1) | WP_031463348.1 | WP_031463348.1 type I-C CRISPR-associated endonuclease Cas1c |
| 159(1) | WP_165147455.1 | WP_165147455.1 type I-C CRISPR-associated endonuclease Cas1c |
| 159(1) | WP_305166258.1 | WP_305166258.1 hypothetical protein |
| 159(1) | WP_076156840.1 | WP_076156840.1 type I-C CRISPR-associated endonuclease Cas1c |
| 159(1) | WP_076294870.1 | WP_076294870.1 type I-C CRISPR-associated endonuclease Cas1c |
| 159(2) | WP_017427975.1 | WP_017427975.1 type I-C CRISPR-associated endonuclease Cas1c |
| 159(1) | WP_165079976.1 | WP_165079976.1 type I-C CRISPR-associated endonuclease Cas1c |
| 159(1) | WP_025719041.1 | WP_025719041.1 type I-C CRISPR-associated endonuclease Cas1c |
| 159(1) | WP_028542681.1 | WP_028542681.1 type I-C CRISPR-associated endonuclease Cas1c |
| 159(1) | WP_076264225.1 | WP_076264225.1 type I-C CRISPR-associated endonuclease Cas1c |
| 159(1) | WP_134909984.1 | WP_134909984.1 type I-C CRISPR-associated endonuclease Cas1c |
| 159(1) | WP_254325788.1 | WP_254325788.1 type I-C CRISPR-associated endonuclease Cas1c |
| 159(1) | WP_094155398.1 | WP_094155398.1 type I-C CRISPR-associated endonuclease Cas1c |
| 159(1) | WP_220031165.1 | WP_220031165.1 type I-C CRISPR-associated endonuclease Cas1c |
| 159(1) | WP_254327972.1 | WP_254327972.1 type I-C CRISPR-associated endonuclease Cas1c |
| 159(1) | WP_192510483.1 | WP_192510483.1 type I-C CRISPR-associated endonuclease Cas1c |

| | | |
|---|---|---|
| 159(1) | WP_063208755.1 | WP_063208755.1 type I-C CRISPR-associated endonuclease Cas1c |
| 160(9) | WP_001221976.1 | WP_001221976.1 glycerate kinase |
| 160(1) | WP_144611191.1 | WP_144611191.1 glycerate kinase |
| 160(2) | WP_074565300.1 | WP_074565300.1 glycerate kinase |
| 160(1) | WP_053564624.1 | WP_053564624.1 glycerate kinase |
| 160(1) | WP_166704997.1 | WP_166704997.1 glycerate kinase |
| 160(1) | WP_103655291.1 | WP_103655291.1 glycerate kinase |
| 160(1) | WP_113711508.1 | WP_113711508.1 glycerate kinase |
| 160(1) | WP_212958798.1 | WP_212958798.1 glycerate kinase |
| 160(1) | WP_098103314.1 | WP_098103314.1 glycerate kinase |
| 160(2) | WP_001221973.1 | WP_001221973.1 glycerate kinase |
| 160(1) | WP_242251794.1 | WP_242251794.1 glycerate kinase |
| 160(1) | WP_307192909.1 | WP_307192909.1 glycerate kinase |
| 160(1) | WP_151630600.1 | WP_151630600.1 glycerate kinase |
| 160(1) | WP_222655355.1 | WP_222655355.1 glycerate kinase |
| 160(1) | WP_071736286.1 | WP_071736286.1 glycerate kinase |
| 160(1) | WP_139867374.1 | WP_139867374.1 glycerate kinase |
| 160(5) | WP_001221974.1 | WP_001221974.1 glycerate kinase |
| 161(1) | WP_164744410.1 | WP_164744410.1 M23 family metallopeptidase |
| 161(1) | WP_079506364.1 | WP_079506364.1 M23 family metallopeptidase |
| 161(2) | WP_102264563.1 | WP_102264563.1 M23 family metallopeptidase |
| 161(1) | WP_238187874.1 | WP_238187874.1 M23 family metallopeptidase |
| 161(1) | WP_041964589.1 | WP_041964589.1 M23 family metallopeptidase |
| 161(1) | WP_210364440.1 | WP_210364440.1 M23 family metallopeptidase |
| 161(1) | WP_239617484.1 | WP_239617484.1 M23 family metallopeptidase |
| 161(1) | WP_283904353.1 | WP_283904353.1 M23 family metallopeptidase |
| 161(2) | WP_133332973.1 | WP_133332973.1 M23 family metallopeptidase |
| 161(1) | WP_182301844.1 | WP_182301844.1 M23 family metallopeptidase |
| 161(1) | WP_226602968.1 | WP_226602968.1 M23 family metallopeptidase |
| 161(1) | WP_162463323.1 | WP_162463323.1 M23 family metallopeptidase |
| 161(1) | WP_134750622.1 | WP_134750622.1 M23 family metallopeptidase |
| 161(1) | WP_136372948.1 | WP_136372948.1 M23 family metallopeptidase |
| 161(1) | WP_214902969.1 | WP_214902969.1 M23 family metallopeptidase |
| 161(1) | WP_035423708.1 | WP_035423708.1 M23 family metallopeptidase |
| 161(1) | WP_138755928.1 | WP_138755928.1 M23 family metallopeptidase |
| 161(1) | WP_215032156.1 | WP_215032156.1 M23 family metallopeptidase |
| 161(1) | WP_191272879.1 | WP_191272879.1 M23 family metallopeptidase |
| 161(1) | WP_251466782.1 | WP_251466782.1 M23 family metallopeptidase |
| 161(1) | WP_167831123.1 | WP_167831123.1 M23 family metallopeptidase |
| 161(1) | WP_132008121.1 | WP_132008121.1 peptidoglycan DD-metalloendopeptidase family protein |
| 161(1) | WP_226642848.1 | WP_226642848.1 M23 family metallopeptidase |
| 161(1) | WP_027088418.1 | WP_027088418.1 M23 family metallopeptidase |
| 161(1) | WP_120112279.1 | WP_120112279.1 M23 family metallopeptidase |
| 161(1) | WP_251542589.1 | WP_251542589.1 M23 family metallopeptidase |
| 161(1) | WP_198310838.1 | WP_198310838.1 M23 family metallopeptidase |
| 161(1) | WP_307601013.1 | WP_307601013.1 M23 family metallopeptidase |
| 161(1) | WP_144479088.1 | WP_144479088.1 M23 family metallopeptidase |
| 162(1) | WP_277157937.1 | WP_277157937.1 hypothetical protein |
| 162(1) | WP_277157942.1 | WP_277157942.1 hypothetical protein |

162(1) WP_058271114.1 WP_058271114.1 hypothetical protein
162(1) WP_270472011.1 WP_270472011.1 hypothetical protein
162(1) WP_070638659.1 WP_070638659.1 hypothetical protein
162(1) WP_307738729.1 WP_307738729.1 hypothetical protein
162(1) WP_270472033.1 WP_270472033.1 hypothetical protein
162(1) WP_006292342.1 WP_006292342.1 DUF5082 family protein
162(1) WP_033521264.1 WP_033521264.1 hypothetical protein
162(1) WP_297183837.1 WP_297183837.1 hypothetical protein
162(1) WP_277175582.1 WP_277175582.1 hypothetical protein
162(2) WP_291288268.1 WP_291288268.1 hypothetical protein
162(1) WP_006290897.1 WP_006290897.1 DUF5082 family protein
162(1) WP_277156180.1 WP_277156180.1 hypothetical protein
162(1) WP_212330870.1 WP_212330870.1 hypothetical protein
162(1) WP_034526613.1 WP_034526613.1 hypothetical protein
162(1) WP_273383878.1 WP_273383878.1 hypothetical protein
162(1) WP_303942540.1 WP_303942540.1 hypothetical protein
162(1) WP_281667416.1 WP_281667416.1 DUF5082 family protein
162(2) WP_006290893.1 WP_006290893.1 hypothetical protein
162(2) WP_297037465.1 WP_297037465.1 hypothetical protein
162(2) WP_102372747.1 WP_102372747.1 hypothetical protein
162(1) WP_277142081.1 WP_277142081.1 hypothetical protein
162(1) WP_006720316.1 WP_006720316.1 hypothetical protein
162(1) WP_007147992.1 WP_007147992.1 DUF5082 family protein
162(1) WP_006289747.1 WP_006289747.1 DUF5082 domain-containing protein

163(1) WP_016124501.1 WP_016124501.1 GNAT family N-acetyltransferase
163(1) WP_000984748.1 WP_000984748.1 GNAT family N-acetyltransferase
163(1) WP_065223580.1 WP_065223580.1 GNAT family N-acetyltransferase
163(1) WP_098139247.1 WP_098139247.1 GNAT family N-acetyltransferase
163(1) WP_098054112.1 WP_098054112.1 GNAT family N-acetyltransferase
163(2) WP_000984741.1 WP_000984741.1 GNAT family N-acetyltransferase
163(3) WP_060852180.1 WP_060852180.1 GNAT family N-acetyltransferase
163(1) WP_098226386.1 WP_098226386.1 GNAT family N-acetyltransferase
163(2) WP_000984760.1 WP_000984760.1 GNAT family N-acetyltransferase
163(1) WP_061666635.1 WP_061666635.1 GNAT family N-acetyltransferase
163(1) WP_097954748.1 WP_097954748.1 GNAT family N-acetyltransferase
163(1) WP_097960445.1 WP_097960445.1 GNAT family N-acetyltransferase
163(1) WP_053512316.1 WP_053512316.1 GNAT family N-acetyltransferase
163(1) WP_097796437.1 WP_097796437.1 GNAT family N-acetyltransferase
163(1) WP_098882507.1 WP_098882507.1 GNAT family N-acetyltransferase
163(2) WP_030025445.1 WP_030025445.1 GNAT family N-acetyltransferase
163(1) WP_000984749.1 WP_000984749.1 GNAT family N-acetyltransferase
163(1) WP_098084157.1 WP_098084157.1 GNAT family N-acetyltransferase
163(1) WP_016117684.1 WP_016117684.1 GNAT family N-acetyltransferase
163(1) WP_153598081.1 WP_153598081.1 GNAT family N-acetyltransferase
163(1) WP_086389014.1 WP_086389014.1 GNAT family N-acetyltransferase
163(1) WP_255289398.1 WP_255289398.1 GNAT family N-acetyltransferase
163(1) WP_000984757.1 WP_000984757.1 GNAT family N-acetyltransferase
163(1) WP_046946378.1 WP_046946378.1 GNAT family N-acetyltransferase

164(1) WP_098856770.1 WP_098856770.1 hypothetical protein
164(1) WP_223617607.1 WP_223617607.1 hypothetical protein
164(1) WP_262719886.1 WP_262719886.1 hypothetical protein
164(1) WP_199695254.1 WP_199695254.1 hypothetical protein
164(3) WP_097996672.1 WP_097996672.1 hypothetical protein
164(4) WP_000646607.1 WP_000646607.1 hypothetical protein
164(2) WP_097889502.1 WP_097889502.1 hypothetical protein
164(2) WP_087956296.1 WP_087956296.1 hypothetical protein
164(1) WP_023521819.1 WP_023521819.1 hypothetical protein
164(4) WP_000646589.1 WP_000646589.1 hypothetical protein
164(1) WP_087982580.1 WP_087982580.1 hypothetical protein
164(1) WP_098239617.1 WP_098239617.1 hypothetical protein
164(1) WP_243500459.1 WP_243500459.1 hypothetical protein
164(2) WP_098088241.1 WP_098088241.1 hypothetical protein
164(1) WP_000856121.1 WP_000856121.1 hypothetical protein
164(1) WP_097822666.1 WP_097822666.1 hypothetical protein
164(1) WP_016129280.1 WP_016129280.1 hypothetical protein

165(1) WP_086390669.1 WP_086390669.1 SMI1/KNR4 family protein
165(1) WP_016129282.1 WP_016129282.1 SMI1/KNR4 family protein
165(1) WP_157418136.1 WP_157418136.1 SMI1/KNR4 family protein
165(1) WP_179034775.1 WP_179034775.1 SMI1/KNR4 family protein
165(1) WP_097905175.1 WP_097905175.1 SMI1/KNR4 family protein
165(1) WP_272541588.1 WP_272541588.1 SMI1/KNR4 family protein
165(1) WP_098562450.1 WP_098562450.1 SMI1/KNR4 family protein
165(2) WP_097961545.1 WP_097961545.1 SMI1/KNR4 family protein
165(1) WP_115359666.1 WP_115359666.1 SMI1/KNR4 family protein
165(1) WP_098235626.1 WP_098235626.1 SMI1/KNR4 family protein
165(1) WP_274529057.1 WP_274529057.1 SMI1/KNR4 family protein
165(2) WP_070139645.1 WP_070139645.1 SMI1/KNR4 family protein
165(2) WP_098137760.1 WP_098137760.1 SMI1/KNR4 family protein
165(1) WP_019912158.1 WP_019912158.1 SMI1/KNR4 family protein
165(1) WP_145406449.1 WP_145406449.1 SMI1/KNR4 family protein
165(1) WP_240379737.1 WP_240379737.1 SMI1/KNR4 family protein
165(1) WP_002137245.1 WP_002137245.1 SMI1/KNR4 family protein
165(1) WP_094879048.1 WP_094879048.1 SMI1/KNR4 family protein
165(7) WP_071736275.1 WP_071736275.1 SMI1/KNR4 family protein

166(1) WP_097920536.1 WP_097920536.1 Imm47 family immunity protein
166(1) WP_238484970.1 WP_238484970.1 Imm47 family immunity protein
166(4) WP_103626234.1 WP_103626234.1 Imm47 family immunity protein
166(1) WP_286124793.1 WP_286124793.1 Imm47 family immunity protein
166(1) WP_000862548.1 WP_000862548.1 Imm47 family immunity protein
166(1) WP_060630651.1 WP_060630651.1 Imm47 family immunity protein
166(1) WP_098684016.1 WP_098684016.1 Imm47 family immunity protein
166(1) WP_000862547.1 WP_000862547.1 Imm47 family immunity protein
166(1) WP_223617606.1 WP_223617606.1 Imm47 family immunity protein
166(2) WP_042512826.1 WP_042512826.1 Imm47 family immunity protein
166(3) WP_087952378.1 WP_087952378.1 Imm47 family immunity protein
166(1) WP_284566228.1 WP_284566228.1 Imm47 family immunity protein
166(1) WP_097987486.1 WP_097987486.1 Imm47 family immunity protein
166(1) WP_098868579.1 WP_098868579.1 Imm47 family immunity protein
166(1) WP_000862554.1 WP_000862554.1 Imm47 family immunity protein
166(1) WP_097846503.1 WP_097846503.1 Imm47 family immunity protein

| | | |
|---|---|---|
| 166(1) | WP_000862555.1 | WP_000862555.1 Imm47 family immunity protein |
| 166(1) | WP_000862551.1 | WP_000862551.1 Imm47 family immunity protein |
| 166(2) | WP_098254019.1 | WP_098254019.1 Imm47 family immunity protein |
| 166(2) | WP_139895505.1 | WP_139895505.1 Imm47 family immunity protein |

| | | |
|---|---|---|
| 167(9) | WP_000570745.1 | WP_000570745.1 SLC13 family permease |
| 167(1) | WP_229136359.1 | WP_229136359.1 SLC13 family permease |
| 167(1) | WP_212958796.1 | WP_212958796.1 SLC13 family permease |
| 167(1) | WP_139867372.1 | WP_139867372.1 SLC13 family permease |
| 167(3) | WP_000570742.1 | WP_000570742.1 SLC13 family permease |
| 167(1) | WP_000570743.1 | WP_000570743.1 SLC13 family permease |
| 167(1) | WP_098111642.1 | WP_098111642.1 SLC13 family permease |
| 167(1) | WP_307192908.1 | WP_307192908.1 SLC13 family permease |
| 167(1) | WP_074565299.1 | WP_074565299.1 SLC13 family permease |
| 167(1) | WP_208752236.1 | WP_208752236.1 SLC13 family permease |
| 167(1) | WP_053564623.1 | WP_053564623.1 SLC13 family permease |
| 167(1) | WP_166704998.1 | WP_166704998.1 SLC13 family permease |
| 167(1) | WP_242264954.1 | WP_242264954.1 SLC13 family permease |
| 167(1) | WP_151630599.1 | WP_151630599.1 SLC13 family permease |
| 167(1) | WP_103655292.1 | WP_103655292.1 SLC13 family permease |
| 167(1) | WP_242251796.1 | WP_242251796.1 SLC13 family permease |
| 167(1) | WP_144611192.1 | WP_144611192.1 SLC13 family permease |
| 167(1) | WP_258126017.1 | WP_258126017.1 SLC13 family permease |

| | | |
|---|---|---|
| 168(1) | WP_308615619.1 | WP_308615619.1 hypothetical protein |
| 168(1) | WP_277157938.1 | WP_277157938.1 hypothetical protein |
| 168(1) | WP_291289936.1 | WP_291289936.1 hypothetical protein |
| 168(1) | WP_270472006.1 | WP_270472006.1 hypothetical protein |
| 168(1) | WP_227917890.1 | WP_227917890.1 hypothetical protein |
| 168(1) | WP_270472010.1 | WP_270472010.1 hypothetical protein |
| 168(1) | WP_297183840.1 | WP_297183840.1 hypothetical protein |
| 168(1) | WP_277156181.1 | WP_277156181.1 hypothetical protein |
| 168(1) | WP_212330876.1 | WP_212330876.1 hypothetical protein |
| 168(1) | WP_180965897.1 | WP_180965897.1 hypothetical protein |
| 168(1) | WP_303942538.1 | WP_303942538.1 hypothetical protein |
| 168(1) | WP_070638657.1 | WP_070638657.1 hypothetical protein |
| 168(1) | WP_302153076.1 | WP_302153076.1 hypothetical protein |
| 168(1) | WP_270472009.1 | WP_270472009.1 hypothetical protein |
| 168(1) | WP_277175583.1 | WP_277175583.1 hypothetical protein |
| 168(1) | WP_212330873.1 | WP_212330873.1 hypothetical protein |
| 168(1) | WP_143242402.1 | WP_143242402.1 hypothetical protein |
| 168(1) | WP_273191256.1 | WP_273191256.1 hypothetical protein |
| 168(1) | WP_270472032.1 | WP_270472032.1 hypothetical protein |
| 168(1) | WP_270472007.1 | WP_270472007.1 hypothetical protein |
| 168(1) | WP_212330859.1 | WP_212330859.1 hypothetical protein |
| 168(1) | WP_270472031.1 | WP_270472031.1 hypothetical protein |
| 168(1) | WP_273383876.1 | WP_273383876.1 hypothetical protein |
| 168(1) | WP_058271113.1 | WP_058271113.1 hypothetical protein |
| 168(1) | WP_270472008.1 | WP_270472008.1 hypothetical protein |
| 168(1) | WP_307738728.1 | WP_307738728.1 hypothetical protein |
| 168(1) | WP_291288267.1 | WP_291288267.1 hypothetical protein |
| 168(1) | WP_277142083.1 | WP_277142083.1 hypothetical protein |

| | | |
|---|---|---|
| 169(1) | WP_195891971.1 | WP_195891971.1 branched-chain amino acid ABC transporter permease |
| 169(8) | WP_005546487.1 | WP_005546487.1 ABC transporter permease |
| 169(1) | WP_174479807.1 | WP_174479807.1 branched-chain amino acid ABC transporter permease |
| 169(1) | WP_042462828.1 | WP_042462828.1 branched-chain amino acid ABC transporter permease |
| 169(2) | WP_044355255.1 | WP_044355255.1 ABC transporter permease |
| 169(1) | WP_042219017.1 | WP_042219017.1 branched-chain amino acid ABC transporter permease |
| 169(2) | WP_179093401.1 | WP_179093401.1 branched-chain amino acid ABC transporter permease |
| 169(2) | WP_039296535.1 | WP_039296535.1 branched-chain amino acid ABC transporter permease |
| 169(2) | WP_021261895.1 | WP_021261895.1 ABC transporter permease |
| 169(1) | WP_021253130.1 | WP_021253130.1 ABC transporter permease |
| 169(1) | WP_268623237.1 | WP_268623237.1 ABC transporter permease |
| 169(1) | WP_065291258.1 | WP_065291258.1 ABC transporter permease |
| 169(2) | WP_072727815.1 | WP_072727815.1 ABC transporter permease |
| 169(1) | WP_028533187.1 | WP_028533187.1 ABC transporter permease |
| 169(1) | WP_090510227.1 | WP_090510227.1 ABC transporter permease |

| | | |
|---|---|---|
| 170(2) | WP_003750481.1 | WP_003750481.1 hypothetical protein |
| 170(2) | WP_185474089.1 | WP_185474089.1 hypothetical protein |
| 170(2) | WP_270995162.1 | WP_270995162.1 hypothetical protein |
| 170(2) | WP_185413339.1 | WP_185413339.1 hypothetical protein |
| 170(2) | WP_194334465.1 | WP_194334465.1 hypothetical protein |
| 170(3) | WP_270996241.1 | WP_270996241.1 hypothetical protein |
| 170(1) | WP_185451814.1 | WP_185451814.1 hypothetical protein |
| 170(6) | WP_185335856.1 | WP_185335856.1 hypothetical protein |
| 170(5) | WP_003745059.1 | WP_003745059.1 hypothetical protein |
| 170(2) | WP_200372544.1 | WP_200372544.1 hypothetical protein |

| | | |
|---|---|---|
| 171(1) | WP_248251172.1 | WP_248251172.1 DMT family transporter |
| 171(1) | WP_283892162.1 | WP_283892162.1 DMT family transporter |
| 171(1) | WP_289483775.1 | WP_289483775.1 DMT family transporter |
| 171(1) | WP_289475089.1 | WP_289475089.1 DMT family transporter |
| 171(1) | WP_148450932.1 | WP_148450932.1 DMT family transporter |
| 171(1) | WP_290401981.1 | WP_290401981.1 DMT family transporter |
| 171(1) | WP_251598151.1 | WP_251598151.1 DMT family transporter |
| 171(1) | WP_163853484.1 | WP_163853484.1 DMT family transporter |
| 171(1) | WP_256227705.1 | WP_256227705.1 DMT family transporter |
| 171(1) | WP_301167489.1 | WP_301167489.1 DMT family transporter |
| 171(1) | WP_197937406.1 | WP_197937406.1 EamA family transporter |
| 171(1) | WP_090239199.1 | WP_090239199.1 DMT family transporter |
| 171(1) | WP_211021538.1 | WP_211021538.1 DMT family transporter |
| 171(1) | WP_189597460.1 | WP_189597460.1 DMT family transporter |
| 171(2) | WP_063188082.1 | WP_063188082.1 DMT family transporter |
| 171(1) | WP_120462963.1 | WP_120462963.1 DMT family transporter |
| 171(1) | WP_265408855.1 | WP_265408855.1 DMT family transporter |
| 171(1) | WP_073008288.1 | WP_073008288.1 DMT family transporter |
| 171(1) | WP_179223591.1 | WP_179223591.1 DMT family transporter |
| 171(1) | WP_122971520.1 | WP_122971520.1 DMT family transporter |
| 171(1) | WP_051775648.1 | WP_051775648.1 DMT family transporter |
| 171(1) | WP_062239903.1 | WP_062239903.1 DMT family transporter |
| 171(1) | WP_247083060.1 | WP_247083060.1 DMT family transporter |
| 171(2) | WP_061975561.1 | WP_061975561.1 DMT family transporter |

171(1) WP_251548775.1 WP_251548775.1 DMT family transporter

172(1) WP_110931535.1 WP_110931535.1 hypothetical protein
172(1) WP_212981449.1 WP_212981449.1 hypothetical protein
172(1) WP_212981566.1 WP_212981566.1 hypothetical protein
172(1) WP_207994171.1 WP_207994171.1 hypothetical protein
172(1) WP_222861017.1 WP_222861017.1 hypothetical protein
172(1) WP_307394647.1 WP_307394647.1 hypothetical protein
172(1) WP_110931534.1 WP_110931534.1 hypothetical protein
172(1) WP_212981567.1 WP_212981567.1 hypothetical protein
172(1) WP_307394645.1 WP_307394645.1 hypothetical protein
172(1) WP_024632677.1 WP_024632677.1 hypothetical protein
172(1) WP_055105800.1 WP_055105800.1 hypothetical protein
172(1) WP_017149781.1 WP_017149781.1 hypothetical protein
172(1) WP_110931539.1 WP_110931539.1 hypothetical protein
172(1) WP_211023034.1 WP_211023034.1 hypothetical protein
172(1) WP_055105796.1 WP_055105796.1 hypothetical protein
172(1) WP_256704223.1 WP_256704223.1 hypothetical protein
172(2) WP_068782413.1 WP_068782413.1 hypothetical protein
172(1) WP_110931540.1 WP_110931540.1 hypothetical protein
172(1) WP_307394643.1 WP_307394643.1 hypothetical protein
172(1) WP_211023035.1 WP_211023035.1 hypothetical protein
172(1) WP_017692361.1 WP_017692361.1 hypothetical protein
172(1) WP_110931538.1 WP_110931538.1 hypothetical protein
172(1) WP_212981569.1 WP_212981569.1 hypothetical protein
172(1) WP_166244898.1 WP_166244898.1 DUF1433 domain-containing protein
172(1) WP_211023033.1 WP_211023033.1 hypothetical protein
172(1) WP_212981447.1 WP_212981447.1 hypothetical protein

173(1) WP_088230454.1 WP_088230454.1 GerW family sporulation protein
173(8) WP_000349987.1 WP_000349987.1 GerW family sporulation protein
173(6) WP_000349991.1 WP_000349991.1 GerW family sporulation protein
173(10) WP_000349986.1 WP_000349986.1 GerW family sporulation protein
173(1) WP_048558214.1 WP_048558214.1 GerW family sporulation protein

174(1) WP_225999418.1 WP_225999418.1 CRISPR-associated endonuclease Cas2
174(1) WP_025716887.1 WP_025716887.1 CRISPR-associated endonuclease Cas2
174(1) WP_025364686.1 WP_025364686.1 CRISPR-associated endonuclease Cas2
174(1) WP_094155397.1 WP_094155397.1 CRISPR-associated endonuclease Cas2
174(1) WP_137062834.1 WP_137062834.1 CRISPR-associated endonuclease Cas2
174(1) WP_152381402.1 WP_152381402.1 CRISPR-associated endonuclease Cas2
174(1) WP_017427976.1 WP_017427976.1 CRISPR-associated endonuclease Cas2
174(1) WP_254327973.1 WP_254327973.1 CRISPR-associated endonuclease Cas2
174(1) WP_238588367.1 WP_238588367.1 CRISPR-associated endonuclease Cas2
174(1) WP_068497046.1 WP_068497046.1 CRISPR-associated endonuclease Cas2
174(1) WP_231517282.1 WP_231517282.1 CRISPR-associated endonuclease Cas2
174(1) WP_254325787.1 WP_254325787.1 CRISPR-associated endonuclease Cas2
174(2) WP_014281197.1 WP_014281197.1 CRISPR-associated endonuclease Cas2
174(11) WP_023988276.1 WP_023988276.1 CRISPR-associated endonuclease Cas2
174(1) WP_063208753.1 WP_063208753.1 CRISPR-associated endonuclease Cas2

175(1) WP_255291105.1 WP_255291105.1 hypothetical protein
175(2) WP_254905509.1 WP_254905509.1 hypothetical protein
175(1) WP_269436849.1 WP_269436849.1 hypothetical protein
175(1) WP_098284019.1 WP_098284019.1 hypothetical protein
175(7) WP_254905302.1 WP_254905302.1 hypothetical protein
175(1) WP_000266199.1 WP_000266199.1 hypothetical protein
175(1) WP_268988596.1 WP_268988596.1 hypothetical protein
175(4) WP_238601968.1 WP_238601968.1 hypothetical protein
175(3) WP_000266198.1 WP_000266198.1 hypothetical protein
175(4) WP_255262000.1 WP_255262000.1 hypothetical protein
175(1) WP_086412800.1 WP_086412800.1 hypothetical protein

176(1) WP_021261899.1 WP_021261899.1 DUF2500 domain-containing protein
176(1) WP_197256881.1 WP_197256881.1 hypothetical protein
176(1) WP_165979808.1 WP_165979808.1 DUF2500 domain-containing protein
176(2) WP_021253134.1 WP_021253134.1 DUF2500 domain-containing protein
176(1) WP_289355909.1 WP_289355909.1 DUF2500 domain-containing protein
176(1) WP_182914760.1 WP_182914760.1 DUF2500 domain-containing protein
176(1) WP_164776262.1 WP_164776262.1 DUF2500 domain-containing protein
176(2) WP_005546478.1 WP_005546478.1 DUF2500 domain-containing protein
176(2) WP_165643412.1 WP_165643412.1 DUF2500 domain-containing protein
176(1) WP_174811282.1 WP_174811282.1 DUF2500 family protein
176(1) WP_111153308.1 WP_111153308.1 DUF2500 domain-containing protein
176(1) WP_142543139.1 WP_142543139.1 DUF2500 domain-containing protein
176(1) WP_231515155.1 WP_231515155.1 DUF2500 domain-containing protein
176(2) WP_072727812.1 WP_072727812.1 DUF2500 domain-containing protein
176(1) WP_006674873.1 WP_006674873.1 DUF2500 domain-containing protein
176(1) WP_254659775.1 WP_254659775.1 DUF2500 domain-containing protein
176(1) WP_281477399.1 WP_281477399.1 DUF2500 domain-containing protein
176(4) WP_163977068.1 WP_163977068.1 DUF2500 domain-containing protein
176(1) WP_272045593.1 WP_272045593.1 DUF2500 family protein

177(1) WP_273061458.1 WP_273061458.1 penicillin-binding protein 2
177(1) WP_223592694.1 WP_223592694.1 PBP1A family penicillin-binding protein
177(1) WP_019120058.1 WP_019120058.1 penicillin-binding transpeptidase domain-containing protein
177(1) WP_215032154.1 WP_215032154.1 penicillin-binding transpeptidase domain-containing protein
177(1) WP_226086186.1 WP_226086186.1 penicillin-binding transpeptidase domain-containing protein
177(1) WP_132008124.1 WP_132008124.1 penicillin-binding transpeptidase domain-containing protein
177(1) WP_102264562.1 WP_102264562.1 penicillin-binding transpeptidase domain-containing protein
177(1) WP_283904352.1 WP_283904352.1 penicillin-binding transpeptidase domain-containing protein
177(1) WP_079506366.1 WP_079506366.1 penicillin-binding transpeptidase domain-containing protein
177(1) WP_226642850.1 WP_226642850.1 penicillin-binding transpeptidase domain-containing protein
177(1) WP_251542591.1 WP_251542591.1 penicillin-binding transpeptidase domain-containing protein
177(1) WP_307738713.1 WP_307738713.1 FtsW/RodA/SpoVE family cell cycle protein
177(1) WP_026572733.1 WP_026572733.1 penicillin-binding transpeptidase domain-containing protein
177(1) WP_214902966.1 WP_214902966.1 penicillin-binding transpeptidase domain-containing protein
177(1) WP_210364441.1 WP_210364441.1 penicillin-binding transpeptidase domain-containing protein
177(1) WP_242841133.1 WP_242841133.1 penicillin-binding transpeptidase domain-containing protein
177(2) WP_133332974.1 WP_133332974.1 penicillin-binding transpeptidase domain-containing protein
177(1) WP_144479086.1 WP_144479086.1 penicillin-binding transpeptidase domain-containing protein
177(1) WP_226679113.1 WP_226679113.1 penicillin-binding transpeptidase domain-containing protein
177(1) WP_041964588.1 WP_041964588.1 penicillin-binding transpeptidase domain-containing protein

177(1) WP_306049002.1 WP_306049002.1 NTF2-like N-terminal transpeptidase domain-containing protein
177(1) WP_251466784.1 WP_251466784.1 penicillin-binding transpeptidase domain-containing protein
177(1) WP_198310837.1 WP_198310837.1 penicillin-binding transpeptidase domain-containing protein
177(1) WP_051922779.1 WP_051922779.1 transglycosylase domain-containing protein
177(1) WP_191272877.1 WP_191272877.1 penicillin-binding transpeptidase domain-containing protein

178(4) WP_185363332.1 WP_185363332.1 trifunctional transcriptional activator/DNA repair protein Ada/methylated-DNA--
178(2) WP_185383819.1 WP_185383819.1 trifunctional transcriptional activator/DNA repair protein Ada/methylated-DNA--
178(3) WP_003745063.1 WP_003745063.1 trifunctional transcriptional activator/DNA repair protein Ada/methylated-DNA--
178(2) WP_185350593.1 WP_185350593.1 trifunctional transcriptional activator/DNA repair protein Ada/methylated-DNA--
178(4) WP_185340300.1 WP_185340300.1 trifunctional transcriptional activator/DNA repair protein Ada/methylated-DNA--
178(2) WP_038406389.1 WP_038406389.1 trifunctional transcriptional activator/DNA repair protein Ada/methylated-DNA--
178(2) WP_149051228.1 WP_149051228.1 trifunctional transcriptional activator/DNA repair protein Ada/methylated-DNA--
178(2) WP_185639004.1 WP_185639004.1 trifunctional transcriptional activator/DNA repair protein Ada/methylated-DNA--
178(4) WP_203573363.1 WP_203573363.1 trifunctional transcriptional activator/DNA repair protein Ada/methylated-DNA--

179(1) WP_111503202.1 WP_111503202.1 AI-2E family transporter
179(1) WP_251456535.1 WP_251456535.1 AI-2E family transporter
179(1) WP_076208249.1 WP_076208249.1 AI-2E family transporter
179(2) WP_076104569.1 WP_076104569.1 AI-2E family transporter
179(3) WP_046217706.1 WP_046217706.1 AI-2E family transporter
179(1) WP_094870383.1 WP_094870383.1 AI-2E family transporter
179(2) WP_019392109.1 WP_019392109.1 AI-2E family transporter
179(1) WP_076303949.1 WP_076303949.1 AI-2E family transporter
179(1) WP_076279387.1 WP_076279387.1 AI-2E family transporter
179(1) WP_076309111.1 WP_076309111.1 AI-2E family transporter
179(4) WP_172460052.1 WP_172460052.1 AI-2E family transporter
179(1) WP_076100015.1 WP_076100015.1 AI-2E family transporter
179(1) WP_064505029.1 WP_064505029.1 AI-2E family transporter
179(3) WP_185183367.1 WP_185183367.1 AI-2E family transporter
179(1) WP_019120065.1 WP_019120065.1 AI-2E family transporter
179(1) WP_076135819.1 WP_076135819.1 AI-2E family transporter

180(1) WP_019424163.1 WP_019424163.1 class II aldolase/adducin family protein
180(1) WP_087433590.1 WP_087433590.1 class II aldolase/adducin family protein
180(1) WP_065291286.1 WP_065291286.1 class II aldolase/adducin family protein
180(2) WP_005546495.1 WP_005546495.1 class II aldolase/adducin family protein
180(1) WP_260612182.1 WP_260612182.1 class II aldolase/adducin family protein
180(4) WP_163977062.1 WP_163977062.1 class II aldolase/adducin family protein
180(1) WP_051249879.1 WP_051249879.1 class II aldolase/adducin family protein
180(2) WP_268599735.1 WP_268599735.1 class II aldolase/adducin family protein
180(1) WP_240463839.1 WP_240463839.1 class II aldolase/adducin family protein
180(3) WP_072728476.1 WP_072728476.1 class II aldolase/adducin family protein
180(1) WP_036621124.1 WP_036621124.1 class II aldolase/adducin family protein
180(1) WP_195953739.1 WP_195953739.1 L-ribulose-5-phosphate 4-epimerase
180(1) WP_138189450.1 WP_138189450.1 class II aldolase/adducin family protein
180(1) WP_036667448.1 WP_036667448.1 class II aldolase/adducin family protein
180(2) WP_044355260.1 WP_044355260.1 class II aldolase/adducin family protein
180(1) WP_176392828.1 WP_176392828.1 class II aldolase/adducin family protein
180(1) WP_267934045.1 WP_267934045.1 class II aldolase/adducin family protein

181(1) WP_215570362.1 WP_215570362.1 hypothetical protein
181(2) WP_098768171.1 WP_098768171.1 hypothetical protein
181(1) WP_098991579.1 WP_098991579.1 hypothetical protein
181(1) WP_255287906.1 WP_255287906.1 hypothetical protein
181(1) WP_000912021.1 WP_000912021.1 endonuclease/exonuclease/phosphatase family protein
181(1) WP_240326483.1 WP_240326483.1 hypothetical protein
181(1) WP_215591196.1 WP_215591196.1 hypothetical protein
181(11) WP_000912022.1 WP_000912022.1 endonuclease/exonuclease/phosphatase family protein
181(1) WP_097905190.1 WP_097905190.1 hypothetical protein
181(1) WP_002158056.1 WP_002158056.1 hypothetical protein
181(1) WP_289398690.1 WP_289398690.1 hypothetical protein
181(1) WP_269325814.1 WP_269325814.1 hypothetical protein
181(2) WP_002148351.1 WP_002148351.1 hypothetical protein

182(1) WP_183578476.1 WP_183578476.1 hypothetical protein
182(3) WP_152402389.1 WP_152402389.1 hypothetical protein
182(1) WP_160496921.1 WP_160496921.1 hypothetical protein
182(1) WP_212949038.1 WP_212949038.1 hypothetical protein
182(2) WP_145156812.1 WP_145156812.1 hypothetical protein
182(1) WP_186325900.1 WP_186325900.1 hypothetical protein
182(1) WP_251674660.1 WP_251674660.1 hypothetical protein
182(1) WP_105443564.1 WP_105443564.1 hypothetical protein
182(1) WP_094593883.1 WP_094593883.1 hypothetical protein
182(1) WP_160037495.1 WP_160037495.1 hypothetical protein
182(4) WP_076167850.1 WP_076167850.1 hypothetical protein
182(1) WP_116231169.1 WP_116231169.1 hypothetical protein
182(1) WP_194230396.1 WP_194230396.1 hypothetical protein
182(1) WP_212980182.1 WP_212980182.1 hypothetical protein
182(2) WP_136604824.1 WP_136604824.1 hypothetical protein
182(1) WP_145134807.1 WP_145134807.1 hypothetical protein
182(1) WP_244719674.1 WP_244719674.1 hypothetical protein

183(1) WP_200633295.1 WP_200633295.1 hypothetical protein
183(1) WP_145134821.1 WP_145134821.1 hypothetical protein
183(1) WP_145948995.1 WP_145948995.1 hypothetical protein
183(1) WP_052146792.1 WP_052146792.1 hypothetical protein
183(1) WP_094593869.1 WP_094593869.1 hypothetical protein
183(1) WP_244719597.1 WP_244719597.1 hypothetical protein
183(1) WP_145021747.1 WP_145021747.1 hypothetical protein
183(1) WP_194230409.1 WP_194230409.1 hypothetical protein
183(1) WP_263880221.1 WP_263880221.1 hypothetical protein
183(1) WP_213649974.1 WP_213649974.1 hypothetical protein
183(1) WP_160037480.1 WP_160037480.1 hypothetical protein

183(1) WP_152402369.1 WP_152402369.1 hypothetical protein
183(1) WP_212980189.1 WP_212980189.1 hypothetical protein
183(1) WP_145156802.1 WP_145156802.1 hypothetical protein
183(1) WP_183578434.1 WP_183578434.1 hypothetical protein
183(6) WP_076167878.1 WP_076167878.1 hypothetical protein
183(1) WP_136604809.1 WP_136604809.1 hypothetical protein
183(1) WP_116231180.1 WP_116231180.1 hypothetical protein
183(1) WP_160496937.1 WP_160496937.1 hypothetical protein

184(2) WP_185639005.1 WP_185639005.1 isocitrate lyase/phosphoenolpyruvate mutase family protein
184(2) WP_270993644.1 WP_270993644.1 isocitrate lyase/phosphoenolpyruvate mutase family protein
184(4) WP_185340332.1 WP_185340332.1 isocitrate lyase/phosphoenolpyruvate mutase family protein
184(2) WP_185472570.1 WP_185472570.1 isocitrate lyase/phosphoenolpyruvate mutase family protein
184(4) WP_038406388.1 WP_038406388.1 isocitrate lyase/phosphoenolpyruvate mutase family protein
184(2) WP_046325933.1 WP_046325933.1 isocitrate lyase/phosphoenolpyruvate mutase family protein
184(2) WP_149051227.1 WP_149051227.1 isocitrate lyase/phosphoenolpyruvate mutase family protein
184(2) WP_185451831.1 WP_185451831.1 isocitrate lyase/phosphoenolpyruvate mutase family protein
184(2) WP_194335287.1 WP_194335287.1 isocitrate lyase/phosphoenolpyruvate mutase family protein
184(2) WP_185474096.1 WP_185474096.1 isocitrate lyase/phosphoenolpyruvate mutase family protein

185(1) WP_072336510.1 WP_072336510.1 glutamine--fructose-6-phosphate transaminase (isomerizing)
185(2) WP_010963719.1 WP_010963719.1 6-phospho-3-hexuloisomerase
185(1) WP_204412740.1 WP_204412740.1 MurR/RpiR family transcriptional regulator
185(1) WP_048826623.1 WP_048826623.1 MurR/RpiR family transcriptional regulator
185(1) WP_236533772.1 WP_236533772.1 MurR/RpiR family transcriptional regulator
185(1) WP_205635663.1 WP_205635663.1 SIS domain-containing protein
185(1) WP_205208650.1 WP_205208650.1 SIS domain-containing protein
185(2) WP_046217698.1 WP_046217698.1 SIS domain-containing protein
185(1) WP_197234704.1 WP_197234704.1 SIS domain-containing protein
185(1) WP_197234706.1 WP_197234706.1 SIS domain-containing protein
185(1) WP_306077398.1 WP_306077398.1 SIS domain-containing protein
185(6) WP_037993167.1 WP_037993167.1 MurR/RpiR family transcriptional regulator
185(1) WP_251456512.1 WP_251456512.1 SIS domain-containing protein
185(1) WP_099223962.1 WP_099223962.1 MurR/RpiR family transcriptional regulator
185(1) WP_181731448.1 WP_181731448.1 MurR/RpiR family transcriptional regulator
185(1) WP_251456510.1 WP_251456510.1 SIS domain-containing protein
185(1) WP_181737023.1 WP_181737023.1 MurR/RpiR family transcriptional regulator

186(1) WP_192510488.1 WP_192510488.1 transposase
186(1) WP_281255103.1 WP_281255103.1 IS3 family transposase
186(1) WP_307399133.1 WP_307399133.1 Mu transposase C-terminal domain-containing protein, partial
186(4) WP_223196724.1 WP_223196724.1 IS3 family transposase
186(1) WP_235827637.1 WP_235827637.1 hypothetical protein
186(1) WP_235827636.1 WP_235827636.1 hypothetical protein
186(1) WP_169028883.1 WP_169028883.1 transposase
186(1) WP_235827417.1 WP_235827417.1 hypothetical protein
186(1) WP_087943621.1 WP_087943621.1 IS3-like element ISBth10 family transposase
186(1) WP_101300061.1 WP_101300061.1 IS6 family transposase
186(1) WP_307388998.1 WP_307388998.1 DDE-type integrase/transposase/recombinase
186(1) WP_242971601.1 WP_242971601.1 hypothetical protein
186(1) WP_307199958.1 WP_307199958.1 transposase
186(1) WP_151208713.1 WP_151208713.1 IS3 family transposase
186(1) WP_025685041.1 WP_025685041.1 IS3 family transposase, partial
186(1) WP_139795984.1 WP_139795984.1 integrase core domain-containing protein, partial
186(1) WP_149096531.1 WP_149096531.1 IS3 family transposase, partial
186(1) WP_108745553.1 WP_108745553.1 IS3 family transposase
186(1) WP_235827638.1 WP_235827638.1 Mu transposase C-terminal domain-containing protein
186(1) WP_205001934.1 WP_205001934.1 IS3 family transposase
186(1) WP_143187949.1 WP_143187949.1 IS3 family transposase, partial

187(1) WP_015595767.1 WP_015595767.1 dTDP-4-dehydrorhamnose reductase
187(1) WP_098579107.1 WP_098579107.1 nucleoside-diphosphate sugar epimerase/dehydratase
187(1) WP_309098605.1 WP_309098605.1 UDP-glucose 4-epimerase GalE
187(1) WP_136790004.1 WP_136790004.1 NAD-dependent epimerase/dehydratase family protein
187(1) WP_204445188.1 WP_204445188.1 UDP-glucose 4-epimerase GalE
187(1) WP_047153282.1 WP_047153282.1 L-threonine 3-dehydrogenase
187(1) WP_272801663.1 WP_272801663.1 NAD-dependent epimerase/dehydratase family protein
187(1) WP_015595768.1 WP_015595768.1 dTDP-glucose 4,6-dehydratase
187(1) WP_304216853.1 WP_304216853.1 UDP-glucose 4-epimerase GalE
187(1) WP_099361330.1 WP_099361330.1 UDP-glucose 4-epimerase GalE
187(1) WP_010676874.1 WP_010676874.1 UDP-glucose 4-epimerase GalE
187(1) WP_043069428.1 WP_043069428.1 SDR family oxidoreductase
187(1) WP_040418513.1 WP_040418513.1 nucleoside-diphosphate sugar epimerase/dehydratase
187(1) WP_243495569.1 WP_243495569.1 GDP-L-fucose synthase
187(1) WP_098579109.1 WP_098579109.1 NAD-dependent epimerase/dehydratase family protein
187(1) WP_232699397.1 WP_232699397.1 NAD(P)-dependent oxidoreductase
187(1) WP_224839849.1 WP_224839849.1 UDP-glucose 4-epimerase GalE
187(1) WP_116353261.1 WP_116353261.1 UDP-glucose 4-epimerase GalE
187(1) WP_232278753.1 WP_232278753.1 TIGR01777 family oxidoreductase
187(1) WP_229139911.1 WP_229139911.1 UDP-glucose 4-epimerase GalE
187(1) WP_154135658.1 WP_154135658.1 TIGR01777 family oxidoreductase
187(1) WP_099354895.1 WP_099354895.1 UDP-glucose 4-epimerase GalE
187(1) WP_077619660.1 WP_077619660.1 UDP-glucose 4-epimerase GalE
187(1) WP_221875365.1 WP_221875365.1 UDP-glucose 4-epimerase GalE

188(5) WP_046325934.1 WP_046325934.1 2-hydroxyacid dehydrogenase family protein
188(4) WP_003718156.1 WP_003718156.1 NAD(P)-dependent oxidoreductase
188(2) WP_149051269.1 WP_149051269.1 2-hydroxyacid dehydrogenase family protein
188(1) WP_005604652.1 WP_005604652.1 D-2-hydroxyacid dehydrogenase
188(2) WP_185639002.1 WP_185639002.1 2-hydroxyacid dehydrogenase family protein
188(2) WP_185474087.1 WP_185474087.1 2-hydroxyacid dehydrogenase family protein
188(6) WP_185340301.1 WP_185340301.1 2-hydroxyacid dehydrogenase family protein
188(1) WP_223592754.1 WP_223592754.1 D-glycerate dehydrogenase

189(1) WP_242400029.1 WP_242400029.1 ankyrin repeat domain-containing protein
189(9) WP_000698515.1 WP_000698515.1 ankyrin repeat domain-containing protein
189(1) WP_163255263.1 WP_163255263.1 ankyrin repeat domain-containing protein
189(6) WP_000698516.1 WP_000698516.1 ankyrin repeat domain-containing protein
189(1) WP_149885491.1 WP_149885491.1 ankyrin repeat domain-containing protein
189(3) WP_071736282.1 WP_071736282.1 ankyrin repeat domain-containing protein
189(1) WP_098167590.1 WP_098167590.1 ankyrin repeat domain-containing protein, partial
189(1) WP_240379738.1 WP_240379738.1 ankyrin repeat domain-containing protein

| | | |
|---|---|---|
| 190(1) | WP_063261714.1 | WP_063261714.1 hypothetical protein |
| 190(1) | WP_208906817.1 | WP_208906817.1 hypothetical protein |
| 190(1) | WP_228492698.1 | WP_228492698.1 hypothetical protein |
| 190(1) | WP_046946393.1 | WP_046946393.1 hypothetical protein |
| 190(1) | WP_163061519.1 | WP_163061519.1 hypothetical protein |
| 190(1) | WP_140159132.1 | WP_140159132.1 hypothetical protein |
| 190(1) | WP_071712836.1 | WP_071712836.1 hypothetical protein |
| 190(4) | WP_000473566.1 | WP_000473566.1 hypothetical protein |
| 190(1) | WP_000473564.1 | WP_000473564.1 hypothetical protein |
| 190(6) | WP_065223578.1 | WP_065223578.1 hypothetical protein |
| 190(1) | WP_003300966.1 | WP_003300966.1 hypothetical protein |
| 190(3) | WP_000473563.1 | WP_000473563.1 hypothetical protein |
| 190(1) | WP_257206140.1 | WP_257206140.1 hypothetical protein |
| 191(1) | WP_077852732.1 | WP_077852732.1 hypothetical protein |
| 191(1) | WP_077852736.1 | WP_077852736.1 hypothetical protein |
| 191(1) | WP_077893461.1 | WP_077893461.1 hypothetical protein |
| 191(1) | WP_257675917.1 | WP_257675917.1 hypothetical protein |
| 191(1) | WP_077893459.1 | WP_077893459.1 hypothetical protein |
| 191(1) | WP_077832288.1 | WP_077832288.1 hypothetical protein |
| 191(1) | WP_077893457.1 | WP_077893457.1 hypothetical protein |
| 191(1) | WP_077893463.1 | WP_077893463.1 hypothetical protein |
| 191(1) | WP_077852739.1 | WP_077852739.1 hypothetical protein |
| 191(1) | WP_077852737.1 | WP_077852737.1 hypothetical protein |
| 191(1) | WP_077893464.1 | WP_077893464.1 hypothetical protein |
| 191(1) | WP_077852733.1 | WP_077852733.1 hypothetical protein |
| 191(1) | WP_077832287.1 | WP_077832287.1 hypothetical protein |
| 191(1) | WP_257675919.1 | WP_257675919.1 hypothetical protein |
| 191(1) | WP_077852735.1 | WP_077852735.1 hypothetical protein |
| 191(1) | WP_077832285.1 | WP_077832285.1 hypothetical protein |
| 191(1) | WP_077893458.1 | WP_077893458.1 hypothetical protein |
| 191(1) | WP_077852734.1 | WP_077852734.1 hypothetical protein |
| 191(1) | WP_077893460.1 | WP_077893460.1 hypothetical protein |
| 191(1) | WP_077832286.1 | WP_077832286.1 hypothetical protein |
| 191(2) | WP_077852738.1 | WP_077852738.1 hypothetical protein |
| 191(1) | WP_077893462.1 | WP_077893462.1 hypothetical protein |
| 192(1) | WP_263880213.1 | WP_263880213.1 hypothetical protein |
| 192(1) | WP_212983260.1 | WP_212983260.1 hypothetical protein |
| 192(1) | WP_183578478.1 | WP_183578478.1 hypothetical protein |
| 192(1) | WP_160496920.1 | WP_160496920.1 hypothetical protein |
| 192(2) | WP_145134806.1 | WP_145134806.1 hypothetical protein |
| 192(1) | WP_179280989.1 | WP_179280989.1 hypothetical protein |
| 192(1) | WP_144025353.1 | WP_144025353.1 hypothetical protein |
| 192(1) | WP_213649961.1 | WP_213649961.1 hypothetical protein |
| 192(1) | WP_186325899.1 | WP_186325899.1 hypothetical protein |
| 192(1) | WP_145156813.1 | WP_145156813.1 hypothetical protein |
| 192(1) | WP_167527258.1 | WP_167527258.1 hypothetical protein |
| 192(1) | WP_236575326.1 | WP_236575326.1 hypothetical protein |
| 192(1) | WP_251674662.1 | WP_251674662.1 hypothetical protein |
| 192(1) | WP_212969032.1 | WP_212969032.1 hypothetical protein |
| 192(1) | WP_089546605.1 | WP_089546605.1 hypothetical protein |
| 192(1) | WP_200633299.1 | WP_200633299.1 hypothetical protein |
| 192(1) | WP_212949039.1 | WP_212949039.1 hypothetical protein |
| 192(1) | WP_145949004.1 | WP_145949004.1 hypothetical protein |
| 192(1) | WP_194230395.1 | WP_194230395.1 hypothetical protein |
| 192(1) | WP_116231168.1 | WP_116231168.1 hypothetical protein |
| 192(1) | WP_212980181.1 | WP_212980181.1 hypothetical protein |
| 192(1) | WP_136604825.1 | WP_136604825.1 hypothetical protein |
| 193(1) | WP_209861620.1 | WP_209861620.1 N-acetylglucosamine-specific PTS transporter subunit IIBC |
| 193(1) | WP_306077403.1 | WP_306077403.1 sucrose-specific PTS transporter subunit IIBC |
| 193(2) | WP_068694727.1 | WP_068694727.1 PTS glucose transporter subunit IIA |
| 193(1) | WP_034675321.1 | WP_034675321.1 sucrose-specific PTS transporter subunit IIBC |
| 193(1) | WP_209861622.1 | WP_209861622.1 PTS glucose transporter subunit IIA |
| 193(1) | WP_273061510.1 | WP_273061510.1 PTS glucose transporter subunit IIA |
| 193(1) | WP_213618826.1 | WP_213618826.1 PTS glucose transporter subunit IIA |
| 193(1) | WP_307312520.1 | WP_307312520.1 sucrose-specific PTS transporter subunit IIBC |
| 193(1) | WP_094870396.1 | WP_094870396.1 beta-glucoside-specific PTS transporter subunit IIABC |
| 193(1) | WP_127199783.1 | WP_127199783.1 PTS glucose transporter subunit IIA |
| 193(1) | WP_068619089.1 | WP_068619089.1 PTS glucose transporter subunit IIA |
| 193(1) | WP_099223965.1 | WP_099223965.1 PTS transporter subunit EIIC |
| 193(1) | WP_251375537.1 | WP_251375537.1 N-acetylglucosamine-specific PTS transporter subunit IIBC |
| 193(1) | WP_223067430.1 | WP_223067430.1 N-acetylglucosamine-specific PTS transporter subunit IIBC |
| 193(1) | WP_179603491.1 | WP_179603491.1 sucrose-specific PTS transporter subunit IIBC |
| 193(1) | WP_053403792.1 | WP_053403792.1 PTS transporter subunit IIBC |
| 193(1) | WP_094092631.1 | WP_094092631.1 N-acetylglucosamine-specific PTS transporter subunit IIBC |
| 193(1) | WP_251375538.1 | WP_251375538.1 PTS glucose transporter subunit IIA |
| 193(1) | WP_048826610.1 | WP_048826610.1 glucose-specific PTS transporter subunit IIBC |
| 193(2) | WP_242172391.1 | WP_242172391.1 PTS transporter subunit IIBC |
| 193(1) | WP_127199784.1 | WP_127199784.1 N-acetylglucosamine-specific PTS transporter subunit IIBC |
| 194(4) | WP_001990239.1 | WP_001990239.1 hypothetical protein |
| 194(3) | WP_001030793.1 | WP_001030793.1 hypothetical protein |
| 194(1) | WP_097918215.1 | WP_097918215.1 hypothetical protein |
| 194(5) | WP_030025440.1 | WP_030025440.1 hypothetical protein |
| 194(1) | WP_001030798.1 | WP_001030798.1 hypothetical protein |
| 194(1) | WP_003193391.1 | WP_003193391.1 hypothetical protein |
| 194(3) | WP_001030796.1 | WP_001030796.1 hypothetical protein |
| 194(3) | WP_002194952.1 | WP_002194952.1 hypothetical protein |
| 194(1) | WP_044584391.1 | WP_044584391.1 hypothetical protein |
| 195(1) | WP_242264955.1 | WP_242264955.1 hypothetical protein |
| 195(1) | WP_073540575.1 | WP_073540575.1 hypothetical protein |
| 195(2) | WP_098204522.1 | WP_098204522.1 hypothetical protein |
| 195(3) | WP_071736284.1 | WP_071736284.1 hypothetical protein |
| 195(8) | WP_003268550.1 | WP_003268550.1 hypothetical protein |
| 195(1) | WP_242400032.1 | WP_242400032.1 hypothetical protein |
| 195(1) | WP_000708806.1 | WP_000708806.1 hypothetical protein |
| 195(2) | WP_098254021.1 | WP_098254021.1 hypothetical protein |
| 195(1) | WP_242261456.1 | WP_242261456.1 hypothetical protein |
| 195(1) | WP_000708805.1 | WP_000708805.1 hypothetical protein |
| 195(1) | WP_088023254.1 | WP_088023254.1 hypothetical protein |

196(2) WP_087956304.1 WP_087956304.1 aspartate/glutamate racemase family protein
196(1) WP_147786230.1 WP_147786230.1 aspartate/glutamate racemase family protein
196(1) WP_149238910.1 WP_149238910.1 aspartate/glutamate racemase family protein
196(2) WP_088063242.1 WP_088063242.1 aspartate/glutamate racemase family protein
196(1) WP_122971529.1 WP_122971529.1 amino acid racemase
196(3) WP_000848549.1 WP_000848549.1 aspartate/glutamate racemase family protein
196(1) WP_262725720.1 WP_262725720.1 aspartate/glutamate racemase family protein
196(1) WP_002012596.1 WP_002012596.1 aspartate/glutamate racemase family protein
196(1) WP_060630658.1 WP_060630658.1 aspartate/glutamate racemase family protein
196(1) WP_097852380.1 WP_097852380.1 aspartate/glutamate racemase family protein
196(3) WP_065228775.1 WP_065228775.1 aspartate/glutamate racemase family protein
196(1) WP_061535948.1 WP_061535948.1 aspartate/glutamate racemase family protein
196(1) WP_061679519.1 WP_061679519.1 aspartate/glutamate racemase family protein
196(1) WP_270716439.1 WP_270716439.1 aspartate/glutamate racemase family protein
196(2) WP_000848548.1 WP_000848548.1 aspartate/glutamate racemase family protein

197(1) WP_054972086.1 WP_054972086.1 Rpn family recombination-promoting nuclease/putative transposase
197(1) WP_254366001.1 WP_254366001.1 Rpn family recombination-promoting nuclease/putative transposase
197(1) WP_254365999.1 WP_254365999.1 Rpn family recombination-promoting nuclease/putative transposase
197(1) WP_161782229.1 WP_161782229.1 Rpn family recombination-promoting nuclease/putative transposase
197(1) WP_088830532.1 WP_088830532.1 Rpn family recombination-promoting nuclease/putative transposase
197(1) WP_025848591.1 WP_025848591.1 PD-(D/E)XK nuclease family transposase
197(1) WP_163853480.1 WP_163853480.1 Rpn family recombination-promoting nuclease/putative transposase
197(1) WP_063188074.1 WP_063188074.1 Rpn family recombination-promoting nuclease/putative transposase
197(1) WP_302878980.1 WP_302878980.1 PD-(D/E)XK nuclease family transposase
197(1) WP_216791202.1 WP_216791202.1 Rpn family recombination-promoting nuclease/putative transposase
197(1) WP_208926627.1 WP_208926627.1 Rpn family recombination-promoting nuclease/putative transposase
197(1) WP_090674377.1 WP_090674377.1 Rpn family recombination-promoting nuclease/putative transposase
197(1) WP_051775650.1 WP_051775650.1 hypothetical protein
197(1) WP_251598149.1 WP_251598149.1 Rpn family recombination-promoting nuclease/putative transposase
197(1) WP_301292475.1 WP_301292475.1 Rpn family recombination-promoting nuclease/putative transposase
197(1) WP_281937382.1 WP_281937382.1 Rpn family recombination-promoting nuclease/putative transposase
197(1) WP_127484359.1 WP_127484359.1 PD-(D/E)XK nuclease family transposase
197(1) WP_302878979.1 WP_302878979.1 Rpn family recombination-promoting nuclease/putative transposase
197(1) WP_108534951.1 WP_108534951.1 Rpn family recombination-promoting nuclease/putative transposase
197(1) WP_010496602.1 WP_010496602.1 Rpn family recombination-promoting nuclease/putative transposase
197(1) WP_051775652.1 WP_051775652.1 Rpn family recombination-promoting nuclease/putative transposase
197(1) WP_254366000.1 WP_254366000.1 Rpn family recombination-promoting nuclease/putative transposase

198(1) WP_110931528.1 WP_110931528.1 hypothetical protein
198(1) WP_145939627.1 WP_145939627.1 hypothetical protein
198(1) WP_076210961.1 WP_076210961.1 hypothetical protein
198(1) WP_307199922.1 WP_307199922.1 hypothetical protein
198(1) WP_150364191.1 WP_150364191.1 hypothetical protein
198(5) WP_111273175.1 WP_111273175.1 hypothetical protein
198(1) WP_024632692.1 WP_024632692.1 hypothetical protein
198(1) WP_210087118.1 WP_210087118.1 hypothetical protein
198(1) WP_068696910.1 WP_068696910.1 hypothetical protein
198(1) WP_211082922.1 WP_211082922.1 hypothetical protein
198(1) WP_055105806.1 WP_055105806.1 hypothetical protein
198(1) WP_110899373.1 WP_110899373.1 hypothetical protein
198(1) WP_253440797.1 WP_253440797.1 hypothetical protein
198(1) WP_289376837.1 WP_289376837.1 hypothetical protein
198(1) WP_211023042.1 WP_211023042.1 hypothetical protein
198(2) WP_017692343.1 WP_017692343.1 hypothetical protein
198(1) WP_068782420.1 WP_068782420.1 hypothetical protein

199(2) WP_098204519.1 WP_098204519.1 hypothetical protein
199(1) WP_159146510.1 WP_159146510.1 hypothetical protein
199(2) WP_195431326.1 WP_195431326.1 hypothetical protein
199(1) WP_184182986.1 WP_184182986.1 hypothetical protein
199(2) WP_002148313.1 WP_002148313.1 hypothetical protein
199(1) WP_308723524.1 WP_308723524.1 hypothetical protein
199(4) WP_002171737.1 WP_002171737.1 hypothetical protein
199(2) WP_098768163.1 WP_098768163.1 hypothetical protein
199(2) WP_000041339.1 WP_000041339.1 hypothetical protein
199(5) WP_000041340.1 WP_000041340.1 hypothetical protein

200(14) WP_000867689.1 WP_000867689.1 DUF2185 domain-containing protein
200(1) WP_240379739.1 WP_240379739.1 DUF2185 domain-containing protein
200(1) WP_113732860.1 WP_113732860.1 DUF2185 domain-containing protein
200(3) WP_071736283.1 WP_071736283.1 DUF2185 domain-containing protein
200(2) WP_141524018.1 WP_141524018.1 DUF2185 domain-containing protein

201(1) WP_251481008.1 WP_251481008.1 flagellar hook-basal body protein
201(1) WP_132008184.1 WP_132008184.1 flagellar hook-basal body protein
201(1) WP_121610607.1 WP_121610607.1 flagellar hook-basal body protein
201(1) WP_215032100.1 WP_215032100.1 flagellar hook-basal body protein
201(1) WP_226086167.1 WP_226086167.1 flagellar hook-basal body protein
201(1) WP_102264539.1 WP_102264539.1 flagellar hook-basal body protein
201(1) WP_226679093.1 WP_226679093.1 flagellar hook-basal body protein
201(1) WP_251542633.1 WP_251542633.1 flagellar hook-basal body protein
201(1) WP_214706014.1 WP_214706014.1 flagellar hook-basal body protein
201(1) WP_251481009.1 WP_251481009.1 flagellar hook-basal body protein

201(1) WP_068673748.1 WP_068673748.1 flagellar hook-basal body protein
201(1) WP_226679092.1 WP_226679092.1 flagellar hook-basal body protein
201(1) WP_226642891.1 WP_226642891.1 flagellar hook-basal body protein
201(1) WP_214706015.1 WP_214706015.1 flagellar hook-basal body protein
201(1) WP_215032098.1 WP_215032098.1 flagellar hook-basal body protein
201(1) WP_121610606.1 WP_121610606.1 flagellar hook-basal body protein
201(1) WP_102264540.1 WP_102264540.1 flagellar hook-basal body protein
201(1) WP_226642893.1 WP_226642893.1 flagellar hook-basal body protein
201(1) WP_068673746.1 WP_068673746.1 flagellar hook-basal body protein
201(1) WP_226086166.1 WP_226086166.1 flagellar hook-basal body protein
201(1) WP_041964565.1 WP_041964565.1 flagellar hook-basal body protein

202(1) WP_088230458.1 WP_088230458.1 hypothetical protein
202(4) WP_001016543.1 WP_001016543.1 hypothetical protein
202(2) WP_002083293.1 WP_002083293.1 hypothetical protein
202(1) WP_063261715.1 WP_063261715.1 hypothetical protein
202(2) WP_001177229.1 WP_001177229.1 hypothetical protein
202(4) WP_065223577.1 WP_065223577.1 hypothetical protein
202(1) WP_033693849.1 WP_033693849.1 hypothetical protein
202(1) WP_097998241.1 WP_097998241.1 hypothetical protein
202(2) WP_098291289.1 WP_098291289.1 hypothetical protein
202(1) WP_204335713.1 WP_204335713.1 hypothetical protein
202(1) WP_208906816.1 WP_208906816.1 hypothetical protein
202(1) WP_258113054.1 WP_258113054.1 hypothetical protein

203(1) WP_111153310.1 WP_111153310.1 DUF2569 domain-containing protein
203(1) WP_143801073.1 WP_143801073.1 DUF2569 domain-containing protein
203(1) WP_281477395.1 WP_281477395.1 DUF2569 domain-containing protein
203(1) WP_111153309.1 WP_111153309.1 DUF2569 domain-containing protein
203(1) WP_133378782.1 WP_133378782.1 DUF2569 domain-containing protein
203(1) WP_143801072.1 WP_143801072.1 DUF2569 domain-containing protein
203(1) WP_119794352.1 WP_119794352.1 DUF2569 domain-containing protein
203(1) WP_006674875.1 WP_006674875.1 DUF2569 domain-containing protein
203(1) WP_213427000.1 WP_213427000.1 DUF2569 domain-containing protein
203(1) WP_197256884.1 WP_197256884.1 DUF2569 domain-containing protein
203(1) WP_272827192.1 WP_272827192.1 DUF2569 family protein
203(1) WP_272827193.1 WP_272827193.1 DUF2569 family protein
203(1) WP_087442397.1 WP_087442397.1 DUF2569 domain-containing protein
203(1) WP_272045597.1 WP_272045597.1 DUF2569 domain-containing protein
203(1) WP_087442398.1 WP_087442398.1 DUF2569 domain-containing protein
203(1) WP_281477393.1 WP_281477393.1 DUF2569 domain-containing protein
203(1) WP_133378781.1 WP_133378781.1 DUF2569 domain-containing protein
203(3) WP_168178796.1 WP_168178796.1 DUF2569 domain-containing protein
203(1) WP_272827190.1 WP_272827190.1 DUF2569 family protein

204(1) WP_142543130.1 WP_142543130.1 YafY family protein
204(1) WP_268619933.1 WP_268619933.1 YafY family protein
204(1) WP_021261886.1 WP_021261886.1 YafY family protein
204(1) WP_268599731.1 WP_268599731.1 YafY family protein
204(4) WP_163977059.1 WP_163977059.1 YafY family protein
204(1) WP_305755049.1 WP_305755049.1 YafY family protein
204(1) WP_072728477.1 WP_072728477.1 YafY family protein
204(1) WP_005546504.1 WP_005546504.1 YafY family protein
204(1) WP_021253122.1 WP_021253122.1 YafY family protein
204(1) WP_065291250.1 WP_065291250.1 YafY family protein
204(2) WP_028533182.1 WP_028533182.1 YafY family protein
204(2) WP_044355258.1 WP_044355258.1 YafY family protein
204(1) WP_138186700.1 WP_138186700.1 YafY family protein
204(1) WP_251480990.1 WP_251480990.1 WYL domain-containing protein
204(1) WP_174809200.1 WP_174809200.1 WYL domain-containing protein
204(1) WP_289355905.1 WP_289355905.1 YafY family protein

205(1) WP_074048565.1 WP_074048565.1 DUF5704 domain-containing protein
205(1) WP_199925557.1 WP_199925557.1 DUF5704 domain-containing protein
205(1) WP_068696905.1 WP_068696905.1 hypothetical protein
205(1) WP_024632678.1 WP_024632678.1 DUF5704 domain-containing protein
205(1) WP_247681894.1 WP_247681894.1 DUF5704 domain-containing protein
205(1) WP_147273191.1 WP_147273191.1 hypothetical protein, partial
205(1) WP_255807588.1 WP_255807588.1 glycine-rich protein
205(1) WP_010250284.1 WP_010250284.1 DUF5704 domain-containing protein
205(1) WP_253299357.1 WP_253299357.1 DUF5704 domain-containing protein
205(1) WP_241230832.1 WP_241230832.1 DUF5704 domain-containing protein
205(1) WP_210087106.1 WP_210087106.1 DUF5704 domain-containing protein, partial
205(1) WP_169823439.1 WP_169823439.1 DUF5704 domain-containing protein
205(1) WP_244964939.1 WP_244964939.1 DUF5704 domain-containing protein
205(1) WP_307394662.1 WP_307394662.1 hypothetical protein
205(1) WP_182301847.1 WP_182301847.1 glycine rich domain-containing protein
205(1) WP_166244099.1 WP_166244099.1 DUF5704 domain-containing protein
205(1) WP_068782416.1 WP_068782416.1 DUF5704 domain-containing protein
205(1) WP_256704221.1 WP_256704221.1 DUF5704 domain-containing protein
205(1) WP_237100197.1 WP_237100197.1 DUF5704 domain-containing protein
205(1) WP_068536340.1 WP_068536340.1 DUF5704 domain-containing protein
205(1) WP_110931533.1 WP_110931533.1 DUF5704 domain-containing protein

206(11) WP_051491338.1 WP_051491338.1 ROK family protein
206(1) WP_309098601.1 WP_309098601.1 ROK family protein
206(2) WP_042125636.1 WP_042125636.1 ROK family protein
206(1) WP_154118035.1 WP_154118035.1 ROK family protein
206(4) WP_256989830.1 WP_256989830.1 ROK family protein
206(1) WP_211745844.1 WP_211745844.1 ROK family transcriptional regulator
206(1) WP_204412733.1 WP_204412733.1 ROK family protein

207(2) WP_133332977.1 WP_133332977.1 hypothetical protein
207(1) WP_306077394.1 WP_306077394.1 hypothetical protein
207(1) WP_099361345.1 WP_099361345.1 hypothetical protein
207(1) WP_307312535.1 WP_307312535.1 hypothetical protein
207(1) WP_077619637.1 WP_077619637.1 hypothetical protein
207(1) WP_034675355.1 WP_034675355.1 hypothetical protein
207(1) WP_306077395.1 WP_306077395.1 hypothetical protein
207(1) WP_179160625.1 WP_179160625.1 hypothetical protein
207(1) WP_251466795.1 WP_251466795.1 hypothetical protein
207(1) WP_179603485.1 WP_179603485.1 hypothetical protein
207(1) WP_077619638.1 WP_077619638.1 hypothetical protein
207(1) WP_142759845.1 WP_142759845.1 hypothetical protein

207(1) WP_306077393.1 WP_306077393.1 hypothetical protein
207(1) WP_304216883.1 WP_304216883.1 hypothetical protein
207(1) WP_309098619.1 WP_309098619.1 hypothetical protein
207(1) WP_179603484.1 WP_179603484.1 hypothetical protein
207(1) WP_307312538.1 WP_307312538.1 hypothetical protein
207(1) WP_198310834.1 WP_198310834.1 hypothetical protein
207(1) WP_010676853.1 WP_010676853.1 hypothetical protein
207(1) WP_307192900.1 WP_307192900.1 hypothetical protein

208(1) WP_089546606.1 WP_089546606.1 DoxX family protein
208(1) WP_212969031.1 WP_212969031.1 DoxX family protein
208(1) WP_036707215.1 WP_036707215.1 DoxX family protein
208(1) WP_194230394.1 WP_194230394.1 DoxX family protein
208(2) WP_145156814.1 WP_145156814.1 DoxX family protein
208(1) WP_201452086.1 WP_201452086.1 DoxX family protein
208(4) WP_076167846.1 WP_076167846.1 DoxX family protein
208(1) WP_094593884.1 WP_094593884.1 DoxX family protein
208(1) WP_212980180.1 WP_212980180.1 DoxX family protein
208(1) WP_145949005.1 WP_145949005.1 DoxX family protein
208(1) WP_145134804.1 WP_145134804.1 DoxX family protein
208(1) WP_145021714.1 WP_145021714.1 DoxX family protein
208(3) WP_152402393.1 WP_152402393.1 DoxX family protein
208(1) WP_116231756.1 WP_116231756.1 DoxX family protein

209(1) WP_197234716.1 WP_197234716.1 ADP-ribosylglycohydrolase family protein
209(1) WP_268583854.1 WP_268583854.1 ADP-ribosylglycohydrolase family protein
209(1) WP_242450724.1 WP_242450724.1 ADP-ribosylglycohydrolase family protein
209(1) WP_019392119.1 WP_019392119.1 ADP-ribosylglycohydrolase family protein
209(1) WP_061805370.1 WP_061805370.1 ADP-ribosylglycohydrolase family protein
209(1) WP_225985160.1 WP_225985160.1 ADP-ribosylglycohydrolase family protein
209(1) WP_101953557.1 WP_101953557.1 ADP-ribosylglycohydrolase family protein
209(1) WP_243518763.1 WP_243518763.1 ADP-ribosylglycohydrolase family protein
209(2) WP_046217704.1 WP_046217704.1 ADP-ribosylglycohydrolase family protein
209(1) WP_113753098.1 WP_113753098.1 ADP-ribosylglycohydrolase family protein
209(3) WP_185375605.1 WP_185375605.1 ADP-ribosylglycohydrolase family protein
209(1) WP_124049167.1 WP_124049167.1 ADP-ribosylglycohydrolase family protein
209(1) WP_038245955.1 WP_038245955.1 ADP-ribosylglycohydrolase family protein
209(1) WP_272541590.1 WP_272541590.1 ADP-ribosylglycohydrolase family protein
209(1) WP_251456522.1 WP_251456522.1 ADP-ribosylglycohydrolase family protein
209(2) WP_111143015.1 WP_111143015.1 ADP-ribosylglycohydrolase family protein

210(2) WP_087433618.1 WP_087433618.1 hypothetical protein
210(1) WP_268629204.1 WP_268629204.1 hypothetical protein
210(2) WP_021261872.1 WP_021261872.1 hypothetical protein
210(2) WP_028533173.1 WP_028533173.1 hypothetical protein
210(1) WP_142543120.1 WP_142543120.1 hypothetical protein
210(1) WP_065291239.1 WP_065291239.1 hypothetical protein
210(1) WP_206094025.1 WP_206094025.1 hypothetical protein
210(1) WP_289355897.1 WP_289355897.1 hypothetical protein
210(3) WP_005546529.1 WP_005546529.1 hypothetical protein
210(1) WP_072727832.1 WP_072727832.1 hypothetical protein
210(1) WP_138186681.1 WP_138186681.1 hypothetical protein
210(1) WP_021253107.1 WP_021253107.1 hypothetical protein
210(3) WP_163977049.1 WP_163977049.1 hypothetical protein

211(1) WP_168178795.1 WP_168178795.1 GNAT family N-acetyltransferase
211(1) WP_213468107.1 WP_213468107.1 GNAT family N-acetyltransferase
211(1) WP_281477397.1 WP_281477397.1 GNAT family N-acetyltransferase
211(1) WP_028533190.1 WP_028533190.1 GNAT family N-acetyltransferase
211(1) WP_119794353.1 WP_119794353.1 GNAT family N-acetyltransferase
211(1) WP_293654346.1 WP_293654346.1 GNAT family N-acetyltransferase
211(1) WP_272045595.1 WP_272045595.1 GNAT family N-acetyltransferase
211(1) WP_083189148.1 WP_083189148.1 GNAT family N-acetyltransferase
211(1) WP_213426999.1 WP_213426999.1 GNAT family N-acetyltransferase
211(1) WP_087442399.1 WP_087442399.1 GNAT family N-acetyltransferase
211(1) WP_006674874.1 WP_006674874.1 GNAT family N-acetyltransferase
211(3) WP_163977069.1 WP_163977069.1 GNAT family N-acetyltransferase
211(1) WP_272827194.1 WP_272827194.1 GNAT family N-acetyltransferase
211(1) WP_197256882.1 WP_197256882.1 GNAT family N-acetyltransferase
211(1) WP_244918731.1 WP_244918731.1 hypothetical protein
211(1) WP_240927212.1 WP_240927212.1 GNAT family N-acetyltransferase
211(1) WP_133378780.1 WP_133378780.1 GNAT family N-acetyltransferase
211(1) WP_005546476.1 WP_005546476.1 GNAT family N-acetyltransferase

212(1) WP_062239846.1 WP_062239846.1 DUF5381 family protein
212(1) WP_066419969.1 WP_066419969.1 DUF5381 family protein
212(1) WP_066419963.1 WP_066419963.1 DUF5381 family protein
212(1) WP_270065065.1 WP_270065065.1 DUF5381 family protein
212(1) WP_057897630.1 WP_057897630.1 DUF5381 family protein
212(1) WP_097899494.1 WP_097899494.1 DUF5381 family protein
212(1) WP_090239236.1 WP_090239236.1 DUF5381 family protein
212(2) WP_204412729.1 WP_204412729.1 DUF5381 family protein
212(1) WP_116556142.1 WP_116556142.1 DUF5381 family protein
212(1) WP_283892181.1 WP_283892181.1 DUF5381 family protein
212(1) WP_301167503.1 WP_301167503.1 DUF5381 family protein
212(1) WP_183600892.1 WP_183600892.1 DUF5381 family protein
212(1) WP_290402001.1 WP_290402001.1 DUF5381 family protein
212(1) WP_066419965.1 WP_066419965.1 DUF5381 family protein
212(1) WP_265408838.1 WP_265408838.1 DUF5381 family protein
212(1) WP_091260860.1 WP_091260860.1 DUF5381 family protein
212(2) WP_248251190.1 WP_248251190.1 DUF5381 family protein
212(1) WP_289475134.1 WP_289475134.1 DUF5381 family protein

213(2) WP_087433771.1 WP_087433771.1 GNAT family N-acetyltransferase
213(3) WP_021253106.1 WP_021253106.1 GNAT family N-acetyltransferase
213(1) WP_142543119.1 WP_142543119.1 GNAT family N-acetyltransferase
213(1) WP_021261871.1 WP_021261871.1 GNAT family N-acetyltransferase
213(3) WP_051135124.1 WP_051135124.1 GNAT family N-acetyltransferase
213(1) WP_163977048.1 WP_163977048.1 GNAT family N-acetyltransferase
213(1) WP_305022928.1 WP_305022928.1 GNAT family N-acetyltransferase
213(1) WP_171414508.1 WP_171414508.1 GNAT family N-acetyltransferase
213(2) WP_065291238.1 WP_065291238.1 GNAT family N-acetyltransferase
213(1) WP_289355896.1 WP_289355896.1 GNAT family N-acetyltransferase
213(1) WP_072727833.1 WP_072727833.1 GNAT family N-acetyltransferase

| | | |
|---|---|---|
| 213(2) | WP_241752099.1 | WP_241752099.1 GNAT family N-acetyltransferase |
| 213(1) | WP_138186679.1 | WP_138186679.1 GNAT family N-acetyltransferase |
| 214(1) | WP_202283186.1 | WP_202283186.1 hypothetical protein |
| 214(1) | WP_000025257.1 | WP_000025257.1 hypothetical protein |
| 214(9) | WP_000025254.1 | WP_000025254.1 hypothetical protein |
| 214(2) | WP_000025256.1 | WP_000025256.1 hypothetical protein |
| 214(1) | WP_002075730.1 | WP_002075730.1 hypothetical protein |
| 214(1) | WP_206704966.1 | WP_206704966.1 hypothetical protein |
| 214(1) | WP_241752511.1 | WP_241752511.1 hypothetical protein |
| 214(1) | WP_098365941.1 | WP_098365941.1 hypothetical protein |
| 214(1) | WP_307199970.1 | WP_307199970.1 hypothetical protein |
| 214(1) | WP_208752238.1 | WP_208752238.1 DUF6572 domain-containing protein |
| 214(1) | WP_016129279.1 | WP_016129279.1 hypothetical protein |
| 215(1) | WP_076104566.1 | WP_076104566.1 bifunctional UDP-sugar hydrolase/5'-nucleotidase |
| 215(1) | WP_076303948.1 | WP_076303948.1 bifunctional UDP-sugar hydrolase/5'-nucleotidase |
| 215(1) | WP_076144378.1 | WP_076144378.1 bifunctional UDP-sugar hydrolase/5'-nucleotidase |
| 215(1) | WP_279664545.1 | WP_279664545.1 bifunctional 2',3'-cyclic-nucleotide 2'-phosphodiesterase/3'-nucleotidase |
| 215(1) | WP_155613670.1 | WP_155613670.1 hypothetical protein |
| 215(1) | WP_229522887.1 | WP_229522887.1 hypothetical protein |
| 215(1) | WP_279830783.1 | WP_279830783.1 hypothetical protein |
| 215(1) | WP_076309109.1 | WP_076309109.1 bifunctional UDP-sugar hydrolase/5'-nucleotidase |
| 215(1) | WP_229522851.1 | WP_229522851.1 LamG domain-containing protein |
| 215(1) | WP_223264157.1 | WP_223264157.1 family 43 glycosylhydrolase |
| 215(1) | WP_076279388.1 | WP_076279388.1 bifunctional UDP-sugar hydrolase/5'-nucleotidase |
| 215(1) | WP_036684085.1 | WP_036684085.1 bifunctional UDP-sugar hydrolase/5'-nucleotidase |
| 215(1) | WP_258208937.1 | WP_258208937.1 5'-nucleotidase C-terminal domain-containing protein |
| 215(1) | WP_307349986.1 | WP_307349986.1 hypothetical protein |
| 215(1) | WP_274485158.1 | WP_274485158.1 hypothetical protein |
| 215(1) | WP_196245375.1 | WP_196245375.1 hypothetical protein |
| 215(1) | WP_283655704.1 | WP_283655704.1 family 43 glycosylhydrolase |
| 215(1) | WP_010346273.1 | WP_010346273.1 family 43 glycosylhydrolase |
| 215(1) | WP_039871638.1 | WP_039871638.1 bifunctional UDP-sugar hydrolase/5'-nucleotidase |
| 215(1) | WP_308082230.1 | WP_308082230.1 5'-nucleotidase C-terminal domain-containing protein |
| 216(1) | WP_253440810.1 | WP_253440810.1 hypothetical protein |
| 216(1) | WP_110899379.1 | WP_110899379.1 hypothetical protein |
| 216(1) | WP_248063421.1 | WP_248063421.1 hypothetical protein |
| 216(1) | WP_145153679.1 | WP_145153679.1 hypothetical protein |
| 216(1) | WP_017692337.1 | WP_017692337.1 hypothetical protein |
| 216(1) | WP_211082919.1 | WP_211082919.1 hypothetical protein |
| 216(1) | WP_024632697.1 | WP_024632697.1 hypothetical protein |
| 216(1) | WP_068696920.1 | WP_068696920.1 hypothetical protein |
| 216(1) | WP_184183044.1 | WP_184183044.1 hypothetical protein |
| 216(1) | WP_068782424.1 | WP_068782424.1 hypothetical protein |
| 216(1) | WP_307199931.1 | WP_307199931.1 hypothetical protein |
| 216(2) | WP_216518273.1 | WP_216518273.1 hypothetical protein |
| 216(1) | WP_111273180.1 | WP_111273180.1 hypothetical protein |
| 216(1) | WP_076210952.1 | WP_076210952.1 hypothetical protein |
| 216(1) | WP_308723532.1 | WP_308723532.1 hypothetical protein |
| 216(1) | WP_068536348.1 | WP_068536348.1 hypothetical protein |
| 216(1) | WP_114298753.1 | WP_114298753.1 hypothetical protein |
| 216(1) | WP_150364186.1 | WP_150364186.1 hypothetical protein |
| 216(1) | WP_210087123.1 | WP_210087123.1 hypothetical protein |
| 217(1) | WP_293654344.1 | WP_293654344.1 GNAT family N-acetyltransferase |
| 217(1) | WP_310735670.1 | WP_310735670.1 GNAT family N-acetyltransferase |
| 217(2) | WP_143801069.1 | WP_143801069.1 GNAT family N-acetyltransferase |
| 217(1) | WP_272045589.1 | WP_272045589.1 hypothetical protein |
| 217(1) | WP_197256877.1 | WP_197256877.1 class I SAM-dependent methyltransferase |
| 217(1) | WP_168178793.1 | WP_168178793.1 GNAT family N-acetyltransferase |
| 217(1) | WP_272827195.1 | WP_272827195.1 class I SAM-dependent methyltransferase |
| 217(1) | WP_181454506.1 | WP_181454506.1 class I SAM-dependent methyltransferase |
| 217(1) | WP_213468109.1 | WP_213468109.1 GNAT family N-acetyltransferase |
| 217(1) | WP_165643411.1 | WP_165643411.1 class I SAM-dependent methyltransferase |
| 217(1) | WP_272045585.1 | WP_272045585.1 class I SAM-dependent methyltransferase |
| 217(3) | WP_168178792.1 | WP_168178792.1 class I SAM-dependent methyltransferase |
| 217(1) | WP_213426996.1 | WP_213426996.1 class I SAM-dependent methyltransferase |
| 217(1) | WP_164776261.1 | WP_164776261.1 class I SAM-dependent methyltransferase |
| 217(3) | WP_006674870.1 | WP_006674870.1 class I SAM-dependent methyltransferase |
| 218(1) | WP_128631301.1 | WP_128631301.1 PTS fructose transporter subunit IIABC |
| 218(1) | WP_303942508.1 | WP_303942508.1 PTS sugar transporter subunit IIA |
| 218(1) | WP_251596646.1 | WP_251596646.1 PTS fructose transporter subunit IIABC |
| 218(1) | WP_102372765.1 | WP_102372765.1 PTS sugar transporter subunit IIA |
| 218(1) | WP_128631302.1 | WP_128631302.1 BglG family transcription antiterminator |
| 218(5) | WP_037993174.1 | WP_037993174.1 BglG family transcription antiterminator |
| 218(1) | WP_273191237.1 | WP_273191237.1 PTS sugar transporter subunit IIA |
| 218(1) | WP_289342244.1 | WP_289342244.1 PTS sugar transporter subunit IIA |
| 218(1) | WP_195953736.1 | WP_195953736.1 PTS sugar transporter subunit IIA |
| 218(1) | WP_311284402.1 | WP_311284402.1 BglG family transcription antiterminator |
| 218(1) | WP_033100026.1 | WP_033100026.1 fructose-specific PTS transporter subunit EIIC |
| 218(1) | WP_048826598.1 | WP_048826598.1 BglG family transcription antiterminator |
| 218(1) | WP_273383856.1 | WP_273383856.1 PTS sugar transporter subunit IIA |
| 218(1) | WP_181731443.1 | WP_181731443.1 BglG family transcription antiterminator |
| 218(1) | WP_198050880.1 | WP_198050880.1 BglG family transcription antiterminator |
| 218(1) | WP_121874302.1 | WP_121874302.1 BglG family transcription antiterminator |
| 219(1) | WP_244723821.1 | WP_244723821.1 A24 family peptidase |
| 219(1) | WP_189031050.1 | WP_189031050.1 prepilin peptidase |
| 219(1) | WP_215185458.1 | WP_215185458.1 prepilin peptidase |
| 219(1) | WP_155609767.1 | WP_155609767.1 A24 family peptidase |
| 219(2) | WP_051494131.1 | WP_051494131.1 prepilin peptidase |
| 219(1) | WP_049869157.1 | WP_049869157.1 prepilin peptidase |
| 219(1) | WP_055109408.1 | WP_055109408.1 A24 family peptidase |
| 219(1) | WP_232274485.1 | WP_232274485.1 A24 family peptidase |
| 219(1) | WP_199925770.1 | WP_199925770.1 A24 family peptidase |
| 219(1) | WP_101300017.1 | WP_101300017.1 prepilin peptidase |
| 219(1) | WP_223068031.1 | WP_223068031.1 A24 family peptidase |
| 219(1) | WP_125084794.1 | WP_125084794.1 A24 family peptidase |
| 219(1) | WP_233530994.1 | WP_233530994.1 A24 family peptidase |
| 219(1) | WP_051569329.1 | WP_051569329.1 A24 family peptidase |

219(2) WP_213592887.1 WP_213592887.1 A24 family peptidase
219(1) WP_238655337.1 WP_238655337.1 prepilin peptidase
219(1) WP_105368168.1 WP_105368168.1 prepilin peptidase

220(8) WP_031307660.1 WP_031307660.1 sporulation transcriptional regulator SpoIIID
220(1) WP_068673752.1 WP_068673752.1 sporulation transcriptional regulator SpoIIID
220(1) WP_246860919.1 WP_246860919.1 sporulation transcriptional regulator SpoIIID
220(4) WP_026572748.1 WP_026572748.1 sporulation transcriptional regulator SpoIIID
220(1) WP_079506386.1 WP_079506386.1 sporulation transcriptional regulator SpoIIID
220(2) WP_046522783.1 WP_046522783.1 sporulation transcriptional regulator SpoIIID
220(2) WP_121610589.1 WP_121610589.1 sporulation transcriptional regulator SpoIIID

221(1) WP_138186705.1 WP_138186705.1 S-methyl-5-thioribose kinase
221(2) WP_028533186.1 WP_028533186.1 S-methyl-5-thioribose kinase
221(4) WP_005546489.1 WP_005546489.1 S-methyl-5-thioribose kinase
221(3) WP_163977065.1 WP_163977065.1 S-methyl-5-thioribose kinase
221(1) WP_142543136.1 WP_142543136.1 S-methyl-5-thioribose kinase
221(1) WP_171414529.1 WP_171414529.1 S-methyl-5-thioribose kinase
221(1) WP_268623238.1 WP_268623238.1 S-methyl-5-thioribose kinase
221(3) WP_021261894.1 WP_021261894.1 S-methyl-5-thioribose kinase
221(1) WP_021253129.1 WP_021253129.1 S-methyl-5-thioribose kinase
221(1) WP_072727816.1 WP_072727816.1 S-methyl-5-thioribose kinase
221(1) WP_065291257.1 WP_065291257.1 S-methyl-5-thioribose kinase

222(1) WP_280788640.1 WP_280788640.1 MATE family efflux transporter
222(1) WP_243518734.1 WP_243518734.1 MATE family efflux transporter
222(2) WP_019392101.1 WP_019392101.1 MATE family efflux transporter
222(1) WP_111925412.1 WP_111925412.1 MATE family efflux transporter
222(1) WP_197234733.1 WP_197234733.1 MATE family efflux transporter
222(1) WP_113713137.1 WP_113713137.1 MATE family efflux transporter
222(1) WP_113753071.1 WP_113753071.1 MATE family efflux transporter
222(1) WP_094870388.1 WP_094870388.1 MATE family efflux transporter
222(1) WP_227219161.1 WP_227219161.1 MATE family efflux transporter
222(1) WP_166244048.1 WP_166244048.1 MATE family efflux transporter
222(1) WP_251456545.1 WP_251456545.1 MATE family efflux transporter
222(1) WP_151585748.1 WP_151585748.1 MATE family efflux transporter
222(1) WP_216700669.1 WP_216700669.1 MATE family efflux transporter
222(2) WP_046217711.1 WP_046217711.1 MATE family efflux transporter
222(1) WP_268583872.1 WP_268583872.1 MATE family efflux transporter
222(1) WP_124049182.1 WP_124049182.1 MATE family efflux transporter
222(1) WP_143148621.1 WP_143148621.1 MATE family efflux transporter

223(1) WP_072336469.1 WP_072336469.1 DUF1992 domain-containing protein
223(1) WP_259392006.1 WP_259392006.1 hypothetical protein
223(1) WP_259392007.1 WP_259392007.1 DUF1992 domain-containing protein
223(1) WP_087442407.1 WP_087442407.1 DUF1992 domain-containing protein
223(1) WP_281477411.1 WP_281477411.1 DUF1992 domain-containing protein
223(1) WP_168178788.1 WP_168178788.1 hypothetical protein
223(1) WP_272045578.1 WP_272045578.1 DUF1992 domain-containing protein
223(1) WP_133378774.1 WP_133378774.1 DUF1992 domain-containing protein
223(2) WP_249723319.1 WP_249723319.1 hypothetical protein
223(1) WP_206765610.1 WP_206765610.1 DUF1992 domain-containing protein
223(1) WP_272827198.1 WP_272827198.1 DUF1992 domain-containing protein
223(1) WP_006674866.1 WP_006674866.1 DUF1992 domain-containing protein
223(1) WP_111153303.1 WP_111153303.1 DUF1992 domain-containing protein
223(1) WP_013914184.1 WP_013914184.1 DUF1992 domain-containing protein
223(2) WP_014368237.1 WP_014368237.1 DUF1992 domain-containing protein
223(1) WP_213426994.1 WP_213426994.1 DnaJ family domain-containing protein
223(1) WP_143801064.1 WP_143801064.1 DUF1992 domain-containing protein

224(1) WP_251480984.1 WP_251480984.1 DUF294 nucleotidyltransferase-like domain-containing protein
224(1) WP_132008145.1 WP_132008145.1 DUF294 nucleotidyltransferase-like domain-containing protein
224(1) WP_214902951.1 WP_214902951.1 DUF294 nucleotidyltransferase-like domain-containing protein
224(1) WP_226602946.1 WP_226602946.1 DUF294 nucleotidyltransferase-like domain-containing protein
224(1) WP_102264557.1 WP_102264557.1 DUF294 nucleotidyltransferase-like domain-containing protein
224(1) WP_226679110.1 WP_226679110.1 DUF294 nucleotidyltransferase-like domain-containing protein
224(1) WP_226086180.1 WP_226086180.1 DUF294 nucleotidyltransferase-like domain-containing protein
224(1) WP_121610587.1 WP_121610587.1 DUF294 nucleotidyltransferase-like domain-containing protein
224(1) WP_210364445.1 WP_210364445.1 DUF294 nucleotidyltransferase-like domain-containing protein
224(1) WP_260983631.1 WP_260983631.1 DUF294 nucleotidyltransferase-like domain-containing protein
224(1) WP_251542599.1 WP_251542599.1 DUF294 nucleotidyltransferase-like domain-containing protein
224(1) WP_041964583.1 WP_041964583.1 DUF294 nucleotidyltransferase-like domain-containing protein
224(1) WP_226642862.1 WP_226642862.1 DUF294 nucleotidyltransferase-like domain-containing protein
224(1) WP_215032137.1 WP_215032137.1 DUF294 nucleotidyltransferase-like domain-containing protein
224(1) WP_053361032.1 WP_053361032.1 DUF294 nucleotidyltransferase-like domain-containing protein
224(1) WP_215038335.1 WP_215038335.1 DUF294 nucleotidyltransferase-like domain-containing protein
224(1) WP_046522778.1 WP_046522778.1 DUF294 nucleotidyltransferase-like domain-containing protein
224(1) WP_283904349.1 WP_283904349.1 DUF294 nucleotidyltransferase-like domain-containing protein
224(1) WP_079506376.1 WP_079506376.1 DUF294 nucleotidyltransferase-like domain-containing protein

225(1) WP_102264558.1 WP_102264558.1 cation acetate symporter
225(1) WP_053361033.1 WP_053361033.1 sodium:solute symporter family protein
225(1) WP_226086181.1 WP_226086181.1 sodium:solute symporter family protein
225(1) WP_215038276.1 WP_215038276.1 sodium:solute symporter family protein
225(1) WP_079506374.1 WP_079506374.1 sodium:solute symporter family protein

225(1) WP_041964584.1 WP_041964584.1 sodium:solute symporter family protein
225(1) WP_226642860.1 WP_226642860.1 sodium:solute symporter family protein
225(2) WP_144479078.1 WP_144479078.1 sodium:solute symporter family protein
225(1) WP_226679111.1 WP_226679111.1 sodium:solute symporter family protein
225(1) WP_251480983.1 WP_251480983.1 sodium:solute symporter family protein
225(1) WP_214902954.1 WP_214902954.1 sodium:solute symporter family protein
225(1) WP_210364444.1 WP_210364444.1 sodium:solute symporter family protein
225(1) WP_283904350.1 WP_283904350.1 cation acetate symporter
225(1) WP_121610586.1 WP_121610586.1 sodium:solute symporter family protein
225(1) WP_132008142.1 WP_132008142.1 sodium:solute symporter family protein
225(1) WP_046522777.1 WP_046522777.1 cation acetate symporter
225(1) WP_226602957.1 WP_226602957.1 sodium:solute symporter family protein
225(1) WP_215032139.1 WP_215032139.1 sodium:solute symporter family protein

226(1) WP_171414527.1 WP_171414527.1 S-methyl-5-thioribose-1-phosphate isomerase
226(2) WP_241752092.1 WP_241752092.1 S-methyl-5-thioribose-1-phosphate isomerase
226(2) WP_044355256.1 WP_044355256.1 S-methyl-5-thioribose-1-phosphate isomerase
226(1) WP_163977064.1 WP_163977064.1 S-methyl-5-thioribose-1-phosphate isomerase
226(2) WP_268599736.1 WP_268599736.1 S-methyl-5-thioribose-1-phosphate isomerase
226(1) WP_268623239.1 WP_268623239.1 S-methyl-5-thioribose-1-phosphate isomerase
226(1) WP_138186704.1 WP_138186704.1 S-methyl-5-thioribose-1-phosphate isomerase
226(1) WP_051614155.1 WP_051614155.1 S-methyl-5-thioribose-1-phosphate isomerase
226(1) WP_021253128.1 WP_021253128.1 S-methyl-5-thioribose-1-phosphate isomerase
226(1) WP_065291256.1 WP_065291256.1 S-methyl-5-thioribose-1-phosphate isomerase
226(1) WP_072727817.1 WP_072727817.1 S-methyl-5-thioribose-1-phosphate isomerase
226(1) WP_142543135.1 WP_142543135.1 S-methyl-5-thioribose-1-phosphate isomerase
226(1) WP_005546492.1 WP_005546492.1 S-methyl-5-thioribose-1-phosphate isomerase
226(1) WP_021261893.1 WP_021261893.1 S-methyl-5-thioribose-1-phosphate isomerase
226(1) WP_289355908.1 WP_289355908.1 S-methyl-5-thioribose-1-phosphate isomerase
226(1) WP_090510231.1 WP_090510231.1 S-methyl-5-thioribose-1-phosphate isomerase

227(1) WP_281937385.1 WP_281937385.1 slipin family protein
227(1) WP_127484365.1 WP_127484365.1 slipin family protein
227(1) WP_090674379.1 WP_090674379.1 slipin family protein
227(1) WP_163853483.1 WP_163853483.1 slipin family protein
227(1) WP_036689388.1 WP_036689388.1 slipin family protein
227(1) WP_216791195.1 WP_216791195.1 slipin family protein
227(1) WP_010496600.1 WP_010496600.1 slipin family protein
227(1) WP_088830529.1 WP_088830529.1 slipin family protein
227(1) WP_254365997.1 WP_254365997.1 slipin family protein
227(1) WP_251598150.1 WP_251598150.1 slipin family protein
227(1) WP_302878981.1 WP_302878981.1 slipin family protein
227(1) WP_054972084.1 WP_054972084.1 slipin family protein
227(2) WP_057897617.1 WP_057897617.1 SPFH domain-containing protein
227(3) WP_063188080.1 WP_063188080.1 slipin family protein
227(1) WP_253411061.1 WP_253411061.1 slipin family protein
227(1) WP_025848588.1 WP_025848588.1 slipin family protein

228(1) WP_227917888.1 WP_227917888.1 hypothetical protein
228(1) WP_277166570.1 WP_277166570.1 hypothetical protein
228(1) WP_148138296.1 WP_148138296.1 hypothetical protein
228(1) WP_277157929.1 WP_277157929.1 hypothetical protein
228(1) WP_277156170.1 WP_277156170.1 hypothetical protein
228(1) WP_212330920.1 WP_212330920.1 hypothetical protein
228(1) WP_148362971.1 WP_148362971.1 hypothetical protein
228(1) WP_307738718.1 WP_307738718.1 hypothetical protein
228(1) WP_102372757.1 WP_102372757.1 hypothetical protein
228(1) WP_273191243.1 WP_273191243.1 hypothetical protein
228(1) WP_277142069.1 WP_277142069.1 hypothetical protein
228(1) WP_270472013.1 WP_270472013.1 hypothetical protein
228(1) WP_277175576.1 WP_277175576.1 hypothetical protein
228(1) WP_303942522.1 WP_303942522.1 hypothetical protein
228(1) WP_273383865.1 WP_273383865.1 hypothetical protein
228(1) WP_277156172.1 WP_277156172.1 hypothetical protein
228(1) WP_033521265.1 WP_033521265.1 hypothetical protein
228(1) WP_277157943.1 WP_277157943.1 hypothetical protein
228(1) WP_277157931.1 WP_277157931.1 hypothetical protein

229(8) WP_075702370.1 WP_075702370.1 hypothetical protein
229(4) WP_003745050.1 WP_003745050.1 hypothetical protein
229(2) WP_149051223.1 WP_149051223.1 hypothetical protein
229(2) WP_185639014.1 WP_185639014.1 hypothetical protein
229(2) WP_185331648.1 WP_185331648.1 hypothetical protein

230(1) WP_226642889.1 WP_226642889.1 rod shape-determining protein
230(1) WP_053361014.1 WP_053361014.1 rod shape-determining protein
230(1) WP_041964566.1 WP_041964566.1 rod shape-determining protein
230(1) WP_226679094.1 WP_226679094.1 rod shape-determining protein
230(1) WP_251542631.1 WP_251542631.1 rod shape-determining protein
230(2) WP_121610605.1 WP_121610605.1 rod shape-determining protein
230(1) WP_167831143.1 WP_167831143.1 rod shape-determining protein
230(1) WP_068673750.1 WP_068673750.1 rod shape-determining protein
230(1) WP_102264541.1 WP_102264541.1 rod shape-determining protein
230(1) WP_215032102.1 WP_215032102.1 rod shape-determining protein
230(1) WP_046522793.1 WP_046522793.1 rod shape-determining protein
230(1) WP_251481007.1 WP_251481007.1 rod shape-determining protein
230(1) WP_214706013.1 WP_214706013.1 rod shape-determining protein
230(4) WP_026572749.1 WP_026572749.1 rod shape-determining protein

231(3) WP_163977060.1 WP_163977060.1 VOC family protein
231(1) WP_138186701.1 WP_138186701.1 VOC family protein
231(2) WP_044355257.1 WP_044355257.1 glyoxalase/bleomycin resistance/dioxygenase family protein
231(1) WP_021261887.1 WP_021261887.1 glyoxalase/bleomycin resistance/dioxygenase family protein
231(2) WP_028533183.1 WP_028533183.1 glyoxalase/bleomycin resistance/dioxygenase family protein
231(1) WP_021253123.1 WP_021253123.1 glyoxalase/bleomycin resistance/dioxygenase family protein
231(3) WP_268599732.1 WP_268599732.1 VOC family protein
231(1) WP_065291251.1 WP_065291251.1 glyoxalase/bleomycin resistance/dioxygenase family protein
231(3) WP_072727822.1 WP_072727822.1 glyoxalase
231(1) WP_171414521.1 WP_171414521.1 VOC family protein

232(2)	WP_149051224.1	WP_149051224.1 hypothetical protein
232(10)	WP_046325929.1	WP_046325929.1 hypothetical protein
232(2)	WP_270994671.1	WP_270994671.1 hypothetical protein
232(2)	WP_185331649.1	WP_185331649.1 hypothetical protein
232(2)	WP_185639013.1	WP_185639013.1 hypothetical protein

233(2)	WP_149051222.1	WP_149051222.1 hypothetical protein
233(10)	WP_046325928.1	WP_046325928.1 hypothetical protein
233(2)	WP_185325794.1	WP_185325794.1 hypothetical protein
233(2)	WP_185639015.1	WP_185639015.1 hypothetical protein
233(2)	WP_270995167.1	WP_270995167.1 hypothetical protein

234(2)	WP_185336773.1	WP_185336773.1 hypothetical protein
234(2)	WP_200372596.1	WP_200372596.1 hypothetical protein
234(1)	WP_000435836.1	WP_000435836.1 SMI1/KNR4 family protein
234(11)	WP_000435837.1	WP_000435837.1 SMI1/KNR4 family protein
234(1)	WP_000435838.1	WP_000435838.1 SMI1/KNR4 family protein
234(1)	WP_281103267.1	WP_281103267.1 SMI1/KNR4 family protein

235(1)	WP_226086179.1	WP_226086179.1 3'-5' exonuclease
235(1)	WP_283904348.1	WP_283904348.1 3'-5' exonuclease
235(1)	WP_251480985.1	WP_251480985.1 exonuclease domain-containing protein
235(1)	WP_053361030.1	WP_053361030.1 3'-5' exonuclease
235(1)	WP_102264556.1	WP_102264556.1 exonuclease domain-containing protein
235(1)	WP_079506378.1	WP_079506378.1 3'-5' exonuclease
235(1)	WP_144479076.1	WP_144479076.1 3'-5' exonuclease
235(1)	WP_215032135.1	WP_215032135.1 exonuclease domain-containing protein
235(1)	WP_132008148.1	WP_132008148.1 exonuclease domain-containing protein
235(1)	WP_251542601.1	WP_251542601.1 3'-5' exonuclease
235(1)	WP_041964582.1	WP_041964582.1 exonuclease domain-containing protein
235(1)	WP_214705991.1	WP_214705991.1 3'-5' exonuclease
235(1)	WP_046522782.1	WP_046522782.1 3'-5' exonuclease
235(1)	WP_121610588.1	WP_121610588.1 exonuclease domain-containing protein
235(1)	WP_226642864.1	WP_226642864.1 3'-5' exonuclease
235(1)	WP_226679109.1	WP_226679109.1 exonuclease domain-containing protein
235(1)	WP_214902947.1	WP_214902947.1 3'-5' exonuclease
235(1)	WP_210364446.1	WP_210364446.1 3'-5' exonuclease

236(1)	WP_240550300.1	WP_240550300.1 RluA family pseudouridine synthase
236(1)	WP_283910912.1	WP_283910912.1 RluA family pseudouridine synthase
236(2)	WP_038569539.1	WP_038569539.1 RluA family pseudouridine synthase
236(1)	WP_006720300.1	WP_006720300.1 RluA family pseudouridine synthase
236(3)	WP_076206337.1	WP_076206337.1 RluA family pseudouridine synthase
236(1)	WP_076104001.1	WP_076104001.1 RluA family pseudouridine synthase
236(4)	WP_036684512.1	WP_036684512.1 RluA family pseudouridine synthase
236(1)	WP_076297596.1	WP_076297596.1 RluA family pseudouridine synthase
236(1)	WP_094876493.1	WP_094876493.1 RluA family pseudouridine synthase
236(2)	WP_094870548.1	WP_094870548.1 RluA family pseudouridine synthase
236(1)	WP_042125599.1	WP_042125599.1 RluA family pseudouridine synthase

237(1)	WP_181754681.1	WP_181754681.1 homoserine O-succinyltransferase
237(1)	WP_076281068.1	WP_076281068.1 homoserine O-succinyltransferase
237(6)	WP_036684477.1	WP_036684477.1 homoserine O-succinyltransferase
237(2)	WP_042125640.1	WP_042125640.1 homoserine O-succinyltransferase
237(2)	WP_076141381.1	WP_076141381.1 homoserine O-succinyltransferase
237(1)	WP_283910933.1	WP_283910933.1 homoserine O-succinyltransferase
237(2)	WP_076126410.1	WP_076126410.1 homoserine O-succinyltransferase
237(3)	WP_042186211.1	WP_042186211.1 homoserine O-succinyltransferase

238(1)	WP_216791199.1	WP_216791199.1 DIP1984 family protein
238(3)	WP_063188076.1	WP_063188076.1 DIP1984 family protein
238(1)	WP_163853481.1	WP_163853481.1 DIP1984 family protein
238(1)	WP_036689392.1	WP_036689392.1 DIP1984 family protein
238(1)	WP_127484361.1	WP_127484361.1 DIP1984 family protein
238(1)	WP_253411064.1	WP_253411064.1 DIP1984 family protein
238(1)	WP_054972085.1	WP_054972085.1 DIP1984 family protein
238(1)	WP_302878996.1	WP_302878996.1 DIP1984 family protein
238(1)	WP_025848589.1	WP_025848589.1 DIP1984 family protein
238(1)	WP_088830530.1	WP_088830530.1 DIP1984 family protein
238(1)	WP_010496601.1	WP_010496601.1 DIP1984 family protein
238(1)	WP_245719734.1	WP_245719734.1 hypothetical protein
238(1)	WP_254365998.1	WP_254365998.1 DIP1984 family protein
238(1)	WP_245719735.1	WP_245719735.1 DIP1984 family protein
238(1)	WP_281937383.1	WP_281937383.1 DIP1984 family protein
238(1)	WP_108534950.1	WP_108534950.1 DIP1984 family protein

239(1)	WP_042125642.1	WP_042125642.1 aminotransferase class I/II-fold pyridoxal phosphate-dependent enzyme
239(3)	WP_076103991.1	WP_076103991.1 PLP-dependent transferase
239(3)	WP_094870525.1	WP_094870525.1 aminotransferase class I/II-fold pyridoxal phosphate-dependent enzyme
239(1)	WP_283910934.1	WP_283910934.1 aminotransferase class I/II-fold pyridoxal phosphate-dependent enzyme
239(2)	WP_076141382.1	WP_076141382.1 PLP-dependent transferase
239(1)	WP_076119225.1	WP_076119225.1 aminotransferase class I/II-fold pyridoxal phosphate-dependent enzyme
239(1)	WP_036684476.1	WP_036684476.1 PLP-dependent transferase
239(2)	WP_076101579.1	WP_076101579.1 PLP-dependent transferase
239(1)	WP_307589589.1	WP_307589589.1 aminotransferase class I/II-fold pyridoxal phosphate-dependent enzyme
239(1)	WP_038569551.1	WP_038569551.1 PLP-dependent transferase
239(2)	WP_076301352.1	WP_076301352.1 PLP-dependent transferase

240(1)	WP_226086187.1	WP_226086187.1 VanZ family protein
240(1)	WP_226679114.1	WP_226679114.1 VanZ family protein
240(1)	WP_226642846.1	WP_226642846.1 VanZ family protein
240(1)	WP_102264564.1	WP_102264564.1 VanZ family protein
240(1)	WP_283904354.1	WP_283904354.1 VanZ family protein
240(1)	WP_041964590.1	WP_041964590.1 VanZ family protein
240(1)	WP_198310839.1	WP_198310839.1 VanZ family protein
240(2)	WP_133332972.1	WP_133332972.1 VanZ family protein
240(1)	WP_251466780.1	WP_251466780.1 VanZ family protein
240(1)	WP_215032159.1	WP_215032159.1 VanZ family protein

240(1) WP_144479090.1 WP_144479090.1 VanZ family protein
240(1) WP_308435601.1 WP_308435601.1 VanZ family protein
240(1) WP_081810637.1 WP_081810637.1 VanZ family protein
240(1) WP_282597661.1 WP_282597661.1 VanZ family protein
240(1) WP_079506362.1 WP_079506362.1 VanZ family protein
240(1) WP_251542587.1 WP_251542587.1 VanZ family protein
240(1) WP_214902972.1 WP_214902972.1 VanZ family protein

241(6) WP_038569550.1 WP_038569550.1 magnesium/cobalt transporter CorA
241(4) WP_036684479.1 WP_036684479.1 magnesium/cobalt transporter CorA
241(1) WP_076297588.1 WP_076297588.1 magnesium/cobalt transporter CorA
241(6) WP_042125638.1 WP_042125638.1 magnesium/cobalt transporter CorA

242(10) WP_036684474.1 WP_036684474.1 cyclic dehypoxanthinyl futalosine synthase
242(3) WP_042125644.1 WP_042125644.1 dehypoxanthine futalosine cyclase
242(1) WP_076184682.1 WP_076184682.1 cyclic dehypoxanthinyl futalosine synthase
242(1) WP_094876480.1 WP_094876480.1 cyclic dehypoxanthinyl futalosine synthase
242(2) WP_094870524.1 WP_094870524.1 cyclic dehypoxanthinyl futalosine synthase

243(1) WP_142950819.1 WP_142950819.1 oxalate:formate antiporter
243(7) WP_000014090.1 WP_000014090.1 oxalate:formate antiporter
243(1) WP_050062648.1 WP_050062648.1 oxalate:formate antiporter
243(8) WP_000014087.1 WP_000014087.1 oxalate:formate antiporter

244(2) WP_185639007.1 WP_185639007.1 DUF4064 domain-containing protein
244(2) WP_194334467.1 WP_194334467.1 DUF4064 domain-containing protein
244(2) WP_196318787.1 WP_196318787.1 DUF4064 domain-containing protein
244(6) WP_185337276.1 WP_185337276.1 DUF4064 domain-containing protein
244(3) WP_185432669.1 WP_185432669.1 DUF4064 domain-containing protein
244(1) WP_185335852.1 WP_185335852.1 DUF4064 domain-containing protein
244(1) WP_106496037.1 WP_106496037.1 hypothetical protein

245(1) WP_251542595.1 WP_251542595.1 hypothetical protein
245(1) WP_079506370.1 WP_079506370.1 hypothetical protein
245(1) WP_214902956.1 WP_214902956.1 hypothetical protein
245(2) WP_144479082.1 WP_144479082.1 hypothetical protein
245(1) WP_167831127.1 WP_167831127.1 hypothetical protein
245(1) WP_246860918.1 WP_246860918.1 hypothetical protein
245(1) WP_214705988.1 WP_214705988.1 hypothetical protein
245(2) WP_102264560.1 WP_102264560.1 hypothetical protein
245(1) WP_251480981.1 WP_251480981.1 hypothetical protein
245(1) WP_226602961.1 WP_226602961.1 hypothetical protein
245(1) WP_053361034.1 WP_053361034.1 hypothetical protein
245(1) WP_046522775.1 WP_046522775.1 hypothetical protein
245(1) WP_041964586.1 WP_041964586.1 hypothetical protein
245(1) WP_226642856.1 WP_226642856.1 hypothetical protein
245(1) WP_215032141.1 WP_215032141.1 hypothetical protein

246(1) WP_036689400.1 WP_036689400.1 TIGR02452 family protein
246(1) WP_216791208.1 WP_216791208.1 TIGR02452 family protein
246(1) WP_090674425.1 WP_090674425.1 TIGR02452 family protein
246(1) WP_281937377.1 WP_281937377.1 TIGR02452 family protein
246(1) WP_108534954.1 WP_108534954.1 TIGR02452 family protein
246(1) WP_054972090.1 WP_054972090.1 TIGR02452 family protein
246(1) WP_251598146.1 WP_251598146.1 TIGR02452 family protein
246(1) WP_127459423.1 WP_127459423.1 TIGR02452 family protein
246(1) WP_253411070.1 WP_253411070.1 TIGR02452 family protein
246(1) WP_163853477.1 WP_163853477.1 TIGR02452 family protein
246(1) WP_010496606.1 WP_010496606.1 TIGR02452 family protein
246(1) WP_025848603.1 WP_025848603.1 TIGR02452 family protein
246(1) WP_254366005.1 WP_254366005.1 TIGR02452 family protein
246(1) WP_063188068.1 WP_063188068.1 TIGR02452 family protein
246(1) WP_088830535.1 WP_088830535.1 TIGR02452 family protein
246(2) WP_127484353.1 WP_127484353.1 TIGR02452 family protein

247(1) WP_254366003.1 WP_254366003.1 class I SAM-dependent methyltransferase
247(1) WP_127459421.1 WP_127459421.1 class I SAM-dependent methyltransferase
247(1) WP_010496604.1 WP_010496604.1 class I SAM-dependent methyltransferase
247(1) WP_216791206.1 WP_216791206.1 class I SAM-dependent methyltransferase
247(1) WP_025848597.1 WP_025848597.1 class I SAM-dependent methyltransferase
247(1) WP_251598147.1 WP_251598147.1 class I SAM-dependent methyltransferase
247(1) WP_063188070.1 WP_063188070.1 class I SAM-dependent methyltransferase
247(1) WP_054972088.1 WP_054972088.1 class I SAM-dependent methyltransferase
247(1) WP_108534953.1 WP_108534953.1 class I SAM-dependent methyltransferase
247(2) WP_127484355.1 WP_127484355.1 class I SAM-dependent methyltransferase
247(1) WP_036689397.1 WP_036689397.1 class I SAM-dependent methyltransferase
247(1) WP_088830534.1 WP_088830534.1 class I SAM-dependent methyltransferase
247(1) WP_163853478.1 WP_163853478.1 methyltransferase domain-containing protein
247(1) WP_253411068.1 WP_253411068.1 class I SAM-dependent methyltransferase
247(1) WP_090674374.1 WP_090674374.1 class I SAM-dependent methyltransferase
247(1) WP_281937379.1 WP_281937379.1 class I SAM-dependent methyltransferase

248(1) WP_253411067.1 WP_253411067.1 metallophosphoesterase
248(1) WP_036689394.1 WP_036689394.1 metallophosphoesterase
248(2) WP_127484357.1 WP_127484357.1 metallophosphoesterase
248(1) WP_054972087.1 WP_054972087.1 metallophosphoesterase
248(1) WP_088830533.1 WP_088830533.1 metallophosphoesterase
248(1) WP_010496603.1 WP_010496603.1 metallophosphoesterase
248(1) WP_063188072.1 WP_063188072.1 metallophosphoesterase
248(1) WP_163853479.1 WP_163853479.1 metallophosphoesterase
248(1) WP_251598148.1 WP_251598148.1 metallophosphoesterase
248(1) WP_254366002.1 WP_254366002.1 metallophosphoesterase
248(1) WP_281937381.1 WP_281937381.1 metallophosphoesterase
248(1) WP_025848595.1 WP_025848595.1 metallophosphoesterase
248(1) WP_216791204.1 WP_216791204.1 metallophosphoesterase
248(1) WP_127459419.1 WP_127459419.1 metallophosphoesterase
248(1) WP_090674376.1 WP_090674376.1 metallophosphoesterase
248(1) WP_108534952.1 WP_108534952.1 metallophosphoesterase

249(3) WP_094870549.1 WP_094870549.1 cob(I)yrinic acid a,c-diamide adenosyltransferase
249(1) WP_076301356.1 WP_076301356.1 cob(I)yrinic acid a,c-diamide adenosyltransferase
249(1) WP_216083868.1 WP_216083868.1 cob(I)yrinic acid a,c-diamide adenosyltransferase
249(8) WP_036684514.1 WP_036684514.1 cob(I)yrinic acid a,c-diamide adenosyltransferase
249(1) WP_042125597.1 WP_042125597.1 cob(I)yrinic acid a,c-diamide adenosyltransferase

249(1) WP_076126402.1 WP_076126402.1 cob(I)yrinic acid a,c-diamide adenosyltransferase
249(1) WP_283910911.1 WP_283910911.1 cob(I)yrinic acid a,c-diamide adenosyltransferase
249(1) WP_177576801.1 WP_177576801.1 cob(I)yrinic acid a,c-diamide adenosyltransferase

250(5) WP_036684473.1 WP_036684473.1 SPFH domain-containing protein
250(1) WP_076281067.1 WP_076281067.1 SPFH domain-containing protein
250(3) WP_094870523.1 WP_094870523.1 SPFH domain-containing protein
250(2) WP_042125646.1 WP_042125646.1 SPFH domain-containing protein
250(1) WP_283910935.1 WP_283910935.1 SPFH domain-containing protein
250(5) WP_038569553.1 WP_038569553.1 SPFH domain-containing protein

251(1) WP_076104008.1 WP_076104008.1 HRDC domain-containing protein
251(1) WP_280787391.1 WP_280787391.1 HRDC domain-containing protein
251(5) WP_036684572.1 WP_036684572.1 HRDC domain-containing protein
251(1) WP_094876481.1 WP_094876481.1 HRDC domain-containing protein
251(1) WP_280752672.1 WP_280752672.1 HRDC domain-containing protein
251(1) WP_076281177.1 WP_076281177.1 HRDC domain-containing protein
251(1) WP_076301395.1 WP_076301395.1 HRDC domain-containing protein
251(1) WP_076297589.1 WP_076297589.1 HRDC domain-containing protein
251(1) WP_076119404.1 WP_076119404.1 HRDC domain-containing protein
251(1) WP_094870526.1 WP_094870526.1 HRDC domain-containing protein
251(1) WP_283912085.1 WP_283912085.1 HRDC domain-containing protein
251(1) WP_076271963.1 WP_076271963.1 HRDC domain-containing protein
251(1) WP_042131984.1 WP_042131984.1 HRDC domain-containing protein

252(1) WP_047909602.1 WP_047909602.1 TraR/DksA C4-type zinc finger protein
252(2) WP_138226015.1 WP_138226015.1 TraR/DksA C4-type zinc finger protein
252(1) WP_285748193.1 WP_285748193.1 TraR/DksA C4-type zinc finger protein
252(1) WP_191798540.1 WP_191798540.1 TraR/DksA C4-type zinc finger protein
252(1) WP_205053315.1 WP_205053315.1 TraR/DksA C4-type zinc finger protein
252(1) WP_169505064.1 WP_169505064.1 TraR/DksA C4-type zinc finger protein
252(1) WP_251583314.1 WP_251583314.1 TraR/DksA C4-type zinc finger protein
252(1) WP_155618098.1 WP_155618098.1 TraR/DksA C4-type zinc finger protein
252(1) WP_095266208.1 WP_095266208.1 TraR/DksA C4-type zinc finger protein
252(1) WP_090722889.1 WP_090722889.1 TraR/DksA C4-type zinc finger protein
252(1) WP_216513501.1 WP_216513501.1 TraR/DksA C4-type zinc finger protein
252(1) WP_091155405.1 WP_091155405.1 TraR/DksA C4-type zinc finger protein
252(1) WP_076311041.1 WP_076311041.1 TraR/DksA C4-type zinc finger protein
252(1) WP_054402897.1 WP_054402897.1 TraR/DksA C4-type zinc finger protein
252(1) WP_095398798.1 WP_095398798.1 TraR/DksA C4-type zinc finger protein
252(1) WP_213532669.1 WP_213532669.1 TraR/DksA C4-type zinc finger protein

253(1) WP_054972129.1 WP_054972129.1 NADAR family protein
253(1) WP_088830698.1 WP_088830698.1 NADAR family protein
253(1) WP_029191601.1 WP_029191601.1 NADAR family protein
253(1) WP_216791702.1 WP_216791702.1 NADAR family protein
253(1) WP_036689580.1 WP_036689580.1 NADAR family protein
253(1) WP_204343583.1 WP_204343583.1 NADAR family protein
253(1) WP_254366004.1 WP_254366004.1 NADAR family protein
253(1) WP_285854166.1 WP_285854166.1 NADAR family protein
253(1) WP_163860040.1 WP_163860040.1 NADAR family protein
253(1) WP_063188261.1 WP_063188261.1 NADAR family protein
253(1) WP_281937658.1 WP_281937658.1 NADAR family protein
253(3) WP_025848601.1 WP_025848601.1 NADAR family protein
253(1) WP_108535143.1 WP_108535143.1 NADAR family protein
253(1) WP_090674427.1 WP_090674427.1 NADAR family protein
253(1) WP_253411162.1 WP_253411162.1 NADAR family protein

254(1) WP_174809201.1 WP_174809201.1 N-acetyltransferase
254(1) WP_289355906.1 WP_289355906.1 GNAT family N-acetyltransferase
254(1) WP_142543131.1 WP_142543131.1 N-acetyltransferase
254(1) WP_065291252.1 WP_065291252.1 N-acetyltransferase
254(1) WP_072727821.1 WP_072727821.1 N-acetyltransferase
254(1) WP_163977061.1 WP_163977061.1 GNAT family N-acetyltransferase
254(2) WP_231586856.1 WP_231586856.1 GNAT family N-acetyltransferase
254(1) WP_268599733.1 WP_268599733.1 GNAT family N-acetyltransferase
254(1) WP_005546501.1 WP_005546501.1 hypothetical protein
254(2) WP_175460717.1 WP_175460717.1 hypothetical protein
254(2) WP_197267875.1 WP_197267875.1 GNAT family N-acetyltransferase
254(1) WP_268619932.1 WP_268619932.1 GNAT family N-acetyltransferase
254(1) WP_021261888.1 WP_021261888.1 N-acetyltransferase
254(1) WP_171414523.1 WP_171414523.1 GNAT family N-acetyltransferase

255(1) WP_152403403.1 WP_152403403.1 pirin family protein
255(1) WP_305022914.1 WP_305022914.1 pirin family protein
255(1) WP_251674664.1 WP_251674664.1 pirin family protein
255(1) WP_145949006.1 WP_145949006.1 pirin family protein
255(1) WP_094593885.1 WP_094593885.1 pirin family protein
255(1) WP_145021711.1 WP_145021711.1 pirin family protein
255(2) WP_089550140.1 WP_089550140.1 pirin family protein
255(1) WP_212984643.1 WP_212984643.1 pirin family protein
255(1) WP_116231167.1 WP_116231167.1 pirin family protein
255(1) WP_212969030.1 WP_212969030.1 pirin family protein
255(2) WP_145156815.1 WP_145156815.1 pirin family protein
255(1) WP_183578481.1 WP_183578481.1 pirin family protein
255(2) WP_145137723.1 WP_145137723.1 pirin family protein
255(1) WP_076169520.1 WP_076169520.1 pirin family protein

256(1) WP_245948214.1 WP_245948214.1 iron ABC transporter permease
256(1) WP_021624832.1 WP_021624832.1 iron chelate uptake ABC transporter family permease subunit
256(1) WP_066022465.1 WP_066022465.1 iron ABC transporter permease
256(1) WP_168976281.1 WP_168976281.1 iron chelate uptake ABC transporter family permease subunit
256(1) WP_268821819.1 WP_268821819.1 iron ABC transporter permease
256(1) WP_111147973.1 WP_111147973.1 iron ABC transporter permease
256(2) WP_043067151.1 WP_043067151.1 iron chelate uptake ABC transporter family permease subunit
256(1) WP_305755051.1 WP_305755051.1 iron ABC transporter permease
256(2) WP_040305097.1 WP_040305097.1 ABC transporter permease
256(1) WP_166244879.1 WP_166244879.1 iron ABC transporter permease
256(1) WP_235286186.1 WP_235286186.1 iron ABC transporter permease
256(1) WP_044037307.1 WP_044037307.1 iron ABC transporter permease
256(2) WP_043067150.1 WP_043067150.1 ABC transporter permease

256(1) WP_166244881.1 WP_166244881.1 iron ABC transporter permease

257(1) WP_220028092.1 WP_220028092.1 D-alanine--D-alanine ligase
257(1) WP_274485608.1 WP_274485608.1 D-alanine--D-alanine ligase
257(1) WP_165176687.1 WP_165176687.1 D-alanine--D-alanine ligase
257(2) WP_254348143.1 WP_254348143.1 D-alanine--D-alanine ligase
257(1) WP_071558836.1 WP_071558836.1 D-alanine--D-alanine ligase
257(1) WP_250245215.1 WP_250245215.1 D-alanine--D-alanine ligase
257(1) WP_010346275.1 WP_010346275.1 D-alanine--D-alanine ligase
257(1) WP_134902883.1 WP_134902883.1 D-alanine--D-alanine ligase
257(1) WP_053325283.1 WP_053325283.1 D-alanine--D-alanine ligase
257(1) WP_039273534.1 WP_039273534.1 D-alanine--D-alanine ligase
257(1) WP_307350418.1 WP_307350418.1 D-alanine--D-alanine ligase
257(1) WP_025685039.1 WP_025685039.1 D-alanine--D-alanine ligase
257(1) WP_013309996.1 WP_013309996.1 D-alanine--D-alanine ligase
257(1) WP_069081267.1 WP_069081267.1 D-alanine--D-alanine ligase
257(1) WP_103047824.1 WP_103047824.1 D-alanine--D-alanine ligase
257(1) WP_061829394.1 WP_061829394.1 D-alanine--D-alanine ligase

258(1) WP_263324537.1 WP_263324537.1 aldo/keto reductase
258(1) WP_124049162.1 WP_124049162.1 aldo/keto reductase
258(1) WP_243518769.1 WP_243518769.1 aldo/keto reductase
258(1) WP_076113459.1 WP_076113459.1 aldo/keto reductase
258(1) WP_186580039.1 WP_186580039.1 aldo/keto reductase
258(1) WP_077619667.1 WP_077619667.1 L-glyceraldehyde 3-phosphate reductase
258(1) WP_265408858.1 WP_265408858.1 aldo/keto reductase
258(1) WP_087194502.1 WP_087194502.1 aldo/keto reductase
258(1) WP_229139905.1 WP_229139905.1 L-glyceraldehyde 3-phosphate reductase
258(1) WP_019392123.1 WP_019392123.1 aldo/keto reductase
258(1) WP_268583849.1 WP_268583849.1 aldo/keto reductase
258(1) WP_150457752.1 WP_150457752.1 aldo/keto reductase
258(1) WP_010676878.1 WP_010676878.1 L-glyceraldehyde 3-phosphate reductase
258(1) WP_307589623.1 WP_307589623.1 aldo/keto reductase
258(2) WP_120046148.1 WP_120046148.1 aldo/keto reductase
258(1) WP_183600886.1 WP_183600886.1 aldo/keto reductase

259(1) WP_313957641.1 WP_313957641.1 DUF1961 family protein
259(1) WP_145948992.1 WP_145948992.1 DUF1961 family protein
259(1) WP_263880224.1 WP_263880224.1 YesU family protein
259(1) WP_094593863.1 WP_094593863.1 DUF1961 family protein
259(2) WP_084203763.1 WP_084203763.1 DUF1961 family protein
259(1) WP_116231185.1 WP_116231185.1 DUF1961 family protein
259(1) WP_251674643.1 WP_251674643.1 DUF1961 family protein
259(1) WP_145156798.1 WP_145156798.1 DUF1961 family protein
259(1) WP_145021759.1 WP_145021759.1 DUF1961 family protein
259(1) WP_163872748.1 WP_163872748.1 DUF1961 family protein
259(1) WP_183578415.1 WP_183578415.1 DUF1961 family protein
259(1) WP_212969041.1 WP_212969041.1 DUF1961 family protein
259(1) WP_152402361.1 WP_152402361.1 DUF1961 family protein
259(1) WP_200633292.1 WP_200633292.1 DUF1961 family protein
259(1) WP_212983252.1 WP_212983252.1 DUF1961 family protein
259(1) WP_313957640.1 WP_313957640.1 DUF1961 family protein

260(3) WP_108534967.1 WP_108534967.1 DUF4387 domain-containing protein
260(1) WP_063188037.1 WP_063188037.1 DUF4387 domain-containing protein
260(1) WP_090674352.1 WP_090674352.1 DUF4387 domain-containing protein
260(5) WP_054972102.1 WP_054972102.1 DUF4387 domain-containing protein
260(1) WP_025848630.1 WP_025848630.1 DUF4387 domain-containing protein
260(2) WP_127484335.1 WP_127484335.1 DUF4387 domain-containing protein
260(3) WP_010496621.1 WP_010496621.1 DUF4387 domain-containing protein

261(1) WP_154118042.1 WP_154118042.1 manganese catalase family protein
261(3) WP_076135817.1 WP_076135817.1 manganese catalase family protein
261(1) WP_036692823.1 WP_036692823.1 manganese catalase family protein
261(1) WP_076284176.1 WP_076284176.1 manganese catalase family protein
261(1) WP_039306375.1 WP_039306375.1 manganese catalase family protein
261(1) WP_019912174.1 WP_019912174.1 manganese catalase family protein
261(6) WP_036684067.1 WP_036684067.1 manganese catalase family protein
261(1) WP_042186391.1 WP_042186391.1 manganese catalase family protein
261(1) WP_039871648.1 WP_039871648.1 manganese catalase family protein

262(1) WP_021291842.1 WP_021291842.1 DUF1851 domain-containing protein
262(2) WP_185462002.1 WP_185462002.1 DUF1851 domain-containing protein
262(6) WP_029948785.1 WP_029948785.1 DUF1851 domain-containing protein
262(1) WP_098148067.1 WP_098148067.1 DUF1851 domain-containing protein
262(1) WP_078181472.1 WP_078181472.1 DUF1851 domain-containing protein
262(1) WP_002148296.1 WP_002148296.1 DUF1851 domain-containing protein
262(1) WP_050821999.1 WP_050821999.1 DUF1851 domain-containing protein
262(1) WP_000438858.1 WP_000438858.1 DUF1851 domain-containing protein
262(1) WP_002141604.1 WP_002141604.1 DUF1851 domain-containing protein
262(1) WP_097905174.1 WP_097905174.1 DUF1851 domain-containing protein

263(5) WP_042125648.1 WP_042125648.1 toxin-antitoxin system HicB family antitoxin
263(1) WP_248547674.1 WP_248547674.1 toxin-antitoxin system HicB family antitoxin
263(10) WP_036684471.1 WP_036684471.1 toxin-antitoxin system HicB family antitoxin

264(2) WP_046217701.1 WP_046217701.1 PTS sugar transporter subunit IIA
264(3) WP_077833125.1 WP_077833125.1 mannose/fructose/sorbose PTS transporter subunit IIA
264(2) WP_046217700.1 WP_046217700.1 PTS sugar transporter subunit IIB
264(1) WP_185383821.1 WP_185383821.1 mannose/fructose/sorbose PTS transporter subunit IIA
264(1) WP_197234710.1 WP_197234710.1 PTS sugar transporter subunit IIA
264(1) WP_251456516.1 WP_251456516.1 PTS sugar transporter subunit IIA
264(1) WP_197234708.1 WP_197234708.1 PTS sugar transporter subunit IIB
264(1) WP_251456514.1 WP_251456514.1 PTS sugar transporter subunit IIB
264(4) WP_077833124.1 WP_077833124.1 mannose/fructose/sorbose PTS transporter subunit IIB

265(1) WP_262725719.1 WP_262725719.1 SAM-dependent methyltransferase
265(2) WP_000493264.1 WP_000493264.1 tRNA (N6-threonylcarbamoyladenosine(37)-N6)-methyltransferase TrmO
265(2) WP_087982557.1 WP_087982557.1 SAM-dependent methyltransferase
265(2) WP_088063243.1 WP_088063243.1 SAM-dependent methyltransferase
265(1) WP_048552331.1 WP_048552331.1 tRNA (N6-threonylcarbamoyladenosine(37)-N6)-methyltransferase TrmO
265(1) WP_060630659.1 WP_060630659.1 SAM-dependent methyltransferase

265(1) WP_270728543.1 WP_270728543.1 SAM-dependent methyltransferase
265(3) WP_000493235.1 WP_000493235.1 tRNA (N6-threonylcarbamoyladenosine(37)-N6)-methyltransferase TrmO
265(3) WP_000493251.1 WP_000493251.1 SAM-dependent methyltransferase

266(1) WP_010496617.1 WP_010496617.1 ADP-ribosylglycohydrolase family protein
266(1) WP_088830543.1 WP_088830543.1 ADP-ribosylglycohydrolase family protein
266(1) WP_281937357.1 WP_281937357.1 ADP-ribosylglycohydrolase family protein
266(1) WP_054972098.1 WP_054972098.1 ADP-ribosylglycohydrolase family protein
266(2) WP_025848623.1 WP_025848623.1 ADP-ribosylglycohydrolase family protein
266(1) WP_090674358.1 WP_090674358.1 ADP-ribosylglycohydrolase family protein
266(1) WP_127484339.1 WP_127484339.1 ADP-ribosylglycohydrolase family protein
266(1) WP_251598139.1 WP_251598139.1 ADP-ribosylglycohydrolase family protein
266(1) WP_108534964.1 WP_108534964.1 ADP-ribosylglycohydrolase family protein
266(1) WP_063188043.1 WP_063188043.1 ADP-ribosylglycohydrolase family protein
266(1) WP_216791235.1 WP_216791235.1 ADP-ribosylglycohydrolase family protein
266(1) WP_163853467.1 WP_163853467.1 ADP-ribosylglycohydrolase family protein
266(1) WP_127459442.1 WP_127459442.1 ADP-ribosylglycohydrolase family protein
266(1) WP_036689450.1 WP_036689450.1 ADP-ribosylglycohydrolase family protein
266(1) WP_253411090.1 WP_253411090.1 ADP-ribosylglycohydrolase family protein

267(1) WP_283910913.1 WP_283910913.1 arsenate reductase family protein
267(1) WP_076297595.1 WP_076297595.1 arsenate reductase family protein
267(3) WP_076126405.1 WP_076126405.1 arsenate reductase family protein
267(3) WP_094870547.1 WP_094870547.1 arsenate reductase family protein
267(6) WP_038569540.1 WP_038569540.1 arsenate reductase family protein
267(1) WP_042125601.1 WP_042125601.1 arsenate reductase family protein
267(1) WP_036684509.1 WP_036684509.1 arsenate reductase family protein

268(2) WP_052097441.1 WP_052097441.1 5'-3' exonuclease H3TH domain-containing protein
268(2) WP_051491346.1 WP_051491346.1 5'-3' exonuclease H3TH domain-containing protein
268(2) WP_094870546.1 WP_094870546.1 5'-3' exonuclease H3TH domain-containing protein
268(1) WP_042125603.1 WP_042125603.1 5'-3' exonuclease
268(3) WP_076101643.1 WP_076101643.1 5'-3' exonuclease H3TH domain-containing protein
268(1) WP_283912084.1 WP_283912084.1 5'-3' exonuclease H3TH domain-containing protein
268(1) WP_076297616.1 WP_076297616.1 5'-3' exonuclease H3TH domain-containing protein
268(1) WP_280787399.1 WP_280787399.1 5'-3' exonuclease H3TH domain-containing protein
268(3) WP_076126491.1 WP_076126491.1 5'-3' exonuclease H3TH domain-containing protein

269(1) WP_281937350.1 WP_281937350.1 acyclic terpene utilization AtuA family protein
269(1) WP_302878972.1 WP_302878972.1 acyclic terpene utilization AtuA family protein
269(1) WP_025848631.1 WP_025848631.1 acyclic terpene utilization AtuA family protein
269(1) WP_036689467.1 WP_036689467.1 acyclic terpene utilization AtuA family protein
269(1) WP_253411098.1 WP_253411098.1 acyclic terpene utilization AtuA family protein
269(1) WP_163853466.1 WP_163853466.1 acyclic terpene utilization AtuA family protein
269(1) WP_216791241.1 WP_216791241.1 acyclic terpene utilization AtuA family protein
269(1) WP_010496622.1 WP_010496622.1 acyclic terpene utilization AtuA family protein
269(1) WP_108534968.1 WP_108534968.1 acyclic terpene utilization AtuA family protein
269(1) WP_090674351.1 WP_090674351.1 acyclic terpene utilization AtuA family protein
269(1) WP_251598136.1 WP_251598136.1 acyclic terpene utilization AtuA family protein
269(1) WP_088830546.1 WP_088830546.1 acyclic terpene utilization AtuA family protein
269(1) WP_127459447.1 WP_127459447.1 acyclic terpene utilization AtuA family protein
269(1) WP_127484333.1 WP_127484333.1 acyclic terpene utilization AtuA family protein
269(1) WP_054972103.1 WP_054972103.1 acyclic terpene utilization AtuA family protein
269(1) WP_063188035.1 WP_063188035.1 acyclic terpene utilization AtuA family protein

270(1) WP_174551274.1 WP_174551274.1 ADP-ribosylglycohydrolase family protein
270(1) WP_216791232.1 WP_216791232.1 ADP-ribosylglycohydrolase family protein
270(1) WP_036689434.1 WP_036689434.1 ADP-ribosylglycohydrolase family protein
270(1) WP_010496616.1 WP_010496616.1 ADP-ribosylglycohydrolase family protein
270(1) WP_189597462.1 WP_189597462.1 ADP-ribosylglycohydrolase family protein
270(1) WP_179223592.1 WP_179223592.1 ADP-ribosylglycohydrolase family protein
270(1) WP_253411088.1 WP_253411088.1 ADP-ribosylglycohydrolase family protein
270(1) WP_205510246.1 WP_205510246.1 ADP-ribosylglycohydrolase family protein
270(1) WP_025848622.1 WP_025848622.1 ADP-ribosylglycohydrolase family protein
270(1) WP_281937359.1 WP_281937359.1 ADP-ribosylglycohydrolase family protein
270(1) WP_174807108.1 WP_174807108.1 ADP-ribosylglycohydrolase family protein
270(1) WP_174806896.1 WP_174806896.1 ADP-ribosylglycohydrolase family protein
270(1) WP_237762783.1 WP_237762783.1 ADP-ribosylglycohydrolase family protein
270(1) WP_063188045.1 WP_063188045.1 ADP-ribosylglycohydrolase family protein
270(1) WP_251598140.1 WP_251598140.1 ADP-ribosylglycohydrolase family protein
270(1) WP_302878974.1 WP_302878974.1 ADP-ribosylglycohydrolase family protein

271(1) WP_061805352.1 WP_061805352.1 bile acid:sodium symporter family protein
271(1) WP_243518730.1 WP_243518730.1 bile acid:sodium symporter family protein
271(1) WP_113753069.1 WP_113753069.1 bile acid:sodium symporter family protein
271(1) WP_307312530.1 WP_307312530.1 bile acid:sodium symporter family protein
271(1) WP_151585751.1 WP_151585751.1 bile acid:sodium symporter family protein
271(1) WP_251456547.1 WP_251456547.1 bile acid:sodium symporter family protein
271(1) WP_179603488.1 WP_179603488.1 bile acid:sodium symporter family protein
271(1) WP_111925411.1 WP_111925411.1 bile acid:sodium symporter family protein
271(1) WP_113713136.1 WP_113713136.1 bile acid:sodium symporter family protein
271(1) WP_124049183.1 WP_124049183.1 bile acid:sodium symporter family protein
271(3) WP_046217712.1 WP_046217712.1 bile acid:sodium symporter family protein
271(2) WP_019392099.1 WP_019392099.1 bile acid:sodium symporter family protein
271(1) WP_268583874.1 WP_268583874.1 bile acid:sodium symporter family protein

272(1) WP_134750638.1 WP_134750638.1 hypothetical protein
272(1) WP_273321520.1 WP_273321520.1 hypothetical protein
272(1) WP_212695802.1 WP_212695802.1 hypothetical protein
272(1) WP_182301852.1 WP_182301852.1 hypothetical protein
272(1) WP_307600999.1 WP_307600999.1 hypothetical protein
272(1) WP_120112289.1 WP_120112289.1 hypothetical protein
272(1) WP_293755489.1 WP_293755489.1 hypothetical protein
272(1) WP_212693250.1 WP_212693250.1 hypothetical protein
272(1) WP_027088413.1 WP_027088413.1 hypothetical protein
272(1) WP_113672718.1 WP_113672718.1 hypothetical protein
272(2) WP_281813135.1 WP_281813135.1 hypothetical protein
272(1) WP_138755932.1 WP_138755932.1 hypothetical protein
272(1) WP_239617489.1 WP_239617489.1 hypothetical protein
272(1) WP_238187870.1 WP_238187870.1 hypothetical protein
272(1) WP_127570450.1 WP_127570450.1 hypothetical protein

273(1) WP_238187871.1 WP_238187871.1 hypothetical protein

| | | |
|---|---|---|
| 273(1) | WP_212695803.1 | WP_212695803.1 hypothetical protein |
| 273(1) | WP_212693251.1 | WP_212693251.1 hypothetical protein |
| 273(1) | WP_134750636.1 | WP_134750636.1 hypothetical protein |
| 273(1) | WP_182301851.1 | WP_182301851.1 hypothetical protein |
| 273(1) | WP_138755931.1 | WP_138755931.1 hypothetical protein |
| 273(1) | WP_307601001.1 | WP_307601001.1 hypothetical protein |
| 273(2) | WP_281813137.1 | WP_281813137.1 hypothetical protein |
| 273(1) | WP_239617488.1 | WP_239617488.1 hypothetical protein |
| 273(1) | WP_113672719.1 | WP_113672719.1 hypothetical protein |
| 273(1) | WP_293755487.1 | WP_293755487.1 hypothetical protein |
| 273(1) | WP_273321518.1 | WP_273321518.1 hypothetical protein |
| 273(1) | WP_120112286.1 | WP_120112286.1 hypothetical protein |
| 273(1) | WP_127570449.1 | WP_127570449.1 hypothetical protein |
| 273(1) | WP_027088414.1 | WP_027088414.1 hypothetical protein |
| 274(1) | WP_040056501.1 | WP_040056501.1 toxic anion resistance protein |
| 274(1) | WP_251456537.1 | WP_251456537.1 toxic anion resistance protein |
| 274(1) | WP_019392108.1 | WP_019392108.1 toxic anion resistance protein |
| 274(1) | WP_113753081.1 | WP_113753081.1 toxic anion resistance protein |
| 274(1) | WP_308110127.1 | WP_308110127.1 toxic anion resistance protein |
| 274(1) | WP_124049177.1 | WP_124049177.1 toxic anion resistance protein |
| 274(1) | WP_268583864.1 | WP_268583864.1 toxic anion resistance protein |
| 274(7) | WP_046217707.1 | WP_046217707.1 toxic anion resistance protein |
| 274(1) | WP_045518294.1 | WP_045518294.1 toxic anion resistance protein |
| 274(1) | WP_113713142.1 | WP_113713142.1 toxic anion resistance protein |
| 275(1) | WP_154118043.1 | WP_154118043.1 spore coat protein CotJB |
| 275(1) | WP_039871647.1 | WP_039871647.1 spore coat protein CotJB |
| 275(9) | WP_036684069.1 | WP_036684069.1 spore coat protein CotJB |
| 275(1) | WP_094870385.1 | WP_094870385.1 spore coat protein CotJB |
| 275(1) | WP_036721463.1 | WP_036721463.1 spore coat protein CotJB |
| 275(1) | WP_019912175.1 | WP_019912175.1 spore coat protein CotJB |
| 275(1) | WP_039306379.1 | WP_039306379.1 spore coat protein CotJB |
| 276(1) | WP_238536147.1 | WP_238536147.1 hypothetical protein |
| 276(1) | WP_142285898.1 | WP_142285898.1 hypothetical protein |
| 276(1) | WP_079994270.1 | WP_079994270.1 hypothetical protein |
| 276(1) | WP_087952379.1 | WP_087952379.1 hypothetical protein |
| 276(1) | WP_098768282.1 | WP_098768282.1 hypothetical protein |
| 276(1) | WP_254520527.1 | WP_254520527.1 hypothetical protein |
| 276(1) | WP_074565301.1 | WP_074565301.1 hypothetical protein |
| 276(2) | WP_098148070.1 | WP_098148070.1 hypothetical protein |
| 276(1) | WP_080448181.1 | WP_080448181.1 hypothetical protein |
| 276(1) | WP_141558154.1 | WP_141558154.1 hypothetical protein |
| 276(1) | WP_098266933.1 | WP_098266933.1 hypothetical protein |
| 276(1) | WP_254904971.1 | WP_254904971.1 hypothetical protein |
| 276(2) | WP_103626235.1 | WP_103626235.1 hypothetical protein |
| 277(1) | WP_183578412.1 | WP_183578412.1 hypothetical protein |
| 277(1) | WP_263880225.1 | WP_263880225.1 hypothetical protein |
| 277(1) | WP_251674641.1 | WP_251674641.1 hypothetical protein |
| 277(1) | WP_116231762.1 | WP_116231762.1 hypothetical protein |
| 277(1) | WP_089550136.1 | WP_089550136.1 hypothetical protein |
| 277(1) | WP_163872744.1 | WP_163872744.1 hypothetical protein |
| 277(1) | WP_200633291.1 | WP_200633291.1 hypothetical protein |
| 277(1) | WP_152402359.1 | WP_152402359.1 hypothetical protein |
| 277(1) | WP_094593862.1 | WP_094593862.1 hypothetical protein |
| 277(1) | WP_212969042.1 | WP_212969042.1 hypothetical protein |
| 277(1) | WP_145156797.1 | WP_145156797.1 hypothetical protein |
| 277(1) | WP_212983251.1 | WP_212983251.1 hypothetical protein |
| 277(1) | WP_145021762.1 | WP_145021762.1 hypothetical protein |
| 277(1) | WP_076169526.1 | WP_076169526.1 hypothetical protein |
| 277(1) | WP_145948991.1 | WP_145948991.1 hypothetical protein |
| 278(1) | WP_063219629.1 | WP_063219629.1 SMI1/KNR4 family protein |
| 278(4) | WP_000190066.1 | WP_000190066.1 SMI1/KNR4 family protein |
| 278(1) | WP_193653017.1 | WP_193653017.1 SMI1/KNR4 family protein |
| 278(3) | WP_064477386.1 | WP_064477386.1 SMI1/KNR4 family protein |
| 278(1) | WP_098579099.1 | WP_098579099.1 SMI1/KNR4 family protein |
| 278(1) | WP_243495564.1 | WP_243495564.1 SMI1/KNR4 family protein |
| 278(1) | WP_272801668.1 | WP_272801668.1 SMI1/KNR4 family protein |
| 278(2) | WP_185336771.1 | WP_185336771.1 SMI1/KNR4 family protein |
| 278(1) | WP_000190068.1 | WP_000190068.1 SMI1/KNR4 family protein |
| 279(3) | WP_168178790.1 | WP_168178790.1 BadF/BadG/BcrA/BcrD ATPase family protein |
| 279(1) | WP_006674868.1 | WP_006674868.1 BadF/BadG/BcrA/BcrD ATPase family protein |
| 279(1) | WP_259392004.1 | WP_259392004.1 hypothetical protein |
| 279(1) | WP_133378776.1 | WP_133378776.1 BadF/BadG/BcrA/BcrD ATPase family protein |
| 279(1) | WP_259392005.1 | WP_259392005.1 hypothetical protein |
| 279(1) | WP_197256873.1 | WP_197256873.1 BadF/BadG/BcrA/BcrD ATPase family protein |
| 279(1) | WP_281477406.1 | WP_281477406.1 BadF/BadG/BcrA/BcrD ATPase family protein |
| 279(1) | WP_249724668.1 | WP_249724668.1 BadF/BadG/BcrA/BcrD ATPase family protein |
| 279(1) | WP_272827196.1 | WP_272827196.1 BadF/BadG/BcrA/BcrD ATPase family protein |
| 279(1) | WP_087442405.1 | WP_087442405.1 ATPase |
| 279(1) | WP_111153305.1 | WP_111153305.1 BadF/BadG/BcrA/BcrD ATPase family protein |
| 279(1) | WP_272045582.1 | WP_272045582.1 BadF/BadG/BcrA/BcrD ATPase family protein |
| 279(1) | WP_143801066.1 | WP_143801066.1 BadF/BadG/BcrA/BcrD ATPase family protein |
| 280(1) | WP_021261870.1 | WP_021261870.1 acyl-CoA dehydrogenase |
| 280(1) | WP_072727834.1 | WP_072727834.1 acyl-CoA dehydrogenase |
| 280(1) | WP_171414507.1 | WP_171414507.1 acyl-CoA dehydrogenase |
| 280(2) | WP_005546531.1 | WP_005546531.1 acyl-CoA dehydrogenase |
| 280(1) | WP_289355895.1 | WP_289355895.1 acyl-CoA dehydrogenase |
| 280(1) | WP_268619941.1 | WP_268619941.1 acyl-CoA dehydrogenase |
| 280(2) | WP_197267869.1 | WP_197267869.1 acyl-CoA dehydrogenase |
| 280(1) | WP_268623246.1 | WP_268623246.1 acyl-CoA dehydrogenase |
| 280(1) | WP_138186677.1 | WP_138186677.1 acyl-CoA dehydrogenase |
| 280(1) | WP_163977047.1 | WP_163977047.1 acyl-CoA dehydrogenase |
| 280(1) | WP_028533172.1 | WP_028533172.1 acyl-CoA dehydrogenase |
| 280(1) | WP_065291237.1 | WP_065291237.1 acyl-CoA dehydrogenase |
| 280(1) | WP_142543118.1 | WP_142543118.1 acyl-CoA dehydrogenase |
| 281(1) | WP_119794372.1 | WP_119794372.1 hypothetical protein |
| 281(2) | WP_168178777.1 | WP_168178777.1 polysaccharide pyruvyl transferase family protein |

281(1) WP_133378763.1 WP_133378763.1 polysaccharide pyruvyl transferase family protein
281(1) WP_244194183.1 WP_244194183.1 polysaccharide pyruvyl transferase family protein
281(1) WP_272045558.1 WP_272045558.1 polysaccharide pyruvyl transferase family protein
281(1) WP_281477431.1 WP_281477431.1 polysaccharide pyruvyl transferase family protein
281(1) WP_197256846.1 WP_197256846.1 polysaccharide pyruvyl transferase family protein
281(1) WP_213426982.1 WP_213426982.1 polysaccharide pyruvyl transferase family protein
281(2) WP_240927211.1 WP_240927211.1 polysaccharide pyruvyl transferase family protein
281(1) WP_238535259.1 WP_238535259.1 polysaccharide pyruvyl transferase family protein
281(1) WP_293654339.1 WP_293654339.1 polysaccharide pyruvyl transferase family protein
281(1) WP_111153293.1 WP_111153293.1 polysaccharide pyruvyl transferase family protein
281(1) WP_119794373.1 WP_119794373.1 hypothetical protein

282(2) WP_281813133.1 WP_281813133.1 hypothetical protein
282(1) WP_027088412.1 WP_027088412.1 hypothetical protein
282(1) WP_127570451.1 WP_127570451.1 hypothetical protein
282(1) WP_239617490.1 WP_239617490.1 hypothetical protein
282(1) WP_138755933.1 WP_138755933.1 hypothetical protein
282(1) WP_182301853.1 WP_182301853.1 hypothetical protein
282(1) WP_238187869.1 WP_238187869.1 hypothetical protein
282(1) WP_273321522.1 WP_273321522.1 hypothetical protein
282(2) WP_113672717.1 WP_113672717.1 hypothetical protein
282(1) WP_120112291.1 WP_120112291.1 hypothetical protein
282(1) WP_293755491.1 WP_293755491.1 hypothetical protein
282(1) WP_307600997.1 WP_307600997.1 hypothetical protein
282(1) WP_212695801.1 WP_212695801.1 hypothetical protein

283(1) WP_066415465.1 WP_066415465.1 hypothetical protein
283(1) WP_251638753.1 WP_251638753.1 hydrogenase maturation nickel metallochaperone HypA
283(1) WP_289607503.1 WP_289607503.1 hypothetical protein
283(1) WP_254759307.1 WP_254759307.1 hypothetical protein
283(1) WP_227219207.1 WP_227219207.1 zinc ribbon domain-containing protein
283(1) WP_289342271.1 WP_289342271.1 hypothetical protein
283(1) WP_090761839.1 WP_090761839.1 hypothetical protein
283(1) WP_263324558.1 WP_263324558.1 hypothetical protein
283(1) WP_294953160.1 WP_294953160.1 hypothetical protein
283(3) WP_213117491.1 WP_213117491.1 hypothetical protein
283(1) WP_307284700.1 WP_307284700.1 hypothetical protein
283(1) WP_283952939.1 WP_283952939.1 hypothetical protein
283(1) WP_308082255.1 WP_308082255.1 hypothetical protein

284(1) WP_102264565.1 WP_102264565.1 nuclease-related domain-containing protein
284(1) WP_198310840.1 WP_198310840.1 hypothetical protein
284(1) WP_165976165.1 WP_165976165.1 NERD domain-containing protein
284(1) WP_226642844.1 WP_226642844.1 nuclease-related domain-containing protein
284(1) WP_041964591.1 WP_041964591.1 nuclease-related domain-containing protein
284(1) WP_229139930.1 WP_229139930.1 NERD domain-containing protein
284(1) WP_251542586.1 WP_251542586.1 nuclease-related domain-containing protein
284(1) WP_283904355.1 WP_283904355.1 nuclease-related domain-containing protein
284(1) WP_144479092.1 WP_144479092.1 nuclease-related domain-containing protein
284(1) WP_079506360.1 WP_079506360.1 nuclease-related domain-containing protein
284(1) WP_214902975.1 WP_214902975.1 nuclease-related domain-containing protein
284(1) WP_226679115.1 WP_226679115.1 nuclease-related domain-containing protein
284(1) WP_051590911.1 WP_051590911.1 nuclease-related domain-containing protein
284(1) WP_308912812.1 WP_308912812.1 NERD domain-containing protein
284(1) WP_210364439.1 WP_210364439.1 nuclease-related domain-containing protein

285(1) WP_154118044.1 WP_154118044.1 spore coat associated protein CotJA
285(1) WP_076281044.1 WP_076281044.1 spore coat associated protein CotJA
285(8) WP_036684070.1 WP_036684070.1 spore coat associated protein CotJA
285(1) WP_094870384.1 WP_094870384.1 spore coat associated protein CotJA
285(1) WP_039306384.1 WP_039306384.1 spore coat associated protein CotJA
285(1) WP_039871645.1 WP_039871645.1 spore coat associated protein CotJA
285(1) WP_036692819.1 WP_036692819.1 spore coat associated protein CotJA

286(9) WP_042401005.1 WP_042401005.1 peptide chain release factor 3
286(2) WP_005210434.1 WP_005210434.1 peptide chain release factor 3
286(1) WP_033521284.1 WP_033521284.1 translational GTPase TypA
286(1) WP_189018635.1 WP_189018635.1 GTP-binding protein
286(1) WP_087194496.1 WP_087194496.1 selenocysteine-specific translation elongation factor

287(1) WP_000428102.1 WP_000428102.1 DUF4274 domain-containing protein
287(1) WP_088032803.1 WP_088032803.1 DUF4274 domain-containing protein
287(1) WP_113733522.1 WP_113733522.1 DUF4274 domain-containing protein
287(1) WP_097987490.1 WP_097987490.1 DUF4274 domain-containing protein
287(2) WP_000428100.1 WP_000428100.1 DUF4274 domain-containing protein
287(1) WP_202283185.1 WP_202283185.1 DUF4274 domain-containing protein
287(2) WP_195431331.1 WP_195431331.1 DUF4274 domain-containing protein
287(3) WP_071736285.1 WP_071736285.1 DUF4274 domain-containing protein
287(1) WP_075666792.1 WP_075666792.1 DUF4274 domain-containing protein
287(1) WP_026593053.1 WP_026593053.1 DUF4274 domain-containing protein

288(2) WP_185326090.1 WP_185326090.1 Imm59 family immunity protein
288(4) WP_221633564.1 WP_221633564.1 Imm59 family immunity protein
288(8) WP_185336756.1 WP_185336756.1 Imm59 family immunity protein

289(2) WP_036684072.1 WP_036684072.1 DedA family protein
289(1) WP_247083061.1 WP_247083061.1 DedA family protein
289(1) WP_074108304.1 WP_074108304.1 DedA family protein
289(1) WP_076135821.1 WP_076135821.1 DedA family protein
289(1) WP_074085369.1 WP_074085369.1 DedA family protein
289(6) WP_076100017.1 WP_076100017.1 DedA family protein
289(2) WP_219872447.1 WP_219872447.1 VTT domain-containing protein

290(2) WP_121610585.1 WP_121610585.1 DUF4212 domain-containing protein
290(1) WP_226086182.1 WP_226086182.1 DUF4212 domain-containing protein
290(1) WP_053364460.1 WP_053364460.1 DUF4212 domain-containing protein
290(1) WP_251542597.1 WP_251542597.1 DUF4212 domain-containing protein
290(1) WP_210364443.1 WP_210364443.1 DUF4212 domain-containing protein
290(1) WP_226602959.1 WP_226602959.1 DUF4212 domain-containing protein
290(1) WP_144479080.1 WP_144479080.1 DUF4212 domain-containing protein
290(1) WP_079506372.1 WP_079506372.1 DUF4212 domain-containing protein
290(2) WP_102264559.1 WP_102264559.1 DUF4212 domain-containing protein
290(1) WP_226642858.1 WP_226642858.1 DUF4212 domain-containing protein
290(1) WP_041964585.1 WP_041964585.1 DUF4212 domain-containing protein

290(1) WP_046522776.1 WP_046522776.1 DUF4212 domain-containing protein

291(1) WP_069196265.1 WP_069196265.1 glycoside hydrolase family 9 protein
291(2) WP_003516552.1 WP_003516552.1 glycoside hydrolase family 9 protein
291(1) WP_128705705.1 WP_128705705.1 glycoside hydrolase family 9 protein
291(2) WP_003520969.1 WP_003520969.1 glycoside hydrolase family 9 protein
291(1) WP_038286701.1 WP_038286701.1 glycoside hydrolase family 9 protein
291(1) WP_038286699.1 WP_038286699.1 glycoside hydrolase family 9 protein
291(1) WP_265444525.1 WP_265444525.1 glycosyl hydrolase family 28-related protein
291(2) WP_003516585.1 WP_003516585.1 glycosyl hydrolase family 28-related protein
291(1) WP_126996337.1 WP_126996337.1 glycoside hydrolase family 9 protein
291(1) WP_069193869.1 WP_069193869.1 glycosyl hydrolase family 28-related protein
291(1) WP_251417689.1 WP_251417689.1 glycoside hydrolase family 9 protein

292(1) WP_251456533.1 WP_251456533.1 DUF3243 domain-containing protein
292(1) WP_111925418.1 WP_111925418.1 DUF3243 domain-containing protein
292(2) WP_061805360.1 WP_061805360.1 DUF3243 domain-containing protein
292(1) WP_243518746.1 WP_243518746.1 DUF3243 domain-containing protein
292(1) WP_113753083.1 WP_113753083.1 DUF3243 domain-containing protein
292(1) WP_124049175.1 WP_124049175.1 DUF3243 domain-containing protein
292(1) WP_268583863.1 WP_268583863.1 DUF3243 domain-containing protein
292(5) WP_019392110.1 WP_019392110.1 DUF3243 domain-containing protein
292(1) WP_197234729.1 WP_197234729.1 DUF3243 domain-containing protein

293(3) WP_061805358.1 WP_061805358.1 5-bromo-4-chloroindolyl phosphate hydrolysis family protein
293(1) WP_251456539.1 WP_251456539.1 5-bromo-4-chloroindolyl phosphate hydrolysis family protein
293(1) WP_124049178.1 WP_124049178.1 5-bromo-4-chloroindolyl phosphate hydrolysis family protein
293(1) WP_019392107.1 WP_019392107.1 5-bromo-4-chloroindolyl phosphate hydrolysis family protein
293(1) WP_268583865.1 WP_268583865.1 5-bromo-4-chloroindolyl phosphate hydrolysis family protein
293(4) WP_040056502.1 WP_040056502.1 5-bromo-4-chloroindolyl phosphate hydrolysis family protein
293(1) WP_197234731.1 WP_197234731.1 5-bromo-4-chloroindolyl phosphate hydrolysis family protein
293(1) WP_151585742.1 WP_151585742.1 5-bromo-4-chloroindolyl phosphate hydrolysis family protein
293(1) WP_243518744.1 WP_243518744.1 5-bromo-4-chloroindolyl phosphate hydrolysis family protein

294(3) WP_168178781.1 WP_168178781.1 hypothetical protein
294(1) WP_087442418.1 WP_087442418.1 hypothetical protein
294(1) WP_213426985.1 WP_213426985.1 hypothetical protein
294(2) WP_143801055.1 WP_143801055.1 hypothetical protein
294(1) WP_006674855.1 WP_006674855.1 hypothetical protein
294(1) WP_133378766.1 WP_133378766.1 hypothetical protein
294(2) WP_111153297.1 WP_111153297.1 hypothetical protein
294(1) WP_272045564.1 WP_272045564.1 hypothetical protein
294(1) WP_119794368.1 WP_119794368.1 hypothetical protein
294(1) WP_197256854.1 WP_197256854.1 hypothetical protein

295(1) WP_213427001.1 WP_213427001.1 proline dehydrogenase family protein
295(2) WP_133378783.1 WP_133378783.1 proline dehydrogenase family protein
295(1) WP_197256886.1 WP_197256886.1 proline dehydrogenase family protein
295(2) WP_006674877.1 WP_006674877.1 proline dehydrogenase family protein
295(2) WP_143801074.1 WP_143801074.1 proline dehydrogenase family protein
295(1) WP_119794350.1 WP_119794350.1 proline dehydrogenase family protein
295(1) WP_272045601.1 WP_272045601.1 proline dehydrogenase family protein
295(1) WP_087442395.1 WP_087442395.1 proline dehydrogenase family protein
295(3) WP_168178797.1 WP_168178797.1 proline dehydrogenase family protein

296(1) WP_243518765.1 WP_243518765.1 hypothetical protein
296(1) WP_120046147.1 WP_120046147.1 hypothetical protein
296(1) WP_019392120.1 WP_019392120.1 hypothetical protein
296(1) WP_151585721.1 WP_151585721.1 hypothetical protein
296(1) WP_272541585.1 WP_272541585.1 hypothetical protein
296(1) WP_142673091.1 WP_142673091.1 aspartate/glutamate racemase family protein
296(1) WP_251456520.1 WP_251456520.1 hypothetical protein
296(1) WP_085117890.1 WP_085117890.1 hypothetical protein
296(1) WP_048896846.1 WP_048896846.1 hypothetical protein
296(1) WP_113753108.1 WP_113753108.1 hypothetical protein
296(1) WP_268583853.1 WP_268583853.1 hypothetical protein
296(1) WP_124049165.1 WP_124049165.1 hypothetical protein
296(1) WP_113713152.1 WP_113713152.1 hypothetical protein
296(1) WP_197234714.1 WP_197234714.1 hypothetical protein

297(1) WP_268583869.1 WP_268583869.1 glycosyl hydrolase family 18 protein
297(1) WP_151585745.1 WP_151585745.1 glycosyl hydrolase family 18 protein
297(1) WP_272541594.1 WP_272541594.1 glycosyl hydrolase family 18 protein
297(1) WP_019392103.1 WP_019392103.1 glycosyl hydrolase family 18 protein
297(1) WP_046217709.1 WP_046217709.1 glycosyl hydrolase family 18 protein
297(1) WP_085117879.1 WP_085117879.1 glycosyl hydrolase family 18 protein
297(1) WP_243518740.1 WP_243518740.1 glycosyl hydrolase family 18 protein
297(1) WP_113753075.1 WP_113753075.1 glycosyl hydrolase family 18 protein
297(1) WP_124049180.1 WP_124049180.1 glycosyl hydrolase family 18 protein
297(1) WP_120046143.1 WP_120046143.1 glycosyl hydrolase family 18 protein
297(1) WP_061805355.1 WP_061805355.1 glycosyl hydrolase family 18 protein
297(1) WP_113713139.1 WP_113713139.1 glycosyl hydrolase family 18 protein
297(1) WP_111925414.1 WP_111925414.1 glycosyl hydrolase family 18 protein
297(1) WP_251456543.1 WP_251456543.1 glycosyl hydrolase family 18 protein

298(1) WP_277142071.1 WP_277142071.1 hypothetical protein
298(1) WP_277156171.1 WP_277156171.1 hypothetical protein
298(1) WP_277175578.1 WP_277175578.1 hypothetical protein
298(1) WP_277166569.1 WP_277166569.1 hypothetical protein
298(1) WP_277156182.1 WP_277156182.1 hypothetical protein
298(1) WP_277175575.1 WP_277175575.1 hypothetical protein
298(1) WP_277175584.1 WP_277175584.1 hypothetical protein
298(1) WP_277175573.1 WP_277175573.1 hypothetical protein
298(1) WP_277156174.1 WP_277156174.1 hypothetical protein
298(1) WP_277142086.1 WP_277142086.1 hypothetical protein

| | | |
|---|---|---|
| 298(1) | WP_277175580.1 | WP_277175580.1 hypothetical protein |
| 298(1) | WP_277166572.1 | WP_277166572.1 hypothetical protein |
| 298(1) | WP_277157933.1 | WP_277157933.1 hypothetical protein |
| 298(1) | WP_277157939.1 | WP_277157939.1 hypothetical protein |
| 299(1) | WP_272045560.1 | WP_272045560.1 hypothetical protein |
| 299(1) | WP_087442421.1 | WP_087442421.1 hypothetical protein |
| 299(2) | WP_213468112.1 | WP_213468112.1 hypothetical protein |
| 299(1) | WP_111153294.1 | WP_111153294.1 hypothetical protein |
| 299(2) | WP_143801053.1 | WP_143801053.1 hypothetical protein |
| 299(1) | WP_133378764.1 | WP_133378764.1 hypothetical protein |
| 299(1) | WP_168178778.1 | WP_168178778.1 hypothetical protein |
| 299(1) | WP_197256848.1 | WP_197256848.1 hypothetical protein |
| 299(1) | WP_281477429.1 | WP_281477429.1 hypothetical protein |
| 299(1) | WP_006674852.1 | WP_006674852.1 hypothetical protein |
| 299(1) | WP_213426983.1 | WP_213426983.1 hypothetical protein |
| 299(1) | WP_119794371.1 | WP_119794371.1 hypothetical protein |
| 300(1) | WP_221697949.1 | WP_221697949.1 hypothetical protein |
| 300(2) | WP_195949154.1 | WP_195949154.1 hypothetical protein |
| 300(2) | WP_195951610.1 | WP_195951610.1 hypothetical protein |
| 300(1) | WP_288161212.1 | WP_288161212.1 hypothetical protein |
| 300(6) | WP_042400998.1 | WP_042400998.1 hypothetical protein |
| 300(1) | WP_005210444.1 | WP_005210444.1 hypothetical protein |
| 300(1) | WP_195244536.1 | WP_195244536.1 hypothetical protein |
| 301(1) | WP_195953750.1 | WP_195953750.1 YARHG domain-containing protein |
| 301(1) | WP_055277384.1 | WP_055277384.1 YARHG domain-containing protein |
| 301(1) | WP_005210441.1 | WP_005210441.1 zinc ribbon domain-containing protein |
| 301(1) | WP_005210438.1 | WP_005210438.1 zinc-ribbon domain-containing protein |
| 301(1) | WP_288161219.1 | WP_288161219.1 YARHG domain-containing protein |
| 301(1) | WP_195244532.1 | WP_195244532.1 zinc-ribbon domain-containing protein |
| 301(1) | WP_288218585.1 | WP_288218585.1 YARHG domain-containing protein |
| 301(1) | WP_195949152.1 | WP_195949152.1 YARHG domain-containing protein |
| 301(1) | WP_195244534.1 | WP_195244534.1 YARHG domain-containing protein |
| 301(2) | WP_195951612.1 | WP_195951612.1 YARHG domain-containing protein |
| 301(2) | WP_195952583.1 | WP_195952583.1 YARHG domain-containing protein |
| 301(1) | WP_195955816.1 | WP_195955816.1 YARHG domain-containing protein |
| 302(1) | WP_006674853.1 | WP_006674853.1 hypothetical protein |
| 302(1) | WP_111153295.1 | WP_111153295.1 hypothetical protein |
| 302(1) | WP_272045561.1 | WP_272045561.1 hypothetical protein |
| 302(2) | WP_133378765.1 | WP_133378765.1 hypothetical protein |
| 302(1) | WP_197256850.1 | WP_197256850.1 hypothetical protein |
| 302(1) | WP_119794370.1 | WP_119794370.1 hypothetical protein |
| 302(1) | WP_293654340.1 | WP_293654340.1 hypothetical protein |
| 302(3) | WP_087442420.1 | WP_087442420.1 hypothetical protein |
| 302(2) | WP_168178779.1 | WP_168178779.1 hypothetical protein |
| 302(1) | WP_213426984.1 | WP_213426984.1 hypothetical protein |
| 303(1) | WP_289483776.1 | WP_289483776.1 MoxR family ATPase |
| 303(1) | WP_179160611.1 | WP_179160611.1 MoxR family ATPase |
| 303(2) | WP_053356040.1 | WP_053356040.1 MoxR family ATPase |
| 303(1) | WP_283892164.1 | WP_283892164.1 MoxR family ATPase |
| 303(1) | WP_062239891.1 | WP_062239891.1 MoxR family ATPase |
| 303(1) | WP_265408853.1 | WP_265408853.1 MoxR family ATPase |
| 303(1) | WP_301167491.1 | WP_301167491.1 MoxR family ATPase |
| 303(2) | WP_061975566.1 | WP_061975566.1 MoxR family ATPase |
| 303(1) | WP_185165652.1 | WP_185165652.1 MoxR family ATPase |
| 303(1) | WP_290401987.1 | WP_290401987.1 MoxR family ATPase |
| 303(1) | WP_090239205.1 | WP_090239205.1 MoxR family ATPase |
| 303(1) | WP_289475096.1 | WP_289475096.1 MoxR family ATPase |
| 304(1) | WP_000520843.1 | WP_000520843.1 transglutaminase domain-containing protein |
| 304(1) | WP_283892194.1 | WP_283892194.1 transglutaminase-like domain-containing protein |
| 304(1) | WP_289483791.1 | WP_289483791.1 transglutaminase-like domain-containing protein |
| 304(1) | WP_265408834.1 | WP_265408834.1 transglutaminase-like domain-containing protein |
| 304(1) | WP_248251194.1 | WP_248251194.1 transglutaminase-like domain-containing protein |
| 304(1) | WP_129478231.1 | WP_129478231.1 transglutaminase-like domain-containing protein |
| 304(1) | WP_061975596.1 | WP_061975596.1 transglutaminase-like domain-containing protein |
| 304(1) | WP_289475154.1 | WP_289475154.1 transglutaminase-like domain-containing protein |
| 304(1) | WP_090239253.1 | WP_090239253.1 transglutaminase-like domain-containing protein |
| 304(1) | WP_062239836.1 | WP_062239836.1 transglutaminase-like domain-containing protein |
| 304(1) | WP_185966789.1 | WP_185966789.1 dockerin type I domain-containing protein |
| 304(1) | WP_290402005.1 | WP_290402005.1 transglutaminase-like domain-containing protein |
| 304(1) | WP_256227726.1 | WP_256227726.1 transglutaminase-like domain-containing protein |
| 304(1) | WP_301167510.1 | WP_301167510.1 transglutaminase domain-containing protein |
| 305(1) | WP_268583866.1 | WP_268583866.1 LTA synthase family protein |
| 305(1) | WP_242490873.1 | WP_242490873.1 LTA synthase family protein |
| 305(1) | WP_242447698.1 | WP_242447698.1 LTA synthase family protein |
| 305(2) | WP_046217708.1 | WP_046217708.1 LTA synthase family protein |
| 305(1) | WP_260393147.1 | WP_260393147.1 LTA synthase family protein |
| 305(1) | WP_242450725.1 | WP_242450725.1 LTA synthase family protein |
| 305(1) | WP_251456541.1 | WP_251456541.1 LTA synthase family protein |
| 305(2) | WP_040365577.1 | WP_040365577.1 LTA synthase family protein |
| 305(1) | WP_235606181.1 | WP_235606181.1 LTA synthase family protein |
| 305(1) | WP_225985163.1 | WP_225985163.1 LTA synthase family protein |
| 305(1) | WP_243518742.1 | WP_243518742.1 LTA synthase family protein |
| 306(1) | WP_289475103.1 | WP_289475103.1 transglutaminase domain-containing protein |
| 306(1) | WP_272885664.1 | WP_272885664.1 transglutaminase domain-containing protein |
| 306(1) | WP_256227713.1 | WP_256227713.1 transglutaminase domain-containing protein |
| 306(1) | WP_283892167.1 | WP_283892167.1 transglutaminase domain-containing protein |
| 306(1) | WP_289483779.1 | WP_289483779.1 transglutaminase domain-containing protein |
| 306(1) | WP_301167493.1 | WP_301167493.1 transglutaminaseTgpA domain-containing protein |
| 306(1) | WP_062239883.1 | WP_062239883.1 transglutaminase domain-containing protein |
| 306(1) | WP_090239211.1 | WP_090239211.1 transglutaminaseTgpA domain-containing protein |
| 306(1) | WP_290401989.1 | WP_290401989.1 transglutaminaseTgpA domain-containing protein |
| 306(1) | WP_265408851.1 | WP_265408851.1 transglutaminase domain-containing protein |
| 306(1) | WP_129478252.1 | WP_129478252.1 transglutaminase domain-containing protein |
| 306(1) | WP_061975570.1 | WP_061975570.1 transglutaminase domain-containing protein |
| 306(1) | WP_179160610.1 | WP_179160610.1 transglutaminaseTgpA domain-containing protein |

307(2) WP_053356036.1 WP_053356036.1 glutamine-hydrolyzing GMP synthase
307(1) WP_129478250.1 WP_129478250.1 glutamine-hydrolyzing GMP synthase
307(1) WP_289483780.1 WP_289483780.1 glutamine-hydrolyzing GMP synthase
307(1) WP_061975573.1 WP_061975573.1 glutamine-hydrolyzing GMP synthase
307(2) WP_283892169.1 WP_283892169.1 glutamine-hydrolyzing GMP synthase
307(1) WP_301167494.1 WP_301167494.1 glutamine-hydrolyzing GMP synthase
307(1) WP_179160716.1 WP_179160716.1 glutamine-hydrolyzing GMP synthase
307(1) WP_290401990.1 WP_290401990.1 glutamine-hydrolyzing GMP synthase
307(1) WP_090239214.1 WP_090239214.1 glutamine-hydrolyzing GMP synthase
307(1) WP_265408850.1 WP_265408850.1 glutamine-hydrolyzing GMP synthase
307(1) WP_248251180.1 WP_248251180.1 glutamine-hydrolyzing GMP synthase

308(1) WP_062239879.1 WP_062239879.1 NCS2 family permease
308(1) WP_248251181.1 WP_248251181.1 NCS2 family permease
308(1) WP_290402007.1 WP_290402007.1 NCS2 family permease
308(1) WP_301167495.1 WP_301167495.1 NCS2 family permease
308(1) WP_289475111.1 WP_289475111.1 NCS2 family permease
308(1) WP_129478248.1 WP_129478248.1 NCS2 family permease
308(1) WP_165820922.1 WP_165820922.1 NCS2 family permease
308(1) WP_061975575.1 WP_061975575.1 NCS2 family permease
308(1) WP_256227714.1 WP_256227714.1 NCS2 family permease
308(1) WP_289483781.1 WP_289483781.1 NCS2 family permease
308(1) WP_283892171.1 WP_283892171.1 NCS2 family permease
308(1) WP_265408849.1 WP_265408849.1 NCS2 family permease
308(1) WP_090239217.1 WP_090239217.1 NCS2 family permease

309(1) WP_226679112.1 WP_226679112.1 gamma-glutamyltransferase
309(1) WP_102264561.1 WP_102264561.1 gamma-glutamyltransferase
309(1) WP_226086184.1 WP_226086184.1 gamma-glutamyltransferase
309(1) WP_226642854.1 WP_226642854.1 gamma-glutamyltransferase
309(1) WP_215032144.1 WP_215032144.1 gamma-glutamyltransferase family protein
309(1) WP_251542593.1 WP_251542593.1 gamma-glutamyltransferase family protein
309(1) WP_214902959.1 WP_214902959.1 gamma-glutamyltransferase
309(1) WP_231417852.1 WP_231417852.1 gamma-glutamyltransferase family protein
309(1) WP_251480980.1 WP_251480980.1 gamma-glutamyltransferase
309(1) WP_306049004.1 WP_306049004.1 gamma-glutamyltransferase
309(1) WP_144479084.1 WP_144479084.1 gamma-glutamyltransferase family protein
309(1) WP_283904351.1 WP_283904351.1 gamma-glutamyltransferase
309(1) WP_041964587.1 WP_041964587.1 gamma-glutamyltransferase

310(1) WP_129478254.1 WP_129478254.1 DUF58 domain-containing protein
310(1) WP_256227712.1 WP_256227712.1 DUF58 domain-containing protein
310(1) WP_289475100.1 WP_289475100.1 DUF58 domain-containing protein
310(1) WP_289483777.1 WP_289483777.1 DUF58 domain-containing protein
310(1) WP_265408852.1 WP_265408852.1 DUF58 domain-containing protein
310(1) WP_306430765.1 WP_306430765.1 DUF58 domain-containing protein
310(1) WP_061975568.1 WP_061975568.1 DUF58 domain-containing protein
310(1) WP_248251178.1 WP_248251178.1 DUF58 domain-containing protein
310(1) WP_090239208.1 WP_090239208.1 DUF58 domain-containing protein
310(1) WP_283892165.1 WP_283892165.1 DUF58 domain-containing protein
310(1) WP_062239887.1 WP_062239887.1 DUF58 domain-containing protein
310(1) WP_290401988.1 WP_290401988.1 DUF58 domain-containing protein
310(1) WP_301167492.1 WP_301167492.1 DUF58 domain-containing protein

311(1) WP_289342272.1 WP_289342272.1 4Fe-4S single cluster domain-containing protein
311(1) WP_251638751.1 WP_251638751.1 4Fe-4S single cluster domain-containing protein
311(1) WP_283952938.1 WP_283952938.1 4Fe-4S single cluster domain-containing protein
311(1) WP_038290964.1 WP_038290964.1 anaerobic ribonucleoside-triphosphate reductase activating protein
311(1) WP_217036000.1 WP_217036000.1 4Fe-4S single cluster domain-containing protein
311(1) WP_307589615.1 WP_307589615.1 pyruvate formate-lyase-activating protein
311(1) WP_213143937.1 WP_213143937.1 anaerobic ribonucleoside-triphosphate reductase activating protein
311(1) WP_263324559.1 WP_263324559.1 4Fe-4S single cluster domain-containing protein
311(1) WP_213117471.1 WP_213117471.1 4Fe-4S single cluster domain-containing protein
311(1) WP_003516593.1 WP_003516593.1 anaerobic ribonucleoside-triphosphate reductase activating protein
311(1) WP_284037995.1 WP_284037995.1 4Fe-4S single cluster domain-containing protein
311(1) WP_069193861.1 WP_069193861.1 anaerobic ribonucleoside-triphosphate reductase activating protein
311(1) WP_000238480.1 WP_000238480.1 pyruvate formate-lyase-activating protein

312(1) WP_302878970.1 WP_302878970.1 hypothetical protein
312(1) WP_302878995.1 WP_302878995.1 glycoside hydrolase family 2 TIM barrel-domain containing protein
312(1) WP_063858000.1 WP_063858000.1 beta-glucuronidase
312(1) WP_209365683.1 WP_209365683.1 glycoside hydrolase family 2 TIM barrel-domain containing protein
312(1) WP_127484327.1 WP_127484327.1 beta-glucuronidase
312(1) WP_309098602.1 WP_309098602.1 glycoside hydrolase family 2 TIM barrel-domain containing protein
312(1) WP_275100931.1 WP_275100931.1 glycoside hydrolase family 2 TIM barrel-domain containing protein
312(1) WP_229139909.1 WP_229139909.1 DUF4981 domain-containing protein
312(1) WP_025848634.1 WP_025848634.1 beta-glucuronidase
312(1) WP_074598763.1 WP_074598763.1 glycoside hydrolase family 2
312(1) WP_179223593.1 WP_179223593.1 beta-glucuronidase
312(1) WP_077619662.1 WP_077619662.1 DUF4981 domain-containing protein
312(1) WP_116353263.1 WP_116353263.1 glycoside hydrolase family 2 TIM barrel-domain containing protein

313(2) WP_185387009.1 WP_185387009.1 hypothetical protein
313(2) WP_185365686.1 WP_185365686.1 hypothetical protein
313(1) WP_185436219.1 WP_185436219.1 hypothetical protein
313(3) WP_194358205.1 WP_194358205.1 hypothetical protein
313(2) WP_185639009.1 WP_185639009.1 hypothetical protein
313(2) WP_181249349.1 WP_181249349.1 hypothetical protein

314(2) WP_185463562.1 WP_185463562.1 hypothetical protein
314(4) WP_185340230.1 WP_185340230.1 hypothetical protein
314(2) WP_200372545.1 WP_200372545.1 hypothetical protein
314(4) WP_203573361.1 WP_203573361.1 hypothetical protein

315(6) WP_036684078.1 WP_036684078.1 radical SAM/CxCxxxxC motif protein YfkAB

315(1) WP_154118045.1 WP_154118045.1 radical SAM/CxCxxxxC motif protein YfkAB
315(2) WP_076104567.1 WP_076104567.1 radical SAM/CxCxxxxC motif protein YfkAB
315(1) WP_076208244.1 WP_076208244.1 radical SAM/CxCxxxxC motif protein YfkAB
315(1) WP_039871643.1 WP_039871643.1 radical SAM/CxCxxxxC motif protein YfkAB
315(1) WP_209872764.1 WP_209872764.1 radical SAM/CxCxxxxC motif protein YfkAB

316(1) WP_213649970.1 WP_213649970.1 hypothetical protein
316(1) WP_205001948.1 WP_205001948.1 hypothetical protein
316(1) WP_244719633.1 WP_244719633.1 hypothetical protein
316(1) WP_205001949.1 WP_205001949.1 hypothetical protein
316(1) WP_194230403.1 WP_194230403.1 hypothetical protein
316(1) WP_205001946.1 WP_205001946.1 hypothetical protein
316(1) WP_036707108.1 WP_036707108.1 hypothetical protein
316(1) WP_142255386.1 WP_142255386.1 hypothetical protein
316(1) WP_136604814.1 WP_136604814.1 hypothetical protein
316(1) WP_160496931.1 WP_160496931.1 hypothetical protein
316(1) WP_160037486.1 WP_160037486.1 hypothetical protein
316(1) WP_212980185.1 WP_212980185.1 hypothetical protein

317(1) WP_039871644.1 WP_039871644.1 hemolysin family protein
317(3) WP_076135925.1 WP_076135925.1 hemolysin family protein
317(1) WP_245368173.1 WP_245368173.1 hemolysin family protein
317(3) WP_036684171.1 WP_036684171.1 hemolysin family protein
317(1) WP_076125129.1 WP_076125129.1 hemolysin family protein
317(2) WP_076187132.1 WP_076187132.1 hemolysin family protein
317(1) WP_154118200.1 WP_154118200.1 hemolysin family protein

318(3) WP_194358204.1 WP_194358204.1 hypothetical protein
318(2) WP_270995159.1 WP_270995159.1 hypothetical protein
318(2) WP_185639008.1 WP_185639008.1 hypothetical protein
318(2) WP_185472556.1 WP_185472556.1 hypothetical protein
318(1) WP_185451812.1 WP_185451812.1 hypothetical protein
318(2) WP_194335285.1 WP_194335285.1 hypothetical protein

319(1) WP_034773582.1 WP_034773582.1 UvrD-helicase domain-containing protein
319(1) WP_277177074.1 WP_277177074.1 ATP-dependent helicase
319(1) WP_181754680.1 WP_181754680.1 UvrD-helicase domain-containing protein
319(1) WP_277150890.1 WP_277150890.1 ATP-dependent helicase
319(1) WP_097911854.1 WP_097911854.1 AAA family ATPase
319(1) WP_277157962.1 WP_277157962.1 ATP-dependent helicase
319(1) WP_053403795.1 WP_053403795.1 RNA polymerase recycling motor HelD
319(1) WP_212695777.1 WP_212695777.1 ATP-dependent helicase
319(1) WP_242172401.1 WP_242172401.1 RNA polymerase recycling motor HelD
319(1) WP_277142109.1 WP_277142109.1 ATP-dependent helicase
319(1) WP_251610760.1 WP_251610760.1 RNA polymerase recycling motor HelD
319(1) WP_277156199.1 WP_277156199.1 ATP-dependent helicase

320(1) WP_283892163.1 WP_283892163.1 phage holin family protein
320(3) WP_061975564.1 WP_061975564.1 phage holin family protein
320(1) WP_290401986.1 WP_290401986.1 phage holin family protein
320(1) WP_090239202.1 WP_090239202.1 phage holin family protein
320(1) WP_053356041.1 WP_053356041.1 phage holin family protein
320(1) WP_289475092.1 WP_289475092.1 phage holin family protein
320(3) WP_062239894.1 WP_062239894.1 phage holin family protein
320(1) WP_301167490.1 WP_301167490.1 phage holin family protein

321(1) WP_129478237.1 WP_129478237.1 hypothetical protein
321(1) WP_301167507.1 WP_301167507.1 hypothetical protein
321(1) WP_248251191.1 WP_248251191.1 hypothetical protein
321(1) WP_289475147.1 WP_289475147.1 hypothetical protein
321(1) WP_290402002.1 WP_290402002.1 hypothetical protein
321(1) WP_283892189.1 WP_283892189.1 hypothetical protein
321(1) WP_265408837.1 WP_265408837.1 hypothetical protein
321(2) WP_061975590.1 WP_061975590.1 hypothetical protein
321(1) WP_256227723.1 WP_256227723.1 hypothetical protein
321(1) WP_053356023.1 WP_053356023.1 hypothetical protein
321(1) WP_289483788.1 WP_289483788.1 hypothetical protein

322(2) WP_283892192.1 WP_283892192.1 EcsC family protein
322(1) WP_129478233.1 WP_129478233.1 EcsC family protein
322(1) WP_090239249.1 WP_090239249.1 EcsC family protein
322(1) WP_289487842.1 WP_289487842.1 EcsC family protein
322(1) WP_301167509.1 WP_301167509.1 EcsC family protein
322(1) WP_265408835.1 WP_265408835.1 EcsC family protein
322(1) WP_290402004.1 WP_290402004.1 EcsC family protein
322(1) WP_256227725.1 WP_256227725.1 EcsC family protein
322(1) WP_061975594.1 WP_061975594.1 EcsC family protein
322(1) WP_062239839.1 WP_062239839.1 EcsC family protein
322(1) WP_248251193.1 WP_248251193.1 EcsC family protein

323(1) WP_265410612.1 WP_265410612.1 nicotinate phosphoribosyltransferase
323(1) WP_283895398.1 WP_283895398.1 nicotinate phosphoribosyltransferase
323(1) WP_248254560.1 WP_248254560.1 nicotinate phosphoribosyltransferase
323(1) WP_290402008.1 WP_290402008.1 nicotinate phosphoribosyltransferase
323(1) WP_090239274.1 WP_090239274.1 nicotinate phosphoribosyltransferase
323(1) WP_301167514.1 WP_301167514.1 nicotinate phosphoribosyltransferase
323(1) WP_256227727.1 WP_256227727.1 nicotinate phosphoribosyltransferase
323(2) WP_289479601.1 WP_289479601.1 nicotinate phosphoribosyltransferase
323(2) WP_061975608.1 WP_061975608.1 nicotinate phosphoribosyltransferase
323(1) WP_062240006.1 WP_062240006.1 nicotinate phosphoribosyltransferase

324(1) WP_156327468.1 WP_156327468.1 hypothetical protein
324(1) WP_195951614.1 WP_195951614.1 hypothetical protein
324(1) WP_195959149.1 WP_195959149.1 hypothetical protein
324(1) WP_302579555.1 WP_302579555.1 hypothetical protein
324(1) WP_195953752.1 WP_195953752.1 hypothetical protein
324(1) WP_195959148.1 WP_195959148.1 hypothetical protein
324(1) WP_288218582.1 WP_288218582.1 hypothetical protein
324(1) WP_207743578.1 WP_207743578.1 hypothetical protein
324(1) WP_195955815.1 WP_195955815.1 hypothetical protein
324(1) WP_196048851.1 WP_196048851.1 hypothetical protein
324(1) WP_195949149.1 WP_195949149.1 hypothetical protein
324(1) WP_055277379.1 WP_055277379.1 hypothetical protein

| | | |
|---|---|---|
| 325(1) | WP_251497187.1 | WP_251497187.1 ABC-2 transporter permease |
| 325(1) | WP_096774451.1 | WP_096774451.1 ABC-2 transporter permease |
| 325(1) | WP_145037678.1 | WP_145037678.1 ABC-2 transporter permease |
| 325(3) | WP_192571322.1 | WP_192571322.1 ABC-2 transporter permease |
| 325(1) | WP_009590634.1 | WP_009590634.1 ABC-2 transporter permease |
| 325(1) | WP_108465354.1 | WP_108465354.1 ABC-2 transporter permease |
| 325(1) | WP_241929671.1 | WP_241929671.1 ABC-2 transporter permease |
| 325(1) | WP_158702652.1 | WP_158702652.1 ABC-2 transporter permease |
| 325(1) | WP_302847864.1 | WP_302847864.1 hypothetical protein |
| 325(1) | WP_015736237.1 | WP_015736237.1 ABC-2 transporter permease |
| 326(1) | WP_303942555.1 | WP_303942555.1 phosphoglucosamine mutase |
| 326(1) | WP_089965956.1 | WP_089965956.1 phospho-sugar mutase |
| 326(1) | WP_072336536.1 | WP_072336536.1 phosphoglucosamine mutase |
| 326(2) | WP_102372738.1 | WP_102372738.1 phosphoglucosamine mutase |
| 326(1) | WP_098579102.1 | WP_098579102.1 hypothetical protein |
| 326(1) | WP_141539749.1 | WP_141539749.1 hypothetical protein |
| 326(1) | WP_291289944.1 | WP_291289944.1 phosphoglucosamine mutase |
| 326(3) | WP_291288251.1 | WP_291288251.1 phosphoglucosamine mutase |
| 326(1) | WP_273383893.1 | WP_273383893.1 phosphoglucosamine mutase |
| 327(1) | WP_305166257.1 | WP_305166257.1 hypothetical protein |
| 327(2) | WP_071558841.1 | WP_071558841.1 hypothetical protein |
| 327(1) | WP_254327967.1 | WP_254327967.1 hypothetical protein |
| 327(2) | WP_061829402.1 | WP_061829402.1 hypothetical protein |
| 327(1) | WP_165147470.1 | WP_165147470.1 hypothetical protein |
| 327(1) | WP_137062805.1 | WP_137062805.1 hypothetical protein |
| 327(1) | WP_165079982.1 | WP_165079982.1 hypothetical protein |
| 327(2) | WP_017427969.1 | WP_017427969.1 hypothetical protein |
| 327(1) | WP_025719051.1 | WP_025719051.1 hypothetical protein |
| 328(1) | WP_289376831.1 | WP_289376831.1 DUF3600 domain-containing protein |
| 328(1) | WP_248063413.1 | WP_248063413.1 DUF3600 domain-containing protein |
| 328(1) | WP_184183067.1 | WP_184183067.1 DUF3600 domain-containing protein |
| 328(1) | WP_076210940.1 | WP_076210940.1 DUF3600 domain-containing protein |
| 328(1) | WP_216518278.1 | WP_216518278.1 DUF3600 domain-containing protein |
| 328(1) | WP_244964941.1 | WP_244964941.1 DUF3600 domain-containing protein |
| 328(1) | WP_308723537.1 | WP_308723537.1 DUF3600 domain-containing protein |
| 328(1) | WP_211082914.1 | WP_211082914.1 DUF3600 domain-containing protein |
| 328(1) | WP_017692330.1 | WP_017692330.1 DUF3600 domain-containing protein |
| 328(1) | WP_150364180.1 | WP_150364180.1 DUF3600 domain-containing protein |
| 328(1) | WP_024634135.1 | WP_024634135.1 DUF3600 domain-containing protein |
| 328(1) | WP_253440823.1 | WP_253440823.1 DUF3600 domain-containing protein |
| 329(5) | WP_000761187.1 | WP_000761187.1 hypothetical protein |
| 329(5) | WP_000761174.1 | WP_000761174.1 hypothetical protein |
| 329(1) | WP_000761171.1 | WP_000761171.1 hypothetical protein |
| 330(1) | WP_174820836.1 | WP_174820836.1 PadR family transcriptional regulator |
| 330(1) | WP_177189473.1 | WP_177189473.1 PadR family transcriptional regulator |
| 330(3) | WP_045243837.1 | WP_045243837.1 PadR family transcriptional regulator |
| 330(1) | WP_090887127.1 | WP_090887127.1 PadR family transcriptional regulator |
| 330(2) | WP_126665665.1 | WP_126665665.1 PadR family transcriptional regulator |
| 330(1) | WP_185156694.1 | WP_185156694.1 PadR family transcriptional regulator |
| 330(1) | WP_016821035.1 | WP_016821035.1 PadR family transcriptional regulator |
| 330(1) | WP_107734205.1 | WP_107734205.1 PadR family transcriptional regulator |
| 331(1) | WP_085117895.1 | WP_085117895.1 GMP reductase |
| 331(1) | WP_151585718.1 | WP_151585718.1 GMP reductase |
| 331(1) | WP_197234702.1 | WP_197234702.1 GMP reductase |
| 331(1) | WP_113753113.1 | WP_113753113.1 GMP reductase |
| 331(7) | WP_019392130.1 | WP_019392130.1 GMP reductase |
| 332(2) | WP_133332989.1 | WP_133332989.1 DNA-directed RNA polymerase subunit beta |
| 332(1) | WP_026572750.1 | WP_026572750.1 DNA-directed RNA polymerase subunit beta |
| 332(1) | WP_215032096.1 | WP_215032096.1 DNA-directed RNA polymerase subunit beta |
| 332(1) | WP_226642895.1 | WP_226642895.1 DNA-directed RNA polymerase subunit beta |
| 332(1) | WP_053364458.1 | WP_053364458.1 DNA-directed RNA polymerase subunit beta |
| 332(1) | WP_046522954.1 | WP_046522954.1 DNA-directed RNA polymerase subunit beta |
| 332(1) | WP_198310822.1 | WP_198310822.1 DNA-directed RNA polymerase subunit beta |
| 332(1) | WP_068673745.1 | WP_068673745.1 DNA-directed RNA polymerase subunit beta |
| 332(1) | WP_102264538.1 | WP_102264538.1 DNA-directed RNA polymerase subunit beta |
| 332(1) | WP_226086165.1 | WP_226086165.1 DNA-directed RNA polymerase subunit beta |
| 333(1) | WP_098776424.1 | WP_098776424.1 ABC transporter permease subunit |
| 333(1) | WP_098776425.1 | WP_098776425.1 peptide ABC transporter permease |
| 333(1) | WP_106408056.1 | WP_106408056.1 hypothetical protein |
| 333(1) | WP_273541946.1 | WP_273541946.1 hypothetical protein |
| 333(1) | WP_116500446.1 | WP_116500446.1 hypothetical protein |
| 333(1) | WP_235675260.1 | WP_235675260.1 hypothetical protein |
| 333(1) | WP_232290549.1 | WP_232290549.1 hypothetical protein |
| 333(1) | WP_227590927.1 | WP_227590927.1 hypothetical protein |
| 333(1) | WP_097905173.1 | WP_097905173.1 ABC transporter permease subunit |
| 333(1) | WP_193653014.1 | WP_193653014.1 peptide ABC transporter permease |
| 333(1) | WP_097905172.1 | WP_097905172.1 ABC transporter permease subunit |
| 334(1) | WP_077893588.1 | WP_077893588.1 Cof-type HAD-IIB family hydrolase |
| 334(1) | WP_077833119.1 | WP_077833119.1 Cof-type HAD-IIB family hydrolase |
| 334(1) | WP_283952955.1 | WP_283952955.1 sugar-phosphatase |
| 334(1) | WP_077852425.1 | WP_077852425.1 Cof-type HAD-IIB family hydrolase |
| 334(1) | WP_257675223.1 | WP_257675223.1 Cof-type HAD-IIB family hydrolase |
| 334(3) | WP_213117466.1 | WP_213117466.1 HAD family hydrolase |
| 334(1) | WP_089965928.1 | WP_089965928.1 Cof-type HAD-IIB family hydrolase |
| 334(1) | WP_246427691.1 | WP_246427691.1 HAD-IIB family hydrolase |
| 334(1) | WP_289342252.1 | WP_289342252.1 HAD family hydrolase |
| 335(1) | WP_198050878.1 | WP_198050878.1 glutamate-1-semialdehyde 2,1-aminomutase |
| 335(1) | WP_307192886.1 | WP_307192886.1 aspartate aminotransferase family protein |
| 335(1) | WP_049720634.1 | WP_049720634.1 glutamate-1-semialdehyde 2,1-aminomutase |
| 335(1) | WP_044037254.1 | WP_044037254.1 glutamate-1-semialdehyde 2,1-aminomutase |
| 335(5) | WP_037993166.1 | WP_037993166.1 glutamate-1-semialdehyde 2,1-aminomutase |
| 335(1) | WP_198053924.1 | WP_198053924.1 glutamate-1-semialdehyde 2,1-aminomutase |
| 335(1) | WP_181737021.1 | WP_181737021.1 aspartate aminotransferase family protein |

| | | |
|---|---|---|
| 336(1) | WP_155703940.1 | WP_155703940.1 hypothetical protein |
| 336(1) | WP_138755927.1 | WP_138755927.1 hypothetical protein |
| 336(1) | WP_307601015.1 | WP_307601015.1 hypothetical protein |
| 336(1) | WP_134750620.1 | WP_134750620.1 hypothetical protein |
| 336(1) | WP_238187875.1 | WP_238187875.1 hypothetical protein |
| 336(1) | WP_127570444.1 | WP_127570444.1 hypothetical protein |
| 336(1) | WP_182301843.1 | WP_182301843.1 hypothetical protein |
| 336(1) | WP_027088419.1 | WP_027088419.1 hypothetical protein |
| 336(1) | WP_136372949.1 | WP_136372949.1 hypothetical protein |
| 336(1) | WP_120112277.1 | WP_120112277.1 hypothetical protein |
| 336(1) | WP_239617483.1 | WP_239617483.1 hypothetical protein |
| 337(1) | WP_136372950.1 | WP_136372950.1 hypothetical protein |
| 337(1) | WP_238187876.1 | WP_238187876.1 hypothetical protein |
| 337(1) | WP_127570443.1 | WP_127570443.1 hypothetical protein |
| 337(1) | WP_307601017.1 | WP_307601017.1 hypothetical protein |
| 337(1) | WP_155703939.1 | WP_155703939.1 hypothetical protein |
| 337(1) | WP_182301842.1 | WP_182301842.1 hypothetical protein |
| 337(1) | WP_138755926.1 | WP_138755926.1 hypothetical protein |
| 337(1) | WP_239617482.1 | WP_239617482.1 hypothetical protein |
| 337(1) | WP_120112274.1 | WP_120112274.1 hypothetical protein |
| 337(1) | WP_027088420.1 | WP_027088420.1 hypothetical protein |
| 337(1) | WP_134750616.1 | WP_134750616.1 hypothetical protein |
| 338(1) | WP_239617481.1 | WP_239617481.1 hypothetical protein |
| 338(1) | WP_155703938.1 | WP_155703938.1 hypothetical protein |
| 338(1) | WP_134750614.1 | WP_134750614.1 hypothetical protein |
| 338(1) | WP_138755925.1 | WP_138755925.1 hypothetical protein |
| 338(1) | WP_182301841.1 | WP_182301841.1 hypothetical protein |
| 338(1) | WP_127570442.1 | WP_127570442.1 hypothetical protein |
| 338(1) | WP_120112272.1 | WP_120112272.1 hypothetical protein |
| 338(1) | WP_027088421.1 | WP_027088421.1 hypothetical protein |
| 338(1) | WP_136372951.1 | WP_136372951.1 hypothetical protein |
| 338(1) | WP_238187877.1 | WP_238187877.1 hypothetical protein |
| 338(1) | WP_307601019.1 | WP_307601019.1 hypothetical protein |
| 339(1) | WP_126996287.1 | WP_126996287.1 hypothetical protein |
| 339(2) | WP_185375607.1 | WP_185375607.1 hypothetical protein |
| 339(1) | WP_077320215.1 | WP_077320215.1 hypothetical protein |
| 339(1) | WP_074598751.1 | WP_074598751.1 hypothetical protein |
| 339(2) | WP_185336758.1 | WP_185336758.1 hypothetical protein |
| 339(1) | WP_185350815.1 | WP_185350815.1 hypothetical protein |
| 339(1) | WP_307601048.1 | WP_307601048.1 hypothetical protein |
| 339(1) | WP_077321500.1 | WP_077321500.1 hypothetical protein |
| 339(1) | WP_181603075.1 | WP_181603075.1 hypothetical protein |
| 340(1) | WP_048826594.1 | WP_048826594.1 PTS ascorbate transporter subunit IIC |
| 340(1) | WP_195953737.1 | WP_195953737.1 PTS ascorbate transporter subunit IIC |
| 340(1) | WP_121874304.1 | WP_121874304.1 PTS ascorbate transporter subunit IIC |
| 340(1) | WP_181731622.1 | WP_181731622.1 PTS ascorbate transporter subunit IIC |
| 340(6) | WP_131470899.1 | WP_131470899.1 PTS ascorbate transporter subunit IIC |

| | | |
|---|---|---|
| 341(1) | WP_195953744.1 | WP_195953744.1 MarR family transcriptional regulator |
| 341(9) | WP_042400980.1 | WP_042400980.1 MarR family transcriptional regulator |
| 342(1) | WP_195949156.1 | WP_195949156.1 IS1 family transposase |
| 342(1) | WP_196048855.1 | WP_196048855.1 IS1 family transposase |
| 342(1) | WP_055277394.1 | WP_055277394.1 hypothetical protein |
| 342(6) | WP_042400987.1 | WP_042400987.1 hypothetical protein |
| 342(1) | WP_195953746.1 | WP_195953746.1 IS1 family transposase |
| 343(1) | WP_179231919.1 | WP_179231919.1 hypothetical protein |
| 343(1) | WP_186325902.1 | WP_186325902.1 hypothetical protein |
| 343(1) | WP_181909013.1 | WP_181909013.1 hypothetical protein |
| 343(1) | WP_167527255.1 | WP_167527255.1 hypothetical protein |
| 343(1) | WP_167372840.1 | WP_167372840.1 hypothetical protein |
| 343(1) | WP_212983253.1 | WP_212983253.1 hypothetical protein |
| 343(3) | WP_186320103.1 | WP_186320103.1 hypothetical protein |
| 343(1) | WP_186327521.1 | WP_186327521.1 hypothetical protein |
| 344(1) | WP_143315262.1 | WP_143315262.1 4'-phosphopantetheinyl transferase superfamily protein |
| 344(1) | WP_138614840.1 | WP_138614840.1 4'-phosphopantetheinyl transferase superfamily protein |
| 344(1) | WP_198053922.1 | WP_198053922.1 4'-phosphopantetheinyl transferase superfamily protein |
| 344(3) | WP_037993160.1 | WP_037993160.1 4'-phosphopantetheinyl transferase superfamily protein |
| 344(1) | WP_121874306.1 | WP_121874306.1 4'-phosphopantetheinyl transferase superfamily protein |
| 344(1) | WP_181737009.1 | WP_181737009.1 4'-phosphopantetheinyl transferase superfamily protein |
| 344(1) | WP_236533771.1 | WP_236533771.1 4'-phosphopantetheinyl transferase superfamily protein |
| 344(1) | WP_198050875.1 | WP_198050875.1 4'-phosphopantetheinyl transferase superfamily protein |
| 345(1) | WP_129549129.1 | WP_129549129.1 peptidase M56 |
| 345(1) | WP_216975175.1 | WP_216975175.1 peptidase M56 |
| 345(1) | WP_233901860.1 | WP_233901860.1 hypothetical protein |
| 345(1) | WP_258013844.1 | WP_258013844.1 peptidase M56 |
| 345(1) | WP_137078368.1 | WP_137078368.1 peptidase M56 |
| 345(1) | WP_243537355.1 | WP_243537355.1 peptidase M56 |
| 345(1) | WP_228552643.1 | WP_228552643.1 hypothetical protein |
| 345(2) | WP_013370791.1 | WP_013370791.1 peptidase M56 |
| 345(1) | WP_253806480.1 | WP_253806480.1 peptidase M56 |
| 346(1) | WP_198053921.1 | WP_198053921.1 hypothetical protein |
| 346(4) | WP_131470900.1 | WP_131470900.1 hypothetical protein |
| 346(1) | WP_138614841.1 | WP_138614841.1 hypothetical protein |
| 346(1) | WP_268983094.1 | WP_268983094.1 hypothetical protein |
| 346(1) | WP_037993157.1 | WP_037993157.1 hypothetical protein |
| 346(1) | WP_236533770.1 | WP_236533770.1 hypothetical protein |
| 346(1) | WP_181737007.1 | WP_181737007.1 hypothetical protein |
| 347(1) | WP_058486877.1 | WP_058486877.1 PHP domain-containing protein |
| 347(1) | WP_019127576.1 | WP_019127576.1 PHP domain-containing protein |
| 347(1) | WP_277142115.1 | WP_277142115.1 PHP domain-containing protein |
| 347(1) | WP_277157967.1 | WP_277157967.1 PHP domain-containing protein |
| 347(1) | WP_277150338.1 | WP_277150338.1 PHP domain-containing protein |
| 347(1) | WP_006289743.1 | WP_006289743.1 PHP domain-containing protein |
| 347(1) | WP_239458681.1 | WP_239458681.1 PHP domain-containing protein |

| | | |
|---|---|---|
| 347(1) | WP_102371689.1 | WP_102371689.1 PHP domain-containing protein |
| 347(2) | WP_006290900.1 | WP_006290900.1 PHP domain-containing protein |
| 348(1) | WP_006290902.1 | WP_006290902.1 NADH:flavin oxidoreductase/NADH oxidase |
| 348(1) | WP_006289740.1 | WP_006289740.1 NADH:flavin oxidoreductase/NADH oxidase |
| 348(1) | WP_074108327.1 | WP_074108327.1 NADH:flavin oxidoreductase/NADH oxidase |
| 348(1) | WP_165175570.1 | WP_165175570.1 NADH:flavin oxidoreductase |
| 348(1) | WP_103047821.1 | WP_103047821.1 NADH:flavin oxidoreductase |
| 348(1) | WP_281667413.1 | WP_281667413.1 NADH:flavin oxidoreductase/NADH oxidase |
| 348(1) | WP_134902882.1 | WP_134902882.1 NADH:flavin oxidoreductase |
| 348(1) | WP_074085372.1 | WP_074085372.1 NADH:flavin oxidoreductase/NADH oxidase |
| 348(1) | WP_084686059.1 | WP_084686059.1 NADH:flavin oxidoreductase/NADH oxidase |
| 348(1) | WP_250244826.1 | WP_250244826.1 NADH:flavin oxidoreductase |
| 349(1) | WP_281937344.1 | WP_281937344.1 Zn-dependent hydrolase |
| 349(1) | WP_127459452.1 | WP_127459452.1 Zn-dependent hydrolase |
| 349(1) | WP_258208943.1 | WP_258208943.1 allantoate deiminase |
| 349(1) | WP_010496625.1 | WP_010496625.1 Zn-dependent hydrolase |
| 349(1) | WP_251598134.1 | WP_251598134.1 Zn-dependent hydrolase |
| 349(1) | WP_108534970.1 | WP_108534970.1 Zn-dependent hydrolase |
| 349(1) | WP_036689478.1 | WP_036689478.1 Zn-dependent hydrolase |
| 349(1) | WP_216791250.1 | WP_216791250.1 Zn-dependent hydrolase |
| 349(1) | WP_163853464.1 | WP_163853464.1 Zn-dependent hydrolase |
| 349(1) | WP_253411104.1 | WP_253411104.1 Zn-dependent hydrolase |
| 350(1) | WP_281937346.1 | WP_281937346.1 enolase C-terminal domain-like protein |
| 350(1) | WP_216791247.1 | WP_216791247.1 enolase C-terminal domain-like protein |
| 350(1) | WP_063188030.1 | WP_063188030.1 hypothetical protein |
| 350(1) | WP_253411102.1 | WP_253411102.1 enolase C-terminal domain-like protein |
| 350(1) | WP_010496624.1 | WP_010496624.1 enolase C-terminal domain-like protein |
| 350(1) | WP_251598135.1 | WP_251598135.1 enolase C-terminal domain-like protein |
| 350(1) | WP_189597464.1 | WP_189597464.1 enolase C-terminal domain-like protein |
| 350(1) | WP_163853465.1 | WP_163853465.1 hypothetical protein |
| 350(1) | WP_127459450.1 | WP_127459450.1 enolase C-terminal domain-like protein |
| 350(1) | WP_036689473.1 | WP_036689473.1 hypothetical protein |
| 351(1) | WP_295632047.1 | WP_295632047.1 S-layer homology domain-containing protein |
| 351(1) | WP_089965932.1 | WP_089965932.1 Ig-like domain-containing protein |
| 351(1) | WP_295628317.1 | WP_295628317.1 S-layer homology domain-containing protein |
| 351(1) | WP_289607496.1 | WP_289607496.1 hypothetical protein |
| 351(1) | WP_227219188.1 | WP_227219188.1 Ig-like domain-containing protein |
| 351(1) | WP_005604630.1 | WP_005604630.1 Ig-like domain-containing protein |
| 351(1) | WP_294386445.1 | WP_294386445.1 Ig-like domain-containing protein |
| 351(1) | WP_276538701.1 | WP_276538701.1 Ig-like domain-containing protein |
| 351(1) | WP_195325257.1 | WP_195325257.1 Ig-like domain-containing protein |
| 351(1) | WP_148409441.1 | WP_148409441.1 Ig-like domain-containing protein |
| 352(1) | WP_196048852.1 | WP_196048852.1 class B sortase |
| 352(1) | WP_195953751.1 | WP_195953751.1 class B sortase |
| 352(1) | WP_302579554.1 | WP_302579554.1 class B sortase |
| 352(1) | WP_195949151.1 | WP_195949151.1 class B sortase |
| 352(2) | WP_195951613.1 | WP_195951613.1 class B sortase |
| 352(1) | WP_288218583.1 | WP_288218583.1 class B sortase |
| 352(2) | WP_055277383.1 | WP_055277383.1 class B sortase |
| 352(1) | WP_294953117.1 | WP_294953117.1 class B sortase |
| 353(1) | WP_288218581.1 | WP_288218581.1 thioester domain-containing protein |
| 353(1) | WP_195959150.1 | WP_195959150.1 thioester domain-containing protein |
| 353(1) | WP_195955814.1 | WP_195955814.1 thioester domain-containing protein |
| 353(1) | WP_302579556.1 | WP_302579556.1 thioester domain-containing protein |
| 353(1) | WP_195951615.1 | WP_195951615.1 thioester domain-containing protein |
| 353(1) | WP_005210431.1 | WP_005210431.1 thioester domain-containing protein |
| 353(1) | WP_195244531.1 | WP_195244531.1 thioester domain-containing protein |
| 353(1) | WP_195953753.1 | WP_195953753.1 thioester domain-containing protein |
| 353(1) | WP_195949148.1 | WP_195949148.1 thioester domain-containing protein |
| 353(1) | WP_055277377.1 | WP_055277377.1 thioester domain-containing protein |
| 354(1) | WP_243495566.1 | WP_243495566.1 tyrosine protein phosphatase |
| 354(2) | WP_055277375.1 | WP_055277375.1 capsular biosynthesis protein |
| 354(1) | WP_195244530.1 | WP_195244530.1 capsular biosynthesis protein |
| 354(1) | WP_005210424.1 | WP_005210424.1 hypothetical protein |
| 354(1) | WP_272801666.1 | WP_272801666.1 tyrosine protein phosphatase |
| 354(1) | WP_195953754.1 | WP_195953754.1 capsular biosynthesis protein |
| 354(1) | WP_195951616.1 | WP_195951616.1 capsular biosynthesis protein |
| 354(1) | WP_098579105.1 | WP_098579105.1 CpsB/CapC family capsule biosynthesis tyrosine phosphatase |
| 354(1) | WP_302579557.1 | WP_302579557.1 capsular biosynthesis protein |
| 355(1) | WP_179603490.1 | WP_179603490.1 sucrose-6-phosphate hydrolase |
| 355(1) | WP_129478264.1 | WP_129478264.1 glycoside hydrolase family 32 protein |
| 355(1) | WP_090239185.1 | WP_090239185.1 glycoside hydrolase family 32 protein |
| 355(1) | WP_034675324.1 | WP_034675324.1 sucrose-6-phosphate hydrolase |
| 355(1) | WP_301167484.1 | WP_301167484.1 glycoside hydrolase family 32 protein |
| 355(1) | WP_061975606.1 | WP_061975606.1 glycoside hydrolase family 32 protein |
| 355(1) | WP_307312524.1 | WP_307312524.1 sucrose-6-phosphate hydrolase |
| 355(1) | WP_289475075.1 | WP_289475075.1 glycoside hydrolase family 32 protein |
| 355(1) | WP_306077402.1 | WP_306077402.1 sucrose-6-phosphate hydrolase |
| 355(1) | WP_289483769.1 | WP_289483769.1 glycoside hydrolase family 32 protein |
| 356(1) | WP_182301840.1 | WP_182301840.1 hypothetical protein |
| 356(1) | WP_127570441.1 | WP_127570441.1 hypothetical protein |
| 356(1) | WP_238187878.1 | WP_238187878.1 hypothetical protein |
| 356(1) | WP_307601021.1 | WP_307601021.1 hypothetical protein |
| 356(1) | WP_155703937.1 | WP_155703937.1 hypothetical protein |
| 356(1) | WP_239617480.1 | WP_239617480.1 hypothetical protein |
| 356(1) | WP_134750612.1 | WP_134750612.1 hypothetical protein |
| 356(1) | WP_138755924.1 | WP_138755924.1 hypothetical protein |
| 356(1) | WP_120112269.1 | WP_120112269.1 hypothetical protein |
| 356(1) | WP_136372952.1 | WP_136372952.1 hypothetical protein |
| 357(9) | WP_042401008.1 | WP_042401008.1 nucleotidyltransferase domain-containing protein |
| 358(9) | WP_042401010.1 | WP_042401010.1 nucleotidyltransferase substrate binding protein |

359(4) WP_055277396.1 WP_055277396.1 ATP-binding protein
359(3) WP_042401406.1 WP_042401406.1 ATP-binding protein
359(2) WP_005210458.1 WP_005210458.1 ATP-binding protein

360(6) WP_076124810.1 WP_076124810.1 potassium channel family protein
360(3) WP_076104570.1 WP_076104570.1 potassium channel family protein

361(8) WP_036684073.1 WP_036684073.1 CDP-diacylglycerol--serine O-phosphatidyltransferase
361(1) WP_076208246.1 WP_076208246.1 CDP-diacylglycerol--serine O-phosphatidyltransferase

362(1) WP_074085368.1 WP_074085368.1 undecaprenyl-diphosphate phosphatase
362(2) WP_076124791.1 WP_076124791.1 undecaprenyl-diphosphate phosphatase
362(3) WP_036684081.1 WP_036684081.1 undecaprenyl-diphosphate phosphatase
362(1) WP_076309110.1 WP_076309110.1 undecaprenyl-diphosphate phosphatase
362(1) WP_074102530.1 WP_074102530.1 undecaprenyl-diphosphate phosphatase
362(1) WP_039871640.1 WP_039871640.1 undecaprenyl-diphosphate phosphatase

363(1) WP_169505065.1 WP_169505065.1 DivIVA domain-containing protein
363(2) WP_091155408.1 WP_091155408.1 DivIVA domain-containing protein
363(2) WP_047909601.1 WP_047909601.1 DivIVA domain-containing protein
363(2) WP_076311040.1 WP_076311040.1 DivIVA domain-containing protein
363(2) WP_138226014.1 WP_138226014.1 DivIVA domain-containing protein

364(1) WP_122971526.1 WP_122971526.1 long-chain fatty acid--CoA ligase
364(4) WP_037997349.1 WP_037997349.1 long-chain-fatty-acid--CoA ligase
364(1) WP_181731436.1 WP_181731436.1 long-chain-fatty-acid--CoA ligase
364(2) WP_121874297.1 WP_121874297.1 long-chain-fatty-acid--CoA ligase
364(1) WP_035190295.1 WP_035190295.1 acetoacetate--CoA ligase

365(5) WP_049720631.1 WP_049720631.1 hypothetical protein
365(1) WP_181731452.1 WP_181731452.1 hypothetical protein
365(2) WP_037993162.1 WP_037993162.1 hypothetical protein
365(1) WP_181737013.1 WP_181737013.1 hypothetical protein

366(1) WP_181737017.1 WP_181737017.1 hypothetical protein
366(1) WP_181731451.1 WP_181731451.1 hypothetical protein
366(4) WP_049720632.1 WP_049720632.1 hypothetical protein
366(3) WP_052186600.1 WP_052186600.1 hypothetical protein

367(1) WP_198050877.1 WP_198050877.1 hypothetical protein
367(4) WP_062843249.1 WP_062843249.1 hypothetical protein
367(1) WP_181731450.1 WP_181731450.1 hypothetical protein
367(1) WP_037993163.1 WP_037993163.1 hypothetical protein
367(1) WP_037993165.1 WP_037993165.1 hypothetical protein
367(1) WP_049720633.1 WP_049720633.1 hypothetical protein

368(5) WP_175413164.1 WP_175413164.1 PTS sugar transporter subunit IIB
368(1) WP_048826596.1 WP_048826596.1 PTS sugar transporter subunit IIB
368(2) WP_181729133.1 WP_181729133.1 PTS sugar transporter subunit IIB
368(1) WP_198053926.1 WP_198053926.1 PTS sugar transporter subunit IIB

369(1) WP_000780503.1 WP_000780503.1 PepSY domain-containing protein
369(2) WP_000780498.1 WP_000780498.1 PepSY domain-containing protein
369(1) WP_144611196.1 WP_144611196.1 PepSY domain-containing protein
369(1) WP_166705002.1 WP_166705002.1 PepSY domain-containing protein
369(1) WP_151630595.1 WP_151630595.1 PepSY domain-containing protein
369(1) WP_053564619.1 WP_053564619.1 PepSY domain-containing protein
369(1) WP_212959026.1 WP_212959026.1 PepSY domain-containing protein
369(1) WP_139867362.1 WP_139867362.1 PepSY domain-containing protein

370(1) WP_229522871.1 WP_229522871.1 four-helix bundle copper-binding protein
370(2) WP_010346302.1 WP_010346302.1 four-helix bundle copper-binding protein
370(1) WP_014281165.1 WP_014281165.1 four-helix bundle copper-binding protein
370(2) WP_007430114.1 WP_007430114.1 four-helix bundle copper-binding protein
370(1) WP_137062785.1 WP_137062785.1 four-helix bundle copper-binding protein
370(2) WP_138226039.1 WP_138226039.1 four-helix bundle copper-binding protein

371(3) WP_049720630.1 WP_049720630.1 thioesterase domain-containing protein
371(1) WP_181737011.1 WP_181737011.1 thioesterase
371(1) WP_232337630.1 WP_232337630.1 thioesterase domain-containing protein
371(1) WP_232335482.1 WP_232335482.1 thioesterase domain-containing protein
371(3) WP_037993161.1 WP_037993161.1 thioesterase domain-containing protein

372(1) WP_120112232.1 WP_120112232.1 hypothetical protein
372(1) WP_027088431.1 WP_027088431.1 hypothetical protein
372(1) WP_238187891.1 WP_238187891.1 hypothetical protein
372(1) WP_127570432.1 WP_127570432.1 hypothetical protein
372(1) WP_136372967.1 WP_136372967.1 hypothetical protein
372(1) WP_134750592.1 WP_134750592.1 hypothetical protein
372(1) WP_138755909.1 WP_138755909.1 hypothetical protein
372(1) WP_182301826.1 WP_182301826.1 hypothetical protein
372(1) WP_182301825.1 WP_182301825.1 hypothetical protein

373(1) WP_106496012.1 WP_106496012.1 UDP-N-acetylglucosamine 1-carboxyvinyltransferase
373(2) WP_133332969.1 WP_133332969.1 UDP-N-acetylglucosamine 1-carboxyvinyltransferase
373(1) WP_013658725.1 WP_013658725.1 UDP-N-acetylglucosamine 1-carboxyvinyltransferase
373(1) WP_191272885.1 WP_191272885.1 UDP-N-acetylglucosamine 1-carboxyvinyltransferase
373(1) WP_251466774.1 WP_251466774.1 UDP-N-acetylglucosamine 1-carboxyvinyltransferase
373(1) WP_026572730.1 WP_026572730.1 UDP-N-acetylglucosamine 1-carboxyvinyltransferase
373(1) WP_198310842.1 WP_198310842.1 UDP-N-acetylglucosamine 1-carboxyvinyltransferase
373(1) WP_210364437.1 WP_210364437.1 UDP-N-acetylglucosamine 1-carboxyvinyltransferase

374(1) WP_079506358.1 WP_079506358.1 stage II sporulation protein D
374(1) WP_026572731.1 WP_026572731.1 stage II sporulation protein D
374(2) WP_133332970.1 WP_133332970.1 stage II sporulation protein D
374(1) WP_041964592.1 WP_041964592.1 stage II sporulation protein D
374(1) WP_210364438.1 WP_210364438.1 stage II sporulation protein D
374(1) WP_191272883.1 WP_191272883.1 stage II sporulation protein D
374(1) WP_198310841.1 WP_198310841.1 stage II sporulation protein D
374(1) WP_251466776.1 WP_251466776.1 stage II sporulation protein D

375(1) WP_257675215.1 WP_257675215.1 hypothetical protein
375(1) WP_077833104.1 WP_077833104.1 hypothetical protein

| | | |
|---|---|---|
| 375(1) | WP_026882393.1 | WP_026882393.1 hypothetical protein |
| 375(1) | WP_066020841.1 | WP_066020841.1 hypothetical protein |
| 375(2) | WP_010963736.1 | WP_010963736.1 hypothetical protein |
| 375(1) | WP_066020836.1 | WP_066020836.1 hypothetical protein |
| 375(2) | WP_077852412.1 | WP_077852412.1 hypothetical protein |

| | | |
|---|---|---|
| 376(1) | WP_103047819.1 | WP_103047819.1 helix-turn-helix transcriptional regulator |
| 376(1) | WP_165175569.1 | WP_165175569.1 helix-turn-helix transcriptional regulator |
| 376(1) | WP_258296273.1 | WP_258296273.1 helix-turn-helix transcriptional regulator |
| 376(1) | WP_075154088.1 | WP_075154088.1 helix-turn-helix transcriptional regulator |
| 376(1) | WP_250244827.1 | WP_250244827.1 helix-turn-helix transcriptional regulator |
| 376(1) | WP_071558835.1 | WP_071558835.1 helix-turn-helix transcriptional regulator |
| 376(1) | WP_053325286.1 | WP_053325286.1 helix-turn-helix transcriptional regulator |
| 376(1) | WP_108464073.1 | WP_108464073.1 hypothetical protein |
| 376(1) | WP_190299141.1 | WP_190299141.1 helix-turn-helix transcriptional regulator |

| | | |
|---|---|---|
| 377(2) | WP_232335496.1 | WP_232335496.1 cation acetate symporter |
| 377(1) | WP_019120046.1 | WP_019120046.1 sodium:solute symporter family protein |
| 377(2) | WP_037993156.1 | WP_037993156.1 cation acetate symporter |
| 377(1) | WP_268983099.1 | WP_268983099.1 cation acetate symporter |
| 377(1) | WP_049720628.1 | WP_049720628.1 cation acetate symporter |
| 377(1) | WP_232335149.1 | WP_232335149.1 cation acetate symporter |
| 377(1) | WP_220214995.1 | WP_220214995.1 cation acetate symporter |

| | | |
|---|---|---|
| 378(1) | WP_139378315.1 | WP_139378315.1 hypothetical protein |
| 378(1) | WP_102264555.1 | WP_102264555.1 hypothetical protein |
| 378(1) | WP_214902944.1 | WP_214902944.1 hypothetical protein |
| 378(1) | WP_226086178.1 | WP_226086178.1 hypothetical protein |
| 378(1) | WP_052442080.1 | WP_052442080.1 hypothetical protein |
| 378(1) | WP_226642866.1 | WP_226642866.1 hypothetical protein |
| 378(2) | WP_144479074.1 | WP_144479074.1 hypothetical protein |
| 378(1) | WP_283904347.1 | WP_283904347.1 hypothetical protein |

| | | |
|---|---|---|
| 379(4) | WP_195431300.1 | WP_195431300.1 hypothetical protein |
| 379(1) | WP_287600019.1 | WP_287600019.1 hypothetical protein |
| 379(1) | WP_139796093.1 | WP_139796093.1 hypothetical protein |
| 379(1) | WP_181603067.1 | WP_181603067.1 hypothetical protein |
| 379(1) | WP_185165637.1 | WP_185165637.1 hypothetical protein |
| 379(1) | WP_143315244.1 | WP_143315244.1 hypothetical protein |

| | | |
|---|---|---|
| 380(1) | WP_288218584.1 | WP_288218584.1 hypothetical protein |
| 380(8) | WP_042401012.1 | WP_042401012.1 hypothetical protein |

| | | |
|---|---|---|
| 381(1) | WP_181603069.1 | WP_181603069.1 hypothetical protein |
| 381(1) | WP_291647947.1 | WP_291647947.1 hypothetical protein |
| 381(1) | WP_084117724.1 | WP_084117724.1 prepilin peptidase |
| 381(2) | WP_195431294.1 | WP_195431294.1 hypothetical protein |
| 381(1) | WP_185165635.1 | WP_185165635.1 prepilin peptidase |
| 381(1) | WP_271839931.1 | WP_271839931.1 hypothetical protein |
| 381(1) | WP_143315242.1 | WP_143315242.1 hypothetical protein |
| 381(1) | WP_287600014.1 | WP_287600014.1 hypothetical protein |

| | | |
|---|---|---|
| 382(1) | WP_084117725.1 | WP_084117725.1 hypothetical protein |
| 382(1) | WP_185165636.1 | WP_185165636.1 hypothetical protein |
| 382(1) | WP_181603068.1 | WP_181603068.1 RcpC/CpaB family pilus assembly protein |
| 382(2) | WP_195431297.1 | WP_195431297.1 hypothetical protein |
| 382(1) | WP_287600017.1 | WP_287600017.1 hypothetical protein |
| 382(1) | WP_291647949.1 | WP_291647949.1 hypothetical protein |
| 382(1) | WP_271839929.1 | WP_271839929.1 hypothetical protein |
| 382(1) | WP_143315243.1 | WP_143315243.1 hypothetical protein |

| | | |
|---|---|---|
| 383(2) | WP_195431303.1 | WP_195431303.1 DUF4320 family protein |
| 383(1) | WP_143315245.1 | WP_143315245.1 DUF4320 family protein |
| 383(1) | WP_287600022.1 | WP_287600022.1 DUF4320 family protein |
| 383(1) | WP_181603066.1 | WP_181603066.1 hypothetical protein |
| 383(1) | WP_185165638.1 | WP_185165638.1 hypothetical protein |
| 383(1) | WP_291647956.1 | WP_291647956.1 hypothetical protein |
| 383(1) | WP_271839925.1 | WP_271839925.1 DUF4320 family protein |
| 383(1) | WP_084117727.1 | WP_084117727.1 hypothetical protein |

| | | |
|---|---|---|
| 384(1) | WP_291647958.1 | WP_291647958.1 hypothetical protein |
| 384(2) | WP_195431306.1 | WP_195431306.1 hypothetical protein |
| 384(1) | WP_271839923.1 | WP_271839923.1 hypothetical protein |
| 384(1) | WP_181603065.1 | WP_181603065.1 hypothetical protein |
| 384(1) | WP_287600024.1 | WP_287600024.1 hypothetical protein |
| 384(1) | WP_143315246.1 | WP_143315246.1 hypothetical protein |
| 384(1) | WP_084117728.1 | WP_084117728.1 hypothetical protein |
| 384(1) | WP_185165639.1 | WP_185165639.1 hypothetical protein |

| | | |
|---|---|---|
| 385(2) | WP_271839920.1 | WP_271839920.1 hypothetical protein |
| 385(1) | WP_291647959.1 | WP_291647959.1 hypothetical protein |
| 385(2) | WP_195431309.1 | WP_195431309.1 hypothetical protein |
| 385(1) | WP_185165640.1 | WP_185165640.1 hypothetical protein |
| 385(1) | WP_143315247.1 | WP_143315247.1 type II secretion system F family protein |
| 385(1) | WP_181603064.1 | WP_181603064.1 type II secretion system F family protein |
| 385(1) | WP_176212752.1 | WP_176212752.1 type II secretion system F family protein |

| | | |
|---|---|---|
| 386(1) | WP_084117730.1 | WP_084117730.1 hypothetical protein |
| 386(2) | WP_271839918.1 | WP_271839918.1 hypothetical protein |
| 386(1) | WP_185165641.1 | WP_185165641.1 hypothetical protein |
| 386(1) | WP_181603063.1 | WP_181603063.1 hypothetical protein |
| 386(1) | WP_291647961.1 | WP_291647961.1 hypothetical protein |
| 386(2) | WP_195431312.1 | WP_195431312.1 hypothetical protein |
| 386(1) | WP_185966720.1 | WP_185966720.1 hypothetical protein |

| | | |
|---|---|---|
| 387(2) | WP_230322459.1 | WP_230322459.1 tyrosine-type recombinase/integrase |
| 387(1) | WP_294953126.1 | WP_294953126.1 site-specific integrase |
| 387(1) | WP_033521281.1 | WP_033521281.1 tyrosine recombinase XerC |
| 387(2) | WP_219872444.1 | WP_219872444.1 tyrosine recombinase XerS |
| 387(2) | WP_228463714.1 | WP_228463714.1 tyrosine-type recombinase/integrase |
| 387(1) | WP_098430133.1 | WP_098430133.1 tyrosine-type recombinase/integrase |

| | | |
|---|---|---|
| 388(2) | WP_283655691.1 | WP_283655691.1 type VII secretion protein EssC |
| 388(2) | WP_283655692.1 | WP_283655692.1 FtsK/SpoIIIE domain-containing protein |

388(1) WP_212330891.1 WP_212330891.1 hypothetical protein
388(1) WP_307388996.1 WP_307388996.1 FtsK/SpoIIIE domain-containing protein
388(1) WP_286458025.1 WP_286458025.1 hypothetical protein
388(1) WP_297035094.1 WP_297035094.1 hypothetical protein
388(1) WP_270472028.1 WP_270472028.1 hypothetical protein

389(1) WP_110931536.1 WP_110931536.1 hypothetical protein
389(2) WP_068782415.1 WP_068782415.1 hypothetical protein
389(1) WP_024632675.1 WP_024632675.1 hypothetical protein
389(1) WP_307394650.1 WP_307394650.1 hypothetical protein
389(1) WP_212981564.1 WP_212981564.1 hypothetical protein
389(1) WP_166244895.1 WP_166244895.1 WXG100 family type VII secretion target
389(1) WP_055105798.1 WP_055105798.1 hypothetical protein
389(1) WP_211023031.1 WP_211023031.1 hypothetical protein

390(1) WP_116353262.1 WP_116353262.1 galactokinase
390(1) WP_077619661.1 WP_077619661.1 galactokinase
390(1) WP_010676875.1 WP_010676875.1 galactokinase
390(1) WP_099354894.1 WP_099354894.1 galactokinase
390(1) WP_309098604.1 WP_309098604.1 galactokinase
390(1) WP_229139910.1 WP_229139910.1 galactokinase
390(1) WP_304216851.1 WP_304216851.1 galactokinase
390(1) WP_034526663.1 WP_034526663.1 galactokinase
390(1) WP_099361329.1 WP_099361329.1 galactokinase

391(1) WP_136379684.1 WP_136379684.1 aldose epimerase family protein
391(1) WP_099354896.1 WP_099354896.1 aldose epimerase family protein
391(1) WP_077619658.1 WP_077619658.1 galactose mutarotase
391(1) WP_229139913.1 WP_229139913.1 galactose mutarotase
391(1) WP_309098607.1 WP_309098607.1 aldose epimerase family protein
391(1) WP_304216857.1 WP_304216857.1 aldose epimerase family protein
391(1) WP_010676872.1 WP_010676872.1 aldose epimerase family protein
391(1) WP_116353259.1 WP_116353259.1 aldose epimerase family protein
391(1) WP_099361332.1 WP_099361332.1 aldose epimerase family protein

392(1) WP_312853822.1 WP_312853822.1 DUF488 domain-containing protein
392(1) WP_312853823.1 WP_312853823.1 hypothetical protein
392(4) WP_037993177.1 WP_037993177.1 DUF488 family protein
392(2) WP_049720637.1 WP_049720637.1 DUF488 family protein
392(1) WP_198053927.1 WP_198053927.1 DUF488 family protein

393(1) WP_181731439.1 WP_181731439.1 L-lactate dehydrogenase
393(1) WP_181737033.1 WP_181737033.1 L-lactate dehydrogenase
393(5) WP_049720640.1 WP_049720640.1 L-lactate dehydrogenase
393(1) WP_121874299.1 WP_121874299.1 L-lactate dehydrogenase

394(8) WP_185463136.1 WP_185463136.1 hypothetical protein

395(7) WP_037993231.1 WP_037993231.1 DUF485 domain-containing protein
395(1) WP_198053919.1 WP_198053919.1 DUF485 domain-containing protein

396(1) WP_198053920.1 WP_198053920.1 cation:proton antiporter
396(7) WP_049720629.1 WP_049720629.1 cation:proton antiporter

397(1) WP_198053929.1 WP_198053929.1 lactate permease LctP family transporter
397(3) WP_138614838.1 WP_138614838.1 lactate permease LctP family transporter
397(1) WP_087194497.1 WP_087194497.1 L-lactate permease
397(2) WP_121874300.1 WP_121874300.1 lactate permease LctP family transporter
397(1) WP_049720639.1 WP_049720639.1 lactate permease LctP family transporter

398(2) WP_046217699.1 WP_046217699.1 PTS mannose/fructose/sorbose/N-acetylgalactosamine transporter subunit IIC
398(2) WP_206765229.1 WP_206765229.1 PTS mannose/fructose/sorbose/N-acetylgalactosamine transporter subunit IIC
398(4) WP_077833123.1 WP_077833123.1 PTS mannose/fructose/sorbose transporter subunit IIC

399(2) WP_003516591.1 WP_003516591.1 homocysteine S-methyltransferase family protein
399(1) WP_091260886.1 WP_091260886.1 methionine synthase
399(1) WP_156424117.1 WP_156424117.1 methionine synthase
399(1) WP_069193863.1 WP_069193863.1 homocysteine S-methyltransferase family protein
399(1) WP_038290966.1 WP_038290966.1 homocysteine S-methyltransferase family protein
399(1) WP_220559126.1 WP_220559126.1 bifunctional homocysteine S-methyltransferase/methylenetetrahydrofolate reductase
399(1) WP_091260871.1 WP_091260871.1 bifunctional homocysteine S-methyltransferase/methylenetetrahydrofolate reductase

400(5) WP_049720627.1 WP_049720627.1 DUF2243 domain-containing protein
400(1) WP_232337629.1 WP_232337629.1 DUF2243 domain-containing protein
400(2) WP_232335118.1 WP_232335118.1 DUF2243 domain-containing protein

401(2) WP_077833113.1 WP_077833113.1 hypothetical protein
401(1) WP_066020847.1 WP_066020847.1 hypothetical protein
401(1) WP_234121611.1 WP_234121611.1 hypothetical protein
401(2) WP_010963727.1 WP_010963727.1 hypothetical protein
401(2) WP_077893583.1 WP_077893583.1 hypothetical protein

402(2) WP_036702435.1 WP_036702435.1 deoxynucleoside kinase
402(1) WP_238655335.1 WP_238655335.1 deoxynucleoside kinase
402(1) WP_238655336.1 WP_238655336.1 deoxynucleoside kinase
402(1) WP_076157850.1 WP_076157850.1 deoxynucleoside kinase
402(1) WP_189031053.1 WP_189031053.1 deoxynucleoside kinase
402(1) WP_036702438.1 WP_036702438.1 deoxynucleoside kinase
402(1) WP_189031052.1 WP_189031052.1 deoxynucleoside kinase

403(1) WP_250944678.1 WP_250944678.1 hypothetical protein
403(2) WP_010963731.1 WP_010963731.1 hypothetical protein
403(1) WP_242874735.1 WP_242874735.1 hypothetical protein
403(3) WP_242951946.1 WP_242951946.1 hypothetical protein
403(1) WP_242841131.1 WP_242841131.1 hypothetical protein

404(1) WP_251457588.1 WP_251457588.1 PTS system mannose/fructose/sorbose family transporter subunit IID
404(4) WP_077833122.1 WP_077833122.1 PTS mannose transporter subunit IID

404(2) WP_046217959.1 WP_046217959.1 PTS system mannose/fructose/sorbose family transporter subunit IID
404(1) WP_197235249.1 WP_197235249.1 PTS system mannose/fructose/sorbose family transporter subunit IID

405(1) WP_026593093.1 WP_026593093.1 hypothetical protein
405(1) WP_108464097.1 WP_108464097.1 hypothetical protein
405(1) WP_253440778.1 WP_253440778.1 hypothetical protein
405(1) WP_071719978.1 WP_071719978.1 hypothetical protein
405(1) WP_244525355.1 WP_244525355.1 hypothetical protein
405(2) WP_010963722.1 WP_010963722.1 hypothetical protein
405(1) WP_039871677.1 WP_039871677.1 hypothetical protein

406(1) WP_083180280.1 WP_083180280.1 helix-turn-helix transcriptional regulator
406(1) WP_021291839.1 WP_021291839.1 helix-turn-helix transcriptional regulator
406(1) WP_074048563.1 WP_074048563.1 helix-turn-helix transcriptional regulator
406(1) WP_074598755.1 WP_074598755.1 helix-turn-helix transcriptional regulator
406(1) WP_253440783.1 WP_253440783.1 helix-turn-helix transcriptional regulator
406(1) WP_211082932.1 WP_211082932.1 helix-turn-helix transcriptional regulator
406(1) WP_160807256.1 WP_160807256.1 helix-turn-helix transcriptional regulator
406(1) WP_079506418.1 WP_079506418.1 helix-turn-helix transcriptional regulator

407(1) WP_192510487.1 WP_192510487.1 hypothetical protein
407(1) WP_063208770.1 WP_063208770.1 hypothetical protein
407(1) WP_229522850.1 WP_229522850.1 hypothetical protein
407(1) WP_068940686.1 WP_068940686.1 hypothetical protein
407(1) WP_068496241.1 WP_068496241.1 hypothetical protein
407(1) WP_209941817.1 WP_209941817.1 hypothetical protein
407(1) WP_254325791.1 WP_254325791.1 hypothetical protein
407(1) WP_010346272.1 WP_010346272.1 hypothetical protein

408(1) WP_048826590.1 WP_048826590.1 transketolase
408(1) WP_232274498.1 WP_232274498.1 transketolase
408(1) WP_254759282.1 WP_254759282.1 transketolase
408(1) WP_102710733.1 WP_102710733.1 alpha-ketoacid dehydrogenase subunit beta
408(1) WP_048826592.1 WP_048826592.1 transketolase
408(1) WP_195953741.1 WP_195953741.1 transketolase family protein
408(1) WP_195953768.1 WP_195953768.1 transketolase
408(1) WP_254759283.1 WP_254759283.1 transketolase C-terminal domain-containing protein

409(1) WP_077832296.1 WP_077832296.1 hypothetical protein
409(2) WP_010963379.1 WP_010963379.1 hypothetical protein
409(2) WP_010963380.1 WP_010963380.1 hypothetical protein
409(1) WP_257675923.1 WP_257675923.1 hypothetical protein
409(2) WP_010963372.1 WP_010963372.1 hypothetical protein

410(2) WP_206759992.1 WP_206759992.1 cytochrome c oxidase assembly protein
410(1) WP_232335495.1 WP_232335495.1 cytochrome c oxidase assembly protein
410(1) WP_198053918.1 WP_198053918.1 cytochrome c oxidase assembly protein
410(3) WP_037993154.1 WP_037993154.1 cytochrome c oxidase assembly protein
410(1) WP_268983098.1 WP_268983098.1 cytochrome c oxidase assembly protein

411(2) WP_198050882.1 WP_198050882.1 non-oxidative hydroxyarylic acid decarboxylases subunit B
411(4) WP_037997344.1 WP_037997344.1 non-oxidative hydroxyarylic acid decarboxylases subunit B
411(1) WP_269747554.1 WP_269747554.1 non-oxidative hydroxyarylic acid decarboxylases subunit B
411(1) WP_181603053.1 WP_181603053.1 UbiX family flavin prenyltransferase

412(1) WP_136790003.1 WP_136790003.1 exopolysaccharide biosynthesis polyprenyl glycosylphosphotransferase
412(1) WP_243495568.1 WP_243495568.1 sugar transferase
412(2) WP_005210474.1 WP_005210474.1 sugar transferase
412(1) WP_098579108.1 WP_098579108.1 sugar transferase
412(2) WP_251426565.1 WP_251426565.1 sugar transferase
412(1) WP_272801828.1 WP_272801828.1 exopolysaccharide biosynthesis polyprenyl glycosylphosphotransferase

413(1) WP_251598131.1 WP_251598131.1 transcriptional regulator
413(1) WP_216791255.1 WP_216791255.1 transcriptional regulator
413(1) WP_010496628.1 WP_010496628.1 hypothetical protein
413(1) WP_127459457.1 WP_127459457.1 transcriptional regulator
413(1) WP_108534973.1 WP_108534973.1 transcriptional regulator
413(1) WP_253411110.1 WP_253411110.1 transcriptional regulator
413(1) WP_036689488.1 WP_036689488.1 hypothetical protein
413(1) WP_281937338.1 WP_281937338.1 transcriptional regulator

414(1) WP_281937340.1 WP_281937340.1 serine hydrolase
414(1) WP_163768035.1 WP_163768035.1 serine hydrolase
414(1) WP_036689486.1 WP_036689486.1 serine hydrolase
414(1) WP_253411108.1 WP_253411108.1 serine hydrolase
414(1) WP_189597466.1 WP_189597466.1 serine hydrolase
414(1) WP_251598132.1 WP_251598132.1 serine hydrolase
414(1) WP_216791252.1 WP_216791252.1 serine hydrolase
414(1) WP_010496627.1 WP_010496627.1 serine hydrolase

415(1) WP_127459453.1 WP_127459453.1 permease
415(2) WP_036689482.1 WP_036689482.1 hypothetical protein
415(1) WP_108534971.1 WP_108534971.1 Na+/H+ antiporter NhaC family protein
415(1) WP_010496626.1 WP_010496626.1 hypothetical protein
415(1) WP_281937342.1 WP_281937342.1 hypothetical protein
415(1) WP_251598133.1 WP_251598133.1 hypothetical protein
415(1) WP_253411106.1 WP_253411106.1 hypothetical protein

416(1) WP_216791197.1 WP_216791197.1 hypothetical protein
416(3) WP_063188078.1 WP_063188078.1 hypothetical protein
416(1) WP_127484363.1 WP_127484363.1 hypothetical protein
416(1) WP_108534949.1 WP_108534949.1 hypothetical protein
416(1) WP_163853482.1 WP_163853482.1 hypothetical protein
416(1) WP_281937384.1 WP_281937384.1 hypothetical protein

417(1) WP_268893545.1 WP_268893545.1 transposase
417(1) WP_227779377.1 WP_227779377.1 transposase
417(1) WP_053432792.1 WP_053432792.1 IS1182 family transposase
417(1) WP_283892160.1 WP_283892160.1 IS1182 family transposase
417(1) WP_197260337.1 WP_197260337.1 transposase, partial

417(1) WP_227779350.1 WP_227779350.1 hypothetical protein
417(1) WP_247081727.1 WP_247081727.1 IS1182 family transposase
417(1) WP_236146857.1 WP_236146857.1 hypothetical protein

418(1) WP_293755506.1 WP_293755506.1 hypothetical protein
418(2) WP_281813118.1 WP_281813118.1 hypothetical protein
418(4) WP_113672711.1 WP_113672711.1 hypothetical protein
418(1) WP_212695795.1 WP_212695795.1 hypothetical protein

419(1) WP_164988931.1 WP_164988931.1 hypothetical protein
419(3) WP_283892183.1 WP_283892183.1 hypothetical protein
419(1) WP_139168458.1 WP_139168458.1 hypothetical protein
419(1) WP_301167504.1 WP_301167504.1 hypothetical protein
419(1) WP_156412604.1 WP_156412604.1 hypothetical protein
419(1) WP_116556141.1 WP_116556141.1 hypothetical protein

420(1) WP_303942553.1 WP_303942553.1 prolipoprotein diacylglyceryl transferase
420(1) WP_273383891.1 WP_273383891.1 prolipoprotein diacylglyceryl transferase
420(1) WP_297183861.1 WP_297183861.1 prolipoprotein diacylglyceryl transferase, partial
420(1) WP_291289945.1 WP_291289945.1 prolipoprotein diacylglyceryl transferase
420(2) WP_291288250.1 WP_291288250.1 prolipoprotein diacylglyceryl transferase
420(1) WP_013490145.1 WP_013490145.1 prolipoprotein diacylglyceryl transferase
420(1) WP_102372739.1 WP_102372739.1 prolipoprotein diacylglyceryl transferase

421(1) WP_304942556.1 WP_304942556.1 hypothetical protein
421(1) WP_113672709.1 WP_113672709.1 hypothetical protein
421(2) WP_281813113.1 WP_281813113.1 hypothetical protein
421(2) WP_212693244.1 WP_212693244.1 hypothetical protein
421(1) WP_293755511.1 WP_293755511.1 hypothetical protein
421(1) WP_212695793.1 WP_212695793.1 hypothetical protein

422(1) WP_305022918.1 WP_305022918.1 helix-turn-helix domain-containing protein
422(1) WP_305755029.1 WP_305755029.1 helix-turn-helix domain-containing protein
422(1) WP_236334712.1 WP_236334712.1 helix-turn-helix domain-containing protein
422(2) WP_198056468.1 WP_198056468.1 helix-turn-helix domain-containing protein
422(1) WP_128631273.1 WP_128631273.1 winged helix-turn-helix transcriptional regulator
422(2) WP_039295412.1 WP_039295412.1 helix-turn-helix domain-containing protein

423(1) WP_243518777.1 WP_243518777.1 ribose 5-phosphate isomerase B
423(1) WP_307192910.1 WP_307192910.1 ribose 5-phosphate isomerase B
423(2) WP_019392127.1 WP_019392127.1 ribose 5-phosphate isomerase B
423(1) WP_124049158.1 WP_124049158.1 ribose 5-phosphate isomerase B
423(1) WP_254759289.1 WP_254759289.1 ribose 5-phosphate isomerase B
423(1) WP_099223958.1 WP_099223958.1 ribose 5-phosphate isomerase B
423(1) WP_268583844.1 WP_268583844.1 ribose 5-phosphate isomerase B

424(1) WP_289613148.1 WP_289613148.1 hypothetical protein
424(2) WP_013252389.1 WP_013252389.1 hypothetical protein
424(1) WP_156963125.1 WP_156963125.1 hypothetical protein
424(1) WP_036575531.1 WP_036575531.1 hypothetical protein
424(1) WP_040651298.1 WP_040651298.1 hypothetical protein
424(1) WP_307388969.1 WP_307388969.1 hypothetical protein
424(1) WP_282742035.1 WP_282742035.1 hypothetical protein

425(1) WP_138755923.1 WP_138755923.1 hypothetical protein
425(1) WP_127570440.1 WP_127570440.1 hypothetical protein
425(1) WP_120112267.1 WP_120112267.1 hypothetical protein
425(1) WP_134750610.1 WP_134750610.1 hypothetical protein
425(1) WP_248565447.1 WP_248565447.1 hypothetical protein
425(1) WP_136372953.1 WP_136372953.1 hypothetical protein
425(1) WP_027088422.1 WP_027088422.1 hypothetical protein
425(1) WP_307601022.1 WP_307601022.1 hypothetical protein

426(1) WP_309098606.1 WP_309098606.1 UDP-glucose--hexose-1-phosphate uridylyltransferase
426(1) WP_304216855.1 WP_304216855.1 UDP-glucose--hexose-1-phosphate uridylyltransferase
426(1) WP_010676873.1 WP_010676873.1 UDP-glucose--hexose-1-phosphate uridylyltransferase
426(1) WP_234987243.1 WP_234987243.1 UDP-glucose--hexose-1-phosphate uridylyltransferase
426(1) WP_229139912.1 WP_229139912.1 UDP-glucose--hexose-1-phosphate uridylyltransferase
426(1) WP_099354944.1 WP_099354944.1 UDP-glucose--hexose-1-phosphate uridylyltransferase
426(1) WP_116353260.1 WP_116353260.1 UDP-glucose--hexose-1-phosphate uridylyltransferase
426(1) WP_180271142.1 WP_180271142.1 UDP-glucose--hexose-1-phosphate uridylyltransferase

427(1) WP_127199760.1 WP_127199760.1 DUF3939 domain-containing protein
427(1) WP_094092610.1 WP_094092610.1 DUF3939 domain-containing protein
427(1) WP_306432892.1 WP_306432892.1 DUF3939 domain-containing protein
427(1) WP_209861657.1 WP_209861657.1 DUF3939 domain-containing protein
427(1) WP_108465342.1 WP_108465342.1 DUF3939 domain-containing protein
427(1) WP_068694702.1 WP_068694702.1 DUF3939 domain-containing protein
427(1) WP_223067449.1 WP_223067449.1 DUF3939 domain-containing protein
427(1) WP_316245811.1 WP_316245811.1 DUF3939 domain-containing protein

428(1) WP_066020833.1 WP_066020833.1 arsenate reductase family protein
428(4) WP_077833101.1 WP_077833101.1 arsenate reductase family protein
428(2) WP_010963739.1 WP_010963739.1 ArsC/Spx/MgsR family protein

429(1) WP_071721669.1 WP_071721669.1 glycosyltransferase
429(1) WP_098139277.1 WP_098139277.1 glycosyltransferase
429(1) WP_066020855.1 WP_066020855.1 glycosyltransferase
429(3) WP_097954762.1 WP_097954762.1 glycosyltransferase
429(1) WP_070173063.1 WP_070173063.1 glycosyltransferase

430(1) WP_021253109.1 WP_021253109.1 DUF4184 family protein
430(1) WP_072727830.1 WP_072727830.1 DUF4184 family protein
430(1) WP_138186684.1 WP_138186684.1 DUF4184 family protein
430(1) WP_065291240.1 WP_065291240.1 DUF4184 family protein
430(2) WP_028533175.1 WP_028533175.1 DUF4184 family protein
430(1) WP_021261874.1 WP_021261874.1 DUF4184 family protein

431(3) WP_085499129.1 WP_085499129.1 hypothetical protein
431(1) WP_076157810.1 WP_076157810.1 hypothetical protein
431(1) WP_152392587.1 WP_152392587.1 hypothetical protein
431(2) WP_085499073.1 WP_085499073.1 hypothetical protein

432(5) WP_037997350.1 WP_037997350.1 hypothetical protein

432(1) WP_121874298.1 WP_121874298.1 hypothetical protein
432(1) WP_181731437.1 WP_181731437.1 hypothetical protein

433(3) WP_049720643.1 WP_049720643.1 DNA polymerase IV
433(3) WP_037997352.1 WP_037997352.1 DNA polymerase IV
433(1) WP_181731438.1 WP_181731438.1 DNA polymerase IV

434(1) WP_195244538.1 WP_195244538.1 hypothetical protein
434(1) WP_156327470.1 WP_156327470.1 hypothetical protein
434(1) WP_005210462.1 WP_005210462.1 hypothetical protein
434(1) WP_196048856.1 WP_196048856.1 hypothetical protein
434(3) WP_156327845.1 WP_156327845.1 hypothetical protein

435(6) WP_037997346.1 WP_037997346.1 hemolysin III family protein
435(1) WP_198053930.1 WP_198053930.1 hemolysin III family protein

436(3) WP_026572751.1 WP_026572751.1 3-hydroxyacyl-ACP dehydratase FabZ
436(1) WP_068673743.1 WP_068673743.1 3-hydroxyacyl-ACP dehydratase FabZ
436(1) WP_198310821.1 WP_198310821.1 3-hydroxyacyl-ACP dehydratase FabZ
436(1) WP_046522794.1 WP_046522794.1 3-hydroxyacyl-ACP dehydratase FabZ
436(1) WP_053361013.1 WP_053361013.1 3-hydroxyacyl-ACP dehydratase FabZ

437(2) WP_087433598.1 WP_087433598.1 EutN/CcmL family microcompartment protein
437(1) WP_206094039.1 WP_206094039.1 EutN/CcmL family microcompartment protein
437(2) WP_112331596.1 WP_112331596.1 EutN/CcmL family microcompartment protein
437(2) WP_036705509.1 WP_036705509.1 EutN/CcmL family microcompartment protein

438(1) WP_099354918.1 WP_099354918.1 metallophosphoesterase
438(1) WP_116353241.1 WP_116353241.1 lamin tail domain-containing protein
438(1) WP_077619636.1 WP_077619636.1 metallophosphoesterase
438(1) WP_222861114.1 WP_222861114.1 LamG-like jellyroll fold domain-containing protein
438(1) WP_099361347.1 WP_099361347.1 metallophosphoesterase
438(1) WP_136379708.1 WP_136379708.1 lamin tail domain-containing protein
438(1) WP_010676851.1 WP_010676851.1 lamin tail domain-containing protein

439(3) WP_176392821.1 WP_176392821.1 phosphate propanoyltransferase
439(1) WP_233697167.1 WP_233697167.1 phosphate propanoyltransferase
439(2) WP_112331597.1 WP_112331597.1 phosphate propanoyltransferase
439(1) WP_051004551.1 WP_051004551.1 phosphate propanoyltransferase

440(1) WP_206094031.1 WP_206094031.1 hypothetical protein
440(1) WP_019424139.1 WP_019424139.1 hypothetical protein
440(1) WP_235286211.1 WP_235286211.1 hypothetical protein
440(2) WP_087433613.1 WP_087433613.1 hypothetical protein
440(1) WP_233697181.1 WP_233697181.1 hypothetical protein
440(1) WP_111147791.1 WP_111147791.1 hypothetical protein

441(1) WP_198053931.1 WP_198053931.1 DUF4309 domain-containing protein
441(1) WP_236533775.1 WP_236533775.1 DUF4309 domain-containing protein
441(2) WP_121874296.1 WP_121874296.1 DUF4309 domain-containing protein
441(1) WP_084712947.1 WP_084712947.1 DUF4309 domain-containing protein
441(2) WP_082240755.1 WP_082240755.1 DUF4309 domain-containing protein

442(5) WP_242833267.1 WP_242833267.1 hypothetical protein
442(2) WP_005210461.1 WP_005210461.1 hypothetical protein

443(1) WP_209861623.1 WP_209861623.1 PRD domain-containing protein
443(1) WP_251375539.1 WP_251375539.1 PRD domain-containing protein
443(1) WP_068619091.1 WP_068619091.1 transcription antiterminator
443(1) WP_127199782.1 WP_127199782.1 PRD domain-containing protein
443(2) WP_068694726.1 WP_068694726.1 PRD domain-containing protein
443(1) WP_213618825.1 WP_213618825.1 transcription antiterminator

444(1) WP_251456508.1 WP_251456508.1 hypothetical protein
444(2) WP_239695517.1 WP_239695517.1 hypothetical protein
444(3) WP_061805373.1 WP_061805373.1 hypothetical protein
444(1) WP_242447699.1 WP_242447699.1 hypothetical protein

445(2) WP_235824708.1 WP_235824708.1 YwmB family TATA-box binding protein
445(1) WP_251466772.1 WP_251466772.1 YwmB family TATA-box binding protein
445(1) WP_026572729.1 WP_026572729.1 YwmB family TATA-box binding protein
445(1) WP_232490091.1 WP_232490091.1 YwmB family TATA-box binding protein
445(1) WP_191272887.1 WP_191272887.1 YwmB family TATA-box binding protein
445(1) WP_210364436.1 WP_210364436.1 YwmB family TATA-box binding protein

446(1) WP_253806481.1 WP_253806481.1 hypothetical protein
446(1) WP_155617167.1 WP_155617167.1 hypothetical protein
446(1) WP_196232760.1 WP_196232760.1 hypothetical protein
446(1) WP_013370795.1 WP_013370795.1 transposase
446(1) WP_148266402.1 WP_148266402.1 IS110 family transposase
446(1) WP_170973160.1 WP_170973160.1 transposase
446(1) WP_155742209.1 WP_155742209.1 hypothetical protein

447(1) WP_250930784.1 WP_250930784.1 hypothetical protein
447(1) WP_077893575.1 WP_077893575.1 hypothetical protein
447(1) WP_066020830.1 WP_066020830.1 hypothetical protein
447(1) WP_250931530.1 WP_250931530.1 hypothetical protein
447(1) WP_257675211.1 WP_257675211.1 hypothetical protein
447(2) WP_010963741.1 WP_010963741.1 hypothetical protein

448(1) WP_248251173.1 WP_248251173.1 hypothetical protein
448(1) WP_256227704.1 WP_256227704.1 hypothetical protein
448(1) WP_062239906.1 WP_062239906.1 hypothetical protein
448(1) WP_265408856.1 WP_265408856.1 hypothetical protein
448(1) WP_062239900.1 WP_062239900.1 hypothetical protein
448(1) WP_256227706.1 WP_256227706.1 hypothetical protein
448(1) WP_265408854.1 WP_265408854.1 hypothetical protein

449(1) WP_267903267.1 WP_267903267.1 Imm6 family immunity protein
449(1) WP_113753103.1 WP_113753103.1 Imm6 family immunity protein
449(1) WP_113765178.1 WP_113765178.1 Imm6 family immunity protein
449(1) WP_113713150.1 WP_113713150.1 Imm6 family immunity protein
449(1) WP_076157812.1 WP_076157812.1 Imm6 family immunity protein

449(1) WP_151585727.1 WP_151585727.1 Imm6 family immunity protein
449(1) WP_208752237.1 WP_208752237.1 Imm6 family immunity protein

450(1) WP_263324533.1 WP_263324533.1 2-keto-3-deoxygluconate transporter
450(2) WP_120046149.1 WP_120046149.1 2-keto-3-deoxygluconate permease
450(1) WP_124049161.1 WP_124049161.1 2-keto-3-deoxygluconate permease
450(1) WP_268583847.1 WP_268583847.1 2-keto-3-deoxygluconate permease
450(1) WP_243518771.1 WP_243518771.1 2-keto-3-deoxygluconate permease
450(1) WP_019392124.1 WP_019392124.1 2-keto-3-deoxygluconate permease

451(1) WP_007430138.1 WP_007430138.1 aspartyl-phosphate phosphatase Spo0E family protein
451(1) WP_081047516.1 WP_081047516.1 aspartyl-phosphate phosphatase Spo0E family protein
451(1) WP_142613007.1 WP_142613007.1 aspartyl-phosphate phosphatase Spo0E family protein
451(3) WP_071558834.1 WP_071558834.1 aspartyl-phosphate phosphatase Spo0E family protein
451(1) WP_279628190.1 WP_279628190.1 hypothetical protein

452(1) WP_280788656.1 WP_280788656.1 LysE family transporter
452(3) WP_076279386.1 WP_076279386.1 LysE family transporter
452(3) WP_076135814.1 WP_076135814.1 LysE family translocator

453(1) WP_206106195.1 WP_206106195.1 carbamate kinase
453(1) WP_220084704.1 WP_220084704.1 carbamate kinase
453(1) WP_197256888.1 WP_197256888.1 carbamate kinase
453(1) WP_213468106.1 WP_213468106.1 carbamate kinase
453(1) WP_040729704.1 WP_040729704.1 carbamate kinase
453(1) WP_168178799.1 WP_168178799.1 carbamate kinase
453(1) WP_293654347.1 WP_293654347.1 carbamate kinase

454(1) WP_212695799.1 WP_212695799.1 hypothetical protein
454(1) WP_293755496.1 WP_293755496.1 hypothetical protein
454(1) WP_113672715.1 WP_113672715.1 hypothetical protein
454(1) WP_212693249.1 WP_212693249.1 hypothetical protein
454(2) WP_281813128.1 WP_281813128.1 hypothetical protein
454(1) WP_273321526.1 WP_273321526.1 hypothetical protein

455(1) WP_251548776.1 WP_251548776.1 MerR family transcriptional regulator
455(1) WP_068694720.1 WP_068694720.1 MerR family transcriptional regulator
455(2) WP_295628296.1 WP_295628296.1 MerR family transcriptional regulator
455(1) WP_039306363.1 WP_039306363.1 MerR family transcriptional regulator
455(1) WP_256718229.1 WP_256718229.1 MerR family transcriptional regulator
455(1) WP_209872750.1 WP_209872750.1 MerR family transcriptional regulator

456(1) WP_195974882.1 WP_195974882.1 helix-turn-helix transcriptional regulator
456(1) WP_204445215.1 WP_204445215.1 helix-turn-helix transcriptional regulator
456(1) WP_112331593.1 WP_112331593.1 helix-turn-helix transcriptional regulator
456(1) WP_257497518.1 WP_257497518.1 hypothetical protein
456(1) WP_187374155.1 WP_187374155.1 hypothetical protein
456(1) WP_295632018.1 WP_295632018.1 helix-turn-helix transcriptional regulator
456(1) WP_295628302.1 WP_295628302.1 helix-turn-helix transcriptional regulator

457(1) WP_110893002.1 WP_110893002.1 hypothetical protein
457(1) WP_305022940.1 WP_305022940.1 hypothetical protein
457(1) WP_169505040.1 WP_169505040.1 hypothetical protein
457(1) WP_251375560.1 WP_251375560.1 hypothetical protein
457(1) WP_209861673.1 WP_209861673.1 hypothetical protein
457(2) WP_138226038.1 WP_138226038.1 hypothetical protein

458(1) WP_307349991.1 WP_307349991.1 NAD(P)-binding domain-containing protein
458(1) WP_053403775.1 WP_053403775.1 NADP-dependent phosphogluconate dehydrogenase
458(1) WP_204517530.1 WP_204517530.1 NAD(P)-dependent oxidoreductase
458(1) WP_232274500.1 WP_232274500.1 NAD(P)-dependent oxidoreductase
458(2) WP_242172360.1 WP_242172360.1 NADP-dependent phosphogluconate dehydrogenase
458(1) WP_216390035.1 WP_216390035.1 NAD(P)-dependent oxidoreductase

459(1) WP_307589581.1 WP_307589581.1 MetQ/NlpA family ABC transporter substrate-binding protein
459(1) WP_179037804.1 WP_179037804.1 MetQ/NlpA family ABC transporter substrate-binding protein
459(5) WP_291288258.1 WP_291288258.1 MetQ/NlpA family ABC transporter substrate-binding protein

460(1) WP_307589603.1 WP_307589603.1 hypothetical protein
460(1) WP_183600898.1 WP_183600898.1 hypothetical protein
460(1) WP_038584732.1 WP_038584732.1 hypothetical protein
460(1) WP_232274496.1 WP_232274496.1 hypothetical protein
460(1) WP_111147806.1 WP_111147806.1 NPCBM/NEW2 domain-containing protein
460(1) WP_076113452.1 WP_076113452.1 hypothetical protein
460(1) WP_235286197.1 WP_235286197.1 hypothetical protein

461(1) WP_127570448.1 WP_127570448.1 hypothetical protein
461(1) WP_136372945.1 WP_136372945.1 hypothetical protein
461(1) WP_027088415.1 WP_027088415.1 hypothetical protein
461(1) WP_307601003.1 WP_307601003.1 hypothetical protein
461(1) WP_182301850.1 WP_182301850.1 hypothetical protein
461(1) WP_239617487.1 WP_239617487.1 hypothetical protein
461(1) WP_134750634.1 WP_134750634.1 hypothetical protein

462(1) WP_045518274.1 WP_045518274.1 hypothetical protein
462(1) WP_066091978.1 WP_066091978.1 hypothetical protein
462(1) WP_308110110.1 WP_308110110.1 hypothetical protein
462(1) WP_149871672.1 WP_149871672.1 hypothetical protein
462(1) WP_024027639.1 WP_024027639.1 hypothetical protein
462(1) WP_278187279.1 WP_278187279.1 hypothetical protein
462(1) WP_007083299.1 WP_007083299.1 hypothetical protein

463(1) WP_291289946.1 WP_291289946.1 NifU family protein
463(1) WP_308615629.1 WP_308615629.1 NifU family protein
463(1) WP_291288249.1 WP_291288249.1 NifU family protein
463(1) WP_303942552.1 WP_303942552.1 NifU family protein
463(1) WP_273383889.1 WP_273383889.1 NifU family protein
463(2) WP_102372740.1 WP_102372740.1 NifU family protein

464(1) WP_006292339.1 WP_006292339.1 hypothetical protein
464(1) WP_007147995.1 WP_007147995.1 hypothetical protein

464(1) WP_006290896.1 WP_006290896.1 hypothetical protein
464(2) WP_006289750.1 WP_006289750.1 hypothetical protein
464(1) WP_034526619.1 WP_034526619.1 hypothetical protein

465(1) WP_006292352.1 WP_006292352.1 hypothetical protein
465(1) WP_007147981.1 WP_007147981.1 hypothetical protein
465(1) WP_050752348.1 WP_050752348.1 hypothetical protein
465(1) WP_007147983.1 WP_007147983.1 hypothetical protein
465(1) WP_006292353.1 WP_006292353.1 hypothetical protein
465(1) WP_007147982.1 WP_007147982.1 YcxB family protein

466(1) WP_000540426.1 WP_000540426.1 YjcZ family sporulation protein
466(1) WP_077320197.1 WP_077320197.1 YjcZ family sporulation protein
466(4) WP_021291856.1 WP_021291856.1 YjcZ family sporulation protein

467(1) WP_098508797.1 WP_098508797.1 hypothetical protein
467(4) WP_090976435.1 WP_090976435.1 hypothetical protein
467(1) WP_098562449.1 WP_098562449.1 hypothetical protein

468(2) WP_019687035.1 WP_019687035.1 hypothetical protein
468(3) WP_019687037.1 WP_019687037.1 hypothetical protein
468(1) WP_113050540.1 WP_113050540.1 hypothetical protein

469(1) WP_120046146.1 WP_120046146.1 hypothetical protein
469(1) WP_142674258.1 WP_142674258.1 hypothetical protein
469(1) WP_151585724.1 WP_151585724.1 hypothetical protein
469(1) WP_142673092.1 WP_142673092.1 hypothetical protein
469(1) WP_124049166.1 WP_124049166.1 hypothetical protein
469(1) WP_142936449.1 WP_142936449.1 hypothetical protein

470(1) WP_180235367.1 WP_180235367.1 SecY-interacting protein Syd
470(3) WP_002137246.1 WP_002137246.1 SecY-interacting protein Syd
470(2) WP_002101116.1 WP_002101116.1 SecY-interacting protein Syd

471(6) WP_185336754.1 WP_185336754.1 hypothetical protein

472(6) WP_185336755.1 WP_185336755.1 hypothetical protein

473(1) WP_198050881.1 WP_198050881.1 hypothetical protein
473(2) WP_049720642.1 WP_049720642.1 hypothetical protein
473(3) WP_052186853.1 WP_052186853.1 hypothetical protein

474(1) WP_098303482.1 WP_098303482.1 DegV family protein
474(1) WP_090688933.1 WP_090688933.1 DegV family protein
474(1) WP_017149779.1 WP_017149779.1 DegV family protein
474(1) WP_097899499.1 WP_097899499.1 DegV family protein
474(1) WP_207994169.1 WP_207994169.1 DegV family protein
474(1) WP_208019340.1 WP_208019340.1 DegV family protein

475(2) WP_187374156.1 WP_187374156.1 hypothetical protein
475(1) WP_195974876.1 WP_195974876.1 hypothetical protein
475(1) WP_208926588.1 WP_208926588.1 hypothetical protein
475(1) WP_177576800.1 WP_177576800.1 hypothetical protein
475(1) WP_204445232.1 WP_204445232.1 hypothetical protein

476(2) WP_112331600.1 WP_112331600.1 zinc-binding dehydrogenase
476(1) WP_220559127.1 WP_220559127.1 NAD(P)H-quinone oxidoreductase
476(1) WP_223067445.1 WP_223067445.1 alcohol dehydrogenase catalytic domain-containing protein
476(1) WP_057897611.1 WP_057897611.1 NAD(P)H-quinone oxidoreductase
476(1) WP_181754678.1 WP_181754678.1 NAD(P)-dependent alcohol dehydrogenase

477(1) WP_077320193.1 WP_077320193.1 FixH family protein
477(1) WP_223068059.1 WP_223068059.1 FixH family protein
477(1) WP_038245455.1 WP_038245455.1 FixH family protein
477(3) WP_021291859.1 WP_021291859.1 FixH family protein

478(1) WP_079506382.1 WP_079506382.1 hypothetical protein
478(1) WP_215032133.1 WP_215032133.1 hypothetical protein
478(3) WP_144479072.1 WP_144479072.1 hypothetical protein
478(1) WP_226679108.1 WP_226679108.1 hypothetical protein

479(1) WP_049869152.1 WP_049869152.1 CrcB family protein
479(3) WP_033100008.1 WP_033100008.1 fluoride efflux transporter CrcB
479(1) WP_033100009.1 WP_033100009.1 fluoride efflux transporter CrcB
479(1) WP_227219165.1 WP_227219165.1 fluoride efflux transporter CrcB

480(1) WP_229522872.1 WP_229522872.1 MogA/MoaB family molybdenum cofactor biosynthesis protein
480(1) WP_007430113.1 WP_007430113.1 MogA/MoaB family molybdenum cofactor biosynthesis protein
480(1) WP_014281164.1 WP_014281164.1 MogA/MoaB family molybdenum cofactor biosynthesis protein
480(1) WP_010346303.1 WP_010346303.1 MogA/MoaB family molybdenum cofactor biosynthesis protein
480(1) WP_094155419.1 WP_094155419.1 MogA/MoaB family molybdenum cofactor biosynthesis protein
480(1) WP_068496271.1 WP_068496271.1 MogA/MoaB family molybdenum cofactor biosynthesis protein

481(1) WP_231094859.1 WP_231094859.1 hypothetical protein
481(1) WP_077320198.1 WP_077320198.1 hypothetical protein
481(4) WP_021291855.1 WP_021291855.1 hypothetical protein

482(1) WP_197052014.1 WP_197052014.1 DUF2690 domain-containing protein
482(2) WP_232335117.1 WP_232335117.1 DUF2690 domain-containing protein
482(1) WP_165489726.1 WP_165489726.1 DUF2690 domain-containing protein
482(2) WP_232335483.1 WP_232335483.1 DUF2690 domain-containing protein

483(1) WP_209872758.1 WP_209872758.1 CoA transferase subunit A
483(1) WP_019424151.1 WP_019424151.1 CoA-transferase
483(2) WP_087433601.1 WP_087433601.1 CoA-transferase
483(1) WP_233697170.1 WP_233697170.1 CoA transferase subunit A
483(1) WP_206094037.1 WP_206094037.1 CoA transferase subunit A

484(1) WP_017147605.1 WP_017147605.1 hypothetical protein
484(1) WP_237711211.1 WP_237711211.1 hypothetical protein
484(1) WP_006290626.1 WP_006290626.1 hypothetical protein
484(1) WP_007147996.1 WP_007147996.1 hypothetical protein
484(2) WP_006289751.1 WP_006289751.1 PsbP-related protein

485(2) WP_242172404.1 WP_242172404.1 dicarboxylate/amino acid:cation symporter
485(1) WP_053403796.1 WP_053403796.1 dicarboxylate/amino acid:cation symporter
485(1) WP_043067155.1 WP_043067155.1 L-cystine transporter
485(2) WP_040305106.1 WP_040305106.1 L-cystine transporter

486(1) WP_213618823.1 WP_213618823.1 PQQ-binding-like beta-propeller repeat protein
486(1) WP_068619094.1 WP_068619094.1 PQQ-binding-like beta-propeller repeat protein
486(1) WP_108465360.1 WP_108465360.1 PQQ-binding-like beta-propeller repeat protein
486(1) WP_251375540.1 WP_251375540.1 PQQ-binding-like beta-propeller repeat protein
486(1) WP_209861626.1 WP_209861626.1 PQQ-binding-like beta-propeller repeat protein
486(1) WP_127199781.1 WP_127199781.1 PQQ-binding-like beta-propeller repeat protein

487(1) WP_179160631.1 WP_179160631.1 amino acid ABC transporter substrate-binding protein
487(1) WP_034675391.1 WP_034675391.1 transporter substrate-binding domain-containing protein
487(1) WP_110931555.1 WP_110931555.1 amino acid ABC transporter substrate-binding protein
487(1) WP_120462939.1 WP_120462939.1 amino acid ABC transporter substrate-binding protein
487(1) WP_251417788.1 WP_251417788.1 amino acid ABC transporter substrate-binding protein
487(1) WP_010496598.1 WP_010496598.1 ABC transporter substrate-binding protein

488(1) WP_026572752.1 WP_026572752.1 YwpF family protein
488(1) WP_198310820.1 WP_198310820.1 YwpF family protein
488(1) WP_251466814.1 WP_251466814.1 YwpF-like family protein
488(1) WP_133332990.1 WP_133332990.1 YwpF family protein
488(1) WP_191272842.1 WP_191272842.1 YwpF family protein
488(1) WP_199419786.1 WP_199419786.1 YwpF-like family protein

489(1) WP_122971518.1 WP_122971518.1 DUF421 domain-containing protein
489(1) WP_213144476.1 WP_213144476.1 DUF421 domain-containing protein
489(1) WP_251557370.1 WP_251557370.1 DUF421 domain-containing protein
489(1) WP_251426354.1 WP_251426354.1 DUF421 domain-containing protein
489(1) WP_251557371.1 WP_251557371.1 DUF421 domain-containing protein
489(1) WP_251426355.1 WP_251426355.1 DUF421 domain-containing protein

490(1) WP_026572726.1 WP_026572726.1 general stress protein
490(1) WP_198310844.1 WP_198310844.1 general stress protein
490(1) WP_251610739.1 WP_251610739.1 general stress protein
490(1) WP_242172362.1 WP_242172362.1 general stress protein
490(1) WP_053403776.1 WP_053403776.1 general stress protein
490(1) WP_191272891.1 WP_191272891.1 general stress protein

491(1) WP_042210988.1 WP_042210988.1 alanine dehydrogenase
491(1) WP_242172366.1 WP_242172366.1 alanine dehydrogenase
491(1) WP_042134264.1 WP_042134264.1 alanine dehydrogenase
491(1) WP_076242353.1 WP_076242353.1 alanine dehydrogenase
491(1) WP_251610744.1 WP_251610744.1 alanine dehydrogenase
491(1) WP_053403777.1 WP_053403777.1 alanine dehydrogenase

492(1) WP_255516964.1 WP_255516964.1 hypothetical protein
492(5) WP_255347143.1 WP_255347143.1 hypothetical protein

493(1) WP_034095018.1 WP_034095018.1 hypothetical protein
493(1) WP_235209104.1 WP_235209104.1 hypothetical protein
493(1) WP_142613006.1 WP_142613006.1 hypothetical protein
493(1) WP_258296274.1 WP_258296274.1 hypothetical protein
493(1) WP_023988286.1 WP_023988286.1 hypothetical protein
493(1) WP_174822148.1 WP_174822148.1 hypothetical protein

494(1) WP_076264224.1 WP_076264224.1 hypothetical protein
494(1) WP_258296277.1 WP_258296277.1 hypothetical protein
494(2) WP_028542679.1 WP_028542679.1 hypothetical protein
494(1) WP_071640590.1 WP_071640590.1 hypothetical protein
494(1) WP_220027685.1 WP_220027685.1 hypothetical protein

495(1) WP_223830775.1 WP_223830775.1 hypothetical protein
495(1) WP_258382280.1 WP_258382280.1 YrpD family protein
495(1) WP_233901880.1 WP_233901880.1 YrpD family protein
495(1) WP_231118743.1 WP_231118743.1 hypothetical protein
495(1) WP_258296275.1 WP_258296275.1 hypothetical protein
495(1) WP_254348220.1 WP_254348220.1 YrpD family protein

496(1) WP_124049156.1 WP_124049156.1 6-phosphofructokinase
496(1) WP_019392129.1 WP_019392129.1 6-phosphofructokinase
496(1) WP_243518783.1 WP_243518783.1 6-phosphofructokinase
496(1) WP_254759287.1 WP_254759287.1 6-phosphofructokinase
496(1) WP_268583843.1 WP_268583843.1 6-phosphofructokinase
496(1) WP_120046152.1 WP_120046152.1 6-phosphofructokinase

497(1) WP_243518772.1 WP_243518772.1 four-carbon acid sugar kinase family protein
497(1) WP_019392125.1 WP_019392125.1 four-carbon acid sugar kinase family protein
497(2) WP_120046150.1 WP_120046150.1 four-carbon acid sugar kinase family protein
497(1) WP_268583846.1 WP_268583846.1 four-carbon acid sugar kinase family protein
497(1) WP_124049160.1 WP_124049160.1 four-carbon acid sugar kinase family protein

498(1) WP_243518768.1 WP_243518768.1 triose-phosphate isomerase
498(1) WP_124049163.1 WP_124049163.1 triose-phosphate isomerase
498(2) WP_040056498.1 WP_040056498.1 triose-phosphate isomerase
498(1) WP_268583850.1 WP_268583850.1 triose-phosphate isomerase
498(1) WP_019392122.1 WP_019392122.1 triose-phosphate isomerase

499(1) WP_063223287.1 WP_063223287.1 suppressor of fused domain protein
499(1) WP_012260840.1 WP_012260840.1 ankyrin repeat domain-containing protein
499(1) WP_199677611.1 WP_199677611.1 suppressor of fused domain protein
499(1) WP_272541586.1 WP_272541586.1 suppressor of fused domain protein
499(1) WP_098266654.1 WP_098266654.1 ankyrin repeat domain-containing protein
499(1) WP_017149757.1 WP_017149757.1 ankyrin repeat domain-containing protein

500(1) WP_161490537.1 WP_161490537.1 hypothetical protein
500(1) WP_176854273.1 WP_176854273.1 hypothetical protein

500(2) WP_170867772.1 WP_170867772.1 hypothetical protein
500(1) WP_274566443.1 WP_274566443.1 hypothetical protein
500(1) WP_175480541.1 WP_175480541.1 hypothetical protein

501(1) WP_124049157.1 WP_124049157.1 class II fructose-bisphosphatase
501(1) WP_106496013.1 WP_106496013.1 class II fructose-bisphosphatase
501(1) WP_120046151.1 WP_120046151.1 class II fructose-bisphosphatase
501(1) WP_276517256.1 WP_276517256.1 class II fructose-bisphosphatase
501(1) WP_268584094.1 WP_268584094.1 class II fructose-bisphosphatase
501(1) WP_019392128.1 WP_019392128.1 class II fructose-bisphosphatase

502(1) WP_036695736.1 WP_036695736.1 vitamin K epoxide reductase family protein
502(1) WP_281667414.1 WP_281667414.1 vitamin K epoxide reductase family protein
502(1) WP_006289742.1 WP_006289742.1 vitamin K epoxide reductase family protein
502(1) WP_277145024.1 WP_277145024.1 vitamin K epoxide reductase family protein
502(1) WP_277150336.1 WP_277150336.1 vitamin K epoxide reductase family protein
502(1) WP_277159398.1 WP_277159398.1 vitamin K epoxide reductase family protein

503(2) WP_057897613.1 WP_057897613.1 aconitate hydratase AcnA
503(1) WP_282742041.1 WP_282742041.1 3-isopropylmalate dehydratase large subunit
503(1) WP_278773416.1 WP_278773416.1 3-isopropylmalate dehydratase large subunit
503(1) WP_202898725.1 WP_202898725.1 3-isopropylmalate dehydratase large subunit
503(1) WP_013252395.1 WP_013252395.1 3-isopropylmalate dehydratase large subunit

504(1) WP_287600003.1 WP_287600003.1 hypothetical protein
504(1) WP_283157897.1 WP_283157897.1 hypothetical protein
504(1) WP_271839940.1 WP_271839940.1 hypothetical protein
504(1) WP_291647940.1 WP_291647940.1 hypothetical protein
504(1) WP_195431286.1 WP_195431286.1 hypothetical protein
504(1) WP_185165631.1 WP_185165631.1 hypothetical protein

505(1) WP_271839933.1 WP_271839933.1 hypothetical protein
505(1) WP_185165634.1 WP_185165634.1 hypothetical protein
505(1) WP_291647945.1 WP_291647945.1 hypothetical protein
505(1) WP_287600011.1 WP_287600011.1 hypothetical protein
505(2) WP_195431290.1 WP_195431290.1 hypothetical protein

506(1) WP_289475078.1 WP_289475078.1 glycoside hydrolase family 68 protein
506(1) WP_289483770.1 WP_289483770.1 glycoside hydrolase family 68 protein
506(1) WP_061975554.1 WP_061975554.1 glycoside hydrolase family 68 protein
506(1) WP_301167485.1 WP_301167485.1 glycoside hydrolase family 68 protein
506(1) WP_129478262.1 WP_129478262.1 glycoside hydrolase family 68 protein
506(1) WP_090239187.1 WP_090239187.1 glycoside hydrolase family 68 protein

507(1) WP_152392563.1 WP_152392563.1 threonine synthase
507(1) WP_291289933.1 WP_291289933.1 threonine ammonia-lyase
507(1) WP_297183828.1 WP_297183828.1 threonine ammonia-lyase
507(2) WP_219872439.1 WP_219872439.1 threonine synthase
507(1) WP_302153069.1 WP_302153069.1 threonine ammonia-lyase

508(1) WP_127484325.1 WP_127484325.1 peptidyl-prolyl cis-trans isomerase
508(1) WP_025848636.1 WP_025848636.1 peptidylprolyl isomerase
508(1) WP_054972108.1 WP_054972108.1 peptidylprolyl isomerase
508(1) WP_127484323.1 WP_127484323.1 hypothetical protein
508(1) WP_088830550.1 WP_088830550.1 peptidyl-prolyl cis-trans isomerase
508(1) WP_302878969.1 WP_302878969.1 hypothetical protein

509(1) WP_013252390.1 WP_013252390.1 hypothetical protein
509(1) WP_156900992.1 WP_156900992.1 hypothetical protein
509(1) WP_036575528.1 WP_036575528.1 hypothetical protein
509(1) WP_278773424.1 WP_278773424.1 hypothetical protein
509(1) WP_307388968.1 WP_307388968.1 hypothetical protein
509(1) WP_278591887.1 WP_278591887.1 hypothetical protein

510(1) WP_013252377.1 WP_013252377.1 WG repeat-containing protein
510(1) WP_278773450.1 WP_278773450.1 WG repeat-containing protein
510(1) WP_273061471.1 WP_273061471.1 WG repeat-containing protein
510(1) WP_307388988.1 WP_307388988.1 WG repeat-containing protein
510(1) WP_013252376.1 WP_013252376.1 WG repeat-containing protein
510(1) WP_036575558.1 WP_036575558.1 WG repeat-containing protein

511(1) WP_024027641.1 WP_024027641.1 hypothetical protein
511(1) WP_308110108.1 WP_308110108.1 beta-mannanase
511(1) WP_149871670.1 WP_149871670.1 hypothetical protein
511(1) WP_007083301.1 WP_007083301.1 hypothetical protein
511(1) WP_066148106.1 WP_066148106.1 hypothetical protein
511(1) WP_166245143.1 WP_166245143.1 beta-mannanase

512(1) WP_007083300.1 WP_007083300.1 hypothetical protein
512(1) WP_024027640.1 WP_024027640.1 hypothetical protein
512(1) WP_166245141.1 WP_166245141.1 hypothetical protein
512(1) WP_308110109.1 WP_308110109.1 hypothetical protein
512(1) WP_066091977.1 WP_066091977.1 hypothetical protein
512(1) WP_149871671.1 WP_149871671.1 hypothetical protein

513(1) WP_308912802.1 WP_308912802.1 MGMT family protein
513(1) WP_251466788.1 WP_251466788.1 MGMT family protein
513(1) WP_198310823.1 WP_198310823.1 MGMT family protein
513(1) WP_191272875.1 WP_191272875.1 MGMT family protein
513(1) WP_026572747.1 WP_026572747.1 MGMT family protein
513(1) WP_133332988.1 WP_133332988.1 MGMT family protein

514(1) WP_308912814.1 WP_308912814.1 hypothetical protein
514(1) WP_308912815.1 WP_308912815.1 hypothetical protein
514(1) WP_165976162.1 WP_165976162.1 hypothetical protein
514(1) WP_165976163.1 WP_165976163.1 hypothetical protein
514(1) WP_308912813.1 WP_308912813.1 hypothetical protein
514(1) WP_165976164.1 WP_165976164.1 hypothetical protein

515(5) WP_000798747.1 WP_000798747.1 hypothetical protein

516(3) WP_006289748.1 WP_006289748.1 WXG100 family type VII secretion target
516(1) WP_006292341.1 WP_006292341.1 WXG100 family type VII secretion target
516(1) WP_007147993.1 WP_007147993.1 WXG100 family type VII secretion target

517(1) WP_043067139.1 WP_043067139.1 DUF3951 domain-containing protein
517(3) WP_040305080.1 WP_040305080.1 DUF3951 domain-containing protein
517(1) WP_043069425.1 WP_043069425.1 DUF3951 domain-containing protein

518(1) WP_034526622.1 WP_034526622.1 hypothetical protein
518(1) WP_006292338.1 WP_006292338.1 hypothetical protein
518(1) WP_017147606.1 WP_017147606.1 hypothetical protein
518(1) WP_006289753.1 WP_006289753.1 hypothetical protein
518(1) WP_050954543.1 WP_050954543.1 hypothetical protein

519(3) WP_040101527.1 WP_040101527.1 Rha family transcriptional regulator
519(1) WP_053326560.1 WP_053326560.1 Rha family transcriptional regulator
519(1) WP_007430139.1 WP_007430139.1 hypothetical protein

520(1) WP_066092035.1 WP_066092035.1 Uracil permease
520(1) WP_066092038.1 WP_066092038.1 hypothetical protein
520(1) WP_066092030.1 WP_066092030.1 Uracil permease
520(1) WP_066092033.1 WP_066092033.1 Uracil permease
520(1) WP_066092027.1 WP_066092027.1 Uracil permease

521(3) WP_038245454.1 WP_038245454.1 stage II sporulation protein P
521(1) WP_021291858.1 WP_021291858.1 stage II sporulation protein P
521(1) WP_077320196.1 WP_077320196.1 stage II sporulation protein P

522(2) WP_003516583.1 WP_003516583.1 isoamylase early set domain-containing protein
522(1) WP_050753439.1 WP_050753439.1 hypothetical protein
522(1) WP_038290980.1 WP_038290980.1 isoamylase early set domain-containing protein
522(1) WP_128705701.1 WP_128705701.1 isoamylase early set domain-containing protein

523(1) WP_151585715.1 WP_151585715.1 RraA family protein
523(1) WP_113713153.1 WP_113713153.1 RraA family protein
523(1) WP_040056492.1 WP_040056492.1 RraA family protein
523(1) WP_111925431.1 WP_111925431.1 RraA family protein
523(1) WP_113753115.1 WP_113753115.1 RraA family protein

524(1) WP_080787837.1 WP_080787837.1 aspartyl-phosphate phosphatase Spo0E family protein
524(1) WP_152623316.1 WP_152623316.1 aspartyl-phosphate phosphatase Spo0E family protein
524(1) WP_168976291.1 WP_168976291.1 aspartyl-phosphate phosphatase Spo0E family protein
524(2) WP_167331605.1 WP_167331605.1 aspartyl-phosphate phosphatase Spo0E family protein

525(4) WP_185350819.1 WP_185350819.1 hypothetical protein
525(1) WP_077320212.1 WP_077320212.1 hypothetical protein

526(2) WP_097898661.1 WP_097898661.1 sporulation protein
526(1) WP_033100001.1 WP_033100001.1 sporulation protein
526(1) WP_198056472.1 WP_198056472.1 sporulation protein
526(1) WP_198059695.1 WP_198059695.1 sporulation protein

527(2) WP_213592846.1 WP_213592846.1 transcriptional regulator
527(1) WP_110934059.1 WP_110934059.1 transcriptional regulator
527(1) WP_125084769.1 WP_125084769.1 transcriptional regulator
527(1) WP_155609744.1 WP_155609744.1 transcriptional regulator

528(1) WP_216975176.1 WP_216975176.1 hypothetical protein
528(2) WP_013370793.1 WP_013370793.1 hypothetical protein
528(1) WP_137078370.1 WP_137078370.1 hypothetical protein
528(1) WP_058829517.1 WP_058829517.1 hypothetical protein

529(1) WP_137078371.1 WP_137078371.1 AAA family ATPase
529(1) WP_216975177.1 WP_216975177.1 AAA family ATPase
529(2) WP_013370794.1 WP_013370794.1 AAA family ATPase
529(1) WP_058829518.1 WP_058829518.1 AAA family ATPase

530(2) WP_185606003.1 WP_185606003.1 hypothetical protein
530(3) WP_228469158.1 WP_228469158.1 hypothetical protein

531(1) WP_231417854.1 WP_231417854.1 chromate transporter
531(1) WP_231417853.1 WP_231417853.1 chromate transporter
531(1) WP_035190290.1 WP_035190290.1 chromate transporter
531(1) WP_035190292.1 WP_035190292.1 chromate transporter
531(1) WP_132283076.1 WP_132283076.1 chromate transporter

532(2) WP_098248506.1 WP_098248506.1 zinc ribbon domain-containing protein
532(1) WP_231417862.1 WP_231417862.1 zinc ribbon domain-containing protein
532(1) WP_136789997.1 WP_136789997.1 zinc ribbon domain-containing protein
532(1) WP_132018305.1 WP_132018305.1 zinc ribbon domain-containing protein

533(1) WP_233697163.1 WP_233697163.1 BMC domain-containing protein
533(2) WP_087433592.1 WP_087433592.1 BMC domain-containing protein
533(1) WP_206094041.1 WP_206094041.1 BMC domain-containing protein
533(1) WP_019424161.1 WP_019424161.1 BMC domain-containing protein

534(1) WP_047153337.1 WP_047153337.1 COX aromatic rich motif-containing protein
534(2) WP_242172385.1 WP_242172385.1 cytochrome aa3 quinol oxidase subunit II
534(1) WP_057897637.1 WP_057897637.1 cytochrome c oxidase subunit II
534(1) WP_053403789.1 WP_053403789.1 cytochrome aa3 quinol oxidase subunit II

535(2) WP_053356043.1 WP_053356043.1 cold shock protein CspC
535(2) WP_248251177.1 WP_248251177.1 cold shock domain-containing protein
535(1) WP_231417876.1 WP_231417876.1 cold-shock protein

536(1) WP_251417635.1 WP_251417635.1 1,4-dihydroxy-6-naphthoate synthase
536(1) WP_188535048.1 WP_188535048.1 1,4-dihydroxy-6-naphthoate synthase
536(1) WP_090688970.1 WP_090688970.1 hypothetical protein
536(1) WP_019120051.1 WP_019120051.1 hypothetical protein
536(1) WP_127194951.1 WP_127194951.1 1,4-dihydroxy-6-naphthoate synthase

537(1) WP_257020596.1 WP_257020596.1 hypothetical protein
537(1) WP_144424628.1 WP_144424628.1 hypothetical protein
537(3) WP_063566330.1 WP_063566330.1 hypothetical protein

538(1) WP_215185492.1 WP_215185492.1 hypothetical protein
538(1) WP_215185486.1 WP_215185486.1 hypothetical protein

538(1) WP_189031020.1 WP_189031020.1 hypothetical protein
538(1) WP_257443634.1 WP_257443634.1 hypothetical protein
538(1) WP_189031022.1 WP_189031022.1 hypothetical protein

539(1) WP_215185485.1 WP_215185485.1 hypothetical protein
539(1) WP_189031021.1 WP_189031021.1 hypothetical protein
539(1) WP_215185490.1 WP_215185490.1 hypothetical protein
539(1) WP_189031023.1 WP_189031023.1 hypothetical protein
539(1) WP_257443636.1 WP_257443636.1 hypothetical protein

540(1) WP_077852628.1 WP_077852628.1 hypothetical protein
540(1) WP_077893590.1 WP_077893590.1 hypothetical protein
540(1) WP_257675224.1 WP_257675224.1 hypothetical protein
540(1) WP_077833121.1 WP_077833121.1 hypothetical protein
540(1) WP_077852426.1 WP_077852426.1 hypothetical protein

541(1) WP_121474468.1 WP_121474468.1 hypothetical protein
541(1) WP_018670539.1 WP_018670539.1 hypothetical protein
541(2) WP_104030454.1 WP_104030454.1 hypothetical protein
541(1) WP_258433009.1 WP_258433009.1 hypothetical protein

542(3) WP_087433602.1 WP_087433602.1 CoA-transferase
542(1) WP_267934046.1 WP_267934046.1 CoA-transferase
542(1) WP_019424150.1 WP_019424150.1 CoA-transferase

543(1) WP_040056494.1 WP_040056494.1 4-hydroxythreonine-4-phosphate dehydrogenase PdxA
543(1) WP_243518775.1 WP_243518775.1 4-hydroxythreonine-4-phosphate dehydrogenase PdxA
543(1) WP_124049159.1 WP_124049159.1 4-hydroxythreonine-4-phosphate dehydrogenase PdxA
543(1) WP_268583845.1 WP_268583845.1 4-hydroxythreonine-4-phosphate dehydrogenase PdxA
543(1) WP_019392126.1 WP_019392126.1 4-hydroxythreonine-4-phosphate dehydrogenase PdxA

544(1) WP_199613897.1 WP_199613897.1 hypothetical protein
544(1) WP_243495552.1 WP_243495552.1 hypothetical protein
544(1) WP_268613675.1 WP_268613675.1 hypothetical protein
544(1) WP_244723769.1 WP_244723769.1 hypothetical protein
544(1) WP_048745732.1 WP_048745732.1 hypothetical protein

545(1) WP_034773566.1 WP_034773566.1 sulfurtransferase
545(1) WP_251245817.1 WP_251245817.1 sulfurtransferase
545(1) WP_272537436.1 WP_272537436.1 sulfurtransferase
545(1) WP_240550299.1 WP_240550299.1 sulfurtransferase
545(1) WP_251241785.1 WP_251241785.1 sulfurtransferase

546(1) WP_272801664.1 WP_272801664.1 UTP--glucose-1-phosphate uridylyltransferase GalU
546(1) WP_243495567.1 WP_243495567.1 UTP--glucose-1-phosphate uridylyltransferase GalU
546(1) WP_186580055.1 WP_186580055.1 UTP--glucose-1-phosphate uridylyltransferase GalU
546(1) WP_015595770.1 WP_015595770.1 glucose-1-phosphate thymidylyltransferase RfbA
546(1) WP_098579106.1 WP_098579106.1 UTP--glucose-1-phosphate uridylyltransferase GalU

547(1) WP_212693229.1 WP_212693229.1 hypothetical protein
547(1) WP_209365672.1 WP_209365672.1 hypothetical protein
547(1) WP_074598749.1 WP_074598749.1 hypothetical protein
547(1) WP_245208022.1 WP_245208022.1 hypothetical protein
547(1) WP_273321563.1 WP_273321563.1 hypothetical protein

548(1) WP_274378588.1 WP_274378588.1 hypothetical protein
548(1) WP_098303174.1 WP_098303174.1 hypothetical protein
548(1) WP_012260844.1 WP_012260844.1 hypothetical protein
548(1) WP_238187893.1 WP_238187893.1 hypothetical protein
548(1) WP_090688985.1 WP_090688985.1 hypothetical protein

549(1) WP_242400030.1 WP_242400030.1 hypothetical protein
549(1) WP_275897037.1 WP_275897037.1 hypothetical protein
549(3) WP_000621737.1 WP_000621737.1 hypothetical protein

550(2) WP_277157954.1 WP_277157954.1 serine hydroxymethyltransferase
550(1) WP_277142102.1 WP_277142102.1 serine hydroxymethyltransferase
550(1) WP_277150345.1 WP_277150345.1 serine hydroxymethyltransferase
550(1) WP_277156195.1 WP_277156195.1 serine hydroxymethyltransferase

551(1) WP_277142105.1 WP_277142105.1 DUF3052 domain-containing protein
551(2) WP_277157956.1 WP_277157956.1 DUF3052 domain-containing protein
551(1) WP_277156196.1 WP_277156196.1 DUF3052 domain-containing protein
551(1) WP_277150891.1 WP_277150891.1 DUF3052 domain-containing protein

552(1) WP_277175574.1 WP_277175574.1 hypothetical protein
552(1) WP_277166571.1 WP_277166571.1 hypothetical protein
552(1) WP_277156176.1 WP_277156176.1 hypothetical protein
552(1) WP_277156177.1 WP_277156177.1 hypothetical protein
552(1) WP_277142074.1 WP_277142074.1 hypothetical protein

553(1) WP_277142107.1 WP_277142107.1 zinc-dependent metalloprotease
553(1) WP_277150343.1 WP_277150343.1 zinc-dependent metalloprotease
553(1) WP_277156198.1 WP_277156198.1 zinc-dependent metalloprotease
553(1) WP_277175596.1 WP_277175596.1 zinc-dependent metalloprotease
553(1) WP_277157959.1 WP_277157959.1 zinc-dependent metalloprotease

554(1) WP_280788642.1 WP_280788642.1 amino acid permease
554(1) WP_247083032.1 WP_247083032.1 APC family permease
554(1) WP_205001903.1 WP_205001903.1 amino acid permease
554(1) WP_089965973.1 WP_089965973.1 APC family permease
554(1) WP_251548781.1 WP_251548781.1 APC family permease

555(1) WP_283952965.1 WP_283952965.1 GNAT family N-acetyltransferase
555(1) WP_195571414.1 WP_195571414.1 GNAT family N-acetyltransferase
555(1) WP_169028904.1 WP_169028904.1 GNAT family N-acetyltransferase
555(2) WP_139829224.1 WP_139829224.1 GNAT family N-acetyltransferase

556(2) WP_213117464.1 WP_213117464.1 hypothetical protein
556(1) WP_213143933.1 WP_213143933.1 hypothetical protein
556(1) WP_284036505.1 WP_284036505.1 hypothetical protein
556(1) WP_066022444.1 WP_066022444.1 CBS domain-containing protein

557(1) WP_038245453.1 WP_038245453.1 UPF0182 family protein

| | | |
|---|---|---|
| 557(1) | WP_245799804.1 | WP_245799804.1 UPF0182 family protein |
| 557(1) | WP_286191346.1 | WP_286191346.1 UPF0182 family protein |
| 557(1) | WP_234448072.1 | WP_234448072.1 UPF0182 family protein |
| 557(1) | WP_021291857.1 | WP_021291857.1 UPF0182 family protein |
| 558(2) | WP_055277398.1 | WP_055277398.1 class I mannose-6-phosphate isomerase |
| 558(1) | WP_195952730.1 | WP_195952730.1 class I mannose-6-phosphate isomerase |
| 558(1) | WP_288218590.1 | WP_288218590.1 class I mannose-6-phosphate isomerase |
| 558(1) | WP_195959144.1 | WP_195959144.1 class I mannose-6-phosphate isomerase |
| 559(1) | WP_074570400.1 | WP_074570400.1 single-stranded DNA-binding protein |
| 559(1) | WP_251466815.1 | WP_251466815.1 single-stranded DNA-binding protein |
| 559(1) | WP_033692028.1 | WP_033692028.1 single-stranded DNA-binding protein |
| 559(1) | WP_199419785.1 | WP_199419785.1 single-stranded DNA-binding protein |
| 559(1) | WP_289607505.1 | WP_289607505.1 single-stranded DNA-binding protein |
| 560(1) | WP_152547783.1 | WP_152547783.1 DNRLRE domain-containing protein |
| 560(1) | WP_127484322.1 | WP_127484322.1 DNRLRE domain-containing protein |
| 560(1) | WP_088830551.1 | WP_088830551.1 DNRLRE domain-containing protein |
| 560(1) | WP_157261481.1 | WP_157261481.1 DNRLRE domain-containing protein |
| 560(1) | WP_302878968.1 | WP_302878968.1 DNRLRE domain-containing protein |
| 561(1) | WP_273191238.1 | WP_273191238.1 glycoside hydrolase family 1 protein |
| 561(1) | WP_273383858.1 | WP_273383858.1 glycoside hydrolase family 1 protein |
| 561(1) | WP_094870395.1 | WP_094870395.1 6-phospho-beta-glucosidase |
| 561(1) | WP_102372763.1 | WP_102372763.1 glycoside hydrolase family 1 protein |
| 561(1) | WP_303942512.1 | WP_303942512.1 glycoside hydrolase family 1 protein |
| 562(1) | WP_273383860.1 | WP_273383860.1 PTS transporter subunit EIIC |
| 562(1) | WP_066020857.1 | WP_066020857.1 PTS sugar transporter subunit IIC |
| 562(2) | WP_102372762.1 | WP_102372762.1 PTS transporter subunit EIIC |
| 562(1) | WP_303942513.1 | WP_303942513.1 PTS transporter subunit EIIC |
| 563(1) | WP_102372743.1 | WP_102372743.1 type III pantothenate kinase |
| 563(1) | WP_273383884.1 | WP_273383884.1 type III pantothenate kinase |
| 563(1) | WP_273191263.1 | WP_273191263.1 type III pantothenate kinase |
| 563(1) | WP_115363974.1 | WP_115363974.1 type III pantothenate kinase |
| 563(1) | WP_303942548.1 | WP_303942548.1 type III pantothenate kinase |
| 564(1) | WP_273383886.1 | WP_273383886.1 OPT/YSL family transporter |
| 564(1) | WP_273191265.1 | WP_273191265.1 OPT/YSL family transporter |
| 564(1) | WP_102372742.1 | WP_102372742.1 OPT/YSL family transporter |
| 564(1) | WP_303942550.1 | WP_303942550.1 OPT/YSL family transporter |
| 564(1) | WP_291288248.1 | WP_291288248.1 OPT/YSL family transporter |
| 565(1) | WP_099361328.1 | WP_099361328.1 sulfite exporter TauE/SafE family protein |
| 565(1) | WP_116353268.1 | WP_116353268.1 sulfite exporter TauE/SafE family protein |
| 565(1) | WP_304216850.1 | WP_304216850.1 sulfite exporter TauE/SafE family protein |
| 565(1) | WP_099354893.1 | WP_099354893.1 sulfite exporter TauE/SafE family protein |
| 565(1) | WP_010676876.1 | WP_010676876.1 sulfite exporter TauE/SafE family protein |
| 566(1) | WP_013252392.1 | WP_013252392.1 hypothetical protein |
| 566(1) | WP_307388965.1 | WP_307388965.1 hypothetical protein |
| 566(1) | WP_278773420.1 | WP_278773420.1 hypothetical protein |
| 566(1) | WP_036575523.1 | WP_036575523.1 hypothetical protein |
| 566(1) | WP_282742037.1 | WP_282742037.1 hypothetical protein |
| 567(1) | WP_013252391.1 | WP_013252391.1 hypothetical protein |
| 567(1) | WP_036575525.1 | WP_036575525.1 hypothetical protein |
| 567(2) | WP_278773422.1 | WP_278773422.1 hypothetical protein |
| 567(1) | WP_307388966.1 | WP_307388966.1 hypothetical protein |
| 568(1) | WP_307399167.1 | WP_307399167.1 AAA family ATPase |
| 568(1) | WP_235827623.1 | WP_235827623.1 hypothetical protein |
| 568(1) | WP_105368178.1 | WP_105368178.1 AAA family ATPase |
| 568(1) | WP_235827629.1 | WP_235827629.1 ATP-binding protein |
| 568(1) | WP_101299048.1 | WP_101299048.1 AAA family ATPase |
| 569(1) | WP_179160718.1 | WP_179160718.1 amidohydrolase |
| 569(1) | WP_034675434.1 | WP_034675434.1 amidohydrolase |
| 569(1) | WP_239548978.1 | WP_239548978.1 M20 family metallopeptidase |
| 569(1) | WP_102710736.1 | WP_102710736.1 M20/M25/M40 family metallo-hydrolase |
| 569(1) | WP_307589587.1 | WP_307589587.1 amidohydrolase |
| 570(1) | WP_243537369.1 | WP_243537369.1 hypothetical protein |
| 570(1) | WP_257126964.1 | WP_257126964.1 hypothetical protein |
| 570(1) | WP_258013845.1 | WP_258013845.1 hypothetical protein |
| 570(1) | WP_240664151.1 | WP_240664151.1 hypothetical protein |
| 570(1) | WP_307589613.1 | WP_307589613.1 hypothetical protein |
| 571(1) | WP_307284656.1 | WP_307284656.1 transposase |
| 571(1) | WP_285889798.1 | WP_285889798.1 transposase |
| 571(1) | WP_308082233.1 | WP_308082233.1 transposase |
| 571(2) | WP_169317565.1 | WP_169317565.1 transposase |
| 572(1) | WP_094092632.1 | WP_094092632.1 histidine kinase |
| 572(1) | WP_214705998.1 | WP_214705998.1 histidine kinase |
| 572(1) | WP_308110120.1 | WP_308110120.1 sensor histidine kinase |
| 572(1) | WP_053361027.1 | WP_053361027.1 sensor histidine kinase |
| 572(1) | WP_149871691.1 | WP_149871691.1 sensor histidine kinase |
| 573(2) | WP_010963377.1 | WP_010963377.1 hypothetical protein |
| 573(2) | WP_010963378.1 | WP_010963378.1 hypothetical protein |
| 574(4) | WP_013252394.1 | WP_013252394.1 3-isopropylmalate dehydratase small subunit |
| 575(1) | WP_036642147.1 | WP_036642147.1 BlaI/MecI/CopY family transcriptional regulator |
| 575(3) | WP_006211220.1 | WP_006211220.1 BlaI/MecI/CopY family transcriptional regulator |
| 576(4) | WP_038406448.1 | WP_038406448.1 matrixin family metalloprotease |
| 577(4) | WP_038408428.1 | WP_038408428.1 hypothetical protein |
| 578(3) | WP_053403786.1 | WP_053403786.1 cytochrome aa3 quinol oxidase subunit IV |

578(1) WP_047153279.1 WP_047153279.1 cytochrome o ubiquinol oxidase subunit IV

579(2) WP_003516588.1 WP_003516588.1 hypothetical protein
579(1) WP_069193866.1 WP_069193866.1 DUF1657 domain-containing protein
579(1) WP_038290987.1 WP_038290987.1 hypothetical protein

580(1) WP_069193868.1 WP_069193868.1 hypothetical protein
580(2) WP_003516586.1 WP_003516586.1 hypothetical protein
580(1) WP_038290974.1 WP_038290974.1 hypothetical protein

581(4) WP_074436684.1 WP_074436684.1 YjcZ family sporulation protein

582(1) WP_076242352.1 WP_076242352.1 ABC transporter substrate-binding protein
582(1) WP_042462825.1 WP_042462825.1 ABC transporter substrate-binding protein
582(1) WP_042210987.1 WP_042210987.1 ABC transporter substrate-binding protein
582(1) WP_042134262.1 WP_042134262.1 ABC transporter substrate-binding protein

583(2) WP_010963373.1 WP_010963373.1 hypothetical protein
583(2) WP_077832297.1 WP_077832297.1 hypothetical protein

584(3) WP_077832291.1 WP_077832291.1 hypothetical protein
584(1) WP_077852742.1 WP_077852742.1 hypothetical protein

585(2) WP_077833116.1 WP_077833116.1 histidine ammonia-lyase
585(1) WP_077852422.1 WP_077852422.1 histidine ammonia-lyase
585(1) WP_077893586.1 WP_077893586.1 histidine ammonia-lyase

586(4) WP_111925432.1 WP_111925432.1 HPr family phosphocarrier protein

587(1) WP_113713154.1 WP_113713154.1 hypothetical protein
587(3) WP_040056490.1 WP_040056490.1 hypothetical protein

588(2) WP_113713155.1 WP_113713155.1 hypothetical protein
588(1) WP_113753118.1 WP_113753118.1 hypothetical protein
588(1) WP_111925433.1 WP_111925433.1 hypothetical protein

589(3) WP_110934058.1 WP_110934058.1 membrane-associated protease 1
589(1) WP_125084768.1 WP_125084768.1 membrane-associated protease 1

590(1) WP_038290968.1 WP_038290968.1 vitamin B12 dependent-methionine synthase activation domain-containing protein
590(2) WP_003516590.1 WP_003516590.1 methionine synthase
590(1) WP_128705699.1 WP_128705699.1 methionine synthase

591(2) WP_003516587.1 WP_003516587.1 spore coat protein
591(1) WP_128705700.1 WP_128705700.1 spore coat protein
591(1) WP_038290971.1 WP_038290971.1 spore coat protein

592(1) WP_121610582.1 WP_121610582.1 alpha-glucosidase
592(1) WP_132008139.1 WP_132008139.1 alpha-glucosidase
592(1) WP_046522774.1 WP_046522774.1 alpha-glucosidase
592(1) WP_053361035.1 WP_053361035.1 alpha-glucosidase

593(1) WP_000756466.1 WP_000756466.1 nucleoside hydrolase
593(1) WP_007430144.1 WP_007430144.1 nucleoside hydrolase
593(1) WP_148409454.1 WP_148409454.1 nucleoside hydrolase
593(1) WP_109625253.1 WP_109625253.1 nucleoside hydrolase

594(1) WP_151585712.1 WP_151585712.1 glyoxalase/bleomycin resistance/dioxygenase family protein
594(1) WP_113753120.1 WP_113753120.1 glyoxalase/bleomycin resistance/dioxygenase family protein
594(1) WP_113765175.1 WP_113765175.1 glyoxalase/bleomycin resistance/dioxygenase family protein
594(1) WP_113713156.1 WP_113713156.1 glyoxalase/bleomycin resistance/dioxygenase family protein

595(1) WP_055277408.1 WP_055277408.1 hypothetical protein
595(1) WP_055291901.1 WP_055291901.1 hypothetical protein
595(2) WP_154821248.1 WP_154821248.1 hypothetical protein

596(3) WP_013914201.1 WP_013914201.1 hypothetical protein
596(1) WP_166244918.1 WP_166244918.1 hypothetical protein

597(2) WP_013914202.1 WP_013914202.1 hypothetical protein
597(1) WP_014368249.1 WP_014368249.1 hypothetical protein
597(1) WP_166244920.1 WP_166244920.1 hypothetical protein

598(1) WP_166244922.1 WP_166244922.1 hypothetical protein
598(1) WP_041619014.1 WP_041619014.1 hypothetical protein
598(1) WP_014649152.1 WP_014649152.1 hypothetical protein
598(1) WP_013914203.1 WP_013914203.1 hypothetical protein

599(1) WP_014649153.1 WP_014649153.1 hypothetical protein
599(1) WP_013914204.1 WP_013914204.1 hypothetical protein
599(1) WP_014368251.1 WP_014368251.1 hypothetical protein
599(1) WP_166244924.1 WP_166244924.1 hypothetical protein

600(1) WP_084595920.1 WP_084595920.1 anti-repressor SinI family protein
600(1) WP_182031394.1 WP_182031394.1 DNA-binding anti-repressor SinI
600(1) WP_114894457.1 WP_114894457.1 anti-repressor SinI family protein
600(1) WP_098579100.1 WP_098579100.1 anti-repressor SinI family protein

601(1) WP_142166602.1 WP_142166602.1 XRE family transcriptional regulator
601(1) WP_136789999.1 WP_136789999.1 helix-turn-helix transcriptional regulator
601(1) WP_098579101.1 WP_098579101.1 XRE family transcriptional regulator
601(1) WP_182031396.1 WP_182031396.1 helix-turn-helix transcriptional regulator

602(4) WP_185337288.1 WP_185337288.1 DUF3130 domain-containing protein

603(4) WP_185340302.1 WP_185340302.1 DNA double-strand break repair Rad50 ATPase

604(2) WP_185375747.1 WP_185375747.1 hypothetical protein
604(2) WP_185313123.1 WP_185313123.1 hypothetical protein

605(3) WP_021291861.1 WP_021291861.1 cytochrome c oxidase assembly protein
605(1) WP_188944150.1 WP_188944150.1 cytochrome c oxidase assembly protein

606(1) WP_160496943.1 WP_160496943.1 sporulation membrane protein YtaF
606(2) WP_194230413.1 WP_194230413.1 sporulation membrane protein YtaF
606(1) WP_136604805.1 WP_136604805.1 sporulation membrane protein YtaF

607(2) WP_185462636.1 WP_185462636.1 enhanced serine sensitivity protein SseB C-terminal domain-containing protein
607(2) WP_194333110.1 WP_194333110.1 enhanced serine sensitivity protein SseB C-terminal domain-containing protein

608(3) WP_156558889.1 WP_156558889.1 hypothetical protein
608(1) WP_195542466.1 WP_195542466.1 hypothetical protein

609(1) WP_195571413.1 WP_195571413.1 class I SAM-dependent methyltransferase
609(2) WP_085499125.1 WP_085499125.1 class I SAM-dependent methyltransferase
609(1) WP_169028903.1 WP_169028903.1 class I SAM-dependent methyltransferase

610(1) WP_197225367.1 WP_197225367.1 transposase, partial
610(1) WP_089522627.1 WP_089522627.1 IS256 family transposase
610(1) WP_094366203.1 WP_094366203.1 IS256 family transposase
610(1) WP_187644993.1 WP_187644993.1 transposase, partial

611(2) WP_198056471.1 WP_198056471.1 DUF84 family protein
611(1) WP_093291065.1 WP_093291065.1 inosine/xanthosine triphosphatase
611(1) WP_033100002.1 WP_033100002.1 DUF84 family protein

612(4) WP_203573365.1 WP_203573365.1 hypothetical protein

613(1) WP_019120070.1 WP_019120070.1 cysteine desulfurase family protein
613(1) WP_047153336.1 WP_047153336.1 aminotransferase class V-fold PLP-dependent enzyme
613(1) WP_204445252.1 WP_204445252.1 cysteine desulfurase NifS
613(1) WP_087194491.1 WP_087194491.1 aminotransferase class V-fold PLP-dependent enzyme

614(1) WP_138226040.1 WP_138226040.1 GTP-binding protein
614(1) WP_070638672.1 WP_070638672.1 GTP-binding protein
614(1) WP_058271121.1 WP_058271121.1 GTP-binding protein
614(1) WP_208203477.1 WP_208203477.1 GTP-binding protein

615(1) WP_208926641.1 WP_208926641.1 GHKL domain-containing protein
615(1) WP_210166180.1 WP_210166180.1 ATP-binding protein
615(1) WP_208926632.1 WP_208926632.1 hypothetical protein
615(1) WP_208926638.1 WP_208926638.1 sensor histidine kinase

616(1) WP_206094026.1 WP_206094026.1 class I SAM-dependent methyltransferase
616(1) WP_211745847.1 WP_211745847.1 class I SAM-dependent methyltransferase
616(2) WP_087433617.1 WP_087433617.1 class I SAM-dependent methyltransferase

617(1) WP_213143934.1 WP_213143934.1 hypothetical protein
617(3) WP_213117469.1 WP_213117469.1 hypothetical protein

618(1) WP_213143936.1 WP_213143936.1 hypothetical protein
618(3) WP_213117470.1 WP_213117470.1 hypothetical protein

619(1) WP_127199761.1 WP_127199761.1 alpha/beta hydrolase family protein
619(1) WP_209861655.1 WP_209861655.1 alpha/beta hydrolase family protein
619(1) WP_213618806.1 WP_213618806.1 alpha/beta hydrolase family protein
619(1) WP_068619130.1 WP_068619130.1 alpha/beta hydrolase family protein

620(1) WP_013370797.1 WP_013370797.1 hypothetical protein
620(1) WP_216975178.1 WP_216975178.1 hypothetical protein
620(2) WP_058829521.1 WP_058829521.1 hypothetical protein

621(2) WP_219872442.1 WP_219872442.1 VTT domain-containing protein
621(2) WP_074085364.1 WP_074085364.1 VTT domain-containing protein

622(2) WP_185462635.1 WP_185462635.1 hypothetical protein
622(2) WP_228474575.1 WP_228474575.1 hypothetical protein

623(1) WP_058271100.1 WP_058271100.1 NADPH-dependent glutamate synthase
623(1) WP_231287930.1 WP_231287930.1 FAD-dependent oxidoreductase
623(1) WP_186580043.1 WP_186580043.1 glutamate synthase subunit beta
623(1) WP_070638637.1 WP_070638637.1 NADPH-dependent glutamate synthase

624(1) WP_077852209.1 WP_077852209.1 group II intron reverse transcriptase/maturase
624(1) WP_229137199.1 WP_229137199.1 reverse transcriptase/maturase family protein
624(1) WP_237100198.1 WP_237100198.1 group II intron reverse transcriptase/maturase
624(1) WP_023520826.1 WP_023520826.1 group II intron reverse transcriptase/maturase

625(1) WP_068673732.1 WP_068673732.1 glucose-6-phosphate dehydrogenase
625(2) WP_242172358.1 WP_242172358.1 glucose-6-phosphate dehydrogenase
625(1) WP_053403774.1 WP_053403774.1 glucose-6-phosphate dehydrogenase

626(1) WP_053403787.1 WP_053403787.1 cytochrome aa3 quinol oxidase subunit III
626(2) WP_242172381.1 WP_242172381.1 cytochrome aa3 quinol oxidase subunit III
626(1) WP_047153280.1 WP_047153280.1 cytochrome o ubiquinol oxidase subunit III

627(1) WP_057897638.1 WP_057897638.1 cytochrome c oxidase subunit I
627(1) WP_047153281.1 WP_047153281.1 cytochrome aa3 quinol oxidase subunit I
627(1) WP_242172383.1 WP_242172383.1 cytochrome aa3 quinol oxidase subunit I
627(1) WP_053403788.1 WP_053403788.1 cytochrome aa3 quinol oxidase subunit I

628(1) WP_195244542.1 WP_195244542.1 polysaccharide pyruvyl transferase family protein
628(1) WP_005210480.1 WP_005210480.1 polysaccharide pyruvyl transferase family protein
628(2) WP_242824710.1 WP_242824710.1 polysaccharide pyruvyl transferase family protein

629(1) WP_136479370.1 WP_136479370.1 hypothetical protein
629(1) WP_243467748.1 WP_243467748.1 C39 family peptidase
629(1) WP_235832542.1 WP_235832542.1 hypothetical protein
629(1) WP_003516600.1 WP_003516600.1 hypothetical protein

630(1) WP_098248515.1 WP_098248515.1 zinc ribbon domain-containing protein

630(1) WP_243495555.1 WP_243495555.1 zinc ribbon domain-containing protein
630(1) WP_136789990.1 WP_136789990.1 zinc ribbon domain-containing protein
630(1) WP_142166614.1 WP_142166614.1 zinc ribbon domain-containing protein

631(2) WP_216518282.1 WP_216518282.1 RNA methyltransferase
631(1) WP_248063410.1 WP_248063410.1 RNA methyltransferase
631(1) WP_110899385.1 WP_110899385.1 RNA methyltransferase

632(1) WP_245208025.1 WP_245208025.1 family 43 glycosylhydrolase
632(1) WP_139186930.1 WP_139186930.1 family 43 glycosylhydrolase
632(1) WP_249897324.1 WP_249897324.1 glycoside hydrolase family 43 protein
632(1) WP_221875361.1 WP_221875361.1 glycoside hydrolase family 43 protein

633(1) WP_073008245.1 WP_073008245.1 hypothetical protein
633(1) WP_251417632.1 WP_251417632.1 hypothetical protein
633(1) WP_209365669.1 WP_209365669.1 hypothetical protein
633(1) WP_072336486.1 WP_072336486.1 hypothetical protein

634(1) WP_251466771.1 WP_251466771.1 DUF1146 family protein
634(1) WP_026572728.1 WP_026572728.1 DUF1146 family protein
634(1) WP_133332968.1 WP_133332968.1 DUF1146 family protein
634(1) WP_191273007.1 WP_191273007.1 DUF1146 family protein

635(1) WP_214706012.1 WP_214706012.1 hypothetical protein
635(1) WP_251542629.1 WP_251542629.1 hypothetical protein
635(1) WP_251481006.1 WP_251481006.1 hypothetical protein
635(1) WP_226679095.1 WP_226679095.1 hypothetical protein

636(1) WP_010963742.1 WP_010963742.1 type II CAAX endopeptidase family protein
636(1) WP_252976928.1 WP_252976928.1 CPBP family intramembrane glutamic endopeptidase
636(1) WP_252976929.1 WP_252976929.1 hypothetical protein
636(1) WP_097892028.1 WP_097892028.1 type II CAAX endopeptidase family protein

637(1) WP_211082931.1 WP_211082931.1 replication initiation factor family protein
637(1) WP_253440784.1 WP_253440784.1 replication initiation factor family protein
637(1) WP_055105784.1 WP_055105784.1 hypothetical protein
637(1) WP_068696855.1 WP_068696855.1 hypothetical protein

638(1) WP_253806482.1 WP_253806482.1 hypothetical protein
638(1) WP_254207755.1 WP_254207755.1 hypothetical protein
638(1) WP_249930775.1 WP_249930775.1 hypothetical protein
638(1) WP_014599682.1 WP_014599682.1 hypothetical protein

639(1) WP_077833099.1 WP_077833099.1 family 78 glycoside hydrolase catalytic domain
639(1) WP_257675214.1 WP_257675214.1 sugar hydrolase
639(1) WP_077852409.1 WP_077852409.1 family 78 glycoside hydrolase catalytic domain
639(1) WP_077893577.1 WP_077893577.1 family 78 glycoside hydrolase catalytic domain

640(1) WP_257675221.1 WP_257675221.1 HutP family protein
640(1) WP_077852423.1 WP_077852423.1 hypothetical protein
640(2) WP_077833117.1 WP_077833117.1 HutP family protein

641(1) WP_181737037.1 WP_181737037.1 IclR family transcriptional regulator
641(1) WP_263324535.1 WP_263324535.1 IclR family transcriptional regulator
641(2) WP_057897614.1 WP_057897614.1 IclR family transcriptional regulator

642(1) WP_181737043.1 WP_181737043.1 bifunctional 2-keto-4-hydroxyglutarate aldolase/2-keto-3-deoxy-6-phosphogluconate aldolase
642(1) WP_263324536.1 WP_263324536.1 bifunctional 4-hydroxy-2-oxoglutarate aldolase/2-dehydro-3-deoxy-phosphogluconate aldolase
642(2) WP_010963716.1 WP_010963716.1 bifunctional 4-hydroxy-2-oxoglutarate aldolase/2-dehydro-3-deoxy-phosphogluconate aldolase

643(1) WP_265444522.1 WP_265444522.1 hypothetical protein
643(2) WP_003516589.1 WP_003516589.1 hypothetical protein
643(1) WP_069193865.1 WP_069193865.1 hypothetical protein

644(1) WP_244723772.1 WP_244723772.1 hypothetical protein
644(1) WP_199613898.1 WP_199613898.1 hypothetical protein
644(1) WP_268613676.1 WP_268613676.1 hypothetical protein
644(1) WP_021880429.1 WP_021880429.1 hypothetical protein

645(1) WP_270065067.1 WP_270065067.1 YibE/F family protein
645(1) WP_066419959.1 WP_066419959.1 YibE/F family protein
645(1) WP_270065068.1 WP_270065068.1 YibE/F family protein
645(1) WP_066419961.1 WP_066419961.1 YibE/F family protein

646(2) WP_207738531.1 WP_207738531.1 hypothetical protein
646(1) WP_271839890.1 WP_271839890.1 hypothetical protein
646(1) WP_195343912.1 WP_195343912.1 hypothetical protein

647(1) WP_098579098.1 WP_098579098.1 histidinol-phosphatase HisJ family protein
647(1) WP_243495563.1 WP_243495563.1 histidinol-phosphatase HisJ family protein
647(1) WP_136789998.1 WP_136789998.1 histidinol-phosphatase HisJ family protein
647(1) WP_272801669.1 WP_272801669.1 histidinol-phosphatase HisJ family protein

648(1) WP_243495550.1 WP_243495550.1 HPP family protein
648(1) WP_272801675.1 WP_272801675.1 HPP family protein
648(1) WP_098248520.1 WP_098248520.1 HPP family protein
648(1) WP_028407764.1 WP_028407764.1 HPP family protein

649(1) WP_226887694.1 WP_226887694.1 DUF4179 domain-containing protein
649(1) WP_164965491.1 WP_164965491.1 DUF4179 domain-containing protein
649(1) WP_226887695.1 WP_226887695.1 DUF4179 domain-containing protein
649(1) WP_274485161.1 WP_274485161.1 DUF4179 domain-containing protein

650(1) WP_276867632.1 WP_276867632.1 hypothetical protein
650(1) WP_204666337.1 WP_204666337.1 hypothetical protein
650(2) WP_204666339.1 WP_204666339.1 hypothetical protein

651(1) WP_080787836.1 WP_080787836.1 cyclic lactone autoinducer peptide
651(2) WP_021624812.1 WP_021624812.1 cyclic lactone autoinducer peptide
651(1) WP_276917298.1 WP_276917298.1 cyclic lactone autoinducer peptide

| | | |
|---|---|---|
| 652(1) | WP_276917301.1 | WP_276917301.1 hypothetical protein |
| 652(1) | WP_235497057.1 | WP_235497057.1 hypothetical protein |
| 652(2) | WP_040305087.1 | WP_040305087.1 hypothetical protein |
| 653(1) | WP_277175595.1 | WP_277175595.1 S16 family serine protease |
| 653(1) | WP_277156197.1 | WP_277156197.1 S16 family serine protease |
| 653(1) | WP_277145022.1 | WP_277145022.1 S16 family serine protease |
| 653(1) | WP_277157957.1 | WP_277157957.1 S16 family serine protease |
| 654(1) | WP_277175598.1 | WP_277175598.1 DUF3107 domain-containing protein |
| 654(1) | WP_277157965.1 | WP_277157965.1 DUF3107 domain-containing protein |
| 654(1) | WP_277150341.1 | WP_277150341.1 DUF3107 domain-containing protein |
| 654(1) | WP_277142113.1 | WP_277142113.1 DUF3107 domain-containing protein |
| 655(2) | WP_179085874.1 | WP_179085874.1 Crp/Fnr family transcriptional regulator |
| 655(1) | WP_026592354.1 | WP_026592354.1 transcriptional regulator YeiL |
| 655(1) | WP_280788634.1 | WP_280788634.1 cyclic nucleotide-binding domain-containing protein |
| 656(1) | WP_006290903.1 | WP_006290903.1 quinone-dependent dihydroorotate dehydrogenase |
| 656(1) | WP_281667412.1 | WP_281667412.1 quinone-dependent dihydroorotate dehydrogenase |
| 656(1) | WP_006289739.1 | WP_006289739.1 quinone-dependent dihydroorotate dehydrogenase |
| 656(1) | WP_034526654.1 | WP_034526654.1 quinone-dependent dihydroorotate dehydrogenase |
| 657(1) | WP_273321566.1 | WP_273321566.1 hypothetical protein |
| 657(2) | WP_281813084.1 | WP_281813084.1 hypothetical protein |
| 657(1) | WP_212693228.1 | WP_212693228.1 hypothetical protein |
| 658(1) | WP_278773418.1 | WP_278773418.1 3-isopropylmalate dehydrogenase |
| 658(1) | WP_013252393.1 | WP_013252393.1 3-isopropylmalate dehydrogenase |
| 658(1) | WP_081742451.1 | WP_081742451.1 3-isopropylmalate dehydrogenase |
| 658(1) | WP_282742039.1 | WP_282742039.1 3-isopropylmalate dehydrogenase |
| 659(1) | WP_036575520.1 | WP_036575520.1 2-isopropylmalate synthase |
| 659(2) | WP_013252396.1 | WP_013252396.1 2-isopropylmalate synthase |
| 659(1) | WP_282742042.1 | WP_282742042.1 2-isopropylmalate synthase |
| 660(1) | WP_013252397.1 | WP_013252397.1 phosphoribosylanthranilate isomerase |
| 660(1) | WP_278773412.1 | WP_278773412.1 phosphoribosylanthranilate isomerase |
| 660(1) | WP_282742044.1 | WP_282742044.1 phosphoribosylanthranilate isomerase |
| 660(1) | WP_051417619.1 | WP_051417619.1 phosphoribosylanthranilate isomerase |
| 661(1) | WP_282742046.1 | WP_282742046.1 DegV family protein |
| 661(3) | WP_013252398.1 | WP_013252398.1 DegV family protein |
| 662(1) | WP_283952952.1 | WP_283952952.1 Ger(x)C family spore germination protein |
| 662(1) | WP_213143931.1 | WP_213143931.1 Ger(x)C family spore germination protein |
| 662(1) | WP_170138179.1 | WP_170138179.1 Ger(x)C family spore germination protein |
| 662(1) | WP_019120047.1 | WP_019120047.1 Ger(x)C family spore germination protein |
| 663(1) | WP_019120049.1 | WP_019120049.1 spore germination protein |
| 663(1) | WP_283952953.1 | WP_283952953.1 spore germination protein |
| 663(1) | WP_114298748.1 | WP_114298748.1 spore germination protein |
| 663(1) | WP_213143930.1 | WP_213143930.1 spore germination protein |
| 664(1) | WP_284036502.1 | WP_284036502.1 DUF2997 domain-containing protein |
| 664(2) | WP_213117475.1 | WP_213117475.1 DUF2997 domain-containing protein |
| 664(1) | WP_213143941.1 | WP_213143941.1 DUF2997 domain-containing protein |
| 665(2) | WP_213117472.1 | WP_213117472.1 DUF1257 domain-containing protein |
| 665(1) | WP_213143938.1 | WP_213143938.1 DUF1257 domain-containing protein |
| 665(1) | WP_284036503.1 | WP_284036503.1 DUF1257 domain-containing protein |
| 666(1) | WP_191798536.1 | WP_191798536.1 aminoglycoside phosphotransferase family protein |
| 666(1) | WP_285748189.1 | WP_285748189.1 aminoglycoside phosphotransferase family protein |
| 666(1) | WP_213532665.1 | WP_213532665.1 aminoglycoside phosphotransferase family protein |
| 666(1) | WP_211745820.1 | WP_211745820.1 aminoglycoside phosphotransferase family protein |
| 667(1) | WP_251426371.1 | WP_251426371.1 twin-arginine translocase TatA/TatE family subunit |
| 667(1) | WP_101356560.1 | WP_101356560.1 twin-arginine translocase TatA/TatE family subunit |
| 667(1) | WP_046522779.1 | WP_046522779.1 twin-arginine translocase TatA/TatE family subunit |
| 667(1) | WP_285867211.1 | WP_285867211.1 twin-arginine translocase TatA/TatE family subunit |
| 668(1) | WP_286124779.1 | WP_286124779.1 FAD dependent oxidoreductase |
| 668(1) | WP_208019344.1 | WP_208019344.1 FAD dependent oxidoreductase |
| 668(1) | WP_286017544.1 | WP_286017544.1 FAD dependent oxidoreductase |
| 668(1) | WP_255260045.1 | WP_255260045.1 FAD dependent oxidoreductase |
| 669(1) | WP_288218594.1 | WP_288218594.1 YfhO family protein |
| 669(1) | WP_196048859.1 | WP_196048859.1 YfhO family protein |
| 669(1) | WP_055277415.1 | WP_055277415.1 YfhO family protein |
| 669(1) | WP_195951787.1 | WP_195951787.1 YfhO family protein |
| 670(1) | WP_289376841.1 | WP_289376841.1 hypothetical protein |
| 670(1) | WP_150364199.1 | WP_150364199.1 hypothetical protein |
| 670(1) | WP_099093085.1 | WP_099093085.1 hypothetical protein |
| 670(1) | WP_179016426.1 | WP_179016426.1 hypothetical protein |
| 671(1) | WP_276867645.1 | WP_276867645.1 hypothetical protein |
| 671(1) | WP_289613141.1 | WP_289613141.1 hypothetical protein |
| 671(1) | WP_289613138.1 | WP_289613138.1 hypothetical protein |
| 671(1) | WP_102371678.1 | WP_102371678.1 DUF5344 family protein |
| 672(1) | WP_289613132.1 | WP_289613132.1 DUF5050 domain-containing protein |
| 672(1) | WP_270574778.1 | WP_270574778.1 hypothetical protein, partial |
| 672(1) | WP_289613133.1 | WP_289613133.1 zinc-ribbon domain-containing protein |
| 672(1) | WP_289613351.1 | WP_289613351.1 hypothetical protein |
| 673(1) | WP_188535084.1 | WP_188535084.1 hypothetical protein |
| 673(1) | WP_148450936.1 | WP_148450936.1 hypothetical protein |
| 673(1) | WP_290407037.1 | WP_290407037.1 hypothetical protein |
| 673(1) | WP_213506894.1 | WP_213506894.1 hypothetical protein |
| 674(4) | WP_291288252.1 | WP_291288252.1 YebC/PmpR family DNA-binding transcriptional regulator |

| | | |
|---|---|---|
| 675(4) | WP_291288254.1 | WP_291288254.1 DUF2273 domain-containing protein |
| 676(1) | WP_195343914.1 | WP_195343914.1 hypothetical protein |
| 676(1) | WP_291672194.1 | WP_291672194.1 hypothetical protein |
| 676(1) | WP_195542467.1 | WP_195542467.1 hypothetical protein |
| 676(1) | WP_156558888.1 | WP_156558888.1 hypothetical protein |
| 677(1) | WP_195542471.1 | WP_195542471.1 hypothetical protein |
| 677(1) | WP_195343919.1 | WP_195343919.1 hypothetical protein |
| 677(1) | WP_156558882.1 | WP_156558882.1 hypothetical protein |
| 677(1) | WP_291682712.1 | WP_291682712.1 hypothetical protein |
| 678(1) | WP_185897342.1 | WP_185897342.1 FAD:protein FMN transferase |
| 678(1) | WP_157692483.1 | WP_157692483.1 FAD:protein FMN transferase |
| 678(1) | WP_213618810.1 | WP_213618810.1 FAD:protein FMN transferase |
| 678(1) | WP_295629809.1 | WP_295629809.1 alpha/beta fold hydrolase |
| 679(1) | WP_303942506.1 | WP_303942506.1 aminotransferase class V-fold PLP-dependent enzyme |
| 679(1) | WP_276572619.1 | WP_276572619.1 aminotransferase class V-fold PLP-dependent enzyme |
| 679(1) | WP_273383854.1 | WP_273383854.1 aminotransferase class V-fold PLP-dependent enzyme |
| 679(1) | WP_219722302.1 | WP_219722302.1 aminotransferase class V-fold PLP-dependent enzyme |
| 680(1) | WP_303942510.1 | WP_303942510.1 PTS lactose/cellobiose transporter subunit IIA |
| 680(1) | WP_276572620.1 | WP_276572620.1 PTS lactose/cellobiose transporter subunit IIA |
| 680(2) | WP_102372764.1 | WP_102372764.1 PTS lactose/cellobiose transporter subunit IIA |
| 681(1) | WP_303942516.1 | WP_303942516.1 PTS sugar transporter subunit IIB |
| 681(3) | WP_102372761.1 | WP_102372761.1 PTS sugar transporter subunit IIB |
| 682(1) | WP_303942518.1 | WP_303942518.1 acyl-CoA dehydratase activase-related protein |
| 682(1) | WP_102372759.1 | WP_102372759.1 2-hydroxyacyl-CoA dehydratase |
| 682(1) | WP_273383862.1 | WP_273383862.1 acyl-CoA dehydratase activase-related protein |
| 682(1) | WP_273191241.1 | WP_273191241.1 2-hydroxyacyl-CoA dehydratase |
| 683(1) | WP_102372758.1 | WP_102372758.1 amidohydrolase family protein |
| 683(1) | WP_273191242.1 | WP_273191242.1 amidohydrolase family protein |
| 683(1) | WP_303942520.1 | WP_303942520.1 amidohydrolase family protein |
| 683(1) | WP_273383864.1 | WP_273383864.1 amidohydrolase family protein |
| 684(1) | WP_307196917.1 | WP_307196917.1 amidohydrolase |
| 684(1) | WP_235847134.1 | WP_235847134.1 amidohydrolase |
| 684(1) | WP_093291066.1 | WP_093291066.1 amidohydrolase family protein |
| 684(1) | WP_218781305.1 | WP_218781305.1 amidohydrolase |
| 685(1) | WP_306458554.1 | WP_306458554.1 KGG domain-containing protein |
| 685(1) | WP_257029485.1 | WP_257029485.1 general stress protein |
| 685(1) | WP_307192926.1 | WP_307192926.1 general stress protein |
| 685(1) | WP_307312509.1 | WP_307312509.1 general stress protein |
| 686(1) | WP_278773469.1 | WP_278773469.1 hypothetical protein |
| 686(1) | WP_238317197.1 | WP_238317197.1 hypothetical protein |
| 686(1) | WP_307388986.1 | WP_307388986.1 hypothetical protein |
| 686(1) | WP_013252379.1 | WP_013252379.1 hypothetical protein |
| 687(1) | WP_048826605.1 | WP_048826605.1 pyrroline-5-carboxylate reductase |
| 687(1) | WP_204517531.1 | WP_204517531.1 NAD(P)-binding domain-containing protein |
| 687(1) | WP_307589579.1 | WP_307589579.1 pyrroline-5-carboxylate reductase |
| 687(1) | WP_179034798.1 | WP_179034798.1 NAD(P)-binding domain-containing protein |
| 688(1) | WP_006290908.1 | WP_006290908.1 ATP-binding protein |
| 688(1) | WP_281667407.1 | WP_281667407.1 ATP-binding protein |
| 688(1) | WP_307738732.1 | WP_307738732.1 AAA family ATPase |
| 688(1) | WP_006290633.1 | WP_006290633.1 ATP-binding protein |
| 689(1) | WP_242490874.1 | WP_242490874.1 sortase |
| 689(1) | WP_307843896.1 | WP_307843896.1 sortase |
| 689(1) | WP_242447700.1 | WP_242447700.1 sortase |
| 689(1) | WP_242450723.1 | WP_242450723.1 sortase |
| 690(1) | WP_119601518.1 | WP_119601518.1 hypothetical protein |
| 690(1) | WP_307284663.1 | WP_307284663.1 hypothetical protein |
| 690(1) | WP_308082236.1 | WP_308082236.1 hypothetical protein |
| 690(1) | WP_199419775.1 | WP_199419775.1 hypothetical protein |
| 691(1) | WP_083446673.1 | WP_083446673.1 oligoendopeptidase F |
| 691(1) | WP_251610758.1 | WP_251610758.1 oligoendopeptidase F |
| 691(1) | WP_242172398.1 | WP_242172398.1 oligoendopeptidase F |
| 691(1) | WP_308083997.1 | WP_308083997.1 oligoendopeptidase F |
| 692(1) | WP_213143939.1 | WP_213143939.1 helix-hairpin-helix domain-containing protein |
| 692(1) | WP_307852061.1 | WP_307852061.1 helix-hairpin-helix domain-containing protein |
| 692(1) | WP_308170380.1 | WP_308170380.1 helix-hairpin-helix domain-containing protein |
| 692(1) | WP_284037994.1 | WP_284037994.1 helix-hairpin-helix domain-containing protein |
| 693(1) | WP_308615627.1 | WP_308615627.1 Asp23/Gls24 family envelope stress response protein |
| 693(2) | WP_291289942.1 | WP_291289942.1 Asp23/Gls24 family envelope stress response protein |
| 693(1) | WP_291288255.1 | WP_291288255.1 Asp23/Gls24 family envelope stress response protein |
| 694(1) | WP_291288253.1 | WP_291288253.1 alkaline shock response membrane anchor protein AmaP |
| 694(1) | WP_308615628.1 | WP_308615628.1 hypothetical protein |
| 694(1) | WP_297183859.1 | WP_297183859.1 alkaline shock response membrane anchor protein AmaP |
| 694(1) | WP_291289943.1 | WP_291289943.1 alkaline shock response membrane anchor protein AmaP |
| 695(1) | WP_308912803.1 | WP_308912803.1 hypothetical protein |
| 695(1) | WP_155925437.1 | WP_155925437.1 hypothetical protein |
| 695(1) | WP_198310824.1 | WP_198310824.1 hypothetical protein |
| 695(1) | WP_133332987.1 | WP_133332987.1 hypothetical protein |
| 696(3) | WP_001041138.1 | WP_001041138.1 hypothetical protein |
| 697(3) | WP_001990263.1 | WP_001990263.1 phosphotransferase |
| 698(2) | WP_005210483.1 | WP_005210483.1 DegT/DnrJ/EryC1/StrS family aminotransferase |

698(1) WP_005210419.1 WP_005210419.1 aminotransferase class I/II-fold pyridoxal phosphate-dependent enzyme

699(3) WP_006289734.1 WP_006289734.1 type 1 glutamine amidotransferase

700(3) WP_013914175.1 WP_013914175.1 antibiotic biosynthesis monooxygenase

701(3) WP_013914193.1 WP_013914193.1 hypothetical protein

702(1) WP_014368232.1 WP_014368232.1 methyltransferase domain-containing protein
702(1) WP_014649143.1 WP_014649143.1 methyltransferase domain-containing protein
702(1) WP_013914174.1 WP_013914174.1 methyltransferase domain-containing protein

703(1) WP_013914186.1 WP_013914186.1 hypothetical protein
703(1) WP_014368239.1 WP_014368239.1 STM3941 family protein
703(1) WP_014649149.1 WP_014649149.1 STM3941 family protein

704(3) WP_033100006.1 WP_033100006.1 LLM class oxidoreductase

705(3) WP_033100013.1 WP_033100013.1 hypothetical protein

706(3) WP_034095017.1 WP_034095017.1 hypothetical protein

707(2) WP_010963718.1 WP_010963718.1 orotidine 5'-phosphate decarboxylase / HUMPS family protein
707(1) WP_036626796.1 WP_036626796.1 3-hexulose-6-phosphate synthase

708(1) WP_038569590.1 WP_038569590.1 molybdenum cofactor biosynthesis protein MoaE
708(1) WP_039871635.1 WP_039871635.1 molybdenum cofactor biosynthesis protein MoaE
708(1) WP_036684088.1 WP_036684088.1 molybdenum cofactor biosynthesis protein MoaE

709(1) WP_014368231.1 WP_014368231.1 GNAT family N-acetyltransferase
709(1) WP_041616461.1 WP_041616461.1 GNAT family N-acetyltransferase
709(1) WP_014649142.1 WP_014649142.1 GNAT family N-acetyltransferase

710(1) WP_014649147.1 WP_014649147.1 hypothetical protein
710(2) WP_041616462.1 WP_041616462.1 hypothetical protein

711(2) WP_014368238.1 WP_014368238.1 erythromycin esterase family protein
711(1) WP_041616464.1 WP_041616464.1 erythromycin esterase family protein

712(1) WP_028589297.1 WP_028589297.1 GNAT family N-acetyltransferase
712(1) WP_018883277.1 WP_018883277.1 GNAT family N-acetyltransferase
712(1) WP_051421099.1 WP_051421099.1 GNAT family N-acetyltransferase

713(1) WP_066092018.1 WP_066092018.1 NlpC/P60 family protein
713(1) WP_066092015.1 WP_066092015.1 NlpC/P60 family protein
713(1) WP_024027625.1 WP_024027625.1 NlpC/P60 family protein

714(2) WP_053564627.1 WP_053564627.1 hypothetical protein
714(1) WP_069355513.1 WP_069355513.1 hypothetical protein

715(1) WP_058271118.1 WP_058271118.1 phosphoribosylaminoimidazolecarboxamide formyltransferase
715(1) WP_070638668.1 WP_070638668.1 phosphoribosylaminoimidazolecarboxamide formyltransferase
715(1) WP_066022462.1 WP_066022462.1 phosphoribosylaminoimidazolecarboxamide formyltransferase

716(3) WP_076135812.1 WP_076135812.1 metalloregulator ArsR/SmtB family transcription factor

717(3) WP_077832273.1 WP_077832273.1 ECF transporter S component

718(1) WP_077893469.1 WP_077893469.1 hydroxylamine reductase
718(2) WP_077832279.1 WP_077832279.1 hydroxylamine reductase

719(1) WP_077853495.1 WP_077853495.1 zinc ribbon domain-containing protein
719(1) WP_077832276.1 WP_077832276.1 zinc ribbon domain-containing protein
719(1) WP_077893473.1 WP_077893473.1 zinc ribbon domain-containing protein

720(1) WP_077853496.1 WP_077853496.1 hypothetical protein
720(1) WP_077832275.1 WP_077832275.1 hypothetical protein
720(1) WP_077893474.1 WP_077893474.1 hypothetical protein

721(2) WP_080449845.1 WP_080449845.1 hypothetical protein
721(1) WP_074595493.1 WP_074595493.1 hypothetical protein

722(3) WP_081943720.1 WP_081943720.1 thermonuclease family protein

723(1) WP_093291072.1 WP_093291072.1 ATP-grasp domain-containing protein
723(1) WP_093291067.1 WP_093291067.1 ATP-grasp domain-containing protein
723(1) WP_093291068.1 WP_093291068.1 ATP-grasp domain-containing protein

724(2) WP_099361324.1 WP_099361324.1 Fur-regulated basic protein FbpA
724(1) WP_010676883.1 WP_010676883.1 Fur-regulated basic protein FbpA

725(2) WP_109625254.1 WP_109625254.1 threonine/serine exporter family protein
725(1) WP_068673731.1 WP_068673731.1 threonine/serine exporter family protein

726(1) WP_111503204.1 WP_111503204.1 hypothetical protein
726(1) WP_076135816.1 WP_076135816.1 hypothetical protein
726(1) WP_076208252.1 WP_076208252.1 hypothetical protein

727(1) WP_076135811.1 WP_076135811.1 NAD-dependent epimerase/dehydratase family protein
727(1) WP_076208254.1 WP_076208254.1 NAD-dependent epimerase/dehydratase family protein
727(1) WP_111503206.1 WP_111503206.1 NAD-dependent epimerase/dehydratase family protein

728(1) WP_017149765.1 WP_017149765.1 PoNi-like cognate immunity protein
728(1) WP_130810245.1 WP_130810245.1 DUF1911 domain-containing protein
728(1) WP_098303173.1 WP_098303173.1 PoNi-like cognate immunity protein

729(1) WP_116353256.1 WP_116353256.1 hypothetical protein
729(1) WP_099354899.1 WP_099354899.1 hypothetical protein
729(1) WP_136379691.1 WP_136379691.1 hypothetical protein

730(1) WP_099354920.1 WP_099354920.1 cytochrome c
730(1) WP_116353239.1 WP_116353239.1 cytochrome c
730(1) WP_136379710.1 WP_136379710.1 cytochrome c

731(1) WP_139355784.1 WP_139355784.1 hypothetical protein
731(1) WP_077893472.1 WP_077893472.1 hypothetical protein
731(1) WP_077832277.1 WP_077832277.1 hypothetical protein

732(1) WP_143315259.1 WP_143315259.1
732(2) WP_010963357.1 WP_010963357.1 ferredoxin hydrogenase

733(1) WP_109625255.1 WP_109625255.1 threonine/serine exporter family protein
733(1) WP_148409456.1 WP_148409456.1 threonine/serine exporter family protein
733(1) WP_068673728.1 WP_068673728.1 threonine/serine exporter family protein

734(1) WP_108464098.1 WP_108464098.1 hypothetical protein
734(1) WP_090688974.1 WP_090688974.1 hypothetical protein
734(1) WP_162145048.1 WP_162145048.1 hypothetical protein

735(1) WP_146032338.1 WP_146032338.1 hypothetical protein
735(1) WP_146753362.1 WP_146753362.1 hypothetical protein
735(1) WP_165264001.1 WP_165264001.1 hypothetical protein

736(1) WP_152381384.1 WP_152381384.1 hypothetical protein
736(1) WP_169505039.1 WP_169505039.1 hypothetical protein
736(1) WP_165147500.1 WP_165147500.1 hypothetical protein

737(2) WP_010963358.1 WP_010963358.1 YiiX/YebB-like N1pC/P60 family cysteine hydrolase
737(1) WP_181737019.1 WP_181737019.1 YiiX/YebB-like N1pC/P60 family cysteine hydrolase

738(1) WP_024632674.1 WP_024632674.1 glycoside hydrolase family 88 protein
738(1) WP_076113444.1 WP_076113444.1 glycoside hydrolase family 88 protein
738(1) WP_183600885.1 WP_183600885.1 glycoside hydrolase family 88 protein

739(3) WP_191965775.1 WP_191965775.1 hypothetical protein

740(2) WP_006289737.1 WP_006289737.1 ornithine carbamoyltransferase
740(1) WP_192808052.1 WP_192808052.1 ornithine carbamoyltransferase

741(1) WP_083875352.1 WP_083875352.1 right-handed parallel beta-helix repeat-containing protein
741(1) WP_005210469.1 WP_005210469.1 hypothetical protein
741(1) WP_195244540.1 WP_195244540.1 glycosyl hydrolase family 28-related protein

742(3) WP_195343927.1 WP_195343927.1 TIGR04197 family type VII secretion effector

743(1) WP_195542477.1 WP_195542477.1 hypothetical protein
743(2) WP_156558875.1 WP_156558875.1 hypothetical protein

744(1) WP_198056464.1 WP_198056464.1 PHB depolymerase family esterase
744(2) WP_033100012.1 WP_033100012.1 PHB depolymerase family esterase

745(1) WP_198056470.1 WP_198056470.1 PhzF family phenazine biosynthesis protein
745(1) WP_033100003.1 WP_033100003.1 PhzF family phenazine biosynthesis protein
745(1) WP_198059696.1 WP_198059696.1 PhzF family phenazine biosynthesis protein

746(1) WP_198310843.1 WP_198310843.1 hypothetical protein
746(1) WP_035423706.1 WP_035423706.1 hypothetical protein
746(1) WP_191272889.1 WP_191272889.1 hypothetical protein

747(1) WP_112331589.1 WP_112331589.1 hypothetical protein
747(1) WP_195974880.1 WP_195974880.1 hypothetical protein
747(1) WP_204445225.1 WP_204445225.1 hypothetical protein

748(2) WP_112331582.1 WP_112331582.1 zinc-ribbon domain-containing protein
748(1) WP_204445239.1 WP_204445239.1 hypothetical protein

749(1) WP_204666323.1 WP_204666323.1 cellulase family glycosylhydrolase
749(1) WP_144687303.1 WP_144687303.1 cellulase family glycosylhydrolase
749(1) WP_102371694.1 WP_102371694.1 cellulase family glycosylhydrolase

750(1) WP_040590335.1 WP_040590335.1 hypothetical protein
750(1) WP_022349193.1 WP_022349193.1 hypothetical protein
750(1) WP_204666334.1 WP_204666334.1 hypothetical protein

751(1) WP_204666335.1 WP_204666335.1 DUF5344 family protein
751(1) WP_006292356.1 WP_006292356.1 DUF5344 family protein
751(1) WP_022349194.1 WP_022349194.1 DUF5344 family protein

752(1) WP_206094027.1 WP_206094027.1 methyltransferase domain-containing protein
752(2) WP_087433616.1 WP_087433616.1 methyltransferase domain-containing protein

753(1) WP_211237321.1 WP_211237321.1 hypothetical protein
753(1) WP_018883263.1 WP_018883263.1 hypothetical protein
753(1) WP_211237322.1 WP_211237322.1 hypothetical protein

754(1) WP_211745837.1 WP_211745837.1 CGNR zinc finger domain-containing protein
754(2) WP_170862374.1 WP_170862374.1 CGNR zinc finger domain-containing protein

755(3) WP_213117468.1 WP_213117468.1 hypothetical protein

756(1) WP_068619129.1 WP_068619129.1 GNAT family N-acetyltransferase
756(1) WP_127199762.1 WP_127199762.1 GNAT family N-acetyltransferase
756(1) WP_213618807.1 WP_213618807.1 GNAT family N-acetyltransferase

757(1) WP_068619128.1 WP_068619128.1 hypothetical protein
757(1) WP_213618809.1 WP_213618809.1 hypothetical protein
757(1) WP_127199764.1 WP_127199764.1 hypothetical protein

758(1) WP_068619122.1 WP_068619122.1 ferric reductase-like transmembrane domain-containing protein
758(1) WP_213618811.1 WP_213618811.1 ferric reductase-like transmembrane domain-containing protein

758(1) WP_127199766.1 WP_127199766.1 ferric reductase-like transmembrane domain-containing protein

759(1) WP_216518249.1 WP_216518249.1 hypothetical protein
759(1) WP_150364202.1 WP_150364202.1 hypothetical protein
759(1) WP_145406448.1 WP_145406448.1 hypothetical protein

760(2) WP_219872443.1 WP_219872443.1 FAD-dependent monooxygenase
760(1) WP_139867402.1 WP_139867402.1 FAD-dependent monooxygenase

761(1) WP_074085367.1 WP_074085367.1 5-methyltetrahydropteroyltriglutamate--homocysteine S-methyltransferase
761(1) WP_223067427.1 WP_223067427.1 5-methyltetrahydropteroyltriglutamate--homocysteine S-methyltransferase
761(1) WP_074108303.1 WP_074108303.1 5-methyltetrahydropteroyltriglutamate--homocysteine S-methyltransferase

762(1) WP_136379689.1 WP_136379689.1 beta-galactosidase
762(1) WP_229140037.1 WP_229140037.1 beta-galactosidase
762(1) WP_159440011.1 WP_159440011.1 beta-galactosidase

763(1) WP_136379731.1 WP_136379731.1 glycosyl hydrolase 53 family protein
763(1) WP_229140038.1 WP_229140038.1 glycosyl hydrolase 53 family protein
763(1) WP_150457753.1 WP_150457753.1 glycosyl hydrolase 53 family protein

764(1) WP_231094865.1 WP_231094865.1 MMPL family transporter
764(1) WP_046522795.1 WP_046522795.1 MMPL family transporter
764(1) WP_053361012.1 WP_053361012.1 MMPL family transporter

765(1) WP_231097269.1 WP_231097269.1 Lrp/AsnC family transcriptional regulator
765(1) WP_017553132.1 WP_017553132.1 Lrp/AsnC family transcriptional regulator
765(1) WP_179093399.1 WP_179093399.1 Lrp/AsnC family transcriptional regulator

766(1) WP_087194490.1 WP_087194490.1 selenide, water dikinase SelD
766(1) WP_235286193.1 WP_235286193.1 selenide, water dikinase SelD
766(1) WP_111147810.1 WP_111147810.1 selenide, water dikinase SelD

767(1) WP_182301833.1 WP_182301833.1 hypothetical protein
767(1) WP_010250262.1 WP_010250262.1 DUF4163 domain-containing protein
767(1) WP_239617474.1 WP_239617474.1 hypothetical protein

768(2) WP_021624826.1 WP_021624826.1 hypothetical protein
768(1) WP_240961477.1 WP_240961477.1 hypothetical protein

769(1) WP_053403772.1 WP_053403772.1 YwdI family protein
769(2) WP_242172354.1 WP_242172354.1 YwdI family protein

770(2) WP_242172356.1 WP_242172356.1 uracil-DNA glycosylase
770(1) WP_053403773.1 WP_053403773.1 uracil-DNA glycosylase

771(1) WP_053403794.1 WP_053403794.1 cytosine permease
771(2) WP_242172399.1 WP_242172399.1 cytosine permease

772(1) WP_242951618.1 WP_242951618.1 jacalin-like lectin
772(1) WP_099517108.1 WP_099517108.1 jacalin-like lectin
772(1) WP_242950780.1 WP_242950780.1 jacalin-like lectin

773(1) WP_136789986.1 WP_136789986.1 pyroglutamyl-peptidase I
773(1) WP_191272873.1 WP_191272873.1 pyroglutamyl-peptidase I
773(1) WP_243495547.1 WP_243495547.1 pyroglutamyl-peptidase I

774(1) WP_243495553.1 WP_243495553.1 hypothetical protein
774(2) WP_098248517.1 WP_098248517.1 hypothetical protein

775(1) WP_243495565.1 WP_243495565.1 MraY family glycosyltransferase
775(1) WP_068673724.1 WP_068673724.1 MraY family glycosyltransferase
775(1) WP_041580746.1 WP_041580746.1 MraY family glycosyltransferase

776(1) WP_244525356.1 WP_244525356.1 hypothetical protein
776(2) WP_244525359.1 WP_244525359.1 hypothetical protein

777(1) WP_245368172.1 WP_245368172.1 acetyl-CoA C-acetyltransferase
777(1) WP_209365666.1 WP_209365666.1 thiolase family protein
777(1) WP_122971527.1 WP_122971527.1 acetyl-CoA C-acetyltransferase

778(1) WP_247610933.1 WP_247610933.1 hypothetical protein
778(2) WP_237657536.1 WP_237657536.1 hypothetical protein

779(2) WP_041903714.1 WP_041903714.1 restriction endonuclease
779(1) WP_251245820.1 WP_251245820.1 restriction endonuclease

780(1) WP_120462962.1 WP_120462962.1 hypothetical protein
780(1) WP_251417684.1 WP_251417684.1 hypothetical protein
780(1) WP_126996331.1 WP_126996331.1 hypothetical protein

781(1) WP_126996333.1 WP_126996333.1 DMT family transporter
781(1) WP_251417686.1 WP_251417686.1 DMT family transporter
781(1) WP_120462940.1 WP_120462940.1 EamA family transporter

782(1) WP_110931553.1 WP_110931553.1 TIGR02206 family membrane protein
782(1) WP_126996339.1 WP_126996339.1 TIGR02206 family membrane protein
782(1) WP_251417691.1 WP_251417691.1 TIGR02206 family membrane protein

783(1) WP_110931554.1 WP_110931554.1 mismatch-specific DNA-glycosylase
783(1) WP_120465215.1 WP_120465215.1 mismatch-specific DNA-glycosylase
783(1) WP_251417693.1 WP_251417693.1 mismatch-specific DNA-glycosylase

784(2) WP_251426356.1 WP_251426356.1 alkaline phosphatase
784(1) WP_175492871.1 WP_175492871.1 alkaline phosphatase

785(1) WP_046522781.1 WP_046522781.1 DUF839 domain-containing protein
785(2) WP_251426370.1 WP_251426370.1 DUF839 domain-containing protein

786(2) WP_251426378.1 WP_251426378.1 efflux RND transporter periplasmic adaptor subunit

786(1) WP_068673736.1 WP_068673736.1 efflux RND transporter periplasmic adaptor subunit

787(1) WP_191272846.1 WP_191272846.1 hypothetical protein
787(1) WP_251466813.1 WP_251466813.1 hypothetical protein
787(1) WP_199419787.1 WP_199419787.1 hypothetical protein

788(1) WP_251548770.1 WP_251548770.1 DUF5317 domain-containing protein
788(1) WP_042462823.1 WP_042462823.1 DUF5317 domain-containing protein
788(1) WP_045518289.1 WP_045518289.1 DUF5317 domain-containing protein

789(1) WP_211745836.1 WP_211745836.1 DinB family protein
789(1) WP_251583279.1 WP_251583279.1 DinB family protein
789(1) WP_091155356.1 WP_091155356.1 DinB family protein

790(1) WP_251596648.1 WP_251596648.1 pyruvate kinase
790(1) WP_166244070.1 WP_166244070.1 pyruvate kinase
790(1) WP_128631300.1 WP_128631300.1 pyruvate kinase

791(1) WP_087194492.1 WP_087194492.1 sulfurtransferase-like selenium metabolism protein YedF
791(1) WP_128631299.1 WP_128631299.1 DsrE family protein
791(1) WP_251596650.1 WP_251596650.1 hypothetical protein

792(1) WP_242172351.1 WP_242172351.1 spore coat protein GerQ
792(1) WP_053403770.1 WP_053403770.1 spore coat protein GerQ
792(1) WP_251610735.1 WP_251610735.1 spore coat protein GerQ

793(1) WP_251610737.1 WP_251610737.1 DUF423 domain-containing protein
793(2) WP_053403771.1 WP_053403771.1 DUF423 domain-containing protein

794(1) WP_242172389.1 WP_242172389.1 NAD(P)H-hydrate dehydratase
794(1) WP_053403791.1 WP_053403791.1 bifunctional ADP-dependent NAD(P)H-hydrate dehydratase/NAD(P)H-hydrate epimerase
794(1) WP_251610755.1 WP_251610755.1 NAD(P)H-hydrate dehydratase

795(1) WP_251610756.1 WP_251610756.1 endonuclease/exonuclease/phosphatase family protein
795(1) WP_053403793.1 WP_053403793.1 endonuclease/exonuclease/phosphatase family protein
795(1) WP_242172393.1 WP_242172393.1 endonuclease/exonuclease/phosphatase family protein

796(1) WP_257391743.1 WP_257391743.1 hypothetical protein
796(1) WP_211082930.1 WP_211082930.1 hypothetical protein
796(1) WP_113672699.1 WP_113672699.1 hypothetical protein

797(1) WP_258296276.1 WP_258296276.1 hypothetical protein
797(1) WP_071640592.1 WP_071640592.1 hypothetical protein
797(1) WP_220027686.1 WP_220027686.1 hypothetical protein

798(3) WP_261363789.1 WP_261363789.1 hypothetical protein

799(1) WP_003516592.1 WP_003516592.1 ribonucleoside triphosphate reductase
799(1) WP_069193862.1 WP_069193862.1 ribonucleoside triphosphate reductase
799(1) WP_265444518.1 WP_265444518.1 ribonucleoside triphosphate reductase

800(1) WP_058271094.1 WP_058271094.1 hypothetical protein
800(1) WP_210388947.1 WP_210388947.1 energy-coupling factor transporter transmembrane component T
800(1) WP_268872530.1 WP_268872530.1 energy-coupling factor transporter transmembrane component T

801(1) WP_270472025.1 WP_270472025.1 hypothetical protein
801(1) WP_270472027.1 WP_270472027.1 hypothetical protein
801(1) WP_270472023.1 WP_270472023.1 hypothetical protein

802(2) WP_165643409.1 WP_165643409.1 hypothetical protein
802(1) WP_272045554.1 WP_272045554.1 hypothetical protein

803(1) WP_098579104.1 WP_098579104.1 VanZ family protein
803(1) WP_182030410.1 WP_182030410.1 VanZ family protein
803(1) WP_272801667.1 WP_272801667.1 VanZ family protein

804(1) WP_098248523.1 WP_098248523.1 YjfB family protein
804(1) WP_272801679.1 WP_272801679.1 YjfB family protein
804(1) WP_136789985.1 WP_136789985.1 YjfB family protein

805(1) WP_015595765.1 WP_015595765.1 YigZ family protein
805(1) WP_232274502.1 WP_232274502.1 YigZ family protein
805(1) WP_273061507.1 WP_273061507.1 YigZ family protein

806(1) WP_273321555.1 WP_273321555.1 Imm74 family immunity protein
806(1) WP_113674463.1 WP_113674463.1 Imm74 family immunity protein
806(1) WP_099329540.1 WP_099329540.1 Imm74 family immunity protein

807(1) WP_273321558.1 WP_273321558.1 hypothetical protein
807(1) WP_212693236.1 WP_212693236.1 hypothetical protein
807(1) WP_113672697.1 WP_113672697.1 hypothetical protein

808(1) WP_102372741.1 WP_102372741.1 DUF6020 family protein
808(1) WP_273383887.1 WP_273383887.1 DUF6020 family protein
808(1) WP_273191267.1 WP_273191267.1 DUF6020 family protein

809(1) WP_274485159.1 WP_274485159.1 hypothetical protein
809(1) WP_161626997.1 WP_161626997.1 hypothetical protein
809(1) WP_254349531.1 WP_254349531.1 hypothetical protein

810(1) WP_102371683.1 WP_102371683.1 molecular chaperone HtpG
810(1) WP_204666331.1 WP_204666331.1 molecular chaperone HtpG
810(1) WP_276867652.1 WP_276867652.1 molecular chaperone HtpG

811(1) WP_102371688.1 WP_102371688.1 serine hydrolase
811(1) WP_204666401.1 WP_204666401.1 serine hydrolase
811(1) WP_276867667.1 WP_276867667.1 serine hydrolase

812(1) WP_204666326.1 WP_204666326.1 16S rRNA (adenine(1518)-N(6)/adenine(1519)-N(6))-dimethyltransferase RsmA

812(1) WP_276867675.1 WP_276867675.1 16S rRNA (adenine(1518)-N(6)/adenine(1519)-N(6))-dimethyltransferase RsmA
812(1) WP_102371691.1 WP_102371691.1 16S rRNA (adenine(1518)-N(6)/adenine(1519)-N(6))-dimethyltransferase RsmA

813(1) WP_276867677.1 WP_276867677.1 16S rRNA (cytidine(1402)-2'-O)-methyltransferase
813(1) WP_102371693.1 WP_102371693.1 16S rRNA (cytidine(1402)-2'-O)-methyltransferase
813(1) WP_204666324.1 WP_204666324.1 16S rRNA (cytidine(1402)-2'-O)-methyltransferase

814(1) WP_152595383.1 WP_152595383.1 hypothetical protein
814(1) WP_277157946.1 WP_277157946.1 hypothetical protein
814(1) WP_033521262.1 WP_033521262.1 hypothetical protein

815(1) WP_277157964.1 WP_277157964.1 aminoglycoside phosphotransferase
815(1) WP_277175597.1 WP_277175597.1 aminoglycoside phosphotransferase
815(1) WP_277142111.1 WP_277142111.1 aminoglycoside phosphotransferase

816(1) WP_013914180.1 WP_013914180.1 alpha-xylosidase
816(1) WP_016362247.1 WP_016362247.1 alpha-xylosidase
816(1) WP_277713348.1 WP_277713348.1 alpha-xylosidase

817(1) WP_278773453.1 WP_278773453.1 FtsK/SpoIIIE domain-containing protein
817(1) WP_036575561.1 WP_036575561.1 FtsK/SpoIIIE domain-containing protein
817(1) WP_013252375.1 WP_013252375.1 FtsK/SpoIIIE domain-containing protein

818(1) WP_036575563.1 WP_036575563.1 hypothetical protein
818(1) WP_278773455.1 WP_278773455.1 hypothetical protein
818(1) WP_013252374.1 WP_013252374.1 hypothetical protein

819(1) WP_013252371.1 WP_013252371.1 hypothetical protein
819(1) WP_162148303.1 WP_162148303.1 hypothetical protein
819(1) WP_278773459.1 WP_278773459.1 hypothetical protein

820(1) WP_278773472.1 WP_278773472.1 hypothetical protein
820(1) WP_013252378.1 WP_013252378.1 hypothetical protein
820(1) WP_278773466.1 WP_278773466.1 hypothetical protein

821(1) WP_006290907.1 WP_006290907.1 Fic family protein
821(1) WP_006289733.1 WP_006289733.1 Fic family protein
821(1) WP_281667408.1 WP_281667408.1 Fic family protein

822(1) WP_006289736.1 WP_006289736.1 arginine deiminase
822(1) WP_281667410.1 WP_281667410.1 arginine deiminase
822(1) WP_036695819.1 WP_036695819.1 arginine deiminase

823(1) WP_281667411.1 WP_281667411.1 ribonuclease III
823(2) WP_006289738.1 WP_006289738.1 ribonuclease III

824(1) WP_126996329.1 WP_126996329.1 hypothetical protein
824(1) WP_283904336.1 WP_283904336.1 RNA 2'-phosphotransferase
824(1) WP_068536395.1 WP_068536395.1 RNA 2'-phosphotransferase

825(1) WP_283927241.1 WP_283927241.1 FHA domain-containing protein
825(1) WP_213592842.1 WP_213592842.1 FHA domain-containing protein
825(1) WP_155609742.1 WP_155609742.1 FHA domain-containing protein

826(1) WP_283928356.1 WP_283928356.1 right-handed parallel beta-helix repeat-containing protein
826(2) WP_230873712.1 WP_230873712.1 right-handed parallel beta-helix repeat-containing protein

827(1) WP_217035873.1 WP_217035873.1 SRPBCC family protein
827(1) WP_213117492.1 WP_213117492.1 SRPBCC family protein
827(1) WP_284036491.1 WP_284036491.1 SRPBCC family protein

828(1) WP_284036504.1 WP_284036504.1 hypothetical protein
828(2) WP_213117465.1 WP_213117465.1 hypothetical protein

829(1) WP_285748163.1 WP_285748163.1 ABC-2 transporter permease
829(1) WP_191798516.1 WP_191798516.1 ABC-2 transporter permease
829(1) WP_213532646.1 WP_213532646.1 ABC-2 transporter permease

830(1) WP_286124790.1 WP_286124790.1 hypothetical protein
830(1) WP_073008236.1 WP_073008236.1 hypothetical protein
830(1) WP_026593057.1 WP_026593057.1 hypothetical protein

831(1) WP_068673734.1 WP_068673734.1 efflux RND transporter permease subunit
831(1) WP_289342253.1 WP_289342253.1 efflux RND transporter permease subunit
831(1) WP_251548777.1 WP_251548777.1 efflux RND transporter permease subunit

832(1) WP_148450934.1 WP_148450934.1 protein-glutamine gamma-glutamyltransferase
832(1) WP_290407035.1 WP_290407035.1 protein-glutamine gamma-glutamyltransferase
832(1) WP_211022778.1 WP_211022778.1 protein-glutamine gamma-glutamyltransferase

833(1) WP_191272897.1 WP_191272897.1 protein-glutamate O-methyltransferase CheR
833(2) WP_295628290.1 WP_295628290.1 chemotaxis protein CheB

834(1) WP_256337824.1 WP_256337824.1 hypothetical protein
834(1) WP_302847970.1 WP_302847970.1 hypothetical protein
834(1) WP_256336116.1 WP_256336116.1 hypothetical protein

835(1) WP_036944577.1 WP_036944577.1 accessory Sec system translocase SecA2
835(1) WP_304216840.1 WP_304216840.1 preprotein translocase subunit SecA
835(1) WP_099361322.1 WP_099361322.1 preprotein translocase subunit SecA

836(1) WP_179160633.1 WP_179160633.1 adenine deaminase C-terminal domain-containing protein
836(1) WP_034675346.1 WP_034675346.1 adenine deaminase C-terminal domain-containing protein
836(1) WP_307192903.1 WP_307192903.1 adenine deaminase C-terminal domain-containing protein

837(1) WP_307312514.1 WP_307312514.1 hypothetical protein
837(1) WP_179603493.1 WP_179603493.1 hypothetical protein
837(1) WP_306077405.1 WP_306077405.1 hypothetical protein

838(1) WP_307394666.1 WP_307394666.1 hypothetical protein
838(2) WP_149051231.1 WP_149051231.1 hypothetical protein

839(1) WP_005604657.1 WP_005604657.1 5'-methylthioadenosine/adenosylhomocysteine nucleosidase
839(1) WP_307284646.1 WP_307284646.1 5'-methylthioadenosine/S-adenosylhomocysteine nucleosidase
839(1) WP_308082226.1 WP_308082226.1 5'-methylthioadenosine/S-adenosylhomocysteine nucleosidase

840(1) WP_308082232.1 WP_308082232.1 hypothetical protein
840(1) WP_307284654.1 WP_307284654.1 hypothetical protein
840(1) WP_223592696.1 WP_223592696.1 hypothetical protein

841(1) WP_308082235.1 WP_308082235.1 hypothetical protein
841(1) WP_119601516.1 WP_119601516.1 hypothetical protein
841(1) WP_307284660.1 WP_307284660.1 hypothetical protein

842(2) WP_074085362.1 WP_074085362.1 hypothetical protein
842(1) WP_308083977.1 WP_308083977.1 hypothetical protein

843(2) WP_155609741.1 WP_155609741.1 FHA domain-containing protein
843(1) WP_308083993.1 WP_308083993.1 FHA domain-containing protein

844(1) WP_053361028.1 WP_053361028.1 ABC transporter permease
844(1) WP_149871692.1 WP_149871692.1 ABC transporter permease
844(1) WP_308110122.1 WP_308110122.1 ABC transporter permease

845(1) WP_308110125.1 WP_308110125.1 hypothetical protein
845(1) WP_193747721.1 WP_193747721.1 hypothetical protein
845(1) WP_190284713.1 WP_190284713.1 hypothetical protein

846(1) WP_026572737.1 WP_026572737.1 hypothetical protein
846(1) WP_308912810.1 WP_308912810.1 hypothetical protein
846(1) WP_133332979.1 WP_133332979.1 hypothetical protein

847(1) WP_309098599.1 WP_309098599.1 dihydrodipicolinate synthase family protein
847(1) WP_263324530.1 WP_263324530.1 dihydrodipicolinate synthase family protein
847(1) WP_199613887.1 WP_199613887.1 5-dehydro-4-deoxyglucarate dehydratase

848(1) WP_048826621.1 WP_048826621.1 N-acetylmannosamine-6-phosphate 2-epimerase
848(1) WP_204412744.1 WP_204412744.1 N-acetylmannosamine-6-phosphate 2-epimerase
848(1) WP_309098600.1 WP_309098600.1 N-acetylmannosamine-6-phosphate 2-epimerase

849(1) WP_002031667.1 WP_002031667.1 hypothetical protein
849(1) WP_002075027.1 WP_002075027.1 hypothetical protein

850(2) WP_002118913.1 WP_002118913.1 hypothetical protein

851(2) WP_005210423.1 WP_005210423.1 sigma-70 family RNA polymerase sigma factor

852(2) WP_005210471.1 WP_005210471.1 GNAT family N-acetyltransferase

853(2) WP_005210479.1 WP_005210479.1 oligosaccharide flippase family protein

854(1) WP_006289756.1 WP_006289756.1 CRISPR-associated endonuclease Cas2
854(1) WP_006290890.1 WP_006290890.1 CRISPR-associated endonuclease Cas2

855(1) WP_006292349.1 WP_006292349.1 hypothetical protein
855(1) WP_007147980.1 WP_007147980.1 hypothetical protein

856(2) WP_010963354.1 WP_010963354.1 dCTP deaminase

857(2) WP_010963355.1 WP_010963355.1 alpha/beta hydrolase

858(2) WP_010963356.1 WP_010963356.1 orotate phosphoribosyltransferase

859(2) WP_010963359.1 WP_010963359.1 hypothetical protein

860(2) WP_010963360.1 WP_010963360.1 phosphatidylserine decarboxylase

861(2) WP_010963362.1 WP_010963362.1 AarF/ABC1/UbiB kinase family protein

862(2) WP_010963363.1 WP_010963363.1 phasin family protein

863(2) WP_010963375.1 WP_010963375.1 hypothetical protein

864(2) WP_010963381.1 WP_010963381.1 hypothetical protein

865(2) WP_010963724.1 WP_010963724.1 AsnC family transcriptional regulator

866(2) WP_010963743.1 WP_010963743.1 ECF transporter S component

867(2) WP_013252372.1 WP_013252372.1 hypothetical protein

868(2) WP_013252373.1 WP_013252373.1 hypothetical protein

869(1) WP_015595772.1 WP_015595772.1 glycosyltransferase family 2 protein
869(1) WP_015595773.1 WP_015595773.1 glycosyltransferase family 2 protein

870(2) WP_019424135.1 WP_019424135.1 5'-nucleotidase

871(2) WP_022349200.1 WP_022349200.1 hypothetical protein

872(1) WP_026478111.1 WP_026478111.1 hypothetical protein
872(1) WP_026478112.1 WP_026478112.1 hypothetical protein, partial

873(1) WP_026882392.1 WP_026882392.1 hypothetical protein
873(1) WP_026882389.1 WP_026882389.1 hypothetical protein

874(1) WP_018883265.1 WP_018883265.1 HEAT repeat domain-containing protein
874(1) WP_028589289.1 WP_028589289.1 HEAT repeat domain-containing protein

875(2) WP_038245440.1 WP_038245440.1 hypothetical protein

876(2) WP_040418521.1 WP_040418521.1 acyltransferase

877(1) WP_007147973.1 WP_007147973.1 alpha/beta fold hydrolase
877(1) WP_040590338.1 WP_040590338.1 alpha/beta fold hydrolase

878(1) WP_034675329.1 WP_034675329.1 glycerol-3-phosphate dehydrogenase/oxidase
878(1) WP_049869178.1 WP_049869178.1 glycerol-3-phosphate dehydrogenase/oxidase

879(1) WP_018883286.1 WP_018883286.1 aminoglycoside phosphotransferase family protein
879(1) WP_051289610.1 WP_051289610.1 aminoglycoside phosphotransferase family protein

880(1) WP_043067149.1 WP_043067149.1 esterase-like activity of phytase family protein
880(1) WP_052520274.1 WP_052520274.1 esterase-like activity of phytase family protein

881(1) WP_053361031.1 WP_053361031.1 hypothetical protein
881(1) WP_046522780.1 WP_046522780.1 hypothetical protein

882(1) WP_046522773.1 WP_046522773.1 sugar phosphate isomerase/epimerase
882(1) WP_053361036.1 WP_053361036.1 sugar phosphate isomerase/epimerase

883(2) WP_057897610.1 WP_057897610.1 NAD-dependent malic enzyme

884(2) WP_057897612.1 WP_057897612.1 hypothetical protein

885(2) WP_057897615.1 WP_057897615.1 hypothetical protein

886(2) WP_057897616.1 WP_057897616.1 NfeD family protein

887(1) WP_066092024.1 WP_066092024.1 YheC/YheD family protein
887(1) WP_066092041.1 WP_066092041.1 YheC/YheD family protein

888(1) WP_068673716.1 WP_068673716.1 hypothetical protein
888(1) WP_068673718.1 WP_068673718.1 hypothetical protein

889(2) WP_068782402.1 WP_068782402.1 DUF3349 domain-containing protein

890(1) WP_070638628.1 WP_070638628.1 GNAT family N-acetyltransferase
890(1) WP_058271096.1 WP_058271096.1 GNAT family N-acetyltransferase

891(1) WP_058271099.1 WP_058271099.1 orotate phosphoribosyltransferase
891(1) WP_070638634.1 WP_070638634.1 orotate phosphoribosyltransferase

892(1) WP_070638640.1 WP_070638640.1 sulfide/dihydroorotate dehydrogenase-like FAD/NAD-binding protein
892(1) WP_058271101.1 WP_058271101.1 sulfide/dihydroorotate dehydrogenase-like FAD/NAD-binding protein

893(1) WP_070638666.1 WP_070638666.1 hypothetical protein
893(1) WP_058271117.1 WP_058271117.1 hypothetical protein

894(1) WP_070638674.1 WP_070638674.1 peptide deformylase
894(1) WP_058271122.1 WP_058271122.1 peptide deformylase

895(1) WP_070638791.1 WP_070638791.1 IMP cyclohydrolase
895(1) WP_058271119.1 WP_058271119.1 inosine monophosphate cyclohydrolase

896(1) WP_074085373.1 WP_074085373.1 MarR family transcriptional regulator
896(1) WP_074102535.1 WP_074102535.1 MarR family transcriptional regulator

897(1) WP_074085361.1 WP_074085361.1 hypothetical protein
897(1) WP_074108299.1 WP_074108299.1 hypothetical protein

898(1) WP_074085365.1 WP_074085365.1 peptide-methionine (R)-S-oxide reductase MsrB
898(1) WP_074108301.1 WP_074108301.1 peptide-methionine (R)-S-oxide reductase MsrB

899(1) WP_074108302.1 WP_074108302.1 DUF4349 domain-containing protein
899(1) WP_074085366.1 WP_074085366.1 DUF4349 domain-containing protein

900(1) WP_074108305.1 WP_074108305.1 FtsX-like permease family protein
900(1) WP_074085370.1 WP_074085370.1 FtsX-like permease family protein

901(1) WP_038584761.1 WP_038584761.1 hypothetical protein
901(1) WP_076113442.1 WP_076113442.1 hypothetical protein

902(1) WP_038584758.1 WP_038584758.1 LytTR family DNA-binding domain-containing protein
902(1) WP_076113443.1 WP_076113443.1 LytTR family DNA-binding domain-containing protein

903(1) WP_076113458.1 WP_076113458.1 N,N'-diacetylchitobiose phosphorylase
903(1) WP_038584717.1 WP_038584717.1 N,N'-diacetylchitobiose phosphorylase

904(1) WP_038584714.1 WP_038584714.1 hypothetical protein
904(1) WP_076113463.1 WP_076113463.1 hypothetical protein

905(1) WP_076113466.1 WP_076113466.1 putative DNA modification/repair radical SAM protein
905(1) WP_063848062.1 WP_063848062.1 putative DNA modification/repair radical SAM protein

906(1) WP_076242348.1 WP_076242348.1 MBL fold metallo-hydrolase
906(1) WP_042134250.1 WP_042134250.1 MBL fold metallo-hydrolase

907(1) WP_081442773.1 WP_081442773.1 tRNA guanosine(34) transglycosylase Tgt
907(1) WP_007147979.1 WP_007147979.1 tRNA guanosine(34) transglycosylase Tgt

908(1) WP_058271097.1 WP_058271097.1 hypothetical protein
908(1) WP_083307170.1 WP_083307170.1 hypothetical protein

909(1) WP_083675870.1 WP_083675870.1 TIGR03915 family putative DNA repair protein
909(1) WP_081965640.1 WP_081965640.1 TIGR03915 family putative DNA repair protein

910(1) WP_089522628.1 WP_089522628.1 IS4 family transposase
910(1) WP_089522625.1 WP_089522625.1 IS4 family transposase

911(1) WP_093291058.1 WP_093291058.1 hypothetical protein
911(1) WP_093291044.1 WP_093291044.1 hypothetical protein

912(1) WP_072336501.1 WP_072336501.1 hypothetical protein
912(1) WP_093291061.1 WP_093291061.1 hypothetical protein

913(1) WP_072336504.1 WP_072336504.1 conjugal transfer protein
913(1) WP_093291062.1 WP_093291062.1 conjugal transfer protein

914(1) WP_073008291.1 WP_073008291.1 radical SAM protein
914(1) WP_093291063.1 WP_093291063.1 radical SAM protein

915(1) WP_068694710.1 WP_068694710.1 glycoside hydrolase family 38 C-terminal domain-containing protein
915(1) WP_094092616.1 WP_094092616.1 glycoside hydrolase family 38 C-terminal domain-containing protein

916(1) WP_068694711.1 WP_068694711.1 hypothetical protein
916(1) WP_094092617.1 WP_094092617.1 hypothetical protein

917(1) WP_068694712.1 WP_068694712.1 glutaminase family protein
917(1) WP_094092618.1 WP_094092618.1 glutaminase family protein

918(2) WP_094870387.1 WP_094870387.1 MmcQ/YjbR family DNA-binding protein

919(2) WP_097898659.1 WP_097898659.1 hypothetical protein

920(2) WP_097898664.1 WP_097898664.1 thioredoxin family protein

921(2) WP_097898666.1 WP_097898666.1 hypothetical protein

922(2) WP_097898669.1 WP_097898669.1 YmaF family protein

923(2) WP_099361320.1 WP_099361320.1 ribosome-associated translation inhibitor RaiA

924(2) WP_102997923.1 WP_102997923.1 hypothetical protein

925(1) WP_104030455.1 WP_104030455.1 aromatic amino acid hydroxylase
925(1) WP_104148786.1 WP_104148786.1 aromatic amino acid hydroxylase

926(1) WP_105368171.1 WP_105368171.1 M4 family metallopeptidase
926(1) WP_101300022.1 WP_101300022.1 M4 family metallopeptidase

927(1) WP_006292330.1 WP_006292330.1 class II fructose-bisphosphate aldolase
927(1) WP_106496010.1 WP_106496010.1 fructose-bisphosphate aldolase

928(1) WP_108464095.1 WP_108464095.1 hypothetical protein
928(1) WP_010250289.1 WP_010250289.1 hypothetical protein

929(2) WP_109625231.1 WP_109625231.1 30S ribosomal protein S12

930(2) WP_109625232.1 WP_109625232.1 DNA-directed RNA polymerase subunit beta'

931(2) WP_109625233.1 WP_109625233.1 DNA-directed RNA polymerase subunit beta

932(2) WP_109625234.1 WP_109625234.1 50S ribosomal protein L7/L12

933(2) WP_109625235.1 WP_109625235.1 50S ribosomal protein L10

934(2) WP_109625236.1 WP_109625236.1 carbamoyl-phosphate synthase large subunit

935(2) WP_109625247.1 WP_109625247.1 50S ribosomal protein L1

936(2) WP_109625248.1 WP_109625248.1 50S ribosomal protein L11

937(2) WP_109625249.1 WP_109625249.1 transcription termination/antitermination protein NusG

938(2) WP_109625250.1 WP_109625250.1 preprotein translocase subunit SecE

939(2) WP_112331603.1 WP_112331603.1 diol dehydratase reactivase subunit alpha

940(2) WP_112331604.1 WP_112331604.1 diol dehydratase small subunit

941(1) WP_035190286.1 WP_035190286.1 acetate kinase
941(1) WP_114298747.1 WP_114298747.1 acetate kinase

942(2) WP_121610591.1 WP_121610591.1 hypothetical protein

943(1) WP_122971523.1 WP_122971523.1 gamma-type small acid-soluble spore protein
943(1) WP_122971522.1 WP_122971522.1 hypothetical protein

944(1) WP_127570426.1 WP_127570426.1 transposase
944(1) WP_089522622.1 WP_089522622.1 transposase

945(1) WP_054847129.1 WP_054847129.1 hypothetical protein
945(1) WP_128705696.1 WP_128705696.1 YcxB family protein

946(1) WP_132008181.1 WP_132008181.1 DUF4352 domain-containing protein
946(1) WP_121610604.1 WP_121610604.1 DUF4352 domain-containing protein

947(1) WP_136789988.1 WP_136789988.1 DUF969 domain-containing protein
947(1) WP_043979920.1 WP_043979920.1 DUF969 domain-containing protein

948(1) WP_137698095.1 WP_137698095.1 uracil-DNA glycosylase
948(1) WP_137698094.1 WP_137698094.1 uracil-DNA glycosylase

949(2) WP_141523146.1 WP_141523146.1 phosphatase PAP2 family protein

950(2) WP_141523147.1 WP_141523147.1 cation diffusion facilitator family transporter

951(2) WP_141523148.1 WP_141523148.1 hypothetical protein

952(1) WP_148138297.1 WP_148138297.1 DUF389 domain-containing protein
952(1) WP_058271120.1 WP_058271120.1 DUF389 domain-containing protein

953(1) WP_148409450.1 WP_148409450.1 radical SAM protein
953(1) WP_109625251.1 WP_109625251.1 radical SAM protein

954(1) WP_109625252.1 WP_109625252.1 ECF transporter S component

954(1) WP_148409452.1 WP_148409452.1 ECF transporter S component

955(2) WP_149051232.1 WP_149051232.1 hypothetical protein

956(2) WP_149051233.1 WP_149051233.1 TM2 domain-containing protein

957(1) WP_149871687.1 WP_149871687.1 multidrug efflux SMR transporter
957(1) WP_149871686.1 WP_149871686.1 multidrug efflux SMR transporter

958(1) WP_152392569.1 WP_152392569.1 FTR1 family protein
958(1) WP_134750584.1 WP_134750584.1 FTR1 family iron permease

959(1) WP_152392571.1 WP_152392571.1 iron uptake transporter deferrochelatase/peroxidase subunit
959(1) WP_134750582.1 WP_134750582.1 iron uptake transporter deferrochelatase/peroxidase subunit

960(2) WP_155847254.1 WP_155847254.1 hypothetical protein

961(1) WP_156487456.1 WP_156487456.1 YheC/YheD family protein
961(1) WP_024027626.1 WP_024027626.1 YheC/YheD family protein

962(1) WP_157184217.1 WP_157184217.1 hypothetical protein
962(1) WP_013490122.1 WP_013490122.1 hypothetical protein

963(2) WP_158306558.1 WP_158306558.1 hypothetical protein

964(1) WP_158701619.1 WP_158701619.1 DUF443 family protein
964(1) WP_106496035.1 WP_106496035.1 DUF443 family protein

965(1) WP_163853469.1 WP_163853469.1 hypothetical protein
965(1) WP_063188049.1 WP_063188049.1 hypothetical protein

966(1) WP_165443462.1 WP_165443462.1 hypothetical protein, partial
966(1) WP_165443463.1 WP_165443463.1 hypothetical protein, partial

967(1) WP_168976282.1 WP_168976282.1 M48 family metallopeptidase
967(1) WP_021624830.1 WP_021624830.1 M48 family metallopeptidase

968(1) WP_168976284.1 WP_168976284.1 hypothetical protein
968(1) WP_021624827.1 WP_021624827.1 hypothetical protein

969(2) WP_168976297.1 WP_168976297.1 hypothetical protein

970(2) WP_175493626.1 WP_175493626.1 class I SAM-dependent methyltransferase

971(1) WP_176476000.1 WP_176476000.1 hypothetical protein
971(1) WP_170948037.1 WP_170948037.1 hypothetical protein

972(1) WP_179034800.1 WP_179034800.1 PIG-L family deacetylase
972(1) WP_165097637.1 WP_165097637.1 PIG-L family deacetylase

973(1) WP_179160628.1 WP_179160628.1 MmgE/PrpD family protein
973(1) WP_052106378.1 WP_052106378.1 MmgE/PrpD family protein

974(1) WP_179160632.1 WP_179160632.1 adenylosuccinate lyase
974(1) WP_034675386.1 WP_034675386.1 adenylosuccinate lyase

975(1) WP_034675432.1 WP_034675432.1 GNAT family N-acetyltransferase
975(1) WP_179160719.1 WP_179160719.1 GNAT family N-acetyltransferase

976(1) WP_183600875.1 WP_183600875.1 GbsR/MarR family transcriptional regulator
976(1) WP_166244887.1 WP_166244887.1 GbsR/MarR family transcriptional regulator

977(1) WP_166244893.1 WP_166244893.1 glycine betaine ABC transporter substrate-binding protein
977(1) WP_183600878.1 WP_183600878.1 glycine betaine ABC transporter substrate-binding protein

978(2) WP_185326091.1 WP_185326091.1 helix-turn-helix transcriptional regulator

979(2) WP_185327601.1 WP_185327601.1 replication initiation factor domain-containing protein

980(2) WP_185337285.1 WP_185337285.1 hypothetical protein

981(2) WP_185350588.1 WP_185350588.1 protein-export chaperone SecB

982(2) WP_185365739.1 WP_185365739.1 hypothetical protein

983(2) WP_185413323.1 WP_185413323.1 hypothetical protein

984(2) WP_185413325.1 WP_185413325.1 toxin B

985(2) WP_185413327.1 WP_185413327.1 hypothetical protein

986(2) WP_185413329.1 WP_185413329.1 hypothetical protein

987(2) WP_185413333.1 WP_185413333.1 hypothetical protein

988(1) WP_188535100.1 WP_188535100.1 hypothetical protein
988(1) WP_148450962.1 WP_148450962.1 hypothetical protein

989(1) WP_026572725.1 WP_026572725.1 IDEAL domain-containing protein
989(1) WP_191272893.1 WP_191272893.1 IDEAL domain-containing protein

990(2) WP_194230412.1 WP_194230412.1 hypothetical protein

991(1) WP_195244541.1 WP_195244541.1 GNAT family N-acetyltransferase
991(1) WP_040418522.1 WP_040418522.1 GNAT family N-acetyltransferase

992(1) WP_195953742.1 WP_195953742.1 fructose-6-phosphate aldolase
992(1) WP_106496011.1 WP_106496011.1 fructose-6-phosphate aldolase

993(1) WP_112331586.1 WP_112331586.1 hypothetical protein
993(1) WP_195974878.1 WP_195974878.1 hypothetical protein

994(1) WP_195974883.1 WP_195974883.1 4Fe-4S dicluster domain-containing protein
994(1) WP_112331595.1 WP_112331595.1 4Fe-4S dicluster domain-containing protein

995(1) WP_197937404.1 WP_197937404.1 hypothetical protein
995(1) WP_090688982.1 WP_090688982.1 hypothetical protein

996(1) WP_204445250.1 WP_204445250.1 Fe-S cluster assembly scaffold protein NifU
996(1) WP_039871624.1 WP_039871624.1 SUF system NifU family Fe-S cluster assembly protein

997(1) WP_055109364.1 WP_055109364.1 RDD family protein
997(1) WP_205001914.1 WP_205001914.1 RDD family protein

998(1) WP_205001912.1 WP_205001912.1 hypothetical protein
998(1) WP_205001926.1 WP_205001926.1 hypothetical protein

999(1) WP_207994172.1 WP_207994172.1 PH domain-containing protein
999(1) WP_017149782.1 WP_017149782.1 PH domain-containing protein

1000(1) WP_090688931.1 WP_090688931.1 metallophosphoesterase family protein
1000(1) WP_208019336.1 WP_208019336.1 metallophosphoesterase family protein

1001(1) WP_090688932.1 WP_090688932.1 hypothetical protein
1001(1) WP_208019337.1 WP_208019337.1 hypothetical protein

1002(1) WP_208926643.1 WP_208926643.1 AbrB/MazE/SpoVT family DNA-binding domain-containing protein
1002(1) WP_083426531.1 WP_083426531.1 AbrB/MazE/SpoVT family DNA-binding domain-containing protein

1003(1) WP_209365686.1 WP_209365686.1 helix-turn-helix domain-containing protein
1003(1) WP_074598768.1 WP_074598768.1 helix-turn-helix domain-containing protein

1004(1) WP_209365687.1 WP_209365687.1 hypothetical protein
1004(1) WP_175486786.1 WP_175486786.1 PH domain-containing protein

1005(1) WP_209365688.1 WP_209365688.1 isochorismatase family protein
1005(1) WP_074598770.1 WP_074598770.1 isochorismatase family protein

1006(1) WP_074598771.1 WP_074598771.1 hypothetical protein
1006(1) WP_209365689.1 WP_209365689.1 hypothetical protein

1007(1) WP_148362974.1 WP_148362974.1 aminoacyl-tRNA hydrolase
1007(1) WP_210388948.1 WP_210388948.1 aminoacyl-tRNA hydrolase

1008(1) WP_211745843.1 WP_211745843.1 peptide ABC transporter substrate-binding protein
1008(1) WP_015595775.1 WP_015595775.1 peptide ABC transporter substrate-binding protein

1009(1) WP_113672691.1 WP_113672691.1 ribonuclease toxin immunity protein CdiI
1009(1) WP_212693230.1 WP_212693230.1 ribonuclease toxin immunity protein CdiI

1010(1) WP_120112227.1 WP_120112227.1 hypothetical protein
1010(1) WP_212981554.1 WP_212981554.1 hypothetical protein

1011(1) WP_213532649.1 WP_213532649.1 hypothetical protein
1011(1) WP_191798519.1 WP_191798519.1 hypothetical protein

1012(1) WP_213532651.1 WP_213532651.1 hypothetical protein
1012(1) WP_213532650.1 WP_213532650.1 hypothetical protein

1013(1) WP_214705995.1 WP_214705995.1 hypothetical protein
1013(1) WP_214705994.1 WP_214705994.1 hypothetical protein

1014(1) WP_215032148.1 WP_215032148.1 hypothetical protein
1014(1) WP_214902963.1 WP_214902963.1 hypothetical protein

1015(1) WP_216390039.1 WP_216390039.1 M14 family metallopeptidase
1015(1) WP_216390038.1 WP_216390038.1 M14 family metallopeptidase

1016(1) WP_082420304.1 WP_082420304.1 hypothetical protein
1016(1) WP_216791210.1 WP_216791210.1 hypothetical protein

1017(2) WP_219872438.1 WP_219872438.1 anthranilate synthase component I

1018(2) WP_219872441.1 WP_219872441.1 DUF3817 domain-containing protein

1019(2) WP_219872449.1 WP_219872449.1 glycogen/starch synthase

1020(2) WP_219872450.1 WP_219872450.1 hypothetical protein

1021(1) WP_220559122.1 WP_220559122.1 hypothetical protein
1021(1) WP_091260855.1 WP_091260855.1 hypothetical protein

1022(1) WP_220559123.1 WP_220559123.1 nucleotidyltransferase domain-containing protein
1022(1) WP_217639392.1 WP_217639392.1 nucleotidyltransferase domain-containing protein

1023(2) WP_222860866.1 WP_222860866.1 putative holin-like toxin

1024(1) WP_223068053.1 WP_223068053.1 copper resistance protein CopC
1024(1) WP_152392562.1 WP_152392562.1 copper resistance protein CopC

1025(1) WP_216830397.1 WP_216830397.1 hypothetical protein
1025(1) WP_223592664.1 WP_223592664.1 hypothetical protein

1026(1) WP_223592666.1 WP_223592666.1 hypothetical protein
1026(1) WP_216830396.1 WP_216830396.1 hypothetical protein

1027(1) WP_034526650.1 WP_034526650.1 3-isopropylmalate dehydrogenase
1027(1) WP_223592703.1 WP_223592703.1 tartrate dehydrogenase

1028(1) WP_169238715.1 WP_169238715.1 Glu/Leu/Phe/Val dehydrogenase
1028(1) WP_224839856.1 WP_224839856.1 Glu/Leu/Phe/Val dehydrogenase

1029(2) WP_226086183.1 WP_226086183.1 hypothetical protein

1030(1) WP_226086185.1 WP_226086185.1 hypothetical protein
1030(1) WP_226642852.1 WP_226642852.1 hypothetical protein

1031(1) WP_005604654.1 WP_005604654.1 glucose-6-phosphate isomerase
1031(1) WP_232274501.1 WP_232274501.1 glucose-6-phosphate isomerase

1032(1) WP_019424138.1 WP_019424138.1 GNAT family N-acetyltransferase
1032(1) WP_233697182.1 WP_233697182.1 GNAT family N-acetyltransferase

1033(1) WP_019424136.1 WP_019424136.1 hypothetical protein
1033(1) WP_233697184.1 WP_233697184.1 hypothetical protein

1034(1) WP_019424134.1 WP_019424134.1 hypothetical protein
1034(1) WP_233697185.1 WP_233697185.1 hypothetical protein

1035(1) WP_235286189.1 WP_235286189.1 IDEAL domain-containing protein
1035(1) WP_181438749.1 WP_181438749.1 IDEAL domain-containing protein

1036(1) WP_111147813.1 WP_111147813.1 phosphate/phosphite/phosphonate ABC transporter substrate-binding protein
1036(1) WP_235286190.1 WP_235286190.1 phosphate/phosphite/phosphonate ABC transporter substrate-binding protein

1037(1) WP_235286194.1 WP_235286194.1 tRNA 2-selenouridine(34) synthase MnmH
1037(1) WP_111147809.1 WP_111147809.1 tRNA 2-selenouridine(34) synthase MnmH

1038(1) WP_235286297.1 WP_235286297.1 molybdopterin oxidoreductase family protein
1038(1) WP_111147972.1 WP_111147972.1 molybdopterin oxidoreductase family protein

1039(1) WP_227139377.1 WP_227139377.1 hypothetical protein
1039(1) WP_235369894.1 WP_235369894.1 hypothetical protein

1040(1) WP_235497058.1 WP_235497058.1 hypothetical protein
1040(1) WP_235497059.1 WP_235497059.1 hypothetical protein

1041(1) WP_123067101.1 WP_123067101.1 ElyC/SanA/YdcF family protein
1041(1) WP_236417193.1 WP_236417193.1 YdcF family protein

1042(1) WP_229752664.1 WP_229752664.1 GxGYxYP family putative glycoside hydrolase
1042(1) WP_237167829.1 WP_237167829.1 GxGYxYP family putative glycoside hydrolase

1043(1) WP_006289758.1 WP_006289758.1 hypothetical protein
1043(1) WP_237711208.1 WP_237711208.1 transcriptional regulator

1044(1) WP_006289757.1 WP_006289757.1 hypothetical protein
1044(1) WP_237711209.1 WP_237711209.1 hypothetical protein

1045(1) WP_237711210.1 WP_237711210.1 CRISPR-associated endonuclease Cas1
1045(1) WP_006290625.1 WP_006290625.1 type II CRISPR-associated endonuclease Cas1

1046(1) WP_022349202.1 WP_022349202.1 DUF5082 family protein
1046(1) WP_239458683.1 WP_239458683.1 DUF5082 domain-containing protein

1047(2) WP_241393146.1 WP_241393146.1 hypothetical protein

1048(2) WP_241393178.1 WP_241393178.1 ATP-binding protein

1049(2) WP_242172349.1 WP_242172349.1 cell wall hydrolase

1050(1) WP_084064112.1 WP_084064112.1 hypothetical protein
1050(1) WP_242251785.1 WP_242251785.1 tetratricopeptide repeat protein

1051(1) WP_084064113.1 WP_084064113.1 hypothetical protein
1051(1) WP_242251787.1 WP_242251787.1 hypothetical protein

1052(1) WP_242400480.1 WP_242400480.1 choline-binding protein A
1052(1) WP_000473028.1 WP_000473028.1 hypothetical protein

1053(1) WP_136789987.1 WP_136789987.1 DUF979 domain-containing protein
1053(1) WP_243495548.1 WP_243495548.1 DUF979 domain-containing protein

1054(1) WP_244527440.1 WP_244527440.1 DUF4083 family protein
1054(1) WP_001048096.1 WP_001048096.1 DUF4083 domain-containing protein

1055(1) WP_213649984.1 WP_213649984.1 tetratricopeptide repeat protein
1055(1) WP_244719544.1 WP_244719544.1 tetratricopeptide repeat protein

1056(1) WP_235815079.1 WP_235815079.1 tRNA pseudouridine(38-40) synthase TruA
1056(1) WP_246824640.1 WP_246824640.1 tRNA pseudouridine(38-40) synthase TruA

1057(1) WP_034671240.1 WP_034671240.1 proline--tRNA ligase
1057(1) WP_247083036.1 WP_247083036.1 proline--tRNA ligase

1058(1) WP_247083050.1 WP_247083050.1 hypothetical protein
1058(1) WP_028401645.1 WP_028401645.1 hypothetical protein

1059(1) WP_247083053.1 WP_247083053.1 DUF1128 domain-containing protein
1059(1) WP_028401646.1 WP_028401646.1 DUF1128 domain-containing protein

1060(1) WP_145406545.1 WP_145406545.1 TrkA family potassium uptake protein
1060(1) WP_248063408.1 WP_248063408.1 TrkA family potassium uptake protein

1061(1) WP_249897302.1 WP_249897302.1 mannonate dehydratase
1061(1) WP_181737049.1 WP_181737049.1 mannonate dehydratase

1062(1) WP_251417687.1 WP_251417687.1 hypothetical protein
1062(1) WP_126996335.1 WP_126996335.1 hypothetical protein

1063(2) WP_251426374.1 WP_251426374.1 5-oxoprolinase subunit PxpB

1064(1) WP_251466816.1 WP_251466816.1 hypothetical protein
1064(1) WP_199419784.1 WP_199419784.1 hypothetical protein

1065(1) WP_251466817.1 WP_251466817.1 hypothetical protein

1065(1) WP_199419781.1 WP_199419781.1 hypothetical protein

1066(1) WP_199419803.1 WP_199419803.1 DEAD/DEAH box helicase
1066(1) WP_251466820.1 WP_251466820.1 DEAD/DEAH box helicase

1067(1) WP_251426372.1 WP_251426372.1 biotin-dependent carboxyltransferase family protein
1067(1) WP_251557381.1 WP_251557381.1 biotin-dependent carboxyltransferase family protein

1068(1) WP_251557382.1 WP_251557382.1 5-oxoprolinase subunit PxpA
1068(1) WP_251426373.1 WP_251426373.1 5-oxoprolinase subunit PxpA

1069(1) WP_251426375.1 WP_251426375.1 hypothetical protein
1069(1) WP_251557383.1 WP_251557383.1 hypothetical protein

1070(1) WP_251426376.1 WP_251426376.1 hypothetical protein
1070(1) WP_251557384.1 WP_251557384.1 hypothetical protein

1071(1) WP_251426381.1 WP_251426381.1 HAD family acid phosphatase
1071(1) WP_251557387.1 WP_251557387.1 HAD family acid phosphatase

1072(1) WP_251426382.1 WP_251426382.1 bifunctional diaminohydroxyphosphoribosylaminopyrimidine deaminase/5-amino-6-(5-phosphoribosylamino)uracil reductase RibD
1072(1) WP_251557388.1 WP_251557388.1 bifunctional diaminohydroxyphosphoribosylaminopyrimidine deaminase/5-amino-6-(5-phosphoribosylamino)uracil reductase RibD

1073(1) WP_251596664.1 WP_251596664.1 mechanosensitive ion channel family protein
1073(1) WP_128631292.1 WP_128631292.1 mechanosensitive ion channel family protein

1074(1) WP_251638795.1 WP_251638795.1 spore protease YyaC
1074(1) WP_005604644.1 WP_005604644.1 spore protease YyaC

1075(1) WP_248251174.1 WP_248251174.1 hypothetical protein
1075(1) WP_256227708.1 WP_256227708.1 hypothetical protein

1076(1) WP_257443620.1 WP_257443620.1 hypothetical protein
1076(1) WP_090689970.1 WP_090689970.1 hypothetical protein

1077(1) WP_052947761.1 WP_052947761.1 hypothetical protein
1077(1) WP_257443632.1 WP_257443632.1 hypothetical protein

1078(1) WP_148409448.1 WP_148409448.1 zinc-ribbon domain-containing protein
1078(1) WP_257497520.1 WP_257497520.1 zinc ribbon domain-containing protein

1079(1) WP_258208945.1 WP_258208945.1 ureidoglycolate dehydrogenase
1079(1) WP_052352966.1 WP_052352966.1 Ldh family oxidoreductase

1080(1) WP_062239910.1 WP_062239910.1 DUF4317 domain-containing protein
1080(1) WP_265408857.1 WP_265408857.1 DUF4317 domain-containing protein

1081(1) WP_265444513.1 WP_265444513.1 AAA family ATPase
1081(1) WP_162302499.1 WP_162302499.1 AAA family ATPase

1082(1) WP_267934048.1 WP_267934048.1 GrpB family protein
1082(1) WP_019424137.1 WP_019424137.1 GrpB family protein

1083(1) WP_268058039.1 WP_268058039.1 ComEC/Rec2 family competence protein
1083(1) WP_089522620.1 WP_089522620.1 ComEC/Rec2 family competence protein

1084(1) WP_005604661.1 WP_005604661.1 HIT family protein
1084(1) WP_268058041.1 WP_268058041.1 histidine triad nucleotide-binding protein

1085(1) WP_019120064.1 WP_019120064.1 alanine--tRNA ligase
1085(1) WP_268058043.1 WP_268058043.1 alanine--tRNA ligase

1086(1) WP_235497056.1 WP_235497056.1 hypothetical protein
1086(1) WP_268753653.1 WP_268753653.1 hypothetical protein

1087(2) WP_268876582.1 WP_268876582.1 hypothetical protein

1088(1) WP_152392570.1 WP_152392570.1 EfeM/EfeO family lipoprotein
1088(1) WP_268999034.1 WP_268999034.1 Efem/EfeO family lipoprotein

1089(1) WP_251241787.1 WP_251241787.1 restriction endonuclease subunit S
1089(1) WP_269920250.1 WP_269920250.1 restriction endonuclease subunit S

1090(1) WP_251241786.1 WP_251241786.1 AAA family ATPase
1090(1) WP_269920251.1 WP_269920251.1 AAA family ATPase

1091(1) WP_269920252.1 WP_269920252.1 hypothetical protein
1091(1) WP_040036768.1 WP_040036768.1 hypothetical protein

1092(1) WP_270065069.1 WP_270065069.1 prolyl oligopeptidase family serine peptidase
1092(1) WP_066420586.1 WP_066420586.1 prolyl oligopeptidase family serine peptidase

1093(1) WP_272801677.1 WP_272801677.1 hypothetical protein
1093(1) WP_098248522.1 WP_098248522.1 hypothetical protein

1094(1) WP_275540580.1 WP_275540580.1 hypothetical protein
1094(1) WP_113672700.1 WP_113672700.1 hypothetical protein

1095(2) WP_275541676.1 WP_275541676.1 hypothetical protein

1096(1) WP_276538702.1 WP_276538702.1 50S ribosomal protein L33
1096(1) WP_109625285.1 WP_109625285.1 50S ribosomal protein L33

1097(1) WP_204666340.1 WP_204666340.1 hypothetical protein
1097(1) WP_276867630.1 WP_276867630.1 hypothetical protein

1098(1) WP_204666338.1 WP_204666338.1 hypothetical protein
1098(1) WP_276867634.1 WP_276867634.1 hypothetical protein

1099(1) WP_204666333.1 WP_204666333.1 DUF5082 family protein

| | | |
|---|---|---|
| 1099(1) | WP_276867642.1 | WP_276867642.1 hypothetical protein |
| 1100(1) | WP_276867656.1 | WP_276867656.1 hypothetical protein |
| 1100(1) | WP_204666330.1 | WP_204666330.1 hypothetical protein |
| 1101(1) | WP_204666403.1 | WP_204666403.1 GNAT family N-acetyltransferase |
| 1101(1) | WP_276867658.1 | WP_276867658.1 GNAT family N-acetyltransferase |
| 1102(1) | WP_204666327.1 | WP_204666327.1 TatD family hydrolase |
| 1102(1) | WP_276867672.1 | WP_276867672.1 TatD family hydrolase |
| 1103(1) | WP_276917294.1 | WP_276917294.1 hypothetical protein |
| 1103(1) | WP_168976296.1 | WP_168976296.1 hypothetical protein |
| 1104(1) | WP_277166567.1 | WP_277166567.1 DivIVA domain-containing protein |
| 1104(1) | WP_277150365.1 | WP_277150365.1 DivIVA domain-containing protein |
| 1105(1) | WP_277166575.1 | WP_277166575.1 hypothetical protein |
| 1105(1) | WP_277142060.1 | WP_277142060.1 hypothetical protein |
| 1106(1) | WP_277142062.1 | WP_277142062.1 hypothetical protein |
| 1106(1) | WP_277166576.1 | WP_277166576.1 hypothetical protein |
| 1107(1) | WP_277166578.1 | WP_277166578.1 hypothetical protein |
| 1107(1) | WP_277142064.1 | WP_277142064.1 hypothetical protein |
| 1108(1) | WP_277166579.1 | WP_277166579.1 DUF5344 family protein |
| 1108(1) | WP_277150353.1 | WP_277150353.1 hypothetical protein |
| 1109(1) | WP_277150367.1 | WP_277150367.1 alpha/beta hydrolase |
| 1109(1) | WP_277166630.1 | WP_277166630.1 alpha/beta hydrolase |
| 1110(1) | WP_277350978.1 | WP_277350978.1 hypothetical protein |
| 1110(1) | WP_276917292.1 | WP_276917292.1 hypothetical protein |
| 1111(1) | WP_255253069.1 | WP_255253069.1 hypothetical protein |
| 1111(1) | WP_277548451.1 | WP_277548451.1 hypothetical protein |
| 1112(1) | WP_155982582.1 | WP_155982582.1 hypothetical protein |
| 1112(1) | WP_278773444.1 | WP_278773444.1 hypothetical protein |
| 1113(1) | WP_278773446.1 | WP_278773446.1 hypothetical protein |
| 1113(1) | WP_155982583.1 | WP_155982583.1 hypothetical protein |
| 1114(1) | WP_278773448.1 | WP_278773448.1 hypothetical protein |
| 1114(1) | WP_036575552.1 | WP_036575552.1 hypothetical protein |
| 1115(1) | WP_278773461.1 | WP_278773461.1 hypothetical protein |
| 1115(1) | WP_013252370.1 | WP_013252370.1 hypothetical protein |
| 1116(1) | WP_278773475.1 | WP_278773475.1 hypothetical protein |
| 1116(1) | WP_036575570.1 | WP_036575570.1 hypothetical protein |
| 1117(1) | WP_094870392.1 | WP_094870392.1 glycosyl hydrolase family 65 protein |
| 1117(1) | WP_280788638.1 | WP_280788638.1 glycosyl hydrolase family 65 protein |
| 1118(1) | WP_280788641.1 | WP_280788641.1 DUF3231 family protein |
| 1118(1) | WP_094870386.1 | WP_094870386.1 DUF3231 family protein |
| 1119(1) | WP_281364067.1 | WP_281364067.1 mannitol dehydrogenase family protein |
| 1119(1) | WP_249897301.1 | WP_249897301.1 mannitol dehydrogenase family protein |
| 1120(2) | WP_281423142.1 | WP_281423142.1 serpin family protein |
| 1121(1) | WP_281477403.1 | WP_281477403.1 hypothetical protein |
| 1121(1) | WP_006674871.1 | WP_006674871.1 hypothetical protein |
| 1122(1) | WP_281477435.1 | WP_281477435.1 hypothetical protein |
| 1122(1) | WP_127510900.1 | WP_127510900.1 hypothetical protein |
| 1123(1) | WP_281667404.1 | WP_281667404.1 endonuclease/exonuclease/phosphatase family protein |
| 1123(1) | WP_006290911.1 | WP_006290911.1 endonuclease/exonuclease/phosphatase family protein |
| 1124(1) | WP_036695745.1 | WP_036695745.1 signal recognition particle protein |
| 1124(1) | WP_281667405.1 | WP_281667405.1 signal recognition particle protein |
| 1125(1) | WP_281667406.1 | WP_281667406.1 cysteine--tRNA ligase |
| 1125(1) | WP_237711212.1 | WP_237711212.1 cysteine--tRNA ligase |
| 1126(2) | WP_281813073.1 | WP_281813073.1 hypothetical protein |
| 1127(2) | WP_281813086.1 | WP_281813086.1 hypothetical protein |
| 1128(1) | WP_211023018.1 | WP_211023018.1 hypothetical protein |
| 1128(1) | WP_283904337.1 | WP_283904337.1 hypothetical protein |
| 1129(1) | WP_283927231.1 | WP_283927231.1 hypothetical protein |
| 1129(1) | WP_269058931.1 | WP_269058931.1 hypothetical protein |
| 1130(1) | WP_244996684.1 | WP_244996684.1 J domain-containing protein |
| 1130(1) | WP_283928357.1 | WP_283928357.1 DnaJ domain-containing protein |
| 1131(1) | WP_286017537.1 | WP_286017537.1 MBL fold metallo-hydrolase |
| 1131(1) | WP_208019341.1 | WP_208019341.1 MBL fold metallo-hydrolase |
| 1132(1) | WP_068536392.1 | WP_068536392.1 hypothetical protein |
| 1132(1) | WP_286017558.1 | WP_286017558.1 hypothetical protein |
| 1133(1) | WP_072336481.1 | WP_072336481.1 DUF6445 family protein |
| 1133(1) | WP_286119520.1 | WP_286119520.1 DUF6445 family protein |
| 1134(1) | WP_286124795.1 | WP_286124795.1 IS21-like element helper ATPase IstB |
| 1134(1) | WP_283889660.1 | WP_283889660.1 IS21-like element IS643 family helper ATPase IstB |

1135(1) WP_288218595.1 WP_288218595.1 GtrA family protein
1135(1) WP_196048860.1 WP_196048860.1 GtrA family protein

1136(1) WP_288218596.1 WP_288218596.1 hypothetical protein
1136(1) WP_055277417.1 WP_055277417.1 hypothetical protein

1137(1) WP_160037469.1 WP_160037469.1 polysaccharide deacetylase family protein
1137(1) WP_289342249.1 WP_289342249.1 polysaccharide deacetylase family protein

1138(1) WP_289376840.1 WP_289376840.1 hypothetical protein
1138(1) WP_184182963.1 WP_184182963.1 hypothetical protein

1139(1) WP_289376843.1 WP_289376843.1 hypothetical protein
1139(1) WP_154985935.1 WP_154985935.1 hypothetical protein

1140(1) WP_289342258.1 WP_289342258.1 hypothetical protein
1140(1) WP_289398704.1 WP_289398704.1 hypothetical protein

1141(1) WP_294386438.1 WP_294386438.1 hypothetical protein
1141(1) WP_294386442.1 WP_294386442.1 hypothetical protein

1142(1) WP_294386463.1 WP_294386463.1 hypothetical protein
1142(1) WP_294386466.1 WP_294386466.1 hypothetical protein

1143(2) WP_295628299.1 WP_295628299.1 endopeptidase La

1144(2) WP_295628320.1 WP_295628320.1 hypothetical protein

1145(1) WP_295629808.1 WP_295629808.1 DUF4405 domain-containing protein
1145(1) WP_251466769.1 WP_251466769.1 DUF4405 domain-containing protein

1146(1) WP_099361323.1 WP_099361323.1 peptide chain release factor 2
1146(1) WP_304216842.1 WP_304216842.1 peptide chain release factor 2

1147(1) WP_099361325.1 WP_099361325.1 DUF4230 domain-containing protein
1147(1) WP_304216845.1 WP_304216845.1 DUF4230 domain-containing protein

1148(1) WP_143315260.1 WP_143315260.1 CIA30 family protein
1148(1) WP_305022932.1 WP_305022932.1 glycosyl hydrolase

1149(1) WP_239548980.1 WP_239548980.1 LCP family protein
1149(1) WP_306077400.1 WP_306077400.1 LytR family transcriptional regulator

1150(1) WP_307192884.1 WP_307192884.1 alanine/glycine:cation symporter family protein
1150(1) WP_307192883.1 WP_307192883.1 alanine/glycine:cation symporter family protein

1151(1) WP_307192911.1 WP_307192911.1 GapA-binding peptide SR1P
1151(1) WP_254759288.1 WP_254759288.1 GapA-binding peptide SR1P

1152(1) WP_307199956.1 WP_307199956.1 RNA polymerase sporulation sigma factor SigK
1152(1) WP_019120042.1 WP_019120042.1 RNA polymerase sporulation sigma factor SigK

1153(1) WP_307199962.1 WP_307199962.1 hypothetical protein
1153(1) WP_114298766.1 WP_114298766.1 hypothetical protein

1154(1) WP_307199965.1 WP_307199965.1 Imm30 family immunity protein
1154(1) WP_043067140.1 WP_043067140.1 Imm30 family immunity protein

1155(1) WP_291289929.1 WP_291289929.1 biosynthetic-type acetolactate synthase large subunit
1155(1) WP_307284653.1 WP_307284653.1 pyruvate oxidase

1156(1) WP_257029484.1 WP_257029484.1 integrase
1156(1) WP_307312512.1 WP_307312512.1 integrase

1157(1) WP_307312516.1 WP_307312516.1 hypothetical protein
1157(1) WP_218895891.1 WP_218895891.1 hypothetical protein

1158(1) WP_307388980.1 WP_307388980.1 hypothetical protein
1158(1) WP_013252382.1 WP_013252382.1 hypothetical protein

1159(1) WP_013252381.1 WP_013252381.1 hypothetical protein
1159(1) WP_307388982.1 WP_307388982.1 hypothetical protein

1160(1) WP_036575554.1 WP_036575554.1 hypothetical protein
1160(1) WP_307388992.1 WP_307388992.1 hypothetical protein

1161(1) WP_307394624.1 WP_307394624.1 hypothetical protein
1161(1) WP_145406466.1 WP_145406466.1 hypothetical protein

1162(1) WP_307399092.1 WP_307399092.1 hypothetical protein
1162(1) WP_019912171.1 WP_019912171.1 hypothetical protein

1163(1) WP_005815658.1 WP_005815658.1 SsrA-binding protein SmpB
1163(1) WP_307399098.1 WP_307399098.1 SsrA-binding protein SmpB

1164(1) WP_000195483.1 WP_000195483.1 formate C-acetyltransferase
1164(1) WP_307589617.1 WP_307589617.1 formate C-acetyltransferase

1165(1) WP_307589625.1 WP_307589625.1 hypothetical protein
1165(1) WP_307589621.1 WP_307589621.1 hypothetical protein

1166(1) WP_045518287.1 WP_045518287.1 hypothetical protein
1166(1) WP_307601039.1 WP_307601039.1 hypothetical protein

1167(1) WP_308082229.1 WP_308082229.1 hypothetical protein
1167(1) WP_307284651.1 WP_307284651.1 hypothetical protein

1168(1) WP_307284658.1 WP_307284658.1 hypothetical protein
1168(1) WP_308082234.1 WP_308082234.1 hypothetical protein

1169(1) WP_042462838.1 WP_042462838.1 reverse transcriptase-like protein
1169(1) WP_308082238.1 WP_308082238.1 reverse transcriptase-like protein

1170(1) WP_308082240.1 WP_308082240.1 restriction endonuclease

| | | |
|---|---|---|
| 1170(1) | WP_231417875.1 | WP_231417875.1 restriction endonuclease |
| 1171(1) | WP_308083970.1 | WP_308083970.1 AAA domain-containing protein |
| 1171(1) | WP_189018630.1 | WP_189018630.1 AAA domain-containing protein |
| 1172(1) | WP_308110106.1 | WP_308110106.1 tubulin-like doman-containing protein |
| 1172(1) | WP_166245149.1 | WP_166245149.1 tubulin-like doman-containing protein |
| 1173(1) | WP_308110107.1 | WP_308110107.1 hypothetical protein |
| 1173(1) | WP_166245147.1 | WP_166245147.1 hypothetical protein |
| 1174(1) | WP_308110118.1 | WP_308110118.1 UDP-glucose/GDP-mannose dehydrogenase family protein |
| 1174(1) | WP_149871689.1 | WP_149871689.1 UDP-glucose/GDP-mannose dehydrogenase family protein |
| 1175(1) | WP_238579119.1 | WP_238579119.1 M15 family metallopeptidase |
| 1175(1) | WP_308110126.1 | WP_308110126.1 M15 family metallopeptidase |
| 1176(1) | WP_045518292.1 | WP_045518292.1 multicopper oxidase domain-containing protein |
| 1176(1) | WP_308110941.1 | WP_308110941.1 multicopper oxidase domain-containing protein |
| 1177(1) | WP_291289947.1 | WP_291289947.1 DUF6020 family protein |
| 1177(1) | WP_308615630.1 | WP_308615630.1 glycosyltransferase |
| 1178(2) | WP_309298633.1 | WP_309298633.1 hypothetical protein |
